# An improved de novo assembly and annotation of the tomato reference genome using single-molecule sequencing, Hi-C proximity ligation and optical maps

**DOI:** 10.1101/767764

**Authors:** Prashant S. Hosmani, Mirella Flores-Gonzalez, Henri van de Geest, Florian Maumus, Linda V. Bakker, Elio Schijlen, Jan van Haarst, Jan Cordewener, Gabino Sanchez-Perez, Sander Peters, Zhangjun Fei, James J. Giovannoni, Lukas A. Mueller, Surya Saha

## Abstract

The original Heinz 1706 reference genome was produced by a large team of scientists from across the globe from a variety of input sources that included 454 sequences in addition to full-length BACs, BAC and fosmid ends sequenced with Sanger technology. We present here the latest tomato reference genome (SL4.0) assembled *de novo* from PacBio long reads and scaffolded using Hi-C contact maps. The assembly was validated using Bionano optical maps and 10X linked-read sequences. This assembly is highly contiguous with fewer gaps compared to previous genome builds and almost all scaffolds have been anchored and oriented to the 12 tomato chromosomes. We have found more repeats compared to the previous versions and one of the largest repeat classes identified are the LTR retrotransposons. We also describe updates to the reference genome and annotation since the last publication. The corresponding ITAG4.0 annotation has 4,794 novel genes along with 29,281 genes preserved from ITAG2.4. Most of the updated genes have extensions in the 5’ and 3’ UTRs resulting in doubling of annotated UTRs per gene. The genome and annotation can be accessed using SGN through BLAST database, Pathway database (SolCyc), Apollo, JBrowse genome browser and FTP available at https://solgenomics.net.

## Introduction

Tomato (*Solanum lycopersicum*) is one of the most valuable vegetable crops as well as an important model for processes such as fruit ripening (Tanksley et al. 1992; Klee and Giovannoni 2011), plant defense (Rosli and Martin 2015), and secondary metabolism (Schilmiller et al. 2010). The previous tomato genome sequence was published by an international consortium in 2012 (The Tomato Genome Consortium 2012), culminating years of effort to characterize the genome. Tanksley *et al*. published the first genetic map in 1992 (Tanksley et al. 1992), as well as an improved map with higher marker densities in 2000 called the F2-2000 map, based on conserved ortholog set (COS) markers (Fulton et al. 2002). In 2003, three years after the completion of the first plant genome (Initiative and The Arabidopsis Genome Initiative 2000), the tomato genome sequencing project was launched as an international collaboration. At the outset, it followed a BAC-by-BAC sequencing approach, mirroring the strategy that was used for Arabidopsis. Many resources were established during that phase, including BAC libraries, BAC-end sequences, and fosmid libraries. With the remarkable progress in sequencing technology around that time, a whole genome shotgun approach became feasible around 2010, using the 454 sequencing technology (Margulies et al. 2005). Illumina short reads (Voelkerding, Dames, and Durtschi 2009) were used to correct the homopolymer errors in 454 sequences. In addition, FISH data was generated to size the inter-scaffold gaps (Chang et al. 2007), resulting in re-ordering and re-orientation of a limited number of scaffolds, providing a major improvement in scaffold order. Altogether these resulted in the assembly SL2.40 as well as the ITAG2.30 annotation that was published in 2012 (The Tomato Genome Consortium 2012).

After the publication, the US partners of the genome project continued targeted full BAC sequencing for filling intra-scaffold gaps. The project initially focused on chromosomes 1 and 10, which had been assigned to the US partners in the BAC by BAC approach. However, later, other chromosomes were also covered and these BACs were integrated into the assembly, which resulted in a more complete assembly SL3.0. Furthermore, the tomato SL3.0 reference genome has accurate scaffold order and orientation that has been manually validated using FISH data (Shearer et al. 2014) and optical maps. However, a large number of inter and intra-scaffold gaps (81.71Mb) remain that can limit the resolution of genetic and genomic analysis besides confounding genome annotation. There are also 3136 contigs in SL3.0 assembly that were not placed in chromosomal locations based on the available evidence at the time. Rapid technical advancements in single-molecule sequencing and chromosome conformation capture technologies have enabled the assembly of high-quality genome assemblies for plants (Zhang et al. 2018; Edger et al. 2019). An improved reference genome for Heinz 1706 will aid quantitative trait analysis and genome-wide breeding efforts, permit accurate comparative analysis especially among the Solanaceae family and enable better mapping of repeat regions. Therefore, we used PacBio single-molecule sequences for *de novo* assembly followed by scaffolding based on Hi-C contact maps to generate a highly contiguous reference assembly for tomato with much fewer gaps (44Kb) and unplaced contigs in addition to better assembly of repetitive regions. The chromosomal length scaffolds were manually validated with Bionano optical maps and 10X sequences, and short read sequences were used to correct sequencing errors. This new assembly is highly collinear with previous versions, but contains fewer gaps and undefined bases (Ns). We also report an updated annotation for tomato, ITAG4.0, which has improved gene structure as well as a number of new genes.

## Results and Discussion

### Improvements in the legacy genome assembly

We integrated 1,069 full-length (phase htgs3) BACs into the SL2.50 genome to cover gaps and replace short whole genome shotgun contigs from 454 sequences. This resulted in removal of 11,699,806 bases (11Mb) of contig gaps in the SL3.0 genome. The reduction in contig gaps varied from 3.17% to 49.07% per chromosome based on the density of BACs. Chromosomes 1 and 10 benefited the most from the BAC integrations as they had the highest numbers of BAC integrated at 329 and 439 BACs respectively.

The contiguity and quality of the assembly was improved and validated using the high-throughput Irys system from Bionano Genomics (San Diego, CA, USA). Bionano optical maps can accurately measure the size of fluorescently labelled restriction fragments, providing an independent validation to verify assemblies. We implemented a modified overlap-layout-consensus strategy by increasing the stringency iteratively (see methods) to increase the scaffold length and coverage. The resulting *de novo* optical map based assembly with 674 cmaps had a total length of 762 Mb and an N50 of 1.48 Mb, with the largest contig size of 5.5 Mb. Using the Bionano Irysview tool, we detected conflicting regions in the alignment between the Bionano optical map assembly and the BAC-integrated assembly. These were manually validated and corrections made to the assembly when supported by sufficient evidence. As a result, two inversions were corrected in Chr12 (Supp. Figure 1), 19 gaps on the genome were resized and Chr00 contigs were integrated in Chr02 and Chr09 in the SL3.0 reference genome.

The whole genome shotgun contigs and many BACs integrated in the genome were sequenced with 454 technology which has a high homopolymer error rate (Gilles et al. 2011; Huse et al. 2007). We identified and corrected 16,723 homopolymer insertion and deletion errors using high quality Illumina sequences. These corrections were used to create final SL3.0 chromosomes. Short contigs on chromosome 0 ranging in length from 2 to 4.8kb were found to be of mitochondrial or bacterial origin during the submission process so these were also removed. The SL3.0 tomato genome reference is available at NCBI (AEKE00000000.3) and the SGN FTP site (ftp://ftp.solgenomics.net/tomato_genome/assembly/build_3.00/).

### *De novo* assembly SL4.0

The SL4.0 reference genome was assembled using Single Molecule Real-Time sequencing. The Canu assembler (Koren et al. 2017) was used to assemble 80x coverage of PacBio long reads, generating 504 contigs. The contigs were error corrected with PacBio reads and Illumina data. Raw assembly with corrected reads resulted in highly continuous assembly of 785 Mbp with contig N50 of 5.5 Mbp (Table 1). Hi-C contact maps were used to order and orient the contigs in 12 super-scaffolds corresponding to 12 chromosomes of tomato. Chromosomes were oriented and numbered according to SL3.0 build. The remaining 152 contigs which were not scaffolded in the 12 chromosomes were stitched together to create chromosome 00 (9.6 Mbp) with a 100 bp gap inserted between adjacent contigs.

**Table 1:** Genome assembly comparisons for different versions along with the size of unplaced contigs (chromosome 00). K-mer comparison from independently generated Illumina data (paired-end reads with 20X coverage) to different genome assembly versions.

| **Genome Assembly versions** | **SL4.0** | **SL3.0** | **SL2.5** |
| --- | --- | --- | --- |
| **Assembly Size (bp)** | 782,520,133 | 828,076,956 | 823,944,041 |
| **Non-N bases** | 782,475,302 | 746,357,470 | 737,636,348 |
| **N's (bp)** | 44,831 | 81,719,486 | 86,307,693 |
| **Chr 00 / unplaced contig size (bp)** | 9,643,350 | 20,852,292 | 21,805,821 |
| **Number of Chr 00 contigs** | 176 | 4,374 | 4,410 |
| **Repeat content (RepeatModeler)** | 64.19% | 56.39% | 56.34% |
| **Repeat content (REPET)** | 71.77% | 61.55% | 60.94% |
| **Assembly completeness estimation based on k-mer's** | 99.24% | 98.96% | 98.83% |

Genome assemblies can be validated for completeness by comparing the k-mer’s present in the raw genomic DNA-Seq reads with those in the genome assembly (Mapleson et al. 2017). The k-mer spectra of SL4.0 genome assembly shows a single homozygous peak at the expected (20x) coverage based on k-mer analysis (Suppl. Figure 2). The tomato genome is highly homozygous and the SL4.0 genome assembly shows single peak representing homozygosity of the genome. K-mer analysis also indicates single large peak representing 99.24% completeness of the assembly (Table 1).

### Comparison of *de novo* assembled SL4.0 with previous reference assemblies

Comparison of SL4.0 with previous assemblies (SL2.5 and SL3.0) using k-mer analysis shows that earlier versions were already nearly complete. However, we have integrated more than 36 Mbp of novel sequence with large reduction in the number of gaps. Analysis of the newly integrated sequence in version 4.0 indicates that it largely consists of repeat rich sequences, which were difficult to assemble previously due to limitations of Illumina and 454 technology. Overall, there was approximately 8% increase in repeat regions in SL4.0 compared to SL3.0.

BUSCO (Simão et al. 2015) measures the completeness of a genome or annotation by benchmarking against a set of single-copy genes conserved within a phylogenetic clade. We evaluated the SL4.0 genome and ITAG4.0 annotation against the Embryophyta and Solanaceae single-copy gene sets (Supp Table 1). Of note, the BUSCO marker gene set was based on the legacy ITAG annotation so these fare well in the analysis. Overall, the BUSCO results and RNA-seq mapping rates (Supp Table 2) are consistent across all the genome builds showing that the gene space is retained in the new genome and annotation. However, DNA-Seq mapping results (Supp Table 2) show marginal increase in mapping rates indicating higher assembly completeness of SL4.0 similar to k-mer analysis results.

Scaffolding in the legacy assembly was based on a variety of data sets including BACs, BAC ends, FISH data, genetic and optical maps. Independently assembled SL4.0 is highly collinear with the SL3.0 as shown in Figure 1. This validates that chromosome conformation capture based Hi-C method is highly reliable and yields similar results to conventional scaffolding methods.

**Figure 1:**
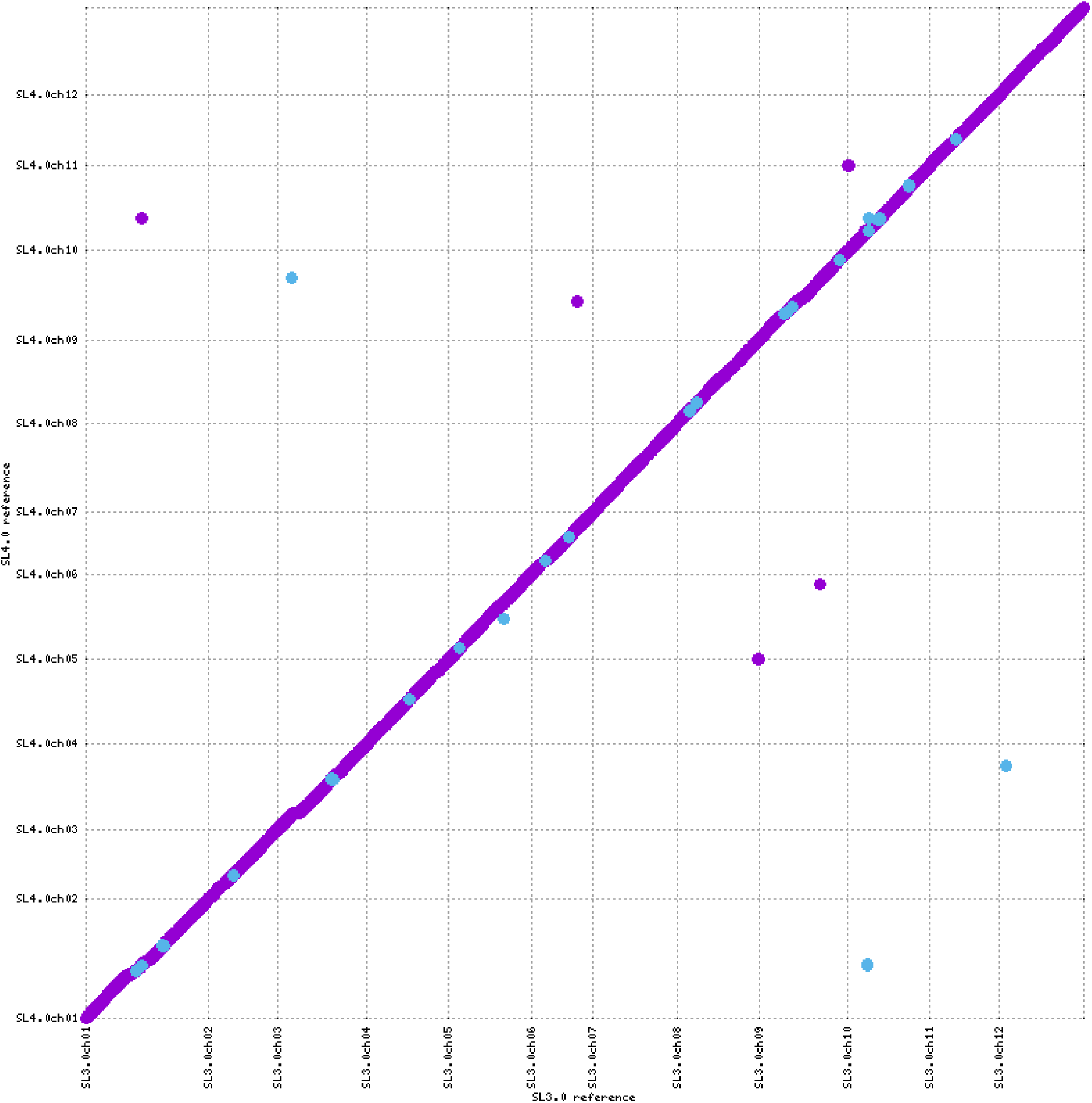
MUMmerplot (Marçais et al. 2018) comparison of all 12 chromosomes from SL3.0 on the X axis with the *de novo* assembled SL4.0 chromosomes on the Y axis. The chromosomes were aligned using NUCmer and the alignments were filtered for unique matches with a 10Kb length cut-off.

### Repeat Identification and Classification

The REPET (Flutre et al. 2011) pipeline identified 71.77% of the SL4.0 reference assembly as repetitive. In comparison, previous assemblies contained around 61% repeats (60.94% and 61.55% in SL2.5 and SL3.0 respectively). The REPET analysis identified repeat classes, shown in Figure 2 with respect to total repeats identified in SL2.5 and SL4.0 assembly versions. The distribution of repeats is similar to previous reports (The Tomato Genome Consortium 2012).

**Figure 2:**
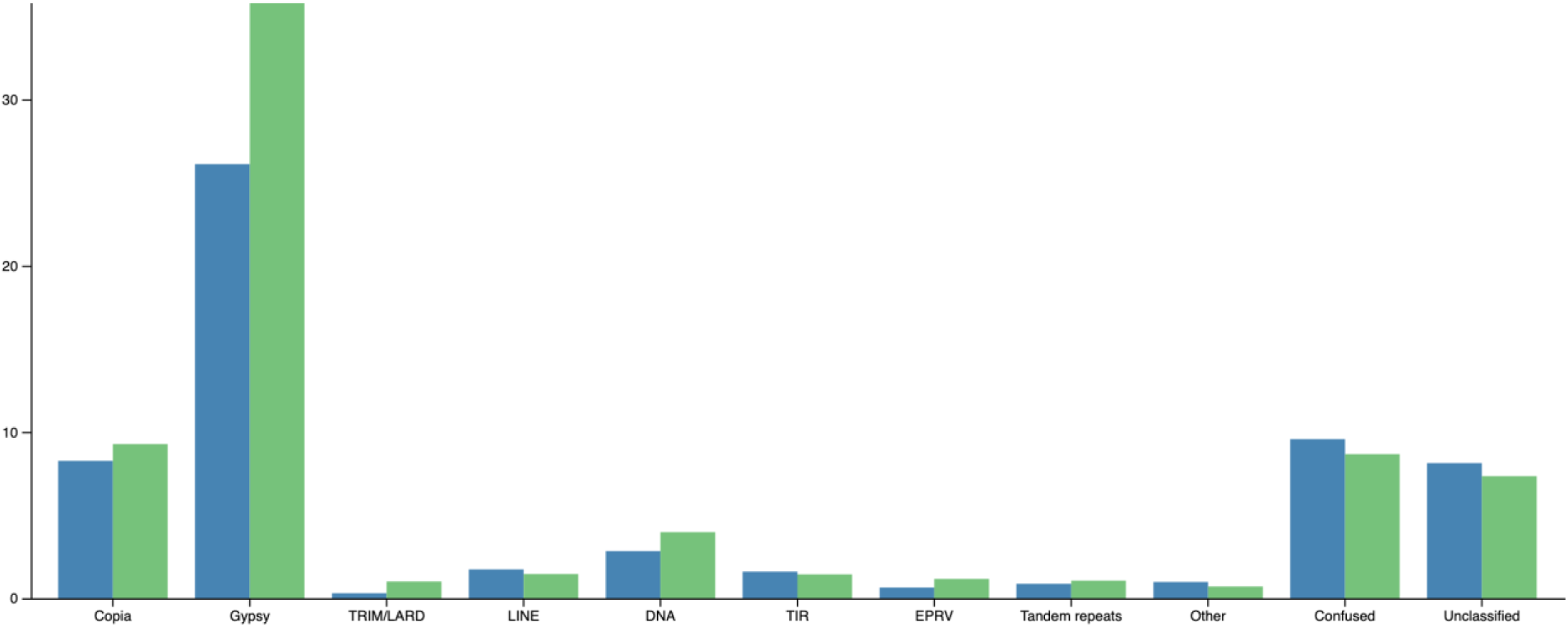
Comprehensive identification and classification of repeats using the REPET pipeline. Repeats identified and classified in SL2.5 are in blue and those in SL4.0 are in green. Repeat classes masking less than one percent of the genome were combined in the “other” category.

In long read based assemblies, complex repeat rich regions are more contiguous than traditional assemblies created from short read approaches. Accordingly, in the SL4.0 assembly, we have identified more repeats compared to the previous assemblies. One of the largest repeat classes identified are the LTR retrotransposons. Long terminal repeat (LTR) transposons are known to be abundant in plants (Galindo-González et al. 2017). Gypsy is the largest class of LTR retrotransposons identified in tomato. In comparison to SL2.50, we have identified 10% more gypsy transposons in SL4.0. Along with identification of more repeats, SL4.0 assembly has fewer unclassified repeats. This could also be the result of a more accurate repeat identification by REPET due to higher contiguity in SL4.0, leading to better characterization and classification.

### Protein coding gene annotation

The International Tomato Annotation Group (ITAG) is a consortium which originally released the annotation (ITAG2.3) of the tomato genome along with the release of the tomato Heinz 1706 SL2.40 reference genome assembly (The Tomato Genome Consortium 2012). The ITAG2.3 annotation contained 34,727 protein-coding gene models predicted using Eugene and partly manually curated by experts. Incorporation of FISH data enabled correction in the orientation and position of a number of scaffolds. This led to changes in the gene coordinates which were included in ITAG2.40 annotation released on February 23, 2014 with 34,725 genes. Annotation was also revised in ITAG3.2 based on improved SL3.0 assembly. The annotation pipeline implemented for ITAG3.2 and ITAG4.0 was similar and is described later in the methods section.

The PacBio based SL4.0 genome described here contains novel sequence information with fewer gaps compared to earlier versions. The availability of new and comprehensive expression profiles of tomato from a range of tissues provides us a rich resource for annotating the new genome. We collected publicly available RNA-Seq datasets generated from a variety of tissues and conditions of wild-type tomato plants (Supp Table 3). These RNA-Seq datasets were enriched in fruit datasets, as fruit development is a major focus of tomato research. RNA-Seq data were selected to represent root, leaf, flower and callus along with various fruit developmental stages. Other important datasets included NBS-LRR gene enriched RNA-Seq and 5’ and 3’ enriched RNA-Seq data.

### Migrating legacy annotation from ITAG2.4

ITAG2.4 protein-coding annotation with its Solyc IDs has been widely used in research across solanaceae community. In our efforts to preserve ITAG nomenclature, we mapped ITAG2.4 gene models to the new SL4.0 reference genome. First, we refined ITAG2.4 annotation by removing known repeat genes along with few other changes based on the community feedback and publications. We integrated manually curated gene annotations and removed 56 genes due to contamination from chloroplast and non-tomato sequences. In a recent report, there are a number of genes from ITAG2.4 that have similarity with known transposable elements (Jouffroy et al. 2016). Based on similarity with known TE, we removed 2,244 TE genes from ITAG2.4 before passing them to our annotation pipeline.The remaining genes were mapped to the repeat masked SL4.0 genome. We mapped ITAG2.4 cDNA to individual chromosomes to avoid cross-mapping to different regions with high stringency. We migrated total of 29,177 genes out of 33,838 genes from 12 chromosomes of ITAG2.4 genes and another 104 on chromosome 00 / unplaced contigs.

### Updating legacy gene models and identifying novel genes

We employed the MAKER annotation pipeline with a custom protocol to update the ITAG2.40 gene models and predict novel genes in SL4.0. A significant proportion (73%) of migrated ITAG2.4 genes were updated using diverse Illumina short read RNA-Seq and PacBio Iso-Seq expression data. This updated set has longer cDNA sequences and previously missing UTR regions. In the next run of MAKER, the updated ITAG2.4 gene models were used as passthrough in the MAKER annotation pipeline along with other gene prediction tools. *Ab-initio* gene predictors Augustus (Hoff and Stanke 2019) and SNAP (Li et al. 2007) enabled discovery of the novel genes. These gene predictors were trained on the reference genome assembly with all available evidence sources including expression and orthology data. The trained gene predictors helped to identify novel genes along with minor changes to the legacy gene models based on RNA-Seq support. The downstream analysis of annotations involved removing newly identified repeat genes based on similarity with the known repeats and incorporating manually curated genes from Apollo annotation editor (http://apollo.sgn.cornell.edu/apollo/) (Lee et al. 2013). This resulted in 34,075 protein coding gene annotation set ITAG4.0. Automated Assignment of Human Readable Descriptions (AHRD) (https://github.com/groupschoof/AHRD) using various plant databases predicted functions of 29,532 ITAG4.0 genes.

ITAG4.0 has 4,794 novel genes (Supp Table 4) along with 29,281 genes preserved from ITAG2.4. Many of the preserved genes between ITAG2.4 and ITAG4.0 have been updated (21,962 out of 29,281) in the current annotation (Supp Table 5). Most of the updated genes have extensions in the 5’ and 3’ UTRs resulting in doubling of UTRs per gene in ITAG4.0 compared to ITAG2.40. The extended UTRs in ITAG4.0 will allow better mapping of short read RNA-Seq data, especially from the cost-effective 3’ RNA-Seq protocol (Tandonnet and Torres 2017), giving a more accurate expression quantification.

### Annotation Edit Distance as a metric of quality for gene models

Comparing annotation sets is an intrinsically complex and challenging task in the absence of a gold standard set. Annotation Edit Distance (AED) provides an objective measure of the annotation quality given an evidence set. The AED is calculated for each gene by the MAKER annotation pipeline, which measures congruence of annotation with the evidence supporting that gene model (Campbell, Holt, et al. 2014). An AED score of 0 is the perfect score indicating concordance of all features in the gene model with the RNA-Seq and/or orthology data, while a score of 1 indicates a lack of any support from genome-independent data. Considering that tomato has a large collection of publicly available RNA-Seq data aggregated over the past decade, AED is a good metric to evaluate improvements in the quality of gene models. The cumulative fraction of transcripts of the AED plot shown in Figure 3 gives a genome wide perspective of the quality of annotation. The latest annotation from arabidopsis (Araport11) is included for comparison (Cheng et al. 2017). The recent update of arabidopsis annotation has also switched to using AED for quality estimation from the traditional 5 star ranking system. Overall, ITAG4.0 protein coding gene annotations have better AED scores compared to ITAG2.4 implying better accord with underlying expression and homology evidence. Importantly, there are more genes with supporting evidence in ITAG4.0 (83%) than ITAG2.4 (76%). We have successfully transferred 29,281 genes from ITAG2.4 to ITAG4.0. 21,962 of 29,281 genes have been updated (Supp Table 5) based on the supporting evidence which has resulted in better AED scores. Figure 3 shows that more than 80% of the ITAG4.0 genes have AED score of less than 0.5 similar to arabidopsis protein coding gene annotation (Araport11) indicating strong evidence support for these genes.

**Figure 3:**
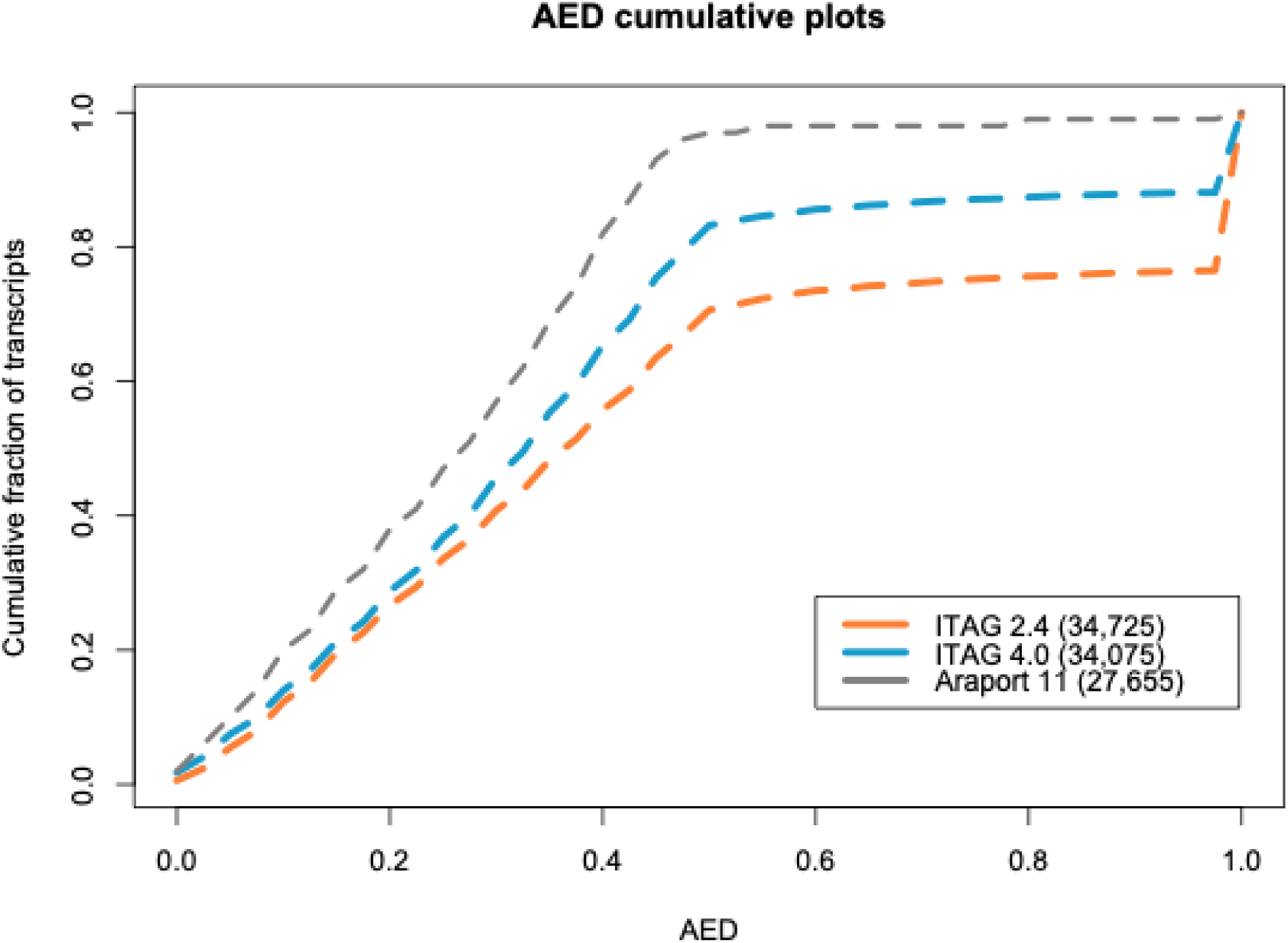
AED cumulative plot shows improvements in the ITAG4.0 compared to ITAG2.4. Genome-wide cumulative fraction of transcripts for ITAG2.4, ITAG4.0 and Araport 11 annotation are shown.

### Novel protein coding genes are involved in the key biological processes

Another important feature of the ITAG4.0 is the identification of novel genes. There are a total of 4,794 newly identified genes using *de novo* gene prediction methods in ITAG4.0. These novel genes have either expression evidence support or supported by homology with SwissProt plant protein database. These newly identified genes belong to various biological processes. Gene ontology (GO) enrichment was performed to see any bias in identification of novel genes. The plot of GO-term enrichment of novel genes (Figure 4) compared to all the genes in ITAG4.0 shows that they are involved in many key biological processes. The prominent classes of GO terms enriched in novel genes are related to stress responses. GO-terms related to stress response include defense response (GO:0006952), response to fungus (GO:0009620), defense response to fungus (GO:0050832), response to biotic stimulus (GO:0009607), response to stress (GO:0006950). Identification of stress related genes correlates with the abundance of expression data related to plants treated with fungal pathogens. Incorporation of RenSeq sequencing method enriches for nucleotide binding-site leucine-rich repeat genes (NB-LRR) (Jupe et al. 2013) has also aided in identification of genes related to disease response.

**Figure 4:**
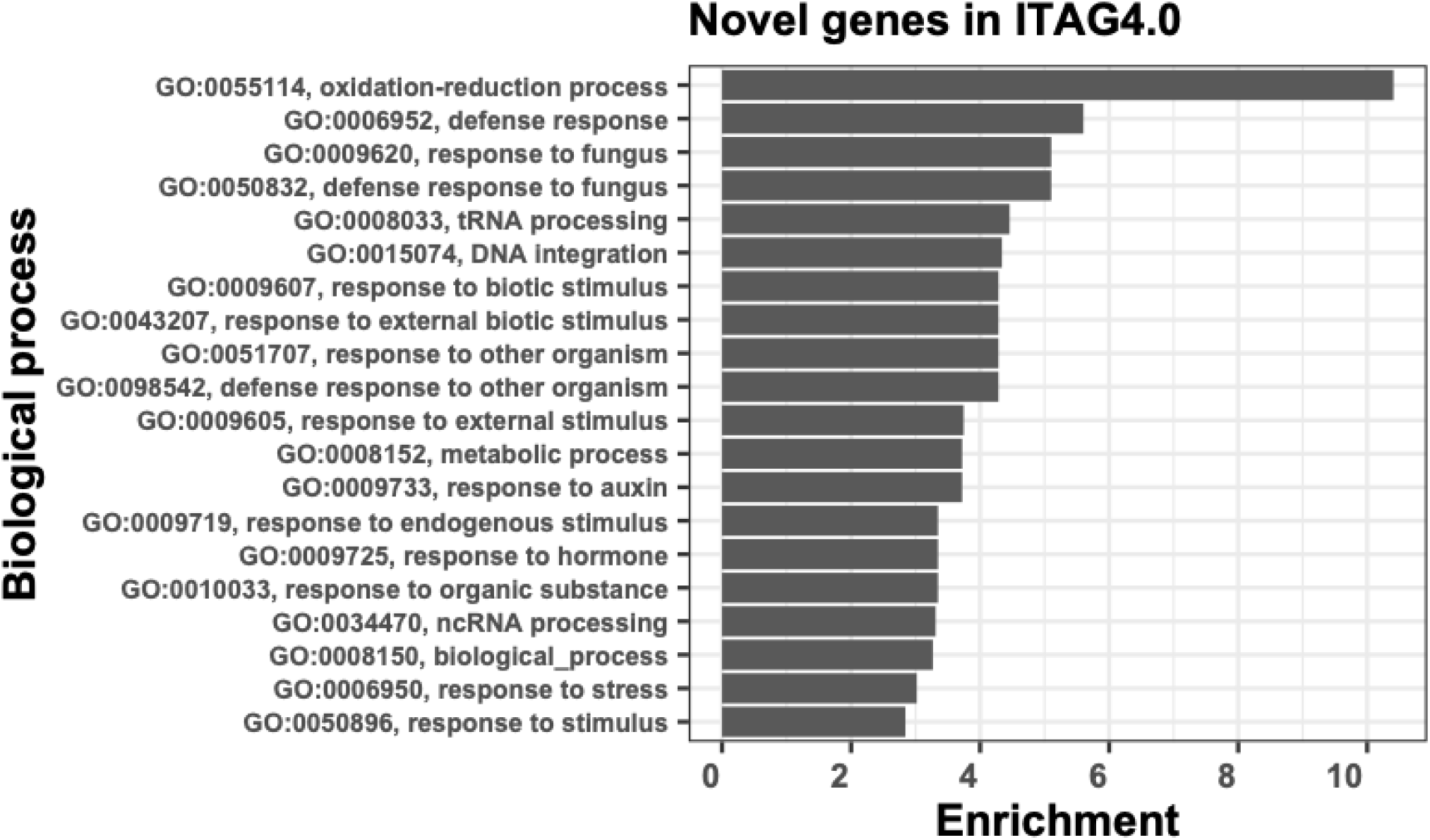
Novel genes in ITAG4.0 are enriched in key biological processes. GO-terms enriched in novel genes are shown as fold enriched in minus log10 of their corresponding P-values.

### Long non-coding RNA identification

Long non-coding RNAs or lncRNA are relatively newly described class of RNAs and thought to be involved in the regulation of many biological processes. LncRNA are defined as transcripts with very low coding potential compared with known protein-coding genes and with length longer than 200 nucleotides. During annotation of protein-coding genes, we have used diverse expression data from Iso-Seq and RNA-seq. The diverse expression data enabled identification of lncRNA’s. Mikado transcriptome discussed above was filtered for known protein-coding genes. The remaining transcripts were the basis for identification of lncRNAs using FEELnc pipeline (Wucher et al. 2017). In total 5,874 lncRNA genes were predicted with 6,694 alternately spliced isoforms.

### Gene family analysis

Gene family analysis was performed with a total of 409,806 genes among 11 species (Supp. Figure 3). We were able to assign 85% genes (348,442) to 7,445 orthogroups. *S. lycopersicum* had 15,933 families and six were unique to tomato. Within the solanaceae group, *S. lycopersicum* shared 14,854 gene families with *S. pennellii* and 13,866 families with *S. tuberosum*. Gene family expansion and contraction analysis were performed on the inferred gene families. We observed more expanded orthogroups in Solanaceae (1,399) than Capsicum (993). We found more contracted families in petunia clade (4,566) compared to Solanaceae (1,778) (Supp. Figure 3). Among the Solanaceae clade, *S. lycopersicum* had fewer contracted gene families (1,496) and expanded gene families (749) compared to *S. tuberosum* (7,046). According to the analysis, 1,642 genes were found to be gained and 2,010 genes lost in *S. lycopersicum* compared to the solanum ancestor.

## Conclusion

Assembly of long read PacBio sequences and scaffolding with Hi-C chromatin capture has yielded a high-quality reference genome assembly of tomato, with major improvement in the contiguity of the genome compared to previous genome builds that were based primarily on 454 sequences. Characterization of repetitive structure of the genome shows that we have identified 10% more repeats compared to previous assemblies. There are numerous improvements in the structure of protein-coding genes in ITAG4.0 annotation Full-length cDNA sequencing using PacBio Iso-Seq has contributed to the accuracy of the current genome annotation. But manual genome curation is still the gold standard for the genome annotation. We provide Apollo annotation editor as a tool to curate the structure and function of genes and the locus editor to add publications and metadata for genes. The latest genome and annotation can be accessed using SGN through BLAST database, Pathway database (SolCyc), JBrowse genome browser and FTP available at https://solgenomics.net (Fernandez-Pozo et al. 2015).

## Acknowledgements

We would like to thank NCBI staff including Valerie Schneider, Anjana Raina and Karen Clark for providing Genome Reference Consortium (https://www.ncbi.nlm.nih.gov/grc) resources and guidance for submission. We would like to thank members of the Solanaceae community for providing expression data sets for annotation of the genome assembly.

## Methods

### SL3.0 and ITAG3.2

#### FISH-based scaffolding

Scaffolds were ordered and assigned to individual chromosomes based on a high density genetic map (F2-2000). Large gaps were sized by Fluorescence In-Situ Hybridization (FISH) (Shearer et al. 2014). A default inter-scaffold gap size of 100Kb was used as a default in cases where order was known but the gap size could not be accurately estimated.

#### BAC-based gap filling

To close smaller, intra-scaffold gaps in the whole genome assembly, we used high quality BAC sequences that had not previously been integrated in the assembly. We selected only completely sequenced BACs without any Ns and with a length greater than 20Kb. The BACs can be downloaded from NCBI or the SGN FTP site (ftp://ftp.solgenomics.net/genomes/Solanum_lycopersicum/Heinz1706/bacs/). The 2764 high throughput genomic sequences (HTGS) phase 3 BACs were separated by chromosome and assembled into larger BAC contigs to remove potential redundancy among the BACs and identify sequences representing longer BAC tiling paths. The assembly step enabled us to align 15 to 50% of the BACs per chromosome. All assembled contigs from BACs were manually validated and only the contigs with a mismatch rate of 1% or less were selected for the next step. Most of the mismatches were at the ends of BACs which can be attributed to sequencing or assembly error when the original BAC was created.

We aligned 500bp BAC ends to the SL2.50 genome with perfect identity to determine the location and orientation. All the regions with integrated BACs were carefully screened manually and the longest BAC was selected where multiple BACs spanned a genomic region. We used the NCBI GRC (https://www.ncbi.nlm.nih.gov/grc) pipeline to integrate selected BACs into the tomato genome. A description of the pipeline and the file formats such as the tiling path file (TPF) and accessioned genome path (AGP) used by the GRC pipeline are available on the GRC website. We created TPF files that included the integrated BACs and substituted out shorter WGS contigs. These files were submitted to the GRC pipeline to validate the overlaps between adjacent WGS contigs and BACs. We identified a number of cases where the ends of the WGS contigs from SL2.50 had errors. The sequence from BACs was selected to fill those regions as the BACs were of higher quality. In a quality control step, we identified E coli transposon insertions in some BACs. Both the IS10 transposon and the flanking target site duplicated region were excised from the BAC sequences before they were integrated into the chromosomes. We also identified and removed chloroplast contamination in chromosomes 1 (63kb), chromosome 9 (7kb) and chromosome 10 (6.5kb). We also removed a 6.7 kb contig at the end of a scaffold in chromosome 0 with origins in a non-tomato plant genome.

#### Homopolymer correction

Sequences produced by 454 pyrosequencing technology are prone to homopolymer errors (Huse et al. 2007; Gilles et al. 2011). We used the Snippy haploid variant calling tool (https://github.com/tseemann/snippy) to iteratively correct insertion and deletion errors caused by runs of three or more homopolymers. The Illumina short reads provided by Syngenta were used as evidence.

#### Bionano Optical Maps

Genome maps were generated for *Solanum lycopersicum* Heinz var 1706 by purifying high-molecular weight DNA by flow cytometry embedded in agarose plugs, nicked and labelled and finally loaded in the Irys system. The DNA backbone was stained using YOYO-1 and locations of fluorescent labels at restriction sites for Nb.BbvC1 and Nt.BspQI along each molecule were detected by image software in the Irys system. In order to uniquely detect molecules in the Irys system an average label density around 12 labels/100 kb is recommended. *In silico* prediction of the label density based on SL2.50 assembly indicated that this optimal label density would be achieved using a dual nicking strategy combining Nb.BbvC1 and Nt.BspQI nicking endonucleases. After the genome maps were generated, we compared the label frequency to the nicking density for each enzyme in regions for which sequence was known. Surprisingly, Nb.BbvC1 labelled less than 1% of the expected sites, probably due to suboptimal conditions. The lack of activity from Nb.BbvC1 prompted us to perform subsequent analyses assuming a single nicking/labelling process using Nt.BspQI only.

Labeled molecules were loaded into the Irys v2 chips for linearization in nanochannel arrays and subsequently visualized. Distinct genetic features intractable to sequencing technologies, such as long arrays of tandem repeats were observed in the raw single molecules. Many of the mismatches with the reference are a result of deep coverage and a mix of alleles from different haploid sequences in the consensus *de novo* assembly. Single molecules were *de novo* assembled into consensus genome maps (cmaps) in IrysView 2.3 using molecules longer than 150kb.

We aligned the *de novo* optical map based assembly to the BAC-integrated assembly in IrysView. 629 cmaps (93.3%) were mapped to the BAC-integrated assembly which covered 92.8% of the optical map assembly and 85.8% of the BAC-integrated assembly. The unaligned (45) cmaps could correspond to the approximately 10% regions in the tomato genome that still remain to be assembled. For structural variation detection in IrysView, alignments between cmaps were obtained using a dynamic programming approach where the scoring function was the likelihood of a pair of intervals being similar. Likelihood is calculated based on a noise model which takes into account fixed sizing error, misaligned sites and optical resolution. An interval whose cumulative likelihood is worse than 0.01% is classified as an outlier region. If such regions occur between highly scoring regions, an insertion or deletion call is made in the outlier region, depending on the relative size of the region on the query and the reference maps.

We identified a number of inconsistencies between the optical map and the genome sequence. For example, the Bionano cmap 15 aligned to six scaffolds in chromosome 12, but where two of the scaffolds were in opposite orientation. On further investigation, we found that there was sufficient coverage of Bionano molecules over the region to support the change in orientation for SL2.50sc04039 and SL2.50sc04057. Therefore, we changed the order and/or orientation for 157 components in SL2.50sc04039 and 808 components in SL2.50sc04057 on chromosome 12. Another example is scaffold SL2.50sc03721 in chromosome 3 had a different orientation. Bionano molecules did not have enough molecules to support this change. Although, in SL4.0 the order and orientation was corrected on scaffold SL2.50sc03721 due to Hi-C evidence. A chromosome 0 contig (NW_004194387.1, 561,203bp) was integrated in 1.4 Mb scaffold gap on chromosome 2 using evidence from Bionano cmaps. Similarly, a contig (NW_004194391, 203,142bp) from chromosome 0 was integrated in a 1.5 Mb scaffold gap on chromosome 9. Bionano evidence was used for resizing nineteen inter scaffold and inter contig gaps from chromosome 1 through chromosome 12.

### SL4.0 reference genome

#### Pacbio data generation and *de novo* assembly

We generated long reads cumulating to 80X coverage of the Heinz 1706 genome from PacBio RS II and Sequel instruments. The Canu v1.5 assembler (Phillippy 2016) was used to correct, trim and assemble the raw PacBio reads with the following parameters: corOutCoverage=40, corMinCoverage=4, minOverlapLength=1000, minReadLength=1000, ovlMerDistinct=0.99, rawErrorRate=0.30, canuIterationMax=1, batThreads=16, cnsMemory=59, minMemory=16G, ovsMemory=20G. Raw assembly with corrected reads resulted in 504 contigs with a total length of 785 Mbp with a contig N50 of 5.5 Mbp. The minimum length was 12,753 bp, maximum length was 28,741,570 bp and the median length was 96,901 bp. The contigs were error corrected with 63X raw PacBio reads iteratively with the Arrow pipeline (https://github.com/PacificBiosciences/GenomicConsensus) for two rounds. The PacBio error corrected contigs were then further error corrected with accurate Illumina short reads with 100X coverage of the genome using Pilon (Walker et al. 2014). Pilon was run iteratively for two rounds with the following parameters: --threads 32 --changes --fix bases --diploid --mindepth 0.6. We used the minimum depth of coverage required to correct a base as 0.6 to capture the major allele at that locus in case it was heterozygous for this diploid genome.

#### Hi-C scaffolding

Young seedlings were sent to Dovetail Genomics (USA) who generated a Hi-C library and sequenced it with Illumina. PBjelly (English et al. 2012) was first used to scaffold the contigs with PacBio reads. The HiRise assembler (Dovetail, Inc) was then used to join the scaffolds into super-scaffolds using connections determined from the Hi-C data. The 12 super-scaffolds were oriented and numbered according to corresponding 12 chromosomes in the SL3.0 build. The remaining contigs (12.5 Mbp and 202 contigs) which were not scaffolded in the 12 chromosomes were screened for chloroplast, mitochondrial and duplicated sequences. The final set of 152 unassembled contigs were stitched together to create chromosome 00 (9.6 Mbp) with a 100 bp gap inserted between two contigs.

### ITAG3.2 annotation

Known 2,246 Transposable elements (TEs) identified in the ITAG2.40 by Jouffroy et al (Jouffroy et al. 2016) were removed before mapping ITAG2.40 models to SL3.0. Along with TEs, genomic contamination of Arabidopsis and Chloroplast regions (56 genes) were also removed. We also incorporated user curated structural annotations from tomato Apollo genome annotation editor hosted at solgenomics.net. Mapping of remaining 32,425 ITAG2.40 cDNAs on SL3.0 was performed using GMAP tool (Wu and Watanabe 2005). Best aligning path with more that 90% identity and 90% coverage was retained. Total of 31,309 genes were mapped on repeat masked SL3.0 genome assembly. 1,116 genes did not get mapped on to repeat masked SL3.0 genome assembly. The procedure for update of legacy (ITAG2.4) gene models and identification of novel genes is similar to the one followed for ITAG4.0 and has been discussed in detail below.

### Repeat Identification and classification

To maximize gene predictions on the reference genome, repeat elements were identified and classified based on RepeatModeler and RepeatMasker based method. Briefly, repeat library was constructed by RepeatModeler (Smit, AFA, Hubley, R & Green, P. 2013-2015) and repeats with similarity with curated SwissProt plant proteins were filtered out (Campbell, Law, et al. 2014). This custom library masked 64.19% of the SL4.0 reference genome. To test the sensitivity of repeat libraries generated using RepeatModeler and REPET (Flutre et al. 2011), we mapped RNA-Seq data to repeat masked genomes (data not shown). Overall, the mapping rate was slightly better for RepeatModeler generated repeat library along with better mapping of ITAG2.40 genes. For repeat analysis we provide REPET based classification which provides comprehensive annotation of repeats useful for repeat studies. And for identification of protein-coding genes RepeatModeler generated library was used to mask SL4.0 reference genome.

### Iso-Seq sequencing and data analysis

Breaker and mature green stages of tomato fruit were harvested and RNA was extracted for sequencing with PacBio SMRT technology. High quality full-length transcripts obtained from SMRT analysis pipeline were mapped to SL4.0 genome assembly using GMAP (Wu and Watanabe 2005) and collapsed redundant isoforms using Cupcake ToFU pipeline. Genome mapping based clustering of full-length transcripts resulted in 17,112 genes and 110,577 isoforms.

### RNA-Seq expression data

Publicly available RNA-Seq data was obtained from NCBI SRA using SRA tools (version 2.8.2-1). RNA-Seq libraries were filtered for transgenic or mutant line expression data. Data sets with less than 80% mapping rate were not used. Other RNA-Seq data sources include RenSeq enriched for NB-LRR (Andolfo et al. 2014) and untranslated region (UTR) enriched expression data (Zhong et al. 2013; Tzfadia et al. 2018). All the RNA-Seq data sets were mapped on the repeat masked SL4.0 genome and a transcriptome was assembled. In total, we mapped more than 12 billion RNA-Seq reads to generate the genome guided transcriptome with a mapping rate of approximately 85%. The mapping was performed using HISAT2 (Kim, Langmead, and Salzberg 2015) and the transcriptome was generated using StringTie (Pertea et al. 2015). Single exon unstranded transcripts were removed to reduce transcript mapping to repetitive elements and pseudogenes. The final transcriptome contained 75,053 transcripts with 4.4 exons per gene and an average transcript length of 1,995 bp compared to 1209 bp in ITAG2.40.

### Transcriptome assembly

Iso-Seq and RNA-Seq based transcriptomes were combined and picked transcripts using Mikado (Venturini et al. 2018). Above described Iso-Seq transcriptome and StringTie transcriptomes were provided to Mikado along with reliable splice junctions obtained by Portcullis (Mapleson et al. 2018). Blast alignments to SwissProt plant proteins (“SwissProt Database” 2008) and Pfam domains (El-Gebali et al. 2019) identified using hmmscan (Eddy 1998) were also provided to Mikado pipeline. Based on homology and identification of protein domains, transcripts were classified as protein-coding and non-coding genes. Overall, Mikado generated 32,998 protein-coding genes and 41,708 transcripts. Protein-coding transcripts identified based on expression data through Mikado were integrated in MAKER annotation pipeline.

### ITAG4.0 protein-coding gene annotation

Gene identification was performed on the SL4.0 genome after masking repeats. Annotation pipeline MAKER (version 3.1) (Campbell, Holt, et al. 2014) was employed iteratively to improve legacy gene models and to predict novel gene models. For protein evidence, curated proteins from SwissProt were provided to MAKER annotation pipeline. For transcript evidence, Mikado generated transcriptome was provided to MAKER which comprised of Iso-Seq data and publically available RNA-Seq data. ITAG2.40 transcripts were mapped to the new SL4.0 reference genome assembly using GMAP (version: gmap-2017-06-20) (Wu and Watanabe 2005). Mapped genes models were updated (parameter: pred_gff) using available expression data including Iso-Seq and RNA-Seq based transcriptome before models from other gene prediction methods were added.

### Novel gene identification

Identification of novel genes in the new reference genome was performed with the help of gene prediction algorithms trained on all available RNA-Seq data sets. Augustus training was done with BRAKER1 pipeline (Hoff et al. 2019) with support from mapped RNA-Seq reads. SNAP gene predictor was trained within MAKER using maker documentation. Both the gene prediction methods were used to find novel genes. Updated ITAG gene models were used as passthrough (parameter: model_gff) during the annotation process. The newly identified gene models were screened for potential repeats. Gene Ontology enrichment for novel genes was performed based on R/Bioconductor package topGO. Overrepresented GO-terms were calculated using Fisher’s exact test.

### Functional annotation

Functional annotation has been done with Araport11, SwissProt and TrEMBL plant databases using BLAST (Altschul et al. 1990). Blast results were input to Automated Assignment of Human Readable Descriptions (AHRD) (https://github.com/groupschoof/AHRD) and get functional descriptions. We also run InterproScan (Zdobnov and Apweiler 2001) and BLAST (Altschul et al. 1990) to get GO terms (G. O. Consortium and Gene Ontology Consortium 2006), Pfam domains (El-Gebali et al. 2019). AHRD results together with Interproscan results were used to assign functional descriptions to ITAG4.0.

### Long non-coding RNA prediction

Prediction of long non-coding RNA (lncRNA) was based on transcript generated using Mikado software (Venturini et al. 2018). For Mikado, an extensive expression data was used including full-length Iso-Seq data. Mikado resulted in 153K non-coding genes. To identify lncRNA within 153K transcripts, FEELnc pipeline was used with default parameters and further filtered based on genomic overlap with ITAG4.0 protein-coding genes.

### Gene family analysis

*Solanum lyocpersicum* gene family analysis has been performed with Orthofinder (Emms and Kelly, n.d.), Interproscan (Zdobnov and Apweiler 2001), KinFin (Laetsch and Blaxter 2017) and CAFE software (De Bie et al. 2006). Orthologous protein family analysis has been done using *Oryza Sativa, Arabidopsis thaliana, Vitis vinifera, Ipomoea Nil, Petunia axilliaris, Petunia inflata, Capsicum chinense, Capsicum annuum, Solanum tuberosum and Solanum pennellii* proteins downloaded from Phytozome (https://phytozome.jgi.doe.gov/).

