## Supplementary Figures for "An improved de novo assembly and annotation of the tomato reference genome using single-molecule sequencing, Hi-C proximity ligation and optical maps"

**Supplementary Table 4:** List of ITAG4.0 novel genes

| **Solyc ID** |
|---|
| Solyc00g500001.1.1 |
| Solyc00g500002.1.1 |
| Solyc00g500003.1.1 |
| Solyc00g500004.1.1 |
| Solyc00g500005.1.1 |
| Solyc00g500006.1.1 |
| Solyc00g500007.1.1 |
| Solyc00g500008.1.1 |
| Solyc00g500009.1.1 |
| Solyc00g500010.1.1 |
| Solyc00g500011.1.1 |
| Solyc00g500012.1.1 |
| Solyc00g500013.1.1 |
| Solyc00g500014.1.1 |
| Solyc00g500015.1.1 |
| Solyc00g500016.1.1 |
| Solyc00g500017.1.1 |
| Solyc00g500018.1.1 |
| Solyc00g500019.1.1 |
| Solyc00g500020.1.1 |
| Solyc00g500021.1.1 |
| Solyc00g500022.1.1 |
| Solyc00g500023.1.1 |
| Solyc00g500024.1.1 |
| Solyc00g500025.1.1 |
| Solyc00g500026.1.1 |
| Solyc00g500027.1.1 |
| Solyc00g500028.1.1 |
| Solyc00g500029.1.1 |
| Solyc00g500030.1.1 |
| Solyc00g500031.1.1 |
| Solyc00g500032.1.1 |
| Solyc00g500033.1.1 |
| Solyc00g500034.1.1 |
| Solyc00g500035.1.1 |
| Solyc00g500036.1.1 |
| Solyc00g500037.1.1 |
| Solyc00g500038.1.1 |
| Solyc00g500039.1.1 |
| Solyc00g500040.1.1 |
| Solyc00g500041.1.1 |

| |
|---|
| Solyc00g500042.1.1 |
| Solyc00g500043.1.1 |
| Solyc00g500044.1.1 |
| Solyc00g500045.1.1 |
| Solyc00g500046.1.1 |
| Solyc00g500047.1.1 |
| Solyc00g500048.1.1 |
| Solyc00g500049.1.1 |
| Solyc00g500050.1.1 |
| Solyc00g500051.1.1 |
| Solyc00g500052.1.1 |
| Solyc00g500053.1.1 |
| Solyc00g500054.1.1 |
| Solyc00g500055.1.1 |
| Solyc00g500056.1.1 |
| Solyc00g500057.1.1 |
| Solyc00g500058.1.1 |
| Solyc00g500059.1.1 |
| Solyc00g500060.1.1 |
| Solyc00g500061.1.1 |
| Solyc00g500062.1.1 |
| Solyc00g500063.1.1 |
| Solyc00g500064.1.1 |
| Solyc00g500065.1.1 |
| Solyc00g500066.1.1 |
| Solyc00g500067.1.1 |
| Solyc00g500068.1.1 |
| Solyc00g500069.1.1 |
| Solyc00g500070.1.1 |
| Solyc00g500071.1.1 |
| Solyc00g500072.1.1 |
| Solyc00g500073.1.1 |
| Solyc00g500074.1.1 |
| Solyc00g500075.1.1 |
| Solyc00g500076.1.1 |
| Solyc00g500077.1.1 |
| Solyc00g500078.1.1 |
| Solyc00g500079.1.1 |
| Solyc00g500080.1.1 |
| Solyc00g500081.1.1 |
| Solyc00g500082.1.1 |
| Solyc00g500083.1.1 |
| Solyc00g500084.1.1 |

| |
|---|
| Solyc00g500085.1.1 |
| Solyc00g500086.1.1 |
| Solyc00g500087.1.1 |
| Solyc00g500088.1.1 |
| Solyc00g500089.1.1 |
| Solyc00g500090.1.1 |
| Solyc00g500091.1.1 |
| Solyc00g500092.1.1 |
| Solyc00g500093.1.1 |
| Solyc00g500094.1.1 |
| Solyc00g500095.1.1 |
| Solyc00g500096.1.1 |
| Solyc00g500097.1.1 |
| Solyc00g500098.1.1 |
| Solyc00g500099.1.1 |
| Solyc00g500100.1.1 |
| Solyc00g500101.1.1 |
| Solyc00g500102.1.1 |
| Solyc00g500103.1.1 |
| Solyc00g500104.1.1 |
| Solyc00g500105.1.1 |
| Solyc00g500106.1.1 |
| Solyc00g500107.1.1 |
| Solyc00g500108.1.1 |
| Solyc00g500109.1.1 |
| Solyc00g500110.1.1 |
| Solyc00g500111.1.1 |
| Solyc00g500112.1.1 |
| Solyc00g500113.1.1 |
| Solyc00g500114.1.1 |
| Solyc00g500115.1.1 |
| Solyc00g500116.1.1 |
| Solyc00g500117.1.1 |
| Solyc00g500118.1.1 |
| Solyc00g500119.1.1 |
| Solyc00g500120.1.1 |
| Solyc00g500121.1.1 |
| Solyc00g500122.1.1 |
| Solyc00g500123.1.1 |
| Solyc00g500124.1.1 |
| Solyc00g500125.1.1 |
| Solyc00g500126.1.1 |
| Solyc00g500127.1.1 |

| |
|---|
| Solyc00g500128.1.1 |
| Solyc00g500129.1.1 |
| Solyc00g500130.1.1 |
| Solyc00g500131.1.1 |
| Solyc00g500132.1.1 |
| Solyc00g500133.1.1 |
| Solyc00g500134.1.1 |
| Solyc00g500135.1.1 |
| Solyc00g500136.1.1 |
| Solyc00g500137.1.1 |
| Solyc00g500138.1.1 |
| Solyc00g500139.1.1 |
| Solyc00g500140.1.1 |
| Solyc00g500141.1.1 |
| Solyc00g500142.1.1 |
| Solyc00g500143.1.1 |
| Solyc00g500144.1.1 |
| Solyc00g500145.1.1 |
| Solyc00g500146.1.1 |
| Solyc00g500147.1.1 |
| Solyc00g500148.1.1 |
| Solyc00g500149.1.1 |
| Solyc00g500150.1.1 |
| Solyc00g500151.1.1 |
| Solyc00g500152.1.1 |
| Solyc00g500153.1.1 |
| Solyc00g500154.1.1 |
| Solyc00g500155.1.1 |
| Solyc00g500156.1.1 |
| Solyc00g500157.1.1 |
| Solyc00g500158.1.1 |
| Solyc00g500159.1.1 |
| Solyc00g500160.1.1 |
| Solyc00g500161.1.1 |
| Solyc00g500162.1.1 |
| Solyc00g500163.1.1 |
| Solyc00g500164.1.1 |
| Solyc00g500165.1.1 |
| Solyc00g500166.1.1 |
| Solyc00g500167.1.1 |
| Solyc00g500168.1.1 |
| Solyc00g500169.1.1 |
| Solyc00g500170.1.1 |

| |
|---|
| Solyc00g500171.1.1 |
| Solyc00g500172.1.1 |
| Solyc00g500173.1.1 |
| Solyc00g500174.1.1 |
| Solyc00g500175.1.1 |
| Solyc00g500176.1.1 |
| Solyc00g500177.1.1 |
| Solyc00g500178.1.1 |
| Solyc00g500179.1.1 |
| Solyc00g500180.1.1 |
| Solyc00g500181.1.1 |
| Solyc00g500182.1.1 |
| Solyc00g500183.1.1 |
| Solyc00g500184.1.1 |
| Solyc00g500185.1.1 |
| Solyc00g500186.1.1 |
| Solyc00g500187.1.1 |
| Solyc00g500188.1.1 |
| Solyc00g500189.1.1 |
| Solyc00g500190.1.1 |
| Solyc00g500191.1.1 |
| Solyc00g500192.1.1 |
| Solyc00g500193.1.1 |
| Solyc00g500194.1.1 |
| Solyc00g500195.1.1 |
| Solyc00g500196.1.1 |
| Solyc00g500197.1.1 |
| Solyc00g500198.1.1 |
| Solyc00g500199.1.1 |
| Solyc00g500200.1.1 |
| Solyc00g500201.1.1 |
| Solyc00g500202.1.1 |
| Solyc00g500203.1.1 |
| Solyc00g500204.1.1 |
| Solyc00g500205.1.1 |
| Solyc00g500206.1.1 |
| Solyc00g500207.1.1 |
| Solyc00g500208.1.1 |
| Solyc00g500209.1.1 |
| Solyc00g500210.1.1 |
| Solyc00g500211.1.1 |
| Solyc00g500212.1.1 |
| Solyc00g500213.1.1 |

| |
|---|
| Solyc00g500214.1.1 |
| Solyc00g500215.1.1 |
| Solyc00g500216.1.1 |
| Solyc00g500217.1.1 |
| Solyc00g500218.1.1 |
| Solyc00g500219.1.1 |
| Solyc00g500220.1.1 |
| Solyc00g500221.1.1 |
| Solyc00g500222.1.1 |
| Solyc00g500223.1.1 |
| Solyc00g500224.1.1 |
| Solyc00g500225.1.1 |
| Solyc00g500226.1.1 |
| Solyc00g500227.1.1 |
| Solyc00g500228.1.1 |
| Solyc00g500229.1.1 |
| Solyc00g500230.1.1 |
| Solyc00g500231.1.1 |
| Solyc00g500232.1.1 |
| Solyc00g500233.1.1 |
| Solyc00g500234.1.1 |
| Solyc00g500235.1.1 |
| Solyc00g500236.1.1 |
| Solyc00g500237.1.1 |
| Solyc00g500238.1.1 |
| Solyc00g500239.1.1 |
| Solyc00g500240.1.1 |
| Solyc00g500241.1.1 |
| Solyc00g500242.1.1 |
| Solyc00g500243.1.1 |
| Solyc00g500244.1.1 |
| Solyc00g500245.1.1 |
| Solyc00g500246.1.1 |
| Solyc00g500247.1.1 |
| Solyc00g500248.1.1 |
| Solyc00g500249.1.1 |
| Solyc00g500250.1.1 |
| Solyc00g500251.1.1 |
| Solyc00g500252.1.1 |
| Solyc00g500253.1.1 |
| Solyc00g500254.1.1 |
| Solyc00g500255.1.1 |
| Solyc00g500256.1.1 |

| |
|---|
| Solyc00g500257.1.1 |
| Solyc00g500258.1.1 |
| Solyc00g500259.1.1 |
| Solyc00g500260.1.1 |
| Solyc00g500261.1.1 |
| Solyc00g500262.1.1 |
| Solyc00g500263.1.1 |
| Solyc00g500264.1.1 |
| Solyc00g500265.1.1 |
| Solyc00g500266.1.1 |
| Solyc00g500267.1.1 |
| Solyc00g500268.1.1 |
| Solyc00g500269.1.1 |
| Solyc00g500270.1.1 |
| Solyc00g500271.1.1 |
| Solyc00g500272.1.1 |
| Solyc00g500273.1.1 |
| Solyc00g500274.1.1 |
| Solyc00g500275.1.1 |
| Solyc00g500276.1.1 |
| Solyc00g500277.1.1 |
| Solyc00g500278.1.1 |
| Solyc00g500279.1.1 |
| Solyc00g500280.1.1 |
| Solyc00g500281.1.1 |
| Solyc00g500282.1.1 |
| Solyc00g500283.1.1 |
| Solyc00g500284.1.1 |
| Solyc00g500285.1.1 |
| Solyc00g500286.1.1 |
| Solyc00g500287.1.1 |
| Solyc00g500288.1.1 |
| Solyc00g500289.1.1 |
| Solyc00g500290.1.1 |
| Solyc00g500291.1.1 |
| Solyc00g500292.1.1 |
| Solyc00g500293.1.1 |
| Solyc00g500294.1.1 |
| Solyc00g500295.1.1 |
| Solyc00g500296.1.1 |
| Solyc00g500297.1.1 |
| Solyc00g500298.1.1 |
| Solyc00g500299.1.1 |

| |
|---|
| Solyc00g500300.1.1 |
| Solyc00g500301.1.1 |
| Solyc00g500302.1.1 |
| Solyc00g500303.1.1 |
| Solyc00g500304.1.1 |
| Solyc00g500305.1.1 |
| Solyc00g500306.1.1 |
| Solyc00g500307.1.1 |
| Solyc00g500308.1.1 |
| Solyc00g500309.1.1 |
| Solyc00g500310.1.1 |
| Solyc00g500311.1.1 |
| Solyc00g500312.1.1 |
| Solyc00g500313.1.1 |
| Solyc00g500314.1.1 |
| Solyc00g500315.1.1 |
| Solyc00g500316.1.1 |
| Solyc00g500317.1.1 |
| Solyc00g500318.1.1 |
| Solyc00g500319.1.1 |
| Solyc00g500320.1.1 |
| Solyc00g500321.1.1 |
| Solyc00g500322.1.1 |
| Solyc00g500323.1.1 |
| Solyc00g500324.1.1 |
| Solyc00g500325.1.1 |
| Solyc00g500326.1.1 |
| Solyc00g500327.1.1 |
| Solyc00g500328.1.1 |
| Solyc00g500329.1.1 |
| Solyc00g500330.1.1 |
| Solyc00g500331.1.1 |
| Solyc00g500332.1.1 |
| Solyc00g500333.1.1 |
| Solyc00g500334.1.1 |
| Solyc00g500335.1.1 |
| Solyc00g500336.1.1 |
| Solyc00g500337.1.1 |
| Solyc00g500338.1.1 |
| Solyc00g500339.1.1 |
| Solyc00g500340.1.1 |
| Solyc00g500341.1.1 |
| Solyc00g500342.1.1 |

| |
|---|
| Solyc00g500343.1.1 |
| Solyc00g500344.1.1 |
| Solyc00g500345.1.1 |
| Solyc00g500346.1.1 |
| Solyc00g500347.1.1 |
| Solyc00g500348.1.1 |
| Solyc00g500349.1.1 |
| Solyc00g500350.1.1 |
| Solyc00g500351.1.1 |
| Solyc00g500352.1.1 |
| Solyc00g500353.1.1 |
| Solyc00g500354.1.1 |
| Solyc00g500355.1.1 |
| Solyc00g500356.1.1 |
| Solyc00g500357.1.1 |
| Solyc00g500358.1.1 |
| Solyc00g500359.1.1 |
| Solyc00g500360.1.1 |
| Solyc00g500361.1.1 |
| Solyc00g500362.1.1 |
| Solyc00g500363.1.1 |
| Solyc00g500364.1.1 |
| Solyc00g500365.1.1 |
| Solyc00g500366.1.1 |
| Solyc00g500367.1.1 |
| Solyc00g500368.1.1 |
| Solyc00g500369.1.1 |
| Solyc00g500370.1.1 |
| Solyc00g500371.1.1 |
| Solyc00g500372.1.1 |
| Solyc00g500373.1.1 |
| Solyc00g500374.1.1 |
| Solyc00g500375.1.1 |
| Solyc00g500376.1.1 |
| Solyc00g500377.1.1 |
| Solyc00g500378.1.1 |
| Solyc00g500379.1.1 |
| Solyc00g500380.1.1 |
| Solyc00g500381.1.1 |
| Solyc00g500382.1.1 |
| Solyc00g500383.1.1 |
| Solyc00g500384.1.1 |
| Solyc00g500385.1.1 |

| |
|---|
| Solyc00g500386.1.1 |
| Solyc00g500387.1.1 |
| Solyc00g500388.1.1 |
| Solyc00g500389.1.1 |
| Solyc00g500390.1.1 |
| Solyc00g500391.1.1 |
| Solyc00g500392.1.1 |
| Solyc00g500393.1.1 |
| Solyc00g500394.1.1 |
| Solyc00g500395.1.1 |
| Solyc00g500396.1.1 |
| Solyc00g500397.1.1 |
| Solyc00g500398.1.1 |
| Solyc00g500399.1.1 |
| Solyc00g500400.1.1 |
| Solyc00g500401.1.1 |
| Solyc00g500402.1.1 |
| Solyc00g500403.1.1 |
| Solyc00g500404.1.1 |
| Solyc00g500405.1.1 |
| Solyc00g500406.1.1 |
| Solyc00g500407.1.1 |
| Solyc00g500408.1.1 |
| Solyc00g500409.1.1 |
| Solyc01g004000.1.1 |
| Solyc01g004002.1.1 |
| Solyc01g004004.1.1 |
| Solyc01g004006.1.1 |
| Solyc01g004008.1.1 |
| Solyc01g005253.1.1 |
| Solyc01g005257.1.1 |
| Solyc01g005783.1.1 |
| Solyc01g005787.1.1 |
| Solyc01g005985.1.1 |
| Solyc01g006055.1.1 |
| Solyc01g006555.1.1 |
| Solyc01g006585.1.1 |
| Solyc01g006845.1.1 |
| Solyc01g007263.1.1 |
| Solyc01g007267.1.1 |
| Solyc01g008425.1.1 |
| Solyc01g008473.1.1 |
| Solyc01g008475.1.1 |

| |
|---|
| Solyc01g008477.1.1 |
| Solyc01g008479.1.1 |
| Solyc01g009145.1.1 |
| Solyc01g009165.1.1 |
| Solyc01g009245.1.1 |
| Solyc01g009305.1.1 |
| Solyc01g009443.1.1 |
| Solyc01g009447.1.1 |
| Solyc01g009505.1.1 |
| Solyc01g009565.1.1 |
| Solyc01g009705.1.1 |
| Solyc01g009735.1.1 |
| Solyc01g009795.1.1 |
| Solyc01g009975.1.1 |
| Solyc01g010063.1.1 |
| Solyc01g010065.1.1 |
| Solyc01g010067.1.1 |
| Solyc01g010175.1.1 |
| Solyc01g010185.1.1 |
| Solyc01g010215.1.1 |
| Solyc01g010365.1.1 |
| Solyc01g010405.1.1 |
| Solyc01g010415.1.1 |
| Solyc01g010705.1.1 |
| Solyc01g010755.1.1 |
| Solyc01g010785.1.1 |
| Solyc01g011005.1.1 |
| Solyc01g011105.1.1 |
| Solyc01g011113.1.1 |
| Solyc01g011115.1.1 |
| Solyc01g011117.1.1 |
| Solyc01g011273.1.1 |
| Solyc01g011277.1.1 |
| Solyc01g011352.1.1 |
| Solyc01g011354.1.1 |
| Solyc01g011356.1.1 |
| Solyc01g011358.1.1 |
| Solyc01g011413.1.1 |
| Solyc01g011417.1.1 |
| Solyc01g011435.1.1 |
| Solyc01g011465.1.1 |
| Solyc01g011495.1.1 |
| Solyc01g012585.1.1 |

| |
|---|
| Solyc01g013883.1.1 |
| Solyc01g013887.1.1 |
| Solyc01g013975.1.1 |
| Solyc01g013985.1.1 |
| Solyc01g013995.2.1 |
| Solyc01g014055.1.1 |
| Solyc01g014105.1.1 |
| Solyc01g014275.1.1 |
| Solyc01g014315.1.1 |
| Solyc01g014355.1.1 |
| Solyc01g014363.1.1 |
| Solyc01g014367.1.1 |
| Solyc01g014373.1.1 |
| Solyc01g014377.1.1 |
| Solyc01g014533.1.1 |
| Solyc01g014725.2.1 |
| Solyc01g014745.1.1 |
| Solyc01g014963.1.1 |
| Solyc01g014967.1.1 |
| Solyc01g015035.1.1 |
| Solyc01g015133.1.1 |
| Solyc01g015137.1.1 |
| Solyc01g015185.1.1 |
| Solyc01g015293.1.1 |
| Solyc01g015297.1.1 |
| Solyc01g016305.1.1 |
| Solyc01g016355.1.1 |
| Solyc01g016412.1.1 |
| Solyc01g016414.1.1 |
| Solyc01g016416.1.1 |
| Solyc01g016418.1.1 |
| Solyc01g016545.1.1 |
| Solyc01g016565.1.1 |
| Solyc01g016605.1.1 |
| Solyc01g016665.1.1 |
| Solyc01g016685.1.1 |
| Solyc01g016725.1.1 |
| Solyc01g017085.1.1 |
| Solyc01g017145.1.1 |
| Solyc01g017535.1.1 |
| Solyc01g017565.1.1 |
| Solyc01g017675.1.1 |
| Solyc01g017725.1.1 |

| |
|---|
| Solyc01g017795.1.1 |
| Solyc01g017825.1.1 |
| Solyc01g017873.1.1 |
| Solyc01g017877.1.1 |
| Solyc01g018015.1.1 |
| Solyc01g018045.1.1 |
| Solyc01g020113.1.1 |
| Solyc01g020117.1.1 |
| Solyc01g020245.1.1 |
| Solyc01g020323.1.1 |
| Solyc01g020327.1.1 |
| Solyc01g020343.1.1 |
| Solyc01g020347.1.1 |
| Solyc01g020353.1.1 |
| Solyc01g020357.1.1 |
| Solyc01g020372.1.1 |
| Solyc01g020495.1.1 |
| Solyc01g020521.1.1 |
| Solyc01g020522.1.1 |
| Solyc01g020524.1.1 |
| Solyc01g020526.1.1 |
| Solyc01g020528.1.1 |
| Solyc01g020575.1.1 |
| Solyc01g021685.1.1 |
| Solyc01g021695.1.1 |
| Solyc01g021745.1.1 |
| Solyc01g022745.1.1 |
| Solyc01g022785.1.1 |
| Solyc01g028803.1.1 |
| Solyc01g028807.1.1 |
| Solyc01g028865.1.1 |
| Solyc01g028875.1.1 |
| Solyc01g028965.1.1 |
| Solyc01g028985.1.1 |
| Solyc01g033995.1.1 |
| Solyc01g034075.1.1 |
| Solyc01g034145.1.1 |
| Solyc01g034213.1.1 |
| Solyc01g034217.1.1 |
| Solyc01g038231.1.1 |
| Solyc01g038232.1.1 |
| Solyc01g038233.1.1 |
| Solyc01g038234.1.1 |

| |
|---|
| Solyc01g038236.1.1 |
| Solyc01g038238.1.1 |
| Solyc01g044275.1.1 |
| Solyc01g044365.1.1 |
| Solyc01g044375.1.1 |
| Solyc01g044523.1.1 |
| Solyc01g044527.1.1 |
| Solyc01g044551.1.1 |
| Solyc01g044552.1.1 |
| Solyc01g044554.1.1 |
| Solyc01g044556.1.1 |
| Solyc01g044558.1.1 |
| Solyc01g050045.1.1 |
| Solyc01g055165.1.1 |
| Solyc01g056285.1.1 |
| Solyc01g056575.1.1 |
| Solyc01g056653.1.1 |
| Solyc01g056657.1.1 |
| Solyc01g056665.1.1 |
| Solyc01g056695.1.1 |
| Solyc01g057003.1.1 |
| Solyc01g057007.1.1 |
| Solyc01g057113.1.1 |
| Solyc01g057117.1.1 |
| Solyc01g057133.1.1 |
| Solyc01g057137.1.1 |
| Solyc01g057165.1.1 |
| Solyc01g057185.1.1 |
| Solyc01g057255.1.1 |
| Solyc01g057395.1.1 |
| Solyc01g057435.1.1 |
| Solyc01g057445.1.1 |
| Solyc01g057523.1.1 |
| Solyc01g057527.1.1 |
| Solyc01g057585.1.1 |
| Solyc01g057675.1.1 |
| Solyc01g057703.1.1 |
| Solyc01g057705.1.1 |
| Solyc01g057707.1.1 |
| Solyc01g057785.1.1 |
| Solyc01g057853.1.1 |
| Solyc01g057855.1.1 |
| Solyc01g057857.1.1 |

| |
|---|
| Solyc01g057903.1.1 |
| Solyc01g057907.1.1 |
| Solyc01g058003.1.1 |
| Solyc01g058007.1.1 |
| Solyc01g058175.1.1 |
| Solyc01g058205.1.1 |
| Solyc01g058225.1.1 |
| Solyc01g058245.1.1 |
| Solyc01g058325.1.1 |
| Solyc01g058385.1.1 |
| Solyc01g058535.1.1 |
| Solyc01g058593.1.1 |
| Solyc01g058597.1.1 |
| Solyc01g058665.1.1 |
| Solyc01g058673.1.1 |
| Solyc01g058677.1.1 |
| Solyc01g058685.1.1 |
| Solyc01g058705.1.1 |
| Solyc01g059765.1.1 |
| Solyc01g059885.1.1 |
| Solyc01g059965.1.1 |
| Solyc01g060015.1.1 |
| Solyc01g060053.1.1 |
| Solyc01g060055.1.1 |
| Solyc01g060057.1.1 |
| Solyc01g060183.1.1 |
| Solyc01g060187.1.1 |
| Solyc01g060375.1.1 |
| Solyc01g060395.1.1 |
| Solyc01g065545.1.1 |
| Solyc01g065585.1.1 |
| Solyc01g065625.1.1 |
| Solyc01g065643.1.1 |
| Solyc01g065647.1.1 |
| Solyc01g065725.1.1 |
| Solyc01g065845.1.1 |
| Solyc01g065903.1.1 |
| Solyc01g065907.1.1 |
| Solyc01g065983.1.1 |
| Solyc01g065987.1.1 |
| Solyc01g066055.1.1 |
| Solyc01g066065.1.1 |
| Solyc01g066115.1.1 |

| |
|---|
| Solyc01g066145.1.1 |
| Solyc01g066205.1.1 |
| Solyc01g066235.1.1 |
| Solyc01g066293.1.1 |
| Solyc01g066297.1.1 |
| Solyc01g066345.1.1 |
| Solyc01g066425.1.1 |
| Solyc01g066445.1.1 |
| Solyc01g066455.1.1 |
| Solyc01g066457.1.1 |
| Solyc01g066575.1.1 |
| Solyc01g066612.1.1 |
| Solyc01g066614.1.1 |
| Solyc01g066616.1.1 |
| Solyc01g066618.1.1 |
| Solyc01g066625.1.1 |
| Solyc01g066775.1.1 |
| Solyc01g066895.1.1 |
| Solyc01g066905.1.1 |
| Solyc01g066915.1.1 |
| Solyc01g067005.1.1 |
| Solyc01g067045.1.1 |
| Solyc01g067145.1.1 |
| Solyc01g067173.1.1 |
| Solyc01g067177.1.1 |
| Solyc01g067255.1.1 |
| Solyc01g067295.1.1 |
| Solyc01g067403.1.1 |
| Solyc01g067407.1.1 |
| Solyc01g067443.1.1 |
| Solyc01g067445.1.1 |
| Solyc01g067447.1.1 |
| Solyc01g067755.1.1 |
| Solyc01g067895.1.1 |
| Solyc01g067965.1.1 |
| Solyc01g068045.1.1 |
| Solyc01g068075.1.1 |
| Solyc01g068293.1.1 |
| Solyc01g068297.1.1 |
| Solyc01g068305.1.1 |
| Solyc01g068435.1.1 |
| Solyc01g073905.1.1 |
| Solyc01g073985.1.1 |

| |
|---|
| Solyc01g079115.1.1 |
| Solyc01g079155.1.1 |
| Solyc01g079285.1.1 |
| Solyc01g079535.1.1 |
| Solyc01g079615.1.1 |
| Solyc01g079805.1.1 |
| Solyc01g079855.1.1 |
| Solyc01g080055.1.1 |
| Solyc01g080075.1.1 |
| Solyc01g080785.1.1 |
| Solyc01g081033.1.1 |
| Solyc01g081037.1.1 |
| Solyc01g081125.1.1 |
| Solyc01g081185.1.1 |
| Solyc01g081455.1.1 |
| Solyc01g081582.1.1 |
| Solyc01g081584.1.1 |
| Solyc01g081586.1.1 |
| Solyc01g081588.1.1 |
| Solyc01g081605.1.1 |
| Solyc01g086815.1.1 |
| Solyc01g086975.1.1 |
| Solyc01g087005.1.1 |
| Solyc01g087015.1.1 |
| Solyc01g087075.1.1 |
| Solyc01g087095.2.1 |
| Solyc01g087525.1.1 |
| Solyc01g088013.1.1 |
| Solyc01g088017.1.1 |
| Solyc01g088105.1.1 |
| Solyc01g088275.1.1 |
| Solyc01g088395.1.1 |
| Solyc01g088575.1.1 |
| Solyc01g088635.1.1 |
| Solyc01g089853.1.1 |
| Solyc01g089857.1.1 |
| Solyc01g089915.1.1 |
| Solyc01g090345.1.1 |
| Solyc01g090515.1.1 |
| Solyc01g090523.1.1 |
| Solyc01g090527.1.1 |
| Solyc01g090693.1.1 |
| Solyc01g090697.1.1 |

| |
|---|
| Solyc01g090965.1.1 |
| Solyc01g091865.1.1 |
| Solyc01g093965.2.1 |
| Solyc01g094705.1.1 |
| Solyc01g094723.1.1 |
| Solyc01g094727.1.1 |
| Solyc01g094915.1.1 |
| Solyc01g095295.1.1 |
| Solyc01g095515.1.1 |
| Solyc01g095895.1.1 |
| Solyc01g095905.1.1 |
| Solyc01g096405.1.1 |
| Solyc01g098555.1.1 |
| Solyc01g099035.1.1 |
| Solyc01g099045.1.1 |
| Solyc01g099565.1.1 |
| Solyc01g100103.1.1 |
| Solyc01g100107.1.1 |
| Solyc01g100915.1.1 |
| Solyc01g101015.1.1 |
| Solyc01g101195.1.1 |
| Solyc01g101225.1.1 |
| Solyc01g102895.1.1 |
| Solyc01g103045.1.1 |
| Solyc01g103595.1.1 |
| Solyc01g103873.1.1 |
| Solyc01g103875.1.1 |
| Solyc01g103877.1.1 |
| Solyc01g104443.1.1 |
| Solyc01g104447.1.1 |
| Solyc01g104595.1.1 |
| Solyc01g105035.1.1 |
| Solyc01g105383.1.1 |
| Solyc01g105387.1.1 |
| Solyc01g105475.1.1 |
| Solyc01g105763.1.1 |
| Solyc01g105767.1.1 |
| Solyc01g106055.1.1 |
| Solyc01g106575.1.1 |
| Solyc01g106605.1.1 |
| Solyc01g106953.1.1 |
| Solyc01g106955.1.1 |
| Solyc01g106957.1.1 |

| |
|---|
| Solyc01g106963.1.1 |
| Solyc01g106967.1.1 |
| Solyc01g107475.1.1 |
| Solyc01g107825.1.1 |
| Solyc01g107925.1.1 |
| Solyc01g108235.1.1 |
| Solyc01g108412.1.1 |
| Solyc01g108414.1.1 |
| Solyc01g108416.1.1 |
| Solyc01g108418.1.1 |
| Solyc01g108513.1.1 |
| Solyc01g108517.1.1 |
| Solyc01g108775.1.1 |
| Solyc01g108835.1.1 |
| Solyc01g109125.1.1 |
| Solyc01g109135.1.1 |
| Solyc01g109275.1.1 |
| Solyc01g109615.1.1 |
| Solyc01g109895.1.1 |
| Solyc01g110555.1.1 |
| Solyc01g110647.1.1 |
| Solyc01g110686.1.1 |
| Solyc01g110825.1.1 |
| Solyc01g110903.1.1 |
| Solyc01g110907.1.1 |
| Solyc01g111825.1.1 |
| Solyc01g112025.1.1 |
| Solyc01g112105.1.1 |
| Solyc01g112315.1.1 |
| Solyc01g150100.1.1 |
| Solyc01g150101.1.1 |
| Solyc01g150102.1.1 |
| Solyc01g150103.1.1 |
| Solyc01g150104.1.1 |
| Solyc01g150105.1.1 |
| Solyc01g150106.1.1 |
| Solyc01g150107.1.1 |
| Solyc01g150108.1.1 |
| Solyc01g150109.1.1 |
| Solyc01g150110.1.1 |
| Solyc01g150111.1.1 |
| Solyc01g150112.1.1 |
| Solyc01g150113.1.1 |

| |
|---|
| Solyc01g150114.1.1 |
| Solyc01g150115.1.1 |
| Solyc01g150116.1.1 |
| Solyc01g150117.1.1 |
| Solyc01g150118.1.1 |
| Solyc01g150119.1.1 |
| Solyc01g150120.1.1 |
| Solyc01g150121.1.1 |
| Solyc01g150122.1.1 |
| Solyc01g150123.1.1 |
| Solyc01g150124.1.1 |
| Solyc01g150125.1.1 |
| Solyc01g150126.1.1 |
| Solyc01g150127.1.1 |
| Solyc01g150128.1.1 |
| Solyc01g150129.1.1 |
| Solyc01g150130.1.1 |
| Solyc01g150131.1.1 |
| Solyc01g150132.1.1 |
| Solyc01g150133.1.1 |
| Solyc01g150134.1.1 |
| Solyc01g150135.1.1 |
| Solyc01g150136.1.1 |
| Solyc01g150137.1.1 |
| Solyc01g150138.1.1 |
| Solyc01g150139.1.1 |
| Solyc01g150140.1.1 |
| Solyc01g150141.1.1 |
| Solyc01g150142.1.1 |
| Solyc01g150143.1.1 |
| Solyc01g150144.1.1 |
| Solyc01g150145.1.1 |
| Solyc01g150146.1.1 |
| Solyc01g150147.1.1 |
| Solyc01g150148.1.1 |
| Solyc01g150149.1.1 |
| Solyc01g150150.1.1 |
| Solyc01g150151.1.1 |
| Solyc01g150152.1.1 |
| Solyc01g150153.1.1 |
| Solyc01g150154.1.1 |
| Solyc01g150155.1.1 |
| Solyc01g150156.1.1 |

| |
|---|
| Solyc01g150157.1.1 |
| Solyc01g150158.1.1 |
| Solyc01g150159.1.1 |
| Solyc01g150160.1.1 |
| Solyc01g150161.1.1 |
| Solyc01g150162.1.1 |
| Solyc01g150163.1.1 |
| Solyc01g150164.1.1 |
| Solyc01g150165.1.1 |
| Solyc01g150166.1.1 |
| Solyc01g150167.1.1 |
| Solyc01g150168.1.1 |
| Solyc01g150169.1.1 |
| Solyc01g150170.1.1 |
| Solyc01g150171.1.1 |
| Solyc01g150172.1.1 |
| Solyc01g150173.1.1 |
| Solyc01g150174.1.1 |
| Solyc01g150175.1.1 |
| Solyc01g150176.1.1 |
| Solyc01g150177.1.1 |
| Solyc02g004000.1.1 |
| Solyc02g004003.1.1 |
| Solyc02g004005.1.1 |
| Solyc02g004007.1.1 |
| Solyc02g005003.1.1 |
| Solyc02g005007.1.1 |
| Solyc02g005145.1.1 |
| Solyc02g005305.1.1 |
| Solyc02g005325.1.1 |
| Solyc02g005335.1.1 |
| Solyc02g005525.1.1 |
| Solyc02g005601.1.1 |
| Solyc02g005602.1.1 |
| Solyc02g005603.1.1 |
| Solyc02g005604.1.1 |
| Solyc02g005606.1.1 |
| Solyc02g005607.1.1 |
| Solyc02g005608.1.1 |
| Solyc02g011845.1.1 |
| Solyc02g011995.1.1 |
| Solyc02g012025.1.1 |
| Solyc02g014035.1.1 |

| |
|---|
| Solyc02g014115.1.1 |
| Solyc02g014385.1.1 |
| Solyc02g014525.1.1 |
| Solyc02g014543.1.1 |
| Solyc02g014547.1.1 |
| Solyc02g014773.1.1 |
| Solyc02g014871.1.1 |
| Solyc02g014872.1.1 |
| Solyc02g014874.1.1 |
| Solyc02g014876.1.1 |
| Solyc02g014878.1.1 |
| Solyc02g021067.1.1 |
| Solyc02g021105.1.1 |
| Solyc02g021163.1.1 |
| Solyc02g021167.1.1 |
| Solyc02g021235.1.1 |
| Solyc02g021365.1.1 |
| Solyc02g021685.1.1 |
| Solyc02g021785.1.1 |
| Solyc02g022935.1.1 |
| Solyc02g024045.1.1 |
| Solyc02g030133.1.1 |
| Solyc02g030135.1.1 |
| Solyc02g030225.1.1 |
| Solyc02g030243.1.1 |
| Solyc02g030247.1.1 |
| Solyc02g030313.1.1 |
| Solyc02g030317.1.1 |
| Solyc02g030375.1.1 |
| Solyc02g030461.1.1 |
| Solyc02g030462.1.1 |
| Solyc02g030468.1.1 |
| Solyc02g030523.1.1 |
| Solyc02g030525.1.1 |
| Solyc02g030527.1.1 |
| Solyc02g030565.1.1 |
| Solyc02g030625.1.1 |
| Solyc02g030633.1.1 |
| Solyc02g030637.1.1 |
| Solyc02g031755.1.1 |
| Solyc02g031985.1.1 |
| Solyc02g032035.1.1 |
| Solyc02g032185.1.1 |

| |
|---|
| Solyc02g032235.1.1 |
| Solyc02g032355.1.1 |
| Solyc02g032523.1.1 |
| Solyc02g032527.1.1 |
| Solyc02g032555.1.1 |
| Solyc02g032585.1.1 |
| Solyc02g032663.1.1 |
| Solyc02g032667.1.1 |
| Solyc02g032695.1.1 |
| Solyc02g032845.1.1 |
| Solyc02g033035.1.1 |
| Solyc02g033055.1.1 |
| Solyc02g036185.1.1 |
| Solyc02g036215.1.1 |
| Solyc02g037545.1.1 |
| Solyc02g037555.1.1 |
| Solyc02g038705.1.1 |
| Solyc02g038723.1.1 |
| Solyc02g038727.1.1 |
| Solyc02g038800.1.1 |
| Solyc02g038801.1.1 |
| Solyc02g038802.1.1 |
| Solyc02g038803.1.1 |
| Solyc02g038804.1.1 |
| Solyc02g038805.1.1 |
| Solyc02g038806.1.1 |
| Solyc02g038807.1.1 |
| Solyc02g038808.1.1 |
| Solyc02g038809.1.1 |
| Solyc02g038810.1.1 |
| Solyc02g038811.1.1 |
| Solyc02g038812.1.1 |
| Solyc02g038813.1.1 |
| Solyc02g038814.1.1 |
| Solyc02g038815.1.1 |
| Solyc02g038816.1.1 |
| Solyc02g038817.1.1 |
| Solyc02g038818.1.1 |
| Solyc02g038819.1.1 |
| Solyc02g038820.1.1 |
| Solyc02g038825.1.1 |
| Solyc02g043813.1.1 |
| Solyc02g043817.1.1 |

| |
|---|
| Solyc02g043885.1.1 |
| Solyc02g043965.1.1 |
| Solyc02g044005.1.1 |
| Solyc02g049108.1.1 |
| Solyc02g050195.1.1 |
| Solyc02g050263.1.1 |
| Solyc02g050265.1.1 |
| Solyc02g050267.1.1 |
| Solyc02g055395.1.1 |
| Solyc02g055533.1.1 |
| Solyc02g055537.1.1 |
| Solyc02g060585.1.1 |
| Solyc02g060595.1.1 |
| Solyc02g060605.1.1 |
| Solyc02g061605.1.1 |
| Solyc02g061665.1.1 |
| Solyc02g061825.1.1 |
| Solyc02g062015.1.1 |
| Solyc02g062045.1.1 |
| Solyc02g062195.1.1 |
| Solyc02g062255.1.1 |
| Solyc02g062313.1.1 |
| Solyc02g062317.1.1 |
| Solyc02g062435.1.1 |
| Solyc02g062505.1.1 |
| Solyc02g062535.1.1 |
| Solyc02g062565.1.1 |
| Solyc02g062615.1.1 |
| Solyc02g062725.1.1 |
| Solyc02g062813.1.1 |
| Solyc02g062815.1.1 |
| Solyc02g062817.1.1 |
| Solyc02g062915.1.1 |
| Solyc02g062953.1.1 |
| Solyc02g062957.1.1 |
| Solyc02g063075.1.1 |
| Solyc02g063085.1.1 |
| Solyc02g063133.1.1 |
| Solyc02g063137.1.1 |
| Solyc02g063355.1.1 |
| Solyc02g063373.1.1 |
| Solyc02g063377.1.1 |
| Solyc02g063455.1.1 |

| |
|---|
| Solyc02g063523.1.1 |
| Solyc02g063527.1.1 |
| Solyc02g063543.1.1 |
| Solyc02g063547.1.1 |
| Solyc02g064775.1.1 |
| Solyc02g064945.1.1 |
| Solyc02g065112.1.1 |
| Solyc02g065114.1.1 |
| Solyc02g065116.1.1 |
| Solyc02g065118.1.1 |
| Solyc02g065253.1.1 |
| Solyc02g065257.1.1 |
| Solyc02g065753.1.1 |
| Solyc02g065757.1.1 |
| Solyc02g065765.1.1 |
| Solyc02g066803.1.1 |
| Solyc02g066807.1.1 |
| Solyc02g066853.1.1 |
| Solyc02g066855.1.1 |
| Solyc02g066857.1.1 |
| Solyc02g066905.1.1 |
| Solyc02g066993.1.1 |
| Solyc02g066997.1.1 |
| Solyc02g067015.1.1 |
| Solyc02g067085.1.1 |
| Solyc02g067103.1.1 |
| Solyc02g067107.1.1 |
| Solyc02g067605.1.1 |
| Solyc02g067655.1.1 |
| Solyc02g067905.1.1 |
| Solyc02g067935.1.1 |
| Solyc02g068195.1.1 |
| Solyc02g068295.1.1 |
| Solyc02g068315.1.1 |
| Solyc02g068415.1.1 |
| Solyc02g068485.1.1 |
| Solyc02g068615.1.1 |
| Solyc02g068793.1.1 |
| Solyc02g068797.1.1 |
| Solyc02g069053.1.1 |
| Solyc02g069055.1.1 |
| Solyc02g069057.1.1 |
| Solyc02g069065.1.1 |

| |
|---|
| Solyc02g069175.1.1 |
| Solyc02g069585.1.1 |
| Solyc02g069715.1.1 |
| Solyc02g070385.1.1 |
| Solyc02g070705.1.1 |
| Solyc02g070755.1.1 |
| Solyc02g070875.1.1 |
| Solyc02g071375.1.1 |
| Solyc02g071383.1.1 |
| Solyc02g071387.1.1 |
| Solyc02g071435.1.1 |
| Solyc02g071535.1.1 |
| Solyc02g072105.1.1 |
| Solyc02g072405.1.1 |
| Solyc02g072423.1.1 |
| Solyc02g072427.1.1 |
| Solyc02g076635.1.1 |
| Solyc02g077865.1.1 |
| Solyc02g078285.1.1 |
| Solyc02g078645.1.1 |
| Solyc02g079365.1.1 |
| Solyc02g079415.1.1 |
| Solyc02g079715.1.1 |
| Solyc02g080145.1.1 |
| Solyc02g080275.1.1 |
| Solyc02g080955.1.1 |
| Solyc02g081015.1.1 |
| Solyc02g081083.1.1 |
| Solyc02g081085.1.1 |
| Solyc02g081087.1.1 |
| Solyc02g081345.1.1 |
| Solyc02g082035.1.1 |
| Solyc02g082125.1.1 |
| Solyc02g082315.2.1 |
| Solyc02g082375.1.1 |
| Solyc02g082633.1.1 |
| Solyc02g082635.1.1 |
| Solyc02g082637.1.1 |
| Solyc02g082725.1.1 |
| Solyc02g082733.1.1 |
| Solyc02g082737.1.1 |
| Solyc02g082775.1.1 |
| Solyc02g083005.1.1 |

| |
|---|
| Solyc02g083075.1.1 |
| Solyc02g083285.2.1 |
| Solyc02g083525.1.1 |
| Solyc02g083765.1.1 |
| Solyc02g083785.1.1 |
| Solyc02g083835.1.1 |
| Solyc02g084005.1.1 |
| Solyc02g084031.1.1 |
| Solyc02g084032.1.1 |
| Solyc02g084034.1.1 |
| Solyc02g084036.1.1 |
| Solyc02g084038.1.1 |
| Solyc02g084083.1.1 |
| Solyc02g084085.1.1 |
| Solyc02g084087.1.1 |
| Solyc02g084113.1.1 |
| Solyc02g084173.1.1 |
| Solyc02g084177.1.1 |
| Solyc02g084245.1.1 |
| Solyc02g084303.1.1 |
| Solyc02g084305.1.1 |
| Solyc02g084307.1.1 |
| Solyc02g084605.1.1 |
| Solyc02g085005.1.1 |
| Solyc02g085145.1.1 |
| Solyc02g085165.1.1 |
| Solyc02g085615.1.1 |
| Solyc02g085725.1.1 |
| Solyc02g085933.1.1 |
| Solyc02g085937.1.1 |
| Solyc02g085965.1.1 |
| Solyc02g086215.1.1 |
| Solyc02g086223.1.1 |
| Solyc02g086227.1.1 |
| Solyc02g086415.1.1 |
| Solyc02g086451.1.1 |
| Solyc02g086452.1.1 |
| Solyc02g086454.1.1 |
| Solyc02g086456.1.1 |
| Solyc02g086458.1.1 |
| Solyc02g086565.1.1 |
| Solyc02g087365.1.1 |
| Solyc02g087373.1.1 |

| |
|---|
| Solyc02g087377.1.1 |
| Solyc02g087475.1.1 |
| Solyc02g087485.1.1 |
| Solyc02g087975.1.1 |
| Solyc02g088735.1.1 |
| Solyc02g088905.1.1 |
| Solyc02g088953.1.1 |
| Solyc02g088955.1.1 |
| Solyc02g088957.1.1 |
| Solyc02g089065.1.1 |
| Solyc02g089175.2.1 |
| Solyc02g089295.1.1 |
| Solyc02g089465.1.1 |
| Solyc02g089655.1.1 |
| Solyc02g090045.1.1 |
| Solyc02g090393.1.1 |
| Solyc02g090397.1.1 |
| Solyc02g090545.1.1 |
| Solyc02g090643.1.1 |
| Solyc02g090647.1.1 |
| Solyc02g091193.1.1 |
| Solyc02g091197.1.1 |
| Solyc02g091535.1.1 |
| Solyc02g091665.1.1 |
| Solyc02g091725.1.1 |
| Solyc02g091835.1.1 |
| Solyc02g091985.1.1 |
| Solyc02g092075.1.1 |
| Solyc02g092225.1.1 |
| Solyc02g092375.1.1 |
| Solyc02g092475.1.1 |
| Solyc02g092525.1.1 |
| Solyc02g092783.1.1 |
| Solyc02g092845.1.1 |
| Solyc02g093195.1.1 |
| Solyc02g093285.1.1 |
| Solyc02g093525.1.1 |
| Solyc02g093545.1.1 |
| Solyc02g093775.1.1 |
| Solyc02g094125.1.1 |
| Solyc02g094585.1.1 |
| Solyc02g150100.1.1 |
| Solyc02g150101.1.1 |

| |
|---|
| Solyc02g150102.1.1 |
| Solyc02g150103.1.1 |
| Solyc02g150104.1.1 |
| Solyc02g150105.1.1 |
| Solyc02g150106.1.1 |
| Solyc02g150107.1.1 |
| Solyc02g150108.1.1 |
| Solyc02g150109.1.1 |
| Solyc02g150110.1.1 |
| Solyc02g150111.1.1 |
| Solyc02g150112.1.1 |
| Solyc02g150113.1.1 |
| Solyc02g150114.1.1 |
| Solyc02g150115.1.1 |
| Solyc02g150116.1.1 |
| Solyc02g150117.1.1 |
| Solyc02g150118.1.1 |
| Solyc02g150119.1.1 |
| Solyc02g150120.1.1 |
| Solyc02g150121.1.1 |
| Solyc02g150122.1.1 |
| Solyc02g150123.1.1 |
| Solyc02g150124.1.1 |
| Solyc02g150125.1.1 |
| Solyc02g150126.1.1 |
| Solyc02g150127.1.1 |
| Solyc02g150128.1.1 |
| Solyc02g150129.1.1 |
| Solyc02g150130.1.1 |
| Solyc02g150131.1.1 |
| Solyc02g150132.1.1 |
| Solyc02g150133.1.1 |
| Solyc02g150134.1.1 |
| Solyc02g150135.1.1 |
| Solyc02g150136.1.1 |
| Solyc02g150137.1.1 |
| Solyc02g150138.1.1 |
| Solyc02g150139.1.1 |
| Solyc02g150140.1.1 |
| Solyc02g150141.1.1 |
| Solyc02g150142.1.1 |
| Solyc02g150143.1.1 |
| Solyc02g150144.1.1 |

| |
|---|
| Solyc02g150145.1.1 |
| Solyc02g150146.1.1 |
| Solyc02g150147.1.1 |
| Solyc02g150148.1.1 |
| Solyc02g150149.1.1 |
| Solyc02g150150.1.1 |
| Solyc03g004000.1.1 |
| Solyc03g004003.1.1 |
| Solyc03g004007.1.1 |
| Solyc03g005213.1.1 |
| Solyc03g005217.1.1 |
| Solyc03g005393.1.1 |
| Solyc03g005395.1.1 |
| Solyc03g005397.1.1 |
| Solyc03g005455.1.1 |
| Solyc03g005555.1.1 |
| Solyc03g005943.1.1 |
| Solyc03g005947.1.1 |
| Solyc03g006043.1.1 |
| Solyc03g006047.1.1 |
| Solyc03g006177.2.1 |
| Solyc03g006365.1.1 |
| Solyc03g006452.1.1 |
| Solyc03g006454.1.1 |
| Solyc03g006456.1.1 |
| Solyc03g006458.1.1 |
| Solyc03g006545.1.1 |
| Solyc03g006712.1.1 |
| Solyc03g006714.1.1 |
| Solyc03g006715.2.1 |
| Solyc03g006716.1.1 |
| Solyc03g006718.1.1 |
| Solyc03g006805.1.1 |
| Solyc03g006903.1.1 |
| Solyc03g006907.1.1 |
| Solyc03g006915.1.1 |
| Solyc03g007485.1.1 |
| Solyc03g007775.2.1 |
| Solyc03g008013.1.1 |
| Solyc03g008015.1.1 |
| Solyc03g008017.1.1 |
| Solyc03g013293.1.1 |
| Solyc03g013295.1.1 |

| |
|---|
| Solyc03g013297.1.1 |
| Solyc03g013304.2.1 |
| Solyc03g013305.1.1 |
| Solyc03g013335.1.1 |
| Solyc03g013375.1.1 |
| Solyc03g013615.1.1 |
| Solyc03g025113.1.1 |
| Solyc03g025117.1.1 |
| Solyc03g025125.1.1 |
| Solyc03g025645.1.1 |
| Solyc03g026015.1.1 |
| Solyc03g026115.1.1 |
| Solyc03g026373.1.1 |
| Solyc03g026375.1.1 |
| Solyc03g026411.1.1 |
| Solyc03g026412.1.1 |
| Solyc03g026414.1.1 |
| Solyc03g026416.1.1 |
| Solyc03g026418.1.1 |
| Solyc03g031483.1.1 |
| Solyc03g031487.1.1 |
| Solyc03g031555.1.1 |
| Solyc03g031895.1.1 |
| Solyc03g031955.1.1 |
| Solyc03g032065.1.1 |
| Solyc03g032075.1.1 |
| Solyc03g032215.1.1 |
| Solyc03g032243.1.1 |
| Solyc03g032247.1.1 |
| Solyc03g033295.1.1 |
| Solyc03g033515.1.1 |
| Solyc03g033755.1.1 |
| Solyc03g033795.1.1 |
| Solyc03g034055.1.1 |
| Solyc03g034057.1.1 |
| Solyc03g034155.1.1 |
| Solyc03g034203.1.1 |
| Solyc03g034205.1.1 |
| Solyc03g034207.1.1 |
| Solyc03g034332.1.1 |
| Solyc03g034334.1.1 |
| Solyc03g034336.1.1 |
| Solyc03g034338.1.1 |

| |
|---|
| Solyc03g034392.1.1 |
| Solyc03g034394.1.1 |
| Solyc03g034396.1.1 |
| Solyc03g034398.1.1 |
| Solyc03g034415.1.1 |
| Solyc03g034455.1.1 |
| Solyc03g036477.2.1 |
| Solyc03g043585.1.1 |
| Solyc03g043735.1.1 |
| Solyc03g043895.1.1 |
| Solyc03g044024.1.1 |
| Solyc03g044205.1.1 |
| Solyc03g044233.1.1 |
| Solyc03g044235.1.1 |
| Solyc03g044237.1.1 |
| Solyc03g044355.1.1 |
| Solyc03g044485.1.1 |
| Solyc03g044545.1.1 |
| Solyc03g044625.1.1 |
| Solyc03g044662.1.1 |
| Solyc03g044664.1.1 |
| Solyc03g044666.1.1 |
| Solyc03g044668.1.1 |
| Solyc03g044725.1.1 |
| Solyc03g044742.1.1 |
| Solyc03g044744.1.1 |
| Solyc03g044746.1.1 |
| Solyc03g044748.1.1 |
| Solyc03g044793.1.1 |
| Solyc03g044797.1.1 |
| Solyc03g044815.1.1 |
| Solyc03g044833.1.1 |
| Solyc03g044837.1.1 |
| Solyc03g044915.1.1 |
| Solyc03g044933.1.1 |
| Solyc03g044937.1.1 |
| Solyc03g046475.1.1 |
| Solyc03g046492.1.1 |
| Solyc03g046494.1.1 |
| Solyc03g046496.1.1 |
| Solyc03g046498.1.1 |
| Solyc03g046591.1.1 |
| Solyc03g046592.1.1 |

| |
|---|
| Solyc03g046593.1.1 |
| Solyc03g046595.1.1 |
| Solyc03g051663.1.1 |
| Solyc03g051665.1.1 |
| Solyc03g051667.1.1 |
| Solyc03g051923.1.1 |
| Solyc03g051927.1.1 |
| Solyc03g053085.1.1 |
| Solyc03g053135.1.1 |
| Solyc03g058305.1.1 |
| Solyc03g058343.1.1 |
| Solyc03g058347.1.1 |
| Solyc03g058433.1.1 |
| Solyc03g058437.1.1 |
| Solyc03g058805.1.1 |
| Solyc03g058855.1.1 |
| Solyc03g058915.1.1 |
| Solyc03g059123.1.1 |
| Solyc03g059203.1.1 |
| Solyc03g059205.1.1 |
| Solyc03g059207.1.1 |
| Solyc03g059315.1.1 |
| Solyc03g059393.1.1 |
| Solyc03g059395.1.1 |
| Solyc03g059397.1.1 |
| Solyc03g059425.1.1 |
| Solyc03g061653.1.1 |
| Solyc03g061657.1.1 |
| Solyc03g062665.1.1 |
| Solyc03g062725.1.1 |
| Solyc03g062835.1.1 |
| Solyc03g062895.1.1 |
| Solyc03g063353.1.1 |
| Solyc03g063357.1.1 |
| Solyc03g063505.1.1 |
| Solyc03g063603.1.1 |
| Solyc03g063607.1.1 |
| Solyc03g063653.1.1 |
| Solyc03g063657.1.1 |
| Solyc03g065233.1.1 |
| Solyc03g065235.1.1 |
| Solyc03g065237.1.1 |
| Solyc03g065255.1.1 |

| |
|---|
| Solyc03g065353.1.1 |
| Solyc03g065355.1.1 |
| Solyc03g065357.1.1 |
| Solyc03g070465.1.1 |
| Solyc03g070485.1.1 |
| Solyc03g071535.1.1 |
| Solyc03g071623.1.1 |
| Solyc03g071627.1.1 |
| Solyc03g071735.1.1 |
| Solyc03g071855.1.1 |
| Solyc03g077905.1.1 |
| Solyc03g077985.1.1 |
| Solyc03g078105.1.1 |
| Solyc03g078325.1.1 |
| Solyc03g078355.1.1 |
| Solyc03g078385.1.1 |
| Solyc03g078425.1.1 |
| Solyc03g078505.1.1 |
| Solyc03g078533.1.1 |
| Solyc03g078537.1.1 |
| Solyc03g078623.1.1 |
| Solyc03g078625.1.1 |
| Solyc03g078627.1.1 |
| Solyc03g078635.1.1 |
| Solyc03g078783.1.1 |
| Solyc03g078787.1.1 |
| Solyc03g078815.1.1 |
| Solyc03g079883.1.1 |
| Solyc03g079887.1.1 |
| Solyc03g079915.1.1 |
| Solyc03g080113.1.1 |
| Solyc03g080117.1.1 |
| Solyc03g080175.1.1 |
| Solyc03g080195.1.1 |
| Solyc03g081345.1.1 |
| Solyc03g082375.1.1 |
| Solyc03g082435.1.1 |
| Solyc03g082445.1.1 |
| Solyc03g082515.1.1 |
| Solyc03g082805.1.1 |
| Solyc03g082845.1.1 |
| Solyc03g082945.1.1 |
| Solyc03g083215.1.1 |

| |
|---|
| Solyc03g083275.1.1 |
| Solyc03g083445.1.1 |
| Solyc03g083485.1.1 |
| Solyc03g083725.1.1 |
| Solyc03g083735.1.1 |
| Solyc03g083965.1.1 |
| Solyc03g083991.1.1 |
| Solyc03g083992.1.1 |
| Solyc03g083993.1.1 |
| Solyc03g083994.1.1 |
| Solyc03g083995.1.1 |
| Solyc03g083996.1.1 |
| Solyc03g083998.1.1 |
| Solyc03g091035.1.1 |
| Solyc03g093055.1.1 |
| Solyc03g093075.1.1 |
| Solyc03g093185.1.1 |
| Solyc03g093343.1.1 |
| Solyc03g093347.1.1 |
| Solyc03g093613.1.1 |
| Solyc03g093617.1.1 |
| Solyc03g093645.1.1 |
| Solyc03g093773.1.1 |
| Solyc03g093777.1.1 |
| Solyc03g093935.1.1 |
| Solyc03g094125.1.1 |
| Solyc03g094155.1.1 |
| Solyc03g094173.1.1 |
| Solyc03g094175.1.1 |
| Solyc03g094177.1.1 |
| Solyc03g095375.1.1 |
| Solyc03g095425.1.1 |
| Solyc03g095515.1.1 |
| Solyc03g095533.1.1 |
| Solyc03g095537.1.1 |
| Solyc03g095775.1.1 |
| Solyc03g095785.1.1 |
| Solyc03g095787.1.1 |
| Solyc03g095845.1.1 |
| Solyc03g095903.1.1 |
| Solyc03g095907.1.1 |
| Solyc03g095973.1.1 |
| Solyc03g095977.1.1 |

| |
|---|
| Solyc03g096005.1.1 |
| Solyc03g096007.1.1 |
| Solyc03g096195.1.1 |
| Solyc03g096225.1.1 |
| Solyc03g096303.1.1 |
| Solyc03g096305.1.1 |
| Solyc03g096307.1.1 |
| Solyc03g096355.1.1 |
| Solyc03g096385.1.1 |
| Solyc03g096433.1.1 |
| Solyc03g096437.1.1 |
| Solyc03g096545.1.1 |
| Solyc03g096605.1.1 |
| Solyc03g096705.1.1 |
| Solyc03g097703.1.1 |
| Solyc03g097707.1.1 |
| Solyc03g097855.1.1 |
| Solyc03g098175.1.1 |
| Solyc03g098535.1.1 |
| Solyc03g098545.1.1 |
| Solyc03g098735.1.1 |
| Solyc03g098745.1.1 |
| Solyc03g098795.1.1 |
| Solyc03g110975.1.1 |
| Solyc03g111415.1.1 |
| Solyc03g111705.1.1 |
| Solyc03g111815.1.1 |
| Solyc03g111885.1.1 |
| Solyc03g111953.1.1 |
| Solyc03g111957.1.1 |
| Solyc03g111993.1.1 |
| Solyc03g111997.1.1 |
| Solyc03g112013.1.1 |
| Solyc03g112017.1.1 |
| Solyc03g112425.1.1 |
| Solyc03g112465.1.1 |
| Solyc03g112735.1.1 |
| Solyc03g112995.1.1 |
| Solyc03g113035.1.1 |
| Solyc03g113825.1.1 |
| Solyc03g114033.1.1 |
| Solyc03g114037.1.1 |
| Solyc03g114175.1.1 |

| |
|---|
| Solyc03g114233.1.1 |
| Solyc03g114237.1.1 |
| Solyc03g114705.1.1 |
| Solyc03g114915.1.1 |
| Solyc03g115165.1.1 |
| Solyc03g115345.1.1 |
| Solyc03g115465.1.1 |
| Solyc03g115655.1.1 |
| Solyc03g115665.1.1 |
| Solyc03g115815.1.1 |
| Solyc03g115985.1.1 |
| Solyc03g116173.1.1 |
| Solyc03g116175.1.1 |
| Solyc03g116177.1.1 |
| Solyc03g116625.1.1 |
| Solyc03g116973.1.1 |
| Solyc03g116977.1.1 |
| Solyc03g117255.1.1 |
| Solyc03g117345.1.1 |
| Solyc03g117845.1.1 |
| Solyc03g117955.1.1 |
| Solyc03g118005.1.1 |
| Solyc03g118135.1.1 |
| Solyc03g118255.1.1 |
| Solyc03g118305.1.1 |
| Solyc03g119485.1.1 |
| Solyc03g119723.1.1 |
| Solyc03g119727.1.1 |
| Solyc03g119905.1.1 |
| Solyc03g120085.1.1 |
| Solyc03g120155.1.1 |
| Solyc03g120585.1.1 |
| Solyc03g121365.1.1 |
| Solyc03g121425.1.1 |
| Solyc03g121825.1.1 |
| Solyc03g122375.1.1 |
| Solyc03g124061.1.1 |
| Solyc03g124062.1.1 |
| Solyc03g124064.1.1 |
| Solyc03g124066.1.1 |
| Solyc03g124068.1.1 |
| Solyc03g150100.1.1 |
| Solyc03g150101.1.1 |

| |
|---|
| Solyc03g150102.1.1 |
| Solyc03g150103.1.1 |
| Solyc03g150104.1.1 |
| Solyc03g150105.1.1 |
| Solyc03g150106.1.1 |
| Solyc03g150107.1.1 |
| Solyc03g150108.1.1 |
| Solyc03g150109.1.1 |
| Solyc03g150110.1.1 |
| Solyc03g150111.1.1 |
| Solyc03g150112.1.1 |
| Solyc03g150113.1.1 |
| Solyc03g150114.1.1 |
| Solyc03g150115.1.1 |
| Solyc03g150116.1.1 |
| Solyc03g150117.1.1 |
| Solyc03g150118.1.1 |
| Solyc03g150119.1.1 |
| Solyc03g150120.1.1 |
| Solyc03g150121.1.1 |
| Solyc03g150122.1.1 |
| Solyc03g150123.1.1 |
| Solyc03g150124.1.1 |
| Solyc03g150125.1.1 |
| Solyc03g150126.1.1 |
| Solyc03g150127.1.1 |
| Solyc03g150128.1.1 |
| Solyc03g150129.1.1 |
| Solyc03g150130.1.1 |
| Solyc03g150131.1.1 |
| Solyc03g150132.1.1 |
| Solyc03g150133.1.1 |
| Solyc03g150134.1.1 |
| Solyc03g150135.1.1 |
| Solyc03g150136.1.1 |
| Solyc03g150137.1.1 |
| Solyc03g150138.1.1 |
| Solyc03g150139.1.1 |
| Solyc03g150140.1.1 |
| Solyc03g150141.1.1 |
| Solyc03g150142.1.1 |
| Solyc03g150143.1.1 |
| Solyc03g150144.1.1 |

| |
|---|
| Solyc03g150145.1.1 |
| Solyc03g150146.1.1 |
| Solyc03g150147.1.1 |
| Solyc03g150148.1.1 |
| Solyc03g150149.1.1 |
| Solyc03g150150.1.1 |
| Solyc03g150151.1.1 |
| Solyc03g150152.1.1 |
| Solyc03g150153.1.1 |
| Solyc03g150154.1.1 |
| Solyc03g150155.1.1 |
| Solyc03g150156.1.1 |
| Solyc03g150157.1.1 |
| Solyc03g150158.1.1 |
| Solyc03g150159.1.1 |
| Solyc03g150160.1.1 |
| Solyc03g150161.1.1 |
| Solyc04g004000.1.1 |
| Solyc04g004005.1.1 |
| Solyc04g005015.1.1 |
| Solyc04g005555.1.1 |
| Solyc04g005653.1.1 |
| Solyc04g005657.1.1 |
| Solyc04g006983.1.1 |
| Solyc04g006987.1.1 |
| Solyc04g007015.1.1 |
| Solyc04g007243.1.1 |
| Solyc04g007247.1.1 |
| Solyc04g007295.1.1 |
| Solyc04g007425.1.1 |
| Solyc04g007585.1.1 |
| Solyc04g007765.1.1 |
| Solyc04g007823.1.1 |
| Solyc04g007825.2.1 |
| Solyc04g007855.1.1 |
| Solyc04g007925.1.1 |
| Solyc04g008135.1.1 |
| Solyc04g008205.1.1 |
| Solyc04g008245.1.1 |
| Solyc04g008303.1.1 |
| Solyc04g008307.1.1 |
| Solyc04g008403.1.1 |
| Solyc04g008407.1.1 |

| |
|---|
| Solyc04g009215.1.1 |
| Solyc04g009305.1.1 |
| Solyc04g009453.1.1 |
| Solyc04g009457.1.1 |
| Solyc04g009525.1.1 |
| Solyc04g010045.1.1 |
| Solyc04g010155.2.1 |
| Solyc04g010185.1.1 |
| Solyc04g010285.1.1 |
| Solyc04g011333.1.1 |
| Solyc04g011335.1.1 |
| Solyc04g011337.1.1 |
| Solyc04g011463.1.1 |
| Solyc04g011467.1.1 |
| Solyc04g011615.1.1 |
| Solyc04g011735.1.1 |
| Solyc04g011915.1.1 |
| Solyc04g011965.1.1 |
| Solyc04g012185.1.1 |
| Solyc04g014515.1.1 |
| Solyc04g014578.2.1 |
| Solyc04g014613.1.1 |
| Solyc04g014617.1.1 |
| Solyc04g014645.1.1 |
| Solyc04g014835.1.1 |
| Solyc04g014853.1.1 |
| Solyc04g014855.1.1 |
| Solyc04g014883.1.1 |
| Solyc04g014887.1.1 |
| Solyc04g014995.1.1 |
| Solyc04g015305.1.1 |
| Solyc04g015635.2.1 |
| Solyc04g015705.1.1 |
| Solyc04g015935.1.1 |
| Solyc04g015975.1.1 |
| Solyc04g016085.1.1 |
| Solyc04g016175.2.1 |
| Solyc04g016185.1.1 |
| Solyc04g016335.1.1 |
| Solyc04g016595.1.1 |
| Solyc04g017625.1.1 |
| Solyc04g017665.1.1 |
| Solyc04g017723.1.1 |

| |
|---|
| Solyc04g017873.1.1 |
| Solyc04g017877.1.1 |
| Solyc04g017925.1.1 |
| Solyc04g017945.1.1 |
| Solyc04g018005.1.1 |
| Solyc04g018045.1.1 |
| Solyc04g018055.1.1 |
| Solyc04g018085.1.1 |
| Solyc04g018155.1.1 |
| Solyc04g018255.1.1 |
| Solyc04g019313.1.1 |
| Solyc04g019317.1.1 |
| Solyc04g024435.1.1 |
| Solyc04g024605.1.1 |
| Solyc04g024963.1.1 |
| Solyc04g024965.1.1 |
| Solyc04g024967.1.1 |
| Solyc04g025005.1.1 |
| Solyc04g025125.1.1 |
| Solyc04g025283.1.1 |
| Solyc04g025287.1.1 |
| Solyc04g025375.1.1 |
| Solyc04g025455.1.1 |
| Solyc04g025653.1.1 |
| Solyc04g025657.1.1 |
| Solyc04g025755.1.1 |
| Solyc04g025915.1.1 |
| Solyc04g025933.1.1 |
| Solyc04g025937.1.1 |
| Solyc04g025995.1.1 |
| Solyc04g026005.1.1 |
| Solyc04g026023.2.1 |
| Solyc04g026045.1.1 |
| Solyc04g026155.1.1 |
| Solyc04g028445.1.1 |
| Solyc04g028465.1.1 |
| Solyc04g039625.1.1 |
| Solyc04g039693.1.1 |
| Solyc04g039695.1.1 |
| Solyc04g039697.1.1 |
| Solyc04g039705.1.1 |
| Solyc04g039733.1.1 |
| Solyc04g039737.1.1 |

| |
|---|
| Solyc04g039775.1.1 |
| Solyc04g039845.1.1 |
| Solyc04g040053.1.1 |
| Solyc04g040057.1.1 |
| Solyc04g040115.1.1 |
| Solyc04g040145.1.1 |
| Solyc04g045425.1.1 |
| Solyc04g045525.1.1 |
| Solyc04g045565.1.1 |
| Solyc04g047885.1.1 |
| Solyc04g048905.1.1 |
| Solyc04g048945.1.1 |
| Solyc04g049005.1.1 |
| Solyc04g049075.1.1 |
| Solyc04g049105.1.1 |
| Solyc04g049222.1.1 |
| Solyc04g049228.1.1 |
| Solyc04g049305.1.1 |
| Solyc04g049403.1.1 |
| Solyc04g049405.1.1 |
| Solyc04g049407.1.1 |
| Solyc04g049495.1.1 |
| Solyc04g049625.1.1 |
| Solyc04g049773.1.1 |
| Solyc04g049777.1.1 |
| Solyc04g049795.1.1 |
| Solyc04g049807.1.1 |
| Solyc04g049873.1.1 |
| Solyc04g049875.1.1 |
| Solyc04g049877.1.1 |
| Solyc04g049893.1.1 |
| Solyc04g049897.1.1 |
| Solyc04g049925.1.1 |
| Solyc04g050015.1.1 |
| Solyc04g050175.1.1 |
| Solyc04g050205.1.1 |
| Solyc04g050365.1.1 |
| Solyc04g050403.1.1 |
| Solyc04g050407.1.1 |
| Solyc04g050445.1.1 |
| Solyc04g050495.1.1 |
| Solyc04g050545.1.1 |
| Solyc04g050573.1.1 |

| |
|---|
| Solyc04g050575.1.1 |
| Solyc04g050577.1.1 |
| Solyc04g050622.1.1 |
| Solyc04g050623.1.1 |
| Solyc04g050625.1.1 |
| Solyc04g050627.1.1 |
| Solyc04g050675.1.1 |
| Solyc04g050685.1.1 |
| Solyc04g050745.1.1 |
| Solyc04g050755.1.1 |
| Solyc04g050795.1.1 |
| Solyc04g050822.1.1 |
| Solyc04g050824.1.1 |
| Solyc04g050826.1.1 |
| Solyc04g050828.1.1 |
| Solyc04g050863.1.1 |
| Solyc04g050865.1.1 |
| Solyc04g050867.1.1 |
| Solyc04g050933.1.1 |
| Solyc04g050937.1.1 |
| Solyc04g050955.1.1 |
| Solyc04g051095.1.1 |
| Solyc04g051155.1.1 |
| Solyc04g051183.1.1 |
| Solyc04g051187.1.1 |
| Solyc04g051235.1.1 |
| Solyc04g051241.1.1 |
| Solyc04g051242.1.1 |
| Solyc04g051244.1.1 |
| Solyc04g051246.1.1 |
| Solyc04g051248.1.1 |
| Solyc04g051275.1.1 |
| Solyc04g051293.1.1 |
| Solyc04g051295.1.1 |
| Solyc04g051297.1.1 |
| Solyc04g051427.2.1 |
| Solyc04g051475.1.1 |
| Solyc04g051625.1.1 |
| Solyc04g051765.1.1 |
| Solyc04g051783.1.1 |
| Solyc04g051787.1.1 |
| Solyc04g051885.1.1 |
| Solyc04g052985.1.1 |

| |
|---|
| Solyc04g053033.1.1 |
| Solyc04g053037.1.1 |
| Solyc04g053091.1.1 |
| Solyc04g053094.1.1 |
| Solyc04g053102.1.1 |
| Solyc04g053104.1.1 |
| Solyc04g053106.1.1 |
| Solyc04g053108.1.1 |
| Solyc04g053145.1.1 |
| Solyc04g053147.1.1 |
| Solyc04g054143.1.1 |
| Solyc04g054147.1.1 |
| Solyc04g054153.1.1 |
| Solyc04g054195.1.1 |
| Solyc04g054223.1.1 |
| Solyc04g054227.1.1 |
| Solyc04g054251.1.1 |
| Solyc04g054252.1.1 |
| Solyc04g054254.1.1 |
| Solyc04g054256.1.1 |
| Solyc04g054258.1.1 |
| Solyc04g054521.1.1 |
| Solyc04g054522.1.1 |
| Solyc04g054523.1.1 |
| Solyc04g054524.1.1 |
| Solyc04g054525.1.1 |
| Solyc04g054526.1.1 |
| Solyc04g054527.1.1 |
| Solyc04g054528.1.1 |
| Solyc04g054529.1.1 |
| Solyc04g054533.1.1 |
| Solyc04g054535.1.1 |
| Solyc04g054537.1.1 |
| Solyc04g054682.1.1 |
| Solyc04g054684.1.1 |
| Solyc04g054686.1.1 |
| Solyc04g054688.1.1 |
| Solyc04g054725.1.1 |
| Solyc04g054735.1.1 |
| Solyc04g054855.1.1 |
| Solyc04g054865.1.1 |
| Solyc04g054875.1.1 |
| Solyc04g054895.1.1 |

| |
|---|
| Solyc04g054915.1.1 |
| Solyc04g055225.1.1 |
| Solyc04g055235.1.1 |
| Solyc04g055253.1.1 |
| Solyc04g055255.1.1 |
| Solyc04g055257.1.1 |
| Solyc04g056365.1.1 |
| Solyc04g056425.1.1 |
| Solyc04g056525.1.1 |
| Solyc04g056535.1.1 |
| Solyc04g056713.1.1 |
| Solyc04g056717.1.1 |
| Solyc04g056723.1.1 |
| Solyc04g056727.1.1 |
| Solyc04g056747.1.1 |
| Solyc04g058205.1.1 |
| Solyc04g063245.1.1 |
| Solyc04g063313.1.1 |
| Solyc04g063317.1.1 |
| Solyc04g063415.1.1 |
| Solyc04g064545.1.1 |
| Solyc04g064805.1.1 |
| Solyc04g064952.1.1 |
| Solyc04g064954.1.1 |
| Solyc04g064956.1.1 |
| Solyc04g064958.1.1 |
| Solyc04g070995.1.1 |
| Solyc04g071083.1.1 |
| Solyc04g071087.1.1 |
| Solyc04g071135.1.1 |
| Solyc04g071265.1.1 |
| Solyc04g071295.1.1 |
| Solyc04g071375.1.1 |
| Solyc04g071523.1.1 |
| Solyc04g071675.1.1 |
| Solyc04g071775.1.1 |
| Solyc04g071803.1.1 |
| Solyc04g071805.1.1 |
| Solyc04g072031.1.1 |
| Solyc04g072032.1.1 |
| Solyc04g072033.1.1 |
| Solyc04g072034.1.1 |
| Solyc04g072035.1.1 |

| |
|---|
| Solyc04g072036.1.1 |
| Solyc04g072037.1.1 |
| Solyc04g072038.1.1 |
| Solyc04g072039.1.1 |
| Solyc04g072043.1.1 |
| Solyc04g072045.1.1 |
| Solyc04g072047.1.1 |
| Solyc04g072135.1.1 |
| Solyc04g072435.1.1 |
| Solyc04g072505.1.1 |
| Solyc04g072603.1.1 |
| Solyc04g072607.1.1 |
| Solyc04g072725.1.1 |
| Solyc04g073953.1.1 |
| Solyc04g073957.1.1 |
| Solyc04g074163.1.1 |
| Solyc04g074167.1.1 |
| Solyc04g074195.1.1 |
| Solyc04g074535.1.1 |
| Solyc04g074865.1.1 |
| Solyc04g074925.1.1 |
| Solyc04g074963.1.1 |
| Solyc04g074967.1.1 |
| Solyc04g075012.1.1 |
| Solyc04g075014.1.1 |
| Solyc04g075016.1.1 |
| Solyc04g075018.1.1 |
| Solyc04g076033.1.1 |
| Solyc04g076037.1.1 |
| Solyc04g076155.1.1 |
| Solyc04g076523.1.1 |
| Solyc04g076845.1.1 |
| Solyc04g077535.1.1 |
| Solyc04g077755.1.1 |
| Solyc04g078075.1.1 |
| Solyc04g078085.1.1 |
| Solyc04g078095.1.1 |
| Solyc04g078195.1.1 |
| Solyc04g078325.1.1 |
| Solyc04g078975.1.1 |
| Solyc04g080395.1.1 |
| Solyc04g080873.1.1 |
| Solyc04g080877.1.1 |

| |
|---|
| Solyc04g080915.1.1 |
| Solyc04g081055.1.1 |
| Solyc04g081145.1.1 |
| Solyc04g081675.1.1 |
| Solyc04g081755.2.1 |
| Solyc04g082155.1.1 |
| Solyc04g082235.1.1 |
| Solyc04g082355.1.1 |
| Solyc04g082475.1.1 |
| Solyc04g082745.1.1 |
| Solyc04g082845.2.1 |
| Solyc04g082865.1.1 |
| Solyc04g082915.1.1 |
| Solyc04g150100.1.1 |
| Solyc04g150101.1.1 |
| Solyc04g150102.1.1 |
| Solyc04g150103.1.1 |
| Solyc04g150104.1.1 |
| Solyc04g150105.1.1 |
| Solyc04g150106.1.1 |
| Solyc04g150107.1.1 |
| Solyc04g150108.1.1 |
| Solyc04g150109.1.1 |
| Solyc04g150110.1.1 |
| Solyc04g150111.1.1 |
| Solyc04g150112.1.1 |
| Solyc04g150113.1.1 |
| Solyc04g150114.1.1 |
| Solyc04g150115.1.1 |
| Solyc04g150116.1.1 |
| Solyc04g150117.1.1 |
| Solyc04g150118.1.1 |
| Solyc04g150119.1.1 |
| Solyc04g150120.1.1 |
| Solyc04g150121.1.1 |
| Solyc04g150122.1.1 |
| Solyc04g150123.1.1 |
| Solyc04g150124.1.1 |
| Solyc04g150125.1.1 |
| Solyc04g150126.1.1 |
| Solyc04g150127.1.1 |
| Solyc04g150128.1.1 |
| Solyc04g150129.1.1 |

| |
|---|
| Solyc04g150130.1.1 |
| Solyc04g150131.1.1 |
| Solyc04g150132.1.1 |
| Solyc04g150133.1.1 |
| Solyc04g150134.1.1 |
| Solyc04g150135.1.1 |
| Solyc04g150136.1.1 |
| Solyc04g150137.1.1 |
| Solyc04g150138.1.1 |
| Solyc04g150139.1.1 |
| Solyc04g150140.1.1 |
| Solyc04g150141.1.1 |
| Solyc04g150142.1.1 |
| Solyc04g150143.1.1 |
| Solyc04g150144.1.1 |
| Solyc04g150145.1.1 |
| Solyc04g150146.1.1 |
| Solyc04g150147.1.1 |
| Solyc04g150148.1.1 |
| Solyc04g150150.1.1 |
| Solyc04g150151.1.1 |
| Solyc04g150152.1.1 |
| Solyc04g150153.1.1 |
| Solyc04g150154.1.1 |
| Solyc04g150155.1.1 |
| Solyc04g150156.1.1 |
| Solyc04g150157.1.1 |
| Solyc04g150158.1.1 |
| Solyc04g150159.1.1 |
| Solyc04g150160.1.1 |
| Solyc04g150161.1.1 |
| Solyc04g150162.1.1 |
| Solyc04g150163.1.1 |
| Solyc04g150164.1.1 |
| Solyc04g150166.1.1 |
| Solyc04g150167.1.1 |
| Solyc04g150168.1.1 |
| Solyc04g150169.1.1 |
| Solyc04g150170.1.1 |
| Solyc04g150171.1.1 |
| Solyc04g150172.1.1 |
| Solyc04g150173.1.1 |
| Solyc05g004000.1.1 |

| |
|---|
| Solyc05g004003.1.1 |
| Solyc05g004007.1.1 |
| Solyc05g005265.1.1 |
| Solyc05g005685.1.1 |
| Solyc05g005865.1.1 |
| Solyc05g006215.1.1 |
| Solyc05g006255.1.1 |
| Solyc05g006285.1.1 |
| Solyc05g006325.1.1 |
| Solyc05g006465.1.1 |
| Solyc05g006475.1.1 |
| Solyc05g006605.1.1 |
| Solyc05g006813.1.1 |
| Solyc05g006817.1.1 |
| Solyc05g006855.1.1 |
| Solyc05g006965.1.1 |
| Solyc05g007245.1.1 |
| Solyc05g007895.1.1 |
| Solyc05g008105.1.1 |
| Solyc05g008145.1.1 |
| Solyc05g008155.1.1 |
| Solyc05g008495.1.1 |
| Solyc05g008625.1.1 |
| Solyc05g008895.1.1 |
| Solyc05g009253.1.1 |
| Solyc05g009257.1.1 |
| Solyc05g009275.1.1 |
| Solyc05g009405.1.1 |
| Solyc05g009515.1.1 |
| Solyc05g009583.1.1 |
| Solyc05g009587.1.1 |
| Solyc05g009745.1.1 |
| Solyc05g009865.1.1 |
| Solyc05g009915.1.1 |
| Solyc05g010015.1.1 |
| Solyc05g010045.1.1 |
| Solyc05g010175.1.1 |
| Solyc05g010345.1.1 |
| Solyc05g010425.1.1 |
| Solyc05g010512.1.1 |
| Solyc05g010514.1.1 |
| Solyc05g010516.1.1 |
| Solyc05g010518.1.1 |

| |
|---|
| Solyc05g010555.1.1 |
| Solyc05g010685.1.1 |
| Solyc05g010725.1.1 |
| Solyc05g010735.1.1 |
| Solyc05g010755.1.1 |
| Solyc05g010813.1.1 |
| Solyc05g010817.1.1 |
| Solyc05g011825.2.1 |
| Solyc05g011883.1.1 |
| Solyc05g011887.1.1 |
| Solyc05g012172.1.1 |
| Solyc05g012173.1.1 |
| Solyc05g012177.1.1 |
| Solyc05g012325.1.1 |
| Solyc05g012745.1.1 |
| Solyc05g012827.1.1 |
| Solyc05g012955.1.1 |
| Solyc05g013005.1.1 |
| Solyc05g013225.1.1 |
| Solyc05g013535.1.1 |
| Solyc05g013613.1.1 |
| Solyc05g013617.1.1 |
| Solyc05g013843.1.1 |
| Solyc05g013847.1.1 |
| Solyc05g013905.1.1 |
| Solyc05g014051.1.1 |
| Solyc05g014052.1.1 |
| Solyc05g014054.1.1 |
| Solyc05g014056.1.1 |
| Solyc05g014058.1.1 |
| Solyc05g014175.1.1 |
| Solyc05g014177.2.1 |
| Solyc05g014275.2.1 |
| Solyc05g014375.1.1 |
| Solyc05g014505.1.1 |
| Solyc05g014795.1.1 |
| Solyc05g014843.1.1 |
| Solyc05g014847.1.1 |
| Solyc05g015005.1.1 |
| Solyc05g015075.1.1 |
| Solyc05g015437.1.1 |
| Solyc05g015473.1.1 |
| Solyc05g015477.1.1 |

| |
|---|
| Solyc05g015493.1.1 |
| Solyc05g015495.1.1 |
| Solyc05g015497.1.1 |
| Solyc05g015532.1.1 |
| Solyc05g015534.1.1 |
| Solyc05g015536.1.1 |
| Solyc05g015538.1.1 |
| Solyc05g015555.1.1 |
| Solyc05g015651.1.1 |
| Solyc05g015652.1.1 |
| Solyc05g015653.1.1 |
| Solyc05g015654.1.1 |
| Solyc05g015735.1.1 |
| Solyc05g015815.1.1 |
| Solyc05g015825.1.1 |
| Solyc05g015865.1.1 |
| Solyc05g015885.1.1 |
| Solyc05g016343.1.1 |
| Solyc05g016347.1.1 |
| Solyc05g016415.1.1 |
| Solyc05g016465.1.1 |
| Solyc05g016713.1.1 |
| Solyc05g016715.1.1 |
| Solyc05g016717.1.1 |
| Solyc05g017793.1.1 |
| Solyc05g017795.1.1 |
| Solyc05g017797.1.1 |
| Solyc05g017847.1.1 |
| Solyc05g017863.1.1 |
| Solyc05g017867.1.1 |
| Solyc05g017955.1.1 |
| Solyc05g018065.1.1 |
| Solyc05g018155.1.1 |
| Solyc05g018185.1.1 |
| Solyc05g018215.1.1 |
| Solyc05g018245.1.1 |
| Solyc05g018403.1.1 |
| Solyc05g018405.1.1 |
| Solyc05g018407.1.1 |
| Solyc05g018413.1.1 |
| Solyc05g018417.1.1 |
| Solyc05g018435.1.1 |
| Solyc05g018482.1.1 |

| |
|---|
| Solyc05g018484.1.1 |
| Solyc05g018486.1.1 |
| Solyc05g018488.1.1 |
| Solyc05g018563.1.1 |
| Solyc05g018567.1.1 |
| Solyc05g018575.1.1 |
| Solyc05g018657.1.1 |
| Solyc05g018743.1.1 |
| Solyc05g018747.1.1 |
| Solyc05g018775.1.1 |
| Solyc05g018865.1.1 |
| Solyc05g021165.1.1 |
| Solyc05g021245.1.1 |
| Solyc05g021405.1.1 |
| Solyc05g021415.1.1 |
| Solyc05g023635.1.1 |
| Solyc05g023725.1.1 |
| Solyc05g023733.1.1 |
| Solyc05g023737.1.1 |
| Solyc05g023755.1.1 |
| Solyc05g023825.1.1 |
| Solyc05g023841.1.1 |
| Solyc05g023842.1.1 |
| Solyc05g023844.1.1 |
| Solyc05g023846.1.1 |
| Solyc05g023848.1.1 |
| Solyc05g023895.1.1 |
| Solyc05g023995.1.1 |
| Solyc05g024235.1.1 |
| Solyc05g024315.1.1 |
| Solyc05g024385.1.1 |
| Solyc05g024453.1.1 |
| Solyc05g024457.1.1 |
| Solyc05g025513.1.1 |
| Solyc05g025517.1.1 |
| Solyc05g025605.1.1 |
| Solyc05g025625.1.1 |
| Solyc05g025665.1.1 |
| Solyc05g025725.1.1 |
| Solyc05g025785.1.1 |
| Solyc05g025815.1.1 |
| Solyc05g025903.1.1 |
| Solyc05g025907.1.1 |

| |
|---|
| Solyc05g025925.1.1 |
| Solyc05g025953.1.1 |
| Solyc05g025955.1.1 |
| Solyc05g025985.1.1 |
| Solyc05g026153.1.1 |
| Solyc05g026205.1.1 |
| Solyc05g026213.1.1 |
| Solyc05g026215.1.1 |
| Solyc05g026217.1.1 |
| Solyc05g026245.1.1 |
| Solyc05g026485.1.1 |
| Solyc05g026595.1.1 |
| Solyc05g026603.1.1 |
| Solyc05g026607.1.1 |
| Solyc05g032735.1.1 |
| Solyc05g032825.1.1 |
| Solyc05g040065.1.1 |
| Solyc05g041113.1.1 |
| Solyc05g041145.1.1 |
| Solyc05g041173.1.1 |
| Solyc05g041177.1.1 |
| Solyc05g041625.1.1 |
| Solyc05g041785.1.1 |
| Solyc05g042015.1.1 |
| Solyc05g042155.1.1 |
| Solyc05g042185.1.1 |
| Solyc05g043243.1.1 |
| Solyc05g043247.1.1 |
| Solyc05g043405.1.1 |
| Solyc05g044513.1.1 |
| Solyc05g044517.1.1 |
| Solyc05g044545.1.1 |
| Solyc05g044555.1.1 |
| Solyc05g045675.1.1 |
| Solyc05g045822.1.1 |
| Solyc05g045824.1.1 |
| Solyc05g045826.1.1 |
| Solyc05g045828.1.1 |
| Solyc05g045927.1.1 |
| Solyc05g045953.1.1 |
| Solyc05g045957.1.1 |
| Solyc05g046215.1.1 |
| Solyc05g046225.1.1 |

| |
|---|
| Solyc05g047485.1.1 |
| Solyc05g047643.1.1 |
| Solyc05g047645.1.1 |
| Solyc05g047647.1.1 |
| Solyc05g048755.1.1 |
| Solyc05g050003.1.1 |
| Solyc05g050007.1.1 |
| Solyc05g050053.1.1 |
| Solyc05g050057.1.1 |
| Solyc05g050065.1.1 |
| Solyc05g050213.1.1 |
| Solyc05g050217.1.1 |
| Solyc05g050365.1.1 |
| Solyc05g050515.1.1 |
| Solyc05g050555.1.1 |
| Solyc05g050685.1.1 |
| Solyc05g050825.1.1 |
| Solyc05g050883.1.1 |
| Solyc05g050887.1.1 |
| Solyc05g050895.1.1 |
| Solyc05g051153.1.1 |
| Solyc05g051157.1.1 |
| Solyc05g051345.1.1 |
| Solyc05g051385.1.1 |
| Solyc05g051425.2.1 |
| Solyc05g051495.1.1 |
| Solyc05g051545.1.1 |
| Solyc05g051583.1.1 |
| Solyc05g051595.1.1 |
| Solyc05g051735.1.1 |
| Solyc05g051815.1.1 |
| Solyc05g051825.1.1 |
| Solyc05g051875.1.1 |
| Solyc05g052015.1.1 |
| Solyc05g052265.1.1 |
| Solyc05g052275.1.1 |
| Solyc05g052715.1.1 |
| Solyc05g053133.1.1 |
| Solyc05g053137.1.1 |
| Solyc05g053155.1.1 |
| Solyc05g053465.1.1 |
| Solyc05g053555.1.1 |
| Solyc05g053835.2.1 |

| |
|---|
| Solyc05g053865.1.1 |
| Solyc05g054025.1.1 |
| Solyc05g054205.1.1 |
| Solyc05g054315.1.1 |
| Solyc05g054405.1.1 |
| Solyc05g054545.1.1 |
| Solyc05g055343.2.1 |
| Solyc05g055425.1.1 |
| Solyc05g055535.1.1 |
| Solyc05g055575.1.1 |
| Solyc05g056473.1.1 |
| Solyc05g056477.1.1 |
| Solyc05g150100.1.1 |
| Solyc05g150101.1.1 |
| Solyc05g150102.1.1 |
| Solyc05g150103.1.1 |
| Solyc05g150104.1.1 |
| Solyc05g150105.1.1 |
| Solyc05g150106.1.1 |
| Solyc05g150107.1.1 |
| Solyc05g150108.1.1 |
| Solyc05g150109.1.1 |
| Solyc05g150110.1.1 |
| Solyc05g150111.1.1 |
| Solyc05g150112.1.1 |
| Solyc05g150113.1.1 |
| Solyc05g150114.1.1 |
| Solyc05g150115.1.1 |
| Solyc05g150116.1.1 |
| Solyc05g150117.1.1 |
| Solyc05g150118.1.1 |
| Solyc05g150119.1.1 |
| Solyc05g150120.1.1 |
| Solyc05g150121.1.1 |
| Solyc05g150122.1.1 |
| Solyc05g150123.1.1 |
| Solyc05g150124.1.1 |
| Solyc05g150125.1.1 |
| Solyc05g150126.1.1 |
| Solyc05g150127.1.1 |
| Solyc05g150128.1.1 |
| Solyc05g150129.1.1 |
| Solyc05g150130.1.1 |

| |
|---|
| Solyc05g150131.1.1 |
| Solyc05g150132.1.1 |
| Solyc05g150133.1.1 |
| Solyc05g150134.1.1 |
| Solyc05g150135.1.1 |
| Solyc05g150136.1.1 |
| Solyc05g150137.1.1 |
| Solyc05g150138.1.1 |
| Solyc05g150139.1.1 |
| Solyc05g150140.1.1 |
| Solyc05g150141.1.1 |
| Solyc05g150142.1.1 |
| Solyc05g150143.1.1 |
| Solyc05g150144.1.1 |
| Solyc05g150145.1.1 |
| Solyc05g150146.1.1 |
| Solyc05g150147.1.1 |
| Solyc05g150148.1.1 |
| Solyc05g150149.1.1 |
| Solyc05g150150.1.1 |
| Solyc05g150151.1.1 |
| Solyc05g150152.1.1 |
| Solyc05g150153.1.1 |
| Solyc06g004000.1.1 |
| Solyc06g004005.1.1 |
| Solyc06g005385.1.1 |
| Solyc06g005395.1.1 |
| Solyc06g005465.1.1 |
| Solyc06g005525.1.1 |
| Solyc06g005635.1.1 |
| Solyc06g005795.2.1 |
| Solyc06g005825.1.1 |
| Solyc06g005965.1.1 |
| Solyc06g006013.1.1 |
| Solyc06g006015.1.1 |
| Solyc06g006017.1.1 |
| Solyc06g006022.1.1 |
| Solyc06g006024.1.1 |
| Solyc06g006026.1.1 |
| Solyc06g006028.1.1 |
| Solyc06g007275.1.1 |
| Solyc06g007395.1.1 |
| Solyc06g007493.1.1 |

| |
|---|
| Solyc06g007497.1.1 |
| Solyc06g007545.1.1 |
| Solyc06g008053.1.1 |
| Solyc06g008057.1.1 |
| Solyc06g008362.1.1 |
| Solyc06g008364.1.1 |
| Solyc06g008366.1.1 |
| Solyc06g008368.1.1 |
| Solyc06g008535.1.1 |
| Solyc06g008625.1.1 |
| Solyc06g008667.1.1 |
| Solyc06g008765.1.1 |
| Solyc06g008785.1.1 |
| Solyc06g009015.1.1 |
| Solyc06g009195.1.1 |
| Solyc06g009335.1.1 |
| Solyc06g009585.1.1 |
| Solyc06g009645.1.1 |
| Solyc06g009785.1.1 |
| Solyc06g009803.1.1 |
| Solyc06g009807.1.1 |
| Solyc06g009815.1.1 |
| Solyc06g009905.1.1 |
| Solyc06g009975.1.1 |
| Solyc06g009983.1.1 |
| Solyc06g009987.1.1 |
| Solyc06g010045.1.1 |
| Solyc06g010275.1.1 |
| Solyc06g011275.1.1 |
| Solyc06g011403.1.1 |
| Solyc06g011405.1.1 |
| Solyc06g011407.1.1 |
| Solyc06g011455.1.1 |
| Solyc06g011575.1.1 |
| Solyc06g011661.1.1 |
| Solyc06g011662.1.1 |
| Solyc06g011664.1.1 |
| Solyc06g011665.1.1 |
| Solyc06g011666.1.1 |
| Solyc06g011668.1.1 |
| Solyc06g011669.1.1 |
| Solyc06g011670.1.1 |
| Solyc06g016673.1.1 |

| |
|---|
| Solyc06g016677.1.1 |
| Solyc06g016685.1.1 |
| Solyc06g016725.1.1 |
| Solyc06g016823.1.1 |
| Solyc06g016827.1.1 |
| Solyc06g017973.1.1 |
| Solyc06g017975.1.1 |
| Solyc06g017977.1.1 |
| Solyc06g031712.1.1 |
| Solyc06g031714.1.1 |
| Solyc06g031716.1.1 |
| Solyc06g031718.1.1 |
| Solyc06g033783.1.1 |
| Solyc06g033787.1.1 |
| Solyc06g033815.1.1 |
| Solyc06g033825.1.1 |
| Solyc06g033895.1.1 |
| Solyc06g033955.1.1 |
| Solyc06g034025.1.1 |
| Solyc06g034055.1.1 |
| Solyc06g034073.1.1 |
| Solyc06g034077.1.1 |
| Solyc06g034233.1.1 |
| Solyc06g034237.1.1 |
| Solyc06g034255.1.1 |
| Solyc06g034395.1.1 |
| Solyc06g035495.1.1 |
| Solyc06g035625.1.1 |
| Solyc06g035723.1.1 |
| Solyc06g035727.1.1 |
| Solyc06g035875.1.1 |
| Solyc06g035883.1.1 |
| Solyc06g035887.1.1 |
| Solyc06g036213.1.1 |
| Solyc06g036217.1.1 |
| Solyc06g036255.1.1 |
| Solyc06g036555.1.1 |
| Solyc06g036725.1.1 |
| Solyc06g036805.1.1 |
| Solyc06g042945.1.1 |
| Solyc06g042955.1.1 |
| Solyc06g043035.1.1 |
| Solyc06g043065.1.1 |

| |
|---|
| Solyc06g043145.1.1 |
| Solyc06g043153.1.1 |
| Solyc06g043157.1.1 |
| Solyc06g048575.1.1 |
| Solyc06g048683.1.1 |
| Solyc06g048705.1.1 |
| Solyc06g048713.1.1 |
| Solyc06g048735.1.1 |
| Solyc06g048813.1.1 |
| Solyc06g048817.1.1 |
| Solyc06g048825.1.1 |
| Solyc06g048905.1.1 |
| Solyc06g048943.1.1 |
| Solyc06g048947.1.1 |
| Solyc06g049033.1.1 |
| Solyc06g049037.1.1 |
| Solyc06g050135.1.1 |
| Solyc06g050225.1.1 |
| Solyc06g050293.1.1 |
| Solyc06g050297.1.1 |
| Solyc06g050303.1.1 |
| Solyc06g050307.1.1 |
| Solyc06g050375.1.1 |
| Solyc06g050565.1.1 |
| Solyc06g050825.1.1 |
| Solyc06g050837.1.1 |
| Solyc06g050925.1.1 |
| Solyc06g051335.1.1 |
| Solyc06g051425.1.1 |
| Solyc06g051525.1.1 |
| Solyc06g051575.1.1 |
| Solyc06g051793.1.1 |
| Solyc06g051797.1.1 |
| Solyc06g052045.1.1 |
| Solyc06g052085.1.1 |
| Solyc06g052115.1.1 |
| Solyc06g052143.1.1 |
| Solyc06g052145.1.1 |
| Solyc06g052147.1.1 |
| Solyc06g053195.1.1 |
| Solyc06g053375.1.1 |
| Solyc06g053385.2.1 |
| Solyc06g053387.1.1 |

| |
|---|
| Solyc06g053665.1.1 |
| Solyc06g053695.1.1 |
| Solyc06g053785.1.1 |
| Solyc06g054185.1.1 |
| Solyc06g054195.1.1 |
| Solyc06g054315.1.1 |
| Solyc06g054685.1.1 |
| Solyc06g060015.1.1 |
| Solyc06g060105.1.1 |
| Solyc06g060195.1.1 |
| Solyc06g060215.1.1 |
| Solyc06g060445.1.1 |
| Solyc06g060623.1.1 |
| Solyc06g060627.1.1 |
| Solyc06g060675.1.1 |
| Solyc06g060735.1.1 |
| Solyc06g060785.1.1 |
| Solyc06g060885.1.1 |
| Solyc06g060955.1.1 |
| Solyc06g061005.1.1 |
| Solyc06g061015.1.1 |
| Solyc06g061023.1.1 |
| Solyc06g061027.1.1 |
| Solyc06g061225.1.1 |
| Solyc06g062295.1.1 |
| Solyc06g062445.1.1 |
| Solyc06g062765.1.1 |
| Solyc06g064533.1.1 |
| Solyc06g064537.1.1 |
| Solyc06g064673.1.1 |
| Solyc06g064677.1.1 |
| Solyc06g065073.1.1 |
| Solyc06g065077.1.1 |
| Solyc06g065255.1.1 |
| Solyc06g065445.1.1 |
| Solyc06g066065.1.1 |
| Solyc06g066245.1.1 |
| Solyc06g066345.1.1 |
| Solyc06g066475.1.1 |
| Solyc06g066853.1.1 |
| Solyc06g066857.1.1 |
| Solyc06g068055.1.1 |
| Solyc06g068165.1.1 |

| |
|---|
| Solyc06g068295.1.1 |
| Solyc06g068605.1.1 |
| Solyc06g068815.1.1 |
| Solyc06g068853.1.1 |
| Solyc06g068857.1.1 |
| Solyc06g069135.1.1 |
| Solyc06g069155.1.1 |
| Solyc06g069245.1.1 |
| Solyc06g069565.2.1 |
| Solyc06g071335.1.1 |
| Solyc06g071475.1.1 |
| Solyc06g071945.1.1 |
| Solyc06g072013.1.1 |
| Solyc06g072017.1.1 |
| Solyc06g072075.1.1 |
| Solyc06g072085.1.1 |
| Solyc06g072145.1.1 |
| Solyc06g072253.1.1 |
| Solyc06g072257.1.1 |
| Solyc06g072385.1.1 |
| Solyc06g072695.1.1 |
| Solyc06g072725.1.1 |
| Solyc06g072845.1.1 |
| Solyc06g073155.1.1 |
| Solyc06g073583.1.1 |
| Solyc06g073587.1.1 |
| Solyc06g073615.1.1 |
| Solyc06g073665.1.1 |
| Solyc06g073825.1.1 |
| Solyc06g074155.1.1 |
| Solyc06g074365.1.1 |
| Solyc06g074585.1.1 |
| Solyc06g075345.1.1 |
| Solyc06g075567.1.1 |
| Solyc06g075605.1.1 |
| Solyc06g075675.1.1 |
| Solyc06g075745.1.1 |
| Solyc06g075775.1.1 |
| Solyc06g076175.1.1 |
| Solyc06g076825.1.1 |
| Solyc06g076965.1.1 |
| Solyc06g082225.1.1 |
| Solyc06g082275.1.1 |

| |
|---|
| Solyc06g082353.1.1 |
| Solyc06g082357.1.1 |
| Solyc06g082595.1.1 |
| Solyc06g082793.1.1 |
| Solyc06g082797.1.1 |
| Solyc06g083005.1.1 |
| Solyc06g083505.1.1 |
| Solyc06g083705.1.1 |
| Solyc06g084325.1.1 |
| Solyc06g084625.1.1 |
| Solyc06g084825.1.1 |
| Solyc06g150100.1.1 |
| Solyc06g150101.1.1 |
| Solyc06g150102.1.1 |
| Solyc06g150103.1.1 |
| Solyc06g150104.1.1 |
| Solyc06g150105.1.1 |
| Solyc06g150106.1.1 |
| Solyc06g150107.1.1 |
| Solyc06g150108.1.1 |
| Solyc06g150109.1.1 |
| Solyc06g150110.1.1 |
| Solyc06g150111.1.1 |
| Solyc06g150112.1.1 |
| Solyc06g150113.1.1 |
| Solyc06g150114.1.1 |
| Solyc06g150115.1.1 |
| Solyc06g150116.1.1 |
| Solyc06g150117.1.1 |
| Solyc06g150118.1.1 |
| Solyc06g150119.1.1 |
| Solyc06g150120.1.1 |
| Solyc06g150121.1.1 |
| Solyc06g150122.1.1 |
| Solyc06g150123.1.1 |
| Solyc06g150124.1.1 |
| Solyc06g150125.1.1 |
| Solyc06g150126.1.1 |
| Solyc06g150127.1.1 |
| Solyc06g150128.1.1 |
| Solyc06g150129.1.1 |
| Solyc06g150130.1.1 |
| Solyc06g150131.1.1 |

| |
|---|
| Solyc06g150132.1.1 |
| Solyc06g150133.1.1 |
| Solyc06g150134.1.1 |
| Solyc06g150135.1.1 |
| Solyc06g150136.1.1 |
| Solyc06g150137.1.1 |
| Solyc06g150138.1.1 |
| Solyc06g150139.1.1 |
| Solyc07g005033.1.1 |
| Solyc07g005037.1.1 |
| Solyc07g005205.1.1 |
| Solyc07g005235.1.1 |
| Solyc07g005305.1.1 |
| Solyc07g005343.1.1 |
| Solyc07g005347.1.1 |
| Solyc07g005475.1.1 |
| Solyc07g005655.1.1 |
| Solyc07g005715.1.1 |
| Solyc07g005775.1.1 |
| Solyc07g005945.1.1 |
| Solyc07g005975.1.1 |
| Solyc07g006075.1.1 |
| Solyc07g006265.1.1 |
| Solyc07g006395.2.1 |
| Solyc07g006605.1.1 |
| Solyc07g006745.1.1 |
| Solyc07g006765.1.1 |
| Solyc07g006855.2.1 |
| Solyc07g006935.1.1 |
| Solyc07g007335.1.1 |
| Solyc07g007355.1.1 |
| Solyc07g007705.1.1 |
| Solyc07g007735.1.1 |
| Solyc07g007755.1.1 |
| Solyc07g008075.1.1 |
| Solyc07g008205.1.1 |
| Solyc07g008373.1.1 |
| Solyc07g008375.1.1 |
| Solyc07g008377.1.1 |
| Solyc07g008475.1.1 |
| Solyc07g008545.1.1 |
| Solyc07g008673.1.1 |
| Solyc07g008677.1.1 |

| |
|---|
| Solyc07g009133.1.1 |
| Solyc07g009137.1.1 |
| Solyc07g009233.1.1 |
| Solyc07g009237.1.1 |
| Solyc07g009265.1.1 |
| Solyc07g009405.1.1 |
| Solyc07g009435.1.1 |
| Solyc07g009437.1.1 |
| Solyc07g009443.1.1 |
| Solyc07g009447.1.1 |
| Solyc07g009473.1.1 |
| Solyc07g009477.1.1 |
| Solyc07g009545.1.1 |
| Solyc07g009563.1.1 |
| Solyc07g009567.1.1 |
| Solyc07g009583.1.1 |
| Solyc07g009585.1.1 |
| Solyc07g009587.1.1 |
| Solyc07g014583.1.1 |
| Solyc07g014587.1.1 |
| Solyc07g014595.1.1 |
| Solyc07g014707.1.1 |
| Solyc07g015765.1.1 |
| Solyc07g015805.1.1 |
| Solyc07g015873.1.1 |
| Solyc07g015877.1.1 |
| Solyc07g015905.1.1 |
| Solyc07g015983.1.1 |
| Solyc07g015987.1.1 |
| Solyc07g016005.1.1 |
| Solyc07g016095.1.1 |
| Solyc07g016113.1.1 |
| Solyc07g016117.1.1 |
| Solyc07g016145.1.1 |
| Solyc07g016215.1.1 |
| Solyc07g017225.1.1 |
| Solyc07g017255.1.1 |
| Solyc07g017405.1.1 |
| Solyc07g017545.1.1 |
| Solyc07g017605.1.1 |
| Solyc07g017675.1.1 |
| Solyc07g017785.1.1 |
| Solyc07g017805.1.1 |

| |
|---|
| Solyc07g017825.1.1 |
| Solyc07g017903.1.1 |
| Solyc07g017905.1.1 |
| Solyc07g017907.1.1 |
| Solyc07g017975.1.1 |
| Solyc07g018005.1.1 |
| Solyc07g018023.1.1 |
| Solyc07g018027.1.1 |
| Solyc07g018075.1.1 |
| Solyc07g018135.1.1 |
| Solyc07g018143.1.1 |
| Solyc07g018145.1.1 |
| Solyc07g018147.1.1 |
| Solyc07g018325.1.1 |
| Solyc07g018393.1.1 |
| Solyc07g018397.1.1 |
| Solyc07g018433.1.1 |
| Solyc07g018437.1.1 |
| Solyc07g019433.1.1 |
| Solyc07g019437.1.1 |
| Solyc07g019465.1.1 |
| Solyc07g019493.1.1 |
| Solyc07g019497.2.1 |
| Solyc07g019498.1.1 |
| Solyc07g019577.1.1 |
| Solyc07g019625.1.1 |
| Solyc07g020735.1.1 |
| Solyc07g020795.1.1 |
| Solyc07g020865.1.1 |
| Solyc07g020915.1.1 |
| Solyc07g020925.1.1 |
| Solyc07g020965.1.1 |
| Solyc07g021105.1.1 |
| Solyc07g021115.1.1 |
| Solyc07g021155.1.1 |
| Solyc07g021205.1.1 |
| Solyc07g021465.1.1 |
| Solyc07g021505.1.1 |
| Solyc07g021515.1.1 |
| Solyc07g022805.1.1 |
| Solyc07g022815.1.1 |
| Solyc07g022915.1.1 |
| Solyc07g024045.1.1 |

| |
|---|
| Solyc07g025095.1.1 |
| Solyc07g025115.1.1 |
| Solyc07g025385.1.1 |
| Solyc07g025453.1.1 |
| Solyc07g025457.1.1 |
| Solyc07g026605.1.1 |
| Solyc07g026705.1.1 |
| Solyc07g026865.1.1 |
| Solyc07g026873.1.1 |
| Solyc07g026877.1.1 |
| Solyc07g026893.1.1 |
| Solyc07g026897.1.1 |
| Solyc07g026915.1.1 |
| Solyc07g032023.1.1 |
| Solyc07g032027.1.1 |
| Solyc07g032105.1.1 |
| Solyc07g032155.1.1 |
| Solyc07g032395.1.1 |
| Solyc07g032595.1.1 |
| Solyc07g032623.1.1 |
| Solyc07g032627.1.1 |
| Solyc07g032683.1.1 |
| Solyc07g032687.1.1 |
| Solyc07g032781.1.1 |
| Solyc07g032782.1.1 |
| Solyc07g032784.1.1 |
| Solyc07g032786.1.1 |
| Solyc07g032788.1.1 |
| Solyc07g032789.1.1 |
| Solyc07g039343.1.1 |
| Solyc07g039347.1.1 |
| Solyc07g039455.1.1 |
| Solyc07g039615.1.1 |
| Solyc07g040661.1.1 |
| Solyc07g040663.2.1 |
| Solyc07g040855.1.1 |
| Solyc07g040905.1.1 |
| Solyc07g040965.1.1 |
| Solyc07g041027.1.1 |
| Solyc07g041035.1.1 |
| Solyc07g041215.1.1 |
| Solyc07g041235.1.1 |
| Solyc07g041365.1.1 |

| |
|---|
| Solyc07g041465.1.1 |
| Solyc07g041555.1.1 |
| Solyc07g041665.1.1 |
| Solyc07g041683.1.1 |
| Solyc07g041687.1.1 |
| Solyc07g041755.1.1 |
| Solyc07g041835.1.1 |
| Solyc07g041925.1.1 |
| Solyc07g042025.1.1 |
| Solyc07g042105.1.1 |
| Solyc07g042235.1.1 |
| Solyc07g042295.1.1 |
| Solyc07g042315.1.1 |
| Solyc07g042317.1.1 |
| Solyc07g042335.1.1 |
| Solyc07g042465.1.1 |
| Solyc07g042493.1.1 |
| Solyc07g042497.1.1 |
| Solyc07g042515.1.1 |
| Solyc07g042553.1.1 |
| Solyc07g042557.1.1 |
| Solyc07g042597.1.1 |
| Solyc07g042635.1.1 |
| Solyc07g042913.1.1 |
| Solyc07g042915.1.1 |
| Solyc07g042917.1.1 |
| Solyc07g042995.1.1 |
| Solyc07g043025.1.1 |
| Solyc07g043205.1.1 |
| Solyc07g043285.1.1 |
| Solyc07g043485.1.1 |
| Solyc07g043655.1.1 |
| Solyc07g043665.1.1 |
| Solyc07g043685.1.1 |
| Solyc07g043713.1.1 |
| Solyc07g043717.1.1 |
| Solyc07g044725.1.1 |
| Solyc07g044785.1.1 |
| Solyc07g044815.1.1 |
| Solyc07g044985.1.1 |
| Solyc07g045123.1.1 |
| Solyc07g045125.1.1 |
| Solyc07g045127.1.1 |

| |
|---|
| Solyc07g045255.1.1 |
| Solyc07g045353.1.1 |
| Solyc07g045455.1.1 |
| Solyc07g045525.1.1 |
| Solyc07g045565.1.1 |
| Solyc07g047745.1.1 |
| Solyc07g047823.1.1 |
| Solyc07g047827.1.1 |
| Solyc07g047915.1.1 |
| Solyc07g049135.1.1 |
| Solyc07g049215.1.1 |
| Solyc07g049405.1.1 |
| Solyc07g049495.1.1 |
| Solyc07g049505.1.1 |
| Solyc07g049525.1.1 |
| Solyc07g049563.1.1 |
| Solyc07g049565.1.1 |
| Solyc07g049567.1.1 |
| Solyc07g049635.1.1 |
| Solyc07g049645.1.1 |
| Solyc07g049655.1.1 |
| Solyc07g049805.1.1 |
| Solyc07g051845.1.1 |
| Solyc07g051883.1.1 |
| Solyc07g051887.1.1 |
| Solyc07g052075.1.1 |
| Solyc07g052165.1.1 |
| Solyc07g052365.1.1 |
| Solyc07g052445.1.1 |
| Solyc07g052485.1.1 |
| Solyc07g052555.1.1 |
| Solyc07g052785.1.1 |
| Solyc07g052813.1.1 |
| Solyc07g052817.1.1 |
| Solyc07g052885.1.1 |
| Solyc07g052913.1.1 |
| Solyc07g052917.1.1 |
| Solyc07g052985.1.1 |
| Solyc07g053025.1.1 |
| Solyc07g053035.1.1 |
| Solyc07g053155.1.1 |
| Solyc07g053225.1.1 |
| Solyc07g053493.1.1 |

| |
|---|
| Solyc07g053497.1.1 |
| Solyc07g053615.1.1 |
| Solyc07g053735.1.1 |
| Solyc07g054205.1.1 |
| Solyc07g054265.1.1 |
| Solyc07g054445.1.1 |
| Solyc07g054475.1.1 |
| Solyc07g054635.1.1 |
| Solyc07g055215.1.1 |
| Solyc07g055415.1.1 |
| Solyc07g055573.1.1 |
| Solyc07g055577.1.1 |
| Solyc07g056325.1.1 |
| Solyc07g056613.1.1 |
| Solyc07g056617.1.1 |
| Solyc07g056685.1.1 |
| Solyc07g056701.1.1 |
| Solyc07g056702.1.1 |
| Solyc07g056704.1.1 |
| Solyc07g056705.1.1 |
| Solyc07g056706.1.1 |
| Solyc07g056708.1.1 |
| Solyc07g061715.1.1 |
| Solyc07g061805.1.1 |
| Solyc07g061865.1.1 |
| Solyc07g061895.1.1 |
| Solyc07g061935.1.1 |
| Solyc07g062115.1.1 |
| Solyc07g062345.1.1 |
| Solyc07g062425.1.1 |
| Solyc07g062543.1.1 |
| Solyc07g062547.1.1 |
| Solyc07g062685.2.1 |
| Solyc07g062715.1.1 |
| Solyc07g063145.1.1 |
| Solyc07g063345.1.1 |
| Solyc07g063465.1.1 |
| Solyc07g063575.1.1 |
| Solyc07g063655.1.1 |
| Solyc07g064625.1.1 |
| Solyc07g065275.1.1 |
| Solyc07g065563.1.1 |
| Solyc07g065567.1.1 |

| |
|---|
| Solyc07g065745.1.1 |
| Solyc07g065845.1.1 |
| Solyc07g065923.1.1 |
| Solyc07g065927.1.1 |
| Solyc07g150100.1.1 |
| Solyc07g150101.1.1 |
| Solyc07g150102.1.1 |
| Solyc07g150103.1.1 |
| Solyc07g150104.1.1 |
| Solyc07g150105.1.1 |
| Solyc07g150106.1.1 |
| Solyc07g150107.1.1 |
| Solyc07g150108.1.1 |
| Solyc07g150109.1.1 |
| Solyc07g150110.1.1 |
| Solyc07g150111.1.1 |
| Solyc07g150112.1.1 |
| Solyc07g150113.1.1 |
| Solyc07g150114.1.1 |
| Solyc07g150115.1.1 |
| Solyc07g150116.1.1 |
| Solyc07g150117.1.1 |
| Solyc07g150118.1.1 |
| Solyc07g150119.1.1 |
| Solyc07g150120.1.1 |
| Solyc07g150121.1.1 |
| Solyc07g150122.1.1 |
| Solyc07g150123.1.1 |
| Solyc07g150124.1.1 |
| Solyc07g150125.1.1 |
| Solyc07g150126.1.1 |
| Solyc07g150127.1.1 |
| Solyc07g150128.1.1 |
| Solyc07g150129.1.1 |
| Solyc07g150130.1.1 |
| Solyc07g150131.1.1 |
| Solyc07g150132.1.1 |
| Solyc07g150133.1.1 |
| Solyc07g150134.1.1 |
| Solyc07g150135.1.1 |
| Solyc07g150136.1.1 |
| Solyc07g150137.1.1 |
| Solyc07g150138.1.1 |

| |
|---|
| Solyc07g150139.1.1 |
| Solyc07g150140.1.1 |
| Solyc07g150141.1.1 |
| Solyc07g150142.1.1 |
| Solyc07g150143.1.1 |
| Solyc07g150144.1.1 |
| Solyc07g150145.1.1 |
| Solyc07g150146.1.1 |
| Solyc07g150147.1.1 |
| Solyc07g150148.1.1 |
| Solyc07g150149.1.1 |
| Solyc07g150150.1.1 |
| Solyc07g150151.1.1 |
| Solyc07g150152.1.1 |
| Solyc08g004000.1.1 |
| Solyc08g004005.1.1 |
| Solyc08g005245.1.1 |
| Solyc08g005275.1.1 |
| Solyc08g005333.1.1 |
| Solyc08g005335.1.1 |
| Solyc08g005337.1.1 |
| Solyc08g005635.1.1 |
| Solyc08g005655.1.1 |
| Solyc08g005673.1.1 |
| Solyc08g005677.1.1 |
| Solyc08g005705.1.1 |
| Solyc08g005985.1.1 |
| Solyc08g006253.1.1 |
| Solyc08g006255.1.1 |
| Solyc08g006257.1.1 |
| Solyc08g006295.1.1 |
| Solyc08g006483.1.1 |
| Solyc08g006487.1.1 |
| Solyc08g006765.1.1 |
| Solyc08g006903.1.1 |
| Solyc08g006907.1.1 |
| Solyc08g007015.1.1 |
| Solyc08g007225.1.1 |
| Solyc08g007255.1.1 |
| Solyc08g007375.1.1 |
| Solyc08g007715.1.1 |
| Solyc08g007805.1.1 |
| Solyc08g007835.1.1 |

| |
|---|
| Solyc08g007863.1.1 |
| Solyc08g007867.1.1 |
| Solyc08g008015.1.1 |
| Solyc08g008045.1.1 |
| Solyc08g008085.1.1 |
| Solyc08g008305.1.1 |
| Solyc08g008453.1.1 |
| Solyc08g008457.1.1 |
| Solyc08g008495.1.1 |
| Solyc08g013803.1.1 |
| Solyc08g013805.2.1 |
| Solyc08g013807.1.1 |
| Solyc08g014015.1.1 |
| Solyc08g014035.1.1 |
| Solyc08g014105.1.1 |
| Solyc08g014175.1.1 |
| Solyc08g014245.1.1 |
| Solyc08g014393.1.1 |
| Solyc08g014397.1.1 |
| Solyc08g014455.1.1 |
| Solyc08g014515.1.1 |
| Solyc08g014565.1.1 |
| Solyc08g014605.1.1 |
| Solyc08g014612.1.1 |
| Solyc08g014614.1.1 |
| Solyc08g014616.1.1 |
| Solyc08g014618.1.1 |
| Solyc08g015631.1.1 |
| Solyc08g015632.1.1 |
| Solyc08g015636.1.1 |
| Solyc08g015638.1.1 |
| Solyc08g015713.1.1 |
| Solyc08g015715.1.1 |
| Solyc08g015717.1.1 |
| Solyc08g015755.1.1 |
| Solyc08g015785.1.1 |
| Solyc08g015833.1.1 |
| Solyc08g015837.1.1 |
| Solyc08g015905.1.1 |
| Solyc08g015965.1.1 |
| Solyc08g016075.1.1 |
| Solyc08g016313.1.1 |
| Solyc08g016315.1.1 |

| |
|---|
| Solyc08g016317.1.1 |
| Solyc08g016415.1.1 |
| Solyc08g016433.1.1 |
| Solyc08g016437.1.1 |
| Solyc08g016563.1.1 |
| Solyc08g016567.1.1 |
| Solyc08g016583.1.1 |
| Solyc08g016587.1.1 |
| Solyc08g016673.1.1 |
| Solyc08g016677.1.1 |
| Solyc08g016725.1.1 |
| Solyc08g016763.1.1 |
| Solyc08g016767.1.1 |
| Solyc08g016775.1.1 |
| Solyc08g016791.1.1 |
| Solyc08g016792.1.1 |
| Solyc08g016794.1.1 |
| Solyc08g016796.1.1 |
| Solyc08g016798.1.1 |
| Solyc08g016799.1.1 |
| Solyc08g016800.1.1 |
| Solyc08g016801.1.1 |
| Solyc08g016802.1.1 |
| Solyc08g016803.1.1 |
| Solyc08g016804.1.1 |
| Solyc08g016805.1.1 |
| Solyc08g016806.1.1 |
| Solyc08g016807.1.1 |
| Solyc08g016808.1.1 |
| Solyc08g021795.1.1 |
| Solyc08g022025.1.1 |
| Solyc08g022133.1.1 |
| Solyc08g022137.1.1 |
| Solyc08g023345.1.1 |
| Solyc08g023495.1.1 |
| Solyc08g023645.1.1 |
| Solyc08g028693.1.1 |
| Solyc08g028697.1.1 |
| Solyc08g028705.1.1 |
| Solyc08g028785.1.1 |
| Solyc08g028875.1.1 |
| Solyc08g028973.1.1 |
| Solyc08g029075.1.1 |

| |
|---|
| Solyc08g029235.1.1 |
| Solyc08g029342.1.1 |
| Solyc08g029344.1.1 |
| Solyc08g029346.1.1 |
| Solyc08g029385.1.1 |
| Solyc08g036413.1.1 |
| Solyc08g036417.1.1 |
| Solyc08g036503.1.1 |
| Solyc08g036505.2.1 |
| Solyc08g036507.1.1 |
| Solyc08g036533.1.1 |
| Solyc08g036535.1.1 |
| Solyc08g036537.1.1 |
| Solyc08g036575.1.1 |
| Solyc08g041685.1.1 |
| Solyc08g041825.1.1 |
| Solyc08g041865.1.1 |
| Solyc08g042025.1.1 |
| Solyc08g042065.1.1 |
| Solyc08g043143.1.1 |
| Solyc08g043147.1.1 |
| Solyc08g044243.1.1 |
| Solyc08g044247.1.1 |
| Solyc08g044305.1.1 |
| Solyc08g044333.1.1 |
| Solyc08g044337.1.1 |
| Solyc08g044345.1.1 |
| Solyc08g044445.1.1 |
| Solyc08g045733.1.1 |
| Solyc08g045737.1.1 |
| Solyc08g048315.1.1 |
| Solyc08g048395.1.1 |
| Solyc08g048455.1.1 |
| Solyc08g048503.1.1 |
| Solyc08g048507.1.1 |
| Solyc08g059655.1.1 |
| Solyc08g059715.1.1 |
| Solyc08g060815.1.1 |
| Solyc08g060925.1.1 |
| Solyc08g060945.1.1 |
| Solyc08g060973.1.1 |
| Solyc08g060977.1.1 |
| Solyc08g061025.1.1 |

| |
|---|
| Solyc08g061145.1.1 |
| Solyc08g061245.1.1 |
| Solyc08g061275.1.1 |
| Solyc08g061455.1.1 |
| Solyc08g061665.1.1 |
| Solyc08g061695.1.1 |
| Solyc08g061735.1.1 |
| Solyc08g061745.1.1 |
| Solyc08g061755.1.1 |
| Solyc08g061843.1.1 |
| Solyc08g061847.1.1 |
| Solyc08g061905.1.1 |
| Solyc08g061935.1.1 |
| Solyc08g061955.1.1 |
| Solyc08g062433.1.1 |
| Solyc08g062435.1.1 |
| Solyc08g062437.1.1 |
| Solyc08g062503.1.1 |
| Solyc08g062507.1.1 |
| Solyc08g062655.1.1 |
| Solyc08g062675.1.1 |
| Solyc08g062725.1.1 |
| Solyc08g062783.1.1 |
| Solyc08g062785.1.1 |
| Solyc08g062865.1.1 |
| Solyc08g062935.1.1 |
| Solyc08g063035.1.1 |
| Solyc08g063055.1.1 |
| Solyc08g065145.1.1 |
| Solyc08g065163.1.1 |
| Solyc08g065193.1.1 |
| Solyc08g065385.1.1 |
| Solyc08g065443.1.1 |
| Solyc08g065447.1.1 |
| Solyc08g065543.1.1 |
| Solyc08g065547.1.1 |
| Solyc08g065623.1.1 |
| Solyc08g065627.1.1 |
| Solyc08g065675.1.1 |
| Solyc08g065695.1.1 |
| Solyc08g065705.1.1 |
| Solyc08g065765.1.1 |
| Solyc08g066063.1.1 |

| |
|---|
| Solyc08g066067.1.1 |
| Solyc08g066335.1.1 |
| Solyc08g066355.1.1 |
| Solyc08g066405.1.1 |
| Solyc08g066485.1.1 |
| Solyc08g066495.1.1 |
| Solyc08g066625.1.1 |
| Solyc08g066705.1.1 |
| Solyc08g066923.1.1 |
| Solyc08g066927.1.1 |
| Solyc08g067025.1.1 |
| Solyc08g067125.1.1 |
| Solyc08g067127.1.1 |
| Solyc08g067165.1.1 |
| Solyc08g067405.1.1 |
| Solyc08g067765.1.1 |
| Solyc08g067793.1.1 |
| Solyc08g067795.2.1 |
| Solyc08g067796.1.1 |
| Solyc08g067797.1.1 |
| Solyc08g067935.1.1 |
| Solyc08g068075.1.1 |
| Solyc08g068145.1.1 |
| Solyc08g068665.1.1 |
| Solyc08g068713.1.1 |
| Solyc08g068717.1.1 |
| Solyc08g069095.1.1 |
| Solyc08g069231.1.1 |
| Solyc08g069232.1.1 |
| Solyc08g069233.1.1 |
| Solyc08g069234.1.1 |
| Solyc08g069235.1.1 |
| Solyc08g069236.1.1 |
| Solyc08g069237.1.1 |
| Solyc08g069238.1.1 |
| Solyc08g069239.1.1 |
| Solyc08g069240.1.1 |
| Solyc08g074307.2.1 |
| Solyc08g074525.1.1 |
| Solyc08g074655.1.1 |
| Solyc08g074682.1.1 |
| Solyc08g074683.1.1 |
| Solyc08g074795.1.1 |

| |
|---|
| Solyc08g074835.1.1 |
| Solyc08g074895.1.1 |
| Solyc08g074913.1.1 |
| Solyc08g074917.1.1 |
| Solyc08g074955.1.1 |
| Solyc08g075043.1.1 |
| Solyc08g075047.1.1 |
| Solyc08g075125.1.1 |
| Solyc08g075235.1.1 |
| Solyc08g075593.1.1 |
| Solyc08g075597.1.1 |
| Solyc08g076235.1.1 |
| Solyc08g076683.1.1 |
| Solyc08g076687.1.1 |
| Solyc08g076883.1.1 |
| Solyc08g076887.1.1 |
| Solyc08g077385.1.1 |
| Solyc08g077825.1.1 |
| Solyc08g078095.1.1 |
| Solyc08g078345.1.1 |
| Solyc08g078843.1.1 |
| Solyc08g078847.1.1 |
| Solyc08g078885.1.1 |
| Solyc08g079055.1.1 |
| Solyc08g079315.1.1 |
| Solyc08g080105.1.1 |
| Solyc08g080453.1.1 |
| Solyc08g080457.1.1 |
| Solyc08g080745.2.1 |
| Solyc08g081255.1.1 |
| Solyc08g081455.1.1 |
| Solyc08g081505.1.1 |
| Solyc08g081535.1.1 |
| Solyc08g081765.1.1 |
| Solyc08g081945.1.1 |
| Solyc08g082353.1.1 |
| Solyc08g082357.1.1 |
| Solyc08g082485.1.1 |
| Solyc08g082695.1.1 |
| Solyc08g082755.1.1 |
| Solyc08g083285.1.1 |
| Solyc08g083385.1.1 |
| Solyc08g150100.1.1 |

| |
|---|
| Solyc08g150101.1.1 |
| Solyc08g150102.1.1 |
| Solyc08g150103.1.1 |
| Solyc08g150104.1.1 |
| Solyc08g150105.1.1 |
| Solyc08g150106.1.1 |
| Solyc08g150107.1.1 |
| Solyc08g150108.1.1 |
| Solyc08g150109.1.1 |
| Solyc08g150110.1.1 |
| Solyc08g150111.1.1 |
| Solyc08g150112.1.1 |
| Solyc08g150113.1.1 |
| Solyc08g150114.1.1 |
| Solyc08g150115.1.1 |
| Solyc08g150116.1.1 |
| Solyc08g150117.1.1 |
| Solyc08g150118.1.1 |
| Solyc08g150119.1.1 |
| Solyc08g150120.1.1 |
| Solyc08g150121.1.1 |
| Solyc08g150122.1.1 |
| Solyc08g150123.1.1 |
| Solyc08g150124.1.1 |
| Solyc08g150125.1.1 |
| Solyc08g150126.1.1 |
| Solyc08g150127.1.1 |
| Solyc08g150128.1.1 |
| Solyc08g150129.1.1 |
| Solyc08g150130.1.1 |
| Solyc08g150131.1.1 |
| Solyc08g150132.1.1 |
| Solyc08g150133.1.1 |
| Solyc08g150134.1.1 |
| Solyc08g150135.1.1 |
| Solyc08g150136.1.1 |
| Solyc08g150137.1.1 |
| Solyc08g150138.1.1 |
| Solyc08g150139.1.1 |
| Solyc08g150140.1.1 |
| Solyc08g150141.1.1 |
| Solyc08g150142.1.1 |
| Solyc08g150143.1.1 |

| |
|---|
| Solyc08g150144.1.1 |
| Solyc08g150145.1.1 |
| Solyc08g150146.1.1 |
| Solyc08g150147.1.1 |
| Solyc08g150148.1.1 |
| Solyc08g150149.1.1 |
| Solyc08g150150.1.1 |
| Solyc09g004000.1.1 |
| Solyc09g005035.1.1 |
| Solyc09g005095.1.1 |
| Solyc09g005135.1.1 |
| Solyc09g005143.1.1 |
| Solyc09g005147.1.1 |
| Solyc09g005425.1.1 |
| Solyc09g005745.1.1 |
| Solyc09g006005.1.1 |
| Solyc09g007055.1.1 |
| Solyc09g008083.1.1 |
| Solyc09g008087.1.1 |
| Solyc09g008145.1.1 |
| Solyc09g008155.1.1 |
| Solyc09g008175.1.1 |
| Solyc09g008203.1.1 |
| Solyc09g008207.1.1 |
| Solyc09g008335.1.1 |
| Solyc09g008505.2.1 |
| Solyc09g008507.1.1 |
| Solyc09g008913.1.1 |
| Solyc09g009105.1.1 |
| Solyc09g009173.1.1 |
| Solyc09g009177.1.1 |
| Solyc09g009565.1.1 |
| Solyc09g009583.1.1 |
| Solyc09g009587.1.1 |
| Solyc09g009703.1.1 |
| Solyc09g009707.1.1 |
| Solyc09g009725.1.1 |
| Solyc09g009731.1.1 |
| Solyc09g009732.1.1 |
| Solyc09g009733.1.1 |
| Solyc09g009734.1.1 |
| Solyc09g009736.1.1 |
| Solyc09g009738.1.1 |

| |
|---|
| Solyc09g009895.1.1 |
| Solyc09g010285.1.1 |
| Solyc09g010485.1.1 |
| Solyc09g010495.1.1 |
| Solyc09g010565.1.1 |
| Solyc09g010605.1.1 |
| Solyc09g010995.1.1 |
| Solyc09g010997.1.1 |
| Solyc09g011125.1.1 |
| Solyc09g011223.1.1 |
| Solyc09g011227.1.1 |
| Solyc09g011455.1.1 |
| Solyc09g011645.1.1 |
| Solyc09g011665.1.1 |
| Solyc09g011695.1.1 |
| Solyc09g011865.1.1 |
| Solyc09g014215.2.1 |
| Solyc09g014525.1.1 |
| Solyc09g014565.1.1 |
| Solyc09g014641.1.1 |
| Solyc09g014642.1.1 |
| Solyc09g014643.1.1 |
| Solyc09g014644.1.1 |
| Solyc09g014645.1.1 |
| Solyc09g014646.1.1 |
| Solyc09g014647.1.1 |
| Solyc09g014648.1.1 |
| Solyc09g014649.1.1 |
| Solyc09g014705.1.1 |
| Solyc09g014765.1.1 |
| Solyc09g014785.1.1 |
| Solyc09g014805.1.1 |
| Solyc09g014935.1.1 |
| Solyc09g014945.1.1 |
| Solyc09g014993.1.1 |
| Solyc09g014997.1.1 |
| Solyc09g015033.1.1 |
| Solyc09g015037.1.1 |
| Solyc09g015065.1.1 |
| Solyc09g015085.1.1 |
| Solyc09g015135.1.1 |
| Solyc09g015145.1.1 |
| Solyc09g015223.1.1 |

| |
|---|
| Solyc09g015225.1.1 |
| Solyc09g015227.1.1 |
| Solyc09g015245.1.1 |
| Solyc09g015453.1.1 |
| Solyc09g015457.1.1 |
| Solyc09g015535.1.1 |
| Solyc09g015585.1.1 |
| Solyc09g015615.1.1 |
| Solyc09g015705.1.1 |
| Solyc09g016955.1.1 |
| Solyc09g018185.1.1 |
| Solyc09g018205.1.1 |
| Solyc09g018215.1.1 |
| Solyc09g018233.1.1 |
| Solyc09g018235.1.1 |
| Solyc09g018237.1.1 |
| Solyc09g018285.1.1 |
| Solyc09g018373.1.1 |
| Solyc09g018375.1.1 |
| Solyc09g018377.1.1 |
| Solyc09g018555.1.1 |
| Solyc09g018595.1.1 |
| Solyc09g018755.1.1 |
| Solyc09g018805.1.1 |
| Solyc09g020013.1.1 |
| Solyc09g020017.1.1 |
| Solyc09g020195.1.1 |
| Solyc09g025215.1.1 |
| Solyc09g025235.1.1 |
| Solyc09g030375.1.1 |
| Solyc09g030455.1.1 |
| Solyc09g030515.1.1 |
| Solyc09g031513.1.1 |
| Solyc09g031517.1.1 |
| Solyc09g031528.1.1 |
| Solyc09g031543.1.1 |
| Solyc09g031573.1.1 |
| Solyc09g031577.1.1 |
| Solyc09g031595.1.1 |
| Solyc09g031655.1.1 |
| Solyc09g037095.1.1 |
| Solyc09g037135.1.1 |
| Solyc09g037185.1.1 |

| |
|---|
| Solyc09g042575.1.1 |
| Solyc09g042665.1.1 |
| Solyc09g042673.1.1 |
| Solyc09g042677.1.1 |
| Solyc09g042705.1.1 |
| Solyc09g042723.1.1 |
| Solyc09g042725.1.1 |
| Solyc09g042727.1.1 |
| Solyc09g047935.1.1 |
| Solyc09g050061.1.1 |
| Solyc09g050062.1.1 |
| Solyc09g050063.1.1 |
| Solyc09g050064.1.1 |
| Solyc09g050065.1.1 |
| Solyc09g050066.1.1 |
| Solyc09g050067.1.1 |
| Solyc09g050068.1.1 |
| Solyc09g050069.1.1 |
| Solyc09g055185.1.1 |
| Solyc09g055265.1.1 |
| Solyc09g055425.2.1 |
| Solyc09g055427.1.1 |
| Solyc09g055475.1.1 |
| Solyc09g055545.1.1 |
| Solyc09g055713.1.1 |
| Solyc09g055717.1.1 |
| Solyc09g055755.1.1 |
| Solyc09g055805.1.1 |
| Solyc09g055953.1.1 |
| Solyc09g055955.1.1 |
| Solyc09g055957.1.1 |
| Solyc09g055995.1.1 |
| Solyc09g056175.1.1 |
| Solyc09g056222.1.1 |
| Solyc09g056224.1.1 |
| Solyc09g056226.1.1 |
| Solyc09g056228.1.1 |
| Solyc09g056233.1.1 |
| Solyc09g056237.1.1 |
| Solyc09g056393.1.1 |
| Solyc09g056397.1.1 |
| Solyc09g056453.1.1 |
| Solyc09g056457.1.1 |

| |
|---|
| Solyc09g057505.1.1 |
| Solyc09g057603.1.1 |
| Solyc09g057607.1.1 |
| Solyc09g057883.1.1 |
| Solyc09g058993.1.1 |
| Solyc09g058997.1.1 |
| Solyc09g059105.1.1 |
| Solyc09g059125.1.1 |
| Solyc09g059153.1.1 |
| Solyc09g059155.1.1 |
| Solyc09g059157.1.1 |
| Solyc09g059245.1.1 |
| Solyc09g059335.1.1 |
| Solyc09g059423.1.1 |
| Solyc09g059427.1.1 |
| Solyc09g059475.1.1 |
| Solyc09g059515.1.1 |
| Solyc09g059565.1.1 |
| Solyc09g059575.1.1 |
| Solyc09g059673.1.1 |
| Solyc09g059677.1.1 |
| Solyc09g059742.1.1 |
| Solyc09g059744.1.1 |
| Solyc09g059746.1.1 |
| Solyc09g059748.1.1 |
| Solyc09g059835.1.1 |
| Solyc09g059915.1.1 |
| Solyc09g059993.1.1 |
| Solyc09g059997.1.1 |
| Solyc09g060115.1.1 |
| Solyc09g060175.1.1 |
| Solyc09g060215.1.1 |
| Solyc09g061345.1.1 |
| Solyc09g061465.1.1 |
| Solyc09g061515.1.1 |
| Solyc09g061545.1.1 |
| Solyc09g061555.1.1 |
| Solyc09g061615.1.1 |
| Solyc09g061625.1.1 |
| Solyc09g061633.1.1 |
| Solyc09g061637.1.1 |
| Solyc09g061715.1.1 |
| Solyc09g061791.1.1 |

| |
|---|
| Solyc09g061792.1.1 |
| Solyc09g061794.1.1 |
| Solyc09g061796.1.1 |
| Solyc09g061798.1.1 |
| Solyc09g063157.2.1 |
| Solyc09g064165.1.1 |
| Solyc09g064243.1.1 |
| Solyc09g064245.1.1 |
| Solyc09g064247.1.1 |
| Solyc09g064273.1.1 |
| Solyc09g064645.1.1 |
| Solyc09g064695.1.1 |
| Solyc09g064755.1.1 |
| Solyc09g064815.1.1 |
| Solyc09g064885.1.1 |
| Solyc09g064915.1.1 |
| Solyc09g064925.1.1 |
| Solyc09g065025.1.1 |
| Solyc09g065105.1.1 |
| Solyc09g065335.1.1 |
| Solyc09g065355.1.1 |
| Solyc09g065373.1.1 |
| Solyc09g065377.1.1 |
| Solyc09g065387.1.1 |
| Solyc09g065475.1.1 |
| Solyc09g065485.1.1 |
| Solyc09g065525.1.1 |
| Solyc09g065565.1.1 |
| Solyc09g065665.1.1 |
| Solyc09g065755.1.1 |
| Solyc09g065885.1.1 |
| Solyc09g066005.1.1 |
| Solyc09g066035.1.1 |
| Solyc09g066085.1.1 |
| Solyc09g066323.1.1 |
| Solyc09g066325.1.1 |
| Solyc09g066327.1.1 |
| Solyc09g066405.1.1 |
| Solyc09g072593.1.1 |
| Solyc09g072595.1.1 |
| Solyc09g072597.1.1 |
| Solyc09g072753.1.1 |
| Solyc09g072757.1.1 |

| |
|---|
| Solyc09g072775.1.1 |
| Solyc09g072865.1.1 |
| Solyc09g072935.1.1 |
| Solyc09g073015.1.1 |
| Solyc09g074205.1.1 |
| Solyc09g074235.1.1 |
| Solyc09g074475.1.1 |
| Solyc09g074575.1.1 |
| Solyc09g074675.1.1 |
| Solyc09g074785.1.1 |
| Solyc09g074805.1.1 |
| Solyc09g075015.1.1 |
| Solyc09g075043.1.1 |
| Solyc09g075045.1.1 |
| Solyc09g075047.1.1 |
| Solyc09g075155.1.1 |
| Solyc09g075965.1.1 |
| Solyc09g075973.1.1 |
| Solyc09g075977.1.1 |
| Solyc09g076053.1.1 |
| Solyc09g076057.1.1 |
| Solyc09g082115.1.1 |
| Solyc09g082845.2.1 |
| Solyc09g082855.1.1 |
| Solyc09g083003.1.1 |
| Solyc09g083007.1.1 |
| Solyc09g083135.1.1 |
| Solyc09g083163.1.1 |
| Solyc09g083165.1.1 |
| Solyc09g083167.1.1 |
| Solyc09g083305.1.1 |
| Solyc09g083435.1.1 |
| Solyc09g089573.1.1 |
| Solyc09g089577.1.1 |
| Solyc09g089585.1.1 |
| Solyc09g089975.1.1 |
| Solyc09g090005.1.1 |
| Solyc09g090225.1.1 |
| Solyc09g090385.1.1 |
| Solyc09g090445.1.1 |
| Solyc09g090645.1.1 |
| Solyc09g090685.1.1 |
| Solyc09g090875.1.1 |

| |
|---|
| Solyc09g091415.1.1 |
| Solyc09g091455.1.1 |
| Solyc09g091615.1.1 |
| Solyc09g091883.1.1 |
| Solyc09g091887.1.1 |
| Solyc09g091965.1.1 |
| Solyc09g092025.1.1 |
| Solyc09g092145.1.1 |
| Solyc09g092235.1.1 |
| Solyc09g092283.1.1 |
| Solyc09g092287.1.1 |
| Solyc09g092645.1.1 |
| Solyc09g092725.1.1 |
| Solyc09g092763.1.1 |
| Solyc09g092765.1.1 |
| Solyc09g092767.1.1 |
| Solyc09g098115.1.1 |
| Solyc09g098117.1.1 |
| Solyc09g098385.1.1 |
| Solyc09g098425.1.1 |
| Solyc09g098601.1.1 |
| Solyc09g098602.1.1 |
| Solyc09g098603.1.1 |
| Solyc09g098604.1.1 |
| Solyc09g098605.1.1 |
| Solyc09g098606.1.1 |
| Solyc09g098607.1.1 |
| Solyc09g098608.1.1 |
| Solyc09g150100.1.1 |
| Solyc09g150101.1.1 |
| Solyc09g150102.1.1 |
| Solyc09g150103.1.1 |
| Solyc09g150104.1.1 |
| Solyc09g150105.1.1 |
| Solyc09g150106.1.1 |
| Solyc09g150107.1.1 |
| Solyc09g150108.1.1 |
| Solyc09g150109.1.1 |
| Solyc09g150110.1.1 |
| Solyc09g150111.1.1 |
| Solyc09g150112.1.1 |
| Solyc09g150113.1.1 |
| Solyc09g150114.1.1 |

| |
|---|
| Solyc09g150115.1.1 |
| Solyc09g150116.1.1 |
| Solyc09g150117.1.1 |
| Solyc09g150118.1.1 |
| Solyc09g150119.1.1 |
| Solyc09g150120.1.1 |
| Solyc09g150121.1.1 |
| Solyc09g150122.1.1 |
| Solyc09g150123.1.1 |
| Solyc09g150125.1.1 |
| Solyc09g150126.1.1 |
| Solyc09g150127.1.1 |
| Solyc09g150128.1.1 |
| Solyc09g150129.1.1 |
| Solyc09g150130.1.1 |
| Solyc09g150131.1.1 |
| Solyc09g150132.1.1 |
| Solyc09g150133.1.1 |
| Solyc09g150134.1.1 |
| Solyc09g150135.1.1 |
| Solyc09g150136.1.1 |
| Solyc09g150137.1.1 |
| Solyc09g150138.1.1 |
| Solyc09g150139.1.1 |
| Solyc09g150140.1.1 |
| Solyc09g150141.1.1 |
| Solyc09g150142.1.1 |
| Solyc09g150143.1.1 |
| Solyc09g150144.1.1 |
| Solyc09g150145.1.1 |
| Solyc09g150146.1.1 |
| Solyc09g150147.1.1 |
| Solyc09g150148.1.1 |
| Solyc09g150149.1.1 |
| Solyc09g150150.1.1 |
| Solyc09g150151.1.1 |
| Solyc09g150152.1.1 |
| Solyc09g150153.1.1 |
| Solyc09g150154.1.1 |
| Solyc09g150155.1.1 |
| Solyc09g150156.1.1 |
| Solyc09g150157.1.1 |
| Solyc09g150158.1.1 |

| |
|---|
| Solyc10g004000.1.1 |
| Solyc10g005125.1.1 |
| Solyc10g005335.1.1 |
| Solyc10g005485.1.1 |
| Solyc10g005905.1.1 |
| Solyc10g006095.1.1 |
| Solyc10g006173.1.1 |
| Solyc10g006177.1.1 |
| Solyc10g006275.1.1 |
| Solyc10g006615.1.1 |
| Solyc10g007073.1.1 |
| Solyc10g007075.1.1 |
| Solyc10g007077.1.1 |
| Solyc10g007156.2.1 |
| Solyc10g007635.1.1 |
| Solyc10g008205.2.1 |
| Solyc10g008243.1.1 |
| Solyc10g008247.1.1 |
| Solyc10g008263.1.1 |
| Solyc10g008267.1.1 |
| Solyc10g008445.1.1 |
| Solyc10g008455.1.1 |
| Solyc10g008665.1.1 |
| Solyc10g009365.1.1 |
| Solyc10g009433.1.1 |
| Solyc10g009437.1.1 |
| Solyc10g009487.1.1 |
| Solyc10g009595.1.1 |
| Solyc10g011685.1.1 |
| Solyc10g011765.1.1 |
| Solyc10g011823.1.1 |
| Solyc10g011827.1.1 |
| Solyc10g011925.1.1 |
| Solyc10g012135.1.1 |
| Solyc10g012143.1.1 |
| Solyc10g012147.1.1 |
| Solyc10g012213.1.1 |
| Solyc10g012217.1.1 |
| Solyc10g012245.1.1 |
| Solyc10g012295.1.1 |
| Solyc10g012435.1.1 |
| Solyc10g017725.1.1 |
| Solyc10g017753.1.1 |

| |
|---|
| Solyc10g017757.1.1 |
| Solyc10g017835.1.1 |
| Solyc10g017965.1.1 |
| Solyc10g017985.1.1 |
| Solyc10g017995.1.1 |
| Solyc10g018015.1.1 |
| Solyc10g018027.1.1 |
| Solyc10g018035.1.1 |
| Solyc10g018165.1.1 |
| Solyc10g018193.1.1 |
| Solyc10g018197.1.1 |
| Solyc10g018203.1.1 |
| Solyc10g018207.1.1 |
| Solyc10g018305.1.1 |
| Solyc10g018325.1.1 |
| Solyc10g018353.1.1 |
| Solyc10g018425.1.1 |
| Solyc10g018595.1.1 |
| Solyc10g018645.1.1 |
| Solyc10g018713.1.1 |
| Solyc10g018717.1.1 |
| Solyc10g018773.1.1 |
| Solyc10g018777.1.1 |
| Solyc10g018813.1.1 |
| Solyc10g018905.1.1 |
| Solyc10g018953.1.1 |
| Solyc10g018957.1.1 |
| Solyc10g019015.1.1 |
| Solyc10g019145.1.1 |
| Solyc10g019175.1.1 |
| Solyc10g019195.1.1 |
| Solyc10g019205.1.1 |
| Solyc10g019223.1.1 |
| Solyc10g019227.1.1 |
| Solyc10g024365.1.1 |
| Solyc10g024383.1.1 |
| Solyc10g024387.1.1 |
| Solyc10g024495.1.1 |
| Solyc10g032563.1.1 |
| Solyc10g032565.1.1 |
| Solyc10g032567.1.1 |
| Solyc10g038045.1.1 |
| Solyc10g038175.1.1 |

| |
|---|
| Solyc10g038183.1.1 |
| Solyc10g038185.1.1 |
| Solyc10g038187.1.1 |
| Solyc10g039193.1.1 |
| Solyc10g039196.1.1 |
| Solyc10g039197.2.1 |
| Solyc10g039215.1.1 |
| Solyc10g039235.1.1 |
| Solyc10g039293.1.1 |
| Solyc10g039297.1.1 |
| Solyc10g039383.1.1 |
| Solyc10g039387.1.1 |
| Solyc10g044443.1.1 |
| Solyc10g044447.1.1 |
| Solyc10g044525.1.1 |
| Solyc10g044535.1.1 |
| Solyc10g044565.1.1 |
| Solyc10g044585.1.1 |
| Solyc10g044683.1.1 |
| Solyc10g044687.1.1 |
| Solyc10g044965.1.1 |
| Solyc10g044982.1.1 |
| Solyc10g044984.1.1 |
| Solyc10g044986.1.1 |
| Solyc10g044988.1.1 |
| Solyc10g045103.1.1 |
| Solyc10g045155.1.1 |
| Solyc10g045157.1.1 |
| Solyc10g045233.1.1 |
| Solyc10g045237.1.1 |
| Solyc10g045385.1.1 |
| Solyc10g045435.1.1 |
| Solyc10g045437.1.1 |
| Solyc10g045453.1.1 |
| Solyc10g045525.1.1 |
| Solyc10g045723.1.1 |
| Solyc10g045727.1.1 |
| Solyc10g045765.1.1 |
| Solyc10g045773.1.1 |
| Solyc10g045777.1.1 |
| Solyc10g046775.1.1 |
| Solyc10g046803.1.1 |
| Solyc10g046807.1.1 |

| |
|---|
| Solyc10g046935.1.1 |
| Solyc10g047175.1.1 |
| Solyc10g047335.1.1 |
| Solyc10g047595.1.1 |
| Solyc10g047653.1.1 |
| Solyc10g047657.1.1 |
| Solyc10g047745.1.1 |
| Solyc10g047763.1.1 |
| Solyc10g047767.1.1 |
| Solyc10g047821.1.1 |
| Solyc10g047822.1.1 |
| Solyc10g047824.1.1 |
| Solyc10g047825.1.1 |
| Solyc10g047826.1.1 |
| Solyc10g047828.1.1 |
| Solyc10g047865.1.1 |
| Solyc10g047955.1.1 |
| Solyc10g048005.1.1 |
| Solyc10g048015.1.1 |
| Solyc10g048195.1.1 |
| Solyc10g049213.1.1 |
| Solyc10g049217.1.1 |
| Solyc10g049265.1.1 |
| Solyc10g049283.1.1 |
| Solyc10g049287.1.1 |
| Solyc10g049305.1.1 |
| Solyc10g049335.1.1 |
| Solyc10g049365.1.1 |
| Solyc10g049415.1.1 |
| Solyc10g049425.1.1 |
| Solyc10g049472.1.1 |
| Solyc10g049474.1.1 |
| Solyc10g049476.1.1 |
| Solyc10g049478.1.1 |
| Solyc10g049515.1.1 |
| Solyc10g049585.1.1 |
| Solyc10g049605.1.1 |
| Solyc10g049663.1.1 |
| Solyc10g049667.1.1 |
| Solyc10g049715.1.1 |
| Solyc10g049755.1.1 |
| Solyc10g049773.1.1 |
| Solyc10g049775.1.1 |

| |
|---|
| Solyc10g049777.1.1 |
| Solyc10g049815.1.1 |
| Solyc10g049913.1.1 |
| Solyc10g049917.1.1 |
| Solyc10g049945.1.1 |
| Solyc10g049993.1.1 |
| Solyc10g049997.1.1 |
| Solyc10g050055.1.1 |
| Solyc10g050065.1.1 |
| Solyc10g050085.1.1 |
| Solyc10g050105.1.1 |
| Solyc10g050223.1.1 |
| Solyc10g050227.1.1 |
| Solyc10g050323.1.1 |
| Solyc10g050325.2.1 |
| Solyc10g050445.1.1 |
| Solyc10g050453.1.1 |
| Solyc10g050457.1.1 |
| Solyc10g050513.1.1 |
| Solyc10g050517.1.1 |
| Solyc10g050773.1.1 |
| Solyc10g050777.1.1 |
| Solyc10g050895.1.1 |
| Solyc10g050993.1.1 |
| Solyc10g051035.1.1 |
| Solyc10g051125.1.1 |
| Solyc10g051165.1.1 |
| Solyc10g051215.1.1 |
| Solyc10g051255.1.1 |
| Solyc10g051373.1.1 |
| Solyc10g051377.1.1 |
| Solyc10g052515.1.1 |
| Solyc10g052565.1.1 |
| Solyc10g052603.1.1 |
| Solyc10g052607.1.1 |
| Solyc10g052685.1.1 |
| Solyc10g052712.1.1 |
| Solyc10g052714.1.1 |
| Solyc10g052716.1.1 |
| Solyc10g052718.1.1 |
| Solyc10g052813.1.1 |
| Solyc10g052817.1.1 |
| Solyc10g052823.1.1 |

| |
|---|
| Solyc10g052827.1.1 |
| Solyc10g052843.1.1 |
| Solyc10g052847.1.1 |
| Solyc10g054015.1.1 |
| Solyc10g054203.1.1 |
| Solyc10g054207.1.1 |
| Solyc10g054413.1.1 |
| Solyc10g054417.1.1 |
| Solyc10g054455.1.1 |
| Solyc10g054625.1.1 |
| Solyc10g054673.1.1 |
| Solyc10g054675.1.1 |
| Solyc10g054677.1.1 |
| Solyc10g054723.1.1 |
| Solyc10g054727.1.1 |
| Solyc10g055035.1.1 |
| Solyc10g055193.1.1 |
| Solyc10g055197.1.1 |
| Solyc10g055265.1.1 |
| Solyc10g055375.1.1 |
| Solyc10g055423.1.1 |
| Solyc10g055427.1.1 |
| Solyc10g055545.1.1 |
| Solyc10g055593.1.1 |
| Solyc10g055597.1.1 |
| Solyc10g055655.1.1 |
| Solyc10g055685.1.1 |
| Solyc10g055725.1.1 |
| Solyc10g061845.1.1 |
| Solyc10g061883.1.1 |
| Solyc10g061885.1.1 |
| Solyc10g061887.1.1 |
| Solyc10g061955.1.1 |
| Solyc10g062085.2.1 |
| Solyc10g062235.1.1 |
| Solyc10g068355.1.1 |
| Solyc10g073353.1.1 |
| Solyc10g073357.1.1 |
| Solyc10g074433.1.1 |
| Solyc10g074437.1.1 |
| Solyc10g074443.1.1 |
| Solyc10g074447.1.1 |
| Solyc10g074465.1.1 |

| |
|---|
| Solyc10g074475.1.1 |
| Solyc10g074643.1.1 |
| Solyc10g074647.1.1 |
| Solyc10g074785.1.1 |
| Solyc10g074822.1.1 |
| Solyc10g074824.1.1 |
| Solyc10g074826.1.1 |
| Solyc10g074828.1.1 |
| Solyc10g075065.1.1 |
| Solyc10g075103.1.1 |
| Solyc10g075107.1.1 |
| Solyc10g075111.1.1 |
| Solyc10g075112.1.1 |
| Solyc10g075114.1.1 |
| Solyc10g075116.1.1 |
| Solyc10g075118.1.1 |
| Solyc10g075173.1.1 |
| Solyc10g075177.1.1 |
| Solyc10g076245.1.1 |
| Solyc10g076355.1.1 |
| Solyc10g076405.1.1 |
| Solyc10g076415.1.1 |
| Solyc10g076435.1.1 |
| Solyc10g076515.1.1 |
| Solyc10g076565.1.1 |
| Solyc10g076915.1.1 |
| Solyc10g076945.1.1 |
| Solyc10g076995.1.1 |
| Solyc10g077065.1.1 |
| Solyc10g077105.1.1 |
| Solyc10g077135.1.1 |
| Solyc10g078175.1.1 |
| Solyc10g078275.1.1 |
| Solyc10g078325.1.1 |
| Solyc10g078715.1.1 |
| Solyc10g078775.1.1 |
| Solyc10g079125.1.1 |
| Solyc10g079155.1.1 |
| Solyc10g079755.1.1 |
| Solyc10g079795.1.1 |
| Solyc10g080725.1.1 |
| Solyc10g080782.1.1 |
| Solyc10g080784.1.1 |

| |
|---|
| Solyc10g080786.1.1 |
| Solyc10g080788.1.1 |
| Solyc10g081205.1.1 |
| Solyc10g081575.1.1 |
| Solyc10g081605.1.1 |
| Solyc10g081705.1.1 |
| Solyc10g081745.1.1 |
| Solyc10g081875.1.1 |
| Solyc10g082061.1.1 |
| Solyc10g082062.1.1 |
| Solyc10g082063.1.1 |
| Solyc10g082064.1.1 |
| Solyc10g082066.1.1 |
| Solyc10g082068.1.1 |
| Solyc10g083063.1.1 |
| Solyc10g083067.1.1 |
| Solyc10g083395.1.1 |
| Solyc10g084125.1.1 |
| Solyc10g084535.1.1 |
| Solyc10g084605.1.1 |
| Solyc10g085205.1.1 |
| Solyc10g085215.1.1 |
| Solyc10g085895.1.1 |
| Solyc10g086065.1.1 |
| Solyc10g086137.2.1 |
| Solyc10g086535.1.1 |
| Solyc10g086565.2.1 |
| Solyc10g086575.1.1 |
| Solyc10g086635.1.1 |
| Solyc10g086783.1.1 |
| Solyc10g086785.1.1 |
| Solyc10g086787.1.1 |
| Solyc10g087015.1.1 |
| Solyc10g087035.1.1 |
| Solyc10g150100.1.1 |
| Solyc10g150101.1.1 |
| Solyc10g150102.1.1 |
| Solyc10g150103.1.1 |
| Solyc10g150104.1.1 |
| Solyc10g150105.1.1 |
| Solyc10g150106.1.1 |
| Solyc10g150107.1.1 |
| Solyc10g150108.1.1 |

| |
|---|
| Solyc10g150109.1.1 |
| Solyc10g150110.1.1 |
| Solyc10g150111.1.1 |
| Solyc10g150112.1.1 |
| Solyc10g150113.1.1 |
| Solyc10g150114.1.1 |
| Solyc10g150115.1.1 |
| Solyc10g150116.1.1 |
| Solyc10g150117.1.1 |
| Solyc10g150118.1.1 |
| Solyc10g150119.1.1 |
| Solyc10g150120.1.1 |
| Solyc10g150121.1.1 |
| Solyc10g150122.1.1 |
| Solyc10g150123.1.1 |
| Solyc10g150124.1.1 |
| Solyc10g150125.1.1 |
| Solyc10g150126.1.1 |
| Solyc10g150127.1.1 |
| Solyc10g150128.1.1 |
| Solyc10g150129.1.1 |
| Solyc10g150130.1.1 |
| Solyc10g150131.1.1 |
| Solyc10g150132.1.1 |
| Solyc10g150133.1.1 |
| Solyc10g150134.1.1 |
| Solyc10g150135.1.1 |
| Solyc10g150136.1.1 |
| Solyc10g150137.1.1 |
| Solyc10g150138.1.1 |
| Solyc10g150139.1.1 |
| Solyc10g150140.1.1 |
| Solyc10g150141.1.1 |
| Solyc10g150142.1.1 |
| Solyc10g150143.1.1 |
| Solyc10g150144.1.1 |
| Solyc10g150145.1.1 |
| Solyc10g150146.1.1 |
| Solyc10g150147.1.1 |
| Solyc10g150148.1.1 |
| Solyc10g150149.1.1 |
| Solyc10g150150.1.1 |
| Solyc10g150151.1.1 |

| |
|---|
| Solyc11g004000.1.1 |
| Solyc11g004005.1.1 |
| Solyc11g005525.1.1 |
| Solyc11g005615.1.1 |
| Solyc11g005635.1.1 |
| Solyc11g005675.1.1 |
| Solyc11g005845.1.1 |
| Solyc11g005935.1.1 |
| Solyc11g006205.1.1 |
| Solyc11g006455.1.1 |
| Solyc11g006485.1.1 |
| Solyc11g006505.1.1 |
| Solyc11g006635.1.1 |
| Solyc11g006675.1.1 |
| Solyc11g006783.1.1 |
| Solyc11g006785.1.1 |
| Solyc11g006787.1.1 |
| Solyc11g006805.1.1 |
| Solyc11g006955.1.1 |
| Solyc11g007585.1.1 |
| Solyc11g007825.1.1 |
| Solyc11g007915.1.1 |
| Solyc11g007945.1.1 |
| Solyc11g008223.1.1 |
| Solyc11g008227.1.1 |
| Solyc11g008245.1.1 |
| Solyc11g008905.1.1 |
| Solyc11g010245.1.1 |
| Solyc11g010735.1.1 |
| Solyc11g010813.1.1 |
| Solyc11g010815.1.1 |
| Solyc11g010817.1.1 |
| Solyc11g010845.1.1 |
| Solyc11g011275.1.1 |
| Solyc11g011373.1.1 |
| Solyc11g011377.1.1 |
| Solyc11g011415.1.1 |
| Solyc11g011495.1.1 |
| Solyc11g011533.1.1 |
| Solyc11g011537.1.1 |
| Solyc11g011585.1.1 |
| Solyc11g011673.1.1 |
| Solyc11g011677.1.1 |

| |
|---|
| Solyc11g011755.1.1 |
| Solyc11g012195.1.1 |
| Solyc11g012325.1.1 |
| Solyc11g012385.1.1 |
| Solyc11g012645.1.1 |
| Solyc11g012665.1.1 |
| Solyc11g012705.2.1 |
| Solyc11g012815.1.1 |
| Solyc11g012923.1.1 |
| Solyc11g012927.1.1 |
| Solyc11g012995.1.1 |
| Solyc11g013215.1.1 |
| Solyc11g013353.1.1 |
| Solyc11g013357.1.1 |
| Solyc11g013633.1.1 |
| Solyc11g013635.1.1 |
| Solyc11g013637.1.1 |
| Solyc11g013745.1.1 |
| Solyc11g013855.1.1 |
| Solyc11g016945.1.1 |
| Solyc11g016993.1.1 |
| Solyc11g016997.1.1 |
| Solyc11g017105.1.1 |
| Solyc11g017305.1.1 |
| Solyc11g017345.1.1 |
| Solyc11g017377.1.1 |
| Solyc11g017475.1.1 |
| Solyc11g018623.1.1 |
| Solyc11g018625.1.1 |
| Solyc11g018627.1.1 |
| Solyc11g018715.1.1 |
| Solyc11g018772.1.1 |
| Solyc11g018774.1.1 |
| Solyc11g018776.1.1 |
| Solyc11g018778.1.1 |
| Solyc11g018805.1.1 |
| Solyc11g018855.1.1 |
| Solyc11g019965.1.1 |
| Solyc11g020015.1.1 |
| Solyc11g020105.1.1 |
| Solyc11g020213.1.1 |
| Solyc11g020215.1.1 |
| Solyc11g020245.1.1 |

| |
|---|
| Solyc11g020363.1.1 |
| Solyc11g020365.1.1 |
| Solyc11g020367.1.1 |
| Solyc11g020433.1.1 |
| Solyc11g020437.1.1 |
| Solyc11g020475.1.1 |
| Solyc11g020503.1.1 |
| Solyc11g020507.1.1 |
| Solyc11g020575.1.1 |
| Solyc11g020785.1.1 |
| Solyc11g020823.1.1 |
| Solyc11g020827.1.1 |
| Solyc11g020843.1.1 |
| Solyc11g020847.1.1 |
| Solyc11g020873.1.1 |
| Solyc11g020877.1.1 |
| Solyc11g020905.1.1 |
| Solyc11g020955.1.1 |
| Solyc11g020975.1.1 |
| Solyc11g020995.1.1 |
| Solyc11g021063.1.1 |
| Solyc11g021067.1.1 |
| Solyc11g021195.1.1 |
| Solyc11g021213.1.1 |
| Solyc11g021215.1.1 |
| Solyc11g021217.1.1 |
| Solyc11g021233.1.1 |
| Solyc11g021237.1.1 |
| Solyc11g021363.1.1 |
| Solyc11g021365.2.1 |
| Solyc11g022465.1.1 |
| Solyc11g027887.2.1 |
| Solyc11g027895.1.1 |
| Solyc11g027897.1.1 |
| Solyc11g027933.1.1 |
| Solyc11g027935.1.1 |
| Solyc11g027937.1.1 |
| Solyc11g028025.1.1 |
| Solyc11g028105.1.1 |
| Solyc11g028165.1.1 |
| Solyc11g028213.1.1 |
| Solyc11g028217.1.1 |
| Solyc11g028325.1.1 |

| |
|---|
| Solyc11g028355.1.1 |
| Solyc11g030383.1.1 |
| Solyc11g030387.1.1 |
| Solyc11g030553.1.1 |
| Solyc11g030555.1.1 |
| Solyc11g030557.1.1 |
| Solyc11g030645.1.1 |
| Solyc11g030915.1.1 |
| Solyc11g030935.1.1 |
| Solyc11g032025.1.1 |
| Solyc11g032045.1.1 |
| Solyc11g032065.1.1 |
| Solyc11g033281.1.1 |
| Solyc11g033282.1.1 |
| Solyc11g033283.1.1 |
| Solyc11g033284.1.1 |
| Solyc11g033286.1.1 |
| Solyc11g033288.1.1 |
| Solyc11g039535.1.1 |
| Solyc11g039573.1.1 |
| Solyc11g039577.1.1 |
| Solyc11g039655.1.1 |
| Solyc11g039713.1.1 |
| Solyc11g039717.1.1 |
| Solyc11g039765.1.1 |
| Solyc11g039785.1.1 |
| Solyc11g039805.1.1 |
| Solyc11g039835.1.1 |
| Solyc11g040065.1.1 |
| Solyc11g040085.1.1 |
| Solyc11g040173.1.1 |
| Solyc11g040223.1.1 |
| Solyc11g040225.1.1 |
| Solyc11g040227.1.1 |
| Solyc11g042455.1.1 |
| Solyc11g042805.1.1 |
| Solyc11g042853.1.1 |
| Solyc11g042857.1.1 |
| Solyc11g042873.1.1 |
| Solyc11g042877.1.1 |
| Solyc11g044235.1.1 |
| Solyc11g044315.1.1 |
| Solyc11g044323.1.1 |

| |
|---|
| Solyc11g044325.1.1 |
| Solyc11g044327.1.1 |
| Solyc11g044453.1.1 |
| Solyc11g044457.1.1 |
| Solyc11g044515.1.1 |
| Solyc11g044605.1.1 |
| Solyc11g044643.1.1 |
| Solyc11g044645.1.1 |
| Solyc11g044647.1.1 |
| Solyc11g044665.1.1 |
| Solyc11g044805.1.1 |
| Solyc11g044875.1.1 |
| Solyc11g044933.1.1 |
| Solyc11g044937.1.1 |
| Solyc11g044943.1.1 |
| Solyc11g044947.1.1 |
| Solyc11g045125.1.1 |
| Solyc11g045215.1.1 |
| Solyc11g045225.1.1 |
| Solyc11g045332.1.1 |
| Solyc11g045334.1.1 |
| Solyc11g045336.1.1 |
| Solyc11g045338.1.1 |
| Solyc11g045435.1.1 |
| Solyc11g045615.1.1 |
| Solyc11g045675.1.1 |
| Solyc11g050735.1.1 |
| Solyc11g050855.1.1 |
| Solyc11g051015.1.1 |
| Solyc11g051123.1.1 |
| Solyc11g051127.1.1 |
| Solyc11g051171.1.1 |
| Solyc11g051172.1.1 |
| Solyc11g051173.1.1 |
| Solyc11g051174.1.1 |
| Solyc11g051176.1.1 |
| Solyc11g051178.1.1 |
| Solyc11g056293.1.1 |
| Solyc11g056297.1.1 |
| Solyc11g056595.1.1 |
| Solyc11g056635.1.1 |
| Solyc11g061755.1.1 |
| Solyc11g061785.1.1 |

| |
|---|
| Solyc11g061813.1.1 |
| Solyc11g061817.1.1 |
| Solyc11g061835.1.1 |
| Solyc11g061855.1.1 |
| Solyc11g061985.1.1 |
| Solyc11g062065.1.1 |
| Solyc11g062155.1.1 |
| Solyc11g063573.1.1 |
| Solyc11g063577.1.1 |
| Solyc11g063743.1.1 |
| Solyc11g063747.1.1 |
| Solyc11g064775.1.1 |
| Solyc11g064845.1.1 |
| Solyc11g064945.1.1 |
| Solyc11g065205.1.1 |
| Solyc11g065365.1.1 |
| Solyc11g065555.1.1 |
| Solyc11g065835.1.1 |
| Solyc11g065935.1.1 |
| Solyc11g066005.1.1 |
| Solyc11g066163.1.1 |
| Solyc11g066165.1.1 |
| Solyc11g066167.1.1 |
| Solyc11g066255.1.1 |
| Solyc11g066325.1.1 |
| Solyc11g066685.1.1 |
| Solyc11g067343.1.1 |
| Solyc11g067345.1.1 |
| Solyc11g067347.1.1 |
| Solyc11g068515.1.1 |
| Solyc11g068625.1.1 |
| Solyc11g068893.1.1 |
| Solyc11g068897.1.1 |
| Solyc11g068905.1.1 |
| Solyc11g069055.1.1 |
| Solyc11g069093.1.1 |
| Solyc11g069097.1.1 |
| Solyc11g069115.1.1 |
| Solyc11g069625.1.1 |
| Solyc11g069735.1.1 |
| Solyc11g069765.1.1 |
| Solyc11g070175.1.1 |
| Solyc11g070183.1.1 |

| |
|---|
| Solyc11g070187.1.1 |
| Solyc11g071223.1.1 |
| Solyc11g071227.1.1 |
| Solyc11g071295.1.1 |
| Solyc11g072475.1.1 |
| Solyc11g072605.1.1 |
| Solyc11g072623.1.1 |
| Solyc11g072627.1.1 |
| Solyc11g072765.1.1 |
| Solyc11g072825.1.1 |
| Solyc11g072935.1.1 |
| Solyc11g073055.1.1 |
| Solyc11g073075.1.1 |
| Solyc11g073231.1.1 |
| Solyc11g073232.1.1 |
| Solyc11g073234.1.1 |
| Solyc11g073236.1.1 |
| Solyc11g073238.1.1 |
| Solyc11g073265.1.1 |
| Solyc11g073315.1.1 |
| Solyc11g150100.1.1 |
| Solyc11g150101.1.1 |
| Solyc11g150102.1.1 |
| Solyc11g150103.1.1 |
| Solyc11g150104.1.1 |
| Solyc11g150105.1.1 |
| Solyc11g150106.1.1 |
| Solyc11g150107.1.1 |
| Solyc11g150108.1.1 |
| Solyc11g150109.1.1 |
| Solyc11g150110.1.1 |
| Solyc11g150111.1.1 |
| Solyc11g150112.1.1 |
| Solyc11g150113.1.1 |
| Solyc11g150114.1.1 |
| Solyc11g150115.1.1 |
| Solyc11g150116.1.1 |
| Solyc11g150117.1.1 |
| Solyc11g150118.1.1 |
| Solyc11g150119.1.1 |
| Solyc11g150120.1.1 |
| Solyc11g150121.1.1 |
| Solyc11g150122.1.1 |

| |
|---|
| Solyc11g150123.1.1 |
| Solyc11g150124.1.1 |
| Solyc11g150125.1.1 |
| Solyc11g150126.1.1 |
| Solyc11g150127.1.1 |
| Solyc11g150128.1.1 |
| Solyc11g150129.1.1 |
| Solyc11g150130.1.1 |
| Solyc11g150131.1.1 |
| Solyc11g150132.1.1 |
| Solyc11g150133.1.1 |
| Solyc11g150134.1.1 |
| Solyc11g150135.1.1 |
| Solyc11g150136.1.1 |
| Solyc11g150137.1.1 |
| Solyc11g150138.1.1 |
| Solyc11g150139.1.1 |
| Solyc11g150140.1.1 |
| Solyc11g150141.1.1 |
| Solyc11g150142.1.1 |
| Solyc11g150143.1.1 |
| Solyc11g150144.1.1 |
| Solyc11g150145.1.1 |
| Solyc11g150146.1.1 |
| Solyc12g005465.2.1 |
| Solyc12g005615.1.1 |
| Solyc12g005645.1.1 |
| Solyc12g005783.1.1 |
| Solyc12g005787.1.1 |
| Solyc12g005795.1.1 |
| Solyc12g005955.1.1 |
| Solyc12g006003.1.1 |
| Solyc12g006007.1.1 |
| Solyc12g006293.1.1 |
| Solyc12g006297.1.1 |
| Solyc12g006313.1.1 |
| Solyc12g006317.1.1 |
| Solyc12g006505.1.1 |
| Solyc12g006575.1.1 |
| Solyc12g006695.1.1 |
| Solyc12g006705.1.1 |
| Solyc12g006805.1.1 |
| Solyc12g006973.1.1 |

| |
|---|
| Solyc12g006977.1.1 |
| Solyc12g006997.1.1 |
| Solyc12g007045.1.1 |
| Solyc12g007285.1.1 |
| Solyc12g008335.1.1 |
| Solyc12g009245.1.1 |
| Solyc12g009345.1.1 |
| Solyc12g009535.1.1 |
| Solyc12g009565.1.1 |
| Solyc12g009695.1.1 |
| Solyc12g009875.1.1 |
| Solyc12g010585.1.1 |
| Solyc12g010595.1.1 |
| Solyc12g010655.1.1 |
| Solyc12g010755.1.1 |
| Solyc12g011023.1.1 |
| Solyc12g011027.1.1 |
| Solyc12g011033.1.1 |
| Solyc12g011037.1.1 |
| Solyc12g011085.1.1 |
| Solyc12g011215.1.1 |
| Solyc12g011233.1.1 |
| Solyc12g011237.1.1 |
| Solyc12g011255.1.1 |
| Solyc12g011455.1.1 |
| Solyc12g013535.1.1 |
| Solyc12g013655.1.1 |
| Solyc12g013895.1.1 |
| Solyc12g013935.1.1 |
| Solyc12g014545.1.1 |
| Solyc12g014595.1.1 |
| Solyc12g014615.1.1 |
| Solyc12g015923.1.1 |
| Solyc12g015927.1.1 |
| Solyc12g015985.1.1 |
| Solyc12g016143.1.1 |
| Solyc12g016147.1.1 |
| Solyc12g017263.1.1 |
| Solyc12g017265.1.1 |
| Solyc12g017267.1.1 |
| Solyc12g017395.1.1 |
| Solyc12g017475.1.1 |
| Solyc12g017545.1.1 |

| |
|---|
| Solyc12g017575.1.1 |
| Solyc12g017675.2.1 |
| Solyc12g017705.1.1 |
| Solyc12g017883.1.1 |
| Solyc12g017887.1.1 |
| Solyc12g019055.1.1 |
| Solyc12g019144.1.1 |
| Solyc12g019235.1.1 |
| Solyc12g019335.1.1 |
| Solyc12g019343.1.1 |
| Solyc12g019347.1.1 |
| Solyc12g019473.1.1 |
| Solyc12g019477.1.1 |
| Solyc12g019645.1.1 |
| Solyc12g019715.1.1 |
| Solyc12g019795.1.1 |
| Solyc12g019913.1.1 |
| Solyc12g019917.1.1 |
| Solyc12g019945.1.1 |
| Solyc12g019955.1.1 |
| Solyc12g019973.1.1 |
| Solyc12g019977.1.1 |
| Solyc12g020035.1.1 |
| Solyc12g021193.1.1 |
| Solyc12g021197.1.1 |
| Solyc12g021215.1.1 |
| Solyc12g021265.1.1 |
| Solyc12g021323.1.1 |
| Solyc12g021327.1.1 |
| Solyc12g021363.1.1 |
| Solyc12g021367.1.1 |
| Solyc12g026475.1.1 |
| Solyc12g027565.1.1 |
| Solyc12g027655.1.1 |
| Solyc12g027705.1.1 |
| Solyc12g027805.1.1 |
| Solyc12g027893.1.1 |
| Solyc12g027897.1.1 |
| Solyc12g033105.1.1 |
| Solyc12g035192.1.1 |
| Solyc12g035194.1.1 |
| Solyc12g035196.1.1 |
| Solyc12g035198.1.1 |

| |
|---|
| Solyc12g035245.1.1 |
| Solyc12g035523.1.1 |
| Solyc12g035525.1.1 |
| Solyc12g035527.1.1 |
| Solyc12g035815.1.1 |
| Solyc12g035825.1.1 |
| Solyc12g035845.1.1 |
| Solyc12g035895.1.1 |
| Solyc12g036263.1.1 |
| Solyc12g036267.1.1 |
| Solyc12g036283.1.1 |
| Solyc12g036287.1.1 |
| Solyc12g036445.1.1 |
| Solyc12g036455.1.1 |
| Solyc12g036475.1.1 |
| Solyc12g036483.1.1 |
| Solyc12g036485.1.1 |
| Solyc12g036487.1.1 |
| Solyc12g036673.2.1 |
| Solyc12g036715.1.1 |
| Solyc12g036825.1.1 |
| Solyc12g038015.1.1 |
| Solyc12g038125.1.1 |
| Solyc12g038225.1.1 |
| Solyc12g038343.1.1 |
| Solyc12g038347.1.1 |
| Solyc12g038425.1.1 |
| Solyc12g038453.1.1 |
| Solyc12g038457.1.1 |
| Solyc12g038523.1.1 |
| Solyc12g038527.1.1 |
| Solyc12g038553.1.1 |
| Solyc12g038557.1.1 |
| Solyc12g038875.1.1 |
| Solyc12g038985.1.1 |
| Solyc12g039185.1.1 |
| Solyc12g039205.1.1 |
| Solyc12g040255.1.1 |
| Solyc12g040285.1.1 |
| Solyc12g040365.1.1 |
| Solyc12g040395.1.1 |
| Solyc12g040535.1.1 |
| Solyc12g040575.1.1 |

| |
|---|
| Solyc12g040642.1.1 |
| Solyc12g040644.1.1 |
| Solyc12g040646.1.1 |
| Solyc12g040648.1.1 |
| Solyc12g040755.1.1 |
| Solyc12g040813.1.1 |
| Solyc12g040815.1.1 |
| Solyc12g040817.1.1 |
| Solyc12g040875.1.1 |
| Solyc12g041895.1.1 |
| Solyc12g042005.1.1 |
| Solyc12g042015.1.1 |
| Solyc12g042035.1.1 |
| Solyc12g042055.1.1 |
| Solyc12g042123.1.1 |
| Solyc12g042127.1.1 |
| Solyc12g042215.1.1 |
| Solyc12g042395.1.1 |
| Solyc12g042485.1.1 |
| Solyc12g042505.1.1 |
| Solyc12g042545.1.1 |
| Solyc12g042565.1.1 |
| Solyc12g042603.1.1 |
| Solyc12g042607.1.1 |
| Solyc12g042625.1.1 |
| Solyc12g042695.1.1 |
| Solyc12g042705.1.1 |
| Solyc12g042793.1.1 |
| Solyc12g042797.1.1 |
| Solyc12g042845.1.1 |
| Solyc12g042855.1.1 |
| Solyc12g042963.1.1 |
| Solyc12g042964.1.1 |
| Solyc12g042965.2.1 |
| Solyc12g043095.1.1 |
| Solyc12g044435.1.1 |
| Solyc12g044455.1.1 |
| Solyc12g044585.1.1 |
| Solyc12g044693.1.1 |
| Solyc12g044695.1.1 |
| Solyc12g044697.1.1 |
| Solyc12g044775.1.1 |
| Solyc12g044795.1.1 |

| |
|---|
| Solyc12g044952.1.1 |
| Solyc12g044954.1.1 |
| Solyc12g044956.1.1 |
| Solyc12g044958.1.1 |
| Solyc12g045033.1.1 |
| Solyc12g045035.1.1 |
| Solyc12g045037.1.1 |
| Solyc12g049095.1.1 |
| Solyc12g049175.1.1 |
| Solyc12g049255.1.1 |
| Solyc12g049365.1.1 |
| Solyc12g049475.1.1 |
| Solyc12g049611.1.1 |
| Solyc12g049612.1.1 |
| Solyc12g049614.1.1 |
| Solyc12g049616.1.1 |
| Solyc12g049618.1.1 |
| Solyc12g055683.1.1 |
| Solyc12g055687.1.1 |
| Solyc12g055855.2.1 |
| Solyc12g056025.1.1 |
| Solyc12g056115.1.1 |
| Solyc12g056123.1.1 |
| Solyc12g056127.1.1 |
| Solyc12g056285.1.1 |
| Solyc12g056305.1.1 |
| Solyc12g056345.1.1 |
| Solyc12g056585.1.1 |
| Solyc12g056615.1.1 |
| Solyc12g056671.1.1 |
| Solyc12g056672.1.1 |
| Solyc12g056674.1.1 |
| Solyc12g056675.1.1 |
| Solyc12g056676.1.1 |
| Solyc12g056678.1.1 |
| Solyc12g056745.1.1 |
| Solyc12g056865.1.1 |
| Solyc12g056993.1.1 |
| Solyc12g056995.2.1 |
| Solyc12g062175.1.1 |
| Solyc12g062203.1.1 |
| Solyc12g062207.1.1 |
| Solyc12g062227.1.1 |

| |
|---|
| Solyc12g062285.1.1 |
| Solyc12g062345.1.1 |
| Solyc12g062425.1.1 |
| Solyc12g062443.1.1 |
| Solyc12g062447.1.1 |
| Solyc12g062525.1.1 |
| Solyc12g062563.1.1 |
| Solyc12g062567.1.1 |
| Solyc12g062705.1.1 |
| Solyc12g062733.2.1 |
| Solyc12g062822.1.1 |
| Solyc12g062824.1.1 |
| Solyc12g062826.1.1 |
| Solyc12g062828.1.1 |
| Solyc12g062855.1.1 |
| Solyc12g062935.1.1 |
| Solyc12g068075.1.1 |
| Solyc12g070123.1.1 |
| Solyc12g070127.1.1 |
| Solyc12g070153.1.1 |
| Solyc12g070155.1.1 |
| Solyc12g070156.2.1 |
| Solyc12g070157.1.1 |
| Solyc12g077353.1.1 |
| Solyc12g077355.1.1 |
| Solyc12g077357.1.1 |
| Solyc12g077395.1.1 |
| Solyc12g077523.1.1 |
| Solyc12g077525.1.1 |
| Solyc12g077527.1.1 |
| Solyc12g077623.1.1 |
| Solyc12g077627.1.1 |
| Solyc12g082763.1.1 |
| Solyc12g082765.1.1 |
| Solyc12g082767.1.1 |
| Solyc12g082791.1.1 |
| Solyc12g082792.1.1 |
| Solyc12g082793.1.1 |
| Solyc12g082794.1.1 |
| Solyc12g082795.1.1 |
| Solyc12g082796.1.1 |
| Solyc12g082797.1.1 |
| Solyc12g082798.1.1 |

| |
|---|
| Solyc12g082799.1.1 |
| Solyc12g087895.1.1 |
| Solyc12g088005.1.1 |
| Solyc12g088195.1.1 |
| Solyc12g088665.1.1 |
| Solyc12g089145.1.1 |
| Solyc12g089205.1.1 |
| Solyc12g089385.1.1 |
| Solyc12g094385.1.1 |
| Solyc12g094435.1.1 |
| Solyc12g096105.1.1 |
| Solyc12g096145.1.1 |
| Solyc12g096175.1.1 |
| Solyc12g096195.1.1 |
| Solyc12g096295.1.1 |
| Solyc12g098095.1.1 |
| Solyc12g098225.1.1 |
| Solyc12g098463.1.1 |
| Solyc12g098467.1.1 |
| Solyc12g098545.1.1 |
| Solyc12g098603.1.1 |
| Solyc12g098607.1.1 |
| Solyc12g098805.1.1 |
| Solyc12g098823.1.1 |
| Solyc12g098827.1.1 |
| Solyc12g098945.1.1 |
| Solyc12g099025.1.1 |
| Solyc12g099395.1.1 |
| Solyc12g099465.1.1 |
| Solyc12g099655.1.1 |
| Solyc12g099705.1.1 |
| Solyc12g100363.1.1 |
| Solyc12g100367.1.1 |
| Solyc12g150100.1.1 |
| Solyc12g150101.1.1 |
| Solyc12g150102.1.1 |
| Solyc12g150103.1.1 |
| Solyc12g150104.1.1 |
| Solyc12g150105.1.1 |
| Solyc12g150106.1.1 |
| Solyc12g150107.1.1 |
| Solyc12g150108.1.1 |
| Solyc12g150109.1.1 |

| |
|---|
| Solyc12g150110.1.1 |
| Solyc12g150111.1.1 |
| Solyc12g150112.1.1 |
| Solyc12g150113.1.1 |
| Solyc12g150114.1.1 |
| Solyc12g150115.1.1 |
| Solyc12g150116.1.1 |
| Solyc12g150117.1.1 |
| Solyc12g150118.1.1 |
| Solyc12g150119.1.1 |
| Solyc12g150120.1.1 |
| Solyc12g150121.1.1 |
| Solyc12g150122.1.1 |
| Solyc12g150123.1.1 |
| Solyc12g150124.1.1 |
| Solyc12g150125.1.1 |
| Solyc12g150126.1.1 |
| Solyc12g150127.1.1 |
| Solyc12g150128.1.1 |
| Solyc12g150129.1.1 |
| Solyc12g150130.1.1 |
| Solyc12g150131.1.1 |
| Solyc12g150132.1.1 |
