## Supplementary Table 3 for "An improved de novo assembly and annotation of the tomato reference genome using single-molecule sequencing, Hi-C proximity ligation and optical maps"

| **Suplementary Table 3:** SRA data from a variety of tissues and conditions of wild-type tomato plants |
|---|
| **SRA ID** |
| SRR926182 |
| SRR924101 |
| SRR926185 |
| SRR924102 |
| SRR567999 |
| SRR788904 |
| SRR788859 |
| SRR567998 |
| SRR567997 |
| SRR949057 |
| SRR963469 |
| SRR948546 |
| SRR948545 |
| SRR988676 |
| SRR963795 |
| SRR993740 |
| SRR988680 |
| SRR768851 |
| SRR654026 |
| SRR768823 |
| SRR993743 |
| SRR653965 |
| SRR1210486 |
| SRR1210490 |
| SRR1210489 |
| SRR1210487 |
| SRR1210488 |
| SRR1210493 |
| SRR1210492 |
| SRR1210491 |
| SRR654013 |
| SRR654006 |
| SRR653970 |
| SRR950390 |
| SRR988673 |
| SRR653990 |
| SRR943813 |
| SRR768837 |
| SRR768850 |
| SRR943820 |
| SRR768824 |

| |
|---|
| SRR988675 |
| SRR943814 |
| SRR653974 |
| SRR768845 |
| SRR768836 |
| SRR988681 |
| SRR653997 |
| SRR768832 |
| SRR768814 |
| SRR943821 |
| SRR768863 |
| SRR943822 |
| SRR654009 |
| SRR653989 |
| SRR768841 |
| SRR654021 |
| SRR988682 |
| SRR768818 |
| SRR768862 |
| SRR943815 |
| SRR768826 |
| SRR943825 |
| SRR768860 |
| SRR653995 |
| SRR1379924 |
| SRR1379927 |
| SRR768859 |
| SRR768846 |
| SRR768815 |
| SRR653932 |
| SRR768833 |
| SRR1380025 |
| SRR768825 |
| SRR768861 |
| SRR768857 |
| SRR768855 |
| SRR768835 |
| SRR768822 |
| SRR654008 |
| SRR768829 |
| SRR768858 |
| SRR988677 |
| SRR653986 |

| |
|---|
| SRR863042 |
| SRR943828 |
| SRR653973 |
| SRR768852 |
| SRR653987 |
| SRR943830 |
| SRR768819 |
| SRR768834 |
| SRR653993 |
| SRR768842 |
| SRR768847 |
| SRR768838 |
| SRR653933 |
| SRR654018 |
| SRR768849 |
| SRR768828 |
| SRR768817 |
| SRR653959 |
| SRR943819 |
| SRR653991 |
| SRR768844 |
| SRR654022 |
| SRR768840 |
| SRR653984 |
| SRR653996 |
| SRR768856 |
| SRR768853 |
| SRR768848 |
| SRR1379921 |
| SRR863093 |
| SRR653960 |
| SRR653998 |
| SRR654019 |
| SRR943823 |
| SRR653962 |
| SRR943827 |
| SRR943824 |
| SRR768854 |
| SRR863087 |
| SRR653992 |
| SRR768827 |
| SRR654000 |
| SRR654025 |

| |
|---|
| SRR653964 |
| SRR653934 |
| SRR654010 |
| SRR654004 |
| SRR653969 |
| SRR653988 |
| SRR1379926 |
| SRR654017 |
| SRR768831 |
| SRR768839 |
| SRR653935 |
| SRR768816 |
| SRR768864 |
| SRR1379911 |
| SRR768843 |
| SRR653971 |
| SRR1379923 |
| SRR654020 |
| SRR654012 |
| SRR1223239 |
| SRR943816 |
| SRR653983 |
| SRR654005 |
| SRR1379912 |
| SRR654024 |
| SRR1379914 |
| SRR863046 |
| SRR1379917 |
| SRR654003 |
| SRR1379922 |
| SRR654016 |
| SRR653961 |
| SRR1223225 |
| SRR654002 |
| SRR1379915 |
| SRR1379925 |
| SRR1379919 |
| SRR943829 |
| SRR863090 |
| SRR653972 |
| SRR653999 |
| SRR863069 |
| SRR654011 |

| |
|---|
| SRR654015 |
| SRR653936 |
| SRR1223238 |
| SRR1223226 |
| SRR1379916 |
| SRR943826 |
| SRR654023 |
| SRR1379928 |
| SRR653968 |
| SRR1379913 |
| SRR943818 |
| SRR863044 |
| SRR863076 |
| SRR1013186 |
| SRR863058 |
| SRR654007 |
| SRR1379920 |
| SRR863073 |
| SRR1379918 |
| SRR863061 |
| SRR863064 |
| SRR863071 |
| SRR863070 |
| SRR863082 |
| SRR654001 |
| SRR653963 |
| SRR653985 |
| SRR1013172 |
| SRR1013135 |
| SRR863075 |
| SRR863079 |
| SRR1013329 |
| SRR863074 |
| SRR863065 |
| SRR1013227 |
| SRR654014 |
| SRR1013183 |
| SRR863048 |
| SRR863084 |
| SRR863077 |
| SRR863060 |
| SRR863045 |
| SRR1013297 |

| |
|---|
| SRR863083 |
| SRR943817 |
| SRR863080 |
| SRR1013163 |
| SRR863047 |
| SRR1013235 |
| SRR863094 |
| SRR1013309 |
| SRR1013072 |
| SRR1013328 |
| SRR863063 |
| SRR1013285 |
| SRR863062 |
| SRR863068 |
| SRR1013036 |
| SRR1013223 |
| SRR863059 |
| SRR863088 |
| SRR863067 |
| SRR1013061 |
| SRR1013232 |
| SRR863050 |
| SRR1013301 |
| SRR1013255 |
| SRR1013068 |
| SRR1013159 |
| SRR1013113 |
| SRR1013306 |
| SRR1013125 |
| SRR863081 |
| SRR1013092 |
| SRR1013277 |
| SRR1013104 |
| SRR863057 |
| SRR1013090 |
| SRR1013112 |
| SRR1013120 |
| SRR1013207 |
| SRR1013269 |
| SRR1013133 |
| SRR1013175 |
| SRR1013116 |
| SRR653994 |

| |
|---|
| SRR1013076 |
| SRR1013111 |
| SRR1013278 |
| SRR1013312 |
| SRR1013321 |
| SRR1013305 |
| SRR1013187 |
| SRR863086 |
| SRR1013260 |
| SRR1013248 |
| SRR1013084 |
| SRR1013325 |
| SRR1013243 |
| SRR1013237 |
| SRR1013086 |
| SRR1013298 |
| SRR1013323 |
| SRR1013290 |
| SRR1013247 |
| SRR863078 |
| SRR1013158 |
| SRR1013096 |
| SRR1013109 |
| SRR863054 |
| SRR1013156 |
| SRR1013147 |
| SRR1013121 |
| SRR1013151 |
| SRR1013139 |
| SRR1013195 |
| SRR1013191 |
| SRR1013293 |
| SRR1013081 |
| SRR863066 |
| SRR1013289 |
| SRR1013127 |
| SRR1013075 |
| SRR567663 |
| SRR1013272 |
| SRR1013038 |
| SRR1013313 |
| SRR1013062 |
| SRR1013327 |

| |
|---|
| SRR1013326 |
| SRR1013044 |
| SRR1013129 |
| SRR1013258 |
| SRR863092 |
| SRR567664 |
| SRR1013063 |
| SRR1013069 |
| SRR1013087 |
| SRR1013335 |
| SRR1013101 |
| SRR863072 |
| SRR863043 |
| SRR1013281 |
| SRR1013131 |
| SRR1013213 |
| SRR1013039 |
| SRR1013132 |
| SRR1013331 |
| SRR1013130 |
| SRR1013173 |
| SRR1013201 |
| SRR1013246 |
| SRR1013088 |
| SRR1013205 |
| SRR1013189 |
| SRR1013287 |
| SRR1013110 |
| SRR1013177 |
| SRR1013148 |
| SRR1013155 |
| SRR567667 |
| SRR1013048 |
| SRR1013145 |
| SRR1013266 |
| SRR1013094 |
| SRR1013307 |
| SRR567665 |
| SRR1013265 |
| SRR1013170 |
| SRR1013152 |
| SRR1013046 |
| SRR1013052 |

| |
|---|
| SRR567668 |
| SRR1013169 |
| SRR1013276 |
| SRR1013124 |
| SRR1013193 |
| SRR567666 |
| SRR863085 |
| SRR1013299 |
| SRR1013165 |
| SRR1013263 |
| SRR1013054 |
| SRR1013089 |
| SRR1013334 |
| SRR1013136 |
| SRR1013114 |
| SRR1013042 |
| SRR1013181 |
| SRR1013171 |
| SRR1013197 |
| SRR1013264 |
| SRR1013053 |
| SRR1013303 |
| SRR1013150 |
| SRR1013282 |
| SRR1013142 |
| SRR1013199 |
| SRR1013270 |
| SRR863091 |
| SRR863055 |
| SRR1013065 |
| SRR1013102 |
| SRR1013211 |
| SRR567679 |
| SRR1013060 |
| SRR1013174 |
| SRR1013252 |
| SRR1013045 |
| SRR1013108 |
| SRR1013071 |
| SRR1013311 |
| SRR1013241 |
| SRR1013203 |
| SRR567689 |

| |
|---|
| SRR1013209 |
| SRR1013122 |
| SRR1013261 |
| SRR567690 |
| SRR1013245 |
| SRR1013200 |
| SRR863052 |
| SRR1013082 |
| SRR1013070 |
| SRR1013041 |
| SRR1013047 |
| SRR1013168 |
| SRR1013279 |
| SRR863089 |
| SRR1013226 |
| SRR567680 |
| SRR1013236 |
| SRR1013202 |
| SRR567675 |
| SRR1013215 |
| SRR1013161 |
| SRR1013067 |
| SRR1013244 |
| SRR1013128 |
| SRR1013224 |
| SRR1013250 |
| SRR1013049 |
| SRR1013262 |
| SRR1013256 |
| SRR1013283 |
| SRR1013085 |
| SRR1013315 |
| SRR1013188 |
| SRR567669 |
| SRR1013184 |
| SRR567693 |
| SRR1013318 |
| SRR1013274 |
| SRR1013115 |
| SRR1013105 |
| SRR1013254 |
| SRR1013295 |
| SRR1013157 |

| |
|---|
| SRR567694 |
| SRR1013064 |
| SRR1013055 |
| SRR1013316 |
| SRR1013218 |
| SRR1013239 |
| SRR567670 |
| SRR1013066 |
| SRR1013176 |
| SRR567676 |
| SRR1013100 |
| SRR1013319 |
| SRR1013234 |
| SRR1013267 |
| SRR1013080 |
| SRR1013185 |
| SRR1013098 |
| SRR567671 |
| SRR1013300 |
| SRR1013078 |
| SRR1013221 |
| SRR1013141 |
| SRR1013238 |
| SRR1013083 |
| SRR1013275 |
| SRR567691 |
| SRR1013249 |
| SRR1013330 |
| SRR863049 |
| SRR1013091 |
| SRR567692 |
| SRR567672 |
| SRR1013231 |
| SRR1013106 |
| SRR1013294 |
| SRR1013149 |
| SRR1013190 |
| SRR1013107 |
| SRR1013208 |
| SRR567681 |
| SRR1013242 |
| SRR1013134 |
| SRR1013144 |

| |
|---|
| SRR1013212 |
| SRR567683 |
| SRR1013198 |
| SRR1013059 |
| SRR1013233 |
| SRR1013332 |
| SRR1013058 |
| SRR567687 |
| SRR1013118 |
| SRR567684 |
| SRR1013310 |
| SRR1013268 |
| SRR1013043 |
| SRR1013225 |
| SRR567688 |
| SRR567682 |
| SRR1013166 |
| SRR1013308 |
| SRR1013182 |
| SRR1013143 |
| SRR1013280 |
| SRR567661 |
| SRR1013304 |
| SRR1013194 |
| SRR1013077 |
| SRR567659 |
| SRR1013220 |
| SRR1013162 |
| SRR567662 |
| SRR1013119 |
| SRR1013138 |
| SRR1013095 |
| SRR1013154 |
| SRR567660 |
| SRR1013292 |
| SRR1013164 |
| SRR1013160 |
| SRR1013214 |
| SRR1013240 |
| SRR1013228 |
| SRR1013219 |
| SRR1013273 |
| SRR1013251 |

| |
|---|
| SRR567685 |
| SRR567673 |
| SRR1013216 |
| SRR1013222 |
| SRR567686 |
| SRR1013317 |
| SRR1013229 |
| SRR1013259 |
| SRR1013179 |
| SRR1013079 |
| SRR1013099 |
| SRR567677 |
| SRR1013284 |
| SRR1013291 |
| SRR567674 |
| SRR1013180 |
| SRR1013153 |
| SRR1013320 |
| SRR1013117 |
| SRR1013204 |
| SRR567678 |
| SRR863051 |
| SRR863053 |
| SRR1013057 |
| SRR1013103 |
| SRR1013074 |
| SRR1013302 |
| SRR1013140 |
| SRR1013093 |
| SRR1013126 |
| SRR1013230 |
| SRR1013333 |
| SRR1013146 |
| SRR863056 |
| SRR1013217 |
| SRR1013192 |
| SRR1013137 |
| SRR1013035 |
| SRR1013271 |
| SRR1013073 |
| SRR1013253 |
| SRR1013196 |
| SRR1013206 |

| |
|---|
| SRR1013040 |
| SRR1013050 |
| SRR1013288 |
| SRR1013123 |
| SRR1013056 |
| SRR1013097 |
| SRR1013314 |
| SRR1013167 |
| SRR1013322 |
| SRR1013037 |
| SRR1013324 |
| SRR1013257 |
| SRR1013051 |
| SRR1013210 |
| SRR1013178 |
| SRR1013286 |
| SRR1013296 |
| SRR1264702 |
| ERR1893565 |
| ERR1910500 |
| ERR1910501 |
| ERR1910502 |
| ERR1910503 |
| ERR2003411 |
| ERR2003412 |
| ERR2003413 |
| ERR2003414 |
| ERR2003415 |
| ERR2003420 |
| ERR2003421 |
| ERR2003422 |
| ERR2003423 |
| ERR2003424 |
| ERR2003425 |
| ERR2003426 |
| ERR2003427 |
| ERR2003431 |
| ERR2003432 |
| ERR2003433 |
| ERR2003434 |
| ERR2003435 |
| ERR2003436 |
| SRR5166984 |

| |
|---|
| SRR5436196 |
| SRR5436211 |
| SRR5436212 |
| SRR5444986 |
| SRR5444987 |
| SRR5444988 |
| SRR5444989 |
| SRR5452442 |
| SRR5452443 |
| SRR5452444 |
| SRR5452445 |
| SRR5452451 |
| SRR5452452 |
| SRR5483420 |
| SRR5483421 |
| SRR5483422 |
| SRR5483423 |
| SRR5483424 |
| SRR5483425 |
| SRR5520670 |
| SRR5520671 |
| SRR5520672 |
| SRR5520673 |
| SRR5520674 |
| SRR5520675 |
| SRR5520676 |
| SRR5520677 |
| SRR5520678 |
| SRR5520679 |
| SRR5520680 |
| SRR5520681 |
| SRR5581484 |
| SRR5581485 |
| SRR5724242 |
| SRR5724243 |
| SRR5724244 |
| SRR5724247 |
| SRR5724248 |
| SRR5724265 |
| SRR5724266 |
| SRR5724327 |
| SRR5724328 |
| SRR5724329 |

| |
|---|
| SRR5724330 |
| SRR5724331 |
| SRR5724332 |
| SRR5724333 |
| SRR5724334 |
| SRR5724337 |
| SRR5724338 |
| SRR5724339 |
| SRR5724340 |
| SRR5724345 |
| SRR5724347 |
| SRR5868455 |
| SRR5868456 |
| SRR5868457 |
| SRR5868458 |
| SRR5868459 |
| SRR5868460 |
| SRR5868461 |
| SRR5868462 |
| SRR5868463 |
| SRR5868464 |
| SRR5868465 |
| SRR5868466 |
| SRR5868467 |
| SRR5868468 |
| SRR5868469 |
| SRR5868470 |
| SRR5868471 |
| SRR5868472 |
| SRR5868473 |
| SRR5868474 |
| SRR5868475 |
| SRR5868476 |
| SRR5868477 |
| SRR5868478 |
| SRR5868479 |
| SRR5868480 |
| SRR5868481 |
| SRR5868482 |
| SRR5868483 |
| SRR5868484 |
| SRR5868485 |
| SRR5868486 |

| |
|---|
| SRR5868487 |
| SRR5868488 |
| SRR5868489 |
| SRR5932903 |
| SRR5932904 |
| SRR5932905 |
| SRR5932906 |
| SRR5932907 |
| SRR5932909 |
| SRR5932996 |
| SRR5932997 |
| SRR5932998 |
| SRR5932999 |
| SRR5933000 |
| SRR5933001 |
| SRR5933002 |
| SRR5933003 |
| SRR5933004 |
| SRR5933005 |
| SRR5933006 |
| SRR5933007 |
| SRR5933008 |
| SRR5933009 |
| SRR5933011 |
| SRR5933035 |
| SRR5933037 |
| SRR5933038 |
| SRR5933039 |
| SRR5933040 |
| SRR5933041 |
| SRR5933042 |
| SRR5933043 |
| SRR5933044 |
| SRR5933045 |
| SRR5933046 |
| SRR5933047 |
| SRR5933048 |
| SRR5933049 |
