## Supplementary Table 4 for "An improved de novo assembly and annotation of the tomato reference genome using single-molecule sequencing, Hi-C proximity ligation and optical maps"

### Supplementary files

Supplementary Table 1: Completeness analysis of different genome builds using the BUSCO tool and the Embryophyta and Solanaceae datasets.

| BUSCO | Embryophyta (1,375 BUSCOs) | Solanaceae (3,052 BUSCOs) |
|---|---|---|
| SL4.0 genome | 97.5% | 95.8% |
| SL3.0 genome | 97.7% | 96.5% |
| SL2.5 genome | 97.8% | 96.4% |
| ITAG4.0 annotation | 95.7% | 94.8% |
| ITAG2.4 annotation | 97.6% | 97.1% |

Supplementary Table 2: DNA-seq and RNA-seq mapping rates for different genome builds. DNA-seq paired-end reads were aligned with bowtie2. RNA-seq paired-end reads were aligned with hisat2.

| Genome build | DNA-seq mapping rate | RNA-seq mapping rate |
|---|---|---|
| SL4.0 | 92.70% | 94.45% |
| SL3.0 | 89.68% | 94.41% |
| SL2.5 | 89.89% | 94.46% |

Supplementary Figure 1: Alignment of Bionano cmaps to chromosome 12 in the IrysView tool shows the inversions in scaffolds SL2.50sc04039 and SL2.50sc04057. These were corrected in SL3.0 build.

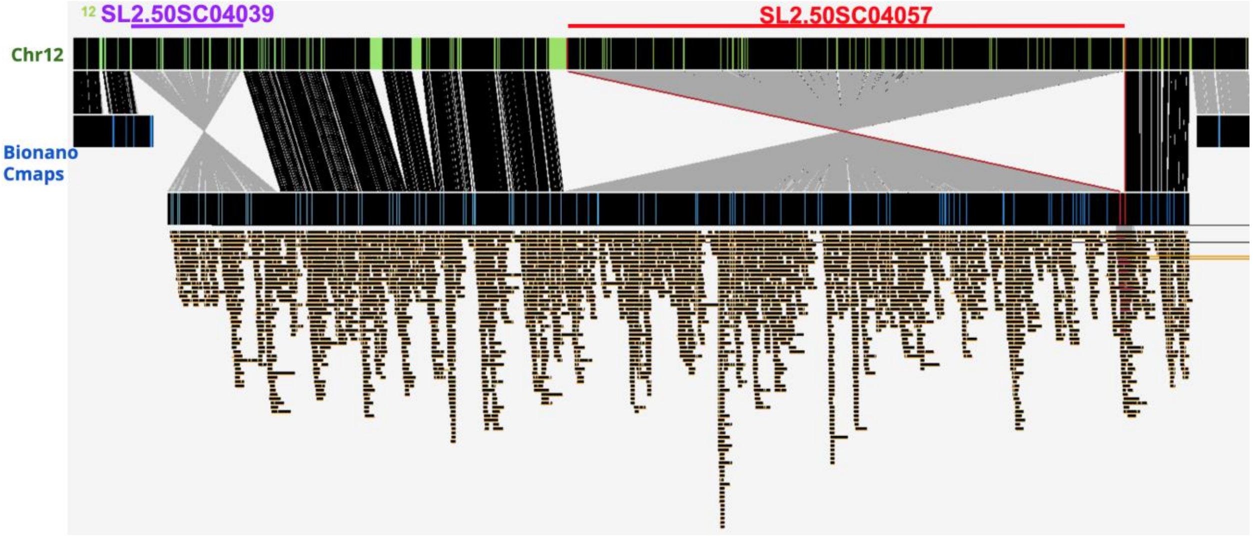


Supplementary Figure 2: K-mer spectra for SL4.0 genome assembly using 20x paired-end Illumina reads.

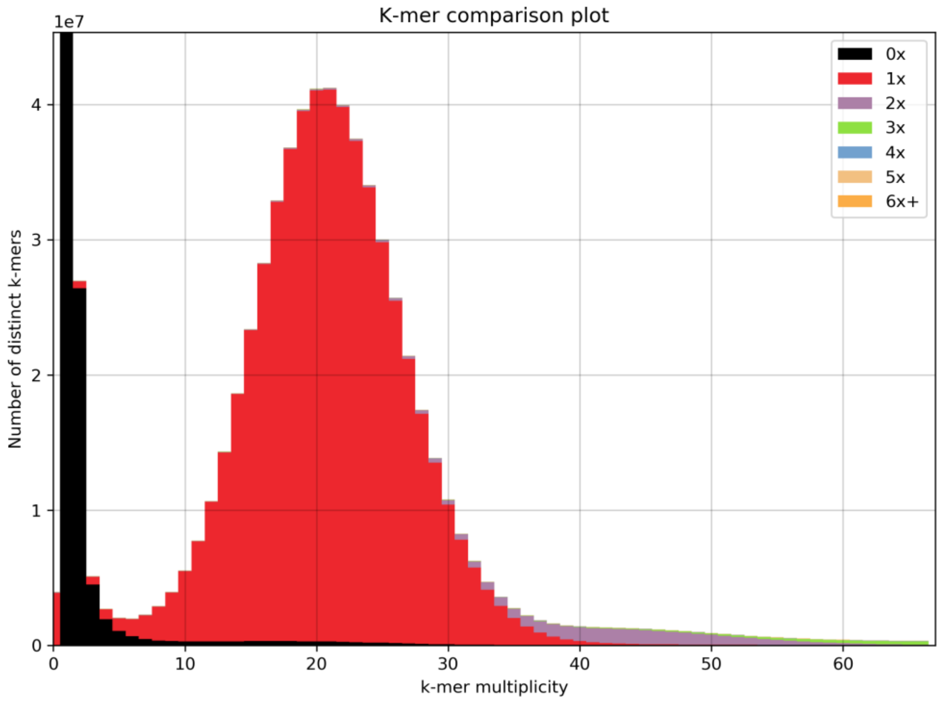

Supplementary Figure 3: Expanded and contracted gene families for 11 species. Green numbers refer to number of expanded gene families and red numbers refer to number of contracted gene families.

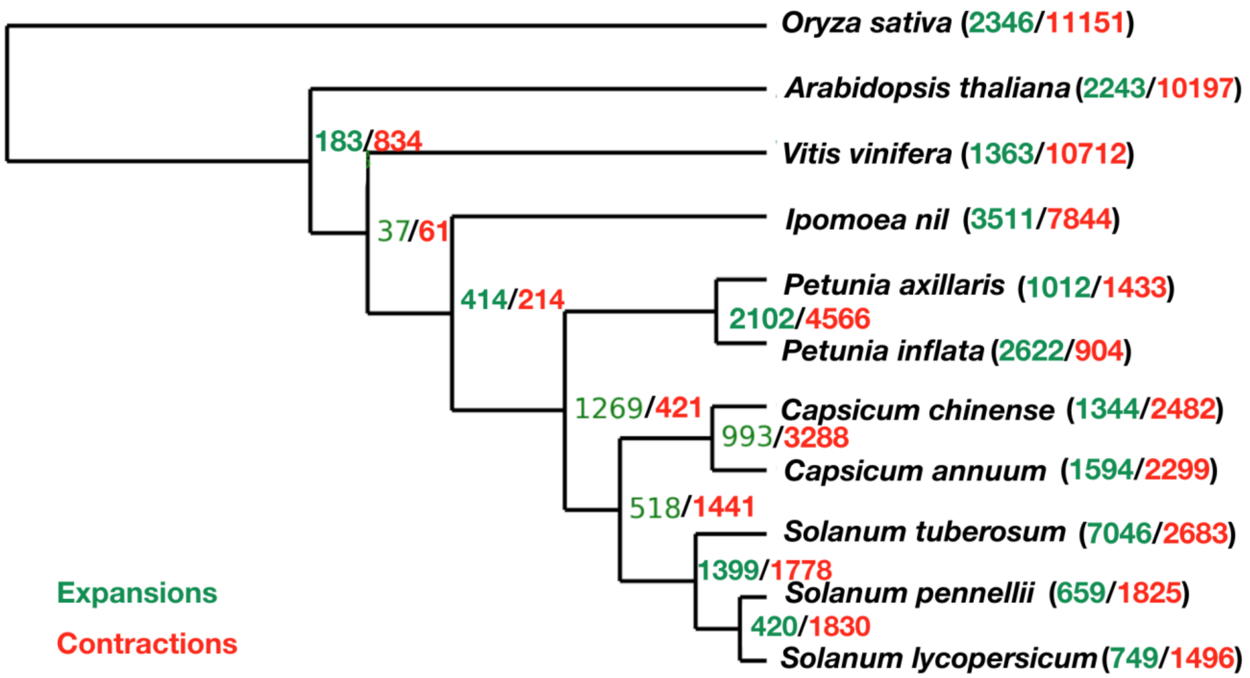
