## Supplementary Table 5 for "An improved de novo assembly and annotation of the tomato reference genome using single-molecule sequencing, Hi-C proximity ligation and optical maps"

| **Supplementary Table 5:** List of updated genes in ITAG4.0 | |
|---|---|
| **ITAG2.4 genes** | **ITAG4.0 updated genes** |
| Solyc01g005000.2 | Solyc01g005000.3 |
| Solyc01g005020.2 | Solyc01g005020.3 |
| Solyc01g005040.2 | Solyc01g005040.3 |
| Solyc01g005060.2 | Solyc01g005060.3 |
| Solyc01g005080.2 | Solyc01g005080.3 |
| Solyc01g005090.2 | Solyc01g005090.3 |
| Solyc01g005110.2 | Solyc01g005110.3 |
| Solyc01g005120.2 | Solyc01g005120.3 |
| Solyc01g005140.2 | Solyc01g005140.3 |
| Solyc01g005150.2 | Solyc01g005150.3 |
| Solyc01g005220.2 | Solyc01g005220.3 |
| Solyc01g005280.2 | Solyc01g005280.3 |
| Solyc01g005310.2 | Solyc01g005310.3 |
| Solyc01g005330.2 | Solyc01g005330.3 |
| Solyc01g005340.2 | Solyc01g005340.3 |
| Solyc01g005370.2 | Solyc01g005370.3 |
| Solyc01g005380.2 | Solyc01g005380.3 |
| Solyc01g005390.2 | Solyc01g005390.3 |
| Solyc01g005400.2 | Solyc01g005400.3 |
| Solyc01g005430.2 | Solyc01g005430.3 |
| Solyc01g005450.2 | Solyc01g005450.3 |
| Solyc01g005470.2 | Solyc01g005470.3 |
| Solyc01g005490.2 | Solyc01g005490.3 |
| Solyc01g005500.2 | Solyc01g005500.3 |
| Solyc01g005510.2 | Solyc01g005510.3 |
| Solyc01g005550.2 | Solyc01g005550.3 |
| Solyc01g005560.2 | Solyc01g005560.3 |
| Solyc01g005620.2 | Solyc01g005620.3 |
| Solyc01g005670.2 | Solyc01g005670.3 |
| Solyc01g005690.2 | Solyc01g005690.3 |
| Solyc01g005700.2 | Solyc01g005700.3 |
| Solyc01g005720.2 | Solyc01g005720.3 |
| Solyc01g005730.2 | Solyc01g005730.3 |
| Solyc01g005740.2 | Solyc01g005740.3 |
| Solyc01g005760.2 | Solyc01g005760.3 |
| Solyc01g005770.2 | Solyc01g005770.3 |
| Solyc01g005775.1 | Solyc01g150100.1 |
| Solyc01g005800.2 | Solyc01g005800.3 |
| Solyc01g005810.2 | Solyc01g005810.3 |
| Solyc01g005840.2 | Solyc01g005840.3 |
| Solyc01g005850.1 | Solyc01g005850.2 |

| | |
|---|---|
| Solyc01g005910.2 | Solyc01g005910.3 |
| Solyc01g005920.2 | Solyc01g005920.3 |
| Solyc01g005960.2 | Solyc01g005960.3 |
| Solyc01g005970.2 | Solyc01g005970.3 |
| Solyc01g006000.2 | Solyc01g006000.3 |
| Solyc01g006010.2 | Solyc01g006010.3 |
| Solyc01g006040.2 | Solyc01g006040.3 |
| Solyc01g006080.1 | Solyc01g006080.2 |
| Solyc01g006090.2 | Solyc01g006090.3 |
| Solyc01g006110.2 | Solyc01g006110.3 |
| Solyc01g006150.2 | Solyc01g006150.3 |
| Solyc01g006190.2 | Solyc01g006190.3 |
| Solyc01g006200.2 | Solyc01g006200.3 |
| Solyc01g006220.2 | Solyc01g006220.3 |
| Solyc01g006240.2 | Solyc01g006240.3 |
| Solyc01g006250.1 | Solyc01g006250.2 |
| Solyc01g006280.2 | Solyc01g006280.3 |
| Solyc01g006300.2 | Solyc01g006300.3 |
| Solyc01g006310.2 | Solyc01g006310.3 |
| Solyc01g006340.2 | Solyc01g006340.3 |
| Solyc01g006370.2 | Solyc01g006370.3 |
| Solyc01g006420.2 | Solyc01g006420.3 |
| Solyc01g006430.2 | Solyc01g006430.3 |
| Solyc01g006440.1 | Solyc01g006440.2 |
| Solyc01g006480.2 | Solyc01g006480.3 |
| Solyc01g006510.2 | Solyc01g006510.3 |
| Solyc01g006520.2 | Solyc01g006520.3 |
| Solyc01g006550.2 | Solyc01g006550.3 |
| Solyc01g006610.1 | Solyc01g006610.2 |
| Solyc01g006620.2 | Solyc01g006620.3 |
| Solyc01g006630.2 | Solyc01g006630.3 |
| Solyc01g006650.1 | Solyc01g006650.2 |
| Solyc01g006760.2 | Solyc01g006760.3 |
| Solyc01g006770.1 | Solyc01g006770.2 |
| Solyc01g006830.2 | Solyc01g006830.3 |
| Solyc01g006840.2 | Solyc01g006840.3 |
| Solyc01g006860.2 | Solyc01g006860.3 |
| Solyc01g006890.2 | Solyc01g006890.3 |
| Solyc01g006900.2 | Solyc01g006900.3 |
| Solyc01g006930.2 | Solyc01g006930.3 |
| Solyc01g006950.2 | Solyc01g006950.3 |
| Solyc01g006960.2 | Solyc01g006960.3 |
| Solyc01g007030.2 | Solyc01g007030.3 |

| | |
|---|---|
| Solyc01g007060.2 | Solyc01g007060.3 |
| Solyc01g007070.2 | Solyc01g007070.3 |
| Solyc01g007080.2 | Solyc01g007080.3 |
| Solyc01g007100.2 | Solyc01g007100.3 |
| Solyc01g007110.2 | Solyc01g007110.3 |
| Solyc01g007120.2 | Solyc01g007120.3 |
| Solyc01g007130.2 | Solyc01g007130.3 |
| Solyc01g007150.2 | Solyc01g007150.3 |
| Solyc01g007180.2 | Solyc01g007180.3 |
| Solyc01g007190.2 | Solyc01g007190.3 |
| Solyc01g007210.2 | Solyc01g007210.3 |
| Solyc01g007240.2 | Solyc01g007240.3 |
| Solyc01g007260.2 | Solyc01g007260.3 |
| Solyc01g007750.2 | Solyc01g007750.3 |
| Solyc01g007760.2 | Solyc01g007760.3 |
| Solyc01g007800.2 | Solyc01g007800.3 |
| Solyc01g007830.1 | Solyc01g007830.2 |
| Solyc01g007840.2 | Solyc01g007840.3 |
| Solyc01g007860.2 | Solyc01g007860.3 |
| Solyc01g007895.1 | Solyc01g150101.1 |
| Solyc01g007910.2 | Solyc01g007910.3 |
| Solyc01g007960.2 | Solyc01g007960.3 |
| Solyc01g007970.1 | Solyc01g007970.2 |
| Solyc01g008000.2 | Solyc01g008000.3 |
| Solyc01g008005.1 | Solyc01g150102.1 |
| Solyc01g008010.2 | Solyc01g008010.3 |
| Solyc01g008060.2 | Solyc01g008060.3 |
| Solyc01g008070.2 | Solyc01g008070.3 |
| Solyc01g008080.2 | Solyc01g008080.3 |
| Solyc01g008100.2 | Solyc01g008100.3 |
| Solyc01g008170.2 | Solyc01g008170.3 |
| Solyc01g008180.2 | Solyc01g008180.3 |
| Solyc01g008200.2 | Solyc01g008200.3 |
| Solyc01g008220.2 | Solyc01g008220.3 |
| Solyc01g008250.2 | Solyc01g008250.3 |
| Solyc01g008270.2 | Solyc01g008270.3 |
| Solyc01g008290.2 | Solyc01g008290.3 |
| Solyc01g008300.1 | Solyc01g008300.2 |
| Solyc01g008320.2 | Solyc01g008320.3 |
| Solyc01g008340.2 | Solyc01g008340.3 |
| Solyc01g008360.2 | Solyc01g008360.3 |
| Solyc01g008370.2 | Solyc01g008370.3 |
| Solyc01g008400.2 | Solyc01g008400.3 |

| | |
|---|---|
| Solyc01g008420.2 | Solyc01g008420.3 |
| Solyc01g008440.2 | Solyc01g008440.3 |
| Solyc01g008460.2 | Solyc01g008460.3 |
| Solyc01g008470.2 | Solyc01g008470.3 |
| Solyc01g008471.1 | Solyc01g150103.1 |
| Solyc01g008472.1 | Solyc01g150104.1 |
| Solyc01g008474.1 | Solyc01g150105.1 |
| Solyc01g008476.1 | Solyc01g150106.1 |
| Solyc01g008478.1 | Solyc01g150107.1 |
| Solyc01g008510.2 | Solyc01g008510.3 |
| Solyc01g008520.2 | Solyc01g008520.3 |
| Solyc01g008540.3 | Solyc01g008540.4 |
| Solyc01g008560.2 | Solyc01g008560.3 |
| Solyc01g008650.2 | Solyc01g008650.3 |
| Solyc01g008690.2 | Solyc01g008690.3 |
| Solyc01g008710.4 | Solyc01g008710.5.1 |
| Solyc01g008730.2 | Solyc01g008730.3 |
| Solyc01g008740.1 | Solyc01g008740.2 |
| Solyc01g008770.2 | Solyc01g008770.3 |
| Solyc01g008820.2 | Solyc01g008820.3 |
| Solyc01g008830.1 | Solyc01g008830.2 |
| Solyc01g008840.2 | Solyc01g008840.3 |
| Solyc01g008850.2 | Solyc01g008850.3 |
| Solyc01g008960.2 | Solyc01g008960.3 |
| Solyc01g008980.2 | Solyc01g008980.3 |
| Solyc01g009000.2 | Solyc01g009000.3 |
| Solyc01g009010.2 | Solyc01g009010.3 |
| Solyc01g009050.2 | Solyc01g009050.3 |
| Solyc01g009060.2 | Solyc01g009060.3 |
| Solyc01g009070.2 | Solyc01g009070.3 |
| Solyc01g009090.2 | Solyc01g009090.3 |
| Solyc01g009120.2 | Solyc01g009120.3 |
| Solyc01g009210.1 | Solyc01g009210.2 |
| Solyc01g009220.2 | Solyc01g009220.3 |
| Solyc01g009300.2 | Solyc01g009300.3 |
| Solyc01g009310.2 | Solyc01g009310.3 |
| Solyc01g009320.1 | Solyc01g009320.2 |
| Solyc01g009370.1 | Solyc01g009370.2 |
| Solyc01g009380.2 | Solyc01g009380.3 |
| Solyc01g009400.2 | Solyc01g009400.3 |
| Solyc01g009410.2 | Solyc01g009410.3 |
| Solyc01g009470.1 | Solyc01g009470.2 |
| Solyc01g009560.1 | Solyc01g009560.2 |

| | |
|---|---|
| Solyc01g009580.1 | Solyc01g009580.2 |
| Solyc01g009590.2 | Solyc01g009590.3 |
| Solyc01g009610.2 | Solyc01g009610.3 |
| Solyc01g009620.2 | Solyc01g009620.3 |
| Solyc01g009630.2 | Solyc01g009630.3 |
| Solyc01g009720.1 | Solyc01g009720.2 |
| Solyc01g009750.2 | Solyc01g009750.3 |
| Solyc01g009760.2 | Solyc01g009760.3 |
| Solyc01g009770.2 | Solyc01g009770.3 |
| Solyc01g009780.2 | Solyc01g009780.3 |
| Solyc01g009810.2 | Solyc01g009810.3 |
| Solyc01g009830.2 | Solyc01g009830.3 |
| Solyc01g009860.2 | Solyc01g009860.3 |
| Solyc01g009870.2 | Solyc01g009870.3 |
| Solyc01g009900.2 | Solyc01g009900.3 |
| Solyc01g009970.2 | Solyc01g009970.3 |
| Solyc01g009990.2 | Solyc01g009990.3 |
| Solyc01g010000.2 | Solyc01g010000.3 |
| Solyc01g010020.2 | Solyc01g010020.3 |
| Solyc01g010030.2 | Solyc01g010030.3 |
| Solyc01g010050.2 | Solyc01g010050.3 |
| Solyc01g010080.2 | Solyc01g010080.3 |
| Solyc01g010083.1 | Solyc01g150108.1 |
| Solyc01g010085.1 | Solyc01g150109.1 |
| Solyc01g010087.1 | Solyc01g150110.1 |
| Solyc01g010150.2 | Solyc01g010150.3 |
| Solyc01g010160.2 | Solyc01g010160.3 |
| Solyc01g010170.1 | Solyc01g010170.2 |
| Solyc01g010180.2 | Solyc01g010180.3 |
| Solyc01g010220.2 | Solyc01g010220.3 |
| Solyc01g010240.2 | Solyc01g010240.3 |
| Solyc01g010260.2 | Solyc01g010260.3 |
| Solyc01g010270.2 | Solyc01g010270.3 |
| Solyc01g010360.2 | Solyc01g010360.3 |
| Solyc01g010370.2 | Solyc01g010370.3 |
| Solyc01g010390.2 | Solyc01g010390.3 |
| Solyc01g010410.2 | Solyc01g010410.3 |
| Solyc01g010420.2 | Solyc01g010420.3 |
| Solyc01g010480.2 | Solyc01g010480.3 |
| Solyc01g010493.1 | Solyc01g150111.1 |
| Solyc01g010497.1 | Solyc01g150112.1 |
| Solyc01g010520.2 | Solyc01g010520.3 |
| Solyc01g010540.2 | Solyc01g010540.3 |

| | |
|---|---|
| Solyc01g010580.2 | Solyc01g010580.3 |
| Solyc01g010640.2 | Solyc01g010640.3 |
| Solyc01g010650.2 | Solyc01g010650.3 |
| Solyc01g010660.2 | Solyc01g010660.3 |
| Solyc01g010700.2 | Solyc01g010700.3 |
| Solyc01g010760.2 | Solyc01g010760.3 |
| Solyc01g010820.2 | Solyc01g010820.3 |
| Solyc01g010860.2 | Solyc01g010860.3 |
| Solyc01g010870.2 | Solyc01g010870.3 |
| Solyc01g010880.2 | Solyc01g010880.3 |
| Solyc01g010900.2 | Solyc01g010900.3 |
| Solyc01g010910.1 | Solyc01g010910.2 |
| Solyc01g010920.1 | Solyc01g010920.2 |
| Solyc01g010940.2 | Solyc01g010940.3 |
| Solyc01g010950.2 | Solyc01g010950.3 |
| Solyc01g010970.2 | Solyc01g010970.3 |
| Solyc01g011000.2 | Solyc01g011000.3 |
| Solyc01g011020.1 | Solyc01g011020.2 |
| Solyc01g011030.2 | Solyc01g011030.3 |
| Solyc01g011040.2 | Solyc01g011040.3 |
| Solyc01g011050.2 | Solyc01g011050.3 |
| Solyc01g011080.2 | Solyc01g011080.3 |
| Solyc01g011175.1 | Solyc01g150113.1 |
| Solyc01g011200.2 | Solyc01g011200.3 |
| Solyc01g011230.2 | Solyc01g011230.3 |
| Solyc01g011270.1 | Solyc01g011270.2 |
| Solyc01g011310.1 | Solyc01g011310.2 |
| Solyc01g011320.1 | Solyc01g011320.2 |
| Solyc01g011337.1 | Solyc01g150115.1 |
| Solyc01g011340.2 | Solyc01g011340.3 |
| Solyc01g011350.2 | Solyc01g011350.3 |
| Solyc01g011410.1 | Solyc01g011410.2 |
| Solyc01g011520.1 | Solyc01g011520.2 |
| Solyc01g012570.1 | Solyc01g012570.2 |
| Solyc01g013950.1 | Solyc01g013950.2 |
| Solyc01g013995.1 | Solyc01g013995.2 |
| Solyc01g014065.1 | Solyc01g150116.1 |
| Solyc01g014100.2 | Solyc01g014100.3 |
| Solyc01g014130.1 | Solyc01g014130.2 |
| Solyc01g014140.2 | Solyc01g014140.3 |
| Solyc01g014180.2 | Solyc01g014180.3 |
| Solyc01g014270.2 | Solyc01g014270.3 |
| Solyc01g014310.1 | Solyc01g014310.2 |

| | |
|---|---|
| Solyc01g014370.1 | Solyc01g014370.2 |
| Solyc01g014390.2 | Solyc01g014390.3 |
| Solyc01g014400.1 | Solyc01g014400.2 |
| Solyc01g014440.1 | Solyc01g014440.2 |
| Solyc01g014480.2 | Solyc01g014480.3 |
| Solyc01g014510.1 | Solyc01g014510.2 |
| Solyc01g014537.1 | Solyc01g150117.1 |
| Solyc01g014725.1 | Solyc01g014725.2 |
| Solyc01g014730.1 | Solyc01g014730.2 |
| Solyc01g014820.1 | Solyc01g014820.2 |
| Solyc01g014840.2 | Solyc01g014840.3 |
| Solyc01g014860.2 | Solyc01g014860.3 |
| Solyc01g014910.1 | Solyc01g014910.2 |
| Solyc01g014930.1 | Solyc01g014930.2 |
| Solyc01g014940.1 | Solyc01g014940.2 |
| Solyc01g015000.2 | Solyc01g015000.3 |
| Solyc01g015080.2 | Solyc01g015080.3 |
| Solyc01g015085.1 | Solyc01g150118.1 |
| Solyc01g016470.2 | Solyc01g016470.3 |
| Solyc01g016550.1 | Solyc01g016550.2 |
| Solyc01g016600.1 | Solyc01g016600.2 |
| Solyc01g016645.1 | Solyc01g150119.1 |
| Solyc01g016670.1 | Solyc01g016670.2 |
| Solyc01g016700.2 | Solyc01g016700.3 |
| Solyc01g016800.1 | Solyc01g016800.2 |
| Solyc01g016805.1 | Solyc01g150120.1 |
| Solyc01g016850.1 | Solyc01g016850.2 |
| Solyc01g016940.1 | Solyc01g016940.2 |
| Solyc01g016970.1 | Solyc01g016970.2 |
| Solyc01g016985.1 | Solyc01g150121.1 |
| Solyc01g017060.1 | Solyc01g017060.2 |
| Solyc01g017080.1 | Solyc01g017080.2 |
| Solyc01g017100.1 | Solyc01g017100.2 |
| Solyc01g017110.1 | Solyc01g017110.2 |
| Solyc01g017160.1 | Solyc01g017160.2 |
| Solyc01g017200.1 | Solyc01g017200.2 |
| Solyc01g017220.1 | Solyc01g017220.2 |
| Solyc01g017230.1 | Solyc01g017230.2 |
| Solyc01g017300.1 | Solyc01g017300.2 |
| Solyc01g017370.1 | Solyc01g017370.2 |
| Solyc01g017540.1 | Solyc01g017540.2 |
| Solyc01g017600.2 | Solyc01g017600.3 |
| Solyc01g017670.1 | Solyc01g017670.2 |

| | |
|---|---|
| Solyc01g017720.1 | Solyc01g017720.2 |
| Solyc01g017740.1 | Solyc01g017740.2 |
| Solyc01g017793.1 | Solyc01g150122.1 |
| Solyc01g017797.1 | Solyc01g150123.1 |
| Solyc01g017860.2 | Solyc01g017860.3 |
| Solyc01g017870.1 | Solyc01g017870.2 |
| Solyc01g018010.1 | Solyc01g018010.2 |
| Solyc01g020110.1 | Solyc01g020110.2 |
| Solyc01g020130.2 | Solyc01g020130.3 |
| Solyc01g020150.2 | Solyc01g020150.3 |
| Solyc01g020170.1 | Solyc01g020170.2 |
| Solyc01g020280.1 | Solyc01g020280.2 |
| Solyc01g020285.1 | Solyc01g150124.1 |
| Solyc01g020340.1 | Solyc01g020340.2 |
| Solyc01g020370.1 | Solyc01g020370.2 |
| Solyc01g020374.1 | Solyc01g150125.1 |
| Solyc01g020376.1 | Solyc01g150126.1 |
| Solyc01g020378.1 | Solyc01g150127.1 |
| Solyc01g020430.1 | Solyc01g020430.2 |
| Solyc01g020440.2 | Solyc01g020440.3 |
| Solyc01g020445.1 | Solyc01g150128.1 |
| Solyc01g020500.1 | Solyc01g020500.2 |
| Solyc01g022740.2 | Solyc01g022740.3 |
| Solyc01g022750.2 | Solyc01g022750.3 |
| Solyc01g034010.1 | Solyc01g034010.2 |
| Solyc01g034020.2 | Solyc01g034020.3 |
| Solyc01g034155.1 | Solyc01g150129.1 |
| Solyc01g028910.1 | Solyc01g028910.2 |
| Solyc01g028900.2 | Solyc01g028900.3 |
| Solyc01g028870.2 | Solyc01g028870.3 |
| Solyc01g028860.2 | Solyc01g028860.3 |
| Solyc01g028830.2 | Solyc01g028830.3 |
| Solyc01g028810.2 | Solyc01g028810.3 |
| Solyc01g044270.2 | Solyc01g044270.3 |
| Solyc01g044340.2 | Solyc01g044340.3 |
| Solyc01g044350.2 | Solyc01g044350.3 |
| Solyc01g044520.1 | Solyc01g044520.2 |
| Solyc01g049650.2 | Solyc01g049650.3 |
| Solyc01g049660.1 | Solyc01g049660.2 |
| Solyc01g049680.2 | Solyc01g049680.3 |
| Solyc01g049770.2 | Solyc01g049770.3 |
| Solyc01g049845.1 | Solyc01g150130.1 |
| Solyc01g049880.2 | Solyc01g049880.3 |

| | |
|---|---|
| Solyc01g049890.2 | Solyc01g049890.3 |
| Solyc01g049920.1 | Solyc01g049920.2 |
| Solyc01g049925.1 | Solyc01g150131.1 |
| Solyc01g050010.2 | Solyc01g050010.3 |
| Solyc01g056280.2 | Solyc01g056280.3 |
| Solyc01g056310.2 | Solyc01g056310.3 |
| Solyc01g056330.1 | Solyc01g056330.2 |
| Solyc01g056370.2 | Solyc01g056370.3 |
| Solyc01g056430.1 | Solyc01g056430.2 |
| Solyc01g056480.2 | Solyc01g056480.3 |
| Solyc01g056485.1 | Solyc01g150132.1 |
| Solyc01g056530.1 | Solyc01g056530.2 |
| Solyc01g056570.2 | Solyc01g056570.3 |
| Solyc01g056655.1 | Solyc01g150133.1 |
| Solyc01g056660.1 | Solyc01g056660.2 |
| Solyc01g056760.2 | Solyc01g056760.3 |
| Solyc01g056780.2 | Solyc01g056780.3 |
| Solyc01g056790.2 | Solyc01g056790.3 |
| Solyc01g056830.2 | Solyc01g056830.3 |
| Solyc01g056850.2 | Solyc01g056850.3 |
| Solyc01g056885.1 | Solyc01g150134.1 |
| Solyc01g056940.3 | Solyc01g056940.4 |
| Solyc01g057000.2 | Solyc01g057000.3 |
| Solyc01g057020.2 | Solyc01g057020.3 |
| Solyc01g057120.1 | Solyc01g057120.2 |
| Solyc01g057180.1 | Solyc01g057180.2 |
| Solyc01g057210.2 | Solyc01g057210.3 |
| Solyc01g057220.2 | Solyc01g057220.3 |
| Solyc01g057250.1 | Solyc01g057250.2 |
| Solyc01g057300.1 | Solyc01g057300.2 |
| Solyc01g057340.2 | Solyc01g057340.3 |
| Solyc01g057355.1 | Solyc01g150135.1 |
| Solyc01g057430.2 | Solyc01g057430.3 |
| Solyc01g057440.2 | Solyc01g057440.3 |
| Solyc01g057510.1 | Solyc01g057510.2 |
| Solyc01g057535.1 | Solyc01g150136.1 |
| Solyc01g057570.2 | Solyc01g057570.3 |
| Solyc01g057600.2 | Solyc01g057600.3 |
| Solyc01g057690.1 | Solyc01g057690.2 |
| Solyc01g057770.2 | Solyc01g057770.3 |
| Solyc01g057780.2 | Solyc01g057780.3 |
| Solyc01g057830.2 | Solyc01g057830.3 |
| Solyc01g057840.2 | Solyc01g057840.3 |

| | |
|---|---|
| Solyc01g057900.2 | Solyc01g057900.3 |
| Solyc01g057910.2 | Solyc01g057910.3 |
| Solyc01g058000.2 | Solyc01g058000.3 |
| Solyc01g058010.2 | Solyc01g058010.3 |
| Solyc01g058030.1 | Solyc01g058030.2 |
| Solyc01g058040.1 | Solyc01g058040.2 |
| Solyc01g058140.2 | Solyc01g058140.3 |
| Solyc01g058160.2 | Solyc01g058160.3 |
| Solyc01g058200.2 | Solyc01g058200.3 |
| Solyc01g058250.1 | Solyc01g058250.2 |
| Solyc01g058270.2 | Solyc01g058270.3 |
| Solyc01g058363.1 | Solyc01g150137.1 |
| Solyc01g058367.1 | Solyc01g150138.1 |
| Solyc01g058390.2 | Solyc01g058390.3 |
| Solyc01g058450.2 | Solyc01g058450.3 |
| Solyc01g058540.2 | Solyc01g058540.3 |
| Solyc01g058670.2 | Solyc01g058670.3 |
| Solyc01g058680.1 | Solyc01g058680.2 |
| Solyc01g059830.2 | Solyc01g059830.3 |
| Solyc01g059850.2 | Solyc01g059850.3 |
| Solyc01g059860.2 | Solyc01g059860.3 |
| Solyc01g059880.2 | Solyc01g059880.3 |
| Solyc01g059930.2 | Solyc01g059930.3 |
| Solyc01g060030.2 | Solyc01g060030.3 |
| Solyc01g060040.2 | Solyc01g060040.3 |
| Solyc01g060070.2 | Solyc01g060070.3 |
| Solyc01g060130.2 | Solyc01g060130.3 |
| Solyc01g060215.1 | Solyc01g150139.1 |
| Solyc01g060355.1 | Solyc01g150140.1 |
| Solyc01g060370.1 | Solyc01g060370.2 |
| Solyc01g060460.1 | Solyc01g060460.2 |
| Solyc01g060470.2 | Solyc01g060470.3 |
| Solyc01g065500.2 | Solyc01g065500.3 |
| Solyc01g065510.2 | Solyc01g065510.3 |
| Solyc01g065530.2 | Solyc01g065530.3 |
| Solyc01g065540.2 | Solyc01g065540.3 |
| Solyc01g065550.2 | Solyc01g065550.3 |
| Solyc01g065690.2 | Solyc01g065690.3 |
| Solyc01g065700.2 | Solyc01g065700.3 |
| Solyc01g065710.2 | Solyc01g065710.3 |
| Solyc01g065740.2 | Solyc01g065740.3 |
| Solyc01g065810.1 | Solyc01g065810.2 |
| Solyc01g065960.1 | Solyc01g065960.2 |

| | |
|---|---|
| Solyc01g065985.1 | Solyc01g150141.1 |
| Solyc01g066050.2 | Solyc01g066050.3 |
| Solyc01g066070.2 | Solyc01g066070.3 |
| Solyc01g066100.1 | Solyc01g066100.2 |
| Solyc01g066110.1 | Solyc01g066110.2 |
| Solyc01g066140.1 | Solyc01g066140.2 |
| Solyc01g066203.1 | Solyc01g150142.1 |
| Solyc01g066207.1 | Solyc01g150143.1 |
| Solyc01g066270.2 | Solyc01g066270.3 |
| Solyc01g066360.2 | Solyc01g066360.3 |
| Solyc01g066420.2 | Solyc01g066420.3 |
| Solyc01g066430.2 | Solyc01g066430.3 |
| Solyc01g066450.2 | Solyc01g066450.3 |
| Solyc01g066453.1 | Solyc01g150144.1 |
| Solyc01g066480.2 | Solyc01g066480.3 |
| Solyc01g066520.2 | Solyc01g066520.3 |
| Solyc01g066560.2 | Solyc01g066560.3 |
| Solyc01g066570.2 | Solyc01g066570.3 |
| Solyc01g066590.2 | Solyc01g066590.3 |
| Solyc01g066650.1 | Solyc01g066650.2 |
| Solyc01g066680.2 | Solyc01g066680.3 |
| Solyc01g066690.2 | Solyc01g066690.3 |
| Solyc01g066720.2 | Solyc01g066720.3 |
| Solyc01g066740.2 | Solyc01g066740.3 |
| Solyc01g066770.2 | Solyc01g066770.3 |
| Solyc01g066830.2 | Solyc01g066830.3 |
| Solyc01g066840.2 | Solyc01g066840.3 |
| Solyc01g066865.1 | Solyc01g150145.1 |
| Solyc01g066870.2 | Solyc01g066870.3 |
| Solyc01g066880.2 | Solyc01g066880.3 |
| Solyc01g066890.1 | Solyc01g066890.2 |
| Solyc01g066910.2 | Solyc01g066910.3 |
| Solyc01g066940.2 | Solyc01g066940.3 |
| Solyc01g066980.2 | Solyc01g066980.3 |
| Solyc01g067010.2 | Solyc01g067010.3 |
| Solyc01g067040.1 | Solyc01g067040.2 |
| Solyc01g067070.2 | Solyc01g067070.3 |
| Solyc01g067140.1 | Solyc01g067140.2 |
| Solyc01g067250.2 | Solyc01g067250.3 |
| Solyc01g067290.2 | Solyc01g067290.3 |
| Solyc01g067300.2 | Solyc01g067300.3 |
| Solyc01g067330.2 | Solyc01g067330.3 |
| Solyc01g067360.2 | Solyc01g067360.3 |

| | |
|---|---|
| Solyc01g067370.2 | Solyc01g067370.3 |
| Solyc01g067400.2 | Solyc01g067400.3 |
| Solyc01g067500.2 | Solyc01g067500.3 |
| Solyc01g067510.2 | Solyc01g067510.3 |
| Solyc01g067520.2 | Solyc01g067520.3 |
| Solyc01g067530.2 | Solyc01g067530.3 |
| Solyc01g067540.1 | Solyc01g067540.2 |
| Solyc01g067550.2 | Solyc01g067550.3 |
| Solyc01g067560.2 | Solyc01g067560.3 |
| Solyc01g067570.2 | Solyc01g067570.3 |
| Solyc01g067590.2 | Solyc01g067590.3 |
| Solyc01g067600.2 | Solyc01g067600.3 |
| Solyc01g067620.2 | Solyc01g067620.3 |
| Solyc01g067630.2 | Solyc01g067630.3 |
| Solyc01g067640.2 | Solyc01g067640.3 |
| Solyc01g067650.1 | Solyc01g067650.2 |
| Solyc01g067660.2 | Solyc01g067660.3 |
| Solyc01g067680.2 | Solyc01g067680.3 |
| Solyc01g067700.2 | Solyc01g067700.3 |
| Solyc01g067710.2 | Solyc01g067710.3 |
| Solyc01g067730.2 | Solyc01g067730.3 |
| Solyc01g067740.2 | Solyc01g067740.3 |
| Solyc01g067750.2 | Solyc01g067750.3 |
| Solyc01g067790.2 | Solyc01g067790.3 |
| Solyc01g067800.2 | Solyc01g067800.3 |
| Solyc01g067830.2 | Solyc01g067830.3 |
| Solyc01g067860.2 | Solyc01g067860.3 |
| Solyc01g067870.2 | Solyc01g067870.3 |
| Solyc01g067900.2 | Solyc01g067900.3 |
| Solyc01g067910.2 | Solyc01g067910.3 |
| Solyc01g068010.1 | Solyc01g068010.2 |
| Solyc01g068030.2 | Solyc01g068030.3 |
| Solyc01g068050.2 | Solyc01g068050.3 |
| Solyc01g068150.2 | Solyc01g068150.3 |
| Solyc01g068170.2 | Solyc01g068170.3 |
| Solyc01g068180.2 | Solyc01g068180.3 |
| Solyc01g068210.2 | Solyc01g068210.3 |
| Solyc01g068240.2 | Solyc01g068240.3 |
| Solyc01g068270.2 | Solyc01g068270.3 |
| Solyc01g068280.2 | Solyc01g068280.3 |
| Solyc01g068340.2 | Solyc01g068340.3 |
| Solyc01g068350.2 | Solyc01g068350.3 |
| Solyc01g068380.2 | Solyc01g068380.3 |

| | |
|---|---|
| Solyc01g068390.2 | Solyc01g068390.3 |
| Solyc01g068400.2 | Solyc01g068400.3 |
| Solyc01g068440.1 | Solyc01g068440.2 |
| Solyc01g068530.2 | Solyc01g068530.3 |
| Solyc01g068560.2 | Solyc01g068560.3 |
| Solyc01g068630.2 | Solyc01g068630.3 |
| Solyc01g068640.2 | Solyc01g068640.3 |
| Solyc01g079100.2 | Solyc01g079100.3 |
| Solyc01g079120.2 | Solyc01g079120.3 |
| Solyc01g079130.1 | Solyc01g079130.2 |
| Solyc01g079170.2 | Solyc01g079170.3 |
| Solyc01g079210.2 | Solyc01g079210.3 |
| Solyc01g079240.2 | Solyc01g079240.3 |
| Solyc01g079250.2 | Solyc01g079250.3 |
| Solyc01g079310.2 | Solyc01g079310.3 |
| Solyc01g079330.2 | Solyc01g079330.3 |
| Solyc01g079340.2 | Solyc01g079340.3 |
| Solyc01g079350.2 | Solyc01g079350.3 |
| Solyc01g079400.2 | Solyc01g079400.3 |
| Solyc01g079410.2 | Solyc01g079410.3 |
| Solyc01g079450.2 | Solyc01g079450.3 |
| Solyc01g079480.2 | Solyc01g079480.3 |
| Solyc01g079510.2 | Solyc01g079510.3 |
| Solyc01g079520.2 | Solyc01g079520.3 |
| Solyc01g079560.2 | Solyc01g079560.3 |
| Solyc01g079570.2 | Solyc01g079570.3 |
| Solyc01g079580.2 | Solyc01g079580.3 |
| Solyc01g079590.2 | Solyc01g079590.3 |
| Solyc01g079610.2 | Solyc01g079610.3 |
| Solyc01g079650.2 | Solyc01g079650.3 |
| Solyc01g079680.2 | Solyc01g079680.3 |
| Solyc01g079700.2 | Solyc01g079700.3 |
| Solyc01g079710.2 | Solyc01g079710.3 |
| Solyc01g079720.2 | Solyc01g079720.3 |
| Solyc01g079750.2 | Solyc01g079750.3 |
| Solyc01g079760.2 | Solyc01g079760.3 |
| Solyc01g079780.2 | Solyc01g079780.3 |
| Solyc01g079820.2 | Solyc01g079820.3 |
| Solyc01g079840.2 | Solyc01g079840.3 |
| Solyc01g079850.2 | Solyc01g079850.3 |
| Solyc01g079860.2 | Solyc01g079860.3 |
| Solyc01g079880.2 | Solyc01g079880.3 |
| Solyc01g079890.2 | Solyc01g079890.3 |

| | |
|---|---|
| Solyc01g079920.2 | Solyc01g079920.3 |
| Solyc01g079950.2 | Solyc01g079950.3 |
| Solyc01g079980.2 | Solyc01g079980.3 |
| Solyc01g079990.1 | Solyc01g079990.2 |
| Solyc01g080040.2 | Solyc01g080040.3 |
| Solyc01g080080.2 | Solyc01g080080.3 |
| Solyc01g080110.2 | Solyc01g080110.3 |
| Solyc01g080140.2 | Solyc01g080140.3 |
| Solyc01g080150.2 | Solyc01g080150.3 |
| Solyc01g080170.2 | Solyc01g080170.3 |
| Solyc01g080180.2 | Solyc01g080180.3 |
| Solyc01g080190.2 | Solyc01g080190.3 |
| Solyc01g080200.2 | Solyc01g080200.3 |
| Solyc01g080210.2 | Solyc01g080210.3 |
| Solyc01g080220.2 | Solyc01g080220.3 |
| Solyc01g080230.2 | Solyc01g080230.3 |
| Solyc01g080250.2 | Solyc01g080250.3 |
| Solyc01g080260.2 | Solyc01g080260.3 |
| Solyc01g080270.2 | Solyc01g080270.3 |
| Solyc01g080280.2 | Solyc01g080280.3 |
| Solyc01g080290.2 | Solyc01g080290.3 |
| Solyc01g080340.2 | Solyc01g080340.3 |
| Solyc01g080350.2 | Solyc01g080350.3 |
| Solyc01g080360.2 | Solyc01g080360.3 |
| Solyc01g080370.2 | Solyc01g080370.3 |
| Solyc01g080400.2 | Solyc01g080400.3 |
| Solyc01g080410.2 | Solyc01g080410.3 |
| Solyc01g080440.1 | Solyc01g080440.2 |
| Solyc01g080460.2 | Solyc01g080460.3 |
| Solyc01g080470.2 | Solyc01g080470.3 |
| Solyc01g080520.2 | Solyc01g080520.3 |
| Solyc01g080530.2 | Solyc01g080530.3 |
| Solyc01g080540.2 | Solyc01g080540.3 |
| Solyc01g080610.2 | Solyc01g080610.3 |
| Solyc01g080620.2 | Solyc01g080620.3 |
| Solyc01g080640.2 | Solyc01g080640.3 |
| Solyc01g080660.2 | Solyc01g080660.3 |
| Solyc01g080670.2 | Solyc01g080670.3 |
| Solyc01g080680.2 | Solyc01g080680.3 |
| Solyc01g080690.2 | Solyc01g080690.3 |
| Solyc01g080750.2 | Solyc01g080750.3 |
| Solyc01g080770.2 | Solyc01g080770.3 |
| Solyc01g080790.2 | Solyc01g080790.3 |

| | |
|---|---|
| Solyc01g080800.2 | Solyc01g080800.3 |
| Solyc01g080810.2 | Solyc01g080810.3 |
| Solyc01g080830.2 | Solyc01g080830.3 |
| Solyc01g080840.2 | Solyc01g080840.3 |
| Solyc01g080860.2 | Solyc01g080860.3 |
| Solyc01g080870.2 | Solyc01g080870.3 |
| Solyc01g080920.2 | Solyc01g080920.3 |
| Solyc01g080940.2 | Solyc01g080940.3 |
| Solyc01g080950.2 | Solyc01g080950.3 |
| Solyc01g080970.2 | Solyc01g080970.3 |
| Solyc01g080980.1 | Solyc01g080980.2 |
| Solyc01g080990.2 | Solyc01g080990.3 |
| Solyc01g081000.2 | Solyc01g081000.3 |
| Solyc01g081060.3 | Solyc01g081060.4 |
| Solyc01g081070.2 | Solyc01g081070.3 |
| Solyc01g081090.1 | Solyc01g081090.2 |
| Solyc01g081150.2 | Solyc01g081150.3 |
| Solyc01g081180.1 | Solyc01g081180.2 |
| Solyc01g081220.1 | Solyc01g081220.2 |
| Solyc01g081240.2 | Solyc01g081240.3 |
| Solyc01g081250.2 | Solyc01g081250.3 |
| Solyc01g081260.1 | Solyc01g081260.2 |
| Solyc01g081280.2 | Solyc01g081280.3 |
| Solyc01g081310.2 | Solyc01g081310.3 |
| Solyc01g081330.2 | Solyc01g081330.3 |
| Solyc01g081400.2 | Solyc01g081400.3 |
| Solyc01g081440.2 | Solyc01g081440.3 |
| Solyc01g081450.2 | Solyc01g081450.3 |
| Solyc01g081460.2 | Solyc01g081460.3 |
| Solyc01g081480.2 | Solyc01g081480.3 |
| Solyc01g081490.2 | Solyc01g081490.3 |
| Solyc01g081520.2 | Solyc01g081520.3 |
| Solyc01g081570.2 | Solyc01g081570.3 |
| Solyc01g081580.1 | Solyc01g081580.2 |
| Solyc01g081600.2 | Solyc01g081600.3 |
| Solyc01g081610.2 | Solyc01g081610.3 |
| Solyc01g081620.2 | Solyc01g081620.3 |
| Solyc01g073650.2 | Solyc01g073650.3 |
| Solyc01g073660.2 | Solyc01g073660.3 |
| Solyc01g073670.2 | Solyc01g073670.3 |
| Solyc01g073700.2 | Solyc01g073700.3 |
| Solyc01g073860.2 | Solyc01g073860.3 |
| Solyc01g073890.2 | Solyc01g073890.3 |

| | |
|---|---|
| Solyc01g073920.2 | Solyc01g073920.3 |
| Solyc01g073930.2 | Solyc01g073930.3 |
| Solyc01g073940.2 | Solyc01g073940.3 |
| Solyc01g073970.2 | Solyc01g073970.3 |
| Solyc01g074000.2 | Solyc01g074000.3 |
| Solyc01g074020.2 | Solyc01g074020.3 |
| Solyc01g074030.2 | Solyc01g074030.3 |
| Solyc01g074040.1 | Solyc01g074040.2 |
| Solyc01g086650.2 | Solyc01g086650.3 |
| Solyc01g086670.2 | Solyc01g086670.3 |
| Solyc01g086720.2 | Solyc01g086720.3 |
| Solyc01g086740.2 | Solyc01g086740.3 |
| Solyc01g086830.3 | Solyc01g086830.4 |
| Solyc01g086850.2 | Solyc01g086850.3 |
| Solyc01g086870.2 | Solyc01g086870.3 |
| Solyc01g086890.2 | Solyc01g086890.3 |
| Solyc01g086920.2 | Solyc01g086920.3 |
| Solyc01g086960.2 | Solyc01g086960.3 |
| Solyc01g086990.2 | Solyc01g086990.3 |
| Solyc01g087000.2 | Solyc01g087000.3 |
| Solyc01g087010.2 | Solyc01g087010.3 |
| Solyc01g087030.2 | Solyc01g087030.3 |
| Solyc01g087050.2 | Solyc01g087050.3 |
| Solyc01g087090.2 | Solyc01g087090.3 |
| Solyc01g087095.1 | Solyc01g087095.2 |
| Solyc01g087120.2 | Solyc01g087120.3 |
| Solyc01g087150.2 | Solyc01g087150.3 |
| Solyc01g087170.2 | Solyc01g087170.3 |
| Solyc01g087200.2 | Solyc01g087200.3 |
| Solyc01g087210.2 | Solyc01g087210.3 |
| Solyc01g087230.2 | Solyc01g087230.3 |
| Solyc01g087240.2 | Solyc01g087240.3 |
| Solyc01g087250.2 | Solyc01g087250.3 |
| Solyc01g087270.2 | Solyc01g087270.3 |
| Solyc01g087280.1 | Solyc01g087280.2 |
| Solyc01g087290.2 | Solyc01g087290.3 |
| Solyc01g087300.2 | Solyc01g087300.3 |
| Solyc01g087320.2 | Solyc01g087320.3 |
| Solyc01g087340.2 | Solyc01g087340.3 |
| Solyc01g087350.2 | Solyc01g087350.3 |
| Solyc01g087430.2 | Solyc01g087430.3 |
| Solyc01g087440.2 | Solyc01g087440.3 |
| Solyc01g087450.2 | Solyc01g087450.3 |

| | |
|---|---|
| Solyc01g087470.2 | Solyc01g087470.3 |
| Solyc01g087500.2 | Solyc01g087500.3 |
| Solyc01g087520.2 | Solyc01g087520.3 |
| Solyc01g087530.2 | Solyc01g087530.3 |
| Solyc01g087550.2 | Solyc01g087550.3 |
| Solyc01g087560.2 | Solyc01g087560.3 |
| Solyc01g087590.2 | Solyc01g087590.3 |
| Solyc01g087610.2 | Solyc01g087610.3 |
| Solyc01g087620.2 | Solyc01g087620.3 |
| Solyc01g087630.1 | Solyc01g087630.2 |
| Solyc01g087640.2 | Solyc01g087640.3 |
| Solyc01g087670.2 | Solyc01g087670.3 |
| Solyc01g087840.2 | Solyc01g087840.3 |
| Solyc01g087890.2 | Solyc01g087890.3 |
| Solyc01g087900.2 | Solyc01g087900.3 |
| Solyc01g087920.1 | Solyc01g087920.2 |
| Solyc01g087950.2 | Solyc01g087950.3 |
| Solyc01g087970.2 | Solyc01g087970.3 |
| Solyc01g087990.2 | Solyc01g087990.3 |
| Solyc01g088020.2 | Solyc01g088020.3 |
| Solyc01g088040.2 | Solyc01g088040.3 |
| Solyc01g088050.2 | Solyc01g088050.3 |
| Solyc01g088060.2 | Solyc01g088060.3 |
| Solyc01g088080.2 | Solyc01g088080.3 |
| Solyc01g088090.2 | Solyc01g088090.3 |
| Solyc01g088150.2 | Solyc01g088150.3 |
| Solyc01g088230.2 | Solyc01g088230.3 |
| Solyc01g088200.3 | Solyc01g088200.4 |
| Solyc01g088260.2 | Solyc01g088260.3 |
| Solyc01g088270.2 | Solyc01g088270.3 |
| Solyc01g088290.1 | Solyc01g088290.2 |
| Solyc01g088300.2 | Solyc01g088300.3 |
| Solyc01g088310.2 | Solyc01g088310.3 |
| Solyc01g088320.1 | Solyc01g088320.2 |
| Solyc01g088340.1 | Solyc01g088340.2 |
| Solyc01g088370.2 | Solyc01g088370.3 |
| Solyc01g088390.2 | Solyc01g088390.3 |
| Solyc01g088410.2 | Solyc01g088410.3 |
| Solyc01g088460.2 | Solyc01g088460.3 |
| Solyc01g088470.2 | Solyc01g088470.3 |
| Solyc01g088480.2 | Solyc01g088480.3 |
| Solyc01g088490.2 | Solyc01g088490.3 |
| Solyc01g088500.2 | Solyc01g088500.3 |

| | |
|---|---|
| Solyc01g088505.1 | Solyc01g150146.1 |
| Solyc01g088520.2 | Solyc01g088520.3 |
| Solyc01g088550.2 | Solyc01g088550.3 |
| Solyc01g088560.2 | Solyc01g088560.3 |
| Solyc01g088570.2 | Solyc01g088570.3 |
| Solyc01g088620.2 | Solyc01g088620.3 |
| Solyc01g088660.2 | Solyc01g088660.3 |
| Solyc01g088680.2 | Solyc01g088680.3 |
| Solyc01g088690.2 | Solyc01g088690.3 |
| Solyc01g088710.2 | Solyc01g088710.3 |
| Solyc01g088720.2 | Solyc01g088720.3 |
| Solyc01g088730.2 | Solyc01g088730.3 |
| Solyc01g088750.2 | Solyc01g088750.3 |
| Solyc01g088770.2 | Solyc01g088770.3 |
| Solyc01g088780.2 | Solyc01g088780.3 |
| Solyc01g089870.2 | Solyc01g089870.3 |
| Solyc01g089880.2 | Solyc01g089880.3 |
| Solyc01g089890.2 | Solyc01g089890.3 |
| Solyc01g089920.2 | Solyc01g089920.3 |
| Solyc01g089930.1 | Solyc01g089930.2 |
| Solyc01g089950.2 | Solyc01g089950.3 |
| Solyc01g089960.2 | Solyc01g089960.3 |
| Solyc01g089970.2 | Solyc01g089970.3 |
| Solyc01g090010.1 | Solyc01g090010.2 |
| Solyc01g090030.1 | Solyc01g090030.2 |
| Solyc01g090040.2 | Solyc01g090040.3 |
| Solyc01g090070.1 | Solyc01g090070.2 |
| Solyc01g090110.1 | Solyc01g090110.2 |
| Solyc01g090120.2 | Solyc01g090120.3 |
| Solyc01g090190.2 | Solyc01g090190.3 |
| Solyc01g090230.2 | Solyc01g090230.3 |
| Solyc01g090310.2 | Solyc01g090310.3 |
| Solyc01g090340.2 | Solyc01g090340.3 |
| Solyc01g090350.2 | Solyc01g090350.3 |
| Solyc01g090360.2 | Solyc01g090360.3 |
| Solyc01g090410.2 | Solyc01g090410.3 |
| Solyc01g090430.2 | Solyc01g090430.3 |
| Solyc01g090450.2 | Solyc01g090450.3 |
| Solyc01g090460.2 | Solyc01g090460.3 |
| Solyc01g090470.2 | Solyc01g090470.3 |
| Solyc01g090520.2 | Solyc01g090520.3 |
| Solyc01g090550.2 | Solyc01g090550.3 |
| Solyc01g090590.2 | Solyc01g090590.3 |

| | |
|---|---|
| Solyc01g090610.2 | Solyc01g090610.3 |
| Solyc01g090660.2 | Solyc01g090660.3 |
| Solyc01g090670.2 | Solyc01g090670.3 |
| Solyc01g090710.2 | Solyc01g090710.3 |
| Solyc01g090730.2 | Solyc01g090730.3 |
| Solyc01g090750.2 | Solyc01g090750.3 |
| Solyc01g090780.2 | Solyc01g090780.3 |
| Solyc01g090800.2 | Solyc01g090800.3 |
| Solyc01g090840.2 | Solyc01g090840.3 |
| Solyc01g090850.2 | Solyc01g090850.3 |
| Solyc01g090860.2 | Solyc01g090860.3 |
| Solyc01g090880.2 | Solyc01g090880.3 |
| Solyc01g090890.2 | Solyc01g090890.3 |
| Solyc01g090910.2 | Solyc01g090910.3 |
| Solyc01g090920.2 | Solyc01g090920.3 |
| Solyc01g090940.2 | Solyc01g090940.3 |
| Solyc01g090960.1 | Solyc01g090960.2 |
| Solyc01g090990.2 | Solyc01g090990.3 |
| Solyc01g091010.2 | Solyc01g091010.3 |
| Solyc01g091020.2 | Solyc01g091020.3 |
| Solyc01g091030.2 | Solyc01g091030.3 |
| Solyc01g091040.2 | Solyc01g091040.3 |
| Solyc01g091060.1 | Solyc01g091060.2 |
| Solyc01g091070.2 | Solyc01g091070.3 |
| Solyc01g091080.2 | Solyc01g091080.3 |
| Solyc01g091110.2 | Solyc01g091110.3 |
| Solyc01g091120.2 | Solyc01g091120.3 |
| Solyc01g091145.1 | Solyc01g150147.1 |
| Solyc01g091190.2 | Solyc01g091190.3 |
| Solyc01g091210.2 | Solyc01g091210.3 |
| Solyc01g091220.2 | Solyc01g091220.3 |
| Solyc01g091230.2 | Solyc01g091230.3 |
| Solyc01g091250.2 | Solyc01g091250.3 |
| Solyc01g091280.2 | Solyc01g091280.3 |
| Solyc01g091300.2 | Solyc01g091300.3 |
| Solyc01g091310.2 | Solyc01g091310.3 |
| Solyc01g091320.2 | Solyc01g091320.3 |
| Solyc01g091340.2 | Solyc01g091340.3 |
| Solyc01g091400.2 | Solyc01g091400.3 |
| Solyc01g091410.2 | Solyc01g091410.3 |
| Solyc01g091420.1 | Solyc01g091420.2 |
| Solyc01g091430.1 | Solyc01g091430.2 |
| Solyc01g091450.2 | Solyc01g091450.3 |

| | |
|---|---|
| Solyc01g091460.2 | Solyc01g091460.3 |
| Solyc01g091470.2 | Solyc01g091470.3 |
| Solyc01g091490.2 | Solyc01g091490.3 |
| Solyc01g091500.2 | Solyc01g091500.3 |
| Solyc01g091520.2 | Solyc01g091520.3 |
| Solyc01g091550.2 | Solyc01g091550.3 |
| Solyc01g091570.2 | Solyc01g091570.3 |
| Solyc01g091580.2 | Solyc01g091580.3 |
| Solyc01g091590.2 | Solyc01g091590.3 |
| Solyc01g091620.2 | Solyc01g091620.3 |
| Solyc01g091630.2 | Solyc01g091630.3 |
| Solyc01g091640.2 | Solyc01g091640.3 |
| Solyc01g091650.2 | Solyc01g091650.3 |
| Solyc01g091660.2 | Solyc01g091660.3 |
| Solyc01g091670.2 | Solyc01g091670.3 |
| Solyc01g091690.2 | Solyc01g091690.3 |
| Solyc01g091720.2 | Solyc01g091720.3 |
| Solyc01g091730.2 | Solyc01g091730.3 |
| Solyc01g091750.2 | Solyc01g091750.3 |
| Solyc01g091760.2 | Solyc01g091760.3 |
| Solyc01g091800.2 | Solyc01g091800.3 |
| Solyc01g091820.2 | Solyc01g091820.3 |
| Solyc01g091850.2 | Solyc01g091850.3 |
| Solyc01g091870.2 | Solyc01g091870.3 |
| Solyc01g091880.2 | Solyc01g091880.3 |
| Solyc01g091890.2 | Solyc01g091890.3 |
| Solyc01g091900.2 | Solyc01g091900.3 |
| Solyc01g091930.2 | Solyc01g091930.3 |
| Solyc01g091940.2 | Solyc01g091940.3 |
| Solyc01g092950.2 | Solyc01g092950.3 |
| Solyc01g093965.1 | Solyc01g093965.2 |
| Solyc01g093960.2 | Solyc01g093960.3 |
| Solyc01g093990.2 | Solyc01g093990.3 |
| Solyc01g094000.2 | Solyc01g094000.3 |
| Solyc01g094010.2 | Solyc01g094010.3 |
| Solyc01g094020.2 | Solyc01g094020.3 |
| Solyc01g094030.2 | Solyc01g094030.3 |
| Solyc01g094040.2 | Solyc01g094040.3 |
| Solyc01g094070.2 | Solyc01g094070.3 |
| Solyc01g094110.2 | Solyc01g094110.3 |
| Solyc01g094120.2 | Solyc01g094120.3 |
| Solyc01g094140.2 | Solyc01g094140.3 |
| Solyc01g094150.2 | Solyc01g094150.3 |

| | |
|---|---|
| Solyc01g094170.2 | Solyc01g094170.3 |
| Solyc01g094190.2 | Solyc01g094190.3 |
| Solyc01g094220.2 | Solyc01g094220.3 |
| Solyc01g094240.2 | Solyc01g094240.3 |
| Solyc01g094270.2 | Solyc01g094270.3 |
| Solyc01g094280.2 | Solyc01g094280.3 |
| Solyc01g094290.2 | Solyc01g094290.3 |
| Solyc01g094320.2 | Solyc01g094320.3 |
| Solyc01g094330.2 | Solyc01g094330.3 |
| Solyc01g094340.2 | Solyc01g094340.3 |
| Solyc01g094360.2 | Solyc01g094360.3 |
| Solyc01g094370.2 | Solyc01g094370.3 |
| Solyc01g094400.2 | Solyc01g094400.3 |
| Solyc01g094410.2 | Solyc01g094410.3 |
| Solyc01g094420.2 | Solyc01g094420.3 |
| Solyc01g094460.2 | Solyc01g094460.3 |
| Solyc01g094500.2 | Solyc01g094500.3 |
| Solyc01g094510.1 | Solyc01g094510.2 |
| Solyc01g094520.2 | Solyc01g094520.3 |
| Solyc01g094550.2 | Solyc01g094550.3 |
| Solyc01g094570.2 | Solyc01g094570.3 |
| Solyc01g094580.2 | Solyc01g094580.3 |
| Solyc01g094620.2 | Solyc01g094620.3 |
| Solyc01g094660.2 | Solyc01g094660.3 |
| Solyc01g094720.2 | Solyc01g094720.3 |
| Solyc01g094760.2 | Solyc01g094760.3 |
| Solyc01g094790.2 | Solyc01g094790.3 |
| Solyc01g094810.2 | Solyc01g094810.3 |
| Solyc01g094820.2 | Solyc01g094820.3 |
| Solyc01g094840.2 | Solyc01g094840.3 |
| Solyc01g094880.2 | Solyc01g094880.3 |
| Solyc01g094910.2 | Solyc01g094910.3 |
| Solyc01g094920.2 | Solyc01g094920.3 |
| Solyc01g094930.2 | Solyc01g094930.3 |
| Solyc01g094950.2 | Solyc01g094950.3 |
| Solyc01g094960.2 | Solyc01g094960.3 |
| Solyc01g095000.2 | Solyc01g095000.3 |
| Solyc01g095030.2 | Solyc01g095030.3 |
| Solyc01g095040.2 | Solyc01g095040.3 |
| Solyc01g095050.2 | Solyc01g095050.3 |
| Solyc01g095080.2 | Solyc01g095080.3 |
| Solyc01g095090.2 | Solyc01g095090.3 |
| Solyc01g095120.2 | Solyc01g095120.3 |

| | |
|---|---|
| Solyc01g095150.2 | Solyc01g095150.3 |
| Solyc01g095160.2 | Solyc01g095160.3 |
| Solyc01g095190.1 | Solyc01g095190.2 |
| Solyc01g095200.2 | Solyc01g095200.3 |
| Solyc01g095210.2 | Solyc01g095210.3 |
| Solyc01g095240.2 | Solyc01g095240.3 |
| Solyc01g095270.2 | Solyc01g095270.3 |
| Solyc01g095300.2 | Solyc01g095300.3 |
| Solyc01g095350.1 | Solyc01g095350.2 |
| Solyc01g095370.2 | Solyc01g095370.3 |
| Solyc01g095390.2 | Solyc01g095390.3 |
| Solyc01g095410.2 | Solyc01g095410.3 |
| Solyc01g095430.2 | Solyc01g095430.3 |
| Solyc01g095460.2 | Solyc01g095460.3 |
| Solyc01g095470.2 | Solyc01g095470.3 |
| Solyc01g095490.2 | Solyc01g095490.3 |
| Solyc01g095500.2 | Solyc01g095500.3 |
| Solyc01g095510.2 | Solyc01g095510.3 |
| Solyc01g095540.2 | Solyc01g095540.3 |
| Solyc01g095560.2 | Solyc01g095560.3 |
| Solyc01g095570.2 | Solyc01g095570.3 |
| Solyc01g095580.2 | Solyc01g095580.3 |
| Solyc01g095590.2 | Solyc01g095590.3 |
| Solyc01g095620.2 | Solyc01g095620.3 |
| Solyc01g095630.2 | Solyc01g095630.3 |
| Solyc01g095640.1 | Solyc01g095640.2 |
| Solyc01g095700.2 | Solyc01g095700.3 |
| Solyc01g095710.2 | Solyc01g095710.3 |
| Solyc01g095715.1 | Solyc01g150148.1 |
| Solyc01g095720.2 | Solyc01g095720.3 |
| Solyc01g095760.2 | Solyc01g095760.3 |
| Solyc01g095770.2 | Solyc01g095770.3 |
| Solyc01g095800.2 | Solyc01g095800.3 |
| Solyc01g095810.2 | Solyc01g095810.3 |
| Solyc01g095830.2 | Solyc01g095830.3 |
| Solyc01g095840.1 | Solyc01g095840.2 |
| Solyc01g095860.2 | Solyc01g095860.3 |
| Solyc01g095970.2 | Solyc01g095970.3 |
| Solyc01g095980.2 | Solyc01g095980.3 |
| Solyc01g095990.2 | Solyc01g095990.3 |
| Solyc01g096070.2 | Solyc01g096070.3 |
| Solyc01g096080.2 | Solyc01g096080.3 |
| Solyc01g096090.3 | Solyc01g096090.4 |

| | |
|---|---|
| Solyc01g096110.2 | Solyc01g096110.3 |
| Solyc01g096120.2 | Solyc01g096120.3 |
| Solyc01g096130.2 | Solyc01g096130.3 |
| Solyc01g096140.2 | Solyc01g096140.3 |
| Solyc01g096190.2 | Solyc01g096190.3 |
| Solyc01g096220.2 | Solyc01g096220.3 |
| Solyc01g096270.2 | Solyc01g096270.3 |
| Solyc01g096280.1 | Solyc01g096280.2 |
| Solyc01g096320.2 | Solyc01g096320.3 |
| Solyc01g096340.2 | Solyc01g096340.3 |
| Solyc01g096350.2 | Solyc01g096350.3 |
| Solyc01g096360.2 | Solyc01g096360.3 |
| Solyc01g096420.2 | Solyc01g096420.3 |
| Solyc01g096435.1 | Solyc01g150149.1 |
| Solyc01g096470.2 | Solyc01g096470.3 |
| Solyc01g096490.2 | Solyc01g096490.3 |
| Solyc01g096500.2 | Solyc01g096500.3 |
| Solyc01g096520.2 | Solyc01g096520.3 |
| Solyc01g096530.2 | Solyc01g096530.3 |
| Solyc01g096550.2 | Solyc01g096550.3 |
| Solyc01g096570.2 | Solyc01g096570.3 |
| Solyc01g096580.2 | Solyc01g096580.3 |
| Solyc01g096590.2 | Solyc01g096590.3 |
| Solyc01g096630.2 | Solyc01g096630.3 |
| Solyc01g096650.2 | Solyc01g096650.3 |
| Solyc01g096760.2 | Solyc01g096760.3 |
| Solyc01g096800.2 | Solyc01g096800.3 |
| Solyc01g096810.2 | Solyc01g096810.3 |
| Solyc01g096830.2 | Solyc01g096830.3 |
| Solyc01g096840.2 | Solyc01g096840.3 |
| Solyc01g096900.2 | Solyc01g096900.3 |
| Solyc01g096910.2 | Solyc01g096910.3 |
| Solyc01g096930.2 | Solyc01g096930.3 |
| Solyc01g096960.2 | Solyc01g096960.3 |
| Solyc01g096970.2 | Solyc01g096970.3 |
| Solyc01g096980.1 | Solyc01g096980.2 |
| Solyc01g097000.2 | Solyc01g097000.3 |
| Solyc01g097010.2 | Solyc01g097010.3 |
| Solyc01g097030.2 | Solyc01g097030.3 |
| Solyc01g097060.2 | Solyc01g097060.3 |
| Solyc01g097110.1 | Solyc01g097110.2 |
| Solyc01g097120.2 | Solyc01g097120.3 |
| Solyc01g097160.2 | Solyc01g097160.3 |

| | |
|---|---|
| Solyc01g097170.2 | Solyc01g097170.3 |
| Solyc01g097240.2 | Solyc01g097240.3 |
| Solyc01g097310.2 | Solyc01g097310.3 |
| Solyc01g097320.2 | Solyc01g097320.3 |
| Solyc01g097330.2 | Solyc01g097330.3 |
| Solyc01g097340.2 | Solyc01g097340.3 |
| Solyc01g097380.1 | Solyc01g097380.2 |
| Solyc01g097440.2 | Solyc01g097440.3 |
| Solyc01g097450.2 | Solyc01g097450.3 |
| Solyc01g097460.2 | Solyc01g097460.3 |
| Solyc01g097600.1 | Solyc01g097600.2 |
| Solyc01g097610.2 | Solyc01g097610.3 |
| Solyc01g097620.1 | Solyc01g097620.2 |
| Solyc01g097640.2 | Solyc01g097640.3 |
| Solyc01g097650.2 | Solyc01g097650.3 |
| Solyc01g097740.2 | Solyc01g097740.3 |
| Solyc01g097760.2 | Solyc01g097760.3 |
| Solyc01g097790.2 | Solyc01g097790.3 |
| Solyc01g097800.2 | Solyc01g097800.3 |
| Solyc01g097810.2 | Solyc01g097810.3 |
| Solyc01g097840.2 | Solyc01g097840.3 |
| Solyc01g097870.2 | Solyc01g097870.3 |
| Solyc01g097920.2 | Solyc01g097920.3 |
| Solyc01g097930.2 | Solyc01g097930.3 |
| Solyc01g097950.2 | Solyc01g097950.3 |
| Solyc01g097960.2 | Solyc01g097960.3 |
| Solyc01g097980.2 | Solyc01g097980.3 |
| Solyc01g097990.2 | Solyc01g097990.3 |
| Solyc01g098000.2 | Solyc01g098000.3 |
| Solyc01g098040.2 | Solyc01g098040.3 |
| Solyc01g098080.2 | Solyc01g098080.3 |
| Solyc01g098090.2 | Solyc01g098090.3 |
| Solyc01g098150.2 | Solyc01g098150.3 |
| Solyc01g098160.2 | Solyc01g098160.3 |
| Solyc01g098170.2 | Solyc01g098170.3 |
| Solyc01g098180.2 | Solyc01g098180.3 |
| Solyc01g098190.2 | Solyc01g098190.3 |
| Solyc01g098200.2 | Solyc01g098200.3 |
| Solyc01g098210.2 | Solyc01g098210.3 |
| Solyc01g098220.1 | Solyc01g098220.2 |
| Solyc01g098230.2 | Solyc01g098230.3 |
| Solyc01g098340.2 | Solyc01g098340.3 |
| Solyc01g098350.2 | Solyc01g098350.3 |

| | |
|---|---|
| Solyc01g098390.2 | Solyc01g098390.3 |
| Solyc01g098400.2 | Solyc01g098400.3 |
| Solyc01g098430.2 | Solyc01g098430.3 |
| Solyc01g098490.2 | Solyc01g098490.3 |
| Solyc01g098500.2 | Solyc01g098500.3 |
| Solyc01g098530.2 | Solyc01g098530.3 |
| Solyc01g098540.2 | Solyc01g098540.3 |
| Solyc01g098550.2 | Solyc01g098550.3 |
| Solyc01g098560.2 | Solyc01g098560.3 |
| Solyc01g098580.2 | Solyc01g098580.3 |
| Solyc01g098590.2 | Solyc01g098590.3 |
| Solyc01g098600.2 | Solyc01g098600.3 |
| Solyc01g098630.2 | Solyc01g098630.3 |
| Solyc01g098640.2 | Solyc01g098640.3 |
| Solyc01g098710.2 | Solyc01g098710.3 |
| Solyc01g098720.2 | Solyc01g098720.3 |
| Solyc01g098750.2 | Solyc01g098750.3 |
| Solyc01g098780.2 | Solyc01g098780.3 |
| Solyc01g098790.1 | Solyc01g098790.2 |
| Solyc01g098820.2 | Solyc01g098820.3 |
| Solyc01g098840.2 | Solyc01g098840.3 |
| Solyc01g098850.2 | Solyc01g098850.3 |
| Solyc01g098860.2 | Solyc01g098860.3 |
| Solyc01g098880.2 | Solyc01g098880.3 |
| Solyc01g098900.2 | Solyc01g098900.3 |
| Solyc01g098910.2 | Solyc01g098910.3 |
| Solyc01g098930.2 | Solyc01g098930.3 |
| Solyc01g098940.2 | Solyc01g098940.3 |
| Solyc01g099010.2 | Solyc01g099010.3 |
| Solyc01g099020.2 | Solyc01g099020.3 |
| Solyc01g099050.2 | Solyc01g099050.3 |
| Solyc01g099090.2 | Solyc01g099090.3 |
| Solyc01g099130.2 | Solyc01g099130.3 |
| Solyc01g099150.4 | Solyc01g099150.5.1 |
| Solyc01g099200.2 | Solyc01g099200.3 |
| Solyc01g099210.2 | Solyc01g099210.3 |
| Solyc01g099220.2 | Solyc01g099220.3 |
| Solyc01g099230.2 | Solyc01g099230.3 |
| Solyc01g099260.2 | Solyc01g099260.3 |
| Solyc01g099280.2 | Solyc01g099280.3 |
| Solyc01g099290.2 | Solyc01g099290.3 |
| Solyc01g099300.2 | Solyc01g099300.3 |
| Solyc01g099340.2 | Solyc01g099340.3 |

| | |
|---|---|
| Solyc01g099350.2 | Solyc01g099350.3 |
| Solyc01g099360.2 | Solyc01g099360.3 |
| Solyc01g099370.2 | Solyc01g099370.3 |
| Solyc01g099410.2 | Solyc01g099410.3 |
| Solyc01g099415.1 | Solyc01g150150.1 |
| Solyc01g099470.2 | Solyc01g099470.3 |
| Solyc01g099520.2 | Solyc01g099520.3 |
| Solyc01g099530.2 | Solyc01g099530.3 |
| Solyc01g099560.2 | Solyc01g099560.3 |
| Solyc01g099620.2 | Solyc01g099620.3 |
| Solyc01g099650.2 | Solyc01g099650.3 |
| Solyc01g099670.2 | Solyc01g099670.3 |
| Solyc01g099680.2 | Solyc01g099680.3 |
| Solyc01g099690.2 | Solyc01g099690.3 |
| Solyc01g099760.2 | Solyc01g099760.3 |
| Solyc01g099770.2 | Solyc01g099770.3 |
| Solyc01g099780.2 | Solyc01g099780.3 |
| Solyc01g099790.2 | Solyc01g099790.3 |
| Solyc01g099810.2 | Solyc01g099810.3 |
| Solyc01g099820.2 | Solyc01g099820.3 |
| Solyc01g099830.2 | Solyc01g099830.3 |
| Solyc01g099840.2 | Solyc01g099840.3 |
| Solyc01g099850.2 | Solyc01g099850.3 |
| Solyc01g099860.2 | Solyc01g099860.3 |
| Solyc01g099890.2 | Solyc01g099890.3 |
| Solyc01g099940.2 | Solyc01g099940.3 |
| Solyc01g099990.2 | Solyc01g099990.3 |
| Solyc01g100000.2 | Solyc01g100000.3 |
| Solyc01g100010.2 | Solyc01g100010.3 |
| Solyc01g100030.4 | Solyc01g100030.5.1 |
| Solyc01g100040.2 | Solyc01g100040.3 |
| Solyc01g100050.2 | Solyc01g100050.3 |
| Solyc01g100070.2 | Solyc01g100070.3 |
| Solyc01g100080.2 | Solyc01g100080.3 |
| Solyc01g100130.2 | Solyc01g100130.3 |
| Solyc01g100160.2 | Solyc01g100160.3 |
| Solyc01g100170.2 | Solyc01g100170.3 |
| Solyc01g100180.3 | Solyc01g100180.4 |
| Solyc01g100200.2 | Solyc01g100200.3 |
| Solyc01g100210.2 | Solyc01g100210.3 |
| Solyc01g100220.2 | Solyc01g100220.3 |
| Solyc01g100230.2 | Solyc01g100230.3 |
| Solyc01g100240.2 | Solyc01g100240.3 |

| | |
|---|---|
| Solyc01g100250.2 | Solyc01g100250.3 |
| Solyc01g100300.2 | Solyc01g100300.3 |
| Solyc01g100320.2 | Solyc01g100320.3 |
| Solyc01g100330.2 | Solyc01g100330.3 |
| Solyc01g100350.2 | Solyc01g100350.3 |
| Solyc01g100370.2 | Solyc01g100370.3 |
| Solyc01g100390.2 | Solyc01g100390.3 |
| Solyc01g100400.2 | Solyc01g100400.3 |
| Solyc01g100410.2 | Solyc01g100410.3 |
| Solyc01g100420.2 | Solyc01g100420.3 |
| Solyc01g100430.2 | Solyc01g100430.3 |
| Solyc01g100455.1 | Solyc01g150151.1 |
| Solyc01g100460.2 | Solyc01g100460.3 |
| Solyc01g100500.2 | Solyc01g100500.3 |
| Solyc01g100520.2 | Solyc01g100520.3 |
| Solyc01g100530.2 | Solyc01g100530.3 |
| Solyc01g100580.2 | Solyc01g100580.3 |
| Solyc01g100590.2 | Solyc01g100590.3 |
| Solyc01g100610.2 | Solyc01g100610.3 |
| Solyc01g100650.2 | Solyc01g100650.3 |
| Solyc01g100720.2 | Solyc01g100720.3 |
| Solyc01g100730.2 | Solyc01g100730.3 |
| Solyc01g100780.2 | Solyc01g100780.3 |
| Solyc01g100800.1 | Solyc01g100800.2 |
| Solyc01g100810.2 | Solyc01g100810.3 |
| Solyc01g100820.2 | Solyc01g100820.3 |
| Solyc01g100830.1 | Solyc01g100830.2 |
| Solyc01g100840.2 | Solyc01g100840.3 |
| Solyc01g100850.2 | Solyc01g100850.3 |
| Solyc01g100860.2 | Solyc01g100860.3 |
| Solyc01g100870.2 | Solyc01g100870.3 |
| Solyc01g100910.2 | Solyc01g100910.3 |
| Solyc01g100920.2 | Solyc01g100920.3 |
| Solyc01g100930.2 | Solyc01g100930.3 |
| Solyc01g100960.2 | Solyc01g100960.3 |
| Solyc01g100970.2 | Solyc01g100970.3 |
| Solyc01g100990.2 | Solyc01g100990.3 |
| Solyc01g101000.2 | Solyc01g101000.3 |
| Solyc01g101040.2 | Solyc01g101040.3 |
| Solyc01g101050.2 | Solyc01g101050.3 |
| Solyc01g101070.2 | Solyc01g101070.3 |
| Solyc01g101090.2 | Solyc01g101090.3 |
| Solyc01g101110.2 | Solyc01g101110.3 |

| | |
|---|---|
| Solyc01g101130.2 | Solyc01g101130.3 |
| Solyc01g101250.2 | Solyc01g101250.3 |
| Solyc01g102260.2 | Solyc01g102260.3 |
| Solyc01g102280.2 | Solyc01g102280.3 |
| Solyc01g102285.1 | Solyc01g150152.1 |
| Solyc01g102290.2 | Solyc01g102290.3 |
| Solyc01g102300.2 | Solyc01g102300.3 |
| Solyc01g102310.2 | Solyc01g102310.3 |
| Solyc01g102320.1 | Solyc01g102320.2 |
| Solyc01g102330.2 | Solyc01g102330.3 |
| Solyc01g102340.2 | Solyc01g102340.3 |
| Solyc01g102350.2 | Solyc01g102350.3 |
| Solyc01g102380.4 | Solyc01g102380.5.1 |
| Solyc01g102390.3 | Solyc01g102390.4 |
| Solyc01g102410.2 | Solyc01g102410.3 |
| Solyc01g102490.2 | Solyc01g102490.3 |
| Solyc01g102530.2 | Solyc01g102530.3 |
| Solyc01g102540.2 | Solyc01g102540.3 |
| Solyc01g102580.2 | Solyc01g102580.3 |
| Solyc01g102590.2 | Solyc01g102590.3 |
| Solyc01g102610.2 | Solyc01g102610.3 |
| Solyc01g102670.2 | Solyc01g102670.3 |
| Solyc01g102720.2 | Solyc01g102720.3 |
| Solyc01g102730.2 | Solyc01g102730.3 |
| Solyc01g102740.2 | Solyc01g102740.3 |
| Solyc01g102750.2 | Solyc01g102750.3 |
| Solyc01g102760.2 | Solyc01g102760.3 |
| Solyc01g102790.2 | Solyc01g102790.3 |
| Solyc01g102800.2 | Solyc01g102800.3 |
| Solyc01g102810.2 | Solyc01g102810.3 |
| Solyc01g102830.2 | Solyc01g102830.3 |
| Solyc01g102840.2 | Solyc01g102840.3 |
| Solyc01g102920.2 | Solyc01g102920.3 |
| Solyc01g102940.2 | Solyc01g102940.3 |
| Solyc01g102950.2 | Solyc01g102950.3 |
| Solyc01g102960.2 | Solyc01g102960.3 |
| Solyc01g102970.2 | Solyc01g102970.3 |
| Solyc01g102980.2 | Solyc01g102980.3 |
| Solyc01g102990.2 | Solyc01g102990.3 |
| Solyc01g103030.2 | Solyc01g103030.3 |
| Solyc01g103050.2 | Solyc01g103050.3 |
| Solyc01g103080.2 | Solyc01g103080.3 |
| Solyc01g103100.2 | Solyc01g103100.3 |

| | |
|---|---|
| Solyc01g103110.2 | Solyc01g103110.3 |
| Solyc01g103120.2 | Solyc01g103120.3 |
| Solyc01g103130.1 | Solyc01g103130.2 |
| Solyc01g103140.2 | Solyc01g103140.3 |
| Solyc01g103150.2 | Solyc01g103150.3 |
| Solyc01g103160.2 | Solyc01g103160.3 |
| Solyc01g103190.2 | Solyc01g103190.3 |
| Solyc01g103220.2 | Solyc01g103220.3 |
| Solyc01g103230.2 | Solyc01g103230.3 |
| Solyc01g103275.1 | Solyc01g150153.1 |
| Solyc01g103330.2 | Solyc01g103330.3 |
| Solyc01g103350.2 | Solyc01g103350.3 |
| Solyc01g103370.2 | Solyc01g103370.3 |
| Solyc01g103390.2 | Solyc01g103390.3 |
| Solyc01g103410.2 | Solyc01g103410.3 |
| Solyc01g103440.2 | Solyc01g103440.3 |
| Solyc01g103450.2 | Solyc01g103450.3 |
| Solyc01g103480.2 | Solyc01g103480.3 |
| Solyc01g103490.2 | Solyc01g103490.3 |
| Solyc01g103530.2 | Solyc01g103530.3 |
| Solyc01g103540.2 | Solyc01g103540.3 |
| Solyc01g103560.2 | Solyc01g103560.3 |
| Solyc01g103570.2 | Solyc01g103570.3 |
| Solyc01g103580.2 | Solyc01g103580.3 |
| Solyc01g103650.2 | Solyc01g103650.3 |
| Solyc01g103680.2 | Solyc01g103680.3 |
| Solyc01g103700.2 | Solyc01g103700.3 |
| Solyc01g103770.2 | Solyc01g103770.3 |
| Solyc01g103790.2 | Solyc01g103790.3 |
| Solyc01g103800.2 | Solyc01g103800.3 |
| Solyc01g103803.1 | Solyc01g150154.1 |
| Solyc01g103807.1 | Solyc01g150155.1 |
| Solyc01g103890.2 | Solyc01g103890.3 |
| Solyc01g103920.2 | Solyc01g103920.3 |
| Solyc01g103930.2 | Solyc01g103930.3 |
| Solyc01g103940.2 | Solyc01g103940.3 |
| Solyc01g103960.2 | Solyc01g103960.3 |
| Solyc01g103990.2 | Solyc01g103990.3 |
| Solyc01g104000.2 | Solyc01g104000.3 |
| Solyc01g104030.2 | Solyc01g104030.3 |
| Solyc01g104060.2 | Solyc01g104060.3 |
| Solyc01g104070.2 | Solyc01g104070.3 |
| Solyc01g104100.2 | Solyc01g104100.3 |

| | |
|---|---|
| Solyc01g104140.2 | Solyc01g104140.3 |
| Solyc01g104170.2 | Solyc01g104170.3 |
| Solyc01g104220.2 | Solyc01g104220.3 |
| Solyc01g104240.2 | Solyc01g104240.3 |
| Solyc01g104260.2 | Solyc01g104260.3 |
| Solyc01g104300.2 | Solyc01g104300.3 |
| Solyc01g104310.2 | Solyc01g104310.3 |
| Solyc01g104340.1 | Solyc01g104340.2 |
| Solyc01g104380.2 | Solyc01g104380.3 |
| Solyc01g104390.1 | Solyc01g104390.2 |
| Solyc01g104400.2 | Solyc01g104400.3 |
| Solyc01g104431.1 | Solyc01g150156.1 |
| Solyc01g104432.1 | Solyc01g150157.1 |
| Solyc01g104433.1 | Solyc01g150158.1 |
| Solyc01g104434.1 | Solyc01g150159.1 |
| Solyc01g104435.1 | Solyc01g150160.1 |
| Solyc01g104436.1 | Solyc01g150161.1 |
| Solyc01g104438.1 | Solyc01g150162.1 |
| Solyc01g104470.2 | Solyc01g104470.3 |
| Solyc01g104490.2 | Solyc01g104490.3 |
| Solyc01g104510.2 | Solyc01g104510.3 |
| Solyc01g104525.1 | Solyc01g150163.1 |
| Solyc01g104530.2 | Solyc01g104530.3 |
| Solyc01g104550.2 | Solyc01g104550.3 |
| Solyc01g104580.2 | Solyc01g104580.3 |
| Solyc01g104590.2 | Solyc01g104590.3 |
| Solyc01g104630.2 | Solyc01g104630.3 |
| Solyc01g104640.2 | Solyc01g104640.3 |
| Solyc01g104650.2 | Solyc01g104650.3 |
| Solyc01g104660.2 | Solyc01g104660.3 |
| Solyc01g104670.2 | Solyc01g104670.3 |
| Solyc01g104725.1 | Solyc01g150165.1 |
| Solyc01g104740.2 | Solyc01g104740.3 |
| Solyc01g104780.2 | Solyc01g104780.3 |
| Solyc01g104850.2 | Solyc01g104850.3 |
| Solyc01g104860.2 | Solyc01g104860.3 |
| Solyc01g104870.2 | Solyc01g104870.3 |
| Solyc01g104880.2 | Solyc01g104880.3 |
| Solyc01g104900.2 | Solyc01g104900.3 |
| Solyc01g104920.2 | Solyc01g104920.3 |
| Solyc01g104940.1 | Solyc01g104940.2 |
| Solyc01g104970.2 | Solyc01g104970.3 |
| Solyc01g105000.2 | Solyc01g105000.3 |

| | |
|---|---|
| Solyc01g105010.2 | Solyc01g105010.3 |
| Solyc01g105020.2 | Solyc01g105020.3 |
| Solyc01g105030.2 | Solyc01g105030.3 |
| Solyc01g105040.2 | Solyc01g105040.3 |
| Solyc01g105050.2 | Solyc01g105050.3 |
| Solyc01g105060.2 | Solyc01g105060.3 |
| Solyc01g105070.2 | Solyc01g105070.3 |
| Solyc01g105090.2 | Solyc01g105090.3 |
| Solyc01g105100.2 | Solyc01g105100.3 |
| Solyc01g105140.2 | Solyc01g105140.3 |
| Solyc01g105180.2 | Solyc01g105180.3 |
| Solyc01g105220.2 | Solyc01g105220.3 |
| Solyc01g105240.2 | Solyc01g105240.3 |
| Solyc01g105280.2 | Solyc01g105280.3 |
| Solyc01g105310.2 | Solyc01g105310.3 |
| Solyc01g105330.2 | Solyc01g105330.3 |
| Solyc01g105360.2 | Solyc01g105360.3 |
| Solyc01g105510.2 | Solyc01g105510.3 |
| Solyc01g105520.2 | Solyc01g105520.3 |
| Solyc01g105560.2 | Solyc01g105560.3 |
| Solyc01g105670.2 | Solyc01g105670.3 |
| Solyc01g105700.2 | Solyc01g105700.3 |
| Solyc01g105710.2 | Solyc01g105710.3 |
| Solyc01g105800.3 | Solyc01g105800.4 |
| Solyc01g105820.2 | Solyc01g105820.3 |
| Solyc01g105890.2 | Solyc01g105890.3 |
| Solyc01g105910.1 | Solyc01g105910.2 |
| Solyc01g105920.2 | Solyc01g105920.3 |
| Solyc01g105930.1 | Solyc01g105930.2 |
| Solyc01g105940.2 | Solyc01g105940.3 |
| Solyc01g105950.2 | Solyc01g105950.3 |
| Solyc01g105970.2 | Solyc01g105970.3 |
| Solyc01g105990.2 | Solyc01g105990.3 |
| Solyc01g106010.2 | Solyc01g106010.3 |
| Solyc01g106020.2 | Solyc01g106020.3 |
| Solyc01g106040.2 | Solyc01g106040.3 |
| Solyc01g106060.2 | Solyc01g106060.3 |
| Solyc01g106090.2 | Solyc01g106090.3 |
| Solyc01g106140.1 | Solyc01g106140.2 |
| Solyc01g106173.1 | Solyc01g150166.1 |
| Solyc01g106175.1 | Solyc01g150167.1 |
| Solyc01g106177.1 | Solyc01g150168.1 |
| Solyc01g106195.1 | Solyc01g150169.1 |

| | |
|---|---|
| Solyc01g106210.2 | Solyc01g106210.3 |
| Solyc01g106260.2 | Solyc01g106260.3 |
| Solyc01g106280.2 | Solyc01g106280.3 |
| Solyc01g106290.2 | Solyc01g106290.3 |
| Solyc01g106300.2 | Solyc01g106300.3 |
| Solyc01g106310.2 | Solyc01g106310.3 |
| Solyc01g106330.2 | Solyc01g106330.3 |
| Solyc01g106340.2 | Solyc01g106340.3 |
| Solyc01g106360.2 | Solyc01g106360.3 |
| Solyc01g106370.2 | Solyc01g106370.3 |
| Solyc01g106390.2 | Solyc01g106390.3 |
| Solyc01g106400.2 | Solyc01g106400.3 |
| Solyc01g106420.2 | Solyc01g106420.3 |
| Solyc01g106430.2 | Solyc01g106430.3 |
| Solyc01g106440.2 | Solyc01g106440.3 |
| Solyc01g106450.2 | Solyc01g106450.3 |
| Solyc01g106470.2 | Solyc01g106470.3 |
| Solyc01g106570.2 | Solyc01g106570.3 |
| Solyc01g106650.2 | Solyc01g106650.3 |
| Solyc01g106690.2 | Solyc01g106690.3 |
| Solyc01g106700.2 | Solyc01g106700.3 |
| Solyc01g106750.1 | Solyc01g106750.2 |
| Solyc01g106780.2 | Solyc01g106780.3 |
| Solyc01g106790.2 | Solyc01g106790.3 |
| Solyc01g106820.2 | Solyc01g106820.3 |
| Solyc01g106840.1 | Solyc01g106840.2 |
| Solyc01g106880.2 | Solyc01g106880.3 |
| Solyc01g106900.2 | Solyc01g106900.3 |
| Solyc01g106980.1 | Solyc01g106980.2 |
| Solyc01g106990.1 | Solyc01g106990.2 |
| Solyc01g107010.2 | Solyc01g107010.3 |
| Solyc01g107040.2 | Solyc01g107040.3 |
| Solyc01g107050.2 | Solyc01g107050.3 |
| Solyc01g107080.2 | Solyc01g107080.3 |
| Solyc01g107120.2 | Solyc01g107120.3 |
| Solyc01g107150.2 | Solyc01g107150.3 |
| Solyc01g107160.2 | Solyc01g107160.3 |
| Solyc01g107180.2 | Solyc01g107180.3 |
| Solyc01g107190.2 | Solyc01g107190.3 |
| Solyc01g107250.2 | Solyc01g107250.3 |
| Solyc01g107290.2 | Solyc01g107290.3 |
| Solyc01g107320.2 | Solyc01g107320.3 |
| Solyc01g107350.2 | Solyc01g107350.3 |

| | |
|---|---|
| Solyc01g107370.2 | Solyc01g107370.3 |
| Solyc01g107380.2 | Solyc01g107380.3 |
| Solyc01g107410.2 | Solyc01g107410.3 |
| Solyc01g107490.1 | Solyc01g107490.2 |
| Solyc01g107510.2 | Solyc01g107510.3 |
| Solyc01g107520.2 | Solyc01g107520.3 |
| Solyc01g107550.2 | Solyc01g107550.3 |
| Solyc01g107560.2 | Solyc01g107560.3 |
| Solyc01g107590.2 | Solyc01g107590.3 |
| Solyc01g107600.2 | Solyc01g107600.3 |
| Solyc01g107640.2 | Solyc01g107640.3 |
| Solyc01g107650.2 | Solyc01g107650.3 |
| Solyc01g107660.2 | Solyc01g107660.3 |
| Solyc01g107720.2 | Solyc01g107720.3 |
| Solyc01g107730.2 | Solyc01g107730.3 |
| Solyc01g107740.2 | Solyc01g107740.3 |
| Solyc01g107750.2 | Solyc01g107750.3 |
| Solyc01g107860.2 | Solyc01g107860.3 |
| Solyc01g107870.2 | Solyc01g107870.3 |
| Solyc01g107880.2 | Solyc01g107880.3 |
| Solyc01g107890.2 | Solyc01g107890.3 |
| Solyc01g107900.2 | Solyc01g107900.3 |
| Solyc01g107930.2 | Solyc01g107930.3 |
| Solyc01g107940.2 | Solyc01g107940.3 |
| Solyc01g107950.1 | Solyc01g107950.2 |
| Solyc01g107980.2 | Solyc01g107980.3 |
| Solyc01g107990.2 | Solyc01g107990.3 |
| Solyc01g108000.2 | Solyc01g108000.3 |
| Solyc01g108010.2 | Solyc01g108010.3 |
| Solyc01g108020.2 | Solyc01g108020.3 |
| Solyc01g108035.1 | Solyc01g150170.1 |
| Solyc01g108050.2 | Solyc01g108050.3 |
| Solyc01g108060.1 | Solyc01g108060.2 |
| Solyc01g108090.1 | Solyc01g108090.2 |
| Solyc01g108120.2 | Solyc01g108120.3 |
| Solyc01g108150.2 | Solyc01g108150.3 |
| Solyc01g108160.2 | Solyc01g108160.3 |
| Solyc01g108170.2 | Solyc01g108170.3 |
| Solyc01g108190.2 | Solyc01g108190.3 |
| Solyc01g108200.2 | Solyc01g108200.3 |
| Solyc01g108220.2 | Solyc01g108220.3 |
| Solyc01g108230.2 | Solyc01g108230.3 |
| Solyc01g108240.2 | Solyc01g108240.3 |

| | |
|---|---|
| Solyc01g108250.2 | Solyc01g108250.3 |
| Solyc01g108280.2 | Solyc01g108280.3 |
| Solyc01g108290.2 | Solyc01g108290.3 |
| Solyc01g108300.2 | Solyc01g108300.3 |
| Solyc01g108320.2 | Solyc01g108320.3 |
| Solyc01g108340.2 | Solyc01g108340.3 |
| Solyc01g108380.1 | Solyc01g108380.2 |
| Solyc01g108390.2 | Solyc01g108390.3 |
| Solyc01g108400.2 | Solyc01g108400.3 |
| Solyc01g108430.2 | Solyc01g108430.3 |
| Solyc01g108450.2 | Solyc01g108450.3 |
| Solyc01g108490.2 | Solyc01g108490.3 |
| Solyc01g108510.2 | Solyc01g108510.3 |
| Solyc01g108520.1 | Solyc01g108520.2 |
| Solyc01g108530.2 | Solyc01g108530.3 |
| Solyc01g108540.2 | Solyc01g108540.3 |
| Solyc01g108560.2 | Solyc01g108560.4 |
| Solyc01g108570.2 | Solyc01g108570.3 |
| Solyc01g108580.2 | Solyc01g108580.3 |
| Solyc01g108600.2 | Solyc01g108600.3 |
| Solyc01g108610.2 | Solyc01g108610.3 |
| Solyc01g108630.2 | Solyc01g108630.3 |
| Solyc01g108710.2 | Solyc01g108710.3 |
| Solyc01g108720.1 | Solyc01g108720.2 |
| Solyc01g108730.1 | Solyc01g108730.2 |
| Solyc01g108740.2 | Solyc01g108740.3 |
| Solyc01g108810.2 | Solyc01g108810.3 |
| Solyc01g108840.2 | Solyc01g108840.3 |
| Solyc01g108860.2 | Solyc01g108860.3 |
| Solyc01g108930.2 | Solyc01g108930.3 |
| Solyc01g108940.1 | Solyc01g108940.2 |
| Solyc01g108990.2 | Solyc01g108990.3 |
| Solyc01g109080.2 | Solyc01g109080.3 |
| Solyc01g109110.2 | Solyc01g109110.3 |
| Solyc01g109120.2 | Solyc01g109120.3 |
| Solyc01g109130.2 | Solyc01g109130.3 |
| Solyc01g109140.2 | Solyc01g109140.3 |
| Solyc01g109170.2 | Solyc01g109170.3 |
| Solyc01g109230.2 | Solyc01g109230.3 |
| Solyc01g109260.2 | Solyc01g109260.3 |
| Solyc01g109280.2 | Solyc01g109280.3 |
| Solyc01g109290.2 | Solyc01g109290.3 |
| Solyc01g109300.2 | Solyc01g109300.3 |

| | |
|---|---|
| Solyc01g109310.2 | Solyc01g109310.3 |
| Solyc01g109330.2 | Solyc01g109330.3 |
| Solyc01g109350.2 | Solyc01g109350.3 |
| Solyc01g109360.2 | Solyc01g109360.3 |
| Solyc01g109370.2 | Solyc01g109370.3 |
| Solyc01g109380.2 | Solyc01g109380.3 |
| Solyc01g109390.2 | Solyc01g109390.3 |
| Solyc01g109400.2 | Solyc01g109400.3 |
| Solyc01g109420.2 | Solyc01g109420.3 |
| Solyc01g109430.2 | Solyc01g109430.3 |
| Solyc01g109450.2 | Solyc01g109450.3 |
| Solyc01g109460.2 | Solyc01g109460.3 |
| Solyc01g109470.2 | Solyc01g109470.3 |
| Solyc01g109505.1 | Solyc01g150171.1 |
| Solyc01g109510.2 | Solyc01g109510.3 |
| Solyc01g109520.2 | Solyc01g109520.3 |
| Solyc01g109530.2 | Solyc01g109530.3 |
| Solyc01g109560.2 | Solyc01g109560.3 |
| Solyc01g109590.2 | Solyc01g109590.3 |
| Solyc01g109620.2 | Solyc01g109620.3 |
| Solyc01g109640.2 | Solyc01g109640.3 |
| Solyc01g109650.2 | Solyc01g109650.3 |
| Solyc01g109670.2 | Solyc01g109670.3 |
| Solyc01g109700.2 | Solyc01g109700.3 |
| Solyc01g109710.2 | Solyc01g109710.3 |
| Solyc01g109720.2 | Solyc01g109720.3 |
| Solyc01g109740.2 | Solyc01g109740.3 |
| Solyc01g109760.2 | Solyc01g109760.3 |
| Solyc01g109770.2 | Solyc01g109770.3 |
| Solyc01g109790.2 | Solyc01g109790.3 |
| Solyc01g109830.2 | Solyc01g109830.3 |
| Solyc01g109850.2 | Solyc01g109850.3 |
| Solyc01g109860.2 | Solyc01g109860.3 |
| Solyc01g109870.2 | Solyc01g109870.3 |
| Solyc01g109880.2 | Solyc01g109880.3 |
| Solyc01g109890.2 | Solyc01g109890.3 |
| Solyc01g109940.2 | Solyc01g109940.3 |
| Solyc01g109960.2 | Solyc01g109960.3 |
| Solyc01g109970.2 | Solyc01g109970.3 |
| Solyc01g110000.2 | Solyc01g110000.3 |
| Solyc01g110010.2 | Solyc01g110010.3 |
| Solyc01g110030.2 | Solyc01g110030.3 |
| Solyc01g110060.2 | Solyc01g110060.3 |

| | |
|---|---|
| Solyc01g110080.1 | Solyc01g110080.2 |
| Solyc01g110110.2 | Solyc01g110110.3 |
| Solyc01g110120.2 | Solyc01g110120.3 |
| Solyc01g110130.2 | Solyc01g110130.3 |
| Solyc01g110200.1 | Solyc01g110200.2 |
| Solyc01g110270.2 | Solyc01g110270.3 |
| Solyc01g110280.2 | Solyc01g110280.3 |
| Solyc01g110290.2 | Solyc01g110290.3 |
| Solyc01g110310.2 | Solyc01g110310.3 |
| Solyc01g110360.2 | Solyc01g110360.3 |
| Solyc01g110380.2 | Solyc01g110380.3 |
| Solyc01g110410.2 | Solyc01g110410.3 |
| Solyc01g110440.4 | Solyc01g110440.5.1 |
| Solyc01g110430.2 | Solyc01g110430.3 |
| Solyc01g110450.2 | Solyc01g110450.3 |
| Solyc01g110470.2 | Solyc01g110470.3 |
| Solyc01g110500.2 | Solyc01g110500.3 |
| Solyc01g110510.2 | Solyc01g110510.3 |
| Solyc01g110520.2 | Solyc01g110520.3 |
| Solyc01g110530.2 | Solyc01g110530.3 |
| Solyc01g110540.2 | Solyc01g110540.3 |
| Solyc01g110550.2 | Solyc01g110550.3 |
| Solyc01g110570.2 | Solyc01g110570.3 |
| Solyc01g110640.1 | Solyc01g110640.2 |
| Solyc01g110643.1 | Solyc01g150172.1 |
| Solyc01g110670.2 | Solyc01g110670.3 |
| Solyc01g110682.1 | Solyc01g150173.1 |
| Solyc01g110684.1 | Solyc01g150174.1 |
| Solyc01g110688.1 | Solyc01g150175.1 |
| Solyc01g110710.2 | Solyc01g110710.3 |
| Solyc01g110790.2 | Solyc01g110790.3 |
| Solyc01g110800.2 | Solyc01g110800.3 |
| Solyc01g110810.2 | Solyc01g110810.3 |
| Solyc01g110840.2 | Solyc01g110840.3 |
| Solyc01g110870.2 | Solyc01g110870.3 |
| Solyc01g110940.2 | Solyc01g110940.3 |
| Solyc01g110950.2 | Solyc01g110950.3 |
| Solyc01g110970.2 | Solyc01g110970.3 |
| Solyc01g111020.2 | Solyc01g111020.3 |
| Solyc01g111030.1 | Solyc01g111030.2 |
| Solyc01g111070.2 | Solyc01g111070.3 |
| Solyc01g111075.1 | Solyc01g150176.1 |
| Solyc01g111090.2 | Solyc01g111090.3 |

| | |
|---|---|
| Solyc01g111140.2 | Solyc01g111140.3 |
| Solyc01g111160.2 | Solyc01g111160.3 |
| Solyc01g111170.2 | Solyc01g111170.3 |
| Solyc01g111200.2 | Solyc01g111200.3 |
| Solyc01g111210.2 | Solyc01g111210.3 |
| Solyc01g111220.2 | Solyc01g111220.3 |
| Solyc01g111230.2 | Solyc01g111230.3 |
| Solyc01g111250.2 | Solyc01g111250.3 |
| Solyc01g111270.2 | Solyc01g111270.3 |
| Solyc01g111280.2 | Solyc01g111280.3 |
| Solyc01g111310.2 | Solyc01g111310.3 |
| Solyc01g111320.2 | Solyc01g111320.3 |
| Solyc01g111350.2 | Solyc01g111350.3 |
| Solyc01g111360.2 | Solyc01g111360.3 |
| Solyc01g111370.2 | Solyc01g111370.3 |
| Solyc01g111380.2 | Solyc01g111380.3 |
| Solyc01g111410.2 | Solyc01g111410.3 |
| Solyc01g111420.2 | Solyc01g111420.3 |
| Solyc01g111450.2 | Solyc01g111450.3 |
| Solyc01g111455.1 | Solyc01g150177.1 |
| Solyc01g111460.2 | Solyc01g111460.3 |
| Solyc01g111500.2 | Solyc01g111500.3 |
| Solyc01g111510.2 | Solyc01g111510.3 |
| Solyc01g111520.2 | Solyc01g111520.3 |
| Solyc01g111530.2 | Solyc01g111530.3 |
| Solyc01g111550.2 | Solyc01g111550.3 |
| Solyc01g111560.2 | Solyc01g111560.3 |
| Solyc01g111570.2 | Solyc01g111570.3 |
| Solyc01g111580.2 | Solyc01g111580.3 |
| Solyc01g111600.2 | Solyc01g111600.3 |
| Solyc01g111610.2 | Solyc01g111610.3 |
| Solyc01g111620.2 | Solyc01g111620.3 |
| Solyc01g111630.2 | Solyc01g111630.3 |
| Solyc01g111640.2 | Solyc01g111640.3 |
| Solyc01g111650.2 | Solyc01g111650.3 |
| Solyc01g111690.2 | Solyc01g111690.3 |
| Solyc01g111700.1 | Solyc01g111700.2 |
| Solyc01g111730.2 | Solyc01g111730.3 |
| Solyc01g111740.2 | Solyc01g111740.3 |
| Solyc01g111790.1 | Solyc01g111790.2 |
| Solyc01g111810.2 | Solyc01g111810.3 |
| Solyc01g111830.2 | Solyc01g111830.3 |
| Solyc01g111840.2 | Solyc01g111840.3 |

| | |
|---|---|
| Solyc01g111850.2 | Solyc01g111850.3 |
| Solyc01g111890.2 | Solyc01g111890.3 |
| Solyc01g111910.2 | Solyc01g111910.3 |
| Solyc01g111920.2 | Solyc01g111920.3 |
| Solyc01g111960.2 | Solyc01g111960.3 |
| Solyc01g111980.2 | Solyc01g111980.3 |
| Solyc01g111990.2 | Solyc01g111990.3 |
| Solyc01g112020.2 | Solyc01g112020.3 |
| Solyc01g112050.2 | Solyc01g112050.3 |
| Solyc01g112080.2 | Solyc01g112080.3 |
| Solyc01g112090.2 | Solyc01g112090.3 |
| Solyc01g112100.2 | Solyc01g112100.3 |
| Solyc01g112150.2 | Solyc01g112150.3 |
| Solyc01g112170.2 | Solyc01g112170.3 |
| Solyc01g112190.2 | Solyc01g112190.3 |
| Solyc01g112200.2 | Solyc01g112200.3 |
| Solyc01g112210.2 | Solyc01g112210.3 |
| Solyc01g112280.2 | Solyc01g112280.3 |
| Solyc01g112320.2 | Solyc01g112320.3 |
| Solyc01g112340.2 | Solyc01g112340.3 |
| Solyc01g112350.2 | Solyc01g112350.3 |
| Solyc02g043860.2 | Solyc02g043860.3 |
| Solyc02g055565.1 | Solyc02g150122.1 |
| Solyc02g055550.2 | Solyc02g055550.3 |
| Solyc02g055505.1 | Solyc02g150121.1 |
| Solyc02g055390.1 | Solyc02g055390.2 |
| Solyc02g049102.1 | Solyc02g150117.1 |
| Solyc02g049104.1 | Solyc02g150118.1 |
| Solyc02g049106.1 | Solyc02g150119.1 |
| Solyc02g050190.1 | Solyc02g050190.2 |
| Solyc02g050200.1 | Solyc02g050200.2 |
| Solyc02g050210.1 | Solyc02g050210.2 |
| Solyc02g050320.1 | Solyc02g050320.2 |
| Solyc02g050335.1 | Solyc02g150120.1 |
| Solyc02g005110.2 | Solyc02g005110.3 |
| Solyc02g005140.2 | Solyc02g005140.3 |
| Solyc02g005180.2 | Solyc02g005180.3 |
| Solyc02g005210.2 | Solyc02g005210.3 |
| Solyc02g005320.1 | Solyc02g005320.2 |
| Solyc02g005350.2 | Solyc02g005350.3 |
| Solyc02g005430.1 | Solyc02g005430.2 |
| Solyc02g005440.1 | Solyc02g005440.2 |
| Solyc02g005490.2 | Solyc02g005490.3 |

| | |
|---|---|
| Solyc02g005605.1 | Solyc02g150100.1 |
| Solyc02g011725.1 | Solyc02g150101.1 |
| Solyc02g011790.1 | Solyc02g011790.2 |
| Solyc02g011870.1 | Solyc02g011870.2 |
| Solyc02g011920.1 | Solyc02g011920.2 |
| Solyc02g011975.1 | Solyc02g150102.1 |
| Solyc02g014050.1 | Solyc02g014050.2 |
| Solyc02g014070.1 | Solyc02g014070.2 |
| Solyc02g014073.1 | Solyc02g150103.1 |
| Solyc02g014077.1 | Solyc02g150104.1 |
| Solyc02g014090.2 | Solyc02g014090.3 |
| Solyc02g014170.2 | Solyc02g014170.3 |
| Solyc02g014180.1 | Solyc02g014180.2 |
| Solyc02g014220.2 | Solyc02g014220.3 |
| Solyc02g014300.1 | Solyc02g014300.2 |
| Solyc02g014325.1 | Solyc02g150105.1 |
| Solyc02g014360.2 | Solyc02g014360.3 |
| Solyc02g014380.2 | Solyc02g014380.3 |
| Solyc02g014430.2 | Solyc02g014430.3 |
| Solyc02g014560.2 | Solyc02g014560.3 |
| Solyc02g014670.1 | Solyc02g014670.2 |
| Solyc02g014730.2 | Solyc02g014730.3 |
| Solyc02g014740.1 | Solyc02g014740.2 |
| Solyc02g014750.1 | Solyc02g014750.2 |
| Solyc02g014770.2 | Solyc02g014770.3 |
| Solyc02g014777.1 | Solyc02g150106.1 |
| Solyc02g014800.1 | Solyc02g014800.2 |
| Solyc02g014840.2 | Solyc02g014840.3 |
| Solyc02g014845.1 | Solyc02g150107.1 |
| Solyc02g014860.2 | Solyc02g014860.3 |
| Solyc02g014873.1 | Solyc02g150108.1 |
| Solyc02g020880.1 | Solyc02g020880.2 |
| Solyc02g020940.2 | Solyc02g020940.3 |
| Solyc02g020960.1 | Solyc02g020960.2 |
| Solyc02g021000.2 | Solyc02g021000.3 |
| Solyc02g021010.1 | Solyc02g021010.2 |
| Solyc02g021063.1 | Solyc02g150109.1 |
| Solyc02g021100.2 | Solyc02g021100.3 |
| Solyc02g021140.2 | Solyc02g021140.3 |
| Solyc02g021160.1 | Solyc02g021160.2 |
| Solyc02g021240.2 | Solyc02g021240.3 |
| Solyc02g021270.2 | Solyc02g021270.3 |
| Solyc02g021360.2 | Solyc02g021360.3 |

| | |
|---|---|
| Solyc02g021400.1 | Solyc02g021400.2 |
| Solyc02g021420.2 | Solyc02g021420.3 |
| Solyc02g021475.1 | Solyc02g150110.1 |
| Solyc02g021550.2 | Solyc02g021550.3 |
| Solyc02g021590.2 | Solyc02g021590.3 |
| Solyc02g021620.2 | Solyc02g021620.3 |
| Solyc02g021640.2 | Solyc02g021640.3 |
| Solyc02g021650.2 | Solyc02g021650.3 |
| Solyc02g021660.1 | Solyc02g021660.2 |
| Solyc02g021670.1 | Solyc02g021670.2 |
| Solyc02g021680.2 | Solyc02g021680.3 |
| Solyc02g021760.2 | Solyc02g021760.3 |
| Solyc02g022830.2 | Solyc02g022830.3 |
| Solyc02g022900.2 | Solyc02g022900.3 |
| Solyc02g022930.2 | Solyc02g022930.3 |
| Solyc02g023950.2 | Solyc02g023950.3 |
| Solyc02g023970.2 | Solyc02g023970.3 |
| Solyc02g023980.2 | Solyc02g023980.3 |
| Solyc02g023990.2 | Solyc02g023990.3 |
| Solyc02g024020.2 | Solyc02g024020.3 |
| Solyc02g024050.2 | Solyc02g024050.3 |
| Solyc02g024070.2 | Solyc02g024070.3 |
| Solyc02g024075.1 | Solyc02g150111.1 |
| Solyc02g026080.1 | Solyc02g026080.2 |
| Solyc02g030137.1 | Solyc02g150112.1 |
| Solyc02g030260.1 | Solyc02g030260.2 |
| Solyc02g030280.1 | Solyc02g030280.2 |
| Solyc02g030300.2 | Solyc02g030300.3 |
| Solyc02g030450.2 | Solyc02g030450.3 |
| Solyc02g030464.1 | Solyc02g150113.1 |
| Solyc02g030466.1 | Solyc02g150114.1 |
| Solyc02g030490.1 | Solyc02g030490.2 |
| Solyc02g030540.1 | Solyc02g030540.2 |
| Solyc02g030600.1 | Solyc02g030600.2 |
| Solyc02g030605.1 | Solyc02g150115.1 |
| Solyc02g030630.1 | Solyc02g030630.2 |
| Solyc02g030640.1 | Solyc02g030640.2 |
| Solyc02g031710.2 | Solyc02g031710.3 |
| Solyc02g031740.2 | Solyc02g031740.3 |
| Solyc02g031750.2 | Solyc02g031750.3 |
| Solyc02g031760.1 | Solyc02g031760.2 |
| Solyc02g031840.2 | Solyc02g031840.3 |
| Solyc02g031860.2 | Solyc02g031860.3 |

| | |
|---|---|
| Solyc02g031960.2 | Solyc02g031960.3 |
| Solyc02g031980.2 | Solyc02g031980.3 |
| Solyc02g032080.1 | Solyc02g032080.2 |
| Solyc02g032120.2 | Solyc02g032120.3 |
| Solyc02g032300.2 | Solyc02g032300.3 |
| Solyc02g032330.2 | Solyc02g032330.3 |
| Solyc02g032340.1 | Solyc02g032340.2 |
| Solyc02g032410.2 | Solyc02g032410.3 |
| Solyc02g032450.2 | Solyc02g032450.3 |
| Solyc02g032490.2 | Solyc02g032490.3 |
| Solyc02g032510.2 | Solyc02g032510.3 |
| Solyc02g032630.1 | Solyc02g032630.2 |
| Solyc02g032650.2 | Solyc02g032650.3 |
| Solyc02g032660.2 | Solyc02g032660.3 |
| Solyc02g032680.1 | Solyc02g032680.2 |
| Solyc02g032780.1 | Solyc02g032780.2 |
| Solyc02g032790.1 | Solyc02g032790.2 |
| Solyc02g032820.2 | Solyc02g032820.3 |
| Solyc02g032850.2 | Solyc02g032850.3 |
| Solyc02g032860.2 | Solyc02g032860.3 |
| Solyc02g032920.2 | Solyc02g032920.3 |
| Solyc02g032930.2 | Solyc02g032930.3 |
| Solyc02g032950.2 | Solyc02g032950.3 |
| Solyc02g033030.2 | Solyc02g033030.3 |
| Solyc02g033080.1 | Solyc02g033080.2 |
| Solyc02g036280.2 | Solyc02g036280.3 |
| Solyc02g036300.2 | Solyc02g036300.3 |
| Solyc02g036320.1 | Solyc02g036320.2 |
| Solyc02g036330.2 | Solyc02g036330.3 |
| Solyc02g036350.2 | Solyc02g036350.3 |
| Solyc02g036370.2 | Solyc02g036370.3 |
| Solyc02g036380.2 | Solyc02g036380.3 |
| Solyc02g036430.1 | Solyc02g036430.2 |
| Solyc02g036440.2 | Solyc02g036440.3 |
| Solyc02g036450.2 | Solyc02g036450.3 |
| Solyc02g037510.2 | Solyc02g037510.3 |
| Solyc02g037530.2 | Solyc02g037530.3 |
| Solyc02g037550.2 | Solyc02g037550.3 |
| Solyc02g037595.1 | Solyc02g150116.1 |
| Solyc02g038720.1 | Solyc02g038720.2 |
| Solyc02g038750.2 | Solyc02g038750.3 |
| Solyc02g060590.1 | Solyc02g060590.2 |
| Solyc02g061620.1 | Solyc02g061620.2 |

| | |
|---|---|
| Solyc02g061640.1 | Solyc02g061640.2 |
| Solyc02g061690.1 | Solyc02g061690.2 |
| Solyc02g061710.1 | Solyc02g061710.2 |
| Solyc02g061730.1 | Solyc02g061730.2 |
| Solyc02g061780.2 | Solyc02g061780.3 |
| Solyc02g061810.2 | Solyc02g061810.3 |
| Solyc02g061840.2 | Solyc02g061840.3 |
| Solyc02g061920.1 | Solyc02g061920.2 |
| Solyc02g061960.2 | Solyc02g061960.3 |
| Solyc02g061980.2 | Solyc02g061980.3 |
| Solyc02g061990.2 | Solyc02g061990.3 |
| Solyc02g062020.2 | Solyc02g062020.3 |
| Solyc02g062040.2 | Solyc02g062040.3 |
| Solyc02g062070.1 | Solyc02g062070.2 |
| Solyc02g062120.2 | Solyc02g062120.3 |
| Solyc02g062180.2 | Solyc02g062180.3 |
| Solyc02g062190.1 | Solyc02g062190.2 |
| Solyc02g062220.1 | Solyc02g062220.2 |
| Solyc02g062240.2 | Solyc02g062240.3 |
| Solyc02g062250.2 | Solyc02g062250.3 |
| Solyc02g062280.2 | Solyc02g062280.3 |
| Solyc02g062300.2 | Solyc02g062300.3 |
| Solyc02g062320.2 | Solyc02g062320.3 |
| Solyc02g062340.2 | Solyc02g062340.3 |
| Solyc02g062350.2 | Solyc02g062350.3 |
| Solyc02g062390.2 | Solyc02g062390.3 |
| Solyc02g062400.2 | Solyc02g062400.3 |
| Solyc02g062410.1 | Solyc02g062410.2 |
| Solyc02g062430.2 | Solyc02g062430.3 |
| Solyc02g062500.2 | Solyc02g062500.3 |
| Solyc02g062510.2 | Solyc02g062510.3 |
| Solyc02g062520.2 | Solyc02g062520.3 |
| Solyc02g062540.2 | Solyc02g062540.3 |
| Solyc02g062550.2 | Solyc02g062550.3 |
| Solyc02g062560.2 | Solyc02g062560.3 |
| Solyc02g062580.2 | Solyc02g062580.3 |
| Solyc02g062590.2 | Solyc02g062590.3 |
| Solyc02g062600.2 | Solyc02g062600.3 |
| Solyc02g062610.2 | Solyc02g062610.3 |
| Solyc02g062620.2 | Solyc02g062620.3 |
| Solyc02g062650.2 | Solyc02g062650.3 |
| Solyc02g062670.2 | Solyc02g062670.3 |
| Solyc02g062680.2 | Solyc02g062680.3 |

| | |
|---|---|
| Solyc02g062690.2 | Solyc02g062690.3 |
| Solyc02g062700.2 | Solyc02g062700.3 |
| Solyc02g062740.2 | Solyc02g062740.3 |
| Solyc02g062790.2 | Solyc02g062790.3 |
| Solyc02g062830.1 | Solyc02g062830.2 |
| Solyc02g062860.2 | Solyc02g062860.3 |
| Solyc02g062870.2 | Solyc02g062870.3 |
| Solyc02g062920.2 | Solyc02g062920.3 |
| Solyc02g062940.2 | Solyc02g062940.3 |
| Solyc02g062950.2 | Solyc02g062950.3 |
| Solyc02g062965.1 | Solyc02g150123.1 |
| Solyc02g062970.2 | Solyc02g062970.3 |
| Solyc02g063010.2 | Solyc02g063010.3 |
| Solyc02g063070.2 | Solyc02g063070.3 |
| Solyc02g063090.2 | Solyc02g063090.3 |
| Solyc02g063100.1 | Solyc02g063100.2 |
| Solyc02g063150.2 | Solyc02g063150.3 |
| Solyc02g063170.2 | Solyc02g063170.3 |
| Solyc02g063220.2 | Solyc02g063220.3 |
| Solyc02g063250.2 | Solyc02g063250.3 |
| Solyc02g063255.1 | Solyc02g150124.1 |
| Solyc02g063290.2 | Solyc02g063290.3 |
| Solyc02g063330.2 | Solyc02g063330.3 |
| Solyc02g063410.2 | Solyc02g063410.3 |
| Solyc02g063420.2 | Solyc02g063420.3 |
| Solyc02g063440.2 | Solyc02g063440.3 |
| Solyc02g063490.2 | Solyc02g063490.3 |
| Solyc02g063520.2 | Solyc02g063520.3 |
| Solyc02g064570.1 | Solyc02g064570.2 |
| Solyc02g064580.2 | Solyc02g064580.3 |
| Solyc02g064590.1 | Solyc02g064590.2 |
| Solyc02g064650.2 | Solyc02g064650.3 |
| Solyc02g064690.2 | Solyc02g064690.3 |
| Solyc02g064700.2 | Solyc02g064700.3 |
| Solyc02g064720.2 | Solyc02g064720.3 |
| Solyc02g064730.2 | Solyc02g064730.3 |
| Solyc02g064740.2 | Solyc02g064740.3 |
| Solyc02g064795.1 | Solyc02g150125.1 |
| Solyc02g064800.2 | Solyc02g064800.3 |
| Solyc02g064850.1 | Solyc02g064850.2 |
| Solyc02g064950.2 | Solyc02g064950.3 |
| Solyc02g064960.2 | Solyc02g064960.3 |
| Solyc02g064990.2 | Solyc02g064990.3 |

| | |
|---|---|
| Solyc02g065020.1 | Solyc02g065020.2 |
| Solyc02g065060.2 | Solyc02g065060.3 |
| Solyc02g065070.2 | Solyc02g065070.3 |
| Solyc02g065080.2 | Solyc02g065080.3 |
| Solyc02g065110.2 | Solyc02g065110.3 |
| Solyc02g065150.1 | Solyc02g065150.2 |
| Solyc02g065160.1 | Solyc02g065160.2 |
| Solyc02g065170.2 | Solyc02g065170.3 |
| Solyc02g065180.2 | Solyc02g065180.3 |
| Solyc02g065190.2 | Solyc02g065190.3 |
| Solyc02g065230.2 | Solyc02g065230.3 |
| Solyc02g065240.2 | Solyc02g065240.3 |
| Solyc02g065250.1 | Solyc02g065250.2 |
| Solyc02g065280.2 | Solyc02g065280.3 |
| Solyc02g065320.1 | Solyc02g065320.2 |
| Solyc02g065350.2 | Solyc02g065350.3 |
| Solyc02g065380.2 | Solyc02g065380.3 |
| Solyc02g065400.2 | Solyc02g065400.3 |
| Solyc02g065420.2 | Solyc02g065420.3 |
| Solyc02g065580.2 | Solyc02g065580.3 |
| Solyc02g065590.2 | Solyc02g065590.3 |
| Solyc02g065600.2 | Solyc02g065600.3 |
| Solyc02g065610.2 | Solyc02g065610.3 |
| Solyc02g065620.2 | Solyc02g065620.3 |
| Solyc02g065640.2 | Solyc02g065640.3 |
| Solyc02g065680.2 | Solyc02g065680.3 |
| Solyc02g065690.2 | Solyc02g065690.3 |
| Solyc02g065700.2 | Solyc02g065700.3 |
| Solyc02g065720.2 | Solyc02g065720.3 |
| Solyc02g065730.1 | Solyc02g065730.2 |
| Solyc02g065740.2 | Solyc02g065740.3 |
| Solyc02g065790.2 | Solyc02g065790.3 |
| Solyc02g065805.1 | Solyc02g150126.1 |
| Solyc02g066840.1 | Solyc02g066840.2 |
| Solyc02g066880.1 | Solyc02g066880.2 |
| Solyc02g066890.1 | Solyc02g066890.2 |
| Solyc02g066910.2 | Solyc02g066910.3 |
| Solyc02g066920.2 | Solyc02g066920.3 |
| Solyc02g066930.2 | Solyc02g066930.3 |
| Solyc02g066960.2 | Solyc02g066960.3 |
| Solyc02g067010.2 | Solyc02g067010.3 |
| Solyc02g067040.2 | Solyc02g067040.3 |
| Solyc02g067050.2 | Solyc02g067050.3 |

| | |
|---|---|
| Solyc02g067060.2 | Solyc02g067060.3 |
| Solyc02g067100.2 | Solyc02g067100.3 |
| Solyc02g067180.2 | Solyc02g067180.3 |
| Solyc02g067210.2 | Solyc02g067210.3 |
| Solyc02g067230.2 | Solyc02g067230.3 |
| Solyc02g067290.2 | Solyc02g067290.3 |
| Solyc02g067350.2 | Solyc02g067350.3 |
| Solyc02g067360.2 | Solyc02g067360.3 |
| Solyc02g067390.2 | Solyc02g067390.3 |
| Solyc02g067410.1 | Solyc02g067410.2 |
| Solyc02g067420.2 | Solyc02g067420.3 |
| Solyc02g067430.2 | Solyc02g067430.3 |
| Solyc02g067440.2 | Solyc02g067440.3 |
| Solyc02g067460.2 | Solyc02g067460.3 |
| Solyc02g067480.2 | Solyc02g067480.3 |
| Solyc02g067490.2 | Solyc02g067490.3 |
| Solyc02g067510.2 | Solyc02g067510.3 |
| Solyc02g067540.2 | Solyc02g067540.3 |
| Solyc02g067550.1 | Solyc02g067550.2 |
| Solyc02g067570.2 | Solyc02g067570.3 |
| Solyc02g067580.2 | Solyc02g067580.3 |
| Solyc02g067590.1 | Solyc02g067590.2 |
| Solyc02g067690.2 | Solyc02g067690.3 |
| Solyc02g067730.1 | Solyc02g067730.2 |
| Solyc02g067770.2 | Solyc02g067770.3 |
| Solyc02g067780.2 | Solyc02g067780.3 |
| Solyc02g067830.2 | Solyc02g067830.3 |
| Solyc02g067840.2 | Solyc02g067840.3 |
| Solyc02g067870.2 | Solyc02g067870.3 |
| Solyc02g067890.2 | Solyc02g067890.3 |
| Solyc02g067900.2 | Solyc02g067900.3 |
| Solyc02g067930.2 | Solyc02g067930.3 |
| Solyc02g067980.2 | Solyc02g067980.3 |
| Solyc02g067990.2 | Solyc02g067990.3 |
| Solyc02g068040.2 | Solyc02g068040.3 |
| Solyc02g068060.2 | Solyc02g068060.3 |
| Solyc02g068080.2 | Solyc02g068080.3 |
| Solyc02g068090.2 | Solyc02g068090.3 |
| Solyc02g068100.2 | Solyc02g068100.3 |
| Solyc02g068120.2 | Solyc02g068120.3 |
| Solyc02g068140.2 | Solyc02g068140.3 |
| Solyc02g068180.2 | Solyc02g068180.3 |
| Solyc02g068230.2 | Solyc02g068230.3 |

| | |
|---|---|
| Solyc02g068300.2 | Solyc02g068300.3 |
| Solyc02g068330.1 | Solyc02g068330.2 |
| Solyc02g068340.2 | Solyc02g068340.3 |
| Solyc02g068370.2 | Solyc02g068370.3 |
| Solyc02g068380.2 | Solyc02g068380.3 |
| Solyc02g068390.2 | Solyc02g068390.3 |
| Solyc02g068440.2 | Solyc02g068440.3 |
| Solyc02g068450.2 | Solyc02g068450.3 |
| Solyc02g068480.2 | Solyc02g068480.3 |
| Solyc02g068500.2 | Solyc02g068500.3 |
| Solyc02g068510.1 | Solyc02g068510.2 |
| Solyc02g068540.1 | Solyc02g068540.2 |
| Solyc02g068550.1 | Solyc02g068550.2 |
| Solyc02g068560.2 | Solyc02g068560.3 |
| Solyc02g068590.2 | Solyc02g068590.3 |
| Solyc02g068600.2 | Solyc02g068600.3 |
| Solyc02g068610.1 | Solyc02g068610.2 |
| Solyc02g068640.2 | Solyc02g068640.3 |
| Solyc02g068650.1 | Solyc02g068650.2 |
| Solyc02g068690.2 | Solyc02g068690.3 |
| Solyc02g068700.2 | Solyc02g068700.3 |
| Solyc02g068710.2 | Solyc02g068710.3 |
| Solyc02g068720.2 | Solyc02g068720.3 |
| Solyc02g068730.2 | Solyc02g068730.3 |
| Solyc02g068740.2 | Solyc02g068740.3 |
| Solyc02g068795.1 | Solyc02g150127.1 |
| Solyc02g068825.1 | Solyc02g150128.1 |
| Solyc02g068880.1 | Solyc02g068880.2 |
| Solyc02g068890.2 | Solyc02g068890.3 |
| Solyc02g068900.2 | Solyc02g068900.3 |
| Solyc02g068910.2 | Solyc02g068910.3 |
| Solyc02g068920.2 | Solyc02g068920.3 |
| Solyc02g068970.1 | Solyc02g068970.2 |
| Solyc02g068990.2 | Solyc02g068990.3 |
| Solyc02g069020.2 | Solyc02g069020.3 |
| Solyc02g069030.2 | Solyc02g069030.3 |
| Solyc02g069070.2 | Solyc02g069070.3 |
| Solyc02g069090.2 | Solyc02g069090.3 |
| Solyc02g069120.2 | Solyc02g069120.3 |
| Solyc02g069130.2 | Solyc02g069130.3 |
| Solyc02g069150.2 | Solyc02g069150.3 |
| Solyc02g069160.2 | Solyc02g069160.3 |
| Solyc02g069220.1 | Solyc02g069220.2 |

| | |
|---|---|
| Solyc02g069230.2 | Solyc02g069230.3 |
| Solyc02g069260.2 | Solyc02g069260.3 |
| Solyc02g069290.2 | Solyc02g069290.3 |
| Solyc02g069310.2 | Solyc02g069310.3 |
| Solyc02g069330.1 | Solyc02g069330.2 |
| Solyc02g069370.2 | Solyc02g069370.3 |
| Solyc02g069380.2 | Solyc02g069380.3 |
| Solyc02g069390.1 | Solyc02g069390.2 |
| Solyc02g069400.2 | Solyc02g069400.3 |
| Solyc02g069440.2 | Solyc02g069440.3 |
| Solyc02g069450.2 | Solyc02g069450.3 |
| Solyc02g069460.2 | Solyc02g069460.3 |
| Solyc02g069560.2 | Solyc02g069560.3 |
| Solyc02g069570.2 | Solyc02g069570.3 |
| Solyc02g069580.2 | Solyc02g069580.3 |
| Solyc02g069600.2 | Solyc02g069600.3 |
| Solyc02g069610.2 | Solyc02g069610.3 |
| Solyc02g069630.2 | Solyc02g069630.3 |
| Solyc02g069640.2 | Solyc02g069640.3 |
| Solyc02g069680.2 | Solyc02g069680.3 |
| Solyc02g069720.2 | Solyc02g069720.3 |
| Solyc02g069730.2 | Solyc02g069730.3 |
| Solyc02g069760.1 | Solyc02g069760.2 |
| Solyc02g069770.2 | Solyc02g069770.3 |
| Solyc02g069780.2 | Solyc02g069780.3 |
| Solyc02g069790.2 | Solyc02g069790.3 |
| Solyc02g069840.2 | Solyc02g069840.3 |
| Solyc02g069850.2 | Solyc02g069850.3 |
| Solyc02g069910.1 | Solyc02g069910.2 |
| Solyc02g069950.2 | Solyc02g069950.3 |
| Solyc02g070050.2 | Solyc02g070050.3 |
| Solyc02g070055.1 | Solyc02g150129.1 |
| Solyc02g070100.2 | Solyc02g070100.3 |
| Solyc02g070210.2 | Solyc02g070210.3 |
| Solyc02g070220.2 | Solyc02g070220.3 |
| Solyc02g070230.1 | Solyc02g070230.2 |
| Solyc02g070240.2 | Solyc02g070240.3 |
| Solyc02g070270.1 | Solyc02g070270.2 |
| Solyc02g070280.2 | Solyc02g070280.3 |
| Solyc02g070290.2 | Solyc02g070290.3 |
| Solyc02g070310.2 | Solyc02g070310.3 |
| Solyc02g070340.2 | Solyc02g070340.3 |
| Solyc02g070390.2 | Solyc02g070390.3 |

| | |
|---|---|
| Solyc02g070400.2 | Solyc02g070400.3 |
| Solyc02g070410.1 | Solyc02g070410.2 |
| Solyc02g070430.2 | Solyc02g070430.3 |
| Solyc02g070440.2 | Solyc02g070440.3 |
| Solyc02g070450.1 | Solyc02g070450.2 |
| Solyc02g070460.2 | Solyc02g070460.3 |
| Solyc02g070510.2 | Solyc02g070510.3 |
| Solyc02g070520.2 | Solyc02g070520.3 |
| Solyc02g070540.2 | Solyc02g070540.3 |
| Solyc02g070550.2 | Solyc02g070550.3 |
| Solyc02g070560.2 | Solyc02g070560.3 |
| Solyc02g070610.2 | Solyc02g070610.3 |
| Solyc02g070630.2 | Solyc02g070630.3 |
| Solyc02g070640.2 | Solyc02g070640.3 |
| Solyc02g070660.2 | Solyc02g070660.3 |
| Solyc02g070770.2 | Solyc02g070770.3 |
| Solyc02g070790.2 | Solyc02g070790.3 |
| Solyc02g070920.2 | Solyc02g070920.3 |
| Solyc02g071020.2 | Solyc02g071020.3 |
| Solyc02g071030.1 | Solyc02g071030.2 |
| Solyc02g071040.2 | Solyc02g071040.3 |
| Solyc02g071060.2 | Solyc02g071060.3 |
| Solyc02g071070.2 | Solyc02g071070.3 |
| Solyc02g071090.2 | Solyc02g071090.3 |
| Solyc02g071100.2 | Solyc02g071100.3 |
| Solyc02g071110.2 | Solyc02g071110.3 |
| Solyc02g071120.2 | Solyc02g071120.3 |
| Solyc02g071140.2 | Solyc02g071140.3 |
| Solyc02g071150.2 | Solyc02g071150.3 |
| Solyc02g071170.2 | Solyc02g071170.3 |
| Solyc02g071180.2 | Solyc02g071180.3 |
| Solyc02g071210.2 | Solyc02g071210.3 |
| Solyc02g071220.2 | Solyc02g071220.3 |
| Solyc02g071240.2 | Solyc02g071240.3 |
| Solyc02g071250.2 | Solyc02g071250.3 |
| Solyc02g071260.2 | Solyc02g071260.3 |
| Solyc02g071280.2 | Solyc02g071280.3 |
| Solyc02g071285.1 | Solyc02g150130.1 |
| Solyc02g071310.2 | Solyc02g071310.3 |
| Solyc02g071340.1 | Solyc02g071340.2 |
| Solyc02g071350.2 | Solyc02g071350.3 |
| Solyc02g071400.2 | Solyc02g071400.3 |
| Solyc02g071410.2 | Solyc02g071410.3 |

| | |
|---|---|
| Solyc02g071430.2 | Solyc02g071430.3 |
| Solyc02g071440.2 | Solyc02g071440.3 |
| Solyc02g071450.2 | Solyc02g071450.3 |
| Solyc02g071480.2 | Solyc02g071480.3 |
| Solyc02g071510.2 | Solyc02g071510.3 |
| Solyc02g071520.2 | Solyc02g071520.3 |
| Solyc02g071530.1 | Solyc02g071530.2 |
| Solyc02g071610.2 | Solyc02g071610.3 |
| Solyc02g071620.2 | Solyc02g071620.3 |
| Solyc02g071670.2 | Solyc02g071670.3 |
| Solyc02g071690.1 | Solyc02g071690.2 |
| Solyc02g071700.2 | Solyc02g071700.3 |
| Solyc02g071710.2 | Solyc02g071710.3 |
| Solyc02g071740.2 | Solyc02g071740.3 |
| Solyc02g071750.2 | Solyc02g071750.3 |
| Solyc02g071770.2 | Solyc02g071770.3 |
| Solyc02g071780.2 | Solyc02g071780.3 |
| Solyc02g071800.2 | Solyc02g071800.3 |
| Solyc02g071880.2 | Solyc02g071880.3 |
| Solyc02g071890.2 | Solyc02g071890.3 |
| Solyc02g071900.2 | Solyc02g071900.3 |
| Solyc02g071920.2 | Solyc02g071920.3 |
| Solyc02g071930.1 | Solyc02g071930.2 |
| Solyc02g071960.1 | Solyc02g071960.2 |
| Solyc02g071980.2 | Solyc02g071980.3 |
| Solyc02g071990.2 | Solyc02g071990.3 |
| Solyc02g072000.2 | Solyc02g072000.3 |
| Solyc02g072090.1 | Solyc02g072090.2 |
| Solyc02g072100.2 | Solyc02g072100.3 |
| Solyc02g072120.2 | Solyc02g072120.3 |
| Solyc02g072130.2 | Solyc02g072130.3 |
| Solyc02g072150.2 | Solyc02g072150.3 |
| Solyc02g072180.2 | Solyc02g072180.3 |
| Solyc02g072230.2 | Solyc02g072230.3 |
| Solyc02g072240.2 | Solyc02g072240.3 |
| Solyc02g072270.2 | Solyc02g072270.3 |
| Solyc02g072310.2 | Solyc02g072310.3 |
| Solyc02g072490.2 | Solyc02g072490.3 |
| Solyc02g072500.2 | Solyc02g072500.3 |
| Solyc02g072520.2 | Solyc02g072520.3 |
| Solyc02g072540.2 | Solyc02g072540.3 |
| Solyc02g072570.1 | Solyc02g072570.2 |
| Solyc02g076660.2 | Solyc02g076660.3 |

| | |
|---|---|
| Solyc02g076670.2 | Solyc02g076670.3 |
| Solyc02g076680.2 | Solyc02g076680.3 |
| Solyc02g076690.2 | Solyc02g076690.3 |
| Solyc02g076710.2 | Solyc02g076710.3 |
| Solyc02g076720.2 | Solyc02g076720.3 |
| Solyc02g076730.2 | Solyc02g076730.3 |
| Solyc02g076760.2 | Solyc02g076760.3 |
| Solyc02g076780.2 | Solyc02g076780.3 |
| Solyc02g076840.2 | Solyc02g076840.3 |
| Solyc02g076850.1 | Solyc02g076850.2 |
| Solyc02g076860.2 | Solyc02g076860.3 |
| Solyc02g076910.2 | Solyc02g076910.3 |
| Solyc02g076920.2 | Solyc02g076920.3 |
| Solyc02g076930.1 | Solyc02g076930.2 |
| Solyc02g076940.1 | Solyc02g076940.2 |
| Solyc02g076980.4 | Solyc02g076980.5.1 |
| Solyc02g076990.2 | Solyc02g076990.3 |
| Solyc02g077000.2 | Solyc02g077000.3 |
| Solyc02g077020.2 | Solyc02g077020.3 |
| Solyc02g077030.2 | Solyc02g077030.3 |
| Solyc02g077060.1 | Solyc02g077060.2 |
| Solyc02g077070.2 | Solyc02g077070.3 |
| Solyc02g077080.2 | Solyc02g077080.3 |
| Solyc02g077100.2 | Solyc02g077100.3 |
| Solyc02g077110.2 | Solyc02g077110.3 |
| Solyc02g077130.1 | Solyc02g077130.2 |
| Solyc02g077140.2 | Solyc02g077140.3 |
| Solyc02g077160.2 | Solyc02g077160.3 |
| Solyc02g077200.2 | Solyc02g077200.3 |
| Solyc02g077220.2 | Solyc02g077220.3 |
| Solyc02g077260.2 | Solyc02g077260.3 |
| Solyc02g077290.1 | Solyc02g077290.2 |
| Solyc02g077300.1 | Solyc02g077300.2 |
| Solyc02g077330.2 | Solyc02g077330.3 |
| Solyc02g077380.1 | Solyc02g077380.2 |
| Solyc02g077390.1 | Solyc02g077390.2 |
| Solyc02g077410.2 | Solyc02g077410.3 |
| Solyc02g077420.2 | Solyc02g077420.3 |
| Solyc02g077440.2 | Solyc02g077440.3 |
| Solyc02g077450.2 | Solyc02g077450.3 |
| Solyc02g077490.2 | Solyc02g077490.3 |
| Solyc02g077520.2 | Solyc02g077520.3 |
| Solyc02g077530.1 | Solyc02g077530.2 |

| | |
|---|---|
| Solyc02g077560.2 | Solyc02g077560.3 |
| Solyc02g077600.2 | Solyc02g077600.3 |
| Solyc02g077610.2 | Solyc02g077610.3 |
| Solyc02g077630.2 | Solyc02g077630.3 |
| Solyc02g077650.1 | Solyc02g077650.2 |
| Solyc02g077660.2 | Solyc02g077660.3 |
| Solyc02g077670.2 | Solyc02g077670.3 |
| Solyc02g077730.2 | Solyc02g077730.3 |
| Solyc02g077740.2 | Solyc02g077740.3 |
| Solyc02g077760.1 | Solyc02g077760.2 |
| Solyc02g077770.2 | Solyc02g077770.3 |
| Solyc02g077780.2 | Solyc02g077780.3 |
| Solyc02g077800.1 | Solyc02g077800.2 |
| Solyc02g077840.1 | Solyc02g077840.2 |
| Solyc02g077850.2 | Solyc02g077850.3 |
| Solyc02g077880.2 | Solyc02g077880.3 |
| Solyc02g077990.2 | Solyc02g077990.3 |
| Solyc02g078020.2 | Solyc02g078020.3 |
| Solyc02g078040.2 | Solyc02g078040.3 |
| Solyc02g078100.2 | Solyc02g078100.3 |
| Solyc02g078120.1 | Solyc02g078120.2 |
| Solyc02g078130.2 | Solyc02g078130.3 |
| Solyc02g078140.2 | Solyc02g078140.3 |
| Solyc02g078160.2 | Solyc02g078160.3 |
| Solyc02g078180.2 | Solyc02g078180.3 |
| Solyc02g078190.2 | Solyc02g078190.3 |
| Solyc02g078210.2 | Solyc02g078210.3 |
| Solyc02g078230.1 | Solyc02g078230.2 |
| Solyc02g078260.2 | Solyc02g078260.3 |
| Solyc02g078270.2 | Solyc02g078270.3 |
| Solyc02g078300.2 | Solyc02g078300.3 |
| Solyc02g078330.2 | Solyc02g078330.3 |
| Solyc02g078350.2 | Solyc02g078350.3 |
| Solyc02g078380.2 | Solyc02g078380.3 |
| Solyc02g078390.2 | Solyc02g078390.3 |
| Solyc02g078400.2 | Solyc02g078400.3 |
| Solyc02g078440.2 | Solyc02g078440.3 |
| Solyc02g078450.2 | Solyc02g078450.3 |
| Solyc02g078460.2 | Solyc02g078460.3 |
| Solyc02g078480.2 | Solyc02g078480.3 |
| Solyc02g078490.2 | Solyc02g078490.3 |
| Solyc02g078530.2 | Solyc02g078530.3 |
| Solyc02g078550.2 | Solyc02g078550.3 |

| | |
|---|---|
| Solyc02g078570.2 | Solyc02g078570.3 |
| Solyc02g078600.2 | Solyc02g078600.3 |
| Solyc02g078640.2 | Solyc02g078640.3 |
| Solyc02g078670.2 | Solyc02g078670.3 |
| Solyc02g078690.1 | Solyc02g078690.2 |
| Solyc02g078700.2 | Solyc02g078700.3 |
| Solyc02g078750.2 | Solyc02g078750.3 |
| Solyc02g078780.2 | Solyc02g078780.3 |
| Solyc02g078820.2 | Solyc02g078820.3 |
| Solyc02g078860.2 | Solyc02g078860.3 |
| Solyc02g078900.2 | Solyc02g078900.3 |
| Solyc02g078910.2 | Solyc02g078910.3 |
| Solyc02g078940.2 | Solyc02g078940.3 |
| Solyc02g078980.2 | Solyc02g078980.3 |
| Solyc02g079030.2 | Solyc02g079030.3 |
| Solyc02g079040.2 | Solyc02g079040.3 |
| Solyc02g079050.2 | Solyc02g079050.3 |
| Solyc02g079060.2 | Solyc02g079060.3 |
| Solyc02g079080.1 | Solyc02g079080.2 |
| Solyc02g079100.2 | Solyc02g079100.3 |
| Solyc02g079110.2 | Solyc02g079110.3 |
| Solyc02g079130.2 | Solyc02g079130.3 |
| Solyc02g079140.1 | Solyc02g079140.2 |
| Solyc02g079150.1 | Solyc02g079150.2 |
| Solyc02g079170.2 | Solyc02g079170.3 |
| Solyc02g079190.2 | Solyc02g079190.3 |
| Solyc02g079210.2 | Solyc02g079210.3 |
| Solyc02g079230.2 | Solyc02g079230.3 |
| Solyc02g079260.1 | Solyc02g079260.2 |
| Solyc02g079280.2 | Solyc02g079280.3 |
| Solyc02g079290.2 | Solyc02g079290.3 |
| Solyc02g079300.2 | Solyc02g079300.3 |
| Solyc02g079320.2 | Solyc02g079320.3 |
| Solyc02g079330.2 | Solyc02g079330.3 |
| Solyc02g079340.2 | Solyc02g079340.3 |
| Solyc02g079350.2 | Solyc02g079350.3 |
| Solyc02g079370.2 | Solyc02g079370.3 |
| Solyc02g079380.1 | Solyc02g079380.2 |
| Solyc02g079400.2 | Solyc02g079400.3 |
| Solyc02g079490.2 | Solyc02g079490.3 |
| Solyc02g079540.1 | Solyc02g079540.2 |
| Solyc02g079590.2 | Solyc02g079590.3 |
| Solyc02g079600.2 | Solyc02g079600.3 |

| | |
|---|---|
| Solyc02g079640.2 | Solyc02g079640.3 |
| Solyc02g079760.2 | Solyc02g079760.3 |
| Solyc02g079770.2 | Solyc02g079770.3 |
| Solyc02g079790.2 | Solyc02g079790.3 |
| Solyc02g079840.1 | Solyc02g079840.2 |
| Solyc02g079880.2 | Solyc02g079880.3 |
| Solyc02g079890.1 | Solyc02g079890.2 |
| Solyc02g079920.2 | Solyc02g079920.3 |
| Solyc02g079930.2 | Solyc02g079930.3 |
| Solyc02g079940.2 | Solyc02g079940.3 |
| Solyc02g079950.2 | Solyc02g079950.3 |
| Solyc02g079960.2 | Solyc02g079960.3 |
| Solyc02g079980.2 | Solyc02g079980.3 |
| Solyc02g079990.2 | Solyc02g079990.3 |
| Solyc02g080030.1 | Solyc02g080030.2 |
| Solyc02g080070.2 | Solyc02g080070.3 |
| Solyc02g080080.2 | Solyc02g080080.3 |
| Solyc02g080190.2 | Solyc02g080190.3 |
| Solyc02g080220.2 | Solyc02g080220.3 |
| Solyc02g080230.2 | Solyc02g080230.3 |
| Solyc02g080240.1 | Solyc02g080240.2 |
| Solyc02g080270.2 | Solyc02g080270.3 |
| Solyc02g080290.2 | Solyc02g080290.3 |
| Solyc02g080300.2 | Solyc02g080300.3 |
| Solyc02g080340.2 | Solyc02g080340.3 |
| Solyc02g080370.2 | Solyc02g080370.3 |
| Solyc02g080410.2 | Solyc02g080410.3 |
| Solyc02g080430.2 | Solyc02g080430.3 |
| Solyc02g080440.2 | Solyc02g080440.3 |
| Solyc02g080470.2 | Solyc02g080470.3 |
| Solyc02g080480.2 | Solyc02g080480.3 |
| Solyc02g080490.2 | Solyc02g080490.3 |
| Solyc02g080530.2 | Solyc02g080530.3 |
| Solyc02g080570.2 | Solyc02g080570.3 |
| Solyc02g080590.2 | Solyc02g080590.3 |
| Solyc02g080620.2 | Solyc02g080620.3 |
| Solyc02g080650.2 | Solyc02g080650.3 |
| Solyc02g080660.2 | Solyc02g080660.3 |
| Solyc02g080670.2 | Solyc02g080670.3 |
| Solyc02g080710.1 | Solyc02g080710.2 |
| Solyc02g080730.1 | Solyc02g080730.2 |
| Solyc02g080755.1 | Solyc02g150131.1 |
| Solyc02g080810.2 | Solyc02g080810.3 |

| | |
|---|---|
| Solyc02g080820.2 | Solyc02g080820.3 |
| Solyc02g080890.2 | Solyc02g080890.3 |
| Solyc02g080900.2 | Solyc02g080900.3 |
| Solyc02g080940.2 | Solyc02g080940.3 |
| Solyc02g080990.1 | Solyc02g080990.2 |
| Solyc02g080995.1 | Solyc02g150132.1 |
| Solyc02g081080.2 | Solyc02g081080.3 |
| Solyc02g081140.2 | Solyc02g081140.3 |
| Solyc02g081160.2 | Solyc02g081160.3 |
| Solyc02g081180.1 | Solyc02g081180.2 |
| Solyc02g081210.2 | Solyc02g081210.3 |
| Solyc02g081300.2 | Solyc02g081300.3 |
| Solyc02g081310.2 | Solyc02g081310.3 |
| Solyc02g081340.2 | Solyc02g081340.3 |
| Solyc02g081380.2 | Solyc02g081380.3 |
| Solyc02g081390.2 | Solyc02g081390.3 |
| Solyc02g081400.2 | Solyc02g081400.3 |
| Solyc02g081410.2 | Solyc02g081410.3 |
| Solyc02g081420.2 | Solyc02g081420.3 |
| Solyc02g081430.2 | Solyc02g081430.3 |
| Solyc02g081500.2 | Solyc02g081500.3 |
| Solyc02g081520.2 | Solyc02g081520.3 |
| Solyc02g081535.1 | Solyc02g150133.1 |
| Solyc02g081550.2 | Solyc02g081550.3 |
| Solyc02g081580.2 | Solyc02g081580.3 |
| Solyc02g081590.2 | Solyc02g081590.3 |
| Solyc02g081620.2 | Solyc02g081620.3 |
| Solyc02g081650.2 | Solyc02g081650.3 |
| Solyc02g081660.2 | Solyc02g081660.3 |
| Solyc02g081730.2 | Solyc02g081730.3 |
| Solyc02g081740.1 | Solyc02g081740.2 |
| Solyc02g081760.1 | Solyc02g081760.2 |
| Solyc02g081770.1 | Solyc02g081770.2 |
| Solyc02g081820.2 | Solyc02g081820.3 |
| Solyc02g081880.2 | Solyc02g081880.3 |
| Solyc02g081910.1 | Solyc02g081910.2 |
| Solyc02g081920.2 | Solyc02g081920.3 |
| Solyc02g081930.2 | Solyc02g081930.3 |
| Solyc02g081940.2 | Solyc02g081940.3 |
| Solyc02g081960.2 | Solyc02g081960.3 |
| Solyc02g081980.2 | Solyc02g081980.3 |
| Solyc02g081990.2 | Solyc02g081990.3 |
| Solyc02g082000.2 | Solyc02g082000.3 |

| | |
|---|---|
| Solyc02g082020.2 | Solyc02g082020.3 |
| Solyc02g082030.2 | Solyc02g082030.3 |
| Solyc02g082040.2 | Solyc02g082040.3 |
| Solyc02g082050.2 | Solyc02g082050.3 |
| Solyc02g082060.1 | Solyc02g082060.2 |
| Solyc02g082090.2 | Solyc02g082090.3 |
| Solyc02g082120.2 | Solyc02g082120.3 |
| Solyc02g082130.1 | Solyc02g082130.2 |
| Solyc02g082140.2 | Solyc02g082140.3 |
| Solyc02g082160.1 | Solyc02g082160.2 |
| Solyc02g082180.2 | Solyc02g082180.3 |
| Solyc02g082190.2 | Solyc02g082190.3 |
| Solyc02g082220.2 | Solyc02g082220.3 |
| Solyc02g082260.2 | Solyc02g082260.3 |
| Solyc02g082270.2 | Solyc02g082270.3 |
| Solyc02g082280.2 | Solyc02g082280.3 |
| Solyc02g082300.2 | Solyc02g082300.3 |
| Solyc02g082310.1 | Solyc02g082310.2 |
| Solyc02g082315.1 | Solyc02g082315.2 |
| Solyc02g082340.2 | Solyc02g082340.3 |
| Solyc02g082350.2 | Solyc02g082350.3 |
| Solyc02g082420.2 | Solyc02g082420.3 |
| Solyc02g082440.1 | Solyc02g082440.2 |
| Solyc02g082450.2 | Solyc02g082450.3 |
| Solyc02g082460.2 | Solyc02g082460.3 |
| Solyc02g082480.2 | Solyc02g082480.3 |
| Solyc02g082550.2 | Solyc02g082550.3 |
| Solyc02g082590.2 | Solyc02g082590.3 |
| Solyc02g082660.2 | Solyc02g082660.3 |
| Solyc02g082670.2 | Solyc02g082670.3 |
| Solyc02g082690.2 | Solyc02g082690.3 |
| Solyc02g082700.2 | Solyc02g082700.3 |
| Solyc02g082730.2 | Solyc02g082730.3 |
| Solyc02g082760.2 | Solyc02g082760.3 |
| Solyc02g082763.1 | Solyc02g150134.1 |
| Solyc02g082767.1 | Solyc02g150135.1 |
| Solyc02g082770.2 | Solyc02g082770.3 |
| Solyc02g082790.1 | Solyc02g082790.2 |
| Solyc02g082800.2 | Solyc02g082800.3 |
| Solyc02g082810.2 | Solyc02g082810.3 |
| Solyc02g082820.2 | Solyc02g082820.3 |
| Solyc02g082840.2 | Solyc02g082840.3 |
| Solyc02g082850.2 | Solyc02g082850.3 |

| | |
|---|---|
| Solyc02g082860.2 | Solyc02g082860.3 |
| Solyc02g082900.2 | Solyc02g082900.3 |
| Solyc02g082930.2 | Solyc02g082930.3 |
| Solyc02g082940.2 | Solyc02g082940.3 |
| Solyc02g082950.2 | Solyc02g082950.3 |
| Solyc02g082960.2 | Solyc02g082960.3 |
| Solyc02g083140.2 | Solyc02g083140.3 |
| Solyc02g083150.2 | Solyc02g083150.3 |
| Solyc02g083170.2 | Solyc02g083170.3 |
| Solyc02g083190.1 | Solyc02g083190.2 |
| Solyc02g083210.1 | Solyc02g083210.2 |
| Solyc02g083220.1 | Solyc02g083220.2 |
| Solyc02g083250.2 | Solyc02g083250.3 |
| Solyc02g083260.2 | Solyc02g083260.3 |
| Solyc02g083285.1 | Solyc02g083285.2 |
| Solyc02g083300.1 | Solyc02g083300.2 |
| Solyc02g083310.2 | Solyc02g083310.3 |
| Solyc02g083320.2 | Solyc02g083320.3 |
| Solyc02g083330.1 | Solyc02g083330.2 |
| Solyc02g083400.2 | Solyc02g083400.3 |
| Solyc02g083420.2 | Solyc02g083420.3 |
| Solyc02g083440.2 | Solyc02g083440.3 |
| Solyc02g083450.2 | Solyc02g083450.3 |
| Solyc02g083460.2 | Solyc02g083460.3 |
| Solyc02g083470.2 | Solyc02g083470.3 |
| Solyc02g083480.2 | Solyc02g083480.3 |
| Solyc02g083490.2 | Solyc02g083490.3 |
| Solyc02g083510.2 | Solyc02g083510.3 |
| Solyc02g083550.1 | Solyc02g083550.2 |
| Solyc02g083560.2 | Solyc02g083560.3 |
| Solyc02g083570.2 | Solyc02g083570.3 |
| Solyc02g083620.2 | Solyc02g083620.3 |
| Solyc02g083630.2 | Solyc02g083630.3 |
| Solyc02g083710.2 | Solyc02g083710.3 |
| Solyc02g083740.1 | Solyc02g083740.2 |
| Solyc02g083815.1 | Solyc02g150136.1 |
| Solyc02g083830.1 | Solyc02g083830.2 |
| Solyc02g083850.2 | Solyc02g083850.3 |
| Solyc02g083860.2 | Solyc02g083860.3 |
| Solyc02g083870.2 | Solyc02g083870.3 |
| Solyc02g083880.2 | Solyc02g083880.3 |
| Solyc02g083900.2 | Solyc02g083900.3 |
| Solyc02g083940.2 | Solyc02g083940.3 |

| | |
|---|---|
| Solyc02g083950.2 | Solyc02g083950.3 |
| Solyc02g083980.2 | Solyc02g083980.3 |
| Solyc02g083990.2 | Solyc02g083990.3 |
| Solyc02g084000.2 | Solyc02g084000.3 |
| Solyc02g084020.2 | Solyc02g084020.3 |
| Solyc02g084030.2 | Solyc02g084030.3 |
| Solyc02g084117.1 | Solyc02g150137.1 |
| Solyc02g084150.1 | Solyc02g084150.2 |
| Solyc02g084170.1 | Solyc02g084170.2 |
| Solyc02g084240.2 | Solyc02g084240.3 |
| Solyc02g084265.1 | Solyc02g150138.1 |
| Solyc02g084280.2 | Solyc02g084280.3 |
| Solyc02g084345.1 | Solyc02g150139.1 |
| Solyc02g084350.2 | Solyc02g084350.3 |
| Solyc02g084360.2 | Solyc02g084360.3 |
| Solyc02g084370.1 | Solyc02g084370.2 |
| Solyc02g084380.2 | Solyc02g084380.3 |
| Solyc02g084390.2 | Solyc02g084390.3 |
| Solyc02g084410.2 | Solyc02g084410.3 |
| Solyc02g084420.2 | Solyc02g084420.3 |
| Solyc02g084430.2 | Solyc02g084430.3 |
| Solyc02g084510.2 | Solyc02g084510.3 |
| Solyc02g084520.2 | Solyc02g084520.3 |
| Solyc02g084530.1 | Solyc02g084530.2 |
| Solyc02g084540.2 | Solyc02g084540.3 |
| Solyc02g084620.2 | Solyc02g084620.3 |
| Solyc02g084630.2 | Solyc02g084630.3 |
| Solyc02g084650.2 | Solyc02g084650.3 |
| Solyc02g084660.2 | Solyc02g084660.3 |
| Solyc02g084680.2 | Solyc02g084680.3 |
| Solyc02g084690.2 | Solyc02g084690.3 |
| Solyc02g084700.2 | Solyc02g084700.3 |
| Solyc02g084710.2 | Solyc02g084710.3 |
| Solyc02g084720.2 | Solyc02g084720.3 |
| Solyc02g084730.2 | Solyc02g084730.3 |
| Solyc02g084760.2 | Solyc02g084760.3 |
| Solyc02g084770.2 | Solyc02g084770.3 |
| Solyc02g084790.2 | Solyc02g084790.3 |
| Solyc02g084820.2 | Solyc02g084820.3 |
| Solyc02g084840.2 | Solyc02g084840.3 |
| Solyc02g084850.2 | Solyc02g084850.3 |
| Solyc02g084870.2 | Solyc02g084870.3 |
| Solyc02g084880.2 | Solyc02g084880.3 |

| | |
|---|---|
| Solyc02g084920.2 | Solyc02g084920.3 |
| Solyc02g084930.2 | Solyc02g084930.3 |
| Solyc02g084950.2 | Solyc02g084950.3 |
| Solyc02g084980.2 | Solyc02g084980.3 |
| Solyc02g084990.2 | Solyc02g084990.3 |
| Solyc02g085040.2 | Solyc02g085040.3 |
| Solyc02g085050.2 | Solyc02g085050.3 |
| Solyc02g085060.1 | Solyc02g085060.2 |
| Solyc02g085100.2 | Solyc02g085100.3 |
| Solyc02g085120.2 | Solyc02g085120.3 |
| Solyc02g085180.4 | Solyc02g085180.5.1 |
| Solyc02g085190.1 | Solyc02g085190.2 |
| Solyc02g085230.2 | Solyc02g085230.3 |
| Solyc02g085290.2 | Solyc02g085290.3 |
| Solyc02g085370.2 | Solyc02g085370.3 |
| Solyc02g085380.2 | Solyc02g085380.3 |
| Solyc02g085400.2 | Solyc02g085400.3 |
| Solyc02g085470.2 | Solyc02g085470.3 |
| Solyc02g085480.3 | Solyc02g085480.4 |
| Solyc02g085490.1 | Solyc02g085490.2 |
| Solyc02g085560.2 | Solyc02g085560.3 |
| Solyc02g085620.2 | Solyc02g085620.3 |
| Solyc02g085630.2 | Solyc02g085630.3 |
| Solyc02g085670.2 | Solyc02g085670.3 |
| Solyc02g085730.2 | Solyc02g085730.3 |
| Solyc02g085780.2 | Solyc02g085780.3 |
| Solyc02g085790.2 | Solyc02g085790.3 |
| Solyc02g085800.1 | Solyc02g085800.2 |
| Solyc02g085810.2 | Solyc02g085810.3 |
| Solyc02g085820.2 | Solyc02g085820.3 |
| Solyc02g085870.2 | Solyc02g085870.3 |
| Solyc02g085930.2 | Solyc02g085930.3 |
| Solyc02g086000.1 | Solyc02g086000.2 |
| Solyc02g086020.1 | Solyc02g086020.2 |
| Solyc02g086030.1 | Solyc02g086030.2 |
| Solyc02g086040.1 | Solyc02g086040.2 |
| Solyc02g086050.2 | Solyc02g086050.3 |
| Solyc02g086220.2 | Solyc02g086220.3 |
| Solyc02g086230.1 | Solyc02g086230.2 |
| Solyc02g086240.2 | Solyc02g086240.3 |
| Solyc02g086290.1 | Solyc02g086290.2 |
| Solyc02g086310.1 | Solyc02g086310.2 |
| Solyc02g086350.2 | Solyc02g086350.3 |

| | |
|---|---|
| Solyc02g086360.2 | Solyc02g086360.3 |
| Solyc02g086445.1 | Solyc02g150140.1 |
| Solyc02g086470.2 | Solyc02g086470.3 |
| Solyc02g086490.2 | Solyc02g086490.3 |
| Solyc02g086500.2 | Solyc02g086500.3 |
| Solyc02g086520.2 | Solyc02g086520.3 |
| Solyc02g086550.2 | Solyc02g086550.3 |
| Solyc02g086560.2 | Solyc02g086560.3 |
| Solyc02g086585.1 | Solyc02g150141.1 |
| Solyc02g086590.2 | Solyc02g086590.3 |
| Solyc02g086600.2 | Solyc02g086600.3 |
| Solyc02g086620.2 | Solyc02g086620.3 |
| Solyc02g086650.2 | Solyc02g086650.3 |
| Solyc02g086690.2 | Solyc02g086690.3 |
| Solyc02g086710.2 | Solyc02g086710.3 |
| Solyc02g086770.2 | Solyc02g086770.3 |
| Solyc02g086820.2 | Solyc02g086820.3 |
| Solyc02g086830.2 | Solyc02g086830.3 |
| Solyc02g086840.2 | Solyc02g086840.3 |
| Solyc02g086890.2 | Solyc02g086890.3 |
| Solyc02g086920.1 | Solyc02g086920.2 |
| Solyc02g086940.2 | Solyc02g086940.3 |
| Solyc02g086980.2 | Solyc02g086980.3 |
| Solyc02g087110.2 | Solyc02g087110.3 |
| Solyc02g087140.2 | Solyc02g087140.3 |
| Solyc02g087160.2 | Solyc02g087160.3 |
| Solyc02g087200.1 | Solyc02g087200.2 |
| Solyc02g087210.2 | Solyc02g087210.3 |
| Solyc02g087230.2 | Solyc02g087230.3 |
| Solyc02g087280.2 | Solyc02g087280.3 |
| Solyc02g087290.2 | Solyc02g087290.3 |
| Solyc02g087310.2 | Solyc02g087310.3 |
| Solyc02g087320.1 | Solyc02g087320.2 |
| Solyc02g087330.2 | Solyc02g087330.3 |
| Solyc02g087340.2 | Solyc02g087340.3 |
| Solyc02g087510.2 | Solyc02g087510.3 |
| Solyc02g087520.2 | Solyc02g087520.3 |
| Solyc02g087530.1 | Solyc02g087530.2 |
| Solyc02g087550.2 | Solyc02g087550.3 |
| Solyc02g087580.2 | Solyc02g087580.3 |
| Solyc02g087600.2 | Solyc02g087600.3 |
| Solyc02g087620.2 | Solyc02g087620.3 |
| Solyc02g087630.2 | Solyc02g087630.3 |

| | |
|---|---|
| Solyc02g087640.2 | Solyc02g087640.3 |
| Solyc02g087665.1 | Solyc02g150142.1 |
| Solyc02g087670.2 | Solyc02g087670.3 |
| Solyc02g087710.2 | Solyc02g087710.3 |
| Solyc02g087730.2 | Solyc02g087730.3 |
| Solyc02g087740.2 | Solyc02g087740.3 |
| Solyc02g087750.2 | Solyc02g087750.3 |
| Solyc02g087760.2 | Solyc02g087760.3 |
| Solyc02g087770.2 | Solyc02g087770.3 |
| Solyc02g087780.2 | Solyc02g087780.3 |
| Solyc02g087790.1 | Solyc02g087790.2 |
| Solyc02g087800.2 | Solyc02g087800.3 |
| Solyc02g087810.2 | Solyc02g087810.3 |
| Solyc02g087840.2 | Solyc02g087840.3 |
| Solyc02g087860.2 | Solyc02g087860.3 |
| Solyc02g087870.2 | Solyc02g087870.3 |
| Solyc02g087880.2 | Solyc02g087880.3 |
| Solyc02g087900.2 | Solyc02g087900.3 |
| Solyc02g087920.2 | Solyc02g087920.3 |
| Solyc02g087930.2 | Solyc02g087930.3 |
| Solyc02g087950.2 | Solyc02g087950.3 |
| Solyc02g087960.2 | Solyc02g087960.3 |
| Solyc02g087980.2 | Solyc02g087980.3 |
| Solyc02g088000.2 | Solyc02g088000.3 |
| Solyc02g088010.2 | Solyc02g088010.3 |
| Solyc02g088070.2 | Solyc02g088070.3 |
| Solyc02g088080.1 | Solyc02g088080.2 |
| Solyc02g088100.2 | Solyc02g088100.3 |
| Solyc02g088170.1 | Solyc02g088170.2 |
| Solyc02g088180.2 | Solyc02g088180.3 |
| Solyc02g088210.2 | Solyc02g088210.3 |
| Solyc02g088220.2 | Solyc02g088220.3 |
| Solyc02g088230.2 | Solyc02g088230.3 |
| Solyc02g088280.2 | Solyc02g088280.3 |
| Solyc02g088300.2 | Solyc02g088300.3 |
| Solyc02g088395.1 | Solyc02g150143.1 |
| Solyc02g088425.1 | Solyc02g150144.1 |
| Solyc02g088490.2 | Solyc02g088490.3 |
| Solyc02g088540.2 | Solyc02g088540.3 |
| Solyc02g088570.1 | Solyc02g088570.2 |
| Solyc02g088620.2 | Solyc02g088620.3 |
| Solyc02g088710.2 | Solyc02g088710.3 |
| Solyc02g088730.1 | Solyc02g088730.2 |

| | |
|---|---|
| Solyc02g088750.1 | Solyc02g088750.2 |
| Solyc02g088770.2 | Solyc02g088770.3 |
| Solyc02g088790.2 | Solyc02g088790.3 |
| Solyc02g088800.1 | Solyc02g088800.2 |
| Solyc02g088840.2 | Solyc02g088840.3 |
| Solyc02g088850.2 | Solyc02g088850.3 |
| Solyc02g088860.1 | Solyc02g088860.2 |
| Solyc02g088890.1 | Solyc02g088890.2 |
| Solyc02g088900.2 | Solyc02g088900.3 |
| Solyc02g088910.2 | Solyc02g088910.3 |
| Solyc02g088940.2 | Solyc02g088940.3 |
| Solyc02g088983.1 | Solyc02g150145.1 |
| Solyc02g088987.1 | Solyc02g150146.1 |
| Solyc02g089020.1 | Solyc02g089020.2 |
| Solyc02g089040.2 | Solyc02g089040.3 |
| Solyc02g089060.2 | Solyc02g089060.3 |
| Solyc02g089070.2 | Solyc02g089070.3 |
| Solyc02g089090.2 | Solyc02g089090.3 |
| Solyc02g089110.2 | Solyc02g089110.3 |
| Solyc02g089140.2 | Solyc02g089140.3 |
| Solyc02g089160.2 | Solyc02g089160.3 |
| Solyc02g089175.1 | Solyc02g089175.2 |
| Solyc02g089190.1 | Solyc02g089190.2 |
| Solyc02g089230.2 | Solyc02g089230.3 |
| Solyc02g089240.2 | Solyc02g089240.3 |
| Solyc02g089280.2 | Solyc02g089280.3 |
| Solyc02g089290.2 | Solyc02g089290.3 |
| Solyc02g089300.2 | Solyc02g089300.3 |
| Solyc02g089320.2 | Solyc02g089320.3 |
| Solyc02g089330.2 | Solyc02g089330.3 |
| Solyc02g089350.2 | Solyc02g089350.3 |
| Solyc02g089410.2 | Solyc02g089410.3 |
| Solyc02g089440.2 | Solyc02g089440.3 |
| Solyc02g089510.2 | Solyc02g089510.3 |
| Solyc02g089520.1 | Solyc02g089520.2 |
| Solyc02g089530.2 | Solyc02g089530.3 |
| Solyc02g089540.2 | Solyc02g089540.3 |
| Solyc02g089550.2 | Solyc02g089550.3 |
| Solyc02g089570.2 | Solyc02g089570.3 |
| Solyc02g089590.2 | Solyc02g089590.3 |
| Solyc02g089600.2 | Solyc02g089600.3 |
| Solyc02g089610.1 | Solyc02g089610.2 |
| Solyc02g089615.1 | Solyc02g150147.1 |

| | |
|---|---|
| Solyc02g089620.2 | Solyc02g089620.3 |
| Solyc02g089630.2 | Solyc02g089630.3 |
| Solyc02g089640.2 | Solyc02g089640.3 |
| Solyc02g089690.2 | Solyc02g089690.3 |
| Solyc02g089710.2 | Solyc02g089710.3 |
| Solyc02g089740.2 | Solyc02g089740.3 |
| Solyc02g089750.2 | Solyc02g089750.4 |
| Solyc02g089760.2 | Solyc02g089760.3 |
| Solyc02g089800.2 | Solyc02g089800.3 |
| Solyc02g089810.2 | Solyc02g089810.3 |
| Solyc02g089830.1 | Solyc02g089830.2 |
| Solyc02g089840.2 | Solyc02g089840.3 |
| Solyc02g089850.2 | Solyc02g089850.3 |
| Solyc02g089950.1 | Solyc02g089950.2 |
| Solyc02g089970.2 | Solyc02g089970.3 |
| Solyc02g090000.2 | Solyc02g090000.3 |
| Solyc02g090040.1 | Solyc02g090040.2 |
| Solyc02g090070.2 | Solyc02g090070.3 |
| Solyc02g090090.2 | Solyc02g090090.3 |
| Solyc02g090110.2 | Solyc02g090110.3 |
| Solyc02g090160.2 | Solyc02g090160.3 |
| Solyc02g090170.2 | Solyc02g090170.3 |
| Solyc02g090180.2 | Solyc02g090180.3 |
| Solyc02g090200.2 | Solyc02g090200.3 |
| Solyc02g090210.2 | Solyc02g090210.3 |
| Solyc02g090220.2 | Solyc02g090220.3 |
| Solyc02g090290.2 | Solyc02g090290.3 |
| Solyc02g090300.2 | Solyc02g090300.3 |
| Solyc02g090330.2 | Solyc02g090330.3 |
| Solyc02g090360.2 | Solyc02g090360.3 |
| Solyc02g090430.2 | Solyc02g090430.3 |
| Solyc02g090440.2 | Solyc02g090440.3 |
| Solyc02g090450.2 | Solyc02g090450.3 |
| Solyc02g090490.2 | Solyc02g090490.3 |
| Solyc02g090500.2 | Solyc02g090500.3 |
| Solyc02g090510.2 | Solyc02g090510.3 |
| Solyc02g090560.2 | Solyc02g090560.3 |
| Solyc02g090600.1 | Solyc02g090600.2 |
| Solyc02g090620.2 | Solyc02g090620.3 |
| Solyc02g090670.2 | Solyc02g090670.3 |
| Solyc02g090680.2 | Solyc02g090680.3 |
| Solyc02g090710.2 | Solyc02g090710.3 |
| Solyc02g090720.2 | Solyc02g090720.3 |

| | |
|---|---|
| Solyc02g090730.2 | Solyc02g090730.3 |
| Solyc02g090740.2 | Solyc02g090740.3 |
| Solyc02g090820.2 | Solyc02g090820.3 |
| Solyc02g090840.2 | Solyc02g090840.3 |
| Solyc02g090850.2 | Solyc02g090850.3 |
| Solyc02g090860.2 | Solyc02g090860.3 |
| Solyc02g090900.2 | Solyc02g090900.3 |
| Solyc02g090910.2 | Solyc02g090910.3 |
| Solyc02g090930.2 | Solyc02g090930.3 |
| Solyc02g090940.2 | Solyc02g090940.3 |
| Solyc02g091050.2 | Solyc02g091050.3 |
| Solyc02g091070.2 | Solyc02g091070.3 |
| Solyc02g091080.1 | Solyc02g091080.2 |
| Solyc02g091100.2 | Solyc02g091100.3 |
| Solyc02g091110.2 | Solyc02g091110.3 |
| Solyc02g091130.2 | Solyc02g091130.3 |
| Solyc02g091140.2 | Solyc02g091140.3 |
| Solyc02g091270.2 | Solyc02g091270.3 |
| Solyc02g091280.2 | Solyc02g091280.3 |
| Solyc02g091290.2 | Solyc02g091290.3 |
| Solyc02g091300.2 | Solyc02g091300.3 |
| Solyc02g091310.1 | Solyc02g091310.2 |
| Solyc02g091410.2 | Solyc02g091410.3 |
| Solyc02g091420.2 | Solyc02g091420.3 |
| Solyc02g091440.1 | Solyc02g091440.2 |
| Solyc02g091460.2 | Solyc02g091460.3 |
| Solyc02g091470.2 | Solyc02g091470.3 |
| Solyc02g091490.2 | Solyc02g091490.4 |
| Solyc02g091510.2 | Solyc02g091510.3 |
| Solyc02g091520.2 | Solyc02g091520.3 |
| Solyc02g091550.1 | Solyc02g091550.2 |
| Solyc02g091560.2 | Solyc02g091560.3 |
| Solyc02g091640.2 | Solyc02g091640.3 |
| Solyc02g091655.1 | Solyc02g150148.1 |
| Solyc02g091660.2 | Solyc02g091660.3 |
| Solyc02g091690.2 | Solyc02g091690.3 |
| Solyc02g091700.2 | Solyc02g091700.3 |
| Solyc02g091710.2 | Solyc02g091710.3 |
| Solyc02g091720.1 | Solyc02g091720.2 |
| Solyc02g091750.2 | Solyc02g091750.3 |
| Solyc02g091790.2 | Solyc02g091790.3 |
| Solyc02g091810.1 | Solyc02g091810.2 |
| Solyc02g091830.2 | Solyc02g091830.3 |

| | |
|---|---|
| Solyc02g091840.2 | Solyc02g091840.3 |
| Solyc02g091860.2 | Solyc02g091860.3 |
| Solyc02g091870.2 | Solyc02g091870.3 |
| Solyc02g091910.1 | Solyc02g091910.2 |
| Solyc02g091920.2 | Solyc02g091920.3 |
| Solyc02g091930.2 | Solyc02g091930.3 |
| Solyc02g091980.1 | Solyc02g091980.2 |
| Solyc02g091990.2 | Solyc02g091990.3 |
| Solyc02g092050.2 | Solyc02g092050.3 |
| Solyc02g092070.2 | Solyc02g092070.3 |
| Solyc02g092080.1 | Solyc02g092080.2 |
| Solyc02g092110.2 | Solyc02g092110.3 |
| Solyc02g092120.2 | Solyc02g092120.3 |
| Solyc02g092150.2 | Solyc02g092150.3 |
| Solyc02g092160.2 | Solyc02g092160.3 |
| Solyc02g092230.2 | Solyc02g092230.3 |
| Solyc02g092250.2 | Solyc02g092250.3 |
| Solyc02g092260.2 | Solyc02g092260.3 |
| Solyc02g092270.2 | Solyc02g092270.3 |
| Solyc02g092330.2 | Solyc02g092330.3 |
| Solyc02g092350.2 | Solyc02g092350.3 |
| Solyc02g092360.2 | Solyc02g092360.3 |
| Solyc02g092410.2 | Solyc02g092410.3 |
| Solyc02g092440.2 | Solyc02g092440.3 |
| Solyc02g092450.2 | Solyc02g092450.3 |
| Solyc02g092460.2 | Solyc02g092460.3 |
| Solyc02g092470.2 | Solyc02g092470.3 |
| Solyc02g092490.2 | Solyc02g092490.3 |
| Solyc02g092510.2 | Solyc02g092510.3 |
| Solyc02g092515.1 | Solyc02g150149.1 |
| Solyc02g092550.2 | Solyc02g092550.3 |
| Solyc02g092560.2 | Solyc02g092560.3 |
| Solyc02g092580.2 | Solyc02g092580.3 |
| Solyc02g092600.2 | Solyc02g092600.3 |
| Solyc02g092610.2 | Solyc02g092610.3 |
| Solyc02g092620.2 | Solyc02g092620.3 |
| Solyc02g092650.2 | Solyc02g092650.3 |
| Solyc02g092660.2 | Solyc02g092660.3 |
| Solyc02g092710.2 | Solyc02g092710.3 |
| Solyc02g092750.2 | Solyc02g092750.3 |
| Solyc02g092760.2 | Solyc02g092760.3 |
| Solyc02g092770.2 | Solyc02g092770.3 |
| Solyc02g092790.2 | Solyc02g092790.3 |

| | |
|---|---|
| Solyc02g092800.2 | Solyc02g092800.3 |
| Solyc02g092850.1 | Solyc02g092850.2 |
| Solyc02g092860.2 | Solyc02g092860.3 |
| Solyc02g092870.1 | Solyc02g092870.2 |
| Solyc02g092880.2 | Solyc02g092880.3 |
| Solyc02g092910.2 | Solyc02g092910.3 |
| Solyc02g092940.2 | Solyc02g092940.3 |
| Solyc02g092950.2 | Solyc02g092950.3 |
| Solyc02g092970.2 | Solyc02g092970.3 |
| Solyc02g092980.2 | Solyc02g092980.3 |
| Solyc02g093000.2 | Solyc02g093000.3 |
| Solyc02g093010.2 | Solyc02g093010.3 |
| Solyc02g093020.2 | Solyc02g093020.3 |
| Solyc02g093030.2 | Solyc02g093030.3 |
| Solyc02g093050.2 | Solyc02g093050.3 |
| Solyc02g093060.2 | Solyc02g093060.3 |
| Solyc02g093070.2 | Solyc02g093070.3 |
| Solyc02g093080.2 | Solyc02g093080.3 |
| Solyc02g093100.2 | Solyc02g093100.3 |
| Solyc02g093110.2 | Solyc02g093110.3 |
| Solyc02g093150.2 | Solyc02g093150.3 |
| Solyc02g093160.2 | Solyc02g093160.3 |
| Solyc02g093180.2 | Solyc02g093180.3 |
| Solyc02g093190.2 | Solyc02g093190.3 |
| Solyc02g093210.2 | Solyc02g093210.3 |
| Solyc02g093220.2 | Solyc02g093220.3 |
| Solyc02g093250.2 | Solyc02g093250.3 |
| Solyc02g093290.2 | Solyc02g093290.3 |
| Solyc02g093300.2 | Solyc02g093300.3 |
| Solyc02g093310.2 | Solyc02g093310.3 |
| Solyc02g093340.2 | Solyc02g093340.3 |
| Solyc02g093360.2 | Solyc02g093360.3 |
| Solyc02g093430.2 | Solyc02g093430.3 |
| Solyc02g093490.2 | Solyc02g093490.3 |
| Solyc02g093510.2 | Solyc02g093510.3 |
| Solyc02g093520.2 | Solyc02g093520.3 |
| Solyc02g093550.2 | Solyc02g093550.3 |
| Solyc02g093590.2 | Solyc02g093590.3 |
| Solyc02g093600.2 | Solyc02g093600.3 |
| Solyc02g093610.2 | Solyc02g093610.3 |
| Solyc02g093620.1 | Solyc02g093620.2 |
| Solyc02g093630.2 | Solyc02g093630.3 |
| Solyc02g093640.2 | Solyc02g093640.3 |

| | |
|---|---|
| Solyc02g093670.2 | Solyc02g093670.3 |
| Solyc02g093680.2 | Solyc02g093680.3 |
| Solyc02g093690.2 | Solyc02g093690.3 |
| Solyc02g093700.2 | Solyc02g093700.3 |
| Solyc02g093720.2 | Solyc02g093720.3 |
| Solyc02g093740.2 | Solyc02g093740.3 |
| Solyc02g093750.2 | Solyc02g093750.3 |
| Solyc02g093770.2 | Solyc02g093770.3 |
| Solyc02g093800.2 | Solyc02g093800.3 |
| Solyc02g093810.2 | Solyc02g093810.3 |
| Solyc02g093840.2 | Solyc02g093840.3 |
| Solyc02g093850.2 | Solyc02g093850.3 |
| Solyc02g093860.2 | Solyc02g093860.3 |
| Solyc02g093880.2 | Solyc02g093880.3 |
| Solyc02g093900.2 | Solyc02g093900.3 |
| Solyc02g093910.2 | Solyc02g093910.3 |
| Solyc02g093940.2 | Solyc02g093940.3 |
| Solyc02g093950.2 | Solyc02g093950.3 |
| Solyc02g093960.2 | Solyc02g093960.3 |
| Solyc02g093980.2 | Solyc02g093980.3 |
| Solyc02g094010.1 | Solyc02g094010.2 |
| Solyc02g094030.2 | Solyc02g094030.3 |
| Solyc02g094040.2 | Solyc02g094040.3 |
| Solyc02g094120.2 | Solyc02g094120.3 |
| Solyc02g094130.2 | Solyc02g094130.3 |
| Solyc02g094170.2 | Solyc02g094170.3 |
| Solyc02g094180.2 | Solyc02g094180.3 |
| Solyc02g094190.2 | Solyc02g094190.3 |
| Solyc02g094200.2 | Solyc02g094200.3 |
| Solyc02g094220.1 | Solyc02g094220.2 |
| Solyc02g094255.1 | Solyc02g150150.1 |
| Solyc02g094270.1 | Solyc02g094270.2 |
| Solyc02g094320.2 | Solyc02g094320.3 |
| Solyc02g094330.2 | Solyc02g094330.3 |
| Solyc02g094350.2 | Solyc02g094350.3 |
| Solyc02g094360.2 | Solyc02g094360.3 |
| Solyc02g094390.2 | Solyc02g094390.3 |
| Solyc02g094420.2 | Solyc02g094420.3 |
| Solyc02g094460.1 | Solyc02g094460.2 |
| Solyc02g094470.2 | Solyc02g094470.3 |
| Solyc02g094500.2 | Solyc02g094500.3 |
| Solyc02g094520.2 | Solyc02g094520.3 |
| Solyc02g094560.2 | Solyc02g094560.3 |

| | |
|---|---|
| Solyc02g094570.2 | Solyc02g094570.3 |
| Solyc02g094580.2 | Solyc02g094580.3 |
| Solyc02g094590.2 | Solyc02g094590.3 |
| Solyc03g005020.2 | Solyc03g005020.3 |
| Solyc03g005050.2 | Solyc03g005050.3 |
| Solyc03g005120.2 | Solyc03g005120.3 |
| Solyc03g005140.1 | Solyc03g005140.2 |
| Solyc03g005150.2 | Solyc03g005150.3 |
| Solyc03g005170.2 | Solyc03g005170.3 |
| Solyc03g005190.2 | Solyc03g005190.3 |
| Solyc03g005200.2 | Solyc03g005200.3 |
| Solyc03g005230.2 | Solyc03g005230.3 |
| Solyc03g005240.1 | Solyc03g005240.2 |
| Solyc03g005260.2 | Solyc03g005260.3 |
| Solyc03g005270.2 | Solyc03g005270.3 |
| Solyc03g005290.1 | Solyc03g005290.2 |
| Solyc03g005300.2 | Solyc03g005300.3 |
| Solyc03g005320.2 | Solyc03g005320.3 |
| Solyc03g005350.2 | Solyc03g005350.3 |
| Solyc03g005380.2 | Solyc03g005380.3 |
| Solyc03g005390.2 | Solyc03g005390.3 |
| Solyc03g005420.1 | Solyc03g005420.2 |
| Solyc03g005440.2 | Solyc03g005440.3 |
| Solyc03g005450.2 | Solyc03g005450.3 |
| Solyc03g005470.2 | Solyc03g005470.3 |
| Solyc03g005510.1 | Solyc03g005510.2 |
| Solyc03g005550.2 | Solyc03g005550.3 |
| Solyc03g005570.2 | Solyc03g005570.3 |
| Solyc03g005590.1 | Solyc03g005590.2 |
| Solyc03g005600.2 | Solyc03g005600.3 |
| Solyc03g005630.2 | Solyc03g005630.3 |
| Solyc03g005670.2 | Solyc03g005670.3 |
| Solyc03g005680.2 | Solyc03g005680.3 |
| Solyc03g005690.2 | Solyc03g005690.3 |
| Solyc03g005720.2 | Solyc03g005720.3 |
| Solyc03g005750.2 | Solyc03g005750.3 |
| Solyc03g005820.2 | Solyc03g005820.3 |
| Solyc03g005880.2 | Solyc03g005880.3 |
| Solyc03g005910.2 | Solyc03g005910.3 |
| Solyc03g005930.2 | Solyc03g005930.3 |
| Solyc03g005960.2 | Solyc03g005960.3 |
| Solyc03g005980.2 | Solyc03g005980.3 |
| Solyc03g005990.2 | Solyc03g005990.3 |

| | |
|---|---|
| Solyc03g006000.2 | Solyc03g006000.3 |
| Solyc03g006020.2 | Solyc03g006020.3 |
| Solyc03g006040.2 | Solyc03g006040.3 |
| Solyc03g006060.2 | Solyc03g006060.3 |
| Solyc03g006070.2 | Solyc03g006070.3 |
| Solyc03g006110.2 | Solyc03g006110.3 |
| Solyc03g006130.2 | Solyc03g006130.3 |
| Solyc03g006140.2 | Solyc03g006140.3 |
| Solyc03g006177.1 | Solyc03g006177.2 |
| Solyc03g006210.1 | Solyc03g006210.2 |
| Solyc03g006240.2 | Solyc03g006240.3 |
| Solyc03g006270.2 | Solyc03g006270.3 |
| Solyc03g006350.2 | Solyc03g006350.3 |
| Solyc03g006360.2 | Solyc03g006360.3 |
| Solyc03g006370.2 | Solyc03g006370.3 |
| Solyc03g006390.1 | Solyc03g006390.2 |
| Solyc03g006400.2 | Solyc03g006400.3 |
| Solyc03g006410.2 | Solyc03g006410.3 |
| Solyc03g006430.2 | Solyc03g006430.3 |
| Solyc03g006450.1 | Solyc03g006450.2 |
| Solyc03g006470.2 | Solyc03g006470.3 |
| Solyc03g006490.2 | Solyc03g006490.3 |
| Solyc03g006500.2 | Solyc03g006500.3 |
| Solyc03g006520.2 | Solyc03g006520.3 |
| Solyc03g006530.2 | Solyc03g006530.3 |
| Solyc03g006560.2 | Solyc03g006560.3 |
| Solyc03g006570.2 | Solyc03g006570.3 |
| Solyc03g006580.2 | Solyc03g006580.3 |
| Solyc03g006610.2 | Solyc03g006610.3 |
| Solyc03g006620.2 | Solyc03g006620.3 |
| Solyc03g006630.2 | Solyc03g006630.3 |
| Solyc03g006640.2 | Solyc03g006640.3 |
| Solyc03g006650.1 | Solyc03g006650.2 |
| Solyc03g006660.2 | Solyc03g006660.3 |
| Solyc03g006680.2 | Solyc03g006680.3 |
| Solyc03g006700.2 | Solyc03g006700.3 |
| Solyc03g006810.2 | Solyc03g006810.3 |
| Solyc03g006820.2 | Solyc03g006820.3 |
| Solyc03g006830.2 | Solyc03g006830.3 |
| Solyc03g006860.2 | Solyc03g006860.3 |
| Solyc03g006870.2 | Solyc03g006870.3 |
| Solyc03g006880.2 | Solyc03g006880.3 |
| Solyc03g006905.1 | Solyc03g150100.1 |

| | |
|---|---|
| Solyc03g006910.2 | Solyc03g006910.3 |
| Solyc03g007030.2 | Solyc03g007030.3 |
| Solyc03g007050.2 | Solyc03g007050.3 |
| Solyc03g007060.2 | Solyc03g007060.3 |
| Solyc03g007070.2 | Solyc03g007070.3 |
| Solyc03g007100.2 | Solyc03g007100.3 |
| Solyc03g007110.2 | Solyc03g007110.3 |
| Solyc03g007120.2 | Solyc03g007120.3 |
| Solyc03g007140.2 | Solyc03g007140.3 |
| Solyc03g007150.2 | Solyc03g007150.3 |
| Solyc03g007170.2 | Solyc03g007170.3 |
| Solyc03g007190.2 | Solyc03g007190.3 |
| Solyc03g007200.1 | Solyc03g007200.2 |
| Solyc03g007210.2 | Solyc03g007210.3 |
| Solyc03g007220.2 | Solyc03g007220.3 |
| Solyc03g007240.2 | Solyc03g007240.3 |
| Solyc03g007260.2 | Solyc03g007260.3 |
| Solyc03g007270.2 | Solyc03g007270.3 |
| Solyc03g007310.2 | Solyc03g007310.3 |
| Solyc03g007330.2 | Solyc03g007330.3 |
| Solyc03g007360.2 | Solyc03g007360.3 |
| Solyc03g007370.2 | Solyc03g007370.3 |
| Solyc03g007380.1 | Solyc03g007380.2 |
| Solyc03g007400.1 | Solyc03g007400.2 |
| Solyc03g007430.2 | Solyc03g007430.3 |
| Solyc03g007450.1 | Solyc03g007450.2 |
| Solyc03g007480.2 | Solyc03g007480.3 |
| Solyc03g007490.2 | Solyc03g007490.3 |
| Solyc03g007530.2 | Solyc03g007530.3 |
| Solyc03g007600.2 | Solyc03g007600.3 |
| Solyc03g007620.2 | Solyc03g007620.3 |
| Solyc03g007680.2 | Solyc03g007680.3 |
| Solyc03g007740.2 | Solyc03g007740.3 |
| Solyc03g007750.2 | Solyc03g007750.3 |
| Solyc03g007760.2 | Solyc03g007760.3 |
| Solyc03g007770.2 | Solyc03g007770.3 |
| Solyc03g007790.2 | Solyc03g007790.3 |
| Solyc03g007800.2 | Solyc03g007800.3 |
| Solyc03g007810.2 | Solyc03g007810.3 |
| Solyc03g007820.2 | Solyc03g007820.3 |
| Solyc03g007830.2 | Solyc03g007830.3 |
| Solyc03g007870.2 | Solyc03g007870.3 |
| Solyc03g007890.2 | Solyc03g007890.3 |

| | |
|---|---|
| Solyc03g007900.2 | Solyc03g007900.3 |
| Solyc03g007950.2 | Solyc03g007950.3 |
| Solyc03g007960.2 | Solyc03g007960.3 |
| Solyc03g007970.1 | Solyc03g007970.2 |
| Solyc03g007990.1 | Solyc03g007990.2 |
| Solyc03g008010.2 | Solyc03g008010.3 |
| Solyc03g025270.2 | Solyc03g025270.3 |
| Solyc03g025290.2 | Solyc03g025290.3 |
| Solyc03g025310.2 | Solyc03g025310.3 |
| Solyc03g025350.2 | Solyc03g025350.3 |
| Solyc03g025360.2 | Solyc03g025360.3 |
| Solyc03g025370.2 | Solyc03g025370.3 |
| Solyc03g025380.2 | Solyc03g025380.3 |
| Solyc03g025390.2 | Solyc03g025390.3 |
| Solyc03g025400.2 | Solyc03g025400.3 |
| Solyc03g025420.2 | Solyc03g025420.3 |
| Solyc03g025440.2 | Solyc03g025440.3 |
| Solyc03g025490.2 | Solyc03g025490.3 |
| Solyc03g025510.2 | Solyc03g025510.3 |
| Solyc03g025540.2 | Solyc03g025540.3 |
| Solyc03g025550.2 | Solyc03g025550.3 |
| Solyc03g025600.2 | Solyc03g025600.3 |
| Solyc03g025630.2 | Solyc03g025630.3 |
| Solyc03g025640.1 | Solyc03g025640.2 |
| Solyc03g025650.1 | Solyc03g025650.2 |
| Solyc03g025670.2 | Solyc03g025670.3 |
| Solyc03g025680.2 | Solyc03g025680.3 |
| Solyc03g025700.1 | Solyc03g025700.2 |
| Solyc03g025710.2 | Solyc03g025710.3 |
| Solyc03g025720.2 | Solyc03g025720.3 |
| Solyc03g025730.2 | Solyc03g025730.3 |
| Solyc03g025780.2 | Solyc03g025780.3 |
| Solyc03g025800.2 | Solyc03g025800.3 |
| Solyc03g025840.2 | Solyc03g025840.3 |
| Solyc03g025850.2 | Solyc03g025850.3 |
| Solyc03g025860.2 | Solyc03g025860.3 |
| Solyc03g025870.2 | Solyc03g025870.3 |
| Solyc03g025880.2 | Solyc03g025880.3 |
| Solyc03g025890.1 | Solyc03g025890.2 |
| Solyc03g025900.2 | Solyc03g025900.3 |
| Solyc03g025923.1 | Solyc03g150102.1 |
| Solyc03g025925.1 | Solyc03g150103.1 |
| Solyc03g025927.1 | Solyc03g150104.1 |

| | |
|---|---|
| Solyc03g025970.2 | Solyc03g025970.3 |
| Solyc03g025980.2 | Solyc03g025980.3 |
| Solyc03g026020.2 | Solyc03g026020.3 |
| Solyc03g026050.2 | Solyc03g026050.3 |
| Solyc03g026060.2 | Solyc03g026060.3 |
| Solyc03g026080.2 | Solyc03g026080.3 |
| Solyc03g026120.2 | Solyc03g026120.3 |
| Solyc03g026130.2 | Solyc03g026130.3 |
| Solyc03g026150.2 | Solyc03g026150.3 |
| Solyc03g026190.2 | Solyc03g026190.3 |
| Solyc03g026220.2 | Solyc03g026220.3 |
| Solyc03g026260.2 | Solyc03g026260.3 |
| Solyc03g026290.2 | Solyc03g026290.3 |
| Solyc03g026350.2 | Solyc03g026350.3 |
| Solyc03g026377.1 | Solyc03g150105.1 |
| Solyc03g031450.2 | Solyc03g031450.3 |
| Solyc03g031470.2 | Solyc03g031470.3 |
| Solyc03g031480.2 | Solyc03g031480.3 |
| Solyc03g031540.1 | Solyc03g031540.2 |
| Solyc03g031600.2 | Solyc03g031600.3 |
| Solyc03g031620.2 | Solyc03g031620.3 |
| Solyc03g031630.2 | Solyc03g031630.3 |
| Solyc03g031640.1 | Solyc03g031640.2 |
| Solyc03g031670.2 | Solyc03g031670.3 |
| Solyc03g031690.2 | Solyc03g031690.3 |
| Solyc03g031720.2 | Solyc03g031720.3 |
| Solyc03g031730.2 | Solyc03g031730.3 |
| Solyc03g031750.1 | Solyc03g031750.2 |
| Solyc03g031770.2 | Solyc03g031770.3 |
| Solyc03g031800.2 | Solyc03g031800.3 |
| Solyc03g031810.2 | Solyc03g031810.3 |
| Solyc03g031840.2 | Solyc03g031840.3 |
| Solyc03g031860.2 | Solyc03g031860.3 |
| Solyc03g031880.2 | Solyc03g031880.3 |
| Solyc03g031890.2 | Solyc03g031890.3 |
| Solyc03g031900.1 | Solyc03g031900.2 |
| Solyc03g031910.2 | Solyc03g031910.3 |
| Solyc03g031920.2 | Solyc03g031920.3 |
| Solyc03g031960.1 | Solyc03g031960.2 |
| Solyc03g032040.2 | Solyc03g032040.3 |
| Solyc03g032070.2 | Solyc03g032070.3 |
| Solyc03g032150.2 | Solyc03g032150.3 |
| Solyc03g032160.2 | Solyc03g032160.3 |

| | |
|---|---|
| Solyc03g032170.1 | Solyc03g032170.2 |
| Solyc03g032180.2 | Solyc03g032180.3 |
| Solyc03g032190.2 | Solyc03g032190.3 |
| Solyc03g032210.2 | Solyc03g032210.3 |
| Solyc03g032220.2 | Solyc03g032220.3 |
| Solyc03g033330.2 | Solyc03g033330.3 |
| Solyc03g033340.2 | Solyc03g033340.3 |
| Solyc03g033350.2 | Solyc03g033350.3 |
| Solyc03g033370.2 | Solyc03g033370.3 |
| Solyc03g033380.2 | Solyc03g033380.3 |
| Solyc03g033390.2 | Solyc03g033390.3 |
| Solyc03g033460.2 | Solyc03g033460.3 |
| Solyc03g033470.2 | Solyc03g033470.3 |
| Solyc03g033510.2 | Solyc03g033510.3 |
| Solyc03g033540.2 | Solyc03g033540.3 |
| Solyc03g033550.2 | Solyc03g033550.3 |
| Solyc03g033560.2 | Solyc03g033560.3 |
| Solyc03g033580.2 | Solyc03g033580.3 |
| Solyc03g033600.2 | Solyc03g033600.3 |
| Solyc03g033620.2 | Solyc03g033620.3 |
| Solyc03g033630.2 | Solyc03g033630.3 |
| Solyc03g033640.2 | Solyc03g033640.3 |
| Solyc03g033690.1 | Solyc03g033690.2 |
| Solyc03g033710.2 | Solyc03g033710.3 |
| Solyc03g033750.1 | Solyc03g033750.2 |
| Solyc03g033850.2 | Solyc03g033850.3 |
| Solyc03g034000.2 | Solyc03g034000.3 |
| Solyc03g034030.2 | Solyc03g034030.3 |
| Solyc03g034050.2 | Solyc03g034050.3 |
| Solyc03g034090.2 | Solyc03g034090.3 |
| Solyc03g034120.2 | Solyc03g034120.3 |
| Solyc03g034150.1 | Solyc03g034150.2 |
| Solyc03g034180.2 | Solyc03g034180.3 |
| Solyc03g034190.2 | Solyc03g034190.3 |
| Solyc03g034200.2 | Solyc03g034200.3 |
| Solyc03g034220.2 | Solyc03g034220.3 |
| Solyc03g034270.2 | Solyc03g034270.3 |
| Solyc03g034280.2 | Solyc03g034280.3 |
| Solyc03g034290.1 | Solyc03g034290.2 |
| Solyc03g034320.2 | Solyc03g034320.3 |
| Solyc03g034400.2 | Solyc03g034400.3 |
| Solyc03g034413.1 | Solyc03g150106.1 |
| Solyc03g034417.1 | Solyc03g150107.1 |

| | |
|---|---|
| Solyc03g036477.1 | Solyc03g036477.2 |
| Solyc03g043640.2 | Solyc03g043640.3 |
| Solyc03g043660.2 | Solyc03g043660.3 |
| Solyc03g043665.1 | Solyc03g150108.1 |
| Solyc03g043700.2 | Solyc03g043700.3 |
| Solyc03g043750.2 | Solyc03g043750.3 |
| Solyc03g043780.1 | Solyc03g043780.2 |
| Solyc03g043795.1 | Solyc03g150109.1 |
| Solyc03g043850.2 | Solyc03g043850.3 |
| Solyc03g043880.2 | Solyc03g043880.3 |
| Solyc03g043890.2 | Solyc03g043890.3 |
| Solyc03g043910.2 | Solyc03g043910.3 |
| Solyc03g043960.2 | Solyc03g043960.3 |
| Solyc03g044022.1 | Solyc03g150110.1 |
| Solyc03g044026.1 | Solyc03g150111.1 |
| Solyc03g044028.1 | Solyc03g150112.1 |
| Solyc03g044085.1 | Solyc03g150113.1 |
| Solyc03g044140.2 | Solyc03g044140.3 |
| Solyc03g044190.2 | Solyc03g044190.3 |
| Solyc03g044200.2 | Solyc03g044200.3 |
| Solyc03g044260.2 | Solyc03g044260.3 |
| Solyc03g044270.2 | Solyc03g044270.3 |
| Solyc03g044273.1 | Solyc03g150115.1 |
| Solyc03g044277.1 | Solyc03g150116.1 |
| Solyc03g044300.2 | Solyc03g044300.3 |
| Solyc03g044370.2 | Solyc03g044370.3 |
| Solyc03g044380.2 | Solyc03g044380.3 |
| Solyc03g044420.1 | Solyc03g044420.2 |
| Solyc03g044430.2 | Solyc03g044430.3 |
| Solyc03g044440.1 | Solyc03g044440.2 |
| Solyc03g044480.2 | Solyc03g044480.3 |
| Solyc03g044510.2 | Solyc03g044510.3 |
| Solyc03g044520.1 | Solyc03g044520.2 |
| Solyc03g044600.1 | Solyc03g044600.2 |
| Solyc03g044660.2 | Solyc03g044660.3 |
| Solyc03g044720.1 | Solyc03g044720.2 |
| Solyc03g044790.2 | Solyc03g044790.3 |
| Solyc03g044800.1 | Solyc03g044800.2 |
| Solyc03g044810.1 | Solyc03g044810.2 |
| Solyc03g044900.2 | Solyc03g044900.3 |
| Solyc03g044950.1 | Solyc03g044950.2 |
| Solyc03g044960.1 | Solyc03g044960.2 |
| Solyc03g045030.1 | Solyc03g045030.2 |

| | |
|---|---|
| Solyc03g045060.2 | Solyc03g045060.3 |
| Solyc03g045110.2 | Solyc03g045110.3 |
| Solyc03g046205.1 | Solyc03g150117.1 |
| Solyc03g046275.1 | Solyc03g150118.1 |
| Solyc03g046340.2 | Solyc03g046340.3 |
| Solyc03g046350.2 | Solyc03g046350.3 |
| Solyc03g046410.2 | Solyc03g046410.3 |
| Solyc03g046450.2 | Solyc03g046450.3 |
| Solyc03g046470.2 | Solyc03g046470.3 |
| Solyc03g046540.1 | Solyc03g046540.2 |
| Solyc03g046590.2 | Solyc03g046590.3 |
| Solyc03g046594.1 | Solyc03g150119.1 |
| Solyc03g046596.1 | Solyc03g150120.1 |
| Solyc03g046598.1 | Solyc03g150121.1 |
| Solyc03g070380.2 | Solyc03g070380.3 |
| Solyc03g070390.2 | Solyc03g070390.3 |
| Solyc03g071570.2 | Solyc03g071570.3 |
| Solyc03g071590.2 | Solyc03g071590.3 |
| Solyc03g071620.1 | Solyc03g071620.2 |
| Solyc03g071660.1 | Solyc03g071660.2 |
| Solyc03g071680.1 | Solyc03g071680.2 |
| Solyc03g071750.2 | Solyc03g071750.3 |
| Solyc03g071795.1 | Solyc03g150133.1 |
| Solyc03g071800.1 | Solyc03g071800.2 |
| Solyc03g071820.1 | Solyc03g071820.2 |
| Solyc03g071840.1 | Solyc03g071840.2 |
| Solyc03g051625.1 | Solyc03g150122.1 |
| Solyc03g051645.1 | Solyc03g150123.1 |
| Solyc03g051790.2 | Solyc03g051790.3 |
| Solyc03g051970.2 | Solyc03g051970.3 |
| Solyc03g052980.2 | Solyc03g052980.3 |
| Solyc03g052990.2 | Solyc03g052990.3 |
| Solyc03g053110.2 | Solyc03g053110.3 |
| Solyc03g053130.2 | Solyc03g053130.3 |
| Solyc03g058190.2 | Solyc03g058190.3 |
| Solyc03g058213.1 | Solyc03g150124.1 |
| Solyc03g058215.1 | Solyc03g150125.1 |
| Solyc03g058217.1 | Solyc03g150126.1 |
| Solyc03g058330.2 | Solyc03g058330.3 |
| Solyc03g058370.1 | Solyc03g058370.2 |
| Solyc03g058390.1 | Solyc03g058390.2 |
| Solyc03g058550.1 | Solyc03g058550.2 |
| Solyc03g058640.1 | Solyc03g058640.2 |

| | |
|---|---|
| Solyc03g058853.1 | Solyc03g150127.1 |
| Solyc03g058857.1 | Solyc03g150128.1 |
| Solyc03g058860.2 | Solyc03g058860.3 |
| Solyc03g058910.2 | Solyc03g058910.3 |
| Solyc03g058920.2 | Solyc03g058920.3 |
| Solyc03g058950.2 | Solyc03g058950.3 |
| Solyc03g059127.1 | Solyc03g150129.1 |
| Solyc03g059200.1 | Solyc03g059200.2 |
| Solyc03g059250.2 | Solyc03g059250.3 |
| Solyc03g059310.2 | Solyc03g059310.3 |
| Solyc03g059350.2 | Solyc03g059350.3 |
| Solyc03g059390.2 | Solyc03g059390.3 |
| Solyc03g059500.2 | Solyc03g059500.3 |
| Solyc03g061655.1 | Solyc03g150130.1 |
| Solyc03g062650.2 | Solyc03g062650.3 |
| Solyc03g062680.2 | Solyc03g062680.3 |
| Solyc03g062720.2 | Solyc03g062720.3 |
| Solyc03g062790.2 | Solyc03g062790.3 |
| Solyc03g062830.2 | Solyc03g062830.3 |
| Solyc03g062890.2 | Solyc03g062890.3 |
| Solyc03g062940.2 | Solyc03g062940.3 |
| Solyc03g063010.1 | Solyc03g063010.2 |
| Solyc03g063060.2 | Solyc03g063060.3 |
| Solyc03g063100.1 | Solyc03g063100.2 |
| Solyc03g063110.2 | Solyc03g063110.3 |
| Solyc03g063150.2 | Solyc03g063150.3 |
| Solyc03g063210.2 | Solyc03g063210.3 |
| Solyc03g063220.1 | Solyc03g063220.2 |
| Solyc03g063240.2 | Solyc03g063240.3 |
| Solyc03g063280.2 | Solyc03g063280.3 |
| Solyc03g063290.2 | Solyc03g063290.3 |
| Solyc03g063320.2 | Solyc03g063320.3 |
| Solyc03g063340.2 | Solyc03g063340.3 |
| Solyc03g063370.2 | Solyc03g063370.3 |
| Solyc03g063420.1 | Solyc03g063420.2 |
| Solyc03g063450.1 | Solyc03g063450.2 |
| Solyc03g063560.2 | Solyc03g063560.3 |
| Solyc03g063563.1 | Solyc03g150131.1 |
| Solyc03g063567.1 | Solyc03g150132.1 |
| Solyc03g063600.2 | Solyc03g063600.4 |
| Solyc03g063750.2 | Solyc03g063750.3 |
| Solyc03g063880.2 | Solyc03g063880.3 |
| Solyc03g063970.2 | Solyc03g063970.3 |

| | |
|---|---|
| Solyc03g064020.2 | Solyc03g064020.3 |
| Solyc03g064030.2 | Solyc03g064030.3 |
| Solyc03g065110.1 | Solyc03g065110.2 |
| Solyc03g065340.2 | Solyc03g065340.3 |
| Solyc03g104810.2 | Solyc03g104810.3 |
| Solyc03g013340.2 | Solyc03g013340.3 |
| Solyc03g013310.2 | Solyc03g013310.3 |
| Solyc03g013304.1 | Solyc03g013304.2 |
| Solyc03g013110.1 | Solyc03g013110.2 |
| Solyc03g013060.1 | Solyc03g013060.2 |
| Solyc03g013030.2 | Solyc03g013030.3 |
| Solyc03g077890.1 | Solyc03g077890.2 |
| Solyc03g077930.2 | Solyc03g077930.3 |
| Solyc03g078000.2 | Solyc03g078000.3 |
| Solyc03g078050.1 | Solyc03g078050.2 |
| Solyc03g078100.2 | Solyc03g078100.3 |
| Solyc03g078150.2 | Solyc03g078150.3 |
| Solyc03g078230.2 | Solyc03g078230.3 |
| Solyc03g078260.2 | Solyc03g078260.3 |
| Solyc03g078270.1 | Solyc03g078270.2 |
| Solyc03g078290.2 | Solyc03g078290.3 |
| Solyc03g078350.2 | Solyc03g078350.3 |
| Solyc03g078390.2 | Solyc03g078390.3 |
| Solyc03g078400.2 | Solyc03g078400.3 |
| Solyc03g078405.1 | Solyc03g150134.1 |
| Solyc03g078440.2 | Solyc03g078440.3 |
| Solyc03g078470.2 | Solyc03g078470.3 |
| Solyc03g078500.2 | Solyc03g078500.3 |
| Solyc03g078535.1 | Solyc03g150135.1 |
| Solyc03g078570.2 | Solyc03g078570.3 |
| Solyc03g078580.2 | Solyc03g078580.3 |
| Solyc03g078610.2 | Solyc03g078610.3 |
| Solyc03g078630.2 | Solyc03g078630.3 |
| Solyc03g078770.2 | Solyc03g078770.3 |
| Solyc03g078820.1 | Solyc03g078820.2 |
| Solyc03g079860.1 | Solyc03g079860.2 |
| Solyc03g079865.1 | Solyc03g150136.1 |
| Solyc03g079900.2 | Solyc03g079900.3 |
| Solyc03g079910.2 | Solyc03g079910.3 |
| Solyc03g079920.2 | Solyc03g079920.3 |
| Solyc03g079930.2 | Solyc03g079930.3 |
| Solyc03g079940.2 | Solyc03g079940.3 |
| Solyc03g079980.1 | Solyc03g079980.2 |

| | |
|---|---|
| Solyc03g080010.2 | Solyc03g080010.3 |
| Solyc03g080050.2 | Solyc03g080050.3 |
| Solyc03g080110.2 | Solyc03g080110.3 |
| Solyc03g080140.2 | Solyc03g080140.3 |
| Solyc03g080150.2 | Solyc03g080150.3 |
| Solyc03g080160.2 | Solyc03g080160.3 |
| Solyc03g080190.2 | Solyc03g080190.3 |
| Solyc03g081190.1 | Solyc03g081190.2 |
| Solyc03g081240.2 | Solyc03g081240.3 |
| Solyc03g081270.1 | Solyc03g081270.2 |
| Solyc03g081320.2 | Solyc03g081320.3 |
| Solyc03g082390.2 | Solyc03g082390.3 |
| Solyc03g082400.2 | Solyc03g082400.3 |
| Solyc03g082410.2 | Solyc03g082410.3 |
| Solyc03g082420.2 | Solyc03g082420.3 |
| Solyc03g082450.2 | Solyc03g082450.3 |
| Solyc03g082550.2 | Solyc03g082550.3 |
| Solyc03g082560.2 | Solyc03g082560.3 |
| Solyc03g082575.1 | Solyc03g150137.1 |
| Solyc03g082580.2 | Solyc03g082580.3 |
| Solyc03g082630.2 | Solyc03g082630.3 |
| Solyc03g082660.2 | Solyc03g082660.3 |
| Solyc03g082670.2 | Solyc03g082670.3 |
| Solyc03g082690.2 | Solyc03g082690.3 |
| Solyc03g082720.2 | Solyc03g082720.3 |
| Solyc03g082740.2 | Solyc03g082740.3 |
| Solyc03g082800.1 | Solyc03g082800.2 |
| Solyc03g082820.2 | Solyc03g082820.3 |
| Solyc03g082830.1 | Solyc03g082830.2 |
| Solyc03g082840.2 | Solyc03g082840.3 |
| Solyc03g082850.2 | Solyc03g082850.3 |
| Solyc03g082860.2 | Solyc03g082860.3 |
| Solyc03g082890.2 | Solyc03g082890.3 |
| Solyc03g082900.2 | Solyc03g082900.3 |
| Solyc03g082940.2 | Solyc03g082940.3 |
| Solyc03g082980.2 | Solyc03g082980.3 |
| Solyc03g082990.2 | Solyc03g082990.3 |
| Solyc03g083000.2 | Solyc03g083000.3 |
| Solyc03g083010.2 | Solyc03g083010.3 |
| Solyc03g083020.2 | Solyc03g083020.3 |
| Solyc03g083100.2 | Solyc03g083100.3 |
| Solyc03g083140.2 | Solyc03g083140.3 |
| Solyc03g083170.2 | Solyc03g083170.3 |

| | |
|---|---|
| Solyc03g083210.2 | Solyc03g083210.3 |
| Solyc03g083250.2 | Solyc03g083250.3 |
| Solyc03g083300.2 | Solyc03g083300.3 |
| Solyc03g083330.2 | Solyc03g083330.3 |
| Solyc03g083350.2 | Solyc03g083350.3 |
| Solyc03g083360.2 | Solyc03g083360.3 |
| Solyc03g083370.2 | Solyc03g083370.3 |
| Solyc03g083400.2 | Solyc03g083400.3 |
| Solyc03g083410.2 | Solyc03g083410.3 |
| Solyc03g083430.2 | Solyc03g083430.3 |
| Solyc03g083460.2 | Solyc03g083460.3 |
| Solyc03g083470.2 | Solyc03g083470.3 |
| Solyc03g083490.2 | Solyc03g083490.3 |
| Solyc03g083500.2 | Solyc03g083500.3 |
| Solyc03g083530.2 | Solyc03g083530.3 |
| Solyc03g083540.2 | Solyc03g083540.3 |
| Solyc03g083570.2 | Solyc03g083570.3 |
| Solyc03g083580.2 | Solyc03g083580.3 |
| Solyc03g083590.2 | Solyc03g083590.3 |
| Solyc03g083785.1 | Solyc03g150138.1 |
| Solyc03g083840.2 | Solyc03g083840.3 |
| Solyc03g083843.1 | Solyc03g150139.1 |
| Solyc03g083847.1 | Solyc03g150140.1 |
| Solyc03g083890.1 | Solyc03g083890.2 |
| Solyc03g083960.2 | Solyc03g083960.3 |
| Solyc03g083970.2 | Solyc03g083970.3 |
| Solyc03g093040.2 | Solyc03g093040.3 |
| Solyc03g093080.2 | Solyc03g093080.3 |
| Solyc03g093120.4 | Solyc03g093120.5.1 |
| Solyc03g093130.2 | Solyc03g093130.3 |
| Solyc03g093140.2 | Solyc03g093140.3 |
| Solyc03g093160.2 | Solyc03g093160.3 |
| Solyc03g093200.2 | Solyc03g093200.3 |
| Solyc03g093210.2 | Solyc03g093210.3 |
| Solyc03g093250.1 | Solyc03g093250.2 |
| Solyc03g093260.2 | Solyc03g093260.3 |
| Solyc03g093270.2 | Solyc03g093270.3 |
| Solyc03g093300.2 | Solyc03g093300.3 |
| Solyc03g093310.2 | Solyc03g093310.3 |
| Solyc03g093320.1 | Solyc03g093320.2 |
| Solyc03g093335.1 | Solyc03g150141.1 |
| Solyc03g093360.2 | Solyc03g093360.3 |
| Solyc03g093370.1 | Solyc03g093370.2 |

| | |
|---|---|
| Solyc03g093380.1 | Solyc03g093380.2 |
| Solyc03g093410.2 | Solyc03g093410.3 |
| Solyc03g093430.2 | Solyc03g093430.3 |
| Solyc03g093450.2 | Solyc03g093450.3 |
| Solyc03g093460.2 | Solyc03g093460.3 |
| Solyc03g093470.2 | Solyc03g093470.3 |
| Solyc03g093490.2 | Solyc03g093490.3 |
| Solyc03g093500.2 | Solyc03g093500.3 |
| Solyc03g093520.2 | Solyc03g093520.3 |
| Solyc03g093625.1 | Solyc03g150142.1 |
| Solyc03g093650.2 | Solyc03g093650.3 |
| Solyc03g093690.2 | Solyc03g093690.3 |
| Solyc03g093790.2 | Solyc03g093790.3 |
| Solyc03g093830.2 | Solyc03g093830.3 |
| Solyc03g093850.1 | Solyc03g093850.2 |
| Solyc03g093880.2 | Solyc03g093880.3 |
| Solyc03g093890.2 | Solyc03g093890.3 |
| Solyc03g093940.1 | Solyc03g093940.2 |
| Solyc03g093970.2 | Solyc03g093970.3 |
| Solyc03g094000.2 | Solyc03g094000.3 |
| Solyc03g094010.2 | Solyc03g094010.3 |
| Solyc03g094020.2 | Solyc03g094020.3 |
| Solyc03g094090.2 | Solyc03g094090.3 |
| Solyc03g094120.2 | Solyc03g094120.3 |
| Solyc03g094160.2 | Solyc03g094160.3 |
| Solyc03g095180.2 | Solyc03g095180.3 |
| Solyc03g095210.1 | Solyc03g095210.2 |
| Solyc03g095220.2 | Solyc03g095220.3 |
| Solyc03g095250.2 | Solyc03g095250.3 |
| Solyc03g095260.2 | Solyc03g095260.3 |
| Solyc03g095290.2 | Solyc03g095290.3 |
| Solyc03g095310.2 | Solyc03g095310.3 |
| Solyc03g095330.2 | Solyc03g095330.3 |
| Solyc03g095360.2 | Solyc03g095360.3 |
| Solyc03g095370.2 | Solyc03g095370.3 |
| Solyc03g095383.1 | Solyc03g150143.1 |
| Solyc03g095387.1 | Solyc03g150144.1 |
| Solyc03g095410.2 | Solyc03g095410.3 |
| Solyc03g095450.2 | Solyc03g095450.3 |
| Solyc03g095470.2 | Solyc03g095470.3 |
| Solyc03g095490.2 | Solyc03g095490.3 |
| Solyc03g095500.2 | Solyc03g095500.3 |
| Solyc03g095510.2 | Solyc03g095510.3 |

| | |
|---|---|
| Solyc03g095550.1 | Solyc03g095550.2 |
| Solyc03g095650.2 | Solyc03g095650.3 |
| Solyc03g095680.1 | Solyc03g095680.2 |
| Solyc03g095690.2 | Solyc03g095690.3 |
| Solyc03g095720.2 | Solyc03g095720.3 |
| Solyc03g095750.1 | Solyc03g095750.2 |
| Solyc03g095760.2 | Solyc03g095760.3 |
| Solyc03g095770.2 | Solyc03g095770.3 |
| Solyc03g095783.1 | Solyc03g150145.1 |
| Solyc03g095810.2 | Solyc03g095810.3 |
| Solyc03g095840.2 | Solyc03g095840.3 |
| Solyc03g095860.2 | Solyc03g095860.3 |
| Solyc03g095950.2 | Solyc03g095950.3 |
| Solyc03g095970.2 | Solyc03g095970.3 |
| Solyc03g095980.2 | Solyc03g095980.3 |
| Solyc03g096000.2 | Solyc03g096000.3 |
| Solyc03g096003.1 | Solyc03g150146.1 |
| Solyc03g096030.2 | Solyc03g096030.3 |
| Solyc03g096040.2 | Solyc03g096040.3 |
| Solyc03g096050.2 | Solyc03g096050.3 |
| Solyc03g096055.1 | Solyc03g150147.1 |
| Solyc03g096070.2 | Solyc03g096070.3 |
| Solyc03g096100.1 | Solyc03g096100.2 |
| Solyc03g096130.1 | Solyc03g096130.2 |
| Solyc03g096140.1 | Solyc03g096140.2 |
| Solyc03g096190.1 | Solyc03g096190.2 |
| Solyc03g096220.2 | Solyc03g096220.3 |
| Solyc03g096250.2 | Solyc03g096250.3 |
| Solyc03g096290.2 | Solyc03g096290.3 |
| Solyc03g096300.2 | Solyc03g096300.3 |
| Solyc03g096350.2 | Solyc03g096350.3 |
| Solyc03g096370.2 | Solyc03g096370.3 |
| Solyc03g096380.2 | Solyc03g096380.3 |
| Solyc03g096390.2 | Solyc03g096390.3 |
| Solyc03g096540.2 | Solyc03g096540.3 |
| Solyc03g096610.1 | Solyc03g096610.2 |
| Solyc03g096650.1 | Solyc03g096650.2 |
| Solyc03g096660.1 | Solyc03g096660.2 |
| Solyc03g096670.2 | Solyc03g096670.3 |
| Solyc03g096680.1 | Solyc03g096680.2 |
| Solyc03g096720.1 | Solyc03g096720.2 |
| Solyc03g096730.2 | Solyc03g096730.3 |
| Solyc03g096790.2 | Solyc03g096790.3 |

| | |
|---|---|
| Solyc03g096810.2 | Solyc03g096810.3 |
| Solyc03g096860.2 | Solyc03g096860.3 |
| Solyc03g096870.2 | Solyc03g096870.3 |
| Solyc03g096910.2 | Solyc03g096910.3 |
| Solyc03g096920.2 | Solyc03g096920.3 |
| Solyc03g097000.2 | Solyc03g097000.3 |
| Solyc03g097010.2 | Solyc03g097010.3 |
| Solyc03g097030.2 | Solyc03g097030.3 |
| Solyc03g097050.2 | Solyc03g097050.3 |
| Solyc03g097070.2 | Solyc03g097070.3 |
| Solyc03g097120.2 | Solyc03g097120.3 |
| Solyc03g097130.2 | Solyc03g097130.3 |
| Solyc03g097140.2 | Solyc03g097140.3 |
| Solyc03g097150.2 | Solyc03g097150.3 |
| Solyc03g097170.2 | Solyc03g097170.3 |
| Solyc03g097190.2 | Solyc03g097190.3 |
| Solyc03g097210.2 | Solyc03g097210.3 |
| Solyc03g097240.2 | Solyc03g097240.3 |
| Solyc03g097270.2 | Solyc03g097270.3 |
| Solyc03g097310.2 | Solyc03g097310.3 |
| Solyc03g097320.2 | Solyc03g097320.3 |
| Solyc03g097360.2 | Solyc03g097360.3 |
| Solyc03g097390.2 | Solyc03g097390.3 |
| Solyc03g097420.2 | Solyc03g097420.3 |
| Solyc03g097440.2 | Solyc03g097440.3 |
| Solyc03g097460.2 | Solyc03g097460.3 |
| Solyc03g097470.2 | Solyc03g097470.3 |
| Solyc03g097500.2 | Solyc03g097500.3 |
| Solyc03g097520.2 | Solyc03g097520.3 |
| Solyc03g097530.2 | Solyc03g097530.3 |
| Solyc03g097540.1 | Solyc03g097540.2 |
| Solyc03g097550.2 | Solyc03g097550.3 |
| Solyc03g097560.2 | Solyc03g097560.3 |
| Solyc03g097570.2 | Solyc03g097570.3 |
| Solyc03g097600.2 | Solyc03g097600.3 |
| Solyc03g097610.2 | Solyc03g097610.3 |
| Solyc03g097630.2 | Solyc03g097630.3 |
| Solyc03g097640.2 | Solyc03g097640.3 |
| Solyc03g097650.2 | Solyc03g097650.3 |
| Solyc03g097660.2 | Solyc03g097660.3 |
| Solyc03g097690.2 | Solyc03g097690.3 |
| Solyc03g097700.2 | Solyc03g097700.3 |
| Solyc03g097730.1 | Solyc03g097730.2 |

| | |
|---|---|
| Solyc03g097780.2 | Solyc03g097780.3 |
| Solyc03g097820.1 | Solyc03g097820.2 |
| Solyc03g097830.2 | Solyc03g097830.3 |
| Solyc03g097840.2 | Solyc03g097840.3 |
| Solyc03g097900.2 | Solyc03g097900.3 |
| Solyc03g097930.2 | Solyc03g097930.3 |
| Solyc03g097940.2 | Solyc03g097940.3 |
| Solyc03g097970.2 | Solyc03g097970.3 |
| Solyc03g097990.2 | Solyc03g097990.3 |
| Solyc03g098000.2 | Solyc03g098000.3 |
| Solyc03g098010.2 | Solyc03g098010.3 |
| Solyc03g098030.2 | Solyc03g098030.3 |
| Solyc03g098050.2 | Solyc03g098050.3 |
| Solyc03g098070.2 | Solyc03g098070.3 |
| Solyc03g098090.1 | Solyc03g098090.2 |
| Solyc03g098140.2 | Solyc03g098140.3 |
| Solyc03g098150.2 | Solyc03g098150.3 |
| Solyc03g098160.2 | Solyc03g098160.3 |
| Solyc03g098170.2 | Solyc03g098170.3 |
| Solyc03g098180.2 | Solyc03g098180.3 |
| Solyc03g098190.2 | Solyc03g098190.3 |
| Solyc03g098230.2 | Solyc03g098230.3 |
| Solyc03g098240.2 | Solyc03g098240.3 |
| Solyc03g098360.1 | Solyc03g098360.2 |
| Solyc03g098400.1 | Solyc03g098400.2 |
| Solyc03g098420.2 | Solyc03g098420.3 |
| Solyc03g098450.2 | Solyc03g098450.3 |
| Solyc03g098460.2 | Solyc03g098460.3 |
| Solyc03g098470.2 | Solyc03g098470.3 |
| Solyc03g098540.1 | Solyc03g098540.2 |
| Solyc03g098560.1 | Solyc03g098560.2 |
| Solyc03g098590.2 | Solyc03g098590.3 |
| Solyc03g098600.2 | Solyc03g098600.3 |
| Solyc03g098610.2 | Solyc03g098610.3 |
| Solyc03g098620.2 | Solyc03g098620.3 |
| Solyc03g098650.2 | Solyc03g098650.3 |
| Solyc03g098660.2 | Solyc03g098660.3 |
| Solyc03g098680.2 | Solyc03g098680.3 |
| Solyc03g098720.2 | Solyc03g098720.3 |
| Solyc03g098750.2 | Solyc03g098750.3 |
| Solyc03g020060.2 | Solyc03g020060.3 |
| Solyc03g020050.2 | Solyc03g020050.3 |
| Solyc03g020040.2 | Solyc03g020040.3 |

| | |
|---|---|
| Solyc03g020030.2 | Solyc03g020030.3 |
| Solyc03g020015.1 | Solyc03g150101.1 |
| Solyc03g019980.1 | Solyc03g019980.2 |
| Solyc03g019960.2 | Solyc03g019960.3 |
| Solyc03g019950.2 | Solyc03g019950.3 |
| Solyc03g019900.2 | Solyc03g019900.3 |
| Solyc03g019890.2 | Solyc03g019890.3 |
| Solyc03g019880.2 | Solyc03g019880.3 |
| Solyc03g019860.2 | Solyc03g019860.3 |
| Solyc03g019850.2 | Solyc03g019850.3 |
| Solyc03g019820.2 | Solyc03g019820.3 |
| Solyc03g019810.2 | Solyc03g019810.3 |
| Solyc03g019790.2 | Solyc03g019790.3 |
| Solyc03g019750.2 | Solyc03g019750.3 |
| Solyc03g019680.2 | Solyc03g019680.3 |
| Solyc03g019670.2 | Solyc03g019670.3 |
| Solyc03g019650.2 | Solyc03g019650.3 |
| Solyc03g110860.2 | Solyc03g110860.3 |
| Solyc03g110870.2 | Solyc03g110870.3 |
| Solyc03g110930.2 | Solyc03g110930.3 |
| Solyc03g110940.2 | Solyc03g110940.3 |
| Solyc03g110960.2 | Solyc03g110960.3 |
| Solyc03g111000.2 | Solyc03g111000.3 |
| Solyc03g111020.2 | Solyc03g111020.3 |
| Solyc03g111040.1 | Solyc03g111040.2 |
| Solyc03g111050.2 | Solyc03g111050.3 |
| Solyc03g111060.2 | Solyc03g111060.3 |
| Solyc03g111070.2 | Solyc03g111070.3 |
| Solyc03g111080.2 | Solyc03g111080.3 |
| Solyc03g111140.2 | Solyc03g111140.3 |
| Solyc03g111160.2 | Solyc03g111160.3 |
| Solyc03g111170.2 | Solyc03g111170.3 |
| Solyc03g111180.2 | Solyc03g111180.3 |
| Solyc03g111200.2 | Solyc03g111200.3 |
| Solyc03g111240.1 | Solyc03g111240.2 |
| Solyc03g111250.2 | Solyc03g111250.3 |
| Solyc03g111260.2 | Solyc03g111260.3 |
| Solyc03g111275.1 | Solyc03g150148.1 |
| Solyc03g111290.1 | Solyc03g111290.2 |
| Solyc03g111310.2 | Solyc03g111310.3 |
| Solyc03g111350.2 | Solyc03g111350.3 |
| Solyc03g111360.2 | Solyc03g111360.3 |
| Solyc03g111370.2 | Solyc03g111370.3 |

| | |
|---|---|
| Solyc03g111380.2 | Solyc03g111380.3 |
| Solyc03g111410.2 | Solyc03g111410.3 |
| Solyc03g111500.2 | Solyc03g111500.3 |
| Solyc03g111510.2 | Solyc03g111510.3 |
| Solyc03g111540.1 | Solyc03g111540.2 |
| Solyc03g111580.2 | Solyc03g111580.3 |
| Solyc03g111610.2 | Solyc03g111610.3 |
| Solyc03g111670.2 | Solyc03g111670.3 |
| Solyc03g111680.2 | Solyc03g111680.3 |
| Solyc03g111700.2 | Solyc03g111700.3 |
| Solyc03g111710.2 | Solyc03g111710.3 |
| Solyc03g111720.2 | Solyc03g111720.3 |
| Solyc03g111730.2 | Solyc03g111730.3 |
| Solyc03g111740.2 | Solyc03g111740.3 |
| Solyc03g111760.2 | Solyc03g111760.3 |
| Solyc03g111770.2 | Solyc03g111770.3 |
| Solyc03g111840.2 | Solyc03g111840.3 |
| Solyc03g111850.2 | Solyc03g111850.3 |
| Solyc03g111870.2 | Solyc03g111870.3 |
| Solyc03g111880.2 | Solyc03g111880.3 |
| Solyc03g111883.1 | Solyc03g150149.1 |
| Solyc03g111887.1 | Solyc03g150150.1 |
| Solyc03g111920.1 | Solyc03g111920.2 |
| Solyc03g111950.2 | Solyc03g111950.3 |
| Solyc03g111975.1 | Solyc03g150151.1 |
| Solyc03g112060.2 | Solyc03g112060.3 |
| Solyc03g112075.1 | Solyc03g150152.1 |
| Solyc03g112090.2 | Solyc03g112090.3 |
| Solyc03g112100.2 | Solyc03g112100.3 |
| Solyc03g112160.2 | Solyc03g112160.3 |
| Solyc03g112270.2 | Solyc03g112270.3 |
| Solyc03g112280.2 | Solyc03g112280.3 |
| Solyc03g112300.2 | Solyc03g112300.3 |
| Solyc03g112330.2 | Solyc03g112330.3 |
| Solyc03g112390.2 | Solyc03g112390.3 |
| Solyc03g112410.1 | Solyc03g112410.2 |
| Solyc03g112460.2 | Solyc03g112460.3 |
| Solyc03g112510.1 | Solyc03g112510.2 |
| Solyc03g112520.2 | Solyc03g112520.3 |
| Solyc03g112530.2 | Solyc03g112530.3 |
| Solyc03g112550.2 | Solyc03g112550.3 |
| Solyc03g112580.2 | Solyc03g112580.3 |
| Solyc03g112630.2 | Solyc03g112630.3 |

| | |
|---|---|
| Solyc03g112640.2 | Solyc03g112640.3 |
| Solyc03g112650.2 | Solyc03g112650.3 |
| Solyc03g112660.2 | Solyc03g112660.3 |
| Solyc03g112740.1 | Solyc03g112740.2 |
| Solyc03g112755.1 | Solyc03g150153.1 |
| Solyc03g112770.2 | Solyc03g112770.3 |
| Solyc03g112790.2 | Solyc03g112790.3 |
| Solyc03g112800.1 | Solyc03g112800.2 |
| Solyc03g112810.2 | Solyc03g112810.3 |
| Solyc03g112850.2 | Solyc03g112850.3 |
| Solyc03g112860.2 | Solyc03g112860.3 |
| Solyc03g112870.2 | Solyc03g112870.3 |
| Solyc03g112900.2 | Solyc03g112900.3 |
| Solyc03g112910.2 | Solyc03g112910.3 |
| Solyc03g112930.2 | Solyc03g112930.3 |
| Solyc03g113000.2 | Solyc03g113000.3 |
| Solyc03g113020.2 | Solyc03g113020.3 |
| Solyc03g113060.2 | Solyc03g113060.3 |
| Solyc03g113100.2 | Solyc03g113100.3 |
| Solyc03g113130.2 | Solyc03g113130.3 |
| Solyc03g113140.2 | Solyc03g113140.3 |
| Solyc03g113150.2 | Solyc03g113150.3 |
| Solyc03g113180.2 | Solyc03g113180.3 |
| Solyc03g113210.2 | Solyc03g113210.3 |
| Solyc03g113220.2 | Solyc03g113220.3 |
| Solyc03g113240.2 | Solyc03g113240.3 |
| Solyc03g113270.2 | Solyc03g113270.3 |
| Solyc03g113280.2 | Solyc03g113280.3 |
| Solyc03g113330.2 | Solyc03g113330.3 |
| Solyc03g113340.2 | Solyc03g113340.3 |
| Solyc03g113350.2 | Solyc03g113350.3 |
| Solyc03g113370.2 | Solyc03g113370.3 |
| Solyc03g113390.2 | Solyc03g113390.3 |
| Solyc03g113410.1 | Solyc03g113410.2 |
| Solyc03g113430.2 | Solyc03g113430.3 |
| Solyc03g113440.2 | Solyc03g113440.3 |
| Solyc03g113490.2 | Solyc03g113490.3 |
| Solyc03g113520.2 | Solyc03g113520.3 |
| Solyc03g113530.2 | Solyc03g113530.3 |
| Solyc03g113540.2 | Solyc03g113540.3 |
| Solyc03g113590.2 | Solyc03g113590.3 |
| Solyc03g113620.2 | Solyc03g113620.3 |
| Solyc03g113630.2 | Solyc03g113630.3 |

| | |
|---|---|
| Solyc03g113700.2 | Solyc03g113700.3 |
| Solyc03g113720.2 | Solyc03g113720.3 |
| Solyc03g113730.2 | Solyc03g113730.3 |
| Solyc03g113760.2 | Solyc03g113760.3 |
| Solyc03g113770.2 | Solyc03g113770.3 |
| Solyc03g113780.2 | Solyc03g113780.3 |
| Solyc03g113800.2 | Solyc03g113800.3 |
| Solyc03g113845.1 | Solyc03g150154.1 |
| Solyc03g113860.1 | Solyc03g113860.2 |
| Solyc03g113910.2 | Solyc03g113910.3 |
| Solyc03g113960.3 | Solyc03g113960.4 |
| Solyc03g113980.2 | Solyc03g113980.3 |
| Solyc03g113990.2 | Solyc03g113990.3 |
| Solyc03g114000.2 | Solyc03g114000.3 |
| Solyc03g114020.2 | Solyc03g114020.3 |
| Solyc03g114030.2 | Solyc03g114030.3 |
| Solyc03g114040.2 | Solyc03g114040.3 |
| Solyc03g114050.2 | Solyc03g114050.3 |
| Solyc03g114060.2 | Solyc03g114060.3 |
| Solyc03g114070.2 | Solyc03g114070.3 |
| Solyc03g114080.1 | Solyc03g114080.2 |
| Solyc03g114120.2 | Solyc03g114120.3 |
| Solyc03g114140.2 | Solyc03g114140.3 |
| Solyc03g114150.2 | Solyc03g114150.3 |
| Solyc03g114170.2 | Solyc03g114170.3 |
| Solyc03g114210.2 | Solyc03g114210.3 |
| Solyc03g114230.1 | Solyc03g114230.2 |
| Solyc03g114280.2 | Solyc03g114280.3 |
| Solyc03g114310.2 | Solyc03g114310.3 |
| Solyc03g114330.2 | Solyc03g114330.3 |
| Solyc03g114340.2 | Solyc03g114340.3 |
| Solyc03g114400.2 | Solyc03g114400.3 |
| Solyc03g114410.2 | Solyc03g114410.3 |
| Solyc03g114450.2 | Solyc03g114450.3 |
| Solyc03g114460.2 | Solyc03g114460.3 |
| Solyc03g114520.2 | Solyc03g114520.3 |
| Solyc03g114540.1 | Solyc03g114540.2 |
| Solyc03g114550.1 | Solyc03g114550.2 |
| Solyc03g114590.2 | Solyc03g114590.3 |
| Solyc03g114680.2 | Solyc03g114680.3 |
| Solyc03g114700.1 | Solyc03g114700.2 |
| Solyc03g114720.2 | Solyc03g114720.3 |
| Solyc03g114730.2 | Solyc03g114730.3 |

| | |
|---|---|
| Solyc03g114750.2 | Solyc03g114750.3 |
| Solyc03g114760.2 | Solyc03g114760.3 |
| Solyc03g114770.2 | Solyc03g114770.3 |
| Solyc03g114780.2 | Solyc03g114780.3 |
| Solyc03g114790.2 | Solyc03g114790.3 |
| Solyc03g114810.2 | Solyc03g114810.3 |
| Solyc03g114820.2 | Solyc03g114820.3 |
| Solyc03g114830.2 | Solyc03g114830.3 |
| Solyc03g114840.2 | Solyc03g114840.3 |
| Solyc03g114900.2 | Solyc03g114900.3 |
| Solyc03g114930.2 | Solyc03g114930.3 |
| Solyc03g114940.2 | Solyc03g114940.3 |
| Solyc03g114970.2 | Solyc03g114970.3 |
| Solyc03g115000.2 | Solyc03g115000.3 |
| Solyc03g115020.2 | Solyc03g115020.3 |
| Solyc03g115025.1 | Solyc03g150155.1 |
| Solyc03g115050.2 | Solyc03g115050.3 |
| Solyc03g115075.1 | Solyc03g150156.1 |
| Solyc03g115160.2 | Solyc03g115160.3 |
| Solyc03g115200.2 | Solyc03g115200.3 |
| Solyc03g115230.2 | Solyc03g115230.3 |
| Solyc03g115290.2 | Solyc03g115290.3 |
| Solyc03g115300.1 | Solyc03g115300.2 |
| Solyc03g115330.1 | Solyc03g115330.2 |
| Solyc03g115360.2 | Solyc03g115360.3 |
| Solyc03g115370.2 | Solyc03g115370.3 |
| Solyc03g115400.1 | Solyc03g115400.2 |
| Solyc03g115410.2 | Solyc03g115410.3 |
| Solyc03g115470.2 | Solyc03g115470.3 |
| Solyc03g115500.2 | Solyc03g115500.3 |
| Solyc03g115540.1 | Solyc03g115540.2 |
| Solyc03g115550.2 | Solyc03g115550.3 |
| Solyc03g115560.2 | Solyc03g115560.3 |
| Solyc03g115610.2 | Solyc03g115610.3 |
| Solyc03g115630.2 | Solyc03g115630.3 |
| Solyc03g115640.2 | Solyc03g115640.3 |
| Solyc03g115660.2 | Solyc03g115660.3 |
| Solyc03g115700.2 | Solyc03g115700.3 |
| Solyc03g115720.2 | Solyc03g115720.3 |
| Solyc03g115730.2 | Solyc03g115730.3 |
| Solyc03g115740.1 | Solyc03g115740.2 |
| Solyc03g115760.2 | Solyc03g115760.3 |
| Solyc03g115770.2 | Solyc03g115770.3 |

| | |
|---|---|
| Solyc03g115780.1 | Solyc03g115780.2 |
| Solyc03g115810.2 | Solyc03g115810.3 |
| Solyc03g115820.2 | Solyc03g115820.3 |
| Solyc03g115830.1 | Solyc03g115830.2 |
| Solyc03g115850.2 | Solyc03g115850.3 |
| Solyc03g115860.2 | Solyc03g115860.3 |
| Solyc03g115870.2 | Solyc03g115870.3 |
| Solyc03g115890.2 | Solyc03g115890.3 |
| Solyc03g115920.2 | Solyc03g115920.3 |
| Solyc03g115950.2 | Solyc03g115950.3 |
| Solyc03g115960.2 | Solyc03g115960.3 |
| Solyc03g115970.1 | Solyc03g115970.2 |
| Solyc03g116020.2 | Solyc03g116020.3 |
| Solyc03g116100.2 | Solyc03g116100.3 |
| Solyc03g116110.2 | Solyc03g116110.3 |
| Solyc03g116130.1 | Solyc03g116130.2 |
| Solyc03g116140.2 | Solyc03g116140.3 |
| Solyc03g116170.2 | Solyc03g116170.3 |
| Solyc03g116190.1 | Solyc03g116190.2 |
| Solyc03g116200.1 | Solyc03g116200.2 |
| Solyc03g116240.2 | Solyc03g116240.3 |
| Solyc03g116243.1 | Solyc03g150157.1 |
| Solyc03g116247.1 | Solyc03g150158.1 |
| Solyc03g116250.1 | Solyc03g116250.2 |
| Solyc03g116260.2 | Solyc03g116260.3 |
| Solyc03g116270.2 | Solyc03g116270.3 |
| Solyc03g116280.2 | Solyc03g116280.3 |
| Solyc03g116320.2 | Solyc03g116320.3 |
| Solyc03g116340.2 | Solyc03g116340.3 |
| Solyc03g116350.2 | Solyc03g116350.3 |
| Solyc03g116360.1 | Solyc03g116360.2 |
| Solyc03g116370.2 | Solyc03g116370.3 |
| Solyc03g116430.1 | Solyc03g116430.2 |
| Solyc03g116530.2 | Solyc03g116530.3 |
| Solyc03g116570.2 | Solyc03g116570.3 |
| Solyc03g116590.2 | Solyc03g116590.3 |
| Solyc03g116600.2 | Solyc03g116600.3 |
| Solyc03g116610.2 | Solyc03g116610.3 |
| Solyc03g116620.2 | Solyc03g116620.3 |
| Solyc03g116670.2 | Solyc03g116670.3 |
| Solyc03g116680.2 | Solyc03g116680.3 |
| Solyc03g116690.2 | Solyc03g116690.3 |
| Solyc03g116710.2 | Solyc03g116710.3 |

| | |
|---|---|
| Solyc03g116740.2 | Solyc03g116740.3 |
| Solyc03g116750.2 | Solyc03g116750.3 |
| Solyc03g116760.2 | Solyc03g116760.3 |
| Solyc03g116780.2 | Solyc03g116780.3 |
| Solyc03g116850.2 | Solyc03g116850.3 |
| Solyc03g116860.2 | Solyc03g116860.3 |
| Solyc03g116870.2 | Solyc03g116870.3 |
| Solyc03g116880.2 | Solyc03g116880.3 |
| Solyc03g116890.2 | Solyc03g116890.3 |
| Solyc03g116900.2 | Solyc03g116900.3 |
| Solyc03g116910.2 | Solyc03g116910.3 |
| Solyc03g116930.2 | Solyc03g116930.3 |
| Solyc03g116950.1 | Solyc03g116950.2 |
| Solyc03g117000.2 | Solyc03g117000.3 |
| Solyc03g117030.1 | Solyc03g117030.2 |
| Solyc03g117040.2 | Solyc03g117040.3 |
| Solyc03g117100.2 | Solyc03g117100.3 |
| Solyc03g117130.2 | Solyc03g117130.3 |
| Solyc03g117150.2 | Solyc03g117150.3 |
| Solyc03g117180.2 | Solyc03g117180.3 |
| Solyc03g117260.1 | Solyc03g117260.2 |
| Solyc03g117370.2 | Solyc03g117370.3 |
| Solyc03g117425.1 | Solyc03g150159.1 |
| Solyc03g117430.2 | Solyc03g117430.3 |
| Solyc03g117440.2 | Solyc03g117440.3 |
| Solyc03g117470.2 | Solyc03g117470.3 |
| Solyc03g117560.2 | Solyc03g117560.3 |
| Solyc03g117570.2 | Solyc03g117570.3 |
| Solyc03g117590.2 | Solyc03g117590.3 |
| Solyc03g117600.2 | Solyc03g117600.3 |
| Solyc03g117610.1 | Solyc03g117610.2 |
| Solyc03g117690.2 | Solyc03g117690.3 |
| Solyc03g117720.2 | Solyc03g117720.3 |
| Solyc03g117730.2 | Solyc03g117730.3 |
| Solyc03g117750.2 | Solyc03g117750.3 |
| Solyc03g117770.2 | Solyc03g117770.3 |
| Solyc03g117790.1 | Solyc03g117790.2 |
| Solyc03g117830.1 | Solyc03g117830.2 |
| Solyc03g117860.2 | Solyc03g117860.3 |
| Solyc03g117870.2 | Solyc03g117870.3 |
| Solyc03g117890.2 | Solyc03g117890.3 |
| Solyc03g117900.2 | Solyc03g117900.3 |
| Solyc03g117910.2 | Solyc03g117910.3 |

| | |
|---|---|
| Solyc03g117940.2 | Solyc03g117940.3 |
| Solyc03g117950.2 | Solyc03g117950.3 |
| Solyc03g117980.2 | Solyc03g117980.3 |
| Solyc03g118020.2 | Solyc03g118020.3 |
| Solyc03g118030.2 | Solyc03g118030.3 |
| Solyc03g118040.2 | Solyc03g118040.3 |
| Solyc03g118070.2 | Solyc03g118070.3 |
| Solyc03g118090.2 | Solyc03g118090.3 |
| Solyc03g118120.2 | Solyc03g118120.3 |
| Solyc03g118130.2 | Solyc03g118130.3 |
| Solyc03g118140.2 | Solyc03g118140.3 |
| Solyc03g118150.2 | Solyc03g118150.3 |
| Solyc03g118170.2 | Solyc03g118170.3 |
| Solyc03g118180.2 | Solyc03g118180.3 |
| Solyc03g118210.2 | Solyc03g118210.3 |
| Solyc03g118290.2 | Solyc03g118290.3 |
| Solyc03g118340.2 | Solyc03g118340.3 |
| Solyc03g118370.2 | Solyc03g118370.3 |
| Solyc03g118390.2 | Solyc03g118390.3 |
| Solyc03g118410.2 | Solyc03g118410.3 |
| Solyc03g118430.2 | Solyc03g118430.3 |
| Solyc03g118500.2 | Solyc03g118500.3 |
| Solyc03g118510.2 | Solyc03g118510.3 |
| Solyc03g118530.2 | Solyc03g118530.3 |
| Solyc03g118550.2 | Solyc03g118550.3 |
| Solyc03g118580.2 | Solyc03g118580.3 |
| Solyc03g118585.1 | Solyc03g150160.1 |
| Solyc03g118620.2 | Solyc03g118620.3 |
| Solyc03g118640.2 | Solyc03g118640.3 |
| Solyc03g118650.2 | Solyc03g118650.3 |
| Solyc03g118660.2 | Solyc03g118660.3 |
| Solyc03g118670.2 | Solyc03g118670.3 |
| Solyc03g118680.2 | Solyc03g118680.3 |
| Solyc03g118710.2 | Solyc03g118710.3 |
| Solyc03g118720.2 | Solyc03g118720.3 |
| Solyc03g118740.2 | Solyc03g118740.3 |
| Solyc03g118780.2 | Solyc03g118780.3 |
| Solyc03g118800.2 | Solyc03g118800.3 |
| Solyc03g118830.2 | Solyc03g118830.3 |
| Solyc03g118840.2 | Solyc03g118840.3 |
| Solyc03g118860.2 | Solyc03g118860.3 |
| Solyc03g118880.2 | Solyc03g118880.3 |
| Solyc03g118920.2 | Solyc03g118920.3 |

| | |
|---|---|
| Solyc03g118970.2 | Solyc03g118970.3 |
| Solyc03g119010.2 | Solyc03g119010.3 |
| Solyc03g119040.2 | Solyc03g119040.3 |
| Solyc03g119060.2 | Solyc03g119060.3 |
| Solyc03g119100.1 | Solyc03g119100.2 |
| Solyc03g119120.2 | Solyc03g119120.3 |
| Solyc03g119130.2 | Solyc03g119130.3 |
| Solyc03g119160.2 | Solyc03g119160.3 |
| Solyc03g119170.2 | Solyc03g119170.3 |
| Solyc03g119190.2 | Solyc03g119190.3 |
| Solyc03g119220.2 | Solyc03g119220.3 |
| Solyc03g119240.2 | Solyc03g119240.3 |
| Solyc03g119280.2 | Solyc03g119280.3 |
| Solyc03g119290.2 | Solyc03g119290.3 |
| Solyc03g119310.2 | Solyc03g119310.3 |
| Solyc03g119330.2 | Solyc03g119330.3 |
| Solyc03g119340.2 | Solyc03g119340.3 |
| Solyc03g119360.2 | Solyc03g119360.3 |
| Solyc03g119370.1 | Solyc03g119370.2 |
| Solyc03g119400.2 | Solyc03g119400.3 |
| Solyc03g119420.2 | Solyc03g119420.3 |
| Solyc03g119440.2 | Solyc03g119440.3 |
| Solyc03g119450.2 | Solyc03g119450.3 |
| Solyc03g119470.1 | Solyc03g119470.2 |
| Solyc03g119510.2 | Solyc03g119510.3 |
| Solyc03g119530.2 | Solyc03g119530.3 |
| Solyc03g119540.2 | Solyc03g119540.3 |
| Solyc03g119650.2 | Solyc03g119650.3 |
| Solyc03g119660.1 | Solyc03g119660.2 |
| Solyc03g119670.2 | Solyc03g119670.3 |
| Solyc03g119700.2 | Solyc03g119700.3 |
| Solyc03g119730.2 | Solyc03g119730.3 |
| Solyc03g119740.2 | Solyc03g119740.3 |
| Solyc03g119760.2 | Solyc03g119760.3 |
| Solyc03g119790.2 | Solyc03g119790.3 |
| Solyc03g119840.2 | Solyc03g119840.3 |
| Solyc03g119850.1 | Solyc03g119850.2 |
| Solyc03g119860.2 | Solyc03g119860.3 |
| Solyc03g119870.2 | Solyc03g119870.3 |
| Solyc03g119880.2 | Solyc03g119880.3 |
| Solyc03g119960.2 | Solyc03g119960.3 |
| Solyc03g119970.2 | Solyc03g119970.3 |
| Solyc03g119980.2 | Solyc03g119980.3 |

| | |
|---|---|
| Solyc03g120000.2 | Solyc03g120000.3 |
| Solyc03g120040.2 | Solyc03g120040.3 |
| Solyc03g120050.2 | Solyc03g120050.3 |
| Solyc03g120100.2 | Solyc03g120100.3 |
| Solyc03g120110.2 | Solyc03g120110.3 |
| Solyc03g120120.2 | Solyc03g120120.3 |
| Solyc03g120140.2 | Solyc03g120140.3 |
| Solyc03g120150.2 | Solyc03g120150.3 |
| Solyc03g120190.2 | Solyc03g120190.3 |
| Solyc03g120200.1 | Solyc03g120200.2 |
| Solyc03g120230.2 | Solyc03g120230.3 |
| Solyc03g120240.2 | Solyc03g120240.3 |
| Solyc03g120250.2 | Solyc03g120250.3 |
| Solyc03g120290.2 | Solyc03g120290.3 |
| Solyc03g120310.2 | Solyc03g120310.3 |
| Solyc03g120340.2 | Solyc03g120340.3 |
| Solyc03g120350.2 | Solyc03g120350.3 |
| Solyc03g120380.2 | Solyc03g120380.3 |
| Solyc03g120390.2 | Solyc03g120390.3 |
| Solyc03g120420.2 | Solyc03g120420.3 |
| Solyc03g120430.2 | Solyc03g120430.3 |
| Solyc03g120520.1 | Solyc03g120520.2 |
| Solyc03g120530.2 | Solyc03g120530.3 |
| Solyc03g120540.2 | Solyc03g120540.3 |
| Solyc03g120560.2 | Solyc03g120560.3 |
| Solyc03g120570.2 | Solyc03g120570.3 |
| Solyc03g120580.2 | Solyc03g120580.3 |
| Solyc03g120610.2 | Solyc03g120610.3 |
| Solyc03g120620.2 | Solyc03g120620.3 |
| Solyc03g120640.2 | Solyc03g120640.3 |
| Solyc03g120670.2 | Solyc03g120670.3 |
| Solyc03g120680.2 | Solyc03g120680.3 |
| Solyc03g120690.2 | Solyc03g120690.3 |
| Solyc03g120700.2 | Solyc03g120700.3 |
| Solyc03g120720.2 | Solyc03g120720.3 |
| Solyc03g120730.2 | Solyc03g120730.3 |
| Solyc03g120750.2 | Solyc03g120750.3 |
| Solyc03g120780.2 | Solyc03g120780.3 |
| Solyc03g120870.2 | Solyc03g120870.3 |
| Solyc03g120890.2 | Solyc03g120890.3 |
| Solyc03g120970.2 | Solyc03g120970.3 |
| Solyc03g120980.2 | Solyc03g120980.3 |
| Solyc03g120990.2 | Solyc03g120990.3 |

| | |
|---|---|
| Solyc03g121000.2 | Solyc03g121000.3 |
| Solyc03g121020.2 | Solyc03g121020.3 |
| Solyc03g121030.1 | Solyc03g121030.2 |
| Solyc03g121050.2 | Solyc03g121050.3 |
| Solyc03g121070.2 | Solyc03g121070.3 |
| Solyc03g121080.2 | Solyc03g121080.3 |
| Solyc03g121110.2 | Solyc03g121110.3 |
| Solyc03g121130.2 | Solyc03g121130.3 |
| Solyc03g121170.2 | Solyc03g121170.3 |
| Solyc03g121230.2 | Solyc03g121230.3 |
| Solyc03g121250.2 | Solyc03g121250.3 |
| Solyc03g121260.2 | Solyc03g121260.3 |
| Solyc03g121270.2 | Solyc03g121270.3 |
| Solyc03g121320.2 | Solyc03g121320.3 |
| Solyc03g121330.2 | Solyc03g121330.3 |
| Solyc03g121350.2 | Solyc03g121350.3 |
| Solyc03g121370.2 | Solyc03g121370.3 |
| Solyc03g121380.2 | Solyc03g121380.3 |
| Solyc03g121410.2 | Solyc03g121410.3 |
| Solyc03g121450.2 | Solyc03g121450.3 |
| Solyc03g121480.2 | Solyc03g121480.3 |
| Solyc03g121500.2 | Solyc03g121500.3 |
| Solyc03g121530.2 | Solyc03g121530.3 |
| Solyc03g121540.2 | Solyc03g121540.3 |
| Solyc03g121550.2 | Solyc03g121550.3 |
| Solyc03g121570.2 | Solyc03g121570.3 |
| Solyc03g121580.2 | Solyc03g121580.3 |
| Solyc03g121590.2 | Solyc03g121590.3 |
| Solyc03g121610.2 | Solyc03g121610.3 |
| Solyc03g121630.2 | Solyc03g121630.3 |
| Solyc03g121640.2 | Solyc03g121640.3 |
| Solyc03g121650.2 | Solyc03g121650.3 |
| Solyc03g121660.2 | Solyc03g121660.3 |
| Solyc03g121730.2 | Solyc03g121730.3 |
| Solyc03g121745.1 | Solyc03g150161.1 |
| Solyc03g121760.2 | Solyc03g121760.3 |
| Solyc03g121770.1 | Solyc03g121770.2 |
| Solyc03g121820.2 | Solyc03g121820.3 |
| Solyc03g121840.2 | Solyc03g121840.3 |
| Solyc03g121870.2 | Solyc03g121870.3 |
| Solyc03g121920.2 | Solyc03g121920.3 |
| Solyc03g121930.2 | Solyc03g121930.3 |
| Solyc03g121940.2 | Solyc03g121940.3 |

| | |
|---|---|
| Solyc03g121950.2 | Solyc03g121950.3 |
| Solyc03g121960.2 | Solyc03g121960.3 |
| Solyc03g121980.2 | Solyc03g121980.3 |
| Solyc03g122080.2 | Solyc03g122080.3 |
| Solyc03g122090.2 | Solyc03g122090.3 |
| Solyc03g122130.2 | Solyc03g122130.3 |
| Solyc03g122140.2 | Solyc03g122140.3 |
| Solyc03g122150.2 | Solyc03g122150.3 |
| Solyc03g122190.2 | Solyc03g122190.3 |
| Solyc03g122230.1 | Solyc03g122230.2 |
| Solyc03g122240.2 | Solyc03g122240.3 |
| Solyc03g122250.2 | Solyc03g122250.3 |
| Solyc03g122260.2 | Solyc03g122260.3 |
| Solyc03g122280.2 | Solyc03g122280.3 |
| Solyc03g122340.2 | Solyc03g122340.3 |
| Solyc03g122350.2 | Solyc03g122350.3 |
| Solyc03g122360.2 | Solyc03g122360.3 |
| Solyc03g123370.2 | Solyc03g123370.3 |
| Solyc03g123440.2 | Solyc03g123440.3 |
| Solyc03g123480.1 | Solyc03g123480.2 |
| Solyc03g123540.2 | Solyc03g123540.3 |
| Solyc03g123560.2 | Solyc03g123560.3 |
| Solyc03g123570.2 | Solyc03g123570.3 |
| Solyc03g123630.3 | Solyc03g123630.4 |
| Solyc03g123640.2 | Solyc03g123640.3 |
| Solyc03g123660.2 | Solyc03g123660.3 |
| Solyc03g123670.2 | Solyc03g123670.3 |
| Solyc03g123710.2 | Solyc03g123710.3 |
| Solyc03g123750.2 | Solyc03g123750.3 |
| Solyc03g123760.2 | Solyc03g123760.3 |
| Solyc03g123820.1 | Solyc03g123820.2 |
| Solyc03g123870.2 | Solyc03g123870.3 |
| Solyc03g123880.2 | Solyc03g123880.3 |
| Solyc03g123900.2 | Solyc03g123900.3 |
| Solyc03g123910.2 | Solyc03g123910.3 |
| Solyc03g123920.2 | Solyc03g123920.3 |
| Solyc03g123960.2 | Solyc03g123960.3 |
| Solyc03g123990.2 | Solyc03g123990.3 |
| Solyc03g124010.2 | Solyc03g124010.3 |
| Solyc03g124040.2 | Solyc03g124040.3 |
| Solyc03g124050.2 | Solyc03g124050.3 |
| Solyc03g124060.2 | Solyc03g124060.3 |
| Solyc03g006715.1 | Solyc03g006715.2 |

| | |
|---|---|
| Solyc04g005020.2 | Solyc04g005020.3 |
| Solyc04g005060.2 | Solyc04g005060.3 |
| Solyc04g005070.2 | Solyc04g005070.3 |
| Solyc04g005080.2 | Solyc04g005080.3 |
| Solyc04g005100.2 | Solyc04g005100.3 |
| Solyc04g005130.2 | Solyc04g005130.3 |
| Solyc04g005205.1 | Solyc04g150100.1 |
| Solyc04g005230.2 | Solyc04g005230.3 |
| Solyc04g005240.1 | Solyc04g005240.2 |
| Solyc04g005260.2 | Solyc04g005260.3 |
| Solyc04g005310.2 | Solyc04g005310.3 |
| Solyc04g005320.2 | Solyc04g005320.3 |
| Solyc04g005330.2 | Solyc04g005330.3 |
| Solyc04g005350.2 | Solyc04g005350.3 |
| Solyc04g005380.2 | Solyc04g005380.3 |
| Solyc04g005450.2 | Solyc04g005450.3 |
| Solyc04g005470.2 | Solyc04g005470.3 |
| Solyc04g005490.2 | Solyc04g005490.3 |
| Solyc04g005510.2 | Solyc04g005510.3 |
| Solyc04g005520.2 | Solyc04g005520.3 |
| Solyc04g005540.2 | Solyc04g005540.3 |
| Solyc04g005550.1 | Solyc04g005550.2 |
| Solyc04g005560.2 | Solyc04g005560.3 |
| Solyc04g005590.2 | Solyc04g005590.3 |
| Solyc04g005610.2 | Solyc04g005610.3 |
| Solyc04g005620.2 | Solyc04g005620.3 |
| Solyc04g005630.2 | Solyc04g005630.3 |
| Solyc04g005640.2 | Solyc04g005640.3 |
| Solyc04g005720.2 | Solyc04g005720.3 |
| Solyc04g005730.2 | Solyc04g005730.3 |
| Solyc04g005790.2 | Solyc04g005790.3 |
| Solyc04g005800.2 | Solyc04g005800.3 |
| Solyc04g005810.2 | Solyc04g005810.3 |
| Solyc04g005830.2 | Solyc04g005830.3 |
| Solyc04g005860.2 | Solyc04g005860.3 |
| Solyc04g005900.2 | Solyc04g005900.3 |
| Solyc04g005910.2 | Solyc04g005910.3 |
| Solyc04g006930.2 | Solyc04g006930.3 |
| Solyc04g006960.2 | Solyc04g006960.3 |
| Solyc04g006990.2 | Solyc04g006990.3 |
| Solyc04g007000.1 | Solyc04g007000.2 |
| Solyc04g007010.2 | Solyc04g007010.3 |
| Solyc04g007070.2 | Solyc04g007070.3 |

| | |
|---|---|
| Solyc04g007115.1 | Solyc04g150101.1 |
| Solyc04g007150.2 | Solyc04g007150.3 |
| Solyc04g007170.2 | Solyc04g007170.3 |
| Solyc04g007210.2 | Solyc04g007210.3 |
| Solyc04g007230.2 | Solyc04g007230.3 |
| Solyc04g007270.2 | Solyc04g007270.3 |
| Solyc04g007280.2 | Solyc04g007280.3 |
| Solyc04g007310.1 | Solyc04g007310.2 |
| Solyc04g007350.2 | Solyc04g007350.3 |
| Solyc04g007370.2 | Solyc04g007370.3 |
| Solyc04g007430.1 | Solyc04g007430.2 |
| Solyc04g007470.2 | Solyc04g007470.3 |
| Solyc04g007490.2 | Solyc04g007490.3 |
| Solyc04g007520.2 | Solyc04g007520.3 |
| Solyc04g007530.2 | Solyc04g007530.3 |
| Solyc04g007560.2 | Solyc04g007560.3 |
| Solyc04g007570.1 | Solyc04g007570.2 |
| Solyc04g007610.2 | Solyc04g007610.3 |
| Solyc04g007630.1 | Solyc04g007630.2 |
| Solyc04g007640.2 | Solyc04g007640.3 |
| Solyc04g007690.2 | Solyc04g007690.3 |
| Solyc04g007710.2 | Solyc04g007710.3 |
| Solyc04g007720.2 | Solyc04g007720.3 |
| Solyc04g007760.2 | Solyc04g007760.3 |
| Solyc04g007763.1 | Solyc04g150102.1 |
| Solyc04g007767.1 | Solyc04g150103.1 |
| Solyc04g007770.2 | Solyc04g007770.3 |
| Solyc04g007780.2 | Solyc04g007780.3 |
| Solyc04g007790.2 | Solyc04g007790.3 |
| Solyc04g007795.1 | Solyc04g150104.1 |
| Solyc04g007820.2 | Solyc04g007820.3 |
| Solyc04g007825.1 | Solyc04g007825.2 |
| Solyc04g007850.2 | Solyc04g007850.3 |
| Solyc04g007860.2 | Solyc04g007860.3 |
| Solyc04g007870.2 | Solyc04g007870.3 |
| Solyc04g007890.2 | Solyc04g007890.3 |
| Solyc04g007900.2 | Solyc04g007900.3 |
| Solyc04g007910.2 | Solyc04g007910.3 |
| Solyc04g007970.2 | Solyc04g007970.3 |
| Solyc04g007980.2 | Solyc04g007980.3 |
| Solyc04g008000.2 | Solyc04g008000.3 |
| Solyc04g008020.2 | Solyc04g008020.3 |
| Solyc04g008030.1 | Solyc04g008030.2 |

| | |
|---|---|
| Solyc04g008060.2 | Solyc04g008060.3 |
| Solyc04g008075.1 | Solyc04g150105.1 |
| Solyc04g008110.2 | Solyc04g008110.3 |
| Solyc04g008210.1 | Solyc04g008210.2 |
| Solyc04g008230.2 | Solyc04g008230.3 |
| Solyc04g008240.2 | Solyc04g008240.3 |
| Solyc04g008260.2 | Solyc04g008260.3 |
| Solyc04g008470.2 | Solyc04g008470.3 |
| Solyc04g008480.1 | Solyc04g008480.2 |
| Solyc04g008490.1 | Solyc04g008490.2 |
| Solyc04g008510.2 | Solyc04g008510.3 |
| Solyc04g008520.2 | Solyc04g008520.3 |
| Solyc04g008530.1 | Solyc04g008530.2 |
| Solyc04g008540.2 | Solyc04g008540.3 |
| Solyc04g008580.2 | Solyc04g008580.3 |
| Solyc04g008590.2 | Solyc04g008590.3 |
| Solyc04g008610.2 | Solyc04g008610.3 |
| Solyc04g008620.2 | Solyc04g008620.3 |
| Solyc04g008630.2 | Solyc04g008630.3 |
| Solyc04g008640.2 | Solyc04g008640.3 |
| Solyc04g008670.1 | Solyc04g008670.2 |
| Solyc04g008690.2 | Solyc04g008690.3 |
| Solyc04g008700.2 | Solyc04g008700.3 |
| Solyc04g008730.2 | Solyc04g008730.3 |
| Solyc04g008740.2 | Solyc04g008740.3 |
| Solyc04g008750.2 | Solyc04g008750.3 |
| Solyc04g008770.2 | Solyc04g008770.3 |
| Solyc04g008810.2 | Solyc04g008810.3 |
| Solyc04g008820.2 | Solyc04g008820.3 |
| Solyc04g008840.2 | Solyc04g008840.3 |
| Solyc04g008870.2 | Solyc04g008870.3 |
| Solyc04g008890.2 | Solyc04g008890.3 |
| Solyc04g008930.1 | Solyc04g008930.2 |
| Solyc04g008960.2 | Solyc04g008960.3 |
| Solyc04g008970.2 | Solyc04g008970.3 |
| Solyc04g008980.2 | Solyc04g008980.3 |
| Solyc04g008990.2 | Solyc04g008990.3 |
| Solyc04g009010.1 | Solyc04g009010.2 |
| Solyc04g009030.2 | Solyc04g009030.3 |
| Solyc04g009090.1 | Solyc04g009090.2 |
| Solyc04g009190.2 | Solyc04g009190.3 |
| Solyc04g009200.2 | Solyc04g009200.3 |
| Solyc04g009220.1 | Solyc04g009220.2 |

| | |
|---|---|
| Solyc04g009240.1 | Solyc04g009240.2 |
| Solyc04g009320.2 | Solyc04g009320.3 |
| Solyc04g009340.2 | Solyc04g009340.3 |
| Solyc04g009350.2 | Solyc04g009350.3 |
| Solyc04g009360.2 | Solyc04g009360.3 |
| Solyc04g009390.1 | Solyc04g009390.2 |
| Solyc04g009410.2 | Solyc04g009410.3 |
| Solyc04g009420.2 | Solyc04g009420.3 |
| Solyc04g009430.2 | Solyc04g009430.3 |
| Solyc04g009440.2 | Solyc04g009440.3 |
| Solyc04g009520.2 | Solyc04g009520.3 |
| Solyc04g009550.2 | Solyc04g009550.3 |
| Solyc04g009560.2 | Solyc04g009560.3 |
| Solyc04g009580.2 | Solyc04g009580.3 |
| Solyc04g009630.2 | Solyc04g009630.3 |
| Solyc04g009680.1 | Solyc04g009680.2 |
| Solyc04g009690.1 | Solyc04g009690.2 |
| Solyc04g009700.2 | Solyc04g009700.3 |
| Solyc04g009710.1 | Solyc04g009710.2 |
| Solyc04g009760.1 | Solyc04g009760.2 |
| Solyc04g009770.2 | Solyc04g009770.3 |
| Solyc04g009800.2 | Solyc04g009800.3 |
| Solyc04g009810.2 | Solyc04g009810.3 |
| Solyc04g009820.2 | Solyc04g009820.3 |
| Solyc04g009830.2 | Solyc04g009830.3 |
| Solyc04g009910.2 | Solyc04g009910.3 |
| Solyc04g009920.2 | Solyc04g009920.3 |
| Solyc04g009950.2 | Solyc04g009950.3 |
| Solyc04g009960.2 | Solyc04g009960.3 |
| Solyc04g009990.2 | Solyc04g009990.3 |
| Solyc04g010000.2 | Solyc04g010000.3 |
| Solyc04g010040.2 | Solyc04g010040.3 |
| Solyc04g010080.2 | Solyc04g010080.3 |
| Solyc04g010090.2 | Solyc04g010090.3 |
| Solyc04g010100.1 | Solyc04g010100.2 |
| Solyc04g010110.2 | Solyc04g010110.3 |
| Solyc04g010120.2 | Solyc04g010120.3 |
| Solyc04g010240.2 | Solyc04g010240.3 |
| Solyc04g010250.2 | Solyc04g010250.3 |
| Solyc04g010260.2 | Solyc04g010260.3 |
| Solyc04g010280.1 | Solyc04g010280.2 |
| Solyc04g010330.2 | Solyc04g010330.3 |
| Solyc04g010333.1 | Solyc04g150106.1 |

| | |
|---|---|
| Solyc04g010337.1 | Solyc04g150107.1 |
| Solyc04g011340.2 | Solyc04g011340.3 |
| Solyc04g011350.2 | Solyc04g011350.3 |
| Solyc04g011360.2 | Solyc04g011360.3 |
| Solyc04g011400.2 | Solyc04g011400.3 |
| Solyc04g011430.2 | Solyc04g011430.3 |
| Solyc04g011480.2 | Solyc04g011480.3 |
| Solyc04g011495.1 | Solyc04g150108.1 |
| Solyc04g011500.2 | Solyc04g011500.3 |
| Solyc04g011520.2 | Solyc04g011520.3 |
| Solyc04g011530.2 | Solyc04g011530.3 |
| Solyc04g011570.2 | Solyc04g011570.3 |
| Solyc04g011590.2 | Solyc04g011590.3 |
| Solyc04g011620.2 | Solyc04g011620.3 |
| Solyc04g011640.2 | Solyc04g011640.3 |
| Solyc04g011650.2 | Solyc04g011650.3 |
| Solyc04g011670.2 | Solyc04g011670.3 |
| Solyc04g011700.1 | Solyc04g011700.2 |
| Solyc04g011720.2 | Solyc04g011720.3 |
| Solyc04g011730.2 | Solyc04g011730.3 |
| Solyc04g011750.2 | Solyc04g011750.3 |
| Solyc04g011760.2 | Solyc04g011760.3 |
| Solyc04g011925.1 | Solyc04g150109.1 |
| Solyc04g011960.1 | Solyc04g011960.2 |
| Solyc04g012010.2 | Solyc04g012010.3 |
| Solyc04g012030.2 | Solyc04g012030.3 |
| Solyc04g012040.2 | Solyc04g012040.3 |
| Solyc04g012050.2 | Solyc04g012050.3 |
| Solyc04g012060.2 | Solyc04g012060.3 |
| Solyc04g012110.1 | Solyc04g012110.2 |
| Solyc04g012120.2 | Solyc04g012120.3 |
| Solyc04g014210.2 | Solyc04g014210.3 |
| Solyc04g014250.2 | Solyc04g014250.3 |
| Solyc04g014260.1 | Solyc04g014260.2 |
| Solyc04g014270.2 | Solyc04g014270.3 |
| Solyc04g014360.2 | Solyc04g014360.3 |
| Solyc04g014380.2 | Solyc04g014380.3 |
| Solyc04g014390.2 | Solyc04g014390.3 |
| Solyc04g014430.1 | Solyc04g014430.2 |
| Solyc04g014470.2 | Solyc04g014470.3 |
| Solyc04g014480.2 | Solyc04g014480.3 |
| Solyc04g014500.2 | Solyc04g014500.3 |
| Solyc04g014510.2 | Solyc04g014510.3 |

| | |
|---|---|
| Solyc04g014550.2 | Solyc04g014550.3 |
| Solyc04g014570.2 | Solyc04g014570.3 |
| Solyc04g014578.1 | Solyc04g014578.2 |
| Solyc04g014600.2 | Solyc04g014600.3 |
| Solyc04g014640.1 | Solyc04g014640.2 |
| Solyc04g014670.2 | Solyc04g014670.3 |
| Solyc04g014690.2 | Solyc04g014690.3 |
| Solyc04g014760.2 | Solyc04g014760.3 |
| Solyc04g014800.2 | Solyc04g014800.3 |
| Solyc04g014810.2 | Solyc04g014810.3 |
| Solyc04g014840.2 | Solyc04g014840.3 |
| Solyc04g014857.1 | Solyc04g150110.1 |
| Solyc04g014870.2 | Solyc04g014870.3 |
| Solyc04g014940.1 | Solyc04g014940.2 |
| Solyc04g014970.1 | Solyc04g014970.2 |
| Solyc04g015000.1 | Solyc04g015000.2 |
| Solyc04g015020.2 | Solyc04g015020.3 |
| Solyc04g015040.2 | Solyc04g015040.3 |
| Solyc04g015050.2 | Solyc04g015050.3 |
| Solyc04g015070.2 | Solyc04g015070.3 |
| Solyc04g015100.2 | Solyc04g015100.3 |
| Solyc04g015110.2 | Solyc04g015110.3 |
| Solyc04g015120.2 | Solyc04g015120.3 |
| Solyc04g015130.2 | Solyc04g015130.3 |
| Solyc04g015160.2 | Solyc04g015160.3 |
| Solyc04g015170.2 | Solyc04g015170.3 |
| Solyc04g015180.2 | Solyc04g015180.3 |
| Solyc04g015190.2 | Solyc04g015190.3 |
| Solyc04g015200.2 | Solyc04g015200.3 |
| Solyc04g015270.2 | Solyc04g015270.3 |
| Solyc04g015280.2 | Solyc04g015280.3 |
| Solyc04g015290.2 | Solyc04g015290.3 |
| Solyc04g015300.2 | Solyc04g015300.3 |
| Solyc04g015330.2 | Solyc04g015330.3 |
| Solyc04g015340.2 | Solyc04g015340.3 |
| Solyc04g015350.2 | Solyc04g015350.3 |
| Solyc04g015360.2 | Solyc04g015360.3 |
| Solyc04g015370.2 | Solyc04g015370.3 |
| Solyc04g015450.2 | Solyc04g015450.3 |
| Solyc04g015470.2 | Solyc04g015470.3 |
| Solyc04g015500.2 | Solyc04g015500.3 |
| Solyc04g015520.2 | Solyc04g015520.3 |
| Solyc04g015570.2 | Solyc04g015570.3 |

| | |
|---|---|
| Solyc04g015580.2 | Solyc04g015580.3 |
| Solyc04g015600.2 | Solyc04g015600.3 |
| Solyc04g015610.2 | Solyc04g015610.3 |
| Solyc04g015620.2 | Solyc04g015620.3 |
| Solyc04g015635.1 | Solyc04g015635.2 |
| Solyc04g015680.2 | Solyc04g015680.3 |
| Solyc04g015720.1 | Solyc04g015720.2 |
| Solyc04g015750.2 | Solyc04g015750.3 |
| Solyc04g015830.2 | Solyc04g015830.3 |
| Solyc04g015850.2 | Solyc04g015850.3 |
| Solyc04g015940.2 | Solyc04g015940.3 |
| Solyc04g015960.2 | Solyc04g015960.3 |
| Solyc04g015970.2 | Solyc04g015970.3 |
| Solyc04g015990.1 | Solyc04g015990.2 |
| Solyc04g016000.2 | Solyc04g016000.3 |
| Solyc04g016015.1 | Solyc04g150112.1 |
| Solyc04g016050.2 | Solyc04g016050.3 |
| Solyc04g016080.1 | Solyc04g016080.2 |
| Solyc04g016130.2 | Solyc04g016130.3 |
| Solyc04g016150.1 | Solyc04g016150.2 |
| Solyc04g016180.2 | Solyc04g016180.3 |
| Solyc04g016175.1 | Solyc04g016175.2 |
| Solyc04g016240.2 | Solyc04g016240.3 |
| Solyc04g016250.2 | Solyc04g016250.3 |
| Solyc04g016260.2 | Solyc04g016260.3 |
| Solyc04g016270.1 | Solyc04g016270.2 |
| Solyc04g016280.2 | Solyc04g016280.3 |
| Solyc04g016290.2 | Solyc04g016290.3 |
| Solyc04g016330.2 | Solyc04g016330.3 |
| Solyc04g016360.2 | Solyc04g016360.3 |
| Solyc04g016370.2 | Solyc04g016370.3 |
| Solyc04g016380.2 | Solyc04g016380.3 |
| Solyc04g016390.1 | Solyc04g016390.2 |
| Solyc04g016410.2 | Solyc04g016410.3 |
| Solyc04g016430.2 | Solyc04g016430.3 |
| Solyc04g016460.2 | Solyc04g016460.3 |
| Solyc04g016480.2 | Solyc04g016480.3 |
| Solyc04g016510.2 | Solyc04g016510.3 |
| Solyc04g016520.2 | Solyc04g016520.3 |
| Solyc04g016530.2 | Solyc04g016530.3 |
| Solyc04g016540.1 | Solyc04g016540.2 |
| Solyc04g016560.1 | Solyc04g016560.2 |
| Solyc04g016590.2 | Solyc04g016590.3 |

| | |
|---|---|
| Solyc04g017670.2 | Solyc04g017670.3 |
| Solyc04g017680.2 | Solyc04g017680.3 |
| Solyc04g017690.2 | Solyc04g017690.3 |
| Solyc04g017710.2 | Solyc04g017710.3 |
| Solyc04g017720.2 | Solyc04g017720.3 |
| Solyc04g017727.1 | Solyc04g150114.1 |
| Solyc04g017965.1 | Solyc04g150115.1 |
| Solyc04g019315.1 | Solyc04g150116.1 |
| Solyc04g040220.2 | Solyc04g040220.3 |
| Solyc04g040200.2 | Solyc04g040200.3 |
| Solyc04g040180.2 | Solyc04g040180.3 |
| Solyc04g040170.2 | Solyc04g040170.3 |
| Solyc04g040140.1 | Solyc04g040140.2 |
| Solyc04g040110.2 | Solyc04g040110.3 |
| Solyc04g040095.1 | Solyc04g150123.1 |
| Solyc04g040010.2 | Solyc04g040010.3 |
| Solyc04g039980.2 | Solyc04g039980.3 |
| Solyc04g039730.2 | Solyc04g039730.3 |
| Solyc04g039670.2 | Solyc04g039670.3 |
| Solyc04g028470.1 | Solyc04g028470.2 |
| Solyc04g026360.2 | Solyc04g026360.3 |
| Solyc04g026350.2 | Solyc04g026350.3 |
| Solyc04g026340.1 | Solyc04g026340.2 |
| Solyc04g026210.2 | Solyc04g026210.3 |
| Solyc04g026200.2 | Solyc04g026200.3 |
| Solyc04g026150.1 | Solyc04g026150.2 |
| Solyc04g026030.2 | Solyc04g026030.3 |
| Solyc04g026020.2 | Solyc04g026020.3 |
| Solyc04g026023.1 | Solyc04g026023.2 |
| Solyc04g026000.1 | Solyc04g026000.2 |
| Solyc04g025990.2 | Solyc04g025990.3 |
| Solyc04g025940.2 | Solyc04g025940.3 |
| Solyc04g025825.1 | Solyc04g150122.1 |
| Solyc04g025750.2 | Solyc04g025750.3 |
| Solyc04g025740.1 | Solyc04g025740.2 |
| Solyc04g025650.2 | Solyc04g025650.3 |
| Solyc04g025610.1 | Solyc04g025610.2 |
| Solyc04g025585.1 | Solyc04g150121.1 |
| Solyc04g025540.2 | Solyc04g025540.3 |
| Solyc04g025530.2 | Solyc04g025530.3 |
| Solyc04g025535.1 | Solyc04g150120.1 |
| Solyc04g025460.1 | Solyc04g025460.2 |
| Solyc04g025440.2 | Solyc04g025440.3 |

| | |
|---|---|
| Solyc04g025400.2 | Solyc04g025400.3 |
| Solyc04g025320.1 | Solyc04g025320.2 |
| Solyc04g025300.2 | Solyc04g025300.3 |
| Solyc04g025260.2 | Solyc04g025260.3 |
| Solyc04g025160.2 | Solyc04g025160.3 |
| Solyc04g025120.1 | Solyc04g025120.2 |
| Solyc04g025000.1 | Solyc04g025000.2 |
| Solyc04g024950.1 | Solyc04g024950.2 |
| Solyc04g024940.2 | Solyc04g024940.3 |
| Solyc04g024885.1 | Solyc04g150118.1 |
| Solyc04g024840.2 | Solyc04g024840.3 |
| Solyc04g024810.1 | Solyc04g024810.2 |
| Solyc04g024785.1 | Solyc04g150117.1 |
| Solyc04g024590.2 | Solyc04g024590.3 |
| Solyc04g024580.1 | Solyc04g024580.2 |
| Solyc04g024530.2 | Solyc04g024530.3 |
| Solyc04g024510.2 | Solyc04g024510.3 |
| Solyc04g024430.2 | Solyc04g024430.3 |
| Solyc04g024420.2 | Solyc04g024420.3 |
| Solyc04g045280.1 | Solyc04g045280.2 |
| Solyc04g045285.1 | Solyc04g150124.1 |
| Solyc04g045330.1 | Solyc04g045330.2 |
| Solyc04g045340.2 | Solyc04g045340.3 |
| Solyc04g045345.1 | Solyc04g150125.1 |
| Solyc04g045440.2 | Solyc04g045440.3 |
| Solyc04g045480.2 | Solyc04g045480.3 |
| Solyc04g045520.2 | Solyc04g045520.3 |
| Solyc04g045590.2 | Solyc04g045590.3 |
| Solyc04g045630.2 | Solyc04g045630.3 |
| Solyc04g045650.2 | Solyc04g045650.3 |
| Solyc04g047750.2 | Solyc04g047750.3 |
| Solyc04g047760.1 | Solyc04g047760.2 |
| Solyc04g047770.2 | Solyc04g047770.3 |
| Solyc04g048900.2 | Solyc04g048900.3 |
| Solyc04g048970.1 | Solyc04g048970.2 |
| Solyc04g049010.2 | Solyc04g049010.3 |
| Solyc04g049070.2 | Solyc04g049070.3 |
| Solyc04g049090.2 | Solyc04g049090.3 |
| Solyc04g049135.1 | Solyc04g150126.1 |
| Solyc04g049160.2 | Solyc04g049160.3 |
| Solyc04g049180.2 | Solyc04g049180.3 |
| Solyc04g049224.1 | Solyc04g150127.1 |
| Solyc04g049226.1 | Solyc04g150128.1 |

| | |
|---|---|
| Solyc04g049300.1 | Solyc04g049300.2 |
| Solyc04g049330.2 | Solyc04g049330.3 |
| Solyc04g049340.2 | Solyc04g049340.3 |
| Solyc04g049360.2 | Solyc04g049360.3 |
| Solyc04g049390.2 | Solyc04g049390.3 |
| Solyc04g049400.2 | Solyc04g049400.3 |
| Solyc04g049440.1 | Solyc04g049440.2 |
| Solyc04g049445.1 | Solyc04g150129.1 |
| Solyc04g049450.2 | Solyc04g049450.3 |
| Solyc04g049485.1 | Solyc04g150130.1 |
| Solyc04g049490.2 | Solyc04g049490.3 |
| Solyc04g049550.2 | Solyc04g049550.3 |
| Solyc04g049630.2 | Solyc04g049630.3 |
| Solyc04g049690.2 | Solyc04g049690.3 |
| Solyc04g049710.2 | Solyc04g049710.3 |
| Solyc04g049750.2 | Solyc04g049750.3 |
| Solyc04g049790.2 | Solyc04g049790.3 |
| Solyc04g049800.2 | Solyc04g049800.3 |
| Solyc04g049803.1 | Solyc04g150131.1 |
| Solyc04g049850.2 | Solyc04g049850.3 |
| Solyc04g049960.2 | Solyc04g049960.3 |
| Solyc04g049990.1 | Solyc04g049990.2 |
| Solyc04g050010.1 | Solyc04g050010.2 |
| Solyc04g050040.2 | Solyc04g050040.3 |
| Solyc04g050050.2 | Solyc04g050050.3 |
| Solyc04g050140.2 | Solyc04g050140.3 |
| Solyc04g050180.2 | Solyc04g050180.3 |
| Solyc04g050220.1 | Solyc04g050220.2 |
| Solyc04g050225.1 | Solyc04g150132.1 |
| Solyc04g050240.2 | Solyc04g050240.3 |
| Solyc04g050245.1 | Solyc04g150133.1 |
| Solyc04g050305.1 | Solyc04g150134.1 |
| Solyc04g050310.2 | Solyc04g050310.3 |
| Solyc04g050345.1 | Solyc04g150135.1 |
| Solyc04g050360.1 | Solyc04g050360.2 |
| Solyc04g050440.2 | Solyc04g050440.3 |
| Solyc04g050510.2 | Solyc04g050510.3 |
| Solyc04g050520.2 | Solyc04g050520.3 |
| Solyc04g050570.2 | Solyc04g050570.3 |
| Solyc04g050620.2 | Solyc04g050620.3 |
| Solyc04g050680.1 | Solyc04g050680.2 |
| Solyc04g050730.2 | Solyc04g050730.3 |
| Solyc04g050750.1 | Solyc04g050750.2 |

| | |
|---|---|
| Solyc04g050770.1 | Solyc04g050770.2 |
| Solyc04g050780.1 | Solyc04g050780.2 |
| Solyc04g050895.1 | Solyc04g150136.1 |
| Solyc04g050930.2 | Solyc04g050930.3 |
| Solyc04g050935.1 | Solyc04g150137.1 |
| Solyc04g050950.2 | Solyc04g050950.3 |
| Solyc04g050960.2 | Solyc04g050960.3 |
| Solyc04g050970.2 | Solyc04g050970.3 |
| Solyc04g050990.2 | Solyc04g050990.3 |
| Solyc04g051020.1 | Solyc04g051020.2 |
| Solyc04g051045.1 | Solyc04g150138.1 |
| Solyc04g051060.1 | Solyc04g051060.2 |
| Solyc04g051080.1 | Solyc04g051080.2 |
| Solyc04g051090.1 | Solyc04g051090.2 |
| Solyc04g051100.2 | Solyc04g051100.3 |
| Solyc04g051150.2 | Solyc04g051150.3 |
| Solyc04g051190.2 | Solyc04g051190.3 |
| Solyc04g051200.1 | Solyc04g051200.2 |
| Solyc04g051270.1 | Solyc04g051270.2 |
| Solyc04g051340.2 | Solyc04g051340.3 |
| Solyc04g051350.2 | Solyc04g051350.3 |
| Solyc04g051360.2 | Solyc04g051360.3 |
| Solyc04g051400.1 | Solyc04g051400.2 |
| Solyc04g051420.2 | Solyc04g051420.3 |
| Solyc04g051427.1 | Solyc04g051427.2 |
| Solyc04g051450.2 | Solyc04g051450.3 |
| Solyc04g051490.2 | Solyc04g051490.3 |
| Solyc04g051500.2 | Solyc04g051500.3 |
| Solyc04g051540.2 | Solyc04g051540.3 |
| Solyc04g051590.2 | Solyc04g051590.3 |
| Solyc04g051610.2 | Solyc04g051610.3 |
| Solyc04g051640.2 | Solyc04g051640.3 |
| Solyc04g051650.2 | Solyc04g051650.3 |
| Solyc04g051660.2 | Solyc04g051660.3 |
| Solyc04g051670.2 | Solyc04g051670.3 |
| Solyc04g051700.2 | Solyc04g051700.3 |
| Solyc04g051720.2 | Solyc04g051720.3 |
| Solyc04g051780.2 | Solyc04g051780.3 |
| Solyc04g051800.2 | Solyc04g051800.3 |
| Solyc04g051860.2 | Solyc04g051860.3 |
| Solyc04g053020.1 | Solyc04g053020.2 |
| Solyc04g053040.2 | Solyc04g053040.3 |
| Solyc04g053055.1 | Solyc04g150140.1 |

| | |
|---|---|
| Solyc04g053070.1 | Solyc04g053070.2 |
| Solyc04g053090.1 | Solyc04g053090.2 |
| Solyc04g053092.1 | Solyc04g150141.1 |
| Solyc04g053096.1 | Solyc04g150142.1 |
| Solyc04g053098.1 | Solyc04g150143.1 |
| Solyc04g053120.2 | Solyc04g053120.3 |
| Solyc04g054145.1 | Solyc04g150144.1 |
| Solyc04g054150.1 | Solyc04g054150.2 |
| Solyc04g054157.1 | Solyc04g150145.1 |
| Solyc04g054190.2 | Solyc04g054190.3 |
| Solyc04g054200.2 | Solyc04g054200.3 |
| Solyc04g054205.1 | Solyc04g150146.1 |
| Solyc04g054220.2 | Solyc04g054220.3 |
| Solyc04g054225.1 | Solyc04g150147.1 |
| Solyc04g054280.1 | Solyc04g054280.2 |
| Solyc04g054310.2 | Solyc04g054310.3 |
| Solyc04g054330.2 | Solyc04g054330.3 |
| Solyc04g054400.2 | Solyc04g054400.3 |
| Solyc04g054410.2 | Solyc04g054410.3 |
| Solyc04g054430.1 | Solyc04g054430.2 |
| Solyc04g054470.2 | Solyc04g054470.3 |
| Solyc04g054480.2 | Solyc04g054480.3 |
| Solyc04g054490.2 | Solyc04g054490.3 |
| Solyc04g054500.2 | Solyc04g054500.3 |
| Solyc04g054520.2 | Solyc04g054520.3 |
| Solyc04g054690.2 | Solyc04g054690.3 |
| Solyc04g054700.2 | Solyc04g054700.3 |
| Solyc04g054710.2 | Solyc04g054710.3 |
| Solyc04g054730.2 | Solyc04g054730.3 |
| Solyc04g054740.2 | Solyc04g054740.3 |
| Solyc04g054745.1 | Solyc04g150150.1 |
| Solyc04g054760.2 | Solyc04g054760.3 |
| Solyc04g054780.2 | Solyc04g054780.3 |
| Solyc04g054790.1 | Solyc04g054790.2 |
| Solyc04g054800.2 | Solyc04g054800.3 |
| Solyc04g054810.2 | Solyc04g054810.3 |
| Solyc04g054820.2 | Solyc04g054820.3 |
| Solyc04g054880.2 | Solyc04g054880.3 |
| Solyc04g054890.2 | Solyc04g054890.3 |
| Solyc04g054900.2 | Solyc04g054900.3 |
| Solyc04g054930.2 | Solyc04g054930.3 |
| Solyc04g054950.2 | Solyc04g054950.3 |
| Solyc04g054980.2 | Solyc04g054980.3 |

| | |
|---|---|
| Solyc04g055000.2 | Solyc04g055000.3 |
| Solyc04g055010.2 | Solyc04g055010.3 |
| Solyc04g055020.1 | Solyc04g055020.2 |
| Solyc04g055030.1 | Solyc04g055030.2 |
| Solyc04g055080.2 | Solyc04g055080.3 |
| Solyc04g055110.2 | Solyc04g055110.3 |
| Solyc04g055120.2 | Solyc04g055120.3 |
| Solyc04g055160.2 | Solyc04g055160.3 |
| Solyc04g055170.2 | Solyc04g055170.3 |
| Solyc04g055180.2 | Solyc04g055180.3 |
| Solyc04g055200.2 | Solyc04g055200.3 |
| Solyc04g055220.1 | Solyc04g055220.2 |
| Solyc04g055230.1 | Solyc04g055230.2 |
| Solyc04g056260.1 | Solyc04g056260.2 |
| Solyc04g056290.2 | Solyc04g056290.3 |
| Solyc04g056310.2 | Solyc04g056310.3 |
| Solyc04g056370.2 | Solyc04g056370.3 |
| Solyc04g056380.2 | Solyc04g056380.3 |
| Solyc04g056390.2 | Solyc04g056390.3 |
| Solyc04g056410.2 | Solyc04g056410.3 |
| Solyc04g056460.1 | Solyc04g056460.2 |
| Solyc04g056480.1 | Solyc04g056480.2 |
| Solyc04g056491.1 | Solyc04g150151.1 |
| Solyc04g056492.1 | Solyc04g150152.1 |
| Solyc04g056494.1 | Solyc04g150153.1 |
| Solyc04g056496.1 | Solyc04g150154.1 |
| Solyc04g056498.1 | Solyc04g150155.1 |
| Solyc04g056505.1 | Solyc04g150156.1 |
| Solyc04g056530.1 | Solyc04g056530.2 |
| Solyc04g056540.2 | Solyc04g056540.3 |
| Solyc04g056570.2 | Solyc04g056570.3 |
| Solyc04g056580.2 | Solyc04g056580.3 |
| Solyc04g056590.2 | Solyc04g056590.3 |
| Solyc04g056620.1 | Solyc04g056620.2 |
| Solyc04g056630.2 | Solyc04g056630.3 |
| Solyc04g083140.1 | Solyc04g083140.2 |
| Solyc04g056680.1 | Solyc04g056680.2 |
| Solyc04g056690.1 | Solyc04g056690.2 |
| Solyc04g056743.1 | Solyc04g150157.1 |
| Solyc04g083150.1 | Solyc04g083150.2 |
| Solyc04g057880.2 | Solyc04g057880.3 |
| Solyc04g057890.2 | Solyc04g057890.3 |
| Solyc04g057950.1 | Solyc04g057950.2 |

| | |
|---|---|
| Solyc04g057980.2 | Solyc04g057980.3 |
| Solyc04g057993.1 | Solyc04g150158.1 |
| Solyc04g057997.1 | Solyc04g150159.1 |
| Solyc04g058000.1 | Solyc04g058000.2 |
| Solyc04g058070.2 | Solyc04g058070.3 |
| Solyc04g058080.2 | Solyc04g058080.3 |
| Solyc04g058090.2 | Solyc04g058090.3 |
| Solyc04g058100.2 | Solyc04g058100.3 |
| Solyc04g058110.2 | Solyc04g058110.3 |
| Solyc04g058130.2 | Solyc04g058130.3 |
| Solyc04g058140.2 | Solyc04g058140.3 |
| Solyc04g058150.2 | Solyc04g058150.3 |
| Solyc04g058160.1 | Solyc04g058160.2 |
| Solyc04g058190.2 | Solyc04g058190.3 |
| Solyc04g058200.2 | Solyc04g058200.3 |
| Solyc04g063210.2 | Solyc04g063210.3 |
| Solyc04g063230.2 | Solyc04g063230.3 |
| Solyc04g063240.2 | Solyc04g063240.3 |
| Solyc04g063270.1 | Solyc04g063270.2 |
| Solyc04g063280.2 | Solyc04g063280.3 |
| Solyc04g063290.2 | Solyc04g063290.3 |
| Solyc04g063350.2 | Solyc04g063350.3 |
| Solyc04g063360.2 | Solyc04g063360.3 |
| Solyc04g063380.2 | Solyc04g063380.3 |
| Solyc04g064460.1 | Solyc04g064460.2 |
| Solyc04g064470.1 | Solyc04g064470.2 |
| Solyc04g064480.2 | Solyc04g064480.3 |
| Solyc04g064490.2 | Solyc04g064490.3 |
| Solyc04g064510.2 | Solyc04g064510.3 |
| Solyc04g064570.2 | Solyc04g064570.3 |
| Solyc04g064610.2 | Solyc04g064610.3 |
| Solyc04g064630.2 | Solyc04g064630.3 |
| Solyc04g064670.2 | Solyc04g064670.3 |
| Solyc04g064710.2 | Solyc04g064710.3 |
| Solyc04g064720.2 | Solyc04g064720.3 |
| Solyc04g064730.1 | Solyc04g064730.2 |
| Solyc04g064750.1 | Solyc04g064750.2 |
| Solyc04g064760.2 | Solyc04g064760.3 |
| Solyc04g064790.2 | Solyc04g064790.3 |
| Solyc04g064820.2 | Solyc04g064820.3 |
| Solyc04g064850.2 | Solyc04g064850.3 |
| Solyc04g064870.2 | Solyc04g064870.3 |
| Solyc04g064890.2 | Solyc04g064890.3 |

| | |
|---|---|
| Solyc04g064940.2 | Solyc04g064940.3 |
| Solyc04g071075.1 | Solyc04g150160.1 |
| Solyc04g071080.1 | Solyc04g071080.2 |
| Solyc04g071085.1 | Solyc04g150161.1 |
| Solyc04g071120.2 | Solyc04g071120.3 |
| Solyc04g071140.2 | Solyc04g071140.3 |
| Solyc04g071150.2 | Solyc04g071150.3 |
| Solyc04g071160.2 | Solyc04g071160.3 |
| Solyc04g071165.1 | Solyc04g150162.1 |
| Solyc04g071260.2 | Solyc04g071260.3 |
| Solyc04g071285.1 | Solyc04g150163.1 |
| Solyc04g071350.2 | Solyc04g071350.3 |
| Solyc04g071410.1 | Solyc04g071410.2 |
| Solyc04g071430.1 | Solyc04g071430.2 |
| Solyc04g071440.2 | Solyc04g071440.3 |
| Solyc04g071470.2 | Solyc04g071470.3 |
| Solyc04g071490.2 | Solyc04g071490.3 |
| Solyc04g071510.2 | Solyc04g071510.3 |
| Solyc04g071520.2 | Solyc04g071520.3 |
| Solyc04g071530.1 | Solyc04g071530.2 |
| Solyc04g071600.2 | Solyc04g071600.3 |
| Solyc04g071680.1 | Solyc04g071680.2 |
| Solyc04g071700.2 | Solyc04g071700.3 |
| Solyc04g071730.2 | Solyc04g071730.3 |
| Solyc04g071760.2 | Solyc04g071760.3 |
| Solyc04g071770.2 | Solyc04g071770.3 |
| Solyc04g071780.2 | Solyc04g071780.3 |
| Solyc04g071800.2 | Solyc04g071800.3 |
| Solyc04g071807.1 | Solyc04g150164.1 |
| Solyc04g071860.2 | Solyc04g071860.3 |
| Solyc04g071880.2 | Solyc04g071880.3 |
| Solyc04g071890.2 | Solyc04g071890.3 |
| Solyc04g071920.1 | Solyc04g071920.2 |
| Solyc04g071940.2 | Solyc04g071940.3 |
| Solyc04g071950.1 | Solyc04g071950.2 |
| Solyc04g071960.2 | Solyc04g071960.3 |
| Solyc04g071980.2 | Solyc04g071980.3 |
| Solyc04g071990.2 | Solyc04g071990.3 |
| Solyc04g072010.2 | Solyc04g072010.3 |
| Solyc04g072020.2 | Solyc04g072020.3 |
| Solyc04g072070.2 | Solyc04g072070.3 |
| Solyc04g072130.1 | Solyc04g072130.2 |
| Solyc04g072145.1 | Solyc04g150166.1 |

| | |
|---|---|
| Solyc04g072160.2 | Solyc04g072160.3 |
| Solyc04g072220.2 | Solyc04g072220.3 |
| Solyc04g072240.2 | Solyc04g072240.3 |
| Solyc04g072280.2 | Solyc04g072280.3 |
| Solyc04g072290.2 | Solyc04g072290.3 |
| Solyc04g072310.2 | Solyc04g072310.3 |
| Solyc04g072360.1 | Solyc04g072360.2 |
| Solyc04g072380.2 | Solyc04g072380.3 |
| Solyc04g072390.2 | Solyc04g072390.3 |
| Solyc04g072410.2 | Solyc04g072410.3 |
| Solyc04g072425.1 | Solyc04g150167.1 |
| Solyc04g072450.2 | Solyc04g072450.3 |
| Solyc04g072460.2 | Solyc04g072460.3 |
| Solyc04g072480.2 | Solyc04g072480.3 |
| Solyc04g072510.2 | Solyc04g072510.3 |
| Solyc04g072540.2 | Solyc04g072540.3 |
| Solyc04g072560.2 | Solyc04g072560.3 |
| Solyc04g072570.2 | Solyc04g072570.3 |
| Solyc04g072640.2 | Solyc04g072640.3 |
| Solyc04g072680.1 | Solyc04g072680.2 |
| Solyc04g072690.2 | Solyc04g072690.3 |
| Solyc04g072740.2 | Solyc04g072740.3 |
| Solyc04g072750.2 | Solyc04g072750.3 |
| Solyc04g072760.2 | Solyc04g072760.3 |
| Solyc04g072790.1 | Solyc04g072790.2 |
| Solyc04g072840.2 | Solyc04g072840.3 |
| Solyc04g072860.2 | Solyc04g072860.3 |
| Solyc04g072870.1 | Solyc04g072870.2 |
| Solyc04g072880.2 | Solyc04g072880.3 |
| Solyc04g072910.2 | Solyc04g072910.3 |
| Solyc04g073950.1 | Solyc04g073950.2 |
| Solyc04g073960.2 | Solyc04g073960.3 |
| Solyc04g073980.2 | Solyc04g073980.3 |
| Solyc04g073990.2 | Solyc04g073990.3 |
| Solyc04g074000.2 | Solyc04g074000.3 |
| Solyc04g074030.2 | Solyc04g074030.3 |
| Solyc04g074040.2 | Solyc04g074040.3 |
| Solyc04g074070.1 | Solyc04g074070.2 |
| Solyc04g074080.2 | Solyc04g074080.3 |
| Solyc04g074130.2 | Solyc04g074130.3 |
| Solyc04g074230.2 | Solyc04g074230.3 |
| Solyc04g074240.2 | Solyc04g074240.3 |
| Solyc04g074270.2 | Solyc04g074270.3 |

| | |
|---|---|
| Solyc04g074290.2 | Solyc04g074290.3 |
| Solyc04g074300.2 | Solyc04g074300.3 |
| Solyc04g074330.2 | Solyc04g074330.3 |
| Solyc04g074340.2 | Solyc04g074340.3 |
| Solyc04g074350.2 | Solyc04g074350.3 |
| Solyc04g074370.1 | Solyc04g074370.2 |
| Solyc04g074390.2 | Solyc04g074390.3 |
| Solyc04g074410.1 | Solyc04g074410.2 |
| Solyc04g074430.1 | Solyc04g074430.2 |
| Solyc04g074480.2 | Solyc04g074480.3 |
| Solyc04g074490.2 | Solyc04g074490.3 |
| Solyc04g074500.2 | Solyc04g074500.3 |
| Solyc04g074510.2 | Solyc04g074510.3 |
| Solyc04g074530.2 | Solyc04g074530.3 |
| Solyc04g074690.2 | Solyc04g074690.3 |
| Solyc04g074710.2 | Solyc04g074710.3 |
| Solyc04g074750.2 | Solyc04g074750.3 |
| Solyc04g074800.2 | Solyc04g074800.3 |
| Solyc04g074840.2 | Solyc04g074840.3 |
| Solyc04g074870.2 | Solyc04g074870.3 |
| Solyc04g074900.2 | Solyc04g074900.3 |
| Solyc04g074910.2 | Solyc04g074910.3 |
| Solyc04g074950.2 | Solyc04g074950.3 |
| Solyc04g074980.2 | Solyc04g074980.3 |
| Solyc04g074990.2 | Solyc04g074990.3 |
| Solyc04g076020.2 | Solyc04g076020.3 |
| Solyc04g076040.2 | Solyc04g076040.3 |
| Solyc04g076060.2 | Solyc04g076060.3 |
| Solyc04g076080.2 | Solyc04g076080.3 |
| Solyc04g076090.2 | Solyc04g076090.3 |
| Solyc04g076120.2 | Solyc04g076120.3 |
| Solyc04g076130.2 | Solyc04g076130.3 |
| Solyc04g076140.2 | Solyc04g076140.3 |
| Solyc04g076200.2 | Solyc04g076200.3 |
| Solyc04g076210.2 | Solyc04g076210.3 |
| Solyc04g076220.2 | Solyc04g076220.3 |
| Solyc04g076230.2 | Solyc04g076230.3 |
| Solyc04g076250.2 | Solyc04g076250.3 |
| Solyc04g076290.2 | Solyc04g076290.3 |
| Solyc04g076390.2 | Solyc04g076390.3 |
| Solyc04g076400.2 | Solyc04g076400.3 |
| Solyc04g076430.2 | Solyc04g076430.3 |
| Solyc04g076480.2 | Solyc04g076480.3 |

| | |
|---|---|
| Solyc04g076510.2 | Solyc04g076510.3 |
| Solyc04g076527.1 | Solyc04g150168.1 |
| Solyc04g076530.2 | Solyc04g076530.3 |
| Solyc04g076560.2 | Solyc04g076560.3 |
| Solyc04g076590.1 | Solyc04g076590.2 |
| Solyc04g076630.2 | Solyc04g076630.3 |
| Solyc04g076650.1 | Solyc04g076650.2 |
| Solyc04g076660.2 | Solyc04g076660.3 |
| Solyc04g076670.2 | Solyc04g076670.3 |
| Solyc04g076740.2 | Solyc04g076740.3 |
| Solyc04g076750.1 | Solyc04g076750.2 |
| Solyc04g076760.1 | Solyc04g076760.2 |
| Solyc04g076770.2 | Solyc04g076770.3 |
| Solyc04g076790.2 | Solyc04g076790.3 |
| Solyc04g076800.2 | Solyc04g076800.3 |
| Solyc04g076850.2 | Solyc04g076850.3 |
| Solyc04g076860.2 | Solyc04g076860.3 |
| Solyc04g076880.2 | Solyc04g076880.3 |
| Solyc04g076900.1 | Solyc04g076900.2 |
| Solyc04g076910.2 | Solyc04g076910.3 |
| Solyc04g076920.2 | Solyc04g076920.3 |
| Solyc04g076940.2 | Solyc04g076940.3 |
| Solyc04g076950.2 | Solyc04g076950.3 |
| Solyc04g076970.2 | Solyc04g076970.3 |
| Solyc04g076980.2 | Solyc04g076980.3 |
| Solyc04g076990.2 | Solyc04g076990.3 |
| Solyc04g077000.2 | Solyc04g077000.3 |
| Solyc04g077020.2 | Solyc04g077020.3 |
| Solyc04g077030.2 | Solyc04g077030.3 |
| Solyc04g077040.2 | Solyc04g077040.3 |
| Solyc04g077050.2 | Solyc04g077050.3 |
| Solyc04g077060.2 | Solyc04g077060.3 |
| Solyc04g077110.2 | Solyc04g077110.3 |
| Solyc04g077140.2 | Solyc04g077140.3 |
| Solyc04g077150.2 | Solyc04g077150.3 |
| Solyc04g077170.2 | Solyc04g077170.3 |
| Solyc04g077210.2 | Solyc04g077210.3 |
| Solyc04g077220.2 | Solyc04g077220.3 |
| Solyc04g077245.1 | Solyc04g150169.1 |
| Solyc04g077260.2 | Solyc04g077260.3 |
| Solyc04g077280.2 | Solyc04g077280.3 |
| Solyc04g077300.2 | Solyc04g077300.3 |
| Solyc04g077310.2 | Solyc04g077310.3 |

| | |
|---|---|
| Solyc04g077320.1 | Solyc04g077320.2 |
| Solyc04g077340.2 | Solyc04g077340.3 |
| Solyc04g077360.2 | Solyc04g077360.3 |
| Solyc04g077370.1 | Solyc04g077370.2 |
| Solyc04g077460.2 | Solyc04g077460.3 |
| Solyc04g077470.2 | Solyc04g077470.3 |
| Solyc04g077480.2 | Solyc04g077480.3 |
| Solyc04g077490.2 | Solyc04g077490.3 |
| Solyc04g077520.2 | Solyc04g077520.3 |
| Solyc04g077580.2 | Solyc04g077580.3 |
| Solyc04g077600.2 | Solyc04g077600.3 |
| Solyc04g077630.2 | Solyc04g077630.3 |
| Solyc04g077650.2 | Solyc04g077650.3 |
| Solyc04g077660.2 | Solyc04g077660.3 |
| Solyc04g077670.2 | Solyc04g077670.3 |
| Solyc04g077680.2 | Solyc04g077680.3 |
| Solyc04g077780.2 | Solyc04g077780.3 |
| Solyc04g077790.2 | Solyc04g077790.3 |
| Solyc04g077850.2 | Solyc04g077850.3 |
| Solyc04g077870.2 | Solyc04g077870.3 |
| Solyc04g077900.1 | Solyc04g077900.2 |
| Solyc04g077920.2 | Solyc04g077920.3 |
| Solyc04g077990.2 | Solyc04g077990.3 |
| Solyc04g078000.2 | Solyc04g078000.3 |
| Solyc04g078020.1 | Solyc04g078020.2 |
| Solyc04g078040.2 | Solyc04g078040.3 |
| Solyc04g078050.2 | Solyc04g078050.3 |
| Solyc04g078090.2 | Solyc04g078090.3 |
| Solyc04g078120.2 | Solyc04g078120.3 |
| Solyc04g078140.2 | Solyc04g078140.3 |
| Solyc04g078170.1 | Solyc04g078170.2 |
| Solyc04g078280.2 | Solyc04g078280.3 |
| Solyc04g078270.2 | Solyc04g078270.3 |
| Solyc04g078300.2 | Solyc04g078300.3 |
| Solyc04g078310.2 | Solyc04g078310.3 |
| Solyc04g078340.2 | Solyc04g078340.3 |
| Solyc04g078370.2 | Solyc04g078370.3 |
| Solyc04g078400.2 | Solyc04g078400.3 |
| Solyc04g078460.2 | Solyc04g078460.3 |
| Solyc04g078470.2 | Solyc04g078470.3 |
| Solyc04g078480.2 | Solyc04g078480.3 |
| Solyc04g078520.2 | Solyc04g078520.3 |
| Solyc04g078550.2 | Solyc04g078550.3 |

| | |
|---|---|
| Solyc04g078565.1 | Solyc04g150170.1 |
| Solyc04g078590.2 | Solyc04g078590.3 |
| Solyc04g078620.2 | Solyc04g078620.3 |
| Solyc04g078630.2 | Solyc04g078630.3 |
| Solyc04g078660.1 | Solyc04g078660.2 |
| Solyc04g078670.2 | Solyc04g078670.3 |
| Solyc04g078690.2 | Solyc04g078690.3 |
| Solyc04g078700.2 | Solyc04g078700.3 |
| Solyc04g078750.2 | Solyc04g078750.3 |
| Solyc04g078780.2 | Solyc04g078780.3 |
| Solyc04g078790.2 | Solyc04g078790.3 |
| Solyc04g078800.2 | Solyc04g078800.3 |
| Solyc04g078830.2 | Solyc04g078830.3 |
| Solyc04g078840.2 | Solyc04g078840.3 |
| Solyc04g078900.2 | Solyc04g078900.3 |
| Solyc04g078910.2 | Solyc04g078910.3 |
| Solyc04g078920.2 | Solyc04g078920.3 |
| Solyc04g078940.2 | Solyc04g078940.3 |
| Solyc04g079060.2 | Solyc04g079060.3 |
| Solyc04g079100.2 | Solyc04g079100.3 |
| Solyc04g079120.2 | Solyc04g079120.3 |
| Solyc04g079180.2 | Solyc04g079180.3 |
| Solyc04g079200.2 | Solyc04g079200.3 |
| Solyc04g079210.2 | Solyc04g079210.3 |
| Solyc04g079240.2 | Solyc04g079240.3 |
| Solyc04g079250.2 | Solyc04g079250.3 |
| Solyc04g079260.2 | Solyc04g079260.3 |
| Solyc04g079290.2 | Solyc04g079290.3 |
| Solyc04g079300.2 | Solyc04g079300.3 |
| Solyc04g079330.1 | Solyc04g079330.2 |
| Solyc04g079340.2 | Solyc04g079340.3 |
| Solyc04g079390.2 | Solyc04g079390.3 |
| Solyc04g079400.2 | Solyc04g079400.3 |
| Solyc04g079410.2 | Solyc04g079410.3 |
| Solyc04g079420.2 | Solyc04g079420.3 |
| Solyc04g079440.2 | Solyc04g079440.3 |
| Solyc04g079470.2 | Solyc04g079470.3 |
| Solyc04g079500.1 | Solyc04g079500.2 |
| Solyc04g079505.1 | Solyc04g150171.1 |
| Solyc04g079510.1 | Solyc04g079510.2 |
| Solyc04g079520.2 | Solyc04g079520.3 |
| Solyc04g079530.2 | Solyc04g079530.3 |
| Solyc04g079560.2 | Solyc04g079560.3 |

| | |
|---|---|
| Solyc04g079590.2 | Solyc04g079590.3 |
| Solyc04g079600.2 | Solyc04g079600.3 |
| Solyc04g079640.2 | Solyc04g079640.3 |
| Solyc04g079690.2 | Solyc04g079690.3 |
| Solyc04g079700.2 | Solyc04g079700.3 |
| Solyc04g079770.2 | Solyc04g079770.3 |
| Solyc04g079790.2 | Solyc04g079790.3 |
| Solyc04g079820.2 | Solyc04g079820.3 |
| Solyc04g079850.1 | Solyc04g079850.2 |
| Solyc04g079870.1 | Solyc04g079870.2 |
| Solyc04g079900.2 | Solyc04g079900.3 |
| Solyc04g079930.2 | Solyc04g079930.3 |
| Solyc04g079940.2 | Solyc04g079940.3 |
| Solyc04g079970.2 | Solyc04g079970.3 |
| Solyc04g079980.2 | Solyc04g079980.3 |
| Solyc04g080030.2 | Solyc04g080030.3 |
| Solyc04g080040.2 | Solyc04g080040.3 |
| Solyc04g080060.2 | Solyc04g080060.3 |
| Solyc04g080080.2 | Solyc04g080080.3 |
| Solyc04g080100.2 | Solyc04g080100.3 |
| Solyc04g080130.2 | Solyc04g080130.3 |
| Solyc04g080160.2 | Solyc04g080160.3 |
| Solyc04g080170.2 | Solyc04g080170.3 |
| Solyc04g080190.2 | Solyc04g080190.3 |
| Solyc04g080220.2 | Solyc04g080220.3 |
| Solyc04g080230.2 | Solyc04g080230.3 |
| Solyc04g080250.2 | Solyc04g080250.3 |
| Solyc04g080280.2 | Solyc04g080280.3 |
| Solyc04g080290.2 | Solyc04g080290.3 |
| Solyc04g080330.2 | Solyc04g080330.3 |
| Solyc04g080340.2 | Solyc04g080340.3 |
| Solyc04g080370.2 | Solyc04g080370.3 |
| Solyc04g080380.2 | Solyc04g080380.3 |
| Solyc04g080430.2 | Solyc04g080430.3 |
| Solyc04g080435.1 | Solyc04g150172.1 |
| Solyc04g080460.2 | Solyc04g080460.3 |
| Solyc04g080500.2 | Solyc04g080500.3 |
| Solyc04g080510.2 | Solyc04g080510.3 |
| Solyc04g080530.2 | Solyc04g080530.3 |
| Solyc04g080570.2 | Solyc04g080570.3 |
| Solyc04g080580.2 | Solyc04g080580.3 |
| Solyc04g080600.2 | Solyc04g080600.3 |
| Solyc04g080610.2 | Solyc04g080610.3 |

| | |
|---|---|
| Solyc04g080620.2 | Solyc04g080620.3 |
| Solyc04g080670.2 | Solyc04g080670.3 |
| Solyc04g080730.2 | Solyc04g080730.3 |
| Solyc04g080760.2 | Solyc04g080760.3 |
| Solyc04g080770.2 | Solyc04g080770.3 |
| Solyc04g080790.2 | Solyc04g080790.3 |
| Solyc04g080850.2 | Solyc04g080850.3 |
| Solyc04g080900.2 | Solyc04g080900.3 |
| Solyc04g080990.1 | Solyc04g080990.2 |
| Solyc04g081000.2 | Solyc04g081000.3 |
| Solyc04g081010.1 | Solyc04g081010.2 |
| Solyc04g081020.2 | Solyc04g081020.3 |
| Solyc04g081030.2 | Solyc04g081030.3 |
| Solyc04g081050.1 | Solyc04g081050.2 |
| Solyc04g081060.2 | Solyc04g081060.3 |
| Solyc04g081070.2 | Solyc04g081070.3 |
| Solyc04g081090.2 | Solyc04g081090.3 |
| Solyc04g081100.2 | Solyc04g081100.3 |
| Solyc04g081160.2 | Solyc04g081160.3 |
| Solyc04g081170.2 | Solyc04g081170.3 |
| Solyc04g081190.2 | Solyc04g081190.3 |
| Solyc04g081200.2 | Solyc04g081200.3 |
| Solyc04g081220.2 | Solyc04g081220.3 |
| Solyc04g081230.2 | Solyc04g081230.3 |
| Solyc04g081260.2 | Solyc04g081260.3 |
| Solyc04g081280.2 | Solyc04g081280.3 |
| Solyc04g081290.2 | Solyc04g081290.3 |
| Solyc04g081310.2 | Solyc04g081310.3 |
| Solyc04g081340.2 | Solyc04g081340.3 |
| Solyc04g081350.2 | Solyc04g081350.3 |
| Solyc04g081360.2 | Solyc04g081360.3 |
| Solyc04g081400.2 | Solyc04g081400.3 |
| Solyc04g081410.2 | Solyc04g081410.3 |
| Solyc04g081430.2 | Solyc04g081430.3 |
| Solyc04g081440.2 | Solyc04g081440.3 |
| Solyc04g081490.2 | Solyc04g081490.3 |
| Solyc04g081500.2 | Solyc04g081500.3 |
| Solyc04g081550.2 | Solyc04g081550.3 |
| Solyc04g081560.2 | Solyc04g081560.3 |
| Solyc04g081570.2 | Solyc04g081570.3 |
| Solyc04g081630.1 | Solyc04g081630.2 |
| Solyc04g081650.2 | Solyc04g081650.3 |
| Solyc04g081690.1 | Solyc04g081690.2 |

| | |
|---|---|
| Solyc04g081700.2 | Solyc04g081700.3 |
| Solyc04g081710.1 | Solyc04g081710.2 |
| Solyc04g081720.1 | Solyc04g081720.2 |
| Solyc04g081740.2 | Solyc04g081740.3 |
| Solyc04g081755.1 | Solyc04g081755.2 |
| Solyc04g081780.2 | Solyc04g081780.3 |
| Solyc04g081790.2 | Solyc04g081790.3 |
| Solyc04g081820.2 | Solyc04g081820.3 |
| Solyc04g081860.2 | Solyc04g081860.3 |
| Solyc04g081920.2 | Solyc04g081920.3 |
| Solyc04g081930.2 | Solyc04g081930.3 |
| Solyc04g081940.2 | Solyc04g081940.3 |
| Solyc04g081970.2 | Solyc04g081970.3 |
| Solyc04g081990.2 | Solyc04g081990.3 |
| Solyc04g082020.2 | Solyc04g082020.3 |
| Solyc04g082040.2 | Solyc04g082040.3 |
| Solyc04g082050.2 | Solyc04g082050.3 |
| Solyc04g082090.2 | Solyc04g082090.3 |
| Solyc04g082120.2 | Solyc04g082120.3 |
| Solyc04g082140.2 | Solyc04g082140.3 |
| Solyc04g082250.2 | Solyc04g082250.3 |
| Solyc04g082270.2 | Solyc04g082270.3 |
| Solyc04g082280.2 | Solyc04g082280.3 |
| Solyc04g082320.2 | Solyc04g082320.3 |
| Solyc04g082330.2 | Solyc04g082330.3 |
| Solyc04g082340.2 | Solyc04g082340.3 |
| Solyc04g082350.1 | Solyc04g082350.2 |
| Solyc04g082360.1 | Solyc04g082360.2 |
| Solyc04g082390.2 | Solyc04g082390.3 |
| Solyc04g082400.2 | Solyc04g082400.3 |
| Solyc04g082420.2 | Solyc04g082420.3 |
| Solyc04g082430.2 | Solyc04g082430.3 |
| Solyc04g082450.2 | Solyc04g082450.3 |
| Solyc04g082480.2 | Solyc04g082480.3 |
| Solyc04g082490.2 | Solyc04g082490.3 |
| Solyc04g082510.2 | Solyc04g082510.3 |
| Solyc04g082530.2 | Solyc04g082530.3 |
| Solyc04g082540.1 | Solyc04g082540.2 |
| Solyc04g082550.1 | Solyc04g082550.2 |
| Solyc04g082570.2 | Solyc04g082570.3 |
| Solyc04g082580.2 | Solyc04g082580.3 |
| Solyc04g082600.2 | Solyc04g082600.3 |
| Solyc04g082610.2 | Solyc04g082610.3 |

| | |
|---|---|
| Solyc04g082620.2 | Solyc04g082620.3 |
| Solyc04g082630.2 | Solyc04g082630.3 |
| Solyc04g082650.2 | Solyc04g082650.3 |
| Solyc04g082680.2 | Solyc04g082680.3 |
| Solyc04g082700.2 | Solyc04g082700.3 |
| Solyc04g082720.2 | Solyc04g082720.3 |
| Solyc04g082730.1 | Solyc04g082730.2 |
| Solyc04g082750.2 | Solyc04g082750.3 |
| Solyc04g082760.2 | Solyc04g082760.3 |
| Solyc04g082780.2 | Solyc04g082780.3 |
| Solyc04g082830.2 | Solyc04g082830.3 |
| Solyc04g082845.1 | Solyc04g082845.2 |
| Solyc04g082890.2 | Solyc04g082890.3 |
| Solyc04g082940.2 | Solyc04g082940.3 |
| Solyc04g082980.1 | Solyc04g082980.2 |
| Solyc04g082990.2 | Solyc04g082990.3 |
| Solyc05g005010.2 | Solyc05g005010.3 |
| Solyc05g005050.2 | Solyc05g005050.3 |
| Solyc05g005060.2 | Solyc05g005060.3 |
| Solyc05g005080.2 | Solyc05g005080.3 |
| Solyc05g005160.2 | Solyc05g005160.3 |
| Solyc05g005180.2 | Solyc05g005180.3 |
| Solyc05g005190.2 | Solyc05g005190.3 |
| Solyc05g005200.2 | Solyc05g005200.3 |
| Solyc05g005210.2 | Solyc05g005210.3 |
| Solyc05g005230.2 | Solyc05g005230.3 |
| Solyc05g005240.1 | Solyc05g005240.2 |
| Solyc05g005250.2 | Solyc05g005250.3 |
| Solyc05g005330.2 | Solyc05g005330.3 |
| Solyc05g005340.2 | Solyc05g005340.3 |
| Solyc05g005385.1 | Solyc05g150100.1 |
| Solyc05g005400.1 | Solyc05g005400.2 |
| Solyc05g005405.1 | Solyc05g150101.1 |
| Solyc05g005430.1 | Solyc05g005430.2 |
| Solyc05g005450.1 | Solyc05g005450.2 |
| Solyc05g005460.2 | Solyc05g005460.3 |
| Solyc05g005510.2 | Solyc05g005510.3 |
| Solyc05g005530.2 | Solyc05g005530.3 |
| Solyc05g005580.2 | Solyc05g005580.3 |
| Solyc05g005680.2 | Solyc05g005680.3 |
| Solyc05g005740.2 | Solyc05g005740.3 |
| Solyc05g005750.2 | Solyc05g005750.3 |
| Solyc05g005770.2 | Solyc05g005770.3 |

| | |
|---|---|
| Solyc05g005780.2 | Solyc05g005780.3 |
| Solyc05g005830.2 | Solyc05g005830.3 |
| Solyc05g005870.2 | Solyc05g005870.3 |
| Solyc05g005940.2 | Solyc05g005940.3 |
| Solyc05g005950.2 | Solyc05g005950.3 |
| Solyc05g005960.2 | Solyc05g005960.3 |
| Solyc05g005970.2 | Solyc05g005970.3 |
| Solyc05g005990.2 | Solyc05g005990.3 |
| Solyc05g006000.2 | Solyc05g006000.3 |
| Solyc05g006020.2 | Solyc05g006020.3 |
| Solyc05g006030.2 | Solyc05g006030.3 |
| Solyc05g006050.2 | Solyc05g006050.3 |
| Solyc05g006120.2 | Solyc05g006120.3 |
| Solyc05g006180.2 | Solyc05g006180.3 |
| Solyc05g006190.1 | Solyc05g006190.2 |
| Solyc05g006210.2 | Solyc05g006210.3 |
| Solyc05g006220.2 | Solyc05g006220.3 |
| Solyc05g006230.1 | Solyc05g006230.2 |
| Solyc05g006290.1 | Solyc05g006290.2 |
| Solyc05g006330.1 | Solyc05g006330.2 |
| Solyc05g006340.2 | Solyc05g006340.3 |
| Solyc05g006343.1 | Solyc05g150102.1 |
| Solyc05g006347.1 | Solyc05g150103.1 |
| Solyc05g006360.1 | Solyc05g006360.2 |
| Solyc05g006370.1 | Solyc05g006370.2 |
| Solyc05g006410.2 | Solyc05g006410.3 |
| Solyc05g006420.2 | Solyc05g006420.3 |
| Solyc05g006510.1 | Solyc05g006510.2 |
| Solyc05g006540.2 | Solyc05g006540.3 |
| Solyc05g006590.2 | Solyc05g006590.3 |
| Solyc05g006620.2 | Solyc05g006620.3 |
| Solyc05g006650.2 | Solyc05g006650.3 |
| Solyc05g006670.1 | Solyc05g006670.2 |
| Solyc05g006690.2 | Solyc05g006690.3 |
| Solyc05g006710.2 | Solyc05g006710.3 |
| Solyc05g006750.2 | Solyc05g006750.3 |
| Solyc05g006780.2 | Solyc05g006780.3 |
| Solyc05g006790.1 | Solyc05g006790.2 |
| Solyc05g006800.1 | Solyc05g006800.2 |
| Solyc05g006830.2 | Solyc05g006830.3 |
| Solyc05g006833.1 | Solyc05g150104.1 |
| Solyc05g006837.1 | Solyc05g150105.1 |
| Solyc05g006870.2 | Solyc05g006870.3 |

| | |
|---|---|
| Solyc05g006930.2 | Solyc05g006930.3 |
| Solyc05g006940.2 | Solyc05g006940.3 |
| Solyc05g006980.2 | Solyc05g006980.3 |
| Solyc05g006990.2 | Solyc05g006990.3 |
| Solyc05g007040.1 | Solyc05g007040.2 |
| Solyc05g007100.2 | Solyc05g007100.3 |
| Solyc05g007140.2 | Solyc05g007140.3 |
| Solyc05g007160.2 | Solyc05g007160.3 |
| Solyc05g007170.2 | Solyc05g007170.3 |
| Solyc05g007180.2 | Solyc05g007180.3 |
| Solyc05g007250.2 | Solyc05g007250.3 |
| Solyc05g007260.2 | Solyc05g007260.3 |
| Solyc05g007280.2 | Solyc05g007280.3 |
| Solyc05g007300.2 | Solyc05g007300.3 |
| Solyc05g007420.1 | Solyc05g007420.2 |
| Solyc05g007450.2 | Solyc05g007450.3 |
| Solyc05g007460.2 | Solyc05g007460.3 |
| Solyc05g007490.2 | Solyc05g007490.3 |
| Solyc05g007560.2 | Solyc05g007560.3 |
| Solyc05g007630.2 | Solyc05g007630.3 |
| Solyc05g007640.2 | Solyc05g007640.3 |
| Solyc05g007690.1 | Solyc05g007690.2 |
| Solyc05g007710.2 | Solyc05g007710.3 |
| Solyc05g007760.1 | Solyc05g007760.2 |
| Solyc05g007770.2 | Solyc05g007770.3 |
| Solyc05g007780.2 | Solyc05g007780.3 |
| Solyc05g007790.2 | Solyc05g007790.3 |
| Solyc05g007810.2 | Solyc05g007810.3 |
| Solyc05g007830.2 | Solyc05g007830.3 |
| Solyc05g007840.2 | Solyc05g007840.3 |
| Solyc05g007850.1 | Solyc05g007850.2 |
| Solyc05g007910.2 | Solyc05g007910.3 |
| Solyc05g007920.2 | Solyc05g007920.3 |
| Solyc05g007930.2 | Solyc05g007930.3 |
| Solyc05g007940.2 | Solyc05g007940.3 |
| Solyc05g007980.2 | Solyc05g007980.3 |
| Solyc05g008000.2 | Solyc05g008000.3 |
| Solyc05g008010.2 | Solyc05g008010.3 |
| Solyc05g008090.2 | Solyc05g008090.3 |
| Solyc05g008150.2 | Solyc05g008150.3 |
| Solyc05g008180.2 | Solyc05g008180.3 |
| Solyc05g008190.2 | Solyc05g008190.3 |
| Solyc05g008230.2 | Solyc05g008230.3 |

| | |
|---|---|
| Solyc05g008250.1 | Solyc05g008250.2 |
| Solyc05g008260.2 | Solyc05g008260.3 |
| Solyc05g008270.2 | Solyc05g008270.3 |
| Solyc05g008280.1 | Solyc05g008280.2 |
| Solyc05g008340.2 | Solyc05g008340.3 |
| Solyc05g008350.2 | Solyc05g008350.3 |
| Solyc05g008370.1 | Solyc05g008370.2 |
| Solyc05g008390.2 | Solyc05g008390.3 |
| Solyc05g008450.2 | Solyc05g008450.3 |
| Solyc05g008490.1 | Solyc05g008490.2 |
| Solyc05g008510.2 | Solyc05g008510.3 |
| Solyc05g008520.2 | Solyc05g008520.3 |
| Solyc05g008530.2 | Solyc05g008530.3 |
| Solyc05g008540.2 | Solyc05g008540.3 |
| Solyc05g008565.1 | Solyc05g150106.1 |
| Solyc05g008580.2 | Solyc05g008580.3 |
| Solyc05g008600.2 | Solyc05g008600.3 |
| Solyc05g008610.2 | Solyc05g008610.3 |
| Solyc05g008670.2 | Solyc05g008670.3 |
| Solyc05g008680.1 | Solyc05g008680.2 |
| Solyc05g008690.1 | Solyc05g008690.2 |
| Solyc05g008720.1 | Solyc05g008720.2 |
| Solyc05g008750.2 | Solyc05g008750.3 |
| Solyc05g008770.2 | Solyc05g008770.3 |
| Solyc05g008830.2 | Solyc05g008830.3 |
| Solyc05g008840.1 | Solyc05g008840.2 |
| Solyc05g008870.2 | Solyc05g008870.3 |
| Solyc05g008890.2 | Solyc05g008890.3 |
| Solyc05g008910.2 | Solyc05g008910.3 |
| Solyc05g008920.2 | Solyc05g008920.3 |
| Solyc05g008930.2 | Solyc05g008930.3 |
| Solyc05g008940.2 | Solyc05g008940.3 |
| Solyc05g008960.2 | Solyc05g008960.3 |
| Solyc05g008990.2 | Solyc05g008990.3 |
| Solyc05g009040.2 | Solyc05g009040.3 |
| Solyc05g009050.2 | Solyc05g009050.3 |
| Solyc05g009110.1 | Solyc05g009110.2 |
| Solyc05g009120.2 | Solyc05g009120.3 |
| Solyc05g009140.2 | Solyc05g009140.3 |
| Solyc05g009160.2 | Solyc05g009160.3 |
| Solyc05g009220.2 | Solyc05g009220.3 |
| Solyc05g009240.2 | Solyc05g009240.3 |
| Solyc05g009310.2 | Solyc05g009310.3 |

| | |
|---|---|
| Solyc05g009330.2 | Solyc05g009330.3 |
| Solyc05g009390.2 | Solyc05g009390.3 |
| Solyc05g009430.2 | Solyc05g009430.3 |
| Solyc05g009440.1 | Solyc05g009440.2 |
| Solyc05g009460.2 | Solyc05g009460.3 |
| Solyc05g009490.2 | Solyc05g009490.3 |
| Solyc05g009500.2 | Solyc05g009500.3 |
| Solyc05g009510.1 | Solyc05g009510.2 |
| Solyc05g009530.2 | Solyc05g009530.3 |
| Solyc05g009540.2 | Solyc05g009540.3 |
| Solyc05g009550.2 | Solyc05g009550.3 |
| Solyc05g009570.2 | Solyc05g009570.3 |
| Solyc05g009580.2 | Solyc05g009580.3 |
| Solyc05g009590.2 | Solyc05g009590.3 |
| Solyc05g009670.2 | Solyc05g009670.3 |
| Solyc05g009690.2 | Solyc05g009690.3 |
| Solyc05g009720.2 | Solyc05g009720.3 |
| Solyc05g009780.2 | Solyc05g009780.3 |
| Solyc05g009810.1 | Solyc05g009810.2 |
| Solyc05g009840.2 | Solyc05g009840.3 |
| Solyc05g009880.2 | Solyc05g009880.3 |
| Solyc05g009910.2 | Solyc05g009910.3 |
| Solyc05g009930.2 | Solyc05g009930.3 |
| Solyc05g009950.2 | Solyc05g009950.3 |
| Solyc05g010050.2 | Solyc05g010050.3 |
| Solyc05g010070.2 | Solyc05g010070.3 |
| Solyc05g010080.2 | Solyc05g010080.3 |
| Solyc05g010090.1 | Solyc05g010090.2 |
| Solyc05g010140.2 | Solyc05g010140.3 |
| Solyc05g010150.2 | Solyc05g010150.3 |
| Solyc05g010160.2 | Solyc05g010160.3 |
| Solyc05g010170.1 | Solyc05g010170.2 |
| Solyc05g010190.1 | Solyc05g010190.2 |
| Solyc05g010200.1 | Solyc05g010200.2 |
| Solyc05g010220.1 | Solyc05g010220.2 |
| Solyc05g010280.2 | Solyc05g010280.3 |
| Solyc05g010310.2 | Solyc05g010310.3 |
| Solyc05g010340.2 | Solyc05g010340.3 |
| Solyc05g010360.1 | Solyc05g010360.2 |
| Solyc05g010400.2 | Solyc05g010400.3 |
| Solyc05g010420.1 | Solyc05g010420.2 |
| Solyc05g010440.1 | Solyc05g010440.2 |
| Solyc05g010530.2 | Solyc05g010530.3 |

| | |
|---|---|
| Solyc05g010570.2 | Solyc05g010570.3 |
| Solyc05g010565.1 | Solyc05g150107.1 |
| Solyc05g010600.2 | Solyc05g010600.3 |
| Solyc05g010610.2 | Solyc05g010610.3 |
| Solyc05g010640.2 | Solyc05g010640.3 |
| Solyc05g010650.2 | Solyc05g010650.3 |
| Solyc05g010670.2 | Solyc05g010670.3 |
| Solyc05g010680.1 | Solyc05g010680.2 |
| Solyc05g010730.1 | Solyc05g010730.3 |
| Solyc05g010740.1 | Solyc05g010740.2 |
| Solyc05g010750.2 | Solyc05g010750.3 |
| Solyc05g010765.1 | Solyc05g150108.1 |
| Solyc05g010800.2 | Solyc05g010800.3 |
| Solyc05g011825.1 | Solyc05g011825.2 |
| Solyc05g011840.2 | Solyc05g011840.3 |
| Solyc05g011920.2 | Solyc05g011920.3 |
| Solyc05g011970.2 | Solyc05g011970.3 |
| Solyc05g011980.3 | Solyc05g011980.4 |
| Solyc05g012000.2 | Solyc05g012000.3 |
| Solyc05g012010.2 | Solyc05g012010.3 |
| Solyc05g012020.3 | Solyc05g012020.4 |
| Solyc05g012070.2 | Solyc05g012070.3 |
| Solyc05g012080.2 | Solyc05g012080.3 |
| Solyc05g012100.2 | Solyc05g012100.3 |
| Solyc05g012120.2 | Solyc05g012120.3 |
| Solyc05g012140.2 | Solyc05g012140.3 |
| Solyc05g012160.2 | Solyc05g012160.3 |
| Solyc05g012180.2 | Solyc05g012180.3 |
| Solyc05g012190.2 | Solyc05g012190.3 |
| Solyc05g012200.2 | Solyc05g012200.3 |
| Solyc05g012220.2 | Solyc05g012220.3 |
| Solyc05g012260.2 | Solyc05g012260.3 |
| Solyc05g012270.2 | Solyc05g012270.3 |
| Solyc05g012280.2 | Solyc05g012280.3 |
| Solyc05g012290.2 | Solyc05g012290.3 |
| Solyc05g012360.2 | Solyc05g012360.3 |
| Solyc05g012370.2 | Solyc05g012370.3 |
| Solyc05g012380.2 | Solyc05g012380.3 |
| Solyc05g012410.1 | Solyc05g012410.2 |
| Solyc05g012490.1 | Solyc05g012490.2 |
| Solyc05g012500.2 | Solyc05g012500.3 |
| Solyc05g012510.2 | Solyc05g012510.3 |
| Solyc05g012520.2 | Solyc05g012520.3 |

| | |
|---|---|
| Solyc05g012570.2 | Solyc05g012570.3 |
| Solyc05g012610.2 | Solyc05g012610.3 |
| Solyc05g012630.1 | Solyc05g012630.2 |
| Solyc05g012670.1 | Solyc05g012670.2 |
| Solyc05g012700.2 | Solyc05g012700.3 |
| Solyc05g012720.2 | Solyc05g012720.3 |
| Solyc05g012740.1 | Solyc05g012740.2 |
| Solyc05g012770.2 | Solyc05g012770.3 |
| Solyc05g012805.1 | Solyc05g150109.1 |
| Solyc05g012823.1 | Solyc05g150110.1 |
| Solyc05g012860.1 | Solyc05g012860.2 |
| Solyc05g012880.1 | Solyc05g012880.2 |
| Solyc05g013010.2 | Solyc05g013010.3 |
| Solyc05g013040.2 | Solyc05g013040.3 |
| Solyc05g013050.2 | Solyc05g013050.3 |
| Solyc05g013070.2 | Solyc05g013070.3 |
| Solyc05g013100.1 | Solyc05g013100.2 |
| Solyc05g013110.1 | Solyc05g013110.2 |
| Solyc05g013130.2 | Solyc05g013130.3 |
| Solyc05g013210.2 | Solyc05g013210.3 |
| Solyc05g013270.1 | Solyc05g013270.2 |
| Solyc05g013290.1 | Solyc05g013290.2 |
| Solyc05g013340.2 | Solyc05g013340.3 |
| Solyc05g013360.2 | Solyc05g013360.3 |
| Solyc05g013380.2 | Solyc05g013380.3 |
| Solyc05g013410.2 | Solyc05g013410.3 |
| Solyc05g013440.2 | Solyc05g013440.3 |
| Solyc05g013560.2 | Solyc05g013560.3 |
| Solyc05g013570.2 | Solyc05g013570.3 |
| Solyc05g013670.2 | Solyc05g013670.3 |
| Solyc05g013680.2 | Solyc05g013680.3 |
| Solyc05g013710.2 | Solyc05g013710.3 |
| Solyc05g013720.2 | Solyc05g013720.3 |
| Solyc05g013730.2 | Solyc05g013730.3 |
| Solyc05g013740.2 | Solyc05g013740.3 |
| Solyc05g013750.2 | Solyc05g013750.3 |
| Solyc05g013900.1 | Solyc05g013900.2 |
| Solyc05g013930.1 | Solyc05g013930.2 |
| Solyc05g013960.2 | Solyc05g013960.3 |
| Solyc05g013980.2 | Solyc05g013980.3 |
| Solyc05g013985.1 | Solyc05g150111.1 |
| Solyc05g013990.2 | Solyc05g013990.3 |
| Solyc05g014030.2 | Solyc05g014030.3 |

| | |
|---|---|
| Solyc05g014050.2 | Solyc05g014050.3 |
| Solyc05g014110.2 | Solyc05g014110.3 |
| Solyc05g014130.2 | Solyc05g014130.3 |
| Solyc05g014170.1 | Solyc05g014170.2 |
| Solyc05g014177.1 | Solyc05g014177.2 |
| Solyc05g014200.2 | Solyc05g014200.3 |
| Solyc05g014230.2 | Solyc05g014230.3 |
| Solyc05g014250.2 | Solyc05g014250.3 |
| Solyc05g014260.2 | Solyc05g014260.3 |
| Solyc05g014275.1 | Solyc05g014275.2 |
| Solyc05g014340.2 | Solyc05g014340.3 |
| Solyc05g014370.2 | Solyc05g014370.3 |
| Solyc05g014470.2 | Solyc05g014470.3 |
| Solyc05g014510.2 | Solyc05g014510.3 |
| Solyc05g014540.2 | Solyc05g014540.3 |
| Solyc05g014560.2 | Solyc05g014560.3 |
| Solyc05g014580.2 | Solyc05g014580.3 |
| Solyc05g014590.2 | Solyc05g014590.3 |
| Solyc05g014630.1 | Solyc05g014630.2 |
| Solyc05g014640.2 | Solyc05g014640.3 |
| Solyc05g014700.2 | Solyc05g014700.3 |
| Solyc05g014760.2 | Solyc05g014760.3 |
| Solyc05g014765.1 | Solyc05g150112.1 |
| Solyc05g014790.2 | Solyc05g014790.3 |
| Solyc05g014840.1 | Solyc05g014840.2 |
| Solyc05g014890.1 | Solyc05g014890.2 |
| Solyc05g014895.1 | Solyc05g150113.1 |
| Solyc05g014970.2 | Solyc05g014970.3 |
| Solyc05g015050.2 | Solyc05g015050.3 |
| Solyc05g015093.1 | Solyc05g150114.1 |
| Solyc05g015097.1 | Solyc05g150115.1 |
| Solyc05g015150.2 | Solyc05g015150.3 |
| Solyc05g015160.2 | Solyc05g015160.3 |
| Solyc05g015220.2 | Solyc05g015220.3 |
| Solyc05g015290.1 | Solyc05g015290.2 |
| Solyc05g015310.2 | Solyc05g015310.3 |
| Solyc05g015320.2 | Solyc05g015320.3 |
| Solyc05g015370.2 | Solyc05g015370.3 |
| Solyc05g015390.2 | Solyc05g015390.3 |
| Solyc05g015433.1 | Solyc05g150116.1 |
| Solyc05g015460.2 | Solyc05g015460.3 |
| Solyc05g015480.2 | Solyc05g015480.3 |
| Solyc05g015510.2 | Solyc05g015510.3 |

| | |
|---|---|
| Solyc05g015520.2 | Solyc05g015520.3 |
| Solyc05g015530.2 | Solyc05g015530.3 |
| Solyc05g015550.1 | Solyc05g015550.2 |
| Solyc05g015610.2 | Solyc05g015610.3 |
| Solyc05g015630.2 | Solyc05g015630.3 |
| Solyc05g015650.2 | Solyc05g015650.3 |
| Solyc05g015656.1 | Solyc05g150117.1 |
| Solyc05g015658.1 | Solyc05g150118.1 |
| Solyc05g015720.1 | Solyc05g015720.2 |
| Solyc05g015750.2 | Solyc05g015750.3 |
| Solyc05g015790.1 | Solyc05g015790.2 |
| Solyc05g015800.2 | Solyc05g015800.3 |
| Solyc05g015840.2 | Solyc05g015840.3 |
| Solyc05g015880.2 | Solyc05g015880.3 |
| Solyc05g015920.2 | Solyc05g015920.3 |
| Solyc05g015940.1 | Solyc05g015940.2 |
| Solyc05g015945.1 | Solyc05g150119.1 |
| Solyc05g015950.2 | Solyc05g015950.3 |
| Solyc05g015955.1 | Solyc05g150120.1 |
| Solyc05g016030.2 | Solyc05g016030.3 |
| Solyc05g016040.1 | Solyc05g016040.2 |
| Solyc05g016070.1 | Solyc05g016070.2 |
| Solyc05g016120.1 | Solyc05g016120.2 |
| Solyc05g016180.2 | Solyc05g016180.3 |
| Solyc05g016230.2 | Solyc05g016230.3 |
| Solyc05g016250.2 | Solyc05g016250.3 |
| Solyc05g016260.2 | Solyc05g016260.3 |
| Solyc05g016320.1 | Solyc05g016320.2 |
| Solyc05g016330.2 | Solyc05g016330.3 |
| Solyc05g016370.1 | Solyc05g016370.2 |
| Solyc05g016410.2 | Solyc05g016410.3 |
| Solyc05g016440.2 | Solyc05g016440.3 |
| Solyc05g016460.1 | Solyc05g016460.2 |
| Solyc05g016690.2 | Solyc05g016690.3 |
| Solyc05g017843.1 | Solyc05g150122.1 |
| Solyc05g017860.2 | Solyc05g017860.3 |
| Solyc05g017900.2 | Solyc05g017900.3 |
| Solyc05g017930.1 | Solyc05g017930.2 |
| Solyc05g017940.2 | Solyc05g017940.3 |
| Solyc05g017950.2 | Solyc05g017950.3 |
| Solyc05g017980.2 | Solyc05g017980.3 |
| Solyc05g018120.1 | Solyc05g018120.2 |
| Solyc05g018130.2 | Solyc05g018130.3 |

| | |
|---|---|
| Solyc05g018150.2 | Solyc05g018150.3 |
| Solyc05g018180.2 | Solyc05g018180.3 |
| Solyc05g018210.2 | Solyc05g018210.3 |
| Solyc05g018300.2 | Solyc05g018300.3 |
| Solyc05g018320.2 | Solyc05g018320.3 |
| Solyc05g018390.2 | Solyc05g018390.3 |
| Solyc05g018395.1 | Solyc05g150123.1 |
| Solyc05g018500.2 | Solyc05g018500.3 |
| Solyc05g018510.2 | Solyc05g018510.3 |
| Solyc05g018520.2 | Solyc05g018520.3 |
| Solyc05g018570.2 | Solyc05g018570.3 |
| Solyc05g018590.2 | Solyc05g018590.3 |
| Solyc05g018600.2 | Solyc05g018600.3 |
| Solyc05g018615.1 | Solyc05g150124.1 |
| Solyc05g018640.2 | Solyc05g018640.3 |
| Solyc05g018653.1 | Solyc05g150125.1 |
| Solyc05g018770.1 | Solyc05g018770.2 |
| Solyc05g018830.1 | Solyc05g018830.2 |
| Solyc05g018910.1 | Solyc05g018910.2 |
| Solyc05g018920.1 | Solyc05g018920.2 |
| Solyc05g019980.1 | Solyc05g019980.2 |
| Solyc05g020005.1 | Solyc05g150126.1 |
| Solyc05g020040.1 | Solyc05g020040.2 |
| Solyc05g021070.1 | Solyc05g021070.2 |
| Solyc05g021105.1 | Solyc05g150127.1 |
| Solyc05g021145.1 | Solyc05g150128.1 |
| Solyc05g021160.1 | Solyc05g021160.2 |
| Solyc05g021190.1 | Solyc05g021190.2 |
| Solyc05g021370.1 | Solyc05g021370.2 |
| Solyc05g021400.1 | Solyc05g021400.2 |
| Solyc05g021485.1 | Solyc05g150129.1 |
| Solyc05g021500.1 | Solyc05g021500.2 |
| Solyc05g021510.1 | Solyc05g021510.2 |
| Solyc05g021535.1 | Solyc05g150130.1 |
| Solyc05g021580.2 | Solyc05g021580.3 |
| Solyc05g023605.1 | Solyc05g150131.1 |
| Solyc05g023700.1 | Solyc05g023700.2 |
| Solyc05g023710.2 | Solyc05g023710.3 |
| Solyc05g023740.2 | Solyc05g023740.3 |
| Solyc05g023750.2 | Solyc05g023750.3 |
| Solyc05g023760.2 | Solyc05g023760.3 |
| Solyc05g023800.2 | Solyc05g023800.3 |
| Solyc05g023820.2 | Solyc05g023820.3 |

| | |
|---|---|
| Solyc05g023835.1 | Solyc05g150132.1 |
| Solyc05g023880.1 | Solyc05g023880.2 |
| Solyc05g023990.1 | Solyc05g023990.2 |
| Solyc05g024010.2 | Solyc05g024010.3 |
| Solyc05g024030.1 | Solyc05g024030.2 |
| Solyc05g024160.2 | Solyc05g024160.3 |
| Solyc05g024180.2 | Solyc05g024180.3 |
| Solyc05g024190.2 | Solyc05g024190.3 |
| Solyc05g024195.1 | Solyc05g150133.1 |
| Solyc05g024200.1 | Solyc05g024200.2 |
| Solyc05g024260.2 | Solyc05g024260.3 |
| Solyc05g024290.2 | Solyc05g024290.3 |
| Solyc05g025500.2 | Solyc05g025500.3 |
| Solyc05g025510.2 | Solyc05g025510.3 |
| Solyc05g025560.1 | Solyc05g025560.2 |
| Solyc05g025650.2 | Solyc05g025650.3 |
| Solyc05g025730.1 | Solyc05g025730.2 |
| Solyc05g025750.2 | Solyc05g025750.3 |
| Solyc05g025790.2 | Solyc05g025790.3 |
| Solyc05g025810.2 | Solyc05g025810.3 |
| Solyc05g025900.2 | Solyc05g025900.3 |
| Solyc05g025910.2 | Solyc05g025910.3 |
| Solyc05g025920.2 | Solyc05g025920.3 |
| Solyc05g025957.1 | Solyc05g150134.1 |
| Solyc05g026130.2 | Solyc05g026130.3 |
| Solyc05g026157.1 | Solyc05g150135.1 |
| Solyc05g026180.1 | Solyc05g026180.2 |
| Solyc05g026210.1 | Solyc05g026210.2 |
| Solyc05g026315.1 | Solyc05g150136.1 |
| Solyc05g026350.1 | Solyc05g026350.2 |
| Solyc05g026470.1 | Solyc05g026470.2 |
| Solyc05g026480.1 | Solyc05g026480.2 |
| Solyc05g026490.2 | Solyc05g026490.3 |
| Solyc05g026520.1 | Solyc05g026520.2 |
| Solyc05g026550.2 | Solyc05g026550.3 |
| Solyc05g026560.2 | Solyc05g026560.3 |
| Solyc05g026570.2 | Solyc05g026570.3 |
| Solyc05g026590.1 | Solyc05g026590.2 |
| Solyc05g026600.2 | Solyc05g026600.3 |
| Solyc05g032880.2 | Solyc05g032880.3 |
| Solyc05g032850.2 | Solyc05g032850.3 |
| Solyc05g032750.2 | Solyc05g032750.3 |
| Solyc05g039915.1 | Solyc05g150137.1 |

| | |
|---|---|
| Solyc05g039950.1 | Solyc05g039950.2 |
| Solyc05g040020.1 | Solyc05g040020.2 |
| Solyc05g041117.1 | Solyc05g150138.1 |
| Solyc05g041320.1 | Solyc05g041320.2 |
| Solyc05g041340.1 | Solyc05g041340.2 |
| Solyc05g041430.2 | Solyc05g041430.3 |
| Solyc05g041530.2 | Solyc05g041530.3 |
| Solyc05g041623.1 | Solyc05g150139.1 |
| Solyc05g041627.1 | Solyc05g150140.1 |
| Solyc05g041720.1 | Solyc05g041720.2 |
| Solyc05g041870.1 | Solyc05g041870.2 |
| Solyc05g041880.1 | Solyc05g041880.2 |
| Solyc05g041910.2 | Solyc05g041910.3 |
| Solyc05g041920.2 | Solyc05g041920.3 |
| Solyc05g041995.1 | Solyc05g150141.1 |
| Solyc05g042030.2 | Solyc05g042030.3 |
| Solyc05g042040.2 | Solyc05g042040.3 |
| Solyc05g042090.1 | Solyc05g042090.2 |
| Solyc05g042150.1 | Solyc05g042150.2 |
| Solyc05g043330.2 | Solyc05g043330.3 |
| Solyc05g043390.1 | Solyc05g043390.2 |
| Solyc05g043400.1 | Solyc05g043400.2 |
| Solyc05g043420.1 | Solyc05g043420.2 |
| Solyc05g044465.1 | Solyc05g150142.1 |
| Solyc05g044480.2 | Solyc05g044480.3 |
| Solyc05g044490.2 | Solyc05g044490.3 |
| Solyc05g044510.1 | Solyc05g044510.2 |
| Solyc05g044570.1 | Solyc05g044570.2 |
| Solyc05g044630.2 | Solyc05g044630.3 |
| Solyc05g045660.2 | Solyc05g045660.3 |
| Solyc05g045870.1 | Solyc05g045870.2 |
| Solyc05g046020.2 | Solyc05g046020.3 |
| Solyc05g046040.1 | Solyc05g046040.2 |
| Solyc05g046120.2 | Solyc05g046120.3 |
| Solyc05g046140.2 | Solyc05g046140.3 |
| Solyc05g046145.1 | Solyc05g150143.1 |
| Solyc05g046150.2 | Solyc05g046150.3 |
| Solyc05g046200.2 | Solyc05g046200.3 |
| Solyc05g046205.1 | Solyc05g150144.1 |
| Solyc05g046283.1 | Solyc05g150145.1 |
| Solyc05g046287.1 | Solyc05g150146.1 |
| Solyc05g046290.2 | Solyc05g046290.3 |
| Solyc05g046300.2 | Solyc05g046300.3 |

| | |
|---|---|
| Solyc05g046340.1 | Solyc05g046340.2 |
| Solyc05g046360.2 | Solyc05g046360.3 |
| Solyc05g047460.2 | Solyc05g047460.3 |
| Solyc05g047480.1 | Solyc05g047480.2 |
| Solyc05g047530.2 | Solyc05g047530.3 |
| Solyc05g047610.2 | Solyc05g047610.3 |
| Solyc05g047640.2 | Solyc05g047640.3 |
| Solyc05g047690.2 | Solyc05g047690.3 |
| Solyc05g047710.2 | Solyc05g047710.3 |
| Solyc05g047712.1 | Solyc05g150147.1 |
| Solyc05g047714.1 | Solyc05g150148.1 |
| Solyc05g047716.1 | Solyc05g150149.1 |
| Solyc05g047718.1 | Solyc05g150150.1 |
| Solyc05g048740.2 | Solyc05g048740.3 |
| Solyc05g048760.2 | Solyc05g048760.3 |
| Solyc05g048770.2 | Solyc05g048770.3 |
| Solyc05g048820.2 | Solyc05g048820.3 |
| Solyc05g048830.2 | Solyc05g048830.3 |
| Solyc05g048860.2 | Solyc05g048860.3 |
| Solyc05g049870.2 | Solyc05g049870.3 |
| Solyc05g049900.2 | Solyc05g049900.3 |
| Solyc05g049950.2 | Solyc05g049950.3 |
| Solyc05g049970.2 | Solyc05g049970.3 |
| Solyc05g049980.2 | Solyc05g049980.3 |
| Solyc05g050010.2 | Solyc05g050010.3 |
| Solyc05g050015.1 | Solyc05g150151.1 |
| Solyc05g050020.2 | Solyc05g050020.3 |
| Solyc05g050110.2 | Solyc05g050110.3 |
| Solyc05g050120.2 | Solyc05g050120.3 |
| Solyc05g050220.2 | Solyc05g050220.3 |
| Solyc05g050240.2 | Solyc05g050240.3 |
| Solyc05g050280.2 | Solyc05g050280.3 |
| Solyc05g050330.2 | Solyc05g050330.3 |
| Solyc05g050350.1 | Solyc05g050350.2 |
| Solyc05g050360.2 | Solyc05g050360.3 |
| Solyc05g050430.1 | Solyc05g050430.2 |
| Solyc05g050510.2 | Solyc05g050510.3 |
| Solyc05g050530.2 | Solyc05g050530.3 |
| Solyc05g050570.2 | Solyc05g050570.3 |
| Solyc05g050580.2 | Solyc05g050580.3 |
| Solyc05g050590.2 | Solyc05g050590.3 |
| Solyc05g050600.2 | Solyc05g050600.3 |
| Solyc05g050650.1 | Solyc05g050650.2 |

| | |
|---|---|
| Solyc05g050670.1 | Solyc05g050670.2 |
| Solyc05g050690.1 | Solyc05g050690.2 |
| Solyc05g050710.2 | Solyc05g050710.3 |
| Solyc05g050740.2 | Solyc05g050740.3 |
| Solyc05g050760.1 | Solyc05g050760.2 |
| Solyc05g050780.2 | Solyc05g050780.3 |
| Solyc05g050800.2 | Solyc05g050800.3 |
| Solyc05g050820.2 | Solyc05g050820.3 |
| Solyc05g050840.2 | Solyc05g050840.3 |
| Solyc05g050850.2 | Solyc05g050850.3 |
| Solyc05g050860.1 | Solyc05g050860.2 |
| Solyc05g050910.2 | Solyc05g050910.3 |
| Solyc05g050920.2 | Solyc05g050920.3 |
| Solyc05g050930.1 | Solyc05g050930.2 |
| Solyc05g050980.2 | Solyc05g050980.3 |
| Solyc05g051000.2 | Solyc05g051000.3 |
| Solyc05g051030.2 | Solyc05g051030.3 |
| Solyc05g051050.2 | Solyc05g051050.3 |
| Solyc05g051060.2 | Solyc05g051060.3 |
| Solyc05g051070.2 | Solyc05g051070.3 |
| Solyc05g051100.1 | Solyc05g051100.2 |
| Solyc05g051210.2 | Solyc05g051210.3 |
| Solyc05g051220.2 | Solyc05g051220.3 |
| Solyc05g051250.2 | Solyc05g051250.3 |
| Solyc05g051260.2 | Solyc05g051260.3 |
| Solyc05g051270.2 | Solyc05g051270.3 |
| Solyc05g051280.2 | Solyc05g051280.3 |
| Solyc05g051320.1 | Solyc05g051320.2 |
| Solyc05g051350.1 | Solyc05g051350.2 |
| Solyc05g051390.2 | Solyc05g051390.3 |
| Solyc05g051410.2 | Solyc05g051410.3 |
| Solyc05g051420.1 | Solyc05g051420.2 |
| Solyc05g051425.1 | Solyc05g051425.2 |
| Solyc05g051450.1 | Solyc05g051450.2 |
| Solyc05g051460.2 | Solyc05g051460.3 |
| Solyc05g051490.2 | Solyc05g051490.3 |
| Solyc05g051520.2 | Solyc05g051520.3 |
| Solyc05g051550.1 | Solyc05g051550.2 |
| Solyc05g051570.2 | Solyc05g051570.3 |
| Solyc05g051580.2 | Solyc05g051580.3 |
| Solyc05g051590.2 | Solyc05g051590.3 |
| Solyc05g051620.2 | Solyc05g051620.3 |
| Solyc05g051690.2 | Solyc05g051690.3 |

| | |
|---|---|
| Solyc05g051700.2 | Solyc05g051700.3 |
| Solyc05g051750.2 | Solyc05g051750.3 |
| Solyc05g051790.2 | Solyc05g051790.3 |
| Solyc05g051800.2 | Solyc05g051800.3 |
| Solyc05g051820.2 | Solyc05g051820.3 |
| Solyc05g051850.2 | Solyc05g051850.3 |
| Solyc05g051870.2 | Solyc05g051870.3 |
| Solyc05g051900.2 | Solyc05g051900.3 |
| Solyc05g051920.2 | Solyc05g051920.3 |
| Solyc05g051940.2 | Solyc05g051940.3 |
| Solyc05g051990.2 | Solyc05g051990.3 |
| Solyc05g052020.2 | Solyc05g052020.3 |
| Solyc05g052070.2 | Solyc05g052070.3 |
| Solyc05g052100.2 | Solyc05g052100.3 |
| Solyc05g052110.2 | Solyc05g052110.3 |
| Solyc05g052130.2 | Solyc05g052130.3 |
| Solyc05g052140.2 | Solyc05g052140.3 |
| Solyc05g052180.2 | Solyc05g052180.3 |
| Solyc05g052220.2 | Solyc05g052220.3 |
| Solyc05g052240.2 | Solyc05g052240.3 |
| Solyc05g052250.2 | Solyc05g052250.3 |
| Solyc05g052260.2 | Solyc05g052260.3 |
| Solyc05g052280.2 | Solyc05g052280.3 |
| Solyc05g052310.2 | Solyc05g052310.3 |
| Solyc05g052350.2 | Solyc05g052350.3 |
| Solyc05g052360.1 | Solyc05g052360.2 |
| Solyc05g052370.2 | Solyc05g052370.3 |
| Solyc05g052390.2 | Solyc05g052390.3 |
| Solyc05g052400.2 | Solyc05g052400.3 |
| Solyc05g052440.1 | Solyc05g052440.2 |
| Solyc05g052450.2 | Solyc05g052450.3 |
| Solyc05g052470.2 | Solyc05g052470.3 |
| Solyc05g052490.2 | Solyc05g052490.3 |
| Solyc05g052520.2 | Solyc05g052520.3 |
| Solyc05g052540.1 | Solyc05g052540.2 |
| Solyc05g052570.2 | Solyc05g052570.3 |
| Solyc05g052580.2 | Solyc05g052580.3 |
| Solyc05g052610.2 | Solyc05g052610.3 |
| Solyc05g052640.2 | Solyc05g052640.3 |
| Solyc05g052710.2 | Solyc05g052710.3 |
| Solyc05g052750.2 | Solyc05g052750.3 |
| Solyc05g052790.2 | Solyc05g052790.3 |
| Solyc05g052800.2 | Solyc05g052800.3 |

| | |
|---|---|
| Solyc05g052820.2 | Solyc05g052820.3 |
| Solyc05g052830.2 | Solyc05g052830.3 |
| Solyc05g052850.2 | Solyc05g052850.3 |
| Solyc05g052860.2 | Solyc05g052860.3 |
| Solyc05g052890.2 | Solyc05g052890.3 |
| Solyc05g052900.2 | Solyc05g052900.3 |
| Solyc05g052930.2 | Solyc05g052930.3 |
| Solyc05g052940.2 | Solyc05g052940.3 |
| Solyc05g053020.2 | Solyc05g053020.3 |
| Solyc05g053040.2 | Solyc05g053040.3 |
| Solyc05g053090.1 | Solyc05g053090.2 |
| Solyc05g053100.2 | Solyc05g053100.3 |
| Solyc05g053130.2 | Solyc05g053130.3 |
| Solyc05g053140.2 | Solyc05g053140.3 |
| Solyc05g053150.1 | Solyc05g053150.2 |
| Solyc05g053160.2 | Solyc05g053160.3 |
| Solyc05g053170.2 | Solyc05g053170.3 |
| Solyc05g053180.2 | Solyc05g053180.3 |
| Solyc05g053190.2 | Solyc05g053190.3 |
| Solyc05g053220.2 | Solyc05g053220.3 |
| Solyc05g053230.2 | Solyc05g053230.3 |
| Solyc05g053260.2 | Solyc05g053260.3 |
| Solyc05g053280.2 | Solyc05g053280.3 |
| Solyc05g053290.2 | Solyc05g053290.3 |
| Solyc05g053300.2 | Solyc05g053300.3 |
| Solyc05g053310.2 | Solyc05g053310.3 |
| Solyc05g053330.2 | Solyc05g053330.3 |
| Solyc05g053350.2 | Solyc05g053350.3 |
| Solyc05g053370.2 | Solyc05g053370.3 |
| Solyc05g053430.2 | Solyc05g053430.3 |
| Solyc05g053450.2 | Solyc05g053450.3 |
| Solyc05g053470.2 | Solyc05g053470.3 |
| Solyc05g053520.2 | Solyc05g053520.3 |
| Solyc05g053540.2 | Solyc05g053540.3 |
| Solyc05g053550.2 | Solyc05g053550.3 |
| Solyc05g053600.2 | Solyc05g053600.3 |
| Solyc05g053620.2 | Solyc05g053620.3 |
| Solyc05g053650.2 | Solyc05g053650.3 |
| Solyc05g053670.2 | Solyc05g053670.3 |
| Solyc05g053690.1 | Solyc05g053690.2 |
| Solyc05g053740.2 | Solyc05g053740.3 |
| Solyc05g053800.2 | Solyc05g053800.3 |
| Solyc05g053810.2 | Solyc05g053810.3 |

| | |
|---|---|
| Solyc05g053820.2 | Solyc05g053820.3 |
| Solyc05g053830.2 | Solyc05g053830.3 |
| Solyc05g053835.1 | Solyc05g053835.2 |
| Solyc05g053850.2 | Solyc05g053850.3 |
| Solyc05g053870.2 | Solyc05g053870.3 |
| Solyc05g053940.2 | Solyc05g053940.3 |
| Solyc05g053960.2 | Solyc05g053960.3 |
| Solyc05g054030.2 | Solyc05g054030.3 |
| Solyc05g054060.2 | Solyc05g054060.3 |
| Solyc05g054080.2 | Solyc05g054080.3 |
| Solyc05g054090.2 | Solyc05g054090.3 |
| Solyc05g054100.2 | Solyc05g054100.3 |
| Solyc05g054110.2 | Solyc05g054110.3 |
| Solyc05g054140.2 | Solyc05g054140.3 |
| Solyc05g054150.2 | Solyc05g054150.3 |
| Solyc05g054180.2 | Solyc05g054180.3 |
| Solyc05g054210.2 | Solyc05g054210.3 |
| Solyc05g054220.2 | Solyc05g054220.3 |
| Solyc05g054240.2 | Solyc05g054240.3 |
| Solyc05g054260.2 | Solyc05g054260.3 |
| Solyc05g054290.1 | Solyc05g054290.2 |
| Solyc05g054350.2 | Solyc05g054350.3 |
| Solyc05g054380.1 | Solyc05g054380.2 |
| Solyc05g054400.2 | Solyc05g054400.3 |
| Solyc05g054460.2 | Solyc05g054460.3 |
| Solyc05g054470.2 | Solyc05g054470.3 |
| Solyc05g054480.2 | Solyc05g054480.3 |
| Solyc05g054490.2 | Solyc05g054490.3 |
| Solyc05g054510.1 | Solyc05g054510.2 |
| Solyc05g054560.2 | Solyc05g054560.3 |
| Solyc05g054570.2 | Solyc05g054570.3 |
| Solyc05g054580.2 | Solyc05g054580.3 |
| Solyc05g054600.2 | Solyc05g054600.3 |
| Solyc05g054660.1 | Solyc05g054660.2 |
| Solyc05g054670.2 | Solyc05g054670.3 |
| Solyc05g054690.2 | Solyc05g054690.3 |
| Solyc05g054700.2 | Solyc05g054700.3 |
| Solyc05g054710.2 | Solyc05g054710.3 |
| Solyc05g054720.2 | Solyc05g054720.3 |
| Solyc05g054730.2 | Solyc05g054730.3 |
| Solyc05g054740.2 | Solyc05g054740.3 |
| Solyc05g054750.2 | Solyc05g054750.3 |
| Solyc05g054830.2 | Solyc05g054830.3 |

| | |
|---|---|
| Solyc05g054840.2 | Solyc05g054840.3 |
| Solyc05g054870.2 | Solyc05g054870.3 |
| Solyc05g054880.2 | Solyc05g054880.3 |
| Solyc05g054900.2 | Solyc05g054900.3 |
| Solyc05g054910.2 | Solyc05g054910.3 |
| Solyc05g054930.2 | Solyc05g054930.3 |
| Solyc05g054940.2 | Solyc05g054940.3 |
| Solyc05g054970.2 | Solyc05g054970.3 |
| Solyc05g054980.2 | Solyc05g054980.3 |
| Solyc05g054990.2 | Solyc05g054990.3 |
| Solyc05g055030.1 | Solyc05g055030.2 |
| Solyc05g055040.2 | Solyc05g055040.3 |
| Solyc05g055060.2 | Solyc05g055060.3 |
| Solyc05g055080.1 | Solyc05g055080.2 |
| Solyc05g055100.1 | Solyc05g055100.2 |
| Solyc05g055160.2 | Solyc05g055160.3 |
| Solyc05g055170.2 | Solyc05g055170.3 |
| Solyc05g055180.2 | Solyc05g055180.3 |
| Solyc05g055185.1 | Solyc05g150152.1 |
| Solyc05g055230.1 | Solyc05g055230.2 |
| Solyc05g055240.2 | Solyc05g055240.3 |
| Solyc05g055280.1 | Solyc05g055280.2 |
| Solyc05g055290.2 | Solyc05g055290.3 |
| Solyc05g055310.2 | Solyc05g055310.3 |
| Solyc05g055320.2 | Solyc05g055320.3 |
| Solyc05g055330.2 | Solyc05g055330.3 |
| Solyc05g055343.1 | Solyc05g055343.2 |
| Solyc05g055350.2 | Solyc05g055350.3 |
| Solyc05g055390.2 | Solyc05g055390.3 |
| Solyc05g055450.2 | Solyc05g055450.3 |
| Solyc05g055480.2 | Solyc05g055480.3 |
| Solyc05g055510.2 | Solyc05g055510.3 |
| Solyc05g055580.1 | Solyc05g055580.2 |
| Solyc05g055590.1 | Solyc05g055590.2 |
| Solyc05g055630.2 | Solyc05g055630.3 |
| Solyc05g055650.2 | Solyc05g055650.3 |
| Solyc05g055700.2 | Solyc05g055700.3 |
| Solyc05g055710.2 | Solyc05g055710.3 |
| Solyc05g055750.2 | Solyc05g055750.3 |
| Solyc05g055760.2 | Solyc05g055760.3 |
| Solyc05g055770.2 | Solyc05g055770.3 |
| Solyc05g055820.2 | Solyc05g055820.3 |
| Solyc05g055830.2 | Solyc05g055830.3 |

| | |
|---|---|
| Solyc05g055850.2 | Solyc05g055850.3 |
| Solyc05g055870.2 | Solyc05g055870.3 |
| Solyc05g055890.2 | Solyc05g055890.3 |
| Solyc05g055930.1 | Solyc05g055930.2 |
| Solyc05g055940.2 | Solyc05g055940.3 |
| Solyc05g056000.2 | Solyc05g056000.3 |
| Solyc05g056010.2 | Solyc05g056010.3 |
| Solyc05g056020.2 | Solyc05g056020.3 |
| Solyc05g056030.2 | Solyc05g056030.3 |
| Solyc05g056040.2 | Solyc05g056040.3 |
| Solyc05g056050.2 | Solyc05g056050.3 |
| Solyc05g056090.2 | Solyc05g056090.3 |
| Solyc05g056100.2 | Solyc05g056100.3 |
| Solyc05g056110.2 | Solyc05g056110.3 |
| Solyc05g056120.2 | Solyc05g056120.3 |
| Solyc05g056160.2 | Solyc05g056160.3 |
| Solyc05g056170.2 | Solyc05g056170.3 |
| Solyc05g056220.2 | Solyc05g056220.3 |
| Solyc05g056240.2 | Solyc05g056240.3 |
| Solyc05g056250.2 | Solyc05g056250.3 |
| Solyc05g056270.2 | Solyc05g056270.3 |
| Solyc05g056310.2 | Solyc05g056310.3 |
| Solyc05g056320.2 | Solyc05g056320.3 |
| Solyc05g056325.1 | Solyc05g150153.1 |
| Solyc05g056340.2 | Solyc05g056340.3 |
| Solyc05g056350.2 | Solyc05g056350.3 |
| Solyc05g056360.1 | Solyc05g056360.2 |
| Solyc05g056400.2 | Solyc05g056400.3 |
| Solyc05g056450.2 | Solyc05g056450.3 |
| Solyc05g056480.2 | Solyc05g056480.3 |
| Solyc05g056550.2 | Solyc05g056550.3 |
| Solyc05g056560.2 | Solyc05g056560.3 |
| Solyc05g056570.2 | Solyc05g056570.3 |
| Solyc05g056580.2 | Solyc05g056580.3 |
| Solyc06g005000.2 | Solyc06g005000.3 |
| Solyc06g005060.2 | Solyc06g005060.3 |
| Solyc06g005090.2 | Solyc06g005090.3 |
| Solyc06g005100.2 | Solyc06g005100.3 |
| Solyc06g005110.2 | Solyc06g005110.3 |
| Solyc06g005130.2 | Solyc06g005130.3 |
| Solyc06g005140.2 | Solyc06g005140.3 |
| Solyc06g005150.2 | Solyc06g005150.3 |
| Solyc06g005170.2 | Solyc06g005170.3 |

| | |
|---|---|
| Solyc06g005200.2 | Solyc06g005200.3 |
| Solyc06g005260.2 | Solyc06g005260.3 |
| Solyc06g005290.2 | Solyc06g005290.3 |
| Solyc06g005310.2 | Solyc06g005310.3 |
| Solyc06g005330.2 | Solyc06g005330.3 |
| Solyc06g005350.2 | Solyc06g005350.3 |
| Solyc06g005360.2 | Solyc06g005360.3 |
| Solyc06g005370.2 | Solyc06g005370.3 |
| Solyc06g005380.2 | Solyc06g005380.3 |
| Solyc06g005410.2 | Solyc06g005410.3 |
| Solyc06g005480.2 | Solyc06g005480.3 |
| Solyc06g005490.2 | Solyc06g005490.3 |
| Solyc06g005530.1 | Solyc06g005530.2 |
| Solyc06g005620.2 | Solyc06g005620.3 |
| Solyc06g005710.2 | Solyc06g005710.3 |
| Solyc06g005750.2 | Solyc06g005750.3 |
| Solyc06g005795.1 | Solyc06g005795.2 |
| Solyc06g005790.2 | Solyc06g005790.3 |
| Solyc06g005800.2 | Solyc06g005800.3 |
| Solyc06g005810.2 | Solyc06g005810.3 |
| Solyc06g005820.2 | Solyc06g005820.3 |
| Solyc06g005850.2 | Solyc06g005850.3 |
| Solyc06g005890.2 | Solyc06g005890.3 |
| Solyc06g005940.2 | Solyc06g005940.3 |
| Solyc06g005950.2 | Solyc06g005950.3 |
| Solyc06g006000.2 | Solyc06g006000.3 |
| Solyc06g006020.2 | Solyc06g006020.3 |
| Solyc06g006080.2 | Solyc06g006080.3 |
| Solyc06g006100.2 | Solyc06g006100.3 |
| Solyc06g006110.2 | Solyc06g006110.3 |
| Solyc06g007110.2 | Solyc06g007110.3 |
| Solyc06g007120.2 | Solyc06g007120.3 |
| Solyc06g007130.4 | Solyc06g007130.5.1 |
| Solyc06g007140.3 | Solyc06g007140.4 |
| Solyc06g007180.2 | Solyc06g007180.3 |
| Solyc06g007210.2 | Solyc06g007210.3 |
| Solyc06g007230.1 | Solyc06g007230.2 |
| Solyc06g007300.1 | Solyc06g007300.2 |
| Solyc06g007320.2 | Solyc06g007320.3 |
| Solyc06g007330.2 | Solyc06g007330.3 |
| Solyc06g007340.2 | Solyc06g007340.3 |
| Solyc06g007355.1 | Solyc06g150100.1 |
| Solyc06g007360.2 | Solyc06g007360.3 |

| | |
|---|---|
| Solyc06g007450.1 | Solyc06g007450.2 |
| Solyc06g007460.2 | Solyc06g007460.3 |
| Solyc06g007470.2 | Solyc06g007470.3 |
| Solyc06g007480.2 | Solyc06g007480.3 |
| Solyc06g007510.2 | Solyc06g007510.3 |
| Solyc06g007540.2 | Solyc06g007540.3 |
| Solyc06g007570.2 | Solyc06g007570.3 |
| Solyc06g007590.2 | Solyc06g007590.3 |
| Solyc06g007620.2 | Solyc06g007620.3 |
| Solyc06g007640.2 | Solyc06g007640.3 |
| Solyc06g007650.1 | Solyc06g007650.2 |
| Solyc06g007660.1 | Solyc06g007660.2 |
| Solyc06g007770.2 | Solyc06g007770.3 |
| Solyc06g007780.2 | Solyc06g007780.3 |
| Solyc06g007850.2 | Solyc06g007850.3 |
| Solyc06g007870.2 | Solyc06g007870.3 |
| Solyc06g007890.2 | Solyc06g007890.3 |
| Solyc06g007930.2 | Solyc06g007930.3 |
| Solyc06g007970.2 | Solyc06g007970.3 |
| Solyc06g008010.2 | Solyc06g008010.3 |
| Solyc06g008020.2 | Solyc06g008020.3 |
| Solyc06g008030.2 | Solyc06g008030.3 |
| Solyc06g008040.2 | Solyc06g008040.3 |
| Solyc06g008080.1 | Solyc06g008080.2 |
| Solyc06g008090.2 | Solyc06g008090.3 |
| Solyc06g008100.2 | Solyc06g008100.3 |
| Solyc06g008150.2 | Solyc06g008150.3 |
| Solyc06g008170.2 | Solyc06g008170.3 |
| Solyc06g008195.1 | Solyc06g150101.1 |
| Solyc06g008200.2 | Solyc06g008200.3 |
| Solyc06g008230.2 | Solyc06g008230.3 |
| Solyc06g008240.2 | Solyc06g008240.3 |
| Solyc06g008250.2 | Solyc06g008250.3 |
| Solyc06g008260.2 | Solyc06g008260.3 |
| Solyc06g008310.2 | Solyc06g008310.3 |
| Solyc06g008330.1 | Solyc06g008330.2 |
| Solyc06g008350.2 | Solyc06g008350.3 |
| Solyc06g008360.2 | Solyc06g008360.3 |
| Solyc06g008490.2 | Solyc06g008490.3 |
| Solyc06g008500.2 | Solyc06g008500.3 |
| Solyc06g008510.2 | Solyc06g008510.3 |
| Solyc06g008580.2 | Solyc06g008580.3 |
| Solyc06g008590.2 | Solyc06g008590.3 |

| | |
|---|---|
| Solyc06g008600.2 | Solyc06g008600.3 |
| Solyc06g008610.2 | Solyc06g008610.3 |
| Solyc06g008660.2 | Solyc06g008660.3 |
| Solyc06g008663.1 | Solyc06g150102.1 |
| Solyc06g008680.2 | Solyc06g008680.3 |
| Solyc06g008700.2 | Solyc06g008700.3 |
| Solyc06g008730.2 | Solyc06g008730.3 |
| Solyc06g008740.2 | Solyc06g008740.3 |
| Solyc06g008810.2 | Solyc06g008810.3 |
| Solyc06g008820.2 | Solyc06g008820.3 |
| Solyc06g008840.2 | Solyc06g008840.3 |
| Solyc06g008850.2 | Solyc06g008850.3 |
| Solyc06g008860.2 | Solyc06g008860.3 |
| Solyc06g008880.2 | Solyc06g008880.3 |
| Solyc06g008920.2 | Solyc06g008920.3 |
| Solyc06g008930.2 | Solyc06g008930.3 |
| Solyc06g008950.1 | Solyc06g008950.2 |
| Solyc06g008960.2 | Solyc06g008960.3 |
| Solyc06g008980.2 | Solyc06g008980.3 |
| Solyc06g009050.2 | Solyc06g009050.3 |
| Solyc06g009070.2 | Solyc06g009070.3 |
| Solyc06g009080.1 | Solyc06g009080.2 |
| Solyc06g009110.2 | Solyc06g009110.3 |
| Solyc06g009130.2 | Solyc06g009130.3 |
| Solyc06g009140.2 | Solyc06g009140.3 |
| Solyc06g009170.2 | Solyc06g009170.3 |
| Solyc06g009200.2 | Solyc06g009200.3 |
| Solyc06g009250.1 | Solyc06g009250.2 |
| Solyc06g009260.2 | Solyc06g009260.3 |
| Solyc06g009270.2 | Solyc06g009270.3 |
| Solyc06g009330.2 | Solyc06g009330.3 |
| Solyc06g009390.2 | Solyc06g009390.3 |
| Solyc06g009400.2 | Solyc06g009400.3 |
| Solyc06g009490.2 | Solyc06g009490.3 |
| Solyc06g009520.2 | Solyc06g009520.3 |
| Solyc06g009530.2 | Solyc06g009530.3 |
| Solyc06g009600.1 | Solyc06g009600.2 |
| Solyc06g009640.1 | Solyc06g009640.2 |
| Solyc06g009650.2 | Solyc06g009650.3 |
| Solyc06g009660.2 | Solyc06g009660.3 |
| Solyc06g009670.2 | Solyc06g009670.3 |
| Solyc06g009693.1 | Solyc06g150103.1 |
| Solyc06g009697.1 | Solyc06g150104.1 |

| | |
|---|---|
| Solyc06g009820.2 | Solyc06g009820.3 |
| Solyc06g009830.2 | Solyc06g009830.3 |
| Solyc06g009840.2 | Solyc06g009840.3 |
| Solyc06g009890.1 | Solyc06g009890.2 |
| Solyc06g009920.1 | Solyc06g009920.2 |
| Solyc06g009970.2 | Solyc06g009970.3 |
| Solyc06g010000.2 | Solyc06g010000.3 |
| Solyc06g010040.2 | Solyc06g010040.3 |
| Solyc06g010065.1 | Solyc06g150105.1 |
| Solyc06g010100.1 | Solyc06g010100.2 |
| Solyc06g010160.1 | Solyc06g010160.2 |
| Solyc06g010170.2 | Solyc06g010170.3 |
| Solyc06g010190.1 | Solyc06g010190.2 |
| Solyc06g010250.2 | Solyc06g010250.3 |
| Solyc06g010260.2 | Solyc06g010260.3 |
| Solyc06g011280.2 | Solyc06g011280.3 |
| Solyc06g011320.1 | Solyc06g011320.2 |
| Solyc06g011350.2 | Solyc06g011350.3 |
| Solyc06g011530.2 | Solyc06g011530.3 |
| Solyc06g011570.2 | Solyc06g011570.3 |
| Solyc06g011663.1 | Solyc06g150106.1 |
| Solyc06g011667.1 | Solyc06g150107.1 |
| Solyc06g016750.2 | Solyc06g016750.3 |
| Solyc06g016770.2 | Solyc06g016770.3 |
| Solyc06g017920.1 | Solyc06g017920.2 |
| Solyc06g018110.2 | Solyc06g018110.3 |
| Solyc06g019170.2 | Solyc06g019170.3 |
| Solyc06g030490.2 | Solyc06g030490.3 |
| Solyc06g030640.1 | Solyc06g030640.2 |
| Solyc06g031710.2 | Solyc06g031710.3 |
| Solyc06g052090.1 | Solyc06g052090.2 |
| Solyc06g033820.1 | Solyc06g033820.2 |
| Solyc06g033850.2 | Solyc06g033850.3 |
| Solyc06g033860.1 | Solyc06g033860.2 |
| Solyc06g033910.1 | Solyc06g033910.2 |
| Solyc06g033960.1 | Solyc06g033960.2 |
| Solyc06g034000.1 | Solyc06g034000.2 |
| Solyc06g034020.2 | Solyc06g034020.3 |
| Solyc06g034030.2 | Solyc06g034030.3 |
| Solyc06g034110.2 | Solyc06g034110.3 |
| Solyc06g034160.2 | Solyc06g034160.3 |
| Solyc06g034180.1 | Solyc06g034180.2 |
| Solyc06g034220.2 | Solyc06g034220.3 |

| | |
|---|---|
| Solyc06g034230.1 | Solyc06g034230.2 |
| Solyc06g034290.2 | Solyc06g034290.3 |
| Solyc06g034350.1 | Solyc06g034350.2 |
| Solyc06g034360.1 | Solyc06g034360.2 |
| Solyc06g034410.2 | Solyc06g034410.3 |
| Solyc06g035410.1 | Solyc06g035410.2 |
| Solyc06g035430.2 | Solyc06g035430.3 |
| Solyc06g035440.2 | Solyc06g035440.3 |
| Solyc06g035455.1 | Solyc06g150108.1 |
| Solyc06g035490.2 | Solyc06g035490.3 |
| Solyc06g035520.2 | Solyc06g035520.3 |
| Solyc06g035530.2 | Solyc06g035530.3 |
| Solyc06g035550.2 | Solyc06g035550.3 |
| Solyc06g035555.1 | Solyc06g150109.1 |
| Solyc06g035580.2 | Solyc06g035580.3 |
| Solyc06g035600.2 | Solyc06g035600.3 |
| Solyc06g035610.2 | Solyc06g035610.3 |
| Solyc06g035620.2 | Solyc06g035620.3 |
| Solyc06g035690.2 | Solyc06g035690.3 |
| Solyc06g035720.2 | Solyc06g035720.3 |
| Solyc06g035790.2 | Solyc06g035790.3 |
| Solyc06g035870.2 | Solyc06g035870.3 |
| Solyc06g035920.2 | Solyc06g035920.3 |
| Solyc06g035940.2 | Solyc06g035940.3 |
| Solyc06g035960.2 | Solyc06g035960.3 |
| Solyc06g035970.2 | Solyc06g035970.3 |
| Solyc06g036070.2 | Solyc06g036070.3 |
| Solyc06g036100.2 | Solyc06g036100.3 |
| Solyc06g036120.2 | Solyc06g036120.3 |
| Solyc06g036180.2 | Solyc06g036180.3 |
| Solyc06g036200.1 | Solyc06g036200.2 |
| Solyc06g036240.1 | Solyc06g036240.2 |
| Solyc06g036260.2 | Solyc06g036260.3 |
| Solyc06g036290.2 | Solyc06g036290.3 |
| Solyc06g036310.2 | Solyc06g036310.3 |
| Solyc06g036350.2 | Solyc06g036350.3 |
| Solyc06g036430.2 | Solyc06g036430.3 |
| Solyc06g036440.1 | Solyc06g036440.2 |
| Solyc06g036480.1 | Solyc06g036480.2 |
| Solyc06g036690.1 | Solyc06g036690.2 |
| Solyc06g043260.1 | Solyc06g043260.2 |
| Solyc06g043113.1 | Solyc06g150110.1 |
| Solyc06g043117.1 | Solyc06g150111.1 |

| | |
|---|---|
| Solyc06g048373.1 | Solyc06g150112.1 |
| Solyc06g048377.1 | Solyc06g150113.1 |
| Solyc06g048420.1 | Solyc06g048420.2 |
| Solyc06g048440.2 | Solyc06g048440.3 |
| Solyc06g048460.1 | Solyc06g048460.2 |
| Solyc06g048480.2 | Solyc06g048480.3 |
| Solyc06g048520.2 | Solyc06g048520.3 |
| Solyc06g048540.2 | Solyc06g048540.3 |
| Solyc06g048560.1 | Solyc06g048560.2 |
| Solyc06g048570.2 | Solyc06g048570.3 |
| Solyc06g048590.2 | Solyc06g048590.3 |
| Solyc06g048600.2 | Solyc06g048600.3 |
| Solyc06g048610.2 | Solyc06g048610.3 |
| Solyc06g048620.2 | Solyc06g048620.3 |
| Solyc06g048630.2 | Solyc06g048630.3 |
| Solyc06g048687.1 | Solyc06g150114.1 |
| Solyc06g048710.1 | Solyc06g048710.2 |
| Solyc06g048717.1 | Solyc06g150115.1 |
| Solyc06g048730.2 | Solyc06g048730.3 |
| Solyc06g048750.2 | Solyc06g048750.3 |
| Solyc06g048770.2 | Solyc06g048770.3 |
| Solyc06g048840.2 | Solyc06g048840.3 |
| Solyc06g048910.1 | Solyc06g048910.2 |
| Solyc06g048950.2 | Solyc06g048950.3 |
| Solyc06g048970.2 | Solyc06g048970.3 |
| Solyc06g049010.1 | Solyc06g049010.2 |
| Solyc06g049050.2 | Solyc06g049050.3 |
| Solyc06g049070.2 | Solyc06g049070.3 |
| Solyc06g049080.2 | Solyc06g049080.3 |
| Solyc06g049090.2 | Solyc06g049090.3 |
| Solyc06g050110.1 | Solyc06g050110.2 |
| Solyc06g050120.2 | Solyc06g050120.3 |
| Solyc06g050130.2 | Solyc06g050130.3 |
| Solyc06g050200.1 | Solyc06g050200.2 |
| Solyc06g050210.2 | Solyc06g050210.3 |
| Solyc06g050310.2 | Solyc06g050310.3 |
| Solyc06g050340.2 | Solyc06g050340.3 |
| Solyc06g050350.2 | Solyc06g050350.3 |
| Solyc06g050360.2 | Solyc06g050360.3 |
| Solyc06g050430.2 | Solyc06g050430.3 |
| Solyc06g050440.2 | Solyc06g050440.3 |
| Solyc06g050460.1 | Solyc06g050460.2 |
| Solyc06g050510.2 | Solyc06g050510.3 |

| | |
|---|---|
| Solyc06g050530.2 | Solyc06g050530.3 |
| Solyc06g050560.2 | Solyc06g050560.3 |
| Solyc06g050590.2 | Solyc06g050590.3 |
| Solyc06g050620.2 | Solyc06g050620.3 |
| Solyc06g050630.2 | Solyc06g050630.3 |
| Solyc06g050645.1 | Solyc06g150116.1 |
| Solyc06g050690.2 | Solyc06g050690.3 |
| Solyc06g050720.2 | Solyc06g050720.3 |
| Solyc06g050770.2 | Solyc06g050770.3 |
| Solyc06g050790.2 | Solyc06g050790.3 |
| Solyc06g050810.2 | Solyc06g050810.3 |
| Solyc06g050830.1 | Solyc06g050830.2 |
| Solyc06g050833.1 | Solyc06g150117.1 |
| Solyc06g050840.2 | Solyc06g050840.3 |
| Solyc06g050860.2 | Solyc06g050860.3 |
| Solyc06g050870.2 | Solyc06g050870.3 |
| Solyc06g050880.2 | Solyc06g050880.3 |
| Solyc06g050900.2 | Solyc06g050900.3 |
| Solyc06g050930.2 | Solyc06g050930.3 |
| Solyc06g050950.2 | Solyc06g050950.3 |
| Solyc06g050980.2 | Solyc06g050980.3 |
| Solyc06g050990.2 | Solyc06g050990.3 |
| Solyc06g051030.2 | Solyc06g051030.3 |
| Solyc06g051070.1 | Solyc06g051070.2 |
| Solyc06g051200.2 | Solyc06g051200.3 |
| Solyc06g051210.2 | Solyc06g051210.3 |
| Solyc06g051230.2 | Solyc06g051230.4 |
| Solyc06g051270.2 | Solyc06g051270.3 |
| Solyc06g051300.2 | Solyc06g051300.3 |
| Solyc06g051310.2 | Solyc06g051310.3 |
| Solyc06g051320.2 | Solyc06g051320.3 |
| Solyc06g051360.2 | Solyc06g051360.3 |
| Solyc06g051400.2 | Solyc06g051400.3 |
| Solyc06g051420.2 | Solyc06g051420.3 |
| Solyc06g051450.1 | Solyc06g051450.2 |
| Solyc06g051520.2 | Solyc06g051520.3 |
| Solyc06g051600.1 | Solyc06g051600.2 |
| Solyc06g051630.4 | Solyc06g051630.5.1 |
| Solyc06g051720.2 | Solyc06g051720.3 |
| Solyc06g051750.2 | Solyc06g051750.3 |
| Solyc06g051760.2 | Solyc06g051760.3 |
| Solyc06g051800.2 | Solyc06g051800.3 |
| Solyc06g084770.1 | Solyc06g084770.2 |

| | |
|---|---|
| Solyc06g051900.2 | Solyc06g051900.3 |
| Solyc06g051930.2 | Solyc06g051930.3 |
| Solyc06g051960.2 | Solyc06g051960.3 |
| Solyc06g051970.2 | Solyc06g051970.3 |
| Solyc06g052010.1 | Solyc06g052010.2 |
| Solyc06g052030.2 | Solyc06g052030.3 |
| Solyc06g052070.2 | Solyc06g052070.3 |
| Solyc06g052130.1 | Solyc06g052130.2 |
| Solyc06g053140.2 | Solyc06g053140.3 |
| Solyc06g053160.2 | Solyc06g053160.3 |
| Solyc06g053190.2 | Solyc06g053190.3 |
| Solyc06g053200.2 | Solyc06g053200.3 |
| Solyc06g053220.2 | Solyc06g053220.3 |
| Solyc06g053250.1 | Solyc06g053250.2 |
| Solyc06g053300.2 | Solyc06g053300.3 |
| Solyc06g053310.2 | Solyc06g053310.3 |
| Solyc06g053340.2 | Solyc06g053340.3 |
| Solyc06g053350.2 | Solyc06g053350.3 |
| Solyc06g053380.2 | Solyc06g053380.3 |
| Solyc06g053385.1 | Solyc06g053385.2 |
| Solyc06g053400.2 | Solyc06g053400.3 |
| Solyc06g053470.2 | Solyc06g053470.3 |
| Solyc06g053480.2 | Solyc06g053480.3 |
| Solyc06g053510.2 | Solyc06g053510.3 |
| Solyc06g053530.2 | Solyc06g053530.3 |
| Solyc06g053540.2 | Solyc06g053540.3 |
| Solyc06g053550.2 | Solyc06g053550.3 |
| Solyc06g053610.2 | Solyc06g053610.3 |
| Solyc06g053630.2 | Solyc06g053630.3 |
| Solyc06g053660.2 | Solyc06g053660.3 |
| Solyc06g053710.2 | Solyc06g053710.3 |
| Solyc06g053750.2 | Solyc06g053750.3 |
| Solyc06g053760.2 | Solyc06g053760.3 |
| Solyc06g053790.2 | Solyc06g053790.3 |
| Solyc06g053805.1 | Solyc06g150118.1 |
| Solyc06g053810.2 | Solyc06g053810.3 |
| Solyc06g053830.2 | Solyc06g053830.3 |
| Solyc06g053840.2 | Solyc06g053840.3 |
| Solyc06g053860.2 | Solyc06g053860.3 |
| Solyc06g053930.2 | Solyc06g053930.3 |
| Solyc06g053960.2 | Solyc06g053960.3 |
| Solyc06g053980.2 | Solyc06g053980.3 |
| Solyc06g054010.2 | Solyc06g054010.3 |

| | |
|---|---|
| Solyc06g054020.2 | Solyc06g054020.3 |
| Solyc06g054070.2 | Solyc06g054070.3 |
| Solyc06g054080.2 | Solyc06g054080.3 |
| Solyc06g054090.1 | Solyc06g054090.2 |
| Solyc06g054100.1 | Solyc06g054100.2 |
| Solyc06g054120.1 | Solyc06g054120.2 |
| Solyc06g054160.1 | Solyc06g054160.2 |
| Solyc06g054240.2 | Solyc06g054240.3 |
| Solyc06g054270.2 | Solyc06g054270.3 |
| Solyc06g054313.1 | Solyc06g150119.1 |
| Solyc06g054317.1 | Solyc06g150120.1 |
| Solyc06g054330.2 | Solyc06g054330.3 |
| Solyc06g054360.1 | Solyc06g054360.2 |
| Solyc06g054370.1 | Solyc06g054370.2 |
| Solyc06g054390.2 | Solyc06g054390.3 |
| Solyc06g054400.2 | Solyc06g054400.3 |
| Solyc06g054420.2 | Solyc06g054420.3 |
| Solyc06g054480.2 | Solyc06g054480.3 |
| Solyc06g054490.2 | Solyc06g054490.3 |
| Solyc06g054520.1 | Solyc06g054520.2 |
| Solyc06g054530.2 | Solyc06g054530.3 |
| Solyc06g054540.2 | Solyc06g054540.3 |
| Solyc06g054550.2 | Solyc06g054550.3 |
| Solyc06g054560.2 | Solyc06g054560.3 |
| Solyc06g054610.1 | Solyc06g054610.2 |
| Solyc06g054640.1 | Solyc06g054640.2 |
| Solyc06g054650.2 | Solyc06g054650.3 |
| Solyc06g054670.1 | Solyc06g054670.2 |
| Solyc06g054680.1 | Solyc06g054680.2 |
| Solyc06g059710.2 | Solyc06g059710.3 |
| Solyc06g059720.2 | Solyc06g059720.3 |
| Solyc06g059750.2 | Solyc06g059750.3 |
| Solyc06g059760.2 | Solyc06g059760.3 |
| Solyc06g059770.1 | Solyc06g059770.2 |
| Solyc06g059800.2 | Solyc06g059800.3 |
| Solyc06g059850.1 | Solyc06g059850.2 |
| Solyc06g059880.2 | Solyc06g059880.3 |
| Solyc06g059980.2 | Solyc06g059980.3 |
| Solyc06g059995.1 | Solyc06g150121.1 |
| Solyc06g060070.2 | Solyc06g060070.3 |
| Solyc06g060090.2 | Solyc06g060090.3 |
| Solyc06g060100.2 | Solyc06g060100.3 |
| Solyc06g060110.2 | Solyc06g060110.3 |

| | |
|---|---|
| Solyc06g060120.2 | Solyc06g060120.3 |
| Solyc06g060150.2 | Solyc06g060150.3 |
| Solyc06g060170.2 | Solyc06g060170.3 |
| Solyc06g060190.2 | Solyc06g060190.3 |
| Solyc06g060200.1 | Solyc06g060200.2 |
| Solyc06g060230.2 | Solyc06g060230.3 |
| Solyc06g060250.2 | Solyc06g060250.3 |
| Solyc06g060260.2 | Solyc06g060260.3 |
| Solyc06g060310.2 | Solyc06g060310.3 |
| Solyc06g060340.2 | Solyc06g060340.3 |
| Solyc06g060360.2 | Solyc06g060360.3 |
| Solyc06g060370.2 | Solyc06g060370.3 |
| Solyc06g060380.2 | Solyc06g060380.3 |
| Solyc06g060390.2 | Solyc06g060390.3 |
| Solyc06g060400.2 | Solyc06g060400.3 |
| Solyc06g060410.2 | Solyc06g060410.3 |
| Solyc06g060420.2 | Solyc06g060420.3 |
| Solyc06g060425.1 | Solyc06g150122.1 |
| Solyc06g060440.1 | Solyc06g060440.2 |
| Solyc06g060450.2 | Solyc06g060450.3 |
| Solyc06g060490.2 | Solyc06g060490.3 |
| Solyc06g060510.1 | Solyc06g060510.2 |
| Solyc06g060540.2 | Solyc06g060540.3 |
| Solyc06g060550.2 | Solyc06g060550.3 |
| Solyc06g060570.2 | Solyc06g060570.3 |
| Solyc06g060590.2 | Solyc06g060590.3 |
| Solyc06g060680.1 | Solyc06g060680.2 |
| Solyc06g060690.2 | Solyc06g060690.3 |
| Solyc06g060720.2 | Solyc06g060720.3 |
| Solyc06g060730.2 | Solyc06g060730.3 |
| Solyc06g060750.2 | Solyc06g060750.3 |
| Solyc06g060760.2 | Solyc06g060760.3 |
| Solyc06g060810.2 | Solyc06g060810.3 |
| Solyc06g060830.2 | Solyc06g060830.3 |
| Solyc06g060850.2 | Solyc06g060850.3 |
| Solyc06g060910.1 | Solyc06g060910.2 |
| Solyc06g060940.1 | Solyc06g060940.2 |
| Solyc06g060970.1 | Solyc06g060970.2 |
| Solyc06g061000.2 | Solyc06g061000.3 |
| Solyc06g061020.2 | Solyc06g061020.3 |
| Solyc06g061030.2 | Solyc06g061030.3 |
| Solyc06g061070.2 | Solyc06g061070.3 |
| Solyc06g061080.2 | Solyc06g061080.3 |

| | |
|---|---|
| Solyc06g061090.2 | Solyc06g061090.3 |
| Solyc06g061110.1 | Solyc06g061110.2 |
| Solyc06g061150.2 | Solyc06g061150.3 |
| Solyc06g061170.2 | Solyc06g061170.3 |
| Solyc06g061180.1 | Solyc06g061180.2 |
| Solyc06g061220.1 | Solyc06g061220.2 |
| Solyc06g061230.2 | Solyc06g061230.3 |
| Solyc06g061260.1 | Solyc06g061260.2 |
| Solyc06g061270.1 | Solyc06g061270.2 |
| Solyc06g061280.2 | Solyc06g061280.3 |
| Solyc06g062300.1 | Solyc06g062300.2 |
| Solyc06g062310.2 | Solyc06g062310.3 |
| Solyc06g062340.2 | Solyc06g062340.3 |
| Solyc06g062360.2 | Solyc06g062360.3 |
| Solyc06g062380.2 | Solyc06g062380.3 |
| Solyc06g062390.2 | Solyc06g062390.3 |
| Solyc06g062400.1 | Solyc06g062400.2 |
| Solyc06g062430.2 | Solyc06g062430.3 |
| Solyc06g062450.2 | Solyc06g062450.3 |
| Solyc06g062460.2 | Solyc06g062460.3 |
| Solyc06g062490.2 | Solyc06g062490.3 |
| Solyc06g062510.2 | Solyc06g062510.3 |
| Solyc06g062560.1 | Solyc06g062560.2 |
| Solyc06g062580.2 | Solyc06g062580.3 |
| Solyc06g062600.2 | Solyc06g062600.3 |
| Solyc06g062610.1 | Solyc06g062610.2 |
| Solyc06g062670.2 | Solyc06g062670.3 |
| Solyc06g062700.2 | Solyc06g062700.3 |
| Solyc06g062720.1 | Solyc06g062720.2 |
| Solyc06g062730.1 | Solyc06g062730.2 |
| Solyc06g062740.1 | Solyc06g062740.2 |
| Solyc06g062750.2 | Solyc06g062750.3 |
| Solyc06g062760.2 | Solyc06g062760.3 |
| Solyc06g062790.2 | Solyc06g062790.3 |
| Solyc06g062805.1 | Solyc06g150123.1 |
| Solyc06g062850.2 | Solyc06g062850.3 |
| Solyc06g062880.1 | Solyc06g062880.2 |
| Solyc06g062920.2 | Solyc06g062920.3 |
| Solyc06g062980.2 | Solyc06g062980.3 |
| Solyc06g063000.2 | Solyc06g063000.3 |
| Solyc06g063030.2 | Solyc06g063030.3 |
| Solyc06g063060.2 | Solyc06g063060.3 |
| Solyc06g063070.2 | Solyc06g063070.3 |

| | |
|---|---|
| Solyc06g063085.1 | Solyc06g150124.1 |
| Solyc06g063100.2 | Solyc06g063100.3 |
| Solyc06g063170.2 | Solyc06g063170.3 |
| Solyc06g063190.2 | Solyc06g063190.3 |
| Solyc06g063200.1 | Solyc06g063200.2 |
| Solyc06g063210.2 | Solyc06g063210.3 |
| Solyc06g063220.2 | Solyc06g063220.3 |
| Solyc06g063240.2 | Solyc06g063240.3 |
| Solyc06g063250.2 | Solyc06g063250.3 |
| Solyc06g063290.2 | Solyc06g063290.3 |
| Solyc06g063330.2 | Solyc06g063330.3 |
| Solyc06g063340.2 | Solyc06g063340.3 |
| Solyc06g063345.1 | Solyc06g150125.1 |
| Solyc06g063390.2 | Solyc06g063390.3 |
| Solyc06g063400.2 | Solyc06g063400.3 |
| Solyc06g063430.1 | Solyc06g063430.2 |
| Solyc06g064440.2 | Solyc06g064440.3 |
| Solyc06g064450.2 | Solyc06g064450.3 |
| Solyc06g064500.2 | Solyc06g064500.3 |
| Solyc06g064525.1 | Solyc06g150126.1 |
| Solyc06g064550.2 | Solyc06g064550.3 |
| Solyc06g064570.2 | Solyc06g064570.3 |
| Solyc06g064580.1 | Solyc06g064580.2 |
| Solyc06g064620.2 | Solyc06g064620.3 |
| Solyc06g064630.2 | Solyc06g064630.3 |
| Solyc06g064640.2 | Solyc06g064640.3 |
| Solyc06g064690.1 | Solyc06g064690.2 |
| Solyc06g064760.1 | Solyc06g064760.2 |
| Solyc06g064800.1 | Solyc06g064800.2 |
| Solyc06g064820.2 | Solyc06g064820.3 |
| Solyc06g064830.2 | Solyc06g064830.3 |
| Solyc06g064850.1 | Solyc06g064850.2 |
| Solyc06g064880.2 | Solyc06g064880.3 |
| Solyc06g064900.2 | Solyc06g064900.3 |
| Solyc06g064910.2 | Solyc06g064910.3 |
| Solyc06g064940.2 | Solyc06g064940.3 |
| Solyc06g064960.2 | Solyc06g064960.3 |
| Solyc06g064980.2 | Solyc06g064980.3 |
| Solyc06g065020.2 | Solyc06g065020.3 |
| Solyc06g065030.2 | Solyc06g065030.3 |
| Solyc06g065050.1 | Solyc06g065050.2 |
| Solyc06g065100.2 | Solyc06g065100.3 |
| Solyc06g065180.2 | Solyc06g065180.3 |

| | |
|---|---|
| Solyc06g065210.2 | Solyc06g065210.4 |
| Solyc06g065230.2 | Solyc06g065230.3 |
| Solyc06g065240.2 | Solyc06g065240.3 |
| Solyc06g065260.2 | Solyc06g065260.3 |
| Solyc06g065310.2 | Solyc06g065310.3 |
| Solyc06g065335.1 | Solyc06g150127.1 |
| Solyc06g065360.2 | Solyc06g065360.3 |
| Solyc06g065370.2 | Solyc06g065370.3 |
| Solyc06g065400.2 | Solyc06g065400.3 |
| Solyc06g065410.2 | Solyc06g065410.3 |
| Solyc06g065420.1 | Solyc06g065420.2 |
| Solyc06g065430.2 | Solyc06g065430.3 |
| Solyc06g065450.2 | Solyc06g065450.3 |
| Solyc06g065490.2 | Solyc06g065490.3 |
| Solyc06g065520.2 | Solyc06g065520.3 |
| Solyc06g065540.2 | Solyc06g065540.3 |
| Solyc06g065550.1 | Solyc06g065550.2 |
| Solyc06g065560.1 | Solyc06g065560.2 |
| Solyc06g065570.2 | Solyc06g065570.3 |
| Solyc06g065580.2 | Solyc06g065580.3 |
| Solyc06g065610.2 | Solyc06g065610.3 |
| Solyc06g065620.1 | Solyc06g065620.2 |
| Solyc06g065630.2 | Solyc06g065630.3 |
| Solyc06g065640.2 | Solyc06g065640.3 |
| Solyc06g065650.2 | Solyc06g065650.3 |
| Solyc06g065680.2 | Solyc06g065680.3 |
| Solyc06g065690.2 | Solyc06g065690.3 |
| Solyc06g065700.2 | Solyc06g065700.3 |
| Solyc06g065710.2 | Solyc06g065710.3 |
| Solyc06g065730.2 | Solyc06g065730.3 |
| Solyc06g065800.2 | Solyc06g065800.3 |
| Solyc06g065820.2 | Solyc06g065820.3 |
| Solyc06g065840.2 | Solyc06g065840.3 |
| Solyc06g065850.2 | Solyc06g065850.3 |
| Solyc06g065910.2 | Solyc06g065910.3 |
| Solyc06g065980.2 | Solyc06g065980.3 |
| Solyc06g066010.2 | Solyc06g066010.3 |
| Solyc06g066020.2 | Solyc06g066020.3 |
| Solyc06g066040.1 | Solyc06g066040.2 |
| Solyc06g066050.2 | Solyc06g066050.3 |
| Solyc06g066060.2 | Solyc06g066060.3 |
| Solyc06g066090.2 | Solyc06g066090.3 |
| Solyc06g066100.2 | Solyc06g066100.3 |

| | |
|---|---|
| Solyc06g066120.2 | Solyc06g066120.3 |
| Solyc06g066160.2 | Solyc06g066160.3 |
| Solyc06g066190.2 | Solyc06g066190.3 |
| Solyc06g066200.1 | Solyc06g066200.2 |
| Solyc06g084820.1 | Solyc06g084820.2 |
| Solyc06g066310.2 | Solyc06g066310.3 |
| Solyc06g066320.2 | Solyc06g066320.3 |
| Solyc06g066330.2 | Solyc06g066330.3 |
| Solyc06g066400.2 | Solyc06g066400.3 |
| Solyc06g066410.2 | Solyc06g066410.3 |
| Solyc06g066430.2 | Solyc06g066430.3 |
| Solyc06g066440.2 | Solyc06g066440.3 |
| Solyc06g066460.2 | Solyc06g066460.3 |
| Solyc06g066550.2 | Solyc06g066550.3 |
| Solyc06g066600.2 | Solyc06g066600.3 |
| Solyc06g066640.2 | Solyc06g066640.3 |
| Solyc06g066680.2 | Solyc06g066680.3 |
| Solyc06g066730.2 | Solyc06g066730.3 |
| Solyc06g066750.2 | Solyc06g066750.3 |
| Solyc06g066760.2 | Solyc06g066760.3 |
| Solyc06g066780.2 | Solyc06g066780.3 |
| Solyc06g066790.2 | Solyc06g066790.3 |
| Solyc06g066810.2 | Solyc06g066810.3 |
| Solyc06g066840.2 | Solyc06g066840.3 |
| Solyc06g066860.1 | Solyc06g066860.2 |
| Solyc06g067860.2 | Solyc06g067860.3 |
| Solyc06g067870.2 | Solyc06g067870.3 |
| Solyc06g067880.2 | Solyc06g067880.3 |
| Solyc06g067890.2 | Solyc06g067890.3 |
| Solyc06g067910.2 | Solyc06g067910.3 |
| Solyc06g067920.2 | Solyc06g067920.3 |
| Solyc06g067930.1 | Solyc06g067930.2 |
| Solyc06g067950.2 | Solyc06g067950.3 |
| Solyc06g067960.2 | Solyc06g067960.3 |
| Solyc06g067980.2 | Solyc06g067980.3 |
| Solyc06g067990.2 | Solyc06g067990.3 |
| Solyc06g068000.1 | Solyc06g068000.2 |
| Solyc06g068010.2 | Solyc06g068010.3 |
| Solyc06g068030.2 | Solyc06g068030.3 |
| Solyc06g068040.2 | Solyc06g068040.3 |
| Solyc06g068050.2 | Solyc06g068050.3 |
| Solyc06g068070.1 | Solyc06g068070.2 |
| Solyc06g068090.2 | Solyc06g068090.3 |

| | |
|---|---|
| Solyc06g068130.2 | Solyc06g068130.3 |
| Solyc06g068140.2 | Solyc06g068140.3 |
| Solyc06g068150.2 | Solyc06g068150.3 |
| Solyc06g068160.2 | Solyc06g068160.3 |
| Solyc06g068170.1 | Solyc06g068170.2 |
| Solyc06g068190.1 | Solyc06g068190.2 |
| Solyc06g068215.1 | Solyc06g150129.1 |
| Solyc06g068220.2 | Solyc06g068220.3 |
| Solyc06g068240.2 | Solyc06g068240.3 |
| Solyc06g068270.2 | Solyc06g068270.3 |
| Solyc06g068280.2 | Solyc06g068280.3 |
| Solyc06g068320.2 | Solyc06g068320.3 |
| Solyc06g068350.1 | Solyc06g068350.2 |
| Solyc06g068360.2 | Solyc06g068360.3 |
| Solyc06g068370.2 | Solyc06g068370.3 |
| Solyc06g068400.2 | Solyc06g068400.3 |
| Solyc06g068410.2 | Solyc06g068410.3 |
| Solyc06g068460.2 | Solyc06g068460.3 |
| Solyc06g068470.2 | Solyc06g068470.3 |
| Solyc06g068480.2 | Solyc06g068480.3 |
| Solyc06g068490.2 | Solyc06g068490.3 |
| Solyc06g068500.2 | Solyc06g068500.3 |
| Solyc06g068520.2 | Solyc06g068520.3 |
| Solyc06g068540.2 | Solyc06g068540.3 |
| Solyc06g068560.2 | Solyc06g068560.3 |
| Solyc06g068600.2 | Solyc06g068600.3 |
| Solyc06g068670.2 | Solyc06g068670.3 |
| Solyc06g068680.2 | Solyc06g068680.3 |
| Solyc06g068710.2 | Solyc06g068710.3 |
| Solyc06g068720.2 | Solyc06g068720.3 |
| Solyc06g068750.2 | Solyc06g068750.3 |
| Solyc06g068760.2 | Solyc06g068760.3 |
| Solyc06g068770.2 | Solyc06g068770.3 |
| Solyc06g068780.2 | Solyc06g068780.3 |
| Solyc06g068800.2 | Solyc06g068800.3 |
| Solyc06g068820.2 | Solyc06g068820.3 |
| Solyc06g068830.1 | Solyc06g068830.2 |
| Solyc06g068840.2 | Solyc06g068840.3 |
| Solyc06g068860.2 | Solyc06g068860.3 |
| Solyc06g068870.2 | Solyc06g068870.3 |
| Solyc06g068880.2 | Solyc06g068880.3 |
| Solyc06g068910.2 | Solyc06g068910.3 |
| Solyc06g068920.2 | Solyc06g068920.3 |

| | |
|---|---|
| Solyc06g068930.1 | Solyc06g068930.2 |
| Solyc06g068970.2 | Solyc06g068970.3 |
| Solyc06g069010.2 | Solyc06g069010.3 |
| Solyc06g069080.2 | Solyc06g069080.3 |
| Solyc06g069090.2 | Solyc06g069090.3 |
| Solyc06g069120.2 | Solyc06g069120.3 |
| Solyc06g069180.2 | Solyc06g069180.3 |
| Solyc06g069210.2 | Solyc06g069210.3 |
| Solyc06g069230.2 | Solyc06g069230.3 |
| Solyc06g069240.1 | Solyc06g069240.2 |
| Solyc06g069250.1 | Solyc06g069250.2 |
| Solyc06g069290.2 | Solyc06g069290.3 |
| Solyc06g069310.2 | Solyc06g069310.3 |
| Solyc06g069320.3 | Solyc06g069320.4 |
| Solyc06g069330.2 | Solyc06g069330.3 |
| Solyc06g069360.2 | Solyc06g069360.3 |
| Solyc06g069385.1 | Solyc06g150130.1 |
| Solyc06g069410.2 | Solyc06g069410.3 |
| Solyc06g069420.2 | Solyc06g069420.3 |
| Solyc06g069430.2 | Solyc06g069430.3 |
| Solyc06g069440.2 | Solyc06g069440.3 |
| Solyc06g069470.2 | Solyc06g069470.3 |
| Solyc06g069480.2 | Solyc06g069480.3 |
| Solyc06g069490.2 | Solyc06g069490.3 |
| Solyc06g069500.2 | Solyc06g069500.3 |
| Solyc06g069520.2 | Solyc06g069520.3 |
| Solyc06g069530.2 | Solyc06g069530.3 |
| Solyc06g069565.1 | Solyc06g069565.2 |
| Solyc06g069570.2 | Solyc06g069570.3 |
| Solyc06g069580.2 | Solyc06g069580.3 |
| Solyc06g069590.2 | Solyc06g069590.3 |
| Solyc06g069640.2 | Solyc06g069640.3 |
| Solyc06g069660.1 | Solyc06g069660.2 |
| Solyc06g069710.2 | Solyc06g069710.3 |
| Solyc06g069730.2 | Solyc06g069730.3 |
| Solyc06g069760.2 | Solyc06g069760.3 |
| Solyc06g069770.2 | Solyc06g069770.3 |
| Solyc06g069800.1 | Solyc06g069800.2 |
| Solyc06g069850.2 | Solyc06g069850.3 |
| Solyc06g069870.2 | Solyc06g069870.3 |
| Solyc06g069900.2 | Solyc06g069900.3 |
| Solyc06g070900.2 | Solyc06g070900.3 |
| Solyc06g070910.2 | Solyc06g070910.3 |

| | |
|---|---|
| Solyc06g070920.2 | Solyc06g070920.3 |
| Solyc06g070930.2 | Solyc06g070930.3 |
| Solyc06g070950.1 | Solyc06g070950.2 |
| Solyc06g070980.2 | Solyc06g070980.3 |
| Solyc06g070990.2 | Solyc06g070990.3 |
| Solyc06g071000.2 | Solyc06g071000.3 |
| Solyc06g071020.2 | Solyc06g071020.3 |
| Solyc06g071030.2 | Solyc06g071030.3 |
| Solyc06g071110.2 | Solyc06g071110.3 |
| Solyc06g071120.2 | Solyc06g071120.3 |
| Solyc06g071130.2 | Solyc06g071130.3 |
| Solyc06g071160.2 | Solyc06g071160.3 |
| Solyc06g071180.2 | Solyc06g071180.3 |
| Solyc06g071200.2 | Solyc06g071200.3 |
| Solyc06g071210.2 | Solyc06g071210.3 |
| Solyc06g071230.2 | Solyc06g071230.3 |
| Solyc06g071270.2 | Solyc06g071270.3 |
| Solyc06g071290.2 | Solyc06g071290.3 |
| Solyc06g071320.2 | Solyc06g071320.3 |
| Solyc06g071380.2 | Solyc06g071380.3 |
| Solyc06g071390.2 | Solyc06g071390.3 |
| Solyc06g071400.2 | Solyc06g071400.3 |
| Solyc06g071410.2 | Solyc06g071410.3 |
| Solyc06g071420.2 | Solyc06g071420.3 |
| Solyc06g071430.2 | Solyc06g071430.3 |
| Solyc06g071460.2 | Solyc06g071460.3 |
| Solyc06g071480.2 | Solyc06g071480.3 |
| Solyc06g071500.2 | Solyc06g071500.3 |
| Solyc06g071510.2 | Solyc06g071510.3 |
| Solyc06g071550.2 | Solyc06g071550.3 |
| Solyc06g071560.2 | Solyc06g071560.3 |
| Solyc06g071570.2 | Solyc06g071570.3 |
| Solyc06g071580.2 | Solyc06g071580.3 |
| Solyc06g071590.2 | Solyc06g071590.3 |
| Solyc06g071600.2 | Solyc06g071600.3 |
| Solyc06g071620.2 | Solyc06g071620.3 |
| Solyc06g071630.1 | Solyc06g071630.2 |
| Solyc06g071640.2 | Solyc06g071640.3 |
| Solyc06g071690.2 | Solyc06g071690.3 |
| Solyc06g071730.2 | Solyc06g071730.3 |
| Solyc06g071750.2 | Solyc06g071750.3 |
| Solyc06g071790.2 | Solyc06g071790.3 |
| Solyc06g071820.2 | Solyc06g071820.3 |

| | |
|---|---|
| Solyc06g071850.2 | Solyc06g071850.3 |
| Solyc06g071860.2 | Solyc06g071860.3 |
| Solyc06g071870.2 | Solyc06g071870.3 |
| Solyc06g071890.2 | Solyc06g071890.3 |
| Solyc06g071900.2 | Solyc06g071900.3 |
| Solyc06g071910.2 | Solyc06g071910.3 |
| Solyc06g071980.2 | Solyc06g071980.3 |
| Solyc06g072010.1 | Solyc06g072010.2 |
| Solyc06g072070.1 | Solyc06g072070.2 |
| Solyc06g072080.2 | Solyc06g072080.3 |
| Solyc06g072120.2 | Solyc06g072120.3 |
| Solyc06g072160.2 | Solyc06g072160.3 |
| Solyc06g072250.2 | Solyc06g072250.3 |
| Solyc06g072280.2 | Solyc06g072280.3 |
| Solyc06g072290.2 | Solyc06g072290.3 |
| Solyc06g072310.2 | Solyc06g072310.3 |
| Solyc06g072330.2 | Solyc06g072330.3 |
| Solyc06g072340.2 | Solyc06g072340.3 |
| Solyc06g072350.2 | Solyc06g072350.3 |
| Solyc06g072360.2 | Solyc06g072360.3 |
| Solyc06g072370.2 | Solyc06g072370.3 |
| Solyc06g072380.2 | Solyc06g072380.3 |
| Solyc06g072420.2 | Solyc06g072420.3 |
| Solyc06g072440.2 | Solyc06g072440.3 |
| Solyc06g072490.2 | Solyc06g072490.3 |
| Solyc06g072520.1 | Solyc06g072520.2 |
| Solyc06g072530.2 | Solyc06g072530.3 |
| Solyc06g072540.1 | Solyc06g072540.2 |
| Solyc06g072550.2 | Solyc06g072550.3 |
| Solyc06g072560.1 | Solyc06g072560.2 |
| Solyc06g072580.2 | Solyc06g072580.3 |
| Solyc06g072600.2 | Solyc06g072600.3 |
| Solyc06g072610.2 | Solyc06g072610.3 |
| Solyc06g072670.2 | Solyc06g072670.3 |
| Solyc06g072700.2 | Solyc06g072700.3 |
| Solyc06g072710.2 | Solyc06g072710.3 |
| Solyc06g072720.2 | Solyc06g072720.3 |
| Solyc06g072750.2 | Solyc06g072750.3 |
| Solyc06g072770.2 | Solyc06g072770.3 |
| Solyc06g072780.1 | Solyc06g072780.2 |
| Solyc06g072790.2 | Solyc06g072790.3 |
| Solyc06g072820.2 | Solyc06g072820.3 |
| Solyc06g072880.1 | Solyc06g072880.2 |

| | |
|---|---|
| Solyc06g072970.2 | Solyc06g072970.3 |
| Solyc06g073010.2 | Solyc06g073010.3 |
| Solyc06g073020.2 | Solyc06g073020.3 |
| Solyc06g073060.2 | Solyc06g073060.3 |
| Solyc06g073070.1 | Solyc06g073070.2 |
| Solyc06g073090.2 | Solyc06g073090.3 |
| Solyc06g073110.2 | Solyc06g073110.3 |
| Solyc06g073140.1 | Solyc06g073140.2 |
| Solyc06g073150.2 | Solyc06g073150.3 |
| Solyc06g073180.2 | Solyc06g073180.3 |
| Solyc06g073190.2 | Solyc06g073190.3 |
| Solyc06g073240.2 | Solyc06g073240.3 |
| Solyc06g073245.1 | Solyc06g150132.1 |
| Solyc06g073260.2 | Solyc06g073260.3 |
| Solyc06g073280.2 | Solyc06g073280.3 |
| Solyc06g073310.2 | Solyc06g073310.3 |
| Solyc06g073320.2 | Solyc06g073320.3 |
| Solyc06g073340.2 | Solyc06g073340.3 |
| Solyc06g073360.2 | Solyc06g073360.3 |
| Solyc06g073380.2 | Solyc06g073380.3 |
| Solyc06g073390.2 | Solyc06g073390.3 |
| Solyc06g073400.2 | Solyc06g073400.3 |
| Solyc06g073410.2 | Solyc06g073410.3 |
| Solyc06g073420.2 | Solyc06g073420.3 |
| Solyc06g073470.2 | Solyc06g073470.3 |
| Solyc06g073480.2 | Solyc06g073480.3 |
| Solyc06g073500.2 | Solyc06g073500.3 |
| Solyc06g073510.2 | Solyc06g073510.3 |
| Solyc06g073520.2 | Solyc06g073520.3 |
| Solyc06g073550.2 | Solyc06g073550.3 |
| Solyc06g073585.1 | Solyc06g150133.1 |
| Solyc06g073590.2 | Solyc06g073590.3 |
| Solyc06g073610.2 | Solyc06g073610.3 |
| Solyc06g073630.2 | Solyc06g073630.3 |
| Solyc06g073660.2 | Solyc06g073660.3 |
| Solyc06g073700.2 | Solyc06g073700.3 |
| Solyc06g073730.1 | Solyc06g073730.2 |
| Solyc06g073740.2 | Solyc06g073740.3 |
| Solyc06g073760.2 | Solyc06g073760.3 |
| Solyc06g073790.2 | Solyc06g073790.3 |
| Solyc06g073800.2 | Solyc06g073800.3 |
| Solyc06g073840.2 | Solyc06g073840.3 |
| Solyc06g073860.1 | Solyc06g073860.2 |

| | |
|---|---|
| Solyc06g073880.2 | Solyc06g073880.3 |
| Solyc06g073890.2 | Solyc06g073890.3 |
| Solyc06g073900.2 | Solyc06g073900.3 |
| Solyc06g073920.2 | Solyc06g073920.3 |
| Solyc06g073950.2 | Solyc06g073950.3 |
| Solyc06g074010.2 | Solyc06g074010.3 |
| Solyc06g074040.1 | Solyc06g074040.2 |
| Solyc06g074060.1 | Solyc06g074060.2 |
| Solyc06g074070.2 | Solyc06g074070.3 |
| Solyc06g074080.2 | Solyc06g074080.3 |
| Solyc06g074090.2 | Solyc06g074090.3 |
| Solyc06g074110.2 | Solyc06g074110.3 |
| Solyc06g074120.2 | Solyc06g074120.3 |
| Solyc06g074150.2 | Solyc06g074150.3 |
| Solyc06g074170.2 | Solyc06g074170.3 |
| Solyc06g074190.2 | Solyc06g074190.3 |
| Solyc06g074210.2 | Solyc06g074210.3 |
| Solyc06g074220.2 | Solyc06g074220.3 |
| Solyc06g074230.2 | Solyc06g074230.3 |
| Solyc06g074250.2 | Solyc06g074250.3 |
| Solyc06g074320.2 | Solyc06g074320.3 |
| Solyc06g074330.2 | Solyc06g074330.3 |
| Solyc06g074340.2 | Solyc06g074340.3 |
| Solyc06g074350.2 | Solyc06g074350.3 |
| Solyc06g074390.2 | Solyc06g074390.3 |
| Solyc06g074470.2 | Solyc06g074470.3 |
| Solyc06g074480.2 | Solyc06g074480.3 |
| Solyc06g074500.1 | Solyc06g074500.2 |
| Solyc06g074550.2 | Solyc06g074550.3 |
| Solyc06g074570.1 | Solyc06g074570.2 |
| Solyc06g074590.1 | Solyc06g074590.2 |
| Solyc06g074600.1 | Solyc06g074600.2 |
| Solyc06g074640.1 | Solyc06g074640.2 |
| Solyc06g074650.2 | Solyc06g074650.3 |
| Solyc06g074740.2 | Solyc06g074740.3 |
| Solyc06g074760.1 | Solyc06g074760.2 |
| Solyc06g074790.1 | Solyc06g074790.2 |
| Solyc06g074810.2 | Solyc06g074810.3 |
| Solyc06g074815.1 | Solyc06g150134.1 |
| Solyc06g074820.2 | Solyc06g074820.3 |
| Solyc06g074840.2 | Solyc06g074840.3 |
| Solyc06g074850.2 | Solyc06g074850.3 |
| Solyc06g074860.1 | Solyc06g074860.2 |

| | |
|---|---|
| Solyc06g074905.1 | Solyc06g150135.1 |
| Solyc06g074910.2 | Solyc06g074910.3 |
| Solyc06g074930.1 | Solyc06g074930.2 |
| Solyc06g074980.2 | Solyc06g074980.3 |
| Solyc06g075000.2 | Solyc06g075000.3 |
| Solyc06g075020.2 | Solyc06g075020.3 |
| Solyc06g075050.1 | Solyc06g075050.2 |
| Solyc06g075060.1 | Solyc06g075060.2 |
| Solyc06g075110.2 | Solyc06g075110.3 |
| Solyc06g075140.2 | Solyc06g075140.3 |
| Solyc06g075150.2 | Solyc06g075150.3 |
| Solyc06g075160.2 | Solyc06g075160.3 |
| Solyc06g075250.2 | Solyc06g075250.3 |
| Solyc06g075260.2 | Solyc06g075260.3 |
| Solyc06g075280.2 | Solyc06g075280.3 |
| Solyc06g075320.2 | Solyc06g075320.3 |
| Solyc06g075360.2 | Solyc06g075360.3 |
| Solyc06g075370.2 | Solyc06g075370.3 |
| Solyc06g075400.2 | Solyc06g075400.3 |
| Solyc06g075450.1 | Solyc06g075450.2 |
| Solyc06g075520.2 | Solyc06g075520.3 |
| Solyc06g075540.2 | Solyc06g075540.3 |
| Solyc06g075550.2 | Solyc06g075550.3 |
| Solyc06g075560.1 | Solyc06g075560.2 |
| Solyc06g075570.1 | Solyc06g075570.2 |
| Solyc06g075600.2 | Solyc06g075600.3 |
| Solyc06g075620.2 | Solyc06g075620.3 |
| Solyc06g075640.1 | Solyc06g075640.2 |
| Solyc06g075650.2 | Solyc06g075650.3 |
| Solyc06g075690.2 | Solyc06g075690.3 |
| Solyc06g075740.1 | Solyc06g075740.2 |
| Solyc06g075790.2 | Solyc06g075790.3 |
| Solyc06g075810.2 | Solyc06g075810.3 |
| Solyc06g075830.1 | Solyc06g075830.2 |
| Solyc06g075920.1 | Solyc06g075920.2 |
| Solyc06g075950.1 | Solyc06g075950.2 |
| Solyc06g075980.2 | Solyc06g075980.3 |
| Solyc06g076020.2 | Solyc06g076020.3 |
| Solyc06g076030.2 | Solyc06g076030.3 |
| Solyc06g076040.2 | Solyc06g076040.3 |
| Solyc06g076090.2 | Solyc06g076090.3 |
| Solyc06g076100.2 | Solyc06g076100.3 |
| Solyc06g076130.2 | Solyc06g076130.3 |

| | |
|---|---|
| Solyc06g076150.2 | Solyc06g076150.3 |
| Solyc06g076200.1 | Solyc06g076200.2 |
| Solyc06g076220.2 | Solyc06g076220.3 |
| Solyc06g076260.2 | Solyc06g076260.3 |
| Solyc06g076270.2 | Solyc06g076270.3 |
| Solyc06g076300.2 | Solyc06g076300.3 |
| Solyc06g076330.2 | Solyc06g076330.3 |
| Solyc06g076340.2 | Solyc06g076340.3 |
| Solyc06g076350.2 | Solyc06g076350.3 |
| Solyc06g076390.2 | Solyc06g076390.3 |
| Solyc06g076400.2 | Solyc06g076400.3 |
| Solyc06g076430.2 | Solyc06g076430.3 |
| Solyc06g076450.2 | Solyc06g076450.3 |
| Solyc06g076470.2 | Solyc06g076470.3 |
| Solyc06g076490.2 | Solyc06g076490.3 |
| Solyc06g076510.2 | Solyc06g076510.3 |
| Solyc06g076570.3 | Solyc06g076570.4 |
| Solyc06g076590.2 | Solyc06g076590.3 |
| Solyc06g076600.1 | Solyc06g076600.2 |
| Solyc06g076610.2 | Solyc06g076610.3 |
| Solyc06g076620.2 | Solyc06g076620.3 |
| Solyc06g076630.2 | Solyc06g076630.3 |
| Solyc06g076640.2 | Solyc06g076640.3 |
| Solyc06g076660.2 | Solyc06g076660.3 |
| Solyc06g076670.2 | Solyc06g076670.3 |
| Solyc06g076750.2 | Solyc06g076750.3 |
| Solyc06g076760.1 | Solyc06g076760.2 |
| Solyc06g076770.2 | Solyc06g076770.3 |
| Solyc06g076780.2 | Solyc06g076780.3 |
| Solyc06g076800.2 | Solyc06g076800.3 |
| Solyc06g076830.1 | Solyc06g076830.2 |
| Solyc06g076850.2 | Solyc06g076850.3 |
| Solyc06g076880.2 | Solyc06g076880.3 |
| Solyc06g076920.2 | Solyc06g076920.3 |
| Solyc06g076930.1 | Solyc06g076930.2 |
| Solyc06g076940.2 | Solyc06g076940.3 |
| Solyc06g076950.2 | Solyc06g076950.3 |
| Solyc06g076960.1 | Solyc06g076960.2 |
| Solyc06g076970.2 | Solyc06g076970.3 |
| Solyc06g081990.1 | Solyc06g081990.2 |
| Solyc06g082030.2 | Solyc06g082030.3 |
| Solyc06g082040.1 | Solyc06g082040.2 |
| Solyc06g082050.1 | Solyc06g082050.2 |

| | |
|---|---|
| Solyc06g082060.2 | Solyc06g082060.3 |
| Solyc06g082070.2 | Solyc06g082070.3 |
| Solyc06g082080.2 | Solyc06g082080.3 |
| Solyc06g082130.2 | Solyc06g082130.3 |
| Solyc06g082140.2 | Solyc06g082140.3 |
| Solyc06g082155.1 | Solyc06g150136.1 |
| Solyc06g082190.2 | Solyc06g082190.3 |
| Solyc06g082270.1 | Solyc06g082270.2 |
| Solyc06g082280.2 | Solyc06g082280.3 |
| Solyc06g082310.2 | Solyc06g082310.3 |
| Solyc06g082340.1 | Solyc06g082340.2 |
| Solyc06g082380.2 | Solyc06g082380.3 |
| Solyc06g082430.1 | Solyc06g082430.2 |
| Solyc06g082470.2 | Solyc06g082470.3 |
| Solyc06g082480.2 | Solyc06g082480.3 |
| Solyc06g082500.2 | Solyc06g082500.3 |
| Solyc06g082510.2 | Solyc06g082510.3 |
| Solyc06g082520.2 | Solyc06g082520.3 |
| Solyc06g082535.1 | Solyc06g150137.1 |
| Solyc06g082640.2 | Solyc06g082640.3 |
| Solyc06g082670.2 | Solyc06g082670.3 |
| Solyc06g082740.2 | Solyc06g082740.3 |
| Solyc06g082750.2 | Solyc06g082750.3 |
| Solyc06g082770.2 | Solyc06g082770.3 |
| Solyc06g082810.2 | Solyc06g082810.3 |
| Solyc06g082880.1 | Solyc06g082880.2 |
| Solyc06g082900.2 | Solyc06g082900.3 |
| Solyc06g082910.2 | Solyc06g082910.3 |
| Solyc06g082920.2 | Solyc06g082920.3 |
| Solyc06g082930.2 | Solyc06g082930.3 |
| Solyc06g082940.2 | Solyc06g082940.3 |
| Solyc06g082960.2 | Solyc06g082960.3 |
| Solyc06g082970.2 | Solyc06g082970.3 |
| Solyc06g082980.2 | Solyc06g082980.3 |
| Solyc06g083030.2 | Solyc06g083030.3 |
| Solyc06g083040.2 | Solyc06g083040.3 |
| Solyc06g083050.2 | Solyc06g083050.3 |
| Solyc06g083070.2 | Solyc06g083070.3 |
| Solyc06g083080.2 | Solyc06g083080.3 |
| Solyc06g083100.1 | Solyc06g083100.2 |
| Solyc06g083110.1 | Solyc06g083110.2 |
| Solyc06g083150.2 | Solyc06g083150.3 |
| Solyc06g083170.2 | Solyc06g083170.3 |

| | |
|---|---|
| Solyc06g083210.2 | Solyc06g083210.3 |
| Solyc06g083220.2 | Solyc06g083220.3 |
| Solyc06g083230.2 | Solyc06g083230.3 |
| Solyc06g083260.2 | Solyc06g083260.3 |
| Solyc06g083270.2 | Solyc06g083270.3 |
| Solyc06g083280.1 | Solyc06g083280.2 |
| Solyc06g083310.2 | Solyc06g083310.3 |
| Solyc06g083340.2 | Solyc06g083340.3 |
| Solyc06g083350.2 | Solyc06g083350.3 |
| Solyc06g083360.2 | Solyc06g083360.3 |
| Solyc06g083380.2 | Solyc06g083380.3 |
| Solyc06g083415.1 | Solyc06g150138.1 |
| Solyc06g083440.2 | Solyc06g083440.3 |
| Solyc06g083520.2 | Solyc06g083520.3 |
| Solyc06g083530.2 | Solyc06g083530.3 |
| Solyc06g083570.2 | Solyc06g083570.3 |
| Solyc06g083620.2 | Solyc06g083620.3 |
| Solyc06g083650.2 | Solyc06g083650.3 |
| Solyc06g083680.2 | Solyc06g083680.3 |
| Solyc06g083690.2 | Solyc06g083690.3 |
| Solyc06g083720.1 | Solyc06g083720.2 |
| Solyc06g083730.2 | Solyc06g083730.3 |
| Solyc06g083750.2 | Solyc06g083750.3 |
| Solyc06g083760.2 | Solyc06g083760.3 |
| Solyc06g083780.2 | Solyc06g083780.3 |
| Solyc06g083790.2 | Solyc06g083790.3 |
| Solyc06g083810.2 | Solyc06g083810.3 |
| Solyc06g083820.2 | Solyc06g083820.3 |
| Solyc06g083870.2 | Solyc06g083870.3 |
| Solyc06g083900.2 | Solyc06g083900.3 |
| Solyc06g083910.2 | Solyc06g083910.3 |
| Solyc06g083930.1 | Solyc06g083930.2 |
| Solyc06g083990.2 | Solyc06g083990.3 |
| Solyc06g084030.2 | Solyc06g084030.3 |
| Solyc06g084040.2 | Solyc06g084040.3 |
| Solyc06g084045.1 | Solyc06g150139.1 |
| Solyc06g084060.2 | Solyc06g084060.3 |
| Solyc06g084070.2 | Solyc06g084070.3 |
| Solyc06g084130.2 | Solyc06g084130.3 |
| Solyc06g084170.2 | Solyc06g084170.3 |
| Solyc06g084180.2 | Solyc06g084180.3 |
| Solyc06g084190.2 | Solyc06g084190.3 |
| Solyc06g084210.2 | Solyc06g084210.3 |

| | |
|---|---|
| Solyc06g084230.2 | Solyc06g084230.3 |
| Solyc06g084240.1 | Solyc06g084240.2 |
| Solyc06g084260.2 | Solyc06g084260.3 |
| Solyc06g084270.2 | Solyc06g084270.3 |
| Solyc06g084290.2 | Solyc06g084290.3 |
| Solyc06g084330.2 | Solyc06g084330.3 |
| Solyc06g084350.2 | Solyc06g084350.3 |
| Solyc06g084390.2 | Solyc06g084390.3 |
| Solyc06g084400.2 | Solyc06g084400.3 |
| Solyc06g084440.2 | Solyc06g084440.3 |
| Solyc06g084480.2 | Solyc06g084480.3 |
| Solyc06g084520.2 | Solyc06g084520.3 |
| Solyc06g084530.2 | Solyc06g084530.3 |
| Solyc06g084540.2 | Solyc06g084540.3 |
| Solyc06g084550.2 | Solyc06g084550.3 |
| Solyc07g005000.2 | Solyc07g005000.3 |
| Solyc07g005020.2 | Solyc07g005020.3 |
| Solyc07g005050.2 | Solyc07g005050.3 |
| Solyc07g005060.2 | Solyc07g005060.3 |
| Solyc07g005120.2 | Solyc07g005120.3 |
| Solyc07g005130.2 | Solyc07g005130.3 |
| Solyc07g005145.1 | Solyc07g150100.1 |
| Solyc07g005180.2 | Solyc07g005180.3 |
| Solyc07g005190.2 | Solyc07g005190.3 |
| Solyc07g005200.2 | Solyc07g005200.3 |
| Solyc07g005210.2 | Solyc07g005210.3 |
| Solyc07g005280.2 | Solyc07g005280.3 |
| Solyc07g005320.2 | Solyc07g005320.3 |
| Solyc07g005330.2 | Solyc07g005330.3 |
| Solyc07g005340.1 | Solyc07g005340.2 |
| Solyc07g005360.2 | Solyc07g005360.3 |
| Solyc07g005390.2 | Solyc07g005390.3 |
| Solyc07g005400.2 | Solyc07g005400.3 |
| Solyc07g005420.1 | Solyc07g005420.2 |
| Solyc07g005450.2 | Solyc07g005450.3 |
| Solyc07g005480.2 | Solyc07g005480.3 |
| Solyc07g005510.2 | Solyc07g005510.3 |
| Solyc07g005560.2 | Solyc07g005560.3 |
| Solyc07g005590.2 | Solyc07g005590.3 |
| Solyc07g005630.2 | Solyc07g005630.3 |
| Solyc07g005640.2 | Solyc07g005640.3 |
| Solyc07g005700.2 | Solyc07g005700.3 |
| Solyc07g005710.2 | Solyc07g005710.3 |

| | |
|---|---|
| Solyc07g005720.2 | Solyc07g005720.3 |
| Solyc07g005750.2 | Solyc07g005750.3 |
| Solyc07g005760.2 | Solyc07g005760.3 |
| Solyc07g005780.1 | Solyc07g005780.2 |
| Solyc07g005810.2 | Solyc07g005810.3 |
| Solyc07g005820.2 | Solyc07g005820.3 |
| Solyc07g005830.2 | Solyc07g005830.3 |
| Solyc07g005860.2 | Solyc07g005860.3 |
| Solyc07g005870.2 | Solyc07g005870.3 |
| Solyc07g005880.2 | Solyc07g005880.3 |
| Solyc07g005890.2 | Solyc07g005890.3 |
| Solyc07g005900.2 | Solyc07g005900.3 |
| Solyc07g005910.2 | Solyc07g005910.3 |
| Solyc07g005960.2 | Solyc07g005960.3 |
| Solyc07g005970.2 | Solyc07g005970.3 |
| Solyc07g005980.1 | Solyc07g005980.2 |
| Solyc07g005990.2 | Solyc07g005990.3 |
| Solyc07g006000.2 | Solyc07g006000.3 |
| Solyc07g006010.2 | Solyc07g006010.3 |
| Solyc07g006020.2 | Solyc07g006020.3 |
| Solyc07g006045.1 | Solyc07g150101.1 |
| Solyc07g006060.2 | Solyc07g006060.3 |
| Solyc07g006120.2 | Solyc07g006120.3 |
| Solyc07g006160.2 | Solyc07g006160.3 |
| Solyc07g006180.2 | Solyc07g006180.3 |
| Solyc07g006190.2 | Solyc07g006190.3 |
| Solyc07g006200.2 | Solyc07g006200.3 |
| Solyc07g006220.1 | Solyc07g006220.2 |
| Solyc07g006260.2 | Solyc07g006260.3 |
| Solyc07g006280.2 | Solyc07g006280.3 |
| Solyc07g006330.1 | Solyc07g006330.2 |
| Solyc07g006340.2 | Solyc07g006340.3 |
| Solyc07g006350.2 | Solyc07g006350.3 |
| Solyc07g006380.2 | Solyc07g006380.3 |
| Solyc07g006395.1 | Solyc07g006395.2 |
| Solyc07g006500.2 | Solyc07g006500.3 |
| Solyc07g006510.2 | Solyc07g006510.3 |
| Solyc07g006520.2 | Solyc07g006520.3 |
| Solyc07g006530.2 | Solyc07g006530.3 |
| Solyc07g006540.2 | Solyc07g006540.3 |
| Solyc07g006560.2 | Solyc07g006560.3 |
| Solyc07g006580.2 | Solyc07g006580.3 |
| Solyc07g006610.2 | Solyc07g006610.3 |

| | |
|---|---|
| Solyc07g006640.2 | Solyc07g006640.3 |
| Solyc07g006690.2 | Solyc07g006690.3 |
| Solyc07g006710.1 | Solyc07g006710.2 |
| Solyc07g006740.2 | Solyc07g006740.3 |
| Solyc07g006750.2 | Solyc07g006750.3 |
| Solyc07g006780.2 | Solyc07g006780.3 |
| Solyc07g006790.2 | Solyc07g006790.3 |
| Solyc07g006810.2 | Solyc07g006810.3 |
| Solyc07g006860.2 | Solyc07g006860.3 |
| Solyc07g006855.1 | Solyc07g006855.2 |
| Solyc07g006900.1 | Solyc07g006900.2 |
| Solyc07g006920.2 | Solyc07g006920.3 |
| Solyc07g006950.2 | Solyc07g006950.3 |
| Solyc07g006960.2 | Solyc07g006960.3 |
| Solyc07g007000.1 | Solyc07g007000.2 |
| Solyc07g007020.2 | Solyc07g007020.3 |
| Solyc07g007120.2 | Solyc07g007120.3 |
| Solyc07g007140.2 | Solyc07g007140.3 |
| Solyc07g007160.2 | Solyc07g007160.3 |
| Solyc07g007190.2 | Solyc07g007190.3 |
| Solyc07g007210.2 | Solyc07g007210.3 |
| Solyc07g007220.2 | Solyc07g007220.3 |
| Solyc07g007240.2 | Solyc07g007240.3 |
| Solyc07g007270.2 | Solyc07g007270.3 |
| Solyc07g007300.2 | Solyc07g007300.3 |
| Solyc07g007310.2 | Solyc07g007310.3 |
| Solyc07g007340.2 | Solyc07g007340.3 |
| Solyc07g007370.2 | Solyc07g007370.3 |
| Solyc07g007390.2 | Solyc07g007390.3 |
| Solyc07g007400.2 | Solyc07g007400.3 |
| Solyc07g007420.2 | Solyc07g007420.3 |
| Solyc07g007430.2 | Solyc07g007430.3 |
| Solyc07g007450.2 | Solyc07g007450.3 |
| Solyc07g007480.1 | Solyc07g007480.2 |
| Solyc07g007490.1 | Solyc07g007490.2 |
| Solyc07g007500.2 | Solyc07g007500.3 |
| Solyc07g007550.2 | Solyc07g007550.3 |
| Solyc07g007560.2 | Solyc07g007560.3 |
| Solyc07g007600.2 | Solyc07g007600.3 |
| Solyc07g007620.2 | Solyc07g007620.3 |
| Solyc07g007630.2 | Solyc07g007630.3 |
| Solyc07g007670.2 | Solyc07g007670.3 |
| Solyc07g007750.2 | Solyc07g007750.3 |

| | |
|---|---|
| Solyc07g007770.1 | Solyc07g007770.2 |
| Solyc07g007780.2 | Solyc07g007780.3 |
| Solyc07g007790.2 | Solyc07g007790.3 |
| Solyc07g007800.2 | Solyc07g007800.3 |
| Solyc07g007810.2 | Solyc07g007810.3 |
| Solyc07g007820.2 | Solyc07g007820.3 |
| Solyc07g007870.2 | Solyc07g007870.3 |
| Solyc07g007890.2 | Solyc07g007890.3 |
| Solyc07g007930.2 | Solyc07g007930.3 |
| Solyc07g007945.1 | Solyc07g150102.1 |
| Solyc07g007960.2 | Solyc07g007960.3 |
| Solyc07g007970.2 | Solyc07g007970.3 |
| Solyc07g007980.2 | Solyc07g007980.3 |
| Solyc07g008020.2 | Solyc07g008020.3 |
| Solyc07g008103.1 | Solyc07g150103.1 |
| Solyc07g008107.1 | Solyc07g150104.1 |
| Solyc07g008130.2 | Solyc07g008130.3 |
| Solyc07g008140.2 | Solyc07g008140.3 |
| Solyc07g008150.2 | Solyc07g008150.3 |
| Solyc07g008160.2 | Solyc07g008160.3 |
| Solyc07g008200.2 | Solyc07g008200.3 |
| Solyc07g008240.2 | Solyc07g008240.3 |
| Solyc07g008250.2 | Solyc07g008250.3 |
| Solyc07g008260.2 | Solyc07g008260.3 |
| Solyc07g008270.2 | Solyc07g008270.3 |
| Solyc07g008310.2 | Solyc07g008310.3 |
| Solyc07g008320.2 | Solyc07g008320.3 |
| Solyc07g008330.2 | Solyc07g008330.3 |
| Solyc07g008350.2 | Solyc07g008350.3 |
| Solyc07g008360.1 | Solyc07g008360.2 |
| Solyc07g008380.1 | Solyc07g008380.2 |
| Solyc07g008420.2 | Solyc07g008420.3 |
| Solyc07g008440.2 | Solyc07g008440.3 |
| Solyc07g008450.2 | Solyc07g008450.3 |
| Solyc07g008460.2 | Solyc07g008460.3 |
| Solyc07g008470.2 | Solyc07g008470.3 |
| Solyc07g008500.1 | Solyc07g008500.2 |
| Solyc07g008520.2 | Solyc07g008520.3 |
| Solyc07g008540.2 | Solyc07g008540.3 |
| Solyc07g008560.2 | Solyc07g008560.3 |
| Solyc07g008570.2 | Solyc07g008570.3 |
| Solyc07g008660.2 | Solyc07g008660.3 |
| Solyc07g008670.2 | Solyc07g008670.3 |

| | |
|---|---|
| Solyc07g008690.2 | Solyc07g008690.3 |
| Solyc07g008700.2 | Solyc07g008700.3 |
| Solyc07g008730.1 | Solyc07g008730.2 |
| Solyc07g008820.2 | Solyc07g008820.3 |
| Solyc07g008840.2 | Solyc07g008840.3 |
| Solyc07g008850.2 | Solyc07g008850.3 |
| Solyc07g008860.2 | Solyc07g008860.3 |
| Solyc07g008880.2 | Solyc07g008880.3 |
| Solyc07g008910.2 | Solyc07g008910.3 |
| Solyc07g008940.2 | Solyc07g008940.3 |
| Solyc07g008980.2 | Solyc07g008980.3 |
| Solyc07g009020.1 | Solyc07g009020.2 |
| Solyc07g009030.2 | Solyc07g009030.3 |
| Solyc07g009040.2 | Solyc07g009040.3 |
| Solyc07g009050.2 | Solyc07g009050.3 |
| Solyc07g009100.2 | Solyc07g009100.3 |
| Solyc07g009140.2 | Solyc07g009140.3 |
| Solyc07g009150.2 | Solyc07g009150.3 |
| Solyc07g009160.2 | Solyc07g009160.3 |
| Solyc07g009220.1 | Solyc07g009220.2 |
| Solyc07g009230.2 | Solyc07g009230.3 |
| Solyc07g009235.1 | Solyc07g150105.1 |
| Solyc07g009260.2 | Solyc07g009260.3 |
| Solyc07g009280.2 | Solyc07g009280.3 |
| Solyc07g009290.2 | Solyc07g009290.3 |
| Solyc07g009320.2 | Solyc07g009320.3 |
| Solyc07g009440.1 | Solyc07g009440.2 |
| Solyc07g009460.2 | Solyc07g009460.3 |
| Solyc07g009490.2 | Solyc07g009490.3 |
| Solyc07g009520.1 | Solyc07g009520.2 |
| Solyc07g009580.1 | Solyc07g009580.2 |
| Solyc07g014580.2 | Solyc07g014580.3 |
| Solyc07g014640.2 | Solyc07g014640.3 |
| Solyc07g014650.2 | Solyc07g014650.3 |
| Solyc07g014670.2 | Solyc07g014670.3 |
| Solyc07g014690.2 | Solyc07g014690.3 |
| Solyc07g014703.1 | Solyc07g150106.1 |
| Solyc07g014730.2 | Solyc07g014730.3 |
| Solyc07g014740.2 | Solyc07g014740.3 |
| Solyc07g014755.1 | Solyc07g150107.1 |
| Solyc07g015840.1 | Solyc07g015840.2 |
| Solyc07g015860.2 | Solyc07g015860.3 |
| Solyc07g015940.2 | Solyc07g015940.3 |

| | |
|---|---|
| Solyc07g015990.1 | Solyc07g015990.2 |
| Solyc07g016050.2 | Solyc07g016050.3 |
| Solyc07g016065.1 | Solyc07g150108.1 |
| Solyc07g016160.2 | Solyc07g016160.3 |
| Solyc07g016180.2 | Solyc07g016180.3 |
| Solyc07g016200.2 | Solyc07g016200.3 |
| Solyc07g017220.2 | Solyc07g017220.3 |
| Solyc07g017230.2 | Solyc07g017230.3 |
| Solyc07g017293.1 | Solyc07g150109.1 |
| Solyc07g017297.1 | Solyc07g150110.1 |
| Solyc07g017343.1 | Solyc07g150111.1 |
| Solyc07g017347.1 | Solyc07g150112.1 |
| Solyc07g017400.2 | Solyc07g017400.3 |
| Solyc07g017430.2 | Solyc07g017430.3 |
| Solyc07g017490.2 | Solyc07g017490.3 |
| Solyc07g017495.1 | Solyc07g150113.1 |
| Solyc07g017500.2 | Solyc07g017500.3 |
| Solyc07g017510.2 | Solyc07g017510.3 |
| Solyc07g017535.1 | Solyc07g150114.1 |
| Solyc07g017600.2 | Solyc07g017600.3 |
| Solyc07g017610.2 | Solyc07g017610.3 |
| Solyc07g017730.2 | Solyc07g017730.3 |
| Solyc07g017770.2 | Solyc07g017770.3 |
| Solyc07g017860.2 | Solyc07g017860.3 |
| Solyc07g017870.1 | Solyc07g017870.2 |
| Solyc07g017900.2 | Solyc07g017900.3 |
| Solyc07g017935.1 | Solyc07g150115.1 |
| Solyc07g017950.2 | Solyc07g017950.3 |
| Solyc07g017955.1 | Solyc07g150116.1 |
| Solyc07g017990.2 | Solyc07g017990.3 |
| Solyc07g018000.2 | Solyc07g018000.3 |
| Solyc07g018010.2 | Solyc07g018010.3 |
| Solyc07g018130.1 | Solyc07g018130.2 |
| Solyc07g018140.2 | Solyc07g018140.3 |
| Solyc07g018210.2 | Solyc07g018210.3 |
| Solyc07g018220.2 | Solyc07g018220.3 |
| Solyc07g018230.1 | Solyc07g018230.2 |
| Solyc07g018270.2 | Solyc07g018270.3 |
| Solyc07g018300.2 | Solyc07g018300.3 |
| Solyc07g018340.2 | Solyc07g018340.3 |
| Solyc07g018360.2 | Solyc07g018360.3 |
| Solyc07g018385.1 | Solyc07g150117.1 |
| Solyc07g018430.2 | Solyc07g018430.3 |

| | |
|---|---|
| Solyc07g019440.2 | Solyc07g019440.3 |
| Solyc07g019450.2 | Solyc07g019450.3 |
| Solyc07g019460.2 | Solyc07g019460.3 |
| Solyc07g019497.1 | Solyc07g019497.2 |
| Solyc07g019530.2 | Solyc07g019530.3 |
| Solyc07g019573.1 | Solyc07g150118.1 |
| Solyc07g019635.1 | Solyc07g150119.1 |
| Solyc07g020710.2 | Solyc07g020710.3 |
| Solyc07g020800.2 | Solyc07g020800.3 |
| Solyc07g020860.2 | Solyc07g020860.3 |
| Solyc07g020910.1 | Solyc07g020910.2 |
| Solyc07g020960.2 | Solyc07g020960.3 |
| Solyc07g021040.1 | Solyc07g021040.2 |
| Solyc07g021335.1 | Solyc07g150120.1 |
| Solyc07g021415.1 | Solyc07g150121.1 |
| Solyc07g021463.1 | Solyc07g150122.1 |
| Solyc07g021467.1 | Solyc07g150123.1 |
| Solyc07g021540.2 | Solyc07g021540.3 |
| Solyc07g021550.2 | Solyc07g021550.3 |
| Solyc07g021630.2 | Solyc07g021630.3 |
| Solyc07g021640.1 | Solyc07g021640.2 |
| Solyc07g021660.1 | Solyc07g021660.2 |
| Solyc07g021670.1 | Solyc07g021670.2 |
| Solyc07g021700.2 | Solyc07g021700.3 |
| Solyc07g021710.2 | Solyc07g021710.3 |
| Solyc07g021740.1 | Solyc07g021740.2 |
| Solyc07g022830.2 | Solyc07g022830.3 |
| Solyc07g022880.2 | Solyc07g022880.3 |
| Solyc07g022910.2 | Solyc07g022910.3 |
| Solyc07g022920.2 | Solyc07g022920.3 |
| Solyc07g024000.2 | Solyc07g024000.3 |
| Solyc07g024010.1 | Solyc07g024010.2 |
| Solyc07g024020.1 | Solyc07g024020.2 |
| Solyc07g024030.2 | Solyc07g024030.3 |
| Solyc07g024040.1 | Solyc07g024040.2 |
| Solyc07g025170.2 | Solyc07g025170.3 |
| Solyc07g025180.1 | Solyc07g025180.2 |
| Solyc07g025210.1 | Solyc07g025210.2 |
| Solyc07g025230.1 | Solyc07g025230.2 |
| Solyc07g025255.1 | Solyc07g150124.1 |
| Solyc07g025370.2 | Solyc07g025370.3 |
| Solyc07g025450.1 | Solyc07g025450.2 |
| Solyc07g025520.2 | Solyc07g025520.3 |

| | |
|---|---|
| Solyc07g026650.2 | Solyc07g026650.3 |
| Solyc07g026770.2 | Solyc07g026770.3 |
| Solyc07g026810.2 | Solyc07g026810.3 |
| Solyc07g026960.2 | Solyc07g026960.3 |
| Solyc07g032080.2 | Solyc07g032080.3 |
| Solyc07g032100.2 | Solyc07g032100.3 |
| Solyc07g032110.2 | Solyc07g032110.3 |
| Solyc07g032150.2 | Solyc07g032150.3 |
| Solyc07g032170.2 | Solyc07g032170.3 |
| Solyc07g032180.2 | Solyc07g032180.3 |
| Solyc07g032220.2 | Solyc07g032220.3 |
| Solyc07g032230.2 | Solyc07g032230.3 |
| Solyc07g032240.2 | Solyc07g032240.3 |
| Solyc07g032390.1 | Solyc07g032390.2 |
| Solyc07g032480.2 | Solyc07g032480.3 |
| Solyc07g032490.2 | Solyc07g032490.3 |
| Solyc07g032510.2 | Solyc07g032510.3 |
| Solyc07g032615.1 | Solyc07g150125.1 |
| Solyc07g032783.1 | Solyc07g150126.1 |
| Solyc07g032785.1 | Solyc07g150127.1 |
| Solyc07g039260.1 | Solyc07g039260.2 |
| Solyc07g039410.2 | Solyc07g039410.3 |
| Solyc07g039450.2 | Solyc07g039450.3 |
| Solyc07g039570.2 | Solyc07g039570.3 |
| Solyc07g040663.1 | Solyc07g040663.2 |
| Solyc07g040667.1 | Solyc07g150130.1 |
| Solyc07g040680.2 | Solyc07g040680.3 |
| Solyc07g040690.2 | Solyc07g040690.3 |
| Solyc07g040710.2 | Solyc07g040710.3 |
| Solyc07g040720.2 | Solyc07g040720.3 |
| Solyc07g040740.2 | Solyc07g040740.3 |
| Solyc07g040750.1 | Solyc07g040750.2 |
| Solyc07g040785.1 | Solyc07g150131.1 |
| Solyc07g040910.2 | Solyc07g040910.3 |
| Solyc07g040940.2 | Solyc07g040940.3 |
| Solyc07g040950.2 | Solyc07g040950.3 |
| Solyc07g040980.2 | Solyc07g040980.3 |
| Solyc07g041010.2 | Solyc07g041010.3 |
| Solyc07g041020.2 | Solyc07g041020.3 |
| Solyc07g041023.1 | Solyc07g150132.1 |
| Solyc07g041070.2 | Solyc07g041070.3 |
| Solyc07g041080.2 | Solyc07g041080.3 |
| Solyc07g041170.1 | Solyc07g041170.2 |

| | |
|---|---|
| Solyc07g041190.2 | Solyc07g041190.3 |
| Solyc07g041200.2 | Solyc07g041200.3 |
| Solyc07g041210.2 | Solyc07g041210.3 |
| Solyc07g041230.2 | Solyc07g041230.3 |
| Solyc07g041300.1 | Solyc07g041300.2 |
| Solyc07g041310.2 | Solyc07g041310.3 |
| Solyc07g041340.1 | Solyc07g041340.2 |
| Solyc07g041500.2 | Solyc07g041500.3 |
| Solyc07g041640.2 | Solyc07g041640.3 |
| Solyc07g041680.1 | Solyc07g041680.2 |
| Solyc07g041750.2 | Solyc07g041750.3 |
| Solyc07g041760.2 | Solyc07g041760.3 |
| Solyc07g041780.2 | Solyc07g041780.3 |
| Solyc07g041840.1 | Solyc07g041840.2 |
| Solyc07g041870.2 | Solyc07g041870.3 |
| Solyc07g041895.1 | Solyc07g150133.1 |
| Solyc07g041900.2 | Solyc07g041900.3 |
| Solyc07g041930.2 | Solyc07g041930.3 |
| Solyc07g041940.2 | Solyc07g041940.3 |
| Solyc07g041960.1 | Solyc07g041960.2 |
| Solyc07g041980.2 | Solyc07g041980.3 |
| Solyc07g042020.2 | Solyc07g042020.3 |
| Solyc07g042030.1 | Solyc07g042030.2 |
| Solyc07g042150.2 | Solyc07g042150.3 |
| Solyc07g042160.2 | Solyc07g042160.3 |
| Solyc07g042170.2 | Solyc07g042170.3 |
| Solyc07g042190.2 | Solyc07g042190.3 |
| Solyc07g042220.1 | Solyc07g042220.2 |
| Solyc07g042250.2 | Solyc07g042250.3 |
| Solyc07g042330.1 | Solyc07g042330.2 |
| Solyc07g042440.2 | Solyc07g042440.3 |
| Solyc07g042460.1 | Solyc07g042460.2 |
| Solyc07g042470.2 | Solyc07g042470.3 |
| Solyc07g042480.2 | Solyc07g042480.3 |
| Solyc07g042510.2 | Solyc07g042510.3 |
| Solyc07g042550.2 | Solyc07g042550.3 |
| Solyc07g042570.2 | Solyc07g042570.3 |
| Solyc07g042580.2 | Solyc07g042580.3 |
| Solyc07g042593.1 | Solyc07g150134.1 |
| Solyc07g042610.1 | Solyc07g042610.2 |
| Solyc07g042680.2 | Solyc07g042680.3 |
| Solyc07g042740.2 | Solyc07g042740.3 |
| Solyc07g042820.2 | Solyc07g042820.3 |

| | |
|---|---|
| Solyc07g042890.2 | Solyc07g042890.3 |
| Solyc07g042930.2 | Solyc07g042930.3 |
| Solyc07g043110.1 | Solyc07g043110.2 |
| Solyc07g043130.2 | Solyc07g043130.3 |
| Solyc07g043133.1 | Solyc07g150135.1 |
| Solyc07g043137.1 | Solyc07g150136.1 |
| Solyc07g043200.1 | Solyc07g043200.2 |
| Solyc07g043230.2 | Solyc07g043230.3 |
| Solyc07g043270.2 | Solyc07g043270.3 |
| Solyc07g043310.2 | Solyc07g043310.3 |
| Solyc07g043365.1 | Solyc07g150137.1 |
| Solyc07g043420.2 | Solyc07g043420.3 |
| Solyc07g043460.2 | Solyc07g043460.3 |
| Solyc07g043510.2 | Solyc07g043510.3 |
| Solyc07g043530.1 | Solyc07g043530.2 |
| Solyc07g043550.2 | Solyc07g043550.3 |
| Solyc07g043570.2 | Solyc07g043570.3 |
| Solyc07g043590.2 | Solyc07g043590.3 |
| Solyc07g043610.2 | Solyc07g043610.3 |
| Solyc07g043620.2 | Solyc07g043620.3 |
| Solyc07g043630.1 | Solyc07g043630.2 |
| Solyc07g043640.2 | Solyc07g043640.3 |
| Solyc07g043660.2 | Solyc07g043660.3 |
| Solyc07g043680.2 | Solyc07g043680.3 |
| Solyc07g043690.1 | Solyc07g043690.2 |
| Solyc07g044710.1 | Solyc07g044710.2 |
| Solyc07g044750.2 | Solyc07g044750.3 |
| Solyc07g044760.2 | Solyc07g044760.3 |
| Solyc07g044780.2 | Solyc07g044780.3 |
| Solyc07g044810.2 | Solyc07g044810.3 |
| Solyc07g044830.2 | Solyc07g044830.3 |
| Solyc07g044840.2 | Solyc07g044840.3 |
| Solyc07g044850.2 | Solyc07g044850.3 |
| Solyc07g044860.2 | Solyc07g044860.3 |
| Solyc07g044880.2 | Solyc07g044880.3 |
| Solyc07g044930.2 | Solyc07g044930.3 |
| Solyc07g044980.2 | Solyc07g044980.3 |
| Solyc07g044990.2 | Solyc07g044990.3 |
| Solyc07g045010.2 | Solyc07g045010.3 |
| Solyc07g045040.2 | Solyc07g045040.3 |
| Solyc07g045050.2 | Solyc07g045050.3 |
| Solyc07g045070.2 | Solyc07g045070.3 |
| Solyc07g045080.2 | Solyc07g045080.3 |

| | |
|---|---|
| Solyc07g045090.2 | Solyc07g045090.3 |
| Solyc07g045120.1 | Solyc07g045120.2 |
| Solyc07g045160.2 | Solyc07g045160.3 |
| Solyc07g045220.2 | Solyc07g045220.3 |
| Solyc07g045240.2 | Solyc07g045240.3 |
| Solyc07g045270.1 | Solyc07g045270.2 |
| Solyc07g045290.2 | Solyc07g045290.3 |
| Solyc07g045320.2 | Solyc07g045320.3 |
| Solyc07g045330.2 | Solyc07g045330.3 |
| Solyc07g045370.2 | Solyc07g045370.3 |
| Solyc07g045380.2 | Solyc07g045380.3 |
| Solyc07g045410.1 | Solyc07g045410.2 |
| Solyc07g045460.2 | Solyc07g045460.3 |
| Solyc07g045480.2 | Solyc07g045480.3 |
| Solyc07g045530.1 | Solyc07g045530.2 |
| Solyc07g047610.2 | Solyc07g047610.3 |
| Solyc07g047625.1 | Solyc07g150138.1 |
| Solyc07g047670.2 | Solyc07g047670.3 |
| Solyc07g047710.2 | Solyc07g047710.3 |
| Solyc07g047720.2 | Solyc07g047720.3 |
| Solyc07g047740.2 | Solyc07g047740.3 |
| Solyc07g047750.2 | Solyc07g047750.3 |
| Solyc07g047780.2 | Solyc07g047780.3 |
| Solyc07g047790.2 | Solyc07g047790.3 |
| Solyc07g047800.2 | Solyc07g047800.3 |
| Solyc07g047830.2 | Solyc07g047830.3 |
| Solyc07g047850.2 | Solyc07g047850.3 |
| Solyc07g047900.2 | Solyc07g047900.3 |
| Solyc07g047910.1 | Solyc07g047910.2 |
| Solyc07g047960.2 | Solyc07g047960.3 |
| Solyc07g048000.2 | Solyc07g048000.3 |
| Solyc07g048070.2 | Solyc07g048070.3 |
| Solyc07g048090.1 | Solyc07g048090.2 |
| Solyc07g048120.2 | Solyc07g048120.3 |
| Solyc07g049120.1 | Solyc07g049120.2 |
| Solyc07g049170.1 | Solyc07g049170.2 |
| Solyc07g049180.2 | Solyc07g049180.3 |
| Solyc07g049210.2 | Solyc07g049210.3 |
| Solyc07g049240.2 | Solyc07g049240.3 |
| Solyc07g049280.2 | Solyc07g049280.3 |
| Solyc07g049290.2 | Solyc07g049290.3 |
| Solyc07g049300.2 | Solyc07g049300.3 |
| Solyc07g049310.2 | Solyc07g049310.3 |

| | |
|---|---|
| Solyc07g049325.1 | Solyc07g150139.1 |
| Solyc07g049360.2 | Solyc07g049360.3 |
| Solyc07g049385.1 | Solyc07g150140.1 |
| Solyc07g049410.2 | Solyc07g049410.3 |
| Solyc07g049430.2 | Solyc07g049430.3 |
| Solyc07g049440.2 | Solyc07g049440.3 |
| Solyc07g049450.2 | Solyc07g049450.3 |
| Solyc07g049460.2 | Solyc07g049460.3 |
| Solyc07g049480.2 | Solyc07g049480.3 |
| Solyc07g049500.2 | Solyc07g049500.3 |
| Solyc07g049520.1 | Solyc07g049520.2 |
| Solyc07g049530.2 | Solyc07g049530.3 |
| Solyc07g049540.1 | Solyc07g049540.2 |
| Solyc07g049550.2 | Solyc07g049550.3 |
| Solyc07g049560.2 | Solyc07g049560.3 |
| Solyc07g049570.2 | Solyc07g049570.3 |
| Solyc07g049580.2 | Solyc07g049580.3 |
| Solyc07g049590.2 | Solyc07g049590.3 |
| Solyc07g049640.2 | Solyc07g049640.3 |
| Solyc07g049690.2 | Solyc07g049690.3 |
| Solyc07g049710.2 | Solyc07g049710.3 |
| Solyc07g049730.2 | Solyc07g049730.3 |
| Solyc07g049750.2 | Solyc07g049750.3 |
| Solyc07g049760.1 | Solyc07g049760.2 |
| Solyc07g051810.1 | Solyc07g051810.2 |
| Solyc07g051820.2 | Solyc07g051820.3 |
| Solyc07g051850.2 | Solyc07g051850.3 |
| Solyc07g051940.3 | Solyc07g051940.4 |
| Solyc07g051950.2 | Solyc07g051950.3 |
| Solyc07g051990.1 | Solyc07g051990.2 |
| Solyc07g052020.1 | Solyc07g052020.2 |
| Solyc07g052025.1 | Solyc07g150141.1 |
| Solyc07g052050.1 | Solyc07g052050.2 |
| Solyc07g052230.2 | Solyc07g052230.3 |
| Solyc07g052240.2 | Solyc07g052240.3 |
| Solyc07g052255.1 | Solyc07g150142.1 |
| Solyc07g052260.2 | Solyc07g052260.3 |
| Solyc07g052270.2 | Solyc07g052270.3 |
| Solyc07g052275.1 | Solyc07g150143.1 |
| Solyc07g052300.2 | Solyc07g052300.3 |
| Solyc07g052320.2 | Solyc07g052320.3 |
| Solyc07g052350.2 | Solyc07g052350.3 |
| Solyc07g052380.2 | Solyc07g052380.3 |

| | |
|---|---|
| Solyc07g052390.2 | Solyc07g052390.3 |
| Solyc07g052400.2 | Solyc07g052400.3 |
| Solyc07g052430.2 | Solyc07g052430.3 |
| Solyc07g052440.2 | Solyc07g052440.3 |
| Solyc07g052470.2 | Solyc07g052470.3 |
| Solyc07g052480.2 | Solyc07g052480.3 |
| Solyc07g052600.2 | Solyc07g052600.3 |
| Solyc07g052610.2 | Solyc07g052610.3 |
| Solyc07g052640.2 | Solyc07g052640.3 |
| Solyc07g052650.1 | Solyc07g052650.2 |
| Solyc07g052670.2 | Solyc07g052670.3 |
| Solyc07g052700.2 | Solyc07g052700.3 |
| Solyc07g052730.2 | Solyc07g052730.3 |
| Solyc07g052740.2 | Solyc07g052740.3 |
| Solyc07g052780.2 | Solyc07g052780.3 |
| Solyc07g052870.1 | Solyc07g052870.2 |
| Solyc07g052910.2 | Solyc07g052910.3 |
| Solyc07g052930.2 | Solyc07g052930.3 |
| Solyc07g052950.2 | Solyc07g052950.3 |
| Solyc07g052970.2 | Solyc07g052970.3 |
| Solyc07g052980.2 | Solyc07g052980.3 |
| Solyc07g053060.2 | Solyc07g053060.3 |
| Solyc07g053110.2 | Solyc07g053110.3 |
| Solyc07g053140.2 | Solyc07g053140.3 |
| Solyc07g053150.2 | Solyc07g053150.3 |
| Solyc07g053180.2 | Solyc07g053180.3 |
| Solyc07g053210.2 | Solyc07g053210.3 |
| Solyc07g053220.1 | Solyc07g053220.2 |
| Solyc07g053230.2 | Solyc07g053230.3 |
| Solyc07g053240.2 | Solyc07g053240.3 |
| Solyc07g053250.2 | Solyc07g053250.3 |
| Solyc07g053260.2 | Solyc07g053260.3 |
| Solyc07g053270.2 | Solyc07g053270.3 |
| Solyc07g053290.2 | Solyc07g053290.3 |
| Solyc07g053310.2 | Solyc07g053310.3 |
| Solyc07g053330.1 | Solyc07g053330.2 |
| Solyc07g053375.1 | Solyc07g150144.1 |
| Solyc07g053380.1 | Solyc07g053380.2 |
| Solyc07g053490.2 | Solyc07g053490.3 |
| Solyc07g053520.1 | Solyc07g053520.2 |
| Solyc07g053600.2 | Solyc07g053600.3 |
| Solyc07g053610.2 | Solyc07g053610.3 |
| Solyc07g053660.2 | Solyc07g053660.3 |

| | |
|---|---|
| Solyc07g053670.2 | Solyc07g053670.3 |
| Solyc07g053680.1 | Solyc07g053680.2 |
| Solyc07g053690.2 | Solyc07g053690.3 |
| Solyc07g053710.2 | Solyc07g053710.3 |
| Solyc07g053720.2 | Solyc07g053720.3 |
| Solyc07g053770.1 | Solyc07g053770.2 |
| Solyc07g053790.1 | Solyc07g053790.2 |
| Solyc07g053800.2 | Solyc07g053800.3 |
| Solyc07g053810.2 | Solyc07g053810.3 |
| Solyc07g053820.2 | Solyc07g053820.3 |
| Solyc07g053830.2 | Solyc07g053830.3 |
| Solyc07g053850.2 | Solyc07g053850.3 |
| Solyc07g053860.2 | Solyc07g053860.3 |
| Solyc07g053890.2 | Solyc07g053890.3 |
| Solyc07g053910.2 | Solyc07g053910.3 |
| Solyc07g053920.2 | Solyc07g053920.3 |
| Solyc07g053960.2 | Solyc07g053960.3 |
| Solyc07g053970.2 | Solyc07g053970.3 |
| Solyc07g053980.2 | Solyc07g053980.3 |
| Solyc07g054000.2 | Solyc07g054000.3 |
| Solyc07g054010.1 | Solyc07g054010.2 |
| Solyc07g054020.1 | Solyc07g054020.2 |
| Solyc07g054090.2 | Solyc07g054090.3 |
| Solyc07g054120.1 | Solyc07g054120.2 |
| Solyc07g054140.2 | Solyc07g054140.3 |
| Solyc07g054160.2 | Solyc07g054160.3 |
| Solyc07g054170.2 | Solyc07g054170.3 |
| Solyc07g054210.2 | Solyc07g054210.3 |
| Solyc07g054270.2 | Solyc07g054270.3 |
| Solyc07g054300.2 | Solyc07g054300.3 |
| Solyc07g054315.1 | Solyc07g150146.1 |
| Solyc07g054320.2 | Solyc07g054320.3 |
| Solyc07g054370.2 | Solyc07g054370.3 |
| Solyc07g054380.1 | Solyc07g054380.2 |
| Solyc07g054440.2 | Solyc07g054440.3 |
| Solyc07g054490.2 | Solyc07g054490.3 |
| Solyc07g054580.2 | Solyc07g054580.3 |
| Solyc07g054600.2 | Solyc07g054600.3 |
| Solyc07g054610.2 | Solyc07g054610.3 |
| Solyc07g054680.2 | Solyc07g054680.3 |
| Solyc07g054700.2 | Solyc07g054700.3 |
| Solyc07g054710.1 | Solyc07g054710.2 |
| Solyc07g054720.1 | Solyc07g054720.2 |

| | |
|---|---|
| Solyc07g054740.2 | Solyc07g054740.3 |
| Solyc07g054920.1 | Solyc07g054920.2 |
| Solyc07g054930.2 | Solyc07g054930.3 |
| Solyc07g054940.1 | Solyc07g054940.2 |
| Solyc07g054950.1 | Solyc07g054950.2 |
| Solyc07g054960.1 | Solyc07g054960.2 |
| Solyc07g055000.1 | Solyc07g055000.2 |
| Solyc07g055010.2 | Solyc07g055010.3 |
| Solyc07g055020.2 | Solyc07g055020.3 |
| Solyc07g055050.2 | Solyc07g055050.3 |
| Solyc07g055060.2 | Solyc07g055060.3 |
| Solyc07g055065.1 | Solyc07g150147.1 |
| Solyc07g055080.2 | Solyc07g055080.3 |
| Solyc07g055140.2 | Solyc07g055140.3 |
| Solyc07g055150.2 | Solyc07g055150.3 |
| Solyc07g055180.2 | Solyc07g055180.3 |
| Solyc07g055190.2 | Solyc07g055190.3 |
| Solyc07g055210.2 | Solyc07g055210.3 |
| Solyc07g055217.1 | Solyc07g150148.1 |
| Solyc07g055250.2 | Solyc07g055250.3 |
| Solyc07g055260.2 | Solyc07g055260.3 |
| Solyc07g055290.2 | Solyc07g055290.3 |
| Solyc07g055300.2 | Solyc07g055300.3 |
| Solyc07g055330.2 | Solyc07g055330.3 |
| Solyc07g055340.2 | Solyc07g055340.3 |
| Solyc07g055400.1 | Solyc07g055400.2 |
| Solyc07g055410.1 | Solyc07g055410.2 |
| Solyc07g055440.1 | Solyc07g055440.2 |
| Solyc07g055450.1 | Solyc07g055450.2 |
| Solyc07g055460.2 | Solyc07g055460.3 |
| Solyc07g055470.2 | Solyc07g055470.3 |
| Solyc07g055480.1 | Solyc07g055480.2 |
| Solyc07g055495.1 | Solyc07g150149.1 |
| Solyc07g055530.2 | Solyc07g055530.3 |
| Solyc07g055550.1 | Solyc07g055550.2 |
| Solyc07g055620.1 | Solyc07g055620.2 |
| Solyc07g055650.1 | Solyc07g055650.2 |
| Solyc07g055710.2 | Solyc07g055710.3 |
| Solyc07g055780.2 | Solyc07g055780.3 |
| Solyc07g055810.2 | Solyc07g055810.3 |
| Solyc07g055820.1 | Solyc07g055820.2 |
| Solyc07g055830.2 | Solyc07g055830.3 |
| Solyc07g055840.2 | Solyc07g055840.3 |

| | |
|---|---|
| Solyc07g055930.2 | Solyc07g055930.3 |
| Solyc07g055950.2 | Solyc07g055950.3 |
| Solyc07g055960.2 | Solyc07g055960.3 |
| Solyc07g055980.2 | Solyc07g055980.3 |
| Solyc07g056010.2 | Solyc07g056010.3 |
| Solyc07g056020.2 | Solyc07g056020.3 |
| Solyc07g056030.2 | Solyc07g056030.3 |
| Solyc07g056040.2 | Solyc07g056040.3 |
| Solyc07g056070.1 | Solyc07g056070.2 |
| Solyc07g056110.2 | Solyc07g056110.3 |
| Solyc07g056120.2 | Solyc07g056120.3 |
| Solyc07g056140.2 | Solyc07g056140.3 |
| Solyc07g056180.1 | Solyc07g056180.2 |
| Solyc07g056190.2 | Solyc07g056190.3 |
| Solyc07g056200.2 | Solyc07g056200.3 |
| Solyc07g056220.2 | Solyc07g056220.3 |
| Solyc07g056230.2 | Solyc07g056230.3 |
| Solyc07g056250.2 | Solyc07g056250.3 |
| Solyc07g056270.2 | Solyc07g056270.3 |
| Solyc07g056280.2 | Solyc07g056280.3 |
| Solyc07g056290.1 | Solyc07g056290.2 |
| Solyc07g056310.2 | Solyc07g056310.3 |
| Solyc07g056330.2 | Solyc07g056330.3 |
| Solyc07g056350.2 | Solyc07g056350.3 |
| Solyc07g056370.2 | Solyc07g056370.3 |
| Solyc07g056410.2 | Solyc07g056410.3 |
| Solyc07g056500.3 | Solyc07g056500.4 |
| Solyc07g056510.2 | Solyc07g056510.3 |
| Solyc07g056520.2 | Solyc07g056520.3 |
| Solyc07g056530.2 | Solyc07g056530.3 |
| Solyc07g056540.2 | Solyc07g056540.3 |
| Solyc07g056550.2 | Solyc07g056550.3 |
| Solyc07g056560.2 | Solyc07g056560.3 |
| Solyc07g056580.2 | Solyc07g056580.3 |
| Solyc07g056590.2 | Solyc07g056590.3 |
| Solyc07g056610.1 | Solyc07g056610.2 |
| Solyc07g056630.2 | Solyc07g056630.3 |
| Solyc07g056670.2 | Solyc07g056670.3 |
| Solyc07g056690.1 | Solyc07g056690.2 |
| Solyc07g056700.1 | Solyc07g056700.2 |
| Solyc07g056703.1 | Solyc07g150150.1 |
| Solyc07g061700.1 | Solyc07g061700.2 |
| Solyc07g061710.1 | Solyc07g061710.2 |

| | |
|---|---|
| Solyc07g061720.2 | Solyc07g061720.3 |
| Solyc07g061730.2 | Solyc07g061730.3 |
| Solyc07g061790.2 | Solyc07g061790.3 |
| Solyc07g061820.1 | Solyc07g061820.2 |
| Solyc07g061900.2 | Solyc07g061900.3 |
| Solyc07g061905.1 | Solyc07g150151.1 |
| Solyc07g061960.2 | Solyc07g061960.3 |
| Solyc07g061970.2 | Solyc07g061970.3 |
| Solyc07g061980.1 | Solyc07g061980.2 |
| Solyc07g061990.2 | Solyc07g061990.3 |
| Solyc07g062000.1 | Solyc07g062000.2 |
| Solyc07g062020.1 | Solyc07g062020.2 |
| Solyc07g062030.2 | Solyc07g062030.3 |
| Solyc07g062040.2 | Solyc07g062040.3 |
| Solyc07g062045.1 | Solyc07g150152.1 |
| Solyc07g062050.2 | Solyc07g062050.3 |
| Solyc07g062060.2 | Solyc07g062060.3 |
| Solyc07g062070.2 | Solyc07g062070.3 |
| Solyc07g062110.2 | Solyc07g062110.3 |
| Solyc07g062130.2 | Solyc07g062130.3 |
| Solyc07g062140.2 | Solyc07g062140.3 |
| Solyc07g062150.2 | Solyc07g062150.3 |
| Solyc07g062160.2 | Solyc07g062160.3 |
| Solyc07g062200.1 | Solyc07g062200.2 |
| Solyc07g062220.2 | Solyc07g062220.3 |
| Solyc07g062260.2 | Solyc07g062260.3 |
| Solyc07g062270.2 | Solyc07g062270.3 |
| Solyc07g062280.2 | Solyc07g062280.3 |
| Solyc07g062290.2 | Solyc07g062290.3 |
| Solyc07g062300.2 | Solyc07g062300.3 |
| Solyc07g062330.1 | Solyc07g062330.2 |
| Solyc07g062350.1 | Solyc07g062350.2 |
| Solyc07g062400.2 | Solyc07g062400.3 |
| Solyc07g062430.2 | Solyc07g062430.3 |
| Solyc07g062450.2 | Solyc07g062450.3 |
| Solyc07g062480.1 | Solyc07g062480.2 |
| Solyc07g062500.2 | Solyc07g062500.3 |
| Solyc07g062520.2 | Solyc07g062520.3 |
| Solyc07g062530.2 | Solyc07g062530.3 |
| Solyc07g062540.2 | Solyc07g062540.3 |
| Solyc07g062560.2 | Solyc07g062560.3 |
| Solyc07g062570.2 | Solyc07g062570.3 |
| Solyc07g062620.2 | Solyc07g062620.3 |

| | |
|---|---|
| Solyc07g062685.1 | Solyc07g062685.2 |
| Solyc07g062700.2 | Solyc07g062700.3 |
| Solyc07g062780.2 | Solyc07g062780.3 |
| Solyc07g062840.2 | Solyc07g062840.3 |
| Solyc07g062860.2 | Solyc07g062860.3 |
| Solyc07g062870.2 | Solyc07g062870.3 |
| Solyc07g062880.2 | Solyc07g062880.3 |
| Solyc07g062910.2 | Solyc07g062910.3 |
| Solyc07g062930.2 | Solyc07g062930.3 |
| Solyc07g062970.2 | Solyc07g062970.3 |
| Solyc07g062990.1 | Solyc07g062990.2 |
| Solyc07g063010.2 | Solyc07g063010.3 |
| Solyc07g063040.2 | Solyc07g063040.3 |
| Solyc07g063150.2 | Solyc07g063150.3 |
| Solyc07g063180.2 | Solyc07g063180.3 |
| Solyc07g063190.2 | Solyc07g063190.3 |
| Solyc07g063210.2 | Solyc07g063210.3 |
| Solyc07g063230.2 | Solyc07g063230.3 |
| Solyc07g063270.2 | Solyc07g063270.3 |
| Solyc07g063290.2 | Solyc07g063290.3 |
| Solyc07g063300.2 | Solyc07g063300.3 |
| Solyc07g063310.2 | Solyc07g063310.3 |
| Solyc07g063320.2 | Solyc07g063320.3 |
| Solyc07g063340.2 | Solyc07g063340.3 |
| Solyc07g063350.2 | Solyc07g063350.3 |
| Solyc07g063390.2 | Solyc07g063390.3 |
| Solyc07g063400.2 | Solyc07g063400.3 |
| Solyc07g063410.2 | Solyc07g063410.3 |
| Solyc07g063420.2 | Solyc07g063420.3 |
| Solyc07g063430.2 | Solyc07g063430.3 |
| Solyc07g063440.2 | Solyc07g063440.3 |
| Solyc07g063480.2 | Solyc07g063480.3 |
| Solyc07g063510.2 | Solyc07g063510.3 |
| Solyc07g063520.2 | Solyc07g063520.3 |
| Solyc07g063550.2 | Solyc07g063550.3 |
| Solyc07g063600.2 | Solyc07g063600.3 |
| Solyc07g063620.2 | Solyc07g063620.3 |
| Solyc07g063630.2 | Solyc07g063630.3 |
| Solyc07g063700.1 | Solyc07g063700.2 |
| Solyc07g063710.1 | Solyc07g063710.2 |
| Solyc07g063750.2 | Solyc07g063750.3 |
| Solyc07g063770.2 | Solyc07g063770.3 |
| Solyc07g063780.1 | Solyc07g063780.2 |

| | |
|---|---|
| Solyc07g063800.1 | Solyc07g063800.2 |
| Solyc07g063810.1 | Solyc07g063810.2 |
| Solyc07g063820.2 | Solyc07g063820.3 |
| Solyc07g063830.2 | Solyc07g063830.3 |
| Solyc07g063850.2 | Solyc07g063850.3 |
| Solyc07g063900.2 | Solyc07g063900.3 |
| Solyc07g063920.2 | Solyc07g063920.3 |
| Solyc07g063940.1 | Solyc07g063940.2 |
| Solyc07g063950.2 | Solyc07g063950.3 |
| Solyc07g063960.2 | Solyc07g063960.3 |
| Solyc07g063970.2 | Solyc07g063970.3 |
| Solyc07g064010.2 | Solyc07g064010.3 |
| Solyc07g064040.2 | Solyc07g064040.3 |
| Solyc07g064050.2 | Solyc07g064050.3 |
| Solyc07g064060.2 | Solyc07g064060.3 |
| Solyc07g064090.2 | Solyc07g064090.3 |
| Solyc07g064110.2 | Solyc07g064110.3 |
| Solyc07g064120.1 | Solyc07g064120.2 |
| Solyc07g064130.1 | Solyc07g064130.2 |
| Solyc07g064160.2 | Solyc07g064160.3 |
| Solyc07g064180.3 | Solyc07g064180.4 |
| Solyc07g064250.2 | Solyc07g064250.3 |
| Solyc07g064270.2 | Solyc07g064270.3 |
| Solyc07g064280.2 | Solyc07g064280.3 |
| Solyc07g064290.2 | Solyc07g064290.3 |
| Solyc07g064310.1 | Solyc07g064310.2 |
| Solyc07g064330.2 | Solyc07g064330.3 |
| Solyc07g064340.2 | Solyc07g064340.3 |
| Solyc07g064400.1 | Solyc07g064400.2 |
| Solyc07g064500.2 | Solyc07g064500.3 |
| Solyc07g064510.2 | Solyc07g064510.3 |
| Solyc07g064520.2 | Solyc07g064520.3 |
| Solyc07g064550.2 | Solyc07g064550.3 |
| Solyc07g064580.2 | Solyc07g064580.3 |
| Solyc07g064590.2 | Solyc07g064590.3 |
| Solyc07g064600.2 | Solyc07g064600.3 |
| Solyc07g064610.2 | Solyc07g064610.3 |
| Solyc07g064620.1 | Solyc07g064620.2 |
| Solyc07g064640.2 | Solyc07g064640.3 |
| Solyc07g064670.2 | Solyc07g064670.3 |
| Solyc07g064680.2 | Solyc07g064680.3 |
| Solyc07g064710.2 | Solyc07g064710.3 |
| Solyc07g064720.2 | Solyc07g064720.3 |

| | |
|---|---|
| Solyc07g064730.1 | Solyc07g064730.2 |
| Solyc07g064740.2 | Solyc07g064740.3 |
| Solyc07g064760.1 | Solyc07g064760.2 |
| Solyc07g064770.1 | Solyc07g064770.2 |
| Solyc07g064780.1 | Solyc07g064780.2 |
| Solyc07g064800.2 | Solyc07g064800.3 |
| Solyc07g064810.2 | Solyc07g064810.3 |
| Solyc07g064830.2 | Solyc07g064830.3 |
| Solyc07g064850.2 | Solyc07g064850.3 |
| Solyc07g064860.1 | Solyc07g064860.2 |
| Solyc07g064870.2 | Solyc07g064870.3 |
| Solyc07g064900.2 | Solyc07g064900.3 |
| Solyc07g064910.2 | Solyc07g064910.3 |
| Solyc07g064920.2 | Solyc07g064920.3 |
| Solyc07g064930.2 | Solyc07g064930.3 |
| Solyc07g064940.2 | Solyc07g064940.3 |
| Solyc07g064950.2 | Solyc07g064950.3 |
| Solyc07g064960.1 | Solyc07g064960.2 |
| Solyc07g064980.2 | Solyc07g064980.3 |
| Solyc07g064990.2 | Solyc07g064990.3 |
| Solyc07g065000.2 | Solyc07g065000.3 |
| Solyc07g065020.2 | Solyc07g065020.3 |
| Solyc07g065030.2 | Solyc07g065030.3 |
| Solyc07g065040.2 | Solyc07g065040.3 |
| Solyc07g065070.1 | Solyc07g065070.2 |
| Solyc07g065120.2 | Solyc07g065120.3 |
| Solyc07g065130.2 | Solyc07g065130.3 |
| Solyc07g065170.2 | Solyc07g065170.3 |
| Solyc07g065180.2 | Solyc07g065180.3 |
| Solyc07g065200.2 | Solyc07g065200.3 |
| Solyc07g065250.2 | Solyc07g065250.3 |
| Solyc07g065290.2 | Solyc07g065290.3 |
| Solyc07g065310.2 | Solyc07g065310.3 |
| Solyc07g065330.2 | Solyc07g065330.3 |
| Solyc07g065350.1 | Solyc07g065350.2 |
| Solyc07g065360.1 | Solyc07g065360.2 |
| Solyc07g065450.2 | Solyc07g065450.3 |
| Solyc07g065490.2 | Solyc07g065490.3 |
| Solyc07g065500.1 | Solyc07g065500.2 |
| Solyc07g065550.2 | Solyc07g065550.3 |
| Solyc07g065600.2 | Solyc07g065600.3 |
| Solyc07g065640.2 | Solyc07g065640.3 |
| Solyc07g065680.2 | Solyc07g065680.3 |

| | |
|---|---|
| Solyc07g065690.2 | Solyc07g065690.3 |
| Solyc07g065710.2 | Solyc07g065710.3 |
| Solyc07g065770.2 | Solyc07g065770.3 |
| Solyc07g065780.1 | Solyc07g065780.2 |
| Solyc07g065800.2 | Solyc07g065800.3 |
| Solyc07g065820.2 | Solyc07g065820.3 |
| Solyc07g065850.2 | Solyc07g065850.3 |
| Solyc07g065860.2 | Solyc07g065860.3 |
| Solyc07g065870.2 | Solyc07g065870.3 |
| Solyc07g065890.2 | Solyc07g065890.3 |
| Solyc07g065900.2 | Solyc07g065900.3 |
| Solyc07g065930.2 | Solyc07g065930.3 |
| Solyc07g065980.2 | Solyc07g065980.3 |
| Solyc07g066000.2 | Solyc07g066000.3 |
| Solyc07g066010.2 | Solyc07g066010.3 |
| Solyc07g066060.2 | Solyc07g066060.3 |
| Solyc07g066070.2 | Solyc07g066070.3 |
| Solyc07g066080.2 | Solyc07g066080.3 |
| Solyc07g066100.2 | Solyc07g066100.3 |
| Solyc07g066120.2 | Solyc07g066120.3 |
| Solyc07g066210.2 | Solyc07g066210.3 |
| Solyc07g066220.2 | Solyc07g066220.3 |
| Solyc07g066240.2 | Solyc07g066240.3 |
| Solyc07g066260.2 | Solyc07g066260.3 |
| Solyc07g066280.2 | Solyc07g066280.3 |
| Solyc07g066290.2 | Solyc07g066290.3 |
| Solyc07g066310.2 | Solyc07g066310.3 |
| Solyc07g066320.2 | Solyc07g066320.3 |
| Solyc07g066330.2 | Solyc07g066330.3 |
| Solyc07g066430.2 | Solyc07g066430.3 |
| Solyc07g066460.2 | Solyc07g066460.3 |
| Solyc07g066470.2 | Solyc07g066470.3 |
| Solyc07g066480.2 | Solyc07g066480.3 |
| Solyc07g066580.2 | Solyc07g066580.3 |
| Solyc07g066600.2 | Solyc07g066600.3 |
| Solyc07g066610.2 | Solyc07g066610.3 |
| Solyc07g066620.2 | Solyc07g066620.3 |
| Solyc08g005020.2 | Solyc08g005020.3 |
| Solyc08g005080.1 | Solyc08g005080.2 |
| Solyc08g005090.1 | Solyc08g005090.2 |
| Solyc08g005100.2 | Solyc08g005100.3 |
| Solyc08g005120.2 | Solyc08g005120.3 |
| Solyc08g005140.2 | Solyc08g005140.3 |

| | |
|---|---|
| Solyc08g005150.2 | Solyc08g005150.3 |
| Solyc08g005170.2 | Solyc08g005170.3 |
| Solyc08g005180.1 | Solyc08g005180.2 |
| Solyc08g005200.2 | Solyc08g005200.3 |
| Solyc08g005210.2 | Solyc08g005210.3 |
| Solyc08g005230.2 | Solyc08g005230.3 |
| Solyc08g005260.1 | Solyc08g005260.2 |
| Solyc08g005270.2 | Solyc08g005270.3 |
| Solyc08g005280.1 | Solyc08g005280.2 |
| Solyc08g005290.2 | Solyc08g005290.3 |
| Solyc08g005320.2 | Solyc08g005320.3 |
| Solyc08g005330.1 | Solyc08g005330.2 |
| Solyc08g005370.1 | Solyc08g005370.2 |
| Solyc08g005380.1 | Solyc08g005380.2 |
| Solyc08g005410.2 | Solyc08g005410.3 |
| Solyc08g005420.2 | Solyc08g005420.3 |
| Solyc08g005430.2 | Solyc08g005430.3 |
| Solyc08g005440.2 | Solyc08g005440.3 |
| Solyc08g005460.2 | Solyc08g005460.3 |
| Solyc08g005480.2 | Solyc08g005480.3 |
| Solyc08g005490.2 | Solyc08g005490.3 |
| Solyc08g005520.2 | Solyc08g005520.3 |
| Solyc08g005525.1 | Solyc08g150101.1 |
| Solyc08g005540.1 | Solyc08g005540.2 |
| Solyc08g005580.2 | Solyc08g005580.3 |
| Solyc08g005610.2 | Solyc08g005610.3 |
| Solyc08g005620.2 | Solyc08g005620.3 |
| Solyc08g005630.2 | Solyc08g005630.3 |
| Solyc08g005650.2 | Solyc08g005650.3 |
| Solyc08g005710.2 | Solyc08g005710.3 |
| Solyc08g005725.1 | Solyc08g150102.1 |
| Solyc08g005770.2 | Solyc08g005770.3 |
| Solyc08g005790.2 | Solyc08g005790.3 |
| Solyc08g005860.2 | Solyc08g005860.3 |
| Solyc08g005870.1 | Solyc08g005870.2 |
| Solyc08g005890.2 | Solyc08g005890.3 |
| Solyc08g005910.2 | Solyc08g005910.3 |
| Solyc08g005920.2 | Solyc08g005920.3 |
| Solyc08g005960.1 | Solyc08g005960.2 |
| Solyc08g005970.2 | Solyc08g005970.3 |
| Solyc08g005980.2 | Solyc08g005980.3 |
| Solyc08g006010.2 | Solyc08g006010.3 |
| Solyc08g006040.2 | Solyc08g006040.3 |

| | |
|---|---|
| Solyc08g006050.1 | Solyc08g006050.2 |
| Solyc08g006060.2 | Solyc08g006060.3 |
| Solyc08g006090.2 | Solyc08g006090.3 |
| Solyc08g006120.2 | Solyc08g006120.3 |
| Solyc08g006150.2 | Solyc08g006150.3 |
| Solyc08g006250.1 | Solyc08g006250.2 |
| Solyc08g006280.1 | Solyc08g006280.2 |
| Solyc08g006310.2 | Solyc08g006310.3 |
| Solyc08g006330.2 | Solyc08g006330.3 |
| Solyc08g006350.2 | Solyc08g006350.3 |
| Solyc08g006460.2 | Solyc08g006460.3 |
| Solyc08g006510.2 | Solyc08g006510.3 |
| Solyc08g006560.2 | Solyc08g006560.3 |
| Solyc08g006580.2 | Solyc08g006580.3 |
| Solyc08g006590.1 | Solyc08g006590.2 |
| Solyc08g006630.2 | Solyc08g006630.3 |
| Solyc08g006640.2 | Solyc08g006640.3 |
| Solyc08g006650.2 | Solyc08g006650.3 |
| Solyc08g006660.1 | Solyc08g006660.2 |
| Solyc08g006670.2 | Solyc08g006670.3 |
| Solyc08g006680.2 | Solyc08g006680.3 |
| Solyc08g006700.2 | Solyc08g006700.3 |
| Solyc08g006710.2 | Solyc08g006710.3 |
| Solyc08g006740.2 | Solyc08g006740.3 |
| Solyc08g006770.2 | Solyc08g006770.3 |
| Solyc08g006780.2 | Solyc08g006780.3 |
| Solyc08g006830.2 | Solyc08g006830.3 |
| Solyc08g006840.2 | Solyc08g006840.3 |
| Solyc08g006860.2 | Solyc08g006860.3 |
| Solyc08g006880.2 | Solyc08g006880.3 |
| Solyc08g006890.2 | Solyc08g006890.3 |
| Solyc08g006900.2 | Solyc08g006900.3 |
| Solyc08g006930.2 | Solyc08g006930.3 |
| Solyc08g006960.2 | Solyc08g006960.3 |
| Solyc08g006980.2 | Solyc08g006980.3 |
| Solyc08g007000.2 | Solyc08g007000.3 |
| Solyc08g007040.2 | Solyc08g007040.3 |
| Solyc08g007050.1 | Solyc08g007050.2 |
| Solyc08g007120.2 | Solyc08g007120.3 |
| Solyc08g007160.2 | Solyc08g007160.3 |
| Solyc08g007180.2 | Solyc08g007180.3 |
| Solyc08g007190.2 | Solyc08g007190.3 |
| Solyc08g007220.2 | Solyc08g007220.3 |

| | |
|---|---|
| Solyc08g007290.2 | Solyc08g007290.3 |
| Solyc08g007310.2 | Solyc08g007310.3 |
| Solyc08g007320.2 | Solyc08g007320.3 |
| Solyc08g007330.2 | Solyc08g007330.3 |
| Solyc08g007340.2 | Solyc08g007340.3 |
| Solyc08g007350.1 | Solyc08g007350.2 |
| Solyc08g007360.2 | Solyc08g007360.3 |
| Solyc08g007370.2 | Solyc08g007370.3 |
| Solyc08g007420.2 | Solyc08g007420.3 |
| Solyc08g007430.1 | Solyc08g007430.2 |
| Solyc08g007435.1 | Solyc08g150103.1 |
| Solyc08g007440.1 | Solyc08g007440.2 |
| Solyc08g007460.2 | Solyc08g007460.3 |
| Solyc08g007470.1 | Solyc08g007470.2 |
| Solyc08g007480.1 | Solyc08g007480.2 |
| Solyc08g007510.2 | Solyc08g007510.3 |
| Solyc08g007540.2 | Solyc08g007540.3 |
| Solyc08g007590.2 | Solyc08g007590.3 |
| Solyc08g007600.1 | Solyc08g007600.2 |
| Solyc08g007620.1 | Solyc08g007620.2 |
| Solyc08g007760.2 | Solyc08g007760.3 |
| Solyc08g007780.2 | Solyc08g007780.3 |
| Solyc08g007790.2 | Solyc08g007790.3 |
| Solyc08g007800.2 | Solyc08g007800.3 |
| Solyc08g007890.2 | Solyc08g007890.3 |
| Solyc08g007920.1 | Solyc08g007920.2 |
| Solyc08g007930.1 | Solyc08g007930.2 |
| Solyc08g007950.2 | Solyc08g007950.3 |
| Solyc08g007970.2 | Solyc08g007970.3 |
| Solyc08g007980.2 | Solyc08g007980.3 |
| Solyc08g007990.2 | Solyc08g007990.3 |
| Solyc08g008000.1 | Solyc08g008000.2 |
| Solyc08g008030.2 | Solyc08g008030.3 |
| Solyc08g008140.2 | Solyc08g008140.3 |
| Solyc08g008210.2 | Solyc08g008210.3 |
| Solyc08g008220.2 | Solyc08g008220.3 |
| Solyc08g008230.2 | Solyc08g008230.3 |
| Solyc08g008240.2 | Solyc08g008240.3 |
| Solyc08g008280.2 | Solyc08g008280.3 |
| Solyc08g008285.1 | Solyc08g150104.1 |
| Solyc08g008290.2 | Solyc08g008290.3 |
| Solyc08g008310.2 | Solyc08g008310.3 |
| Solyc08g008320.2 | Solyc08g008320.3 |

| | |
|---|---|
| Solyc08g008370.2 | Solyc08g008370.3 |
| Solyc08g008410.2 | Solyc08g008410.3 |
| Solyc08g008420.2 | Solyc08g008420.3 |
| Solyc08g008440.1 | Solyc08g008440.2 |
| Solyc08g008470.1 | Solyc08g008470.2 |
| Solyc08g008510.2 | Solyc08g008510.3 |
| Solyc08g008520.2 | Solyc08g008520.3 |
| Solyc08g008580.2 | Solyc08g008580.3 |
| Solyc08g008600.2 | Solyc08g008600.3 |
| Solyc08g008610.2 | Solyc08g008610.3 |
| Solyc08g008640.2 | Solyc08g008640.3 |
| Solyc08g008670.2 | Solyc08g008670.3 |
| Solyc08g013670.2 | Solyc08g013670.3 |
| Solyc08g013730.2 | Solyc08g013730.3 |
| Solyc08g013750.2 | Solyc08g013750.3 |
| Solyc08g013790.1 | Solyc08g013790.2 |
| Solyc08g013805.1 | Solyc08g013805.2 |
| Solyc08g013820.2 | Solyc08g013820.3 |
| Solyc08g013840.2 | Solyc08g013840.3 |
| Solyc08g013845.1 | Solyc08g150105.1 |
| Solyc08g013860.2 | Solyc08g013860.3 |
| Solyc08g013900.2 | Solyc08g013900.3 |
| Solyc08g013920.2 | Solyc08g013920.3 |
| Solyc08g013940.2 | Solyc08g013940.3 |
| Solyc08g013990.2 | Solyc08g013990.3 |
| Solyc08g014000.2 | Solyc08g014000.3 |
| Solyc08g014040.2 | Solyc08g014040.3 |
| Solyc08g014050.1 | Solyc08g014050.2 |
| Solyc08g014120.2 | Solyc08g014120.3 |
| Solyc08g014130.2 | Solyc08g014130.3 |
| Solyc08g014243.1 | Solyc08g150106.1 |
| Solyc08g014247.1 | Solyc08g150107.1 |
| Solyc08g014280.2 | Solyc08g014280.3 |
| Solyc08g014310.2 | Solyc08g014310.3 |
| Solyc08g014330.2 | Solyc08g014330.3 |
| Solyc08g014340.2 | Solyc08g014340.3 |
| Solyc08g014360.1 | Solyc08g014360.2 |
| Solyc08g014430.2 | Solyc08g014430.3 |
| Solyc08g014510.2 | Solyc08g014510.3 |
| Solyc08g014550.2 | Solyc08g014550.3 |
| Solyc08g014570.2 | Solyc08g014570.3 |
| Solyc08g014590.1 | Solyc08g014590.2 |
| Solyc08g015630.2 | Solyc08g015630.3 |

| | |
|---|---|
| Solyc08g015634.1 | Solyc08g150108.1 |
| Solyc08g015650.2 | Solyc08g015650.3 |
| Solyc08g015660.2 | Solyc08g015660.3 |
| Solyc08g015710.1 | Solyc08g015710.2 |
| Solyc08g015780.2 | Solyc08g015780.3 |
| Solyc08g015810.1 | Solyc08g015810.2 |
| Solyc08g015860.2 | Solyc08g015860.3 |
| Solyc08g015870.2 | Solyc08g015870.3 |
| Solyc08g015940.2 | Solyc08g015940.3 |
| Solyc08g015960.2 | Solyc08g015960.3 |
| Solyc08g016000.1 | Solyc08g016000.2 |
| Solyc08g016010.2 | Solyc08g016010.3 |
| Solyc08g016080.2 | Solyc08g016080.3 |
| Solyc08g016160.2 | Solyc08g016160.3 |
| Solyc08g016170.2 | Solyc08g016170.3 |
| Solyc08g016180.2 | Solyc08g016180.3 |
| Solyc08g016210.2 | Solyc08g016210.3 |
| Solyc08g016310.2 | Solyc08g016310.3 |
| Solyc08g016420.2 | Solyc08g016420.3 |
| Solyc08g016500.2 | Solyc08g016500.3 |
| Solyc08g016510.2 | Solyc08g016510.3 |
| Solyc08g016620.2 | Solyc08g016620.3 |
| Solyc08g016660.1 | Solyc08g016660.2 |
| Solyc08g016670.2 | Solyc08g016670.3 |
| Solyc08g016750.2 | Solyc08g016750.3 |
| Solyc08g016793.1 | Solyc08g150109.1 |
| Solyc08g016795.1 | Solyc08g150110.1 |
| Solyc08g016797.1 | Solyc08g150111.1 |
| Solyc08g054575.1 | Solyc08g150120.1 |
| Solyc08g036640.2 | Solyc08g036640.3 |
| Solyc08g036615.1 | Solyc08g150117.1 |
| Solyc08g036505.1 | Solyc08g036505.2 |
| Solyc08g048550.2 | Solyc08g048550.3 |
| Solyc08g048540.1 | Solyc08g048540.2 |
| Solyc08g048500.2 | Solyc08g048500.3 |
| Solyc08g048435.1 | Solyc08g150119.1 |
| Solyc08g048410.1 | Solyc08g048410.2 |
| Solyc08g048230.1 | Solyc08g048230.2 |
| Solyc08g045880.1 | Solyc08g045880.2 |
| Solyc08g045870.2 | Solyc08g045870.3 |
| Solyc08g045850.2 | Solyc08g045850.3 |
| Solyc08g045800.1 | Solyc08g045800.2 |
| Solyc08g045640.2 | Solyc08g045640.3 |

| | |
|---|---|
| Solyc08g044550.1 | Solyc08g044550.2 |
| Solyc08g043180.2 | Solyc08g043180.3 |
| Solyc08g042100.2 | Solyc08g042100.3 |
| Solyc08g042080.2 | Solyc08g042080.3 |
| Solyc08g042070.1 | Solyc08g042070.2 |
| Solyc08g042060.1 | Solyc08g042060.2 |
| Solyc08g042050.2 | Solyc08g042050.3 |
| Solyc08g042000.2 | Solyc08g042000.3 |
| Solyc08g041980.2 | Solyc08g041980.3 |
| Solyc08g041925.1 | Solyc08g150118.1 |
| Solyc08g041880.1 | Solyc08g041880.2 |
| Solyc08g041870.2 | Solyc08g041870.3 |
| Solyc08g041820.2 | Solyc08g041820.3 |
| Solyc08g041780.2 | Solyc08g041780.3 |
| Solyc08g041760.1 | Solyc08g041760.2 |
| Solyc08g041670.1 | Solyc08g041670.2 |
| Solyc08g023660.2 | Solyc08g023660.3 |
| Solyc08g023650.2 | Solyc08g023650.3 |
| Solyc08g023630.2 | Solyc08g023630.3 |
| Solyc08g023570.1 | Solyc08g023570.2 |
| Solyc08g023470.2 | Solyc08g023470.3 |
| Solyc08g023440.2 | Solyc08g023440.3 |
| Solyc08g022220.1 | Solyc08g022220.2 |
| Solyc08g022215.1 | Solyc08g150113.1 |
| Solyc08g022150.1 | Solyc08g022150.2 |
| Solyc08g022070.2 | Solyc08g022070.3 |
| Solyc08g022075.1 | Solyc08g150112.1 |
| Solyc08g022060.1 | Solyc08g022060.2 |
| Solyc08g022050.1 | Solyc08g022050.2 |
| Solyc08g022030.2 | Solyc08g022030.3 |
| Solyc08g021980.1 | Solyc08g021980.2 |
| Solyc08g021960.2 | Solyc08g021960.3 |
| Solyc08g021820.2 | Solyc08g021820.3 |
| Solyc08g029348.1 | Solyc08g150116.1 |
| Solyc08g029280.1 | Solyc08g029280.2 |
| Solyc08g029270.1 | Solyc08g029270.2 |
| Solyc08g029240.2 | Solyc08g029240.3 |
| Solyc08g029200.2 | Solyc08g029200.3 |
| Solyc08g029190.2 | Solyc08g029190.3 |
| Solyc08g029170.2 | Solyc08g029170.3 |
| Solyc08g029130.2 | Solyc08g029130.3 |
| Solyc08g029110.2 | Solyc08g029110.3 |
| Solyc08g029070.1 | Solyc08g029070.2 |

| | |
|---|---|
| Solyc08g029000.2 | Solyc08g029000.3 |
| Solyc08g028970.2 | Solyc08g028970.3 |
| Solyc08g028975.1 | Solyc08g150114.1 |
| Solyc08g028977.1 | Solyc08g150115.1 |
| Solyc08g028700.1 | Solyc08g028700.2 |
| Solyc08g059760.2 | Solyc08g059760.3 |
| Solyc08g059765.1 | Solyc08g150121.1 |
| Solyc08g059800.2 | Solyc08g059800.3 |
| Solyc08g060810.2 | Solyc08g060810.3 |
| Solyc08g060860.2 | Solyc08g060860.3 |
| Solyc08g061090.2 | Solyc08g061090.3 |
| Solyc08g061100.2 | Solyc08g061100.3 |
| Solyc08g061130.2 | Solyc08g061130.3 |
| Solyc08g061140.2 | Solyc08g061140.3 |
| Solyc08g061210.2 | Solyc08g061210.3 |
| Solyc08g061220.2 | Solyc08g061220.3 |
| Solyc08g061250.2 | Solyc08g061250.3 |
| Solyc08g061273.1 | Solyc08g150122.1 |
| Solyc08g061277.1 | Solyc08g150123.1 |
| Solyc08g061320.2 | Solyc08g061320.3 |
| Solyc08g061480.2 | Solyc08g061480.3 |
| Solyc08g061490.2 | Solyc08g061490.3 |
| Solyc08g061520.2 | Solyc08g061520.3 |
| Solyc08g061560.2 | Solyc08g061560.3 |
| Solyc08g061595.1 | Solyc08g150124.1 |
| Solyc08g061630.2 | Solyc08g061630.3 |
| Solyc08g061820.2 | Solyc08g061820.3 |
| Solyc08g061880.1 | Solyc08g061880.2 |
| Solyc08g061903.1 | Solyc08g150125.1 |
| Solyc08g061907.1 | Solyc08g150126.1 |
| Solyc08g061970.2 | Solyc08g061970.3 |
| Solyc08g062040.1 | Solyc08g062040.2 |
| Solyc08g062190.2 | Solyc08g062190.3 |
| Solyc08g062210.2 | Solyc08g062210.3 |
| Solyc08g062220.2 | Solyc08g062220.3 |
| Solyc08g062280.1 | Solyc08g062280.2 |
| Solyc08g062330.2 | Solyc08g062330.3 |
| Solyc08g062340.2 | Solyc08g062340.3 |
| Solyc08g062360.2 | Solyc08g062360.3 |
| Solyc08g062370.2 | Solyc08g062370.3 |
| Solyc08g062420.2 | Solyc08g062420.3 |
| Solyc08g062500.2 | Solyc08g062500.3 |
| Solyc08g062585.1 | Solyc08g150127.1 |

| | |
|---|---|
| Solyc08g062620.2 | Solyc08g062620.3 |
| Solyc08g062630.2 | Solyc08g062630.3 |
| Solyc08g062640.2 | Solyc08g062640.3 |
| Solyc08g062660.2 | Solyc08g062660.3 |
| Solyc08g062670.2 | Solyc08g062670.3 |
| Solyc08g062700.2 | Solyc08g062700.3 |
| Solyc08g062770.2 | Solyc08g062770.3 |
| Solyc08g062780.1 | Solyc08g062780.2 |
| Solyc08g062787.1 | Solyc08g150128.1 |
| Solyc08g062800.2 | Solyc08g062800.3 |
| Solyc08g062820.2 | Solyc08g062820.3 |
| Solyc08g062850.1 | Solyc08g062850.2 |
| Solyc08g062860.2 | Solyc08g062860.3 |
| Solyc08g062920.2 | Solyc08g062920.3 |
| Solyc08g062965.1 | Solyc08g150129.1 |
| Solyc08g063010.2 | Solyc08g063010.3 |
| Solyc08g063050.2 | Solyc08g063050.3 |
| Solyc08g063070.2 | Solyc08g063070.3 |
| Solyc08g063080.2 | Solyc08g063080.3 |
| Solyc08g063123.1 | Solyc08g150130.1 |
| Solyc08g063127.1 | Solyc08g150131.1 |
| Solyc08g065160.2 | Solyc08g065160.3 |
| Solyc08g065167.1 | Solyc08g150132.1 |
| Solyc08g065190.2 | Solyc08g065190.3 |
| Solyc08g065197.1 | Solyc08g150133.1 |
| Solyc08g065220.2 | Solyc08g065220.3 |
| Solyc08g065265.1 | Solyc08g150134.1 |
| Solyc08g065320.2 | Solyc08g065320.3 |
| Solyc08g065330.2 | Solyc08g065330.3 |
| Solyc08g065360.2 | Solyc08g065360.3 |
| Solyc08g065380.2 | Solyc08g065380.3 |
| Solyc08g065420.2 | Solyc08g065420.3 |
| Solyc08g065430.2 | Solyc08g065430.3 |
| Solyc08g065480.2 | Solyc08g065480.3 |
| Solyc08g065490.2 | Solyc08g065490.3 |
| Solyc08g065500.1 | Solyc08g065500.2 |
| Solyc08g065540.2 | Solyc08g065540.3 |
| Solyc08g065610.2 | Solyc08g065610.3 |
| Solyc08g065640.2 | Solyc08g065640.3 |
| Solyc08g065710.1 | Solyc08g065710.2 |
| Solyc08g065730.1 | Solyc08g065730.2 |
| Solyc08g065760.2 | Solyc08g065760.3 |
| Solyc08g065780.1 | Solyc08g065780.2 |

| | |
|---|---|
| Solyc08g065790.2 | Solyc08g065790.3 |
| Solyc08g065840.2 | Solyc08g065840.3 |
| Solyc08g065860.2 | Solyc08g065860.3 |
| Solyc08g065880.2 | Solyc08g065880.3 |
| Solyc08g065890.2 | Solyc08g065890.3 |
| Solyc08g065970.2 | Solyc08g065970.3 |
| Solyc08g065990.2 | Solyc08g065990.3 |
| Solyc08g066000.1 | Solyc08g066000.2 |
| Solyc08g066030.2 | Solyc08g066030.3 |
| Solyc08g066050.2 | Solyc08g066050.3 |
| Solyc08g066060.1 | Solyc08g066060.2 |
| Solyc08g066100.2 | Solyc08g066100.3 |
| Solyc08g066110.2 | Solyc08g066110.3 |
| Solyc08g066140.2 | Solyc08g066140.3 |
| Solyc08g066160.2 | Solyc08g066160.3 |
| Solyc08g066170.2 | Solyc08g066170.3 |
| Solyc08g066180.2 | Solyc08g066180.3 |
| Solyc08g066190.1 | Solyc08g066190.2 |
| Solyc08g066200.2 | Solyc08g066200.3 |
| Solyc08g066210.2 | Solyc08g066210.3 |
| Solyc08g066230.1 | Solyc08g066230.2 |
| Solyc08g066240.2 | Solyc08g066240.3 |
| Solyc08g066260.2 | Solyc08g066260.3 |
| Solyc08g066290.2 | Solyc08g066290.3 |
| Solyc08g066350.1 | Solyc08g066350.2 |
| Solyc08g066360.2 | Solyc08g066360.3 |
| Solyc08g066365.1 | Solyc08g150135.1 |
| Solyc08g066390.2 | Solyc08g066390.3 |
| Solyc08g066440.2 | Solyc08g066440.3 |
| Solyc08g066490.2 | Solyc08g066490.3 |
| Solyc08g066510.2 | Solyc08g066510.3 |
| Solyc08g066520.2 | Solyc08g066520.3 |
| Solyc08g066540.2 | Solyc08g066540.3 |
| Solyc08g066550.2 | Solyc08g066550.3 |
| Solyc08g066560.2 | Solyc08g066560.3 |
| Solyc08g066590.2 | Solyc08g066590.3 |
| Solyc08g066610.2 | Solyc08g066610.3 |
| Solyc08g066620.2 | Solyc08g066620.3 |
| Solyc08g066640.2 | Solyc08g066640.3 |
| Solyc08g066650.2 | Solyc08g066650.3 |
| Solyc08g066670.2 | Solyc08g066670.3 |
| Solyc08g066680.2 | Solyc08g066680.3 |
| Solyc08g066700.2 | Solyc08g066700.3 |

| | |
|---|---|
| Solyc08g066710.1 | Solyc08g066710.2 |
| Solyc08g066730.2 | Solyc08g066730.3 |
| Solyc08g066740.2 | Solyc08g066740.3 |
| Solyc08g066750.2 | Solyc08g066750.3 |
| Solyc08g066770.2 | Solyc08g066770.3 |
| Solyc08g066780.1 | Solyc08g066780.2 |
| Solyc08g066810.2 | Solyc08g066810.3 |
| Solyc08g066820.2 | Solyc08g066820.3 |
| Solyc08g066840.2 | Solyc08g066840.3 |
| Solyc08g066850.2 | Solyc08g066850.3 |
| Solyc08g066920.1 | Solyc08g066920.2 |
| Solyc08g066930.1 | Solyc08g066930.2 |
| Solyc08g066950.2 | Solyc08g066950.3 |
| Solyc08g066960.2 | Solyc08g066960.3 |
| Solyc08g067000.2 | Solyc08g067000.3 |
| Solyc08g067030.2 | Solyc08g067030.3 |
| Solyc08g067050.2 | Solyc08g067050.3 |
| Solyc08g067060.1 | Solyc08g067060.2 |
| Solyc08g067070.2 | Solyc08g067070.3 |
| Solyc08g067120.2 | Solyc08g067120.3 |
| Solyc08g067123.1 | Solyc08g150136.1 |
| Solyc08g067160.2 | Solyc08g067160.3 |
| Solyc08g067185.1 | Solyc08g150137.1 |
| Solyc08g067200.1 | Solyc08g067200.2 |
| Solyc08g067210.2 | Solyc08g067210.3 |
| Solyc08g067290.2 | Solyc08g067290.3 |
| Solyc08g067350.1 | Solyc08g067350.2 |
| Solyc08g067360.2 | Solyc08g067360.3 |
| Solyc08g067410.1 | Solyc08g067410.2 |
| Solyc08g067440.1 | Solyc08g067440.2 |
| Solyc08g067470.2 | Solyc08g067470.3 |
| Solyc08g067560.1 | Solyc08g067560.2 |
| Solyc08g067610.2 | Solyc08g067610.3 |
| Solyc08g067650.1 | Solyc08g067650.2 |
| Solyc08g067670.2 | Solyc08g067670.3 |
| Solyc08g067690.1 | Solyc08g067690.2 |
| Solyc08g067730.1 | Solyc08g067730.2 |
| Solyc08g067770.2 | Solyc08g067770.3 |
| Solyc08g083450.1 | Solyc08g083450.2 |
| Solyc08g067795.1 | Solyc08g067795.2 |
| Solyc08g067810.2 | Solyc08g067810.3 |
| Solyc08g067820.2 | Solyc08g067820.3 |
| Solyc08g067830.2 | Solyc08g067830.3 |

| | |
|---|---|
| Solyc08g067840.2 | Solyc08g067840.3 |
| Solyc08g067910.2 | Solyc08g067910.3 |
| Solyc08g067920.2 | Solyc08g067920.3 |
| Solyc08g067940.2 | Solyc08g067940.3 |
| Solyc08g067950.2 | Solyc08g067950.3 |
| Solyc08g067960.2 | Solyc08g067960.3 |
| Solyc08g067970.2 | Solyc08g067970.3 |
| Solyc08g068050.2 | Solyc08g068050.3 |
| Solyc08g068070.2 | Solyc08g068070.3 |
| Solyc08g068090.2 | Solyc08g068090.3 |
| Solyc08g068100.2 | Solyc08g068100.3 |
| Solyc08g068160.1 | Solyc08g068160.2 |
| Solyc08g068180.2 | Solyc08g068180.3 |
| Solyc08g068190.2 | Solyc08g068190.3 |
| Solyc08g068210.2 | Solyc08g068210.3 |
| Solyc08g068220.2 | Solyc08g068220.3 |
| Solyc08g068270.2 | Solyc08g068270.3 |
| Solyc08g068280.1 | Solyc08g068280.2 |
| Solyc08g068290.2 | Solyc08g068290.3 |
| Solyc08g068300.2 | Solyc08g068300.3 |
| Solyc08g068310.2 | Solyc08g068310.3 |
| Solyc08g068320.2 | Solyc08g068320.3 |
| Solyc08g068330.2 | Solyc08g068330.3 |
| Solyc08g068340.2 | Solyc08g068340.3 |
| Solyc08g068370.2 | Solyc08g068370.3 |
| Solyc08g068380.2 | Solyc08g068380.3 |
| Solyc08g068390.2 | Solyc08g068390.3 |
| Solyc08g068420.2 | Solyc08g068420.3 |
| Solyc08g068490.2 | Solyc08g068490.3 |
| Solyc08g068500.1 | Solyc08g068500.2 |
| Solyc08g068510.1 | Solyc08g068510.2 |
| Solyc08g068520.2 | Solyc08g068520.3 |
| Solyc08g068600.2 | Solyc08g068600.3 |
| Solyc08g068610.2 | Solyc08g068610.3 |
| Solyc08g068640.2 | Solyc08g068640.4 |
| Solyc08g068740.1 | Solyc08g068740.2 |
| Solyc08g068810.2 | Solyc08g068810.3 |
| Solyc08g068820.2 | Solyc08g068820.3 |
| Solyc08g068840.2 | Solyc08g068840.3 |
| Solyc08g068850.2 | Solyc08g068850.3 |
| Solyc08g068860.2 | Solyc08g068860.3 |
| Solyc08g068870.2 | Solyc08g068870.3 |
| Solyc08g068960.2 | Solyc08g068960.3 |

| | |
|---|---|
| Solyc08g068970.2 | Solyc08g068970.3 |
| Solyc08g068990.1 | Solyc08g068990.2 |
| Solyc08g069040.2 | Solyc08g069040.3 |
| Solyc08g069050.2 | Solyc08g069050.3 |
| Solyc08g069110.2 | Solyc08g069110.3 |
| Solyc08g069125.1 | Solyc08g150139.1 |
| Solyc08g069160.2 | Solyc08g069160.3 |
| Solyc08g069180.2 | Solyc08g069180.3 |
| Solyc08g069200.1 | Solyc08g069200.2 |
| Solyc08g069220.1 | Solyc08g069220.2 |
| Solyc08g074240.2 | Solyc08g074240.3 |
| Solyc08g074260.2 | Solyc08g074260.3 |
| Solyc08g074290.2 | Solyc08g074290.3 |
| Solyc08g074307.1 | Solyc08g074307.2 |
| Solyc08g074360.2 | Solyc08g074360.3 |
| Solyc08g074390.2 | Solyc08g074390.3 |
| Solyc08g074420.2 | Solyc08g074420.3 |
| Solyc08g074450.2 | Solyc08g074450.3 |
| Solyc08g074520.1 | Solyc08g074520.2 |
| Solyc08g074550.2 | Solyc08g074550.3 |
| Solyc08g074560.2 | Solyc08g074560.3 |
| Solyc08g074670.2 | Solyc08g074670.3 |
| Solyc08g074690.2 | Solyc08g074690.3 |
| Solyc08g074715.1 | Solyc08g150140.1 |
| Solyc08g074730.1 | Solyc08g074730.2 |
| Solyc08g074740.2 | Solyc08g074740.3 |
| Solyc08g074760.2 | Solyc08g074760.3 |
| Solyc08g074780.2 | Solyc08g074780.3 |
| Solyc08g074790.2 | Solyc08g074790.3 |
| Solyc08g074800.2 | Solyc08g074800.3 |
| Solyc08g074865.1 | Solyc08g150141.1 |
| Solyc08g074910.2 | Solyc08g074910.3 |
| Solyc08g074950.1 | Solyc08g074950.2 |
| Solyc08g074960.2 | Solyc08g074960.3 |
| Solyc08g075020.2 | Solyc08g075020.3 |
| Solyc08g075030.2 | Solyc08g075030.3 |
| Solyc08g075080.2 | Solyc08g075080.3 |
| Solyc08g075120.2 | Solyc08g075120.3 |
| Solyc08g075160.2 | Solyc08g075160.3 |
| Solyc08g075165.1 | Solyc08g150142.1 |
| Solyc08g075210.1 | Solyc08g075210.2 |
| Solyc08g075240.2 | Solyc08g075240.3 |
| Solyc08g075340.2 | Solyc08g075340.3 |

| | |
|---|---|
| Solyc08g075353.1 | Solyc08g150143.1 |
| Solyc08g075357.1 | Solyc08g150144.1 |
| Solyc08g075370.2 | Solyc08g075370.3 |
| Solyc08g075380.2 | Solyc08g075380.3 |
| Solyc08g075400.2 | Solyc08g075400.3 |
| Solyc08g075410.2 | Solyc08g075410.3 |
| Solyc08g075430.2 | Solyc08g075430.3 |
| Solyc08g075450.2 | Solyc08g075450.3 |
| Solyc08g075460.2 | Solyc08g075460.3 |
| Solyc08g075470.2 | Solyc08g075470.3 |
| Solyc08g075530.2 | Solyc08g075530.3 |
| Solyc08g075540.4 | Solyc08g075540.5.1 |
| Solyc08g075550.3 | Solyc08g075550.4 |
| Solyc08g075590.1 | Solyc08g075590.2 |
| Solyc08g075630.2 | Solyc08g075630.3 |
| Solyc08g075650.2 | Solyc08g075650.3 |
| Solyc08g075670.1 | Solyc08g075670.2 |
| Solyc08g075690.2 | Solyc08g075690.3 |
| Solyc08g075700.2 | Solyc08g075700.3 |
| Solyc08g075710.2 | Solyc08g075710.3 |
| Solyc08g075730.2 | Solyc08g075730.3 |
| Solyc08g075760.2 | Solyc08g075760.3 |
| Solyc08g075810.2 | Solyc08g075810.3 |
| Solyc08g075840.2 | Solyc08g075840.3 |
| Solyc08g075850.2 | Solyc08g075850.3 |
| Solyc08g075870.2 | Solyc08g075870.3 |
| Solyc08g075890.2 | Solyc08g075890.3 |
| Solyc08g075960.2 | Solyc08g075960.3 |
| Solyc08g075970.2 | Solyc08g075970.3 |
| Solyc08g075980.1 | Solyc08g075980.2 |
| Solyc08g075990.2 | Solyc08g075990.3 |
| Solyc08g076030.2 | Solyc08g076030.3 |
| Solyc08g076040.1 | Solyc08g076040.2 |
| Solyc08g076060.2 | Solyc08g076060.3 |
| Solyc08g076080.2 | Solyc08g076080.3 |
| Solyc08g076090.2 | Solyc08g076090.3 |
| Solyc08g076120.2 | Solyc08g076120.3 |
| Solyc08g076140.2 | Solyc08g076140.3 |
| Solyc08g076160.2 | Solyc08g076160.3 |
| Solyc08g076210.2 | Solyc08g076210.3 |
| Solyc08g076220.2 | Solyc08g076220.3 |
| Solyc08g076290.2 | Solyc08g076290.3 |
| Solyc08g076300.2 | Solyc08g076300.3 |

| | |
|---|---|
| Solyc08g076310.2 | Solyc08g076310.3 |
| Solyc08g076320.2 | Solyc08g076320.3 |
| Solyc08g076340.2 | Solyc08g076340.3 |
| Solyc08g076350.2 | Solyc08g076350.3 |
| Solyc08g076370.2 | Solyc08g076370.3 |
| Solyc08g076390.2 | Solyc08g076390.3 |
| Solyc08g076400.2 | Solyc08g076400.3 |
| Solyc08g076420.2 | Solyc08g076420.3 |
| Solyc08g076430.2 | Solyc08g076430.3 |
| Solyc08g076450.2 | Solyc08g076450.3 |
| Solyc08g076460.2 | Solyc08g076460.3 |
| Solyc08g076480.2 | Solyc08g076480.3 |
| Solyc08g076510.2 | Solyc08g076510.3 |
| Solyc08g076530.2 | Solyc08g076530.3 |
| Solyc08g076540.2 | Solyc08g076540.3 |
| Solyc08g076570.2 | Solyc08g076570.3 |
| Solyc08g076590.2 | Solyc08g076590.3 |
| Solyc08g076645.1 | Solyc08g150145.1 |
| Solyc08g076650.2 | Solyc08g076650.3 |
| Solyc08g076660.2 | Solyc08g076660.3 |
| Solyc08g076670.2 | Solyc08g076670.3 |
| Solyc08g076685.1 | Solyc08g150146.1 |
| Solyc08g076710.2 | Solyc08g076710.3 |
| Solyc08g076780.1 | Solyc08g076780.2 |
| Solyc08g076790.2 | Solyc08g076790.3 |
| Solyc08g076820.2 | Solyc08g076820.3 |
| Solyc08g076840.2 | Solyc08g076840.3 |
| Solyc08g076850.2 | Solyc08g076850.3 |
| Solyc08g076860.2 | Solyc08g076860.3 |
| Solyc08g076880.2 | Solyc08g076880.3 |
| Solyc08g076910.2 | Solyc08g076910.3 |
| Solyc08g076920.2 | Solyc08g076920.3 |
| Solyc08g076950.2 | Solyc08g076950.3 |
| Solyc08g076970.2 | Solyc08g076970.3 |
| Solyc08g076990.2 | Solyc08g076990.3 |
| Solyc08g077050.2 | Solyc08g077050.3 |
| Solyc08g077060.2 | Solyc08g077060.3 |
| Solyc08g077070.2 | Solyc08g077070.3 |
| Solyc08g077090.2 | Solyc08g077090.3 |
| Solyc08g077100.2 | Solyc08g077100.3 |
| Solyc08g077150.2 | Solyc08g077150.3 |
| Solyc08g077170.2 | Solyc08g077170.3 |
| Solyc08g077180.2 | Solyc08g077180.3 |

| | |
|---|---|
| Solyc08g077220.2 | Solyc08g077220.3 |
| Solyc08g077230.2 | Solyc08g077230.3 |
| Solyc08g077240.2 | Solyc08g077240.3 |
| Solyc08g077260.2 | Solyc08g077260.3 |
| Solyc08g077300.2 | Solyc08g077300.3 |
| Solyc08g077310.2 | Solyc08g077310.3 |
| Solyc08g077330.2 | Solyc08g077330.3 |
| Solyc08g077350.2 | Solyc08g077350.3 |
| Solyc08g077360.2 | Solyc08g077360.3 |
| Solyc08g077420.2 | Solyc08g077420.3 |
| Solyc08g077440.2 | Solyc08g077440.3 |
| Solyc08g077460.2 | Solyc08g077460.3 |
| Solyc08g077490.1 | Solyc08g077490.2 |
| Solyc08g077520.2 | Solyc08g077520.3 |
| Solyc08g077530.2 | Solyc08g077530.3 |
| Solyc08g077540.1 | Solyc08g077540.2 |
| Solyc08g077550.2 | Solyc08g077550.3 |
| Solyc08g077580.1 | Solyc08g077580.2 |
| Solyc08g077630.2 | Solyc08g077630.3 |
| Solyc08g077680.2 | Solyc08g077680.3 |
| Solyc08g077690.2 | Solyc08g077690.3 |
| Solyc08g077700.2 | Solyc08g077700.3 |
| Solyc08g077710.2 | Solyc08g077710.3 |
| Solyc08g077720.2 | Solyc08g077720.3 |
| Solyc08g077730.2 | Solyc08g077730.3 |
| Solyc08g077760.2 | Solyc08g077760.3 |
| Solyc08g077770.2 | Solyc08g077770.3 |
| Solyc08g077780.2 | Solyc08g077780.3 |
| Solyc08g077790.2 | Solyc08g077790.3 |
| Solyc08g077810.1 | Solyc08g077810.2 |
| Solyc08g077840.2 | Solyc08g077840.3 |
| Solyc08g077850.1 | Solyc08g077850.2 |
| Solyc08g077880.2 | Solyc08g077880.3 |
| Solyc08g077890.2 | Solyc08g077890.3 |
| Solyc08g077900.2 | Solyc08g077900.3 |
| Solyc08g077910.2 | Solyc08g077910.3 |
| Solyc08g077920.2 | Solyc08g077920.3 |
| Solyc08g077930.2 | Solyc08g077930.3 |
| Solyc08g077940.1 | Solyc08g077940.2 |
| Solyc08g077960.2 | Solyc08g077960.3 |
| Solyc08g077990.2 | Solyc08g077990.3 |
| Solyc08g078000.2 | Solyc08g078000.3 |
| Solyc08g078010.2 | Solyc08g078010.3 |

| | |
|---|---|
| Solyc08g078060.2 | Solyc08g078060.3 |
| Solyc08g078120.1 | Solyc08g078120.2 |
| Solyc08g078140.2 | Solyc08g078140.3 |
| Solyc08g078160.2 | Solyc08g078160.3 |
| Solyc08g078190.1 | Solyc08g078190.2 |
| Solyc08g078200.1 | Solyc08g078200.2 |
| Solyc08g078210.2 | Solyc08g078210.3 |
| Solyc08g078270.2 | Solyc08g078270.3 |
| Solyc08g078310.2 | Solyc08g078310.3 |
| Solyc08g078320.2 | Solyc08g078320.3 |
| Solyc08g078340.2 | Solyc08g078340.3 |
| Solyc08g078380.2 | Solyc08g078380.3 |
| Solyc08g078390.3 | Solyc08g078390.4 |
| Solyc08g078410.1 | Solyc08g078410.2 |
| Solyc08g078420.1 | Solyc08g078420.2 |
| Solyc08g078450.2 | Solyc08g078450.3 |
| Solyc08g078460.2 | Solyc08g078460.3 |
| Solyc08g078510.2 | Solyc08g078510.3 |
| Solyc08g078540.2 | Solyc08g078540.3 |
| Solyc08g078600.2 | Solyc08g078600.3 |
| Solyc08g078630.3 | Solyc08g078630.4 |
| Solyc08g078650.2 | Solyc08g078650.3 |
| Solyc08g078660.2 | Solyc08g078660.3 |
| Solyc08g078690.2 | Solyc08g078690.3 |
| Solyc08g078720.2 | Solyc08g078720.3 |
| Solyc08g078820.2 | Solyc08g078820.3 |
| Solyc08g078840.2 | Solyc08g078840.3 |
| Solyc08g078850.2 | Solyc08g078850.3 |
| Solyc08g078880.2 | Solyc08g078880.3 |
| Solyc08g078920.1 | Solyc08g078920.2 |
| Solyc08g078950.2 | Solyc08g078950.3 |
| Solyc08g079020.2 | Solyc08g079020.3 |
| Solyc08g079050.1 | Solyc08g079050.2 |
| Solyc08g079100.2 | Solyc08g079100.3 |
| Solyc08g079160.2 | Solyc08g079160.3 |
| Solyc08g079170.2 | Solyc08g079170.3 |
| Solyc08g079180.2 | Solyc08g079180.3 |
| Solyc08g079260.2 | Solyc08g079260.3 |
| Solyc08g079270.2 | Solyc08g079270.3 |
| Solyc08g079280.2 | Solyc08g079280.3 |
| Solyc08g079300.2 | Solyc08g079300.3 |
| Solyc08g079330.1 | Solyc08g079330.2 |
| Solyc08g083520.1 | Solyc08g083520.2 |

| | |
|---|---|
| Solyc08g079370.1 | Solyc08g079370.2 |
| Solyc08g083500.1 | Solyc08g083500.2 |
| Solyc08g079420.2 | Solyc08g079420.3 |
| Solyc08g079430.2 | Solyc08g079430.3 |
| Solyc08g079450.2 | Solyc08g079450.3 |
| Solyc08g079460.2 | Solyc08g079460.3 |
| Solyc08g079470.2 | Solyc08g079470.3 |
| Solyc08g079480.2 | Solyc08g079480.3 |
| Solyc08g079500.2 | Solyc08g079500.3 |
| Solyc08g079510.2 | Solyc08g079510.3 |
| Solyc08g079520.2 | Solyc08g079520.3 |
| Solyc08g079530.2 | Solyc08g079530.3 |
| Solyc08g079570.2 | Solyc08g079570.3 |
| Solyc08g079610.2 | Solyc08g079610.3 |
| Solyc08g079620.2 | Solyc08g079620.3 |
| Solyc08g079630.2 | Solyc08g079630.3 |
| Solyc08g079650.2 | Solyc08g079650.3 |
| Solyc08g079660.2 | Solyc08g079660.3 |
| Solyc08g079670.2 | Solyc08g079670.3 |
| Solyc08g079680.2 | Solyc08g079680.3 |
| Solyc08g079710.2 | Solyc08g079710.3 |
| Solyc08g079720.2 | Solyc08g079720.3 |
| Solyc08g079730.1 | Solyc08g079730.2 |
| Solyc08g079740.2 | Solyc08g079740.3 |
| Solyc08g079750.2 | Solyc08g079750.3 |
| Solyc08g079770.2 | Solyc08g079770.3 |
| Solyc08g079780.1 | Solyc08g079780.2 |
| Solyc08g079820.2 | Solyc08g079820.3 |
| Solyc08g079970.1 | Solyc08g079970.2 |
| Solyc08g080000.2 | Solyc08g080000.3 |
| Solyc08g080090.2 | Solyc08g080090.3 |
| Solyc08g080110.2 | Solyc08g080110.3 |
| Solyc08g080120.2 | Solyc08g080120.3 |
| Solyc08g080130.2 | Solyc08g080130.3 |
| Solyc08g080170.2 | Solyc08g080170.3 |
| Solyc08g080240.2 | Solyc08g080240.3 |
| Solyc08g080270.2 | Solyc08g080270.3 |
| Solyc08g080310.1 | Solyc08g080310.2 |
| Solyc08g080320.2 | Solyc08g080320.3 |
| Solyc08g080340.2 | Solyc08g080340.3 |
| Solyc08g080360.2 | Solyc08g080360.3 |
| Solyc08g080370.2 | Solyc08g080370.3 |
| Solyc08g080390.2 | Solyc08g080390.3 |

| | |
|---|---|
| Solyc08g080400.1 | Solyc08g080400.2 |
| Solyc08g080410.2 | Solyc08g080410.3 |
| Solyc08g080430.1 | Solyc08g080430.2 |
| Solyc08g080470.2 | Solyc08g080470.3 |
| Solyc08g080480.2 | Solyc08g080480.3 |
| Solyc08g080490.2 | Solyc08g080490.3 |
| Solyc08g080500.2 | Solyc08g080500.3 |
| Solyc08g080540.2 | Solyc08g080540.3 |
| Solyc08g080590.2 | Solyc08g080590.3 |
| Solyc08g080640.1 | Solyc08g080640.2 |
| Solyc08g080650.1 | Solyc08g080650.3 |
| Solyc08g080730.2 | Solyc08g080730.3 |
| Solyc08g080740.2 | Solyc08g080740.3 |
| Solyc08g080745.1 | Solyc08g080745.2 |
| Solyc08g080760.1 | Solyc08g080760.2 |
| Solyc08g080770.2 | Solyc08g080770.3 |
| Solyc08g080830.2 | Solyc08g080830.3 |
| Solyc08g080870.2 | Solyc08g080870.3 |
| Solyc08g080890.2 | Solyc08g080890.3 |
| Solyc08g080900.2 | Solyc08g080900.3 |
| Solyc08g080910.2 | Solyc08g080910.3 |
| Solyc08g080920.2 | Solyc08g080920.3 |
| Solyc08g080940.2 | Solyc08g080940.3 |
| Solyc08g080990.2 | Solyc08g080990.3 |
| Solyc08g081000.2 | Solyc08g081000.3 |
| Solyc08g081010.2 | Solyc08g081010.3 |
| Solyc08g081040.2 | Solyc08g081040.3 |
| Solyc08g081060.2 | Solyc08g081060.3 |
| Solyc08g081070.2 | Solyc08g081070.3 |
| Solyc08g081080.2 | Solyc08g081080.3 |
| Solyc08g081160.2 | Solyc08g081160.3 |
| Solyc08g081180.2 | Solyc08g081180.3 |
| Solyc08g081190.2 | Solyc08g081190.3 |
| Solyc08g081200.2 | Solyc08g081200.3 |
| Solyc08g081240.2 | Solyc08g081240.3 |
| Solyc08g081270.2 | Solyc08g081270.3 |
| Solyc08g081280.2 | Solyc08g081280.3 |
| Solyc08g081320.2 | Solyc08g081320.3 |
| Solyc08g081360.2 | Solyc08g081360.3 |
| Solyc08g081380.2 | Solyc08g081380.3 |
| Solyc08g081410.2 | Solyc08g081410.3 |
| Solyc08g081430.2 | Solyc08g081430.3 |
| Solyc08g081460.1 | Solyc08g081460.2 |

| | |
|---|---|
| Solyc08g081470.2 | Solyc08g081470.3 |
| Solyc08g081480.2 | Solyc08g081480.3 |
| Solyc08g081530.2 | Solyc08g081530.3 |
| Solyc08g081570.2 | Solyc08g081570.3 |
| Solyc08g081630.1 | Solyc08g081630.2 |
| Solyc08g081640.2 | Solyc08g081640.3 |
| Solyc08g081680.1 | Solyc08g081680.2 |
| Solyc08g081690.2 | Solyc08g081690.3 |
| Solyc08g081740.2 | Solyc08g081740.3 |
| Solyc08g081750.2 | Solyc08g081750.3 |
| Solyc08g081770.2 | Solyc08g081770.3 |
| Solyc08g081810.2 | Solyc08g081810.3 |
| Solyc08g081830.2 | Solyc08g081830.3 |
| Solyc08g081840.2 | Solyc08g081840.3 |
| Solyc08g081870.2 | Solyc08g081870.3 |
| Solyc08g081910.2 | Solyc08g081910.3 |
| Solyc08g081930.2 | Solyc08g081930.3 |
| Solyc08g081940.2 | Solyc08g081940.3 |
| Solyc08g081970.2 | Solyc08g081970.3 |
| Solyc08g081990.2 | Solyc08g081990.3 |
| Solyc08g082010.2 | Solyc08g082010.3 |
| Solyc08g082020.2 | Solyc08g082020.3 |
| Solyc08g082060.2 | Solyc08g082060.3 |
| Solyc08g082070.2 | Solyc08g082070.3 |
| Solyc08g082080.2 | Solyc08g082080.3 |
| Solyc08g082100.2 | Solyc08g082100.3 |
| Solyc08g082130.2 | Solyc08g082130.3 |
| Solyc08g082150.1 | Solyc08g082150.2 |
| Solyc08g082165.1 | Solyc08g150147.1 |
| Solyc08g082180.2 | Solyc08g082180.3 |
| Solyc08g082190.2 | Solyc08g082190.3 |
| Solyc08g082200.2 | Solyc08g082200.3 |
| Solyc08g082250.2 | Solyc08g082250.3 |
| Solyc08g082260.1 | Solyc08g082260.2 |
| Solyc08g082270.2 | Solyc08g082270.3 |
| Solyc08g082300.2 | Solyc08g082300.3 |
| Solyc08g082310.2 | Solyc08g082310.3 |
| Solyc08g082330.2 | Solyc08g082330.3 |
| Solyc08g082340.2 | Solyc08g082340.3 |
| Solyc08g082345.1 | Solyc08g150148.1 |
| Solyc08g082440.2 | Solyc08g082440.3 |
| Solyc08g082450.2 | Solyc08g082450.3 |
| Solyc08g082480.2 | Solyc08g082480.3 |

| | |
|---|---|
| Solyc08g082500.1 | Solyc08g082500.2 |
| Solyc08g082540.2 | Solyc08g082540.3 |
| Solyc08g082560.2 | Solyc08g082560.3 |
| Solyc08g082580.2 | Solyc08g082580.3 |
| Solyc08g082590.2 | Solyc08g082590.3 |
| Solyc08g082620.2 | Solyc08g082620.3 |
| Solyc08g082630.2 | Solyc08g082630.3 |
| Solyc08g082680.2 | Solyc08g082680.3 |
| Solyc08g082700.2 | Solyc08g082700.3 |
| Solyc08g082710.2 | Solyc08g082710.3 |
| Solyc08g082750.2 | Solyc08g082750.3 |
| Solyc08g082770.2 | Solyc08g082770.3 |
| Solyc08g082780.1 | Solyc08g082780.2 |
| Solyc08g082790.2 | Solyc08g082790.3 |
| Solyc08g082800.2 | Solyc08g082800.3 |
| Solyc08g082830.2 | Solyc08g082830.3 |
| Solyc08g082850.2 | Solyc08g082850.3 |
| Solyc08g082870.2 | Solyc08g082870.3 |
| Solyc08g082890.2 | Solyc08g082890.3 |
| Solyc08g082910.1 | Solyc08g082910.2 |
| Solyc08g082920.2 | Solyc08g082920.3 |
| Solyc08g082960.2 | Solyc08g082960.3 |
| Solyc08g082990.2 | Solyc08g082990.3 |
| Solyc08g083000.1 | Solyc08g083000.2 |
| Solyc08g083060.2 | Solyc08g083060.3 |
| Solyc08g083070.2 | Solyc08g083070.3 |
| Solyc08g083080.2 | Solyc08g083080.3 |
| Solyc08g083120.2 | Solyc08g083120.3 |
| Solyc08g083175.1 | Solyc08g150149.1 |
| Solyc08g083190.2 | Solyc08g083190.3 |
| Solyc08g083210.2 | Solyc08g083210.3 |
| Solyc08g083215.1 | Solyc08g150150.1 |
| Solyc08g083240.2 | Solyc08g083240.3 |
| Solyc08g083260.2 | Solyc08g083260.3 |
| Solyc08g083280.2 | Solyc08g083280.3 |
| Solyc08g083310.2 | Solyc08g083310.3 |
| Solyc08g083360.2 | Solyc08g083360.3 |
| Solyc09g005110.2 | Solyc09g005110.3 |
| Solyc09g005120.2 | Solyc09g005120.3 |
| Solyc09g005230.2 | Solyc09g005230.3 |
| Solyc09g005250.2 | Solyc09g005250.3 |
| Solyc09g005270.2 | Solyc09g005270.3 |
| Solyc09g005280.2 | Solyc09g005280.3 |

| | |
|---|---|
| Solyc09g005300.2 | Solyc09g005300.3 |
| Solyc09g005310.2 | Solyc09g005310.3 |
| Solyc09g005320.2 | Solyc09g005320.3 |
| Solyc09g005330.2 | Solyc09g005330.3 |
| Solyc09g005340.2 | Solyc09g005340.3 |
| Solyc09g005350.2 | Solyc09g005350.3 |
| Solyc09g005360.2 | Solyc09g005360.3 |
| Solyc09g005380.1 | Solyc09g005380.2 |
| Solyc09g005390.2 | Solyc09g005390.3 |
| Solyc09g005400.2 | Solyc09g005400.3 |
| Solyc09g005470.2 | Solyc09g005470.3 |
| Solyc09g005480.2 | Solyc09g005480.3 |
| Solyc09g005500.2 | Solyc09g005500.3 |
| Solyc09g005530.2 | Solyc09g005530.3 |
| Solyc09g005550.2 | Solyc09g005550.3 |
| Solyc09g005590.2 | Solyc09g005590.3 |
| Solyc09g005600.2 | Solyc09g005600.3 |
| Solyc09g005630.2 | Solyc09g005630.3 |
| Solyc09g005640.2 | Solyc09g005640.3 |
| Solyc09g005690.2 | Solyc09g005690.3 |
| Solyc09g005700.2 | Solyc09g005700.3 |
| Solyc09g005720.2 | Solyc09g005720.3 |
| Solyc09g005750.2 | Solyc09g005750.3 |
| Solyc09g005760.2 | Solyc09g005760.3 |
| Solyc09g005765.1 | Solyc09g150100.1 |
| Solyc09g005840.1 | Solyc09g005840.2 |
| Solyc09g005850.2 | Solyc09g005850.3 |
| Solyc09g005860.2 | Solyc09g005860.3 |
| Solyc09g005950.2 | Solyc09g005950.3 |
| Solyc09g007030.2 | Solyc09g007030.3 |
| Solyc09g007080.2 | Solyc09g007080.3 |
| Solyc09g007100.2 | Solyc09g007100.3 |
| Solyc09g007110.2 | Solyc09g007110.3 |
| Solyc09g007130.2 | Solyc09g007130.3 |
| Solyc09g007150.2 | Solyc09g007150.3 |
| Solyc09g007180.2 | Solyc09g007180.3 |
| Solyc09g007190.2 | Solyc09g007190.3 |
| Solyc09g007230.2 | Solyc09g007230.3 |
| Solyc09g007270.2 | Solyc09g007270.3 |
| Solyc09g007310.2 | Solyc09g007310.3 |
| Solyc09g007350.2 | Solyc09g007350.3 |
| Solyc09g007360.2 | Solyc09g007360.3 |
| Solyc09g007380.2 | Solyc09g007380.3 |

| | |
|---|---|
| Solyc09g007390.2 | Solyc09g007390.3 |
| Solyc09g007410.2 | Solyc09g007410.3 |
| Solyc09g007440.2 | Solyc09g007440.3 |
| Solyc09g007450.2 | Solyc09g007450.3 |
| Solyc09g007470.1 | Solyc09g007470.2 |
| Solyc09g007490.2 | Solyc09g007490.3 |
| Solyc09g007500.2 | Solyc09g007500.3 |
| Solyc09g007520.2 | Solyc09g007520.3 |
| Solyc09g007530.2 | Solyc09g007530.3 |
| Solyc09g007535.1 | Solyc09g150101.1 |
| Solyc09g007540.2 | Solyc09g007540.3 |
| Solyc09g007550.2 | Solyc09g007550.3 |
| Solyc09g007620.1 | Solyc09g007620.2 |
| Solyc09g007630.2 | Solyc09g007630.3 |
| Solyc09g007670.2 | Solyc09g007670.3 |
| Solyc09g007700.2 | Solyc09g007700.3 |
| Solyc09g007710.2 | Solyc09g007710.3 |
| Solyc09g007770.2 | Solyc09g007770.3 |
| Solyc09g007820.1 | Solyc09g007820.2 |
| Solyc09g007830.2 | Solyc09g007830.3 |
| Solyc09g007840.2 | Solyc09g007840.3 |
| Solyc09g007850.2 | Solyc09g007850.3 |
| Solyc09g007870.2 | Solyc09g007870.3 |
| Solyc09g007940.2 | Solyc09g007940.3 |
| Solyc09g007990.2 | Solyc09g007990.3 |
| Solyc09g008000.2 | Solyc09g008000.3 |
| Solyc09g008010.2 | Solyc09g008010.3 |
| Solyc09g008040.2 | Solyc09g008040.3 |
| Solyc09g008050.2 | Solyc09g008050.3 |
| Solyc09g008080.2 | Solyc09g008080.3 |
| Solyc09g008090.2 | Solyc09g008090.3 |
| Solyc09g008130.2 | Solyc09g008130.3 |
| Solyc09g008140.2 | Solyc09g008140.3 |
| Solyc09g008190.2 | Solyc09g008190.3 |
| Solyc09g008230.2 | Solyc09g008230.3 |
| Solyc09g008240.2 | Solyc09g008240.3 |
| Solyc09g008280.1 | Solyc09g008280.2 |
| Solyc09g008310.2 | Solyc09g008310.3 |
| Solyc09g008330.2 | Solyc09g008330.3 |
| Solyc09g008350.2 | Solyc09g008350.3 |
| Solyc09g008380.2 | Solyc09g008380.3 |
| Solyc09g008400.2 | Solyc09g008400.3 |
| Solyc09g008410.2 | Solyc09g008410.3 |

| | |
|---|---|
| Solyc09g008420.2 | Solyc09g008420.3 |
| Solyc09g008450.2 | Solyc09g008450.3 |
| Solyc09g008460.2 | Solyc09g008460.3 |
| Solyc09g008505.1 | Solyc09g008505.2 |
| Solyc09g008600.2 | Solyc09g008600.3 |
| Solyc09g008630.2 | Solyc09g008630.3 |
| Solyc09g008650.2 | Solyc09g008650.3 |
| Solyc09g008660.2 | Solyc09g008660.3 |
| Solyc09g008670.2 | Solyc09g008670.3 |
| Solyc09g008700.1 | Solyc09g008700.2 |
| Solyc09g008730.2 | Solyc09g008730.3 |
| Solyc09g008770.2 | Solyc09g008770.3 |
| Solyc09g008800.2 | Solyc09g008800.3 |
| Solyc09g008810.2 | Solyc09g008810.3 |
| Solyc09g008820.2 | Solyc09g008820.3 |
| Solyc09g008830.2 | Solyc09g008830.3 |
| Solyc09g008850.2 | Solyc09g008850.3 |
| Solyc09g098620.1 | Solyc09g098620.2 |
| Solyc09g008910.1 | Solyc09g008910.2 |
| Solyc09g008920.2 | Solyc09g008920.3 |
| Solyc09g008960.1 | Solyc09g008960.2 |
| Solyc09g009020.2 | Solyc09g009020.3 |
| Solyc09g009040.2 | Solyc09g009040.3 |
| Solyc09g009090.2 | Solyc09g009090.3 |
| Solyc09g009140.2 | Solyc09g009140.3 |
| Solyc09g009175.1 | Solyc09g150102.1 |
| Solyc09g009180.2 | Solyc09g009180.3 |
| Solyc09g009230.1 | Solyc09g009230.2 |
| Solyc09g009250.2 | Solyc09g009250.3 |
| Solyc09g009260.2 | Solyc09g009260.3 |
| Solyc09g009270.1 | Solyc09g009270.2 |
| Solyc09g009290.2 | Solyc09g009290.3 |
| Solyc09g009380.2 | Solyc09g009380.3 |
| Solyc09g009390.2 | Solyc09g009390.3 |
| Solyc09g009410.2 | Solyc09g009410.3 |
| Solyc09g009430.2 | Solyc09g009430.3 |
| Solyc09g009500.2 | Solyc09g009500.3 |
| Solyc09g009520.2 | Solyc09g009520.3 |
| Solyc09g009560.1 | Solyc09g009560.2 |
| Solyc09g009630.2 | Solyc09g009630.3 |
| Solyc09g009640.2 | Solyc09g009640.3 |
| Solyc09g009650.2 | Solyc09g009650.3 |
| Solyc09g009660.2 | Solyc09g009660.3 |

| | |
|---|---|
| Solyc09g009670.2 | Solyc09g009670.3 |
| Solyc09g009690.2 | Solyc09g009690.3 |
| Solyc09g009720.1 | Solyc09g009720.2 |
| Solyc09g009770.2 | Solyc09g009770.3 |
| Solyc09g009800.2 | Solyc09g009800.3 |
| Solyc09g009880.2 | Solyc09g009880.3 |
| Solyc09g009930.1 | Solyc09g009930.2 |
| Solyc09g009940.2 | Solyc09g009940.3 |
| Solyc09g009960.2 | Solyc09g009960.3 |
| Solyc09g009970.2 | Solyc09g009970.3 |
| Solyc09g010060.2 | Solyc09g010060.3 |
| Solyc09g010090.4 | Solyc09g010090.5.1 |
| Solyc09g010100.2 | Solyc09g010100.3 |
| Solyc09g010110.2 | Solyc09g010110.3 |
| Solyc09g010170.2 | Solyc09g010170.3 |
| Solyc09g010180.2 | Solyc09g010180.3 |
| Solyc09g010190.2 | Solyc09g010190.3 |
| Solyc09g010210.2 | Solyc09g010210.3 |
| Solyc09g010230.1 | Solyc09g010230.2 |
| Solyc09g010280.2 | Solyc09g010280.3 |
| Solyc09g010300.2 | Solyc09g010300.3 |
| Solyc09g010320.2 | Solyc09g010320.3 |
| Solyc09g010360.2 | Solyc09g010360.3 |
| Solyc09g010370.2 | Solyc09g010370.3 |
| Solyc09g010380.2 | Solyc09g010380.3 |
| Solyc09g010400.2 | Solyc09g010400.3 |
| Solyc09g010440.2 | Solyc09g010440.3 |
| Solyc09g010460.2 | Solyc09g010460.3 |
| Solyc09g010470.2 | Solyc09g010470.3 |
| Solyc09g010500.2 | Solyc09g010500.3 |
| Solyc09g010520.2 | Solyc09g010520.3 |
| Solyc09g010530.2 | Solyc09g010530.3 |
| Solyc09g010630.2 | Solyc09g010630.3 |
| Solyc09g010660.2 | Solyc09g010660.3 |
| Solyc09g010680.2 | Solyc09g010680.3 |
| Solyc09g010720.2 | Solyc09g010720.3 |
| Solyc09g010760.1 | Solyc09g010760.2 |
| Solyc09g010770.2 | Solyc09g010770.3 |
| Solyc09g010780.2 | Solyc09g010780.3 |
| Solyc09g010790.1 | Solyc09g010790.2 |
| Solyc09g010800.4 | Solyc09g010800.5.1 |
| Solyc09g010810.2 | Solyc09g010810.3 |
| Solyc09g010820.2 | Solyc09g010820.3 |

| | |
|---|---|
| Solyc09g010880.2 | Solyc09g010880.3 |
| Solyc09g010913.1 | Solyc09g150103.1 |
| Solyc09g010917.1 | Solyc09g150104.1 |
| Solyc09g010950.2 | Solyc09g010950.3 |
| Solyc09g010960.2 | Solyc09g010960.3 |
| Solyc09g010990.2 | Solyc09g010990.3 |
| Solyc09g011010.2 | Solyc09g011010.3 |
| Solyc09g011030.2 | Solyc09g011030.3 |
| Solyc09g011045.1 | Solyc09g150105.1 |
| Solyc09g011080.2 | Solyc09g011080.3 |
| Solyc09g011170.2 | Solyc09g011170.3 |
| Solyc09g011200.1 | Solyc09g011200.2 |
| Solyc09g011220.2 | Solyc09g011220.3 |
| Solyc09g011240.2 | Solyc09g011240.3 |
| Solyc09g011250.2 | Solyc09g011250.3 |
| Solyc09g011260.1 | Solyc09g011260.2 |
| Solyc09g011300.2 | Solyc09g011300.3 |
| Solyc09g011310.2 | Solyc09g011310.3 |
| Solyc09g011320.2 | Solyc09g011320.3 |
| Solyc09g011340.2 | Solyc09g011340.3 |
| Solyc09g011380.2 | Solyc09g011380.3 |
| Solyc09g011420.2 | Solyc09g011420.3 |
| Solyc09g011470.2 | Solyc09g011470.3 |
| Solyc09g011480.2 | Solyc09g011480.3 |
| Solyc09g011490.2 | Solyc09g011490.3 |
| Solyc09g011500.2 | Solyc09g011500.3 |
| Solyc09g011520.2 | Solyc09g011520.3 |
| Solyc09g011550.2 | Solyc09g011550.3 |
| Solyc09g011560.2 | Solyc09g011560.3 |
| Solyc09g011600.2 | Solyc09g011600.3 |
| Solyc09g011610.2 | Solyc09g011610.3 |
| Solyc09g011620.1 | Solyc09g011620.2 |
| Solyc09g011640.4 | Solyc09g011640.5.1 |
| Solyc09g011660.2 | Solyc09g011660.3 |
| Solyc09g011720.2 | Solyc09g011720.3 |
| Solyc09g011750.2 | Solyc09g011750.3 |
| Solyc09g011760.2 | Solyc09g011760.3 |
| Solyc09g011770.2 | Solyc09g011770.3 |
| Solyc09g011780.2 | Solyc09g011780.3 |
| Solyc09g011790.2 | Solyc09g011790.3 |
| Solyc09g011800.2 | Solyc09g011800.3 |
| Solyc09g011810.2 | Solyc09g011810.3 |
| Solyc09g011880.2 | Solyc09g011880.3 |

| | |
|---|---|
| Solyc09g011910.2 | Solyc09g011910.3 |
| Solyc09g011920.2 | Solyc09g011920.3 |
| Solyc09g011930.2 | Solyc09g011930.3 |
| Solyc09g011935.1 | Solyc09g150106.1 |
| Solyc09g011960.1 | Solyc09g011960.2 |
| Solyc09g013070.2 | Solyc09g013070.3 |
| Solyc09g013080.2 | Solyc09g013080.3 |
| Solyc09g013100.2 | Solyc09g013100.3 |
| Solyc09g013110.2 | Solyc09g013110.3 |
| Solyc09g013130.2 | Solyc09g013130.3 |
| Solyc09g013160.2 | Solyc09g013160.3 |
| Solyc09g014170.2 | Solyc09g014170.3 |
| Solyc09g014220.2 | Solyc09g014220.3 |
| Solyc09g014230.1 | Solyc09g014230.2 |
| Solyc09g014250.2 | Solyc09g014250.3 |
| Solyc09g014300.2 | Solyc09g014300.3 |
| Solyc09g014310.2 | Solyc09g014310.3 |
| Solyc09g014350.2 | Solyc09g014350.3 |
| Solyc09g014380.2 | Solyc09g014380.3 |
| Solyc09g014400.2 | Solyc09g014400.3 |
| Solyc09g014410.2 | Solyc09g014410.3 |
| Solyc09g014470.1 | Solyc09g014470.2 |
| Solyc09g014500.2 | Solyc09g014500.3 |
| Solyc09g014510.2 | Solyc09g014510.3 |
| Solyc09g014520.2 | Solyc09g014520.3 |
| Solyc09g014550.2 | Solyc09g014550.3 |
| Solyc09g014560.1 | Solyc09g014560.2 |
| Solyc09g014580.2 | Solyc09g014580.3 |
| Solyc09g014590.2 | Solyc09g014590.3 |
| Solyc09g014620.2 | Solyc09g014620.3 |
| Solyc09g014623.1 | Solyc09g150107.1 |
| Solyc09g014627.1 | Solyc09g150108.1 |
| Solyc09g014640.1 | Solyc09g014640.2 |
| Solyc09g014730.2 | Solyc09g014730.3 |
| Solyc09g014740.2 | Solyc09g014740.3 |
| Solyc09g014770.2 | Solyc09g014770.3 |
| Solyc09g014780.2 | Solyc09g014780.3 |
| Solyc09g014820.2 | Solyc09g014820.3 |
| Solyc09g014860.2 | Solyc09g014860.3 |
| Solyc09g014890.2 | Solyc09g014890.3 |
| Solyc09g014900.2 | Solyc09g014900.3 |
| Solyc09g014950.1 | Solyc09g014950.2 |
| Solyc09g014995.1 | Solyc09g150109.1 |

| | |
|---|---|
| Solyc09g015070.2 | Solyc09g015070.3 |
| Solyc09g015080.2 | Solyc09g015080.3 |
| Solyc09g015110.1 | Solyc09g015110.2 |
| Solyc09g015155.1 | Solyc09g150110.1 |
| Solyc09g015190.1 | Solyc09g015190.2 |
| Solyc09g015330.1 | Solyc09g015330.2 |
| Solyc09g015385.1 | Solyc09g150111.1 |
| Solyc09g015430.2 | Solyc09g015430.3 |
| Solyc09g015475.1 | Solyc09g150112.1 |
| Solyc09g015490.2 | Solyc09g015490.3 |
| Solyc09g015505.1 | Solyc09g150113.1 |
| Solyc09g015520.2 | Solyc09g015520.3 |
| Solyc09g015630.1 | Solyc09g015630.2 |
| Solyc09g015670.2 | Solyc09g015670.3 |
| Solyc09g015770.2 | Solyc09g015770.3 |
| Solyc09g015780.1 | Solyc09g015780.2 |
| Solyc09g015820.2 | Solyc09g015820.3 |
| Solyc09g015930.2 | Solyc09g015930.3 |
| Solyc09g016950.2 | Solyc09g016950.3 |
| Solyc09g017980.1 | Solyc09g017980.2 |
| Solyc09g018010.2 | Solyc09g018010.3 |
| Solyc09g018020.2 | Solyc09g018020.3 |
| Solyc09g018080.2 | Solyc09g018080.3 |
| Solyc09g018160.2 | Solyc09g018160.3 |
| Solyc09g018260.1 | Solyc09g018260.2 |
| Solyc09g018440.1 | Solyc09g018440.2 |
| Solyc09g018450.2 | Solyc09g018450.3 |
| Solyc09g018510.2 | Solyc09g018510.3 |
| Solyc09g018630.2 | Solyc09g018630.3 |
| Solyc09g018650.2 | Solyc09g018650.3 |
| Solyc09g018720.2 | Solyc09g018720.3 |
| Solyc09g018750.2 | Solyc09g018750.3 |
| Solyc09g018790.2 | Solyc09g018790.3 |
| Solyc09g018800.1 | Solyc09g018800.2 |
| Solyc09g018803.1 | Solyc09g150114.1 |
| Solyc09g018807.1 | Solyc09g150115.1 |
| Solyc09g018850.2 | Solyc09g018850.3 |
| Solyc09g019970.2 | Solyc09g019970.3 |
| Solyc09g019980.2 | Solyc09g019980.3 |
| Solyc09g020000.2 | Solyc09g020000.3 |
| Solyc09g020010.1 | Solyc09g020010.2 |
| Solyc09g020110.2 | Solyc09g020110.3 |
| Solyc09g020130.2 | Solyc09g020130.3 |

| | |
|---|---|
| Solyc09g020140.2 | Solyc09g020140.3 |
| Solyc09g020190.2 | Solyc09g020190.3 |
| Solyc09g037030.1 | Solyc09g037030.2 |
| Solyc09g037050.1 | Solyc09g037050.2 |
| Solyc09g037070.1 | Solyc09g037070.2 |
| Solyc09g037130.1 | Solyc09g037130.2 |
| Solyc09g042230.1 | Solyc09g042230.2 |
| Solyc09g042250.1 | Solyc09g042250.2 |
| Solyc09g042260.2 | Solyc09g042260.3 |
| Solyc09g042263.1 | Solyc09g150120.1 |
| Solyc09g042267.1 | Solyc09g150121.1 |
| Solyc09g042300.1 | Solyc09g042300.2 |
| Solyc09g042305.1 | Solyc09g150122.1 |
| Solyc09g042330.1 | Solyc09g042330.2 |
| Solyc09g042430.1 | Solyc09g042430.2 |
| Solyc09g042650.1 | Solyc09g042650.2 |
| Solyc09g042710.2 | Solyc09g042710.3 |
| Solyc09g042750.2 | Solyc09g042750.3 |
| Solyc09g042755.1 | Solyc09g150123.1 |
| Solyc09g042760.1 | Solyc09g042760.2 |
| Solyc09g031970.2 | Solyc09g031970.3 |
| Solyc09g031930.2 | Solyc09g031930.3 |
| Solyc09g031910.1 | Solyc09g031910.2 |
| Solyc09g031875.1 | Solyc09g150119.1 |
| Solyc09g031810.1 | Solyc09g031810.2 |
| Solyc09g031780.2 | Solyc09g031780.3 |
| Solyc09g031770.2 | Solyc09g031770.3 |
| Solyc09g031705.1 | Solyc09g150118.1 |
| Solyc09g031650.2 | Solyc09g031650.3 |
| Solyc09g031580.2 | Solyc09g031580.3 |
| Solyc09g031547.1 | Solyc09g150117.1 |
| Solyc09g030450.2 | Solyc09g030450.3 |
| Solyc09g030390.2 | Solyc09g030390.3 |
| Solyc09g030385.1 | Solyc09g150116.1 |
| Solyc09g030370.2 | Solyc09g030370.3 |
| Solyc09g047840.2 | Solyc09g047840.3 |
| Solyc09g047900.1 | Solyc09g047900.2 |
| Solyc09g047920.1 | Solyc09g047920.2 |
| Solyc09g050050.1 | Solyc09g050050.2 |
| Solyc09g055150.1 | Solyc09g055150.2 |
| Solyc09g055230.2 | Solyc09g055230.3 |
| Solyc09g055250.2 | Solyc09g055250.3 |
| Solyc09g055310.2 | Solyc09g055310.3 |

| | |
|---|---|
| Solyc09g055350.2 | Solyc09g055350.3 |
| Solyc09g055425.1 | Solyc09g055425.2 |
| Solyc09g055450.1 | Solyc09g055450.2 |
| Solyc09g055540.1 | Solyc09g055540.2 |
| Solyc09g055570.2 | Solyc09g055570.3 |
| Solyc09g055590.2 | Solyc09g055590.3 |
| Solyc09g055710.2 | Solyc09g055710.3 |
| Solyc09g055715.1 | Solyc09g150125.1 |
| Solyc09g055850.2 | Solyc09g055850.3 |
| Solyc09g055910.2 | Solyc09g055910.3 |
| Solyc09g055930.2 | Solyc09g055930.3 |
| Solyc09g055940.2 | Solyc09g055940.3 |
| Solyc09g055950.1 | Solyc09g055950.2 |
| Solyc09g055990.1 | Solyc09g055990.2 |
| Solyc09g056040.3 | Solyc09g056040.4 |
| Solyc09g056100.1 | Solyc09g056100.2 |
| Solyc09g056170.2 | Solyc09g056170.3 |
| Solyc09g056180.2 | Solyc09g056180.3 |
| Solyc09g056200.2 | Solyc09g056200.3 |
| Solyc09g056280.1 | Solyc09g056280.2 |
| Solyc09g056300.1 | Solyc09g056300.2 |
| Solyc09g056340.2 | Solyc09g056340.3 |
| Solyc09g056360.2 | Solyc09g056360.3 |
| Solyc09g056435.1 | Solyc09g150126.1 |
| Solyc09g057530.1 | Solyc09g057530.2 |
| Solyc09g057555.1 | Solyc09g150127.1 |
| Solyc09g057630.2 | Solyc09g057630.3 |
| Solyc09g057650.2 | Solyc09g057650.3 |
| Solyc09g057660.2 | Solyc09g057660.3 |
| Solyc09g057670.2 | Solyc09g057670.3 |
| Solyc09g057680.2 | Solyc09g057680.3 |
| Solyc09g057880.2 | Solyc09g057880.3 |
| Solyc09g057887.1 | Solyc09g150128.1 |
| Solyc09g057903.1 | Solyc09g150129.1 |
| Solyc09g057907.1 | Solyc09g150130.1 |
| Solyc09g057910.2 | Solyc09g057910.3 |
| Solyc09g057920.2 | Solyc09g057920.3 |
| Solyc09g057940.1 | Solyc09g057940.2 |
| Solyc09g057945.1 | Solyc09g150131.1 |
| Solyc09g057950.2 | Solyc09g057950.3 |
| Solyc09g058980.2 | Solyc09g058980.3 |
| Solyc09g059040.2 | Solyc09g059040.3 |
| Solyc09g059100.1 | Solyc09g059100.2 |

| | |
|---|---|
| Solyc09g059240.2 | Solyc09g059240.3 |
| Solyc09g059270.2 | Solyc09g059270.3 |
| Solyc09g059310.2 | Solyc09g059310.3 |
| Solyc09g059340.1 | Solyc09g059340.2 |
| Solyc09g059345.1 | Solyc09g150132.1 |
| Solyc09g059430.2 | Solyc09g059430.3 |
| Solyc09g059473.1 | Solyc09g150133.1 |
| Solyc09g059477.1 | Solyc09g150134.1 |
| Solyc09g059500.2 | Solyc09g059500.3 |
| Solyc09g059510.2 | Solyc09g059510.3 |
| Solyc09g059520.2 | Solyc09g059520.3 |
| Solyc09g063157.1 | Solyc09g063157.2 |
| Solyc09g059570.2 | Solyc09g059570.3 |
| Solyc09g059610.2 | Solyc09g059610.3 |
| Solyc09g059675.1 | Solyc09g150135.1 |
| Solyc09g059710.1 | Solyc09g059710.2 |
| Solyc09g059805.1 | Solyc09g150136.1 |
| Solyc09g059820.1 | Solyc09g059820.2 |
| Solyc09g059960.2 | Solyc09g059960.3 |
| Solyc09g059970.2 | Solyc09g059970.3 |
| Solyc09g060070.1 | Solyc09g060070.2 |
| Solyc09g060090.2 | Solyc09g060090.3 |
| Solyc09g060140.2 | Solyc09g060140.3 |
| Solyc09g060160.1 | Solyc09g060160.2 |
| Solyc09g061230.2 | Solyc09g061230.3 |
| Solyc09g061260.1 | Solyc09g061260.2 |
| Solyc09g061290.2 | Solyc09g061290.3 |
| Solyc09g061310.2 | Solyc09g061310.3 |
| Solyc09g061380.2 | Solyc09g061380.3 |
| Solyc09g061395.1 | Solyc09g150137.1 |
| Solyc09g061430.2 | Solyc09g061430.3 |
| Solyc09g061530.1 | Solyc09g061530.2 |
| Solyc09g061550.1 | Solyc09g061550.2 |
| Solyc09g061620.2 | Solyc09g061620.3 |
| Solyc09g061630.1 | Solyc09g061630.2 |
| Solyc09g061680.2 | Solyc09g061680.3 |
| Solyc09g061685.1 | Solyc09g150138.1 |
| Solyc09g061700.2 | Solyc09g061700.3 |
| Solyc09g061710.2 | Solyc09g061710.3 |
| Solyc09g061793.1 | Solyc09g150139.1 |
| Solyc09g061880.1 | Solyc09g061880.2 |
| Solyc09g061890.2 | Solyc09g061890.3 |
| Solyc09g061910.1 | Solyc09g061910.2 |

| | |
|---|---|
| Solyc09g061960.2 | Solyc09g061960.3 |
| Solyc09g062990.1 | Solyc09g062990.2 |
| Solyc09g063050.1 | Solyc09g063050.2 |
| Solyc09g063060.2 | Solyc09g063060.3 |
| Solyc09g063070.2 | Solyc09g063070.3 |
| Solyc09g063130.2 | Solyc09g063130.3 |
| Solyc09g063150.2 | Solyc09g063150.3 |
| Solyc09g064160.2 | Solyc09g064160.3 |
| Solyc09g064240.2 | Solyc09g064240.3 |
| Solyc09g064270.2 | Solyc09g064270.3 |
| Solyc09g064277.1 | Solyc09g150140.1 |
| Solyc09g064310.2 | Solyc09g064310.3 |
| Solyc09g064320.2 | Solyc09g064320.3 |
| Solyc09g064330.1 | Solyc09g064330.2 |
| Solyc09g064365.1 | Solyc09g150141.1 |
| Solyc09g064380.2 | Solyc09g064380.3 |
| Solyc09g064400.1 | Solyc09g064400.2 |
| Solyc09g064450.2 | Solyc09g064450.3 |
| Solyc09g064470.2 | Solyc09g064470.3 |
| Solyc09g064500.2 | Solyc09g064500.3 |
| Solyc09g064510.2 | Solyc09g064510.3 |
| Solyc09g064520.2 | Solyc09g064520.3 |
| Solyc09g064530.2 | Solyc09g064530.3 |
| Solyc09g064540.2 | Solyc09g064540.3 |
| Solyc09g064555.1 | Solyc09g150142.1 |
| Solyc09g064600.2 | Solyc09g064600.3 |
| Solyc09g064610.2 | Solyc09g064610.3 |
| Solyc09g064630.2 | Solyc09g064630.3 |
| Solyc09g064720.2 | Solyc09g064720.3 |
| Solyc09g064730.2 | Solyc09g064730.3 |
| Solyc09g064780.2 | Solyc09g064780.3 |
| Solyc09g064790.2 | Solyc09g064790.3 |
| Solyc09g064850.2 | Solyc09g064850.3 |
| Solyc09g064870.2 | Solyc09g064870.3 |
| Solyc09g064920.2 | Solyc09g064920.3 |
| Solyc09g064930.2 | Solyc09g064930.3 |
| Solyc09g064960.1 | Solyc09g064960.2 |
| Solyc09g065030.2 | Solyc09g065030.3 |
| Solyc09g065050.1 | Solyc09g065050.2 |
| Solyc09g065070.1 | Solyc09g065070.2 |
| Solyc09g065080.1 | Solyc09g065080.2 |
| Solyc09g065090.1 | Solyc09g065090.2 |
| Solyc09g065170.1 | Solyc09g065170.2 |

| | |
|---|---|
| Solyc09g065180.2 | Solyc09g065180.3 |
| Solyc09g065240.2 | Solyc09g065240.3 |
| Solyc09g065250.1 | Solyc09g065250.2 |
| Solyc09g065260.1 | Solyc09g065260.2 |
| Solyc09g065270.2 | Solyc09g065270.3 |
| Solyc09g065330.2 | Solyc09g065330.3 |
| Solyc09g065340.2 | Solyc09g065340.3 |
| Solyc09g065383.1 | Solyc09g150143.1 |
| Solyc09g065400.1 | Solyc09g065400.2 |
| Solyc09g065405.1 | Solyc09g150144.1 |
| Solyc09g065420.2 | Solyc09g065420.3 |
| Solyc09g065445.1 | Solyc09g150145.1 |
| Solyc09g065453.1 | Solyc09g150146.1 |
| Solyc09g065457.1 | Solyc09g150147.1 |
| Solyc09g065470.2 | Solyc09g065470.3 |
| Solyc09g065480.1 | Solyc09g065480.2 |
| Solyc09g065490.1 | Solyc09g065490.2 |
| Solyc09g065530.1 | Solyc09g065530.2 |
| Solyc09g065540.2 | Solyc09g065540.3 |
| Solyc09g065560.2 | Solyc09g065560.3 |
| Solyc09g065590.2 | Solyc09g065590.3 |
| Solyc09g065600.2 | Solyc09g065600.3 |
| Solyc09g065610.1 | Solyc09g065610.2 |
| Solyc09g065620.2 | Solyc09g065620.3 |
| Solyc09g065630.2 | Solyc09g065630.3 |
| Solyc09g065640.2 | Solyc09g065640.3 |
| Solyc09g065650.2 | Solyc09g065650.3 |
| Solyc09g065655.1 | Solyc09g150148.1 |
| Solyc09g065670.2 | Solyc09g065670.3 |
| Solyc09g065700.2 | Solyc09g065700.3 |
| Solyc09g065730.2 | Solyc09g065730.3 |
| Solyc09g065740.1 | Solyc09g065740.2 |
| Solyc09g065750.2 | Solyc09g065750.3 |
| Solyc09g065780.2 | Solyc09g065780.3 |
| Solyc09g065790.1 | Solyc09g065790.2 |
| Solyc09g065810.2 | Solyc09g065810.3 |
| Solyc09g065840.2 | Solyc09g065840.3 |
| Solyc09g065860.2 | Solyc09g065860.3 |
| Solyc09g065900.2 | Solyc09g065900.3 |
| Solyc09g065920.2 | Solyc09g065920.3 |
| Solyc09g065950.2 | Solyc09g065950.3 |
| Solyc09g065955.1 | Solyc09g150149.1 |
| Solyc09g065990.2 | Solyc09g065990.3 |

| | |
|---|---|
| Solyc09g066010.2 | Solyc09g066010.3 |
| Solyc09g066030.1 | Solyc09g066030.2 |
| Solyc09g066100.2 | Solyc09g066100.3 |
| Solyc09g066110.2 | Solyc09g066110.3 |
| Solyc09g066140.2 | Solyc09g066140.3 |
| Solyc09g066150.1 | Solyc09g066150.2 |
| Solyc09g066155.1 | Solyc09g150150.1 |
| Solyc09g066180.1 | Solyc09g066180.2 |
| Solyc09g066270.2 | Solyc09g066270.3 |
| Solyc09g066290.2 | Solyc09g066290.3 |
| Solyc09g066310.2 | Solyc09g066310.3 |
| Solyc09g066320.1 | Solyc09g066320.2 |
| Solyc09g066330.1 | Solyc09g066330.2 |
| Solyc09g066400.1 | Solyc09g066400.2 |
| Solyc09g066420.2 | Solyc09g066420.3 |
| Solyc09g066430.2 | Solyc09g066430.3 |
| Solyc09g025270.2 | Solyc09g025270.3 |
| Solyc09g025240.2 | Solyc09g025240.3 |
| Solyc09g025220.1 | Solyc09g025220.2 |
| Solyc09g025210.2 | Solyc09g025210.3 |
| Solyc09g025200.1 | Solyc09g025200.2 |
| Solyc09g072530.1 | Solyc09g072530.2 |
| Solyc09g072540.1 | Solyc09g072540.2 |
| Solyc09g072570.2 | Solyc09g072570.3 |
| Solyc09g072590.2 | Solyc09g072590.3 |
| Solyc09g072600.1 | Solyc09g072600.2 |
| Solyc09g072610.2 | Solyc09g072610.3 |
| Solyc09g072630.2 | Solyc09g072630.3 |
| Solyc09g072640.2 | Solyc09g072640.3 |
| Solyc09g072670.2 | Solyc09g072670.3 |
| Solyc09g072680.1 | Solyc09g072680.2 |
| Solyc09g072725.1 | Solyc09g150151.1 |
| Solyc09g072730.1 | Solyc09g072730.2 |
| Solyc09g072790.2 | Solyc09g072790.3 |
| Solyc09g072800.2 | Solyc09g072800.3 |
| Solyc09g072810.2 | Solyc09g072810.3 |
| Solyc09g072860.1 | Solyc09g072860.2 |
| Solyc09g072880.2 | Solyc09g072880.3 |
| Solyc09g072885.1 | Solyc09g150152.1 |
| Solyc09g072890.1 | Solyc09g072890.2 |
| Solyc09g072970.2 | Solyc09g072970.3 |
| Solyc09g073000.2 | Solyc09g073000.3 |
| Solyc09g073040.2 | Solyc09g073040.3 |

| | |
|---|---|
| Solyc09g074050.2 | Solyc09g074050.3 |
| Solyc09g074060.2 | Solyc09g074060.3 |
| Solyc09g074090.2 | Solyc09g074090.3 |
| Solyc09g074180.2 | Solyc09g074180.3 |
| Solyc09g074185.1 | Solyc09g150153.1 |
| Solyc09g074190.1 | Solyc09g074190.2 |
| Solyc09g074210.2 | Solyc09g074210.3 |
| Solyc09g074250.2 | Solyc09g074250.3 |
| Solyc09g074270.2 | Solyc09g074270.3 |
| Solyc09g074310.2 | Solyc09g074310.3 |
| Solyc09g074320.2 | Solyc09g074320.3 |
| Solyc09g074330.2 | Solyc09g074330.3 |
| Solyc09g074340.2 | Solyc09g074340.3 |
| Solyc09g074400.2 | Solyc09g074400.3 |
| Solyc09g074440.2 | Solyc09g074440.3 |
| Solyc09g074470.2 | Solyc09g074470.3 |
| Solyc09g074510.2 | Solyc09g074510.3 |
| Solyc09g074520.2 | Solyc09g074520.3 |
| Solyc09g074530.2 | Solyc09g074530.3 |
| Solyc09g074550.2 | Solyc09g074550.3 |
| Solyc09g074560.2 | Solyc09g074560.3 |
| Solyc09g074600.1 | Solyc09g074600.2 |
| Solyc09g074610.2 | Solyc09g074610.3 |
| Solyc09g074630.2 | Solyc09g074630.3 |
| Solyc09g074640.2 | Solyc09g074640.3 |
| Solyc09g074650.2 | Solyc09g074650.3 |
| Solyc09g074660.2 | Solyc09g074660.3 |
| Solyc09g074680.2 | Solyc09g074680.3 |
| Solyc09g074690.2 | Solyc09g074690.3 |
| Solyc09g074700.1 | Solyc09g074700.2 |
| Solyc09g074750.2 | Solyc09g074750.3 |
| Solyc09g074765.1 | Solyc09g150154.1 |
| Solyc09g074780.2 | Solyc09g074780.3 |
| Solyc09g074830.2 | Solyc09g074830.3 |
| Solyc09g074855.1 | Solyc09g150155.1 |
| Solyc09g074920.2 | Solyc09g074920.3 |
| Solyc09g074930.2 | Solyc09g074930.3 |
| Solyc09g074950.2 | Solyc09g074950.3 |
| Solyc09g074970.2 | Solyc09g074970.3 |
| Solyc09g075000.2 | Solyc09g075000.3 |
| Solyc09g075010.2 | Solyc09g075010.3 |
| Solyc09g075020.2 | Solyc09g075020.3 |
| Solyc09g075030.2 | Solyc09g075030.3 |

| | |
|---|---|
| Solyc09g075040.2 | Solyc09g075040.3 |
| Solyc09g075060.2 | Solyc09g075060.3 |
| Solyc09g075070.2 | Solyc09g075070.3 |
| Solyc09g075080.2 | Solyc09g075080.3 |
| Solyc09g075100.2 | Solyc09g075100.3 |
| Solyc09g075120.2 | Solyc09g075120.3 |
| Solyc09g075130.2 | Solyc09g075130.3 |
| Solyc09g075140.2 | Solyc09g075140.3 |
| Solyc09g075150.2 | Solyc09g075150.3 |
| Solyc09g075180.2 | Solyc09g075180.3 |
| Solyc09g075190.2 | Solyc09g075190.3 |
| Solyc09g075210.2 | Solyc09g075210.3 |
| Solyc09g075280.1 | Solyc09g075280.2 |
| Solyc09g075290.2 | Solyc09g075290.3 |
| Solyc09g075300.2 | Solyc09g075300.3 |
| Solyc09g075370.2 | Solyc09g075370.3 |
| Solyc09g075420.2 | Solyc09g075420.3 |
| Solyc09g075430.2 | Solyc09g075430.3 |
| Solyc09g075450.2 | Solyc09g075450.3 |
| Solyc09g075460.2 | Solyc09g075460.3 |
| Solyc09g075480.2 | Solyc09g075480.3 |
| Solyc09g075500.2 | Solyc09g075500.3 |
| Solyc09g075530.2 | Solyc09g075530.3 |
| Solyc09g075550.2 | Solyc09g075550.3 |
| Solyc09g075610.2 | Solyc09g075610.3 |
| Solyc09g075650.2 | Solyc09g075650.3 |
| Solyc09g075730.2 | Solyc09g075730.3 |
| Solyc09g075780.1 | Solyc09g075780.2 |
| Solyc09g075790.2 | Solyc09g075790.3 |
| Solyc09g075810.2 | Solyc09g075810.3 |
| Solyc09g075820.2 | Solyc09g075820.3 |
| Solyc09g075830.2 | Solyc09g075830.3 |
| Solyc09g075850.2 | Solyc09g075850.3 |
| Solyc09g075860.2 | Solyc09g075860.3 |
| Solyc09g075870.1 | Solyc09g075870.2 |
| Solyc09g075880.2 | Solyc09g075880.3 |
| Solyc09g075890.2 | Solyc09g075890.3 |
| Solyc09g075900.2 | Solyc09g075900.3 |
| Solyc09g075930.1 | Solyc09g075930.2 |
| Solyc09g075940.2 | Solyc09g075940.3 |
| Solyc09g075970.2 | Solyc09g075970.3 |
| Solyc09g075980.2 | Solyc09g075980.3 |
| Solyc09g075990.2 | Solyc09g075990.3 |

| | |
|---|---|
| Solyc09g076000.2 | Solyc09g076000.3 |
| Solyc09g076010.2 | Solyc09g076010.3 |
| Solyc09g076020.2 | Solyc09g076020.3 |
| Solyc09g076040.2 | Solyc09g076040.3 |
| Solyc09g082060.2 | Solyc09g082060.3 |
| Solyc09g082100.2 | Solyc09g082100.3 |
| Solyc09g082250.2 | Solyc09g082250.3 |
| Solyc09g082280.2 | Solyc09g082280.3 |
| Solyc09g082290.2 | Solyc09g082290.3 |
| Solyc09g082300.2 | Solyc09g082300.3 |
| Solyc09g082310.1 | Solyc09g082310.2 |
| Solyc09g082320.2 | Solyc09g082320.3 |
| Solyc09g082330.1 | Solyc09g082330.2 |
| Solyc09g082395.1 | Solyc09g150156.1 |
| Solyc09g082400.1 | Solyc09g082400.2 |
| Solyc09g082460.2 | Solyc09g082460.3 |
| Solyc09g082480.1 | Solyc09g082480.2 |
| Solyc09g082510.2 | Solyc09g082510.3 |
| Solyc09g082520.2 | Solyc09g082520.3 |
| Solyc09g082540.2 | Solyc09g082540.3 |
| Solyc09g082550.2 | Solyc09g082550.3 |
| Solyc09g082570.2 | Solyc09g082570.3 |
| Solyc09g082620.2 | Solyc09g082620.3 |
| Solyc09g082630.2 | Solyc09g082630.3 |
| Solyc09g082660.2 | Solyc09g082660.3 |
| Solyc09g082670.2 | Solyc09g082670.3 |
| Solyc09g082680.2 | Solyc09g082680.3 |
| Solyc09g082690.2 | Solyc09g082690.3 |
| Solyc09g082710.2 | Solyc09g082710.3 |
| Solyc09g082720.2 | Solyc09g082720.3 |
| Solyc09g082760.2 | Solyc09g082760.3 |
| Solyc09g082770.2 | Solyc09g082770.3 |
| Solyc09g082780.2 | Solyc09g082780.3 |
| Solyc09g082800.2 | Solyc09g082800.3 |
| Solyc09g082820.2 | Solyc09g082820.3 |
| Solyc09g082845.1 | Solyc09g082845.2 |
| Solyc09g082860.2 | Solyc09g082860.3 |
| Solyc09g082890.1 | Solyc09g082890.2 |
| Solyc09g082900.2 | Solyc09g082900.3 |
| Solyc09g098610.1 | Solyc09g098610.2 |
| Solyc09g082910.1 | Solyc09g082910.2 |
| Solyc09g082940.2 | Solyc09g082940.3 |
| Solyc09g082950.1 | Solyc09g082950.2 |

| | |
|---|---|
| Solyc09g082970.2 | Solyc09g082970.3 |
| Solyc09g082980.2 | Solyc09g082980.3 |
| Solyc09g082990.2 | Solyc09g082990.3 |
| Solyc09g083000.2 | Solyc09g083000.3 |
| Solyc09g083050.2 | Solyc09g083050.3 |
| Solyc09g083105.1 | Solyc09g150157.1 |
| Solyc09g083120.2 | Solyc09g083120.3 |
| Solyc09g083130.2 | Solyc09g083130.3 |
| Solyc09g083160.2 | Solyc09g083160.3 |
| Solyc09g083170.2 | Solyc09g083170.3 |
| Solyc09g083180.2 | Solyc09g083180.3 |
| Solyc09g083190.2 | Solyc09g083190.3 |
| Solyc09g083210.2 | Solyc09g083210.3 |
| Solyc09g083220.2 | Solyc09g083220.3 |
| Solyc09g083280.2 | Solyc09g083280.3 |
| Solyc09g083290.2 | Solyc09g083290.3 |
| Solyc09g083330.2 | Solyc09g083330.3 |
| Solyc09g083350.2 | Solyc09g083350.3 |
| Solyc09g083360.2 | Solyc09g083360.3 |
| Solyc09g083370.2 | Solyc09g083370.3 |
| Solyc09g083400.2 | Solyc09g083400.3 |
| Solyc09g083420.2 | Solyc09g083420.3 |
| Solyc09g084450.2 | Solyc09g084450.3 |
| Solyc09g084460.2 | Solyc09g084460.3 |
| Solyc09g084470.2 | Solyc09g084470.3 |
| Solyc09g084480.2 | Solyc09g084480.3 |
| Solyc09g089500.2 | Solyc09g089500.3 |
| Solyc09g089530.2 | Solyc09g089530.3 |
| Solyc09g089610.2 | Solyc09g089610.3 |
| Solyc09g089630.2 | Solyc09g089630.3 |
| Solyc09g089670.2 | Solyc09g089670.3 |
| Solyc09g089690.2 | Solyc09g089690.3 |
| Solyc09g089700.2 | Solyc09g089700.3 |
| Solyc09g089710.2 | Solyc09g089710.3 |
| Solyc09g089730.2 | Solyc09g089730.3 |
| Solyc09g089740.2 | Solyc09g089740.3 |
| Solyc09g089770.2 | Solyc09g089770.3 |
| Solyc09g089780.2 | Solyc09g089780.3 |
| Solyc09g089790.2 | Solyc09g089790.3 |
| Solyc09g089800.1 | Solyc09g089800.2 |
| Solyc09g089820.1 | Solyc09g089820.2 |
| Solyc09g089850.2 | Solyc09g089850.3 |
| Solyc09g089870.2 | Solyc09g089870.3 |

| | |
|---|---|
| Solyc09g089880.2 | Solyc09g089880.3 |
| Solyc09g089990.2 | Solyc09g089990.3 |
| Solyc09g090030.1 | Solyc09g090030.2 |
| Solyc09g090040.2 | Solyc09g090040.3 |
| Solyc09g090060.2 | Solyc09g090060.3 |
| Solyc09g090090.1 | Solyc09g090090.2 |
| Solyc09g090100.2 | Solyc09g090100.3 |
| Solyc09g090110.2 | Solyc09g090110.3 |
| Solyc09g090130.2 | Solyc09g090130.3 |
| Solyc09g090160.2 | Solyc09g090160.3 |
| Solyc09g090190.2 | Solyc09g090190.3 |
| Solyc09g090210.2 | Solyc09g090210.3 |
| Solyc09g090220.2 | Solyc09g090220.3 |
| Solyc09g090230.1 | Solyc09g090230.2 |
| Solyc09g090270.2 | Solyc09g090270.3 |
| Solyc09g090300.2 | Solyc09g090300.3 |
| Solyc09g090320.1 | Solyc09g090320.2 |
| Solyc09g090330.2 | Solyc09g090330.3 |
| Solyc09g090350.2 | Solyc09g090350.3 |
| Solyc09g090360.2 | Solyc09g090360.3 |
| Solyc09g090370.2 | Solyc09g090370.3 |
| Solyc09g090410.2 | Solyc09g090410.3 |
| Solyc09g090430.2 | Solyc09g090430.3 |
| Solyc09g090460.2 | Solyc09g090460.3 |
| Solyc09g090470.2 | Solyc09g090470.3 |
| Solyc09g090500.2 | Solyc09g090500.3 |
| Solyc09g090510.2 | Solyc09g090510.3 |
| Solyc09g090520.2 | Solyc09g090520.3 |
| Solyc09g090550.2 | Solyc09g090550.3 |
| Solyc09g090580.2 | Solyc09g090580.3 |
| Solyc09g090600.2 | Solyc09g090600.3 |
| Solyc09g090610.2 | Solyc09g090610.3 |
| Solyc09g090640.2 | Solyc09g090640.3 |
| Solyc09g090650.2 | Solyc09g090650.3 |
| Solyc09g090660.2 | Solyc09g090660.3 |
| Solyc09g090670.2 | Solyc09g090670.3 |
| Solyc09g090680.2 | Solyc09g090680.3 |
| Solyc09g090710.2 | Solyc09g090710.3 |
| Solyc09g090720.2 | Solyc09g090720.3 |
| Solyc09g090730.1 | Solyc09g090730.2 |
| Solyc09g090750.2 | Solyc09g090750.3 |
| Solyc09g090770.1 | Solyc09g090770.2 |
| Solyc09g090790.2 | Solyc09g090790.3 |

| | |
|---|---|
| Solyc09g090800.1 | Solyc09g090800.2 |
| Solyc09g090840.2 | Solyc09g090840.3 |
| Solyc09g090850.2 | Solyc09g090850.3 |
| Solyc09g090870.2 | Solyc09g090870.3 |
| Solyc09g090930.2 | Solyc09g090930.3 |
| Solyc09g090940.2 | Solyc09g090940.3 |
| Solyc09g090980.2 | Solyc09g090980.3 |
| Solyc09g091020.2 | Solyc09g091020.3 |
| Solyc09g091030.2 | Solyc09g091030.3 |
| Solyc09g091040.2 | Solyc09g091040.3 |
| Solyc09g091060.2 | Solyc09g091060.3 |
| Solyc09g091090.1 | Solyc09g091090.2 |
| Solyc09g091120.2 | Solyc09g091120.3 |
| Solyc09g091180.2 | Solyc09g091180.3 |
| Solyc09g091190.2 | Solyc09g091190.3 |
| Solyc09g091200.2 | Solyc09g091200.3 |
| Solyc09g091260.2 | Solyc09g091260.3 |
| Solyc09g091390.2 | Solyc09g091390.3 |
| Solyc09g091410.1 | Solyc09g091410.2 |
| Solyc09g091440.2 | Solyc09g091440.3 |
| Solyc09g091450.2 | Solyc09g091450.3 |
| Solyc09g091460.2 | Solyc09g091460.3 |
| Solyc09g091470.2 | Solyc09g091470.3 |
| Solyc09g091500.2 | Solyc09g091500.3 |
| Solyc09g091510.2 | Solyc09g091510.3 |
| Solyc09g091520.1 | Solyc09g091520.2 |
| Solyc09g091530.1 | Solyc09g091530.2 |
| Solyc09g091540.1 | Solyc09g091540.2 |
| Solyc09g091550.2 | Solyc09g091550.3 |
| Solyc09g091560.2 | Solyc09g091560.3 |
| Solyc09g091580.2 | Solyc09g091580.3 |
| Solyc09g091590.2 | Solyc09g091590.3 |
| Solyc09g091610.2 | Solyc09g091610.3 |
| Solyc09g091650.2 | Solyc09g091650.3 |
| Solyc09g091660.2 | Solyc09g091660.3 |
| Solyc09g091670.2 | Solyc09g091670.3 |
| Solyc09g091750.2 | Solyc09g091750.3 |
| Solyc09g091780.2 | Solyc09g091780.3 |
| Solyc09g091790.2 | Solyc09g091790.3 |
| Solyc09g091870.1 | Solyc09g091870.2 |
| Solyc09g091910.1 | Solyc09g091910.2 |
| Solyc09g091920.2 | Solyc09g091920.3 |
| Solyc09g091930.2 | Solyc09g091930.3 |

| | |
|---|---|
| Solyc09g091960.2 | Solyc09g091960.3 |
| Solyc09g091980.2 | Solyc09g091980.3 |
| Solyc09g091990.2 | Solyc09g091990.3 |
| Solyc09g092040.2 | Solyc09g092040.3 |
| Solyc09g092130.2 | Solyc09g092130.3 |
| Solyc09g092140.2 | Solyc09g092140.3 |
| Solyc09g092180.2 | Solyc09g092180.3 |
| Solyc09g092240.2 | Solyc09g092240.3 |
| Solyc09g092250.2 | Solyc09g092250.3 |
| Solyc09g092270.2 | Solyc09g092270.3 |
| Solyc09g092300.2 | Solyc09g092300.3 |
| Solyc09g092320.2 | Solyc09g092320.3 |
| Solyc09g092360.2 | Solyc09g092360.3 |
| Solyc09g092380.2 | Solyc09g092380.3 |
| Solyc09g092400.1 | Solyc09g092400.2 |
| Solyc09g092420.1 | Solyc09g092420.2 |
| Solyc09g092430.2 | Solyc09g092430.3 |
| Solyc09g092450.2 | Solyc09g092450.3 |
| Solyc09g092460.3 | Solyc09g092460.4 |
| Solyc09g092490.2 | Solyc09g092490.3 |
| Solyc09g092520.2 | Solyc09g092520.3 |
| Solyc09g092585.1 | Solyc09g150158.1 |
| Solyc09g092590.1 | Solyc09g092590.2 |
| Solyc09g092600.2 | Solyc09g092600.3 |
| Solyc09g092620.2 | Solyc09g092620.3 |
| Solyc09g092640.2 | Solyc09g092640.3 |
| Solyc09g092670.2 | Solyc09g092670.3 |
| Solyc09g092690.2 | Solyc09g092690.3 |
| Solyc09g092700.2 | Solyc09g092700.3 |
| Solyc09g092720.2 | Solyc09g092720.3 |
| Solyc09g092750.2 | Solyc09g092750.3 |
| Solyc09g097760.2 | Solyc09g097760.3 |
| Solyc09g097770.2 | Solyc09g097770.3 |
| Solyc09g082810.2 | Solyc09g082810.3 |
| Solyc09g097810.2 | Solyc09g097810.3 |
| Solyc09g097820.2 | Solyc09g097820.3 |
| Solyc09g097840.1 | Solyc09g097840.2 |
| Solyc09g097900.2 | Solyc09g097900.3 |
| Solyc09g097960.2 | Solyc09g097960.3 |
| Solyc09g097970.2 | Solyc09g097970.3 |
| Solyc09g098010.2 | Solyc09g098010.3 |
| Solyc09g098770.1 | Solyc09g098770.2 |
| Solyc09g098040.2 | Solyc09g098040.3 |

| | |
|---|---|
| Solyc09g098060.2 | Solyc09g098060.3 |
| Solyc09g098070.2 | Solyc09g098070.3 |
| Solyc09g098120.2 | Solyc09g098120.3 |
| Solyc09g098150.2 | Solyc09g098150.3 |
| Solyc09g098160.2 | Solyc09g098160.3 |
| Solyc09g098190.2 | Solyc09g098190.3 |
| Solyc09g098210.1 | Solyc09g098210.2 |
| Solyc09g098220.2 | Solyc09g098220.3 |
| Solyc09g098230.2 | Solyc09g098230.3 |
| Solyc09g098280.2 | Solyc09g098280.3 |
| Solyc09g098290.2 | Solyc09g098290.3 |
| Solyc09g098300.2 | Solyc09g098300.3 |
| Solyc09g098310.2 | Solyc09g098310.3 |
| Solyc09g098330.2 | Solyc09g098330.3 |
| Solyc09g098340.2 | Solyc09g098340.3 |
| Solyc09g098390.2 | Solyc09g098390.3 |
| Solyc09g098510.2 | Solyc09g098510.3 |
| Solyc09g098540.2 | Solyc09g098540.3 |
| Solyc09g098580.2 | Solyc09g098580.3 |
| Solyc10g005180.2 | Solyc10g005180.3 |
| Solyc10g005195.1 | Solyc10g150100.1 |
| Solyc10g005200.2 | Solyc10g005200.3 |
| Solyc10g005220.2 | Solyc10g005220.3 |
| Solyc10g005240.2 | Solyc10g005240.3 |
| Solyc10g005250.2 | Solyc10g005250.3 |
| Solyc10g005260.2 | Solyc10g005260.3 |
| Solyc10g005300.2 | Solyc10g005300.3 |
| Solyc10g005310.2 | Solyc10g005310.3 |
| Solyc10g005320.2 | Solyc10g005320.3 |
| Solyc10g005330.2 | Solyc10g005330.3 |
| Solyc10g005340.2 | Solyc10g005340.3 |
| Solyc10g005350.2 | Solyc10g005350.3 |
| Solyc10g005360.3 | Solyc10g005360.4 |
| Solyc10g005370.2 | Solyc10g005370.3 |
| Solyc10g005400.2 | Solyc10g005400.3 |
| Solyc10g005450.1 | Solyc10g005450.2 |
| Solyc10g005460.2 | Solyc10g005460.3 |
| Solyc10g005480.2 | Solyc10g005480.3 |
| Solyc10g005500.2 | Solyc10g005500.3 |
| Solyc10g005510.2 | Solyc10g005510.3 |
| Solyc10g005520.2 | Solyc10g005520.3 |
| Solyc10g005530.2 | Solyc10g005530.3 |
| Solyc10g005560.2 | Solyc10g005560.3 |

| | |
|---|---|
| Solyc10g005620.2 | Solyc10g005620.3 |
| Solyc10g005630.2 | Solyc10g005630.3 |
| Solyc10g005650.2 | Solyc10g005650.3 |
| Solyc10g005660.2 | Solyc10g005660.3 |
| Solyc10g005680.1 | Solyc10g005680.2 |
| Solyc10g005690.2 | Solyc10g005690.3 |
| Solyc10g005710.1 | Solyc10g005710.2 |
| Solyc10g005730.2 | Solyc10g005730.3 |
| Solyc10g005750.2 | Solyc10g005750.3 |
| Solyc10g005780.2 | Solyc10g005780.3 |
| Solyc10g005790.2 | Solyc10g005790.3 |
| Solyc10g005800.2 | Solyc10g005800.3 |
| Solyc10g005830.2 | Solyc10g005830.3 |
| Solyc10g005860.2 | Solyc10g005860.3 |
| Solyc10g005880.2 | Solyc10g005880.3 |
| Solyc10g005890.2 | Solyc10g005890.3 |
| Solyc10g005900.2 | Solyc10g005900.3 |
| Solyc10g005930.1 | Solyc10g005930.2 |
| Solyc10g005950.2 | Solyc10g005950.3 |
| Solyc10g006010.2 | Solyc10g006010.3 |
| Solyc10g006020.2 | Solyc10g006020.3 |
| Solyc10g006030.2 | Solyc10g006030.3 |
| Solyc10g006050.2 | Solyc10g006050.3 |
| Solyc10g006070.2 | Solyc10g006070.3 |
| Solyc10g006100.2 | Solyc10g006100.3 |
| Solyc10g006110.2 | Solyc10g006110.3 |
| Solyc10g006120.1 | Solyc10g006120.2 |
| Solyc10g006140.2 | Solyc10g006140.3 |
| Solyc10g006150.2 | Solyc10g006150.3 |
| Solyc10g006230.2 | Solyc10g006230.3 |
| Solyc10g006240.2 | Solyc10g006240.3 |
| Solyc10g006250.2 | Solyc10g006250.3 |
| Solyc10g006260.2 | Solyc10g006260.3 |
| Solyc10g006290.2 | Solyc10g006290.3 |
| Solyc10g006310.2 | Solyc10g006310.3 |
| Solyc10g006320.2 | Solyc10g006320.3 |
| Solyc10g006330.2 | Solyc10g006330.3 |
| Solyc10g006340.2 | Solyc10g006340.3 |
| Solyc10g006350.2 | Solyc10g006350.3 |
| Solyc10g006370.2 | Solyc10g006370.3 |
| Solyc10g006390.2 | Solyc10g006390.3 |
| Solyc10g006400.2 | Solyc10g006400.3 |
| Solyc10g006410.2 | Solyc10g006410.3 |

| | |
|---|---|
| Solyc10g006440.2 | Solyc10g006440.3 |
| Solyc10g006450.2 | Solyc10g006450.3 |
| Solyc10g006460.2 | Solyc10g006460.3 |
| Solyc10g006470.2 | Solyc10g006470.3 |
| Solyc10g006480.1 | Solyc10g006480.2 |
| Solyc10g006490.2 | Solyc10g006490.3 |
| Solyc10g006510.2 | Solyc10g006510.3 |
| Solyc10g006520.2 | Solyc10g006520.3 |
| Solyc10g006580.2 | Solyc10g006580.3 |
| Solyc10g006640.2 | Solyc10g006640.3 |
| Solyc10g006650.2 | Solyc10g006650.3 |
| Solyc10g006660.2 | Solyc10g006660.3 |
| Solyc10g006680.1 | Solyc10g006680.2 |
| Solyc10g006760.2 | Solyc10g006760.3 |
| Solyc10g006880.2 | Solyc10g006880.3 |
| Solyc10g006900.2 | Solyc10g006900.3 |
| Solyc10g006920.2 | Solyc10g006920.3 |
| Solyc10g006970.2 | Solyc10g006970.3 |
| Solyc10g006980.2 | Solyc10g006980.3 |
| Solyc10g006990.2 | Solyc10g006990.3 |
| Solyc10g007000.2 | Solyc10g007000.3 |
| Solyc10g007030.1 | Solyc10g007030.2 |
| Solyc10g007040.2 | Solyc10g007040.3 |
| Solyc10g007050.2 | Solyc10g007050.3 |
| Solyc10g007100.2 | Solyc10g007100.3 |
| Solyc10g007103.1 | Solyc10g150101.1 |
| Solyc10g007107.1 | Solyc10g150102.1 |
| Solyc10g007110.2 | Solyc10g007110.3 |
| Solyc10g007140.2 | Solyc10g007140.3 |
| Solyc10g007156.1 | Solyc10g007156.2 |
| Solyc10g007160.1 | Solyc10g007160.2 |
| Solyc10g007170.1 | Solyc10g007170.2 |
| Solyc10g007200.2 | Solyc10g007200.3 |
| Solyc10g007230.1 | Solyc10g007230.2 |
| Solyc10g007260.2 | Solyc10g007260.3 |
| Solyc10g007300.1 | Solyc10g007300.2 |
| Solyc10g007320.2 | Solyc10g007320.3 |
| Solyc10g007370.1 | Solyc10g007370.2 |
| Solyc10g007390.2 | Solyc10g007390.3 |
| Solyc10g007410.3 | Solyc10g007410.4 |
| Solyc10g007440.2 | Solyc10g007440.3 |
| Solyc10g007450.1 | Solyc10g007450.2 |
| Solyc10g007510.1 | Solyc10g007510.2 |

| | |
|---|---|
| Solyc10g007530.1 | Solyc10g007530.2 |
| Solyc10g007540.1 | Solyc10g007540.2 |
| Solyc10g007550.2 | Solyc10g007550.3 |
| Solyc10g007560.1 | Solyc10g007560.2 |
| Solyc10g007600.2 | Solyc10g007600.3 |
| Solyc10g007650.2 | Solyc10g007650.3 |
| Solyc10g007660.2 | Solyc10g007660.3 |
| Solyc10g007690.2 | Solyc10g007690.3 |
| Solyc10g007700.2 | Solyc10g007700.3 |
| Solyc10g007710.2 | Solyc10g007710.3 |
| Solyc10g007740.2 | Solyc10g007740.3 |
| Solyc10g007770.2 | Solyc10g007770.3 |
| Solyc10g007790.2 | Solyc10g007790.3 |
| Solyc10g007800.2 | Solyc10g007800.3 |
| Solyc10g007810.2 | Solyc10g007810.3 |
| Solyc10g007820.2 | Solyc10g007820.3 |
| Solyc10g007840.2 | Solyc10g007840.3 |
| Solyc10g007850.2 | Solyc10g007850.3 |
| Solyc10g007860.2 | Solyc10g007860.3 |
| Solyc10g007870.2 | Solyc10g007870.3 |
| Solyc10g007970.1 | Solyc10g007970.2 |
| Solyc10g007980.2 | Solyc10g007980.3 |
| Solyc10g007990.2 | Solyc10g007990.3 |
| Solyc10g008020.2 | Solyc10g008020.3 |
| Solyc10g008100.2 | Solyc10g008100.3 |
| Solyc10g008140.2 | Solyc10g008140.3 |
| Solyc10g008160.2 | Solyc10g008160.3 |
| Solyc10g008180.1 | Solyc10g008180.2 |
| Solyc10g008190.2 | Solyc10g008190.3 |
| Solyc10g008205.1 | Solyc10g008205.2 |
| Solyc10g008210.2 | Solyc10g008210.3 |
| Solyc10g008240.2 | Solyc10g008240.3 |
| Solyc10g008260.1 | Solyc10g008260.2 |
| Solyc10g008270.2 | Solyc10g008270.3 |
| Solyc10g008280.2 | Solyc10g008280.3 |
| Solyc10g008300.2 | Solyc10g008300.3 |
| Solyc10g008390.2 | Solyc10g008390.3 |
| Solyc10g008440.2 | Solyc10g008440.3 |
| Solyc10g008450.1 | Solyc10g008450.2 |
| Solyc10g008460.2 | Solyc10g008460.3 |
| Solyc10g008470.2 | Solyc10g008470.3 |
| Solyc10g008480.2 | Solyc10g008480.3 |
| Solyc10g008490.2 | Solyc10g008490.3 |

| | |
|---|---|
| Solyc10g008515.1 | Solyc10g150103.1 |
| Solyc10g008520.2 | Solyc10g008520.3 |
| Solyc10g008550.2 | Solyc10g008550.3 |
| Solyc10g008560.2 | Solyc10g008560.3 |
| Solyc10g008570.2 | Solyc10g008570.3 |
| Solyc10g008620.2 | Solyc10g008620.3 |
| Solyc10g008630.2 | Solyc10g008630.3 |
| Solyc10g008640.2 | Solyc10g008640.3 |
| Solyc10g008650.1 | Solyc10g008650.2 |
| Solyc10g008660.1 | Solyc10g008660.2 |
| Solyc10g008670.2 | Solyc10g008670.3 |
| Solyc10g008710.2 | Solyc10g008710.3 |
| Solyc10g008740.2 | Solyc10g008740.3 |
| Solyc10g008750.2 | Solyc10g008750.3 |
| Solyc10g008760.2 | Solyc10g008760.3 |
| Solyc10g008770.2 | Solyc10g008770.3 |
| Solyc10g008790.2 | Solyc10g008790.3 |
| Solyc10g008810.2 | Solyc10g008810.3 |
| Solyc10g008840.2 | Solyc10g008840.3 |
| Solyc10g008870.2 | Solyc10g008870.3 |
| Solyc10g008880.2 | Solyc10g008880.3 |
| Solyc10g008890.2 | Solyc10g008890.3 |
| Solyc10g008900.2 | Solyc10g008900.3 |
| Solyc10g008950.2 | Solyc10g008950.3 |
| Solyc10g009020.2 | Solyc10g009020.3 |
| Solyc10g009050.2 | Solyc10g009050.3 |
| Solyc10g009070.2 | Solyc10g009070.3 |
| Solyc10g009090.2 | Solyc10g009090.3 |
| Solyc10g009120.1 | Solyc10g009120.2 |
| Solyc10g009140.2 | Solyc10g009140.3 |
| Solyc10g009150.2 | Solyc10g009150.3 |
| Solyc10g009160.2 | Solyc10g009160.3 |
| Solyc10g009170.1 | Solyc10g009170.2 |
| Solyc10g009180.2 | Solyc10g009180.3 |
| Solyc10g009200.2 | Solyc10g009200.3 |
| Solyc10g009220.2 | Solyc10g009220.3 |
| Solyc10g009240.2 | Solyc10g009240.3 |
| Solyc10g009270.2 | Solyc10g009270.3 |
| Solyc10g009330.2 | Solyc10g009330.3 |
| Solyc10g009390.2 | Solyc10g009390.3 |
| Solyc10g009420.2 | Solyc10g009420.3 |
| Solyc10g009430.2 | Solyc10g009430.3 |
| Solyc10g009490.1 | Solyc10g009490.2 |

| | |
|---|---|
| Solyc10g009503.1 | Solyc10g150104.1 |
| Solyc10g009507.1 | Solyc10g150105.1 |
| Solyc10g009550.2 | Solyc10g009550.3 |
| Solyc10g009570.2 | Solyc10g009570.3 |
| Solyc10g009610.1 | Solyc10g009610.2 |
| Solyc10g011650.2 | Solyc10g011650.3 |
| Solyc10g011660.2 | Solyc10g011660.3 |
| Solyc10g011670.2 | Solyc10g011670.3 |
| Solyc10g011690.2 | Solyc10g011690.3 |
| Solyc10g011730.2 | Solyc10g011730.3 |
| Solyc10g011740.2 | Solyc10g011740.3 |
| Solyc10g011750.2 | Solyc10g011750.3 |
| Solyc10g011760.2 | Solyc10g011760.3 |
| Solyc10g011790.2 | Solyc10g011790.3 |
| Solyc10g011870.2 | Solyc10g011870.3 |
| Solyc10g011900.2 | Solyc10g011900.3 |
| Solyc10g011980.2 | Solyc10g011980.3 |
| Solyc10g012050.2 | Solyc10g012050.3 |
| Solyc10g012060.2 | Solyc10g012060.3 |
| Solyc10g012090.1 | Solyc10g012090.2 |
| Solyc10g012140.1 | Solyc10g012140.2 |
| Solyc10g012160.2 | Solyc10g012160.3 |
| Solyc10g012170.2 | Solyc10g012170.3 |
| Solyc10g012240.2 | Solyc10g012240.3 |
| Solyc10g012270.1 | Solyc10g012270.2 |
| Solyc10g012370.2 | Solyc10g012370.3 |
| Solyc10g012380.1 | Solyc10g012380.2 |
| Solyc10g012430.2 | Solyc10g012430.3 |
| Solyc10g017510.2 | Solyc10g017510.3 |
| Solyc10g017520.2 | Solyc10g017520.3 |
| Solyc10g017530.2 | Solyc10g017530.3 |
| Solyc10g017570.2 | Solyc10g017570.3 |
| Solyc10g017580.2 | Solyc10g017580.3 |
| Solyc10g017590.2 | Solyc10g017590.3 |
| Solyc10g017600.2 | Solyc10g017600.3 |
| Solyc10g017620.2 | Solyc10g017620.3 |
| Solyc10g017750.1 | Solyc10g017750.2 |
| Solyc10g017810.1 | Solyc10g017810.2 |
| Solyc10g017850.2 | Solyc10g017850.3 |
| Solyc10g017880.1 | Solyc10g017880.2 |
| Solyc10g017950.1 | Solyc10g017950.2 |
| Solyc10g017960.1 | Solyc10g017960.2 |
| Solyc10g017990.1 | Solyc10g017990.2 |

| | |
|---|---|
| Solyc10g018010.1 | Solyc10g018010.2 |
| Solyc10g018023.1 | Solyc10g150106.1 |
| Solyc10g018050.1 | Solyc10g018050.2 |
| Solyc10g018060.1 | Solyc10g018060.2 |
| Solyc10g018120.1 | Solyc10g018120.2 |
| Solyc10g018140.1 | Solyc10g018140.2 |
| Solyc10g018150.1 | Solyc10g018150.2 |
| Solyc10g018160.1 | Solyc10g018160.2 |
| Solyc10g018190.1 | Solyc10g018190.2 |
| Solyc10g018200.1 | Solyc10g018200.2 |
| Solyc10g018250.1 | Solyc10g018250.2 |
| Solyc10g018290.1 | Solyc10g018290.2 |
| Solyc10g018350.1 | Solyc10g018350.2 |
| Solyc10g018357.1 | Solyc10g150107.1 |
| Solyc10g018510.1 | Solyc10g018510.2 |
| Solyc10g018530.1 | Solyc10g018530.2 |
| Solyc10g018540.1 | Solyc10g018540.2 |
| Solyc10g018560.1 | Solyc10g018560.2 |
| Solyc10g018590.1 | Solyc10g018590.2 |
| Solyc10g018600.1 | Solyc10g018600.2 |
| Solyc10g018610.1 | Solyc10g018610.2 |
| Solyc10g018705.1 | Solyc10g150108.1 |
| Solyc10g018817.1 | Solyc10g150109.1 |
| Solyc10g018850.1 | Solyc10g018850.2 |
| Solyc10g018870.1 | Solyc10g018870.2 |
| Solyc10g018920.1 | Solyc10g018920.2 |
| Solyc10g019040.1 | Solyc10g019040.2 |
| Solyc10g019050.1 | Solyc10g019050.2 |
| Solyc10g019095.1 | Solyc10g150110.1 |
| Solyc10g019100.1 | Solyc10g019100.2 |
| Solyc10g019110.1 | Solyc10g019110.2 |
| Solyc10g019160.1 | Solyc10g019160.2 |
| Solyc10g019170.1 | Solyc10g019170.2 |
| Solyc10g019190.1 | Solyc10g019190.2 |
| Solyc10g019230.1 | Solyc10g019230.2 |
| Solyc10g019280.1 | Solyc10g019280.2 |
| Solyc10g024320.1 | Solyc10g024320.2 |
| Solyc10g024340.1 | Solyc10g024340.2 |
| Solyc10g024360.1 | Solyc10g024360.2 |
| Solyc10g024410.1 | Solyc10g024410.2 |
| Solyc10g024490.1 | Solyc10g024490.2 |
| Solyc10g037930.1 | Solyc10g037930.2 |
| Solyc10g037950.1 | Solyc10g037950.2 |

| | |
|---|---|
| Solyc10g037980.1 | Solyc10g037980.2 |
| Solyc10g038020.1 | Solyc10g038020.2 |
| Solyc10g038060.1 | Solyc10g038060.2 |
| Solyc10g038150.1 | Solyc10g038150.2 |
| Solyc10g038190.1 | Solyc10g038190.2 |
| Solyc10g039190.1 | Solyc10g039190.2 |
| Solyc10g039195.1 | Solyc10g150111.1 |
| Solyc10g039197.1 | Solyc10g039197.2 |
| Solyc10g039250.1 | Solyc10g039250.2 |
| Solyc10g039270.1 | Solyc10g039270.2 |
| Solyc10g039280.1 | Solyc10g039280.2 |
| Solyc10g039290.1 | Solyc10g039290.2 |
| Solyc10g039360.1 | Solyc10g039360.2 |
| Solyc10g039363.1 | Solyc10g150112.1 |
| Solyc10g039367.1 | Solyc10g150113.1 |
| Solyc10g061960.1 | Solyc10g061960.2 |
| Solyc10g061970.1 | Solyc10g061970.2 |
| Solyc10g062020.1 | Solyc10g062020.2 |
| Solyc10g062085.1 | Solyc10g062085.2 |
| Solyc10g062110.1 | Solyc10g062110.2 |
| Solyc10g062180.1 | Solyc10g062180.2 |
| Solyc10g062200.1 | Solyc10g062200.2 |
| Solyc10g062220.1 | Solyc10g062220.2 |
| Solyc10g062340.1 | Solyc10g062340.2 |
| Solyc10g044440.1 | Solyc10g044440.2 |
| Solyc10g044480.1 | Solyc10g044480.2 |
| Solyc10g044510.1 | Solyc10g044510.2 |
| Solyc10g044680.1 | Solyc10g044680.2 |
| Solyc10g044700.1 | Solyc10g044700.2 |
| Solyc10g044710.1 | Solyc10g044710.2 |
| Solyc10g044730.1 | Solyc10g044730.2 |
| Solyc10g044740.1 | Solyc10g044740.2 |
| Solyc10g044780.1 | Solyc10g044780.2 |
| Solyc10g044840.1 | Solyc10g044840.2 |
| Solyc10g044910.1 | Solyc10g044910.2 |
| Solyc10g044920.1 | Solyc10g044920.2 |
| Solyc10g045080.1 | Solyc10g045080.2 |
| Solyc10g045100.1 | Solyc10g045100.2 |
| Solyc10g045107.1 | Solyc10g150114.1 |
| Solyc10g045153.1 | Solyc10g150115.1 |
| Solyc10g045240.1 | Solyc10g045240.2 |
| Solyc10g045270.1 | Solyc10g045270.2 |
| Solyc10g045290.1 | Solyc10g045290.2 |

| | |
|---|---|
| Solyc10g045310.1 | Solyc10g045310.2 |
| Solyc10g045320.1 | Solyc10g045320.2 |
| Solyc10g045380.1 | Solyc10g045380.2 |
| Solyc10g045395.1 | Solyc10g150116.1 |
| Solyc10g045430.1 | Solyc10g045430.2 |
| Solyc10g045433.1 | Solyc10g150117.1 |
| Solyc10g045457.1 | Solyc10g150118.1 |
| Solyc10g045490.1 | Solyc10g045490.2 |
| Solyc10g045520.1 | Solyc10g045520.2 |
| Solyc10g045540.1 | Solyc10g045540.2 |
| Solyc10g045550.1 | Solyc10g045550.2 |
| Solyc10g045555.1 | Solyc10g150119.1 |
| Solyc10g045600.1 | Solyc10g045600.2 |
| Solyc10g045640.1 | Solyc10g045640.2 |
| Solyc10g045740.1 | Solyc10g045740.2 |
| Solyc10g046830.1 | Solyc10g046830.2 |
| Solyc10g046970.1 | Solyc10g046970.2 |
| Solyc10g046990.1 | Solyc10g046990.2 |
| Solyc10g047000.1 | Solyc10g047000.2 |
| Solyc10g047050.1 | Solyc10g047050.2 |
| Solyc10g047060.1 | Solyc10g047060.2 |
| Solyc10g047083.1 | Solyc10g150120.1 |
| Solyc10g047087.1 | Solyc10g150121.1 |
| Solyc10g047090.1 | Solyc10g047090.2 |
| Solyc10g047110.1 | Solyc10g047110.2 |
| Solyc10g047120.1 | Solyc10g047120.2 |
| Solyc10g047130.1 | Solyc10g047130.2 |
| Solyc10g047140.1 | Solyc10g047140.2 |
| Solyc10g047170.1 | Solyc10g047170.2 |
| Solyc10g047240.1 | Solyc10g047240.2 |
| Solyc10g047270.1 | Solyc10g047270.2 |
| Solyc10g047290.1 | Solyc10g047290.2 |
| Solyc10g047300.1 | Solyc10g047300.2 |
| Solyc10g047320.1 | Solyc10g047320.2 |
| Solyc10g047340.1 | Solyc10g047340.2 |
| Solyc10g047350.1 | Solyc10g047350.2 |
| Solyc10g047360.1 | Solyc10g047360.2 |
| Solyc10g047370.1 | Solyc10g047370.2 |
| Solyc10g047500.1 | Solyc10g047500.2 |
| Solyc10g047530.1 | Solyc10g047530.2 |
| Solyc10g047570.1 | Solyc10g047570.2 |
| Solyc10g047630.1 | Solyc10g047630.2 |
| Solyc10g047670.1 | Solyc10g047670.2 |

| | |
|---|---|
| Solyc10g047685.1 | Solyc10g150122.1 |
| Solyc10g047823.1 | Solyc10g150123.1 |
| Solyc10g047827.1 | Solyc10g150124.1 |
| Solyc10g047900.1 | Solyc10g047900.2 |
| Solyc10g047930.1 | Solyc10g047930.2 |
| Solyc10g047950.1 | Solyc10g047950.2 |
| Solyc10g048010.1 | Solyc10g048010.2 |
| Solyc10g048030.1 | Solyc10g048030.2 |
| Solyc10g048065.1 | Solyc10g150125.1 |
| Solyc10g048070.1 | Solyc10g048070.2 |
| Solyc10g048140.1 | Solyc10g048140.2 |
| Solyc10g049255.1 | Solyc10g150126.1 |
| Solyc10g049260.1 | Solyc10g049260.2 |
| Solyc10g049270.1 | Solyc10g049270.2 |
| Solyc10g049280.1 | Solyc10g049280.2 |
| Solyc10g049360.1 | Solyc10g049360.2 |
| Solyc10g049370.1 | Solyc10g049370.2 |
| Solyc10g049420.1 | Solyc10g049420.2 |
| Solyc10g049450.1 | Solyc10g049450.2 |
| Solyc10g049580.1 | Solyc10g049580.2 |
| Solyc10g049620.1 | Solyc10g049620.2 |
| Solyc10g049630.1 | Solyc10g049630.2 |
| Solyc10g049640.1 | Solyc10g049640.2 |
| Solyc10g049660.1 | Solyc10g049660.2 |
| Solyc10g049710.1 | Solyc10g049710.2 |
| Solyc10g049720.1 | Solyc10g049720.2 |
| Solyc10g049800.1 | Solyc10g049800.2 |
| Solyc10g049910.1 | Solyc10g049910.2 |
| Solyc10g049970.1 | Solyc10g049970.2 |
| Solyc10g050030.1 | Solyc10g050030.2 |
| Solyc10g050060.1 | Solyc10g050060.2 |
| Solyc10g050095.1 | Solyc10g150127.1 |
| Solyc10g050160.1 | Solyc10g050160.2 |
| Solyc10g050320.1 | Solyc10g050320.2 |
| Solyc10g050325.1 | Solyc10g050325.2 |
| Solyc10g050370.1 | Solyc10g050370.2 |
| Solyc10g050440.1 | Solyc10g050440.2 |
| Solyc10g050450.1 | Solyc10g050450.2 |
| Solyc10g050500.1 | Solyc10g050500.2 |
| Solyc10g050530.1 | Solyc10g050530.2 |
| Solyc10g050540.1 | Solyc10g050540.2 |
| Solyc10g050690.1 | Solyc10g050690.2 |
| Solyc10g050720.1 | Solyc10g050720.2 |

| | |
|---|---|
| Solyc10g050730.1 | Solyc10g050730.2 |
| Solyc10g050735.1 | Solyc10g150128.1 |
| Solyc10g050850.1 | Solyc10g050850.2 |
| Solyc10g050880.1 | Solyc10g050880.2 |
| Solyc10g050890.1 | Solyc10g050890.2 |
| Solyc10g050920.1 | Solyc10g050920.2 |
| Solyc10g050990.1 | Solyc10g050990.2 |
| Solyc10g050997.1 | Solyc10g150129.1 |
| Solyc10g051020.1 | Solyc10g051020.2 |
| Solyc10g051050.2 | Solyc10g051050.3 |
| Solyc10g051160.1 | Solyc10g051160.2 |
| Solyc10g051170.1 | Solyc10g051170.2 |
| Solyc10g051200.1 | Solyc10g051200.2 |
| Solyc10g051270.1 | Solyc10g051270.2 |
| Solyc10g051273.1 | Solyc10g150130.1 |
| Solyc10g051277.1 | Solyc10g150131.1 |
| Solyc10g051300.1 | Solyc10g051300.2 |
| Solyc10g051310.1 | Solyc10g051310.2 |
| Solyc10g051340.1 | Solyc10g051340.2 |
| Solyc10g051350.1 | Solyc10g051350.2 |
| Solyc10g052490.1 | Solyc10g052490.2 |
| Solyc10g052500.1 | Solyc10g052500.2 |
| Solyc10g052510.1 | Solyc10g052510.2 |
| Solyc10g052520.1 | Solyc10g052520.2 |
| Solyc10g052585.1 | Solyc10g150132.1 |
| Solyc10g052605.1 | Solyc10g150133.1 |
| Solyc10g052660.3 | Solyc10g052660.4 |
| Solyc10g052670.1 | Solyc10g052670.2 |
| Solyc10g052875.1 | Solyc10g150134.1 |
| Solyc10g053890.1 | Solyc10g053890.2 |
| Solyc10g053900.1 | Solyc10g053900.2 |
| Solyc10g053960.1 | Solyc10g053960.2 |
| Solyc10g053970.1 | Solyc10g053970.2 |
| Solyc10g053980.1 | Solyc10g053980.2 |
| Solyc10g054030.1 | Solyc10g054030.2 |
| Solyc10g054090.1 | Solyc10g054090.2 |
| Solyc10g054105.1 | Solyc10g150135.1 |
| Solyc10g054280.1 | Solyc10g054280.2 |
| Solyc10g054320.1 | Solyc10g054320.2 |
| Solyc10g054350.1 | Solyc10g054350.2 |
| Solyc10g054420.1 | Solyc10g054420.2 |
| Solyc10g054490.1 | Solyc10g054490.2 |
| Solyc10g054560.1 | Solyc10g054560.2 |

| | |
|---|---|
| Solyc10g054570.1 | Solyc10g054570.2 |
| Solyc10g054590.1 | Solyc10g054590.2 |
| Solyc10g054660.1 | Solyc10g054660.2 |
| Solyc10g054733.1 | Solyc10g150136.1 |
| Solyc10g054737.1 | Solyc10g150137.1 |
| Solyc10g054780.1 | Solyc10g054780.2 |
| Solyc10g054790.1 | Solyc10g054790.2 |
| Solyc10g054800.1 | Solyc10g054800.2 |
| Solyc10g054810.1 | Solyc10g054810.2 |
| Solyc10g054820.1 | Solyc10g054820.2 |
| Solyc10g054840.1 | Solyc10g054840.2 |
| Solyc10g054850.1 | Solyc10g054850.2 |
| Solyc10g054860.1 | Solyc10g054860.2 |
| Solyc10g054900.1 | Solyc10g054900.2 |
| Solyc10g055050.1 | Solyc10g055050.2 |
| Solyc10g055085.1 | Solyc10g150138.1 |
| Solyc10g055110.1 | Solyc10g055110.2 |
| Solyc10g055130.1 | Solyc10g055130.2 |
| Solyc10g055240.1 | Solyc10g055240.2 |
| Solyc10g055260.1 | Solyc10g055260.2 |
| Solyc10g055300.1 | Solyc10g055300.2 |
| Solyc10g055370.1 | Solyc10g055370.2 |
| Solyc10g055390.1 | Solyc10g055390.2 |
| Solyc10g055410.1 | Solyc10g055410.2 |
| Solyc10g055420.1 | Solyc10g055420.2 |
| Solyc10g055450.1 | Solyc10g055450.2 |
| Solyc10g055460.1 | Solyc10g055460.2 |
| Solyc10g055470.1 | Solyc10g055470.2 |
| Solyc10g055540.1 | Solyc10g055540.2 |
| Solyc10g055560.1 | Solyc10g055560.2 |
| Solyc10g055580.1 | Solyc10g055580.2 |
| Solyc10g055620.1 | Solyc10g055620.2 |
| Solyc10g055630.1 | Solyc10g055630.2 |
| Solyc10g055640.1 | Solyc10g055640.2 |
| Solyc10g055650.1 | Solyc10g055650.2 |
| Solyc10g055670.1 | Solyc10g055670.2 |
| Solyc10g055700.1 | Solyc10g055700.2 |
| Solyc10g055710.1 | Solyc10g055710.2 |
| Solyc10g055720.1 | Solyc10g055720.2 |
| Solyc10g055740.1 | Solyc10g055740.2 |
| Solyc10g055760.1 | Solyc10g055760.2 |
| Solyc10g055770.1 | Solyc10g055770.2 |
| Solyc10g055800.1 | Solyc10g055800.2 |

| | |
|---|---|
| Solyc10g055810.1 | Solyc10g055810.2 |
| Solyc10g073350.1 | Solyc10g073350.2 |
| Solyc10g074430.1 | Solyc10g074430.2 |
| Solyc10g074440.1 | Solyc10g074440.2 |
| Solyc10g074500.1 | Solyc10g074500.2 |
| Solyc10g074530.1 | Solyc10g074530.2 |
| Solyc10g074560.1 | Solyc10g074560.2 |
| Solyc10g074570.1 | Solyc10g074570.2 |
| Solyc10g074590.1 | Solyc10g074590.2 |
| Solyc10g074650.1 | Solyc10g074650.2 |
| Solyc10g074680.1 | Solyc10g074680.2 |
| Solyc10g074700.1 | Solyc10g074700.2 |
| Solyc10g074710.1 | Solyc10g074710.2 |
| Solyc10g074720.2 | Solyc10g074720.3 |
| Solyc10g074740.1 | Solyc10g074740.2 |
| Solyc10g074790.1 | Solyc10g074790.2 |
| Solyc10g074810.1 | Solyc10g074810.2 |
| Solyc10g074860.1 | Solyc10g074860.2 |
| Solyc10g074870.1 | Solyc10g074870.2 |
| Solyc10g074890.1 | Solyc10g074890.2 |
| Solyc10g074900.1 | Solyc10g074900.2 |
| Solyc10g074910.1 | Solyc10g074910.2 |
| Solyc10g074920.1 | Solyc10g074920.2 |
| Solyc10g074940.1 | Solyc10g074940.2 |
| Solyc10g074950.1 | Solyc10g074950.2 |
| Solyc10g074980.1 | Solyc10g074980.2 |
| Solyc10g074990.1 | Solyc10g074990.2 |
| Solyc10g075010.1 | Solyc10g075010.2 |
| Solyc10g075050.1 | Solyc10g075050.2 |
| Solyc10g075060.1 | Solyc10g075060.2 |
| Solyc10g075100.1 | Solyc10g075100.2 |
| Solyc10g075110.1 | Solyc10g075110.2 |
| Solyc10g075150.1 | Solyc10g075150.2 |
| Solyc10g075175.1 | Solyc10g150139.1 |
| Solyc10g076190.1 | Solyc10g076190.2 |
| Solyc10g076210.1 | Solyc10g076210.2 |
| Solyc10g076250.1 | Solyc10g076250.2 |
| Solyc10g076260.1 | Solyc10g076260.2 |
| Solyc10g076290.1 | Solyc10g076290.2 |
| Solyc10g076300.1 | Solyc10g076300.2 |
| Solyc10g076350.1 | Solyc10g076350.2 |
| Solyc10g076380.1 | Solyc10g076380.2 |
| Solyc10g076390.1 | Solyc10g076390.2 |

| | |
|---|---|
| Solyc10g076400.1 | Solyc10g076400.2 |
| Solyc10g076450.1 | Solyc10g076450.2 |
| Solyc10g076460.1 | Solyc10g076460.2 |
| Solyc10g076480.1 | Solyc10g076480.2 |
| Solyc10g076510.1 | Solyc10g076510.2 |
| Solyc10g076530.1 | Solyc10g076530.2 |
| Solyc10g076570.1 | Solyc10g076570.2 |
| Solyc10g076600.1 | Solyc10g076600.2 |
| Solyc10g076610.1 | Solyc10g076610.2 |
| Solyc10g076635.1 | Solyc10g150140.1 |
| Solyc10g076660.1 | Solyc10g076660.2 |
| Solyc10g076690.1 | Solyc10g076690.2 |
| Solyc10g076700.1 | Solyc10g076700.2 |
| Solyc10g076720.1 | Solyc10g076720.2 |
| Solyc10g076750.1 | Solyc10g076750.2 |
| Solyc10g076790.1 | Solyc10g076790.2 |
| Solyc10g076800.1 | Solyc10g076800.2 |
| Solyc10g076820.1 | Solyc10g076820.2 |
| Solyc10g076830.1 | Solyc10g076830.2 |
| Solyc10g076840.1 | Solyc10g076840.2 |
| Solyc10g076860.1 | Solyc10g076860.2 |
| Solyc10g076870.1 | Solyc10g076870.2 |
| Solyc10g076880.1 | Solyc10g076880.2 |
| Solyc10g076910.1 | Solyc10g076910.2 |
| Solyc10g077000.2 | Solyc10g077000.3 |
| Solyc10g077010.1 | Solyc10g077010.2 |
| Solyc10g077020.1 | Solyc10g077020.2 |
| Solyc10g077030.1 | Solyc10g077030.2 |
| Solyc10g077040.1 | Solyc10g077040.2 |
| Solyc10g077050.1 | Solyc10g077050.2 |
| Solyc10g077070.1 | Solyc10g077070.2 |
| Solyc10g077090.1 | Solyc10g077090.2 |
| Solyc10g078160.1 | Solyc10g078160.2 |
| Solyc10g078190.1 | Solyc10g078190.2 |
| Solyc10g078220.1 | Solyc10g078220.2 |
| Solyc10g078240.1 | Solyc10g078240.2 |
| Solyc10g078270.1 | Solyc10g078270.2 |
| Solyc10g078280.2 | Solyc10g078280.3 |
| Solyc10g078300.1 | Solyc10g078300.2 |
| Solyc10g078310.1 | Solyc10g078310.2 |
| Solyc10g078320.1 | Solyc10g078320.2 |
| Solyc10g078330.1 | Solyc10g078330.2 |
| Solyc10g078340.1 | Solyc10g078340.2 |

| | |
|---|---|
| Solyc10g078350.1 | Solyc10g078350.2 |
| Solyc10g078370.1 | Solyc10g078370.2 |
| Solyc10g078380.1 | Solyc10g078380.2 |
| Solyc10g078400.1 | Solyc10g078400.2 |
| Solyc10g078410.1 | Solyc10g078410.2 |
| Solyc10g078430.1 | Solyc10g078430.2 |
| Solyc10g078440.1 | Solyc10g078440.2 |
| Solyc10g078450.1 | Solyc10g078450.2 |
| Solyc10g078460.1 | Solyc10g078460.2 |
| Solyc10g078470.1 | Solyc10g078470.2 |
| Solyc10g078480.1 | Solyc10g078480.2 |
| Solyc10g078490.1 | Solyc10g078490.2 |
| Solyc10g078520.1 | Solyc10g078520.2 |
| Solyc10g078530.1 | Solyc10g078530.2 |
| Solyc10g078550.3 | Solyc10g078550.4 |
| Solyc10g078590.1 | Solyc10g078590.2 |
| Solyc10g078630.1 | Solyc10g078630.2 |
| Solyc10g078660.1 | Solyc10g078660.2 |
| Solyc10g078720.1 | Solyc10g078720.2 |
| Solyc10g078730.1 | Solyc10g078730.2 |
| Solyc10g078740.1 | Solyc10g078740.2 |
| Solyc10g078750.1 | Solyc10g078750.2 |
| Solyc10g078760.1 | Solyc10g078760.2 |
| Solyc10g078770.1 | Solyc10g078770.2 |
| Solyc10g078780.1 | Solyc10g078780.2 |
| Solyc10g078820.1 | Solyc10g078820.2 |
| Solyc10g078830.1 | Solyc10g078830.2 |
| Solyc10g078860.1 | Solyc10g078860.2 |
| Solyc10g078880.1 | Solyc10g078880.2 |
| Solyc10g078890.1 | Solyc10g078890.2 |
| Solyc10g078930.1 | Solyc10g078930.2 |
| Solyc10g078940.1 | Solyc10g078940.2 |
| Solyc10g078950.1 | Solyc10g078950.2 |
| Solyc10g078960.1 | Solyc10g078960.2 |
| Solyc10g079030.1 | Solyc10g079030.2 |
| Solyc10g079050.1 | Solyc10g079050.2 |
| Solyc10g079060.1 | Solyc10g079060.2 |
| Solyc10g079070.1 | Solyc10g079070.2 |
| Solyc10g079080.1 | Solyc10g079080.2 |
| Solyc10g079110.1 | Solyc10g079110.2 |
| Solyc10g079130.1 | Solyc10g079130.2 |
| Solyc10g079140.1 | Solyc10g079140.2 |
| Solyc10g079170.1 | Solyc10g079170.2 |

| | |
|---|---|
| Solyc10g079200.1 | Solyc10g079200.2 |
| Solyc10g079210.1 | Solyc10g079210.2 |
| Solyc10g079240.1 | Solyc10g079240.2 |
| Solyc10g079250.1 | Solyc10g079250.2 |
| Solyc10g079260.1 | Solyc10g079260.2 |
| Solyc10g079280.1 | Solyc10g079280.2 |
| Solyc10g079290.1 | Solyc10g079290.2 |
| Solyc10g079300.1 | Solyc10g079300.2 |
| Solyc10g079310.1 | Solyc10g079310.2 |
| Solyc10g079330.1 | Solyc10g079330.2 |
| Solyc10g079340.1 | Solyc10g079340.2 |
| Solyc10g079380.1 | Solyc10g079380.2 |
| Solyc10g079410.1 | Solyc10g079410.2 |
| Solyc10g079430.1 | Solyc10g079430.2 |
| Solyc10g079440.1 | Solyc10g079440.2 |
| Solyc10g079450.1 | Solyc10g079450.2 |
| Solyc10g079470.2 | Solyc10g079470.3 |
| Solyc10g079490.1 | Solyc10g079490.2 |
| Solyc10g079500.1 | Solyc10g079500.2 |
| Solyc10g079530.1 | Solyc10g079530.2 |
| Solyc10g079580.1 | Solyc10g079580.2 |
| Solyc10g079600.1 | Solyc10g079600.2 |
| Solyc10g079610.1 | Solyc10g079610.2 |
| Solyc10g079620.1 | Solyc10g079620.2 |
| Solyc10g079640.1 | Solyc10g079640.2 |
| Solyc10g079650.1 | Solyc10g079650.2 |
| Solyc10g079670.1 | Solyc10g079670.2 |
| Solyc10g079690.1 | Solyc10g079690.2 |
| Solyc10g079700.1 | Solyc10g079700.2 |
| Solyc10g079720.1 | Solyc10g079720.2 |
| Solyc10g079740.1 | Solyc10g079740.2 |
| Solyc10g079750.1 | Solyc10g079750.2 |
| Solyc10g079820.1 | Solyc10g079820.2 |
| Solyc10g079840.1 | Solyc10g079840.2 |
| Solyc10g079850.1 | Solyc10g079850.2 |
| Solyc10g079860.1 | Solyc10g079860.2 |
| Solyc10g079880.1 | Solyc10g079880.2 |
| Solyc10g079890.1 | Solyc10g079890.2 |
| Solyc10g079910.1 | Solyc10g079910.2 |
| Solyc10g079920.1 | Solyc10g079920.2 |
| Solyc10g079950.1 | Solyc10g079950.2 |
| Solyc10g079970.1 | Solyc10g079970.2 |
| Solyc10g080010.1 | Solyc10g080010.2 |

| | |
|---|---|
| Solyc10g080030.1 | Solyc10g080030.2 |
| Solyc10g080140.1 | Solyc10g080140.2 |
| Solyc10g080150.1 | Solyc10g080150.2 |
| Solyc10g080160.1 | Solyc10g080160.2 |
| Solyc10g080200.1 | Solyc10g080200.2 |
| Solyc10g080210.1 | Solyc10g080210.2 |
| Solyc10g080220.1 | Solyc10g080220.2 |
| Solyc10g080230.1 | Solyc10g080230.2 |
| Solyc10g080330.1 | Solyc10g080330.2 |
| Solyc10g080390.1 | Solyc10g080390.2 |
| Solyc10g080393.1 | Solyc10g150141.1 |
| Solyc10g080397.1 | Solyc10g150142.1 |
| Solyc10g080440.1 | Solyc10g080440.2 |
| Solyc10g080460.1 | Solyc10g080460.2 |
| Solyc10g080480.1 | Solyc10g080480.2 |
| Solyc10g080490.1 | Solyc10g080490.2 |
| Solyc10g080500.1 | Solyc10g080500.2 |
| Solyc10g080530.1 | Solyc10g080530.2 |
| Solyc10g080540.1 | Solyc10g080540.2 |
| Solyc10g080550.1 | Solyc10g080550.2 |
| Solyc10g080560.1 | Solyc10g080560.2 |
| Solyc10g080570.1 | Solyc10g080570.2 |
| Solyc10g080590.1 | Solyc10g080590.2 |
| Solyc10g080630.1 | Solyc10g080630.2 |
| Solyc10g080670.1 | Solyc10g080670.2 |
| Solyc10g080690.1 | Solyc10g080690.2 |
| Solyc10g080700.1 | Solyc10g080700.2 |
| Solyc10g080710.1 | Solyc10g080710.2 |
| Solyc10g080720.1 | Solyc10g080720.2 |
| Solyc10g080740.1 | Solyc10g080740.2 |
| Solyc10g080760.1 | Solyc10g080760.2 |
| Solyc10g080780.1 | Solyc10g080780.2 |
| Solyc10g080810.1 | Solyc10g080810.2 |
| Solyc10g080815.1 | Solyc10g150143.1 |
| Solyc10g080855.1 | Solyc10g150144.1 |
| Solyc10g080870.2 | Solyc10g080870.3 |
| Solyc10g080880.1 | Solyc10g080880.2 |
| Solyc10g080890.1 | Solyc10g080890.2 |
| Solyc10g080920.1 | Solyc10g080920.2 |
| Solyc10g080930.1 | Solyc10g080930.2 |
| Solyc10g080950.1 | Solyc10g080950.2 |
| Solyc10g080970.1 | Solyc10g080970.2 |
| Solyc10g080980.1 | Solyc10g080980.2 |

| | |
|---|---|
| Solyc10g081020.1 | Solyc10g081020.2 |
| Solyc10g081060.1 | Solyc10g081060.2 |
| Solyc10g081130.1 | Solyc10g081130.2 |
| Solyc10g081160.1 | Solyc10g081160.2 |
| Solyc10g081170.1 | Solyc10g081170.2 |
| Solyc10g081180.1 | Solyc10g081180.2 |
| Solyc10g081190.1 | Solyc10g081190.2 |
| Solyc10g081260.1 | Solyc10g081260.2 |
| Solyc10g081270.1 | Solyc10g081270.2 |
| Solyc10g081280.1 | Solyc10g081280.2 |
| Solyc10g081290.1 | Solyc10g081290.2 |
| Solyc10g081310.1 | Solyc10g081310.2 |
| Solyc10g081350.1 | Solyc10g081350.2 |
| Solyc10g081360.1 | Solyc10g081360.2 |
| Solyc10g081380.1 | Solyc10g081380.2 |
| Solyc10g081410.1 | Solyc10g081410.2 |
| Solyc10g081420.1 | Solyc10g081420.2 |
| Solyc10g081430.1 | Solyc10g081430.2 |
| Solyc10g081440.1 | Solyc10g081440.2 |
| Solyc10g081450.1 | Solyc10g081450.2 |
| Solyc10g081460.1 | Solyc10g081460.2 |
| Solyc10g081470.1 | Solyc10g081470.2 |
| Solyc10g081490.1 | Solyc10g081490.2 |
| Solyc10g081510.1 | Solyc10g081510.2 |
| Solyc10g081530.1 | Solyc10g081530.2 |
| Solyc10g081540.1 | Solyc10g081540.2 |
| Solyc10g081550.1 | Solyc10g081550.2 |
| Solyc10g081590.1 | Solyc10g081590.2 |
| Solyc10g081600.1 | Solyc10g081600.2 |
| Solyc10g081610.1 | Solyc10g081610.2 |
| Solyc10g081620.1 | Solyc10g081620.2 |
| Solyc10g081630.1 | Solyc10g081630.2 |
| Solyc10g081650.1 | Solyc10g081650.2 |
| Solyc10g081660.1 | Solyc10g081660.2 |
| Solyc10g081710.1 | Solyc10g081710.2 |
| Solyc10g081720.1 | Solyc10g081720.2 |
| Solyc10g081730.1 | Solyc10g081730.2 |
| Solyc10g081800.1 | Solyc10g081800.2 |
| Solyc10g081890.1 | Solyc10g081890.2 |
| Solyc10g081910.1 | Solyc10g081910.2 |
| Solyc10g081920.1 | Solyc10g081920.2 |
| Solyc10g081940.1 | Solyc10g081940.2 |
| Solyc10g081955.1 | Solyc10g150145.1 |

| | |
|---|---|
| Solyc10g081970.1 | Solyc10g081970.2 |
| Solyc10g081980.1 | Solyc10g081980.2 |
| Solyc10g082020.1 | Solyc10g082020.2 |
| Solyc10g082030.1 | Solyc10g082030.2 |
| Solyc10g082040.1 | Solyc10g082040.2 |
| Solyc10g083160.1 | Solyc10g083160.2 |
| Solyc10g083165.1 | Solyc10g150146.1 |
| Solyc10g083170.1 | Solyc10g083170.2 |
| Solyc10g083180.1 | Solyc10g083180.2 |
| Solyc10g083210.1 | Solyc10g083210.2 |
| Solyc10g083240.1 | Solyc10g083240.2 |
| Solyc10g083250.1 | Solyc10g083250.2 |
| Solyc10g083270.1 | Solyc10g083270.3 |
| Solyc10g083290.3 | Solyc10g083290.4 |
| Solyc10g083310.1 | Solyc10g083310.2 |
| Solyc10g083350.1 | Solyc10g083350.2 |
| Solyc10g083360.1 | Solyc10g083360.2 |
| Solyc10g083380.1 | Solyc10g083380.2 |
| Solyc10g083390.1 | Solyc10g083390.2 |
| Solyc10g083420.1 | Solyc10g083420.2 |
| Solyc10g083430.1 | Solyc10g083430.2 |
| Solyc10g083440.1 | Solyc10g083440.2 |
| Solyc10g083480.1 | Solyc10g083480.2 |
| Solyc10g083500.1 | Solyc10g083500.2 |
| Solyc10g083510.1 | Solyc10g083510.2 |
| Solyc10g083520.1 | Solyc10g083520.2 |
| Solyc10g083590.1 | Solyc10g083590.2 |
| Solyc10g083600.1 | Solyc10g083600.2 |
| Solyc10g083610.1 | Solyc10g083610.2 |
| Solyc10g083620.1 | Solyc10g083620.2 |
| Solyc10g083630.1 | Solyc10g083630.2 |
| Solyc10g083640.1 | Solyc10g083640.2 |
| Solyc10g083690.2 | Solyc10g083690.3 |
| Solyc10g083710.1 | Solyc10g083710.2 |
| Solyc10g083730.1 | Solyc10g083730.2 |
| Solyc10g083800.1 | Solyc10g083800.2 |
| Solyc10g083870.1 | Solyc10g083870.2 |
| Solyc10g083880.1 | Solyc10g083880.2 |
| Solyc10g083890.1 | Solyc10g083890.2 |
| Solyc10g083900.1 | Solyc10g083900.2 |
| Solyc10g083920.1 | Solyc10g083920.2 |
| Solyc10g083930.1 | Solyc10g083930.2 |
| Solyc10g083960.1 | Solyc10g083960.2 |

| | |
|---|---|
| Solyc10g084023.1 | Solyc10g150147.1 |
| Solyc10g084027.1 | Solyc10g150148.1 |
| Solyc10g084040.1 | Solyc10g084040.2 |
| Solyc10g084050.1 | Solyc10g084050.2 |
| Solyc10g084100.1 | Solyc10g084100.2 |
| Solyc10g084110.1 | Solyc10g084110.2 |
| Solyc10g084120.1 | Solyc10g084120.2 |
| Solyc10g084140.1 | Solyc10g084140.2 |
| Solyc10g084150.1 | Solyc10g084150.2 |
| Solyc10g084160.1 | Solyc10g084160.2 |
| Solyc10g084180.1 | Solyc10g084180.2 |
| Solyc10g084200.1 | Solyc10g084200.2 |
| Solyc10g084210.1 | Solyc10g084210.2 |
| Solyc10g084270.1 | Solyc10g084270.2 |
| Solyc10g084280.1 | Solyc10g084280.2 |
| Solyc10g084290.1 | Solyc10g084290.2 |
| Solyc10g084310.1 | Solyc10g084310.2 |
| Solyc10g084340.1 | Solyc10g084340.2 |
| Solyc10g084350.1 | Solyc10g084350.2 |
| Solyc10g084390.1 | Solyc10g084390.2 |
| Solyc10g084400.1 | Solyc10g084400.2 |
| Solyc10g084410.1 | Solyc10g084410.2 |
| Solyc10g084430.1 | Solyc10g084430.2 |
| Solyc10g084600.1 | Solyc10g084600.2 |
| Solyc10g084610.1 | Solyc10g084610.3 |
| Solyc10g084630.1 | Solyc10g084630.2 |
| Solyc10g084660.1 | Solyc10g084660.2 |
| Solyc10g084690.1 | Solyc10g084690.2 |
| Solyc10g084700.1 | Solyc10g084700.2 |
| Solyc10g084710.1 | Solyc10g084710.2 |
| Solyc10g084730.1 | Solyc10g084730.2 |
| Solyc10g084800.1 | Solyc10g084800.2 |
| Solyc10g084820.1 | Solyc10g084820.2 |
| Solyc10g084830.1 | Solyc10g084830.2 |
| Solyc10g084840.1 | Solyc10g084840.2 |
| Solyc10g084850.1 | Solyc10g084850.2 |
| Solyc10g084880.2 | Solyc10g084880.3 |
| Solyc10g084920.1 | Solyc10g084920.2 |
| Solyc10g084930.1 | Solyc10g084930.2 |
| Solyc10g084960.1 | Solyc10g084960.2 |
| Solyc10g084970.1 | Solyc10g084970.2 |
| Solyc10g084990.1 | Solyc10g084990.2 |
| Solyc10g085000.1 | Solyc10g085000.2 |

| | |
|---|---|
| Solyc10g085100.1 | Solyc10g085100.2 |
| Solyc10g085130.1 | Solyc10g085130.2 |
| Solyc10g085140.1 | Solyc10g085140.2 |
| Solyc10g085150.1 | Solyc10g085150.2 |
| Solyc10g085160.1 | Solyc10g085160.2 |
| Solyc10g085180.1 | Solyc10g085180.2 |
| Solyc10g085190.1 | Solyc10g085190.2 |
| Solyc10g085200.1 | Solyc10g085200.2 |
| Solyc10g085210.1 | Solyc10g085210.2 |
| Solyc10g085230.1 | Solyc10g085230.2 |
| Solyc10g085250.1 | Solyc10g085250.2 |
| Solyc10g085320.1 | Solyc10g085320.2 |
| Solyc10g085330.1 | Solyc10g085330.2 |
| Solyc10g085350.1 | Solyc10g085350.2 |
| Solyc10g085380.1 | Solyc10g085380.2 |
| Solyc10g085480.1 | Solyc10g085480.2 |
| Solyc10g085500.1 | Solyc10g085500.2 |
| Solyc10g085580.2 | Solyc10g085580.3 |
| Solyc10g085600.1 | Solyc10g085600.2 |
| Solyc10g085610.1 | Solyc10g085610.2 |
| Solyc10g085620.1 | Solyc10g085620.2 |
| Solyc10g085650.1 | Solyc10g085650.2 |
| Solyc10g085665.1 | Solyc10g150150.1 |
| Solyc10g085670.1 | Solyc10g085670.2 |
| Solyc10g085680.1 | Solyc10g085680.2 |
| Solyc10g085720.1 | Solyc10g085720.2 |
| Solyc10g085740.1 | Solyc10g085740.2 |
| Solyc10g085750.1 | Solyc10g085750.2 |
| Solyc10g085790.1 | Solyc10g085790.2 |
| Solyc10g085800.1 | Solyc10g085800.2 |
| Solyc10g085810.1 | Solyc10g085810.2 |
| Solyc10g085820.1 | Solyc10g085820.2 |
| Solyc10g085830.1 | Solyc10g085830.2 |
| Solyc10g085910.1 | Solyc10g085910.2 |
| Solyc10g085950.1 | Solyc10g085950.2 |
| Solyc10g085960.1 | Solyc10g085960.2 |
| Solyc10g085980.1 | Solyc10g085980.2 |
| Solyc10g086010.1 | Solyc10g086010.2 |
| Solyc10g086030.1 | Solyc10g086030.2 |
| Solyc10g086040.1 | Solyc10g086040.2 |
| Solyc10g086060.1 | Solyc10g086060.2 |
| Solyc10g086070.1 | Solyc10g086070.2 |
| Solyc10g086080.1 | Solyc10g086080.2 |

| | |
|---|---|
| Solyc10g086090.1 | Solyc10g086090.2 |
| Solyc10g086100.1 | Solyc10g086100.2 |
| Solyc10g086120.1 | Solyc10g086120.2 |
| Solyc10g086125.1 | Solyc10g150151.1 |
| Solyc10g086137.1 | Solyc10g086137.2 |
| Solyc10g086150.1 | Solyc10g086150.2 |
| Solyc10g086180.1 | Solyc10g086180.2 |
| Solyc10g086210.1 | Solyc10g086210.2 |
| Solyc10g086220.1 | Solyc10g086220.2 |
| Solyc10g086240.1 | Solyc10g086240.2 |
| Solyc10g086250.1 | Solyc10g086250.2 |
| Solyc10g086270.1 | Solyc10g086270.2 |
| Solyc10g086280.1 | Solyc10g086280.2 |
| Solyc10g086320.1 | Solyc10g086320.2 |
| Solyc10g086330.1 | Solyc10g086330.2 |
| Solyc10g086350.1 | Solyc10g086350.2 |
| Solyc10g086390.1 | Solyc10g086390.2 |
| Solyc10g086400.1 | Solyc10g086400.2 |
| Solyc10g086410.2 | Solyc10g086410.3 |
| Solyc10g086440.1 | Solyc10g086440.2 |
| Solyc10g086460.1 | Solyc10g086460.2 |
| Solyc10g086490.1 | Solyc10g086490.2 |
| Solyc10g086510.1 | Solyc10g086510.2 |
| Solyc10g086520.1 | Solyc10g086520.2 |
| Solyc10g086550.1 | Solyc10g086550.2 |
| Solyc10g086565.1 | Solyc10g086565.2 |
| Solyc10g086580.1 | Solyc10g086580.2 |
| Solyc10g086620.1 | Solyc10g086620.2 |
| Solyc10g086630.1 | Solyc10g086630.2 |
| Solyc10g086660.1 | Solyc10g086660.2 |
| Solyc10g086670.1 | Solyc10g086670.2 |
| Solyc10g086720.1 | Solyc10g086720.2 |
| Solyc10g086730.1 | Solyc10g086730.2 |
| Solyc10g086760.1 | Solyc10g086760.2 |
| Solyc10g086780.1 | Solyc10g086780.2 |
| Solyc10g005020.2 | Solyc10g005020.3 |
| Solyc10g005050.2 | Solyc10g005050.3 |
| Solyc10g005080.2 | Solyc10g005080.3 |
| Solyc10g005090.2 | Solyc10g005090.3 |
| Solyc10g005100.2 | Solyc10g005100.3 |
| Solyc10g005110.2 | Solyc10g005110.3 |
| Solyc10g005120.1 | Solyc10g005120.2 |
| Solyc11g005050.1 | Solyc11g005050.2 |

| | |
|---|---|
| Solyc11g005060.1 | Solyc11g005060.2 |
| Solyc11g005070.1 | Solyc11g005070.2 |
| Solyc11g005090.1 | Solyc11g005090.2 |
| Solyc11g005130.1 | Solyc11g005130.2 |
| Solyc11g005170.1 | Solyc11g005170.2 |
| Solyc11g005200.1 | Solyc11g005200.2 |
| Solyc11g005220.1 | Solyc11g005220.2 |
| Solyc11g005260.1 | Solyc11g005260.2 |
| Solyc11g005300.1 | Solyc11g005300.2 |
| Solyc11g005350.1 | Solyc11g005350.2 |
| Solyc11g005360.1 | Solyc11g005360.2 |
| Solyc11g005370.1 | Solyc11g005370.2 |
| Solyc11g005380.1 | Solyc11g005380.2 |
| Solyc11g005400.1 | Solyc11g005400.2 |
| Solyc11g005410.1 | Solyc11g005410.2 |
| Solyc11g005430.1 | Solyc11g005430.2 |
| Solyc11g005440.1 | Solyc11g005440.2 |
| Solyc11g005460.1 | Solyc11g005460.2 |
| Solyc11g005480.1 | Solyc11g005480.2 |
| Solyc11g005550.1 | Solyc11g005550.2 |
| Solyc11g005600.1 | Solyc11g005600.2 |
| Solyc11g005660.1 | Solyc11g005660.2 |
| Solyc11g005670.1 | Solyc11g005670.2 |
| Solyc11g005680.1 | Solyc11g005680.2 |
| Solyc11g005710.1 | Solyc11g005710.2 |
| Solyc11g005740.1 | Solyc11g005740.2 |
| Solyc11g005750.1 | Solyc11g005750.2 |
| Solyc11g005770.1 | Solyc11g005770.2 |
| Solyc11g005800.1 | Solyc11g005800.2 |
| Solyc11g005830.1 | Solyc11g005830.2 |
| Solyc11g005840.1 | Solyc11g005840.2 |
| Solyc11g005860.1 | Solyc11g005860.2 |
| Solyc11g005870.1 | Solyc11g005870.2 |
| Solyc11g005880.1 | Solyc11g005880.2 |
| Solyc11g005900.1 | Solyc11g005900.2 |
| Solyc11g005910.1 | Solyc11g005910.2 |
| Solyc11g005950.1 | Solyc11g005950.2 |
| Solyc11g005990.1 | Solyc11g005990.2 |
| Solyc11g006020.1 | Solyc11g006020.2 |
| Solyc11g006040.1 | Solyc11g006040.2 |
| Solyc11g006060.1 | Solyc11g006060.2 |
| Solyc11g006130.1 | Solyc11g006130.2 |
| Solyc11g006180.1 | Solyc11g006180.2 |

| | |
|---|---|
| Solyc11g006200.1 | Solyc11g006200.2 |
| Solyc11g006220.1 | Solyc11g006220.2 |
| Solyc11g006240.1 | Solyc11g006240.2 |
| Solyc11g006245.1 | Solyc11g150100.1 |
| Solyc11g006250.1 | Solyc11g006250.2 |
| Solyc11g006260.1 | Solyc11g006260.2 |
| Solyc11g006270.1 | Solyc11g006270.2 |
| Solyc11g006290.1 | Solyc11g006290.2 |
| Solyc11g006300.1 | Solyc11g006300.2 |
| Solyc11g006310.1 | Solyc11g006310.2 |
| Solyc11g006320.1 | Solyc11g006320.2 |
| Solyc11g006340.1 | Solyc11g006340.2 |
| Solyc11g006370.1 | Solyc11g006370.2 |
| Solyc11g006380.1 | Solyc11g006380.2 |
| Solyc11g006390.1 | Solyc11g006390.2 |
| Solyc11g006400.1 | Solyc11g006400.2 |
| Solyc11g006410.1 | Solyc11g006410.2 |
| Solyc11g006420.1 | Solyc11g006420.2 |
| Solyc11g006470.1 | Solyc11g006470.2 |
| Solyc11g006490.1 | Solyc11g006490.2 |
| Solyc11g006500.1 | Solyc11g006500.2 |
| Solyc11g006520.1 | Solyc11g006520.2 |
| Solyc11g006590.1 | Solyc11g006590.2 |
| Solyc11g006710.1 | Solyc11g006710.2 |
| Solyc11g006720.1 | Solyc11g006720.2 |
| Solyc11g006780.1 | Solyc11g006780.2 |
| Solyc11g006800.1 | Solyc11g006800.2 |
| Solyc11g006820.1 | Solyc11g006820.2 |
| Solyc11g006830.1 | Solyc11g006830.2 |
| Solyc11g006840.1 | Solyc11g006840.2 |
| Solyc11g006860.1 | Solyc11g006860.2 |
| Solyc11g006890.1 | Solyc11g006890.2 |
| Solyc11g006920.1 | Solyc11g006920.2 |
| Solyc11g006970.1 | Solyc11g006970.2 |
| Solyc11g006990.1 | Solyc11g006990.2 |
| Solyc11g007000.1 | Solyc11g007000.2 |
| Solyc11g007010.1 | Solyc11g007010.2 |
| Solyc11g007020.1 | Solyc11g007020.2 |
| Solyc11g007040.1 | Solyc11g007040.2 |
| Solyc11g007070.1 | Solyc11g007070.2 |
| Solyc11g007110.1 | Solyc11g007110.2 |
| Solyc11g007120.1 | Solyc11g007120.2 |
| Solyc11g007130.1 | Solyc11g007130.2 |

| | |
|---|---|
| Solyc11g007140.1 | Solyc11g007140.2 |
| Solyc11g007150.1 | Solyc11g007150.2 |
| Solyc11g007160.1 | Solyc11g007160.2 |
| Solyc11g007170.1 | Solyc11g007170.2 |
| Solyc11g007190.1 | Solyc11g007190.2 |
| Solyc11g007220.1 | Solyc11g007220.2 |
| Solyc11g007250.1 | Solyc11g007250.2 |
| Solyc11g007300.1 | Solyc11g007300.2 |
| Solyc11g007320.1 | Solyc11g007320.2 |
| Solyc11g007330.1 | Solyc11g007330.2 |
| Solyc11g007340.1 | Solyc11g007340.2 |
| Solyc11g007500.1 | Solyc11g007500.2 |
| Solyc11g007530.1 | Solyc11g007530.2 |
| Solyc11g007540.1 | Solyc11g007540.2 |
| Solyc11g007560.1 | Solyc11g007560.2 |
| Solyc11g007570.1 | Solyc11g007570.2 |
| Solyc11g007600.1 | Solyc11g007600.2 |
| Solyc11g007610.1 | Solyc11g007610.2 |
| Solyc11g007640.1 | Solyc11g007640.2 |
| Solyc11g007700.1 | Solyc11g007700.2 |
| Solyc11g007710.1 | Solyc11g007710.2 |
| Solyc11g007720.1 | Solyc11g007720.2 |
| Solyc11g007730.1 | Solyc11g007730.2 |
| Solyc11g007740.1 | Solyc11g007740.2 |
| Solyc11g007760.1 | Solyc11g007760.2 |
| Solyc11g007770.1 | Solyc11g007770.2 |
| Solyc11g007780.1 | Solyc11g007780.2 |
| Solyc11g007830.1 | Solyc11g007830.2 |
| Solyc11g007850.1 | Solyc11g007850.2 |
| Solyc11g007890.1 | Solyc11g007890.2 |
| Solyc11g007910.1 | Solyc11g007910.2 |
| Solyc11g007940.1 | Solyc11g007940.2 |
| Solyc11g007960.1 | Solyc11g007960.2 |
| Solyc11g007970.1 | Solyc11g007970.2 |
| Solyc11g007980.1 | Solyc11g007980.2 |
| Solyc11g007990.1 | Solyc11g007990.2 |
| Solyc11g008000.1 | Solyc11g008000.2 |
| Solyc11g008010.1 | Solyc11g008010.2 |
| Solyc11g008020.1 | Solyc11g008020.2 |
| Solyc11g008090.1 | Solyc11g008090.2 |
| Solyc11g008200.1 | Solyc11g008200.2 |
| Solyc11g008250.1 | Solyc11g008250.2 |
| Solyc11g008260.1 | Solyc11g008260.2 |

| | |
|---|---|
| Solyc11g008270.1 | Solyc11g008270.2 |
| Solyc11g008350.1 | Solyc11g008350.2 |
| Solyc11g008390.1 | Solyc11g008390.2 |
| Solyc11g008400.1 | Solyc11g008400.2 |
| Solyc11g008430.1 | Solyc11g008430.2 |
| Solyc11g008450.1 | Solyc11g008450.2 |
| Solyc11g008470.1 | Solyc11g008470.2 |
| Solyc11g008480.1 | Solyc11g008480.2 |
| Solyc11g008500.1 | Solyc11g008500.2 |
| Solyc11g008560.1 | Solyc11g008560.2 |
| Solyc11g008570.1 | Solyc11g008570.2 |
| Solyc11g008590.1 | Solyc11g008590.2 |
| Solyc11g008620.1 | Solyc11g008620.2 |
| Solyc11g008670.1 | Solyc11g008670.2 |
| Solyc11g008680.1 | Solyc11g008680.2 |
| Solyc11g008700.1 | Solyc11g008700.2 |
| Solyc11g008710.1 | Solyc11g008710.2 |
| Solyc11g008730.1 | Solyc11g008730.2 |
| Solyc11g008740.1 | Solyc11g008740.2 |
| Solyc11g008770.1 | Solyc11g008770.2 |
| Solyc11g008810.1 | Solyc11g008810.2 |
| Solyc11g008820.1 | Solyc11g008820.2 |
| Solyc11g008830.1 | Solyc11g008830.2 |
| Solyc11g008850.1 | Solyc11g008850.2 |
| Solyc11g008870.1 | Solyc11g008870.2 |
| Solyc11g008880.1 | Solyc11g008880.2 |
| Solyc11g008890.1 | Solyc11g008890.2 |
| Solyc11g008900.1 | Solyc11g008900.2 |
| Solyc11g008990.1 | Solyc11g008990.2 |
| Solyc11g009000.1 | Solyc11g009000.2 |
| Solyc11g009010.1 | Solyc11g009010.2 |
| Solyc11g009020.1 | Solyc11g009020.2 |
| Solyc11g009040.2 | Solyc11g009040.3 |
| Solyc11g009070.1 | Solyc11g009070.2 |
| Solyc11g009080.1 | Solyc11g009080.2 |
| Solyc11g009090.1 | Solyc11g009090.2 |
| Solyc11g009100.1 | Solyc11g009100.2 |
| Solyc11g010100.1 | Solyc11g010100.2 |
| Solyc11g010120.1 | Solyc11g010120.2 |
| Solyc11g010170.1 | Solyc11g010170.2 |
| Solyc11g010180.1 | Solyc11g010180.2 |
| Solyc11g010220.1 | Solyc11g010220.2 |
| Solyc11g010230.1 | Solyc11g010230.2 |

| | |
|---|---|
| Solyc11g010270.1 | Solyc11g010270.2 |
| Solyc11g010290.1 | Solyc11g010290.2 |
| Solyc11g010300.1 | Solyc11g010300.2 |
| Solyc11g010310.1 | Solyc11g010310.2 |
| Solyc11g010340.1 | Solyc11g010340.2 |
| Solyc11g010350.1 | Solyc11g010350.2 |
| Solyc11g010380.1 | Solyc11g010380.2 |
| Solyc11g010410.1 | Solyc11g010410.2 |
| Solyc11g010430.1 | Solyc11g010430.2 |
| Solyc11g010450.1 | Solyc11g010450.2 |
| Solyc11g010470.1 | Solyc11g010470.2 |
| Solyc11g010480.1 | Solyc11g010480.2 |
| Solyc11g010510.1 | Solyc11g010510.2 |
| Solyc11g010530.1 | Solyc11g010530.2 |
| Solyc11g010550.1 | Solyc11g010550.2 |
| Solyc11g010570.1 | Solyc11g010570.2 |
| Solyc11g010600.1 | Solyc11g010600.2 |
| Solyc11g010610.1 | Solyc11g010610.2 |
| Solyc11g010660.1 | Solyc11g010660.2 |
| Solyc11g010710.1 | Solyc11g010710.2 |
| Solyc11g010800.1 | Solyc11g010800.2 |
| Solyc11g010840.1 | Solyc11g010840.2 |
| Solyc11g010850.1 | Solyc11g010850.2 |
| Solyc11g010870.1 | Solyc11g010870.2 |
| Solyc11g010880.1 | Solyc11g010880.2 |
| Solyc11g010900.1 | Solyc11g010900.2 |
| Solyc11g010930.1 | Solyc11g010930.2 |
| Solyc11g010940.1 | Solyc11g010940.2 |
| Solyc11g010950.1 | Solyc11g010950.2 |
| Solyc11g010960.1 | Solyc11g010960.2 |
| Solyc11g010980.1 | Solyc11g010980.2 |
| Solyc11g011000.1 | Solyc11g011000.2 |
| Solyc11g011020.1 | Solyc11g011020.2 |
| Solyc11g011030.1 | Solyc11g011030.2 |
| Solyc11g011040.1 | Solyc11g011040.2 |
| Solyc11g011050.1 | Solyc11g011050.2 |
| Solyc11g011120.1 | Solyc11g011120.2 |
| Solyc11g011140.1 | Solyc11g011140.2 |
| Solyc11g011150.1 | Solyc11g011150.2 |
| Solyc11g011170.1 | Solyc11g011170.2 |
| Solyc11g011200.1 | Solyc11g011200.2 |
| Solyc11g011210.1 | Solyc11g011210.2 |
| Solyc11g011280.1 | Solyc11g011280.3 |

| | |
|---|---|
| Solyc11g011300.1 | Solyc11g011300.2 |
| Solyc11g011330.1 | Solyc11g011330.2 |
| Solyc11g011340.1 | Solyc11g011340.2 |
| Solyc11g011360.1 | Solyc11g011360.2 |
| Solyc11g011370.1 | Solyc11g011370.2 |
| Solyc11g011380.1 | Solyc11g011380.2 |
| Solyc11g011400.1 | Solyc11g011400.2 |
| Solyc11g011450.1 | Solyc11g011450.2 |
| Solyc11g011460.1 | Solyc11g011460.2 |
| Solyc11g011470.1 | Solyc11g011470.2 |
| Solyc11g011480.1 | Solyc11g011480.2 |
| Solyc11g011490.1 | Solyc11g011490.2 |
| Solyc11g011500.1 | Solyc11g011500.2 |
| Solyc11g011530.1 | Solyc11g011530.2 |
| Solyc11g011580.1 | Solyc11g011580.2 |
| Solyc11g011590.1 | Solyc11g011590.2 |
| Solyc11g011610.1 | Solyc11g011610.2 |
| Solyc11g011840.1 | Solyc11g011840.2 |
| Solyc11g011850.1 | Solyc11g011850.2 |
| Solyc11g011880.1 | Solyc11g011880.2 |
| Solyc11g011890.1 | Solyc11g011890.2 |
| Solyc11g011910.1 | Solyc11g011910.2 |
| Solyc11g011920.1 | Solyc11g011920.2 |
| Solyc11g011960.1 | Solyc11g011960.2 |
| Solyc11g011970.1 | Solyc11g011970.2 |
| Solyc11g012000.1 | Solyc11g012000.2 |
| Solyc11g012010.1 | Solyc11g012010.2 |
| Solyc11g012020.1 | Solyc11g012020.2 |
| Solyc11g012030.1 | Solyc11g012030.2 |
| Solyc11g012040.1 | Solyc11g012040.2 |
| Solyc11g012080.1 | Solyc11g012080.2 |
| Solyc11g012100.1 | Solyc11g012100.2 |
| Solyc11g012110.1 | Solyc11g012110.2 |
| Solyc11g012120.1 | Solyc11g012120.2 |
| Solyc11g012130.1 | Solyc11g012130.2 |
| Solyc11g012140.1 | Solyc11g012140.2 |
| Solyc11g012160.1 | Solyc11g012160.2 |
| Solyc11g012180.1 | Solyc11g012180.2 |
| Solyc11g012270.1 | Solyc11g012270.2 |
| Solyc11g012290.1 | Solyc11g012290.2 |
| Solyc11g012360.1 | Solyc11g012360.2 |
| Solyc11g012400.1 | Solyc11g012400.2 |
| Solyc11g012410.1 | Solyc11g012410.2 |

| | |
|---|---|
| Solyc11g012420.1 | Solyc11g012420.2 |
| Solyc11g012450.1 | Solyc11g012450.2 |
| Solyc11g012470.1 | Solyc11g012470.2 |
| Solyc11g012490.1 | Solyc11g012490.2 |
| Solyc11g012540.1 | Solyc11g012540.2 |
| Solyc11g012580.1 | Solyc11g012580.2 |
| Solyc11g012620.1 | Solyc11g012620.2 |
| Solyc11g012625.1 | Solyc11g150101.1 |
| Solyc11g012660.1 | Solyc11g012660.2 |
| Solyc11g012700.1 | Solyc11g012700.2 |
| Solyc11g012705.1 | Solyc11g012705.2 |
| Solyc11g012710.1 | Solyc11g012710.2 |
| Solyc11g012720.1 | Solyc11g012720.2 |
| Solyc11g012740.1 | Solyc11g012740.2 |
| Solyc11g012760.1 | Solyc11g012760.2 |
| Solyc11g012770.1 | Solyc11g012770.2 |
| Solyc11g012790.1 | Solyc11g012790.2 |
| Solyc11g012810.1 | Solyc11g012810.2 |
| Solyc11g012820.1 | Solyc11g012820.2 |
| Solyc11g012840.1 | Solyc11g012840.2 |
| Solyc11g012850.1 | Solyc11g012850.2 |
| Solyc11g012870.1 | Solyc11g012870.2 |
| Solyc11g012880.1 | Solyc11g012880.2 |
| Solyc11g012910.1 | Solyc11g012910.2 |
| Solyc11g012930.1 | Solyc11g012930.2 |
| Solyc11g012940.1 | Solyc11g012940.2 |
| Solyc11g012950.1 | Solyc11g012950.2 |
| Solyc11g012970.1 | Solyc11g012970.2 |
| Solyc11g013000.1 | Solyc11g013000.2 |
| Solyc11g013010.1 | Solyc11g013010.2 |
| Solyc11g013030.1 | Solyc11g013030.2 |
| Solyc11g013090.1 | Solyc11g013090.2 |
| Solyc11g013100.1 | Solyc11g013100.2 |
| Solyc11g013120.1 | Solyc11g013120.2 |
| Solyc11g013130.1 | Solyc11g013130.2 |
| Solyc11g013170.1 | Solyc11g013170.2 |
| Solyc11g013180.1 | Solyc11g013180.2 |
| Solyc11g013190.1 | Solyc11g013190.2 |
| Solyc11g013200.1 | Solyc11g013200.2 |
| Solyc11g013220.1 | Solyc11g013220.2 |
| Solyc11g013270.1 | Solyc11g013270.2 |
| Solyc11g013300.1 | Solyc11g013300.2 |
| Solyc11g013310.1 | Solyc11g013310.2 |

| | |
|---|---|
| Solyc11g013320.1 | Solyc11g013320.2 |
| Solyc11g013393.1 | Solyc11g150102.1 |
| Solyc11g013397.1 | Solyc11g150103.1 |
| Solyc11g013410.1 | Solyc11g013410.2 |
| Solyc11g013420.1 | Solyc11g013420.2 |
| Solyc11g013440.1 | Solyc11g013440.2 |
| Solyc11g013460.1 | Solyc11g013460.2 |
| Solyc11g013490.1 | Solyc11g013490.2 |
| Solyc11g013550.1 | Solyc11g013550.2 |
| Solyc11g013600.1 | Solyc11g013600.2 |
| Solyc11g013610.1 | Solyc11g013610.2 |
| Solyc11g013700.1 | Solyc11g013700.2 |
| Solyc11g013770.1 | Solyc11g013770.2 |
| Solyc11g013820.1 | Solyc11g013820.2 |
| Solyc11g013830.1 | Solyc11g013830.2 |
| Solyc11g013870.1 | Solyc11g013870.2 |
| Solyc11g015890.1 | Solyc11g015890.2 |
| Solyc11g016970.1 | Solyc11g016970.2 |
| Solyc11g017000.1 | Solyc11g017000.2 |
| Solyc11g017010.1 | Solyc11g017010.2 |
| Solyc11g017040.1 | Solyc11g017040.2 |
| Solyc11g017060.1 | Solyc11g017060.2 |
| Solyc11g017070.1 | Solyc11g017070.2 |
| Solyc11g017073.1 | Solyc11g150104.1 |
| Solyc11g017077.1 | Solyc11g150105.1 |
| Solyc11g017130.1 | Solyc11g017130.2 |
| Solyc11g017200.1 | Solyc11g017200.2 |
| Solyc11g017230.1 | Solyc11g017230.2 |
| Solyc11g017250.1 | Solyc11g017250.2 |
| Solyc11g017270.1 | Solyc11g017270.2 |
| Solyc11g017280.1 | Solyc11g017280.2 |
| Solyc11g017285.1 | Solyc11g150106.1 |
| Solyc11g017300.1 | Solyc11g017300.2 |
| Solyc11g017373.1 | Solyc11g150107.1 |
| Solyc11g017400.1 | Solyc11g017400.2 |
| Solyc11g017420.1 | Solyc11g017420.2 |
| Solyc11g017450.1 | Solyc11g017450.2 |
| Solyc11g017470.1 | Solyc11g017470.2 |
| Solyc11g018510.1 | Solyc11g018510.2 |
| Solyc11g018560.1 | Solyc11g018560.2 |
| Solyc11g018580.1 | Solyc11g018580.2 |
| Solyc11g018610.1 | Solyc11g018610.2 |
| Solyc11g018660.1 | Solyc11g018660.2 |

| | |
|---|---|
| Solyc11g018690.1 | Solyc11g018690.2 |
| Solyc11g018750.1 | Solyc11g018750.2 |
| Solyc11g018770.2 | Solyc11g018770.3 |
| Solyc11g018853.1 | Solyc11g150108.1 |
| Solyc11g018857.1 | Solyc11g150109.1 |
| Solyc11g019920.1 | Solyc11g019920.2 |
| Solyc11g019930.1 | Solyc11g019930.2 |
| Solyc11g019935.1 | Solyc11g150110.1 |
| Solyc11g019950.1 | Solyc11g019950.2 |
| Solyc11g020040.1 | Solyc11g020040.2 |
| Solyc11g020060.1 | Solyc11g020060.2 |
| Solyc11g020135.1 | Solyc11g150111.1 |
| Solyc11g020160.1 | Solyc11g020160.2 |
| Solyc11g020210.1 | Solyc11g020210.2 |
| Solyc11g020217.1 | Solyc11g150112.1 |
| Solyc11g020280.1 | Solyc11g020280.2 |
| Solyc11g020300.1 | Solyc11g020300.2 |
| Solyc11g020345.1 | Solyc11g150113.1 |
| Solyc11g020550.1 | Solyc11g020550.2 |
| Solyc11g020555.1 | Solyc11g150114.1 |
| Solyc11g020590.1 | Solyc11g020590.2 |
| Solyc11g020610.1 | Solyc11g020610.3 |
| Solyc11g020780.1 | Solyc11g020780.2 |
| Solyc11g020810.1 | Solyc11g020810.2 |
| Solyc11g020820.1 | Solyc11g020820.2 |
| Solyc11g020875.1 | Solyc11g150115.1 |
| Solyc11g020960.1 | Solyc11g020960.2 |
| Solyc11g021060.1 | Solyc11g021060.2 |
| Solyc11g021100.1 | Solyc11g021100.2 |
| Solyc11g021180.1 | Solyc11g021180.2 |
| Solyc11g021190.1 | Solyc11g021190.2 |
| Solyc11g021210.1 | Solyc11g021210.2 |
| Solyc11g021220.1 | Solyc11g021220.2 |
| Solyc11g021280.1 | Solyc11g021280.2 |
| Solyc11g021300.1 | Solyc11g021300.2 |
| Solyc11g022385.1 | Solyc11g150116.1 |
| Solyc11g022390.1 | Solyc11g022390.2 |
| Solyc11g022470.1 | Solyc11g022470.2 |
| Solyc11g022600.1 | Solyc11g022600.2 |
| Solyc11g050710.1 | Solyc11g050710.2 |
| Solyc11g050760.1 | Solyc11g050760.2 |
| Solyc11g050880.1 | Solyc11g050880.2 |
| Solyc11g050890.1 | Solyc11g050890.2 |

| | |
|---|---|
| Solyc11g050900.1 | Solyc11g050900.2 |
| Solyc11g051035.1 | Solyc11g150137.1 |
| Solyc11g051060.1 | Solyc11g051060.2 |
| Solyc11g051105.1 | Solyc11g150138.1 |
| Solyc11g051125.1 | Solyc11g150139.1 |
| Solyc11g051160.1 | Solyc11g051160.2 |
| Solyc11g051170.1 | Solyc11g051170.2 |
| Solyc11g027830.1 | Solyc11g027830.2 |
| Solyc11g027840.1 | Solyc11g027840.2 |
| Solyc11g027860.1 | Solyc11g027860.2 |
| Solyc11g027870.1 | Solyc11g027870.2 |
| Solyc11g027880.1 | Solyc11g027880.2 |
| Solyc11g027887.1 | Solyc11g027887.2 |
| Solyc11g027893.1 | Solyc11g150117.1 |
| Solyc11g027920.1 | Solyc11g027920.2 |
| Solyc11g027930.1 | Solyc11g027930.2 |
| Solyc11g027981.1 | Solyc11g150118.1 |
| Solyc11g027982.1 | Solyc11g150119.1 |
| Solyc11g027984.1 | Solyc11g150120.1 |
| Solyc11g027986.1 | Solyc11g150121.1 |
| Solyc11g027988.1 | Solyc11g150122.1 |
| Solyc11g028010.1 | Solyc11g028010.2 |
| Solyc11g028040.1 | Solyc11g028040.2 |
| Solyc11g028070.1 | Solyc11g028070.2 |
| Solyc11g028100.1 | Solyc11g028100.2 |
| Solyc11g028175.1 | Solyc11g150123.1 |
| Solyc11g028183.1 | Solyc11g150124.1 |
| Solyc11g028187.1 | Solyc11g150125.1 |
| Solyc11g028315.1 | Solyc11g150126.1 |
| Solyc11g028330.1 | Solyc11g028330.2 |
| Solyc11g030380.1 | Solyc11g030380.2 |
| Solyc11g030390.1 | Solyc11g030390.2 |
| Solyc11g030475.1 | Solyc11g150127.1 |
| Solyc11g030600.2 | Solyc11g030600.3 |
| Solyc11g030710.1 | Solyc11g030710.2 |
| Solyc11g030720.1 | Solyc11g030720.2 |
| Solyc11g030730.1 | Solyc11g030730.2 |
| Solyc11g030950.1 | Solyc11g030950.2 |
| Solyc11g030953.1 | Solyc11g150128.1 |
| Solyc11g030957.1 | Solyc11g150129.1 |
| Solyc11g031960.1 | Solyc11g031960.2 |
| Solyc11g032050.1 | Solyc11g032050.2 |
| Solyc11g032060.1 | Solyc11g032060.3 |

| | |
|---|---|
| Solyc11g032100.1 | Solyc11g032100.2 |
| Solyc11g032120.1 | Solyc11g032120.2 |
| Solyc11g032200.1 | Solyc11g032200.2 |
| Solyc11g032220.1 | Solyc11g032220.2 |
| Solyc11g033260.1 | Solyc11g033260.2 |
| Solyc11g033270.1 | Solyc11g033270.2 |
| Solyc11g033280.1 | Solyc11g033280.2 |
| Solyc11g045645.1 | Solyc11g150136.1 |
| Solyc11g045610.1 | Solyc11g045610.2 |
| Solyc11g045520.1 | Solyc11g045520.2 |
| Solyc11g045350.1 | Solyc11g045350.2 |
| Solyc11g045330.1 | Solyc11g045330.2 |
| Solyc11g045230.1 | Solyc11g045230.2 |
| Solyc11g045220.1 | Solyc11g045220.2 |
| Solyc11g045130.1 | Solyc11g045130.2 |
| Solyc11g045040.1 | Solyc11g045040.2 |
| Solyc11g045030.1 | Solyc11g045030.2 |
| Solyc11g044910.1 | Solyc11g044910.2 |
| Solyc11g044900.1 | Solyc11g044900.2 |
| Solyc11g044870.1 | Solyc11g044870.2 |
| Solyc11g044840.1 | Solyc11g044840.2 |
| Solyc11g044810.1 | Solyc11g044810.2 |
| Solyc11g044815.1 | Solyc11g150135.1 |
| Solyc11g044740.1 | Solyc11g044740.2 |
| Solyc11g044600.1 | Solyc11g044600.2 |
| Solyc11g044540.1 | Solyc11g044540.2 |
| Solyc11g044530.1 | Solyc11g044530.2 |
| Solyc11g044480.1 | Solyc11g044480.2 |
| Solyc11g044365.1 | Solyc11g150134.1 |
| Solyc11g044320.1 | Solyc11g044320.2 |
| Solyc11g044310.1 | Solyc11g044310.2 |
| Solyc11g043160.1 | Solyc11g043160.2 |
| Solyc11g043150.1 | Solyc11g043150.2 |
| Solyc11g043130.1 | Solyc11g043130.2 |
| Solyc11g043120.1 | Solyc11g043120.2 |
| Solyc11g043025.1 | Solyc11g150133.1 |
| Solyc11g042990.1 | Solyc11g042990.2 |
| Solyc11g042930.1 | Solyc11g042930.2 |
| Solyc11g042890.1 | Solyc11g042890.2 |
| Solyc11g042850.1 | Solyc11g042850.2 |
| Solyc11g042820.1 | Solyc11g042820.2 |
| Solyc11g042810.1 | Solyc11g042810.2 |
| Solyc11g042800.1 | Solyc11g042800.2 |

| | |
|---|---|
| Solyc11g042740.1 | Solyc11g042740.2 |
| Solyc11g042640.1 | Solyc11g042640.2 |
| Solyc11g042610.1 | Solyc11g042610.2 |
| Solyc11g042615.1 | Solyc11g150132.1 |
| Solyc11g042460.1 | Solyc11g042460.2 |
| Solyc11g042430.1 | Solyc11g042430.2 |
| Solyc11g042420.1 | Solyc11g042420.2 |
| Solyc11g040390.1 | Solyc11g040390.2 |
| Solyc11g040380.1 | Solyc11g040380.2 |
| Solyc11g040370.1 | Solyc11g040370.2 |
| Solyc11g040330.1 | Solyc11g040330.2 |
| Solyc11g040300.1 | Solyc11g040300.2 |
| Solyc11g040280.1 | Solyc11g040280.2 |
| Solyc11g040285.1 | Solyc11g150131.1 |
| Solyc11g040260.1 | Solyc11g040260.2 |
| Solyc11g040220.1 | Solyc11g040220.2 |
| Solyc11g040177.1 | Solyc11g150130.1 |
| Solyc11g040150.1 | Solyc11g040150.2 |
| Solyc11g040140.1 | Solyc11g040140.2 |
| Solyc11g040130.1 | Solyc11g040130.2 |
| Solyc11g040110.1 | Solyc11g040110.2 |
| Solyc11g040040.1 | Solyc11g040040.2 |
| Solyc11g039960.1 | Solyc11g039960.2 |
| Solyc11g039930.1 | Solyc11g039930.2 |
| Solyc11g039910.1 | Solyc11g039910.2 |
| Solyc11g039890.1 | Solyc11g039890.2 |
| Solyc11g039880.1 | Solyc11g039880.2 |
| Solyc11g039840.1 | Solyc11g039840.2 |
| Solyc11g039830.1 | Solyc11g039830.2 |
| Solyc11g039800.1 | Solyc11g039800.2 |
| Solyc11g039650.1 | Solyc11g039650.2 |
| Solyc11g039530.1 | Solyc11g039530.2 |
| Solyc11g038350.1 | Solyc11g038350.2 |
| Solyc11g039360.1 | Solyc11g039360.2 |
| Solyc11g056590.1 | Solyc11g056590.2 |
| Solyc11g056650.1 | Solyc11g056650.2 |
| Solyc11g056670.1 | Solyc11g056670.2 |
| Solyc11g061700.1 | Solyc11g061700.2 |
| Solyc11g061750.1 | Solyc11g061750.2 |
| Solyc11g061770.1 | Solyc11g061770.2 |
| Solyc11g061780.1 | Solyc11g061780.2 |
| Solyc11g061960.1 | Solyc11g061960.2 |
| Solyc11g061965.1 | Solyc11g150140.1 |

| | |
|---|---|
| Solyc11g061980.1 | Solyc11g061980.2 |
| Solyc11g062020.1 | Solyc11g062020.2 |
| Solyc11g062130.1 | Solyc11g062130.2 |
| Solyc11g062190.1 | Solyc11g062190.2 |
| Solyc11g062195.1 | Solyc11g150141.1 |
| Solyc11g062210.1 | Solyc11g062210.2 |
| Solyc11g062220.1 | Solyc11g062220.2 |
| Solyc11g062240.1 | Solyc11g062240.2 |
| Solyc11g062250.1 | Solyc11g062250.2 |
| Solyc11g062270.1 | Solyc11g062270.2 |
| Solyc11g062280.1 | Solyc11g062280.2 |
| Solyc11g062310.1 | Solyc11g062310.2 |
| Solyc11g062320.1 | Solyc11g062320.2 |
| Solyc11g062350.1 | Solyc11g062350.2 |
| Solyc11g062400.1 | Solyc11g062400.2 |
| Solyc11g062430.1 | Solyc11g062430.2 |
| Solyc11g062440.1 | Solyc11g062440.2 |
| Solyc11g063460.1 | Solyc11g063460.2 |
| Solyc11g056260.1 | Solyc11g056260.2 |
| Solyc11g063530.1 | Solyc11g063530.2 |
| Solyc11g063540.1 | Solyc11g063540.2 |
| Solyc11g063550.1 | Solyc11g063550.2 |
| Solyc11g063590.1 | Solyc11g063590.2 |
| Solyc11g063620.1 | Solyc11g063620.2 |
| Solyc11g063640.1 | Solyc11g063640.2 |
| Solyc11g064780.1 | Solyc11g064780.2 |
| Solyc11g064790.1 | Solyc11g064790.2 |
| Solyc11g064880.1 | Solyc11g064880.2 |
| Solyc11g064890.1 | Solyc11g064890.2 |
| Solyc11g064900.1 | Solyc11g064900.2 |
| Solyc11g064920.1 | Solyc11g064920.2 |
| Solyc11g064960.1 | Solyc11g064960.2 |
| Solyc11g064990.1 | Solyc11g064990.2 |
| Solyc11g065000.1 | Solyc11g065000.2 |
| Solyc11g065040.1 | Solyc11g065040.2 |
| Solyc11g065070.1 | Solyc11g065070.2 |
| Solyc11g065080.1 | Solyc11g065080.2 |
| Solyc11g065110.1 | Solyc11g065110.2 |
| Solyc11g065125.1 | Solyc11g150142.1 |
| Solyc11g065180.1 | Solyc11g065180.2 |
| Solyc11g065210.1 | Solyc11g065210.3 |
| Solyc11g065220.1 | Solyc11g065220.2 |
| Solyc11g065240.1 | Solyc11g065240.2 |

| | |
|---|---|
| Solyc11g065320.1 | Solyc11g065320.2 |
| Solyc11g065340.1 | Solyc11g065340.2 |
| Solyc11g065350.1 | Solyc11g065350.2 |
| Solyc11g065360.1 | Solyc11g065360.2 |
| Solyc11g065380.1 | Solyc11g065380.2 |
| Solyc11g065400.1 | Solyc11g065400.2 |
| Solyc11g065425.1 | Solyc11g150143.1 |
| Solyc11g065470.1 | Solyc11g065470.2 |
| Solyc11g065490.1 | Solyc11g065490.2 |
| Solyc11g065600.1 | Solyc11g065600.2 |
| Solyc11g065620.1 | Solyc11g065620.2 |
| Solyc11g065640.1 | Solyc11g065640.2 |
| Solyc11g065660.1 | Solyc11g065660.2 |
| Solyc11g065700.1 | Solyc11g065700.2 |
| Solyc11g065720.1 | Solyc11g065720.2 |
| Solyc11g065740.1 | Solyc11g065740.2 |
| Solyc11g065750.1 | Solyc11g065750.2 |
| Solyc11g065760.1 | Solyc11g065760.2 |
| Solyc11g065830.1 | Solyc11g065830.2 |
| Solyc11g065845.1 | Solyc11g150144.1 |
| Solyc11g065880.1 | Solyc11g065880.2 |
| Solyc11g065890.1 | Solyc11g065890.2 |
| Solyc11g065900.1 | Solyc11g065900.2 |
| Solyc11g065920.1 | Solyc11g065920.2 |
| Solyc11g065940.1 | Solyc11g065940.2 |
| Solyc11g065990.1 | Solyc11g065990.2 |
| Solyc11g066020.1 | Solyc11g066020.2 |
| Solyc11g066040.1 | Solyc11g066040.2 |
| Solyc11g066100.1 | Solyc11g066100.2 |
| Solyc11g066110.1 | Solyc11g066110.2 |
| Solyc11g066140.1 | Solyc11g066140.2 |
| Solyc11g066250.1 | Solyc11g066250.2 |
| Solyc11g066260.1 | Solyc11g066260.2 |
| Solyc11g066290.1 | Solyc11g066290.2 |
| Solyc11g066320.1 | Solyc11g066320.2 |
| Solyc11g066330.1 | Solyc11g066330.2 |
| Solyc11g066350.1 | Solyc11g066350.2 |
| Solyc11g066360.1 | Solyc11g066360.2 |
| Solyc11g066370.1 | Solyc11g066370.2 |
| Solyc11g066390.1 | Solyc11g066390.2 |
| Solyc11g066410.1 | Solyc11g066410.2 |
| Solyc11g066430.1 | Solyc11g066430.2 |
| Solyc11g066440.1 | Solyc11g066440.2 |

| | |
|---|---|
| Solyc11g066480.1 | Solyc11g066480.2 |
| Solyc11g066500.1 | Solyc11g066500.2 |
| Solyc11g066540.1 | Solyc11g066540.2 |
| Solyc11g066580.2 | Solyc11g066580.3 |
| Solyc11g066590.1 | Solyc11g066590.2 |
| Solyc11g066600.1 | Solyc11g066600.2 |
| Solyc11g066610.1 | Solyc11g066610.2 |
| Solyc11g066620.1 | Solyc11g066620.2 |
| Solyc11g066630.1 | Solyc11g066630.2 |
| Solyc11g066690.1 | Solyc11g066690.2 |
| Solyc11g066700.1 | Solyc11g066700.2 |
| Solyc11g066730.1 | Solyc11g066730.2 |
| Solyc11g066740.1 | Solyc11g066740.2 |
| Solyc11g066790.1 | Solyc11g066790.2 |
| Solyc11g066800.1 | Solyc11g066800.2 |
| Solyc11g066810.1 | Solyc11g066810.2 |
| Solyc11g066820.1 | Solyc11g066820.2 |
| Solyc11g066830.1 | Solyc11g066830.2 |
| Solyc11g066880.1 | Solyc11g066880.2 |
| Solyc11g066900.1 | Solyc11g066900.2 |
| Solyc11g066950.1 | Solyc11g066950.2 |
| Solyc11g066990.1 | Solyc11g066990.2 |
| Solyc11g067010.1 | Solyc11g067010.2 |
| Solyc11g067040.1 | Solyc11g067040.2 |
| Solyc11g067060.1 | Solyc11g067060.3 |
| Solyc11g067090.1 | Solyc11g067090.2 |
| Solyc11g067100.1 | Solyc11g067100.2 |
| Solyc11g067110.1 | Solyc11g067110.2 |
| Solyc11g067140.1 | Solyc11g067140.2 |
| Solyc11g067160.1 | Solyc11g067160.2 |
| Solyc11g067180.1 | Solyc11g067180.2 |
| Solyc11g067200.1 | Solyc11g067200.2 |
| Solyc11g067220.1 | Solyc11g067220.2 |
| Solyc11g067230.1 | Solyc11g067230.2 |
| Solyc11g067240.1 | Solyc11g067240.2 |
| Solyc11g067280.1 | Solyc11g067280.2 |
| Solyc11g067300.1 | Solyc11g067300.2 |
| Solyc11g068350.1 | Solyc11g068350.2 |
| Solyc11g068400.1 | Solyc11g068400.2 |
| Solyc11g068420.1 | Solyc11g068420.2 |
| Solyc11g068440.1 | Solyc11g068440.2 |
| Solyc11g068450.1 | Solyc11g068450.2 |
| Solyc11g068460.1 | Solyc11g068460.2 |

| | |
|---|---|
| Solyc11g068480.1 | Solyc11g068480.2 |
| Solyc11g068500.1 | Solyc11g068500.2 |
| Solyc11g068540.1 | Solyc11g068540.2 |
| Solyc11g068560.1 | Solyc11g068560.2 |
| Solyc11g068610.1 | Solyc11g068610.2 |
| Solyc11g068620.1 | Solyc11g068620.2 |
| Solyc11g068640.1 | Solyc11g068640.2 |
| Solyc11g068650.1 | Solyc11g068650.2 |
| Solyc11g068660.1 | Solyc11g068660.2 |
| Solyc11g068670.1 | Solyc11g068670.2 |
| Solyc11g068690.1 | Solyc11g068690.2 |
| Solyc11g068700.1 | Solyc11g068700.2 |
| Solyc11g068720.1 | Solyc11g068720.2 |
| Solyc11g068730.1 | Solyc11g068730.2 |
| Solyc11g068740.1 | Solyc11g068740.2 |
| Solyc11g068745.1 | Solyc11g150145.1 |
| Solyc11g068750.1 | Solyc11g068750.2 |
| Solyc11g068800.1 | Solyc11g068800.2 |
| Solyc11g068830.1 | Solyc11g068830.2 |
| Solyc11g068870.1 | Solyc11g068870.2 |
| Solyc11g068880.1 | Solyc11g068880.2 |
| Solyc11g068890.1 | Solyc11g068890.2 |
| Solyc11g068960.1 | Solyc11g068960.2 |
| Solyc11g068970.1 | Solyc11g068970.2 |
| Solyc11g068980.1 | Solyc11g068980.2 |
| Solyc11g069000.1 | Solyc11g069000.2 |
| Solyc11g069010.1 | Solyc11g069010.2 |
| Solyc11g069040.2 | Solyc11g069040.3 |
| Solyc11g069050.1 | Solyc11g069050.2 |
| Solyc11g069090.1 | Solyc11g069090.2 |
| Solyc11g069095.1 | Solyc11g150146.1 |
| Solyc11g069130.1 | Solyc11g069130.2 |
| Solyc11g069140.1 | Solyc11g069140.2 |
| Solyc11g069170.1 | Solyc11g069170.2 |
| Solyc11g069180.1 | Solyc11g069180.2 |
| Solyc11g069190.1 | Solyc11g069190.2 |
| Solyc11g069220.1 | Solyc11g069220.2 |
| Solyc11g069240.1 | Solyc11g069240.2 |
| Solyc11g069250.1 | Solyc11g069250.2 |
| Solyc11g069260.1 | Solyc11g069260.2 |
| Solyc11g069270.1 | Solyc11g069270.2 |
| Solyc11g069280.1 | Solyc11g069280.2 |
| Solyc11g069290.1 | Solyc11g069290.2 |

| | |
|---|---|
| Solyc11g069300.1 | Solyc11g069300.2 |
| Solyc11g069340.1 | Solyc11g069340.2 |
| Solyc11g069350.1 | Solyc11g069350.2 |
| Solyc11g069360.1 | Solyc11g069360.2 |
| Solyc11g069380.1 | Solyc11g069380.2 |
| Solyc11g069400.1 | Solyc11g069400.2 |
| Solyc11g069430.1 | Solyc11g069430.2 |
| Solyc11g069460.1 | Solyc11g069460.2 |
| Solyc11g069480.1 | Solyc11g069480.2 |
| Solyc11g069500.1 | Solyc11g069500.2 |
| Solyc11g069520.1 | Solyc11g069520.2 |
| Solyc11g069530.1 | Solyc11g069530.2 |
| Solyc11g069570.1 | Solyc11g069570.2 |
| Solyc11g069580.1 | Solyc11g069580.2 |
| Solyc11g069590.1 | Solyc11g069590.2 |
| Solyc11g069600.1 | Solyc11g069600.2 |
| Solyc11g069640.1 | Solyc11g069640.2 |
| Solyc11g069650.1 | Solyc11g069650.2 |
| Solyc11g069670.1 | Solyc11g069670.2 |
| Solyc11g069700.1 | Solyc11g069700.2 |
| Solyc11g069710.1 | Solyc11g069710.2 |
| Solyc11g069720.1 | Solyc11g069720.2 |
| Solyc11g069770.1 | Solyc11g069770.2 |
| Solyc11g069790.1 | Solyc11g069790.2 |
| Solyc11g069830.1 | Solyc11g069830.2 |
| Solyc11g069910.1 | Solyc11g069910.2 |
| Solyc11g069950.1 | Solyc11g069950.2 |
| Solyc11g069960.1 | Solyc11g069960.2 |
| Solyc11g069990.1 | Solyc11g069990.2 |
| Solyc11g070030.1 | Solyc11g070030.2 |
| Solyc11g070100.1 | Solyc11g070100.2 |
| Solyc11g070120.1 | Solyc11g070120.2 |
| Solyc11g070130.1 | Solyc11g070130.2 |
| Solyc11g070140.1 | Solyc11g070140.2 |
| Solyc11g070150.1 | Solyc11g070150.2 |
| Solyc11g070160.1 | Solyc11g070160.2 |
| Solyc11g070170.1 | Solyc11g070170.2 |
| Solyc11g071230.1 | Solyc11g071230.2 |
| Solyc11g071240.1 | Solyc11g071240.2 |
| Solyc11g071260.1 | Solyc11g071260.2 |
| Solyc11g071270.1 | Solyc11g071270.2 |
| Solyc11g071280.1 | Solyc11g071280.2 |
| Solyc11g071290.1 | Solyc11g071290.2 |

| | |
|---|---|
| Solyc11g071300.1 | Solyc11g071300.2 |
| Solyc11g071320.1 | Solyc11g071320.2 |
| Solyc11g071440.1 | Solyc11g071440.2 |
| Solyc11g071460.1 | Solyc11g071460.2 |
| Solyc11g071490.1 | Solyc11g071490.2 |
| Solyc11g071520.1 | Solyc11g071520.2 |
| Solyc11g071540.1 | Solyc11g071540.2 |
| Solyc11g071550.1 | Solyc11g071550.2 |
| Solyc11g071560.1 | Solyc11g071560.2 |
| Solyc11g071670.1 | Solyc11g071670.2 |
| Solyc11g071680.1 | Solyc11g071680.2 |
| Solyc11g071700.1 | Solyc11g071700.2 |
| Solyc11g071710.1 | Solyc11g071710.2 |
| Solyc11g071720.1 | Solyc11g071720.2 |
| Solyc11g071790.1 | Solyc11g071790.2 |
| Solyc11g071800.1 | Solyc11g071800.2 |
| Solyc11g071810.1 | Solyc11g071810.2 |
| Solyc11g071830.1 | Solyc11g071830.2 |
| Solyc11g071840.1 | Solyc11g071840.2 |
| Solyc11g071860.1 | Solyc11g071860.2 |
| Solyc11g071870.1 | Solyc11g071870.2 |
| Solyc11g071910.1 | Solyc11g071910.2 |
| Solyc11g071930.1 | Solyc11g071930.2 |
| Solyc11g071960.1 | Solyc11g071960.2 |
| Solyc11g071970.1 | Solyc11g071970.2 |
| Solyc11g071990.1 | Solyc11g071990.2 |
| Solyc11g072000.1 | Solyc11g072000.2 |
| Solyc11g072040.1 | Solyc11g072040.2 |
| Solyc11g072090.1 | Solyc11g072090.2 |
| Solyc11g072100.1 | Solyc11g072100.2 |
| Solyc11g072110.1 | Solyc11g072110.2 |
| Solyc11g072120.1 | Solyc11g072120.2 |
| Solyc11g072130.1 | Solyc11g072130.2 |
| Solyc11g072170.1 | Solyc11g072170.2 |
| Solyc11g072190.1 | Solyc11g072190.2 |
| Solyc11g072200.1 | Solyc11g072200.2 |
| Solyc11g072220.1 | Solyc11g072220.2 |
| Solyc11g072240.1 | Solyc11g072240.2 |
| Solyc11g072260.1 | Solyc11g072260.2 |
| Solyc11g072300.1 | Solyc11g072300.2 |
| Solyc11g072310.1 | Solyc11g072310.2 |
| Solyc11g072330.1 | Solyc11g072330.2 |
| Solyc11g072340.1 | Solyc11g072340.2 |

| | |
|---|---|
| Solyc11g072380.1 | Solyc11g072380.2 |
| Solyc11g072390.1 | Solyc11g072390.2 |
| Solyc11g072410.1 | Solyc11g072410.2 |
| Solyc11g072440.1 | Solyc11g072440.2 |
| Solyc11g072450.1 | Solyc11g072450.2 |
| Solyc11g072460.1 | Solyc11g072460.2 |
| Solyc11g072480.1 | Solyc11g072480.2 |
| Solyc11g072490.1 | Solyc11g072490.2 |
| Solyc11g072500.1 | Solyc11g072500.2 |
| Solyc11g072530.1 | Solyc11g072530.2 |
| Solyc11g072550.1 | Solyc11g072550.2 |
| Solyc11g072570.1 | Solyc11g072570.2 |
| Solyc11g072580.1 | Solyc11g072580.2 |
| Solyc11g072630.1 | Solyc11g072630.2 |
| Solyc11g072660.1 | Solyc11g072660.2 |
| Solyc11g072690.1 | Solyc11g072690.2 |
| Solyc11g072700.1 | Solyc11g072700.2 |
| Solyc11g072720.1 | Solyc11g072720.2 |
| Solyc11g072730.1 | Solyc11g072730.2 |
| Solyc11g072770.1 | Solyc11g072770.2 |
| Solyc11g072790.1 | Solyc11g072790.2 |
| Solyc11g072810.1 | Solyc11g072810.2 |
| Solyc11g072850.1 | Solyc11g072850.2 |
| Solyc11g072860.1 | Solyc11g072860.2 |
| Solyc11g072880.1 | Solyc11g072880.2 |
| Solyc11g072920.1 | Solyc11g072920.2 |
| Solyc11g072940.1 | Solyc11g072940.2 |
| Solyc11g072970.1 | Solyc11g072970.2 |
| Solyc11g073050.1 | Solyc11g073050.2 |
| Solyc11g073100.1 | Solyc11g073100.2 |
| Solyc11g073110.1 | Solyc11g073110.2 |
| Solyc11g073120.1 | Solyc11g073120.2 |
| Solyc11g073130.1 | Solyc11g073130.2 |
| Solyc11g073180.1 | Solyc11g073180.2 |
| Solyc11g073190.1 | Solyc11g073190.2 |
| Solyc11g073200.1 | Solyc11g073200.2 |
| Solyc11g073210.1 | Solyc11g073210.2 |
| Solyc11g073240.1 | Solyc11g073240.2 |
| Solyc11g073250.1 | Solyc11g073250.2 |
| Solyc11g073260.1 | Solyc11g073260.2 |
| Solyc11g073280.1 | Solyc11g073280.2 |
| Solyc11g073300.1 | Solyc11g073300.2 |
| Solyc11g073310.1 | Solyc11g073310.2 |

| | |
|---|---|
| Solyc12g005010.1 | Solyc12g005010.2 |
| Solyc12g005020.1 | Solyc12g005020.2 |
| Solyc12g005040.1 | Solyc12g005040.2 |
| Solyc12g005050.1 | Solyc12g005050.2 |
| Solyc12g005070.1 | Solyc12g005070.2 |
| Solyc12g005080.1 | Solyc12g005080.2 |
| Solyc12g005095.1 | Solyc12g150100.1 |
| Solyc12g005110.1 | Solyc12g005110.2 |
| Solyc12g005140.1 | Solyc12g005140.2 |
| Solyc12g005150.1 | Solyc12g005150.2 |
| Solyc12g005170.1 | Solyc12g005170.2 |
| Solyc12g005180.1 | Solyc12g005180.2 |
| Solyc12g005220.1 | Solyc12g005220.2 |
| Solyc12g005250.1 | Solyc12g005250.2 |
| Solyc12g005270.1 | Solyc12g005270.2 |
| Solyc12g005290.1 | Solyc12g005290.2 |
| Solyc12g005300.1 | Solyc12g005300.2 |
| Solyc12g005310.1 | Solyc12g005310.2 |
| Solyc12g005330.1 | Solyc12g005330.2 |
| Solyc12g005350.1 | Solyc12g005350.2 |
| Solyc12g005370.1 | Solyc12g005370.2 |
| Solyc12g005390.1 | Solyc12g005390.2 |
| Solyc12g005410.1 | Solyc12g005410.2 |
| Solyc12g005420.1 | Solyc12g005420.2 |
| Solyc12g005460.1 | Solyc12g005460.2 |
| Solyc12g005465.1 | Solyc12g005465.2 |
| Solyc12g005470.1 | Solyc12g005470.2 |
| Solyc12g005480.1 | Solyc12g005480.2 |
| Solyc12g005580.1 | Solyc12g005580.2 |
| Solyc12g005590.1 | Solyc12g005590.2 |
| Solyc12g005630.1 | Solyc12g005630.2 |
| Solyc12g005640.1 | Solyc12g005640.2 |
| Solyc12g005660.1 | Solyc12g005660.2 |
| Solyc12g005700.1 | Solyc12g005700.2 |
| Solyc12g005730.1 | Solyc12g005730.2 |
| Solyc12g005760.1 | Solyc12g005760.2 |
| Solyc12g005790.1 | Solyc12g005790.2 |
| Solyc12g005840.1 | Solyc12g005840.2 |
| Solyc12g005850.1 | Solyc12g005850.2 |
| Solyc12g005860.1 | Solyc12g005860.2 |
| Solyc12g005880.1 | Solyc12g005880.2 |
| Solyc12g005910.1 | Solyc12g005910.2 |
| Solyc12g005920.1 | Solyc12g005920.2 |

| | |
|---|---|
| Solyc12g005930.1 | Solyc12g005930.2 |
| Solyc12g005940.1 | Solyc12g005940.2 |
| Solyc12g005950.1 | Solyc12g005950.2 |
| Solyc12g005980.1 | Solyc12g005980.2 |
| Solyc12g005990.1 | Solyc12g005990.2 |
| Solyc12g006050.1 | Solyc12g006050.2 |
| Solyc12g006060.1 | Solyc12g006060.2 |
| Solyc12g006080.1 | Solyc12g006080.2 |
| Solyc12g006100.1 | Solyc12g006100.2 |
| Solyc12g006140.1 | Solyc12g006140.2 |
| Solyc12g006170.1 | Solyc12g006170.2 |
| Solyc12g006180.1 | Solyc12g006180.2 |
| Solyc12g006240.1 | Solyc12g006240.2 |
| Solyc12g006270.1 | Solyc12g006270.2 |
| Solyc12g006290.1 | Solyc12g006290.2 |
| Solyc12g006300.1 | Solyc12g006300.2 |
| Solyc12g006310.1 | Solyc12g006310.2 |
| Solyc12g006400.1 | Solyc12g006400.2 |
| Solyc12g006420.1 | Solyc12g006420.2 |
| Solyc12g006450.1 | Solyc12g006450.2 |
| Solyc12g006460.1 | Solyc12g006460.2 |
| Solyc12g006470.1 | Solyc12g006470.2 |
| Solyc12g006480.1 | Solyc12g006480.2 |
| Solyc12g006490.1 | Solyc12g006490.2 |
| Solyc12g006520.1 | Solyc12g006520.2 |
| Solyc12g006530.1 | Solyc12g006530.2 |
| Solyc12g006540.1 | Solyc12g006540.2 |
| Solyc12g006550.1 | Solyc12g006550.2 |
| Solyc12g006570.1 | Solyc12g006570.2 |
| Solyc12g006620.1 | Solyc12g006620.2 |
| Solyc12g006630.1 | Solyc12g006630.2 |
| Solyc12g006640.1 | Solyc12g006640.2 |
| Solyc12g006650.1 | Solyc12g006650.2 |
| Solyc12g006680.1 | Solyc12g006680.2 |
| Solyc12g006690.1 | Solyc12g006690.2 |
| Solyc12g006780.1 | Solyc12g006780.2 |
| Solyc12g006800.1 | Solyc12g006800.2 |
| Solyc12g006830.1 | Solyc12g006830.2 |
| Solyc12g006840.1 | Solyc12g006840.2 |
| Solyc12g006850.1 | Solyc12g006850.2 |
| Solyc12g006860.1 | Solyc12g006860.2 |
| Solyc12g006870.1 | Solyc12g006870.2 |
| Solyc12g006880.1 | Solyc12g006880.2 |

| | |
|---|---|
| Solyc12g006890.1 | Solyc12g006890.2 |
| Solyc12g006920.1 | Solyc12g006920.2 |
| Solyc12g006930.1 | Solyc12g006930.2 |
| Solyc12g006940.1 | Solyc12g006940.2 |
| Solyc12g006950.1 | Solyc12g006950.2 |
| Solyc12g006960.1 | Solyc12g006960.2 |
| Solyc12g006980.1 | Solyc12g006980.2 |
| Solyc12g006990.1 | Solyc12g006990.2 |
| Solyc12g007010.1 | Solyc12g007010.2 |
| Solyc12g007020.1 | Solyc12g007020.2 |
| Solyc12g007070.1 | Solyc12g007070.2 |
| Solyc12g007080.1 | Solyc12g007080.2 |
| Solyc12g007100.1 | Solyc12g007100.2 |
| Solyc12g007110.1 | Solyc12g007110.2 |
| Solyc12g007120.1 | Solyc12g007120.2 |
| Solyc12g007130.1 | Solyc12g007130.2 |
| Solyc12g007160.1 | Solyc12g007160.2 |
| Solyc12g007180.1 | Solyc12g007180.2 |
| Solyc12g007190.1 | Solyc12g007190.2 |
| Solyc12g007200.1 | Solyc12g007200.2 |
| Solyc12g007210.1 | Solyc12g007210.2 |
| Solyc12g007230.1 | Solyc12g007230.2 |
| Solyc12g007260.1 | Solyc12g007260.2 |
| Solyc12g008320.1 | Solyc12g008320.2 |
| Solyc12g008340.1 | Solyc12g008340.2 |
| Solyc12g008360.1 | Solyc12g008360.2 |
| Solyc12g008400.1 | Solyc12g008400.2 |
| Solyc12g008450.1 | Solyc12g008450.2 |
| Solyc12g008460.1 | Solyc12g008460.2 |
| Solyc12g008500.1 | Solyc12g008500.2 |
| Solyc12g008510.1 | Solyc12g008510.2 |
| Solyc12g008520.1 | Solyc12g008520.2 |
| Solyc12g008530.1 | Solyc12g008530.2 |
| Solyc12g008540.1 | Solyc12g008540.2 |
| Solyc12g008580.1 | Solyc12g008580.2 |
| Solyc12g008600.1 | Solyc12g008600.2 |
| Solyc12g008620.1 | Solyc12g008620.2 |
| Solyc12g008630.1 | Solyc12g008630.2 |
| Solyc12g008650.1 | Solyc12g008650.2 |
| Solyc12g008670.1 | Solyc12g008670.2 |
| Solyc12g008710.1 | Solyc12g008710.2 |
| Solyc12g008720.1 | Solyc12g008720.2 |
| Solyc12g008770.1 | Solyc12g008770.2 |

| | |
|---|---|
| Solyc12g008780.1 | Solyc12g008780.2 |
| Solyc12g008810.1 | Solyc12g008810.2 |
| Solyc12g008840.1 | Solyc12g008840.2 |
| Solyc12g008900.1 | Solyc12g008900.2 |
| Solyc12g008910.1 | Solyc12g008910.2 |
| Solyc12g008940.1 | Solyc12g008940.2 |
| Solyc12g008960.1 | Solyc12g008960.2 |
| Solyc12g008990.1 | Solyc12g008990.2 |
| Solyc12g009010.1 | Solyc12g009010.2 |
| Solyc12g009020.1 | Solyc12g009020.2 |
| Solyc12g009030.1 | Solyc12g009030.2 |
| Solyc12g009060.1 | Solyc12g009060.2 |
| Solyc12g009065.1 | Solyc12g150101.1 |
| Solyc12g009075.1 | Solyc12g150102.1 |
| Solyc12g009080.1 | Solyc12g009080.2 |
| Solyc12g009100.1 | Solyc12g009100.2 |
| Solyc12g009120.1 | Solyc12g009120.2 |
| Solyc12g009150.1 | Solyc12g009150.2 |
| Solyc12g009160.1 | Solyc12g009160.2 |
| Solyc12g009170.1 | Solyc12g009170.2 |
| Solyc12g009180.1 | Solyc12g009180.2 |
| Solyc12g009190.1 | Solyc12g009190.2 |
| Solyc12g009200.1 | Solyc12g009200.2 |
| Solyc12g009210.1 | Solyc12g009210.2 |
| Solyc12g009220.1 | Solyc12g009220.2 |
| Solyc12g009300.1 | Solyc12g009300.3 |
| Solyc12g009310.1 | Solyc12g009310.2 |
| Solyc12g009340.1 | Solyc12g009340.2 |
| Solyc12g009380.1 | Solyc12g009380.2 |
| Solyc12g009390.1 | Solyc12g009390.2 |
| Solyc12g009400.1 | Solyc12g009400.2 |
| Solyc12g009410.1 | Solyc12g009410.2 |
| Solyc12g009430.1 | Solyc12g009430.2 |
| Solyc12g009440.1 | Solyc12g009440.2 |
| Solyc12g009470.1 | Solyc12g009470.2 |
| Solyc12g009480.1 | Solyc12g009480.2 |
| Solyc12g009500.1 | Solyc12g009500.2 |
| Solyc12g009520.1 | Solyc12g009520.2 |
| Solyc12g009550.1 | Solyc12g009550.2 |
| Solyc12g009560.1 | Solyc12g009560.2 |
| Solyc12g009580.1 | Solyc12g009580.2 |
| Solyc12g009590.1 | Solyc12g009590.2 |
| Solyc12g009600.1 | Solyc12g009600.2 |

| | |
|---|---|
| Solyc12g009610.1 | Solyc12g009610.2 |
| Solyc12g009620.1 | Solyc12g009620.2 |
| Solyc12g009650.1 | Solyc12g009650.2 |
| Solyc12g009680.1 | Solyc12g009680.2 |
| Solyc12g009770.1 | Solyc12g009770.2 |
| Solyc12g009790.1 | Solyc12g009790.2 |
| Solyc12g009820.1 | Solyc12g009820.2 |
| Solyc12g009830.1 | Solyc12g009830.2 |
| Solyc12g009840.1 | Solyc12g009840.2 |
| Solyc12g009870.1 | Solyc12g009870.2 |
| Solyc12g009940.1 | Solyc12g009940.2 |
| Solyc12g009950.1 | Solyc12g009950.2 |
| Solyc12g009960.1 | Solyc12g009960.2 |
| Solyc12g009980.1 | Solyc12g009980.2 |
| Solyc12g009990.1 | Solyc12g009990.2 |
| Solyc12g010010.1 | Solyc12g010010.2 |
| Solyc12g010040.1 | Solyc12g010040.2 |
| Solyc12g010050.1 | Solyc12g010050.2 |
| Solyc12g010060.1 | Solyc12g010060.2 |
| Solyc12g010115.1 | Solyc12g150103.1 |
| Solyc12g010120.1 | Solyc12g010120.2 |
| Solyc12g010140.1 | Solyc12g010140.2 |
| Solyc12g010170.1 | Solyc12g010170.2 |
| Solyc12g010180.1 | Solyc12g010180.2 |
| Solyc12g010200.1 | Solyc12g010200.2 |
| Solyc12g010210.1 | Solyc12g010210.2 |
| Solyc12g010230.1 | Solyc12g010230.2 |
| Solyc12g010310.1 | Solyc12g010310.2 |
| Solyc12g010320.1 | Solyc12g010320.2 |
| Solyc12g010330.1 | Solyc12g010330.2 |
| Solyc12g010360.1 | Solyc12g010360.2 |
| Solyc12g010390.1 | Solyc12g010390.2 |
| Solyc12g010460.1 | Solyc12g010460.2 |
| Solyc12g010480.1 | Solyc12g010480.2 |
| Solyc12g010510.1 | Solyc12g010510.2 |
| Solyc12g010580.1 | Solyc12g010580.2 |
| Solyc12g010690.1 | Solyc12g010690.2 |
| Solyc12g010700.1 | Solyc12g010700.2 |
| Solyc12g010730.1 | Solyc12g010730.2 |
| Solyc12g010750.1 | Solyc12g010750.2 |
| Solyc12g010780.1 | Solyc12g010780.2 |
| Solyc12g010790.1 | Solyc12g010790.2 |
| Solyc12g010800.1 | Solyc12g010800.2 |

| | |
|---|---|
| Solyc12g010820.1 | Solyc12g010820.2 |
| Solyc12g010840.1 | Solyc12g010840.2 |
| Solyc12g010860.1 | Solyc12g010860.2 |
| Solyc12g010890.1 | Solyc12g010890.2 |
| Solyc12g010910.1 | Solyc12g010910.2 |
| Solyc12g010920.1 | Solyc12g010920.2 |
| Solyc12g010930.1 | Solyc12g010930.2 |
| Solyc12g010950.1 | Solyc12g010950.2 |
| Solyc12g010960.1 | Solyc12g010960.2 |
| Solyc12g011000.1 | Solyc12g011000.2 |
| Solyc12g011010.1 | Solyc12g011010.2 |
| Solyc12g011020.1 | Solyc12g011020.2 |
| Solyc12g011050.1 | Solyc12g011050.2 |
| Solyc12g011100.1 | Solyc12g011100.2 |
| Solyc12g011130.1 | Solyc12g011130.2 |
| Solyc12g011160.1 | Solyc12g011160.2 |
| Solyc12g011190.1 | Solyc12g011190.2 |
| Solyc12g011270.1 | Solyc12g011270.2 |
| Solyc12g011280.1 | Solyc12g011280.2 |
| Solyc12g011290.1 | Solyc12g011290.2 |
| Solyc12g011300.1 | Solyc12g011300.2 |
| Solyc12g011310.1 | Solyc12g011310.2 |
| Solyc12g011320.1 | Solyc12g011320.2 |
| Solyc12g011340.1 | Solyc12g011340.2 |
| Solyc12g011360.1 | Solyc12g011360.2 |
| Solyc12g011420.1 | Solyc12g011420.2 |
| Solyc12g011440.1 | Solyc12g011440.2 |
| Solyc12g011450.1 | Solyc12g011450.2 |
| Solyc12g013530.1 | Solyc12g013530.2 |
| Solyc12g013540.1 | Solyc12g013540.2 |
| Solyc12g013560.1 | Solyc12g013560.2 |
| Solyc12g013570.1 | Solyc12g013570.2 |
| Solyc12g013590.1 | Solyc12g013590.2 |
| Solyc12g013610.1 | Solyc12g013610.2 |
| Solyc12g013620.1 | Solyc12g013620.2 |
| Solyc12g013640.1 | Solyc12g013640.2 |
| Solyc12g013650.1 | Solyc12g013650.2 |
| Solyc12g013670.1 | Solyc12g013670.2 |
| Solyc12g013690.1 | Solyc12g013690.2 |
| Solyc12g013700.1 | Solyc12g013700.2 |
| Solyc12g013710.1 | Solyc12g013710.2 |
| Solyc12g013750.1 | Solyc12g013750.2 |
| Solyc12g013810.1 | Solyc12g013810.2 |

| | |
|---|---|
| Solyc12g013830.1 | Solyc12g013830.2 |
| Solyc12g013850.1 | Solyc12g013850.2 |
| Solyc12g013860.1 | Solyc12g013860.2 |
| Solyc12g013880.1 | Solyc12g013880.2 |
| Solyc12g013910.1 | Solyc12g013910.2 |
| Solyc12g013920.1 | Solyc12g013920.2 |
| Solyc12g013930.1 | Solyc12g013930.2 |
| Solyc12g013950.1 | Solyc12g013950.2 |
| Solyc12g014010.1 | Solyc12g014010.2 |
| Solyc12g014050.1 | Solyc12g014050.2 |
| Solyc12g014060.1 | Solyc12g014060.2 |
| Solyc12g014065.1 | Solyc12g150104.1 |
| Solyc12g014070.1 | Solyc12g014070.2 |
| Solyc12g014080.1 | Solyc12g014080.2 |
| Solyc12g014100.1 | Solyc12g014100.2 |
| Solyc12g014110.1 | Solyc12g014110.2 |
| Solyc12g014130.1 | Solyc12g014130.2 |
| Solyc12g014170.1 | Solyc12g014170.2 |
| Solyc12g014180.1 | Solyc12g014180.2 |
| Solyc12g014190.1 | Solyc12g014190.2 |
| Solyc12g014230.1 | Solyc12g014230.2 |
| Solyc12g014250.1 | Solyc12g014250.2 |
| Solyc12g014280.1 | Solyc12g014280.2 |
| Solyc12g014290.1 | Solyc12g014290.2 |
| Solyc12g014320.1 | Solyc12g014320.2 |
| Solyc12g014355.1 | Solyc12g150105.1 |
| Solyc12g014360.1 | Solyc12g014360.2 |
| Solyc12g014410.1 | Solyc12g014410.2 |
| Solyc12g014420.1 | Solyc12g014420.2 |
| Solyc12g014430.1 | Solyc12g014430.2 |
| Solyc12g014440.1 | Solyc12g014440.2 |
| Solyc12g014450.1 | Solyc12g014450.2 |
| Solyc12g014460.1 | Solyc12g014460.2 |
| Solyc12g014470.1 | Solyc12g014470.2 |
| Solyc12g014540.1 | Solyc12g014540.2 |
| Solyc12g014570.1 | Solyc12g014570.2 |
| Solyc12g014580.1 | Solyc12g014580.2 |
| Solyc12g014600.1 | Solyc12g014600.2 |
| Solyc12g014610.1 | Solyc12g014610.2 |
| Solyc12g015640.1 | Solyc12g015640.2 |
| Solyc12g015660.1 | Solyc12g015660.2 |
| Solyc12g015710.1 | Solyc12g015710.2 |
| Solyc12g015740.1 | Solyc12g015740.2 |

| | |
|---|---|
| Solyc12g015780.1 | Solyc12g015780.2 |
| Solyc12g015800.1 | Solyc12g015800.2 |
| Solyc12g015860.1 | Solyc12g015860.2 |
| Solyc12g015870.1 | Solyc12g015870.2 |
| Solyc12g015880.1 | Solyc12g015880.2 |
| Solyc12g015920.1 | Solyc12g015920.2 |
| Solyc12g015930.1 | Solyc12g015930.2 |
| Solyc12g016190.1 | Solyc12g016190.2 |
| Solyc12g017240.1 | Solyc12g017240.2 |
| Solyc12g017250.1 | Solyc12g017250.2 |
| Solyc12g017260.1 | Solyc12g017260.2 |
| Solyc12g017280.1 | Solyc12g017280.2 |
| Solyc12g017350.1 | Solyc12g017350.2 |
| Solyc12g017410.1 | Solyc12g017410.2 |
| Solyc12g017420.1 | Solyc12g017420.2 |
| Solyc12g017450.1 | Solyc12g017450.2 |
| Solyc12g017510.1 | Solyc12g017510.2 |
| Solyc12g017540.1 | Solyc12g017540.2 |
| Solyc12g017570.1 | Solyc12g017570.2 |
| Solyc12g017630.1 | Solyc12g017630.2 |
| Solyc12g017675.1 | Solyc12g017675.2 |
| Solyc12g017700.1 | Solyc12g017700.2 |
| Solyc12g017760.1 | Solyc12g017760.2 |
| Solyc12g017780.1 | Solyc12g017780.2 |
| Solyc12g017820.1 | Solyc12g017820.2 |
| Solyc12g017835.1 | Solyc12g150106.1 |
| Solyc12g017840.1 | Solyc12g017840.2 |
| Solyc12g017850.1 | Solyc12g017850.2 |
| Solyc12g017870.1 | Solyc12g017870.2 |
| Solyc12g017885.1 | Solyc12g150107.1 |
| Solyc12g017910.1 | Solyc12g017910.2 |
| Solyc12g018990.1 | Solyc12g018990.2 |
| Solyc12g019000.1 | Solyc12g019000.2 |
| Solyc12g019010.1 | Solyc12g019010.2 |
| Solyc12g019050.1 | Solyc12g019050.2 |
| Solyc12g019060.1 | Solyc12g019060.2 |
| Solyc12g019090.1 | Solyc12g019090.2 |
| Solyc12g019100.1 | Solyc12g019100.2 |
| Solyc12g019120.1 | Solyc12g019120.2 |
| Solyc12g019180.1 | Solyc12g019180.2 |
| Solyc12g019200.1 | Solyc12g019200.2 |
| Solyc12g019230.1 | Solyc12g019230.2 |
| Solyc12g019300.1 | Solyc12g019300.2 |

| | |
|---|---|
| Solyc12g019320.1 | Solyc12g019320.2 |
| Solyc12g019460.1 | Solyc12g019460.2 |
| Solyc12g019490.1 | Solyc12g019490.2 |
| Solyc12g019540.1 | Solyc12g019540.2 |
| Solyc12g019570.1 | Solyc12g019570.2 |
| Solyc12g019620.1 | Solyc12g019620.2 |
| Solyc12g019640.1 | Solyc12g019640.2 |
| Solyc12g019690.1 | Solyc12g019690.2 |
| Solyc12g019740.1 | Solyc12g019740.2 |
| Solyc12g019860.1 | Solyc12g019860.2 |
| Solyc12g019890.1 | Solyc12g019890.2 |
| Solyc12g019900.1 | Solyc12g019900.2 |
| Solyc12g019960.1 | Solyc12g019960.2 |
| Solyc12g020050.1 | Solyc12g020050.2 |
| Solyc12g020110.1 | Solyc12g020110.2 |
| Solyc12g020120.1 | Solyc12g020120.2 |
| Solyc12g021130.1 | Solyc12g021130.2 |
| Solyc12g021170.1 | Solyc12g021170.2 |
| Solyc12g021180.1 | Solyc12g021180.2 |
| Solyc12g021190.1 | Solyc12g021190.2 |
| Solyc12g021200.1 | Solyc12g021200.2 |
| Solyc12g021230.1 | Solyc12g021230.2 |
| Solyc12g021280.1 | Solyc12g021280.2 |
| Solyc12g077590.1 | Solyc12g077590.2 |
| Solyc12g077460.1 | Solyc12g077460.2 |
| Solyc12g076350.1 | Solyc12g076350.2 |
| Solyc12g076320.1 | Solyc12g076320.2 |
| Solyc12g026370.1 | Solyc12g026370.2 |
| Solyc12g026380.1 | Solyc12g026380.2 |
| Solyc12g026400.1 | Solyc12g026400.2 |
| Solyc12g026470.1 | Solyc12g026470.2 |
| Solyc12g027550.1 | Solyc12g027550.2 |
| Solyc12g027730.1 | Solyc12g027730.2 |
| Solyc12g068070.1 | Solyc12g068070.2 |
| Solyc12g070100.1 | Solyc12g070100.2 |
| Solyc12g070130.1 | Solyc12g070130.2 |
| Solyc12g070156.1 | Solyc12g070156.2 |
| Solyc12g070270.1 | Solyc12g070270.2 |
| Solyc12g062223.1 | Solyc12g150126.1 |
| Solyc12g062250.1 | Solyc12g062250.2 |
| Solyc12g062260.1 | Solyc12g062260.2 |
| Solyc12g062280.1 | Solyc12g062280.2 |
| Solyc12g062290.1 | Solyc12g062290.2 |

| | |
|---|---|
| Solyc12g062340.1 | Solyc12g062340.2 |
| Solyc12g062420.1 | Solyc12g062420.2 |
| Solyc12g062640.1 | Solyc12g062640.2 |
| Solyc12g062690.1 | Solyc12g062690.2 |
| Solyc12g062733.1 | Solyc12g062733.2 |
| Solyc12g062860.1 | Solyc12g062860.2 |
| Solyc12g062870.1 | Solyc12g062870.2 |
| Solyc12g062875.1 | Solyc12g150127.1 |
| Solyc12g062910.1 | Solyc12g062910.2 |
| Solyc12g032990.1 | Solyc12g032990.2 |
| Solyc12g033130.1 | Solyc12g033130.2 |
| Solyc12g035130.1 | Solyc12g035130.2 |
| Solyc12g035190.1 | Solyc12g035190.2 |
| Solyc12g035225.1 | Solyc12g150108.1 |
| Solyc12g035240.1 | Solyc12g035240.2 |
| Solyc12g035270.1 | Solyc12g035270.2 |
| Solyc12g035360.1 | Solyc12g035360.2 |
| Solyc12g035400.1 | Solyc12g035400.2 |
| Solyc12g035450.1 | Solyc12g035450.2 |
| Solyc12g035470.1 | Solyc12g035470.2 |
| Solyc12g035590.1 | Solyc12g035590.2 |
| Solyc12g035600.1 | Solyc12g035600.2 |
| Solyc12g035620.1 | Solyc12g035620.2 |
| Solyc12g035650.1 | Solyc12g035650.2 |
| Solyc12g035710.1 | Solyc12g035710.2 |
| Solyc12g035790.1 | Solyc12g035790.2 |
| Solyc12g036095.1 | Solyc12g150109.1 |
| Solyc12g036170.1 | Solyc12g036170.2 |
| Solyc12g036200.1 | Solyc12g036200.2 |
| Solyc12g036250.1 | Solyc12g036250.2 |
| Solyc12g036390.1 | Solyc12g036390.2 |
| Solyc12g036470.1 | Solyc12g036470.2 |
| Solyc12g036480.1 | Solyc12g036480.2 |
| Solyc12g036500.1 | Solyc12g036500.2 |
| Solyc12g036560.1 | Solyc12g036560.2 |
| Solyc12g036600.1 | Solyc12g036600.2 |
| Solyc12g036653.1 | Solyc12g150110.1 |
| Solyc12g036657.1 | Solyc12g150111.1 |
| Solyc12g036673.1 | Solyc12g036673.2 |
| Solyc12g036790.1 | Solyc12g036790.2 |
| Solyc12g036820.1 | Solyc12g036820.2 |
| Solyc12g036850.1 | Solyc12g036850.2 |
| Solyc12g036915.1 | Solyc12g150112.1 |

| | |
|---|---|
| Solyc12g037950.1 | Solyc12g037950.2 |
| Solyc12g037970.1 | Solyc12g037970.2 |
| Solyc12g037980.1 | Solyc12g037980.2 |
| Solyc12g037990.1 | Solyc12g037990.2 |
| Solyc12g038145.1 | Solyc12g150114.1 |
| Solyc12g038160.1 | Solyc12g038160.2 |
| Solyc12g038230.1 | Solyc12g038230.2 |
| Solyc12g038340.1 | Solyc12g038340.2 |
| Solyc12g038370.1 | Solyc12g038370.2 |
| Solyc12g038400.1 | Solyc12g038400.2 |
| Solyc12g038410.1 | Solyc12g038410.2 |
| Solyc12g038420.1 | Solyc12g038420.2 |
| Solyc12g038490.1 | Solyc12g038490.2 |
| Solyc12g038510.1 | Solyc12g038510.2 |
| Solyc12g038580.1 | Solyc12g038580.2 |
| Solyc12g038590.1 | Solyc12g038590.2 |
| Solyc12g038600.1 | Solyc12g038600.2 |
| Solyc12g038630.1 | Solyc12g038630.2 |
| Solyc12g038670.1 | Solyc12g038670.2 |
| Solyc12g038750.1 | Solyc12g038750.2 |
| Solyc12g038840.1 | Solyc12g038840.2 |
| Solyc12g038920.1 | Solyc12g038920.2 |
| Solyc12g038980.1 | Solyc12g038980.2 |
| Solyc12g039045.1 | Solyc12g150115.1 |
| Solyc12g039070.1 | Solyc12g039070.2 |
| Solyc12g039100.1 | Solyc12g039100.2 |
| Solyc12g039120.1 | Solyc12g039120.2 |
| Solyc12g039180.1 | Solyc12g039180.2 |
| Solyc12g040210.1 | Solyc12g040210.2 |
| Solyc12g040270.1 | Solyc12g040270.2 |
| Solyc12g040320.1 | Solyc12g040320.2 |
| Solyc12g040340.1 | Solyc12g040340.2 |
| Solyc12g040450.1 | Solyc12g040450.2 |
| Solyc12g040470.1 | Solyc12g040470.2 |
| Solyc12g040480.1 | Solyc12g040480.2 |
| Solyc12g040500.1 | Solyc12g040500.2 |
| Solyc12g040510.1 | Solyc12g040510.2 |
| Solyc12g040520.1 | Solyc12g040520.2 |
| Solyc12g040640.1 | Solyc12g040640.2 |
| Solyc12g040680.1 | Solyc12g040680.2 |
| Solyc12g040790.1 | Solyc12g040790.2 |
| Solyc12g040800.1 | Solyc12g040800.2 |
| Solyc12g040810.1 | Solyc12g040810.2 |

| | |
|---|---|
| Solyc12g040860.1 | Solyc12g040860.2 |
| Solyc12g041890.1 | Solyc12g041890.2 |
| Solyc12g041960.1 | Solyc12g041960.2 |
| Solyc12g042080.1 | Solyc12g042080.2 |
| Solyc12g042200.1 | Solyc12g042200.2 |
| Solyc12g042210.1 | Solyc12g042210.2 |
| Solyc12g042240.1 | Solyc12g042240.2 |
| Solyc12g042300.1 | Solyc12g042300.2 |
| Solyc12g042355.1 | Solyc12g150116.1 |
| Solyc12g042380.1 | Solyc12g042380.2 |
| Solyc12g042430.1 | Solyc12g042430.2 |
| Solyc12g042440.1 | Solyc12g042440.2 |
| Solyc12g042460.1 | Solyc12g042460.2 |
| Solyc12g042470.1 | Solyc12g042470.2 |
| Solyc12g042500.1 | Solyc12g042500.2 |
| Solyc12g042530.1 | Solyc12g042530.2 |
| Solyc12g042550.1 | Solyc12g042550.2 |
| Solyc12g042570.1 | Solyc12g042570.2 |
| Solyc12g042580.1 | Solyc12g042580.2 |
| Solyc12g042590.1 | Solyc12g042590.2 |
| Solyc12g042600.1 | Solyc12g042600.2 |
| Solyc12g042650.1 | Solyc12g042650.2 |
| Solyc12g042655.1 | Solyc12g150117.1 |
| Solyc12g042740.1 | Solyc12g042740.2 |
| Solyc12g042750.1 | Solyc12g042750.2 |
| Solyc12g042780.1 | Solyc12g042780.2 |
| Solyc12g042840.1 | Solyc12g042840.2 |
| Solyc12g042850.1 | Solyc12g042850.2 |
| Solyc12g042890.1 | Solyc12g042890.2 |
| Solyc12g042900.1 | Solyc12g042900.2 |
| Solyc12g042910.1 | Solyc12g042910.2 |
| Solyc12g042920.1 | Solyc12g042920.2 |
| Solyc12g042930.1 | Solyc12g042930.2 |
| Solyc12g042950.1 | Solyc12g042950.2 |
| Solyc12g042965.1 | Solyc12g042965.2 |
| Solyc12g042967.1 | Solyc12g150118.1 |
| Solyc12g042980.1 | Solyc12g042980.2 |
| Solyc12g043020.1 | Solyc12g043020.2 |
| Solyc12g043100.1 | Solyc12g043100.2 |
| Solyc12g043110.1 | Solyc12g043110.2 |
| Solyc12g043120.1 | Solyc12g043120.2 |
| Solyc12g043165.1 | Solyc12g150119.1 |
| Solyc12g044230.1 | Solyc12g044230.2 |

| | |
|---|---|
| Solyc12g044250.1 | Solyc12g044250.2 |
| Solyc12g044260.1 | Solyc12g044260.2 |
| Solyc12g044270.1 | Solyc12g044270.2 |
| Solyc12g044280.1 | Solyc12g044280.2 |
| Solyc12g044290.1 | Solyc12g044290.2 |
| Solyc12g044300.1 | Solyc12g044300.2 |
| Solyc12g044310.1 | Solyc12g044310.2 |
| Solyc12g044320.1 | Solyc12g044320.2 |
| Solyc12g044330.1 | Solyc12g044330.2 |
| Solyc12g044360.1 | Solyc12g044360.2 |
| Solyc12g044410.1 | Solyc12g044410.2 |
| Solyc12g044440.1 | Solyc12g044440.2 |
| Solyc12g044450.1 | Solyc12g044450.2 |
| Solyc12g044520.1 | Solyc12g044520.2 |
| Solyc12g044600.2 | Solyc12g044600.3 |
| Solyc12g044610.1 | Solyc12g044610.2 |
| Solyc12g044630.1 | Solyc12g044630.2 |
| Solyc12g044640.1 | Solyc12g044640.2 |
| Solyc12g044655.1 | Solyc12g150120.1 |
| Solyc12g044670.1 | Solyc12g044670.2 |
| Solyc12g044685.1 | Solyc12g150121.1 |
| Solyc12g044700.1 | Solyc12g044700.2 |
| Solyc12g044720.1 | Solyc12g044720.2 |
| Solyc12g044730.1 | Solyc12g044730.2 |
| Solyc12g044740.1 | Solyc12g044740.2 |
| Solyc12g044780.1 | Solyc12g044780.2 |
| Solyc12g044840.1 | Solyc12g044840.2 |
| Solyc12g044880.1 | Solyc12g044880.2 |
| Solyc12g044890.1 | Solyc12g044890.2 |
| Solyc12g044910.1 | Solyc12g044910.2 |
| Solyc12g044940.1 | Solyc12g044940.2 |
| Solyc12g045020.1 | Solyc12g045020.2 |
| Solyc12g045030.1 | Solyc12g045030.2 |
| Solyc12g049050.1 | Solyc12g049050.2 |
| Solyc12g049120.1 | Solyc12g049120.2 |
| Solyc12g049170.1 | Solyc12g049170.2 |
| Solyc12g049230.1 | Solyc12g049230.2 |
| Solyc12g049235.1 | Solyc12g150122.1 |
| Solyc12g049250.1 | Solyc12g049250.2 |
| Solyc12g049290.1 | Solyc12g049290.2 |
| Solyc12g049300.1 | Solyc12g049300.2 |
| Solyc12g049310.1 | Solyc12g049310.2 |
| Solyc12g049330.1 | Solyc12g049330.2 |

| | |
|---|---|
| Solyc12g049350.1 | Solyc12g049350.2 |
| Solyc12g049360.1 | Solyc12g049360.2 |
| Solyc12g049370.1 | Solyc12g049370.2 |
| Solyc12g049390.1 | Solyc12g049390.2 |
| Solyc12g049400.1 | Solyc12g049400.2 |
| Solyc12g049420.1 | Solyc12g049420.2 |
| Solyc12g049520.1 | Solyc12g049520.2 |
| Solyc12g049530.1 | Solyc12g049530.2 |
| Solyc12g049550.1 | Solyc12g049550.2 |
| Solyc12g049560.1 | Solyc12g049560.2 |
| Solyc12g049580.1 | Solyc12g049580.2 |
| Solyc12g049590.1 | Solyc12g049590.2 |
| Solyc12g049613.1 | Solyc12g150123.1 |
| Solyc12g049670.1 | Solyc12g049670.2 |
| Solyc12g055720.1 | Solyc12g055720.2 |
| Solyc12g055740.1 | Solyc12g055740.2 |
| Solyc12g055760.1 | Solyc12g055760.2 |
| Solyc12g055830.1 | Solyc12g055830.2 |
| Solyc12g055840.1 | Solyc12g055840.2 |
| Solyc12g055850.1 | Solyc12g055850.2 |
| Solyc12g055855.1 | Solyc12g055855.2 |
| Solyc12g055870.1 | Solyc12g055870.2 |
| Solyc12g055990.1 | Solyc12g055990.2 |
| Solyc12g056010.1 | Solyc12g056010.2 |
| Solyc12g056080.1 | Solyc12g056080.2 |
| Solyc12g056090.1 | Solyc12g056090.2 |
| Solyc12g056110.1 | Solyc12g056110.2 |
| Solyc12g056130.1 | Solyc12g056130.2 |
| Solyc12g056140.1 | Solyc12g056140.2 |
| Solyc12g056150.1 | Solyc12g056150.2 |
| Solyc12g056160.1 | Solyc12g056160.2 |
| Solyc12g056190.1 | Solyc12g056190.2 |
| Solyc12g056200.1 | Solyc12g056200.2 |
| Solyc12g056220.1 | Solyc12g056220.2 |
| Solyc12g056230.1 | Solyc12g056230.2 |
| Solyc12g056240.1 | Solyc12g056240.2 |
| Solyc12g056250.1 | Solyc12g056250.2 |
| Solyc12g056270.1 | Solyc12g056270.2 |
| Solyc12g056310.1 | Solyc12g056310.2 |
| Solyc12g056340.1 | Solyc12g056340.2 |
| Solyc12g056350.1 | Solyc12g056350.2 |
| Solyc12g056410.1 | Solyc12g056410.2 |
| Solyc12g056420.1 | Solyc12g056420.2 |

| | |
|---|---|
| Solyc12g056440.1 | Solyc12g056440.2 |
| Solyc12g056450.1 | Solyc12g056450.2 |
| Solyc12g056470.1 | Solyc12g056470.2 |
| Solyc12g056480.1 | Solyc12g056480.2 |
| Solyc12g056530.1 | Solyc12g056530.2 |
| Solyc12g056560.1 | Solyc12g056560.2 |
| Solyc12g056580.1 | Solyc12g056580.2 |
| Solyc12g056590.1 | Solyc12g056590.2 |
| Solyc12g056600.2 | Solyc12g056600.3 |
| Solyc12g056620.1 | Solyc12g056620.2 |
| Solyc12g056625.1 | Solyc12g150124.1 |
| Solyc12g056640.1 | Solyc12g056640.2 |
| Solyc12g056650.1 | Solyc12g056650.2 |
| Solyc12g056673.1 | Solyc12g150125.1 |
| Solyc12g056700.1 | Solyc12g056700.2 |
| Solyc12g056720.1 | Solyc12g056720.2 |
| Solyc12g056740.1 | Solyc12g056740.2 |
| Solyc12g056780.1 | Solyc12g056780.2 |
| Solyc12g056790.1 | Solyc12g056790.2 |
| Solyc12g056800.1 | Solyc12g056800.2 |
| Solyc12g056810.1 | Solyc12g056810.2 |
| Solyc12g056820.1 | Solyc12g056820.2 |
| Solyc12g056840.1 | Solyc12g056840.2 |
| Solyc12g056850.1 | Solyc12g056850.2 |
| Solyc12g056860.1 | Solyc12g056860.2 |
| Solyc12g056900.1 | Solyc12g056900.2 |
| Solyc12g056930.1 | Solyc12g056930.2 |
| Solyc12g056940.1 | Solyc12g056940.2 |
| Solyc12g056950.1 | Solyc12g056950.2 |
| Solyc12g056960.1 | Solyc12g056960.2 |
| Solyc12g056995.1 | Solyc12g056995.2 |
| Solyc12g057020.1 | Solyc12g057020.2 |
| Solyc12g057030.1 | Solyc12g057030.2 |
| Solyc12g057040.1 | Solyc12g057040.2 |
| Solyc12g057050.1 | Solyc12g057050.2 |
| Solyc12g057060.1 | Solyc12g057060.2 |
| Solyc12g057070.1 | Solyc12g057070.2 |
| Solyc12g057080.1 | Solyc12g057080.2 |
| Solyc12g057100.1 | Solyc12g057100.2 |
| Solyc12g087830.1 | Solyc12g087830.2 |
| Solyc12g087870.1 | Solyc12g087870.2 |
| Solyc12g087900.1 | Solyc12g087900.2 |
| Solyc12g087920.1 | Solyc12g087920.2 |

| | |
|---|---|
| Solyc12g087940.1 | Solyc12g087940.2 |
| Solyc12g088000.1 | Solyc12g088000.2 |
| Solyc12g088010.1 | Solyc12g088010.2 |
| Solyc12g088030.1 | Solyc12g088030.2 |
| Solyc12g088040.1 | Solyc12g088040.2 |
| Solyc12g088050.1 | Solyc12g088050.2 |
| Solyc12g088060.1 | Solyc12g088060.2 |
| Solyc12g088070.1 | Solyc12g088070.2 |
| Solyc12g088140.1 | Solyc12g088140.2 |
| Solyc12g088170.1 | Solyc12g088170.2 |
| Solyc12g088190.1 | Solyc12g088190.2 |
| Solyc12g088220.1 | Solyc12g088220.2 |
| Solyc12g088240.1 | Solyc12g088240.2 |
| Solyc12g088250.1 | Solyc12g088250.2 |
| Solyc12g088260.1 | Solyc12g088260.2 |
| Solyc12g088270.1 | Solyc12g088270.2 |
| Solyc12g088290.1 | Solyc12g088290.2 |
| Solyc12g088320.1 | Solyc12g088320.2 |
| Solyc12g088340.1 | Solyc12g088340.2 |
| Solyc12g088360.1 | Solyc12g088360.2 |
| Solyc12g088370.1 | Solyc12g088370.2 |
| Solyc12g088420.1 | Solyc12g088420.2 |
| Solyc12g088440.1 | Solyc12g088440.2 |
| Solyc12g088480.1 | Solyc12g088480.2 |
| Solyc12g088500.1 | Solyc12g088500.2 |
| Solyc12g088520.1 | Solyc12g088520.2 |
| Solyc12g088530.1 | Solyc12g088530.2 |
| Solyc12g088620.1 | Solyc12g088620.2 |
| Solyc12g088630.1 | Solyc12g088630.2 |
| Solyc12g088640.1 | Solyc12g088640.2 |
| Solyc12g088650.1 | Solyc12g088650.2 |
| Solyc12g088660.1 | Solyc12g088660.2 |
| Solyc12g088670.1 | Solyc12g088670.2 |
| Solyc12g088680.1 | Solyc12g088680.2 |
| Solyc12g088690.1 | Solyc12g088690.2 |
| Solyc12g088720.1 | Solyc12g088720.2 |
| Solyc12g088730.1 | Solyc12g088730.2 |
| Solyc12g088740.1 | Solyc12g088740.2 |
| Solyc12g088750.1 | Solyc12g088750.2 |
| Solyc12g088820.1 | Solyc12g088820.2 |
| Solyc12g088860.1 | Solyc12g088860.2 |
| Solyc12g088880.1 | Solyc12g088880.2 |
| Solyc12g088900.1 | Solyc12g088900.2 |

| | |
|---|---|
| Solyc12g088920.1 | Solyc12g088920.2 |
| Solyc12g088960.1 | Solyc12g088960.2 |
| Solyc12g088980.1 | Solyc12g088980.2 |
| Solyc12g088990.1 | Solyc12g088990.2 |
| Solyc12g089000.1 | Solyc12g089000.2 |
| Solyc12g089020.1 | Solyc12g089020.2 |
| Solyc12g089030.1 | Solyc12g089030.2 |
| Solyc12g089060.1 | Solyc12g089060.2 |
| Solyc12g089090.1 | Solyc12g089090.2 |
| Solyc12g089100.1 | Solyc12g089100.2 |
| Solyc12g089130.1 | Solyc12g089130.2 |
| Solyc12g089140.1 | Solyc12g089140.2 |
| Solyc12g089150.1 | Solyc12g089150.2 |
| Solyc12g089180.1 | Solyc12g089180.2 |
| Solyc12g089210.1 | Solyc12g089210.2 |
| Solyc12g089230.1 | Solyc12g089230.2 |
| Solyc12g089240.1 | Solyc12g089240.2 |
| Solyc12g089280.1 | Solyc12g089280.2 |
| Solyc12g089293.1 | Solyc12g150128.1 |
| Solyc12g089295.1 | Solyc12g150129.1 |
| Solyc12g089297.1 | Solyc12g150130.1 |
| Solyc12g089300.1 | Solyc12g089300.2 |
| Solyc12g089310.1 | Solyc12g089310.2 |
| Solyc12g089330.1 | Solyc12g089330.2 |
| Solyc12g089340.1 | Solyc12g089340.2 |
| Solyc12g089370.1 | Solyc12g089370.2 |
| Solyc12g089380.1 | Solyc12g089380.2 |
| Solyc12g094380.1 | Solyc12g094380.2 |
| Solyc12g094400.1 | Solyc12g094400.2 |
| Solyc12g094420.1 | Solyc12g094420.2 |
| Solyc12g094440.1 | Solyc12g094440.2 |
| Solyc12g094460.1 | Solyc12g094460.2 |
| Solyc12g094490.1 | Solyc12g094490.2 |
| Solyc12g094520.1 | Solyc12g094520.2 |
| Solyc12g094550.1 | Solyc12g094550.2 |
| Solyc12g094560.1 | Solyc12g094560.2 |
| Solyc12g094590.1 | Solyc12g094590.2 |
| Solyc12g094630.1 | Solyc12g094630.2 |
| Solyc12g094660.1 | Solyc12g094660.2 |
| Solyc12g094710.1 | Solyc12g094710.2 |
| Solyc12g095750.1 | Solyc12g095750.2 |
| Solyc12g095760.1 | Solyc12g095760.2 |
| Solyc12g095770.1 | Solyc12g095770.2 |

| | |
|---|---|
| Solyc12g095780.1 | Solyc12g095780.2 |
| Solyc12g095790.1 | Solyc12g095790.2 |
| Solyc12g095800.1 | Solyc12g095800.2 |
| Solyc12g095810.1 | Solyc12g095810.2 |
| Solyc12g095840.1 | Solyc12g095840.2 |
| Solyc12g095850.1 | Solyc12g095850.2 |
| Solyc12g095860.1 | Solyc12g095860.2 |
| Solyc12g095880.1 | Solyc12g095880.2 |
| Solyc12g095890.1 | Solyc12g095890.2 |
| Solyc12g095900.1 | Solyc12g095900.2 |
| Solyc12g095910.1 | Solyc12g095910.2 |
| Solyc12g095920.1 | Solyc12g095920.2 |
| Solyc12g095930.1 | Solyc12g095930.2 |
| Solyc12g095980.1 | Solyc12g095980.2 |
| Solyc12g095990.1 | Solyc12g095990.2 |
| Solyc12g096000.1 | Solyc12g096000.2 |
| Solyc12g096050.1 | Solyc12g096050.2 |
| Solyc12g096060.1 | Solyc12g096060.2 |
| Solyc12g096070.1 | Solyc12g096070.2 |
| Solyc12g096080.1 | Solyc12g096080.2 |
| Solyc12g096090.1 | Solyc12g096090.2 |
| Solyc12g096100.1 | Solyc12g096100.2 |
| Solyc12g096120.1 | Solyc12g096120.2 |
| Solyc12g096130.1 | Solyc12g096130.2 |
| Solyc12g096150.1 | Solyc12g096150.2 |
| Solyc12g096160.1 | Solyc12g096160.2 |
| Solyc12g096190.1 | Solyc12g096190.2 |
| Solyc12g096200.1 | Solyc12g096200.2 |
| Solyc12g096210.1 | Solyc12g096210.2 |
| Solyc12g096220.1 | Solyc12g096220.2 |
| Solyc12g096230.1 | Solyc12g096230.2 |
| Solyc12g096250.1 | Solyc12g096250.2 |
| Solyc12g096260.1 | Solyc12g096260.2 |
| Solyc12g096270.1 | Solyc12g096270.2 |
| Solyc12g096280.1 | Solyc12g096280.2 |
| Solyc12g096300.1 | Solyc12g096300.2 |
| Solyc12g096320.1 | Solyc12g096320.2 |
| Solyc12g096340.1 | Solyc12g096340.2 |
| Solyc12g096350.1 | Solyc12g096350.2 |
| Solyc12g096370.1 | Solyc12g096370.2 |
| Solyc12g096390.1 | Solyc12g096390.2 |
| Solyc12g096400.1 | Solyc12g096400.2 |
| Solyc12g096420.1 | Solyc12g096420.2 |

| | |
|---|---|
| Solyc12g096450.1 | Solyc12g096450.2 |
| Solyc12g096470.1 | Solyc12g096470.2 |
| Solyc12g096500.1 | Solyc12g096500.2 |
| Solyc12g096540.1 | Solyc12g096540.2 |
| Solyc12g096550.1 | Solyc12g096550.2 |
| Solyc12g096560.1 | Solyc12g096560.2 |
| Solyc12g096580.1 | Solyc12g096580.2 |
| Solyc12g096600.1 | Solyc12g096600.2 |
| Solyc12g096610.1 | Solyc12g096610.2 |
| Solyc12g096630.1 | Solyc12g096630.2 |
| Solyc12g096650.1 | Solyc12g096650.2 |
| Solyc12g096670.1 | Solyc12g096670.2 |
| Solyc12g096690.1 | Solyc12g096690.2 |
| Solyc12g096700.1 | Solyc12g096700.2 |
| Solyc12g096710.1 | Solyc12g096710.2 |
| Solyc12g096760.1 | Solyc12g096760.2 |
| Solyc12g096780.1 | Solyc12g096780.2 |
| Solyc12g096810.1 | Solyc12g096810.2 |
| Solyc12g096830.1 | Solyc12g096830.2 |
| Solyc12g096900.1 | Solyc12g096900.2 |
| Solyc12g096930.1 | Solyc12g096930.2 |
| Solyc12g096950.1 | Solyc12g096950.2 |
| Solyc12g096960.1 | Solyc12g096960.2 |
| Solyc12g096970.1 | Solyc12g096970.2 |
| Solyc12g096990.1 | Solyc12g096990.2 |
| Solyc12g097000.1 | Solyc12g097000.2 |
| Solyc12g097020.1 | Solyc12g097020.2 |
| Solyc12g097030.1 | Solyc12g097030.2 |
| Solyc12g097050.1 | Solyc12g097050.2 |
| Solyc12g097080.1 | Solyc12g097080.2 |
| Solyc12g098140.1 | Solyc12g098140.2 |
| Solyc12g098150.1 | Solyc12g098150.2 |
| Solyc12g098160.1 | Solyc12g098160.2 |
| Solyc12g098190.1 | Solyc12g098190.2 |
| Solyc12g098200.1 | Solyc12g098200.2 |
| Solyc12g098310.1 | Solyc12g098310.2 |
| Solyc12g098340.1 | Solyc12g098340.2 |
| Solyc12g098360.1 | Solyc12g098360.2 |
| Solyc12g098370.1 | Solyc12g098370.2 |
| Solyc12g098400.1 | Solyc12g098400.2 |
| Solyc12g098410.1 | Solyc12g098410.2 |
| Solyc12g098430.1 | Solyc12g098430.2 |
| Solyc12g098440.1 | Solyc12g098440.2 |

| | |
|---|---|
| Solyc12g098460.1 | Solyc12g098460.2 |
| Solyc12g098490.1 | Solyc12g098490.2 |
| Solyc12g098500.1 | Solyc12g098500.2 |
| Solyc12g098520.1 | Solyc12g098520.2 |
| Solyc12g098530.1 | Solyc12g098530.2 |
| Solyc12g098540.1 | Solyc12g098540.2 |
| Solyc12g098560.1 | Solyc12g098560.2 |
| Solyc12g098570.1 | Solyc12g098570.2 |
| Solyc12g098620.1 | Solyc12g098620.2 |
| Solyc12g098630.1 | Solyc12g098630.2 |
| Solyc12g098650.1 | Solyc12g098650.2 |
| Solyc12g098690.1 | Solyc12g098690.2 |
| Solyc12g098710.1 | Solyc12g098710.2 |
| Solyc12g098740.1 | Solyc12g098740.2 |
| Solyc12g098753.1 | Solyc12g150131.1 |
| Solyc12g098757.1 | Solyc12g150132.1 |
| Solyc12g098770.1 | Solyc12g098770.2 |
| Solyc12g098780.1 | Solyc12g098780.2 |
| Solyc12g098800.1 | Solyc12g098800.2 |
| Solyc12g098810.1 | Solyc12g098810.2 |
| Solyc12g098820.1 | Solyc12g098820.2 |
| Solyc12g098830.1 | Solyc12g098830.2 |
| Solyc12g098850.1 | Solyc12g098850.2 |
| Solyc12g098860.1 | Solyc12g098860.2 |
| Solyc12g098880.1 | Solyc12g098880.2 |
| Solyc12g098890.1 | Solyc12g098890.2 |
| Solyc12g098900.1 | Solyc12g098900.2 |
| Solyc12g098910.1 | Solyc12g098910.2 |
| Solyc12g098920.1 | Solyc12g098920.2 |
| Solyc12g098940.1 | Solyc12g098940.2 |
| Solyc12g098950.1 | Solyc12g098950.2 |
| Solyc12g098960.1 | Solyc12g098960.2 |
| Solyc12g098980.1 | Solyc12g098980.2 |
| Solyc12g098990.1 | Solyc12g098990.2 |
| Solyc12g099040.1 | Solyc12g099040.2 |
| Solyc12g099100.1 | Solyc12g099100.2 |
| Solyc12g099110.1 | Solyc12g099110.2 |
| Solyc12g099130.1 | Solyc12g099130.2 |
| Solyc12g099140.1 | Solyc12g099140.2 |
| Solyc12g099160.1 | Solyc12g099160.2 |
| Solyc12g099200.1 | Solyc12g099200.2 |
| Solyc12g099210.1 | Solyc12g099210.2 |
| Solyc12g099230.1 | Solyc12g099230.2 |

| | |
|---|---|
| Solyc12g099240.1 | Solyc12g099240.2 |
| Solyc12g099250.1 | Solyc12g099250.2 |
| Solyc12g099260.1 | Solyc12g099260.2 |
| Solyc12g099270.1 | Solyc12g099270.2 |
| Solyc12g099280.1 | Solyc12g099280.2 |
| Solyc12g099290.1 | Solyc12g099290.2 |
| Solyc12g099310.1 | Solyc12g099310.2 |
| Solyc12g099320.1 | Solyc12g099320.2 |
| Solyc12g099330.1 | Solyc12g099330.2 |
| Solyc12g099360.1 | Solyc12g099360.2 |
| Solyc12g099370.1 | Solyc12g099370.2 |
| Solyc12g099390.1 | Solyc12g099390.2 |
| Solyc12g099410.1 | Solyc12g099410.2 |
| Solyc12g099420.1 | Solyc12g099420.2 |
| Solyc12g099430.1 | Solyc12g099430.2 |
| Solyc12g099470.1 | Solyc12g099470.2 |
| Solyc12g099480.1 | Solyc12g099480.2 |
| Solyc12g099540.1 | Solyc12g099540.2 |
| Solyc12g099560.1 | Solyc12g099560.2 |
| Solyc12g099580.1 | Solyc12g099580.2 |
| Solyc12g099600.1 | Solyc12g099600.2 |
| Solyc12g099630.1 | Solyc12g099630.2 |
| Solyc12g099650.1 | Solyc12g099650.2 |
| Solyc12g099660.1 | Solyc12g099660.2 |
| Solyc12g099720.1 | Solyc12g099720.2 |
| Solyc12g099740.1 | Solyc12g099740.2 |
| Solyc12g099760.1 | Solyc12g099760.2 |
| Solyc12g099770.1 | Solyc12g099770.2 |
| Solyc12g099800.1 | Solyc12g099800.2 |
| Solyc12g099810.1 | Solyc12g099810.2 |
| Solyc12g099830.1 | Solyc12g099830.2 |
| Solyc12g099860.1 | Solyc12g099860.2 |
| Solyc12g099890.1 | Solyc12g099890.2 |
| Solyc12g099910.1 | Solyc12g099910.2 |
| Solyc12g099930.1 | Solyc12g099930.2 |
| Solyc12g099970.1 | Solyc12g099970.2 |
| Solyc12g099990.1 | Solyc12g099990.2 |
| Solyc12g100030.1 | Solyc12g100030.2 |
| Solyc12g100040.1 | Solyc12g100040.2 |
| Solyc12g100050.1 | Solyc12g100050.2 |
| Solyc12g100120.1 | Solyc12g100120.2 |
| Solyc12g100150.1 | Solyc12g100150.2 |
| Solyc12g100180.1 | Solyc12g100180.2 |

| | |
|---|---|
| Solyc12g100190.1 | Solyc12g100190.2 |
| Solyc12g100200.1 | Solyc12g100200.2 |
| Solyc12g100220.1 | Solyc12g100220.2 |
| Solyc12g100270.1 | Solyc12g100270.2 |
| Solyc12g100300.1 | Solyc12g100300.2 |
| Solyc12g100310.1 | Solyc12g100310.2 |
| Solyc12g100330.1 | Solyc12g100330.2 |
| Solyc01g005010.2 | Solyc01g005010.4 |
| Solyc01g005030.2 | Solyc01g005030.4 |
| Solyc01g005050.2 | Solyc01g005050.4 |
| Solyc01g005160.2 | Solyc01g005160.4 |
| Solyc01g005210.2 | Solyc01g005210.4 |
| Solyc01g005230.2 | Solyc01g005230.4 |
| Solyc01g005240.2 | Solyc01g005240.4 |
| Solyc01g005250.2 | Solyc01g005250.4 |
| Solyc01g005290.2 | Solyc01g005290.4 |
| Solyc01g005300.2 | Solyc01g005300.4 |
| Solyc01g005350.2 | Solyc01g005350.4 |
| Solyc01g005360.2 | Solyc01g005360.4 |
| Solyc01g005410.2 | Solyc01g005410.4 |
| Solyc01g005440.2 | Solyc01g005440.4 |
| Solyc01g005460.2 | Solyc01g005460.4 |
| Solyc01g005520.2 | Solyc01g005520.4 |
| Solyc01g005540.2 | Solyc01g005540.4 |
| Solyc01g005600.2 | Solyc01g005600.4 |
| Solyc01g005610.2 | Solyc01g005610.4 |
| Solyc01g005630.1 | Solyc01g005630.3 |
| Solyc01g005640.2 | Solyc01g005640.4 |
| Solyc01g005650.1 | Solyc01g005650.3 |
| Solyc01g005660.1 | Solyc01g005660.3 |
| Solyc01g005680.2 | Solyc01g005680.4 |
| Solyc01g005820.2 | Solyc01g005820.4 |
| Solyc01g005830.2 | Solyc01g005830.4 |
| Solyc01g005870.1 | Solyc01g005870.3 |
| Solyc01g005940.2 | Solyc01g005940.4 |
| Solyc01g005950.2 | Solyc01g005950.4 |
| Solyc01g006020.2 | Solyc01g006020.4 |
| Solyc01g006100.2 | Solyc01g006100.4 |
| Solyc01g006120.2 | Solyc01g006120.4 |
| Solyc01g006140.2 | Solyc01g006140.4 |
| Solyc01g006170.2 | Solyc01g006170.4 |
| Solyc01g006180.2 | Solyc01g006180.4 |
| Solyc01g006210.2 | Solyc01g006210.4 |

| | |
|---|---|
| Solyc01g006260.2 | Solyc01g006260.4 |
| Solyc01g006290.2 | Solyc01g006290.4 |
| Solyc01g006320.2 | Solyc01g006320.4 |
| Solyc01g006330.2 | Solyc01g006330.4 |
| Solyc01g006350.2 | Solyc01g006350.4 |
| Solyc01g006360.2 | Solyc01g006360.4 |
| Solyc01g006400.2 | Solyc01g006400.4 |
| Solyc01g006410.2 | Solyc01g006410.4 |
| Solyc01g006450.2 | Solyc01g006450.4 |
| Solyc01g006460.1 | Solyc01g006460.3 |
| Solyc01g006490.2 | Solyc01g006490.4 |
| Solyc01g006500.2 | Solyc01g006500.4 |
| Solyc01g006540.2 | Solyc01g006540.4 |
| Solyc01g006580.2 | Solyc01g006580.4 |
| Solyc01g006590.1 | Solyc01g006590.3 |
| Solyc01g006600.1 | Solyc01g006600.3 |
| Solyc01g006660.1 | Solyc01g006660.3 |
| Solyc01g006680.2 | Solyc01g006680.4 |
| Solyc01g006690.2 | Solyc01g006690.4 |
| Solyc01g006710.2 | Solyc01g006710.4 |
| Solyc01g006720.2 | Solyc01g006720.4 |
| Solyc01g006730.2 | Solyc01g006730.4 |
| Solyc01g006740.2 | Solyc01g006740.4 |
| Solyc01g006780.2 | Solyc01g006780.4 |
| Solyc01g006800.2 | Solyc01g006800.4 |
| Solyc01g006810.2 | Solyc01g006810.4 |
| Solyc01g006870.2 | Solyc01g006870.4 |
| Solyc01g006880.2 | Solyc01g006880.4 |
| Solyc01g006910.2 | Solyc01g006910.4 |
| Solyc01g006920.2 | Solyc01g006920.4 |
| Solyc01g006940.2 | Solyc01g006940.4 |
| Solyc01g006970.2 | Solyc01g006970.4 |
| Solyc01g006980.2 | Solyc01g006980.4 |
| Solyc01g006990.2 | Solyc01g006990.4 |
| Solyc01g007000.2 | Solyc01g007000.4 |
| Solyc01g007020.2 | Solyc01g007020.4 |
| Solyc01g007040.2 | Solyc01g007040.4 |
| Solyc01g007050.1 | Solyc01g007050.3 |
| Solyc01g007160.2 | Solyc01g007160.4 |
| Solyc01g007170.2 | Solyc01g007170.4 |
| Solyc01g007220.2 | Solyc01g007220.4 |
| Solyc01g007230.2 | Solyc01g007230.4 |
| Solyc01g007250.2 | Solyc01g007250.4 |

| | |
|---|---|
| Solyc01g007740.2 | Solyc01g007740.4 |
| Solyc01g007780.2 | Solyc01g007780.4 |
| Solyc01g007880.2 | Solyc01g007880.4 |
| Solyc01g007890.1 | Solyc01g007890.3 |
| Solyc01g007900.2 | Solyc01g007900.4 |
| Solyc01g007930.2 | Solyc01g007930.4 |
| Solyc01g007940.2 | Solyc01g007940.4 |
| Solyc01g007950.2 | Solyc01g007950.4 |
| Solyc01g007980.2 | Solyc01g007980.4 |
| Solyc01g007990.2 | Solyc01g007990.4 |
| Solyc01g008020.2 | Solyc01g008020.4 |
| Solyc01g008050.2 | Solyc01g008050.4 |
| Solyc01g008090.2 | Solyc01g008090.4 |
| Solyc01g008110.2 | Solyc01g008110.4 |
| Solyc01g008120.2 | Solyc01g008120.4 |
| Solyc01g008130.2 | Solyc01g008130.4 |
| Solyc01g008140.2 | Solyc01g008140.4 |
| Solyc01g008160.2 | Solyc01g008160.4 |
| Solyc01g008190.2 | Solyc01g008190.4 |
| Solyc01g008230.2 | Solyc01g008230.4 |
| Solyc01g008240.2 | Solyc01g008240.4 |
| Solyc01g008260.2 | Solyc01g008260.4 |
| Solyc01g008280.2 | Solyc01g008280.4 |
| Solyc01g008310.2 | Solyc01g008310.4 |
| Solyc01g008330.2 | Solyc01g008330.4 |
| Solyc01g008390.1 | Solyc01g008390.3 |
| Solyc01g008410.1 | Solyc01g008410.3 |
| Solyc01g008480.2 | Solyc01g008480.4 |
| Solyc01g008490.2 | Solyc01g008490.4 |
| Solyc01g008500.2 | Solyc01g008500.4 |
| Solyc01g008600.2 | Solyc01g008600.4 |
| Solyc01g008610.2 | Solyc01g008610.4 |
| Solyc01g008620.2 | Solyc01g008620.4 |
| Solyc01g008640.2 | Solyc01g008640.4 |
| Solyc01g008660.2 | Solyc01g008660.4 |
| Solyc01g008670.2 | Solyc01g008670.4 |
| Solyc01g008680.2 | Solyc01g008680.4 |
| Solyc01g008760.2 | Solyc01g008760.4 |
| Solyc01g008790.2 | Solyc01g008790.4 |
| Solyc01g008800.1 | Solyc01g008800.3 |
| Solyc01g008880.1 | Solyc01g008880.3 |
| Solyc01g008890.2 | Solyc01g008890.4 |
| Solyc01g008900.2 | Solyc01g008900.4 |

| | |
|---|---|
| Solyc01g008920.2 | Solyc01g008920.4 |
| Solyc01g008950.2 | Solyc01g008950.4 |
| Solyc01g008970.2 | Solyc01g008970.4 |
| Solyc01g008990.2 | Solyc01g008990.4 |
| Solyc01g009020.2 | Solyc01g009020.4 |
| Solyc01g009040.2 | Solyc01g009040.4 |
| Solyc01g009080.2 | Solyc01g009080.4 |
| Solyc01g009100.2 | Solyc01g009100.4 |
| Solyc01g009170.2 | Solyc01g009170.4 |
| Solyc01g009180.2 | Solyc01g009180.4 |
| Solyc01g009190.1 | Solyc01g009190.3 |
| Solyc01g009200.2 | Solyc01g009200.4 |
| Solyc01g009230.2 | Solyc01g009230.4 |
| Solyc01g009240.2 | Solyc01g009240.4 |
| Solyc01g009280.2 | Solyc01g009280.4 |
| Solyc01g009290.2 | Solyc01g009290.4 |
| Solyc01g009340.2 | Solyc01g009340.4 |
| Solyc01g009390.2 | Solyc01g009390.4 |
| Solyc01g009420.2 | Solyc01g009420.4 |
| Solyc01g009440.1 | Solyc01g009440.3 |
| Solyc01g009480.2 | Solyc01g009480.4 |
| Solyc01g009500.2 | Solyc01g009500.4 |
| Solyc01g009520.2 | Solyc01g009520.4 |
| Solyc01g009600.2 | Solyc01g009600.4 |
| Solyc01g009640.1 | Solyc01g009640.3 |
| Solyc01g009660.1 | Solyc01g009660.3 |
| Solyc01g009670.2 | Solyc01g009670.4 |
| Solyc01g009690.1 | Solyc01g009690.3 |
| Solyc01g009700.1 | Solyc01g009700.3 |
| Solyc01g009730.1 | Solyc01g009730.3 |
| Solyc01g009820.2 | Solyc01g009820.4 |
| Solyc01g009840.1 | Solyc01g009840.3 |
| Solyc01g009920.1 | Solyc01g009920.3 |
| Solyc01g009930.1 | Solyc01g009930.3 |
| Solyc01g009960.2 | Solyc01g009960.4 |
| Solyc01g010130.2 | Solyc01g010130.4 |
| Solyc01g010250.2 | Solyc01g010250.4 |
| Solyc01g010290.2 | Solyc01g010290.4 |
| Solyc01g010310.2 | Solyc01g010310.4 |
| Solyc01g010350.2 | Solyc01g010350.4 |
| Solyc01g010430.2 | Solyc01g010430.4 |
| Solyc01g010440.2 | Solyc01g010440.4 |
| Solyc01g010460.2 | Solyc01g010460.4 |

| | |
|---|---|
| Solyc01g010490.2 | Solyc01g010490.4 |
| Solyc01g010500.1 | Solyc01g010500.3 |
| Solyc01g010510.2 | Solyc01g010510.4 |
| Solyc01g010570.2 | Solyc01g010570.4 |
| Solyc01g010600.2 | Solyc01g010600.4 |
| Solyc01g010670.1 | Solyc01g010670.3 |
| Solyc01g010710.2 | Solyc01g010710.4 |
| Solyc01g010720.2 | Solyc01g010720.4 |
| Solyc01g010740.2 | Solyc01g010740.4 |
| Solyc01g010750.2 | Solyc01g010750.4 |
| Solyc01g010780.2 | Solyc01g010780.4 |
| Solyc01g010810.2 | Solyc01g010810.4 |
| Solyc01g010830.2 | Solyc01g010830.4 |
| Solyc01g010890.2 | Solyc01g010890.4 |
| Solyc01g010930.2 | Solyc01g010930.4 |
| Solyc01g010960.1 | Solyc01g010960.3 |
| Solyc01g011090.2 | Solyc01g011090.4 |
| Solyc01g011220.1 | Solyc01g011220.3 |
| Solyc01g011330.1 | Solyc01g011330.3 |
| Solyc01g011430.2 | Solyc01g011430.4 |
| Solyc01g011460.1 | Solyc01g011460.3 |
| Solyc01g011510.2 | Solyc01g011510.4 |
| Solyc01g012670.1 | Solyc01g012670.3 |
| Solyc01g012700.2 | Solyc01g012700.4 |
| Solyc01g012750.1 | Solyc01g012750.3 |
| Solyc01g013890.1 | Solyc01g013890.3 |
| Solyc01g014060.1 | Solyc01g014060.3 |
| Solyc01g014220.1 | Solyc01g014220.3 |
| Solyc01g014230.2 | Solyc01g014230.4 |
| Solyc01g014320.2 | Solyc01g014320.4 |
| Solyc01g014490.2 | Solyc01g014490.4 |
| Solyc01g014540.1 | Solyc01g014540.3 |
| Solyc01g014550.1 | Solyc01g014550.3 |
| Solyc01g014560.2 | Solyc01g014560.4 |
| Solyc01g014720.1 | Solyc01g014720.3 |
| Solyc01g014850.2 | Solyc01g014850.4 |
| Solyc01g014980.2 | Solyc01g014980.4 |
| Solyc01g014990.1 | Solyc01g014990.3 |
| Solyc01g015230.1 | Solyc01g015230.3 |
| Solyc01g016380.2 | Solyc01g016380.4 |
| Solyc01g016390.2 | Solyc01g016390.4 |
| Solyc01g016410.2 | Solyc01g016410.4 |
| Solyc01g016460.2 | Solyc01g016460.4 |

| | |
|---|---|
| Solyc01g016540.1 | Solyc01g016540.3 |
| Solyc01g016640.1 | Solyc01g016640.3 |
| Solyc01g016680.1 | Solyc01g016680.3 |
| Solyc01g017180.1 | Solyc01g017180.3 |
| Solyc01g017330.1 | Solyc01g017330.3 |
| Solyc01g017360.1 | Solyc01g017360.3 |
| Solyc01g017410.1 | Solyc01g017410.3 |
| Solyc01g017460.1 | Solyc01g017460.3 |
| Solyc01g017470.1 | Solyc01g017470.3 |
| Solyc01g017560.1 | Solyc01g017560.3 |
| Solyc01g017640.1 | Solyc01g017640.3 |
| Solyc01g017790.1 | Solyc01g017790.3 |
| Solyc01g017820.1 | Solyc01g017820.3 |
| Solyc01g017950.1 | Solyc01g017950.3 |
| Solyc01g018020.1 | Solyc01g018020.3 |
| Solyc01g018070.1 | Solyc01g018070.3 |
| Solyc01g020100.1 | Solyc01g020100.3 |
| Solyc01g020120.1 | Solyc01g020120.3 |
| Solyc01g020220.1 | Solyc01g020220.3 |
| Solyc01g020300.2 | Solyc01g020300.4 |
| Solyc01g020460.1 | Solyc01g020460.3 |
| Solyc01g020470.1 | Solyc01g020470.3 |
| Solyc01g020520.2 | Solyc01g020520.4 |
| Solyc01g020570.1 | Solyc01g020570.3 |
| Solyc01g021600.2 | Solyc01g021600.4 |
| Solyc01g021620.1 | Solyc01g021620.3 |
| Solyc01g021700.1 | Solyc01g021700.3 |
| Solyc01g028800.1 | Solyc01g028800.3 |
| Solyc01g028930.2 | Solyc01g028930.4 |
| Solyc01g034150.2 | Solyc01g034150.4 |
| Solyc01g044300.1 | Solyc01g044300.3 |
| Solyc01g044360.2 | Solyc01g044360.4 |
| Solyc01g044370.1 | Solyc01g044370.3 |
| Solyc01g044420.2 | Solyc01g044420.4 |
| Solyc01g044490.2 | Solyc01g044490.4 |
| Solyc01g044550.1 | Solyc01g044550.3 |
| Solyc01g049690.2 | Solyc01g049690.4 |
| Solyc01g049810.2 | Solyc01g049810.4 |
| Solyc01g049910.2 | Solyc01g049910.4 |
| Solyc01g050040.2 | Solyc01g050040.4 |
| Solyc01g050070.2 | Solyc01g050070.4 |
| Solyc01g056340.2 | Solyc01g056340.4 |
| Solyc01g056360.2 | Solyc01g056360.4 |

| | |
|---|---|
| Solyc01g056450.1 | Solyc01g056450.3 |
| Solyc01g056580.2 | Solyc01g056580.4 |
| Solyc01g056620.2 | Solyc01g056620.4 |
| Solyc01g056690.2 | Solyc01g056690.4 |
| Solyc01g056720.2 | Solyc01g056720.4 |
| Solyc01g056730.1 | Solyc01g056730.3 |
| Solyc01g056810.2 | Solyc01g056810.4 |
| Solyc01g056880.1 | Solyc01g056880.3 |
| Solyc01g057090.2 | Solyc01g057090.4 |
| Solyc01g057200.2 | Solyc01g057200.4 |
| Solyc01g057260.2 | Solyc01g057260.4 |
| Solyc01g057270.2 | Solyc01g057270.4 |
| Solyc01g057310.2 | Solyc01g057310.4 |
| Solyc01g057320.1 | Solyc01g057320.3 |
| Solyc01g057520.2 | Solyc01g057520.4 |
| Solyc01g057590.1 | Solyc01g057590.3 |
| Solyc01g057680.2 | Solyc01g057680.4 |
| Solyc01g057740.1 | Solyc01g057740.3 |
| Solyc01g057760.2 | Solyc01g057760.4 |
| Solyc01g057850.2 | Solyc01g057850.4 |
| Solyc01g057960.1 | Solyc01g057960.3 |
| Solyc01g058020.2 | Solyc01g058020.4 |
| Solyc01g058170.2 | Solyc01g058170.4 |
| Solyc01g058190.2 | Solyc01g058190.4 |
| Solyc01g058210.1 | Solyc01g058210.3 |
| Solyc01g058220.1 | Solyc01g058220.3 |
| Solyc01g058260.2 | Solyc01g058260.4 |
| Solyc01g058280.2 | Solyc01g058280.4 |
| Solyc01g058400.1 | Solyc01g058400.3 |
| Solyc01g058520.2 | Solyc01g058520.4 |
| Solyc01g058530.2 | Solyc01g058530.4 |
| Solyc01g058720.2 | Solyc01g058720.4 |
| Solyc01g058730.2 | Solyc01g058730.4 |
| Solyc01g059760.1 | Solyc01g059760.3 |
| Solyc01g059840.1 | Solyc01g059840.3 |
| Solyc01g059900.2 | Solyc01g059900.4 |
| Solyc01g059920.2 | Solyc01g059920.4 |
| Solyc01g060010.1 | Solyc01g060010.3 |
| Solyc01g060020.2 | Solyc01g060020.4 |
| Solyc01g060050.1 | Solyc01g060050.3 |
| Solyc01g060080.2 | Solyc01g060080.4 |
| Solyc01g060140.2 | Solyc01g060140.4 |
| Solyc01g060170.2 | Solyc01g060170.4 |

| | |
|---|---|
| Solyc01g060180.2 | Solyc01g060180.4 |
| Solyc01g060200.1 | Solyc01g060200.3 |
| Solyc01g060210.2 | Solyc01g060210.4 |
| Solyc01g060240.2 | Solyc01g060240.4 |
| Solyc01g060280.2 | Solyc01g060280.4 |
| Solyc01g060310.1 | Solyc01g060310.3 |
| Solyc01g060410.2 | Solyc01g060410.4 |
| Solyc01g060490.2 | Solyc01g060490.4 |
| Solyc01g065490.2 | Solyc01g065490.4 |
| Solyc01g065560.2 | Solyc01g065560.4 |
| Solyc01g065580.2 | Solyc01g065580.4 |
| Solyc01g065640.1 | Solyc01g065640.3 |
| Solyc01g065720.2 | Solyc01g065720.4 |
| Solyc01g065840.2 | Solyc01g065840.4 |
| Solyc01g065900.1 | Solyc01g065900.3 |
| Solyc01g065980.2 | Solyc01g065980.4 |
| Solyc01g066020.1 | Solyc01g066020.3 |
| Solyc01g066060.2 | Solyc01g066060.4 |
| Solyc01g066080.1 | Solyc01g066080.3 |
| Solyc01g066090.1 | Solyc01g066090.3 |
| Solyc01g066120.1 | Solyc01g066120.3 |
| Solyc01g066310.2 | Solyc01g066310.4 |
| Solyc01g066340.2 | Solyc01g066340.4 |
| Solyc01g066380.2 | Solyc01g066380.4 |
| Solyc01g066390.2 | Solyc01g066390.4 |
| Solyc01g066510.1 | Solyc01g066510.3 |
| Solyc01g066610.2 | Solyc01g066610.4 |
| Solyc01g066620.2 | Solyc01g066620.4 |
| Solyc01g066700.2 | Solyc01g066700.4 |
| Solyc01g066710.2 | Solyc01g066710.4 |
| Solyc01g066730.2 | Solyc01g066730.4 |
| Solyc01g066810.2 | Solyc01g066810.4 |
| Solyc01g066820.2 | Solyc01g066820.4 |
| Solyc01g066850.2 | Solyc01g066850.4 |
| Solyc01g067000.2 | Solyc01g067000.4 |
| Solyc01g067020.2 | Solyc01g067020.4 |
| Solyc01g067090.2 | Solyc01g067090.4 |
| Solyc01g067100.2 | Solyc01g067100.4 |
| Solyc01g067120.2 | Solyc01g067120.4 |
| Solyc01g067130.2 | Solyc01g067130.4 |
| Solyc01g067160.2 | Solyc01g067160.4 |
| Solyc01g067210.2 | Solyc01g067210.4 |
| Solyc01g067230.2 | Solyc01g067230.4 |

| | |
|---|---|
| Solyc01g067270.2 | Solyc01g067270.4 |
| Solyc01g067340.1 | Solyc01g067340.3 |
| Solyc01g067350.2 | Solyc01g067350.4 |
| Solyc01g067380.2 | Solyc01g067380.4 |
| Solyc01g067390.2 | Solyc01g067390.4 |
| Solyc01g067410.1 | Solyc01g067410.3 |
| Solyc01g067420.2 | Solyc01g067420.4 |
| Solyc01g067440.2 | Solyc01g067440.4 |
| Solyc01g067460.1 | Solyc01g067460.3 |
| Solyc01g067480.2 | Solyc01g067480.4 |
| Solyc01g067490.2 | Solyc01g067490.4 |
| Solyc01g067690.2 | Solyc01g067690.4 |
| Solyc01g067720.2 | Solyc01g067720.4 |
| Solyc01g067880.1 | Solyc01g067880.3 |
| Solyc01g067890.2 | Solyc01g067890.4 |
| Solyc01g067930.2 | Solyc01g067930.4 |
| Solyc01g068000.2 | Solyc01g068000.4 |
| Solyc01g068020.2 | Solyc01g068020.4 |
| Solyc01g068060.2 | Solyc01g068060.4 |
| Solyc01g068080.2 | Solyc01g068080.4 |
| Solyc01g068100.2 | Solyc01g068100.4 |
| Solyc01g068120.2 | Solyc01g068120.4 |
| Solyc01g068140.2 | Solyc01g068140.4 |
| Solyc01g068160.2 | Solyc01g068160.4 |
| Solyc01g068200.2 | Solyc01g068200.4 |
| Solyc01g068260.2 | Solyc01g068260.4 |
| Solyc01g068290.2 | Solyc01g068290.4 |
| Solyc01g068300.1 | Solyc01g068300.3 |
| Solyc01g068320.1 | Solyc01g068320.3 |
| Solyc01g068330.2 | Solyc01g068330.4 |
| Solyc01g068360.2 | Solyc01g068360.4 |
| Solyc01g068370.2 | Solyc01g068370.4 |
| Solyc01g068410.2 | Solyc01g068410.4 |
| Solyc01g068420.2 | Solyc01g068420.4 |
| Solyc01g068430.1 | Solyc01g068430.3 |
| Solyc01g068450.2 | Solyc01g068450.4 |
| Solyc01g068460.2 | Solyc01g068460.4 |
| Solyc01g068480.2 | Solyc01g068480.4 |
| Solyc01g068490.2 | Solyc01g068490.4 |
| Solyc01g068500.2 | Solyc01g068500.4 |
| Solyc01g068580.2 | Solyc01g068580.4 |
| Solyc01g068590.2 | Solyc01g068590.4 |
| Solyc01g068620.2 | Solyc01g068620.4 |

| | |
|---|---|
| Solyc01g073680.2 | Solyc01g073680.4 |
| Solyc01g073730.2 | Solyc01g073730.4 |
| Solyc01g073740.2 | Solyc01g073740.4 |
| Solyc01g073750.2 | Solyc01g073750.4 |
| Solyc01g073820.2 | Solyc01g073820.4 |
| Solyc01g073870.2 | Solyc01g073870.4 |
| Solyc01g073900.2 | Solyc01g073900.4 |
| Solyc01g073910.2 | Solyc01g073910.4 |
| Solyc01g073950.2 | Solyc01g073950.4 |
| Solyc01g073960.2 | Solyc01g073960.4 |
| Solyc01g074010.2 | Solyc01g074010.4 |
| Solyc01g074050.1 | Solyc01g074050.3 |
| Solyc01g079060.2 | Solyc01g079060.4 |
| Solyc01g079090.2 | Solyc01g079090.4 |
| Solyc01g079110.2 | Solyc01g079110.4 |
| Solyc01g079150.2 | Solyc01g079150.4 |
| Solyc01g079160.2 | Solyc01g079160.4 |
| Solyc01g079180.2 | Solyc01g079180.4 |
| Solyc01g079200.2 | Solyc01g079200.4 |
| Solyc01g079220.2 | Solyc01g079220.4 |
| Solyc01g079230.2 | Solyc01g079230.4 |
| Solyc01g079260.2 | Solyc01g079260.4 |
| Solyc01g079300.2 | Solyc01g079300.4 |
| Solyc01g079320.2 | Solyc01g079320.4 |
| Solyc01g079360.2 | Solyc01g079360.4 |
| Solyc01g079370.2 | Solyc01g079370.4 |
| Solyc01g079390.2 | Solyc01g079390.4 |
| Solyc01g079420.2 | Solyc01g079420.4 |
| Solyc01g079470.2 | Solyc01g079470.4 |
| Solyc01g079500.2 | Solyc01g079500.4 |
| Solyc01g079530.2 | Solyc01g079530.4 |
| Solyc01g079600.2 | Solyc01g079600.4 |
| Solyc01g079620.2 | Solyc01g079620.4 |
| Solyc01g079690.2 | Solyc01g079690.4 |
| Solyc01g079790.2 | Solyc01g079790.4 |
| Solyc01g079800.2 | Solyc01g079800.4 |
| Solyc01g079810.2 | Solyc01g079810.4 |
| Solyc01g079870.2 | Solyc01g079870.4 |
| Solyc01g079930.1 | Solyc01g079930.3 |
| Solyc01g079940.2 | Solyc01g079940.4 |
| Solyc01g080030.1 | Solyc01g080030.3 |
| Solyc01g080050.2 | Solyc01g080050.4 |
| Solyc01g080070.2 | Solyc01g080070.4 |

| | |
|---|---|
| Solyc01g080160.2 | Solyc01g080160.4 |
| Solyc01g080240.2 | Solyc01g080240.4 |
| Solyc01g080300.2 | Solyc01g080300.4 |
| Solyc01g080310.2 | Solyc01g080310.4 |
| Solyc01g080320.1 | Solyc01g080320.3 |
| Solyc01g080330.2 | Solyc01g080330.4 |
| Solyc01g080380.2 | Solyc01g080380.4 |
| Solyc01g080430.2 | Solyc01g080430.4 |
| Solyc01g080450.2 | Solyc01g080450.4 |
| Solyc01g080480.2 | Solyc01g080480.4 |
| Solyc01g080550.1 | Solyc01g080550.3 |
| Solyc01g080570.2 | Solyc01g080570.4 |
| Solyc01g080580.2 | Solyc01g080580.4 |
| Solyc01g080600.2 | Solyc01g080600.4 |
| Solyc01g080700.2 | Solyc01g080700.4 |
| Solyc01g080720.2 | Solyc01g080720.4 |
| Solyc01g080850.2 | Solyc01g080850.4 |
| Solyc01g080880.2 | Solyc01g080880.4 |
| Solyc01g080900.2 | Solyc01g080900.4 |
| Solyc01g080930.2 | Solyc01g080930.4 |
| Solyc01g080960.2 | Solyc01g080960.4 |
| Solyc01g081010.2 | Solyc01g081010.4 |
| Solyc01g081030.2 | Solyc01g081030.4 |
| Solyc01g081040.2 | Solyc01g081040.4 |
| Solyc01g081080.1 | Solyc01g081080.3 |
| Solyc01g081100.1 | Solyc01g081100.3 |
| Solyc01g081110.2 | Solyc01g081110.4 |
| Solyc01g081160.2 | Solyc01g081160.4 |
| Solyc01g081170.1 | Solyc01g081170.3 |
| Solyc01g081190.2 | Solyc01g081190.4 |
| Solyc01g081210.2 | Solyc01g081210.4 |
| Solyc01g081290.2 | Solyc01g081290.4 |
| Solyc01g081300.2 | Solyc01g081300.4 |
| Solyc01g081320.2 | Solyc01g081320.4 |
| Solyc01g081340.2 | Solyc01g081340.4 |
| Solyc01g081370.2 | Solyc01g081370.4 |
| Solyc01g081380.2 | Solyc01g081380.4 |
| Solyc01g081390.2 | Solyc01g081390.4 |
| Solyc01g081410.2 | Solyc01g081410.4 |
| Solyc01g081420.1 | Solyc01g081420.3 |
| Solyc01g081430.2 | Solyc01g081430.4 |
| Solyc01g081470.2 | Solyc01g081470.4 |
| Solyc01g081500.2 | Solyc01g081500.4 |

| | |
|---|---|
| Solyc01g081510.2 | Solyc01g081510.4 |
| Solyc01g081530.1 | Solyc01g081530.3 |
| Solyc01g081540.2 | Solyc01g081540.4 |
| Solyc01g081590.2 | Solyc01g081590.4 |
| Solyc01g081630.2 | Solyc01g081630.4 |
| Solyc01g081640.2 | Solyc01g081640.4 |
| Solyc01g086680.2 | Solyc01g086680.4 |
| Solyc01g086700.2 | Solyc01g086700.4 |
| Solyc01g086710.2 | Solyc01g086710.4 |
| Solyc01g086750.2 | Solyc01g086750.4 |
| Solyc01g086760.2 | Solyc01g086760.4 |
| Solyc01g086800.2 | Solyc01g086800.4 |
| Solyc01g086820.2 | Solyc01g086820.4 |
| Solyc01g086880.1 | Solyc01g086880.3 |
| Solyc01g086900.2 | Solyc01g086900.4 |
| Solyc01g086930.2 | Solyc01g086930.4 |
| Solyc01g086950.2 | Solyc01g086950.4 |
| Solyc01g086970.2 | Solyc01g086970.4 |
| Solyc01g086980.2 | Solyc01g086980.4 |
| Solyc01g087060.2 | Solyc01g087060.4 |
| Solyc01g087070.2 | Solyc01g087070.4 |
| Solyc01g087130.2 | Solyc01g087130.4 |
| Solyc01g087140.2 | Solyc01g087140.4 |
| Solyc01g087190.2 | Solyc01g087190.4 |
| Solyc01g087260.2 | Solyc01g087260.4 |
| Solyc01g087330.2 | Solyc01g087330.4 |
| Solyc01g087360.1 | Solyc01g087360.3 |
| Solyc01g087400.2 | Solyc01g087400.4 |
| Solyc01g087410.2 | Solyc01g087410.4 |
| Solyc01g087420.2 | Solyc01g087420.4 |
| Solyc01g087460.1 | Solyc01g087460.3 |
| Solyc01g087490.2 | Solyc01g087490.4 |
| Solyc01g087510.1 | Solyc01g087510.3 |
| Solyc01g087540.2 | Solyc01g087540.4 |
| Solyc01g087650.2 | Solyc01g087650.4 |
| Solyc01g087680.2 | Solyc01g087680.4 |
| Solyc01g087690.1 | Solyc01g087690.3 |
| Solyc01g087720.2 | Solyc01g087720.4 |
| Solyc01g087730.2 | Solyc01g087730.4 |
| Solyc01g087750.2 | Solyc01g087750.4 |
| Solyc01g087770.2 | Solyc01g087770.4 |
| Solyc01g087870.2 | Solyc01g087870.4 |
| Solyc01g087910.2 | Solyc01g087910.4 |

| | |
|---|---|
| Solyc01g087940.1 | Solyc01g087940.3 |
| Solyc01g087960.1 | Solyc01g087960.3 |
| Solyc01g087980.2 | Solyc01g087980.4 |
| Solyc01g088000.2 | Solyc01g088000.4 |
| Solyc01g088030.2 | Solyc01g088030.4 |
| Solyc01g088100.2 | Solyc01g088100.4 |
| Solyc01g088160.2 | Solyc01g088160.4 |
| Solyc01g088240.2 | Solyc01g088240.4 |
| Solyc01g088250.2 | Solyc01g088250.4 |
| Solyc01g088350.2 | Solyc01g088350.4 |
| Solyc01g088360.2 | Solyc01g088360.4 |
| Solyc01g088380.1 | Solyc01g088380.3 |
| Solyc01g088400.2 | Solyc01g088400.4 |
| Solyc01g088430.2 | Solyc01g088430.4 |
| Solyc01g088590.2 | Solyc01g088590.5 |
| Solyc01g088600.2 | Solyc01g088600.4 |
| Solyc01g088610.2 | Solyc01g088610.4 |
| Solyc01g088630.1 | Solyc01g088630.3 |
| Solyc01g088640.2 | Solyc01g088640.4 |
| Solyc01g088650.2 | Solyc01g088650.4 |
| Solyc01g088670.1 | Solyc01g088670.3 |
| Solyc01g088700.2 | Solyc01g088700.4 |
| Solyc01g088740.2 | Solyc01g088740.4 |
| Solyc01g088760.2 | Solyc01g088760.4 |
| Solyc01g088800.2 | Solyc01g088800.4 |
| Solyc01g088810.2 | Solyc01g088810.4 |
| Solyc01g089850.2 | Solyc01g089850.4 |
| Solyc01g089900.2 | Solyc01g089900.4 |
| Solyc01g089910.2 | Solyc01g089910.4 |
| Solyc01g089980.2 | Solyc01g089980.4 |
| Solyc01g089990.1 | Solyc01g089990.3 |
| Solyc01g090020.1 | Solyc01g090020.3 |
| Solyc01g090050.2 | Solyc01g090050.4 |
| Solyc01g090080.1 | Solyc01g090080.3 |
| Solyc01g090090.1 | Solyc01g090090.3 |
| Solyc01g090100.1 | Solyc01g090100.3 |
| Solyc01g090130.1 | Solyc01g090130.3 |
| Solyc01g090140.1 | Solyc01g090140.3 |
| Solyc01g090160.2 | Solyc01g090160.4 |
| Solyc01g090170.2 | Solyc01g090170.4 |
| Solyc01g090180.2 | Solyc01g090180.4 |
| Solyc01g090200.2 | Solyc01g090200.4 |
| Solyc01g090210.2 | Solyc01g090210.4 |

| | |
|---|---|
| Solyc01g090220.2 | Solyc01g090220.4 |
| Solyc01g090270.2 | Solyc01g090270.4 |
| Solyc01g090280.2 | Solyc01g090280.4 |
| Solyc01g090290.2 | Solyc01g090290.4 |
| Solyc01g090320.2 | Solyc01g090320.4 |
| Solyc01g090370.1 | Solyc01g090370.3 |
| Solyc01g090420.2 | Solyc01g090420.4 |
| Solyc01g090480.2 | Solyc01g090480.4 |
| Solyc01g090530.1 | Solyc01g090530.3 |
| Solyc01g090560.2 | Solyc01g090560.4 |
| Solyc01g090600.2 | Solyc01g090600.4 |
| Solyc01g090630.2 | Solyc01g090630.4 |
| Solyc01g090690.2 | Solyc01g090690.4 |
| Solyc01g090760.2 | Solyc01g090760.4 |
| Solyc01g090790.2 | Solyc01g090790.4 |
| Solyc01g090950.2 | Solyc01g090950.4 |
| Solyc01g091000.2 | Solyc01g091000.4 |
| Solyc01g091050.2 | Solyc01g091050.4 |
| Solyc01g091090.2 | Solyc01g091090.4 |
| Solyc01g091130.2 | Solyc01g091130.4 |
| Solyc01g091140.2 | Solyc01g091140.4 |
| Solyc01g091180.2 | Solyc01g091180.4 |
| Solyc01g091200.2 | Solyc01g091200.4 |
| Solyc01g091240.2 | Solyc01g091240.4 |
| Solyc01g091290.2 | Solyc01g091290.4 |
| Solyc01g091330.2 | Solyc01g091330.4 |
| Solyc01g091350.2 | Solyc01g091350.4 |
| Solyc01g091370.2 | Solyc01g091370.4 |
| Solyc01g091380.2 | Solyc01g091380.4 |
| Solyc01g091440.2 | Solyc01g091440.4 |
| Solyc01g091480.2 | Solyc01g091480.4 |
| Solyc01g091510.2 | Solyc01g091510.4 |
| Solyc01g091530.2 | Solyc01g091530.4 |
| Solyc01g091540.1 | Solyc01g091540.3 |
| Solyc01g091600.2 | Solyc01g091600.4 |
| Solyc01g091700.2 | Solyc01g091700.4 |
| Solyc01g091710.2 | Solyc01g091710.4 |
| Solyc01g091740.2 | Solyc01g091740.4 |
| Solyc01g091770.2 | Solyc01g091770.4 |
| Solyc01g091780.2 | Solyc01g091780.4 |
| Solyc01g091840.2 | Solyc01g091840.4 |
| Solyc01g091860.2 | Solyc01g091860.4 |
| Solyc01g091950.2 | Solyc01g091950.4 |

| | |
|---|---|
| Solyc01g094060.2 | Solyc01g094060.4 |
| Solyc01g094080.2 | Solyc01g094080.4 |
| Solyc01g094090.2 | Solyc01g094090.4 |
| Solyc01g094180.2 | Solyc01g094180.4 |
| Solyc01g094200.2 | Solyc01g094200.4 |
| Solyc01g094230.2 | Solyc01g094230.4 |
| Solyc01g094260.2 | Solyc01g094260.4 |
| Solyc01g094350.2 | Solyc01g094350.4 |
| Solyc01g094470.2 | Solyc01g094470.4 |
| Solyc01g094480.2 | Solyc01g094480.4 |
| Solyc01g094530.2 | Solyc01g094530.4 |
| Solyc01g094540.2 | Solyc01g094540.4 |
| Solyc01g094560.2 | Solyc01g094560.4 |
| Solyc01g094590.2 | Solyc01g094590.4 |
| Solyc01g094610.2 | Solyc01g094610.4 |
| Solyc01g094640.2 | Solyc01g094640.4 |
| Solyc01g094650.2 | Solyc01g094650.4 |
| Solyc01g094670.2 | Solyc01g094670.4 |
| Solyc01g094680.2 | Solyc01g094680.4 |
| Solyc01g094690.2 | Solyc01g094690.4 |
| Solyc01g094700.2 | Solyc01g094700.4 |
| Solyc01g094750.2 | Solyc01g094750.4 |
| Solyc01g094800.2 | Solyc01g094800.4 |
| Solyc01g094830.2 | Solyc01g094830.4 |
| Solyc01g094850.1 | Solyc01g094850.3 |
| Solyc01g094860.2 | Solyc01g094860.4 |
| Solyc01g094870.1 | Solyc01g094870.3 |
| Solyc01g094890.2 | Solyc01g094890.4 |
| Solyc01g094940.2 | Solyc01g094940.4 |
| Solyc01g094970.2 | Solyc01g094970.4 |
| Solyc01g095010.2 | Solyc01g095010.4 |
| Solyc01g095060.2 | Solyc01g095060.4 |
| Solyc01g095070.2 | Solyc01g095070.4 |
| Solyc01g095100.2 | Solyc01g095100.4 |
| Solyc01g095110.2 | Solyc01g095110.4 |
| Solyc01g095130.2 | Solyc01g095130.4 |
| Solyc01g095140.2 | Solyc01g095140.4 |
| Solyc01g095230.2 | Solyc01g095230.4 |
| Solyc01g095280.2 | Solyc01g095280.4 |
| Solyc01g095290.2 | Solyc01g095290.4 |
| Solyc01g095320.2 | Solyc01g095320.4 |
| Solyc01g095400.2 | Solyc01g095400.4 |
| Solyc01g095450.2 | Solyc01g095450.4 |

| | |
|---|---|
| Solyc01g095480.2 | Solyc01g095480.4 |
| Solyc01g095600.2 | Solyc01g095600.4 |
| Solyc01g095610.2 | Solyc01g095610.4 |
| Solyc01g095650.2 | Solyc01g095650.4 |
| Solyc01g095670.2 | Solyc01g095670.4 |
| Solyc01g095730.2 | Solyc01g095730.4 |
| Solyc01g095740.2 | Solyc01g095740.4 |
| Solyc01g095790.2 | Solyc01g095790.4 |
| Solyc01g095870.1 | Solyc01g095870.3 |
| Solyc01g095890.2 | Solyc01g095890.4 |
| Solyc01g095900.2 | Solyc01g095900.4 |
| Solyc01g095920.2 | Solyc01g095920.4 |
| Solyc01g095930.2 | Solyc01g095930.4 |
| Solyc01g095960.2 | Solyc01g095960.4 |
| Solyc01g096010.2 | Solyc01g096010.4 |
| Solyc01g096020.2 | Solyc01g096020.4 |
| Solyc01g096030.2 | Solyc01g096030.4 |
| Solyc01g096040.2 | Solyc01g096040.4 |
| Solyc01g096050.2 | Solyc01g096050.4 |
| Solyc01g096060.2 | Solyc01g096060.4 |
| Solyc01g096150.2 | Solyc01g096150.4 |
| Solyc01g096160.2 | Solyc01g096160.4 |
| Solyc01g096170.2 | Solyc01g096170.4 |
| Solyc01g096180.2 | Solyc01g096180.4 |
| Solyc01g096200.2 | Solyc01g096200.4 |
| Solyc01g096210.2 | Solyc01g096210.4 |
| Solyc01g096230.2 | Solyc01g096230.4 |
| Solyc01g096290.2 | Solyc01g096290.4 |
| Solyc01g096300.2 | Solyc01g096300.4 |
| Solyc01g096370.2 | Solyc01g096370.4 |
| Solyc01g096390.2 | Solyc01g096390.4 |
| Solyc01g096400.2 | Solyc01g096400.4 |
| Solyc01g096410.2 | Solyc01g096410.4 |
| Solyc01g096430.2 | Solyc01g096430.4 |
| Solyc01g096440.2 | Solyc01g096440.4 |
| Solyc01g096450.2 | Solyc01g096450.4 |
| Solyc01g096540.2 | Solyc01g096540.4 |
| Solyc01g096560.1 | Solyc01g096560.3 |
| Solyc01g096600.2 | Solyc01g096600.4 |
| Solyc01g096610.2 | Solyc01g096610.4 |
| Solyc01g096620.2 | Solyc01g096620.4 |
| Solyc01g096670.2 | Solyc01g096670.4 |
| Solyc01g096680.2 | Solyc01g096680.4 |

| | |
|---|---|
| Solyc01g096690.1 | Solyc01g096690.3 |
| Solyc01g096700.2 | Solyc01g096700.4 |
| Solyc01g096710.2 | Solyc01g096710.4 |
| Solyc01g096720.2 | Solyc01g096720.4 |
| Solyc01g096740.2 | Solyc01g096740.4 |
| Solyc01g096780.2 | Solyc01g096780.4 |
| Solyc01g096790.2 | Solyc01g096790.4 |
| Solyc01g096820.2 | Solyc01g096820.4 |
| Solyc01g096850.2 | Solyc01g096850.4 |
| Solyc01g096860.1 | Solyc01g096860.3 |
| Solyc01g096870.2 | Solyc01g096870.4 |
| Solyc01g096880.2 | Solyc01g096880.4 |
| Solyc01g096890.2 | Solyc01g096890.4 |
| Solyc01g096940.2 | Solyc01g096940.4 |
| Solyc01g096950.2 | Solyc01g096950.4 |
| Solyc01g096990.2 | Solyc01g096990.4 |
| Solyc01g097020.2 | Solyc01g097020.4 |
| Solyc01g097040.2 | Solyc01g097040.4 |
| Solyc01g097050.2 | Solyc01g097050.4 |
| Solyc01g097070.2 | Solyc01g097070.4 |
| Solyc01g097130.2 | Solyc01g097130.4 |
| Solyc01g097140.2 | Solyc01g097140.4 |
| Solyc01g097150.2 | Solyc01g097150.4 |
| Solyc01g097190.2 | Solyc01g097190.4 |
| Solyc01g097220.1 | Solyc01g097220.3 |
| Solyc01g097230.2 | Solyc01g097230.4 |
| Solyc01g097250.2 | Solyc01g097250.4 |
| Solyc01g097260.2 | Solyc01g097260.4 |
| Solyc01g097290.2 | Solyc01g097290.4 |
| Solyc01g097300.2 | Solyc01g097300.4 |
| Solyc01g097350.2 | Solyc01g097350.4 |
| Solyc01g097390.2 | Solyc01g097390.4 |
| Solyc01g097430.2 | Solyc01g097430.4 |
| Solyc01g097470.2 | Solyc01g097470.4 |
| Solyc01g097500.2 | Solyc01g097500.4 |
| Solyc01g097510.2 | Solyc01g097510.4 |
| Solyc01g097520.2 | Solyc01g097520.4 |
| Solyc01g097530.1 | Solyc01g097530.3 |
| Solyc01g097540.2 | Solyc01g097540.4 |
| Solyc01g097550.2 | Solyc01g097550.4 |
| Solyc01g097630.1 | Solyc01g097630.3 |
| Solyc01g097670.2 | Solyc01g097670.4 |
| Solyc01g097730.2 | Solyc01g097730.4 |

| | |
|---|---|
| Solyc01g097770.2 | Solyc01g097770.4 |
| Solyc01g097820.2 | Solyc01g097820.4 |
| Solyc01g097880.2 | Solyc01g097880.4 |
| Solyc01g097970.2 | Solyc01g097970.4 |
| Solyc01g098010.2 | Solyc01g098010.4 |
| Solyc01g098030.2 | Solyc01g098030.4 |
| Solyc01g098110.2 | Solyc01g098110.4 |
| Solyc01g098140.2 | Solyc01g098140.4 |
| Solyc01g098240.1 | Solyc01g098240.3 |
| Solyc01g098250.2 | Solyc01g098250.4 |
| Solyc01g098280.2 | Solyc01g098280.4 |
| Solyc01g098310.1 | Solyc01g098310.3 |
| Solyc01g098320.2 | Solyc01g098320.4 |
| Solyc01g098380.2 | Solyc01g098380.4 |
| Solyc01g098410.2 | Solyc01g098410.4 |
| Solyc01g098420.1 | Solyc01g098420.3 |
| Solyc01g098440.1 | Solyc01g098440.3 |
| Solyc01g098460.2 | Solyc01g098460.4 |
| Solyc01g098480.1 | Solyc01g098480.3 |
| Solyc01g098610.2 | Solyc01g098610.4 |
| Solyc01g098670.1 | Solyc01g098670.3 |
| Solyc01g098700.2 | Solyc01g098700.4 |
| Solyc01g098740.2 | Solyc01g098740.4 |
| Solyc01g098760.2 | Solyc01g098760.4 |
| Solyc01g098800.2 | Solyc01g098800.4 |
| Solyc01g098810.2 | Solyc01g098810.4 |
| Solyc01g098920.2 | Solyc01g098920.4 |
| Solyc01g098950.1 | Solyc01g098950.3 |
| Solyc01g098960.1 | Solyc01g098960.3 |
| Solyc01g098970.1 | Solyc01g098970.3 |
| Solyc01g098980.2 | Solyc01g098980.4 |
| Solyc01g099030.2 | Solyc01g099030.4 |
| Solyc01g099040.2 | Solyc01g099040.4 |
| Solyc01g099060.1 | Solyc01g099060.3 |
| Solyc01g099080.2 | Solyc01g099080.4 |
| Solyc01g099100.2 | Solyc01g099100.4 |
| Solyc01g099110.2 | Solyc01g099110.4 |
| Solyc01g099120.2 | Solyc01g099120.4 |
| Solyc01g099140.2 | Solyc01g099140.4 |
| Solyc01g099160.2 | Solyc01g099160.4 |
| Solyc01g099170.2 | Solyc01g099170.4 |
| Solyc01g099190.2 | Solyc01g099190.4 |
| Solyc01g099240.2 | Solyc01g099240.4 |

| | |
|---|---|
| Solyc01g099250.1 | Solyc01g099250.3 |
| Solyc01g099310.2 | Solyc01g099310.4 |
| Solyc01g099320.2 | Solyc01g099320.4 |
| Solyc01g099380.2 | Solyc01g099380.4 |
| Solyc01g099400.2 | Solyc01g099400.4 |
| Solyc01g099420.1 | Solyc01g099420.3 |
| Solyc01g099440.2 | Solyc01g099440.4 |
| Solyc01g099450.2 | Solyc01g099450.4 |
| Solyc01g099460.1 | Solyc01g099460.3 |
| Solyc01g099480.2 | Solyc01g099480.4 |
| Solyc01g099570.2 | Solyc01g099570.4 |
| Solyc01g099580.1 | Solyc01g099580.3 |
| Solyc01g099590.2 | Solyc01g099590.4 |
| Solyc01g099600.2 | Solyc01g099600.4 |
| Solyc01g099630.2 | Solyc01g099630.4 |
| Solyc01g099640.2 | Solyc01g099640.4 |
| Solyc01g099660.2 | Solyc01g099660.4 |
| Solyc01g099710.2 | Solyc01g099710.4 |
| Solyc01g099740.2 | Solyc01g099740.4 |
| Solyc01g099800.2 | Solyc01g099800.4 |
| Solyc01g099870.1 | Solyc01g099870.3 |
| Solyc01g099880.2 | Solyc01g099880.4 |
| Solyc01g099900.2 | Solyc01g099900.4 |
| Solyc01g099910.2 | Solyc01g099910.4 |
| Solyc01g099920.2 | Solyc01g099920.4 |
| Solyc01g099960.2 | Solyc01g099960.4 |
| Solyc01g099970.2 | Solyc01g099970.4 |
| Solyc01g099980.2 | Solyc01g099980.4 |
| Solyc01g100020.2 | Solyc01g100020.4 |
| Solyc01g100060.2 | Solyc01g100060.4 |
| Solyc01g100100.2 | Solyc01g100100.4 |
| Solyc01g100110.2 | Solyc01g100110.4 |
| Solyc01g100120.2 | Solyc01g100120.4 |
| Solyc01g100140.2 | Solyc01g100140.4 |
| Solyc01g100190.2 | Solyc01g100190.4 |
| Solyc01g100260.2 | Solyc01g100260.4 |
| Solyc01g100270.2 | Solyc01g100270.4 |
| Solyc01g100280.2 | Solyc01g100280.4 |
| Solyc01g100290.2 | Solyc01g100290.4 |
| Solyc01g100340.2 | Solyc01g100340.4 |
| Solyc01g100360.2 | Solyc01g100360.4 |
| Solyc01g100380.2 | Solyc01g100380.4 |
| Solyc01g100440.1 | Solyc01g100440.3 |

| | |
|---|---|
| Solyc01g100490.2 | Solyc01g100490.4 |
| Solyc01g100510.2 | Solyc01g100510.4 |
| Solyc01g100540.2 | Solyc01g100540.4 |
| Solyc01g100570.2 | Solyc01g100570.4 |
| Solyc01g100600.2 | Solyc01g100600.4 |
| Solyc01g100620.1 | Solyc01g100620.3 |
| Solyc01g100630.1 | Solyc01g100630.3 |
| Solyc01g100640.2 | Solyc01g100640.4 |
| Solyc01g100660.2 | Solyc01g100660.4 |
| Solyc01g100740.2 | Solyc01g100740.4 |
| Solyc01g100880.2 | Solyc01g100880.4 |
| Solyc01g100890.2 | Solyc01g100890.4 |
| Solyc01g100900.2 | Solyc01g100900.4 |
| Solyc01g100980.2 | Solyc01g100980.4 |
| Solyc01g101020.2 | Solyc01g101020.4 |
| Solyc01g101030.2 | Solyc01g101030.4 |
| Solyc01g101060.2 | Solyc01g101060.4 |
| Solyc01g101100.2 | Solyc01g101100.4 |
| Solyc01g101120.2 | Solyc01g101120.4 |
| Solyc01g101140.2 | Solyc01g101140.4 |
| Solyc01g101150.1 | Solyc01g101150.3 |
| Solyc01g101170.2 | Solyc01g101170.4 |
| Solyc01g101180.2 | Solyc01g101180.4 |
| Solyc01g101220.1 | Solyc01g101220.3 |
| Solyc01g101240.2 | Solyc01g101240.4 |
| Solyc01g102360.1 | Solyc01g102360.3 |
| Solyc01g102370.2 | Solyc01g102370.4 |
| Solyc01g102510.2 | Solyc01g102510.4 |
| Solyc01g102520.2 | Solyc01g102520.4 |
| Solyc01g102550.2 | Solyc01g102550.4 |
| Solyc01g102570.2 | Solyc01g102570.4 |
| Solyc01g102620.1 | Solyc01g102620.3 |
| Solyc01g102630.2 | Solyc01g102630.4 |
| Solyc01g102660.2 | Solyc01g102660.4 |
| Solyc01g102680.2 | Solyc01g102680.4 |
| Solyc01g102690.2 | Solyc01g102690.4 |
| Solyc01g102700.2 | Solyc01g102700.4 |
| Solyc01g102820.2 | Solyc01g102820.4 |
| Solyc01g102850.1 | Solyc01g102850.3 |
| Solyc01g102870.1 | Solyc01g102870.3 |
| Solyc01g102880.1 | Solyc01g102880.3 |
| Solyc01g102890.2 | Solyc01g102890.4 |
| Solyc01g102910.2 | Solyc01g102910.4 |

| | |
|---|---|
| Solyc01g103000.1 | Solyc01g103000.3 |
| Solyc01g103010.2 | Solyc01g103010.4 |
| Solyc01g103040.2 | Solyc01g103040.4 |
| Solyc01g103060.2 | Solyc01g103060.4 |
| Solyc01g103090.2 | Solyc01g103090.4 |
| Solyc01g103170.2 | Solyc01g103170.4 |
| Solyc01g103180.2 | Solyc01g103180.4 |
| Solyc01g103240.2 | Solyc01g103240.4 |
| Solyc01g103250.2 | Solyc01g103250.4 |
| Solyc01g103270.2 | Solyc01g103270.4 |
| Solyc01g103310.2 | Solyc01g103310.4 |
| Solyc01g103340.2 | Solyc01g103340.4 |
| Solyc01g103360.2 | Solyc01g103360.4 |
| Solyc01g103380.2 | Solyc01g103380.4 |
| Solyc01g103420.2 | Solyc01g103420.4 |
| Solyc01g103430.2 | Solyc01g103430.4 |
| Solyc01g103510.2 | Solyc01g103510.4 |
| Solyc01g103590.2 | Solyc01g103590.4 |
| Solyc01g103610.2 | Solyc01g103610.4 |
| Solyc01g103620.2 | Solyc01g103620.4 |
| Solyc01g103630.2 | Solyc01g103630.4 |
| Solyc01g103640.2 | Solyc01g103640.4 |
| Solyc01g103660.2 | Solyc01g103660.4 |
| Solyc01g103670.2 | Solyc01g103670.4 |
| Solyc01g103690.2 | Solyc01g103690.4 |
| Solyc01g103710.2 | Solyc01g103710.4 |
| Solyc01g103720.2 | Solyc01g103720.4 |
| Solyc01g103750.2 | Solyc01g103750.4 |
| Solyc01g103780.2 | Solyc01g103780.4 |
| Solyc01g103850.2 | Solyc01g103850.4 |
| Solyc01g103860.2 | Solyc01g103860.4 |
| Solyc01g103950.2 | Solyc01g103950.4 |
| Solyc01g103970.2 | Solyc01g103970.4 |
| Solyc01g103980.2 | Solyc01g103980.4 |
| Solyc01g104040.2 | Solyc01g104040.4 |
| Solyc01g104050.2 | Solyc01g104050.4 |
| Solyc01g104080.2 | Solyc01g104080.4 |
| Solyc01g104090.2 | Solyc01g104090.4 |
| Solyc01g104110.2 | Solyc01g104110.4 |
| Solyc01g104130.2 | Solyc01g104130.4 |
| Solyc01g104150.2 | Solyc01g104150.4 |
| Solyc01g104200.2 | Solyc01g104200.4 |
| Solyc01g104230.2 | Solyc01g104230.4 |

| | |
|---|---|
| Solyc01g104250.2 | Solyc01g104250.4 |
| Solyc01g104270.2 | Solyc01g104270.4 |
| Solyc01g104290.1 | Solyc01g104290.3 |
| Solyc01g104320.2 | Solyc01g104320.4 |
| Solyc01g104370.2 | Solyc01g104370.4 |
| Solyc01g104410.2 | Solyc01g104410.4 |
| Solyc01g104520.1 | Solyc01g104520.3 |
| Solyc01g104570.2 | Solyc01g104570.4 |
| Solyc01g104600.2 | Solyc01g104600.4 |
| Solyc01g104620.2 | Solyc01g104620.4 |
| Solyc01g104680.2 | Solyc01g104680.4 |
| Solyc01g104700.2 | Solyc01g104700.4 |
| Solyc01g104750.2 | Solyc01g104750.4 |
| Solyc01g104770.2 | Solyc01g104770.4 |
| Solyc01g104790.2 | Solyc01g104790.4 |
| Solyc01g104820.2 | Solyc01g104820.4 |
| Solyc01g104840.2 | Solyc01g104840.4 |
| Solyc01g104890.2 | Solyc01g104890.4 |
| Solyc01104930.1 | Solyc01g104930.3 |
| Solyc01g104950.2 | Solyc01g104950.4 |
| Solyc01g104980.2 | Solyc01g104980.4 |
| Solyc01g104990.2 | Solyc01g104990.4 |
| Solyc01g105120.2 | Solyc01g105120.4 |
| Solyc01g105150.2 | Solyc01g105150.4 |
| Solyc01g105160.2 | Solyc01g105160.4 |
| Solyc01g105190.2 | Solyc01g105190.4 |
| Solyc01g105230.2 | Solyc01g105230.4 |
| Solyc01g105270.2 | Solyc01g105270.4 |
| Solyc01g105300.2 | Solyc01g105300.4 |
| Solyc01g105320.2 | Solyc01g105320.4 |
| Solyc01g105340.2 | Solyc01g105340.4 |
| Solyc01g105350.1 | Solyc01g105350.3 |
| Solyc01g105370.2 | Solyc01g105370.4 |
| Solyc01g105410.2 | Solyc01g105410.4 |
| Solyc01g105420.2 | Solyc01g105420.4 |
| Solyc01g105430.2 | Solyc01g105430.4 |
| Solyc01g105450.2 | Solyc01g105450.4 |
| Solyc01g105460.2 | Solyc01g105460.4 |
| Solyc01g105470.2 | Solyc01g105470.4 |
| Solyc01g105490.2 | Solyc01g105490.4 |
| Solyc01g105500.2 | Solyc01g105500.4 |
| Solyc01g105540.2 | Solyc01g105540.4 |
| Solyc01g105620.2 | Solyc01g105620.4 |

| | |
|---|---|
| Solyc01g105630.2 | Solyc01g105630.4 |
| Solyc01g105650.2 | Solyc01g105650.4 |
| Solyc01g105660.2 | Solyc01g105660.4 |
| Solyc01g105680.2 | Solyc01g105680.4 |
| Solyc01g105780.2 | Solyc01g105780.4 |
| Solyc01g105790.1 | Solyc01g105790.3 |
| Solyc01g106000.2 | Solyc01g106000.4 |
| Solyc01g106030.2 | Solyc01g106030.4 |
| Solyc01g106050.2 | Solyc01g106050.4 |
| Solyc01g106080.2 | Solyc01g106080.4 |
| Solyc01g106160.1 | Solyc01g106160.3 |
| Solyc01g106170.2 | Solyc01g106170.4 |
| Solyc01g106180.1 | Solyc01g106180.3 |
| Solyc01g106200.1 | Solyc01g106200.3 |
| Solyc01g106220.2 | Solyc01g106220.4 |
| Solyc01g106250.2 | Solyc01g106250.4 |
| Solyc01g106350.2 | Solyc01g106350.4 |
| Solyc01g106410.1 | Solyc01g106410.3 |
| Solyc01g106460.2 | Solyc01g106460.4 |
| Solyc01g106480.2 | Solyc01g106480.4 |
| Solyc01g106500.2 | Solyc01g106500.4 |
| Solyc01g106590.2 | Solyc01g106590.4 |
| Solyc01g106740.2 | Solyc01g106740.4 |
| Solyc01g106770.2 | Solyc01g106770.4 |
| Solyc01g106800.2 | Solyc01g106800.4 |
| Solyc01g106830.2 | Solyc01g106830.4 |
| Solyc01g106860.2 | Solyc01g106860.4 |
| Solyc01g106870.2 | Solyc01g106870.4 |
| Solyc01g106920.2 | Solyc01g106920.4 |
| Solyc01g106940.2 | Solyc01g106940.4 |
| Solyc01g106970.1 | Solyc01g106970.3 |
| Solyc01g107000.1 | Solyc01g107000.3 |
| Solyc01g107020.2 | Solyc01g107020.4 |
| Solyc01g107070.2 | Solyc01g107070.4 |
| Solyc01g107130.2 | Solyc01g107130.4 |
| Solyc01g107140.1 | Solyc01g107140.3 |
| Solyc01g107200.2 | Solyc01g107200.4 |
| Solyc01g107240.2 | Solyc01g107240.4 |
| Solyc01g107270.2 | Solyc01g107270.4 |
| Solyc01g107300.2 | Solyc01g107300.4 |
| Solyc01g107330.2 | Solyc01g107330.4 |
| Solyc01g107340.2 | Solyc01g107340.4 |
| Solyc01g107360.2 | Solyc01g107360.4 |

| | |
|---|---|
| Solyc01g107390.2 | Solyc01g107390.4 |
| Solyc01g107470.2 | Solyc01g107470.4 |
| Solyc01g107540.2 | Solyc01g107540.4 |
| Solyc01g107580.2 | Solyc01g107580.4 |
| Solyc01g107610.2 | Solyc01g107610.4 |
| Solyc01g107620.2 | Solyc01g107620.4 |
| Solyc01g107630.2 | Solyc01g107630.4 |
| Solyc01g107680.2 | Solyc01g107680.4 |
| Solyc01g107690.2 | Solyc01g107690.4 |
| Solyc01g107700.2 | Solyc01g107700.4 |
| Solyc01g107760.2 | Solyc01g107760.4 |
| Solyc01g107780.2 | Solyc01g107780.4 |
| Solyc01g107800.2 | Solyc01g107800.4 |
| Solyc01g107850.2 | Solyc01g107850.4 |
| Solyc01g107910.2 | Solyc01g107910.4 |
| Solyc01g107960.2 | Solyc01g107960.4 |
| Solyc01g108030.2 | Solyc01g108030.4 |
| Solyc01g108070.2 | Solyc01g108070.4 |
| Solyc01g108080.2 | Solyc01g108080.4 |
| Solyc01g108110.2 | Solyc01g108110.4 |
| Solyc01g108130.2 | Solyc01g108130.4 |
| Solyc01g108140.2 | Solyc01g108140.4 |
| Solyc01g108180.2 | Solyc01g108180.4 |
| Solyc01g108210.2 | Solyc01g108210.4 |
| Solyc01g108260.2 | Solyc01g108260.4 |
| Solyc01g108270.2 | Solyc01g108270.4 |
| Solyc01g108310.2 | Solyc01g108310.4 |
| Solyc01g108350.2 | Solyc01g108350.4 |
| Solyc01g108360.2 | Solyc01g108360.4 |
| Solyc01g108410.1 | Solyc01g108410.3 |
| Solyc01g108470.1 | Solyc01g108470.3 |
| Solyc01g108500.2 | Solyc01g108500.4 |
| Solyc01g108660.2 | Solyc01g108660.4 |
| Solyc01g108670.2 | Solyc01g108670.4 |
| Solyc01g108680.2 | Solyc01g108680.4 |
| Solyc01g108780.2 | Solyc01g108780.4 |
| Solyc01g108790.1 | Solyc01g108790.3 |
| Solyc01g108800.2 | Solyc01g108800.4 |
| Solyc01g108830.1 | Solyc01g108830.3 |
| Solyc01g108880.2 | Solyc01g108880.4 |
| Solyc01g108900.2 | Solyc01g108900.4 |
| Solyc01g108920.2 | Solyc01g108920.4 |
| Solyc01g108950.1 | Solyc01g108950.3 |

| | |
|---|---|
| Solyc01g108970.2 | Solyc01g108970.4 |
| Solyc01g109020.1 | Solyc01g109020.3 |
| Solyc01g109040.2 | Solyc01g109040.4 |
| Solyc01g109060.2 | Solyc01g109060.4 |
| Solyc01g109180.2 | Solyc01g109180.4 |
| Solyc01g109200.2 | Solyc01g109200.4 |
| Solyc01g109220.2 | Solyc01g109220.4 |
| Solyc01g109240.1 | Solyc01g109240.3 |
| Solyc01g109320.2 | Solyc01g109320.4 |
| Solyc01g109340.2 | Solyc01g109340.4 |
| Solyc01g109410.2 | Solyc01g109410.4 |
| Solyc01g109490.2 | Solyc01g109490.4 |
| Solyc01g109500.2 | Solyc01g109500.4 |
| Solyc01g109540.2 | Solyc01g109540.4 |
| Solyc01g109570.2 | Solyc01g109570.4 |
| Solyc01g109580.2 | Solyc01g109580.4 |
| Solyc01g109600.2 | Solyc01g109600.4 |
| Solyc01g109610.2 | Solyc01g109610.4 |
| Solyc01g109630.2 | Solyc01g109630.4 |
| Solyc01g109910.2 | Solyc01g109910.4 |
| Solyc01g109930.2 | Solyc01g109930.4 |
| Solyc01g109950.2 | Solyc01g109950.4 |
| Solyc01g109980.2 | Solyc01g109980.4 |
| Solyc01g109990.2 | Solyc01g109990.4 |
| Solyc01g110020.2 | Solyc01g110020.4 |
| Solyc01g110070.1 | Solyc01g110070.3 |
| Solyc01g110100.2 | Solyc01g110100.4 |
| Solyc01g110140.2 | Solyc01g110140.4 |
| Solyc01g110180.2 | Solyc01g110180.4 |
| Solyc01g110190.2 | Solyc01g110190.4 |
| Solyc01g110210.1 | Solyc01g110210.3 |
| Solyc01g110260.1 | Solyc01g110260.3 |
| Solyc01g110300.2 | Solyc01g110300.4 |
| Solyc01g110320.2 | Solyc01g110320.4 |
| Solyc01g110330.1 | Solyc01g110330.3 |
| Solyc01g110340.2 | Solyc01g110340.4 |
| Solyc01g110370.2 | Solyc01g110370.4 |
| Solyc01g110460.2 | Solyc01g110460.4 |
| Solyc01g110480.2 | Solyc01g110480.4 |
| Solyc01g110490.2 | Solyc01g110490.4 |
| Solyc01g110590.2 | Solyc01g110590.4 |
| Solyc01g110610.2 | Solyc01g110610.4 |
| Solyc01g110630.2 | Solyc01g110630.4 |

| | |
|---|---|
| Solyc01g110660.2 | Solyc01g110660.4 |
| Solyc01g110680.2 | Solyc01g110680.4 |
| Solyc01g110730.2 | Solyc01g110730.4 |
| Solyc01g110820.2 | Solyc01g110820.4 |
| Solyc01g110920.2 | Solyc01g110920.4 |
| Solyc01g110960.2 | Solyc01g110960.4 |
| Solyc01g110980.2 | Solyc01g110980.4 |
| Solyc01g111000.2 | Solyc01g111000.4 |
| Solyc01g111010.2 | Solyc01g111010.4 |
| Solyc01g111040.2 | Solyc01g111040.4 |
| Solyc01g111050.2 | Solyc01g111050.4 |
| Solyc01g111060.2 | Solyc01g111060.4 |
| Solyc01g111100.2 | Solyc01g111100.5 |
| Solyc01g111120.2 | Solyc01g111120.4 |
| Solyc01g111130.2 | Solyc01g111130.4 |
| Solyc01g111240.2 | Solyc01g111240.4 |
| Solyc01g111300.2 | Solyc01g111300.4 |
| Solyc01g111330.2 | Solyc01g111330.4 |
| Solyc01g111340.2 | Solyc01g111340.4 |
| Solyc01g111400.2 | Solyc01g111400.4 |
| Solyc01g111470.2 | Solyc01g111470.4 |
| Solyc01g111540.2 | Solyc01g111540.4 |
| Solyc01g111590.2 | Solyc01g111590.4 |
| Solyc01g111660.2 | Solyc01g111660.4 |
| Solyc01g111680.2 | Solyc01g111680.4 |
| Solyc01g111710.2 | Solyc01g111710.4 |
| Solyc01g111720.2 | Solyc01g111720.4 |
| Solyc01g111750.2 | Solyc01g111750.4 |
| Solyc01g111760.2 | Solyc01g111760.4 |
| Solyc01g111780.2 | Solyc01g111780.4 |
| Solyc01g111800.2 | Solyc01g111800.4 |
| Solyc01g111860.2 | Solyc01g111860.4 |
| Solyc01g111870.1 | Solyc01g111870.3 |
| Solyc01g111880.2 | Solyc01g111880.4 |
| Solyc01g111900.2 | Solyc01g111900.4 |
| Solyc01g111930.2 | Solyc01g111930.4 |
| Solyc01g111940.2 | Solyc01g111940.4 |
| Solyc01g111950.2 | Solyc01g111950.4 |
| Solyc01g111970.2 | Solyc01g111970.4 |
| Solyc01g112000.2 | Solyc01g112000.4 |
| Solyc01g112060.2 | Solyc01g112060.4 |
| Solyc01g112070.2 | Solyc01g112070.4 |
| Solyc01g112120.2 | Solyc01g112120.4 |

| | |
|---|---|
| Solyc01g112130.2 | Solyc01g112130.4 |
| Solyc01g112140.2 | Solyc01g112140.4 |
| Solyc01g112160.2 | Solyc01g112160.4 |
| Solyc01g112180.2 | Solyc01g112180.4 |
| Solyc01g112220.2 | Solyc01g112220.4 |
| Solyc01g112230.2 | Solyc01g112230.4 |
| Solyc01g112240.2 | Solyc01g112240.4 |
| Solyc01g112250.2 | Solyc01g112250.4 |
| Solyc01g112290.2 | Solyc01g112290.4 |
| Solyc01g112310.2 | Solyc01g112310.4 |
| Solyc01g113620.1 | Solyc01g113620.3 |
| Solyc02g005070.1 | Solyc02g005070.3 |
| Solyc02g005200.2 | Solyc02g005200.4 |
| Solyc02g005290.2 | Solyc02g005290.4 |
| Solyc02g005310.1 | Solyc02g005310.3 |
| Solyc02g005340.2 | Solyc02g005340.4 |
| Solyc02g005360.1 | Solyc02g005360.3 |
| Solyc02g005400.1 | Solyc02g005400.3 |
| Solyc02g005410.1 | Solyc02g005410.3 |
| Solyc02g005480.2 | Solyc02g005480.4 |
| Solyc02g005510.2 | Solyc02g005510.4 |
| Solyc02g011680.2 | Solyc02g011680.4 |
| Solyc02g011780.1 | Solyc02g011780.3 |
| Solyc02g011810.1 | Solyc02g011810.3 |
| Solyc02g011820.1 | Solyc02g011820.3 |
| Solyc02g011850.1 | Solyc02g011850.3 |
| Solyc02g011940.1 | Solyc02g011940.3 |
| Solyc02g012010.1 | Solyc02g012010.3 |
| Solyc02g012020.1 | Solyc02g012020.3 |
| Solyc02g014020.1 | Solyc02g014020.3 |
| Solyc02g014030.1 | Solyc02g014030.3 |
| Solyc02g014130.1 | Solyc02g014130.3 |
| Solyc02g014150.2 | Solyc02g014150.4 |
| Solyc02g014190.2 | Solyc02g014190.4 |
| Solyc02g014250.1 | Solyc02g014250.3 |
| Solyc02g014310.2 | Solyc02g014310.4 |
| Solyc02g014350.2 | Solyc02g014350.4 |
| Solyc02g014450.1 | Solyc02g014450.3 |
| Solyc02g014460.2 | Solyc02g014460.4 |
| Solyc02g014470.2 | Solyc02g014470.4 |
| Solyc02g014490.2 | Solyc02g014490.4 |
| Solyc02g014520.2 | Solyc02g014520.4 |
| Solyc02g014530.2 | Solyc02g014530.4 |

| | |
|---|---|
| Solyc02g014540.1 | Solyc02g014540.3 |
| Solyc02g014720.2 | Solyc02g014720.4 |
| Solyc02g014830.2 | Solyc02g014830.4 |
| Solyc02g014870.2 | Solyc02g014870.4 |
| Solyc02g020910.2 | Solyc02g020910.4 |
| Solyc02g020980.2 | Solyc02g020980.4 |
| Solyc02g021020.2 | Solyc02g021020.4 |
| Solyc02g021060.2 | Solyc02g021060.4 |
| Solyc02g021230.2 | Solyc02g021230.4 |
| Solyc02g021260.1 | Solyc02g021260.3 |
| Solyc02g021340.2 | Solyc02g021340.4 |
| Solyc02g021350.2 | Solyc02g021350.4 |
| Solyc02g021410.2 | Solyc02g021410.4 |
| Solyc02g021440.2 | Solyc02g021440.4 |
| Solyc02g021470.2 | Solyc02g021470.4 |
| Solyc02g021480.2 | Solyc02g021480.4 |
| Solyc02g021560.2 | Solyc02g021560.4 |
| Solyc02g021700.2 | Solyc02g021700.4 |
| Solyc02g021780.1 | Solyc02g021780.3 |
| Solyc02g022810.1 | Solyc02g022810.3 |
| Solyc02g022890.1 | Solyc02g022890.3 |
| Solyc02g022910.2 | Solyc02g022910.4 |
| Solyc02g023940.1 | Solyc02g023940.3 |
| Solyc02g024000.2 | Solyc02g024000.4 |
| Solyc02g024060.1 | Solyc02g024060.3 |
| Solyc02g030080.2 | Solyc02g030080.4 |
| Solyc02g030100.2 | Solyc02g030100.4 |
| Solyc02g030120.1 | Solyc02g030120.3 |
| Solyc02g030130.2 | Solyc02g030130.4 |
| Solyc02g030170.2 | Solyc02g030170.4 |
| Solyc02g030210.2 | Solyc02g030210.4 |
| Solyc02g030360.1 | Solyc02g030360.3 |
| Solyc02g030380.1 | Solyc02g030380.3 |
| Solyc02g030480.2 | Solyc02g030480.4 |
| Solyc02g030610.1 | Solyc02g030610.3 |
| Solyc02g030660.1 | Solyc02g030660.3 |
| Solyc02g031700.1 | Solyc02g031700.3 |
| Solyc02g031770.2 | Solyc02g031770.4 |
| Solyc02g031790.1 | Solyc02g031790.3 |
| Solyc02g031890.1 | Solyc02g031890.3 |
| Solyc02g031900.1 | Solyc02g031900.3 |
| Solyc02g031920.2 | Solyc02g031920.4 |
| Solyc02g032020.1 | Solyc02g032020.3 |

| | |
|---|---|
| Solyc02g032100.2 | Solyc02g032100.4 |
| Solyc02g032140.1 | Solyc02g032140.3 |
| Solyc02g032170.2 | Solyc02g032170.4 |
| Solyc02g032180.1 | Solyc02g032180.3 |
| Solyc02g032200.2 | Solyc02g032200.4 |
| Solyc02g032380.1 | Solyc02g032380.3 |
| Solyc02g032480.1 | Solyc02g032480.3 |
| Solyc02g032550.1 | Solyc02g032550.3 |
| Solyc02g032560.1 | Solyc02g032560.3 |
| Solyc02g032580.1 | Solyc02g032580.3 |
| Solyc02g032720.1 | Solyc02g032720.3 |
| Solyc02g032830.1 | Solyc02g032830.3 |
| Solyc02g032870.2 | Solyc02g032870.4 |
| Solyc02g032910.1 | Solyc02g032910.3 |
| Solyc02g032940.2 | Solyc02g032940.4 |
| Solyc02g032970.2 | Solyc02g032970.4 |
| Solyc02g033050.1 | Solyc02g033050.3 |
| Solyc02g033100.1 | Solyc02g033100.3 |
| Solyc02g033110.1 | Solyc02g033110.3 |
| Solyc02g033120.1 | Solyc02g033120.3 |
| Solyc02g036130.1 | Solyc02g036130.3 |
| Solyc02g036170.1 | Solyc02g036170.3 |
| Solyc02g036230.1 | Solyc02g036230.3 |
| Solyc02g036260.2 | Solyc02g036260.4 |
| Solyc02g036270.2 | Solyc02g036270.4 |
| Solyc02g036290.2 | Solyc02g036290.4 |
| Solyc02g036310.2 | Solyc02g036310.4 |
| Solyc02g036470.2 | Solyc02g036470.4 |
| Solyc02g037540.1 | Solyc02g037540.3 |
| Solyc02g037590.1 | Solyc02g037590.3 |
| Solyc02g038640.1 | Solyc02g038640.3 |
| Solyc02g038660.1 | Solyc02g038660.3 |
| Solyc02g038670.1 | Solyc02g038670.3 |
| Solyc02g038700.1 | Solyc02g038700.3 |
| Solyc02g038740.2 | Solyc02g038740.4 |
| Solyc02g038760.2 | Solyc02g038760.4 |
| Solyc02g038790.1 | Solyc02g038790.3 |
| Solyc02g043810.2 | Solyc02g043810.4 |
| Solyc02g044000.1 | Solyc02g044000.3 |
| Solyc02g044020.1 | Solyc02g044020.3 |
| Solyc02g044040.1 | Solyc02g044040.3 |
| Solyc02g050180.2 | Solyc02g050180.4 |
| Solyc02g050240.2 | Solyc02g050240.4 |

| | |
|---|---|
| Solyc02g050280.2 | Solyc02g050280.4 |
| Solyc02g055430.1 | Solyc02g055430.3 |
| Solyc02g055530.2 | Solyc02g055530.4 |
| Solyc02g060580.1 | Solyc02g060580.3 |
| Solyc02g061650.1 | Solyc02g061650.3 |
| Solyc02g061680.1 | Solyc02g061680.3 |
| Solyc02g061720.1 | Solyc02g061720.3 |
| Solyc02g061740.1 | Solyc02g061740.3 |
| Solyc02g061760.2 | Solyc02g061760.4 |
| Solyc02g061770.2 | Solyc02g061770.4 |
| Solyc02g061790.2 | Solyc02g061790.4 |
| Solyc02g061820.2 | Solyc02g061820.4 |
| Solyc02g061850.2 | Solyc02g061850.4 |
| Solyc02g061940.2 | Solyc02g061940.4 |
| Solyc02g061950.2 | Solyc02g061950.4 |
| Solyc02g062000.2 | Solyc02g062000.4 |
| Solyc02g062050.1 | Solyc02g062050.3 |
| Solyc02g062100.2 | Solyc02g062100.4 |
| Solyc02g062110.2 | Solyc02g062110.4 |
| Solyc02g062130.2 | Solyc02g062130.4 |
| Solyc02g062150.1 | Solyc02g062150.3 |
| Solyc02g062210.1 | Solyc02g062210.3 |
| Solyc02g062330.1 | Solyc02g062330.3 |
| Solyc02g062370.2 | Solyc02g062370.4 |
| Solyc02g062380.1 | Solyc02g062380.3 |
| Solyc02g062420.2 | Solyc02g062420.4 |
| Solyc02g062460.2 | Solyc02g062460.4 |
| Solyc02g062490.2 | Solyc02g062490.4 |
| Solyc02g062570.2 | Solyc02g062570.4 |
| Solyc02g062640.2 | Solyc02g062640.4 |
| Solyc02g062720.2 | Solyc02g062720.4 |
| Solyc02g062750.2 | Solyc02g062750.4 |
| Solyc02g062760.2 | Solyc02g062760.4 |
| Solyc02g062780.2 | Solyc02g062780.4 |
| Solyc02g062810.2 | Solyc02g062810.4 |
| Solyc02g062850.1 | Solyc02g062850.3 |
| Solyc02g062890.1 | Solyc02g062890.3 |
| Solyc02g062910.1 | Solyc02g062910.3 |
| Solyc02g062960.2 | Solyc02g062960.4 |
| Solyc02g063000.2 | Solyc02g063000.4 |
| Solyc02g063020.1 | Solyc02g063020.3 |
| Solyc02g063030.2 | Solyc02g063030.4 |
| Solyc02g063060.2 | Solyc02g063060.4 |

| | |
|---|---|
| Solyc02g063130.2 | Solyc02g063130.4 |
| Solyc02g063140.2 | Solyc02g063140.4 |
| Solyc02g063160.2 | Solyc02g063160.4 |
| Solyc02g063180.2 | Solyc02g063180.4 |
| Solyc02g063200.1 | Solyc02g063200.3 |
| Solyc02g063240.2 | Solyc02g063240.4 |
| Solyc02g063260.2 | Solyc02g063260.4 |
| Solyc02g063270.2 | Solyc02g063270.4 |
| Solyc02g063320.2 | Solyc02g063320.4 |
| Solyc02g063390.2 | Solyc02g063390.4 |
| Solyc02g063400.1 | Solyc02g063400.3 |
| Solyc02g063430.2 | Solyc02g063430.4 |
| Solyc02g063450.2 | Solyc02g063450.4 |
| Solyc02g063540.1 | Solyc02g063540.3 |
| Solyc02g064630.2 | Solyc02g064630.4 |
| Solyc02g064640.2 | Solyc02g064640.4 |
| Solyc02g064680.2 | Solyc02g064680.4 |
| Solyc02g064750.2 | Solyc02g064750.4 |
| Solyc02g064760.1 | Solyc02g064760.3 |
| Solyc02g064770.2 | Solyc02g064770.4 |
| Solyc02g064830.2 | Solyc02g064830.4 |
| Solyc02g064860.1 | Solyc02g064860.3 |
| Solyc02g064900.1 | Solyc02g064900.3 |
| Solyc02g064910.1 | Solyc02g064910.3 |
| Solyc02g064970.2 | Solyc02g064970.4 |
| Solyc02g065090.2 | Solyc02g065090.4 |
| Solyc02g065210.2 | Solyc02g065210.4 |
| Solyc02g065220.2 | Solyc02g065220.4 |
| Solyc02g065260.2 | Solyc02g065260.4 |
| Solyc02g065340.1 | Solyc02g065340.3 |
| Solyc02g065360.2 | Solyc02g065360.4 |
| Solyc02g065410.1 | Solyc02g065410.3 |
| Solyc02g065440.1 | Solyc02g065440.3 |
| Solyc02g065490.2 | Solyc02g065490.4 |
| Solyc02g065530.2 | Solyc02g065530.4 |
| Solyc02g065540.1 | Solyc02g065540.3 |
| Solyc02g065550.2 | Solyc02g065550.4 |
| Solyc02g065630.2 | Solyc02g065630.4 |
| Solyc02g065660.2 | Solyc02g065660.4 |
| Solyc02g065670.1 | Solyc02g065670.3 |
| Solyc02g065710.1 | Solyc02g065710.3 |
| Solyc02g065750.1 | Solyc02g065750.3 |
| Solyc02g065760.1 | Solyc02g065760.3 |

| | |
|---|---|
| Solyc02g065800.1 | Solyc02g065800.3 |
| Solyc02g066800.1 | Solyc02g066800.3 |
| Solyc02g066830.2 | Solyc02g066830.4 |
| Solyc02g066900.1 | Solyc02g066900.3 |
| Solyc02g066940.2 | Solyc02g066940.4 |
| Solyc02g066950.2 | Solyc02g066950.4 |
| Solyc02g067030.2 | Solyc02g067030.4 |
| Solyc02g067080.2 | Solyc02g067080.4 |
| Solyc02g067090.2 | Solyc02g067090.4 |
| Solyc02g067160.2 | Solyc02g067160.4 |
| Solyc02g067190.2 | Solyc02g067190.4 |
| Solyc02g067200.2 | Solyc02g067200.4 |
| Solyc02g067220.2 | Solyc02g067220.4 |
| Solyc02g067240.1 | Solyc02g067240.3 |
| Solyc02g067250.1 | Solyc02g067250.3 |
| Solyc02g067270.1 | Solyc02g067270.3 |
| Solyc02g067310.2 | Solyc02g067310.4 |
| Solyc02g067320.1 | Solyc02g067320.3 |
| Solyc02g067340.2 | Solyc02g067340.4 |
| Solyc02g067370.2 | Solyc02g067370.4 |
| Solyc02g067380.2 | Solyc02g067380.4 |
| Solyc02g067400.1 | Solyc02g067400.3 |
| Solyc02g067450.2 | Solyc02g067450.4 |
| Solyc02g067470.2 | Solyc02g067470.4 |
| Solyc02g067520.2 | Solyc02g067520.4 |
| Solyc02g067530.2 | Solyc02g067530.4 |
| Solyc02g067620.1 | Solyc02g067620.3 |
| Solyc02g067660.2 | Solyc02g067660.4 |
| Solyc02g067670.2 | Solyc02g067670.4 |
| Solyc02g067710.1 | Solyc02g067710.3 |
| Solyc02g067750.2 | Solyc02g067750.4 |
| Solyc02g067760.2 | Solyc02g067760.4 |
| Solyc02g067790.2 | Solyc02g067790.4 |
| Solyc02g067800.2 | Solyc02g067800.4 |
| Solyc02g067820.2 | Solyc02g067820.4 |
| Solyc02g067860.2 | Solyc02g067860.4 |
| Solyc02g067880.1 | Solyc02g067880.3 |
| Solyc02g067920.2 | Solyc02g067920.4 |
| Solyc02g067950.2 | Solyc02g067950.4 |
| Solyc02g067960.1 | Solyc02g067960.3 |
| Solyc02g067970.2 | Solyc02g067970.4 |
| Solyc02g068010.2 | Solyc02g068010.4 |
| Solyc02g068050.2 | Solyc02g068050.4 |

| | |
|---|---|
| Solyc02g068070.2 | Solyc02g068070.4 |
| Solyc02g068110.2 | Solyc02g068110.4 |
| Solyc02g068130.2 | Solyc02g068130.4 |
| Solyc02g068150.2 | Solyc02g068150.4 |
| Solyc02g068240.2 | Solyc02g068240.4 |
| Solyc02g068250.2 | Solyc02g068250.4 |
| Solyc02g068260.1 | Solyc02g068260.3 |
| Solyc02g068280.2 | Solyc02g068280.4 |
| Solyc02g068310.2 | Solyc02g068310.4 |
| Solyc02g068360.2 | Solyc02g068360.4 |
| Solyc02g068410.1 | Solyc02g068410.3 |
| Solyc02g068420.2 | Solyc02g068420.4 |
| Solyc02g068430.2 | Solyc02g068430.4 |
| Solyc02g068490.2 | Solyc02g068490.4 |
| Solyc02g068530.2 | Solyc02g068530.4 |
| Solyc02g068660.1 | Solyc02g068660.3 |
| Solyc02g068670.1 | Solyc02g068670.3 |
| Solyc02g068760.2 | Solyc02g068760.4 |
| Solyc02g068770.2 | Solyc02g068770.4 |
| Solyc02g068780.2 | Solyc02g068780.4 |
| Solyc02g068790.2 | Solyc02g068790.4 |
| Solyc02g068820.1 | Solyc02g068820.3 |
| Solyc02g068870.1 | Solyc02g068870.3 |
| Solyc02g068930.2 | Solyc02g068930.4 |
| Solyc02g069010.2 | Solyc02g069010.4 |
| Solyc02g069040.2 | Solyc02g069040.4 |
| Solyc02g069060.2 | Solyc02g069060.4 |
| Solyc02g069080.2 | Solyc02g069080.4 |
| Solyc02g069100.2 | Solyc02g069100.4 |
| Solyc02g069110.2 | Solyc02g069110.4 |
| Solyc02g069140.1 | Solyc02g069140.3 |
| Solyc02g069170.1 | Solyc02g069170.3 |
| Solyc02g069180.2 | Solyc02g069180.4 |
| Solyc02g069190.2 | Solyc02g069190.4 |
| Solyc02g069250.2 | Solyc02g069250.4 |
| Solyc02g069340.2 | Solyc02g069340.4 |
| Solyc02g069350.2 | Solyc02g069350.4 |
| Solyc02g069410.2 | Solyc02g069410.4 |
| Solyc02g069420.2 | Solyc02g069420.4 |
| Solyc02g069430.2 | Solyc02g069430.4 |
| Solyc02g069470.2 | Solyc02g069470.4 |
| Solyc02g069480.1 | Solyc02g069480.3 |
| Solyc02g069490.2 | Solyc02g069490.4 |

| | |
|---|---|
| Solyc02g069540.1 | Solyc02g069540.3 |
| Solyc02g069620.2 | Solyc02g069620.4 |
| Solyc02g069650.1 | Solyc02g069650.3 |
| Solyc02g069670.2 | Solyc02g069670.4 |
| Solyc02g069710.2 | Solyc02g069710.4 |
| Solyc02g069740.2 | Solyc02g069740.4 |
| Solyc02g069750.1 | Solyc02g069750.3 |
| Solyc02g069810.1 | Solyc02g069810.3 |
| Solyc02g069830.2 | Solyc02g069830.4 |
| Solyc02g069860.2 | Solyc02g069860.4 |
| Solyc02g069880.2 | Solyc02g069880.4 |
| Solyc02g069920.2 | Solyc02g069920.4 |
| Solyc02g069940.2 | Solyc02g069940.4 |
| Solyc02g069960.2 | Solyc02g069960.4 |
| Solyc02g069970.2 | Solyc02g069970.4 |
| Solyc02g069980.2 | Solyc02g069980.4 |
| Solyc02g070000.2 | Solyc02g070000.4 |
| Solyc02g070030.2 | Solyc02g070030.4 |
| Solyc02g070080.2 | Solyc02g070080.4 |
| Solyc02g070180.1 | Solyc02g070180.3 |
| Solyc02g070200.1 | Solyc02g070200.3 |
| Solyc02g070250.2 | Solyc02g070250.4 |
| Solyc02g070260.2 | Solyc02g070260.4 |
| Solyc02g070300.2 | Solyc02g070300.4 |
| Solyc02g070330.2 | Solyc02g070330.4 |
| Solyc02g070480.2 | Solyc02g070480.4 |
| Solyc02g070490.2 | Solyc02g070490.4 |
| Solyc02g070530.2 | Solyc02g070530.4 |
| Solyc02g070570.2 | Solyc02g070570.4 |
| Solyc02g070600.1 | Solyc02g070600.3 |
| Solyc02g070620.2 | Solyc02g070620.4 |
| Solyc02g070670.2 | Solyc02g070670.4 |
| Solyc02g070680.2 | Solyc02g070680.4 |
| Solyc02g070700.1 | Solyc02g070700.3 |
| Solyc02g070710.1 | Solyc02g070710.3 |
| Solyc02g070760.2 | Solyc02g070760.4 |
| Solyc02g070780.2 | Solyc02g070780.4 |
| Solyc02g070810.2 | Solyc02g070810.4 |
| Solyc02g070860.2 | Solyc02g070860.4 |
| Solyc02g070870.2 | Solyc02g070870.4 |
| Solyc02g071050.2 | Solyc02g071050.4 |
| Solyc02g071080.1 | Solyc02g071080.3 |
| Solyc02g071130.2 | Solyc02g071130.4 |

| | |
|---|---|
| Solyc02g071160.2 | Solyc02g071160.4 |
| Solyc02g071190.2 | Solyc02g071190.4 |
| Solyc02g071200.2 | Solyc02g071200.4 |
| Solyc02g071270.1 | Solyc02g071270.3 |
| Solyc02g071320.2 | Solyc02g071320.4 |
| Solyc02g071330.2 | Solyc02g071330.4 |
| Solyc02g071360.2 | Solyc02g071360.4 |
| Solyc02g071420.2 | Solyc02g071420.4 |
| Solyc02g071470.2 | Solyc02g071470.4 |
| Solyc02g071490.2 | Solyc02g071490.4 |
| Solyc02g071560.2 | Solyc02g071560.4 |
| Solyc02g071570.2 | Solyc02g071570.4 |
| Solyc02g071580.2 | Solyc02g071580.4 |
| Solyc02g071590.1 | Solyc02g071590.3 |
| Solyc02g071640.1 | Solyc02g071640.3 |
| Solyc02g071660.2 | Solyc02g071660.4 |
| Solyc02g071730.2 | Solyc02g071730.4 |
| Solyc02g071760.2 | Solyc02g071760.4 |
| Solyc02g071790.1 | Solyc02g071790.3 |
| Solyc02g071810.2 | Solyc02g071810.4 |
| Solyc02g071820.2 | Solyc02g071820.4 |
| Solyc02g071860.2 | Solyc02g071860.4 |
| Solyc02g071870.2 | Solyc02g071870.4 |
| Solyc02g072010.1 | Solyc02g072010.3 |
| Solyc02g072020.1 | Solyc02g072020.3 |
| Solyc02g072060.1 | Solyc02g072060.3 |
| Solyc02g072070.2 | Solyc02g072070.4 |
| Solyc02g072160.2 | Solyc02g072160.4 |
| Solyc02g072190.2 | Solyc02g072190.4 |
| Solyc02g072260.2 | Solyc02g072260.4 |
| Solyc02g072300.2 | Solyc02g072300.4 |
| Solyc02g072330.1 | Solyc02g072330.3 |
| Solyc02g072440.2 | Solyc02g072440.4 |
| Solyc02g072450.2 | Solyc02g072450.4 |
| Solyc02g072470.2 | Solyc02g072470.4 |
| Solyc02g072480.2 | Solyc02g072480.4 |
| Solyc02g072530.1 | Solyc02g072530.3 |
| Solyc02g074600.2 | Solyc02g074600.4 |
| Solyc02g075610.2 | Solyc02g075610.4 |
| Solyc02g075620.2 | Solyc02g075620.4 |
| Solyc02g075630.1 | Solyc02g075630.3 |
| Solyc02g076740.1 | Solyc02g076740.3 |
| Solyc02g076750.1 | Solyc02g076750.3 |

| | |
|---|---|
| Solyc02g076790.2 | Solyc02g076790.4 |
| Solyc02g076810.2 | Solyc02g076810.4 |
| Solyc02g076820.2 | Solyc02g076820.4 |
| Solyc02g076880.2 | Solyc02g076880.4 |
| Solyc02g076900.2 | Solyc02g076900.4 |
| Solyc02g077090.2 | Solyc02g077090.4 |
| Solyc02g077170.1 | Solyc02g077170.3 |
| Solyc02g077190.1 | Solyc02g077190.3 |
| Solyc02g077240.2 | Solyc02g077240.4 |
| Solyc02g077270.2 | Solyc02g077270.4 |
| Solyc02g077280.2 | Solyc02g077280.4 |
| Solyc02g077310.2 | Solyc02g077310.4 |
| Solyc02g077320.2 | Solyc02g077320.4 |
| Solyc02g077400.2 | Solyc02g077400.4 |
| Solyc02g077430.2 | Solyc02g077430.4 |
| Solyc02g077460.1 | Solyc02g077460.3 |
| Solyc02g077470.2 | Solyc02g077470.4 |
| Solyc02g077510.2 | Solyc02g077510.4 |
| Solyc02g077540.2 | Solyc02g077540.4 |
| Solyc02g077550.2 | Solyc02g077550.4 |
| Solyc02g077570.1 | Solyc02g077570.3 |
| Solyc02g077680.2 | Solyc02g077680.4 |
| Solyc02g077690.2 | Solyc02g077690.4 |
| Solyc02g077720.2 | Solyc02g077720.4 |
| Solyc02g077750.2 | Solyc02g077750.4 |
| Solyc02g077890.2 | Solyc02g077890.4 |
| Solyc02g077900.2 | Solyc02g077900.4 |
| Solyc02g077910.2 | Solyc02g077910.4 |
| Solyc02g077920.2 | Solyc02g077920.4 |
| Solyc02g077950.1 | Solyc02g077950.3 |
| Solyc02g077960.1 | Solyc02g077960.3 |
| Solyc02g077970.2 | Solyc02g077970.4 |
| Solyc02g077980.1 | Solyc02g077980.3 |
| Solyc02g078030.1 | Solyc02g078030.3 |
| Solyc02g078050.2 | Solyc02g078050.4 |
| Solyc02g078150.2 | Solyc02g078150.4 |
| Solyc02g078170.1 | Solyc02g078170.3 |
| Solyc02g078200.2 | Solyc02g078200.4 |
| Solyc02g078240.2 | Solyc02g078240.4 |
| Solyc02g078250.2 | Solyc02g078250.4 |
| Solyc02g078320.1 | Solyc02g078320.3 |
| Solyc02g078340.2 | Solyc02g078340.4 |
| Solyc02g078360.2 | Solyc02g078360.4 |

| | |
|---|---|
| Solyc02g078470.2 | Solyc02g078470.4 |
| Solyc02g078510.2 | Solyc02g078510.4 |
| Solyc02g078520.2 | Solyc02g078520.4 |
| Solyc02g078540.2 | Solyc02g078540.4 |
| Solyc02g078560.2 | Solyc02g078560.4 |
| Solyc02g078610.2 | Solyc02g078610.4 |
| Solyc02g078650.2 | Solyc02g078650.4 |
| Solyc02g078660.2 | Solyc02g078660.4 |
| Solyc02g078740.2 | Solyc02g078740.4 |
| Solyc02g078790.2 | Solyc02g078790.4 |
| Solyc02g078800.2 | Solyc02g078800.4 |
| Solyc02g078810.2 | Solyc02g078810.4 |
| Solyc02g078830.2 | Solyc02g078830.4 |
| Solyc02g078840.2 | Solyc02g078840.4 |
| Solyc02g078880.2 | Solyc02g078880.4 |
| Solyc02g078920.2 | Solyc02g078920.4 |
| Solyc02g078930.1 | Solyc02g078930.3 |
| Solyc02g078950.2 | Solyc02g078950.4 |
| Solyc02g078960.2 | Solyc02g078960.4 |
| Solyc02g078970.2 | Solyc02g078970.4 |
| Solyc02g078990.2 | Solyc02g078990.4 |
| Solyc02g079000.2 | Solyc02g079000.4 |
| Solyc02g079010.2 | Solyc02g079010.4 |
| Solyc02g079020.2 | Solyc02g079020.4 |
| Solyc02g079070.2 | Solyc02g079070.4 |
| Solyc02g079090.2 | Solyc02g079090.4 |
| Solyc02g079220.2 | Solyc02g079220.4 |
| Solyc02g079250.2 | Solyc02g079250.4 |
| Solyc02g079270.2 | Solyc02g079270.4 |
| Solyc02g079310.1 | Solyc02g079310.3 |
| Solyc02g079360.1 | Solyc02g079360.3 |
| Solyc02g079390.2 | Solyc02g079390.4 |
| Solyc02g079430.2 | Solyc02g079430.4 |
| Solyc02g079440.1 | Solyc02g079440.3 |
| Solyc02g079520.1 | Solyc02g079520.3 |
| Solyc02g079530.2 | Solyc02g079530.4 |
| Solyc02g079550.1 | Solyc02g079550.3 |
| Solyc02g079570.2 | Solyc02g079570.4 |
| Solyc02g079580.2 | Solyc02g079580.4 |
| Solyc02g079620.1 | Solyc02g079620.3 |
| Solyc02g079630.1 | Solyc02g079630.3 |
| Solyc02g079710.2 | Solyc02g079710.4 |
| Solyc02g079730.2 | Solyc02g079730.4 |

| | |
|---|---|
| Solyc02g079750.2 | Solyc02g079750.4 |
| Solyc02g079810.1 | Solyc02g079810.3 |
| Solyc02g079820.2 | Solyc02g079820.4 |
| Solyc02g079850.2 | Solyc02g079850.4 |
| Solyc02g079870.2 | Solyc02g079870.4 |
| Solyc02g079910.1 | Solyc02g079910.3 |
| Solyc02g079970.2 | Solyc02g079970.4 |
| Solyc02g080040.2 | Solyc02g080040.4 |
| Solyc02g080120.1 | Solyc02g080120.3 |
| Solyc02g080130.2 | Solyc02g080130.4 |
| Solyc02g080140.2 | Solyc02g080140.4 |
| Solyc02g080150.1 | Solyc02g080150.3 |
| Solyc02g080160.2 | Solyc02g080160.4 |
| Solyc02g080200.2 | Solyc02g080200.4 |
| Solyc02g080260.2 | Solyc02g080260.4 |
| Solyc02g080320.2 | Solyc02g080320.4 |
| Solyc02g080330.2 | Solyc02g080330.4 |
| Solyc02g080380.2 | Solyc02g080380.4 |
| Solyc02g080420.2 | Solyc02g080420.4 |
| Solyc02g080460.1 | Solyc02g080460.3 |
| Solyc02g080520.2 | Solyc02g080520.4 |
| Solyc02g080610.2 | Solyc02g080610.4 |
| Solyc02g080630.2 | Solyc02g080630.4 |
| Solyc02g080640.2 | Solyc02g080640.4 |
| Solyc02g080700.1 | Solyc02g080700.3 |
| Solyc02g080740.1 | Solyc02g080740.3 |
| Solyc02g080750.1 | Solyc02g080750.3 |
| Solyc02g080790.2 | Solyc02g080790.4 |
| Solyc02g080800.2 | Solyc02g080800.4 |
| Solyc02g080860.1 | Solyc02g080860.3 |
| Solyc02g080870.1 | Solyc02g080870.3 |
| Solyc02g080880.2 | Solyc02g080880.4 |
| Solyc02g080910.2 | Solyc02g080910.4 |
| Solyc02g080920.1 | Solyc02g080920.3 |
| Solyc02g080930.1 | Solyc02g080930.3 |
| Solyc02g080950.1 | Solyc02g080950.3 |
| Solyc02g080960.1 | Solyc02g080960.3 |
| Solyc02g080970.2 | Solyc02g080970.4 |
| Solyc02g081010.1 | Solyc02g081010.3 |
| Solyc02g081030.2 | Solyc02g081030.4 |
| Solyc02g081040.2 | Solyc02g081040.4 |
| Solyc02g081050.2 | Solyc02g081050.4 |
| Solyc02g081060.2 | Solyc02g081060.4 |

| | |
|---|---|
| Solyc02g081070.2 | Solyc02g081070.4 |
| Solyc02g081120.2 | Solyc02g081120.4 |
| Solyc02g081130.1 | Solyc02g081130.3 |
| Solyc02g081170.2 | Solyc02g081170.4 |
| Solyc02g081190.2 | Solyc02g081190.4 |
| Solyc02g081220.1 | Solyc02g081220.3 |
| Solyc02g081230.2 | Solyc02g081230.4 |
| Solyc02g081250.1 | Solyc02g081250.3 |
| Solyc02g081270.2 | Solyc02g081270.4 |
| Solyc02g081290.2 | Solyc02g081290.4 |
| Solyc02g081320.2 | Solyc02g081320.4 |
| Solyc02g081360.2 | Solyc02g081360.4 |
| Solyc02g081460.2 | Solyc02g081460.4 |
| Solyc02g081470.2 | Solyc02g081470.4 |
| Solyc02g081480.2 | Solyc02g081480.4 |
| Solyc02g081510.1 | Solyc02g081510.3 |
| Solyc02g081570.2 | Solyc02g081570.4 |
| Solyc02g081600.2 | Solyc02g081600.4 |
| Solyc02g081610.1 | Solyc02g081610.3 |
| Solyc02g081640.2 | Solyc02g081640.4 |
| Solyc02g081680.2 | Solyc02g081680.4 |
| Solyc02g081780.1 | Solyc02g081780.3 |
| Solyc02g081810.2 | Solyc02g081810.4 |
| Solyc02g081830.2 | Solyc02g081830.4 |
| Solyc02g081850.2 | Solyc02g081850.4 |
| Solyc02g081860.1 | Solyc02g081860.3 |
| Solyc02g081870.2 | Solyc02g081870.4 |
| Solyc02g081890.2 | Solyc02g081890.4 |
| Solyc02g081900.2 | Solyc02g081900.4 |
| Solyc02g081950.2 | Solyc02g081950.4 |
| Solyc02g081970.2 | Solyc02g081970.4 |
| Solyc02g082010.1 | Solyc02g082010.3 |
| Solyc02g082070.2 | Solyc02g082070.4 |
| Solyc02g082100.2 | Solyc02g082100.4 |
| Solyc02g082110.2 | Solyc02g082110.4 |
| Solyc02g082170.1 | Solyc02g082170.3 |
| Solyc02g082200.2 | Solyc02g082200.4 |
| Solyc02g082230.2 | Solyc02g082230.4 |
| Solyc02g082250.2 | Solyc02g082250.4 |
| Solyc02g082290.2 | Solyc02g082290.4 |
| Solyc02g082360.2 | Solyc02g082360.4 |
| Solyc02g082400.2 | Solyc02g082400.4 |
| Solyc02g082410.2 | Solyc02g082410.4 |

| | |
|---|---|
| Solyc02g082430.2 | Solyc02g082430.4 |
| Solyc02g082470.2 | Solyc02g082470.4 |
| Solyc02g082490.2 | Solyc02g082490.4 |
| Solyc02g082520.1 | Solyc02g082520.3 |
| Solyc02g082540.2 | Solyc02g082540.4 |
| Solyc02g082600.2 | Solyc02g082600.4 |
| Solyc02g082720.2 | Solyc02g082720.4 |
| Solyc02g082750.2 | Solyc02g082750.4 |
| Solyc02g082830.1 | Solyc02g082830.3 |
| Solyc02g082870.2 | Solyc02g082870.4 |
| Solyc02g082910.2 | Solyc02g082910.4 |
| Solyc02g082920.2 | Solyc02g082920.4 |
| Solyc02g082990.2 | Solyc02g082990.4 |
| Solyc02g083120.1 | Solyc02g083120.3 |
| Solyc02g083130.2 | Solyc02g083130.4 |
| Solyc02g083160.1 | Solyc02g083160.3 |
| Solyc02g083180.2 | Solyc02g083180.4 |
| Solyc02g083280.2 | Solyc02g083280.4 |
| Solyc02g083290.2 | Solyc02g083290.4 |
| Solyc02g083340.2 | Solyc02g083340.4 |
| Solyc02g083350.2 | Solyc02g083350.4 |
| Solyc02g083390.2 | Solyc02g083390.4 |
| Solyc02g083410.2 | Solyc02g083410.4 |
| Solyc02g083600.2 | Solyc02g083600.4 |
| Solyc02g083610.2 | Solyc02g083610.4 |
| Solyc02g083640.2 | Solyc02g083640.4 |
| Solyc02g083670.2 | Solyc02g083670.4 |
| Solyc02g083700.2 | Solyc02g083700.4 |
| Solyc02g083720.2 | Solyc02g083720.4 |
| Solyc02g083730.2 | Solyc02g083730.4 |
| Solyc02g083760.2 | Solyc02g083760.4 |
| Solyc02g083780.1 | Solyc02g083780.3 |
| Solyc02g083790.2 | Solyc02g083790.4 |
| Solyc02g083800.2 | Solyc02g083800.4 |
| Solyc02g083810.2 | Solyc02g083810.4 |
| Solyc02g083910.2 | Solyc02g083910.4 |
| Solyc02g083960.2 | Solyc02g083960.4 |
| Solyc02g084100.1 | Solyc02g084100.3 |
| Solyc02g084130.2 | Solyc02g084130.4 |
| Solyc02g084140.1 | Solyc02g084140.3 |
| Solyc02g084160.1 | Solyc02g084160.3 |
| Solyc02g084190.1 | Solyc02g084190.3 |
| Solyc02g084200.1 | Solyc02g084200.3 |

| | |
|---|---|
| Solyc02g084230.1 | Solyc02g084230.3 |
| Solyc02g084260.1 | Solyc02g084260.3 |
| Solyc02g084290.1 | Solyc02g084290.3 |
| Solyc02g084340.1 | Solyc02g084340.3 |
| Solyc02g084400.2 | Solyc02g084400.4 |
| Solyc02g084440.2 | Solyc02g084440.4 |
| Solyc02g084450.2 | Solyc02g084450.4 |
| Solyc02g084480.1 | Solyc02g084480.3 |
| Solyc02g084560.2 | Solyc02g084560.4 |
| Solyc02g084570.2 | Solyc02g084570.4 |
| Solyc02g084590.2 | Solyc02g084590.4 |
| Solyc02g084600.2 | Solyc02g084600.4 |
| Solyc02g084640.2 | Solyc02g084640.4 |
| Solyc02g084670.2 | Solyc02g084670.4 |
| Solyc02g084740.2 | Solyc02g084740.4 |
| Solyc02g084750.2 | Solyc02g084750.4 |
| Solyc02g084780.2 | Solyc02g084780.4 |
| Solyc02g084800.2 | Solyc02g084800.4 |
| Solyc02g084830.2 | Solyc02g084830.4 |
| Solyc02g084890.1 | Solyc02g084890.3 |
| Solyc02g084910.2 | Solyc02g084910.4 |
| Solyc02g084970.2 | Solyc02g084970.4 |
| Solyc02g085020.2 | Solyc02g085020.4 |
| Solyc02g085030.2 | Solyc02g085030.4 |
| Solyc02g085110.2 | Solyc02g085110.4 |
| Solyc02g085130.2 | Solyc02g085130.4 |
| Solyc02g085140.2 | Solyc02g085140.4 |
| Solyc02g085150.2 | Solyc02g085150.4 |
| Solyc02g085170.2 | Solyc02g085170.4 |
| Solyc02g085220.2 | Solyc02g085220.4 |
| Solyc02g085240.2 | Solyc02g085240.4 |
| Solyc02g085250.2 | Solyc02g085250.4 |
| Solyc02g085300.2 | Solyc02g085300.4 |
| Solyc02g085350.2 | Solyc02g085350.4 |
| Solyc02g085360.2 | Solyc02g085360.4 |
| Solyc02g085390.2 | Solyc02g085390.4 |
| Solyc02g085420.2 | Solyc02g085420.4 |
| Solyc02g085430.2 | Solyc02g085430.4 |
| Solyc02g085440.2 | Solyc02g085440.4 |
| Solyc02g085500.2 | Solyc02g085500.4 |
| Solyc02g085520.2 | Solyc02g085520.4 |
| Solyc02g085530.1 | Solyc02g085530.3 |
| Solyc02g085550.1 | Solyc02g085550.3 |

| | |
|---|---|
| Solyc02g085570.2 | Solyc02g085570.4 |
| Solyc02g085580.2 | Solyc02g085580.4 |
| Solyc02g085590.2 | Solyc02g085590.4 |
| Solyc02g085610.2 | Solyc02g085610.4 |
| Solyc02g085640.2 | Solyc02g085640.4 |
| Solyc02g085650.2 | Solyc02g085650.4 |
| Solyc02g085690.2 | Solyc02g085690.4 |
| Solyc02g085710.2 | Solyc02g085710.4 |
| Solyc02g085740.2 | Solyc02g085740.4 |
| Solyc02g085750.2 | Solyc02g085750.4 |
| Solyc02g085770.2 | Solyc02g085770.4 |
| Solyc02g085830.2 | Solyc02g085830.4 |
| Solyc02g085860.2 | Solyc02g085860.4 |
| Solyc02g085880.1 | Solyc02g085880.3 |
| Solyc02g085910.2 | Solyc02g085910.4 |
| Solyc02g085940.2 | Solyc02g085940.4 |
| Solyc02g085950.2 | Solyc02g085950.4 |
| Solyc02g086010.1 | Solyc02g086010.3 |
| Solyc02g086060.2 | Solyc02g086060.4 |
| Solyc02g086080.2 | Solyc02g086080.4 |
| Solyc02g086090.2 | Solyc02g086090.4 |
| Solyc02g086100.2 | Solyc02g086100.4 |
| Solyc02g086130.2 | Solyc02g086130.4 |
| Solyc02g086150.2 | Solyc02g086150.4 |
| Solyc02g086160.2 | Solyc02g086160.4 |
| Solyc02g086180.2 | Solyc02g086180.4 |
| Solyc02g086190.1 | Solyc02g086190.3 |
| Solyc02g086200.1 | Solyc02g086200.3 |
| Solyc02g086210.2 | Solyc02g086210.4 |
| Solyc02g086270.2 | Solyc02g086270.4 |
| Solyc02g086300.2 | Solyc02g086300.4 |
| Solyc02g086330.2 | Solyc02g086330.4 |
| Solyc02g086370.2 | Solyc02g086370.4 |
| Solyc02g086400.1 | Solyc02g086400.3 |
| Solyc02g086410.1 | Solyc02g086410.3 |
| Solyc02g086430.2 | Solyc02g086430.4 |
| Solyc02g086440.2 | Solyc02g086440.4 |
| Solyc02g086510.2 | Solyc02g086510.4 |
| Solyc02g086530.2 | Solyc02g086530.4 |
| Solyc02g086540.1 | Solyc02g086540.3 |
| Solyc02g086570.1 | Solyc02g086570.3 |
| Solyc02g086580.2 | Solyc02g086580.4 |
| Solyc02g086610.2 | Solyc02g086610.4 |

| | |
|---|---|
| Solyc02g086630.1 | Solyc02g086630.3 |
| Solyc02g086640.2 | Solyc02g086640.4 |
| Solyc02g086660.2 | Solyc02g086660.4 |
| Solyc02g086670.2 | Solyc02g086670.4 |
| Solyc02g086680.2 | Solyc02g086680.4 |
| Solyc02g086700.2 | Solyc02g086700.4 |
| Solyc02g086730.1 | Solyc02g086730.3 |
| Solyc02g086750.1 | Solyc02g086750.3 |
| Solyc02g086780.2 | Solyc02g086780.4 |
| Solyc02g086790.2 | Solyc02g086790.4 |
| Solyc02g086800.2 | Solyc02g086800.4 |
| Solyc02g086850.1 | Solyc02g086850.3 |
| Solyc02g086860.2 | Solyc02g086860.4 |
| Solyc02g086870.2 | Solyc02g086870.4 |
| Solyc02g086880.2 | Solyc02g086880.4 |
| Solyc02g086900.2 | Solyc02g086900.4 |
| Solyc02g086910.2 | Solyc02g086910.4 |
| Solyc02g086930.2 | Solyc02g086930.4 |
| Solyc02g086970.2 | Solyc02g086970.4 |
| Solyc02g086990.2 | Solyc02g086990.4 |
| Solyc02g087000.2 | Solyc02g087000.4 |
| Solyc02g087060.2 | Solyc02g087060.4 |
| Solyc02g087070.2 | Solyc02g087070.4 |
| Solyc02g087090.2 | Solyc02g087090.4 |
| Solyc02g087120.2 | Solyc02g087120.4 |
| Solyc02g087130.2 | Solyc02g087130.4 |
| Solyc02g087170.2 | Solyc02g087170.4 |
| Solyc02g087180.2 | Solyc02g087180.4 |
| Solyc02g087220.2 | Solyc02g087220.4 |
| Solyc02g087240.2 | Solyc02g087240.4 |
| Solyc02g087250.2 | Solyc02g087250.4 |
| Solyc02g087350.1 | Solyc02g087350.3 |
| Solyc02g087360.2 | Solyc02g087360.4 |
| Solyc02g087390.1 | Solyc02g087390.3 |
| Solyc02g087410.2 | Solyc02g087410.4 |
| Solyc02g087430.2 | Solyc02g087430.4 |
| Solyc02g087440.2 | Solyc02g087440.4 |
| Solyc02g087480.2 | Solyc02g087480.4 |
| Solyc02g087490.1 | Solyc02g087490.3 |
| Solyc02g087500.1 | Solyc02g087500.3 |
| Solyc02g087540.1 | Solyc02g087540.3 |
| Solyc02g087590.1 | Solyc02g087590.3 |
| Solyc02g087660.2 | Solyc02g087660.4 |

| | |
|---|---|
| Solyc02g087830.2 | Solyc02g087830.4 |
| Solyc02g087890.2 | Solyc02g087890.4 |
| Solyc02g087910.1 | Solyc02g087910.3 |
| Solyc02g087990.2 | Solyc02g087990.4 |
| Solyc02g088060.1 | Solyc02g088060.3 |
| Solyc02g088110.2 | Solyc02g088110.4 |
| Solyc02g088120.2 | Solyc02g088120.4 |
| Solyc02g088140.1 | Solyc02g088140.3 |
| Solyc02g088160.2 | Solyc02g088160.4 |
| Solyc02g088190.2 | Solyc02g088190.4 |
| Solyc02g088240.2 | Solyc02g088240.4 |
| Solyc02g088290.2 | Solyc02g088290.4 |
| Solyc02g088320.2 | Solyc02g088320.4 |
| Solyc02g088340.2 | Solyc02g088340.4 |
| Solyc02g088380.2 | Solyc02g088380.4 |
| Solyc02g088390.2 | Solyc02g088390.4 |
| Solyc02g088420.1 | Solyc02g088420.3 |
| Solyc02g088450.2 | Solyc02g088450.4 |
| Solyc02g088460.2 | Solyc02g088460.4 |
| Solyc02g088510.2 | Solyc02g088510.4 |
| Solyc02g088520.2 | Solyc02g088520.4 |
| Solyc02g088550.2 | Solyc02g088550.4 |
| Solyc02g088560.2 | Solyc02g088560.4 |
| Solyc02g088590.2 | Solyc02g088590.4 |
| Solyc02g088600.2 | Solyc02g088600.4 |
| Solyc02g088610.2 | Solyc02g088610.4 |
| Solyc02g088630.2 | Solyc02g088630.4 |
| Solyc02g088660.2 | Solyc02g088660.4 |
| Solyc02g088690.2 | Solyc02g088690.4 |
| Solyc02g088700.2 | Solyc02g088700.4 |
| Solyc02g088720.2 | Solyc02g088720.4 |
| Solyc02g088780.2 | Solyc02g088780.4 |
| Solyc02g088820.2 | Solyc02g088820.4 |
| Solyc02g088920.2 | Solyc02g088920.4 |
| Solyc02g088950.1 | Solyc02g088950.3 |
| Solyc02g088980.1 | Solyc02g088980.3 |
| Solyc02g089050.2 | Solyc02g089050.4 |
| Solyc02g089120.2 | Solyc02g089120.4 |
| Solyc02g089130.2 | Solyc02g089130.4 |
| Solyc02g089150.2 | Solyc02g089150.4 |
| Solyc02g089170.2 | Solyc02g089170.4 |
| Solyc02g089200.2 | Solyc02g089200.4 |
| Solyc02g089210.2 | Solyc02g089210.4 |

| | |
|---|---|
| Solyc02g089220.2 | Solyc02g089220.4 |
| Solyc02g089250.2 | Solyc02g089250.4 |
| Solyc02g089260.2 | Solyc02g089260.4 |
| Solyc02g089340.2 | Solyc02g089340.4 |
| Solyc02g089360.2 | Solyc02g089360.4 |
| Solyc02g089390.1 | Solyc02g089390.3 |
| Solyc02g089400.2 | Solyc02g089400.4 |
| Solyc02g089490.2 | Solyc02g089490.4 |
| Solyc02g089500.2 | Solyc02g089500.4 |
| Solyc02g089560.2 | Solyc02g089560.4 |
| Solyc02g089670.2 | Solyc02g089670.4 |
| Solyc02g089680.2 | Solyc02g089680.4 |
| Solyc02g089700.2 | Solyc02g089700.4 |
| Solyc02g089720.1 | Solyc02g089720.3 |
| Solyc02g089790.2 | Solyc02g089790.4 |
| Solyc02g089820.2 | Solyc02g089820.4 |
| Solyc02g089870.1 | Solyc02g089870.3 |
| Solyc02g089880.2 | Solyc02g089880.4 |
| Solyc02g089890.2 | Solyc02g089890.4 |
| Solyc02g089930.2 | Solyc02g089930.4 |
| Solyc02g089940.2 | Solyc02g089940.4 |
| Solyc02g090010.1 | Solyc02g090010.3 |
| Solyc02g090020.2 | Solyc02g090020.4 |
| Solyc02g090130.2 | Solyc02g090130.4 |
| Solyc02g090230.2 | Solyc02g090230.4 |
| Solyc02g090240.2 | Solyc02g090240.4 |
| Solyc02g090350.2 | Solyc02g090350.4 |
| Solyc02g090370.2 | Solyc02g090370.4 |
| Solyc02g090380.2 | Solyc02g090380.4 |
| Solyc02g090390.2 | Solyc02g090390.4 |
| Solyc02g090400.2 | Solyc02g090400.4 |
| Solyc02g090410.2 | Solyc02g090410.4 |
| Solyc02g090470.2 | Solyc02g090470.4 |
| Solyc02g090480.2 | Solyc02g090480.4 |
| Solyc02g090530.2 | Solyc02g090530.4 |
| Solyc02g090540.2 | Solyc02g090540.4 |
| Solyc02g090570.2 | Solyc02g090570.4 |
| Solyc02g090630.2 | Solyc02g090630.4 |
| Solyc02g090640.2 | Solyc02g090640.4 |
| Solyc02g090660.1 | Solyc02g090660.3 |
| Solyc02g090690.1 | Solyc02g090690.3 |
| Solyc02g090750.2 | Solyc02g090750.4 |
| Solyc02g090810.2 | Solyc02g090810.4 |

| | |
|---|---|
| Solyc02g090890.2 | Solyc02g090890.4 |
| Solyc02g091000.2 | Solyc02g091000.4 |
| Solyc02g091020.1 | Solyc02g091020.3 |
| Solyc02g091030.2 | Solyc02g091030.4 |
| Solyc02g091090.2 | Solyc02g091090.4 |
| Solyc02g091120.2 | Solyc02g091120.4 |
| Solyc02g091150.1 | Solyc02g091150.3 |
| Solyc02g091160.2 | Solyc02g091160.4 |
| Solyc02g091170.2 | Solyc02g091170.4 |
| Solyc02g091190.2 | Solyc02g091190.4 |
| Solyc02g091230.2 | Solyc02g091230.4 |
| Solyc02g091330.2 | Solyc02g091330.4 |
| Solyc02g091340.2 | Solyc02g091340.4 |
| Solyc02g091350.2 | Solyc02g091350.4 |
| Solyc02g091430.2 | Solyc02g091430.4 |
| Solyc02g091530.2 | Solyc02g091530.4 |
| Solyc02g091580.2 | Solyc02g091580.4 |
| Solyc02g091590.2 | Solyc02g091590.4 |
| Solyc02g091650.2 | Solyc02g091650.4 |
| Solyc02g091680.2 | Solyc02g091680.4 |
| Solyc02g091730.1 | Solyc02g091730.3 |
| Solyc02g091770.1 | Solyc02g091770.3 |
| Solyc02g091800.2 | Solyc02g091800.4 |
| Solyc02g091820.1 | Solyc02g091820.3 |
| Solyc02g091880.2 | Solyc02g091880.4 |
| Solyc02g091890.1 | Solyc02g091890.3 |
| Solyc02g091900.2 | Solyc02g091900.4 |
| Solyc02g091950.2 | Solyc02g091950.4 |
| Solyc02g091960.2 | Solyc02g091960.4 |
| Solyc02g091970.2 | Solyc02g091970.4 |
| Solyc02g092010.2 | Solyc02g092010.4 |
| Solyc02g092020.1 | Solyc02g092020.3 |
| Solyc02g092030.1 | Solyc02g092030.3 |
| Solyc02g092090.1 | Solyc02g092090.3 |
| Solyc02g092290.2 | Solyc02g092290.4 |
| Solyc02g092310.2 | Solyc02g092310.4 |
| Solyc02g092380.2 | Solyc02g092380.4 |
| Solyc02g092420.2 | Solyc02g092420.4 |
| Solyc02g092480.2 | Solyc02g092480.4 |
| Solyc02g092520.1 | Solyc02g092520.3 |
| Solyc02g092530.2 | Solyc02g092530.4 |
| Solyc02g092640.2 | Solyc02g092640.4 |
| Solyc02g092730.2 | Solyc02g092730.4 |

| | |
|---|---|
| Solyc02g092780.1 | Solyc02g092780.3 |
| Solyc02g092810.2 | Solyc02g092810.4 |
| Solyc02g092820.2 | Solyc02g092820.4 |
| Solyc02g092920.2 | Solyc02g092920.4 |
| Solyc02g093040.2 | Solyc02g093040.4 |
| Solyc02g093120.2 | Solyc02g093120.4 |
| Solyc02g093130.1 | Solyc02g093130.3 |
| Solyc02g093140.2 | Solyc02g093140.4 |
| Solyc02g093170.2 | Solyc02g093170.4 |
| Solyc02g093230.2 | Solyc02g093230.4 |
| Solyc02g093240.1 | Solyc02g093240.3 |
| Solyc02g093270.2 | Solyc02g093270.4 |
| Solyc02g093320.2 | Solyc02g093320.4 |
| Solyc02g093330.2 | Solyc02g093330.4 |
| Solyc02g093390.2 | Solyc02g093390.4 |
| Solyc02g093400.1 | Solyc02g093400.3 |
| Solyc02g093410.2 | Solyc02g093410.4 |
| Solyc02g093420.2 | Solyc02g093420.4 |
| Solyc02g093440.2 | Solyc02g093440.4 |
| Solyc02g093460.2 | Solyc02g093460.4 |
| Solyc02g093470.2 | Solyc02g093470.4 |
| Solyc02g093480.2 | Solyc02g093480.4 |
| Solyc02g093500.2 | Solyc02g093500.4 |
| Solyc02g093530.2 | Solyc02g093530.4 |
| Solyc02g093540.2 | Solyc02g093540.4 |
| Solyc02g093580.2 | Solyc02g093580.4 |
| Solyc02g093650.2 | Solyc02g093650.4 |
| Solyc02g093730.2 | Solyc02g093730.4 |
| Solyc02g093760.2 | Solyc02g093760.4 |
| Solyc02g093820.2 | Solyc02g093820.4 |
| Solyc02g093830.2 | Solyc02g093830.4 |
| Solyc02g093870.2 | Solyc02g093870.4 |
| Solyc02g093930.2 | Solyc02g093930.4 |
| Solyc02g093970.2 | Solyc02g093970.4 |
| Solyc02g093990.2 | Solyc02g093990.4 |
| Solyc02g094050.2 | Solyc02g094050.4 |
| Solyc02g094090.2 | Solyc02g094090.4 |
| Solyc02g094100.2 | Solyc02g094100.4 |
| Solyc02g094140.2 | Solyc02g094140.4 |
| Solyc02g094150.2 | Solyc02g094150.4 |
| Solyc02g094250.2 | Solyc02g094250.4 |
| Solyc02g094280.2 | Solyc02g094280.4 |
| Solyc02g094300.2 | Solyc02g094300.4 |

| | |
|---|---|
| Solyc02g094310.1 | Solyc02g094310.3 |
| Solyc02g094380.1 | Solyc02g094380.3 |
| Solyc02g094400.2 | Solyc02g094400.4 |
| Solyc02g094410.2 | Solyc02g094410.4 |
| Solyc02g094430.2 | Solyc02g094430.4 |
| Solyc02g094440.2 | Solyc02g094440.4 |
| Solyc02g094480.2 | Solyc02g094480.4 |
| Solyc02g094510.2 | Solyc02g094510.4 |
| Solyc02g094640.2 | Solyc02g094640.4 |
| Solyc02g094860.1 | Solyc02g094860.3 |
| Solyc03g005000.2 | Solyc03g005000.4 |
| Solyc03g005010.2 | Solyc03g005010.4 |
| Solyc03g005060.2 | Solyc03g005060.4 |
| Solyc03g005080.2 | Solyc03g005080.4 |
| Solyc03g005090.2 | Solyc03g005090.4 |
| Solyc03g005130.1 | Solyc03g005130.3 |
| Solyc03g005160.2 | Solyc03g005160.4 |
| Solyc03g005210.2 | Solyc03g005210.4 |
| Solyc03g005220.2 | Solyc03g005220.4 |
| Solyc03g005250.2 | Solyc03g005250.4 |
| Solyc03g005280.2 | Solyc03g005280.4 |
| Solyc03g005310.2 | Solyc03g005310.4 |
| Solyc03g005340.2 | Solyc03g005340.4 |
| Solyc03g005410.2 | Solyc03g005410.4 |
| Solyc03g005430.2 | Solyc03g005430.4 |
| Solyc03g005460.2 | Solyc03g005460.4 |
| Solyc03g005480.2 | Solyc03g005480.4 |
| Solyc03g005490.2 | Solyc03g005490.4 |
| Solyc03g005610.2 | Solyc03g005610.4 |
| Solyc03g005620.2 | Solyc03g005620.4 |
| Solyc03g005660.2 | Solyc03g005660.4 |
| Solyc03g005710.1 | Solyc03g005710.3 |
| Solyc03g005730.2 | Solyc03g005730.4 |
| Solyc03g005740.1 | Solyc03g005740.3 |
| Solyc03g005800.2 | Solyc03g005800.4 |
| Solyc03g005810.2 | Solyc03g005810.4 |
| Solyc03g005830.2 | Solyc03g005830.4 |
| Solyc03g005840.1 | Solyc03g005840.3 |
| Solyc03g005860.2 | Solyc03g005860.4 |
| Solyc03g005870.2 | Solyc03g005870.4 |
| Solyc03g005900.2 | Solyc03g005900.4 |
| Solyc03g005920.2 | Solyc03g005920.4 |
| Solyc03g005940.2 | Solyc03g005940.4 |

| | |
|---|---|
| Solyc03g005970.2 | Solyc03g005970.4 |
| Solyc03g006030.2 | Solyc03g006030.4 |
| Solyc03g006050.2 | Solyc03g006050.4 |
| Solyc03g006080.2 | Solyc03g006080.4 |
| Solyc03g006100.2 | Solyc03g006100.4 |
| Solyc03g006120.2 | Solyc03g006120.4 |
| Solyc03g006170.2 | Solyc03g006170.4 |
| Solyc03g006200.1 | Solyc03g006200.3 |
| Solyc03g006220.2 | Solyc03g006220.4 |
| Solyc03g006260.2 | Solyc03g006260.4 |
| Solyc03g006290.2 | Solyc03g006290.4 |
| Solyc03g006300.1 | Solyc03g006300.3 |
| Solyc03g006310.2 | Solyc03g006310.4 |
| Solyc03g006330.2 | Solyc03g006330.4 |
| Solyc03g006380.2 | Solyc03g006380.4 |
| Solyc03g006420.2 | Solyc03g006420.4 |
| Solyc03g006440.1 | Solyc03g006440.3 |
| Solyc03g006460.2 | Solyc03g006460.4 |
| Solyc03g006480.1 | Solyc03g006480.3 |
| Solyc03g006510.2 | Solyc03g006510.4 |
| Solyc03g006550.2 | Solyc03g006550.4 |
| Solyc03g006670.2 | Solyc03g006670.4 |
| Solyc03g006730.1 | Solyc03g006730.3 |
| Solyc03g006740.1 | Solyc03g006740.3 |
| Solyc03g006760.2 | Solyc03g006760.4 |
| Solyc03g006770.2 | Solyc03g006770.4 |
| Solyc03g006790.1 | Solyc03g006790.3 |
| Solyc03g006840.2 | Solyc03g006840.4 |
| Solyc03g006890.1 | Solyc03g006890.3 |
| Solyc03g006930.1 | Solyc03g006930.3 |
| Solyc03g006940.1 | Solyc03g006940.3 |
| Solyc03g006950.1 | Solyc03g006950.3 |
| Solyc03g006960.2 | Solyc03g006960.4 |
| Solyc03g006980.2 | Solyc03g006980.4 |
| Solyc03g007000.2 | Solyc03g007000.4 |
| Solyc03g007010.2 | Solyc03g007010.4 |
| Solyc03g007040.2 | Solyc03g007040.4 |
| Solyc03g007080.2 | Solyc03g007080.4 |
| Solyc03g007130.2 | Solyc03g007130.4 |
| Solyc03g007230.2 | Solyc03g007230.4 |
| Solyc03g007280.2 | Solyc03g007280.4 |
| Solyc03g007290.2 | Solyc03g007290.4 |
| Solyc03g007320.2 | Solyc03g007320.4 |

| | |
|---|---|
| Solyc03g007390.2 | Solyc03g007390.4 |
| Solyc03g007410.2 | Solyc03g007410.4 |
| Solyc03g007440.2 | Solyc03g007440.4 |
| Solyc03g007460.1 | Solyc03g007460.3 |
| Solyc03g007470.2 | Solyc03g007470.4 |
| Solyc03g007500.2 | Solyc03g007500.4 |
| Solyc03g007510.2 | Solyc03g007510.4 |
| Solyc03g007590.2 | Solyc03g007590.4 |
| Solyc03g007610.2 | Solyc03g007610.4 |
| Solyc03g007660.2 | Solyc03g007660.4 |
| Solyc03g007670.2 | Solyc03g007670.4 |
| Solyc03g007880.1 | Solyc03g007880.3 |
| Solyc03g007910.1 | Solyc03g007910.3 |
| Solyc03g007920.2 | Solyc03g007920.4 |
| Solyc03g007930.2 | Solyc03g007930.4 |
| Solyc03g007940.2 | Solyc03g007940.4 |
| Solyc03g013160.2 | Solyc03g013160.4 |
| Solyc03g013430.1 | Solyc03g013430.3 |
| Solyc03g013440.2 | Solyc03g013440.4 |
| Solyc03g013450.1 | Solyc03g013450.3 |
| Solyc03g013630.2 | Solyc03g013630.4 |
| Solyc03g019660.2 | Solyc03g019660.4 |
| Solyc03g019730.2 | Solyc03g019730.4 |
| Solyc03g019760.2 | Solyc03g019760.4 |
| Solyc03g019770.2 | Solyc03g019770.4 |
| Solyc03g019780.2 | Solyc03g019780.4 |
| Solyc03g019800.2 | Solyc03g019800.4 |
| Solyc03g019830.2 | Solyc03g019830.4 |
| Solyc03g019870.2 | Solyc03g019870.4 |
| Solyc03g019940.2 | Solyc03g019940.4 |
| Solyc03g020070.2 | Solyc03g020070.4 |
| Solyc03g020080.2 | Solyc03g020080.4 |
| Solyc03g025150.2 | Solyc03g025150.4 |
| Solyc03g025160.2 | Solyc03g025160.4 |
| Solyc03g025240.1 | Solyc03g025240.3 |
| Solyc03g025250.2 | Solyc03g025250.4 |
| Solyc03g025280.2 | Solyc03g025280.4 |
| Solyc03g025320.2 | Solyc03g025320.4 |
| Solyc03g025330.2 | Solyc03g025330.4 |
| Solyc03g025410.2 | Solyc03g025410.4 |
| Solyc03g025450.2 | Solyc03g025450.4 |
| Solyc03g025470.2 | Solyc03g025470.4 |
| Solyc03g025480.2 | Solyc03g025480.4 |

| | |
|---|---|
| Solyc03g025500.2 | Solyc03g025500.4 |
| Solyc03g025520.2 | Solyc03g025520.4 |
| Solyc03g025560.2 | Solyc03g025560.4 |
| Solyc03g025610.1 | Solyc03g025610.3 |
| Solyc03g025740.2 | Solyc03g025740.4 |
| Solyc03g025750.2 | Solyc03g025750.4 |
| Solyc03g025760.2 | Solyc03g025760.4 |
| Solyc03g025770.2 | Solyc03g025770.4 |
| Solyc03g025790.2 | Solyc03g025790.4 |
| Solyc03g025810.2 | Solyc03g025810.4 |
| Solyc03g025830.2 | Solyc03g025830.4 |
| Solyc03g025920.2 | Solyc03g025920.4 |
| Solyc03g025930.2 | Solyc03g025930.4 |
| Solyc03g025940.1 | Solyc03g025940.3 |
| Solyc03g025950.2 | Solyc03g025950.4 |
| Solyc03g026000.2 | Solyc03g026000.4 |
| Solyc03g026040.2 | Solyc03g026040.4 |
| Solyc03g026070.2 | Solyc03g026070.4 |
| Solyc03g026090.2 | Solyc03g026090.4 |
| Solyc03g026110.2 | Solyc03g026110.4 |
| Solyc03g026140.2 | Solyc03g026140.4 |
| Solyc03g026160.2 | Solyc03g026160.4 |
| Solyc03g026170.2 | Solyc03g026170.4 |
| Solyc03g026200.2 | Solyc03g026200.4 |
| Solyc03g026210.2 | Solyc03g026210.4 |
| Solyc03g026240.2 | Solyc03g026240.4 |
| Solyc03g026250.1 | Solyc03g026250.3 |
| Solyc03g026300.2 | Solyc03g026300.4 |
| Solyc03g026310.2 | Solyc03g026310.4 |
| Solyc03g026320.2 | Solyc03g026320.4 |
| Solyc03g026340.2 | Solyc03g026340.4 |
| Solyc03g026400.1 | Solyc03g026400.3 |
| Solyc03g026410.1 | Solyc03g026410.3 |
| Solyc03g031440.2 | Solyc03g031440.4 |
| Solyc03g031510.2 | Solyc03g031510.4 |
| Solyc03g031520.2 | Solyc03g031520.4 |
| Solyc03g031530.2 | Solyc03g031530.4 |
| Solyc03g031560.2 | Solyc03g031560.4 |
| Solyc03g031570.2 | Solyc03g031570.4 |
| Solyc03g031580.1 | Solyc03g031580.3 |
| Solyc03g031590.2 | Solyc03g031590.4 |
| Solyc03g031650.2 | Solyc03g031650.4 |
| Solyc03g031700.2 | Solyc03g031700.4 |

| | |
|---|---|
| Solyc03g031710.1 | Solyc03g031710.3 |
| Solyc03g031740.1 | Solyc03g031740.3 |
| Solyc03g031760.2 | Solyc03g031760.4 |
| Solyc03g031780.2 | Solyc03g031780.4 |
| Solyc03g031820.2 | Solyc03g031820.4 |
| Solyc03g031870.1 | Solyc03g031870.3 |
| Solyc03g031940.2 | Solyc03g031940.4 |
| Solyc03g031970.2 | Solyc03g031970.4 |
| Solyc03g031980.2 | Solyc03g031980.4 |
| Solyc03g031990.2 | Solyc03g031990.4 |
| Solyc03g032000.2 | Solyc03g032000.4 |
| Solyc03g032020.2 | Solyc03g032020.4 |
| Solyc03g032030.2 | Solyc03g032030.4 |
| Solyc03g032050.2 | Solyc03g032050.4 |
| Solyc03g032080.2 | Solyc03g032080.4 |
| Solyc03g032130.2 | Solyc03g032130.4 |
| Solyc03g032140.2 | Solyc03g032140.4 |
| Solyc03g032240.2 | Solyc03g032240.4 |
| Solyc03g033300.1 | Solyc03g033300.3 |
| Solyc03g033310.2 | Solyc03g033310.4 |
| Solyc03g033320.2 | Solyc03g033320.4 |
| Solyc03g033360.1 | Solyc03g033360.3 |
| Solyc03g033400.2 | Solyc03g033400.4 |
| Solyc03g033410.2 | Solyc03g033410.4 |
| Solyc03g033420.2 | Solyc03g033420.4 |
| Solyc03g033450.2 | Solyc03g033450.4 |
| Solyc03g033480.2 | Solyc03g033480.4 |
| Solyc03g033490.1 | Solyc03g033490.3 |
| Solyc03g033500.2 | Solyc03g033500.4 |
| Solyc03g033530.2 | Solyc03g033530.4 |
| Solyc03g033660.2 | Solyc03g033660.4 |
| Solyc03g033740.2 | Solyc03g033740.4 |
| Solyc03g033760.1 | Solyc03g033760.3 |
| Solyc03g033790.2 | Solyc03g033790.4 |
| Solyc03g033840.2 | Solyc03g033840.4 |
| Solyc03g033950.2 | Solyc03g033950.4 |
| Solyc03g033980.2 | Solyc03g033980.4 |
| Solyc03g034020.2 | Solyc03g034020.4 |
| Solyc03g034060.2 | Solyc03g034060.4 |
| Solyc03g034080.2 | Solyc03g034080.4 |
| Solyc03g034100.2 | Solyc03g034100.4 |
| Solyc03g034110.2 | Solyc03g034110.4 |
| Solyc03g034130.2 | Solyc03g034130.4 |

| | |
|---|---|
| Solyc03g034140.2 | Solyc03g034140.4 |
| Solyc03g034160.2 | Solyc03g034160.4 |
| Solyc03g034240.2 | Solyc03g034240.4 |
| Solyc03g034450.2 | Solyc03g034450.4 |
| Solyc03g042560.1 | Solyc03g042560.3 |
| Solyc03g043580.1 | Solyc03g043580.3 |
| Solyc03g043600.1 | Solyc03g043600.3 |
| Solyc03g043720.2 | Solyc03g043720.4 |
| Solyc03g043740.2 | Solyc03g043740.4 |
| Solyc03g043760.2 | Solyc03g043760.4 |
| Solyc03g043770.2 | Solyc03g043770.4 |
| Solyc03g043920.2 | Solyc03g043920.4 |
| Solyc03g043930.2 | Solyc03g043930.4 |
| Solyc03g043940.2 | Solyc03g043940.4 |
| Solyc03g043950.2 | Solyc03g043950.4 |
| Solyc03g044010.2 | Solyc03g044010.4 |
| Solyc03g044020.2 | Solyc03g044020.4 |
| Solyc03g044060.2 | Solyc03g044060.4 |
| Solyc03g044090.2 | Solyc03g044090.4 |
| Solyc03g044100.2 | Solyc03g044100.4 |
| Solyc03g044150.2 | Solyc03g044150.4 |
| Solyc03g044160.1 | Solyc03g044160.3 |
| Solyc03g044210.2 | Solyc03g044210.4 |
| Solyc03g044280.1 | Solyc03g044280.3 |
| Solyc03g044350.1 | Solyc03g044350.3 |
| Solyc03g044470.2 | Solyc03g044470.4 |
| Solyc03g044620.2 | Solyc03g044620.4 |
| Solyc03g044710.2 | Solyc03g044710.4 |
| Solyc03g044740.2 | Solyc03g044740.4 |
| Solyc03g044820.2 | Solyc03g044820.4 |
| Solyc03g044830.2 | Solyc03g044830.4 |
| Solyc03g044880.2 | Solyc03g044880.4 |
| Solyc03g044930.1 | Solyc03g044930.3 |
| Solyc03g045020.2 | Solyc03g045020.4 |
| Solyc03g045050.2 | Solyc03g045050.4 |
| Solyc03g045080.1 | Solyc03g045080.3 |
| Solyc03g045140.2 | Solyc03g045140.4 |
| Solyc03g046270.2 | Solyc03g046270.4 |
| Solyc03g046300.1 | Solyc03g046300.3 |
| Solyc03g046320.2 | Solyc03g046320.4 |
| Solyc03g046360.1 | Solyc03g046360.3 |
| Solyc03g046440.1 | Solyc03g046440.3 |
| Solyc03g046490.1 | Solyc03g046490.3 |

| | |
|---|---|
| Solyc03g046570.2 | Solyc03g046570.4 |
| Solyc03g051620.1 | Solyc03g051620.3 |
| Solyc03g051640.1 | Solyc03g051640.3 |
| Solyc03g051660.1 | Solyc03g051660.3 |
| Solyc03g051700.1 | Solyc03g051700.3 |
| Solyc03g051780.1 | Solyc03g051780.3 |
| Solyc03g051810.2 | Solyc03g051810.4 |
| Solyc03g051900.2 | Solyc03g051900.4 |
| Solyc03g051920.2 | Solyc03g051920.4 |
| Solyc03g051950.2 | Solyc03g051950.4 |
| Solyc03g051960.2 | Solyc03g051960.4 |
| Solyc03g053060.1 | Solyc03g053060.3 |
| Solyc03g053080.1 | Solyc03g053080.3 |
| Solyc03g058210.1 | Solyc03g058210.3 |
| Solyc03g058350.2 | Solyc03g058350.4 |
| Solyc03g058400.2 | Solyc03g058400.4 |
| Solyc03g058420.1 | Solyc03g058420.3 |
| Solyc03g058430.1 | Solyc03g058430.3 |
| Solyc03g058450.2 | Solyc03g058450.4 |
| Solyc03g058470.1 | Solyc03g058470.3 |
| Solyc03g058880.2 | Solyc03g058880.4 |
| Solyc03g058890.2 | Solyc03g058890.4 |
| Solyc03g058900.1 | Solyc03g058900.3 |
| Solyc03g058930.2 | Solyc03g058930.4 |
| Solyc03g058940.2 | Solyc03g058940.4 |
| Solyc03g058970.2 | Solyc03g058970.4 |
| Solyc03g059010.2 | Solyc03g059010.4 |
| Solyc03g059020.1 | Solyc03g059020.3 |
| Solyc03g059030.2 | Solyc03g059030.4 |
| Solyc03g059070.2 | Solyc03g059070.4 |
| Solyc03g059080.1 | Solyc03g059080.3 |
| Solyc03g059100.1 | Solyc03g059100.3 |
| Solyc03g059140.2 | Solyc03g059140.4 |
| Solyc03g059160.1 | Solyc03g059160.3 |
| Solyc03g059180.2 | Solyc03g059180.4 |
| Solyc03g059190.2 | Solyc03g059190.4 |
| Solyc03g059260.2 | Solyc03g059260.4 |
| Solyc03g059270.2 | Solyc03g059270.4 |
| Solyc03g059420.2 | Solyc03g059420.4 |
| Solyc03g060510.2 | Solyc03g060510.4 |
| Solyc03g061560.1 | Solyc03g061560.3 |
| Solyc03g061590.2 | Solyc03g061590.4 |
| Solyc03g061600.2 | Solyc03g061600.4 |

| | |
|---|---|
| Solyc03g061650.1 | Solyc03g061650.3 |
| Solyc03g062660.2 | Solyc03g062660.4 |
| Solyc03g062710.2 | Solyc03g062710.4 |
| Solyc03g062740.2 | Solyc03g062740.4 |
| Solyc03g063030.2 | Solyc03g063030.4 |
| Solyc03g063040.2 | Solyc03g063040.4 |
| Solyc03g063140.2 | Solyc03g063140.4 |
| Solyc03g063260.2 | Solyc03g063260.4 |
| Solyc03g063380.1 | Solyc03g063380.3 |
| Solyc03g063680.1 | Solyc03g063680.3 |
| Solyc03g063720.2 | Solyc03g063720.4 |
| Solyc03g063730.2 | Solyc03g063730.4 |
| Solyc03g063760.2 | Solyc03g063760.4 |
| Solyc03g063770.1 | Solyc03g063770.3 |
| Solyc03g064010.2 | Solyc03g064010.4 |
| Solyc03g065200.2 | Solyc03g065200.4 |
| Solyc03g065250.2 | Solyc03g065250.4 |
| Solyc03g065350.1 | Solyc03g065350.3 |
| Solyc03g070400.1 | Solyc03g070400.3 |
| Solyc03g070410.1 | Solyc03g070410.3 |
| Solyc03g070440.2 | Solyc03g070440.4 |
| Solyc03g071550.1 | Solyc03g071550.3 |
| Solyc03g071560.2 | Solyc03g071560.4 |
| Solyc03g071690.2 | Solyc03g071690.4 |
| Solyc03g071720.2 | Solyc03g071720.4 |
| Solyc03g077910.2 | Solyc03g077910.4 |
| Solyc03g077950.1 | Solyc03g077950.3 |
| Solyc03g078020.2 | Solyc03g078020.4 |
| Solyc03g078060.2 | Solyc03g078060.4 |
| Solyc03g078070.1 | Solyc03g078070.3 |
| Solyc03g078080.2 | Solyc03g078080.4 |
| Solyc03g078090.2 | Solyc03g078090.4 |
| Solyc03g078120.2 | Solyc03g078120.4 |
| Solyc03g078200.1 | Solyc03g078200.3 |
| Solyc03g078210.2 | Solyc03g078210.4 |
| Solyc03g078240.1 | Solyc03g078240.3 |
| Solyc03g078280.1 | Solyc03g078280.3 |
| Solyc03g078300.1 | Solyc03g078300.3 |
| Solyc03g078420.2 | Solyc03g078420.4 |
| Solyc03g078480.1 | Solyc03g078480.3 |
| Solyc03g078490.2 | Solyc03g078490.4 |
| Solyc03g078520.2 | Solyc03g078520.4 |
| Solyc03g078600.1 | Solyc03g078600.3 |

| | |
|---|---|
| Solyc03g078650.2 | Solyc03g078650.4 |
| Solyc03g078670.1 | Solyc03g078670.3 |
| Solyc03g078690.2 | Solyc03g078690.4 |
| Solyc03g078710.1 | Solyc03g078710.3 |
| Solyc03g078720.2 | Solyc03g078720.4 |
| Solyc03g078730.1 | Solyc03g078730.3 |
| Solyc03g078780.1 | Solyc03g078780.3 |
| Solyc03g078810.2 | Solyc03g078810.4 |
| Solyc03g079850.2 | Solyc03g079850.4 |
| Solyc03g079880.2 | Solyc03g079880.4 |
| Solyc03g079890.1 | Solyc03g079890.3 |
| Solyc03g079950.2 | Solyc03g079950.4 |
| Solyc03g079960.2 | Solyc03g079960.4 |
| Solyc03g080020.2 | Solyc03g080020.4 |
| Solyc03g080030.2 | Solyc03g080030.4 |
| Solyc03g080070.2 | Solyc03g080070.4 |
| Solyc03g080080.2 | Solyc03g080080.4 |
| Solyc03g080090.2 | Solyc03g080090.4 |
| Solyc03g080100.2 | Solyc03g080100.4 |
| Solyc03g080180.2 | Solyc03g080180.4 |
| Solyc03g081250.1 | Solyc03g081250.3 |
| Solyc03g081260.2 | Solyc03g081260.4 |
| Solyc03g081300.2 | Solyc03g081300.4 |
| Solyc03g082380.2 | Solyc03g082380.4 |
| Solyc03g082470.2 | Solyc03g082470.4 |
| Solyc03g082480.2 | Solyc03g082480.4 |
| Solyc03g082500.2 | Solyc03g082500.4 |
| Solyc03g082570.2 | Solyc03g082570.4 |
| Solyc03g082590.2 | Solyc03g082590.4 |
| Solyc03g082600.2 | Solyc03g082600.4 |
| Solyc03g082620.2 | Solyc03g082620.4 |
| Solyc03g082640.1 | Solyc03g082640.3 |
| Solyc03g082700.2 | Solyc03g082700.4 |
| Solyc03g082710.2 | Solyc03g082710.4 |
| Solyc03g082780.1 | Solyc03g082780.3 |
| Solyc03g082790.2 | Solyc03g082790.4 |
| Solyc03g082870.2 | Solyc03g082870.4 |
| Solyc03g082910.2 | Solyc03g082910.4 |
| Solyc03g082920.2 | Solyc03g082920.4 |
| Solyc03g082970.2 | Solyc03g082970.4 |
| Solyc03g083070.1 | Solyc03g083070.3 |
| Solyc03g083090.2 | Solyc03g083090.4 |
| Solyc03g083130.1 | Solyc03g083130.3 |

| | |
|---|---|
| Solyc03g083160.1 | Solyc03g083160.3 |
| Solyc03g083240.2 | Solyc03g083240.4 |
| Solyc03g083270.2 | Solyc03g083270.4 |
| Solyc03g083280.2 | Solyc03g083280.4 |
| Solyc03g083290.2 | Solyc03g083290.4 |
| Solyc03g083310.2 | Solyc03g083310.4 |
| Solyc03g083320.2 | Solyc03g083320.4 |
| Solyc03g083340.1 | Solyc03g083340.3 |
| Solyc03g083380.2 | Solyc03g083380.4 |
| Solyc03g083390.2 | Solyc03g083390.4 |
| Solyc03g083440.2 | Solyc03g083440.4 |
| Solyc03g083480.2 | Solyc03g083480.4 |
| Solyc03g083510.2 | Solyc03g083510.4 |
| Solyc03g083520.2 | Solyc03g083520.4 |
| Solyc03g083610.2 | Solyc03g083610.4 |
| Solyc03g083650.2 | Solyc03g083650.4 |
| Solyc03g083670.1 | Solyc03g083670.3 |
| Solyc03g083810.2 | Solyc03g083810.4 |
| Solyc03g083820.1 | Solyc03g083820.3 |
| Solyc03g083870.2 | Solyc03g083870.4 |
| Solyc03g083880.2 | Solyc03g083880.4 |
| Solyc03g083900.2 | Solyc03g083900.4 |
| Solyc03g083910.2 | Solyc03g083910.5 |
| Solyc03g093150.1 | Solyc03g093150.3 |
| Solyc03g093230.2 | Solyc03g093230.4 |
| Solyc03g093240.2 | Solyc03g093240.4 |
| Solyc03g093280.1 | Solyc03g093280.3 |
| Solyc03g093330.2 | Solyc03g093330.4 |
| Solyc03g093350.2 | Solyc03g093350.4 |
| Solyc03g093390.2 | Solyc03g093390.4 |
| Solyc03g093440.2 | Solyc03g093440.4 |
| Solyc03g093630.1 | Solyc03g093630.3 |
| Solyc03g093640.2 | Solyc03g093640.4 |
| Solyc03g093680.2 | Solyc03g093680.4 |
| Solyc03g093700.1 | Solyc03g093700.3 |
| Solyc03g093720.1 | Solyc03g093720.3 |
| Solyc03g093750.1 | Solyc03g093750.3 |
| Solyc03g093840.1 | Solyc03g093840.3 |
| Solyc03g093930.2 | Solyc03g093930.4 |
| Solyc03g093990.2 | Solyc03g093990.4 |
| Solyc03g094030.2 | Solyc03g094030.4 |
| Solyc03g094040.2 | Solyc03g094040.4 |
| Solyc03g094050.2 | Solyc03g094050.4 |

| | |
|---|---|
| Solyc03g094060.2 | Solyc03g094060.4 |
| Solyc03g094070.2 | Solyc03g094070.4 |
| Solyc03g094080.2 | Solyc03g094080.4 |
| Solyc03g094110.2 | Solyc03g094110.4 |
| Solyc03g094170.1 | Solyc03g094170.3 |
| Solyc03g095190.2 | Solyc03g095190.4 |
| Solyc03g095300.2 | Solyc03g095300.4 |
| Solyc03g095350.1 | Solyc03g095350.3 |
| Solyc03g095420.1 | Solyc03g095420.3 |
| Solyc03g095610.2 | Solyc03g095610.4 |
| Solyc03g095620.2 | Solyc03g095620.4 |
| Solyc03g095660.1 | Solyc03g095660.3 |
| Solyc03g095710.2 | Solyc03g095710.4 |
| Solyc03g095730.2 | Solyc03g095730.4 |
| Solyc03g095780.1 | Solyc03g095780.3 |
| Solyc03g095820.2 | Solyc03g095820.4 |
| Solyc03g095830.1 | Solyc03g095830.3 |
| Solyc03g095870.1 | Solyc03g095870.3 |
| Solyc03g095900.2 | Solyc03g095900.4 |
| Solyc03g096080.2 | Solyc03g096080.4 |
| Solyc03g096090.2 | Solyc03g096090.4 |
| Solyc03g096110.1 | Solyc03g096110.3 |
| Solyc03g096150.1 | Solyc03g096150.3 |
| Solyc03g096160.1 | Solyc03g096160.3 |
| Solyc03g096180.2 | Solyc03g096180.4 |
| Solyc03g096260.1 | Solyc03g096260.3 |
| Solyc03g096360.2 | Solyc03g096360.4 |
| Solyc03g096450.1 | Solyc03g096450.3 |
| Solyc03g096460.2 | Solyc03g096460.4 |
| Solyc03g096620.1 | Solyc03g096620.3 |
| Solyc03g096640.2 | Solyc03g096640.4 |
| Solyc03g096700.1 | Solyc03g096700.3 |
| Solyc03g096740.2 | Solyc03g096740.4 |
| Solyc03g096800.2 | Solyc03g096800.4 |
| Solyc03g096820.2 | Solyc03g096820.4 |
| Solyc03g096830.2 | Solyc03g096830.4 |
| Solyc03g096850.2 | Solyc03g096850.4 |
| Solyc03g096880.2 | Solyc03g096880.4 |
| Solyc03g096890.1 | Solyc03g096890.3 |
| Solyc03g096900.2 | Solyc03g096900.4 |
| Solyc03g096930.2 | Solyc03g096930.4 |
| Solyc03g096940.2 | Solyc03g096940.4 |
| Solyc03g096950.2 | Solyc03g096950.4 |

| | |
|---|---|
| Solyc03g096990.2 | Solyc03g096990.4 |
| Solyc03g097020.2 | Solyc03g097020.4 |
| Solyc03g097060.2 | Solyc03g097060.4 |
| Solyc03g097080.2 | Solyc03g097080.4 |
| Solyc03g097090.2 | Solyc03g097090.4 |
| Solyc03g097110.2 | Solyc03g097110.4 |
| Solyc03g097160.2 | Solyc03g097160.4 |
| Solyc03g097250.2 | Solyc03g097250.4 |
| Solyc03g097260.2 | Solyc03g097260.4 |
| Solyc03g097280.2 | Solyc03g097280.4 |
| Solyc03g097290.2 | Solyc03g097290.4 |
| Solyc03g097300.2 | Solyc03g097300.4 |
| Solyc03g097350.2 | Solyc03g097350.4 |
| Solyc03g097370.2 | Solyc03g097370.4 |
| Solyc03g097380.2 | Solyc03g097380.4 |
| Solyc03g097450.2 | Solyc03g097450.4 |
| Solyc03g097490.2 | Solyc03g097490.4 |
| Solyc03g097580.2 | Solyc03g097580.4 |
| Solyc03g097590.2 | Solyc03g097590.4 |
| Solyc03g097670.2 | Solyc03g097670.4 |
| Solyc03g097680.2 | Solyc03g097680.4 |
| Solyc03g097720.2 | Solyc03g097720.4 |
| Solyc03g097750.2 | Solyc03g097750.4 |
| Solyc03g097800.2 | Solyc03g097800.4 |
| Solyc03g097870.2 | Solyc03g097870.4 |
| Solyc03g097910.2 | Solyc03g097910.4 |
| Solyc03g097950.2 | Solyc03g097950.4 |
| Solyc03g097980.2 | Solyc03g097980.4 |
| Solyc03g098020.2 | Solyc03g098020.4 |
| Solyc03g098060.1 | Solyc03g098060.3 |
| Solyc03g098100.2 | Solyc03g098100.4 |
| Solyc03g098200.2 | Solyc03g098200.4 |
| Solyc03g098210.2 | Solyc03g098210.4 |
| Solyc03g098220.2 | Solyc03g098220.4 |
| Solyc03g098250.2 | Solyc03g098250.4 |
| Solyc03g098280.2 | Solyc03g098280.4 |
| Solyc03g098290.2 | Solyc03g098290.4 |
| Solyc03g098320.2 | Solyc03g098320.4 |
| Solyc03g098340.2 | Solyc03g098340.4 |
| Solyc03g098430.2 | Solyc03g098430.4 |
| Solyc03g098440.2 | Solyc03g098440.4 |
| Solyc03g098500.2 | Solyc03g098500.4 |
| Solyc03g098630.2 | Solyc03g098630.4 |

| | |
|---|---|
| Solyc03g098640.2 | Solyc03g098640.4 |
| Solyc03g098760.1 | Solyc03g098760.3 |
| Solyc03g104790.1 | Solyc03g104790.3 |
| Solyc03g110840.1 | Solyc03g110840.3 |
| Solyc03g110880.2 | Solyc03g110880.4 |
| Solyc03g110900.2 | Solyc03g110900.4 |
| Solyc03g110910.2 | Solyc03g110910.4 |
| Solyc03g111010.2 | Solyc03g111010.4 |
| Solyc03g111090.2 | Solyc03g111090.4 |
| Solyc03g111150.2 | Solyc03g111150.4 |
| Solyc03g111190.1 | Solyc03g111190.3 |
| Solyc03g111210.2 | Solyc03g111210.4 |
| Solyc03g111230.2 | Solyc03g111230.4 |
| Solyc03g111330.2 | Solyc03g111330.4 |
| Solyc03g111340.2 | Solyc03g111340.4 |
| Solyc03g111390.2 | Solyc03g111390.4 |
| Solyc03g111450.1 | Solyc03g111450.3 |
| Solyc03g111460.1 | Solyc03g111460.3 |
| Solyc03g111470.1 | Solyc03g111470.3 |
| Solyc03g111520.1 | Solyc03g111520.3 |
| Solyc03g111530.2 | Solyc03g111530.4 |
| Solyc03g111550.2 | Solyc03g111550.4 |
| Solyc03g111570.2 | Solyc03g111570.4 |
| Solyc03g111590.2 | Solyc03g111590.4 |
| Solyc03g111600.2 | Solyc03g111600.4 |
| Solyc03g111630.2 | Solyc03g111630.4 |
| Solyc03g111640.2 | Solyc03g111640.4 |
| Solyc03g111650.2 | Solyc03g111650.4 |
| Solyc03g111690.2 | Solyc03g111690.4 |
| Solyc03g111750.1 | Solyc03g111750.3 |
| Solyc03g111790.2 | Solyc03g111790.4 |
| Solyc03g111830.2 | Solyc03g111830.4 |
| Solyc03g111930.2 | Solyc03g111930.4 |
| Solyc03g111970.2 | Solyc03g111970.4 |
| Solyc03g111990.2 | Solyc03g111990.4 |
| Solyc03g112010.2 | Solyc03g112010.4 |
| Solyc03g112030.1 | Solyc03g112030.3 |
| Solyc03g112050.2 | Solyc03g112050.4 |
| Solyc03g112070.2 | Solyc03g112070.4 |
| Solyc03g112120.1 | Solyc03g112120.3 |
| Solyc03g112140.2 | Solyc03g112140.4 |
| Solyc03g112190.2 | Solyc03g112190.4 |
| Solyc03g112230.2 | Solyc03g112230.4 |

| | |
|---|---|
| Solyc03g112320.2 | Solyc03g112320.4 |
| Solyc03g112350.2 | Solyc03g112350.4 |
| Solyc03g112420.1 | Solyc03g112420.3 |
| Solyc03g112450.2 | Solyc03g112450.4 |
| Solyc03g112500.2 | Solyc03g112500.4 |
| Solyc03g112540.2 | Solyc03g112540.4 |
| Solyc03g112560.2 | Solyc03g112560.4 |
| Solyc03g112590.2 | Solyc03g112590.4 |
| Solyc03g112680.1 | Solyc03g112680.3 |
| Solyc03g112720.2 | Solyc03g112720.4 |
| Solyc03g112730.2 | Solyc03g112730.4 |
| Solyc03g112750.2 | Solyc03g112750.4 |
| Solyc03g112820.2 | Solyc03g112820.4 |
| Solyc03g112830.2 | Solyc03g112830.4 |
| Solyc03g112950.2 | Solyc03g112950.4 |
| Solyc03g113010.1 | Solyc03g113010.3 |
| Solyc03g113030.2 | Solyc03g113030.4 |
| Solyc03g113040.2 | Solyc03g113040.4 |
| Solyc03g113070.2 | Solyc03g113070.4 |
| Solyc03g113120.2 | Solyc03g113120.4 |
| Solyc03g113170.1 | Solyc03g113170.3 |
| Solyc03g113230.1 | Solyc03g113230.3 |
| Solyc03g113250.2 | Solyc03g113250.4 |
| Solyc03g113300.2 | Solyc03g113300.4 |
| Solyc03g113360.2 | Solyc03g113360.4 |
| Solyc03g113380.2 | Solyc03g113380.4 |
| Solyc03g113400.2 | Solyc03g113400.4 |
| Solyc03g113420.2 | Solyc03g113420.4 |
| Solyc03g113450.2 | Solyc03g113450.4 |
| Solyc03g113470.1 | Solyc03g113470.3 |
| Solyc03g113500.2 | Solyc03g113500.4 |
| Solyc03g113560.2 | Solyc03g113560.4 |
| Solyc03g113600.2 | Solyc03g113600.4 |
| Solyc03g113610.2 | Solyc03g113610.4 |
| Solyc03g113670.2 | Solyc03g113670.4 |
| Solyc03g113680.2 | Solyc03g113680.4 |
| Solyc03g113710.2 | Solyc03g113710.4 |
| Solyc03g113740.2 | Solyc03g113740.4 |
| Solyc03g113750.1 | Solyc03g113750.3 |
| Solyc03g113790.2 | Solyc03g113790.4 |
| Solyc03g113820.1 | Solyc03g113820.3 |
| Solyc03g113840.1 | Solyc03g113840.3 |
| Solyc03g113880.2 | Solyc03g113880.4 |

| | |
|---|---|
| Solyc03g113900.2 | Solyc03g113900.4 |
| Solyc03g113920.2 | Solyc03g113920.4 |
| Solyc03g113930.1 | Solyc03g113930.3 |
| Solyc03g113940.2 | Solyc03g113940.4 |
| Solyc03g113950.2 | Solyc03g113950.4 |
| Solyc03g113970.2 | Solyc03g113970.4 |
| Solyc03g114010.1 | Solyc03g114010.3 |
| Solyc03g114110.2 | Solyc03g114110.4 |
| Solyc03g114200.2 | Solyc03g114200.4 |
| Solyc03g114240.2 | Solyc03g114240.4 |
| Solyc03g114250.2 | Solyc03g114250.4 |
| Solyc03g114290.2 | Solyc03g114290.4 |
| Solyc03g114300.2 | Solyc03g114300.4 |
| Solyc03g114320.2 | Solyc03g114320.4 |
| Solyc03g114360.2 | Solyc03g114360.4 |
| Solyc03g114370.2 | Solyc03g114370.4 |
| Solyc03g114380.2 | Solyc03g114380.4 |
| Solyc03g114480.2 | Solyc03g114480.4 |
| Solyc03g114490.1 | Solyc03g114490.3 |
| Solyc03g114500.2 | Solyc03g114500.4 |
| Solyc03g114510.2 | Solyc03g114510.4 |
| Solyc03g114530.2 | Solyc03g114530.4 |
| Solyc03g114580.2 | Solyc03g114580.4 |
| Solyc03g114600.2 | Solyc03g114600.4 |
| Solyc03g114640.2 | Solyc03g114640.4 |
| Solyc03g114650.1 | Solyc03g114650.3 |
| Solyc03g114670.2 | Solyc03g114670.4 |
| Solyc03g114690.2 | Solyc03g114690.4 |
| Solyc03g114710.2 | Solyc03g114710.4 |
| Solyc03g114740.2 | Solyc03g114740.4 |
| Solyc03g114850.2 | Solyc03g114850.4 |
| Solyc03g114860.2 | Solyc03g114860.4 |
| Solyc03g114880.1 | Solyc03g114880.3 |
| Solyc03g114890.2 | Solyc03g114890.4 |
| Solyc03g114910.2 | Solyc03g114910.4 |
| Solyc03g114960.2 | Solyc03g114960.4 |
| Solyc03g114980.2 | Solyc03g114980.4 |
| Solyc03g115010.1 | Solyc03g115010.3 |
| Solyc03g115040.2 | Solyc03g115040.4 |
| Solyc03g115110.2 | Solyc03g115110.4 |
| Solyc03g115140.2 | Solyc03g115140.4 |
| Solyc03g115150.2 | Solyc03g115150.4 |
| Solyc03g115180.1 | Solyc03g115180.3 |

| | |
|---|---|
| Solyc03g115190.2 | Solyc03g115190.4 |
| Solyc03g115220.2 | Solyc03g115220.4 |
| Solyc03g115240.2 | Solyc03g115240.4 |
| Solyc03g115250.2 | Solyc03g115250.4 |
| Solyc03g115270.1 | Solyc03g115270.3 |
| Solyc03g115320.1 | Solyc03g115320.3 |
| Solyc03g115340.1 | Solyc03g115340.3 |
| Solyc03g115380.1 | Solyc03g115380.3 |
| Solyc03g115390.2 | Solyc03g115390.4 |
| Solyc03g115420.1 | Solyc03g115420.3 |
| Solyc03g115440.1 | Solyc03g115440.3 |
| Solyc03g115510.1 | Solyc03g115510.3 |
| Solyc03g115530.2 | Solyc03g115530.4 |
| Solyc03g115580.2 | Solyc03g115580.4 |
| Solyc03g115590.2 | Solyc03g115590.4 |
| Solyc03g115600.2 | Solyc03g115600.4 |
| Solyc03g115650.2 | Solyc03g115650.4 |
| Solyc03g115680.2 | Solyc03g115680.4 |
| Solyc03g115840.2 | Solyc03g115840.4 |
| Solyc03g115900.2 | Solyc03g115900.4 |
| Solyc03g115930.1 | Solyc03g115930.3 |
| Solyc03g115940.2 | Solyc03g115940.4 |
| Solyc03g115990.1 | Solyc03g115990.3 |
| Solyc03g116010.1 | Solyc03g116010.3 |
| Solyc03g116030.2 | Solyc03g116030.4 |
| Solyc03g116060.2 | Solyc03g116060.4 |
| Solyc03g116120.1 | Solyc03g116120.3 |
| Solyc03g116150.2 | Solyc03g116150.4 |
| Solyc03g116230.2 | Solyc03g116230.4 |
| Solyc03g116290.1 | Solyc03g116290.3 |
| Solyc03g116380.2 | Solyc03g116380.4 |
| Solyc03g116390.2 | Solyc03g116390.4 |
| Solyc03g116410.1 | Solyc03g116410.3 |
| Solyc03g116440.1 | Solyc03g116440.3 |
| Solyc03g116460.2 | Solyc03g116460.4 |
| Solyc03g116500.2 | Solyc03g116500.4 |
| Solyc03g116550.2 | Solyc03g116550.4 |
| Solyc03g116580.2 | Solyc03g116580.4 |
| Solyc03g116640.2 | Solyc03g116640.4 |
| Solyc03g116650.1 | Solyc03g116650.3 |
| Solyc03g116700.2 | Solyc03g116700.4 |
| Solyc03g116720.1 | Solyc03g116720.3 |
| Solyc03g116730.2 | Solyc03g116730.4 |

| | |
|---|---|
| Solyc03g116790.2 | Solyc03g116790.4 |
| Solyc03g116810.2 | Solyc03g116810.4 |
| Solyc03g116820.2 | Solyc03g116820.4 |
| Solyc03g116830.2 | Solyc03g116830.4 |
| Solyc03g116840.1 | Solyc03g116840.3 |
| Solyc03g116920.1 | Solyc03g116920.3 |
| Solyc03g116940.1 | Solyc03g116940.3 |
| Solyc03g116960.2 | Solyc03g116960.4 |
| Solyc03g116970.1 | Solyc03g116970.3 |
| Solyc03g117010.2 | Solyc03g117010.4 |
| Solyc03g117020.2 | Solyc03g117020.4 |
| Solyc03g117050.2 | Solyc03g117050.4 |
| Solyc03g117060.2 | Solyc03g117060.4 |
| Solyc03g117080.2 | Solyc03g117080.4 |
| Solyc03g117110.2 | Solyc03g117110.4 |
| Solyc03g117120.2 | Solyc03g117120.4 |
| Solyc03g117160.2 | Solyc03g117160.4 |
| Solyc03g117250.2 | Solyc03g117250.4 |
| Solyc03g117300.2 | Solyc03g117300.4 |
| Solyc03g117320.2 | Solyc03g117320.4 |
| Solyc03g117330.2 | Solyc03g117330.4 |
| Solyc03g117360.2 | Solyc03g117360.4 |
| Solyc03g117380.2 | Solyc03g117380.4 |
| Solyc03g117390.1 | Solyc03g117390.3 |
| Solyc03g117400.2 | Solyc03g117400.4 |
| Solyc03g117420.2 | Solyc03g117420.4 |
| Solyc03g117480.2 | Solyc03g117480.4 |
| Solyc03g117490.2 | Solyc03g117490.4 |
| Solyc03g117510.2 | Solyc03g117510.4 |
| Solyc03g117520.1 | Solyc03g117520.3 |
| Solyc03g117540.2 | Solyc03g117540.4 |
| Solyc03g117620.2 | Solyc03g117620.4 |
| Solyc03g117650.2 | Solyc03g117650.4 |
| Solyc03g117670.2 | Solyc03g117670.4 |
| Solyc03g117680.2 | Solyc03g117680.4 |
| Solyc03g117710.1 | Solyc03g117710.3 |
| Solyc03g117760.2 | Solyc03g117760.4 |
| Solyc03g117800.2 | Solyc03g117800.4 |
| Solyc03g117810.2 | Solyc03g117810.4 |
| Solyc03g117850.2 | Solyc03g117850.4 |
| Solyc03g117930.2 | Solyc03g117930.4 |
| Solyc03g117960.2 | Solyc03g117960.4 |
| Solyc03g117970.2 | Solyc03g117970.4 |

| | |
|---|---|
| Solyc03g118010.2 | Solyc03g118010.4 |
| Solyc03g118080.2 | Solyc03g118080.4 |
| Solyc03g118100.2 | Solyc03g118100.4 |
| Solyc03g118160.1 | Solyc03g118160.3 |
| Solyc03g118190.2 | Solyc03g118190.4 |
| Solyc03g118200.2 | Solyc03g118200.4 |
| Solyc03g118220.1 | Solyc03g118220.3 |
| Solyc03g118230.1 | Solyc03g118230.3 |
| Solyc03g118240.2 | Solyc03g118240.4 |
| Solyc03g118250.2 | Solyc03g118250.4 |
| Solyc03g118260.1 | Solyc03g118260.3 |
| Solyc03g118310.2 | Solyc03g118310.4 |
| Solyc03g118330.1 | Solyc03g118330.3 |
| Solyc03g118350.2 | Solyc03g118350.4 |
| Solyc03g118380.1 | Solyc03g118380.3 |
| Solyc03g118420.2 | Solyc03g118420.4 |
| Solyc03g118470.2 | Solyc03g118470.4 |
| Solyc03g118480.2 | Solyc03g118480.4 |
| Solyc03g118490.2 | Solyc03g118490.4 |
| Solyc03g118540.2 | Solyc03g118540.4 |
| Solyc03g118570.2 | Solyc03g118570.4 |
| Solyc03g118610.2 | Solyc03g118610.4 |
| Solyc03g118690.2 | Solyc03g118690.4 |
| Solyc03g118750.2 | Solyc03g118750.4 |
| Solyc03g118760.2 | Solyc03g118760.4 |
| Solyc03g118770.2 | Solyc03g118770.4 |
| Solyc03g118790.1 | Solyc03g118790.3 |
| Solyc03g118820.2 | Solyc03g118820.4 |
| Solyc03g118850.2 | Solyc03g118850.4 |
| Solyc03g118870.2 | Solyc03g118870.4 |
| Solyc03g118890.2 | Solyc03g118890.4 |
| Solyc03g118900.1 | Solyc03g118900.3 |
| Solyc03g118940.2 | Solyc03g118940.4 |
| Solyc03g118950.1 | Solyc03g118950.3 |
| Solyc03g118960.1 | Solyc03g118960.3 |
| Solyc03g119030.2 | Solyc03g119030.4 |
| Solyc03g119050.2 | Solyc03g119050.4 |
| Solyc03g119140.2 | Solyc03g119140.4 |
| Solyc03g119150.2 | Solyc03g119150.4 |
| Solyc03g119200.2 | Solyc03g119200.4 |
| Solyc03g119250.2 | Solyc03g119250.4 |
| Solyc03g119260.2 | Solyc03g119260.4 |
| Solyc03g119300.2 | Solyc03g119300.4 |

| | |
|---|---|
| Solyc03g119380.1 | Solyc03g119380.3 |
| Solyc03g119390.2 | Solyc03g119390.4 |
| Solyc03g119430.2 | Solyc03g119430.4 |
| Solyc03g119460.2 | Solyc03g119460.4 |
| Solyc03g119490.2 | Solyc03g119490.4 |
| Solyc03g119520.2 | Solyc03g119520.4 |
| Solyc03g119550.1 | Solyc03g119550.3 |
| Solyc03g119570.1 | Solyc03g119570.3 |
| Solyc03g119630.2 | Solyc03g119630.4 |
| Solyc03g119640.2 | Solyc03g119640.4 |
| Solyc03g119690.1 | Solyc03g119690.3 |
| Solyc03g119710.2 | Solyc03g119710.4 |
| Solyc03g119720.2 | Solyc03g119720.4 |
| Solyc03g119770.2 | Solyc03g119770.4 |
| Solyc03g119780.2 | Solyc03g119780.4 |
| Solyc03g119800.1 | Solyc03g119800.3 |
| Solyc03g119830.2 | Solyc03g119830.4 |
| Solyc03g119890.2 | Solyc03g119890.4 |
| Solyc03g119900.2 | Solyc03g119900.4 |
| Solyc03g119910.2 | Solyc03g119910.4 |
| Solyc03g119990.2 | Solyc03g119990.4 |
| Solyc03g120010.2 | Solyc03g120010.4 |
| Solyc03g120020.2 | Solyc03g120020.4 |
| Solyc03g120070.2 | Solyc03g120070.4 |
| Solyc03g120080.2 | Solyc03g120080.4 |
| Solyc03g120130.2 | Solyc03g120130.4 |
| Solyc03g120180.1 | Solyc03g120180.3 |
| Solyc03g120210.1 | Solyc03g120210.3 |
| Solyc03g120260.2 | Solyc03g120260.4 |
| Solyc03g120270.2 | Solyc03g120270.4 |
| Solyc03g120440.1 | Solyc03g120440.3 |
| Solyc03g120450.2 | Solyc03g120450.4 |
| Solyc03g120460.2 | Solyc03g120460.4 |
| Solyc03g120470.2 | Solyc03g120470.4 |
| Solyc03g120500.2 | Solyc03g120500.4 |
| Solyc03g120600.2 | Solyc03g120600.4 |
| Solyc03g120630.2 | Solyc03g120630.4 |
| Solyc03g120650.2 | Solyc03g120650.4 |
| Solyc03g120660.2 | Solyc03g120660.4 |
| Solyc03g120760.2 | Solyc03g120760.4 |
| Solyc03g120770.2 | Solyc03g120770.4 |
| Solyc03g120800.2 | Solyc03g120800.4 |
| Solyc03g120810.2 | Solyc03g120810.4 |

| | |
|---|---|
| Solyc03g120820.2 | Solyc03g120820.4 |
| Solyc03g120840.1 | Solyc03g120840.3 |
| Solyc03g120850.2 | Solyc03g120850.4 |
| Solyc03g120880.2 | Solyc03g120880.4 |
| Solyc03g120900.1 | Solyc03g120900.3 |
| Solyc03g120910.2 | Solyc03g120910.4 |
| Solyc03g120940.2 | Solyc03g120940.4 |
| Solyc03g121010.2 | Solyc03g121010.4 |
| Solyc03g121040.2 | Solyc03g121040.4 |
| Solyc03g121060.2 | Solyc03g121060.4 |
| Solyc03g121090.2 | Solyc03g121090.4 |
| Solyc03g121100.2 | Solyc03g121100.4 |
| Solyc03g121120.2 | Solyc03g121120.4 |
| Solyc03g121140.2 | Solyc03g121140.4 |
| Solyc03g121180.2 | Solyc03g121180.4 |
| Solyc03g121190.2 | Solyc03g121190.4 |
| Solyc03g121200.2 | Solyc03g121200.4 |
| Solyc03g121210.2 | Solyc03g121210.4 |
| Solyc03g121280.2 | Solyc03g121280.4 |
| Solyc03g121290.2 | Solyc03g121290.4 |
| Solyc03g121420.2 | Solyc03g121420.4 |
| Solyc03g121440.2 | Solyc03g121440.4 |
| Solyc03g121470.2 | Solyc03g121470.4 |
| Solyc03g121490.2 | Solyc03g121490.4 |
| Solyc03g121510.2 | Solyc03g121510.4 |
| Solyc03g121600.2 | Solyc03g121600.4 |
| Solyc03g121670.2 | Solyc03g121670.4 |
| Solyc03g121680.1 | Solyc03g121680.3 |
| Solyc03g121700.2 | Solyc03g121700.4 |
| Solyc03g121740.1 | Solyc03g121740.3 |
| Solyc03g121800.2 | Solyc03g121800.4 |
| Solyc03g121880.2 | Solyc03g121880.4 |
| Solyc03g121990.2 | Solyc03g121990.4 |
| Solyc03g122000.2 | Solyc03g122000.4 |
| Solyc03g122020.2 | Solyc03g122020.4 |
| Solyc03g122040.2 | Solyc03g122040.4 |
| Solyc03g122060.1 | Solyc03g122060.3 |
| Solyc03g122070.1 | Solyc03g122070.3 |
| Solyc03g122100.1 | Solyc03g122100.3 |
| Solyc03g122120.2 | Solyc03g122120.4 |
| Solyc03g122170.2 | Solyc03g122170.4 |
| Solyc03g122180.2 | Solyc03g122180.4 |
| Solyc03g122270.2 | Solyc03g122270.4 |

| | |
|---|---|
| Solyc03g122300.1 | Solyc03g122300.3 |
| Solyc03g122310.2 | Solyc03g122310.4 |
| Solyc03g123390.2 | Solyc03g123390.4 |
| Solyc03g123430.2 | Solyc03g123430.4 |
| Solyc03g123460.2 | Solyc03g123460.4 |
| Solyc03g123530.2 | Solyc03g123530.4 |
| Solyc03g123590.2 | Solyc03g123590.4 |
| Solyc03g123600.2 | Solyc03g123600.4 |
| Solyc03g123610.2 | Solyc03g123610.4 |
| Solyc03g123650.1 | Solyc03g123650.3 |
| Solyc03g123720.2 | Solyc03g123720.4 |
| Solyc03g123730.2 | Solyc03g123730.4 |
| Solyc03g123740.2 | Solyc03g123740.4 |
| Solyc03g123780.2 | Solyc03g123780.4 |
| Solyc03g123790.2 | Solyc03g123790.4 |
| Solyc03g123800.1 | Solyc03g123800.3 |
| Solyc03g123830.2 | Solyc03g123830.4 |
| Solyc03g123840.2 | Solyc03g123840.4 |
| Solyc03g123860.2 | Solyc03g123860.4 |
| Solyc03g123890.2 | Solyc03g123890.4 |
| Solyc03g123950.2 | Solyc03g123950.4 |
| Solyc03g123980.2 | Solyc03g123980.4 |
| Solyc03g124030.2 | Solyc03g124030.4 |
| Solyc04g005010.2 | Solyc04g005010.4 |
| Solyc04g005030.2 | Solyc04g005030.4 |
| Solyc04g005090.2 | Solyc04g005090.4 |
| Solyc04g005150.2 | Solyc04g005150.4 |
| Solyc04g005160.1 | Solyc04g005160.3 |
| Solyc04g005180.2 | Solyc04g005180.4 |
| Solyc04g005200.2 | Solyc04g005200.4 |
| Solyc04g005210.1 | Solyc04g005210.3 |
| Solyc04g005220.1 | Solyc04g005220.3 |
| Solyc04g005250.2 | Solyc04g005250.4 |
| Solyc04g005280.2 | Solyc04g005280.4 |
| Solyc04g005300.2 | Solyc04g005300.4 |
| Solyc04g005390.1 | Solyc04g005390.3 |
| Solyc04g005410.1 | Solyc04g005410.3 |
| Solyc04g005420.2 | Solyc04g005420.4 |
| Solyc04g005430.2 | Solyc04g005430.4 |
| Solyc04g005570.2 | Solyc04g005570.4 |
| Solyc04g005600.1 | Solyc04g005600.3 |
| Solyc04g005650.1 | Solyc04g005650.3 |
| Solyc04g005660.2 | Solyc04g005660.4 |

| | |
|---|---|
| Solyc04g005670.1 | Solyc04g005670.3 |
| Solyc04g005680.2 | Solyc04g005680.4 |
| Solyc04g005690.2 | Solyc04g005690.4 |
| Solyc04g005710.1 | Solyc04g005710.3 |
| Solyc04g005770.1 | Solyc04g005770.3 |
| Solyc04g005820.2 | Solyc04g005820.4 |
| Solyc04g005890.1 | Solyc04g005890.3 |
| Solyc04g006940.2 | Solyc04g006940.4 |
| Solyc04g006950.2 | Solyc04g006950.4 |
| Solyc04g006970.2 | Solyc04g006970.4 |
| Solyc04g007030.2 | Solyc04g007030.4 |
| Solyc04g007050.2 | Solyc04g007050.4 |
| Solyc04g007060.2 | Solyc04g007060.4 |
| Solyc04g007100.2 | Solyc04g007100.4 |
| Solyc04g007120.2 | Solyc04g007120.4 |
| Solyc04g007190.2 | Solyc04g007190.4 |
| Solyc04g007200.2 | Solyc04g007200.4 |
| Solyc04g007260.2 | Solyc04g007260.4 |
| Solyc04g007290.2 | Solyc04g007290.4 |
| Solyc04g007300.2 | Solyc04g007300.4 |
| Solyc04g007320.1 | Solyc04g007320.3 |
| Solyc04g007330.1 | Solyc04g007330.3 |
| Solyc04g007380.1 | Solyc04g007380.3 |
| Solyc04g007390.2 | Solyc04g007390.4 |
| Solyc04g007400.2 | Solyc04g007400.4 |
| Solyc04g007420.2 | Solyc04g007420.4 |
| Solyc04g007450.2 | Solyc04g007450.4 |
| Solyc04g007460.2 | Solyc04g007460.4 |
| Solyc04g007510.2 | Solyc04g007510.4 |
| Solyc04g007540.1 | Solyc04g007540.3 |
| Solyc04g007550.2 | Solyc04g007550.4 |
| Solyc04g007590.1 | Solyc04g007590.3 |
| Solyc04g007600.2 | Solyc04g007600.4 |
| Solyc04g007650.2 | Solyc04g007650.4 |
| Solyc04g007680.2 | Solyc04g007680.4 |
| Solyc04g007750.2 | Solyc04g007750.4 |
| Solyc04g007800.2 | Solyc04g007800.4 |
| Solyc04g007880.2 | Solyc04g007880.4 |
| Solyc04g007940.2 | Solyc04g007940.4 |
| Solyc04g007950.2 | Solyc04g007950.4 |
| Solyc04g007960.2 | Solyc04g007960.4 |
| Solyc04g008010.2 | Solyc04g008010.4 |
| Solyc04g008040.2 | Solyc04g008040.4 |

| | |
|---|---|
| Solyc04g008050.2 | Solyc04g008050.4 |
| Solyc04g008070.1 | Solyc04g008070.3 |
| Solyc04g008090.2 | Solyc04g008090.4 |
| Solyc04g008100.1 | Solyc04g008100.3 |
| Solyc04g008120.1 | Solyc04g008120.3 |
| Solyc04g008130.1 | Solyc04g008130.3 |
| Solyc04g008150.1 | Solyc04g008150.3 |
| Solyc04g008200.2 | Solyc04g008200.4 |
| Solyc04g008220.2 | Solyc04g008220.4 |
| Solyc04g008270.2 | Solyc04g008270.4 |
| Solyc04g008290.2 | Solyc04g008290.4 |
| Solyc04g008310.1 | Solyc04g008310.3 |
| Solyc04g008320.1 | Solyc04g008320.3 |
| Solyc04g008370.1 | Solyc04g008370.3 |
| Solyc04g008380.2 | Solyc04g008380.4 |
| Solyc04g008390.1 | Solyc04g008390.3 |
| Solyc04g008410.1 | Solyc04g008410.3 |
| Solyc04g008450.2 | Solyc04g008450.4 |
| Solyc04g008460.2 | Solyc04g008460.4 |
| Solyc04g008500.2 | Solyc04g008500.4 |
| Solyc04g008550.2 | Solyc04g008550.4 |
| Solyc04g008570.2 | Solyc04g008570.4 |
| Solyc04g008600.2 | Solyc04g008600.4 |
| Solyc04g008650.2 | Solyc04g008650.4 |
| Solyc04g008660.2 | Solyc04g008660.4 |
| Solyc04g008680.2 | Solyc04g008680.4 |
| Solyc04g008710.2 | Solyc04g008710.4 |
| Solyc04g008720.2 | Solyc04g008720.4 |
| Solyc04g008760.1 | Solyc04g008760.3 |
| Solyc04g008780.2 | Solyc04g008780.4 |
| Solyc04g008860.2 | Solyc04g008860.4 |
| Solyc04g008880.2 | Solyc04g008880.4 |
| Solyc04g008900.2 | Solyc04g008900.4 |
| Solyc04g008910.1 | Solyc04g008910.3 |
| Solyc04g008940.2 | Solyc04g008940.4 |
| Solyc04g008950.2 | Solyc04g008950.4 |
| Solyc04g009020.2 | Solyc04g009020.4 |
| Solyc04g009040.2 | Solyc04g009040.4 |
| Solyc04g009060.2 | Solyc04g009060.4 |
| Solyc04g009120.1 | Solyc04g009120.3 |
| Solyc04g009130.2 | Solyc04g009130.4 |
| Solyc04g009140.2 | Solyc04g009140.4 |
| Solyc04g009160.2 | Solyc04g009160.4 |

| | |
|---|---|
| Solyc04g009170.1 | Solyc04g009170.3 |
| Solyc04g009180.1 | Solyc04g009180.3 |
| Solyc04g009230.2 | Solyc04g009230.4 |
| Solyc04g009260.1 | Solyc04g009260.3 |
| Solyc04g009300.2 | Solyc04g009300.4 |
| Solyc04g009310.2 | Solyc04g009310.4 |
| Solyc04g009330.2 | Solyc04g009330.4 |
| Solyc04g009380.1 | Solyc04g009380.3 |
| Solyc04g009400.2 | Solyc04g009400.4 |
| Solyc04g009470.2 | Solyc04g009470.4 |
| Solyc04g009500.2 | Solyc04g009500.4 |
| Solyc04g009510.2 | Solyc04g009510.4 |
| Solyc04g009530.2 | Solyc04g009530.4 |
| Solyc04g009590.2 | Solyc04g009590.4 |
| Solyc04g009620.2 | Solyc04g009620.4 |
| Solyc04g009650.2 | Solyc04g009650.4 |
| Solyc04g009730.2 | Solyc04g009730.4 |
| Solyc04g009740.2 | Solyc04g009740.4 |
| Solyc04g009790.2 | Solyc04g009790.4 |
| Solyc04g009840.2 | Solyc04g009840.4 |
| Solyc04g009850.2 | Solyc04g009850.4 |
| Solyc04g009860.2 | Solyc04g009860.4 |
| Solyc04g009900.2 | Solyc04g009900.4 |
| Solyc04g009930.1 | Solyc04g009930.3 |
| Solyc04g009940.2 | Solyc04g009940.4 |
| Solyc04g009970.2 | Solyc04g009970.4 |
| Solyc04g009980.2 | Solyc04g009980.4 |
| Solyc04g010010.1 | Solyc04g010010.3 |
| Solyc04g010050.2 | Solyc04g010050.4 |
| Solyc04g010070.2 | Solyc04g010070.4 |
| Solyc04g010180.2 | Solyc04g010180.4 |
| Solyc04g010230.2 | Solyc04g010230.4 |
| Solyc04g010290.2 | Solyc04g010290.4 |
| Solyc04g010300.2 | Solyc04g010300.4 |
| Solyc04g010310.2 | Solyc04g010310.4 |
| Solyc04g011370.2 | Solyc04g011370.4 |
| Solyc04g011380.2 | Solyc04g011380.4 |
| Solyc04g011420.2 | Solyc04g011420.4 |
| Solyc04g011490.2 | Solyc04g011490.4 |
| Solyc04g011510.2 | Solyc04g011510.4 |
| Solyc04g011540.2 | Solyc04g011540.4 |
| Solyc04g011560.2 | Solyc04g011560.4 |
| Solyc04g011580.2 | Solyc04g011580.4 |

| | |
|---|---|
| Solyc04g011600.2 | Solyc04g011600.4 |
| Solyc04g011610.1 | Solyc04g011610.3 |
| Solyc04g011680.1 | Solyc04g011680.3 |
| Solyc04g011690.2 | Solyc04g011690.4 |
| Solyc04g011900.2 | Solyc04g011900.4 |
| Solyc04g011920.1 | Solyc04g011920.3 |
| Solyc04g011940.1 | Solyc04g011940.3 |
| Solyc04g011990.1 | Solyc04g011990.3 |
| Solyc04g012070.1 | Solyc04g012070.3 |
| Solyc04g012080.1 | Solyc04g012080.3 |
| Solyc04g012160.2 | Solyc04g012160.4 |
| Solyc04g012170.2 | Solyc04g012170.4 |
| Solyc04g012180.2 | Solyc04g012180.4 |
| Solyc04g014370.2 | Solyc04g014370.4 |
| Solyc04g014400.2 | Solyc04g014400.4 |
| Solyc04g014410.2 | Solyc04g014410.4 |
| Solyc04g014420.2 | Solyc04g014420.4 |
| Solyc04g014460.2 | Solyc04g014460.4 |
| Solyc04g014540.1 | Solyc04g014540.3 |
| Solyc04g014560.2 | Solyc04g014560.4 |
| Solyc04g014610.2 | Solyc04g014610.4 |
| Solyc04g014680.2 | Solyc04g014680.4 |
| Solyc04g014700.2 | Solyc04g014700.4 |
| Solyc04g014720.2 | Solyc04g014720.4 |
| Solyc04g014730.2 | Solyc04g014730.4 |
| Solyc04g014740.1 | Solyc04g014740.3 |
| Solyc04g014820.2 | Solyc04g014820.4 |
| Solyc04g014830.1 | Solyc04g014830.3 |
| Solyc04g014880.2 | Solyc04g014880.4 |
| Solyc04g014920.1 | Solyc04g014920.3 |
| Solyc04g014930.1 | Solyc04g014930.3 |
| Solyc04g014950.2 | Solyc04g014950.4 |
| Solyc04g014960.2 | Solyc04g014960.4 |
| Solyc04g014990.1 | Solyc04g014990.3 |
| Solyc04g015010.2 | Solyc04g015010.4 |
| Solyc04g015030.2 | Solyc04g015030.4 |
| Solyc04g015060.1 | Solyc04g015060.3 |
| Solyc04g015090.1 | Solyc04g015090.3 |
| Solyc04g015150.2 | Solyc04g015150.4 |
| Solyc04g015210.2 | Solyc04g015210.4 |
| Solyc04g015220.2 | Solyc04g015220.4 |
| Solyc04g015440.2 | Solyc04g015440.4 |
| Solyc04g015460.2 | Solyc04g015460.4 |

| | |
|---|---|
| Solyc04g015490.2 | Solyc04g015490.4 |
| Solyc04g015530.1 | Solyc04g015530.3 |
| Solyc04g015560.2 | Solyc04g015560.4 |
| Solyc04g015590.2 | Solyc04g015590.4 |
| Solyc04g015660.1 | Solyc04g015660.3 |
| Solyc04g015690.2 | Solyc04g015690.4 |
| Solyc04g015760.2 | Solyc04g015760.4 |
| Solyc04g015950.1 | Solyc04g015950.3 |
| Solyc04g016010.2 | Solyc04g016010.4 |
| Solyc04g016160.2 | Solyc04g016160.4 |
| Solyc04g016210.2 | Solyc04g016210.4 |
| Solyc04g016220.2 | Solyc04g016220.4 |
| Solyc04g016230.2 | Solyc04g016230.4 |
| Solyc04g016310.1 | Solyc04g016310.3 |
| Solyc04g016440.2 | Solyc04g016440.4 |
| Solyc04g016470.2 | Solyc04g016470.4 |
| Solyc04g016490.2 | Solyc04g016490.4 |
| Solyc04g016550.2 | Solyc04g016550.4 |
| Solyc04g017620.2 | Solyc04g017620.4 |
| Solyc04g017650.2 | Solyc04g017650.4 |
| Solyc04g017700.2 | Solyc04g017700.4 |
| Solyc04g017750.2 | Solyc04g017750.4 |
| Solyc04g017780.1 | Solyc04g017780.3 |
| Solyc04g017830.1 | Solyc04g017830.3 |
| Solyc04g017890.1 | Solyc04g017890.3 |
| Solyc04g017940.1 | Solyc04g017940.3 |
| Solyc04g017960.1 | Solyc04g017960.3 |
| Solyc04g018000.1 | Solyc04g018000.3 |
| Solyc04g018020.1 | Solyc04g018020.3 |
| Solyc04g018080.2 | Solyc04g018080.4 |
| Solyc04g018100.2 | Solyc04g018100.4 |
| Solyc04g019310.1 | Solyc04g019310.3 |
| Solyc04g024340.2 | Solyc04g024340.4 |
| Solyc04g024560.1 | Solyc04g024560.3 |
| Solyc04g024650.1 | Solyc04g024650.3 |
| Solyc04g024710.2 | Solyc04g024710.4 |
| Solyc04g024880.1 | Solyc04g024880.3 |
| Solyc04g024980.1 | Solyc04g024980.3 |
| Solyc04g024990.1 | Solyc04g024990.3 |
| Solyc04g025170.2 | Solyc04g025170.4 |
| Solyc04g025180.2 | Solyc04g025180.4 |
| Solyc04g025210.2 | Solyc04g025210.4 |
| Solyc04g025280.2 | Solyc04g025280.4 |

| | |
|---|---|
| Solyc04g025290.2 | Solyc04g025290.4 |
| Solyc04g025370.1 | Solyc04g025370.3 |
| Solyc04g025450.1 | Solyc04g025450.3 |
| Solyc04g025480.1 | Solyc04g025480.3 |
| Solyc04g025500.1 | Solyc04g025500.3 |
| Solyc04g025510.2 | Solyc04g025510.4 |
| Solyc04g025560.2 | Solyc04g025560.4 |
| Solyc04g025570.1 | Solyc04g025570.3 |
| Solyc04g025580.1 | Solyc04g025580.3 |
| Solyc04g025670.2 | Solyc04g025670.4 |
| Solyc04g025710.1 | Solyc04g025710.3 |
| Solyc04g025840.1 | Solyc04g025840.3 |
| Solyc04g025870.2 | Solyc04g025870.4 |
| Solyc04g025880.2 | Solyc04g025880.4 |
| Solyc04g025930.1 | Solyc04g025930.3 |
| Solyc04g026080.1 | Solyc04g026080.3 |
| Solyc04g026110.2 | Solyc04g026110.4 |
| Solyc04g026160.1 | Solyc04g026160.3 |
| Solyc04g026170.1 | Solyc04g026170.3 |
| Solyc04g026220.2 | Solyc04g026220.4 |
| Solyc04g026380.2 | Solyc04g026380.4 |
| Solyc04g028400.1 | Solyc04g028400.3 |
| Solyc04g028560.2 | Solyc04g028560.4 |
| Solyc04g028580.1 | Solyc04g028580.3 |
| Solyc04g039650.2 | Solyc04g039650.4 |
| Solyc04g039750.1 | Solyc04g039750.3 |
| Solyc04g039770.1 | Solyc04g039770.3 |
| Solyc04g039790.1 | Solyc04g039790.3 |
| Solyc04g039800.1 | Solyc04g039800.3 |
| Solyc04g039820.1 | Solyc04g039820.3 |
| Solyc04g039830.1 | Solyc04g039830.3 |
| Solyc04g039870.1 | Solyc04g039870.3 |
| Solyc04g039880.1 | Solyc04g039880.3 |
| Solyc04g039910.1 | Solyc04g039910.3 |
| Solyc04g039930.2 | Solyc04g039930.4 |
| Solyc04g039940.1 | Solyc04g039940.3 |
| Solyc04g039950.2 | Solyc04g039950.4 |
| Solyc04g039960.2 | Solyc04g039960.4 |
| Solyc04g040090.2 | Solyc04g040090.4 |
| Solyc04g040160.2 | Solyc04g040160.4 |
| Solyc04g040210.2 | Solyc04g040210.4 |
| Solyc04g045310.2 | Solyc04g045310.4 |
| Solyc04g045410.1 | Solyc04g045410.3 |

| | |
|---|---|
| Solyc04g045430.2 | Solyc04g045430.4 |
| Solyc04g045470.2 | Solyc04g045470.4 |
| Solyc04g045530.2 | Solyc04g045530.4 |
| Solyc04g045560.2 | Solyc04g045560.4 |
| Solyc04g045600.2 | Solyc04g045600.4 |
| Solyc04g045660.2 | Solyc04g045660.4 |
| Solyc04g047690.2 | Solyc04g047690.4 |
| Solyc04g047720.2 | Solyc04g047720.4 |
| Solyc04g047740.1 | Solyc04g047740.3 |
| Solyc04g048940.1 | Solyc04g048940.3 |
| Solyc04g048950.2 | Solyc04g048950.4 |
| Solyc04g049000.1 | Solyc04g049000.3 |
| Solyc04g049020.1 | Solyc04g049020.3 |
| Solyc04g049080.2 | Solyc04g049080.4 |
| Solyc04g049120.2 | Solyc04g049120.4 |
| Solyc04g049130.2 | Solyc04g049130.4 |
| Solyc04g049140.2 | Solyc04g049140.4 |
| Solyc04g049150.1 | Solyc04g049150.3 |
| Solyc04g049170.1 | Solyc04g049170.3 |
| Solyc04g049200.1 | Solyc04g049200.3 |
| Solyc04g049210.1 | Solyc04g049210.3 |
| Solyc04g049220.1 | Solyc04g049220.3 |
| Solyc04g049310.1 | Solyc04g049310.3 |
| Solyc04g049350.2 | Solyc04g049350.4 |
| Solyc04g049380.2 | Solyc04g049380.4 |
| Solyc04g049480.1 | Solyc04g049480.3 |
| Solyc04g049540.2 | Solyc04g049540.4 |
| Solyc04g049640.1 | Solyc04g049640.3 |
| Solyc04g049670.2 | Solyc04g049670.4 |
| Solyc04g049720.2 | Solyc04g049720.4 |
| Solyc04g049770.2 | Solyc04g049770.4 |
| Solyc04g049780.1 | Solyc04g049780.3 |
| Solyc04g049840.2 | Solyc04g049840.4 |
| Solyc04g049890.2 | Solyc04g049890.4 |
| Solyc04g049910.2 | Solyc04g049910.4 |
| Solyc04g049920.2 | Solyc04g049920.4 |
| Solyc04g050000.1 | Solyc04g050000.3 |
| Solyc04g050070.1 | Solyc04g050070.3 |
| Solyc04g050150.2 | Solyc04g050150.4 |
| Solyc04g050170.2 | Solyc04g050170.4 |
| Solyc04g050190.2 | Solyc04g050190.4 |
| Solyc04g050330.1 | Solyc04g050330.3 |
| Solyc04g050340.1 | Solyc04g050340.3 |

| | |
|---|---|
| Solyc04g050380.1 | Solyc04g050380.3 |
| Solyc04g050390.1 | Solyc04g050390.3 |
| Solyc04g050470.2 | Solyc04g050470.4 |
| Solyc04g050480.2 | Solyc04g050480.4 |
| Solyc04g050490.2 | Solyc04g050490.4 |
| Solyc04g050540.2 | Solyc04g050540.4 |
| Solyc04g050550.1 | Solyc04g050550.3 |
| Solyc04g050560.2 | Solyc04g050560.4 |
| Solyc04g050700.1 | Solyc04g050700.3 |
| Solyc04g050720.2 | Solyc04g050720.4 |
| Solyc04g050760.2 | Solyc04g050760.4 |
| Solyc04g050820.1 | Solyc04g050820.3 |
| Solyc04g050850.1 | Solyc04g050850.3 |
| Solyc04g050860.1 | Solyc04g050860.3 |
| Solyc04g051140.1 | Solyc04g051140.3 |
| Solyc04g051230.1 | Solyc04g051230.3 |
| Solyc04g051370.2 | Solyc04g051370.4 |
| Solyc04g051410.2 | Solyc04g051410.4 |
| Solyc04g051460.2 | Solyc04g051460.4 |
| Solyc04g051580.2 | Solyc04g051580.4 |
| Solyc04g051600.2 | Solyc04g051600.4 |
| Solyc04g051620.1 | Solyc04g051620.3 |
| Solyc04g051680.2 | Solyc04g051680.4 |
| Solyc04g051690.2 | Solyc04g051690.4 |
| Solyc04g051830.2 | Solyc04g051830.4 |
| Solyc04g051870.1 | Solyc04g051870.3 |
| Solyc04g051880.1 | Solyc04g051880.3 |
| Solyc04g052890.1 | Solyc04g052890.3 |
| Solyc04g052910.1 | Solyc04g052910.3 |
| Solyc04g052970.1 | Solyc04g052970.3 |
| Solyc04g053050.2 | Solyc04g053050.4 |
| Solyc04g053080.2 | Solyc04g053080.4 |
| Solyc04g053100.1 | Solyc04g053100.3 |
| Solyc04g053130.2 | Solyc04g053130.4 |
| Solyc04g053140.1 | Solyc04g053140.3 |
| Solyc04g054140.2 | Solyc04g054140.4 |
| Solyc04g054170.2 | Solyc04g054170.4 |
| Solyc04g054250.2 | Solyc04g054250.4 |
| Solyc04g054260.2 | Solyc04g054260.4 |
| Solyc04g054270.2 | Solyc04g054270.4 |
| Solyc04g054290.1 | Solyc04g054290.3 |
| Solyc04g054320.2 | Solyc04g054320.4 |
| Solyc04g054440.1 | Solyc04g054440.3 |

| | |
|---|---|
| Solyc04g054510.2 | Solyc04g054510.4 |
| Solyc04g054680.1 | Solyc04g054680.3 |
| Solyc04g054720.2 | Solyc04g054720.4 |
| Solyc04g054830.2 | Solyc04g054830.4 |
| Solyc04g054910.2 | Solyc04g054910.4 |
| Solyc04g054970.1 | Solyc04g054970.3 |
| Solyc04g055060.1 | Solyc04g055060.3 |
| Solyc04g055190.2 | Solyc04g055190.4 |
| Solyc04g056270.2 | Solyc04g056270.4 |
| Solyc04g056280.2 | Solyc04g056280.4 |
| Solyc04g056320.1 | Solyc04g056320.3 |
| Solyc04g056340.2 | Solyc04g056340.4 |
| Solyc04g056360.2 | Solyc04g056360.4 |
| Solyc04g056440.1 | Solyc04g056440.3 |
| Solyc04g056450.2 | Solyc04g056450.4 |
| Solyc04g056490.1 | Solyc04g056490.3 |
| Solyc04g056500.1 | Solyc04g056500.3 |
| Solyc04g056560.2 | Solyc04g056560.4 |
| Solyc04g056600.2 | Solyc04g056600.4 |
| Solyc04g056650.2 | Solyc04g056650.4 |
| Solyc04g056710.1 | Solyc04g056710.3 |
| Solyc04g056730.2 | Solyc04g056730.4 |
| Solyc04g057780.2 | Solyc04g057780.4 |
| Solyc04g057860.1 | Solyc04g057860.3 |
| Solyc04g057910.2 | Solyc04g057910.4 |
| Solyc04g057930.2 | Solyc04g057930.4 |
| Solyc04g057940.2 | Solyc04g057940.4 |
| Solyc04g057960.2 | Solyc04g057960.4 |
| Solyc04g057990.2 | Solyc04g057990.4 |
| Solyc04g058030.1 | Solyc04g058030.3 |
| Solyc04g058040.1 | Solyc04g058040.3 |
| Solyc04g058060.1 | Solyc04g058060.3 |
| Solyc04g058120.1 | Solyc04g058120.3 |
| Solyc04g058180.2 | Solyc04g058180.4 |
| Solyc04g063300.1 | Solyc04g063300.3 |
| Solyc04g063340.1 | Solyc04g063340.3 |
| Solyc04g063370.1 | Solyc04g063370.3 |
| Solyc04g063400.1 | Solyc04g063400.3 |
| Solyc04g063430.1 | Solyc04g063430.3 |
| Solyc04g063440.1 | Solyc04g063440.3 |
| Solyc04g064500.2 | Solyc04g064500.4 |
| Solyc04g064520.2 | Solyc04g064520.4 |
| Solyc04g064540.2 | Solyc04g064540.4 |

| | |
|---|---|
| Solyc04g064560.2 | Solyc04g064560.4 |
| Solyc04g064590.1 | Solyc04g064590.3 |
| Solyc04g064620.2 | Solyc04g064620.4 |
| Solyc04g064640.2 | Solyc04g064640.4 |
| Solyc04g064680.2 | Solyc04g064680.4 |
| Solyc04g064690.2 | Solyc04g064690.4 |
| Solyc04g064700.2 | Solyc04g064700.4 |
| Solyc04g064740.2 | Solyc04g064740.4 |
| Solyc04g064800.2 | Solyc04g064800.4 |
| Solyc04g064810.2 | Solyc04g064810.4 |
| Solyc04g064830.2 | Solyc04g064830.4 |
| Solyc04g064880.2 | Solyc04g064880.4 |
| Solyc04g064920.2 | Solyc04g064920.4 |
| Solyc04g070960.1 | Solyc04g070960.3 |
| Solyc04g070970.2 | Solyc04g070970.4 |
| Solyc04g070980.2 | Solyc04g070980.4 |
| Solyc04g070990.2 | Solyc04g070990.4 |
| Solyc04g071010.2 | Solyc04g071010.4 |
| Solyc04g071040.2 | Solyc04g071040.4 |
| Solyc04g071050.2 | Solyc04g071050.4 |
| Solyc04g071060.1 | Solyc04g071060.3 |
| Solyc04g071200.1 | Solyc04g071200.3 |
| Solyc04g071230.1 | Solyc04g071230.3 |
| Solyc04g071280.1 | Solyc04g071280.3 |
| Solyc04g071290.2 | Solyc04g071290.4 |
| Solyc04g071300.1 | Solyc04g071300.3 |
| Solyc04g071320.2 | Solyc04g071320.4 |
| Solyc04g071330.2 | Solyc04g071330.4 |
| Solyc04g071340.2 | Solyc04g071340.5 |
| Solyc04g071360.2 | Solyc04g071360.4 |
| Solyc04g071400.2 | Solyc04g071400.4 |
| Solyc04g071540.2 | Solyc04g071540.4 |
| Solyc04g071560.2 | Solyc04g071560.4 |
| Solyc04g071610.2 | Solyc04g071610.4 |
| Solyc04g071650.2 | Solyc04g071650.4 |
| Solyc04g071690.2 | Solyc04g071690.4 |
| Solyc04g071710.2 | Solyc04g071710.4 |
| Solyc04g071750.2 | Solyc04g071750.4 |
| Solyc04g071790.1 | Solyc04g071790.3 |
| Solyc04g071900.2 | Solyc04g071900.4 |
| Solyc04g071930.2 | Solyc04g071930.4 |
| Solyc04g071970.2 | Solyc04g071970.4 |
| Solyc04g072000.2 | Solyc04g072000.4 |

| | |
|---|---|
| Solyc04g072060.2 | Solyc04g072060.4 |
| Solyc04g072110.2 | Solyc04g072110.4 |
| Solyc04g072140.2 | Solyc04g072140.4 |
| Solyc04g072150.1 | Solyc04g072150.3 |
| Solyc04g072180.1 | Solyc04g072180.3 |
| Solyc04g072200.1 | Solyc04g072200.3 |
| Solyc04g072230.2 | Solyc04g072230.4 |
| Solyc04g072250.2 | Solyc04g072250.4 |
| Solyc04g072260.1 | Solyc04g072260.3 |
| Solyc04g072320.1 | Solyc04g072320.3 |
| Solyc04g072370.1 | Solyc04g072370.3 |
| Solyc04g072440.2 | Solyc04g072440.4 |
| Solyc04g072470.2 | Solyc04g072470.4 |
| Solyc04g072490.2 | Solyc04g072490.4 |
| Solyc04g072520.1 | Solyc04g072520.3 |
| Solyc04g072550.1 | Solyc04g072550.3 |
| Solyc04g072590.1 | Solyc04g072590.3 |
| Solyc04g072600.1 | Solyc04g072600.3 |
| Solyc04g072660.2 | Solyc04g072660.4 |
| Solyc04g072670.2 | Solyc04g072670.4 |
| Solyc04g072700.2 | Solyc04g072700.4 |
| Solyc04g072720.2 | Solyc04g072720.4 |
| Solyc04g072730.1 | Solyc04g072730.3 |
| Solyc04g072770.2 | Solyc04g072770.4 |
| Solyc04g072780.2 | Solyc04g072780.4 |
| Solyc04g072800.2 | Solyc04g072800.4 |
| Solyc04g072810.2 | Solyc04g072810.4 |
| Solyc04g072830.2 | Solyc04g072830.4 |
| Solyc04g072850.2 | Solyc04g072850.4 |
| Solyc04g072890.2 | Solyc04g072890.4 |
| Solyc04g072920.2 | Solyc04g072920.4 |
| Solyc04g073940.2 | Solyc04g073940.4 |
| Solyc04g073970.2 | Solyc04g073970.4 |
| Solyc04g074010.1 | Solyc04g074010.3 |
| Solyc04g074050.2 | Solyc04g074050.4 |
| Solyc04g074060.1 | Solyc04g074060.3 |
| Solyc04g074090.2 | Solyc04g074090.4 |
| Solyc04g074110.2 | Solyc04g074110.4 |
| Solyc04g074120.2 | Solyc04g074120.4 |
| Solyc04g074150.1 | Solyc04g074150.3 |
| Solyc04g074160.1 | Solyc04g074160.3 |
| Solyc04g074170.1 | Solyc04g074170.3 |
| Solyc04g074220.2 | Solyc04g074220.4 |

| | |
|---|---|
| Solyc04g074250.1 | Solyc04g074250.3 |
| Solyc04g074310.2 | Solyc04g074310.4 |
| Solyc04g074320.1 | Solyc04g074320.3 |
| Solyc04g074360.1 | Solyc04g074360.3 |
| Solyc04g074380.2 | Solyc04g074380.4 |
| Solyc04g074550.2 | Solyc04g074550.4 |
| Solyc04g074630.2 | Solyc04g074630.4 |
| Solyc04g074640.2 | Solyc04g074640.4 |
| Solyc04g074670.2 | Solyc04g074670.4 |
| Solyc04g074700.2 | Solyc04g074700.4 |
| Solyc04g074740.2 | Solyc04g074740.4 |
| Solyc04g074780.2 | Solyc04g074780.4 |
| Solyc04g074790.1 | Solyc04g074790.3 |
| Solyc04g074810.2 | Solyc04g074810.4 |
| Solyc04g074820.1 | Solyc04g074820.3 |
| Solyc04g074930.2 | Solyc04g074930.4 |
| Solyc04g074940.2 | Solyc04g074940.4 |
| Solyc04g074960.2 | Solyc04g074960.4 |
| Solyc04g076030.1 | Solyc04g076030.3 |
| Solyc04g076050.2 | Solyc04g076050.4 |
| Solyc04g076110.2 | Solyc04g076110.4 |
| Solyc04g076180.2 | Solyc04g076180.4 |
| Solyc04g076240.1 | Solyc04g076240.3 |
| Solyc04g076280.2 | Solyc04g076280.4 |
| Solyc04g076300.2 | Solyc04g076300.4 |
| Solyc04g076310.2 | Solyc04g076310.4 |
| Solyc04g076320.2 | Solyc04g076320.4 |
| Solyc04g076330.1 | Solyc04g076330.3 |
| Solyc04g076370.2 | Solyc04g076370.4 |
| Solyc04g076380.2 | Solyc04g076380.4 |
| Solyc04g076420.2 | Solyc04g076420.4 |
| Solyc04g076460.2 | Solyc04g076460.4 |
| Solyc04g076500.2 | Solyc04g076500.4 |
| Solyc04g076520.1 | Solyc04g076520.3 |
| Solyc04g076540.2 | Solyc04g076540.4 |
| Solyc04g076570.2 | Solyc04g076570.4 |
| Solyc04g076580.2 | Solyc04g076580.4 |
| Solyc04g076600.1 | Solyc04g076600.3 |
| Solyc04g076610.1 | Solyc04g076610.3 |
| Solyc04g076620.2 | Solyc04g076620.4 |
| Solyc04g076640.1 | Solyc04g076640.3 |
| Solyc04g076690.2 | Solyc04g076690.4 |
| Solyc04g076720.1 | Solyc04g076720.3 |

| | |
|---|---|
| Solyc04g076780.2 | Solyc04g076780.4 |
| Solyc04g076810.2 | Solyc04g076810.4 |
| Solyc04g076830.2 | Solyc04g076830.4 |
| Solyc04g076870.2 | Solyc04g076870.4 |
| Solyc04g076890.2 | Solyc04g076890.4 |
| Solyc04g076960.2 | Solyc04g076960.4 |
| Solyc04g077010.2 | Solyc04g077010.4 |
| Solyc04g077120.2 | Solyc04g077120.4 |
| Solyc04g077180.1 | Solyc04g077180.3 |
| Solyc04g077190.2 | Solyc04g077190.4 |
| Solyc04g077240.2 | Solyc04g077240.4 |
| Solyc04g077250.2 | Solyc04g077250.4 |
| Solyc04g077270.2 | Solyc04g077270.4 |
| Solyc04g077290.1 | Solyc04g077290.3 |
| Solyc04g077330.2 | Solyc04g077330.4 |
| Solyc04g077410.2 | Solyc04g077410.4 |
| Solyc04g077430.2 | Solyc04g077430.4 |
| Solyc04g077440.2 | Solyc04g077440.4 |
| Solyc04g077510.2 | Solyc04g077510.4 |
| Solyc04g077540.2 | Solyc04g077540.4 |
| Solyc04g077590.2 | Solyc04g077590.4 |
| Solyc04g077610.2 | Solyc04g077610.4 |
| Solyc04g077640.2 | Solyc04g077640.4 |
| Solyc04g077710.1 | Solyc04g077710.3 |
| Solyc04g077860.2 | Solyc04g077860.4 |
| Solyc04g077930.2 | Solyc04g077930.4 |
| Solyc04g077970.2 | Solyc04g077970.4 |
| Solyc04g078070.2 | Solyc04g078070.4 |
| Solyc04g078130.2 | Solyc04g078130.4 |
| Solyc04g078190.2 | Solyc04g078190.4 |
| Solyc04g078210.1 | Solyc04g078210.3 |
| Solyc04g078220.1 | Solyc04g078220.3 |
| Solyc04g078240.1 | Solyc04g078240.3 |
| Solyc04g078250.2 | Solyc04g078250.4 |
| Solyc04g078290.2 | Solyc04g078290.4 |
| Solyc04g078320.2 | Solyc04g078320.4 |
| Solyc04g078380.1 | Solyc04g078380.3 |
| Solyc04g078390.1 | Solyc04g078390.3 |
| Solyc04g078410.2 | Solyc04g078410.4 |
| Solyc04g078430.1 | Solyc04g078430.3 |
| Solyc04g078440.2 | Solyc04g078440.4 |
| Solyc04g078450.2 | Solyc04g078450.4 |
| Solyc04g078510.2 | Solyc04g078510.4 |

| | |
|---|---|
| Solyc04g078530.1 | Solyc04g078530.3 |
| Solyc04g078540.2 | Solyc04g078540.4 |
| Solyc04g078610.2 | Solyc04g078610.4 |
| Solyc04g078640.1 | Solyc04g078640.3 |
| Solyc04g078650.2 | Solyc04g078650.4 |
| Solyc04g078680.2 | Solyc04g078680.4 |
| Solyc04g078710.1 | Solyc04g078710.3 |
| Solyc04g078760.2 | Solyc04g078760.4 |
| Solyc04g078770.2 | Solyc04g078770.4 |
| Solyc04g078810.2 | Solyc04g078810.4 |
| Solyc04g078820.2 | Solyc04g078820.4 |
| Solyc04g078850.2 | Solyc04g078850.4 |
| Solyc04g078860.2 | Solyc04g078860.4 |
| Solyc04g078870.2 | Solyc04g078870.4 |
| Solyc04g078890.2 | Solyc04g078890.4 |
| Solyc04g078930.2 | Solyc04g078930.4 |
| Solyc04g078970.1 | Solyc04g078970.3 |
| Solyc04g078980.2 | Solyc04g078980.4 |
| Solyc04g078990.2 | Solyc04g078990.4 |
| Solyc04g079000.2 | Solyc04g079000.4 |
| Solyc04g079020.2 | Solyc04g079020.4 |
| Solyc04g079040.2 | Solyc04g079040.4 |
| Solyc04g079050.1 | Solyc04g079050.3 |
| Solyc04g079070.2 | Solyc04g079070.4 |
| Solyc04g079090.2 | Solyc04g079090.4 |
| Solyc04g079190.2 | Solyc04g079190.4 |
| Solyc04g079230.2 | Solyc04g079230.4 |
| Solyc04g079270.2 | Solyc04g079270.4 |
| Solyc04g079280.2 | Solyc04g079280.4 |
| Solyc04g079310.2 | Solyc04g079310.4 |
| Solyc04g079320.2 | Solyc04g079320.4 |
| Solyc04g079370.2 | Solyc04g079370.4 |
| Solyc04g079430.2 | Solyc04g079430.4 |
| Solyc04g079450.2 | Solyc04g079450.4 |
| Solyc04g079550.2 | Solyc04g079550.4 |
| Solyc04g079680.2 | Solyc04g079680.4 |
| Solyc04g079710.2 | Solyc04g079710.4 |
| Solyc04g079720.2 | Solyc04g079720.4 |
| Solyc04g079750.2 | Solyc04g079750.4 |
| Solyc04g079760.2 | Solyc04g079760.4 |
| Solyc04g079780.2 | Solyc04g079780.4 |
| Solyc04g079800.2 | Solyc04g079800.4 |
| Solyc04g079840.2 | Solyc04g079840.4 |

| | |
|---|---|
| Solyc04g079880.2 | Solyc04g079880.4 |
| Solyc04g079890.2 | Solyc04g079890.4 |
| Solyc04g079910.2 | Solyc04g079910.4 |
| Solyc04g079920.2 | Solyc04g079920.4 |
| Solyc04g079950.2 | Solyc04g079950.4 |
| Solyc04g079990.2 | Solyc04g079990.4 |
| Solyc04g080010.2 | Solyc04g080010.4 |
| Solyc04g080020.2 | Solyc04g080020.4 |
| Solyc04g080050.2 | Solyc04g080050.4 |
| Solyc04g080090.2 | Solyc04g080090.4 |
| Solyc04g080110.2 | Solyc04g080110.4 |
| Solyc04g080150.1 | Solyc04g080150.3 |
| Solyc04g080240.2 | Solyc04g080240.4 |
| Solyc04g080260.2 | Solyc04g080260.4 |
| Solyc04g080300.2 | Solyc04g080300.4 |
| Solyc04g080350.2 | Solyc04g080350.4 |
| Solyc04g080360.2 | Solyc04g080360.4 |
| Solyc04g080410.1 | Solyc04g080410.3 |
| Solyc04g080420.2 | Solyc04g080420.4 |
| Solyc04g080440.2 | Solyc04g080440.4 |
| Solyc04g080470.2 | Solyc04g080470.4 |
| Solyc04g080490.2 | Solyc04g080490.4 |
| Solyc04g080550.2 | Solyc04g080550.4 |
| Solyc04g080590.2 | Solyc04g080590.4 |
| Solyc04g080630.2 | Solyc04g080630.4 |
| Solyc04g080650.2 | Solyc04g080650.4 |
| Solyc04g080660.2 | Solyc04g080660.4 |
| Solyc04g080680.2 | Solyc04g080680.4 |
| Solyc04g080690.1 | Solyc04g080690.3 |
| Solyc04g080700.2 | Solyc04g080700.4 |
| Solyc04g080710.2 | Solyc04g080710.4 |
| Solyc04g080720.2 | Solyc04g080720.4 |
| Solyc04g080750.2 | Solyc04g080750.4 |
| Solyc04g080800.2 | Solyc04g080800.4 |
| Solyc04g080810.2 | Solyc04g080810.4 |
| Solyc04g080830.2 | Solyc04g080830.4 |
| Solyc04g080840.1 | Solyc04g080840.3 |
| Solyc04g080890.2 | Solyc04g080890.4 |
| Solyc04g080920.1 | Solyc04g080920.3 |
| Solyc04g080930.2 | Solyc04g080930.4 |
| Solyc04g080940.2 | Solyc04g080940.4 |
| Solyc04g080960.2 | Solyc04g080960.4 |
| Solyc04g080970.2 | Solyc04g080970.4 |

| | |
|---|---|
| Solyc04g081210.2 | Solyc04g081210.4 |
| Solyc04g081300.2 | Solyc04g081300.4 |
| Solyc04g081320.2 | Solyc04g081320.4 |
| Solyc04g081330.2 | Solyc04g081330.4 |
| Solyc04g081390.2 | Solyc04g081390.4 |
| Solyc04g081480.1 | Solyc04g081480.3 |
| Solyc04g081510.2 | Solyc04g081510.4 |
| Solyc04g081520.1 | Solyc04g081520.3 |
| Solyc04g081580.2 | Solyc04g081580.4 |
| Solyc04g081590.2 | Solyc04g081590.4 |
| Solyc04g081600.1 | Solyc04g081600.3 |
| Solyc04g081640.1 | Solyc04g081640.3 |
| Solyc04g081660.1 | Solyc04g081660.3 |
| Solyc04g081670.2 | Solyc04g081670.4 |
| Solyc04g081750.2 | Solyc04g081750.4 |
| Solyc04g081770.2 | Solyc04g081770.4 |
| Solyc04g081800.1 | Solyc04g081800.3 |
| Solyc04g081810.2 | Solyc04g081810.4 |
| Solyc04g081840.2 | Solyc04g081840.4 |
| Solyc04g081870.2 | Solyc04g081870.4 |
| Solyc04g081880.2 | Solyc04g081880.4 |
| Solyc04g081900.2 | Solyc04g081900.4 |
| Solyc04g081910.2 | Solyc04g081910.4 |
| Solyc04g081950.2 | Solyc04g081950.4 |
| Solyc04g082000.2 | Solyc04g082000.4 |
| Solyc04g082060.2 | Solyc04g082060.4 |
| Solyc04g082070.2 | Solyc04g082070.4 |
| Solyc04g082110.2 | Solyc04g082110.4 |
| Solyc04g082130.2 | Solyc04g082130.4 |
| Solyc04g082160.2 | Solyc04g082160.4 |
| Solyc04g082220.1 | Solyc04g082220.3 |
| Solyc04g082260.2 | Solyc04g082260.4 |
| Solyc04g082380.2 | Solyc04g082380.4 |
| Solyc04g082440.2 | Solyc04g082440.4 |
| Solyc04g082460.2 | Solyc04g082460.4 |
| Solyc04g082470.1 | Solyc04g082470.3 |
| Solyc04g082500.2 | Solyc04g082500.4 |
| Solyc04g082520.1 | Solyc04g082520.3 |
| Solyc04g082560.2 | Solyc04g082560.4 |
| Solyc04g082670.2 | Solyc04g082670.4 |
| Solyc04g082710.2 | Solyc04g082710.4 |
| Solyc04g082740.2 | Solyc04g082740.4 |
| Solyc04g082770.2 | Solyc04g082770.4 |

| | |
|---|---|
| Solyc04g082790.2 | Solyc04g082790.4 |
| Solyc04g082810.2 | Solyc04g082810.4 |
| Solyc04g082820.2 | Solyc04g082820.4 |
| Solyc04g082840.2 | Solyc04g082840.4 |
| Solyc04g082860.1 | Solyc04g082860.3 |
| Solyc04g082880.2 | Solyc04g082880.4 |
| Solyc04g082900.2 | Solyc04g082900.4 |
| Solyc04g082920.2 | Solyc04g082920.4 |
| Solyc04g082930.1 | Solyc04g082930.3 |
| Solyc04g082950.2 | Solyc04g082950.4 |
| Solyc04g083010.2 | Solyc04g083010.4 |
| Solyc04g083160.1 | Solyc04g083160.2 |
| Solyc05g005000.2 | Solyc05g005000.4 |
| Solyc05g005020.2 | Solyc05g005020.4 |
| Solyc05g005040.2 | Solyc05g005040.4 |
| Solyc05g005090.2 | Solyc05g005090.4 |
| Solyc05g005100.2 | Solyc05g005100.4 |
| Solyc05g005120.2 | Solyc05g005120.4 |
| Solyc05g005130.1 | Solyc05g005130.3 |
| Solyc05g005140.2 | Solyc05g005140.4 |
| Solyc05g005170.2 | Solyc05g005170.4 |
| Solyc05g005260.1 | Solyc05g005260.3 |
| Solyc05g005280.2 | Solyc05g005280.4 |
| Solyc05g005290.2 | Solyc05g005290.4 |
| Solyc05g005310.1 | Solyc05g005310.3 |
| Solyc05g005350.1 | Solyc05g005350.3 |
| Solyc05g005360.1 | Solyc05g005360.3 |
| Solyc05g005370.1 | Solyc05g005370.3 |
| Solyc05g005380.1 | Solyc05g005380.3 |
| Solyc05g005390.1 | Solyc05g005390.3 |
| Solyc05g005420.1 | Solyc05g005420.3 |
| Solyc05g005440.1 | Solyc05g005440.3 |
| Solyc05g005470.2 | Solyc05g005470.4 |
| Solyc05g005490.2 | Solyc05g005490.4 |
| Solyc05g005540.2 | Solyc05g005540.4 |
| Solyc05g005550.2 | Solyc05g005550.4 |
| Solyc05g005590.2 | Solyc05g005590.4 |
| Solyc05g005610.2 | Solyc05g005610.4 |
| Solyc05g005620.2 | Solyc05g005620.4 |
| Solyc05g005630.2 | Solyc05g005630.4 |
| Solyc05g005640.2 | Solyc05g005640.4 |
| Solyc05g005650.1 | Solyc05g005650.3 |
| Solyc05g005700.2 | Solyc05g005700.4 |

| | |
|---|---|
| Solyc05g005710.2 | Solyc05g005710.4 |
| Solyc05g005720.2 | Solyc05g005720.4 |
| Solyc05g005760.2 | Solyc05g005760.4 |
| Solyc05g005790.2 | Solyc05g005790.4 |
| Solyc05g005800.2 | Solyc05g005800.4 |
| Solyc05g005810.2 | Solyc05g005810.4 |
| Solyc05g005820.2 | Solyc05g005820.4 |
| Solyc05g005840.2 | Solyc05g005840.4 |
| Solyc05g005850.2 | Solyc05g005850.4 |
| Solyc05g005860.1 | Solyc05g005860.3 |
| Solyc05g005910.2 | Solyc05g005910.4 |
| Solyc05g005920.2 | Solyc05g005920.4 |
| Solyc05g005930.1 | Solyc05g005930.3 |
| Solyc05g005980.2 | Solyc05g005980.4 |
| Solyc05g006010.2 | Solyc05g006010.4 |
| Solyc05g006070.2 | Solyc05g006070.4 |
| Solyc05g006090.2 | Solyc05g006090.4 |
| Solyc05g006100.2 | Solyc05g006100.4 |
| Solyc05g006130.2 | Solyc05g006130.4 |
| Solyc05g006160.2 | Solyc05g006160.4 |
| Solyc05g006170.2 | Solyc05g006170.4 |
| Solyc05g006200.2 | Solyc05g006200.4 |
| Solyc05g006240.2 | Solyc05g006240.4 |
| Solyc05g006310.1 | Solyc05g006310.3 |
| Solyc05g006320.2 | Solyc05g006320.4 |
| Solyc05g006430.1 | Solyc05g006430.3 |
| Solyc05g006450.1 | Solyc05g006450.3 |
| Solyc05g006460.1 | Solyc05g006460.3 |
| Solyc05g006470.2 | Solyc05g006470.4 |
| Solyc05g006490.2 | Solyc05g006490.4 |
| Solyc05g006500.2 | Solyc05g006500.4 |
| Solyc05g006520.2 | Solyc05g006520.4 |
| Solyc05g006530.2 | Solyc05g006530.4 |
| Solyc05g006550.2 | Solyc05g006550.4 |
| Solyc05g006610.2 | Solyc05g006610.4 |
| Solyc05g006630.2 | Solyc05g006630.4 |
| Solyc05g006640.2 | Solyc05g006640.4 |
| Solyc05g006660.2 | Solyc05g006660.4 |
| Solyc05g006680.2 | Solyc05g006680.4 |
| Solyc05g006700.2 | Solyc05g006700.4 |
| Solyc05g006730.2 | Solyc05g006730.4 |
| Solyc05g006740.2 | Solyc05g006740.4 |
| Solyc05g006760.2 | Solyc05g006760.4 |

| | |
|---|---|
| Solyc05g006770.2 | Solyc05g006770.4 |
| Solyc05g006810.2 | Solyc05g006810.4 |
| Solyc05g006820.2 | Solyc05g006820.4 |
| Solyc05g006850.2 | Solyc05g006850.4 |
| Solyc05g006910.2 | Solyc05g006910.4 |
| Solyc05g006920.2 | Solyc05g006920.4 |
| Solyc05g006950.2 | Solyc05g006950.4 |
| Solyc05g007000.2 | Solyc05g007000.4 |
| Solyc05g007010.1 | Solyc05g007010.3 |
| Solyc05g007020.2 | Solyc05g007020.4 |
| Solyc05g007050.2 | Solyc05g007050.4 |
| Solyc05g007060.2 | Solyc05g007060.4 |
| Solyc05g007080.2 | Solyc05g007080.4 |
| Solyc05g007090.2 | Solyc05g007090.4 |
| Solyc05g007120.2 | Solyc05g007120.4 |
| Solyc05g007130.2 | Solyc05g007130.4 |
| Solyc05g007150.2 | Solyc05g007150.4 |
| Solyc05g007200.2 | Solyc05g007200.4 |
| Solyc05g007220.2 | Solyc05g007220.4 |
| Solyc05g007230.2 | Solyc05g007230.4 |
| Solyc05g007270.2 | Solyc05g007270.4 |
| Solyc05g007350.1 | Solyc05g007350.3 |
| Solyc05g007430.2 | Solyc05g007430.4 |
| Solyc05g007470.2 | Solyc05g007470.4 |
| Solyc05g007500.2 | Solyc05g007500.4 |
| Solyc05g007510.2 | Solyc05g007510.4 |
| Solyc05g007540.1 | Solyc05g007540.3 |
| Solyc05g007550.1 | Solyc05g007550.3 |
| Solyc05g007590.2 | Solyc05g007590.4 |
| Solyc05g007670.2 | Solyc05g007670.4 |
| Solyc05g007730.2 | Solyc05g007730.4 |
| Solyc05g007740.1 | Solyc05g007740.3 |
| Solyc05g007750.2 | Solyc05g007750.4 |
| Solyc05g007800.2 | Solyc05g007800.4 |
| Solyc05g007820.2 | Solyc05g007820.4 |
| Solyc05g007870.1 | Solyc05g007870.3 |
| Solyc05g007880.2 | Solyc05g007880.4 |
| Solyc05g007890.2 | Solyc05g007890.4 |
| Solyc05g007950.2 | Solyc05g007950.4 |
| Solyc05g007960.2 | Solyc05g007960.4 |
| Solyc05g007970.2 | Solyc05g007970.4 |
| Solyc05g007990.2 | Solyc05g007990.4 |
| Solyc05g008020.2 | Solyc05g008020.4 |

| | |
|---|---|
| Solyc05g008040.2 | Solyc05g008040.4 |
| Solyc05g008050.1 | Solyc05g008050.3 |
| Solyc05g008060.2 | Solyc05g008060.4 |
| Solyc05g008070.2 | Solyc05g008070.4 |
| Solyc05g008080.1 | Solyc05g008080.3 |
| Solyc05g008120.2 | Solyc05g008120.4 |
| Solyc05g008130.2 | Solyc05g008130.4 |
| Solyc05g008140.2 | Solyc05g008140.4 |
| Solyc05g008200.2 | Solyc05g008200.4 |
| Solyc05g008210.2 | Solyc05g008210.4 |
| Solyc05g008220.2 | Solyc05g008220.4 |
| Solyc05g008290.2 | Solyc05g008290.4 |
| Solyc05g008300.1 | Solyc05g008300.3 |
| Solyc05g008310.2 | Solyc05g008310.4 |
| Solyc05g008330.1 | Solyc05g008330.3 |
| Solyc05g008400.2 | Solyc05g008400.4 |
| Solyc05g008420.2 | Solyc05g008420.4 |
| Solyc05g008430.2 | Solyc05g008430.4 |
| Solyc05g008440.1 | Solyc05g008440.3 |
| Solyc05g008460.2 | Solyc05g008460.4 |
| Solyc05g008470.1 | Solyc05g008470.3 |
| Solyc05g008480.2 | Solyc05g008480.4 |
| Solyc05g008500.1 | Solyc05g008500.3 |
| Solyc05g008570.1 | Solyc05g008570.3 |
| Solyc05g008590.2 | Solyc05g008590.4 |
| Solyc05g008620.2 | Solyc05g008620.4 |
| Solyc05g008630.2 | Solyc05g008630.4 |
| Solyc05g008700.1 | Solyc05g008700.3 |
| Solyc05g008780.2 | Solyc05g008780.4 |
| Solyc05g008790.2 | Solyc05g008790.4 |
| Solyc05g008800.2 | Solyc05g008800.4 |
| Solyc05g008810.2 | Solyc05g008810.4 |
| Solyc05g008820.2 | Solyc05g008820.4 |
| Solyc05g008850.2 | Solyc05g008850.4 |
| Solyc05g008860.2 | Solyc05g008860.4 |
| Solyc05g008900.2 | Solyc05g008900.4 |
| Solyc05g008950.2 | Solyc05g008950.4 |
| Solyc05g009020.2 | Solyc05g009020.4 |
| Solyc05g009030.2 | Solyc05g009030.4 |
| Solyc05g009070.2 | Solyc05g009070.4 |
| Solyc05g009080.2 | Solyc05g009080.4 |
| Solyc05g009090.1 | Solyc05g009090.3 |
| Solyc05g009100.2 | Solyc05g009100.4 |

| | |
|---|---|
| Solyc05g009130.2 | Solyc05g009130.4 |
| Solyc05g009150.2 | Solyc05g009150.4 |
| Solyc05g009170.1 | Solyc05g009170.3 |
| Solyc05g009190.2 | Solyc05g009190.4 |
| Solyc05g009230.1 | Solyc05g009230.3 |
| Solyc05g009270.2 | Solyc05g009270.4 |
| Solyc05g009280.2 | Solyc05g009280.4 |
| Solyc05g009290.2 | Solyc05g009290.4 |
| Solyc05g009320.2 | Solyc05g009320.4 |
| Solyc05g009350.2 | Solyc05g009350.4 |
| Solyc05g009370.2 | Solyc05g009370.4 |
| Solyc05g009380.2 | Solyc05g009380.4 |
| Solyc05g009400.1 | Solyc05g009400.3 |
| Solyc05g009410.1 | Solyc05g009410.3 |
| Solyc05g009470.2 | Solyc05g009470.4 |
| Solyc05g009560.2 | Solyc05g009560.4 |
| Solyc05g009600.2 | Solyc05g009600.4 |
| Solyc05g009620.2 | Solyc05g009620.4 |
| Solyc05g009630.2 | Solyc05g009630.4 |
| Solyc05g009640.2 | Solyc05g009640.4 |
| Solyc05g009650.2 | Solyc05g009650.4 |
| Solyc05g009660.2 | Solyc05g009660.4 |
| Solyc05g009700.2 | Solyc05g009700.4 |
| Solyc05g009710.2 | Solyc05g009710.4 |
| Solyc05g009770.1 | Solyc05g009770.3 |
| Solyc05g009800.2 | Solyc05g009800.4 |
| Solyc05g009820.2 | Solyc05g009820.4 |
| Solyc05g009850.2 | Solyc05g009850.4 |
| Solyc05g009860.1 | Solyc05g009860.3 |
| Solyc05g009870.2 | Solyc05g009870.4 |
| Solyc05g009900.1 | Solyc05g009900.3 |
| Solyc05g009920.2 | Solyc05g009920.4 |
| Solyc05g009960.2 | Solyc05g009960.4 |
| Solyc05g009970.2 | Solyc05g009970.4 |
| Solyc05g009980.2 | Solyc05g009980.4 |
| Solyc05g009990.2 | Solyc05g009990.4 |
| Solyc05g010030.2 | Solyc05g010030.4 |
| Solyc05g010060.2 | Solyc05g010060.4 |
| Solyc05g010100.2 | Solyc05g010100.4 |
| Solyc05g010120.2 | Solyc05g010120.4 |
| Solyc05g010180.2 | Solyc05g010180.4 |
| Solyc05g010240.2 | Solyc05g010240.4 |
| Solyc05g010250.1 | Solyc05g010250.3 |

| | |
|---|---|
| Solyc05g010260.2 | Solyc05g010260.4 |
| Solyc05g010300.2 | Solyc05g010300.4 |
| Solyc05g010320.2 | Solyc05g010320.4 |
| Solyc05g010330.2 | Solyc05g010330.4 |
| Solyc05g010470.2 | Solyc05g010470.4 |
| Solyc05g010590.2 | Solyc05g010590.4 |
| Solyc05g010620.1 | Solyc05g010620.3 |
| Solyc05g010630.2 | Solyc05g010630.4 |
| Solyc05g010660.2 | Solyc05g010660.4 |
| Solyc05g010710.1 | Solyc05g010710.3 |
| Solyc05g010720.2 | Solyc05g010720.4 |
| Solyc05g010770.2 | Solyc05g010770.4 |
| Solyc05g011810.2 | Solyc05g011810.4 |
| Solyc05g011820.1 | Solyc05g011820.3 |
| Solyc05g011880.1 | Solyc05g011880.3 |
| Solyc05g011910.1 | Solyc05g011910.3 |
| Solyc05g011930.2 | Solyc05g011930.4 |
| Solyc05g011940.2 | Solyc05g011940.4 |
| Solyc05g011950.1 | Solyc05g011950.3 |
| Solyc05g011960.2 | Solyc05g011960.4 |
| Solyc05g011990.2 | Solyc05g011990.4 |
| Solyc05g012040.2 | Solyc05g012040.4 |
| Solyc05g012050.2 | Solyc05g012050.4 |
| Solyc05g012060.2 | Solyc05g012060.4 |
| Solyc05g012090.2 | Solyc05g012090.4 |
| Solyc05g012110.2 | Solyc05g012110.4 |
| Solyc05g012130.2 | Solyc05g012130.4 |
| Solyc05g012150.2 | Solyc05g012150.4 |
| Solyc05g012170.1 | Solyc05g012170.3 |
| Solyc05g012210.2 | Solyc05g012210.4 |
| Solyc05g012250.1 | Solyc05g012250.3 |
| Solyc05g012310.2 | Solyc05g012310.4 |
| Solyc05g012340.2 | Solyc05g012340.4 |
| Solyc05g012350.2 | Solyc05g012350.4 |
| Solyc05g012390.2 | Solyc05g012390.4 |
| Solyc05g012400.2 | Solyc05g012400.4 |
| Solyc05g012420.2 | Solyc05g012420.4 |
| Solyc05g012440.2 | Solyc05g012440.4 |
| Solyc05g012450.2 | Solyc05g012450.4 |
| Solyc05g012460.2 | Solyc05g012460.4 |
| Solyc05g012480.2 | Solyc05g012480.4 |
| Solyc05g012540.2 | Solyc05g012540.4 |
| Solyc05g012550.2 | Solyc05g012550.4 |

| | |
|---|---|
| Solyc05g012560.1 | Solyc05g012560.3 |
| Solyc05g012590.2 | Solyc05g012590.4 |
| Solyc05g012650.2 | Solyc05g012650.4 |
| Solyc05g012660.1 | Solyc05g012660.3 |
| Solyc05g012690.2 | Solyc05g012690.4 |
| Solyc05g012710.2 | Solyc05g012710.4 |
| Solyc05g012760.1 | Solyc05g012760.3 |
| Solyc05g012780.2 | Solyc05g012780.4 |
| Solyc05g012790.2 | Solyc05g012790.4 |
| Solyc05g012810.1 | Solyc05g012810.3 |
| Solyc05g012900.2 | Solyc05g012900.4 |
| Solyc05g012910.2 | Solyc05g012910.4 |
| Solyc05g012920.1 | Solyc05g012920.3 |
| Solyc05g012930.2 | Solyc05g012930.4 |
| Solyc05g012940.2 | Solyc05g012940.4 |
| Solyc05g012950.1 | Solyc05g012950.3 |
| Solyc05g013000.1 | Solyc05g013000.3 |
| Solyc05g013090.1 | Solyc05g013090.3 |
| Solyc05g013140.2 | Solyc05g013140.4 |
| Solyc05g013150.2 | Solyc05g013150.4 |
| Solyc05g013160.2 | Solyc05g013160.4 |
| Solyc05g013180.2 | Solyc05g013180.4 |
| Solyc05g013190.2 | Solyc05g013190.4 |
| Solyc05g013250.1 | Solyc05g013250.3 |
| Solyc05g013260.1 | Solyc05g013260.3 |
| Solyc05g013280.2 | Solyc05g013280.4 |
| Solyc05g013330.2 | Solyc05g013330.4 |
| Solyc05g013350.2 | Solyc05g013350.4 |
| Solyc05g013400.2 | Solyc05g013400.4 |
| Solyc05g013450.2 | Solyc05g013450.4 |
| Solyc05g013460.2 | Solyc05g013460.4 |
| Solyc05g013470.2 | Solyc05g013470.4 |
| Solyc05g013490.2 | Solyc05g013490.4 |
| Solyc05g013500.2 | Solyc05g013500.4 |
| Solyc05g013510.2 | Solyc05g013510.4 |
| Solyc05g013530.2 | Solyc05g013530.4 |
| Solyc05g013580.2 | Solyc05g013580.4 |
| Solyc05g013620.1 | Solyc05g013620.3 |
| Solyc05g013660.2 | Solyc05g013660.4 |
| Solyc05g013690.2 | Solyc05g013690.4 |
| Solyc05g013700.2 | Solyc05g013700.4 |
| Solyc05g013760.2 | Solyc05g013760.4 |
| Solyc05g013770.2 | Solyc05g013770.4 |

| | |
|---|---|
| Solyc05g013810.2 | Solyc05g013810.4 |
| Solyc05g013820.2 | Solyc05g013820.4 |
| Solyc05g013860.2 | Solyc05g013860.4 |
| Solyc05g013870.2 | Solyc05g013870.4 |
| Solyc05g013890.1 | Solyc05g013890.3 |
| Solyc05g013920.2 | Solyc05g013920.4 |
| Solyc05g013940.2 | Solyc05g013940.4 |
| Solyc05g013970.2 | Solyc05g013970.4 |
| Solyc05g014000.2 | Solyc05g014000.4 |
| Solyc05g014080.1 | Solyc05g014080.3 |
| Solyc05g014090.1 | Solyc05g014090.3 |
| Solyc05g014100.1 | Solyc05g014100.3 |
| Solyc05g014140.1 | Solyc05g014140.3 |
| Solyc05g014150.2 | Solyc05g014150.4 |
| Solyc05g014160.2 | Solyc05g014160.4 |
| Solyc05g014210.2 | Solyc05g014210.4 |
| Solyc05g014240.2 | Solyc05g014240.4 |
| Solyc05g014270.2 | Solyc05g014270.4 |
| Solyc05g014280.2 | Solyc05g014280.4 |
| Solyc05g014290.2 | Solyc05g014290.4 |
| Solyc05g014390.2 | Solyc05g014390.4 |
| Solyc05g014400.2 | Solyc05g014400.4 |
| Solyc05g014420.1 | Solyc05g014420.3 |
| Solyc05g014460.1 | Solyc05g014460.3 |
| Solyc05g014480.2 | Solyc05g014480.4 |
| Solyc05g014490.1 | Solyc05g014490.3 |
| Solyc05g014500.1 | Solyc05g014500.3 |
| Solyc05g014620.1 | Solyc05g014620.3 |
| Solyc05g014690.1 | Solyc05g014690.3 |
| Solyc05g014710.2 | Solyc05g014710.4 |
| Solyc05g014720.2 | Solyc05g014720.4 |
| Solyc05g014830.1 | Solyc05g014830.3 |
| Solyc05g014860.1 | Solyc05g014860.3 |
| Solyc05g014870.1 | Solyc05g014870.3 |
| Solyc05g014880.1 | Solyc05g014880.3 |
| Solyc05g014940.1 | Solyc05g014940.3 |
| Solyc05g014980.2 | Solyc05g014980.4 |
| Solyc05g015030.1 | Solyc05g015030.3 |
| Solyc05g015040.2 | Solyc05g015040.4 |
| Solyc05g015060.2 | Solyc05g015060.4 |
| Solyc05g015070.2 | Solyc05g015070.4 |
| Solyc05g015170.2 | Solyc05g015170.4 |
| Solyc05g015210.2 | Solyc05g015210.4 |

| | |
|---|---|
| Solyc05g015280.1 | Solyc05g015280.3 |
| Solyc05g015300.2 | Solyc05g015300.4 |
| Solyc05g015350.2 | Solyc05g015350.4 |
| Solyc05g015410.2 | Solyc05g015410.4 |
| Solyc05g015420.2 | Solyc05g015420.4 |
| Solyc05g015490.2 | Solyc05g015490.4 |
| Solyc05g015500.2 | Solyc05g015500.4 |
| Solyc05g015810.1 | Solyc05g015810.3 |
| Solyc05g015820.2 | Solyc05g015820.4 |
| Solyc05g015850.2 | Solyc05g015850.4 |
| Solyc05g015860.2 | Solyc05g015860.4 |
| Solyc05g015900.2 | Solyc05g015900.4 |
| Solyc05g015930.2 | Solyc05g015930.4 |
| Solyc05g016000.1 | Solyc05g016000.3 |
| Solyc05g016060.2 | Solyc05g016060.4 |
| Solyc05g016130.1 | Solyc05g016130.3 |
| Solyc05g016240.2 | Solyc05g016240.4 |
| Solyc05g016380.2 | Solyc05g016380.4 |
| Solyc05g016390.2 | Solyc05g016390.4 |
| Solyc05g016550.1 | Solyc05g016550.3 |
| Solyc05g016580.1 | Solyc05g016580.3 |
| Solyc05g016620.1 | Solyc05g016620.3 |
| Solyc05g016710.1 | Solyc05g016710.3 |
| Solyc05g017720.1 | Solyc05g017720.3 |
| Solyc05g017760.2 | Solyc05g017760.4 |
| Solyc05g017830.1 | Solyc05g017830.3 |
| Solyc05g017840.2 | Solyc05g017840.4 |
| Solyc05g017890.2 | Solyc05g017890.4 |
| Solyc05g017990.2 | Solyc05g017990.4 |
| Solyc05g018230.2 | Solyc05g018230.4 |
| Solyc05g018240.2 | Solyc05g018240.4 |
| Solyc05g018340.2 | Solyc05g018340.4 |
| Solyc05g018350.2 | Solyc05g018350.4 |
| Solyc05g018410.1 | Solyc05g018410.3 |
| Solyc05g018430.1 | Solyc05g018430.3 |
| Solyc05g018610.1 | Solyc05g018610.3 |
| Solyc05g018650.2 | Solyc05g018650.4 |
| Solyc05g018700.2 | Solyc05g018700.4 |
| Solyc05g018710.1 | Solyc05g018710.3 |
| Solyc05g018750.2 | Solyc05g018750.4 |
| Solyc05g018760.1 | Solyc05g018760.3 |
| Solyc05g018810.2 | Solyc05g018810.4 |
| Solyc05g018870.1 | Solyc05g018870.3 |

| | |
|---|---|
| Solyc05g020020.2 | Solyc05g020020.4 |
| Solyc05g021090.2 | Solyc05g021090.4 |
| Solyc05g021180.1 | Solyc05g021180.3 |
| Solyc05g021390.2 | Solyc05g021390.4 |
| Solyc05g021410.1 | Solyc05g021410.3 |
| Solyc05g021530.1 | Solyc05g021530.3 |
| Solyc05g021570.1 | Solyc05g021570.3 |
| Solyc05g023600.1 | Solyc05g023600.3 |
| Solyc05g023670.1 | Solyc05g023670.3 |
| Solyc05g023680.2 | Solyc05g023680.4 |
| Solyc05g023770.2 | Solyc05g023770.4 |
| Solyc05g023980.1 | Solyc05g023980.3 |
| Solyc05g024120.1 | Solyc05g024120.3 |
| Solyc05g024230.1 | Solyc05g024230.3 |
| Solyc05g024410.2 | Solyc05g024410.4 |
| Solyc05g025590.2 | Solyc05g025590.4 |
| Solyc05g025620.1 | Solyc05g025620.3 |
| Solyc05g025630.1 | Solyc05g025630.3 |
| Solyc05g025680.1 | Solyc05g025680.3 |
| Solyc05g025780.2 | Solyc05g025780.4 |
| Solyc05g025820.2 | Solyc05g025820.4 |
| Solyc05g025870.2 | Solyc05g025870.4 |
| Solyc05g025890.1 | Solyc05g025890.3 |
| Solyc05g025980.1 | Solyc05g025980.3 |
| Solyc05g026050.2 | Solyc05g026050.4 |
| Solyc05g026140.2 | Solyc05g026140.4 |
| Solyc05g026240.1 | Solyc05g026240.3 |
| Solyc05g026250.1 | Solyc05g026250.3 |
| Solyc05g026310.2 | Solyc05g026310.4 |
| Solyc05g026330.1 | Solyc05g026330.3 |
| Solyc05g026360.2 | Solyc05g026360.4 |
| Solyc05g026370.1 | Solyc05g026370.3 |
| Solyc05g026380.1 | Solyc05g026380.3 |
| Solyc05g026510.2 | Solyc05g026510.4 |
| Solyc05g031600.1 | Solyc05g031600.3 |
| Solyc05g032650.1 | Solyc05g032650.3 |
| Solyc05g032660.2 | Solyc05g032660.4 |
| Solyc05g032670.1 | Solyc05g032670.3 |
| Solyc05g032680.2 | Solyc05g032680.4 |
| Solyc05g032710.2 | Solyc05g032710.4 |
| Solyc05g032770.2 | Solyc05g032770.4 |
| Solyc05g032820.1 | Solyc05g032820.3 |
| Solyc05g039990.2 | Solyc05g039990.4 |

| | |
|---|---|
| Solyc05g040010.1 | Solyc05g040010.3 |
| Solyc05g040050.1 | Solyc05g040050.3 |
| Solyc05g040100.2 | Solyc05g040100.4 |
| Solyc05g041100.1 | Solyc05g041100.3 |
| Solyc05g041110.1 | Solyc05g041110.3 |
| Solyc05g041140.2 | Solyc05g041140.4 |
| Solyc05g041160.1 | Solyc05g041160.3 |
| Solyc05g041170.1 | Solyc05g041170.3 |
| Solyc05g041200.2 | Solyc05g041200.4 |
| Solyc05g041220.1 | Solyc05g041220.3 |
| Solyc05g041300.1 | Solyc05g041300.3 |
| Solyc05g041420.2 | Solyc05g041420.4 |
| Solyc05g041440.2 | Solyc05g041440.4 |
| Solyc05g041540.2 | Solyc05g041540.4 |
| Solyc05g041570.1 | Solyc05g041570.3 |
| Solyc05g041590.2 | Solyc05g041590.4 |
| Solyc05g041610.1 | Solyc05g041610.3 |
| Solyc05g041620.1 | Solyc05g041620.3 |
| Solyc05g041640.1 | Solyc05g041640.3 |
| Solyc05g041670.1 | Solyc05g041670.3 |
| Solyc05g041690.1 | Solyc05g041690.3 |
| Solyc05g041710.1 | Solyc05g041710.3 |
| Solyc05g041770.1 | Solyc05g041770.3 |
| Solyc05g041860.2 | Solyc05g041860.4 |
| Solyc05g042050.2 | Solyc05g042050.4 |
| Solyc05g042070.2 | Solyc05g042070.4 |
| Solyc05g042120.2 | Solyc05g042120.4 |
| Solyc05g042180.1 | Solyc05g042180.3 |
| Solyc05g043210.1 | Solyc05g043210.3 |
| Solyc05g043320.1 | Solyc05g043320.3 |
| Solyc05g043350.2 | Solyc05g043350.4 |
| Solyc05g043360.1 | Solyc05g043360.3 |
| Solyc05g043430.2 | Solyc05g043430.4 |
| Solyc05g044470.1 | Solyc05g044470.3 |
| Solyc05g044540.1 | Solyc05g044540.3 |
| Solyc05g044610.1 | Solyc05g044610.3 |
| Solyc05g044620.1 | Solyc05g044620.3 |
| Solyc05g045650.2 | Solyc05g045650.4 |
| Solyc05g045670.2 | Solyc05g045670.4 |
| Solyc05g045710.1 | Solyc05g045710.3 |
| Solyc05g045850.1 | Solyc05g045850.3 |
| Solyc05g045950.1 | Solyc05g045950.3 |
| Solyc05g046000.2 | Solyc05g046000.4 |

| | |
|---|---|
| Solyc05g046010.2 | Solyc05g046010.4 |
| Solyc05g046030.2 | Solyc05g046030.4 |
| Solyc05g046110.2 | Solyc05g046110.4 |
| Solyc05g046130.2 | Solyc05g046130.4 |
| Solyc05g046210.1 | Solyc05g046210.3 |
| Solyc05g046280.1 | Solyc05g046280.3 |
| Solyc05g046330.2 | Solyc05g046330.4 |
| Solyc05g047420.2 | Solyc05g047420.4 |
| Solyc05g047450.1 | Solyc05g047450.3 |
| Solyc05g047510.1 | Solyc05g047510.3 |
| Solyc05g047520.2 | Solyc05g047520.4 |
| Solyc05g047550.2 | Solyc05g047550.4 |
| Solyc05g047580.2 | Solyc05g047580.4 |
| Solyc05g047590.2 | Solyc05g047590.4 |
| Solyc05g047600.2 | Solyc05g047600.4 |
| Solyc05g047680.2 | Solyc05g047680.4 |
| Solyc05g048800.1 | Solyc05g048800.3 |
| Solyc05g048810.2 | Solyc05g048810.4 |
| Solyc05g048850.2 | Solyc05g048850.4 |
| Solyc05g049860.1 | Solyc05g049860.3 |
| Solyc05g049910.1 | Solyc05g049910.3 |
| Solyc05g049960.2 | Solyc05g049960.4 |
| Solyc05g049990.2 | Solyc05g049990.4 |
| Solyc05g050040.1 | Solyc05g050040.3 |
| Solyc05g050090.2 | Solyc05g050090.4 |
| Solyc05g050130.2 | Solyc05g050130.4 |
| Solyc05g050200.2 | Solyc05g050200.4 |
| Solyc05g050210.2 | Solyc05g050210.4 |
| Solyc05g050230.2 | Solyc05g050230.4 |
| Solyc05g050290.1 | Solyc05g050290.3 |
| Solyc05g050300.1 | Solyc05g050300.3 |
| Solyc05g050320.2 | Solyc05g050320.4 |
| Solyc05g050340.2 | Solyc05g050340.4 |
| Solyc05g050380.2 | Solyc05g050380.4 |
| Solyc05g050390.2 | Solyc05g050390.4 |
| Solyc05g050490.2 | Solyc05g050490.4 |
| Solyc05g050520.2 | Solyc05g050520.4 |
| Solyc05g050540.2 | Solyc05g050540.4 |
| Solyc05g050630.2 | Solyc05g050630.4 |
| Solyc05g050680.2 | Solyc05g050680.4 |
| Solyc05g050770.2 | Solyc05g050770.4 |
| Solyc05g050790.1 | Solyc05g050790.3 |
| Solyc05g050810.2 | Solyc05g050810.4 |

| | |
|---|---|
| Solyc05g050830.1 | Solyc05g050830.3 |
| Solyc05g050870.2 | Solyc05g050870.4 |
| Solyc05g050890.1 | Solyc05g050890.3 |
| Solyc05g050900.2 | Solyc05g050900.4 |
| Solyc05g050940.2 | Solyc05g050940.4 |
| Solyc05g050960.2 | Solyc05g050960.4 |
| Solyc05g050970.2 | Solyc05g050970.4 |
| Solyc05g051010.2 | Solyc05g051010.4 |
| Solyc05g051040.2 | Solyc05g051040.4 |
| Solyc05g051080.1 | Solyc05g051080.3 |
| Solyc05g051140.1 | Solyc05g051140.3 |
| Solyc05g051180.1 | Solyc05g051180.3 |
| Solyc05g051230.2 | Solyc05g051230.4 |
| Solyc05g051290.2 | Solyc05g051290.4 |
| Solyc05g051340.1 | Solyc05g051340.3 |
| Solyc05g051380.2 | Solyc05g051380.4 |
| Solyc05g051400.2 | Solyc05g051400.4 |
| Solyc05g051470.1 | Solyc05g051470.3 |
| Solyc05g051500.2 | Solyc05g051500.4 |
| Solyc05g051510.2 | Solyc05g051510.4 |
| Solyc05g051530.2 | Solyc05g051530.4 |
| Solyc05g051540.1 | Solyc05g051540.3 |
| Solyc05g051560.2 | Solyc05g051560.4 |
| Solyc05g051610.1 | Solyc05g051610.3 |
| Solyc05g051630.2 | Solyc05g051630.4 |
| Solyc05g051640.2 | Solyc05g051640.4 |
| Solyc05g051680.2 | Solyc05g051680.4 |
| Solyc05g051780.2 | Solyc05g051780.4 |
| Solyc05g051830.2 | Solyc05g051830.4 |
| Solyc05g051880.2 | Solyc05g051880.4 |
| Solyc05g051970.2 | Solyc05g051970.4 |
| Solyc05g052120.2 | Solyc05g052120.4 |
| Solyc05g052150.2 | Solyc05g052150.4 |
| Solyc05g052160.2 | Solyc05g052160.4 |
| Solyc05g052190.2 | Solyc05g052190.4 |
| Solyc05g052200.2 | Solyc05g052200.4 |
| Solyc05g052270.1 | Solyc05g052270.3 |
| Solyc05g052290.1 | Solyc05g052290.3 |
| Solyc05g052300.2 | Solyc05g052300.4 |
| Solyc05g052320.2 | Solyc05g052320.4 |
| Solyc05g052330.1 | Solyc05g052330.3 |
| Solyc05g052340.2 | Solyc05g052340.4 |
| Solyc05g052410.1 | Solyc05g052410.3 |

| | |
|---|---|
| Solyc05g052420.1 | Solyc05g052420.3 |
| Solyc05g052430.1 | Solyc05g052430.3 |
| Solyc05g052480.2 | Solyc05g052480.4 |
| Solyc05g052500.2 | Solyc05g052500.4 |
| Solyc05g052510.2 | Solyc05g052510.4 |
| Solyc05g052530.1 | Solyc05g052530.3 |
| Solyc05g052600.2 | Solyc05g052600.4 |
| Solyc05g052620.2 | Solyc05g052620.4 |
| Solyc05g052690.2 | Solyc05g052690.4 |
| Solyc05g052700.2 | Solyc05g052700.4 |
| Solyc05g052760.2 | Solyc05g052760.4 |
| Solyc05g052780.2 | Solyc05g052780.4 |
| Solyc05g052810.2 | Solyc05g052810.4 |
| Solyc05g052870.2 | Solyc05g052870.4 |
| Solyc05g052880.2 | Solyc05g052880.4 |
| Solyc05g052910.2 | Solyc05g052910.4 |
| Solyc05g052920.2 | Solyc05g052920.4 |
| Solyc05g052950.2 | Solyc05g052950.4 |
| Solyc05g052960.2 | Solyc05g052960.4 |
| Solyc05g052970.2 | Solyc05g052970.4 |
| Solyc05g052980.2 | Solyc05g052980.4 |
| Solyc05g052990.2 | Solyc05g052990.4 |
| Solyc05g053070.2 | Solyc05g053070.4 |
| Solyc05g053110.1 | Solyc05g053110.3 |
| Solyc05g053200.1 | Solyc05g053200.3 |
| Solyc05g053210.2 | Solyc05g053210.4 |
| Solyc05g053240.2 | Solyc05g053240.4 |
| Solyc05g053250.2 | Solyc05g053250.4 |
| Solyc05g053340.2 | Solyc05g053340.4 |
| Solyc05g053380.2 | Solyc05g053380.4 |
| Solyc05g053390.2 | Solyc05g053390.4 |
| Solyc05g053400.1 | Solyc05g053400.3 |
| Solyc05g053410.2 | Solyc05g053410.4 |
| Solyc05g053440.2 | Solyc05g053440.4 |
| Solyc05g053490.2 | Solyc05g053490.4 |
| Solyc05g053500.2 | Solyc05g053500.4 |
| Solyc05g053560.2 | Solyc05g053560.4 |
| Solyc05g053570.2 | Solyc05g053570.4 |
| Solyc05g053590.2 | Solyc05g053590.4 |
| Solyc05g053660.1 | Solyc05g053660.3 |
| Solyc05g053700.1 | Solyc05g053700.3 |
| Solyc05g053710.2 | Solyc05g053710.4 |
| Solyc05g053720.2 | Solyc05g053720.4 |

| | |
|---|---|
| Solyc05g053750.1 | Solyc05g053750.3 |
| Solyc05g053760.2 | Solyc05g053760.4 |
| Solyc05g053770.2 | Solyc05g053770.4 |
| Solyc05g053780.2 | Solyc05g053780.4 |
| Solyc05g053860.2 | Solyc05g053860.4 |
| Solyc05g053890.1 | Solyc05g053890.3 |
| Solyc05g053900.2 | Solyc05g053900.4 |
| Solyc05g053930.2 | Solyc05g053930.4 |
| Solyc05g053950.2 | Solyc05g053950.4 |
| Solyc05g054010.2 | Solyc05g054010.4 |
| Solyc05g054020.1 | Solyc05g054020.3 |
| Solyc05g054040.2 | Solyc05g054040.4 |
| Solyc05g054050.2 | Solyc05g054050.4 |
| Solyc05g054070.2 | Solyc05g054070.4 |
| Solyc05g054130.2 | Solyc05g054130.4 |
| Solyc05g054170.2 | Solyc05g054170.4 |
| Solyc05g054190.2 | Solyc05g054190.4 |
| Solyc05g054200.2 | Solyc05g054200.4 |
| Solyc05g054230.2 | Solyc05g054230.4 |
| Solyc05g054310.2 | Solyc05g054310.4 |
| Solyc05g054340.2 | Solyc05g054340.4 |
| Solyc05g054360.2 | Solyc05g054360.4 |
| Solyc05g054390.2 | Solyc05g054390.4 |
| Solyc05g054410.2 | Solyc05g054410.4 |
| Solyc05g054420.2 | Solyc05g054420.4 |
| Solyc05g054430.2 | Solyc05g054430.4 |
| Solyc05g054440.2 | Solyc05g054440.4 |
| Solyc05g054520.2 | Solyc05g054520.4 |
| Solyc05g054540.2 | Solyc05g054540.4 |
| Solyc05g054590.2 | Solyc05g054590.4 |
| Solyc05g054620.2 | Solyc05g054620.4 |
| Solyc05g054630.2 | Solyc05g054630.4 |
| Solyc05g054640.2 | Solyc05g054640.4 |
| Solyc05g054760.2 | Solyc05g054760.4 |
| Solyc05g054780.1 | Solyc05g054780.3 |
| Solyc05g054800.1 | Solyc05g054800.3 |
| Solyc05g054810.2 | Solyc05g054810.4 |
| Solyc05g054890.2 | Solyc05g054890.4 |
| Solyc05g054920.2 | Solyc05g054920.4 |
| Solyc05g054960.2 | Solyc05g054960.4 |
| Solyc05g055000.2 | Solyc05g055000.4 |
| Solyc05g055010.2 | Solyc05g055010.4 |
| Solyc05g055020.2 | Solyc05g055020.4 |

| | |
|---|---|
| Solyc05g055050.1 | Solyc05g055050.3 |
| Solyc05g055070.2 | Solyc05g055070.4 |
| Solyc05g055090.2 | Solyc05g055090.4 |
| Solyc05g055110.2 | Solyc05g055110.4 |
| Solyc05g055200.2 | Solyc05g055200.4 |
| Solyc05g055210.1 | Solyc05g055210.3 |
| Solyc05g055260.2 | Solyc05g055260.4 |
| Solyc05g055340.2 | Solyc05g055340.4 |
| Solyc05g055380.1 | Solyc05g055380.3 |
| Solyc05g055400.2 | Solyc05g055400.4 |
| Solyc05g055410.1 | Solyc05g055410.3 |
| Solyc05g055420.2 | Solyc05g055420.4 |
| Solyc05g055430.2 | Solyc05g055430.4 |
| Solyc05g055470.2 | Solyc05g055470.4 |
| Solyc05g055490.1 | Solyc05g055490.3 |
| Solyc05g055530.2 | Solyc05g055530.4 |
| Solyc05g055540.1 | Solyc05g055540.3 |
| Solyc05g055570.1 | Solyc05g055570.3 |
| Solyc05g055600.2 | Solyc05g055600.4 |
| Solyc05g055610.2 | Solyc05g055610.4 |
| Solyc05g055640.2 | Solyc05g055640.4 |
| Solyc05g055690.1 | Solyc05g055690.3 |
| Solyc05g055730.2 | Solyc05g055730.4 |
| Solyc05g055740.2 | Solyc05g055740.4 |
| Solyc05g055780.2 | Solyc05g055780.4 |
| Solyc05g055790.2 | Solyc05g055790.4 |
| Solyc05g055840.2 | Solyc05g055840.4 |
| Solyc05g055860.2 | Solyc05g055860.4 |
| Solyc05g055880.2 | Solyc05g055880.4 |
| Solyc05g055900.1 | Solyc05g055900.3 |
| Solyc05g055910.2 | Solyc05g055910.4 |
| Solyc05g055920.2 | Solyc05g055920.4 |
| Solyc05g055970.2 | Solyc05g055970.4 |
| Solyc05g055980.2 | Solyc05g055980.4 |
| Solyc05g055990.2 | Solyc05g055990.4 |
| Solyc05g056060.2 | Solyc05g056060.4 |
| Solyc05g056130.2 | Solyc05g056130.4 |
| Solyc05g056140.2 | Solyc05g056140.4 |
| Solyc05g056150.2 | Solyc05g056150.4 |
| Solyc05g056210.2 | Solyc05g056210.4 |
| Solyc05g056230.2 | Solyc05g056230.4 |
| Solyc05g056290.2 | Solyc05g056290.4 |
| Solyc05g056300.2 | Solyc05g056300.4 |

| | |
|---|---|
| Solyc05g056370.2 | Solyc05g056370.4 |
| Solyc05g056420.2 | Solyc05g056420.4 |
| Solyc05g056430.1 | Solyc05g056430.3 |
| Solyc05g056440.1 | Solyc05g056440.3 |
| Solyc05g056460.2 | Solyc05g056460.4 |
| Solyc05g056490.2 | Solyc05g056490.4 |
| Solyc05g056520.2 | Solyc05g056520.4 |
| Solyc05g056530.2 | Solyc05g056530.4 |
| Solyc05g056540.2 | Solyc05g056540.4 |
| Solyc05g056590.2 | Solyc05g056590.4 |
| Solyc06g005020.1 | Solyc06g005020.3 |
| Solyc06g005070.1 | Solyc06g005070.3 |
| Solyc06g005080.2 | Solyc06g005080.4 |
| Solyc06g005160.2 | Solyc06g005160.4 |
| Solyc06g005190.2 | Solyc06g005190.4 |
| Solyc06g005230.1 | Solyc06g005230.3 |
| Solyc06g005240.2 | Solyc06g005240.4 |
| Solyc06g005270.1 | Solyc06g005270.3 |
| Solyc06g005300.2 | Solyc06g005300.4 |
| Solyc06g005500.2 | Solyc06g005500.4 |
| Solyc06g005520.2 | Solyc06g005520.4 |
| Solyc06g005550.1 | Solyc06g005550.3 |
| Solyc06g005560.2 | Solyc06g005560.4 |
| Solyc06g005570.2 | Solyc06g005570.4 |
| Solyc06g005580.2 | Solyc06g005580.4 |
| Solyc06g005590.2 | Solyc06g005590.4 |
| Solyc06g005630.1 | Solyc06g005630.3 |
| Solyc06g005670.2 | Solyc06g005670.4 |
| Solyc06g005680.2 | Solyc06g005680.4 |
| Solyc06g005690.1 | Solyc06g005690.3 |
| Solyc06g005720.2 | Solyc06g005720.4 |
| Solyc06g005740.1 | Solyc06g005740.3 |
| Solyc06g005840.2 | Solyc06g005840.4 |
| Solyc06g005860.1 | Solyc06g005860.3 |
| Solyc06g005880.2 | Solyc06g005880.4 |
| Solyc06g005910.2 | Solyc06g005910.4 |
| Solyc06g005920.1 | Solyc06g005920.3 |
| Solyc06g005930.1 | Solyc06g005930.3 |
| Solyc06g005960.2 | Solyc06g005960.4 |
| Solyc06g005980.2 | Solyc06g005980.4 |
| Solyc06g005990.2 | Solyc06g005990.4 |
| Solyc06g006010.1 | Solyc06g006010.3 |
| Solyc06g007150.2 | Solyc06g007150.4 |

| | |
|---|---|
| Solyc06g007160.2 | Solyc06g007160.4 |
| Solyc06g007170.2 | Solyc06g007170.4 |
| Solyc06g007190.2 | Solyc06g007190.4 |
| Solyc06g007200.2 | Solyc06g007200.4 |
| Solyc06g007220.2 | Solyc06g007220.4 |
| Solyc06g007240.2 | Solyc06g007240.4 |
| Solyc06g007250.2 | Solyc06g007250.4 |
| Solyc06g007310.2 | Solyc06g007310.4 |
| Solyc06g007350.2 | Solyc06g007350.4 |
| Solyc06g007390.2 | Solyc06g007390.4 |
| Solyc06g007430.1 | Solyc06g007430.3 |
| Solyc06g007440.2 | Solyc06g007440.4 |
| Solyc06g007490.1 | Solyc06g007490.3 |
| Solyc06g007520.2 | Solyc06g007520.4 |
| Solyc06g007560.1 | Solyc06g007560.3 |
| Solyc06g007670.2 | Solyc06g007670.4 |
| Solyc06g007700.2 | Solyc06g007700.4 |
| Solyc06g007710.2 | Solyc06g007710.4 |
| Solyc06g007760.2 | Solyc06g007760.4 |
| Solyc06g007800.2 | Solyc06g007800.4 |
| Solyc06g007910.2 | Solyc06g007910.4 |
| Solyc06g007920.1 | Solyc06g007920.3 |
| Solyc06g007960.2 | Solyc06g007960.4 |
| Solyc06g007980.2 | Solyc06g007980.4 |
| Solyc06g008050.2 | Solyc06g008050.4 |
| Solyc06g008110.2 | Solyc06g008110.4 |
| Solyc06g008130.2 | Solyc06g008130.4 |
| Solyc06g008160.2 | Solyc06g008160.4 |
| Solyc06g008180.1 | Solyc06g008180.3 |
| Solyc06g008210.2 | Solyc06g008210.4 |
| Solyc06g008220.2 | Solyc06g008220.4 |
| Solyc06g008300.2 | Solyc06g008300.4 |
| Solyc06g008320.2 | Solyc06g008320.4 |
| Solyc06g008340.2 | Solyc06g008340.4 |
| Solyc06g008460.1 | Solyc06g008460.3 |
| Solyc06g008470.1 | Solyc06g008470.3 |
| Solyc06g008520.2 | Solyc06g008520.4 |
| Solyc06g008530.1 | Solyc06g008530.3 |
| Solyc06g008650.2 | Solyc06g008650.4 |
| Solyc06g008710.1 | Solyc06g008710.3 |
| Solyc06g008720.2 | Solyc06g008720.4 |
| Solyc06g008780.1 | Solyc06g008780.3 |
| Solyc06g008890.2 | Solyc06g008890.4 |

| | |
|---|---|
| Solyc06g008900.2 | Solyc06g008900.4 |
| Solyc06g008940.2 | Solyc06g008940.4 |
| Solyc06g008970.2 | Solyc06g008970.4 |
| Solyc06g009040.2 | Solyc06g009040.4 |
| Solyc06g009060.2 | Solyc06g009060.4 |
| Solyc06g009160.2 | Solyc06g009160.4 |
| Solyc06g009180.1 | Solyc06g009180.3 |
| Solyc06g009190.2 | Solyc06g009190.4 |
| Solyc06g009210.2 | Solyc06g009210.4 |
| Solyc06g009220.2 | Solyc06g009220.4 |
| Solyc06g009240.2 | Solyc06g009240.4 |
| Solyc06g009290.2 | Solyc06g009290.4 |
| Solyc06g009340.2 | Solyc06g009340.4 |
| Solyc06g009370.2 | Solyc06g009370.4 |
| Solyc06g009410.1 | Solyc06g009410.3 |
| Solyc06g009420.2 | Solyc06g009420.4 |
| Solyc06g009430.1 | Solyc06g009430.3 |
| Solyc06g009440.1 | Solyc06g009440.3 |
| Solyc06g009480.1 | Solyc06g009480.3 |
| Solyc06g009550.2 | Solyc06g009550.4 |
| Solyc06g009680.2 | Solyc06g009680.4 |
| Solyc06g009710.2 | Solyc06g009710.4 |
| Solyc06g009750.2 | Solyc06g009750.4 |
| Solyc06g009780.2 | Solyc06g009780.4 |
| Solyc06g009790.2 | Solyc06g009790.4 |
| Solyc06g009800.1 | Solyc06g009800.3 |
| Solyc06g009810.2 | Solyc06g009810.4 |
| Solyc06g009850.1 | Solyc06g009850.3 |
| Solyc06g009900.1 | Solyc06g009900.3 |
| Solyc06g009960.1 | Solyc06g009960.3 |
| Solyc06g010010.1 | Solyc06g010010.3 |
| Solyc06g010030.2 | Solyc06g010030.4 |
| Solyc06g010240.1 | Solyc06g010240.3 |
| Solyc06g011290.1 | Solyc06g011290.3 |
| Solyc06g011330.1 | Solyc06g011330.3 |
| Solyc06g011370.2 | Solyc06g011370.4 |
| Solyc06g011450.1 | Solyc06g011450.3 |
| Solyc06g011500.1 | Solyc06g011500.3 |
| Solyc06g011520.2 | Solyc06g011520.4 |
| Solyc06g011540.2 | Solyc06g011540.4 |
| Solyc06g011590.1 | Solyc06g011590.3 |
| Solyc06g011660.1 | Solyc06g011660.3 |
| Solyc06g016720.1 | Solyc06g016720.3 |

| | |
|---|---|
| Solyc06g016790.1 | Solyc06g016790.3 |
| Solyc06g017860.1 | Solyc06g017860.3 |
| Solyc06g017970.2 | Solyc06g017970.4 |
| Solyc06g018050.2 | Solyc06g018050.4 |
| Solyc06g018100.2 | Solyc06g018100.4 |
| Solyc06g019140.1 | Solyc06g019140.3 |
| Solyc06g019200.2 | Solyc06g019200.4 |
| Solyc06g030470.2 | Solyc06g030470.4 |
| Solyc06g030510.2 | Solyc06g030510.4 |
| Solyc06g030540.2 | Solyc06g030540.4 |
| Solyc06g030590.1 | Solyc06g030590.3 |
| Solyc06g030620.1 | Solyc06g030620.3 |
| Solyc06g030660.1 | Solyc06g030660.3 |
| Solyc06g031660.1 | Solyc06g031660.3 |
| Solyc06g031670.1 | Solyc06g031670.3 |
| Solyc06g031690.2 | Solyc06g031690.4 |
| Solyc06g033810.1 | Solyc06g033810.3 |
| Solyc06g033890.1 | Solyc06g033890.3 |
| Solyc06g033900.1 | Solyc06g033900.3 |
| Solyc06g034070.1 | Solyc06g034070.3 |
| Solyc06g034100.2 | Solyc06g034100.4 |
| Solyc06g034120.2 | Solyc06g034120.4 |
| Solyc06g034140.1 | Solyc06g034140.3 |
| Solyc06g034150.2 | Solyc06g034150.4 |
| Solyc06g034310.2 | Solyc06g034310.4 |
| Solyc06g034330.2 | Solyc06g034330.5 |
| Solyc06g034340.1 | Solyc06g034340.3 |
| Solyc06g035450.2 | Solyc06g035450.4 |
| Solyc06g035560.1 | Solyc06g035560.3 |
| Solyc06g035680.1 | Solyc06g035680.3 |
| Solyc06g035760.2 | Solyc06g035760.4 |
| Solyc06g035980.2 | Solyc06g035980.4 |
| Solyc06g036000.2 | Solyc06g036000.4 |
| Solyc06g036050.2 | Solyc06g036050.4 |
| Solyc06g036060.2 | Solyc06g036060.4 |
| Solyc06g036080.2 | Solyc06g036080.4 |
| Solyc06g036130.2 | Solyc06g036130.4 |
| Solyc06g036230.2 | Solyc06g036230.4 |
| Solyc06g036300.2 | Solyc06g036300.4 |
| Solyc06g036340.2 | Solyc06g036340.4 |
| Solyc06g036420.1 | Solyc06g036420.3 |
| Solyc06g036470.1 | Solyc06g036470.3 |
| Solyc06g036490.1 | Solyc06g036490.3 |

| | |
|---|---|
| Solyc06g036580.2 | Solyc06g036580.4 |
| Solyc06g036600.1 | Solyc06g036600.3 |
| Solyc06g036630.1 | Solyc06g036630.3 |
| Solyc06g036700.1 | Solyc06g036700.3 |
| Solyc06g036720.2 | Solyc06g036720.4 |
| Solyc06g043030.2 | Solyc06g043030.4 |
| Solyc06g043140.2 | Solyc06g043140.4 |
| Solyc06g043150.2 | Solyc06g043150.4 |
| Solyc06g043170.2 | Solyc06g043170.4 |
| Solyc06g043250.2 | Solyc06g043250.4 |
| Solyc06g048370.1 | Solyc06g048370.3 |
| Solyc06g048380.1 | Solyc06g048380.3 |
| Solyc06g048390.1 | Solyc06g048390.3 |
| Solyc06g048410.2 | Solyc06g048410.4 |
| Solyc06g048430.2 | Solyc06g048430.4 |
| Solyc06g048470.2 | Solyc06g048470.4 |
| Solyc06g048510.2 | Solyc06g048510.4 |
| Solyc06g048530.2 | Solyc06g048530.4 |
| Solyc06g048670.2 | Solyc06g048670.4 |
| Solyc06g048680.1 | Solyc06g048680.3 |
| Solyc06g048740.1 | Solyc06g048740.3 |
| Solyc06g048810.2 | Solyc06g048810.4 |
| Solyc06g048860.1 | Solyc06g048860.3 |
| Solyc06g048870.1 | Solyc06g048870.3 |
| Solyc06g048890.2 | Solyc06g048890.4 |
| Solyc06g048920.2 | Solyc06g048920.4 |
| Solyc06g048930.2 | Solyc06g048930.4 |
| Solyc06g048940.2 | Solyc06g048940.4 |
| Solyc06g048960.2 | Solyc06g048960.4 |
| Solyc06g049030.2 | Solyc06g049030.4 |
| Solyc06g049040.2 | Solyc06g049040.4 |
| Solyc06g050160.2 | Solyc06g050160.4 |
| Solyc06g050170.2 | Solyc06g050170.4 |
| Solyc06g050180.1 | Solyc06g050180.3 |
| Solyc06g050220.2 | Solyc06g050220.4 |
| Solyc06g050230.2 | Solyc06g050230.4 |
| Solyc06g050300.1 | Solyc06g050300.3 |
| Solyc06g050400.1 | Solyc06g050400.3 |
| Solyc06g050520.1 | Solyc06g050520.3 |
| Solyc06g050550.2 | Solyc06g050550.4 |
| Solyc06g050570.1 | Solyc06g050570.3 |
| Solyc06g050600.2 | Solyc06g050600.4 |
| Solyc06g050610.1 | Solyc06g050610.3 |

| | |
|---|---|
| Solyc06g050710.2 | Solyc06g050710.4 |
| Solyc06g050730.2 | Solyc06g050730.4 |
| Solyc06g050800.2 | Solyc06g050800.4 |
| Solyc06g050820.1 | Solyc06g050820.3 |
| Solyc06g050920.2 | Solyc06g050920.4 |
| Solyc06g051040.2 | Solyc06g051040.4 |
| Solyc06g051060.2 | Solyc06g051060.4 |
| Solyc06g051080.2 | Solyc06g051080.4 |
| Solyc06g051090.1 | Solyc06g051090.3 |
| Solyc06g051120.2 | Solyc06g051120.4 |
| Solyc06g051160.2 | Solyc06g051160.4 |
| Solyc06g051190.1 | Solyc06g051190.3 |
| Solyc06g051260.2 | Solyc06g051260.4 |
| Solyc06g051350.2 | Solyc06g051350.4 |
| Solyc06g051380.1 | Solyc06g051380.3 |
| Solyc06g051410.2 | Solyc06g051410.4 |
| Solyc06g051440.2 | Solyc06g051440.4 |
| Solyc06g051460.2 | Solyc06g051460.4 |
| Solyc06g051540.2 | Solyc06g051540.4 |
| Solyc06g051550.2 | Solyc06g051550.4 |
| Solyc06g051570.2 | Solyc06g051570.4 |
| Solyc06g051650.2 | Solyc06g051650.4 |
| Solyc06g051730.2 | Solyc06g051730.4 |
| Solyc06g051810.2 | Solyc06g051810.4 |
| Solyc06g051820.2 | Solyc06g051820.4 |
| Solyc06g051830.1 | Solyc06g051830.3 |
| Solyc06g051910.1 | Solyc06g051910.3 |
| Solyc06g051920.2 | Solyc06g051920.4 |
| Solyc06g051940.2 | Solyc06g051940.4 |
| Solyc06g051950.2 | Solyc06g051950.4 |
| Solyc06g051980.2 | Solyc06g051980.4 |
| Solyc06g051990.2 | Solyc06g051990.4 |
| Solyc06g052040.2 | Solyc06g052040.4 |
| Solyc06g052050.2 | Solyc06g052050.4 |
| Solyc06g052080.2 | Solyc06g052080.4 |
| Solyc06g052100.1 | Solyc06g052100.3 |
| Solyc06g052140.1 | Solyc06g052140.3 |
| Solyc06g053170.2 | Solyc06g053170.4 |
| Solyc06g053180.2 | Solyc06g053180.4 |
| Solyc06g053210.2 | Solyc06g053210.4 |
| Solyc06g053320.2 | Solyc06g053320.4 |
| Solyc06g053330.2 | Solyc06g053330.4 |
| Solyc06g053360.2 | Solyc06g053360.4 |

| | |
|---|---|
| Solyc06g053370.1 | Solyc06g053370.3 |
| Solyc06g053390.2 | Solyc06g053390.4 |
| Solyc06g053420.2 | Solyc06g053420.4 |
| Solyc06g053430.2 | Solyc06g053430.4 |
| Solyc06g053450.2 | Solyc06g053450.4 |
| Solyc06g053460.1 | Solyc06g053460.3 |
| Solyc06g053490.2 | Solyc06g053490.4 |
| Solyc06g053520.2 | Solyc06g053520.4 |
| Solyc06g053560.2 | Solyc06g053560.4 |
| Solyc06g053590.2 | Solyc06g053590.4 |
| Solyc06g053600.2 | Solyc06g053600.4 |
| Solyc06g053620.2 | Solyc06g053620.4 |
| Solyc06g053650.1 | Solyc06g053650.3 |
| Solyc06g053690.1 | Solyc06g053690.3 |
| Solyc06g053720.1 | Solyc06g053720.3 |
| Solyc06g053740.2 | Solyc06g053740.4 |
| Solyc06g053770.2 | Solyc06g053770.4 |
| Solyc06g053780.2 | Solyc06g053780.4 |
| Solyc06g053800.1 | Solyc06g053800.3 |
| Solyc06g053820.2 | Solyc06g053820.4 |
| Solyc06g053850.2 | Solyc06g053850.4 |
| Solyc06g053870.2 | Solyc06g053870.4 |
| Solyc06g053900.2 | Solyc06g053900.4 |
| Solyc06g053910.2 | Solyc06g053910.4 |
| Solyc06g053950.1 | Solyc06g053950.3 |
| Solyc06g053970.2 | Solyc06g053970.4 |
| Solyc06g054050.1 | Solyc06g054050.3 |
| Solyc06g054110.1 | Solyc06g054110.3 |
| Solyc06g054230.2 | Solyc06g054230.4 |
| Solyc06g054250.2 | Solyc06g054250.4 |
| Solyc06g054290.1 | Solyc06g054290.3 |
| Solyc06g054380.2 | Solyc06g054380.4 |
| Solyc06g054410.2 | Solyc06g054410.4 |
| Solyc06g054510.2 | Solyc06g054510.4 |
| Solyc06g054580.2 | Solyc06g054580.4 |
| Solyc06g054620.2 | Solyc06g054620.4 |
| Solyc06g054630.1 | Solyc06g054630.3 |
| Solyc06g059730.1 | Solyc06g059730.3 |
| Solyc06g059740.2 | Solyc06g059740.4 |
| Solyc06g059790.2 | Solyc06g059790.4 |
| Solyc06g059830.1 | Solyc06g059830.3 |
| Solyc06g059840.2 | Solyc06g059840.4 |
| Solyc06g059930.2 | Solyc06g059930.4 |

| | |
|---|---|
| Solyc06g059960.2 | Solyc06g059960.4 |
| Solyc06g059970.2 | Solyc06g059970.4 |
| Solyc06g059990.2 | Solyc06g059990.4 |
| Solyc06g060010.2 | Solyc06g060010.4 |
| Solyc06g060060.2 | Solyc06g060060.4 |
| Solyc06g060130.2 | Solyc06g060130.4 |
| Solyc06g060140.2 | Solyc06g060140.4 |
| Solyc06g060180.1 | Solyc06g060180.3 |
| Solyc06g060270.2 | Solyc06g060270.4 |
| Solyc06g060290.2 | Solyc06g060290.4 |
| Solyc06g060320.2 | Solyc06g060320.4 |
| Solyc06g060350.2 | Solyc06g060350.4 |
| Solyc06g060580.1 | Solyc06g060580.3 |
| Solyc06g060600.1 | Solyc06g060600.3 |
| Solyc06g060620.2 | Solyc06g060620.4 |
| Solyc06g060670.2 | Solyc06g060670.4 |
| Solyc06g060710.2 | Solyc06g060710.4 |
| Solyc06g060740.1 | Solyc06g060740.3 |
| Solyc06g060800.2 | Solyc06g060800.4 |
| Solyc06g060880.2 | Solyc06g060880.4 |
| Solyc06g060960.1 | Solyc06g060960.3 |
| Solyc06g060980.1 | Solyc06g060980.3 |
| Solyc06g060990.2 | Solyc06g060990.4 |
| Solyc06g061010.2 | Solyc06g061010.4 |
| Solyc06g061040.2 | Solyc06g061040.4 |
| Solyc06g061050.2 | Solyc06g061050.4 |
| Solyc06g061100.2 | Solyc06g061100.4 |
| Solyc06g061120.2 | Solyc06g061120.4 |
| Solyc06g061210.2 | Solyc06g061210.4 |
| Solyc06g061240.2 | Solyc06g061240.4 |
| Solyc06g061250.1 | Solyc06g061250.3 |
| Solyc06g062320.1 | Solyc06g062320.3 |
| Solyc06g062330.1 | Solyc06g062330.3 |
| Solyc06g062350.2 | Solyc06g062350.4 |
| Solyc06g062370.2 | Solyc06g062370.4 |
| Solyc06g062420.2 | Solyc06g062420.4 |
| Solyc06g062440.2 | Solyc06g062440.4 |
| Solyc06g062480.2 | Solyc06g062480.4 |
| Solyc06g062500.2 | Solyc06g062500.4 |
| Solyc06g062530.2 | Solyc06g062530.4 |
| Solyc06g062590.2 | Solyc06g062590.4 |
| Solyc06g062660.2 | Solyc06g062660.4 |
| Solyc06g062680.1 | Solyc06g062680.3 |

| | |
|---|---|
| Solyc06g062690.2 | Solyc06g062690.4 |
| Solyc06g062780.2 | Solyc06g062780.4 |
| Solyc06g062800.2 | Solyc06g062800.4 |
| Solyc06g062820.2 | Solyc06g062820.4 |
| Solyc06g062830.2 | Solyc06g062830.4 |
| Solyc06g062840.2 | Solyc06g062840.4 |
| Solyc06g062860.2 | Solyc06g062860.4 |
| Solyc06g062870.2 | Solyc06g062870.4 |
| Solyc06g062900.2 | Solyc06g062900.4 |
| Solyc06g062940.2 | Solyc06g062940.4 |
| Solyc06g062960.1 | Solyc06g062960.3 |
| Solyc06g062990.1 | Solyc06g062990.3 |
| Solyc06g063020.2 | Solyc06g063020.4 |
| Solyc06g063090.2 | Solyc06g063090.4 |
| Solyc06g063110.1 | Solyc06g063110.3 |
| Solyc06g063120.2 | Solyc06g063120.4 |
| Solyc06g063140.2 | Solyc06g063140.4 |
| Solyc06g063150.2 | Solyc06g063150.4 |
| Solyc06g063180.1 | Solyc06g063180.3 |
| Solyc06g063260.2 | Solyc06g063260.4 |
| Solyc06g063270.2 | Solyc06g063270.4 |
| Solyc06g063300.2 | Solyc06g063300.4 |
| Solyc06g063410.2 | Solyc06g063410.4 |
| Solyc06g063420.2 | Solyc06g063420.4 |
| Solyc06g064460.2 | Solyc06g064460.4 |
| Solyc06g064470.2 | Solyc06g064470.4 |
| Solyc06g064480.2 | Solyc06g064480.4 |
| Solyc06g064520.2 | Solyc06g064520.4 |
| Solyc06g064540.1 | Solyc06g064540.3 |
| Solyc06g064590.1 | Solyc06g064590.3 |
| Solyc06g064650.2 | Solyc06g064650.4 |
| Solyc06g064660.2 | Solyc06g064660.4 |
| Solyc06g064670.1 | Solyc06g064670.3 |
| Solyc06g064740.2 | Solyc06g064740.4 |
| Solyc06g064840.2 | Solyc06g064840.4 |
| Solyc06g064860.1 | Solyc06g064860.3 |
| Solyc06g064870.2 | Solyc06g064870.4 |
| Solyc06g064890.2 | Solyc06g064890.4 |
| Solyc06g064920.1 | Solyc06g064920.3 |
| Solyc06g064930.1 | Solyc06g064930.3 |
| Solyc06g064970.2 | Solyc06g064970.4 |
| Solyc06g065010.2 | Solyc06g065010.4 |
| Solyc06g065040.2 | Solyc06g065040.4 |

| | |
|---|---|
| Solyc06g065140.1 | Solyc06g065140.3 |
| Solyc06g065170.2 | Solyc06g065170.4 |
| Solyc06g065270.2 | Solyc06g065270.4 |
| Solyc06g065290.2 | Solyc06g065290.4 |
| Solyc06g065300.2 | Solyc06g065300.4 |
| Solyc06g065320.2 | Solyc06g065320.4 |
| Solyc06g065330.2 | Solyc06g065330.4 |
| Solyc06g065380.2 | Solyc06g065380.4 |
| Solyc06g065390.2 | Solyc06g065390.4 |
| Solyc06g065470.2 | Solyc06g065470.4 |
| Solyc06g065480.2 | Solyc06g065480.4 |
| Solyc06g065500.2 | Solyc06g065500.4 |
| Solyc06g065510.2 | Solyc06g065510.4 |
| Solyc06g065530.2 | Solyc06g065530.4 |
| Solyc06g065590.2 | Solyc06g065590.4 |
| Solyc06g065670.2 | Solyc06g065670.4 |
| Solyc06g065740.2 | Solyc06g065740.4 |
| Solyc06g065830.1 | Solyc06g065830.3 |
| Solyc06g065900.2 | Solyc06g065900.4 |
| Solyc06g065920.2 | Solyc06g065920.4 |
| Solyc06g065930.2 | Solyc06g065930.4 |
| Solyc06g065940.2 | Solyc06g065940.4 |
| Solyc06g065950.2 | Solyc06g065950.4 |
| Solyc06g066000.1 | Solyc06g066000.3 |
| Solyc06g066030.2 | Solyc06g066030.4 |
| Solyc06g066140.1 | Solyc06g066140.3 |
| Solyc06g066180.2 | Solyc06g066180.4 |
| Solyc06g066230.2 | Solyc06g066230.4 |
| Solyc06g066240.2 | Solyc06g066240.4 |
| Solyc06g066340.2 | Solyc06g066340.4 |
| Solyc06g066360.1 | Solyc06g066360.3 |
| Solyc06g066370.2 | Solyc06g066370.4 |
| Solyc06g066390.1 | Solyc06g066390.3 |
| Solyc06g066420.2 | Solyc06g066420.4 |
| Solyc06g066470.2 | Solyc06g066470.4 |
| Solyc06g066560.1 | Solyc06g066560.3 |
| Solyc06g066570.2 | Solyc06g066570.4 |
| Solyc06g066580.2 | Solyc06g066580.4 |
| Solyc06g066590.2 | Solyc06g066590.4 |
| Solyc06g066610.2 | Solyc06g066610.4 |
| Solyc06g066620.2 | Solyc06g066620.4 |
| Solyc06g066650.2 | Solyc06g066650.4 |
| Solyc06g066660.2 | Solyc06g066660.4 |

| | |
|---|---|
| Solyc06g066670.2 | Solyc06g066670.4 |
| Solyc06g066690.2 | Solyc06g066690.4 |
| Solyc06g066740.1 | Solyc06g066740.3 |
| Solyc06g066820.2 | Solyc06g066820.4 |
| Solyc06g066830.2 | Solyc06g066830.4 |
| Solyc06g067900.1 | Solyc06g067900.3 |
| Solyc06g067940.2 | Solyc06g067940.4 |
| Solyc06g067970.1 | Solyc06g067970.3 |
| Solyc06g068080.2 | Solyc06g068080.4 |
| Solyc06g068120.2 | Solyc06g068120.4 |
| Solyc06g068210.1 | Solyc06g068210.3 |
| Solyc06g068230.2 | Solyc06g068230.4 |
| Solyc06g068260.1 | Solyc06g068260.3 |
| Solyc06g068340.1 | Solyc06g068340.3 |
| Solyc06g068380.2 | Solyc06g068380.4 |
| Solyc06g068430.2 | Solyc06g068430.4 |
| Solyc06g068440.2 | Solyc06g068440.4 |
| Solyc06g068450.1 | Solyc06g068450.3 |
| Solyc06g068550.2 | Solyc06g068550.4 |
| Solyc06g068570.2 | Solyc06g068570.4 |
| Solyc06g068590.2 | Solyc06g068590.4 |
| Solyc06g068610.1 | Solyc06g068610.3 |
| Solyc06g068630.2 | Solyc06g068630.4 |
| Solyc06g068640.1 | Solyc06g068640.3 |
| Solyc06g068650.2 | Solyc06g068650.4 |
| Solyc06g068700.2 | Solyc06g068700.4 |
| Solyc06g068740.2 | Solyc06g068740.4 |
| Solyc06g068790.2 | Solyc06g068790.4 |
| Solyc06g068810.2 | Solyc06g068810.4 |
| Solyc06g068850.2 | Solyc06g068850.4 |
| Solyc06g068900.2 | Solyc06g068900.4 |
| Solyc06g068940.2 | Solyc06g068940.4 |
| Solyc06g068950.2 | Solyc06g068950.4 |
| Solyc06g068980.2 | Solyc06g068980.4 |
| Solyc06g069020.2 | Solyc06g069020.4 |
| Solyc06g069040.2 | Solyc06g069040.4 |
| Solyc06g069100.1 | Solyc06g069100.3 |
| Solyc06g069110.2 | Solyc06g069110.4 |
| Solyc06g069130.2 | Solyc06g069130.4 |
| Solyc06g069190.2 | Solyc06g069190.4 |
| Solyc06g069280.2 | Solyc06g069280.4 |
| Solyc06g069350.1 | Solyc06g069350.3 |
| Solyc06g069370.2 | Solyc06g069370.4 |

| | |
|---|---|
| Solyc06g069380.2 | Solyc06g069380.4 |
| Solyc06g069390.2 | Solyc06g069390.4 |
| Solyc06g069400.2 | Solyc06g069400.4 |
| Solyc06g069450.2 | Solyc06g069450.4 |
| Solyc06g069460.1 | Solyc06g069460.3 |
| Solyc06g069560.1 | Solyc06g069560.3 |
| Solyc06g069600.1 | Solyc06g069600.3 |
| Solyc06g069650.2 | Solyc06g069650.4 |
| Solyc06g069670.2 | Solyc06g069670.4 |
| Solyc06g069690.2 | Solyc06g069690.4 |
| Solyc06g069700.2 | Solyc06g069700.4 |
| Solyc06g069720.2 | Solyc06g069720.4 |
| Solyc06g069780.2 | Solyc06g069780.4 |
| Solyc06g069790.2 | Solyc06g069790.4 |
| Solyc06g069820.2 | Solyc06g069820.4 |
| Solyc06g069860.2 | Solyc06g069860.4 |
| Solyc06g069890.2 | Solyc06g069890.4 |
| Solyc06g070940.2 | Solyc06g070940.4 |
| Solyc06g070960.1 | Solyc06g070960.3 |
| Solyc06g071050.2 | Solyc06g071050.4 |
| Solyc06g071090.1 | Solyc06g071090.3 |
| Solyc06g071100.2 | Solyc06g071100.4 |
| Solyc06g071150.2 | Solyc06g071150.4 |
| Solyc06g071190.2 | Solyc06g071190.4 |
| Solyc06g071240.1 | Solyc06g071240.3 |
| Solyc06g071260.2 | Solyc06g071260.4 |
| Solyc06g071280.2 | Solyc06g071280.4 |
| Solyc06g071310.2 | Solyc06g071310.4 |
| Solyc06g071330.2 | Solyc06g071330.4 |
| Solyc06g071350.2 | Solyc06g071350.4 |
| Solyc06g071440.2 | Solyc06g071440.4 |
| Solyc06g071450.2 | Solyc06g071450.4 |
| Solyc06g071470.2 | Solyc06g071470.4 |
| Solyc06g071490.1 | Solyc06g071490.3 |
| Solyc06g071530.2 | Solyc06g071530.4 |
| Solyc06g071610.1 | Solyc06g071610.3 |
| Solyc06g071650.1 | Solyc06g071650.3 |
| Solyc06g071660.1 | Solyc06g071660.3 |
| Solyc06g071680.2 | Solyc06g071680.4 |
| Solyc06g071770.2 | Solyc06g071770.4 |
| Solyc06g071780.2 | Solyc06g071780.4 |
| Solyc06g071800.2 | Solyc06g071800.4 |
| Solyc06g071840.1 | Solyc06g071840.3 |

| | |
|---|---|
| Solyc06g071880.2 | Solyc06g071880.4 |
| Solyc06g071920.2 | Solyc06g071920.4 |
| Solyc06g071940.1 | Solyc06g071940.3 |
| Solyc06g071950.1 | Solyc06g071950.3 |
| Solyc06g071960.2 | Solyc06g071960.4 |
| Solyc06g071990.2 | Solyc06g071990.4 |
| Solyc06g072000.1 | Solyc06g072000.3 |
| Solyc06g072050.2 | Solyc06g072050.4 |
| Solyc06g072090.1 | Solyc06g072090.3 |
| Solyc06g072100.1 | Solyc06g072100.3 |
| Solyc06g072110.2 | Solyc06g072110.4 |
| Solyc06g072130.2 | Solyc06g072130.4 |
| Solyc06g072140.2 | Solyc06g072140.4 |
| Solyc06g072270.2 | Solyc06g072270.4 |
| Solyc06g072300.2 | Solyc06g072300.4 |
| Solyc06g072320.2 | Solyc06g072320.4 |
| Solyc06g072400.1 | Solyc06g072400.3 |
| Solyc06g072410.1 | Solyc06g072410.3 |
| Solyc06g072430.1 | Solyc06g072430.3 |
| Solyc06g072470.2 | Solyc06g072470.4 |
| Solyc06g072510.2 | Solyc06g072510.4 |
| Solyc06g072570.2 | Solyc06g072570.4 |
| Solyc06g072590.2 | Solyc06g072590.4 |
| Solyc06g072620.2 | Solyc06g072620.4 |
| Solyc06g072630.2 | Solyc06g072630.4 |
| Solyc06g072660.2 | Solyc06g072660.4 |
| Solyc06g072680.2 | Solyc06g072680.4 |
| Solyc06g072690.1 | Solyc06g072690.3 |
| Solyc06g072730.2 | Solyc06g072730.4 |
| Solyc06g072760.2 | Solyc06g072760.4 |
| Solyc06g072800.2 | Solyc06g072800.4 |
| Solyc06g072810.2 | Solyc06g072810.4 |
| Solyc06g072830.2 | Solyc06g072830.4 |
| Solyc06g072890.1 | Solyc06g072890.3 |
| Solyc06g072900.2 | Solyc06g072900.4 |
| Solyc06g072920.1 | Solyc06g072920.3 |
| Solyc06g072950.1 | Solyc06g072950.3 |
| Solyc06g072980.2 | Solyc06g072980.4 |
| Solyc06g073000.2 | Solyc06g073000.4 |
| Solyc06g073030.1 | Solyc06g073030.3 |
| Solyc06g073080.2 | Solyc06g073080.4 |
| Solyc06g073100.2 | Solyc06g073100.4 |
| Solyc06g073120.2 | Solyc06g073120.4 |

| | |
|---|---|
| Solyc06g073130.2 | Solyc06g073130.4 |
| Solyc06g073200.2 | Solyc06g073200.4 |
| Solyc06g073210.2 | Solyc06g073210.4 |
| Solyc06g073230.2 | Solyc06g073230.4 |
| Solyc06g073270.2 | Solyc06g073270.4 |
| Solyc06g073330.2 | Solyc06g073330.4 |
| Solyc06g073370.2 | Solyc06g073370.4 |
| Solyc06g073430.2 | Solyc06g073430.4 |
| Solyc06g073460.2 | Solyc06g073460.4 |
| Solyc06g073530.1 | Solyc06g073530.3 |
| Solyc06g073540.2 | Solyc06g073540.4 |
| Solyc06g073560.2 | Solyc06g073560.4 |
| Solyc06g073570.2 | Solyc06g073570.4 |
| Solyc06g073580.2 | Solyc06g073580.4 |
| Solyc06g073600.2 | Solyc06g073600.4 |
| Solyc06g073640.2 | Solyc06g073640.4 |
| Solyc06g073670.2 | Solyc06g073670.4 |
| Solyc06g073710.2 | Solyc06g073710.4 |
| Solyc06g073720.1 | Solyc06g073720.3 |
| Solyc06g073750.2 | Solyc06g073750.4 |
| Solyc06g073780.2 | Solyc06g073780.4 |
| Solyc06g073810.2 | Solyc06g073810.4 |
| Solyc06g073820.1 | Solyc06g073820.3 |
| Solyc06g073870.2 | Solyc06g073870.4 |
| Solyc06g073910.2 | Solyc06g073910.4 |
| Solyc06g073930.2 | Solyc06g073930.4 |
| Solyc06g073940.2 | Solyc06g073940.4 |
| Solyc06g073960.2 | Solyc06g073960.4 |
| Solyc06g073970.1 | Solyc06g073970.3 |
| Solyc06g073980.2 | Solyc06g073980.4 |
| Solyc06g074050.1 | Solyc06g074050.3 |
| Solyc06g074130.2 | Solyc06g074130.4 |
| Solyc06g074160.1 | Solyc06g074160.3 |
| Solyc06g074180.1 | Solyc06g074180.3 |
| Solyc06g074200.2 | Solyc06g074200.4 |
| Solyc06g074240.1 | Solyc06g074240.3 |
| Solyc06g074300.2 | Solyc06g074300.4 |
| Solyc06g074310.2 | Solyc06g074310.4 |
| Solyc06g074360.2 | Solyc06g074360.4 |
| Solyc06g074370.1 | Solyc06g074370.3 |
| Solyc06g074400.2 | Solyc06g074400.4 |
| Solyc06g074410.2 | Solyc06g074410.4 |
| Solyc06g074430.2 | Solyc06g074430.4 |

| | |
|---|---|
| Solyc06g074450.1 | Solyc06g074450.3 |
| Solyc06g074460.2 | Solyc06g074460.4 |
| Solyc06g074510.2 | Solyc06g074510.4 |
| Solyc06g074630.2 | Solyc06g074630.4 |
| Solyc06g074670.2 | Solyc06g074670.4 |
| Solyc06g074680.2 | Solyc06g074680.4 |
| Solyc06g074700.2 | Solyc06g074700.4 |
| Solyc06g074720.2 | Solyc06g074720.4 |
| Solyc06g074730.2 | Solyc06g074730.4 |
| Solyc06g074770.2 | Solyc06g074770.4 |
| Solyc06g074870.1 | Solyc06g074870.3 |
| Solyc06g074920.1 | Solyc06g074920.3 |
| Solyc06g074940.2 | Solyc06g074940.4 |
| Solyc06g074960.2 | Solyc06g074960.4 |
| Solyc06g074990.1 | Solyc06g074990.3 |
| Solyc06g075010.2 | Solyc06g075010.4 |
| Solyc06g075070.2 | Solyc06g075070.4 |
| Solyc06g075080.2 | Solyc06g075080.4 |
| Solyc06g075090.2 | Solyc06g075090.4 |
| Solyc06g075130.2 | Solyc06g075130.4 |
| Solyc06g075210.2 | Solyc06g075210.4 |
| Solyc06g075270.2 | Solyc06g075270.4 |
| Solyc06g075290.2 | Solyc06g075290.4 |
| Solyc06g075300.1 | Solyc06g075300.3 |
| Solyc06g075310.2 | Solyc06g075310.4 |
| Solyc06g075330.1 | Solyc06g075330.3 |
| Solyc06g075340.2 | Solyc06g075340.4 |
| Solyc06g075350.1 | Solyc06g075350.3 |
| Solyc06g075390.2 | Solyc06g075390.4 |
| Solyc06g075430.1 | Solyc06g075430.3 |
| Solyc06g075440.1 | Solyc06g075440.3 |
| Solyc06g075460.2 | Solyc06g075460.4 |
| Solyc06g075470.1 | Solyc06g075470.3 |
| Solyc06g075480.2 | Solyc06g075480.4 |
| Solyc06g075500.2 | Solyc06g075500.4 |
| Solyc06g075510.2 | Solyc06g075510.4 |
| Solyc06g075580.2 | Solyc06g075580.4 |
| Solyc06g075590.2 | Solyc06g075590.4 |
| Solyc06g075630.2 | Solyc06g075630.4 |
| Solyc06g075660.2 | Solyc06g075660.4 |
| Solyc06g075720.1 | Solyc06g075720.3 |
| Solyc06g075780.1 | Solyc06g075780.3 |
| Solyc06g075860.1 | Solyc06g075860.3 |

| | |
|---|---|
| Solyc06g075970.2 | Solyc06g075970.4 |
| Solyc06g075990.2 | Solyc06g075990.4 |
| Solyc06g076050.2 | Solyc06g076050.4 |
| Solyc06g076080.1 | Solyc06g076080.3 |
| Solyc06g076140.2 | Solyc06g076140.4 |
| Solyc06g076160.2 | Solyc06g076160.4 |
| Solyc06g076170.2 | Solyc06g076170.4 |
| Solyc06g076210.1 | Solyc06g076210.3 |
| Solyc06g076250.2 | Solyc06g076250.4 |
| Solyc06g076280.1 | Solyc06g076280.3 |
| Solyc06g076360.2 | Solyc06g076360.4 |
| Solyc06g076410.2 | Solyc06g076410.4 |
| Solyc06g076440.1 | Solyc06g076440.3 |
| Solyc06g076460.2 | Solyc06g076460.4 |
| Solyc06g076480.2 | Solyc06g076480.4 |
| Solyc06g076550.2 | Solyc06g076550.4 |
| Solyc06g076650.2 | Solyc06g076650.4 |
| Solyc06g076680.2 | Solyc06g076680.4 |
| Solyc06g076690.2 | Solyc06g076690.4 |
| Solyc06g076710.1 | Solyc06g076710.3 |
| Solyc06g076740.1 | Solyc06g076740.3 |
| Solyc06g076840.1 | Solyc06g076840.3 |
| Solyc06g076860.2 | Solyc06g076860.4 |
| Solyc06g076870.2 | Solyc06g076870.4 |
| Solyc06g076900.2 | Solyc06g076900.4 |
| Solyc06g076910.1 | Solyc06g076910.3 |
| Solyc06g076980.2 | Solyc06g076980.4 |
| Solyc06g082000.2 | Solyc06g082000.4 |
| Solyc06g082010.2 | Solyc06g082010.4 |
| Solyc06g082020.2 | Solyc06g082020.4 |
| Solyc06g082090.2 | Solyc06g082090.4 |
| Solyc06g082100.2 | Solyc06g082100.4 |
| Solyc06g082120.2 | Solyc06g082120.4 |
| Solyc06g082150.2 | Solyc06g082150.4 |
| Solyc06g082160.2 | Solyc06g082160.4 |
| Solyc06g082180.1 | Solyc06g082180.3 |
| Solyc06g082200.2 | Solyc06g082200.4 |
| Solyc06g082220.2 | Solyc06g082220.4 |
| Solyc06g082230.2 | Solyc06g082230.4 |
| Solyc06g082260.2 | Solyc06g082260.4 |
| Solyc06g082290.2 | Solyc06g082290.4 |
| Solyc06g082300.2 | Solyc06g082300.4 |
| Solyc06g082350.2 | Solyc06g082350.4 |

| | |
|---|---|
| Solyc06g082360.2 | Solyc06g082360.4 |
| Solyc06g082370.2 | Solyc06g082370.4 |
| Solyc06g082390.2 | Solyc06g082390.4 |
| Solyc06g082420.2 | Solyc06g082420.4 |
| Solyc06g082490.2 | Solyc06g082490.4 |
| Solyc06g082530.1 | Solyc06g082530.3 |
| Solyc06g082560.1 | Solyc06g082560.3 |
| Solyc06g082570.1 | Solyc06g082570.3 |
| Solyc06g082580.2 | Solyc06g082580.4 |
| Solyc06g082610.2 | Solyc06g082610.4 |
| Solyc06g082630.2 | Solyc06g082630.4 |
| Solyc06g082650.2 | Solyc06g082650.4 |
| Solyc06g082660.2 | Solyc06g082660.4 |
| Solyc06g082680.2 | Solyc06g082680.4 |
| Solyc06g082710.2 | Solyc06g082710.4 |
| Solyc06g082760.2 | Solyc06g082760.4 |
| Solyc06g082780.2 | Solyc06g082780.4 |
| Solyc06g082820.2 | Solyc06g082820.4 |
| Solyc06g082840.2 | Solyc06g082840.4 |
| Solyc06g082860.2 | Solyc06g082860.4 |
| Solyc06g082870.2 | Solyc06g082870.4 |
| Solyc06g082890.2 | Solyc06g082890.4 |
| Solyc06g082950.2 | Solyc06g082950.4 |
| Solyc06g082990.2 | Solyc06g082990.4 |
| Solyc06g083020.1 | Solyc06g083020.3 |
| Solyc06g083060.2 | Solyc06g083060.4 |
| Solyc06g083090.2 | Solyc06g083090.4 |
| Solyc06g083130.2 | Solyc06g083130.4 |
| Solyc06g083140.2 | Solyc06g083140.4 |
| Solyc06g083180.2 | Solyc06g083180.4 |
| Solyc06g083190.2 | Solyc06g083190.4 |
| Solyc06g083200.1 | Solyc06g083200.3 |
| Solyc06g083250.2 | Solyc06g083250.4 |
| Solyc06g083300.2 | Solyc06g083300.4 |
| Solyc06g083390.2 | Solyc06g083390.4 |
| Solyc06g083400.1 | Solyc06g083400.3 |
| Solyc06g083410.2 | Solyc06g083410.4 |
| Solyc06g083420.1 | Solyc06g083420.3 |
| Solyc06g083450.2 | Solyc06g083450.4 |
| Solyc06g083460.2 | Solyc06g083460.4 |
| Solyc06g083470.2 | Solyc06g083470.4 |
| Solyc06g083480.2 | Solyc06g083480.4 |
| Solyc06g083500.2 | Solyc06g083500.4 |

| | |
|---|---|
| Solyc06g083540.2 | Solyc06g083540.4 |
| Solyc06g083580.2 | Solyc06g083580.4 |
| Solyc06g083590.2 | Solyc06g083590.4 |
| Solyc06g083600.1 | Solyc06g083600.3 |
| Solyc06g083630.2 | Solyc06g083630.4 |
| Solyc06g083640.2 | Solyc06g083640.4 |
| Solyc06g083660.2 | Solyc06g083660.4 |
| Solyc06g083670.2 | Solyc06g083670.4 |
| Solyc06g083700.2 | Solyc06g083700.4 |
| Solyc06g083710.1 | Solyc06g083710.3 |
| Solyc06g083770.2 | Solyc06g083770.4 |
| Solyc06g083800.2 | Solyc06g083800.4 |
| Solyc06g083830.2 | Solyc06g083830.4 |
| Solyc06g083850.2 | Solyc06g083850.4 |
| Solyc06g083890.2 | Solyc06g083890.4 |
| Solyc06g083920.2 | Solyc06g083920.4 |
| Solyc06g083950.1 | Solyc06g083950.3 |
| Solyc06g083960.1 | Solyc06g083960.3 |
| Solyc06g083980.1 | Solyc06g083980.3 |
| Solyc06g084000.2 | Solyc06g084000.4 |
| Solyc06g084020.2 | Solyc06g084020.4 |
| Solyc06g084050.2 | Solyc06g084050.4 |
| Solyc06g084080.2 | Solyc06g084080.4 |
| Solyc06g084090.2 | Solyc06g084090.4 |
| Solyc06g084120.2 | Solyc06g084120.4 |
| Solyc06g084140.2 | Solyc06g084140.4 |
| Solyc06g084150.2 | Solyc06g084150.4 |
| Solyc06g084160.2 | Solyc06g084160.4 |
| Solyc06g084250.2 | Solyc06g084250.4 |
| Solyc06g084310.2 | Solyc06g084310.4 |
| Solyc06g084360.2 | Solyc06g084360.4 |
| Solyc06g084380.2 | Solyc06g084380.4 |
| Solyc06g084410.2 | Solyc06g084410.4 |
| Solyc06g084430.2 | Solyc06g084430.4 |
| Solyc06g084450.2 | Solyc06g084450.4 |
| Solyc06g084460.2 | Solyc06g084460.4 |
| Solyc06g084510.2 | Solyc06g084510.4 |
| Solyc06g084570.2 | Solyc06g084570.4 |
| Solyc06g084590.2 | Solyc06g084590.4 |
| Solyc06g084610.2 | Solyc06g084610.4 |
| Solyc06g084760.1 | Solyc06g084760.3 |
| Solyc07g005010.2 | Solyc07g005010.4 |
| Solyc07g005030.2 | Solyc07g005030.4 |

| | |
|---|---|
| Solyc07g005090.2 | Solyc07g005090.4 |
| Solyc07g005100.2 | Solyc07g005100.4 |
| Solyc07g005110.2 | Solyc07g005110.4 |
| Solyc07g005140.2 | Solyc07g005140.4 |
| Solyc07g005150.1 | Solyc07g005150.3 |
| Solyc07g005170.1 | Solyc07g005170.3 |
| Solyc07g005240.2 | Solyc07g005240.4 |
| Solyc07g005310.1 | Solyc07g005310.3 |
| Solyc07g005370.2 | Solyc07g005370.4 |
| Solyc07g005380.2 | Solyc07g005380.4 |
| Solyc07g005460.2 | Solyc07g005460.4 |
| Solyc07g005470.2 | Solyc07g005470.4 |
| Solyc07g005490.2 | Solyc07g005490.4 |
| Solyc07g005530.2 | Solyc07g005530.4 |
| Solyc07g005570.2 | Solyc07g005570.4 |
| Solyc07g005580.2 | Solyc07g005580.4 |
| Solyc07g005600.2 | Solyc07g005600.4 |
| Solyc07g005650.2 | Solyc07g005650.4 |
| Solyc07g005660.2 | Solyc07g005660.4 |
| Solyc07g005670.1 | Solyc07g005670.3 |
| Solyc07g005680.2 | Solyc07g005680.4 |
| Solyc07g005730.2 | Solyc07g005730.4 |
| Solyc07g005790.2 | Solyc07g005790.4 |
| Solyc07g005850.1 | Solyc07g005850.3 |
| Solyc07g005920.1 | Solyc07g005920.3 |
| Solyc07g005930.2 | Solyc07g005930.4 |
| Solyc07g005940.2 | Solyc07g005940.4 |
| Solyc07g005950.2 | Solyc07g005950.4 |
| Solyc07g006030.2 | Solyc07g006030.4 |
| Solyc07g006050.1 | Solyc07g006050.3 |
| Solyc07g006090.1 | Solyc07g006090.3 |
| Solyc07g006100.2 | Solyc07g006100.4 |
| Solyc07g006110.2 | Solyc07g006110.4 |
| Solyc07g006130.1 | Solyc07g006130.3 |
| Solyc07g006140.2 | Solyc07g006140.4 |
| Solyc07g006170.2 | Solyc07g006170.4 |
| Solyc07g006240.2 | Solyc07g006240.4 |
| Solyc07g006290.1 | Solyc07g006290.3 |
| Solyc07g006300.2 | Solyc07g006300.4 |
| Solyc07g006390.1 | Solyc07g006390.3 |
| Solyc07g006400.1 | Solyc07g006400.3 |
| Solyc07g006480.2 | Solyc07g006480.4 |
| Solyc07g006570.2 | Solyc07g006570.4 |

| | |
|---|---|
| Solyc07g006620.2 | Solyc07g006620.4 |
| Solyc07g006630.2 | Solyc07g006630.4 |
| Solyc07g006650.2 | Solyc07g006650.4 |
| Solyc07g006720.2 | Solyc07g006720.4 |
| Solyc07g006730.2 | Solyc07g006730.4 |
| Solyc07g006760.2 | Solyc07g006760.4 |
| Solyc07g006820.2 | Solyc07g006820.4 |
| Solyc07g006830.2 | Solyc07g006830.4 |
| Solyc07g006850.1 | Solyc07g006850.3 |
| Solyc07g006870.2 | Solyc07g006870.4 |
| Solyc07g006910.2 | Solyc07g006910.4 |
| Solyc07g006940.1 | Solyc07g006940.3 |
| Solyc07g006970.2 | Solyc07g006970.4 |
| Solyc07g006990.1 | Solyc07g006990.3 |
| Solyc07g007040.2 | Solyc07g007040.4 |
| Solyc07g007060.1 | Solyc07g007060.3 |
| Solyc07g007110.2 | Solyc07g007110.4 |
| Solyc07g007180.2 | Solyc07g007180.4 |
| Solyc07g007200.2 | Solyc07g007200.4 |
| Solyc07g007230.2 | Solyc07g007230.4 |
| Solyc07g007250.2 | Solyc07g007250.4 |
| Solyc07g007260.2 | Solyc07g007260.4 |
| Solyc07g007280.2 | Solyc07g007280.4 |
| Solyc07g007320.1 | Solyc07g007320.3 |
| Solyc07g007360.1 | Solyc07g007360.3 |
| Solyc07g007380.2 | Solyc07g007380.4 |
| Solyc07g007410.2 | Solyc07g007410.4 |
| Solyc07g007460.1 | Solyc07g007460.3 |
| Solyc07g007510.2 | Solyc07g007510.4 |
| Solyc07g007640.1 | Solyc07g007640.3 |
| Solyc07g007650.1 | Solyc07g007650.3 |
| Solyc07g007680.2 | Solyc07g007680.4 |
| Solyc07g007690.2 | Solyc07g007690.4 |
| Solyc07g007700.1 | Solyc07g007700.3 |
| Solyc07g007710.2 | Solyc07g007710.4 |
| Solyc07g007720.2 | Solyc07g007720.4 |
| Solyc07g007730.2 | Solyc07g007730.4 |
| Solyc07g007830.2 | Solyc07g007830.4 |
| Solyc07g007840.1 | Solyc07g007840.3 |
| Solyc07g007920.1 | Solyc07g007920.3 |
| Solyc07g007990.2 | Solyc07g007990.4 |
| Solyc07g008000.2 | Solyc07g008000.4 |
| Solyc07g008010.2 | Solyc07g008010.4 |

| | |
|---|---|
| Solyc07g008030.1 | Solyc07g008030.3 |
| Solyc07g008040.2 | Solyc07g008040.4 |
| Solyc07g008050.2 | Solyc07g008050.4 |
| Solyc07g008090.2 | Solyc07g008090.4 |
| Solyc07g008180.2 | Solyc07g008180.4 |
| Solyc07g008190.2 | Solyc07g008190.4 |
| Solyc07g008210.2 | Solyc07g008210.4 |
| Solyc07g008290.2 | Solyc07g008290.4 |
| Solyc07g008300.1 | Solyc07g008300.3 |
| Solyc07g008340.2 | Solyc07g008340.4 |
| Solyc07g008370.2 | Solyc07g008370.4 |
| Solyc07g008390.1 | Solyc07g008390.3 |
| Solyc07g008400.1 | Solyc07g008400.3 |
| Solyc07g008410.2 | Solyc07g008410.4 |
| Solyc07g008480.1 | Solyc07g008480.3 |
| Solyc07g008490.2 | Solyc07g008490.4 |
| Solyc07g008580.1 | Solyc07g008580.3 |
| Solyc07g008610.1 | Solyc07g008610.3 |
| Solyc07g008640.1 | Solyc07g008640.3 |
| Solyc07g008650.2 | Solyc07g008650.4 |
| Solyc07g008680.2 | Solyc07g008680.4 |
| Solyc07g008740.1 | Solyc07g008740.3 |
| Solyc07g008750.2 | Solyc07g008750.4 |
| Solyc07g008760.2 | Solyc07g008760.4 |
| Solyc07g008790.1 | Solyc07g008790.3 |
| Solyc07g008800.2 | Solyc07g008800.4 |
| Solyc07g008890.1 | Solyc07g008890.3 |
| Solyc07g008900.2 | Solyc07g008900.4 |
| Solyc07g008920.2 | Solyc07g008920.4 |
| Solyc07g008950.2 | Solyc07g008950.4 |
| Solyc07g008970.2 | Solyc07g008970.4 |
| Solyc07g009060.2 | Solyc07g009060.4 |
| Solyc07g009070.2 | Solyc07g009070.4 |
| Solyc07g009080.2 | Solyc07g009080.4 |
| Solyc07g009090.2 | Solyc07g009090.4 |
| Solyc07g009110.1 | Solyc07g009110.3 |
| Solyc07g009130.2 | Solyc07g009130.4 |
| Solyc07g009170.2 | Solyc07g009170.4 |
| Solyc07g009210.1 | Solyc07g009210.3 |
| Solyc07g009330.2 | Solyc07g009330.4 |
| Solyc07g009350.2 | Solyc07g009350.4 |
| Solyc07g009360.1 | Solyc07g009360.3 |
| Solyc07g009380.2 | Solyc07g009380.4 |

| | |
|---|---|
| Solyc07g009410.2 | Solyc07g009410.4 |
| Solyc07g009470.1 | Solyc07g009470.3 |
| Solyc07g009500.1 | Solyc07g009500.3 |
| Solyc07g009560.1 | Solyc07g009560.3 |
| Solyc07g014590.2 | Solyc07g014590.4 |
| Solyc07g014610.2 | Solyc07g014610.4 |
| Solyc07g014660.1 | Solyc07g014660.3 |
| Solyc07g014680.2 | Solyc07g014680.4 |
| Solyc07g014700.2 | Solyc07g014700.4 |
| Solyc07g015810.1 | Solyc07g015810.3 |
| Solyc07g015870.2 | Solyc07g015870.4 |
| Solyc07g015900.1 | Solyc07g015900.3 |
| Solyc07g015910.2 | Solyc07g015910.4 |
| Solyc07g015920.1 | Solyc07g015920.3 |
| Solyc07g015930.1 | Solyc07g015930.3 |
| Solyc07g015970.1 | Solyc07g015970.3 |
| Solyc07g015980.2 | Solyc07g015980.4 |
| Solyc07g016040.1 | Solyc07g016040.3 |
| Solyc07g016060.1 | Solyc07g016060.3 |
| Solyc07g016080.2 | Solyc07g016080.4 |
| Solyc07g016120.1 | Solyc07g016120.3 |
| Solyc07g016150.2 | Solyc07g016150.4 |
| Solyc07g016170.2 | Solyc07g016170.4 |
| Solyc07g017250.2 | Solyc07g017250.4 |
| Solyc07g017280.1 | Solyc07g017280.3 |
| Solyc07g017330.1 | Solyc07g017330.3 |
| Solyc07g017370.1 | Solyc07g017370.3 |
| Solyc07g017470.1 | Solyc07g017470.3 |
| Solyc07g017520.2 | Solyc07g017520.4 |
| Solyc07g017530.2 | Solyc07g017530.4 |
| Solyc07g017540.2 | Solyc07g017540.4 |
| Solyc07g017560.2 | Solyc07g017560.4 |
| Solyc07g017660.1 | Solyc07g017660.3 |
| Solyc07g017700.1 | Solyc07g017700.3 |
| Solyc07g017750.2 | Solyc07g017750.4 |
| Solyc07g017780.2 | Solyc07g017780.4 |
| Solyc07g017800.2 | Solyc07g017800.4 |
| Solyc07g017820.1 | Solyc07g017820.3 |
| Solyc07g017880.2 | Solyc07g017880.4 |
| Solyc07g017960.2 | Solyc07g017960.4 |
| Solyc07g017970.1 | Solyc07g017970.3 |
| Solyc07g018070.2 | Solyc07g018070.4 |
| Solyc07g018090.2 | Solyc07g018090.4 |

| | |
|---|---|
| Solyc07g018180.1 | Solyc07g018180.3 |
| Solyc07g018190.2 | Solyc07g018190.4 |
| Solyc07g018290.2 | Solyc07g018290.4 |
| Solyc07g018320.1 | Solyc07g018320.3 |
| Solyc07g018370.1 | Solyc07g018370.3 |
| Solyc07g018390.1 | Solyc07g018390.3 |
| Solyc07g019510.2 | Solyc07g019510.4 |
| Solyc07g019620.1 | Solyc07g019620.3 |
| Solyc07g019650.2 | Solyc07g019650.4 |
| Solyc07g019670.2 | Solyc07g019670.4 |
| Solyc07g020790.2 | Solyc07g020790.4 |
| Solyc07g020870.1 | Solyc07g020870.3 |
| Solyc07g021020.2 | Solyc07g021020.4 |
| Solyc07g021030.1 | Solyc07g021030.3 |
| Solyc07g021050.1 | Solyc07g021050.3 |
| Solyc07g021120.1 | Solyc07g021120.3 |
| Solyc07g021180.1 | Solyc07g021180.3 |
| Solyc07g021220.1 | Solyc07g021220.3 |
| Solyc07g021330.1 | Solyc07g021330.3 |
| Solyc07g021450.1 | Solyc07g021450.3 |
| Solyc07g021460.1 | Solyc07g021460.3 |
| Solyc07g021490.1 | Solyc07g021490.3 |
| Solyc07g021510.1 | Solyc07g021510.3 |
| Solyc07g021650.1 | Solyc07g021650.3 |
| Solyc07g021730.1 | Solyc07g021730.3 |
| Solyc07g022760.2 | Solyc07g022760.4 |
| Solyc07g022790.2 | Solyc07g022790.4 |
| Solyc07g022860.2 | Solyc07g022860.4 |
| Solyc07g022900.2 | Solyc07g022900.4 |
| Solyc07g023990.1 | Solyc07g023990.3 |
| Solyc07g024070.1 | Solyc07g024070.3 |
| Solyc07g024080.1 | Solyc07g024080.3 |
| Solyc07g025140.2 | Solyc07g025140.4 |
| Solyc07g025250.1 | Solyc07g025250.3 |
| Solyc07g025400.2 | Solyc07g025400.4 |
| Solyc07g025500.1 | Solyc07g025500.3 |
| Solyc07g025510.2 | Solyc07g025510.4 |
| Solyc07g026560.1 | Solyc07g026560.3 |
| Solyc07g026620.2 | Solyc07g026620.4 |
| Solyc07g026670.2 | Solyc07g026670.4 |
| Solyc07g026710.1 | Solyc07g026710.3 |
| Solyc07g026720.1 | Solyc07g026720.3 |
| Solyc07g026860.2 | Solyc07g026860.4 |

| | |
|---|---|
| Solyc07g026880.1 | Solyc07g026880.3 |
| Solyc07g026910.1 | Solyc07g026910.3 |
| Solyc07g026930.1 | Solyc07g026930.3 |
| Solyc07g027020.2 | Solyc07g027020.4 |
| Solyc07g032020.1 | Solyc07g032020.3 |
| Solyc07g032070.1 | Solyc07g032070.3 |
| Solyc07g032090.2 | Solyc07g032090.4 |
| Solyc07g032140.1 | Solyc07g032140.3 |
| Solyc07g032380.2 | Solyc07g032380.4 |
| Solyc07g032420.1 | Solyc07g032420.3 |
| Solyc07g032620.1 | Solyc07g032620.3 |
| Solyc07g032670.1 | Solyc07g032670.3 |
| Solyc07g032730.2 | Solyc07g032730.4 |
| Solyc07g032740.2 | Solyc07g032740.4 |
| Solyc07g037910.1 | Solyc07g037910.3 |
| Solyc07g037920.1 | Solyc07g037920.3 |
| Solyc07g038110.1 | Solyc07g038110.3 |
| Solyc07g038130.1 | Solyc07g038130.3 |
| Solyc07g038160.2 | Solyc07g038160.4 |
| Solyc07g039200.2 | Solyc07g039200.4 |
| Solyc07g039210.2 | Solyc07g039210.4 |
| Solyc07g039330.2 | Solyc07g039330.4 |
| Solyc07g039340.2 | Solyc07g039340.4 |
| Solyc07g039400.1 | Solyc07g039400.3 |
| Solyc07g039500.2 | Solyc07g039500.4 |
| Solyc07g039510.2 | Solyc07g039510.4 |
| Solyc07g039550.2 | Solyc07g039550.4 |
| Solyc07g040660.2 | Solyc07g040660.4 |
| Solyc07g040700.1 | Solyc07g040700.3 |
| Solyc07g040780.2 | Solyc07g040780.4 |
| Solyc07g040880.2 | Solyc07g040880.4 |
| Solyc07g040900.1 | Solyc07g040900.3 |
| Solyc07g040990.2 | Solyc07g040990.4 |
| Solyc07g041000.2 | Solyc07g041000.4 |
| Solyc07g041150.2 | Solyc07g041150.4 |
| Solyc07g041180.2 | Solyc07g041180.4 |
| Solyc07g041260.1 | Solyc07g041260.3 |
| Solyc07g041280.2 | Solyc07g041280.4 |
| Solyc07g041290.1 | Solyc07g041290.3 |
| Solyc07g041320.1 | Solyc07g041320.3 |
| Solyc07g041430.1 | Solyc07g041430.3 |
| Solyc07g041440.1 | Solyc07g041440.3 |
| Solyc07g041470.1 | Solyc07g041470.3 |

| | |
|---|---|
| Solyc07g041490.1 | Solyc07g041490.3 |
| Solyc07g041510.1 | Solyc07g041510.3 |
| Solyc07g041650.1 | Solyc07g041650.3 |
| Solyc07g041660.2 | Solyc07g041660.4 |
| Solyc07g041730.2 | Solyc07g041730.4 |
| Solyc07g041820.1 | Solyc07g041820.3 |
| Solyc07g041830.2 | Solyc07g041830.4 |
| Solyc07g041850.1 | Solyc07g041850.3 |
| Solyc07g041920.2 | Solyc07g041920.4 |
| Solyc07g041970.2 | Solyc07g041970.4 |
| Solyc07g042000.2 | Solyc07g042000.4 |
| Solyc07g042010.2 | Solyc07g042010.4 |
| Solyc07g042080.2 | Solyc07g042080.4 |
| Solyc07g042090.2 | Solyc07g042090.4 |
| Solyc07g042130.2 | Solyc07g042130.4 |
| Solyc07g042180.2 | Solyc07g042180.4 |
| Solyc07g042260.2 | Solyc07g042260.4 |
| Solyc07g042270.2 | Solyc07g042270.4 |
| Solyc07g042280.2 | Solyc07g042280.4 |
| Solyc07g042290.1 | Solyc07g042290.3 |
| Solyc07g042380.2 | Solyc07g042380.4 |
| Solyc07g042390.1 | Solyc07g042390.3 |
| Solyc07g042520.2 | Solyc07g042520.4 |
| Solyc07g042560.2 | Solyc07g042560.4 |
| Solyc07g042590.2 | Solyc07g042590.4 |
| Solyc07g042600.1 | Solyc07g042600.3 |
| Solyc07g042620.2 | Solyc07g042620.4 |
| Solyc07g042630.2 | Solyc07g042630.4 |
| Solyc07g042690.2 | Solyc07g042690.4 |
| Solyc07g042750.2 | Solyc07g042750.4 |
| Solyc07g042810.1 | Solyc07g042810.3 |
| Solyc07g042830.2 | Solyc07g042830.4 |
| Solyc07g042900.2 | Solyc07g042900.4 |
| Solyc07g042970.2 | Solyc07g042970.4 |
| Solyc07g043170.2 | Solyc07g043170.4 |
| Solyc07g043240.2 | Solyc07g043240.4 |
| Solyc07g043260.2 | Solyc07g043260.4 |
| Solyc07g043330.1 | Solyc07g043330.3 |
| Solyc07g043350.1 | Solyc07g043350.3 |
| Solyc07g043390.2 | Solyc07g043390.4 |
| Solyc07g043450.1 | Solyc07g043450.3 |
| Solyc07g043470.1 | Solyc07g043470.3 |
| Solyc07g043560.2 | Solyc07g043560.4 |

| | |
|---|---|
| Solyc07g043580.2 | Solyc07g043580.4 |
| Solyc07g043600.2 | Solyc07g043600.4 |
| Solyc07g043650.1 | Solyc07g043650.3 |
| Solyc07g043700.1 | Solyc07g043700.3 |
| Solyc07g043710.2 | Solyc07g043710.4 |
| Solyc07g044720.1 | Solyc07g044720.3 |
| Solyc07g044730.2 | Solyc07g044730.4 |
| Solyc07g044870.2 | Solyc07g044870.4 |
| Solyc07g045000.2 | Solyc07g045000.4 |
| Solyc07g045030.2 | Solyc07g045030.4 |
| Solyc07g045180.2 | Solyc07g045180.4 |
| Solyc07g045190.1 | Solyc07g045190.3 |
| Solyc07g045230.2 | Solyc07g045230.4 |
| Solyc07g045280.1 | Solyc07g045280.3 |
| Solyc07g045310.2 | Solyc07g045310.4 |
| Solyc07g045340.2 | Solyc07g045340.4 |
| Solyc07g045350.2 | Solyc07g045350.4 |
| Solyc07g045390.1 | Solyc07g045390.3 |
| Solyc07g045420.2 | Solyc07g045420.4 |
| Solyc07g045430.2 | Solyc07g045430.4 |
| Solyc07g045470.1 | Solyc07g045470.3 |
| Solyc07g045490.1 | Solyc07g045490.3 |
| Solyc07g045510.2 | Solyc07g045510.4 |
| Solyc07g045520.2 | Solyc07g045520.4 |
| Solyc07g045540.2 | Solyc07g045540.4 |
| Solyc07g045550.1 | Solyc07g045550.3 |
| Solyc07g045570.1 | Solyc07g045570.3 |
| Solyc07g047620.1 | Solyc07g047620.3 |
| Solyc07g047680.1 | Solyc07g047680.3 |
| Solyc07g047770.2 | Solyc07g047770.4 |
| Solyc07g047820.1 | Solyc07g047820.3 |
| Solyc07g047860.2 | Solyc07g047860.4 |
| Solyc07g047920.1 | Solyc07g047920.3 |
| Solyc07g047940.2 | Solyc07g047940.4 |
| Solyc07g047970.2 | Solyc07g047970.4 |
| Solyc07g047980.1 | Solyc07g047980.3 |
| Solyc07g048010.2 | Solyc07g048010.4 |
| Solyc07g048020.1 | Solyc07g048020.3 |
| Solyc07g048030.2 | Solyc07g048030.4 |
| Solyc07g048050.1 | Solyc07g048050.3 |
| Solyc07g048060.1 | Solyc07g048060.3 |
| Solyc07g048080.2 | Solyc07g048080.4 |
| Solyc07g048100.1 | Solyc07g048100.3 |

| | |
|---|---|
| Solyc07g048110.2 | Solyc07g048110.4 |
| Solyc07g049160.2 | Solyc07g049160.4 |
| Solyc07g049190.2 | Solyc07g049190.4 |
| Solyc07g049220.2 | Solyc07g049220.4 |
| Solyc07g049260.2 | Solyc07g049260.4 |
| Solyc07g049270.2 | Solyc07g049270.4 |
| Solyc07g049320.2 | Solyc07g049320.4 |
| Solyc07g049390.1 | Solyc07g049390.3 |
| Solyc07g049470.2 | Solyc07g049470.4 |
| Solyc07g049490.1 | Solyc07g049490.3 |
| Solyc07g049600.2 | Solyc07g049600.4 |
| Solyc07g049670.2 | Solyc07g049670.4 |
| Solyc07g049680.2 | Solyc07g049680.4 |
| Solyc07g049720.2 | Solyc07g049720.4 |
| Solyc07g049740.2 | Solyc07g049740.4 |
| Solyc07g049780.1 | Solyc07g049780.3 |
| Solyc07g051800.1 | Solyc07g051800.3 |
| Solyc07g051840.2 | Solyc07g051840.4 |
| Solyc07g051930.1 | Solyc07g051930.3 |
| Solyc07g051960.1 | Solyc07g051960.3 |
| Solyc07g051970.1 | Solyc07g051970.3 |
| Solyc07g051980.1 | Solyc07g051980.3 |
| Solyc07g052010.1 | Solyc07g052010.3 |
| Solyc07g052040.1 | Solyc07g052040.3 |
| Solyc07g052070.1 | Solyc07g052070.3 |
| Solyc07g052100.2 | Solyc07g052100.4 |
| Solyc07g052110.2 | Solyc07g052110.4 |
| Solyc07g052120.2 | Solyc07g052120.4 |
| Solyc07g052150.2 | Solyc07g052150.4 |
| Solyc07g052250.2 | Solyc07g052250.4 |
| Solyc07g052330.1 | Solyc07g052330.3 |
| Solyc07g052340.2 | Solyc07g052340.4 |
| Solyc07g052370.2 | Solyc07g052370.4 |
| Solyc07g052410.2 | Solyc07g052410.4 |
| Solyc07g052490.2 | Solyc07g052490.4 |
| Solyc07g052560.1 | Solyc07g052560.3 |
| Solyc07g052570.2 | Solyc07g052570.4 |
| Solyc07g052630.1 | Solyc07g052630.3 |
| Solyc07g052690.2 | Solyc07g052690.4 |
| Solyc07g052720.2 | Solyc07g052720.4 |
| Solyc07g052760.1 | Solyc07g052760.3 |
| Solyc07g052790.1 | Solyc07g052790.3 |
| Solyc07g052800.1 | Solyc07g052800.3 |

| | |
|---|---|
| Solyc07g052810.1 | Solyc07g052810.3 |
| Solyc07g052830.2 | Solyc07g052830.4 |
| Solyc07g052860.1 | Solyc07g052860.3 |
| Solyc07g052880.2 | Solyc07g052880.4 |
| Solyc07g052940.2 | Solyc07g052940.4 |
| Solyc07g052960.1 | Solyc07g052960.3 |
| Solyc07g053010.1 | Solyc07g053010.3 |
| Solyc07g053020.1 | Solyc07g053020.3 |
| Solyc07g053030.2 | Solyc07g053030.4 |
| Solyc07g053040.1 | Solyc07g053040.3 |
| Solyc07g053070.1 | Solyc07g053070.3 |
| Solyc07g053080.2 | Solyc07g053080.4 |
| Solyc07g053100.2 | Solyc07g053100.4 |
| Solyc07g053120.2 | Solyc07g053120.4 |
| Solyc07g053130.2 | Solyc07g053130.4 |
| Solyc07g053160.2 | Solyc07g053160.4 |
| Solyc07g053170.2 | Solyc07g053170.4 |
| Solyc07g053190.2 | Solyc07g053190.4 |
| Solyc07g053200.2 | Solyc07g053200.4 |
| Solyc07g053280.2 | Solyc07g053280.4 |
| Solyc07g053350.2 | Solyc07g053350.4 |
| Solyc07g053360.2 | Solyc07g053360.4 |
| Solyc07g053410.2 | Solyc07g053410.4 |
| Solyc07g053420.2 | Solyc07g053420.4 |
| Solyc07g053430.2 | Solyc07g053430.4 |
| Solyc07g053450.2 | Solyc07g053450.4 |
| Solyc07g053460.2 | Solyc07g053460.4 |
| Solyc07g053470.2 | Solyc07g053470.4 |
| Solyc07g053480.2 | Solyc07g053480.4 |
| Solyc07g053550.1 | Solyc07g053550.3 |
| Solyc07g053570.2 | Solyc07g053570.4 |
| Solyc07g053580.1 | Solyc07g053580.3 |
| Solyc07g053590.2 | Solyc07g053590.4 |
| Solyc07g053630.2 | Solyc07g053630.4 |
| Solyc07g053650.2 | Solyc07g053650.4 |
| Solyc07g053700.2 | Solyc07g053700.4 |
| Solyc07g053730.2 | Solyc07g053730.4 |
| Solyc07g053760.2 | Solyc07g053760.4 |
| Solyc07g053780.2 | Solyc07g053780.4 |
| Solyc07g053870.2 | Solyc07g053870.4 |
| Solyc07g053880.1 | Solyc07g053880.3 |
| Solyc07g053990.2 | Solyc07g053990.4 |
| Solyc07g054040.1 | Solyc07g054040.3 |

| | |
|---|---|
| Solyc07g054050.1 | Solyc07g054050.3 |
| Solyc07g054070.2 | Solyc07g054070.4 |
| Solyc07g054080.1 | Solyc07g054080.3 |
| Solyc07g054100.1 | Solyc07g054100.3 |
| Solyc07g054180.2 | Solyc07g054180.4 |
| Solyc07g054190.2 | Solyc07g054190.4 |
| Solyc07g054200.2 | Solyc07g054200.4 |
| Solyc07g054310.1 | Solyc07g054310.3 |
| Solyc07g054330.2 | Solyc07g054330.4 |
| Solyc07g054400.1 | Solyc07g054400.3 |
| Solyc07g054450.2 | Solyc07g054450.4 |
| Solyc07g054460.2 | Solyc07g054460.4 |
| Solyc07g054480.1 | Solyc07g054480.3 |
| Solyc07g054500.2 | Solyc07g054500.4 |
| Solyc07g054540.2 | Solyc07g054540.4 |
| Solyc07g054620.2 | Solyc07g054620.4 |
| Solyc07g054630.2 | Solyc07g054630.4 |
| Solyc07g054670.2 | Solyc07g054670.4 |
| Solyc07g054820.2 | Solyc07g054820.4 |
| Solyc07g054830.2 | Solyc07g054830.4 |
| Solyc07g054840.2 | Solyc07g054840.4 |
| Solyc07g054870.2 | Solyc07g054870.4 |
| Solyc07g054880.2 | Solyc07g054880.4 |
| Solyc07g054910.1 | Solyc07g054910.3 |
| Solyc07g054980.1 | Solyc07g054980.3 |
| Solyc07g055100.2 | Solyc07g055100.4 |
| Solyc07g055120.2 | Solyc07g055120.4 |
| Solyc07g055130.2 | Solyc07g055130.4 |
| Solyc07g055160.2 | Solyc07g055160.4 |
| Solyc07g055200.2 | Solyc07g055200.4 |
| Solyc07g055220.1 | Solyc07g055220.3 |
| Solyc07g055230.2 | Solyc07g055230.4 |
| Solyc07g055280.2 | Solyc07g055280.4 |
| Solyc07g055320.2 | Solyc07g055320.4 |
| Solyc07g055350.2 | Solyc07g055350.4 |
| Solyc07g055490.2 | Solyc07g055490.4 |
| Solyc07g055560.2 | Solyc07g055560.4 |
| Solyc07g055580.1 | Solyc07g055580.3 |
| Solyc07g055610.1 | Solyc07g055610.3 |
| Solyc07g055630.2 | Solyc07g055630.4 |
| Solyc07g055720.2 | Solyc07g055720.4 |
| Solyc07g055730.2 | Solyc07g055730.4 |
| Solyc07g055750.2 | Solyc07g055750.4 |

| | |
|---|---|
| Solyc07g055760.1 | Solyc07g055760.3 |
| Solyc07g055790.1 | Solyc07g055790.3 |
| Solyc07g055850.2 | Solyc07g055850.4 |
| Solyc07g055860.2 | Solyc07g055860.4 |
| Solyc07g055870.2 | Solyc07g055870.4 |
| Solyc07g055920.2 | Solyc07g055920.4 |
| Solyc07g055940.1 | Solyc07g055940.3 |
| Solyc07g056150.2 | Solyc07g056150.4 |
| Solyc07g056160.2 | Solyc07g056160.4 |
| Solyc07g056170.2 | Solyc07g056170.4 |
| Solyc07g056210.2 | Solyc07g056210.4 |
| Solyc07g056260.2 | Solyc07g056260.4 |
| Solyc07g056320.2 | Solyc07g056320.4 |
| Solyc07g056340.2 | Solyc07g056340.4 |
| Solyc07g056390.2 | Solyc07g056390.4 |
| Solyc07g056650.2 | Solyc07g056650.4 |
| Solyc07g056660.2 | Solyc07g056660.4 |
| Solyc07g061740.2 | Solyc07g061740.4 |
| Solyc07g061750.2 | Solyc07g061750.4 |
| Solyc07g061760.1 | Solyc07g061760.3 |
| Solyc07g061780.2 | Solyc07g061780.4 |
| Solyc07g061800.2 | Solyc07g061800.4 |
| Solyc07g061860.1 | Solyc07g061860.3 |
| Solyc07g061880.1 | Solyc07g061880.3 |
| Solyc07g061920.2 | Solyc07g061920.4 |
| Solyc07g061940.2 | Solyc07g061940.4 |
| Solyc07g061950.2 | Solyc07g061950.4 |
| Solyc07g062080.2 | Solyc07g062080.4 |
| Solyc07g062100.2 | Solyc07g062100.4 |
| Solyc07g062120.2 | Solyc07g062120.4 |
| Solyc07g062170.1 | Solyc07g062170.3 |
| Solyc07g062180.2 | Solyc07g062180.4 |
| Solyc07g062190.2 | Solyc07g062190.4 |
| Solyc07g062210.2 | Solyc07g062210.4 |
| Solyc07g062230.2 | Solyc07g062230.4 |
| Solyc07g062310.2 | Solyc07g062310.4 |
| Solyc07g062320.1 | Solyc07g062320.3 |
| Solyc07g062390.2 | Solyc07g062390.4 |
| Solyc07g062460.2 | Solyc07g062460.4 |
| Solyc07g062470.2 | Solyc07g062470.4 |
| Solyc07g062580.2 | Solyc07g062580.4 |
| Solyc07g062590.2 | Solyc07g062590.4 |
| Solyc07g062600.2 | Solyc07g062600.4 |

| | |
|---|---|
| Solyc07g062610.2 | Solyc07g062610.4 |
| Solyc07g062650.2 | Solyc07g062650.4 |
| Solyc07g062660.2 | Solyc07g062660.4 |
| Solyc07g062680.1 | Solyc07g062680.3 |
| Solyc07g062710.2 | Solyc07g062710.4 |
| Solyc07g062720.1 | Solyc07g062720.3 |
| Solyc07g062810.2 | Solyc07g062810.4 |
| Solyc07g062830.1 | Solyc07g062830.3 |
| Solyc07g062890.2 | Solyc07g062890.4 |
| Solyc07g062940.2 | Solyc07g062940.4 |
| Solyc07g062950.2 | Solyc07g062950.4 |
| Solyc07g062980.2 | Solyc07g062980.4 |
| Solyc07g063000.2 | Solyc07g063000.4 |
| Solyc07g063020.1 | Solyc07g063020.3 |
| Solyc07g063030.1 | Solyc07g063030.3 |
| Solyc07g063080.2 | Solyc07g063080.4 |
| Solyc07g063110.2 | Solyc07g063110.4 |
| Solyc07g063120.2 | Solyc07g063120.4 |
| Solyc07g063130.2 | Solyc07g063130.4 |
| Solyc07g063200.1 | Solyc07g063200.3 |
| Solyc07g063220.2 | Solyc07g063220.4 |
| Solyc07g063240.2 | Solyc07g063240.4 |
| Solyc07g063260.2 | Solyc07g063260.4 |
| Solyc07g063280.1 | Solyc07g063280.3 |
| Solyc07g063330.2 | Solyc07g063330.4 |
| Solyc07g063450.1 | Solyc07g063450.3 |
| Solyc07g063460.1 | Solyc07g063460.3 |
| Solyc07g063500.2 | Solyc07g063500.4 |
| Solyc07g063560.2 | Solyc07g063560.4 |
| Solyc07g063570.2 | Solyc07g063570.4 |
| Solyc07g063590.2 | Solyc07g063590.4 |
| Solyc07g063610.2 | Solyc07g063610.4 |
| Solyc07g063650.2 | Solyc07g063650.4 |
| Solyc07g063680.2 | Solyc07g063680.4 |
| Solyc07g063730.1 | Solyc07g063730.3 |
| Solyc07g063860.2 | Solyc07g063860.4 |
| Solyc07g063880.2 | Solyc07g063880.4 |
| Solyc07g063910.2 | Solyc07g063910.4 |
| Solyc07g063930.2 | Solyc07g063930.4 |
| Solyc07g064020.2 | Solyc07g064020.4 |
| Solyc07g064030.2 | Solyc07g064030.4 |
| Solyc07g064070.1 | Solyc07g064070.3 |
| Solyc07g064150.2 | Solyc07g064150.4 |

| | |
|---|---|
| Solyc07g064200.2 | Solyc07g064200.4 |
| Solyc07g064230.2 | Solyc07g064230.4 |
| Solyc07g064240.2 | Solyc07g064240.4 |
| Solyc07g064360.1 | Solyc07g064360.3 |
| Solyc07g064380.2 | Solyc07g064380.4 |
| Solyc07g064450.2 | Solyc07g064450.4 |
| Solyc07g064470.2 | Solyc07g064470.4 |
| Solyc07g064490.2 | Solyc07g064490.4 |
| Solyc07g064560.2 | Solyc07g064560.4 |
| Solyc07g064630.2 | Solyc07g064630.4 |
| Solyc07g064650.1 | Solyc07g064650.3 |
| Solyc07g064700.2 | Solyc07g064700.4 |
| Solyc07g064790.2 | Solyc07g064790.4 |
| Solyc07g064840.1 | Solyc07g064840.3 |
| Solyc07g064880.2 | Solyc07g064880.4 |
| Solyc07g064970.2 | Solyc07g064970.4 |
| Solyc07g065010.2 | Solyc07g065010.4 |
| Solyc07g065050.2 | Solyc07g065050.4 |
| Solyc07g065140.2 | Solyc07g065140.4 |
| Solyc07g065160.2 | Solyc07g065160.4 |
| Solyc07g065190.2 | Solyc07g065190.4 |
| Solyc07g065210.2 | Solyc07g065210.4 |
| Solyc07g065220.1 | Solyc07g065220.3 |
| Solyc07g065240.1 | Solyc07g065240.3 |
| Solyc07g065260.2 | Solyc07g065260.4 |
| Solyc07g065300.1 | Solyc07g065300.3 |
| Solyc07g065320.2 | Solyc07g065320.4 |
| Solyc07g065370.2 | Solyc07g065370.4 |
| Solyc07g065440.2 | Solyc07g065440.4 |
| Solyc07g065470.2 | Solyc07g065470.4 |
| Solyc07g065510.2 | Solyc07g065510.4 |
| Solyc07g065560.1 | Solyc07g065560.3 |
| Solyc07g065570.1 | Solyc07g065570.3 |
| Solyc07g065580.1 | Solyc07g065580.3 |
| Solyc07g065590.2 | Solyc07g065590.4 |
| Solyc07g065620.2 | Solyc07g065620.4 |
| Solyc07g065630.2 | Solyc07g065630.4 |
| Solyc07g065660.2 | Solyc07g065660.4 |
| Solyc07g065700.2 | Solyc07g065700.4 |
| Solyc07g065760.1 | Solyc07g065760.3 |
| Solyc07g065880.2 | Solyc07g065880.4 |
| Solyc07g065920.2 | Solyc07g065920.4 |
| Solyc07g065950.2 | Solyc07g065950.4 |

| | |
|---|---|
| Solyc07g066030.2 | Solyc07g066030.4 |
| Solyc07g066050.2 | Solyc07g066050.4 |
| Solyc07g066090.2 | Solyc07g066090.4 |
| Solyc07g066170.2 | Solyc07g066170.4 |
| Solyc07g066190.1 | Solyc07g066190.3 |
| Solyc07g066230.2 | Solyc07g066230.4 |
| Solyc07g066270.2 | Solyc07g066270.4 |
| Solyc07g066300.2 | Solyc07g066300.4 |
| Solyc07g066380.2 | Solyc07g066380.4 |
| Solyc07g066410.2 | Solyc07g066410.4 |
| Solyc07g066420.2 | Solyc07g066420.4 |
| Solyc07g066440.2 | Solyc07g066440.4 |
| Solyc07g066450.2 | Solyc07g066450.4 |
| Solyc07g066510.2 | Solyc07g066510.4 |
| Solyc07g066550.2 | Solyc07g066550.4 |
| Solyc07g066570.2 | Solyc07g066570.4 |
| Solyc07g066630.2 | Solyc07g066630.4 |
| Solyc07g066650.2 | Solyc07g066650.4 |
| Solyc07g066660.2 | Solyc07g066660.4 |
| Solyc08g005000.2 | Solyc08g005000.4 |
| Solyc08g005010.2 | Solyc08g005010.4 |
| Solyc08g005030.1 | Solyc08g005030.3 |
| Solyc08g005050.2 | Solyc08g005050.4 |
| Solyc08g005070.2 | Solyc08g005070.4 |
| Solyc08g005190.2 | Solyc08g005190.4 |
| Solyc08g005220.2 | Solyc08g005220.4 |
| Solyc08g005240.1 | Solyc08g005240.3 |
| Solyc08g005300.1 | Solyc08g005300.3 |
| Solyc08g005310.2 | Solyc08g005310.4 |
| Solyc08g005400.2 | Solyc08g005400.4 |
| Solyc08g005450.1 | Solyc08g005450.3 |
| Solyc08g005470.2 | Solyc08g005470.4 |
| Solyc08g005500.2 | Solyc08g005500.4 |
| Solyc08g005510.1 | Solyc08g005510.3 |
| Solyc08g005550.2 | Solyc08g005550.4 |
| Solyc08g005570.2 | Solyc08g005570.4 |
| Solyc08g005590.2 | Solyc08g005590.4 |
| Solyc08g005600.2 | Solyc08g005600.4 |
| Solyc08g005640.2 | Solyc08g005640.4 |
| Solyc08g005680.2 | Solyc08g005680.4 |
| Solyc08g005720.2 | Solyc08g005720.4 |
| Solyc08g005750.1 | Solyc08g005750.3 |
| Solyc08g005780.2 | Solyc08g005780.4 |

| | |
|---|---|
| Solyc08g005800.2 | Solyc08g005800.4 |
| Solyc08g005880.2 | Solyc08g005880.4 |
| Solyc08g005900.2 | Solyc08g005900.4 |
| Solyc08g006020.2 | Solyc08g006020.4 |
| Solyc08g006030.2 | Solyc08g006030.4 |
| Solyc08g006070.2 | Solyc08g006070.4 |
| Solyc08g006110.2 | Solyc08g006110.4 |
| Solyc08g006130.1 | Solyc08g006130.3 |
| Solyc08g006140.1 | Solyc08g006140.3 |
| Solyc08g006160.2 | Solyc08g006160.4 |
| Solyc08g006180.2 | Solyc08g006180.4 |
| Solyc08g006190.1 | Solyc08g006190.3 |
| Solyc08g006200.1 | Solyc08g006200.3 |
| Solyc08g006210.1 | Solyc08g006210.3 |
| Solyc08g006220.2 | Solyc08g006220.4 |
| Solyc08g006230.2 | Solyc08g006230.4 |
| Solyc08g006240.1 | Solyc08g006240.3 |
| Solyc08g006290.1 | Solyc08g006290.3 |
| Solyc08g006320.2 | Solyc08g006320.4 |
| Solyc08g006360.1 | Solyc08g006360.3 |
| Solyc08g006410.2 | Solyc08g006410.4 |
| Solyc08g006430.2 | Solyc08g006430.4 |
| Solyc08g006440.2 | Solyc08g006440.4 |
| Solyc08g006470.2 | Solyc08g006470.4 |
| Solyc08g006500.2 | Solyc08g006500.4 |
| Solyc08g006530.2 | Solyc08g006530.4 |
| Solyc08g006540.2 | Solyc08g006540.4 |
| Solyc08g006550.2 | Solyc08g006550.4 |
| Solyc08g006570.2 | Solyc08g006570.4 |
| Solyc08g006600.2 | Solyc08g006600.4 |
| Solyc08g006720.2 | Solyc08g006720.4 |
| Solyc08g006750.2 | Solyc08g006750.4 |
| Solyc08g006760.2 | Solyc08g006760.4 |
| Solyc08g006790.2 | Solyc08g006790.4 |
| Solyc08g006820.2 | Solyc08g006820.4 |
| Solyc08g006850.2 | Solyc08g006850.4 |
| Solyc08g006870.2 | Solyc08g006870.4 |
| Solyc08g006910.2 | Solyc08g006910.4 |
| Solyc08g006920.1 | Solyc08g006920.3 |
| Solyc08g006940.2 | Solyc08g006940.4 |
| Solyc08g006950.2 | Solyc08g006950.4 |
| Solyc08g006970.2 | Solyc08g006970.4 |
| Solyc08g006990.1 | Solyc08g006990.3 |

| | |
|---|---|
| Solyc08g007010.2 | Solyc08g007010.4 |
| Solyc08g007060.2 | Solyc08g007060.4 |
| Solyc08g007080.2 | Solyc08g007080.4 |
| Solyc08g007090.1 | Solyc08g007090.3 |
| Solyc08g007100.2 | Solyc08g007100.4 |
| Solyc08g007110.2 | Solyc08g007110.4 |
| Solyc08g007130.2 | Solyc08g007130.4 |
| Solyc08g007140.2 | Solyc08g007140.4 |
| Solyc08g007200.2 | Solyc08g007200.4 |
| Solyc08g007210.2 | Solyc08g007210.4 |
| Solyc08g007230.1 | Solyc08g007230.3 |
| Solyc08g007240.2 | Solyc08g007240.4 |
| Solyc08g007250.1 | Solyc08g007250.3 |
| Solyc08g007270.2 | Solyc08g007270.4 |
| Solyc08g007300.1 | Solyc08g007300.3 |
| Solyc08g007380.2 | Solyc08g007380.4 |
| Solyc08g007400.2 | Solyc08g007400.4 |
| Solyc08g007410.2 | Solyc08g007410.4 |
| Solyc08g007520.2 | Solyc08g007520.4 |
| Solyc08g007530.2 | Solyc08g007530.4 |
| Solyc08g007560.1 | Solyc08g007560.3 |
| Solyc08g007570.2 | Solyc08g007570.4 |
| Solyc08g007580.2 | Solyc08g007580.4 |
| Solyc08g007750.2 | Solyc08g007750.4 |
| Solyc08g007770.2 | Solyc08g007770.4 |
| Solyc08g007840.1 | Solyc08g007840.3 |
| Solyc08g007860.2 | Solyc08g007860.4 |
| Solyc08g007900.1 | Solyc08g007900.3 |
| Solyc08g007910.2 | Solyc08g007910.4 |
| Solyc08g007940.2 | Solyc08g007940.4 |
| Solyc08g008010.2 | Solyc08g008010.4 |
| Solyc08g008120.2 | Solyc08g008120.4 |
| Solyc08g008130.2 | Solyc08g008130.4 |
| Solyc08g008160.2 | Solyc08g008160.4 |
| Solyc08g008170.2 | Solyc08g008170.4 |
| Solyc08g008190.2 | Solyc08g008190.4 |
| Solyc08g008200.1 | Solyc08g008200.3 |
| Solyc08g008250.1 | Solyc08g008250.3 |
| Solyc08g008340.2 | Solyc08g008340.4 |
| Solyc08g008380.2 | Solyc08g008380.4 |
| Solyc08g008430.2 | Solyc08g008430.4 |
| Solyc08g008450.1 | Solyc08g008450.3 |
| Solyc08g008480.2 | Solyc08g008480.4 |

| | |
|---|---|
| Solyc08g008490.2 | Solyc08g008490.4 |
| Solyc08g008550.2 | Solyc08g008550.4 |
| Solyc08g008590.2 | Solyc08g008590.4 |
| Solyc08g008620.2 | Solyc08g008620.4 |
| Solyc08g008630.2 | Solyc08g008630.4 |
| Solyc08g008660.2 | Solyc08g008660.4 |
| Solyc08g013680.2 | Solyc08g013680.4 |
| Solyc08g013710.2 | Solyc08g013710.4 |
| Solyc08g013720.2 | Solyc08g013720.4 |
| Solyc08g013740.2 | Solyc08g013740.4 |
| Solyc08g013800.2 | Solyc08g013800.4 |
| Solyc08g013880.2 | Solyc08g013880.4 |
| Solyc08g013910.2 | Solyc08g013910.4 |
| Solyc08g013930.2 | Solyc08g013930.4 |
| Solyc08g013970.1 | Solyc08g013970.3 |
| Solyc08g013980.2 | Solyc08g013980.4 |
| Solyc08g014010.2 | Solyc08g014010.4 |
| Solyc08g014060.2 | Solyc08g014060.4 |
| Solyc08g014080.2 | Solyc08g014080.4 |
| Solyc08g014150.2 | Solyc08g014150.4 |
| Solyc08g014190.2 | Solyc08g014190.4 |
| Solyc08g014230.2 | Solyc08g014230.4 |
| Solyc08g014240.2 | Solyc08g014240.4 |
| Solyc08g014300.2 | Solyc08g014300.4 |
| Solyc08g014380.1 | Solyc08g014380.3 |
| Solyc08g014420.2 | Solyc08g014420.4 |
| Solyc08g014440.2 | Solyc08g014440.4 |
| Solyc08g014450.2 | Solyc08g014450.4 |
| Solyc08g014470.2 | Solyc08g014470.4 |
| Solyc08g014480.2 | Solyc08g014480.4 |
| Solyc08g014560.1 | Solyc08g014560.3 |
| Solyc08g014580.2 | Solyc08g014580.4 |
| Solyc08g014600.2 | Solyc08g014600.4 |
| Solyc08g014610.2 | Solyc08g014610.4 |
| Solyc08g015670.2 | Solyc08g015670.4 |
| Solyc08g015690.2 | Solyc08g015690.4 |
| Solyc08g015730.1 | Solyc08g015730.3 |
| Solyc08g015750.1 | Solyc08g015750.3 |
| Solyc08g015770.2 | Solyc08g015770.4 |
| Solyc08g015840.1 | Solyc08g015840.3 |
| Solyc08g015990.2 | Solyc08g015990.4 |
| Solyc08g016050.2 | Solyc08g016050.4 |
| Solyc08g016070.1 | Solyc08g016070.3 |

| | |
|---|---|
| Solyc08g016090.2 | Solyc08g016090.4 |
| Solyc08g016190.2 | Solyc08g016190.4 |
| Solyc08g016270.1 | Solyc08g016270.3 |
| Solyc08g016560.1 | Solyc08g016560.3 |
| Solyc08g016770.2 | Solyc08g016770.4 |
| Solyc08g021870.1 | Solyc08g021870.3 |
| Solyc08g021920.2 | Solyc08g021920.4 |
| Solyc08g021940.1 | Solyc08g021940.3 |
| Solyc08g021990.1 | Solyc08g021990.3 |
| Solyc08g022080.2 | Solyc08g022080.4 |
| Solyc08g022130.1 | Solyc08g022130.3 |
| Solyc08g022210.2 | Solyc08g022210.4 |
| Solyc08g023340.2 | Solyc08g023340.4 |
| Solyc08g023360.1 | Solyc08g023360.3 |
| Solyc08g023420.1 | Solyc08g023420.3 |
| Solyc08g023460.2 | Solyc08g023460.4 |
| Solyc08g023490.2 | Solyc08g023490.4 |
| Solyc08g023500.2 | Solyc08g023500.4 |
| Solyc08g023510.1 | Solyc08g023510.3 |
| Solyc08g023580.2 | Solyc08g023580.4 |
| Solyc08g023590.2 | Solyc08g023590.4 |
| Solyc08g028690.2 | Solyc08g028690.4 |
| Solyc08g028880.1 | Solyc08g028880.3 |
| Solyc08g028950.1 | Solyc08g028950.3 |
| Solyc08g029050.2 | Solyc08g029050.4 |
| Solyc08g029090.2 | Solyc08g029090.4 |
| Solyc08g029150.1 | Solyc08g029150.3 |
| Solyc08g029220.1 | Solyc08g029220.3 |
| Solyc08g029230.2 | Solyc08g029230.4 |
| Solyc08g029300.1 | Solyc08g029300.3 |
| Solyc08g029360.2 | Solyc08g029360.4 |
| Solyc08g029400.1 | Solyc08g029400.3 |
| Solyc08g036600.1 | Solyc08g036600.3 |
| Solyc08g036610.1 | Solyc08g036610.3 |
| Solyc08g036620.2 | Solyc08g036620.4 |
| Solyc08g036660.2 | Solyc08g036660.4 |
| Solyc08g041750.1 | Solyc08g041750.3 |
| Solyc08g041800.2 | Solyc08g041800.4 |
| Solyc08g041850.2 | Solyc08g041850.4 |
| Solyc08g041890.2 | Solyc08g041890.4 |
| Solyc08g042010.2 | Solyc08g042010.4 |
| Solyc08g042030.1 | Solyc08g042030.3 |
| Solyc08g042040.2 | Solyc08g042040.4 |

| | |
|---|---|
| Solyc08g043140.2 | Solyc08g043140.4 |
| Solyc08g043170.2 | Solyc08g043170.4 |
| Solyc08g044260.2 | Solyc08g044260.4 |
| Solyc08g044270.2 | Solyc08g044270.4 |
| Solyc08g044380.1 | Solyc08g044380.3 |
| Solyc08g044400.1 | Solyc08g044400.3 |
| Solyc08g044510.2 | Solyc08g044510.4 |
| Solyc08g044610.1 | Solyc08g044610.3 |
| Solyc08g045740.1 | Solyc08g045740.3 |
| Solyc08g045780.2 | Solyc08g045780.4 |
| Solyc08g048240.2 | Solyc08g048240.4 |
| Solyc08g048250.2 | Solyc08g048250.4 |
| Solyc08g048290.2 | Solyc08g048290.4 |
| Solyc08g048300.1 | Solyc08g048300.3 |
| Solyc08g048310.1 | Solyc08g048310.3 |
| Solyc08g048370.2 | Solyc08g048370.4 |
| Solyc08g048390.1 | Solyc08g048390.3 |
| Solyc08g048430.2 | Solyc08g048430.4 |
| Solyc08g048450.2 | Solyc08g048450.4 |
| Solyc08g054570.1 | Solyc08g054570.3 |
| Solyc08g059660.1 | Solyc08g059660.3 |
| Solyc08g059710.2 | Solyc08g059710.4 |
| Solyc08g059750.1 | Solyc08g059750.3 |
| Solyc08g060830.2 | Solyc08g060830.4 |
| Solyc08g060900.1 | Solyc08g060900.3 |
| Solyc08g060920.2 | Solyc08g060920.4 |
| Solyc08g060950.1 | Solyc08g060950.3 |
| Solyc08g060970.2 | Solyc08g060970.4 |
| Solyc08g061000.2 | Solyc08g061000.4 |
| Solyc08g061010.2 | Solyc08g061010.4 |
| Solyc08g061060.1 | Solyc08g061060.3 |
| Solyc08g061240.1 | Solyc08g061240.3 |
| Solyc08g061260.2 | Solyc08g061260.4 |
| Solyc08g061390.1 | Solyc08g061390.3 |
| Solyc08g061410.1 | Solyc08g061410.3 |
| Solyc08g061500.1 | Solyc08g061500.3 |
| Solyc08g061570.1 | Solyc08g061570.3 |
| Solyc08g061590.2 | Solyc08g061590.4 |
| Solyc08g061610.2 | Solyc08g061610.4 |
| Solyc08g061650.2 | Solyc08g061650.4 |
| Solyc08g061680.1 | Solyc08g061680.3 |
| Solyc08g061690.1 | Solyc08g061690.3 |
| Solyc08g061710.1 | Solyc08g061710.3 |

| | |
|---|---|
| Solyc08g061730.1 | Solyc08g061730.3 |
| Solyc08g061740.1 | Solyc08g061740.3 |
| Solyc08g061780.1 | Solyc08g061780.3 |
| Solyc08g061790.1 | Solyc08g061790.3 |
| Solyc08g061890.2 | Solyc08g061890.4 |
| Solyc08g061910.2 | Solyc08g061910.4 |
| Solyc08g061920.1 | Solyc08g061920.3 |
| Solyc08g061930.2 | Solyc08g061930.4 |
| Solyc08g061980.2 | Solyc08g061980.4 |
| Solyc08g062000.1 | Solyc08g062000.3 |
| Solyc08g062100.1 | Solyc08g062100.3 |
| Solyc08g062110.1 | Solyc08g062110.3 |
| Solyc08g062180.1 | Solyc08g062180.3 |
| Solyc08g062200.1 | Solyc08g062200.3 |
| Solyc08g062250.1 | Solyc08g062250.3 |
| Solyc08g062290.2 | Solyc08g062290.4 |
| Solyc08g062530.1 | Solyc08g062530.4 |
| Solyc08g062560.2 | Solyc08g062560.4 |
| Solyc08g062600.1 | Solyc08g062600.3 |
| Solyc08g062610.2 | Solyc08g062610.4 |
| Solyc08g062900.1 | Solyc08g062900.3 |
| Solyc08g062910.2 | Solyc08g062910.4 |
| Solyc08g062930.1 | Solyc08g062930.3 |
| Solyc08g062940.2 | Solyc08g062940.4 |
| Solyc08g062950.2 | Solyc08g062950.4 |
| Solyc08g062960.2 | Solyc08g062960.4 |
| Solyc08g063000.2 | Solyc08g063000.4 |
| Solyc08g063040.2 | Solyc08g063040.4 |
| Solyc08g065130.1 | Solyc08g065130.3 |
| Solyc08g065350.2 | Solyc08g065350.4 |
| Solyc08g065410.1 | Solyc08g065410.3 |
| Solyc08g065470.2 | Solyc08g065470.4 |
| Solyc08g065530.1 | Solyc08g065530.3 |
| Solyc08g065590.1 | Solyc08g065590.3 |
| Solyc08g065630.1 | Solyc08g065630.3 |
| Solyc08g065670.2 | Solyc08g065670.4 |
| Solyc08g065680.1 | Solyc08g065680.3 |
| Solyc08g065700.1 | Solyc08g065700.3 |
| Solyc08g065740.1 | Solyc08g065740.3 |
| Solyc08g065820.1 | Solyc08g065820.3 |
| Solyc08g065830.2 | Solyc08g065830.4 |
| Solyc08g065870.2 | Solyc08g065870.4 |
| Solyc08g065900.2 | Solyc08g065900.4 |

| | |
|---|---|
| Solyc08g065940.2 | Solyc08g065940.4 |
| Solyc08g066020.1 | Solyc08g066020.3 |
| Solyc08g066120.2 | Solyc08g066120.4 |
| Solyc08g066220.2 | Solyc08g066220.4 |
| Solyc08g066250.2 | Solyc08g066250.4 |
| Solyc08g066270.1 | Solyc08g066270.3 |
| Solyc08g066300.1 | Solyc08g066300.3 |
| Solyc08g066310.1 | Solyc08g066310.3 |
| Solyc08g066320.2 | Solyc08g066320.4 |
| Solyc08g066330.1 | Solyc08g066330.3 |
| Solyc08g066400.1 | Solyc08g066400.3 |
| Solyc08g066460.1 | Solyc08g066460.3 |
| Solyc08g066480.1 | Solyc08g066480.3 |
| Solyc08g066500.2 | Solyc08g066500.4 |
| Solyc08g066530.2 | Solyc08g066530.4 |
| Solyc08g066570.1 | Solyc08g066570.3 |
| Solyc08g066580.2 | Solyc08g066580.4 |
| Solyc08g066600.2 | Solyc08g066600.4 |
| Solyc08g066690.2 | Solyc08g066690.4 |
| Solyc08g066720.1 | Solyc08g066720.3 |
| Solyc08g066760.2 | Solyc08g066760.4 |
| Solyc08g066790.2 | Solyc08g066790.4 |
| Solyc08g066800.2 | Solyc08g066800.4 |
| Solyc08g066870.2 | Solyc08g066870.4 |
| Solyc08g066890.2 | Solyc08g066890.4 |
| Solyc08g066940.2 | Solyc08g066940.4 |
| Solyc08g066980.2 | Solyc08g066980.4 |
| Solyc08g066990.2 | Solyc08g066990.4 |
| Solyc08g067020.2 | Solyc08g067020.4 |
| Solyc08g067040.2 | Solyc08g067040.4 |
| Solyc08g067080.1 | Solyc08g067080.3 |
| Solyc08g067090.2 | Solyc08g067090.4 |
| Solyc08g067150.2 | Solyc08g067150.4 |
| Solyc08g067170.1 | Solyc08g067170.3 |
| Solyc08g067180.2 | Solyc08g067180.4 |
| Solyc08g067190.2 | Solyc08g067190.4 |
| Solyc08g067230.2 | Solyc08g067230.4 |
| Solyc08g067240.2 | Solyc08g067240.4 |
| Solyc08g067250.2 | Solyc08g067250.4 |
| Solyc08g067260.2 | Solyc08g067260.4 |
| Solyc08g067320.1 | Solyc08g067320.3 |
| Solyc08g067340.2 | Solyc08g067340.4 |
| Solyc08g067390.2 | Solyc08g067390.4 |

| | |
|---|---|
| Solyc08g067420.2 | Solyc08g067420.4 |
| Solyc08g067480.2 | Solyc08g067480.4 |
| Solyc08g067580.1 | Solyc08g067580.3 |
| Solyc08g067740.2 | Solyc08g067740.4 |
| Solyc08g067760.2 | Solyc08g067760.4 |
| Solyc08g067790.2 | Solyc08g067790.4 |
| Solyc08g067800.1 | Solyc08g067800.3 |
| Solyc08g067930.2 | Solyc08g067930.4 |
| Solyc08g067990.1 | Solyc08g067990.3 |
| Solyc08g068000.1 | Solyc08g068000.3 |
| Solyc08g068120.1 | Solyc08g068120.3 |
| Solyc08g068140.2 | Solyc08g068140.4 |
| Solyc08g068150.2 | Solyc08g068150.4 |
| Solyc08g068200.2 | Solyc08g068200.4 |
| Solyc08g068230.2 | Solyc08g068230.4 |
| Solyc08g068240.1 | Solyc08g068240.3 |
| Solyc08g068250.2 | Solyc08g068250.4 |
| Solyc08g068400.2 | Solyc08g068400.4 |
| Solyc08g068410.1 | Solyc08g068410.3 |
| Solyc08g068430.2 | Solyc08g068430.4 |
| Solyc08g068440.2 | Solyc08g068440.4 |
| Solyc08g068530.2 | Solyc08g068530.4 |
| Solyc08g068540.2 | Solyc08g068540.4 |
| Solyc08g068570.2 | Solyc08g068570.4 |
| Solyc08g068580.2 | Solyc08g068580.4 |
| Solyc08g068590.2 | Solyc08g068590.4 |
| Solyc08g068620.1 | Solyc08g068620.3 |
| Solyc08g068630.2 | Solyc08g068630.4 |
| Solyc08g068680.2 | Solyc08g068680.4 |
| Solyc08g068770.1 | Solyc08g068770.3 |
| Solyc08g068780.1 | Solyc08g068780.3 |
| Solyc08g068790.1 | Solyc08g068790.3 |
| Solyc08g068800.2 | Solyc08g068800.4 |
| Solyc08g068920.2 | Solyc08g068920.4 |
| Solyc08g069000.2 | Solyc08g069000.4 |
| Solyc08g069010.2 | Solyc08g069010.4 |
| Solyc08g069030.2 | Solyc08g069030.4 |
| Solyc08g069060.2 | Solyc08g069060.4 |
| Solyc08g069070.1 | Solyc08g069070.3 |
| Solyc08g069080.2 | Solyc08g069080.4 |
| Solyc08g069120.2 | Solyc08g069120.4 |
| Solyc08g069140.2 | Solyc08g069140.4 |
| Solyc08g069170.1 | Solyc08g069170.3 |

| | |
|---|---|
| Solyc08g074250.1 | Solyc08g074250.3 |
| Solyc08g074270.2 | Solyc08g074270.4 |
| Solyc08g074280.2 | Solyc08g074280.4 |
| Solyc08g074330.1 | Solyc08g074330.3 |
| Solyc08g074340.1 | Solyc08g074340.3 |
| Solyc08g074350.1 | Solyc08g074350.3 |
| Solyc08g074370.2 | Solyc08g074370.4 |
| Solyc08g074410.2 | Solyc08g074410.4 |
| Solyc08g074440.2 | Solyc08g074440.4 |
| Solyc08g074460.1 | Solyc08g074460.3 |
| Solyc08g074490.2 | Solyc08g074490.4 |
| Solyc08g074600.2 | Solyc08g074600.4 |
| Solyc08g074650.2 | Solyc08g074650.4 |
| Solyc08g074770.1 | Solyc08g074770.3 |
| Solyc08g074830.2 | Solyc08g074830.4 |
| Solyc08g074850.1 | Solyc08g074850.3 |
| Solyc08g074870.1 | Solyc08g074870.3 |
| Solyc08g074890.2 | Solyc08g074890.4 |
| Solyc08g074980.2 | Solyc08g074980.4 |
| Solyc08g074990.1 | Solyc08g074990.3 |
| Solyc08g075010.2 | Solyc08g075010.4 |
| Solyc08g075050.1 | Solyc08g075050.3 |
| Solyc08g075060.2 | Solyc08g075060.4 |
| Solyc08g075090.2 | Solyc08g075090.4 |
| Solyc08g075100.2 | Solyc08g075100.4 |
| Solyc08g075110.1 | Solyc08g075110.3 |
| Solyc08g075280.2 | Solyc08g075280.4 |
| Solyc08g075300.2 | Solyc08g075300.4 |
| Solyc08g075310.2 | Solyc08g075310.4 |
| Solyc08g075320.2 | Solyc08g075320.4 |
| Solyc08g075330.1 | Solyc08g075330.3 |
| Solyc08g075350.1 | Solyc08g075350.3 |
| Solyc08g075360.1 | Solyc08g075360.3 |
| Solyc08g075390.2 | Solyc08g075390.4 |
| Solyc08g075440.2 | Solyc08g075440.4 |
| Solyc08g075480.2 | Solyc08g075480.4 |
| Solyc08g075490.2 | Solyc08g075490.4 |
| Solyc08g075500.2 | Solyc08g075500.4 |
| Solyc08g075510.2 | Solyc08g075510.4 |
| Solyc08g075570.2 | Solyc08g075570.4 |
| Solyc08g075580.1 | Solyc08g075580.3 |
| Solyc08g075620.1 | Solyc08g075620.3 |
| Solyc08g075640.2 | Solyc08g075640.4 |

| | |
|---|---|
| Solyc08g075720.1 | Solyc08g075720.3 |
| Solyc08g075750.2 | Solyc08g075750.4 |
| Solyc08g075780.2 | Solyc08g075780.4 |
| Solyc08g075790.2 | Solyc08g075790.4 |
| Solyc08g075820.2 | Solyc08g075820.4 |
| Solyc08g075830.2 | Solyc08g075830.4 |
| Solyc08g075880.2 | Solyc08g075880.4 |
| Solyc08g075920.2 | Solyc08g075920.4 |
| Solyc08g075930.2 | Solyc08g075930.4 |
| Solyc08g075940.2 | Solyc08g075940.4 |
| Solyc08g075950.1 | Solyc08g075950.3 |
| Solyc08g076000.2 | Solyc08g076000.4 |
| Solyc08g076010.2 | Solyc08g076010.4 |
| Solyc08g076020.2 | Solyc08g076020.4 |
| Solyc08g076050.2 | Solyc08g076050.4 |
| Solyc08g076100.2 | Solyc08g076100.4 |
| Solyc08g076150.2 | Solyc08g076150.4 |
| Solyc08g076200.2 | Solyc08g076200.4 |
| Solyc08g076250.1 | Solyc08g076250.3 |
| Solyc08g076360.2 | Solyc08g076360.4 |
| Solyc08g076380.1 | Solyc08g076380.3 |
| Solyc08g076410.2 | Solyc08g076410.4 |
| Solyc08g076440.2 | Solyc08g076440.4 |
| Solyc08g076470.2 | Solyc08g076470.4 |
| Solyc08g076550.1 | Solyc08g076550.3 |
| Solyc08g076610.2 | Solyc08g076610.4 |
| Solyc08g076630.1 | Solyc08g076630.3 |
| Solyc08g076680.1 | Solyc08g076680.3 |
| Solyc08g076690.1 | Solyc08g076690.3 |
| Solyc08g076720.2 | Solyc08g076720.4 |
| Solyc08g076730.2 | Solyc08g076730.4 |
| Solyc08g076800.2 | Solyc08g076800.4 |
| Solyc08g076810.2 | Solyc08g076810.4 |
| Solyc08g076980.2 | Solyc08g076980.4 |
| Solyc08g077030.2 | Solyc08g077030.4 |
| Solyc08g077110.2 | Solyc08g077110.4 |
| Solyc08g077200.2 | Solyc08g077200.4 |
| Solyc08g077210.2 | Solyc08g077210.4 |
| Solyc08g077270.2 | Solyc08g077270.4 |
| Solyc08g077320.1 | Solyc08g077320.3 |
| Solyc08g077370.2 | Solyc08g077370.4 |
| Solyc08g077380.2 | Solyc08g077380.4 |
| Solyc08g077400.1 | Solyc08g077400.3 |

| | |
|---|---|
| Solyc08g077410.2 | Solyc08g077410.4 |
| Solyc08g077470.2 | Solyc08g077470.4 |
| Solyc08g077480.2 | Solyc08g077480.4 |
| Solyc08g077500.2 | Solyc08g077500.4 |
| Solyc08g077510.2 | Solyc08g077510.4 |
| Solyc08g077560.2 | Solyc08g077560.4 |
| Solyc08g077570.1 | Solyc08g077570.3 |
| Solyc08g077610.1 | Solyc08g077610.3 |
| Solyc08g077640.1 | Solyc08g077640.3 |
| Solyc08g077670.1 | Solyc08g077670.3 |
| Solyc08g077750.2 | Solyc08g077750.4 |
| Solyc08g077800.2 | Solyc08g077800.4 |
| Solyc08g077820.2 | Solyc08g077820.4 |
| Solyc08g077860.2 | Solyc08g077860.4 |
| Solyc08g077870.2 | Solyc08g077870.4 |
| Solyc08g077980.2 | Solyc08g077980.4 |
| Solyc08g078030.2 | Solyc08g078030.4 |
| Solyc08g078040.2 | Solyc08g078040.4 |
| Solyc08g078070.2 | Solyc08g078070.4 |
| Solyc08g078130.2 | Solyc08g078130.4 |
| Solyc08g078230.2 | Solyc08g078230.4 |
| Solyc08g078240.2 | Solyc08g078240.4 |
| Solyc08g078250.2 | Solyc08g078250.4 |
| Solyc08g078300.2 | Solyc08g078300.4 |
| Solyc08g078330.2 | Solyc08g078330.4 |
| Solyc08g078370.2 | Solyc08g078370.4 |
| Solyc08g078430.2 | Solyc08g078430.4 |
| Solyc08g078470.2 | Solyc08g078470.4 |
| Solyc08g078490.2 | Solyc08g078490.4 |
| Solyc08g078530.2 | Solyc08g078530.4 |
| Solyc08g078570.1 | Solyc08g078570.3 |
| Solyc08g078580.1 | Solyc08g078580.3 |
| Solyc08g078640.1 | Solyc08g078640.3 |
| Solyc08g078680.2 | Solyc08g078680.4 |
| Solyc08g078710.1 | Solyc08g078710.3 |
| Solyc08g078750.2 | Solyc08g078750.4 |
| Solyc08g078860.2 | Solyc08g078860.4 |
| Solyc08g078870.1 | Solyc08g078870.3 |
| Solyc08g078890.1 | Solyc08g078890.3 |
| Solyc08g078960.2 | Solyc08g078960.4 |
| Solyc08g078990.1 | Solyc08g078990.3 |
| Solyc08g079000.2 | Solyc08g079000.4 |
| Solyc08g079060.2 | Solyc08g079060.4 |

| | |
|---|---|
| Solyc08g079080.2 | Solyc08g079080.5 |
| Solyc08g079090.2 | Solyc08g079090.4 |
| Solyc08g079110.2 | Solyc08g079110.4 |
| Solyc08g079120.1 | Solyc08g079120.3 |
| Solyc08g079250.2 | Solyc08g079250.4 |
| Solyc08g079310.2 | Solyc08g079310.4 |
| Solyc08g079350.1 | Solyc08g079350.3 |
| Solyc08g079490.2 | Solyc08g079490.4 |
| Solyc08g079540.2 | Solyc08g079540.4 |
| Solyc08g079640.1 | Solyc08g079640.3 |
| Solyc08g079700.1 | Solyc08g079700.3 |
| Solyc08g079760.2 | Solyc08g079760.4 |
| Solyc08g079800.2 | Solyc08g079800.4 |
| Solyc08g079830.2 | Solyc08g079830.4 |
| Solyc08g079990.1 | Solyc08g079990.3 |
| Solyc08g080030.2 | Solyc08g080030.4 |
| Solyc08g080040.2 | Solyc08g080040.4 |
| Solyc08g080050.2 | Solyc08g080050.4 |
| Solyc08g080060.2 | Solyc08g080060.4 |
| Solyc08g080080.2 | Solyc08g080080.4 |
| Solyc08g080100.2 | Solyc08g080100.4 |
| Solyc08g080140.2 | Solyc08g080140.4 |
| Solyc08g080160.2 | Solyc08g080160.4 |
| Solyc08g080190.2 | Solyc08g080190.4 |
| Solyc08g080200.2 | Solyc08g080200.4 |
| Solyc08g080220.1 | Solyc08g080220.3 |
| Solyc08g080250.2 | Solyc08g080250.4 |
| Solyc08g080260.2 | Solyc08g080260.4 |
| Solyc08g080290.2 | Solyc08g080290.4 |
| Solyc08g080380.2 | Solyc08g080380.4 |
| Solyc08g080420.2 | Solyc08g080420.4 |
| Solyc08g080450.1 | Solyc08g080450.3 |
| Solyc08g080510.2 | Solyc08g080510.4 |
| Solyc08g080520.2 | Solyc08g080520.4 |
| Solyc08g080560.1 | Solyc08g080560.3 |
| Solyc08g080570.2 | Solyc08g080570.4 |
| Solyc08g080580.2 | Solyc08g080580.4 |
| Solyc08g080630.2 | Solyc08g080630.4 |
| Solyc08g080680.2 | Solyc08g080680.4 |
| Solyc08g080690.2 | Solyc08g080690.4 |
| Solyc08g080700.1 | Solyc08g080700.3 |
| Solyc08g080710.1 | Solyc08g080710.3 |
| Solyc08g080750.2 | Solyc08g080750.4 |

| | |
|---|---|
| Solyc08g080780.2 | Solyc08g080780.4 |
| Solyc08g080810.1 | Solyc08g080810.3 |
| Solyc08g080850.2 | Solyc08g080850.4 |
| Solyc08g080960.2 | Solyc08g080960.4 |
| Solyc08g080970.1 | Solyc08g080970.3 |
| Solyc08g080980.2 | Solyc08g080980.4 |
| Solyc08g081050.2 | Solyc08g081050.4 |
| Solyc08g081100.2 | Solyc08g081100.4 |
| Solyc08g081120.2 | Solyc08g081120.4 |
| Solyc08g081140.2 | Solyc08g081140.4 |
| Solyc08g081150.2 | Solyc08g081150.4 |
| Solyc08g081170.2 | Solyc08g081170.4 |
| Solyc08g081210.2 | Solyc08g081210.4 |
| Solyc08g081250.2 | Solyc08g081250.4 |
| Solyc08g081290.2 | Solyc08g081290.4 |
| Solyc08g081300.2 | Solyc08g081300.4 |
| Solyc08g081310.2 | Solyc08g081310.4 |
| Solyc08g081330.2 | Solyc08g081330.4 |
| Solyc08g081340.2 | Solyc08g081340.4 |
| Solyc08g081390.2 | Solyc08g081390.4 |
| Solyc08g081400.2 | Solyc08g081400.4 |
| Solyc08g081420.2 | Solyc08g081420.4 |
| Solyc08g081450.2 | Solyc08g081450.4 |
| Solyc08g081490.2 | Solyc08g081490.4 |
| Solyc08g081500.1 | Solyc08g081500.3 |
| Solyc08g081550.2 | Solyc08g081550.4 |
| Solyc08g081580.2 | Solyc08g081580.4 |
| Solyc08g081590.1 | Solyc08g081590.3 |
| Solyc08g081610.2 | Solyc08g081610.4 |
| Solyc08g081620.2 | Solyc08g081620.4 |
| Solyc08g081650.1 | Solyc08g081650.3 |
| Solyc08g081670.1 | Solyc08g081670.3 |
| Solyc08g081720.2 | Solyc08g081720.4 |
| Solyc08g081730.2 | Solyc08g081730.4 |
| Solyc08g081790.1 | Solyc08g081790.3 |
| Solyc08g081820.2 | Solyc08g081820.4 |
| Solyc08g081880.1 | Solyc08g081880.3 |
| Solyc08g081890.2 | Solyc08g081890.4 |
| Solyc08g081900.2 | Solyc08g081900.4 |
| Solyc08g081920.2 | Solyc08g081920.4 |
| Solyc08g081960.1 | Solyc08g081960.3 |
| Solyc08g081980.2 | Solyc08g081980.4 |
| Solyc08g082040.2 | Solyc08g082040.4 |

| | |
|---|---|
| Solyc08g082050.1 | Solyc08g082050.3 |
| Solyc08g082110.2 | Solyc08g082110.4 |
| Solyc08g082170.2 | Solyc08g082170.4 |
| Solyc08g082210.2 | Solyc08g082210.4 |
| Solyc08g082280.2 | Solyc08g082280.4 |
| Solyc08g082290.2 | Solyc08g082290.4 |
| Solyc08g082350.2 | Solyc08g082350.4 |
| Solyc08g082430.2 | Solyc08g082430.4 |
| Solyc08g082460.2 | Solyc08g082460.4 |
| Solyc08g082490.1 | Solyc08g082490.3 |
| Solyc08g082530.2 | Solyc08g082530.4 |
| Solyc08g082610.2 | Solyc08g082610.4 |
| Solyc08g082670.2 | Solyc08g082670.4 |
| Solyc08g082730.2 | Solyc08g082730.4 |
| Solyc08g082740.2 | Solyc08g082740.4 |
| Solyc08g082810.2 | Solyc08g082810.4 |
| Solyc08g082820.2 | Solyc08g082820.4 |
| Solyc08g082840.1 | Solyc08g082840.3 |
| Solyc08g082860.2 | Solyc08g082860.4 |
| Solyc08g082880.2 | Solyc08g082880.4 |
| Solyc08g082900.2 | Solyc08g082900.4 |
| Solyc08g082950.2 | Solyc08g082950.4 |
| Solyc08g082970.2 | Solyc08g082970.4 |
| Solyc08g083040.2 | Solyc08g083040.4 |
| Solyc08g083100.1 | Solyc08g083100.3 |
| Solyc08g083110.2 | Solyc08g083110.4 |
| Solyc08g083130.2 | Solyc08g083130.4 |
| Solyc08g083140.2 | Solyc08g083140.4 |
| Solyc08g083230.1 | Solyc08g083230.3 |
| Solyc08g083250.2 | Solyc08g083250.4 |
| Solyc08g083270.1 | Solyc08g083270.3 |
| Solyc08g083290.2 | Solyc08g083290.4 |
| Solyc08g083300.2 | Solyc08g083300.4 |
| Solyc08g083320.2 | Solyc08g083320.4 |
| Solyc08g083350.2 | Solyc08g083350.4 |
| Solyc08g083370.2 | Solyc08g083370.4 |
| Solyc08g083510.1 | Solyc08g083510.3 |
| Solyc09g005010.2 | Solyc09g005010.4 |
| Solyc09g005030.1 | Solyc09g005030.3 |
| Solyc09g005060.1 | Solyc09g005060.3 |
| Solyc09g005100.2 | Solyc09g005100.4 |
| Solyc09g005130.2 | Solyc09g005130.4 |
| Solyc09g005170.1 | Solyc09g005170.3 |

| | |
|---|---|
| Solyc09g005180.2 | Solyc09g005180.4 |
| Solyc09g005190.1 | Solyc09g005190.3 |
| Solyc09g005210.2 | Solyc09g005210.4 |
| Solyc09g005220.1 | Solyc09g005220.3 |
| Solyc09g005240.1 | Solyc09g005240.3 |
| Solyc09g005260.2 | Solyc09g005260.4 |
| Solyc09g005410.1 | Solyc09g005410.3 |
| Solyc09g005420.2 | Solyc09g005420.4 |
| Solyc09g005430.1 | Solyc09g005430.3 |
| Solyc09g005540.1 | Solyc09g005540.3 |
| Solyc09g005560.2 | Solyc09g005560.4 |
| Solyc09g005580.2 | Solyc09g005580.4 |
| Solyc09g005610.2 | Solyc09g005610.4 |
| Solyc09g005620.2 | Solyc09g005620.4 |
| Solyc09g005660.2 | Solyc09g005660.4 |
| Solyc09g005670.1 | Solyc09g005670.3 |
| Solyc09g005710.1 | Solyc09g005710.3 |
| Solyc09g005730.2 | Solyc09g005730.4 |
| Solyc09g005740.1 | Solyc09g005740.3 |
| Solyc09g005780.1 | Solyc09g005780.3 |
| Solyc09g005810.2 | Solyc09g005810.4 |
| Solyc09g005910.2 | Solyc09g005910.4 |
| Solyc09g005920.2 | Solyc09g005920.4 |
| Solyc09g005940.2 | Solyc09g005940.4 |
| Solyc09g005960.2 | Solyc09g005960.4 |
| Solyc09g005980.2 | Solyc09g005980.4 |
| Solyc09g005990.2 | Solyc09g005990.4 |
| Solyc09g007090.2 | Solyc09g007090.4 |
| Solyc09g007120.2 | Solyc09g007120.4 |
| Solyc09g007140.2 | Solyc09g007140.4 |
| Solyc09g007160.2 | Solyc09g007160.4 |
| Solyc09g007170.2 | Solyc09g007170.4 |
| Solyc09g007200.2 | Solyc09g007200.4 |
| Solyc09g007210.2 | Solyc09g007210.4 |
| Solyc09g007250.2 | Solyc09g007250.4 |
| Solyc09g007260.2 | Solyc09g007260.4 |
| Solyc09g007280.2 | Solyc09g007280.4 |
| Solyc09g007290.2 | Solyc09g007290.4 |
| Solyc09g007300.1 | Solyc09g007300.3 |
| Solyc09g007320.2 | Solyc09g007320.4 |
| Solyc09g007330.2 | Solyc09g007330.4 |
| Solyc09g007340.2 | Solyc09g007340.4 |
| Solyc09g007370.2 | Solyc09g007370.4 |

| | |
|---|---|
| Solyc09g007420.2 | Solyc09g007420.4 |
| Solyc09g007510.1 | Solyc09g007510.3 |
| Solyc09g007560.2 | Solyc09g007560.4 |
| Solyc09g007570.1 | Solyc09g007570.3 |
| Solyc09g007580.1 | Solyc09g007580.3 |
| Solyc09g007610.1 | Solyc09g007610.3 |
| Solyc09g007640.2 | Solyc09g007640.4 |
| Solyc09g007650.1 | Solyc09g007650.3 |
| Solyc09g007680.2 | Solyc09g007680.4 |
| Solyc09g007720.2 | Solyc09g007720.4 |
| Solyc09g007730.2 | Solyc09g007730.4 |
| Solyc09g007750.2 | Solyc09g007750.4 |
| Solyc09g007760.2 | Solyc09g007760.4 |
| Solyc09g007810.2 | Solyc09g007810.4 |
| Solyc09g007860.2 | Solyc09g007860.4 |
| Solyc09g007900.2 | Solyc09g007900.4 |
| Solyc09g007910.2 | Solyc09g007910.4 |
| Solyc09g007920.2 | Solyc09g007920.4 |
| Solyc09g007970.1 | Solyc09g007970.3 |
| Solyc09g008020.2 | Solyc09g008020.4 |
| Solyc09g008060.2 | Solyc09g008060.4 |
| Solyc09g008070.2 | Solyc09g008070.4 |
| Solyc09g008100.2 | Solyc09g008100.4 |
| Solyc09g008120.2 | Solyc09g008120.4 |
| Solyc09g008150.1 | Solyc09g008150.3 |
| Solyc09g008170.1 | Solyc09g008170.3 |
| Solyc09g008200.2 | Solyc09g008200.4 |
| Solyc09g008250.2 | Solyc09g008250.4 |
| Solyc09g008290.2 | Solyc09g008290.4 |
| Solyc09g008320.2 | Solyc09g008320.4 |
| Solyc09g008390.1 | Solyc09g008390.3 |
| Solyc09g008430.2 | Solyc09g008430.4 |
| Solyc09g008470.2 | Solyc09g008470.4 |
| Solyc09g008480.2 | Solyc09g008480.4 |
| Solyc09g008490.2 | Solyc09g008490.4 |
| Solyc09g008500.1 | Solyc09g008500.3 |
| Solyc09g008520.2 | Solyc09g008520.4 |
| Solyc09g008530.1 | Solyc09g008530.3 |
| Solyc09g008550.2 | Solyc09g008550.4 |
| Solyc09g008560.2 | Solyc09g008560.4 |
| Solyc09g008590.2 | Solyc09g008590.4 |
| Solyc09g008610.2 | Solyc09g008610.4 |
| Solyc09g008620.1 | Solyc09g008620.3 |

| | |
|---|---|
| Solyc09g008690.1 | Solyc09g008690.3 |
| Solyc09g008720.1 | Solyc09g008720.3 |
| Solyc09g008780.2 | Solyc09g008780.4 |
| Solyc09g008790.2 | Solyc09g008790.4 |
| Solyc09g008840.2 | Solyc09g008840.4 |
| Solyc09g008860.2 | Solyc09g008860.4 |
| Solyc09g008930.2 | Solyc09g008930.4 |
| Solyc09g008940.2 | Solyc09g008940.4 |
| Solyc09g009000.2 | Solyc09g009000.4 |
| Solyc09g009010.2 | Solyc09g009010.4 |
| Solyc09g009030.2 | Solyc09g009030.4 |
| Solyc09g009080.2 | Solyc09g009080.4 |
| Solyc09g009100.2 | Solyc09g009100.4 |
| Solyc09g009120.2 | Solyc09g009120.4 |
| Solyc09g009160.1 | Solyc09g009160.3 |
| Solyc09g009170.2 | Solyc09g009170.4 |
| Solyc09g009190.2 | Solyc09g009190.4 |
| Solyc09g009200.1 | Solyc09g009200.3 |
| Solyc09g009210.2 | Solyc09g009210.4 |
| Solyc09g009300.2 | Solyc09g009300.4 |
| Solyc09g009340.1 | Solyc09g009340.3 |
| Solyc09g009370.2 | Solyc09g009370.4 |
| Solyc09g009400.2 | Solyc09g009400.4 |
| Solyc09g009440.1 | Solyc09g009440.3 |
| Solyc09g009460.2 | Solyc09g009460.4 |
| Solyc09g009490.2 | Solyc09g009490.4 |
| Solyc09g009510.2 | Solyc09g009510.4 |
| Solyc09g009530.2 | Solyc09g009530.4 |
| Solyc09g009540.2 | Solyc09g009540.4 |
| Solyc09g009550.2 | Solyc09g009550.4 |
| Solyc09g009570.1 | Solyc09g009570.3 |
| Solyc09g009580.2 | Solyc09g009580.4 |
| Solyc09g009600.1 | Solyc09g009600.3 |
| Solyc09g009610.1 | Solyc09g009610.3 |
| Solyc09g009700.2 | Solyc09g009700.4 |
| Solyc09g009750.1 | Solyc09g009750.3 |
| Solyc09g009780.2 | Solyc09g009780.4 |
| Solyc09g009790.2 | Solyc09g009790.4 |
| Solyc09g009820.2 | Solyc09g009820.4 |
| Solyc09g009850.2 | Solyc09g009850.4 |
| Solyc09g009890.2 | Solyc09g009890.4 |
| Solyc09g009900.2 | Solyc09g009900.4 |
| Solyc09g009910.2 | Solyc09g009910.4 |

| | |
|---|---|
| Solyc09g009950.2 | Solyc09g009950.4 |
| Solyc09g009980.1 | Solyc09g009980.3 |
| Solyc09g010000.2 | Solyc09g010000.4 |
| Solyc09g010030.1 | Solyc09g010030.3 |
| Solyc09g010040.1 | Solyc09g010040.3 |
| Solyc09g010130.2 | Solyc09g010130.4 |
| Solyc09g010140.1 | Solyc09g010140.3 |
| Solyc09g010150.2 | Solyc09g010150.4 |
| Solyc09g010160.1 | Solyc09g010160.3 |
| Solyc09g010200.2 | Solyc09g010200.4 |
| Solyc09g010220.2 | Solyc09g010220.4 |
| Solyc09g010240.1 | Solyc09g010240.3 |
| Solyc09g010260.2 | Solyc09g010260.4 |
| Solyc09g010330.2 | Solyc09g010330.4 |
| Solyc09g010350.2 | Solyc09g010350.4 |
| Solyc09g010420.2 | Solyc09g010420.4 |
| Solyc09g010490.1 | Solyc09g010490.3 |
| Solyc09g010510.2 | Solyc09g010510.4 |
| Solyc09g010560.1 | Solyc09g010560.3 |
| Solyc09g010580.1 | Solyc09g010580.3 |
| Solyc09g010610.1 | Solyc09g010610.3 |
| Solyc09g010670.2 | Solyc09g010670.4 |
| Solyc09g010690.2 | Solyc09g010690.4 |
| Solyc09g010740.2 | Solyc09g010740.4 |
| Solyc09g010750.1 | Solyc09g010750.3 |
| Solyc09g010830.2 | Solyc09g010830.4 |
| Solyc09g010850.2 | Solyc09g010850.4 |
| Solyc09g010860.2 | Solyc09g010860.4 |
| Solyc09g010870.2 | Solyc09g010870.4 |
| Solyc09g010910.1 | Solyc09g010910.3 |
| Solyc09g010930.2 | Solyc09g010930.4 |
| Solyc09g010940.2 | Solyc09g010940.4 |
| Solyc09g010970.2 | Solyc09g010970.4 |
| Solyc09g011090.1 | Solyc09g011090.3 |
| Solyc09g011140.2 | Solyc09g011140.4 |
| Solyc09g011160.2 | Solyc09g011160.4 |
| Solyc09g011210.2 | Solyc09g011210.4 |
| Solyc09g011270.2 | Solyc09g011270.4 |
| Solyc09g011330.1 | Solyc09g011330.3 |
| Solyc09g011350.1 | Solyc09g011350.3 |
| Solyc09g011360.2 | Solyc09g011360.4 |
| Solyc09g011370.1 | Solyc09g011370.3 |
| Solyc09g011410.2 | Solyc09g011410.4 |

| | |
|---|---|
| Solyc09g011450.2 | Solyc09g011450.4 |
| Solyc09g011530.1 | Solyc09g011530.3 |
| Solyc09g011590.2 | Solyc09g011590.4 |
| Solyc09g011650.2 | Solyc09g011650.4 |
| Solyc09g011670.2 | Solyc09g011670.4 |
| Solyc09g011690.2 | Solyc09g011690.4 |
| Solyc09g011710.2 | Solyc09g011710.4 |
| Solyc09g011730.2 | Solyc09g011730.4 |
| Solyc09g011740.2 | Solyc09g011740.4 |
| Solyc09g011820.2 | Solyc09g011820.4 |
| Solyc09g011830.2 | Solyc09g011830.4 |
| Solyc09g011850.2 | Solyc09g011850.4 |
| Solyc09g011860.2 | Solyc09g011860.4 |
| Solyc09g011870.1 | Solyc09g011870.3 |
| Solyc09g011970.1 | Solyc09g011970.3 |
| Solyc09g013090.2 | Solyc09g013090.4 |
| Solyc09g013140.1 | Solyc09g013140.3 |
| Solyc09g013150.2 | Solyc09g013150.4 |
| Solyc09g014160.1 | Solyc09g014160.3 |
| Solyc09g014210.1 | Solyc09g014210.3 |
| Solyc09g014240.2 | Solyc09g014240.4 |
| Solyc09g014320.2 | Solyc09g014320.4 |
| Solyc09g014440.2 | Solyc09g014440.4 |
| Solyc09g014450.2 | Solyc09g014450.4 |
| Solyc09g014480.1 | Solyc09g014480.3 |
| Solyc09g014540.1 | Solyc09g014540.3 |
| Solyc09g014610.2 | Solyc09g014610.4 |
| Solyc09g014690.1 | Solyc09g014690.3 |
| Solyc09g014700.1 | Solyc09g014700.3 |
| Solyc09g014710.2 | Solyc09g014710.4 |
| Solyc09g014720.1 | Solyc09g014720.3 |
| Solyc09g014760.2 | Solyc09g014760.4 |
| Solyc09g014790.2 | Solyc09g014790.4 |
| Solyc09g014800.1 | Solyc09g014800.3 |
| Solyc09g014830.1 | Solyc09g014830.3 |
| Solyc09g014910.2 | Solyc09g014910.4 |
| Solyc09g014940.1 | Solyc09g014940.3 |
| Solyc09g014970.2 | Solyc09g014970.4 |
| Solyc09g014980.2 | Solyc09g014980.4 |
| Solyc09g014990.2 | Solyc09g014990.4 |
| Solyc09g015000.2 | Solyc09g015000.4 |
| Solyc09g015120.2 | Solyc09g015120.4 |
| Solyc09g015130.2 | Solyc09g015130.4 |

| | |
|---|---|
| Solyc09g015170.2 | Solyc09g015170.4 |
| Solyc09g015180.2 | Solyc09g015180.4 |
| Solyc09g015220.1 | Solyc09g015220.3 |
| Solyc09g015310.1 | Solyc09g015310.3 |
| Solyc09g015320.1 | Solyc09g015320.3 |
| Solyc09g015340.1 | Solyc09g015340.3 |
| Solyc09g015350.2 | Solyc09g015350.4 |
| Solyc09g015360.2 | Solyc09g015360.4 |
| Solyc09g015470.2 | Solyc09g015470.4 |
| Solyc09g015600.1 | Solyc09g015600.3 |
| Solyc09g015650.2 | Solyc09g015650.4 |
| Solyc09g015660.2 | Solyc09g015660.4 |
| Solyc09g015700.2 | Solyc09g015700.4 |
| Solyc09g015810.1 | Solyc09g015810.3 |
| Solyc09g015830.1 | Solyc09g015830.3 |
| Solyc09g015840.2 | Solyc09g015840.4 |
| Solyc09g015850.2 | Solyc09g015850.4 |
| Solyc09g015880.2 | Solyc09g015880.4 |
| Solyc09g015890.2 | Solyc09g015890.4 |
| Solyc09g016940.2 | Solyc09g016940.4 |
| Solyc09g017990.1 | Solyc09g017990.3 |
| Solyc09g018030.2 | Solyc09g018030.4 |
| Solyc09g018060.2 | Solyc09g018060.4 |
| Solyc09g018130.1 | Solyc09g018130.3 |
| Solyc09g018170.2 | Solyc09g018170.4 |
| Solyc09g018180.1 | Solyc09g018180.3 |
| Solyc09g018220.1 | Solyc09g018220.3 |
| Solyc09g018330.1 | Solyc09g018330.3 |
| Solyc09g018410.1 | Solyc09g018410.3 |
| Solyc09g018420.1 | Solyc09g018420.3 |
| Solyc09g018490.2 | Solyc09g018490.4 |
| Solyc09g018500.1 | Solyc09g018500.3 |
| Solyc09g018530.1 | Solyc09g018530.3 |
| Solyc09g018690.2 | Solyc09g018690.4 |
| Solyc09g018730.2 | Solyc09g018730.4 |
| Solyc09g018780.2 | Solyc09g018780.4 |
| Solyc09g018900.1 | Solyc09g018900.3 |
| Solyc09g018960.2 | Solyc09g018960.4 |
| Solyc09g025230.2 | Solyc09g025230.4 |
| Solyc09g025250.2 | Solyc09g025250.4 |
| Solyc09g025260.2 | Solyc09g025260.4 |
| Solyc09g025310.2 | Solyc09g025310.4 |
| Solyc09g030360.1 | Solyc09g030360.3 |

| | |
|---|---|
| Solyc09g031550.1 | Solyc09g031550.3 |
| Solyc09g031590.2 | Solyc09g031590.4 |
| Solyc09g031610.2 | Solyc09g031610.4 |
| Solyc09g031680.1 | Solyc09g031680.3 |
| Solyc09g031700.2 | Solyc09g031700.4 |
| Solyc09g031750.2 | Solyc09g031750.4 |
| Solyc09g031790.2 | Solyc09g031790.4 |
| Solyc09g031870.2 | Solyc09g031870.4 |
| Solyc09g031880.1 | Solyc09g031880.3 |
| Solyc09g031890.1 | Solyc09g031890.3 |
| Solyc09g031950.1 | Solyc09g031950.3 |
| Solyc09g037020.1 | Solyc09g037020.3 |
| Solyc09g037060.1 | Solyc09g037060.3 |
| Solyc09g037080.1 | Solyc09g037080.3 |
| Solyc09g037170.1 | Solyc09g037170.3 |
| Solyc09g042290.1 | Solyc09g042290.3 |
| Solyc09g042660.2 | Solyc09g042660.4 |
| Solyc09g042670.2 | Solyc09g042670.4 |
| Solyc09g042700.2 | Solyc09g042700.4 |
| Solyc09g042740.2 | Solyc09g042740.4 |
| Solyc09g042770.2 | Solyc09g042770.4 |
| Solyc09g047870.2 | Solyc09g047870.4 |
| Solyc09g047910.2 | Solyc09g047910.4 |
| Solyc09g048970.1 | Solyc09g048970.3 |
| Solyc09g048980.2 | Solyc09g048980.4 |
| Solyc09g048990.2 | Solyc09g048990.4 |
| Solyc09g050020.1 | Solyc09g050020.3 |
| Solyc09g055180.2 | Solyc09g055180.4 |
| Solyc09g055260.2 | Solyc09g055260.4 |
| Solyc09g055340.2 | Solyc09g055340.4 |
| Solyc09g055460.1 | Solyc09g055460.3 |
| Solyc09g055650.1 | Solyc09g055650.3 |
| Solyc09g055660.1 | Solyc09g055660.3 |
| Solyc09g055700.2 | Solyc09g055700.4 |
| Solyc09g055890.2 | Solyc09g055890.4 |
| Solyc09g055900.2 | Solyc09g055900.4 |
| Solyc09g055920.2 | Solyc09g055920.4 |
| Solyc09g056130.2 | Solyc09g056130.4 |
| Solyc09g056230.1 | Solyc09g056230.3 |
| Solyc09g056260.2 | Solyc09g056260.4 |
| Solyc09g056270.1 | Solyc09g056270.3 |
| Solyc09g056290.1 | Solyc09g056290.3 |
| Solyc09g056350.2 | Solyc09g056350.4 |

| | |
|---|---|
| Solyc09g056430.2 | Solyc09g056430.4 |
| Solyc09g056450.2 | Solyc09g056450.4 |
| Solyc09g057520.1 | Solyc09g057520.3 |
| Solyc09g057580.2 | Solyc09g057580.4 |
| Solyc09g057640.2 | Solyc09g057640.4 |
| Solyc09g057710.2 | Solyc09g057710.4 |
| Solyc09g057760.2 | Solyc09g057760.4 |
| Solyc09g057870.2 | Solyc09g057870.4 |
| Solyc09g057900.1 | Solyc09g057900.3 |
| Solyc09g057930.1 | Solyc09g057930.3 |
| Solyc09g059020.2 | Solyc09g059020.4 |
| Solyc09g059030.2 | Solyc09g059030.4 |
| Solyc09g059070.1 | Solyc09g059070.3 |
| Solyc09g059170.1 | Solyc09g059170.3 |
| Solyc09g059260.2 | Solyc09g059260.4 |
| Solyc09g059280.1 | Solyc09g059280.3 |
| Solyc09g059290.2 | Solyc09g059290.4 |
| Solyc09g059320.1 | Solyc09g059320.3 |
| Solyc09g059330.1 | Solyc09g059330.3 |
| Solyc09g059400.2 | Solyc09g059400.4 |
| Solyc09g059460.2 | Solyc09g059460.4 |
| Solyc09g059600.2 | Solyc09g059600.4 |
| Solyc09g059620.2 | Solyc09g059620.4 |
| Solyc09g059630.1 | Solyc09g059630.3 |
| Solyc09g059640.1 | Solyc09g059640.3 |
| Solyc09g059650.2 | Solyc09g059650.4 |
| Solyc09g059670.2 | Solyc09g059670.4 |
| Solyc09g059740.1 | Solyc09g059740.3 |
| Solyc09g059830.1 | Solyc09g059830.3 |
| Solyc09g059850.1 | Solyc09g059850.3 |
| Solyc09g059980.1 | Solyc09g059980.3 |
| Solyc09g060080.2 | Solyc09g060080.4 |
| Solyc09g060110.2 | Solyc09g060110.4 |
| Solyc09g060150.1 | Solyc09g060150.3 |
| Solyc09g060180.1 | Solyc09g060180.3 |
| Solyc09g061250.2 | Solyc09g061250.4 |
| Solyc09g061270.2 | Solyc09g061270.4 |
| Solyc09g061280.2 | Solyc09g061280.4 |
| Solyc09g061320.2 | Solyc09g061320.4 |
| Solyc09g061330.1 | Solyc09g061330.3 |
| Solyc09g061460.1 | Solyc09g061460.3 |
| Solyc09g061750.1 | Solyc09g061750.3 |
| Solyc09g061840.2 | Solyc09g061840.4 |

| | |
|---|---|
| Solyc09g061860.2 | Solyc09g061860.4 |
| Solyc09g061870.1 | Solyc09g061870.3 |
| Solyc09g061930.2 | Solyc09g061930.4 |
| Solyc09g061940.1 | Solyc09g061940.3 |
| Solyc09g063010.2 | Solyc09g063010.4 |
| Solyc09g063020.2 | Solyc09g063020.4 |
| Solyc09g063030.2 | Solyc09g063030.4 |
| Solyc09g063140.2 | Solyc09g063140.4 |
| Solyc09g064200.2 | Solyc09g064200.4 |
| Solyc09g064230.1 | Solyc09g064230.3 |
| Solyc09g064370.2 | Solyc09g064370.4 |
| Solyc09g064430.2 | Solyc09g064430.4 |
| Solyc09g064440.2 | Solyc09g064440.4 |
| Solyc09g064490.1 | Solyc09g064490.3 |
| Solyc09g064590.2 | Solyc09g064590.4 |
| Solyc09g064620.1 | Solyc09g064620.3 |
| Solyc09g064660.2 | Solyc09g064660.4 |
| Solyc09g064670.1 | Solyc09g064670.3 |
| Solyc09g064680.1 | Solyc09g064680.3 |
| Solyc09g064740.2 | Solyc09g064740.4 |
| Solyc09g064800.1 | Solyc09g064800.3 |
| Solyc09g064810.2 | Solyc09g064810.4 |
| Solyc09g064840.2 | Solyc09g064840.4 |
| Solyc09g064890.2 | Solyc09g064890.4 |
| Solyc09g065000.2 | Solyc09g065000.4 |
| Solyc09g065010.2 | Solyc09g065010.4 |
| Solyc09g065020.2 | Solyc09g065020.4 |
| Solyc09g065100.1 | Solyc09g065100.3 |
| Solyc09g065120.2 | Solyc09g065120.4 |
| Solyc09g065130.2 | Solyc09g065130.4 |
| Solyc09g065190.1 | Solyc09g065190.3 |
| Solyc09g065200.2 | Solyc09g065200.4 |
| Solyc09g065210.2 | Solyc09g065210.4 |
| Solyc09g065280.2 | Solyc09g065280.4 |
| Solyc09g065290.2 | Solyc09g065290.4 |
| Solyc09g065320.1 | Solyc09g065320.3 |
| Solyc09g065360.2 | Solyc09g065360.4 |
| Solyc09g065370.1 | Solyc09g065370.3 |
| Solyc09g065410.2 | Solyc09g065410.4 |
| Solyc09g065430.2 | Solyc09g065430.4 |
| Solyc09g065440.2 | Solyc09g065440.4 |
| Solyc09g065450.2 | Solyc09g065450.4 |
| Solyc09g065520.2 | Solyc09g065520.4 |

| | |
|---|---|
| Solyc09g065550.2 | Solyc09g065550.4 |
| Solyc09g065570.2 | Solyc09g065570.4 |
| Solyc09g065580.2 | Solyc09g065580.4 |
| Solyc09g065660.2 | Solyc09g065660.4 |
| Solyc09g065690.2 | Solyc09g065690.4 |
| Solyc09g065720.2 | Solyc09g065720.4 |
| Solyc09g065760.2 | Solyc09g065760.4 |
| Solyc09g065800.2 | Solyc09g065800.4 |
| Solyc09g065820.2 | Solyc09g065820.4 |
| Solyc09g065830.2 | Solyc09g065830.4 |
| Solyc09g065850.2 | Solyc09g065850.4 |
| Solyc09g065870.2 | Solyc09g065870.4 |
| Solyc09g065880.2 | Solyc09g065880.4 |
| Solyc09g065890.2 | Solyc09g065890.4 |
| Solyc09g065910.1 | Solyc09g065910.3 |
| Solyc09g065940.2 | Solyc09g065940.4 |
| Solyc09g066060.1 | Solyc09g066060.3 |
| Solyc09g066070.2 | Solyc09g066070.4 |
| Solyc09g066080.1 | Solyc09g066080.3 |
| Solyc09g066120.2 | Solyc09g066120.4 |
| Solyc09g066130.2 | Solyc09g066130.4 |
| Solyc09g066210.2 | Solyc09g066210.4 |
| Solyc09g066260.2 | Solyc09g066260.4 |
| Solyc09g066280.2 | Solyc09g066280.4 |
| Solyc09g066300.1 | Solyc09g066300.3 |
| Solyc09g066410.1 | Solyc09g066410.3 |
| Solyc09g066440.2 | Solyc09g066440.4 |
| Solyc09g066450.1 | Solyc09g066450.3 |
| Solyc09g066460.2 | Solyc09g066460.4 |
| Solyc09g066470.2 | Solyc09g066470.4 |
| Solyc09g066490.2 | Solyc09g066490.4 |
| Solyc09g066500.2 | Solyc09g066500.4 |
| Solyc09g072550.1 | Solyc09g072550.3 |
| Solyc09g072560.2 | Solyc09g072560.4 |
| Solyc09g072620.2 | Solyc09g072620.4 |
| Solyc09g072650.2 | Solyc09g072650.4 |
| Solyc09g072660.2 | Solyc09g072660.4 |
| Solyc09g072700.2 | Solyc09g072700.4 |
| Solyc09g072710.2 | Solyc09g072710.4 |
| Solyc09g072720.1 | Solyc09g072720.3 |
| Solyc09g072820.2 | Solyc09g072820.4 |
| Solyc09g072830.2 | Solyc09g072830.4 |
| Solyc09g072900.2 | Solyc09g072900.4 |

| | |
|---|---|
| Solyc09g072910.1 | Solyc09g072910.3 |
| Solyc09g073010.1 | Solyc09g073010.3 |
| Solyc09g073030.2 | Solyc09g073030.4 |
| Solyc09g074080.1 | Solyc09g074080.3 |
| Solyc09g074100.2 | Solyc09g074100.4 |
| Solyc09g074110.2 | Solyc09g074110.4 |
| Solyc09g074120.1 | Solyc09g074120.3 |
| Solyc09g074130.1 | Solyc09g074130.3 |
| Solyc09g074140.1 | Solyc09g074140.3 |
| Solyc09g074150.1 | Solyc09g074150.3 |
| Solyc09g074200.1 | Solyc09g074200.3 |
| Solyc09g074230.2 | Solyc09g074230.4 |
| Solyc09g074360.2 | Solyc09g074360.4 |
| Solyc09g074370.2 | Solyc09g074370.4 |
| Solyc09g074380.2 | Solyc09g074380.4 |
| Solyc09g074390.1 | Solyc09g074390.3 |
| Solyc09g074410.2 | Solyc09g074410.4 |
| Solyc09g074420.2 | Solyc09g074420.4 |
| Solyc09g074430.2 | Solyc09g074430.4 |
| Solyc09g074460.1 | Solyc09g074460.3 |
| Solyc09g074500.1 | Solyc09g074500.3 |
| Solyc09g074670.2 | Solyc09g074670.4 |
| Solyc09g074730.1 | Solyc09g074730.3 |
| Solyc09g074790.1 | Solyc09g074790.3 |
| Solyc09g074820.2 | Solyc09g074820.4 |
| Solyc09g074850.2 | Solyc09g074850.4 |
| Solyc09g074870.2 | Solyc09g074870.4 |
| Solyc09g074880.2 | Solyc09g074880.4 |
| Solyc09g074900.2 | Solyc09g074900.4 |
| Solyc09g074940.1 | Solyc09g074940.3 |
| Solyc09g074990.2 | Solyc09g074990.4 |
| Solyc09g075050.1 | Solyc09g075050.3 |
| Solyc09g075200.2 | Solyc09g075200.4 |
| Solyc09g075330.2 | Solyc09g075330.4 |
| Solyc09g075350.2 | Solyc09g075350.4 |
| Solyc09g075360.2 | Solyc09g075360.4 |
| Solyc09g075390.2 | Solyc09g075390.4 |
| Solyc09g075400.2 | Solyc09g075400.4 |
| Solyc09g075410.2 | Solyc09g075410.4 |
| Solyc09g075440.2 | Solyc09g075440.4 |
| Solyc09g075490.2 | Solyc09g075490.4 |
| Solyc09g075510.2 | Solyc09g075510.4 |
| Solyc09g075620.1 | Solyc09g075620.3 |

| | |
|---|---|
| Solyc09g075640.2 | Solyc09g075640.4 |
| Solyc09g075720.2 | Solyc09g075720.4 |
| Solyc09g075760.2 | Solyc09g075760.4 |
| Solyc09g075770.1 | Solyc09g075770.3 |
| Solyc09g075800.2 | Solyc09g075800.4 |
| Solyc09g075840.2 | Solyc09g075840.4 |
| Solyc09g076030.2 | Solyc09g076030.4 |
| Solyc09g076050.2 | Solyc09g076050.4 |
| Solyc09g082050.2 | Solyc09g082050.4 |
| Solyc09g082110.2 | Solyc09g082110.4 |
| Solyc09g082140.2 | Solyc09g082140.4 |
| Solyc09g082170.2 | Solyc09g082170.4 |
| Solyc09g082210.2 | Solyc09g082210.4 |
| Solyc09g082240.2 | Solyc09g082240.4 |
| Solyc09g082270.2 | Solyc09g082270.4 |
| Solyc09g082360.2 | Solyc09g082360.4 |
| Solyc09g082370.2 | Solyc09g082370.4 |
| Solyc09g082380.2 | Solyc09g082380.4 |
| Solyc09g082390.1 | Solyc09g082390.3 |
| Solyc09g082470.2 | Solyc09g082470.4 |
| Solyc09g082490.2 | Solyc09g082490.4 |
| Solyc09g082500.2 | Solyc09g082500.4 |
| Solyc09g082530.1 | Solyc09g082530.3 |
| Solyc09g082560.2 | Solyc09g082560.4 |
| Solyc09g082580.2 | Solyc09g082580.4 |
| Solyc09g082590.2 | Solyc09g082590.4 |
| Solyc09g082600.1 | Solyc09g082600.3 |
| Solyc09g082650.2 | Solyc09g082650.4 |
| Solyc09g082730.2 | Solyc09g082730.5 |
| Solyc09g082790.2 | Solyc09g082790.4 |
| Solyc09g082830.2 | Solyc09g082830.4 |
| Solyc09g082870.1 | Solyc09g082870.3 |
| Solyc09g083060.2 | Solyc09g083060.4 |
| Solyc09g083080.2 | Solyc09g083080.4 |
| Solyc09g083090.2 | Solyc09g083090.4 |
| Solyc09g083110.1 | Solyc09g083110.3 |
| Solyc09g083150.2 | Solyc09g083150.4 |
| Solyc09g083200.2 | Solyc09g083200.4 |
| Solyc09g083260.2 | Solyc09g083260.4 |
| Solyc09g083270.2 | Solyc09g083270.4 |
| Solyc09g083300.1 | Solyc09g083300.3 |
| Solyc09g083310.1 | Solyc09g083310.3 |
| Solyc09g083340.1 | Solyc09g083340.3 |

| | |
|---|---|
| Solyc09g083380.2 | Solyc09g083380.4 |
| Solyc09g083410.2 | Solyc09g083410.4 |
| Solyc09g083430.1 | Solyc09g083430.3 |
| Solyc09g084490.2 | Solyc09g084490.4 |
| Solyc09g089490.2 | Solyc09g089490.4 |
| Solyc09g089520.2 | Solyc09g089520.4 |
| Solyc09g089540.2 | Solyc09g089540.4 |
| Solyc09g089550.2 | Solyc09g089550.4 |
| Solyc09g089560.2 | Solyc09g089560.4 |
| Solyc09g089570.2 | Solyc09g089570.4 |
| Solyc09g089580.2 | Solyc09g089580.4 |
| Solyc09g089660.2 | Solyc09g089660.4 |
| Solyc09g089680.2 | Solyc09g089680.4 |
| Solyc09g089720.1 | Solyc09g089720.3 |
| Solyc09g089830.2 | Solyc09g089830.4 |
| Solyc09g089840.1 | Solyc09g089840.3 |
| Solyc09g089860.2 | Solyc09g089860.4 |
| Solyc09g089930.1 | Solyc09g089930.3 |
| Solyc09g090020.2 | Solyc09g090020.4 |
| Solyc09g090140.2 | Solyc09g090140.4 |
| Solyc09g090150.2 | Solyc09g090150.4 |
| Solyc09g090200.2 | Solyc09g090200.4 |
| Solyc09g090250.2 | Solyc09g090250.4 |
| Solyc09g090280.1 | Solyc09g090280.3 |
| Solyc09g090310.1 | Solyc09g090310.3 |
| Solyc09g090400.2 | Solyc09g090400.4 |
| Solyc09g090420.2 | Solyc09g090420.4 |
| Solyc09g090490.1 | Solyc09g090490.3 |
| Solyc09g090590.2 | Solyc09g090590.4 |
| Solyc09g090630.1 | Solyc09g090630.3 |
| Solyc09g090740.2 | Solyc09g090740.4 |
| Solyc09g090760.2 | Solyc09g090760.4 |
| Solyc09g090810.1 | Solyc09g090810.3 |
| Solyc09g090830.2 | Solyc09g090830.4 |
| Solyc09g090860.2 | Solyc09g090860.4 |
| Solyc09g090890.1 | Solyc09g090890.3 |
| Solyc09g090900.2 | Solyc09g090900.4 |
| Solyc09g090910.1 | Solyc09g090910.3 |
| Solyc09g090920.2 | Solyc09g090920.4 |
| Solyc09g090960.2 | Solyc09g090960.4 |
| Solyc09g090970.2 | Solyc09g090970.4 |
| Solyc09g091000.2 | Solyc09g091000.4 |
| Solyc09g091050.2 | Solyc09g091050.4 |

| | |
|---|---|
| Solyc09g091070.1 | Solyc09g091070.3 |
| Solyc09g091080.2 | Solyc09g091080.4 |
| Solyc09g091100.2 | Solyc09g091100.4 |
| Solyc09g091130.2 | Solyc09g091130.4 |
| Solyc09g091140.2 | Solyc09g091140.4 |
| Solyc09g091170.2 | Solyc09g091170.4 |
| Solyc09g091210.2 | Solyc09g091210.4 |
| Solyc09g091220.1 | Solyc09g091220.3 |
| Solyc09g091230.2 | Solyc09g091230.4 |
| Solyc09g091240.2 | Solyc09g091240.4 |
| Solyc09g091250.2 | Solyc09g091250.4 |
| Solyc09g091280.2 | Solyc09g091280.4 |
| Solyc09g091400.2 | Solyc09g091400.4 |
| Solyc09g091420.2 | Solyc09g091420.4 |
| Solyc09g091430.2 | Solyc09g091430.4 |
| Solyc09g091480.2 | Solyc09g091480.4 |
| Solyc09g091490.1 | Solyc09g091490.3 |
| Solyc09g091600.2 | Solyc09g091600.4 |
| Solyc09g091700.2 | Solyc09g091700.4 |
| Solyc09g091710.1 | Solyc09g091710.3 |
| Solyc09g091730.2 | Solyc09g091730.4 |
| Solyc09g091740.2 | Solyc09g091740.4 |
| Solyc09g091760.1 | Solyc09g091760.3 |
| Solyc09g091800.2 | Solyc09g091800.4 |
| Solyc09g091840.2 | Solyc09g091840.4 |
| Solyc09g091850.2 | Solyc09g091850.4 |
| Solyc09g091860.2 | Solyc09g091860.4 |
| Solyc09g091880.2 | Solyc09g091880.4 |
| Solyc09g091900.2 | Solyc09g091900.4 |
| Solyc09g091970.2 | Solyc09g091970.4 |
| Solyc09g092020.2 | Solyc09g092020.4 |
| Solyc09g092030.2 | Solyc09g092030.4 |
| Solyc09g092050.2 | Solyc09g092050.4 |
| Solyc09g092060.1 | Solyc09g092060.3 |
| Solyc09g092070.2 | Solyc09g092070.4 |
| Solyc09g092080.2 | Solyc09g092080.4 |
| Solyc09g092090.2 | Solyc09g092090.4 |
| Solyc09g092100.2 | Solyc09g092100.4 |
| Solyc09g092110.2 | Solyc09g092110.4 |
| Solyc09g092120.2 | Solyc09g092120.4 |
| Solyc09g092170.1 | Solyc09g092170.3 |
| Solyc09g092200.1 | Solyc09g092200.3 |
| Solyc09g092210.2 | Solyc09g092210.4 |

| | |
|---|---|
| Solyc09g092260.2 | Solyc09g092260.4 |
| Solyc09g092330.1 | Solyc09g092330.3 |
| Solyc09g092340.2 | Solyc09g092340.4 |
| Solyc09g092410.2 | Solyc09g092410.4 |
| Solyc09g092530.2 | Solyc09g092530.4 |
| Solyc09g092560.2 | Solyc09g092560.4 |
| Solyc09g092580.2 | Solyc09g092580.4 |
| Solyc09g092680.2 | Solyc09g092680.4 |
| Solyc09g097830.2 | Solyc09g097830.4 |
| Solyc09g097860.2 | Solyc09g097860.4 |
| Solyc09g097870.2 | Solyc09g097870.4 |
| Solyc09g097880.2 | Solyc09g097880.4 |
| Solyc09g097910.2 | Solyc09g097910.4 |
| Solyc09g097920.1 | Solyc09g097920.3 |
| Solyc09g097930.2 | Solyc09g097930.4 |
| Solyc09g097950.1 | Solyc09g097950.3 |
| Solyc09g097980.2 | Solyc09g097980.4 |
| Solyc09g098000.2 | Solyc09g098000.4 |
| Solyc09g098030.2 | Solyc09g098030.4 |
| Solyc09g098080.2 | Solyc09g098080.4 |
| Solyc09g098090.2 | Solyc09g098090.4 |
| Solyc09g098100.2 | Solyc09g098100.4 |
| Solyc09g098110.2 | Solyc09g098110.4 |
| Solyc09g098130.1 | Solyc09g098130.3 |
| Solyc09g098170.2 | Solyc09g098170.4 |
| Solyc09g098180.2 | Solyc09g098180.4 |
| Solyc09g098240.2 | Solyc09g098240.4 |
| Solyc09g098260.1 | Solyc09g098260.3 |
| Solyc09g098270.2 | Solyc09g098270.4 |
| Solyc09g098350.2 | Solyc09g098350.4 |
| Solyc09g098360.2 | Solyc09g098360.4 |
| Solyc09g098370.2 | Solyc09g098370.4 |
| Solyc09g098380.1 | Solyc09g098380.3 |
| Solyc09g098400.1 | Solyc09g098400.3 |
| Solyc09g098410.1 | Solyc09g098410.3 |
| Solyc09g098420.1 | Solyc09g098420.3 |
| Solyc09g098440.2 | Solyc09g098440.4 |
| Solyc09g098450.2 | Solyc09g098450.4 |
| Solyc09g098470.1 | Solyc09g098470.3 |
| Solyc09g098490.2 | Solyc09g098490.4 |
| Solyc09g098500.2 | Solyc09g098500.4 |
| Solyc09g098520.2 | Solyc09g098520.4 |
| Solyc09g098550.2 | Solyc09g098550.4 |

| | |
|---|---|
| Solyc09g098590.2 | Solyc09g098590.4 |
| Solyc10g005000.2 | Solyc10g005000.4 |
| Solyc10g005010.2 | Solyc10g005010.4 |
| Solyc10g005030.2 | Solyc10g005030.4 |
| Solyc10g005040.2 | Solyc10g005040.4 |
| Solyc10g005060.2 | Solyc10g005060.4 |
| Solyc10g005150.1 | Solyc10g005150.3 |
| Solyc10g005210.2 | Solyc10g005210.4 |
| Solyc10g005290.2 | Solyc10g005290.4 |
| Solyc10g005390.2 | Solyc10g005390.4 |
| Solyc10g005410.2 | Solyc10g005410.4 |
| Solyc10g005430.2 | Solyc10g005430.4 |
| Solyc10g005440.1 | Solyc10g005440.3 |
| Solyc10g005490.2 | Solyc10g005490.4 |
| Solyc10g005540.2 | Solyc10g005540.4 |
| Solyc10g005550.1 | Solyc10g005550.3 |
| Solyc10g005590.2 | Solyc10g005590.4 |
| Solyc10g005600.2 | Solyc10g005600.4 |
| Solyc10g005640.2 | Solyc10g005640.4 |
| Solyc10g005670.1 | Solyc10g005670.3 |
| Solyc10g005740.1 | Solyc10g005740.3 |
| Solyc10g005760.1 | Solyc10g005760.3 |
| Solyc10g005770.2 | Solyc10g005770.4 |
| Solyc10g005820.1 | Solyc10g005820.3 |
| Solyc10g005840.2 | Solyc10g005840.4 |
| Solyc10g005850.2 | Solyc10g005850.4 |
| Solyc10g005870.2 | Solyc10g005870.4 |
| Solyc10g005920.2 | Solyc10g005920.4 |
| Solyc10g005990.2 | Solyc10g005990.4 |
| Solyc10g006000.2 | Solyc10g006000.4 |
| Solyc10g006080.1 | Solyc10g006080.3 |
| Solyc10g006090.2 | Solyc10g006090.4 |
| Solyc10g006170.2 | Solyc10g006170.4 |
| Solyc10g006190.2 | Solyc10g006190.4 |
| Solyc10g006200.2 | Solyc10g006200.4 |
| Solyc10g006210.1 | Solyc10g006210.3 |
| Solyc10g006220.2 | Solyc10g006220.4 |
| Solyc10g006270.2 | Solyc10g006270.4 |
| Solyc10g006300.2 | Solyc10g006300.4 |
| Solyc10g006380.1 | Solyc10g006380.3 |
| Solyc10g006420.2 | Solyc10g006420.4 |
| Solyc10g006500.2 | Solyc10g006500.4 |
| Solyc10g006530.2 | Solyc10g006530.4 |

| | |
|---|---|
| Solyc10g006540.2 | Solyc10g006540.4 |
| Solyc10g006550.1 | Solyc10g006550.3 |
| Solyc10g006560.2 | Solyc10g006560.4 |
| Solyc10g006590.1 | Solyc10g006590.3 |
| Solyc10g006600.2 | Solyc10g006600.4 |
| Solyc10g006610.2 | Solyc10g006610.4 |
| Solyc10g006630.1 | Solyc10g006630.3 |
| Solyc10g006690.2 | Solyc10g006690.4 |
| Solyc10g006710.2 | Solyc10g006710.4 |
| Solyc10g006720.2 | Solyc10g006720.4 |
| Solyc10g006740.2 | Solyc10g006740.4 |
| Solyc10g006750.2 | Solyc10g006750.4 |
| Solyc10g006770.2 | Solyc10g006770.4 |
| Solyc10g006780.2 | Solyc10g006780.4 |
| Solyc10g006790.2 | Solyc10g006790.4 |
| Solyc10g006800.2 | Solyc10g006800.4 |
| Solyc10g006810.2 | Solyc10g006810.4 |
| Solyc10g006820.2 | Solyc10g006820.4 |
| Solyc10g006830.2 | Solyc10g006830.4 |
| Solyc10g006840.2 | Solyc10g006840.4 |
| Solyc10g006850.2 | Solyc10g006850.4 |
| Solyc10g006860.2 | Solyc10g006860.4 |
| Solyc10g006890.2 | Solyc10g006890.4 |
| Solyc10g006910.1 | Solyc10g006910.3 |
| Solyc10g006930.2 | Solyc10g006930.4 |
| Solyc10g006960.1 | Solyc10g006960.3 |
| Solyc10g007060.2 | Solyc10g007060.4 |
| Solyc10g007070.2 | Solyc10g007070.4 |
| Solyc10g007130.2 | Solyc10g007130.4 |
| Solyc10g007150.2 | Solyc10g007150.4 |
| Solyc10g007220.1 | Solyc10g007220.3 |
| Solyc10g007240.1 | Solyc10g007240.3 |
| Solyc10g007270.2 | Solyc10g007270.4 |
| Solyc10g007280.2 | Solyc10g007280.4 |
| Solyc10g007290.2 | Solyc10g007290.4 |
| Solyc10g007350.2 | Solyc10g007350.4 |
| Solyc10g007360.2 | Solyc10g007360.4 |
| Solyc10g007400.2 | Solyc10g007400.4 |
| Solyc10g007430.1 | Solyc10g007430.3 |
| Solyc10g007460.2 | Solyc10g007460.4 |
| Solyc10g007470.2 | Solyc10g007470.4 |
| Solyc10g007480.2 | Solyc10g007480.4 |
| Solyc10g007500.2 | Solyc10g007500.4 |

| | |
|---|---|
| Solyc10g007520.1 | Solyc10g007520.3 |
| Solyc10g007570.2 | Solyc10g007570.4 |
| Solyc10g007590.2 | Solyc10g007590.4 |
| Solyc10g007610.2 | Solyc10g007610.4 |
| Solyc10g007620.1 | Solyc10g007620.3 |
| Solyc10g007680.2 | Solyc10g007680.4 |
| Solyc10g007720.2 | Solyc10g007720.4 |
| Solyc10g007760.2 | Solyc10g007760.4 |
| Solyc10g007780.1 | Solyc10g007780.3 |
| Solyc10g007880.2 | Solyc10g007880.4 |
| Solyc10g007890.2 | Solyc10g007890.4 |
| Solyc10g007900.2 | Solyc10g007900.4 |
| Solyc10g007930.2 | Solyc10g007930.4 |
| Solyc10g007950.2 | Solyc10g007950.4 |
| Solyc10g008010.2 | Solyc10g008010.4 |
| Solyc10g008030.2 | Solyc10g008030.4 |
| Solyc10g008040.2 | Solyc10g008040.4 |
| Solyc10g008060.2 | Solyc10g008060.4 |
| Solyc10g008070.2 | Solyc10g008070.4 |
| Solyc10g008080.1 | Solyc10g008080.3 |
| Solyc10g008110.2 | Solyc10g008110.4 |
| Solyc10g008120.2 | Solyc10g008120.4 |
| Solyc10g008200.2 | Solyc10g008200.4 |
| Solyc10g008220.2 | Solyc10g008220.4 |
| Solyc10g008230.1 | Solyc10g008230.3 |
| Solyc10g008330.2 | Solyc10g008330.4 |
| Solyc10g008340.2 | Solyc10g008340.4 |
| Solyc10g008500.2 | Solyc10g008500.4 |
| Solyc10g008510.1 | Solyc10g008510.3 |
| Solyc10g008680.1 | Solyc10g008680.3 |
| Solyc10g008700.1 | Solyc10g008700.3 |
| Solyc10g008720.2 | Solyc10g008720.4 |
| Solyc10g008800.2 | Solyc10g008800.4 |
| Solyc10g008820.2 | Solyc10g008820.4 |
| Solyc10g008980.2 | Solyc10g008980.4 |
| Solyc10g008990.2 | Solyc10g008990.4 |
| Solyc10g009000.2 | Solyc10g009000.4 |
| Solyc10g009010.2 | Solyc10g009010.4 |
| Solyc10g009030.2 | Solyc10g009030.4 |
| Solyc10g009080.2 | Solyc10g009080.4 |
| Solyc10g009100.1 | Solyc10g009100.3 |
| Solyc10g009130.2 | Solyc10g009130.4 |
| Solyc10g009210.2 | Solyc10g009210.4 |

| | |
|---|---|
| Solyc10g009280.2 | Solyc10g009280.4 |
| Solyc10g009310.2 | Solyc10g009310.4 |
| Solyc10g009350.2 | Solyc10g009350.4 |
| Solyc10g009360.2 | Solyc10g009360.4 |
| Solyc10g009440.2 | Solyc10g009440.4 |
| Solyc10g009480.2 | Solyc10g009480.4 |
| Solyc10g009500.2 | Solyc10g009500.4 |
| Solyc10g009580.2 | Solyc10g009580.4 |
| Solyc10g009590.2 | Solyc10g009590.4 |
| Solyc10g009620.1 | Solyc10g009620.3 |
| Solyc10g011680.2 | Solyc10g011680.4 |
| Solyc10g011700.2 | Solyc10g011700.4 |
| Solyc10g011770.2 | Solyc10g011770.4 |
| Solyc10g011810.2 | Solyc10g011810.4 |
| Solyc10g011820.2 | Solyc10g011820.4 |
| Solyc10g011840.1 | Solyc10g011840.3 |
| Solyc10g011910.2 | Solyc10g011910.4 |
| Solyc10g011920.1 | Solyc10g011920.3 |
| Solyc10g011950.1 | Solyc10g011950.3 |
| Solyc10g011960.1 | Solyc10g011960.3 |
| Solyc10g012070.2 | Solyc10g012070.4 |
| Solyc10g012080.2 | Solyc10g012080.4 |
| Solyc10g012110.1 | Solyc10g012110.3 |
| Solyc10g012190.1 | Solyc10g012190.3 |
| Solyc10g012210.1 | Solyc10g012210.3 |
| Solyc10g012320.1 | Solyc10g012320.3 |
| Solyc10g012350.1 | Solyc10g012350.3 |
| Solyc10g017490.1 | Solyc10g017490.3 |
| Solyc10g017540.2 | Solyc10g017540.4 |
| Solyc10g017550.2 | Solyc10g017550.4 |
| Solyc10g017630.1 | Solyc10g017630.3 |
| Solyc10g017650.1 | Solyc10g017650.3 |
| Solyc10g017910.1 | Solyc10g017910.3 |
| Solyc10g018020.1 | Solyc10g018020.3 |
| Solyc10g018030.1 | Solyc10g018030.3 |
| Solyc10g018300.1 | Solyc10g018300.3 |
| Solyc10g018330.1 | Solyc10g018330.3 |
| Solyc10g018380.1 | Solyc10g018380.3 |
| Solyc10g018390.1 | Solyc10g018390.3 |
| Solyc10g018420.1 | Solyc10g018420.3 |
| Solyc10g018630.1 | Solyc10g018630.3 |
| Solyc10g018780.1 | Solyc10g018780.3 |
| Solyc10g018980.1 | Solyc10g018980.3 |

| | |
|---|---|
| Solyc10g018990.1 | Solyc10g018990.3 |
| Solyc10g019000.1 | Solyc10g019000.3 |
| Solyc10g019130.1 | Solyc10g019130.3 |
| Solyc10g019200.1 | Solyc10g019200.3 |
| Solyc10g019260.1 | Solyc10g019260.3 |
| Solyc10g019270.1 | Solyc10g019270.3 |
| Solyc10g024330.1 | Solyc10g024330.3 |
| Solyc10g024350.1 | Solyc10g024350.3 |
| Solyc10g024370.1 | Solyc10g024370.3 |
| Solyc10g024420.1 | Solyc10g024420.3 |
| Solyc10g024470.1 | Solyc10g024470.3 |
| Solyc10g036640.1 | Solyc10g036640.3 |
| Solyc10g037880.1 | Solyc10g037880.3 |
| Solyc10g037910.1 | Solyc10g037910.3 |
| Solyc10g037960.1 | Solyc10g037960.3 |
| Solyc10g038000.1 | Solyc10g038000.3 |
| Solyc10g038030.1 | Solyc10g038030.3 |
| Solyc10g038080.1 | Solyc10g038080.3 |
| Solyc10g038120.1 | Solyc10g038120.3 |
| Solyc10g038130.1 | Solyc10g038130.3 |
| Solyc10g038180.1 | Solyc10g038180.3 |
| Solyc10g039210.2 | Solyc10g039210.4 |
| Solyc10g044450.1 | Solyc10g044450.3 |
| Solyc10g044470.1 | Solyc10g044470.3 |
| Solyc10g044520.1 | Solyc10g044520.3 |
| Solyc10g044640.1 | Solyc10g044640.3 |
| Solyc10g044670.1 | Solyc10g044670.3 |
| Solyc10g044690.1 | Solyc10g044690.3 |
| Solyc10g044720.1 | Solyc10g044720.3 |
| Solyc10g044750.1 | Solyc10g044750.3 |
| Solyc10g044900.1 | Solyc10g044900.3 |
| Solyc10g044960.1 | Solyc10g044960.3 |
| Solyc10g044980.1 | Solyc10g044980.3 |
| Solyc10g045090.1 | Solyc10g045090.3 |
| Solyc10g045190.1 | Solyc10g045190.3 |
| Solyc10g045420.1 | Solyc10g045420.3 |
| Solyc10g045500.1 | Solyc10g045500.3 |
| Solyc10g045650.1 | Solyc10g045650.3 |
| Solyc10g045760.1 | Solyc10g045760.3 |
| Solyc10g045770.1 | Solyc10g045770.3 |
| Solyc10g046770.1 | Solyc10g046770.3 |
| Solyc10g046800.1 | Solyc10g046800.3 |
| Solyc10g046820.1 | Solyc10g046820.3 |

| | |
|---|---|
| Solyc10g046930.1 | Solyc10g046930.3 |
| Solyc10g047150.1 | Solyc10g047150.3 |
| Solyc10g047420.1 | Solyc10g047420.3 |
| Solyc10g047590.1 | Solyc10g047590.3 |
| Solyc10g047610.1 | Solyc10g047610.3 |
| Solyc10g047640.1 | Solyc10g047640.3 |
| Solyc10g047650.1 | Solyc10g047650.3 |
| Solyc10g047680.1 | Solyc10g047680.3 |
| Solyc10g047700.1 | Solyc10g047700.3 |
| Solyc10g047810.1 | Solyc10g047810.3 |
| Solyc10g047980.1 | Solyc10g047980.3 |
| Solyc10g047990.1 | Solyc10g047990.3 |
| Solyc10g048000.1 | Solyc10g048000.3 |
| Solyc10g048080.1 | Solyc10g048080.3 |
| Solyc10g048100.1 | Solyc10g048100.3 |
| Solyc10g048120.1 | Solyc10g048120.3 |
| Solyc10g048130.1 | Solyc10g048130.3 |
| Solyc10g048150.1 | Solyc10g048150.3 |
| Solyc10g049210.1 | Solyc10g049210.3 |
| Solyc10g049250.1 | Solyc10g049250.3 |
| Solyc10g049300.1 | Solyc10g049300.3 |
| Solyc10g049330.1 | Solyc10g049330.3 |
| Solyc10g049510.1 | Solyc10g049510.3 |
| Solyc10g049530.1 | Solyc10g049530.3 |
| Solyc10g049550.1 | Solyc10g049550.3 |
| Solyc10g049560.1 | Solyc10g049560.3 |
| Solyc10g049570.1 | Solyc10g049570.3 |
| Solyc10g049600.1 | Solyc10g049600.3 |
| Solyc10g049650.1 | Solyc10g049650.3 |
| Solyc10g049750.1 | Solyc10g049750.3 |
| Solyc10g049850.1 | Solyc10g049850.3 |
| Solyc10g049890.1 | Solyc10g049890.3 |
| Solyc10g050080.1 | Solyc10g050080.3 |
| Solyc10g050170.1 | Solyc10g050170.3 |
| Solyc10g050190.1 | Solyc10g050190.3 |
| Solyc10g050210.1 | Solyc10g050210.3 |
| Solyc10g050230.1 | Solyc10g050230.3 |
| Solyc10g050420.1 | Solyc10g050420.3 |
| Solyc10g050470.1 | Solyc10g050470.3 |
| Solyc10g050550.1 | Solyc10g050550.3 |
| Solyc10g050750.1 | Solyc10g050750.3 |
| Solyc10g050840.1 | Solyc10g050840.3 |
| Solyc10g050860.1 | Solyc10g050860.3 |

| | |
|---|---|
| Solyc10g050870.1 | Solyc10g050870.3 |
| Solyc10g050960.1 | Solyc10g050960.3 |
| Solyc10g051010.1 | Solyc10g051010.3 |
| Solyc10g051090.1 | Solyc10g051090.3 |
| Solyc10g051120.1 | Solyc10g051120.3 |
| Solyc10g051130.1 | Solyc10g051130.3 |
| Solyc10g051140.1 | Solyc10g051140.3 |
| Solyc10g051150.1 | Solyc10g051150.3 |
| Solyc10g051220.1 | Solyc10g051220.3 |
| Solyc10g051260.1 | Solyc10g051260.3 |
| Solyc10g051330.1 | Solyc10g051330.3 |
| Solyc10g051370.1 | Solyc10g051370.3 |
| Solyc10g052580.1 | Solyc10g052580.3 |
| Solyc10g052640.1 | Solyc10g052640.3 |
| Solyc10g053910.1 | Solyc10g053910.3 |
| Solyc10g054040.1 | Solyc10g054040.3 |
| Solyc10g054050.1 | Solyc10g054050.3 |
| Solyc10g054080.1 | Solyc10g054080.3 |
| Solyc10g054100.1 | Solyc10g054100.3 |
| Solyc10g054130.1 | Solyc10g054130.3 |
| Solyc10g054170.1 | Solyc10g054170.3 |
| Solyc10g054270.1 | Solyc10g054270.3 |
| Solyc10g054330.1 | Solyc10g054330.3 |
| Solyc10g054380.1 | Solyc10g054380.3 |
| Solyc10g054390.1 | Solyc10g054390.3 |
| Solyc10g054440.1 | Solyc10g054440.3 |
| Solyc10g054480.1 | Solyc10g054480.3 |
| Solyc10g054670.1 | Solyc10g054670.3 |
| Solyc10g054870.1 | Solyc10g054870.3 |
| Solyc10g054880.1 | Solyc10g054880.3 |
| Solyc10g054950.1 | Solyc10g054950.3 |
| Solyc10g054980.1 | Solyc10g054980.3 |
| Solyc10g054990.1 | Solyc10g054990.3 |
| Solyc10g055000.1 | Solyc10g055000.3 |
| Solyc10g055010.1 | Solyc10g055010.3 |
| Solyc10g055020.1 | Solyc10g055020.3 |
| Solyc10g055040.1 | Solyc10g055040.3 |
| Solyc10g055060.1 | Solyc10g055060.3 |
| Solyc10g055080.1 | Solyc10g055080.3 |
| Solyc10g055160.1 | Solyc10g055160.3 |
| Solyc10g055210.1 | Solyc10g055210.3 |
| Solyc10g055330.1 | Solyc10g055330.3 |
| Solyc10g055550.1 | Solyc10g055550.3 |

| | |
|---|---|
| Solyc10g055570.1 | Solyc10g055570.3 |
| Solyc10g055730.1 | Solyc10g055730.3 |
| Solyc10g055820.1 | Solyc10g055820.3 |
| Solyc10g061840.1 | Solyc10g061840.3 |
| Solyc10g061940.1 | Solyc10g061940.3 |
| Solyc10g061950.1 | Solyc10g061950.3 |
| Solyc10g061990.1 | Solyc10g061990.3 |
| Solyc10g062040.1 | Solyc10g062040.3 |
| Solyc10g062160.1 | Solyc10g062160.3 |
| Solyc10g074380.1 | Solyc10g074380.3 |
| Solyc10g074400.1 | Solyc10g074400.3 |
| Solyc10g074460.1 | Solyc10g074460.3 |
| Solyc10g074470.1 | Solyc10g074470.3 |
| Solyc10g074490.1 | Solyc10g074490.3 |
| Solyc10g074550.1 | Solyc10g074550.3 |
| Solyc10g074600.1 | Solyc10g074600.3 |
| Solyc10g074610.1 | Solyc10g074610.3 |
| Solyc10g074620.1 | Solyc10g074620.3 |
| Solyc10g074630.1 | Solyc10g074630.3 |
| Solyc10g074640.1 | Solyc10g074640.3 |
| Solyc10g074800.1 | Solyc10g074800.3 |
| Solyc10g074850.1 | Solyc10g074850.3 |
| Solyc10g074880.1 | Solyc10g074880.3 |
| Solyc10g075000.1 | Solyc10g075000.3 |
| Solyc10g075020.1 | Solyc10g075020.3 |
| Solyc10g075030.1 | Solyc10g075030.3 |
| Solyc10g075040.1 | Solyc10g075040.3 |
| Solyc10g075070.1 | Solyc10g075070.3 |
| Solyc10g075090.1 | Solyc10g075090.3 |
| Solyc10g075170.1 | Solyc10g075170.3 |
| Solyc10g076200.1 | Solyc10g076200.3 |
| Solyc10g076220.1 | Solyc10g076220.3 |
| Solyc10g076230.1 | Solyc10g076230.3 |
| Solyc10g076240.1 | Solyc10g076240.3 |
| Solyc10g076270.1 | Solyc10g076270.3 |
| Solyc10g076280.1 | Solyc10g076280.3 |
| Solyc10g076320.1 | Solyc10g076320.3 |
| Solyc10g076330.1 | Solyc10g076330.3 |
| Solyc10g076340.1 | Solyc10g076340.3 |
| Solyc10g076370.1 | Solyc10g076370.3 |
| Solyc10g076500.1 | Solyc10g076500.3 |
| Solyc10g076590.1 | Solyc10g076590.3 |
| Solyc10g076650.1 | Solyc10g076650.3 |

| | |
|---|---|
| Solyc10g076670.1 | Solyc10g076670.3 |
| Solyc10g076710.1 | Solyc10g076710.3 |
| Solyc10g076740.1 | Solyc10g076740.3 |
| Solyc10g076760.1 | Solyc10g076760.3 |
| Solyc10g076890.1 | Solyc10g076890.3 |
| Solyc10g076900.1 | Solyc10g076900.3 |
| Solyc10g076920.1 | Solyc10g076920.3 |
| Solyc10g076940.1 | Solyc10g076940.3 |
| Solyc10g076990.1 | Solyc10g076990.3 |
| Solyc10g077060.1 | Solyc10g077060.3 |
| Solyc10g077080.1 | Solyc10g077080.3 |
| Solyc10g077120.1 | Solyc10g077120.3 |
| Solyc10g078150.1 | Solyc10g078150.3 |
| Solyc10g078170.1 | Solyc10g078170.3 |
| Solyc10g078180.1 | Solyc10g078180.3 |
| Solyc10g078230.1 | Solyc10g078230.3 |
| Solyc10g078360.1 | Solyc10g078360.3 |
| Solyc10g078390.1 | Solyc10g078390.3 |
| Solyc10g078510.1 | Solyc10g078510.3 |
| Solyc10g078540.1 | Solyc10g078540.3 |
| Solyc10g078600.1 | Solyc10g078600.3 |
| Solyc10g078650.1 | Solyc10g078650.3 |
| Solyc10g078670.1 | Solyc10g078670.3 |
| Solyc10g078680.1 | Solyc10g078680.3 |
| Solyc10g078690.1 | Solyc10g078690.3 |
| Solyc10g078700.1 | Solyc10g078700.3 |
| Solyc10g078800.1 | Solyc10g078800.3 |
| Solyc10g078910.1 | Solyc10g078910.3 |
| Solyc10g078920.1 | Solyc10g078920.3 |
| Solyc10g079040.1 | Solyc10g079040.3 |
| Solyc10g079090.1 | Solyc10g079090.3 |
| Solyc10g079120.1 | Solyc10g079120.3 |
| Solyc10g079150.1 | Solyc10g079150.3 |
| Solyc10g079190.1 | Solyc10g079190.3 |
| Solyc10g079270.1 | Solyc10g079270.3 |
| Solyc10g079320.1 | Solyc10g079320.3 |
| Solyc10g079350.1 | Solyc10g079350.3 |
| Solyc10g079360.1 | Solyc10g079360.3 |
| Solyc10g079370.1 | Solyc10g079370.3 |
| Solyc10g079400.1 | Solyc10g079400.3 |
| Solyc10g079510.1 | Solyc10g079510.3 |
| Solyc10g079550.1 | Solyc10g079550.3 |
| Solyc10g079570.1 | Solyc10g079570.3 |

| | |
|---|---|
| Solyc10g079660.1 | Solyc10g079660.3 |
| Solyc10g079680.1 | Solyc10g079680.3 |
| Solyc10g079710.1 | Solyc10g079710.3 |
| Solyc10g079730.1 | Solyc10g079730.3 |
| Solyc10g079770.1 | Solyc10g079770.3 |
| Solyc10g079830.1 | Solyc10g079830.3 |
| Solyc10g079870.1 | Solyc10g079870.3 |
| Solyc10g079900.1 | Solyc10g079900.3 |
| Solyc10g079990.1 | Solyc10g079990.3 |
| Solyc10g080020.1 | Solyc10g080020.3 |
| Solyc10g080040.1 | Solyc10g080040.3 |
| Solyc10g080050.1 | Solyc10g080050.3 |
| Solyc10g080070.1 | Solyc10g080070.3 |
| Solyc10g080080.1 | Solyc10g080080.3 |
| Solyc10g080090.1 | Solyc10g080090.3 |
| Solyc10g080100.1 | Solyc10g080100.3 |
| Solyc10g080110.1 | Solyc10g080110.3 |
| Solyc10g080130.1 | Solyc10g080130.3 |
| Solyc10g080180.1 | Solyc10g080180.3 |
| Solyc10g080260.1 | Solyc10g080260.3 |
| Solyc10g080310.1 | Solyc10g080310.3 |
| Solyc10g080320.1 | Solyc10g080320.3 |
| Solyc10g080340.1 | Solyc10g080340.3 |
| Solyc10g080350.1 | Solyc10g080350.3 |
| Solyc10g080410.1 | Solyc10g080410.3 |
| Solyc10g080420.1 | Solyc10g080420.3 |
| Solyc10g080520.1 | Solyc10g080520.3 |
| Solyc10g080600.1 | Solyc10g080600.3 |
| Solyc10g080640.1 | Solyc10g080640.3 |
| Solyc10g080650.1 | Solyc10g080650.3 |
| Solyc10g080660.1 | Solyc10g080660.3 |
| Solyc10g080680.1 | Solyc10g080680.3 |
| Solyc10g080730.1 | Solyc10g080730.3 |
| Solyc10g080750.1 | Solyc10g080750.3 |
| Solyc10g080770.1 | Solyc10g080770.3 |
| Solyc10g080900.1 | Solyc10g080900.3 |
| Solyc10g080940.1 | Solyc10g080940.3 |
| Solyc10g080990.1 | Solyc10g080990.3 |
| Solyc10g081010.1 | Solyc10g081010.3 |
| Solyc10g081030.1 | Solyc10g081030.3 |
| Solyc10g081100.1 | Solyc10g081100.3 |
| Solyc10g081110.1 | Solyc10g081110.3 |
| Solyc10g081120.1 | Solyc10g081120.3 |

| | |
|---|---|
| Solyc10g081140.1 | Solyc10g081140.3 |
| Solyc10g081150.1 | Solyc10g081150.3 |
| Solyc10g081210.1 | Solyc10g081210.3 |
| Solyc10g081220.1 | Solyc10g081220.3 |
| Solyc10g081230.1 | Solyc10g081230.3 |
| Solyc10g081240.1 | Solyc10g081240.3 |
| Solyc10g081250.1 | Solyc10g081250.3 |
| Solyc10g081330.1 | Solyc10g081330.3 |
| Solyc10g081370.1 | Solyc10g081370.3 |
| Solyc10g081560.1 | Solyc10g081560.3 |
| Solyc10g081570.1 | Solyc10g081570.3 |
| Solyc10g081580.1 | Solyc10g081580.3 |
| Solyc10g081640.1 | Solyc10g081640.3 |
| Solyc10g081740.1 | Solyc10g081740.3 |
| Solyc10g081760.1 | Solyc10g081760.3 |
| Solyc10g081770.1 | Solyc10g081770.3 |
| Solyc10g081780.1 | Solyc10g081780.3 |
| Solyc10g081810.1 | Solyc10g081810.3 |
| Solyc10g081820.1 | Solyc10g081820.3 |
| Solyc10g081830.1 | Solyc10g081830.3 |
| Solyc10g081840.1 | Solyc10g081840.3 |
| Solyc10g081850.1 | Solyc10g081850.3 |
| Solyc10g081870.1 | Solyc10g081870.3 |
| Solyc10g081880.1 | Solyc10g081880.3 |
| Solyc10g081900.1 | Solyc10g081900.3 |
| Solyc10g081950.1 | Solyc10g081950.3 |
| Solyc10g082000.1 | Solyc10g082000.3 |
| Solyc10g083080.1 | Solyc10g083080.3 |
| Solyc10g083110.1 | Solyc10g083110.3 |
| Solyc10g083120.1 | Solyc10g083120.3 |
| Solyc10g083150.1 | Solyc10g083150.3 |
| Solyc10g083190.1 | Solyc10g083190.3 |
| Solyc10g083220.1 | Solyc10g083220.3 |
| Solyc10g083280.1 | Solyc10g083280.4 |
| Solyc10g083320.1 | Solyc10g083320.3 |
| Solyc10g083340.1 | Solyc10g083340.3 |
| Solyc10g083450.1 | Solyc10g083450.3 |
| Solyc10g083490.1 | Solyc10g083490.3 |
| Solyc10g083540.1 | Solyc10g083540.3 |
| Solyc10g083570.1 | Solyc10g083570.3 |
| Solyc10g083670.1 | Solyc10g083670.3 |
| Solyc10g083700.2 | Solyc10g083700.4 |
| Solyc10g083720.1 | Solyc10g083720.3 |

| | |
|---|---|
| Solyc10g083740.1 | Solyc10g083740.3 |
| Solyc10g083750.1 | Solyc10g083750.3 |
| Solyc10g083760.1 | Solyc10g083760.3 |
| Solyc10g083770.1 | Solyc10g083770.3 |
| Solyc10g083790.1 | Solyc10g083790.3 |
| Solyc10g083850.1 | Solyc10g083850.3 |
| Solyc10g083860.1 | Solyc10g083860.3 |
| Solyc10g084000.1 | Solyc10g084000.3 |
| Solyc10g084080.1 | Solyc10g084080.3 |
| Solyc10g084090.1 | Solyc10g084090.3 |
| Solyc10g084220.1 | Solyc10g084220.3 |
| Solyc10g084240.1 | Solyc10g084240.3 |
| Solyc10g084260.1 | Solyc10g084260.3 |
| Solyc10g084320.1 | Solyc10g084320.3 |
| Solyc10g084370.1 | Solyc10g084370.3 |
| Solyc10g084440.1 | Solyc10g084440.3 |
| Solyc10g084540.1 | Solyc10g084540.3 |
| Solyc10g084550.1 | Solyc10g084550.3 |
| Solyc10g084560.1 | Solyc10g084560.3 |
| Solyc10g084650.1 | Solyc10g084650.3 |
| Solyc10g084670.1 | Solyc10g084670.3 |
| Solyc10g084740.1 | Solyc10g084740.3 |
| Solyc10g084760.1 | Solyc10g084760.3 |
| Solyc10g084770.1 | Solyc10g084770.3 |
| Solyc10g084780.1 | Solyc10g084780.3 |
| Solyc10g084810.1 | Solyc10g084810.3 |
| Solyc10g084890.1 | Solyc10g084890.3 |
| Solyc10g084910.1 | Solyc10g084910.3 |
| Solyc10g084940.1 | Solyc10g084940.3 |
| Solyc10g084950.1 | Solyc10g084950.3 |
| Solyc10g084980.1 | Solyc10g084980.3 |
| Solyc10g085010.1 | Solyc10g085010.3 |
| Solyc10g085020.1 | Solyc10g085020.3 |
| Solyc10g085050.1 | Solyc10g085050.3 |
| Solyc10g085060.1 | Solyc10g085060.3 |
| Solyc10g085070.1 | Solyc10g085070.3 |
| Solyc10g085090.1 | Solyc10g085090.3 |
| Solyc10g085110.1 | Solyc10g085110.3 |
| Solyc10g085120.1 | Solyc10g085120.3 |
| Solyc10g085170.1 | Solyc10g085170.3 |
| Solyc10g085220.1 | Solyc10g085220.3 |
| Solyc10g085290.1 | Solyc10g085290.3 |
| Solyc10g085360.1 | Solyc10g085360.3 |

| | |
|---|---|
| Solyc10g085370.1 | Solyc10g085370.3 |
| Solyc10g085400.1 | Solyc10g085400.3 |
| Solyc10g085410.1 | Solyc10g085410.3 |
| Solyc10g085420.1 | Solyc10g085420.3 |
| Solyc10g085430.1 | Solyc10g085430.3 |
| Solyc10g085450.1 | Solyc10g085450.3 |
| Solyc10g085460.1 | Solyc10g085460.3 |
| Solyc10g085490.1 | Solyc10g085490.3 |
| Solyc10g085510.1 | Solyc10g085510.3 |
| Solyc10g085550.1 | Solyc10g085550.3 |
| Solyc10g085560.1 | Solyc10g085560.3 |
| Solyc10g085630.1 | Solyc10g085630.3 |
| Solyc10g085660.1 | Solyc10g085660.3 |
| Solyc10g085710.1 | Solyc10g085710.3 |
| Solyc10g085730.1 | Solyc10g085730.3 |
| Solyc10g085900.1 | Solyc10g085900.3 |
| Solyc10g085920.1 | Solyc10g085920.3 |
| Solyc10g085970.1 | Solyc10g085970.3 |
| Solyc10g085990.1 | Solyc10g085990.3 |
| Solyc10g086000.1 | Solyc10g086000.3 |
| Solyc10g086020.1 | Solyc10g086020.3 |
| Solyc10g086130.1 | Solyc10g086130.3 |
| Solyc10g086190.1 | Solyc10g086190.3 |
| Solyc10g086260.1 | Solyc10g086260.3 |
| Solyc10g086290.1 | Solyc10g086290.3 |
| Solyc10g086430.1 | Solyc10g086430.3 |
| Solyc10g086480.1 | Solyc10g086480.3 |
| Solyc10g086540.1 | Solyc10g086540.3 |
| Solyc10g086560.1 | Solyc10g086560.3 |
| Solyc10g086590.1 | Solyc10g086590.3 |
| Solyc10g086600.1 | Solyc10g086600.3 |
| Solyc10g086640.1 | Solyc10g086640.3 |
| Solyc10g086690.1 | Solyc10g086690.3 |
| Solyc10g086700.1 | Solyc10g086700.3 |
| Solyc10g086710.1 | Solyc10g086710.3 |
| Solyc10g086740.1 | Solyc10g086740.3 |
| Solyc10g086770.1 | Solyc10g086770.3 |
| Solyc10g087010.1 | Solyc10g087010.3 |
| Solyc10g087040.1 | Solyc10g087040.3 |
| Solyc11g005010.1 | Solyc11g005010.3 |
| Solyc11g005020.1 | Solyc11g005020.3 |
| Solyc11g005030.1 | Solyc11g005030.3 |
| Solyc11g005100.1 | Solyc11g005100.3 |

| | |
|---|---|
| Solyc11g005110.1 | Solyc11g005110.3 |
| Solyc11g005120.1 | Solyc11g005120.3 |
| Solyc11g005150.1 | Solyc11g005150.3 |
| Solyc11g005190.1 | Solyc11g005190.3 |
| Solyc11g005250.1 | Solyc11g005250.3 |
| Solyc11g005330.1 | Solyc11g005330.2 |
| Solyc11g005340.1 | Solyc11g005340.3 |
| Solyc11g005420.1 | Solyc11g005420.3 |
| Solyc11g005450.1 | Solyc11g005450.3 |
| Solyc11g005520.1 | Solyc11g005520.3 |
| Solyc11g005560.1 | Solyc11g005560.3 |
| Solyc11g005690.1 | Solyc11g005690.3 |
| Solyc11g005730.1 | Solyc11g005730.3 |
| Solyc11g005760.1 | Solyc11g005760.3 |
| Solyc11g005780.1 | Solyc11g005780.3 |
| Solyc11g005790.1 | Solyc11g005790.3 |
| Solyc11g005890.1 | Solyc11g005890.3 |
| Solyc11g005920.1 | Solyc11g005920.3 |
| Solyc11g005940.1 | Solyc11g005940.3 |
| Solyc11g005980.1 | Solyc11g005980.3 |
| Solyc11g006000.1 | Solyc11g006000.3 |
| Solyc11g006010.1 | Solyc11g006010.3 |
| Solyc11g006070.1 | Solyc11g006070.3 |
| Solyc11g006090.1 | Solyc11g006090.3 |
| Solyc11g006160.1 | Solyc11g006160.3 |
| Solyc11g006170.1 | Solyc11g006170.3 |
| Solyc11g006190.1 | Solyc11g006190.3 |
| Solyc11g006210.1 | Solyc11g006210.3 |
| Solyc11g006230.1 | Solyc11g006230.3 |
| Solyc11g006450.1 | Solyc11g006450.3 |
| Solyc11g006460.1 | Solyc11g006460.3 |
| Solyc11g006530.1 | Solyc11g006530.3 |
| Solyc11g006560.1 | Solyc11g006560.3 |
| Solyc11g006570.1 | Solyc11g006570.3 |
| Solyc11g006580.1 | Solyc11g006580.3 |
| Solyc11g006620.1 | Solyc11g006620.3 |
| Solyc11g006640.1 | Solyc11g006640.3 |
| Solyc11g006650.1 | Solyc11g006650.3 |
| Solyc11g006660.1 | Solyc11g006660.3 |
| Solyc11g006680.1 | Solyc11g006680.3 |
| Solyc11g006740.1 | Solyc11g006740.3 |
| Solyc11g006760.1 | Solyc11g006760.3 |
| Solyc11g006770.1 | Solyc11g006770.3 |

| | |
|---|---|
| Solyc11g006850.1 | Solyc11g006850.3 |
| Solyc11g006910.1 | Solyc11g006910.3 |
| Solyc11g006930.1 | Solyc11g006930.3 |
| Solyc11g006940.1 | Solyc11g006940.3 |
| Solyc11g006950.1 | Solyc11g006950.3 |
| Solyc11g007030.1 | Solyc11g007030.3 |
| Solyc11g007090.1 | Solyc11g007090.3 |
| Solyc11g007180.1 | Solyc11g007180.3 |
| Solyc11g007200.1 | Solyc11g007200.3 |
| Solyc11g007210.1 | Solyc11g007210.3 |
| Solyc11g007270.1 | Solyc11g007270.3 |
| Solyc11g007280.1 | Solyc11g007280.3 |
| Solyc11g007310.1 | Solyc11g007310.3 |
| Solyc11g007370.1 | Solyc11g007370.3 |
| Solyc11g007410.1 | Solyc11g007410.3 |
| Solyc11g007430.1 | Solyc11g007430.3 |
| Solyc11g007450.1 | Solyc11g007450.3 |
| Solyc11g007580.1 | Solyc11g007580.3 |
| Solyc11g007590.1 | Solyc11g007590.3 |
| Solyc11g007630.1 | Solyc11g007630.3 |
| Solyc11g007670.1 | Solyc11g007670.3 |
| Solyc11g007690.1 | Solyc11g007690.3 |
| Solyc11g007750.1 | Solyc11g007750.3 |
| Solyc11g007800.1 | Solyc11g007800.3 |
| Solyc11g007900.1 | Solyc11g007900.3 |
| Solyc11g008030.1 | Solyc11g008030.3 |
| Solyc11g008040.1 | Solyc11g008040.3 |
| Solyc11g008080.1 | Solyc11g008080.3 |
| Solyc11g008120.1 | Solyc11g008120.3 |
| Solyc11g008140.1 | Solyc11g008140.3 |
| Solyc11g008150.1 | Solyc11g008150.3 |
| Solyc11g008210.1 | Solyc11g008210.3 |
| Solyc11g008220.1 | Solyc11g008220.3 |
| Solyc11g008280.1 | Solyc11g008280.3 |
| Solyc11g008310.1 | Solyc11g008310.3 |
| Solyc11g008320.1 | Solyc11g008320.3 |
| Solyc11g008330.1 | Solyc11g008330.3 |
| Solyc11g008340.1 | Solyc11g008340.3 |
| Solyc11g008360.1 | Solyc11g008360.3 |
| Solyc11g008380.1 | Solyc11g008380.3 |
| Solyc11g008410.1 | Solyc11g008410.3 |
| Solyc11g008420.1 | Solyc11g008420.3 |
| Solyc11g008440.1 | Solyc11g008440.3 |

| | |
|---|---|
| Solyc11g008490.1 | Solyc11g008490.3 |
| Solyc11g008510.1 | Solyc11g008510.3 |
| Solyc11g008520.1 | Solyc11g008520.3 |
| Solyc11g008530.1 | Solyc11g008530.3 |
| Solyc11g008540.1 | Solyc11g008540.3 |
| Solyc11g008550.1 | Solyc11g008550.3 |
| Solyc11g008580.1 | Solyc11g008580.3 |
| Solyc11g008610.1 | Solyc11g008610.3 |
| Solyc11g008630.1 | Solyc11g008630.3 |
| Solyc11g008720.1 | Solyc11g008720.3 |
| Solyc11g008760.1 | Solyc11g008760.3 |
| Solyc11g008780.1 | Solyc11g008780.3 |
| Solyc11g008790.1 | Solyc11g008790.3 |
| Solyc11g008800.1 | Solyc11g008800.3 |
| Solyc11g008860.1 | Solyc11g008860.3 |
| Solyc11g008930.1 | Solyc11g008930.3 |
| Solyc11g008960.1 | Solyc11g008960.3 |
| Solyc11g009050.1 | Solyc11g009050.3 |
| Solyc11g010110.1 | Solyc11g010110.3 |
| Solyc11g010130.1 | Solyc11g010130.3 |
| Solyc11g010200.1 | Solyc11g010200.3 |
| Solyc11g010240.1 | Solyc11g010240.3 |
| Solyc11g010260.1 | Solyc11g010260.3 |
| Solyc11g010330.1 | Solyc11g010330.3 |
| Solyc11g010360.1 | Solyc11g010360.3 |
| Solyc11g010400.1 | Solyc11g010400.3 |
| Solyc11g010420.1 | Solyc11g010420.3 |
| Solyc11g010440.1 | Solyc11g010440.3 |
| Solyc11g010460.1 | Solyc11g010460.3 |
| Solyc11g010560.1 | Solyc11g010560.3 |
| Solyc11g010590.1 | Solyc11g010590.3 |
| Solyc11g010630.2 | Solyc11g010630.4 |
| Solyc11g010650.2 | Solyc11g010650.4 |
| Solyc11g010680.2 | Solyc11g010680.4 |
| Solyc11g010690.1 | Solyc11g010690.3 |
| Solyc11g010700.1 | Solyc11g010700.3 |
| Solyc11g010730.1 | Solyc11g010730.3 |
| Solyc11g010740.1 | Solyc11g010740.3 |
| Solyc11g010860.1 | Solyc11g010860.3 |
| Solyc11g010890.1 | Solyc11g010890.3 |
| Solyc11g010910.1 | Solyc11g010910.3 |
| Solyc11g010920.1 | Solyc11g010920.3 |
| Solyc11g010990.1 | Solyc11g010990.3 |

| | |
|---|---|
| Solyc11g011080.1 | Solyc11g011080.3 |
| Solyc11g011110.1 | Solyc11g011110.3 |
| Solyc11g011130.1 | Solyc11g011130.3 |
| Solyc11g011180.1 | Solyc11g011180.3 |
| Solyc11g011230.1 | Solyc11g011230.3 |
| Solyc11g011250.1 | Solyc11g011250.3 |
| Solyc11g011290.1 | Solyc11g011290.3 |
| Solyc11g011310.1 | Solyc11g011310.3 |
| Solyc11g011350.1 | Solyc11g011350.3 |
| Solyc11g011390.1 | Solyc11g011390.3 |
| Solyc11g011410.1 | Solyc11g011410.3 |
| Solyc11g011420.1 | Solyc11g011420.3 |
| Solyc11g011560.1 | Solyc11g011560.3 |
| Solyc11g011600.1 | Solyc11g011600.3 |
| Solyc11g011620.1 | Solyc11g011620.3 |
| Solyc11g011630.1 | Solyc11g011630.3 |
| Solyc11g011770.1 | Solyc11g011770.3 |
| Solyc11g011780.1 | Solyc11g011780.3 |
| Solyc11g011790.1 | Solyc11g011790.3 |
| Solyc11g011800.1 | Solyc11g011800.3 |
| Solyc11g011820.1 | Solyc11g011820.3 |
| Solyc11g011870.1 | Solyc11g011870.3 |
| Solyc11g011930.1 | Solyc11g011930.3 |
| Solyc11g011940.1 | Solyc11g011940.3 |
| Solyc11g012070.1 | Solyc11g012070.3 |
| Solyc11g012150.1 | Solyc11g012150.3 |
| Solyc11g012190.1 | Solyc11g012190.3 |
| Solyc11g012200.1 | Solyc11g012200.3 |
| Solyc11g012230.1 | Solyc11g012230.3 |
| Solyc11g012260.1 | Solyc11g012260.3 |
| Solyc11g012320.1 | Solyc11g012320.3 |
| Solyc11g012330.1 | Solyc11g012330.3 |
| Solyc11g012460.1 | Solyc11g012460.3 |
| Solyc11g012480.1 | Solyc11g012480.3 |
| Solyc11g012500.1 | Solyc11g012500.3 |
| Solyc11g012510.1 | Solyc11g012510.3 |
| Solyc11g012530.1 | Solyc11g012530.3 |
| Solyc11g012550.1 | Solyc11g012550.3 |
| Solyc11g012560.1 | Solyc11g012560.3 |
| Solyc11g012570.1 | Solyc11g012570.3 |
| Solyc11g012590.1 | Solyc11g012590.3 |
| Solyc11g012680.1 | Solyc11g012680.3 |
| Solyc11g012690.1 | Solyc11g012690.3 |

| | |
|---|---|
| Solyc11g012730.1 | Solyc11g012730.3 |
| Solyc11g012780.1 | Solyc11g012780.3 |
| Solyc11g012830.1 | Solyc11g012830.3 |
| Solyc11g012860.1 | Solyc11g012860.3 |
| Solyc11g012900.1 | Solyc11g012900.3 |
| Solyc11g013080.1 | Solyc11g013080.3 |
| Solyc11g013210.1 | Solyc11g013210.3 |
| Solyc11g013240.1 | Solyc11g013240.3 |
| Solyc11g013260.1 | Solyc11g013260.3 |
| Solyc11g013370.1 | Solyc11g013370.3 |
| Solyc11g013390.1 | Solyc11g013390.3 |
| Solyc11g013400.1 | Solyc11g013400.3 |
| Solyc11g013480.1 | Solyc11g013480.3 |
| Solyc11g013520.1 | Solyc11g013520.3 |
| Solyc11g013530.1 | Solyc11g013530.3 |
| Solyc11g013570.1 | Solyc11g013570.3 |
| Solyc11g013730.1 | Solyc11g013730.3 |
| Solyc11g013740.1 | Solyc11g013740.3 |
| Solyc11g013750.1 | Solyc11g013750.3 |
| Solyc11g013810.1 | Solyc11g013810.3 |
| Solyc11g013840.1 | Solyc11g013840.3 |
| Solyc11g016990.1 | Solyc11g016990.3 |
| Solyc11g017140.1 | Solyc11g017140.3 |
| Solyc11g017210.1 | Solyc11g017210.3 |
| Solyc11g017240.1 | Solyc11g017240.3 |
| Solyc11g017310.1 | Solyc11g017310.3 |
| Solyc11g017320.1 | Solyc11g017320.3 |
| Solyc11g017360.1 | Solyc11g017360.3 |
| Solyc11g017370.1 | Solyc11g017370.3 |
| Solyc11g017440.1 | Solyc11g017440.3 |
| Solyc11g017460.1 | Solyc11g017460.3 |
| Solyc11g018490.1 | Solyc11g018490.3 |
| Solyc11g018500.1 | Solyc11g018500.3 |
| Solyc11g018550.2 | Solyc11g018550.4 |
| Solyc11g018620.1 | Solyc11g018620.3 |
| Solyc11g018670.1 | Solyc11g018670.3 |
| Solyc11g018700.1 | Solyc11g018700.3 |
| Solyc11g018740.1 | Solyc11g018740.3 |
| Solyc11g018800.1 | Solyc11g018800.3 |
| Solyc11g019860.1 | Solyc11g019860.3 |
| Solyc11g019870.1 | Solyc11g019870.3 |
| Solyc11g019880.1 | Solyc11g019880.3 |
| Solyc11g019890.1 | Solyc11g019890.3 |

| | |
|---|---|
| Solyc11g019970.1 | Solyc11g019970.3 |
| Solyc11g019990.1 | Solyc11g019990.4 |
| Solyc11g020030.1 | Solyc11g020030.3 |
| Solyc11g020080.1 | Solyc11g020080.3 |
| Solyc11g020100.1 | Solyc11g020100.3 |
| Solyc11g020130.1 | Solyc11g020130.3 |
| Solyc11g020180.1 | Solyc11g020180.3 |
| Solyc11g020260.1 | Solyc11g020260.3 |
| Solyc11g020290.1 | Solyc11g020290.3 |
| Solyc11g020360.1 | Solyc11g020360.3 |
| Solyc11g020380.1 | Solyc11g020380.3 |
| Solyc11g020430.1 | Solyc11g020430.3 |
| Solyc11g020450.1 | Solyc11g020450.3 |
| Solyc11g020490.1 | Solyc11g020490.3 |
| Solyc11g020500.1 | Solyc11g020500.3 |
| Solyc11g020520.1 | Solyc11g020520.3 |
| Solyc11g020530.1 | Solyc11g020530.3 |
| Solyc11g020640.1 | Solyc11g020640.3 |
| Solyc11g020760.1 | Solyc11g020760.3 |
| Solyc11g020770.1 | Solyc11g020770.3 |
| Solyc11g020850.1 | Solyc11g020850.3 |
| Solyc11g020870.1 | Solyc11g020870.3 |
| Solyc11g020950.1 | Solyc11g020950.3 |
| Solyc11g020990.1 | Solyc11g020990.3 |
| Solyc11g021040.1 | Solyc11g021040.3 |
| Solyc11g021080.1 | Solyc11g021080.3 |
| Solyc11g021090.1 | Solyc11g021090.3 |
| Solyc11g021110.1 | Solyc11g021110.3 |
| Solyc11g021240.1 | Solyc11g021240.3 |
| Solyc11g021360.1 | Solyc11g021360.3 |
| Solyc11g022380.1 | Solyc11g022380.3 |
| Solyc11g022420.1 | Solyc11g022420.3 |
| Solyc11g022430.1 | Solyc11g022430.3 |
| Solyc11g022460.1 | Solyc11g022460.3 |
| Solyc11g022530.1 | Solyc11g022530.3 |
| Solyc11g022540.1 | Solyc11g022540.3 |
| Solyc11g027810.1 | Solyc11g027810.3 |
| Solyc11g027890.1 | Solyc11g027890.3 |
| Solyc11g027980.1 | Solyc11g027980.3 |
| Solyc11g028020.1 | Solyc11g028020.3 |
| Solyc11g028090.1 | Solyc11g028090.3 |
| Solyc11g028170.1 | Solyc11g028170.3 |
| Solyc11g028310.1 | Solyc11g028310.3 |

| | |
|---|---|
| Solyc11g028340.1 | Solyc11g028340.3 |
| Solyc11g030550.1 | Solyc11g030550.3 |
| Solyc11g030860.1 | Solyc11g030860.3 |
| Solyc11g030890.1 | Solyc11g030890.3 |
| Solyc11g030930.1 | Solyc11g030930.3 |
| Solyc11g030940.1 | Solyc11g030940.3 |
| Solyc11g032040.1 | Solyc11g032040.3 |
| Solyc11g032070.1 | Solyc11g032070.3 |
| Solyc11g032130.1 | Solyc11g032130.3 |
| Solyc11g032230.1 | Solyc11g032230.3 |
| Solyc11g033250.1 | Solyc11g033250.3 |
| Solyc11g039370.1 | Solyc11g039370.3 |
| Solyc11g039410.1 | Solyc11g039410.3 |
| Solyc11g039420.1 | Solyc11g039420.3 |
| Solyc11g039570.1 | Solyc11g039570.3 |
| Solyc11g039590.1 | Solyc11g039590.3 |
| Solyc11g039740.1 | Solyc11g039740.3 |
| Solyc11g039750.1 | Solyc11g039750.3 |
| Solyc11g039810.1 | Solyc11g039810.3 |
| Solyc11g039900.1 | Solyc11g039900.3 |
| Solyc11g039950.1 | Solyc11g039950.3 |
| Solyc11g039980.1 | Solyc11g039980.3 |
| Solyc11g040020.1 | Solyc11g040020.3 |
| Solyc11g040030.1 | Solyc11g040030.3 |
| Solyc11g040050.1 | Solyc11g040050.3 |
| Solyc11g040060.1 | Solyc11g040060.3 |
| Solyc11g040120.1 | Solyc11g040120.3 |
| Solyc11g040170.1 | Solyc11g040170.3 |
| Solyc11g040180.1 | Solyc11g040180.3 |
| Solyc11g040340.1 | Solyc11g040340.3 |
| Solyc11g042500.1 | Solyc11g042500.3 |
| Solyc11g042620.1 | Solyc11g042620.3 |
| Solyc11g042630.1 | Solyc11g042630.3 |
| Solyc11g042710.1 | Solyc11g042710.3 |
| Solyc11g042730.1 | Solyc11g042730.3 |
| Solyc11g042880.1 | Solyc11g042880.3 |
| Solyc11g042920.1 | Solyc11g042920.3 |
| Solyc11g043070.1 | Solyc11g043070.3 |
| Solyc11g043080.1 | Solyc11g043080.3 |
| Solyc11g043110.1 | Solyc11g043110.3 |
| Solyc11g043210.1 | Solyc11g043210.3 |
| Solyc11g044230.1 | Solyc11g044230.3 |
| Solyc11g044250.1 | Solyc11g044250.3 |

| | |
|---|---|
| Solyc11g044270.1 | Solyc11g044270.3 |
| Solyc11g044330.1 | Solyc11g044330.3 |
| Solyc11g044340.1 | Solyc11g044340.3 |
| Solyc11g044360.1 | Solyc11g044360.3 |
| Solyc11g044450.1 | Solyc11g044450.3 |
| Solyc11g044460.1 | Solyc11g044460.3 |
| Solyc11g044470.1 | Solyc11g044470.3 |
| Solyc11g044510.1 | Solyc11g044510.3 |
| Solyc11g044560.1 | Solyc11g044560.3 |
| Solyc11g044570.1 | Solyc11g044570.3 |
| Solyc11g044680.1 | Solyc11g044680.3 |
| Solyc11g044750.1 | Solyc11g044750.3 |
| Solyc11g044800.1 | Solyc11g044800.3 |
| Solyc11g044860.1 | Solyc11g044860.3 |
| Solyc11g044880.1 | Solyc11g044880.3 |
| Solyc11g044960.1 | Solyc11g044960.3 |
| Solyc11g045110.1 | Solyc11g045110.3 |
| Solyc11g045120.1 | Solyc11g045120.3 |
| Solyc11g045160.1 | Solyc11g045160.3 |
| Solyc11g045180.1 | Solyc11g045180.3 |
| Solyc11g045310.1 | Solyc11g045310.3 |
| Solyc11g045340.1 | Solyc11g045340.3 |
| Solyc11g045390.1 | Solyc11g045390.3 |
| Solyc11g045450.1 | Solyc11g045450.3 |
| Solyc11g045460.1 | Solyc11g045460.3 |
| Solyc11g045480.1 | Solyc11g045480.3 |
| Solyc11g045530.1 | Solyc11g045530.3 |
| Solyc11g045670.1 | Solyc11g045670.3 |
| Solyc11g045680.1 | Solyc11g045680.3 |
| Solyc11g045690.1 | Solyc11g045690.3 |
| Solyc11g051010.1 | Solyc11g051010.3 |
| Solyc11g051020.1 | Solyc11g051020.3 |
| Solyc11g051100.1 | Solyc11g051100.3 |
| Solyc11g051120.1 | Solyc11g051120.3 |
| Solyc11g056290.1 | Solyc11g056290.3 |
| Solyc11g061730.1 | Solyc11g061730.3 |
| Solyc11g061760.1 | Solyc11g061760.3 |
| Solyc11g061810.1 | Solyc11g061810.3 |
| Solyc11g061870.1 | Solyc11g061870.3 |
| Solyc11g061890.1 | Solyc11g061890.3 |
| Solyc11g061920.1 | Solyc11g061920.3 |
| Solyc11g062010.1 | Solyc11g062010.3 |
| Solyc11g062060.1 | Solyc11g062060.3 |

| | |
|---|---|
| Solyc11g062070.1 | Solyc11g062070.3 |
| Solyc11g062150.1 | Solyc11g062150.3 |
| Solyc11g062260.1 | Solyc11g062260.3 |
| Solyc11g062290.1 | Solyc11g062290.3 |
| Solyc11g062300.1 | Solyc11g062300.3 |
| Solyc11g062390.1 | Solyc11g062390.3 |
| Solyc11g062410.1 | Solyc11g062410.3 |
| Solyc11g063440.1 | Solyc11g063440.3 |
| Solyc11g063600.1 | Solyc11g063600.3 |
| Solyc11g063610.1 | Solyc11g063610.3 |
| Solyc11g063690.1 | Solyc11g063690.3 |
| Solyc11g063730.1 | Solyc11g063730.3 |
| Solyc11g063740.1 | Solyc11g063740.3 |
| Solyc11g064750.1 | Solyc11g064750.3 |
| Solyc11g064770.1 | Solyc11g064770.3 |
| Solyc11g064800.1 | Solyc11g064800.3 |
| Solyc11g064950.1 | Solyc11g064950.3 |
| Solyc11g065020.1 | Solyc11g065020.3 |
| Solyc11g065030.1 | Solyc11g065030.3 |
| Solyc11g065100.1 | Solyc11g065100.3 |
| Solyc11g065120.1 | Solyc11g065120.3 |
| Solyc11g065160.1 | Solyc11g065160.3 |
| Solyc11g065190.1 | Solyc11g065190.3 |
| Solyc11g065260.1 | Solyc11g065260.3 |
| Solyc11g065310.1 | Solyc11g065310.3 |
| Solyc11g065370.1 | Solyc11g065370.3 |
| Solyc11g065390.1 | Solyc11g065390.3 |
| Solyc11g065420.1 | Solyc11g065420.3 |
| Solyc11g065440.1 | Solyc11g065440.3 |
| Solyc11g065480.1 | Solyc11g065480.3 |
| Solyc11g065690.1 | Solyc11g065690.3 |
| Solyc11g065710.1 | Solyc11g065710.3 |
| Solyc11g065730.1 | Solyc11g065730.3 |
| Solyc11g065780.1 | Solyc11g065780.3 |
| Solyc11g065820.1 | Solyc11g065820.3 |
| Solyc11g065840.1 | Solyc11g065840.3 |
| Solyc11g065870.1 | Solyc11g065870.3 |
| Solyc11g065930.1 | Solyc11g065930.3 |
| Solyc11g065950.1 | Solyc11g065950.3 |
| Solyc11g065960.1 | Solyc11g065960.3 |
| Solyc11g065970.1 | Solyc11g065970.3 |
| Solyc11g065980.1 | Solyc11g065980.3 |
| Solyc11g066060.1 | Solyc11g066060.3 |

| | |
|---|---|
| Solyc11g066120.1 | Solyc11g066120.3 |
| Solyc11g066150.1 | Solyc11g066150.3 |
| Solyc11g066270.1 | Solyc11g066270.3 |
| Solyc11g066280.1 | Solyc11g066280.3 |
| Solyc11g066300.1 | Solyc11g066300.3 |
| Solyc11g066380.1 | Solyc11g066380.3 |
| Solyc11g066510.1 | Solyc11g066510.3 |
| Solyc11g066520.1 | Solyc11g066520.3 |
| Solyc11g066530.1 | Solyc11g066530.3 |
| Solyc11g066650.1 | Solyc11g066650.3 |
| Solyc11g066660.1 | Solyc11g066660.3 |
| Solyc11g066710.1 | Solyc11g066710.3 |
| Solyc11g066720.1 | Solyc11g066720.3 |
| Solyc11g066770.1 | Solyc11g066770.3 |
| Solyc11g066780.1 | Solyc11g066780.3 |
| Solyc11g066840.1 | Solyc11g066840.3 |
| Solyc11g066850.1 | Solyc11g066850.3 |
| Solyc11g066910.1 | Solyc11g066910.3 |
| Solyc11g066960.1 | Solyc11g066960.3 |
| Solyc11g066970.1 | Solyc11g066970.3 |
| Solyc11g067000.1 | Solyc11g067000.3 |
| Solyc11g067020.1 | Solyc11g067020.3 |
| Solyc11g067050.1 | Solyc11g067050.4 |
| Solyc11g067080.1 | Solyc11g067080.3 |
| Solyc11g067170.1 | Solyc11g067170.3 |
| Solyc11g067190.1 | Solyc11g067190.3 |
| Solyc11g067250.1 | Solyc11g067250.3 |
| Solyc11g067260.1 | Solyc11g067260.3 |
| Solyc11g067310.1 | Solyc11g067310.3 |
| Solyc11g068370.1 | Solyc11g068370.3 |
| Solyc11g068430.1 | Solyc11g068430.3 |
| Solyc11g068470.1 | Solyc11g068470.3 |
| Solyc11g068520.1 | Solyc11g068520.3 |
| Solyc11g068710.1 | Solyc11g068710.3 |
| Solyc11g068820.1 | Solyc11g068820.3 |
| Solyc11g068910.1 | Solyc11g068910.3 |
| Solyc11g068930.1 | Solyc11g068930.3 |
| Solyc11g068950.1 | Solyc11g068950.3 |
| Solyc11g068990.1 | Solyc11g068990.3 |
| Solyc11g069020.1 | Solyc11g069020.3 |
| Solyc11g069030.1 | Solyc11g069030.3 |
| Solyc11g069070.1 | Solyc11g069070.3 |
| Solyc11g069150.1 | Solyc11g069150.3 |

| | |
|---|---|
| Solyc11g069160.1 | Solyc11g069160.3 |
| Solyc11g069230.1 | Solyc11g069230.3 |
| Solyc11g069370.1 | Solyc11g069370.3 |
| Solyc11g069470.1 | Solyc11g069470.3 |
| Solyc11g069490.1 | Solyc11g069490.3 |
| Solyc11g069510.1 | Solyc11g069510.3 |
| Solyc11g069540.1 | Solyc11g069540.3 |
| Solyc11g069610.1 | Solyc11g069610.3 |
| Solyc11g069620.1 | Solyc11g069620.3 |
| Solyc11g069690.1 | Solyc11g069690.3 |
| Solyc11g069750.1 | Solyc11g069750.3 |
| Solyc11g069780.1 | Solyc11g069780.3 |
| Solyc11g069810.1 | Solyc11g069810.3 |
| Solyc11g069820.1 | Solyc11g069820.3 |
| Solyc11g069840.1 | Solyc11g069840.3 |
| Solyc11g069850.1 | Solyc11g069850.3 |
| Solyc11g069860.1 | Solyc11g069860.3 |
| Solyc11g069890.1 | Solyc11g069890.3 |
| Solyc11g069900.1 | Solyc11g069900.3 |
| Solyc11g069930.1 | Solyc11g069930.3 |
| Solyc11g070010.1 | Solyc11g070010.3 |
| Solyc11g070040.1 | Solyc11g070040.3 |
| Solyc11g070070.1 | Solyc11g070070.3 |
| Solyc11g070110.1 | Solyc11g070110.3 |
| Solyc11g071200.1 | Solyc11g071200.3 |
| Solyc11g071210.1 | Solyc11g071210.3 |
| Solyc11g071250.1 | Solyc11g071250.3 |
| Solyc11g071330.1 | Solyc11g071330.3 |
| Solyc11g071340.1 | Solyc11g071340.3 |
| Solyc11g071420.1 | Solyc11g071420.3 |
| Solyc11g071450.1 | Solyc11g071450.3 |
| Solyc11g071500.1 | Solyc11g071500.3 |
| Solyc11g071510.1 | Solyc11g071510.3 |
| Solyc11g071570.1 | Solyc11g071570.3 |
| Solyc11g071640.1 | Solyc11g071640.3 |
| Solyc11g071650.1 | Solyc11g071650.3 |
| Solyc11g071690.1 | Solyc11g071690.3 |
| Solyc11g071730.1 | Solyc11g071730.3 |
| Solyc11g071820.1 | Solyc11g071820.3 |
| Solyc11g071880.1 | Solyc11g071880.3 |
| Solyc11g071890.1 | Solyc11g071890.3 |
| Solyc11g071920.1 | Solyc11g071920.3 |
| Solyc11g072010.1 | Solyc11g072010.3 |

| | |
|---|---|
| Solyc11g072030.1 | Solyc11g072030.3 |
| Solyc11g072060.1 | Solyc11g072060.3 |
| Solyc11g072070.1 | Solyc11g072070.3 |
| Solyc11g072140.1 | Solyc11g072140.3 |
| Solyc11g072150.1 | Solyc11g072150.3 |
| Solyc11g072160.1 | Solyc11g072160.3 |
| Solyc11g072180.1 | Solyc11g072180.3 |
| Solyc11g072270.1 | Solyc11g072270.3 |
| Solyc11g072280.1 | Solyc11g072280.3 |
| Solyc11g072290.1 | Solyc11g072290.3 |
| Solyc11g072320.1 | Solyc11g072320.3 |
| Solyc11g072400.1 | Solyc11g072400.3 |
| Solyc11g072470.1 | Solyc11g072470.3 |
| Solyc11g072510.1 | Solyc11g072510.3 |
| Solyc11g072520.1 | Solyc11g072520.3 |
| Solyc11g072540.1 | Solyc11g072540.3 |
| Solyc11g072560.1 | Solyc11g072560.3 |
| Solyc11g072590.1 | Solyc11g072590.3 |
| Solyc11g072600.1 | Solyc11g072600.3 |
| Solyc11g072610.1 | Solyc11g072610.3 |
| Solyc11g072640.1 | Solyc11g072640.3 |
| Solyc11g072650.1 | Solyc11g072650.3 |
| Solyc11g072710.1 | Solyc11g072710.3 |
| Solyc11g072750.1 | Solyc11g072750.3 |
| Solyc11g072800.1 | Solyc11g072800.3 |
| Solyc11g072820.1 | Solyc11g072820.3 |
| Solyc11g072930.1 | Solyc11g072930.3 |
| Solyc11g072950.1 | Solyc11g072950.3 |
| Solyc11g073020.1 | Solyc11g073020.3 |
| Solyc11g073060.1 | Solyc11g073060.3 |
| Solyc11g073080.1 | Solyc11g073080.3 |
| Solyc11g073160.1 | Solyc11g073160.3 |
| Solyc11g073220.1 | Solyc11g073220.3 |
| Solyc12g005030.1 | Solyc12g005030.3 |
| Solyc12g005160.1 | Solyc12g005160.3 |
| Solyc12g005190.1 | Solyc12g005190.3 |
| Solyc12g005200.1 | Solyc12g005200.3 |
| Solyc12g005230.1 | Solyc12g005230.3 |
| Solyc12g005280.1 | Solyc12g005280.3 |
| Solyc12g005320.1 | Solyc12g005320.3 |
| Solyc12g005340.1 | Solyc12g005340.3 |
| Solyc12g005360.1 | Solyc12g005360.3 |
| Solyc12g005380.1 | Solyc12g005380.3 |

| | |
|---|---|
| Solyc12g005400.1 | Solyc12g005400.3 |
| Solyc12g005500.1 | Solyc12g005500.3 |
| Solyc12g005510.1 | Solyc12g005510.3 |
| Solyc12g005550.1 | Solyc12g005550.3 |
| Solyc12g005610.1 | Solyc12g005610.3 |
| Solyc12g005710.1 | Solyc12g005710.3 |
| Solyc12g005770.1 | Solyc12g005770.3 |
| Solyc12g005780.1 | Solyc12g005780.3 |
| Solyc12g005830.1 | Solyc12g005830.3 |
| Solyc12g005870.1 | Solyc12g005870.3 |
| Solyc12g005890.1 | Solyc12g005890.3 |
| Solyc12g005900.1 | Solyc12g005900.3 |
| Solyc12g005960.1 | Solyc12g005960.3 |
| Solyc12g006040.1 | Solyc12g006040.3 |
| Solyc12g006110.1 | Solyc12g006110.3 |
| Solyc12g006120.1 | Solyc12g006120.3 |
| Solyc12g006130.1 | Solyc12g006130.3 |
| Solyc12g006220.1 | Solyc12g006220.3 |
| Solyc12g006250.1 | Solyc12g006250.3 |
| Solyc12g006280.1 | Solyc12g006280.3 |
| Solyc12g006320.1 | Solyc12g006320.3 |
| Solyc12g006340.1 | Solyc12g006340.3 |
| Solyc12g006350.1 | Solyc12g006350.3 |
| Solyc12g006360.1 | Solyc12g006360.3 |
| Solyc12g006410.1 | Solyc12g006410.3 |
| Solyc12g006430.1 | Solyc12g006430.3 |
| Solyc12g006560.1 | Solyc12g006560.3 |
| Solyc12g006590.1 | Solyc12g006590.3 |
| Solyc12g006600.1 | Solyc12g006600.3 |
| Solyc12g006700.1 | Solyc12g006700.3 |
| Solyc12g006710.1 | Solyc12g006710.3 |
| Solyc12g006790.1 | Solyc12g006790.3 |
| Solyc12g007000.1 | Solyc12g007000.3 |
| Solyc12g007030.1 | Solyc12g007030.3 |
| Solyc12g007040.1 | Solyc12g007040.3 |
| Solyc12g007150.1 | Solyc12g007150.3 |
| Solyc12g007170.1 | Solyc12g007170.3 |
| Solyc12g007220.1 | Solyc12g007220.3 |
| Solyc12g007270.1 | Solyc12g007270.3 |
| Solyc12g007280.1 | Solyc12g007280.3 |
| Solyc12g007310.1 | Solyc12g007310.3 |
| Solyc12g008330.1 | Solyc12g008330.3 |
| Solyc12g008350.1 | Solyc12g008350.3 |

| | |
|---|---|
| Solyc12g008370.1 | Solyc12g008370.3 |
| Solyc12g008390.1 | Solyc12g008390.3 |
| Solyc12g008420.1 | Solyc12g008420.3 |
| Solyc12g008430.1 | Solyc12g008430.3 |
| Solyc12g008480.1 | Solyc12g008480.3 |
| Solyc12g008550.1 | Solyc12g008550.3 |
| Solyc12g008570.1 | Solyc12g008570.3 |
| Solyc12g008590.1 | Solyc12g008590.3 |
| Solyc12g008610.1 | Solyc12g008610.3 |
| Solyc12g008640.1 | Solyc12g008640.3 |
| Solyc12g008680.1 | Solyc12g008680.3 |
| Solyc12g008690.1 | Solyc12g008690.3 |
| Solyc12g008700.1 | Solyc12g008700.3 |
| Solyc12g008730.1 | Solyc12g008730.3 |
| Solyc12g008750.1 | Solyc12g008750.3 |
| Solyc12g008760.1 | Solyc12g008760.3 |
| Solyc12g008790.1 | Solyc12g008790.3 |
| Solyc12g008800.1 | Solyc12g008800.3 |
| Solyc12g008830.1 | Solyc12g008830.3 |
| Solyc12g008850.1 | Solyc12g008850.3 |
| Solyc12g008860.1 | Solyc12g008860.3 |
| Solyc12g008890.1 | Solyc12g008890.3 |
| Solyc12g008920.1 | Solyc12g008920.3 |
| Solyc12g008930.1 | Solyc12g008930.3 |
| Solyc12g008970.1 | Solyc12g008970.3 |
| Solyc12g008980.1 | Solyc12g008980.3 |
| Solyc12g009040.1 | Solyc12g009040.3 |
| Solyc12g009050.1 | Solyc12g009050.3 |
| Solyc12g009090.1 | Solyc12g009090.3 |
| Solyc12g009110.1 | Solyc12g009110.3 |
| Solyc12g009130.1 | Solyc12g009130.3 |
| Solyc12g009140.1 | Solyc12g009140.3 |
| Solyc12g009230.1 | Solyc12g009230.3 |
| Solyc12g009250.1 | Solyc12g009250.3 |
| Solyc12g009260.1 | Solyc12g009260.3 |
| Solyc12g009280.1 | Solyc12g009280.3 |
| Solyc12g009370.1 | Solyc12g009370.3 |
| Solyc12g009420.1 | Solyc12g009420.3 |
| Solyc12g009450.1 | Solyc12g009450.3 |
| Solyc12g009490.1 | Solyc12g009490.3 |
| Solyc12g009570.1 | Solyc12g009570.3 |
| Solyc12g009630.1 | Solyc12g009630.3 |
| Solyc12g009660.1 | Solyc12g009660.3 |

| | |
|---|---|
| Solyc12g009720.1 | Solyc12g009720.3 |
| Solyc12g009730.1 | Solyc12g009730.3 |
| Solyc12g009800.1 | Solyc12g009800.3 |
| Solyc12g009810.1 | Solyc12g009810.3 |
| Solyc12g009860.1 | Solyc12g009860.3 |
| Solyc12g009890.1 | Solyc12g009890.3 |
| Solyc12g009930.1 | Solyc12g009930.3 |
| Solyc12g010000.1 | Solyc12g010000.3 |
| Solyc12g010020.1 | Solyc12g010020.3 |
| Solyc12g010030.1 | Solyc12g010030.3 |
| Solyc12g010110.1 | Solyc12g010110.3 |
| Solyc12g010220.1 | Solyc12g010220.3 |
| Solyc12g010340.1 | Solyc12g010340.3 |
| Solyc12g010350.1 | Solyc12g010350.3 |
| Solyc12g010380.1 | Solyc12g010380.3 |
| Solyc12g010410.1 | Solyc12g010410.3 |
| Solyc12g010440.1 | Solyc12g010440.3 |
| Solyc12g010450.1 | Solyc12g010450.3 |
| Solyc12g010470.1 | Solyc12g010470.3 |
| Solyc12g010490.1 | Solyc12g010490.3 |
| Solyc12g010500.1 | Solyc12g010500.3 |
| Solyc12g010520.1 | Solyc12g010520.3 |
| Solyc12g010530.1 | Solyc12g010530.3 |
| Solyc12g010560.1 | Solyc12g010560.3 |
| Solyc12g010590.1 | Solyc12g010590.3 |
| Solyc12g010600.1 | Solyc12g010600.3 |
| Solyc12g010610.1 | Solyc12g010610.3 |
| Solyc12g010680.1 | Solyc12g010680.3 |
| Solyc12g010740.1 | Solyc12g010740.3 |
| Solyc12g010870.1 | Solyc12g010870.3 |
| Solyc12g010980.1 | Solyc12g010980.3 |
| Solyc12g010990.1 | Solyc12g010990.3 |
| Solyc12g011030.1 | Solyc12g011030.3 |
| Solyc12g011060.1 | Solyc12g011060.3 |
| Solyc12g011070.1 | Solyc12g011070.3 |
| Solyc12g011080.1 | Solyc12g011080.3 |
| Solyc12g011090.1 | Solyc12g011090.3 |
| Solyc12g011120.1 | Solyc12g011120.3 |
| Solyc12g011140.1 | Solyc12g011140.3 |
| Solyc12g011170.1 | Solyc12g011170.3 |
| Solyc12g011180.1 | Solyc12g011180.3 |
| Solyc12g011200.1 | Solyc12g011200.3 |
| Solyc12g011330.2 | Solyc12g011330.4 |

| | |
|---|---|
| Solyc12g011350.1 | Solyc12g011350.3 |
| Solyc12g011370.1 | Solyc12g011370.3 |
| Solyc12g011400.1 | Solyc12g011400.3 |
| Solyc12g011430.1 | Solyc12g011430.3 |
| Solyc12g013470.1 | Solyc12g013470.3 |
| Solyc12g013480.1 | Solyc12g013480.3 |
| Solyc12g013520.1 | Solyc12g013520.3 |
| Solyc12g013580.1 | Solyc12g013580.3 |
| Solyc12g013600.1 | Solyc12g013600.3 |
| Solyc12g013660.1 | Solyc12g013660.3 |
| Solyc12g013680.1 | Solyc12g013680.3 |
| Solyc12g013730.1 | Solyc12g013730.3 |
| Solyc12g013740.1 | Solyc12g013740.3 |
| Solyc12g013770.1 | Solyc12g013770.3 |
| Solyc12g013780.1 | Solyc12g013780.3 |
| Solyc12g013790.1 | Solyc12g013790.3 |
| Solyc12g013800.1 | Solyc12g013800.3 |
| Solyc12g013820.1 | Solyc12g013820.3 |
| Solyc12g013840.1 | Solyc12g013840.3 |
| Solyc12g013870.1 | Solyc12g013870.3 |
| Solyc12g013900.1 | Solyc12g013900.3 |
| Solyc12g013960.1 | Solyc12g013960.3 |
| Solyc12g013970.1 | Solyc12g013970.3 |
| Solyc12g013990.1 | Solyc12g013990.3 |
| Solyc12g014000.1 | Solyc12g014000.3 |
| Solyc12g014120.1 | Solyc12g014120.3 |
| Solyc12g014140.1 | Solyc12g014140.3 |
| Solyc12g014150.1 | Solyc12g014150.3 |
| Solyc12g014160.1 | Solyc12g014160.3 |
| Solyc12g014200.1 | Solyc12g014200.3 |
| Solyc12g014210.1 | Solyc12g014210.3 |
| Solyc12g014240.1 | Solyc12g014240.3 |
| Solyc12g014270.1 | Solyc12g014270.3 |
| Solyc12g014300.1 | Solyc12g014300.3 |
| Solyc12g014350.1 | Solyc12g014350.3 |
| Solyc12g014380.1 | Solyc12g014380.3 |
| Solyc12g014390.1 | Solyc12g014390.3 |
| Solyc12g014400.1 | Solyc12g014400.3 |
| Solyc12g014490.1 | Solyc12g014490.3 |
| Solyc12g014500.1 | Solyc12g014500.3 |
| Solyc12g014530.1 | Solyc12g014530.3 |
| Solyc12g014560.1 | Solyc12g014560.3 |
| Solyc12g014590.1 | Solyc12g014590.3 |

| | |
|---|---|
| Solyc12g015650.1 | Solyc12g015650.3 |
| Solyc12g015690.1 | Solyc12g015690.3 |
| Solyc12g015700.1 | Solyc12g015700.3 |
| Solyc12g015770.1 | Solyc12g015770.3 |
| Solyc12g015810.1 | Solyc12g015810.3 |
| Solyc12g015840.1 | Solyc12g015840.3 |
| Solyc12g015970.1 | Solyc12g015970.3 |
| Solyc12g016010.1 | Solyc12g016010.3 |
| Solyc12g016020.1 | Solyc12g016020.3 |
| Solyc12g016030.1 | Solyc12g016030.3 |
| Solyc12g016040.1 | Solyc12g016040.3 |
| Solyc12g016120.1 | Solyc12g016120.3 |
| Solyc12g016220.1 | Solyc12g016220.3 |
| Solyc12g017310.1 | Solyc12g017310.3 |
| Solyc12g017360.1 | Solyc12g017360.3 |
| Solyc12g017370.1 | Solyc12g017370.3 |
| Solyc12g017380.1 | Solyc12g017380.3 |
| Solyc12g017390.1 | Solyc12g017390.3 |
| Solyc12g017400.1 | Solyc12g017400.3 |
| Solyc12g017430.1 | Solyc12g017430.3 |
| Solyc12g017480.1 | Solyc12g017480.3 |
| Solyc12g017660.1 | Solyc12g017660.3 |
| Solyc12g017670.1 | Solyc12g017670.3 |
| Solyc12g017690.1 | Solyc12g017690.3 |
| Solyc12g017800.1 | Solyc12g017800.3 |
| Solyc12g017830.1 | Solyc12g017830.3 |
| Solyc12g017860.1 | Solyc12g017860.3 |
| Solyc12g017880.1 | Solyc12g017880.3 |
| Solyc12g017900.1 | Solyc12g017900.3 |
| Solyc12g017920.1 | Solyc12g017920.3 |
| Solyc12g019080.1 | Solyc12g019080.3 |
| Solyc12g019110.1 | Solyc12g019110.3 |
| Solyc12g019130.1 | Solyc12g019130.3 |
| Solyc12g019140.1 | Solyc12g019140.3 |
| Solyc12g019240.1 | Solyc12g019240.3 |
| Solyc12g019330.1 | Solyc12g019330.3 |
| Solyc12g019410.1 | Solyc12g019410.3 |
| Solyc12g019420.1 | Solyc12g019420.3 |
| Solyc12g019430.1 | Solyc12g019430.3 |
| Solyc12g019440.1 | Solyc12g019440.3 |
| Solyc12g019480.1 | Solyc12g019480.3 |
| Solyc12g019610.1 | Solyc12g019610.3 |
| Solyc12g019750.1 | Solyc12g019750.3 |

| | |
|---|---|
| Solyc12g021250.1 | Solyc12g021250.3 |
| Solyc12g021310.1 | Solyc12g021310.3 |
| Solyc12g021340.1 | Solyc12g021340.3 |
| Solyc12g021360.1 | Solyc12g021360.3 |
| Solyc12g026430.1 | Solyc12g026430.3 |
| Solyc12g027540.1 | Solyc12g027540.3 |
| Solyc12g027580.1 | Solyc12g027580.3 |
| Solyc12g027770.1 | Solyc12g027770.3 |
| Solyc12g027850.1 | Solyc12g027850.3 |
| Solyc12g027880.1 | Solyc12g027880.3 |
| Solyc12g032920.1 | Solyc12g032920.3 |
| Solyc12g032930.1 | Solyc12g032930.3 |
| Solyc12g032950.1 | Solyc12g032950.3 |
| Solyc12g035170.1 | Solyc12g035170.3 |
| Solyc12g035250.1 | Solyc12g035250.3 |
| Solyc12g035310.1 | Solyc12g035310.3 |
| Solyc12g035420.1 | Solyc12g035420.3 |
| Solyc12g035440.1 | Solyc12g035440.3 |
| Solyc12g035460.1 | Solyc12g035460.3 |
| Solyc12g035500.1 | Solyc12g035500.3 |
| Solyc12g035520.1 | Solyc12g035520.3 |
| Solyc12g035670.1 | Solyc12g035670.3 |
| Solyc12g035820.1 | Solyc12g035820.3 |
| Solyc12g035870.1 | Solyc12g035870.3 |
| Solyc12g035890.1 | Solyc12g035890.3 |
| Solyc12g036090.1 | Solyc12g036090.3 |
| Solyc12g036160.1 | Solyc12g036160.3 |
| Solyc12g036220.1 | Solyc12g036220.3 |
| Solyc12g036230.1 | Solyc12g036230.3 |
| Solyc12g036300.1 | Solyc12g036300.3 |
| Solyc12g036320.1 | Solyc12g036320.3 |
| Solyc12g036410.1 | Solyc12g036410.3 |
| Solyc12g036430.1 | Solyc12g036430.3 |
| Solyc12g036450.1 | Solyc12g036450.3 |
| Solyc12g036490.1 | Solyc12g036490.3 |
| Solyc12g036520.1 | Solyc12g036520.3 |
| Solyc12g036550.1 | Solyc12g036550.3 |
| Solyc12g036710.1 | Solyc12g036710.3 |
| Solyc12g036810.1 | Solyc12g036810.3 |
| Solyc12g036860.1 | Solyc12g036860.3 |
| Solyc12g036870.1 | Solyc12g036870.3 |
| Solyc12g036910.1 | Solyc12g036910.3 |
| Solyc12g038020.1 | Solyc12g038020.3 |

| | |
|---|---|
| Solyc12g038120.1 | Solyc12g038120.3 |
| Solyc12g038200.1 | Solyc12g038200.3 |
| Solyc12g038220.1 | Solyc12g038220.3 |
| Solyc12g038430.1 | Solyc12g038430.3 |
| Solyc12g038520.1 | Solyc12g038520.3 |
| Solyc12g038570.1 | Solyc12g038570.3 |
| Solyc12g038810.1 | Solyc12g038810.3 |
| Solyc12g038870.1 | Solyc12g038870.3 |
| Solyc12g038950.1 | Solyc12g038950.3 |
| Solyc12g038970.1 | Solyc12g038970.3 |
| Solyc12g039050.1 | Solyc12g039050.3 |
| Solyc12g039080.1 | Solyc12g039080.3 |
| Solyc12g040280.1 | Solyc12g040280.3 |
| Solyc12g040300.1 | Solyc12g040300.3 |
| Solyc12g040390.1 | Solyc12g040390.3 |
| Solyc12g040740.1 | Solyc12g040740.3 |
| Solyc12g040870.1 | Solyc12g040870.3 |
| Solyc12g041870.1 | Solyc12g041870.3 |
| Solyc12g041950.1 | Solyc12g041950.3 |
| Solyc12g041980.1 | Solyc12g041980.3 |
| Solyc12g042000.1 | Solyc12g042000.3 |
| Solyc12g042020.1 | Solyc12g042020.3 |
| Solyc12g042030.1 | Solyc12g042030.3 |
| Solyc12g042060.1 | Solyc12g042060.3 |
| Solyc12g042070.1 | Solyc12g042070.3 |
| Solyc12g042110.1 | Solyc12g042110.3 |
| Solyc12g042170.1 | Solyc12g042170.3 |
| Solyc12g042230.1 | Solyc12g042230.3 |
| Solyc12g042250.1 | Solyc12g042250.3 |
| Solyc12g042260.1 | Solyc12g042260.3 |
| Solyc12g042270.1 | Solyc12g042270.3 |
| Solyc12g042280.1 | Solyc12g042280.3 |
| Solyc12g042360.1 | Solyc12g042360.3 |
| Solyc12g042370.1 | Solyc12g042370.3 |
| Solyc12g042480.1 | Solyc12g042480.3 |
| Solyc12g042520.1 | Solyc12g042520.3 |
| Solyc12g042560.1 | Solyc12g042560.3 |
| Solyc12g042690.1 | Solyc12g042690.3 |
| Solyc12g042770.1 | Solyc12g042770.3 |
| Solyc12g042830.1 | Solyc12g042830.3 |
| Solyc12g042860.1 | Solyc12g042860.3 |
| Solyc12g042880.1 | Solyc12g042880.3 |
| Solyc12g043000.1 | Solyc12g043000.3 |

| | |
|---|---|
| Solyc12g043030.1 | Solyc12g043030.3 |
| Solyc12g043090.1 | Solyc12g043090.3 |
| Solyc12g043150.1 | Solyc12g043150.3 |
| Solyc12g043160.1 | Solyc12g043160.3 |
| Solyc12g044180.1 | Solyc12g044180.3 |
| Solyc12g044190.1 | Solyc12g044190.3 |
| Solyc12g044200.1 | Solyc12g044200.3 |
| Solyc12g044220.1 | Solyc12g044220.3 |
| Solyc12g044240.1 | Solyc12g044240.3 |
| Solyc12g044370.1 | Solyc12g044370.3 |
| Solyc12g044380.1 | Solyc12g044380.3 |
| Solyc12g044390.1 | Solyc12g044390.3 |
| Solyc12g044400.1 | Solyc12g044400.3 |
| Solyc12g044530.1 | Solyc12g044530.3 |
| Solyc12g044540.1 | Solyc12g044540.3 |
| Solyc12g044580.1 | Solyc12g044580.3 |
| Solyc12g044650.1 | Solyc12g044650.3 |
| Solyc12g044660.1 | Solyc12g044660.3 |
| Solyc12g044690.1 | Solyc12g044690.3 |
| Solyc12g044750.1 | Solyc12g044750.3 |
| Solyc12g044800.1 | Solyc12g044800.3 |
| Solyc12g044810.1 | Solyc12g044810.3 |
| Solyc12g044820.1 | Solyc12g044820.3 |
| Solyc12g044860.1 | Solyc12g044860.3 |
| Solyc12g044900.1 | Solyc12g044900.3 |
| Solyc12g044920.1 | Solyc12g044920.3 |
| Solyc12g044930.1 | Solyc12g044930.3 |
| Solyc12g044950.1 | Solyc12g044950.3 |
| Solyc12g049080.1 | Solyc12g049080.3 |
| Solyc12g049090.1 | Solyc12g049090.3 |
| Solyc12g049100.1 | Solyc12g049100.3 |
| Solyc12g049190.1 | Solyc12g049190.3 |
| Solyc12g049220.1 | Solyc12g049220.3 |
| Solyc12g049270.1 | Solyc12g049270.3 |
| Solyc12g049280.1 | Solyc12g049280.3 |
| Solyc12g049320.1 | Solyc12g049320.3 |
| Solyc12g049340.1 | Solyc12g049340.3 |
| Solyc12g049410.1 | Solyc12g049410.3 |
| Solyc12g049440.1 | Solyc12g049440.3 |
| Solyc12g049470.1 | Solyc12g049470.3 |
| Solyc12g049500.1 | Solyc12g049500.3 |
| Solyc12g049540.1 | Solyc12g049540.3 |
| Solyc12g049570.1 | Solyc12g049570.3 |

| | |
|---|---|
| Solyc12g055730.1 | Solyc12g055730.3 |
| Solyc12g055750.1 | Solyc12g055750.3 |
| Solyc12g055780.1 | Solyc12g055780.3 |
| Solyc12g055820.1 | Solyc12g055820.3 |
| Solyc12g055880.1 | Solyc12g055880.3 |
| Solyc12g055890.1 | Solyc12g055890.3 |
| Solyc12g055910.1 | Solyc12g055910.3 |
| Solyc12g055920.1 | Solyc12g055920.3 |
| Solyc12g055930.1 | Solyc12g055930.3 |
| Solyc12g055970.1 | Solyc12g055970.3 |
| Solyc12g056100.1 | Solyc12g056100.3 |
| Solyc12g056180.1 | Solyc12g056180.3 |
| Solyc12g056210.1 | Solyc12g056210.3 |
| Solyc12g056280.1 | Solyc12g056280.3 |
| Solyc12g056300.1 | Solyc12g056300.3 |
| Solyc12g056400.1 | Solyc12g056400.3 |
| Solyc12g056460.1 | Solyc12g056460.3 |
| Solyc12g056510.1 | Solyc12g056510.3 |
| Solyc12g056520.1 | Solyc12g056520.3 |
| Solyc12g056610.1 | Solyc12g056610.3 |
| Solyc12g056630.1 | Solyc12g056630.3 |
| Solyc12g056660.1 | Solyc12g056660.3 |
| Solyc12g056710.1 | Solyc12g056710.3 |
| Solyc12g056750.1 | Solyc12g056750.3 |
| Solyc12g056770.1 | Solyc12g056770.3 |
| Solyc12g056920.1 | Solyc12g056920.3 |
| Solyc12g057010.1 | Solyc12g057010.3 |
| Solyc12g062470.1 | Solyc12g062470.3 |
| Solyc12g062480.1 | Solyc12g062480.3 |
| Solyc12g062530.1 | Solyc12g062530.3 |
| Solyc12g062540.1 | Solyc12g062540.3 |
| Solyc12g062680.1 | Solyc12g062680.3 |
| Solyc12g062720.1 | Solyc12g062720.3 |
| Solyc12g062730.1 | Solyc12g062730.3 |
| Solyc12g062820.1 | Solyc12g062820.3 |
| Solyc12g062930.1 | Solyc12g062930.3 |
| Solyc12g062940.1 | Solyc12g062940.3 |
| Solyc12g063010.1 | Solyc12g063010.3 |
| Solyc12g063050.1 | Solyc12g063050.3 |
| Solyc12g070200.1 | Solyc12g070200.3 |
| Solyc12g077370.1 | Solyc12g077370.3 |
| Solyc12g077390.1 | Solyc12g077390.3 |
| Solyc12g077540.1 | Solyc12g077540.3 |

| | |
|---|---|
| Solyc12g077660.1 | Solyc12g077660.3 |
| Solyc12g082730.1 | Solyc12g082730.3 |
| Solyc12g082770.1 | Solyc12g082770.3 |
| Solyc12g087810.1 | Solyc12g087810.3 |
| Solyc12g087850.1 | Solyc12g087850.3 |
| Solyc12g087860.1 | Solyc12g087860.3 |
| Solyc12g087890.1 | Solyc12g087890.3 |
| Solyc12g087980.1 | Solyc12g087980.3 |
| Solyc12g088090.1 | Solyc12g088090.3 |
| Solyc12g088110.1 | Solyc12g088110.3 |
| Solyc12g088130.1 | Solyc12g088130.3 |
| Solyc12g088160.1 | Solyc12g088160.3 |
| Solyc12g088200.1 | Solyc12g088200.3 |
| Solyc12g088210.1 | Solyc12g088210.3 |
| Solyc12g088230.1 | Solyc12g088230.3 |
| Solyc12g088280.1 | Solyc12g088280.3 |
| Solyc12g088310.1 | Solyc12g088310.3 |
| Solyc12g088460.1 | Solyc12g088460.3 |
| Solyc12g088470.1 | Solyc12g088470.3 |
| Solyc12g088540.1 | Solyc12g088540.3 |
| Solyc12g088570.1 | Solyc12g088570.3 |
| Solyc12g088790.1 | Solyc12g088790.3 |
| Solyc12g088800.1 | Solyc12g088800.3 |
| Solyc12g088810.1 | Solyc12g088810.3 |
| Solyc12g088830.1 | Solyc12g088830.3 |
| Solyc12g088890.1 | Solyc12g088890.3 |
| Solyc12g088910.1 | Solyc12g088910.3 |
| Solyc12g088940.1 | Solyc12g088940.3 |
| Solyc12g088950.1 | Solyc12g088950.3 |
| Solyc12g089010.1 | Solyc12g089010.3 |
| Solyc12g089040.1 | Solyc12g089040.3 |
| Solyc12g089050.1 | Solyc12g089050.3 |
| Solyc12g089080.1 | Solyc12g089080.3 |
| Solyc12g089160.1 | Solyc12g089160.3 |
| Solyc12g089170.1 | Solyc12g089170.3 |
| Solyc12g089200.1 | Solyc12g089200.3 |
| Solyc12g089220.1 | Solyc12g089220.3 |
| Solyc12g089260.1 | Solyc12g089260.3 |
| Solyc12g089290.1 | Solyc12g089290.3 |
| Solyc12g089320.1 | Solyc12g089320.3 |
| Solyc12g089350.1 | Solyc12g089350.3 |
| Solyc12g094410.1 | Solyc12g094410.3 |
| Solyc12g094450.1 | Solyc12g094450.3 |

| | |
|---|---|
| Solyc12g094470.1 | Solyc12g094470.3 |
| Solyc12g094500.1 | Solyc12g094500.3 |
| Solyc12g094600.1 | Solyc12g094600.3 |
| Solyc12g094610.1 | Solyc12g094610.3 |
| Solyc12g094620.1 | Solyc12g094620.3 |
| Solyc12g094640.1 | Solyc12g094640.3 |
| Solyc12g094650.1 | Solyc12g094650.3 |
| Solyc12g094680.1 | Solyc12g094680.3 |
| Solyc12g094700.1 | Solyc12g094700.3 |
| Solyc12g094720.1 | Solyc12g094720.3 |
| Solyc12g094740.1 | Solyc12g094740.3 |
| Solyc12g095740.1 | Solyc12g095740.3 |
| Solyc12g095820.1 | Solyc12g095820.3 |
| Solyc12g095830.1 | Solyc12g095830.3 |
| Solyc12g095870.1 | Solyc12g095870.3 |
| Solyc12g095960.1 | Solyc12g095960.3 |
| Solyc12g095970.1 | Solyc12g095970.3 |
| Solyc12g096010.1 | Solyc12g096010.3 |
| Solyc12g096020.1 | Solyc12g096020.3 |
| Solyc12g096040.1 | Solyc12g096040.3 |
| Solyc12g096110.1 | Solyc12g096110.3 |
| Solyc12g096140.1 | Solyc12g096140.3 |
| Solyc12g096240.1 | Solyc12g096240.3 |
| Solyc12g096290.1 | Solyc12g096290.3 |
| Solyc12g096330.1 | Solyc12g096330.3 |
| Solyc12g096460.1 | Solyc12g096460.3 |
| Solyc12g096510.1 | Solyc12g096510.3 |
| Solyc12g096520.1 | Solyc12g096520.3 |
| Solyc12g096590.1 | Solyc12g096590.3 |
| Solyc12g096640.1 | Solyc12g096640.3 |
| Solyc12g096660.1 | Solyc12g096660.3 |
| Solyc12g096840.1 | Solyc12g096840.3 |
| Solyc12g096850.1 | Solyc12g096850.3 |
| Solyc12g096860.1 | Solyc12g096860.3 |
| Solyc12g096880.1 | Solyc12g096880.3 |
| Solyc12g096940.1 | Solyc12g096940.3 |
| Solyc12g096980.1 | Solyc12g096980.3 |
| Solyc12g097070.1 | Solyc12g097070.3 |
| Solyc12g097090.1 | Solyc12g097090.3 |
| Solyc12g098100.1 | Solyc12g098100.3 |
| Solyc12g098120.1 | Solyc12g098120.3 |
| Solyc12g098180.1 | Solyc12g098180.3 |
| Solyc12g098210.1 | Solyc12g098210.3 |

| | |
|---|---|
| Solyc12g098280.1 | Solyc12g098280.3 |
| Solyc12g098320.1 | Solyc12g098320.3 |
| Solyc12g098330.1 | Solyc12g098330.3 |
| Solyc12g098380.1 | Solyc12g098380.3 |
| Solyc12g098390.1 | Solyc12g098390.3 |
| Solyc12g098420.1 | Solyc12g098420.3 |
| Solyc12g098590.1 | Solyc12g098590.3 |
| Solyc12g098640.1 | Solyc12g098640.3 |
| Solyc12g098660.1 | Solyc12g098660.3 |
| Solyc12g098670.1 | Solyc12g098670.3 |
| Solyc12g098680.1 | Solyc12g098680.3 |
| Solyc12g098720.1 | Solyc12g098720.3 |
| Solyc12g098730.1 | Solyc12g098730.3 |
| Solyc12g098840.1 | Solyc12g098840.3 |
| Solyc12g098870.1 | Solyc12g098870.3 |
| Solyc12g098930.1 | Solyc12g098930.3 |
| Solyc12g099000.1 | Solyc12g099000.3 |
| Solyc12g099010.1 | Solyc12g099010.3 |
| Solyc12g099080.1 | Solyc12g099080.3 |
| Solyc12g099090.1 | Solyc12g099090.3 |
| Solyc12g099120.1 | Solyc12g099120.3 |
| Solyc12g099170.1 | Solyc12g099170.3 |
| Solyc12g099180.1 | Solyc12g099180.3 |
| Solyc12g099340.1 | Solyc12g099340.3 |
| Solyc12g099380.1 | Solyc12g099380.3 |
| Solyc12g099440.1 | Solyc12g099440.3 |
| Solyc12g099450.1 | Solyc12g099450.3 |
| Solyc12g099510.1 | Solyc12g099510.3 |
| Solyc12g099520.1 | Solyc12g099520.3 |
| Solyc12g099530.1 | Solyc12g099530.3 |
| Solyc12g099570.1 | Solyc12g099570.3 |
| Solyc12g099700.1 | Solyc12g099700.3 |
| Solyc12g099730.1 | Solyc12g099730.3 |
| Solyc12g099790.1 | Solyc12g099790.3 |
| Solyc12g099870.1 | Solyc12g099870.3 |
| Solyc12g099950.1 | Solyc12g099950.3 |
| Solyc12g099980.1 | Solyc12g099980.3 |
| Solyc12g100000.1 | Solyc12g100000.3 |
| Solyc12g100070.1 | Solyc12g100070.3 |
| Solyc12g100140.1 | Solyc12g100140.3 |
| Solyc12g100160.1 | Solyc12g100160.3 |
| Solyc12g100250.1 | Solyc12g100250.3 |
| Solyc12g100280.1 | Solyc12g100280.3 |

| | |
|---|---|
| Solyc12g100290.1 | Solyc12g100290.3 |
| Solyc01g005070.2 | Solyc01g005070.3 |
| Solyc01g005480.2 | Solyc01g005480.3 |
| Solyc01g005530.2 | Solyc01g005530.3 |
| Solyc01g005930.2 | Solyc01g005930.3 |
| Solyc01g006700.2 | Solyc01g006700.3 |
| Solyc01g006790.2 | Solyc01g006790.3 |
| Solyc01g007140.2 | Solyc01g007140.3 |
| Solyc01g007870.2 | Solyc01g007870.3 |
| Solyc01g008780.2 | Solyc01g008780.3 |
| Solyc01g008810.2 | Solyc01g008810.3 |
| Solyc01g009250.2 | Solyc01g009250.3 |
| Solyc01g009430.2 | Solyc01g009430.3 |
| Solyc01g010280.2 | Solyc01g010280.3 |
| Solyc01g011333.1 | Solyc01g150114.1 |
| Solyc01g020320.2 | Solyc01g020320.3 |
| Solyc01g021640.2 | Solyc01g021640.3 |
| Solyc01g034070.1 | Solyc01g034070.2 |
| Solyc01g056520.2 | Solyc01g056520.3 |
| Solyc01g056590.1 | Solyc01g056590.2 |
| Solyc01g056990.2 | Solyc01g056990.3 |
| Solyc01g060450.2 | Solyc01g060450.3 |
| Solyc01g066440.2 | Solyc01g066440.3 |
| Solyc01g066860.2 | Solyc01g066860.3 |
| Solyc01g067080.2 | Solyc01g067080.3 |
| Solyc01g067280.2 | Solyc01g067280.3 |
| Solyc01g067820.2 | Solyc01g067820.3 |
| Solyc01g068190.2 | Solyc01g068190.3 |
| Solyc01g068540.2 | Solyc01g068540.3 |
| Solyc01g079080.2 | Solyc01g079080.3 |
| Solyc01g079630.2 | Solyc01g079630.3 |
| Solyc01g079830.2 | Solyc01g079830.3 |
| Solyc01g080100.2 | Solyc01g080100.3 |
| Solyc01g080510.2 | Solyc01g080510.3 |
| Solyc01g080590.2 | Solyc01g080590.3 |
| Solyc01g081200.2 | Solyc01g081200.3 |
| Solyc01g086860.2 | Solyc01g086860.3 |
| Solyc01g087580.2 | Solyc01g087580.3 |
| Solyc01g087600.2 | Solyc01g087600.3 |
| Solyc01g087880.2 | Solyc01g087880.3 |
| Solyc01g088140.2 | Solyc01g088140.3 |
| Solyc01g090260.4 | Solyc01g090260.5.1 |
| Solyc01g090380.2 | Solyc01g090380.3 |

| | |
|---|---|
| Solyc01g090680.2 | Solyc01g090680.3 |
| Solyc01g090720.2 | Solyc01g090720.3 |
| Solyc01g090740.2 | Solyc01g090740.3 |
| Solyc01g090900.2 | Solyc01g090900.3 |
| Solyc01g091260.2 | Solyc01g091260.3 |
| Solyc01g091270.2 | Solyc01g091270.3 |
| Solyc01g091830.2 | Solyc01g091830.3 |
| Solyc01g094630.2 | Solyc01g094630.3 |
| Solyc01g095020.2 | Solyc01g095020.3 |
| Solyc01g095420.2 | Solyc01g095420.3 |
| Solyc01g096660.2 | Solyc01g096660.3 |
| Solyc01g096920.2 | Solyc01g096920.3 |
| Solyc01g098520.2 | Solyc01g098520.3 |
| Solyc01g098830.2 | Solyc01g098830.3 |
| Solyc01g098870.2 | Solyc01g098870.3 |
| Solyc01g099700.2 | Solyc01g099700.3 |
| Solyc01g099730.2 | Solyc01g099730.3 |
| Solyc01g099930.2 | Solyc01g099930.3 |
| Solyc01g100480.2 | Solyc01g100480.3 |
| Solyc01g100690.2 | Solyc01g100690.3 |
| Solyc01g102440.2 | Solyc01g102440.3 |
| Solyc01g102450.2 | Solyc01g102450.3 |
| Solyc01g102480.2 | Solyc01g102480.3 |
| Solyc01g102500.2 | Solyc01g102500.3 |
| Solyc01g103760.2 | Solyc01g103760.3 |
| Solyc01g104350.2 | Solyc01g104350.3 |
| Solyc01g104540.2 | Solyc01g104540.3 |
| Solyc01g104560.2 | Solyc01g104560.3 |
| Solyc01g104685.1 | Solyc01g150164.1 |
| Solyc01g104710.2 | Solyc01g104710.3 |
| Solyc01g104720.2 | Solyc01g104720.3 |
| Solyc01g104910.2 | Solyc01g104910.3 |
| Solyc01g105290.2 | Solyc01g105290.3 |
| Solyc01g105530.2 | Solyc01g105530.3 |
| Solyc01g105600.2 | Solyc01g105600.3 |
| Solyc01g105980.2 | Solyc01g105980.3 |
| Solyc01g106070.1 | Solyc01g106070.2 |
| Solyc01g106580.2 | Solyc01g106580.3 |
| Solyc01g106950.2 | Solyc01g106950.3 |
| Solyc01g107030.2 | Solyc01g107030.3 |
| Solyc01g107100.2 | Solyc01g107100.3 |
| Solyc01g107280.2 | Solyc01g107280.3 |
| Solyc01g108100.2 | Solyc01g108100.3 |

| | |
|---|---|
| Solyc01g108620.2 | Solyc01g108620.3 |
| Solyc01g108650.2 | Solyc01g108650.3 |
| Solyc01g108890.2 | Solyc01g108890.3 |
| Solyc01g109010.2 | Solyc01g109010.3 |
| Solyc01g109270.2 | Solyc01g109270.3 |
| Solyc01g109680.2 | Solyc01g109680.3 |
| Solyc01g109780.2 | Solyc01g109780.3 |
| Solyc01g111110.2 | Solyc01g111110.3 |
| Solyc01g112010.2 | Solyc01g112010.3 |
| Solyc01g112370.2 | Solyc01g112370.3 |
| Solyc02g021050.2 | Solyc02g021050.3 |
| Solyc02g027080.1 | Solyc02g027080.2 |
| Solyc02g031950.2 | Solyc02g031950.3 |
| Solyc02g033060.2 | Solyc02g033060.3 |
| Solyc02g036460.2 | Solyc02g036460.3 |
| Solyc02g062170.2 | Solyc02g062170.3 |
| Solyc02g062530.2 | Solyc02g062530.3 |
| Solyc02g062770.1 | Solyc02g062770.2 |
| Solyc02g063050.2 | Solyc02g063050.3 |
| Solyc02g063300.2 | Solyc02g063300.3 |
| Solyc02g064710.2 | Solyc02g064710.3 |
| Solyc02g065370.2 | Solyc02g065370.3 |
| Solyc02g068290.1 | Solyc02g068290.2 |
| Solyc02g068400.2 | Solyc02g068400.3 |
| Solyc02g069820.2 | Solyc02g069820.3 |
| Solyc02g071460.1 | Solyc02g071460.2 |
| Solyc02g072040.2 | Solyc02g072040.3 |
| Solyc02g072050.1 | Solyc02g072050.2 |
| Solyc02g072170.2 | Solyc02g072170.3 |
| Solyc02g072510.2 | Solyc02g072510.3 |
| Solyc02g077230.2 | Solyc02g077230.3 |
| Solyc02g078280.2 | Solyc02g078280.3 |
| Solyc02g078430.2 | Solyc02g078430.3 |
| Solyc02g080110.2 | Solyc02g080110.3 |
| Solyc02g080500.2 | Solyc02g080500.3 |
| Solyc02g080780.2 | Solyc02g080780.3 |
| Solyc02g082710.2 | Solyc02g082710.3 |
| Solyc02g083380.2 | Solyc02g083380.3 |
| Solyc02g083500.2 | Solyc02g083500.3 |
| Solyc02g083540.2 | Solyc02g083540.3 |
| Solyc02g083650.2 | Solyc02g083650.3 |
| Solyc02g084220.2 | Solyc02g084220.3 |
| Solyc02g084550.2 | Solyc02g084550.3 |

| | |
|---|---|
| Solyc02g084810.2 | Solyc02g084810.3 |
| Solyc02g085000.2 | Solyc02g085000.3 |
| Solyc02g085920.2 | Solyc02g085920.3 |
| Solyc02g086110.2 | Solyc02g086110.3 |
| Solyc02g086140.1 | Solyc02g086140.2 |
| Solyc02g086170.2 | Solyc02g086170.3 |
| Solyc02g087150.2 | Solyc02g087150.3 |
| Solyc02g087450.2 | Solyc02g087450.3 |
| Solyc02g088260.2 | Solyc02g088260.3 |
| Solyc02g088270.2 | Solyc02g088270.3 |
| Solyc02g088480.2 | Solyc02g088480.3 |
| Solyc02g089100.2 | Solyc02g089100.3 |
| Solyc02g089580.2 | Solyc02g089580.3 |
| Solyc02g089650.2 | Solyc02g089650.3 |
| Solyc02g089660.1 | Solyc02g089660.2 |
| Solyc02g089780.2 | Solyc02g089780.3 |
| Solyc02g089960.2 | Solyc02g089960.3 |
| Solyc02g089980.2 | Solyc02g089980.3 |
| Solyc02g090580.2 | Solyc02g090580.3 |
| Solyc02g091390.2 | Solyc02g091390.3 |
| Solyc02g091940.2 | Solyc02g091940.3 |
| Solyc02g092000.2 | Solyc02g092000.3 |
| Solyc02g092200.2 | Solyc02g092200.3 |
| Solyc02g092240.2 | Solyc02g092240.3 |
| Solyc02g092390.2 | Solyc02g092390.3 |
| Solyc02g092590.2 | Solyc02g092590.3 |
| Solyc02g092700.2 | Solyc02g092700.3 |
| Solyc02g093350.2 | Solyc02g093350.3 |
| Solyc02g093450.2 | Solyc02g093450.3 |
| Solyc02g093790.2 | Solyc02g093790.3 |
| Solyc03g005030.2 | Solyc03g005030.3 |
| Solyc03g005100.2 | Solyc03g005100.3 |
| Solyc03g005110.2 | Solyc03g005110.3 |
| Solyc03g006010.2 | Solyc03g006010.3 |
| Solyc03g006540.2 | Solyc03g006540.3 |
| Solyc03g006590.2 | Solyc03g006590.3 |
| Solyc03g006850.2 | Solyc03g006850.3 |
| Solyc03g006900.1 | Solyc03g006900.2 |
| Solyc03g007420.2 | Solyc03g007420.3 |
| Solyc03g007520.2 | Solyc03g007520.3 |
| Solyc03g007540.2 | Solyc03g007540.3 |
| Solyc03g007710.2 | Solyc03g007710.3 |
| Solyc03g007720.2 | Solyc03g007720.3 |

| | |
|---|---|
| Solyc03g007775.1 | Solyc03g007775.2 |
| Solyc03g025460.2 | Solyc03g025460.3 |
| Solyc03g025530.2 | Solyc03g025530.3 |
| Solyc03g025910.2 | Solyc03g025910.3 |
| Solyc03g031680.2 | Solyc03g031680.3 |
| Solyc03g032100.1 | Solyc03g032100.2 |
| Solyc03g032120.2 | Solyc03g032120.3 |
| Solyc03g033650.2 | Solyc03g033650.3 |
| Solyc03g034230.2 | Solyc03g034230.3 |
| Solyc03g043870.2 | Solyc03g043870.3 |
| Solyc03g044155.1 | Solyc03g150114.1 |
| Solyc03g044630.1 | Solyc03g044630.2 |
| Solyc03g045090.2 | Solyc03g045090.3 |
| Solyc03g046260.1 | Solyc03g046260.2 |
| Solyc03g058340.2 | Solyc03g058340.3 |
| Solyc03g062900.2 | Solyc03g062900.3 |
| Solyc03g078160.2 | Solyc03g078160.3 |
| Solyc03g080170.2 | Solyc03g080170.3 |
| Solyc03g081310.2 | Solyc03g081310.3 |
| Solyc03g083030.2 | Solyc03g083030.3 |
| Solyc03g083050.2 | Solyc03g083050.3 |
| Solyc03g083060.2 | Solyc03g083060.3 |
| Solyc03g093220.2 | Solyc03g093220.3 |
| Solyc03g093480.2 | Solyc03g093480.3 |
| Solyc03g093980.1 | Solyc03g093980.2 |
| Solyc03g096960.2 | Solyc03g096960.3 |
| Solyc03g097040.1 | Solyc03g097040.2 |
| Solyc03g097180.2 | Solyc03g097180.3 |
| Solyc03g097200.2 | Solyc03g097200.3 |
| Solyc03g098120.2 | Solyc03g098120.3 |
| Solyc03g098130.2 | Solyc03g098130.3 |
| Solyc03g020020.2 | Solyc03g020020.3 |
| Solyc03g019910.2 | Solyc03g019910.3 |
| Solyc03g111030.2 | Solyc03g111030.3 |
| Solyc03g111660.2 | Solyc03g111660.3 |
| Solyc03g112670.2 | Solyc03g112670.3 |
| Solyc03g112760.2 | Solyc03g112760.3 |
| Solyc03g112840.1 | Solyc03g112840.2 |
| Solyc03g113090.2 | Solyc03g113090.3 |
| Solyc03g113510.1 | Solyc03g113510.2 |
| Solyc03g115060.2 | Solyc03g115060.3 |
| Solyc03g115620.1 | Solyc03g115620.2 |
| Solyc03g116330.2 | Solyc03g116330.3 |

| | |
|---|---|
| Solyc03g116400.2 | Solyc03g116400.3 |
| Solyc03g116470.1 | Solyc03g116470.2 |
| Solyc03g116630.2 | Solyc03g116630.3 |
| Solyc03g116660.2 | Solyc03g116660.3 |
| Solyc03g116770.2 | Solyc03g116770.3 |
| Solyc03g117090.2 | Solyc03g117090.3 |
| Solyc03g117580.2 | Solyc03g117580.3 |
| Solyc03g117740.2 | Solyc03g117740.3 |
| Solyc03g118050.2 | Solyc03g118050.3 |
| Solyc03g118060.2 | Solyc03g118060.3 |
| Solyc03g118630.2 | Solyc03g118630.3 |
| Solyc03g118700.2 | Solyc03g118700.3 |
| Solyc03g119940.2 | Solyc03g119940.3 |
| Solyc03g120300.2 | Solyc03g120300.3 |
| Solyc03g120410.1 | Solyc03g120410.3 |
| Solyc03g120480.2 | Solyc03g120480.3 |
| Solyc03g120490.2 | Solyc03g120490.3 |
| Solyc03g120830.2 | Solyc03g120830.3 |
| Solyc03g120950.2 | Solyc03g120950.3 |
| Solyc03g121520.2 | Solyc03g121520.3 |
| Solyc03g121560.2 | Solyc03g121560.3 |
| Solyc03g121850.2 | Solyc03g121850.3 |
| Solyc04g005120.2 | Solyc04g005120.3 |
| Solyc04g005290.2 | Solyc04g005290.3 |
| Solyc04g005500.2 | Solyc04g005500.3 |
| Solyc04g005530.2 | Solyc04g005530.3 |
| Solyc04g007700.2 | Solyc04g007700.3 |
| Solyc04g007730.2 | Solyc04g007730.3 |
| Solyc04g007840.2 | Solyc04g007840.3 |
| Solyc04g008560.2 | Solyc04g008560.3 |
| Solyc04g008800.2 | Solyc04g008800.3 |
| Solyc04g009570.2 | Solyc04g009570.3 |
| Solyc04g009600.2 | Solyc04g009600.3 |
| Solyc04g010155.1 | Solyc04g010155.2 |
| Solyc04g010220.2 | Solyc04g010220.3 |
| Solyc04g011910.2 | Solyc04g011910.3 |
| Solyc04g014710.2 | Solyc04g014710.3 |
| Solyc04g014850.2 | Solyc04g014850.3 |
| Solyc04g015400.2 | Solyc04g015400.3 |
| Solyc04g015480.2 | Solyc04g015480.3 |
| Solyc04g015585.1 | Solyc04g150111.1 |
| Solyc04g015710.2 | Solyc04g015710.3 |
| Solyc04g015740.2 | Solyc04g015740.3 |

| | |
|---|---|
| Solyc04g016295.1 | Solyc04g150113.1 |
| Solyc04g016420.2 | Solyc04g016420.3 |
| Solyc04g025475.1 | Solyc04g150119.1 |
| Solyc04g024520.2 | Solyc04g024520.3 |
| Solyc04g024500.2 | Solyc04g024500.3 |
| Solyc04g045620.2 | Solyc04g045620.3 |
| Solyc04g049290.1 | Solyc04g049290.2 |
| Solyc04g049860.1 | Solyc04g049860.2 |
| Solyc04g050790.2 | Solyc04g050790.3 |
| Solyc04g050920.2 | Solyc04g050920.3 |
| Solyc04g051205.1 | Solyc04g150139.1 |
| Solyc04g051710.1 | Solyc04g051710.2 |
| Solyc04g051850.2 | Solyc04g051850.3 |
| Solyc04g054460.2 | Solyc04g054460.3 |
| Solyc04g054475.1 | Solyc04g150148.1 |
| Solyc04g056610.2 | Solyc04g056610.3 |
| Solyc04g057870.2 | Solyc04g057870.3 |
| Solyc04g057970.2 | Solyc04g057970.3 |
| Solyc04g064930.2 | Solyc04g064930.3 |
| Solyc04g071180.2 | Solyc04g071180.3 |
| Solyc04g071310.2 | Solyc04g071310.3 |
| Solyc04g071420.2 | Solyc04g071420.3 |
| Solyc04g071630.2 | Solyc04g071630.3 |
| Solyc04g071660.2 | Solyc04g071660.3 |
| Solyc04g074100.2 | Solyc04g074100.3 |
| Solyc04g074720.2 | Solyc04g074720.3 |
| Solyc04g074760.2 | Solyc04g074760.3 |
| Solyc04g076010.2 | Solyc04g076010.3 |
| Solyc04g076150.2 | Solyc04g076150.3 |
| Solyc04g076260.1 | Solyc04g076260.2 |
| Solyc04g076340.2 | Solyc04g076340.3 |
| Solyc04g076410.2 | Solyc04g076410.3 |
| Solyc04g076550.2 | Solyc04g076550.3 |
| Solyc04g077450.2 | Solyc04g077450.3 |
| Solyc04g077560.2 | Solyc04g077560.3 |
| Solyc04g077690.2 | Solyc04g077690.3 |
| Solyc04g078030.2 | Solyc04g078030.3 |
| Solyc04g078600.2 | Solyc04g078600.3 |
| Solyc04g078880.2 | Solyc04g078880.3 |
| Solyc04g079140.2 | Solyc04g079140.3 |
| Solyc04g079540.2 | Solyc04g079540.3 |
| Solyc04g079740.2 | Solyc04g079740.3 |
| Solyc04g080950.2 | Solyc04g080950.3 |

| | |
|---|---|
| Solyc04g081450.2 | Solyc04g081450.3 |
| Solyc04g081540.2 | Solyc04g081540.3 |
| Solyc04g081730.2 | Solyc04g081730.3 |
| Solyc04g082210.2 | Solyc04g082210.3 |
| Solyc04g082310.2 | Solyc04g082310.3 |
| Solyc04g082590.2 | Solyc04g082590.3 |
| Solyc04g082640.2 | Solyc04g082640.3 |
| Solyc04g082800.2 | Solyc04g082800.3 |
| Solyc04g083015.1 | Solyc04g150173.1 |
| Solyc05g006110.2 | Solyc05g006110.3 |
| Solyc05g006580.2 | Solyc05g006580.3 |
| Solyc05g006900.1 | Solyc05g006900.2 |
| Solyc05g006960.2 | Solyc05g006960.3 |
| Solyc05g007440.2 | Solyc05g007440.3 |
| Solyc05g009210.1 | Solyc05g009210.2 |
| Solyc05g009520.1 | Solyc05g009520.2 |
| Solyc05g012600.2 | Solyc05g012600.3 |
| Solyc05g012640.2 | Solyc05g012640.3 |
| Solyc05g012680.2 | Solyc05g012680.3 |
| Solyc05g012820.1 | Solyc05g012820.2 |
| Solyc05g012850.2 | Solyc05g012850.3 |
| Solyc05g013780.2 | Solyc05g013780.3 |
| Solyc05g014310.2 | Solyc05g014310.3 |
| Solyc05g014320.2 | Solyc05g014320.3 |
| Solyc05g015180.1 | Solyc05g015180.2 |
| Solyc05g015640.2 | Solyc05g015640.3 |
| Solyc05g015710.2 | Solyc05g015710.3 |
| Solyc05g015980.2 | Solyc05g015980.3 |
| Solyc05g016185.1 | Solyc05g150121.1 |
| Solyc05g018490.2 | Solyc05g018490.3 |
| Solyc05g024430.2 | Solyc05g024430.3 |
| Solyc05g026540.2 | Solyc05g026540.3 |
| Solyc05g046270.2 | Solyc05g046270.3 |
| Solyc05g046310.2 | Solyc05g046310.3 |
| Solyc05g046390.2 | Solyc05g046390.3 |
| Solyc05g048750.2 | Solyc05g048750.3 |
| Solyc05g049880.2 | Solyc05g049880.3 |
| Solyc05g049940.2 | Solyc05g049940.3 |
| Solyc05g050000.2 | Solyc05g050000.3 |
| Solyc05g051370.2 | Solyc05g051370.3 |
| Solyc05g051770.2 | Solyc05g051770.3 |
| Solyc05g052210.2 | Solyc05g052210.3 |
| Solyc05g052460.2 | Solyc05g052460.3 |

| | |
|---|---|
| Solyc05g053360.2 | Solyc05g053360.3 |
| Solyc05g053730.2 | Solyc05g053730.3 |
| Solyc05g054770.2 | Solyc05g054770.3 |
| Solyc05g055550.2 | Solyc05g055550.3 |
| Solyc05g056380.2 | Solyc05g056380.3 |
| Solyc05g056510.2 | Solyc05g056510.3 |
| Solyc06g005250.2 | Solyc06g005250.3 |
| Solyc06g005340.2 | Solyc06g005340.3 |
| Solyc06g005600.2 | Solyc06g005600.3 |
| Solyc06g007600.2 | Solyc06g007600.3 |
| Solyc06g008560.1 | Solyc06g008560.2 |
| Solyc06g009120.2 | Solyc06g009120.3 |
| Solyc06g009380.2 | Solyc06g009380.3 |
| Solyc06g011560.2 | Solyc06g011560.3 |
| Solyc06g036030.1 | Solyc06g036030.2 |
| Solyc06g036250.1 | Solyc06g036250.2 |
| Solyc06g036670.1 | Solyc06g036670.2 |
| Solyc06g036780.2 | Solyc06g036780.3 |
| Solyc06g042980.2 | Solyc06g042980.3 |
| Solyc06g042950.2 | Solyc06g042950.3 |
| Solyc06g048850.2 | Solyc06g048850.3 |
| Solyc06g052060.2 | Solyc06g052060.3 |
| Solyc06g053440.2 | Solyc06g053440.3 |
| Solyc06g054590.2 | Solyc06g054590.3 |
| Solyc06g059860.2 | Solyc06g059860.3 |
| Solyc06g060460.2 | Solyc06g060460.3 |
| Solyc06g060560.2 | Solyc06g060560.3 |
| Solyc06g060820.1 | Solyc06g060820.2 |
| Solyc06g060860.1 | Solyc06g060860.2 |
| Solyc06g060920.2 | Solyc06g060920.3 |
| Solyc06g061140.2 | Solyc06g061140.3 |
| Solyc06g063130.2 | Solyc06g063130.3 |
| Solyc06g063230.2 | Solyc06g063230.3 |
| Solyc06g063310.1 | Solyc06g063310.2 |
| Solyc06g063320.2 | Solyc06g063320.3 |
| Solyc06g064810.2 | Solyc06g064810.3 |
| Solyc06g065200.2 | Solyc06g065200.4 |
| Solyc06g065825.1 | Solyc06g150128.1 |
| Solyc06g065870.2 | Solyc06g065870.3 |
| Solyc06g066290.2 | Solyc06g066290.3 |
| Solyc06g066800.1 | Solyc06g066800.2 |
| Solyc06g068110.2 | Solyc06g068110.3 |
| Solyc06g068180.2 | Solyc06g068180.3 |

| | |
|---|---|
| Solyc06g068660.2 | Solyc06g068660.3 |
| Solyc06g069030.2 | Solyc06g069030.3 |
| Solyc06g069510.2 | Solyc06g069510.3 |
| Solyc06g070970.2 | Solyc06g070970.3 |
| Solyc06g071140.2 | Solyc06g071140.3 |
| Solyc06g071250.2 | Solyc06g071250.4 |
| Solyc06g071540.2 | Solyc06g071540.3 |
| Solyc06g072135.1 | Solyc06g150131.1 |
| Solyc06g072840.2 | Solyc06g072840.3 |
| Solyc06g072930.2 | Solyc06g072930.4 |
| Solyc06g073490.1 | Solyc06g073490.2 |
| Solyc06g073990.1 | Solyc06g073990.2 |
| Solyc06g074380.2 | Solyc06g074380.3 |
| Solyc06g074620.2 | Solyc06g074620.3 |
| Solyc06g074690.2 | Solyc06g074690.3 |
| Solyc06g075100.2 | Solyc06g075100.3 |
| Solyc06g076010.2 | Solyc06g076010.3 |
| Solyc06g082830.2 | Solyc06g082830.3 |
| Solyc06g083370.2 | Solyc06g083370.3 |
| Solyc06g083560.2 | Solyc06g083560.3 |
| Solyc06g083610.2 | Solyc06g083610.3 |
| Solyc06g084200.1 | Solyc06g084200.2 |
| Solyc06g084500.2 | Solyc06g084500.3 |
| Solyc07g005410.2 | Solyc07g005410.3 |
| Solyc07g005430.2 | Solyc07g005430.3 |
| Solyc07g005690.2 | Solyc07g005690.3 |
| Solyc07g006040.2 | Solyc07g006040.3 |
| Solyc07g006150.2 | Solyc07g006150.3 |
| Solyc07g006490.2 | Solyc07g006490.3 |
| Solyc07g007170.2 | Solyc07g007170.3 |
| Solyc07g007940.2 | Solyc07g007940.3 |
| Solyc07g008430.1 | Solyc07g008430.2 |
| Solyc07g008780.2 | Solyc07g008780.3 |
| Solyc07g014600.1 | Solyc07g014600.2 |
| Solyc07g017410.2 | Solyc07g017410.3 |
| Solyc07g025380.2 | Solyc07g025380.3 |
| Solyc07g032260.2 | Solyc07g032260.3 |
| Solyc07g032787.1 | Solyc07g150128.1 |
| Solyc07g032790.1 | Solyc07g150129.1 |
| Solyc07g038100.2 | Solyc07g038100.3 |
| Solyc07g038190.2 | Solyc07g038190.3 |
| Solyc07g039380.2 | Solyc07g039380.3 |
| Solyc07g041160.2 | Solyc07g041160.3 |

| | |
|---|---|
| Solyc07g042100.2 | Solyc07g042100.3 |
| Solyc07g042450.2 | Solyc07g042450.3 |
| Solyc07g042710.2 | Solyc07g042710.3 |
| Solyc07g043070.2 | Solyc07g043070.3 |
| Solyc07g043320.2 | Solyc07g043320.3 |
| Solyc07g045150.2 | Solyc07g045150.3 |
| Solyc07g045170.2 | Solyc07g045170.3 |
| Solyc07g049350.2 | Solyc07g049350.3 |
| Solyc07g052360.1 | Solyc07g052360.2 |
| Solyc07g052420.2 | Solyc07g052420.3 |
| Solyc07g052460.2 | Solyc07g052460.3 |
| Solyc07g053525.1 | Solyc07g150145.1 |
| Solyc07g053560.2 | Solyc07g053560.3 |
| Solyc07g054060.2 | Solyc07g054060.3 |
| Solyc07g054430.2 | Solyc07g054430.3 |
| Solyc07g054850.2 | Solyc07g054850.3 |
| Solyc07g055110.2 | Solyc07g055110.3 |
| Solyc07g056240.2 | Solyc07g056240.3 |
| Solyc07g062250.2 | Solyc07g062250.3 |
| Solyc07g062640.2 | Solyc07g062640.3 |
| Solyc07g063060.1 | Solyc07g063060.2 |
| Solyc07g063250.2 | Solyc07g063250.3 |
| Solyc07g063530.2 | Solyc07g063530.3 |
| Solyc07g063540.2 | Solyc07g063540.3 |
| Solyc07g063870.2 | Solyc07g063870.3 |
| Solyc07g064300.2 | Solyc07g064300.3 |
| Solyc07g064420.2 | Solyc07g064420.3 |
| Solyc07g064440.2 | Solyc07g064440.3 |
| Solyc07g064570.2 | Solyc07g064570.3 |
| Solyc07g065480.2 | Solyc07g065480.3 |
| Solyc07g065740.2 | Solyc07g065740.3 |
| Solyc07g066340.2 | Solyc07g066340.3 |
| Solyc07g066370.2 | Solyc07g066370.3 |
| Solyc07g066520.2 | Solyc07g066520.3 |
| Solyc07g066640.2 | Solyc07g066640.3 |
| Solyc08g005015.1 | Solyc08g150100.1 |
| Solyc08g005060.2 | Solyc08g005060.3 |
| Solyc08g005560.2 | Solyc08g005560.3 |
| Solyc08g006000.2 | Solyc08g006000.3 |
| Solyc08g006420.2 | Solyc08g006420.3 |
| Solyc08g006480.2 | Solyc08g006480.3 |
| Solyc08g006610.2 | Solyc08g006610.3 |
| Solyc08g006620.2 | Solyc08g006620.3 |

| | |
|---|---|
| Solyc08g007030.2 | Solyc08g007030.3 |
| Solyc08g007490.2 | Solyc08g007490.3 |
| Solyc08g007500.2 | Solyc08g007500.3 |
| Solyc08g007710.2 | Solyc08g007710.3 |
| Solyc08g008330.2 | Solyc08g008330.3 |
| Solyc08g008350.2 | Solyc08g008350.3 |
| Solyc08g015620.2 | Solyc08g015620.3 |
| Solyc08g042140.2 | Solyc08g042140.3 |
| Solyc08g041770.2 | Solyc08g041770.3 |
| Solyc08g041710.2 | Solyc08g041710.3 |
| Solyc08g023270.2 | Solyc08g023270.3 |
| Solyc08g022120.2 | Solyc08g022120.3 |
| Solyc08g059700.1 | Solyc08g059700.2 |
| Solyc08g061800.2 | Solyc08g061800.3 |
| Solyc08g065450.1 | Solyc08g065450.2 |
| Solyc08g065980.2 | Solyc08g065980.3 |
| Solyc08g067270.2 | Solyc08g067270.3 |
| Solyc08g067630.2 | Solyc08g067630.3 |
| Solyc08g067860.1 | Solyc08g067860.2 |
| Solyc08g067870.1 | Solyc08g067870.2 |
| Solyc08g067885.1 | Solyc08g150138.1 |
| Solyc08g068010.1 | Solyc08g068010.2 |
| Solyc08g068040.2 | Solyc08g068040.3 |
| Solyc08g068060.2 | Solyc08g068060.3 |
| Solyc08g068350.2 | Solyc08g068350.3 |
| Solyc08g068930.2 | Solyc08g068930.3 |
| Solyc08g069230.2 | Solyc08g069230.3 |
| Solyc08g074400.1 | Solyc08g074400.2 |
| Solyc08g075070.2 | Solyc08g075070.3 |
| Solyc08g075150.2 | Solyc08g075150.3 |
| Solyc08g075290.1 | Solyc08g075290.2 |
| Solyc08g075770.2 | Solyc08g075770.3 |
| Solyc08g075860.2 | Solyc08g075860.3 |
| Solyc08g076130.2 | Solyc08g076130.3 |
| Solyc08g076180.2 | Solyc08g076180.3 |
| Solyc08g076190.2 | Solyc08g076190.3 |
| Solyc08g076520.2 | Solyc08g076520.3 |
| Solyc08g076580.2 | Solyc08g076580.3 |
| Solyc08g077040.2 | Solyc08g077040.3 |
| Solyc08g077430.2 | Solyc08g077430.3 |
| Solyc08g078440.2 | Solyc08g078440.3 |
| Solyc08g078480.2 | Solyc08g078480.3 |
| Solyc08g078500.2 | Solyc08g078500.3 |

| | |
|---|---|
| Solyc08g078560.2 | Solyc08g078560.3 |
| Solyc08g078730.2 | Solyc08g078730.3 |
| Solyc08g079600.2 | Solyc08g079600.3 |
| Solyc08g080280.2 | Solyc08g080280.3 |
| Solyc08g081710.2 | Solyc08g081710.3 |
| Solyc08g082160.2 | Solyc08g082160.3 |
| Solyc08g082230.2 | Solyc08g082230.3 |
| Solyc08g082320.2 | Solyc08g082320.3 |
| Solyc08g082390.2 | Solyc08g082390.3 |
| Solyc08g082420.2 | Solyc08g082420.3 |
| Solyc08g083010.2 | Solyc08g083010.3 |
| Solyc08g083200.2 | Solyc08g083200.3 |
| Solyc09g005140.1 | Solyc09g005140.2 |
| Solyc09g005200.2 | Solyc09g005200.3 |
| Solyc09g005570.2 | Solyc09g005570.3 |
| Solyc09g006000.2 | Solyc09g006000.3 |
| Solyc09g007240.2 | Solyc09g007240.3 |
| Solyc09g007480.2 | Solyc09g007480.3 |
| Solyc09g007800.2 | Solyc09g007800.3 |
| Solyc09g008260.2 | Solyc09g008260.3 |
| Solyc09g008270.2 | Solyc09g008270.3 |
| Solyc09g008980.1 | Solyc09g008980.2 |
| Solyc09g009450.1 | Solyc09g009450.2 |
| Solyc09g010410.2 | Solyc09g010410.3 |
| Solyc09g014215.1 | Solyc09g014215.2 |
| Solyc09g015420.2 | Solyc09g015420.3 |
| Solyc09g015450.2 | Solyc09g015450.3 |
| Solyc09g015530.2 | Solyc09g015530.3 |
| Solyc09g018670.2 | Solyc09g018670.3 |
| Solyc09g018890.2 | Solyc09g018890.3 |
| Solyc09g018950.2 | Solyc09g018950.3 |
| Solyc09g042480.1 | Solyc09g042480.2 |
| Solyc09g057590.2 | Solyc09g057590.3 |
| Solyc09g058990.2 | Solyc09g058990.3 |
| Solyc09g059890.1 | Solyc09g059890.2 |
| Solyc09g059950.2 | Solyc09g059950.3 |
| Solyc09g060100.2 | Solyc09g060100.3 |
| Solyc09g061420.2 | Solyc09g061420.3 |
| Solyc09g061500.1 | Solyc09g061500.2 |
| Solyc09g064860.2 | Solyc09g064860.3 |
| Solyc09g065380.1 | Solyc09g065380.2 |
| Solyc09g065510.2 | Solyc09g065510.3 |
| Solyc09g072580.2 | Solyc09g072580.3 |

| | |
|---|---|
| Solyc09g072990.2 | Solyc09g072990.3 |
| Solyc09g074620.2 | Solyc09g074620.3 |
| Solyc09g075660.2 | Solyc09g075660.3 |
| Solyc09g082180.2 | Solyc09g082180.3 |
| Solyc09g083070.2 | Solyc09g083070.3 |
| Solyc09g090120.2 | Solyc09g090120.3 |
| Solyc09g090290.2 | Solyc09g090290.3 |
| Solyc09g090530.3 | Solyc09g090530.4 |
| Solyc09g090540.2 | Solyc09g090540.3 |
| Solyc09g090560.1 | Solyc09g090560.2 |
| Solyc09g091010.2 | Solyc09g091010.3 |
| Solyc09g092220.2 | Solyc09g092220.3 |
| Solyc09g092510.2 | Solyc09g092510.3 |
| Solyc10g005230.2 | Solyc10g005230.3 |
| Solyc10g005470.2 | Solyc10g005470.3 |
| Solyc10g005580.2 | Solyc10g005580.3 |
| Solyc10g005610.1 | Solyc10g005610.2 |
| Solyc10g005700.2 | Solyc10g005700.3 |
| Solyc10g006360.1 | Solyc10g006360.2 |
| Solyc10g007340.2 | Solyc10g007340.3 |
| Solyc10g007490.1 | Solyc10g007490.2 |
| Solyc10g007730.2 | Solyc10g007730.3 |
| Solyc10g008050.1 | Solyc10g008050.2 |
| Solyc10g008130.2 | Solyc10g008130.3 |
| Solyc10g008420.2 | Solyc10g008420.3 |
| Solyc10g008540.2 | Solyc10g008540.3 |
| Solyc10g008600.2 | Solyc10g008600.3 |
| Solyc10g008730.2 | Solyc10g008730.3 |
| Solyc10g008850.1 | Solyc10g008850.2 |
| Solyc10g009320.2 | Solyc10g009320.3 |
| Solyc10g009400.2 | Solyc10g009400.3 |
| Solyc10g012280.1 | Solyc10g012280.2 |
| Solyc10g012400.2 | Solyc10g012400.3 |
| Solyc10g012410.2 | Solyc10g012410.3 |
| Solyc10g018640.1 | Solyc10g018640.2 |
| Solyc10g039380.1 | Solyc10g039380.2 |
| Solyc10g044620.1 | Solyc10g044620.2 |
| Solyc10g045350.1 | Solyc10g045350.2 |
| Solyc10g045630.1 | Solyc10g045630.2 |
| Solyc10g047220.1 | Solyc10g047220.2 |
| Solyc10g048060.1 | Solyc10g048060.2 |
| Solyc10g050220.1 | Solyc10g050220.2 |
| Solyc10g051030.1 | Solyc10g051030.2 |

| | |
|---|---|
| Solyc10g054060.1 | Solyc10g054060.2 |
| Solyc10g054930.1 | Solyc10g054930.2 |
| Solyc10g074580.1 | Solyc10g074580.2 |
| Solyc10g074690.1 | Solyc10g074690.2 |
| Solyc10g076420.1 | Solyc10g076420.2 |
| Solyc10g076930.1 | Solyc10g076930.2 |
| Solyc10g078420.1 | Solyc10g078420.2 |
| Solyc10g080400.1 | Solyc10g080400.2 |
| Solyc10g081480.1 | Solyc10g081480.2 |
| Solyc10g081670.1 | Solyc10g081670.2 |
| Solyc10g083200.1 | Solyc10g083200.2 |
| Solyc10g083530.1 | Solyc10g083530.2 |
| Solyc10g083580.1 | Solyc10g083580.2 |
| Solyc10g083990.1 | Solyc10g083990.2 |
| Solyc10g084060.1 | Solyc10g084060.2 |
| Solyc10g084230.1 | Solyc10g084230.2 |
| Solyc10g084265.1 | Solyc10g150149.1 |
| Solyc10g084420.1 | Solyc10g084420.2 |
| Solyc10g085300.1 | Solyc10g085300.2 |
| Solyc10g085690.1 | Solyc10g085690.2 |
| Solyc10g086360.1 | Solyc10g086360.2 |
| Solyc10g086470.1 | Solyc10g086470.2 |
| Solyc10g086570.2 | Solyc10g086570.3 |
| Solyc11g005000.1 | Solyc11g005000.2 |
| Solyc11g005570.1 | Solyc11g005570.2 |
| Solyc11g005810.1 | Solyc11g005810.2 |
| Solyc11g005930.1 | Solyc11g005930.2 |
| Solyc11g006080.1 | Solyc11g006080.2 |
| Solyc11g006960.1 | Solyc11g006960.2 |
| Solyc11g007060.1 | Solyc11g007060.2 |
| Solyc11g007510.1 | Solyc11g007510.2 |
| Solyc11g007680.1 | Solyc11g007680.2 |
| Solyc11g008910.1 | Solyc11g008910.2 |
| Solyc11g009060.1 | Solyc11g009060.2 |
| Solyc11g011220.1 | Solyc11g011220.2 |
| Solyc11g012060.1 | Solyc11g012060.2 |
| Solyc11g012090.1 | Solyc11g012090.2 |
| Solyc11g012280.1 | Solyc11g012280.2 |
| Solyc11g012610.1 | Solyc11g012610.2 |
| Solyc11g012890.1 | Solyc11g012890.2 |
| Solyc11g013020.1 | Solyc11g013020.2 |
| Solyc11g013110.1 | Solyc11g013110.2 |
| Solyc11g013450.1 | Solyc11g013450.2 |

| | |
|---|---|
| Solyc11g013760.1 | Solyc11g013760.2 |
| Solyc11g013850.1 | Solyc11g013850.2 |
| Solyc11g016940.1 | Solyc11g016940.2 |
| Solyc11g017380.1 | Solyc11g017380.2 |
| Solyc11g017390.1 | Solyc11g017390.2 |
| Solyc11g018810.1 | Solyc11g018810.2 |
| Solyc11g019960.1 | Solyc11g019960.2 |
| Solyc11g020710.1 | Solyc11g020710.2 |
| Solyc11g021365.1 | Solyc11g021365.2 |
| Solyc11g031970.1 | Solyc11g031970.2 |
| Solyc11g045210.1 | Solyc11g045210.2 |
| Solyc11g045100.1 | Solyc11g045100.2 |
| Solyc11g062360.1 | Solyc11g062360.2 |
| Solyc11g062420.1 | Solyc11g062420.2 |
| Solyc11g064830.1 | Solyc11g064830.2 |
| Solyc11g064850.1 | Solyc11g064850.2 |
| Solyc11g065090.1 | Solyc11g065090.2 |
| Solyc11g065270.1 | Solyc11g065270.2 |
| Solyc11g066010.1 | Solyc11g066010.2 |
| Solyc11g066070.1 | Solyc11g066070.2 |
| Solyc11g066080.1 | Solyc11g066080.2 |
| Solyc11g066310.1 | Solyc11g066310.2 |
| Solyc11g066340.1 | Solyc11g066340.2 |
| Solyc11g066450.1 | Solyc11g066450.2 |
| Solyc11g066860.1 | Solyc11g066860.2 |
| Solyc11g066940.1 | Solyc11g066940.2 |
| Solyc11g067030.1 | Solyc11g067030.2 |
| Solyc11g067150.1 | Solyc11g067150.2 |
| Solyc11g068510.1 | Solyc11g068510.2 |
| Solyc11g068680.1 | Solyc11g068680.2 |
| Solyc11g068760.1 | Solyc11g068760.2 |
| Solyc11g068790.1 | Solyc11g068790.2 |
| Solyc11g068810.1 | Solyc11g068810.2 |
| Solyc11g069110.1 | Solyc11g069110.2 |
| Solyc11g069210.1 | Solyc11g069210.2 |
| Solyc11g069410.1 | Solyc11g069410.2 |
| Solyc11g069420.1 | Solyc11g069420.2 |
| Solyc11g069560.1 | Solyc11g069560.2 |
| Solyc11g070050.1 | Solyc11g070050.2 |
| Solyc11g070060.1 | Solyc11g070060.2 |
| Solyc11g071220.1 | Solyc11g071220.2 |
| Solyc11g072250.1 | Solyc11g072250.2 |
| Solyc11g073140.1 | Solyc11g073140.2 |

| | |
|---|---|
| Solyc12g005120.1 | Solyc12g005120.2 |
| Solyc12g005570.1 | Solyc12g005570.2 |
| Solyc12g005650.1 | Solyc12g005650.2 |
| Solyc12g006030.1 | Solyc12g006030.2 |
| Solyc12g006160.1 | Solyc12g006160.2 |
| Solyc12g006440.1 | Solyc12g006440.2 |
| Solyc12g006660.1 | Solyc12g006660.2 |
| Solyc12g007090.1 | Solyc12g007090.2 |
| Solyc12g008490.1 | Solyc12g008490.2 |
| Solyc12g008950.1 | Solyc12g008950.2 |
| Solyc12g009970.1 | Solyc12g009970.2 |
| Solyc12g010400.1 | Solyc12g010400.2 |
| Solyc12g010830.1 | Solyc12g010830.2 |
| Solyc12g013500.1 | Solyc12g013500.2 |
| Solyc12g013550.1 | Solyc12g013550.2 |
| Solyc12g013890.1 | Solyc12g013890.2 |
| Solyc12g014020.1 | Solyc12g014020.2 |
| Solyc12g014090.1 | Solyc12g014090.2 |
| Solyc12g014310.1 | Solyc12g014310.2 |
| Solyc12g014480.1 | Solyc12g014480.2 |
| Solyc12g015630.1 | Solyc12g015630.2 |
| Solyc12g016210.1 | Solyc12g016210.2 |
| Solyc12g017230.1 | Solyc12g017230.2 |
| Solyc12g017470.1 | Solyc12g017470.2 |
| Solyc12g019470.1 | Solyc12g019470.2 |
| Solyc12g019550.1 | Solyc12g019550.2 |
| Solyc12g077440.1 | Solyc12g077440.2 |
| Solyc12g036650.1 | Solyc12g036650.2 |
| Solyc12g036935.1 | Solyc12g150113.1 |
| Solyc12g040360.1 | Solyc12g040360.2 |
| Solyc12g040460.1 | Solyc12g040460.2 |
| Solyc12g041880.1 | Solyc12g041880.2 |
| Solyc12g042340.1 | Solyc12g042340.2 |
| Solyc12g043070.1 | Solyc12g043070.2 |
| Solyc12g044620.1 | Solyc12g044620.2 |
| Solyc12g044850.1 | Solyc12g044850.2 |
| Solyc12g055680.1 | Solyc12g055680.2 |
| Solyc12g055950.1 | Solyc12g055950.2 |
| Solyc12g056030.1 | Solyc12g056030.2 |
| Solyc12g056170.1 | Solyc12g056170.2 |
| Solyc12g056290.1 | Solyc12g056290.2 |
| Solyc12g056760.1 | Solyc12g056760.2 |
| Solyc12g056880.1 | Solyc12g056880.2 |

| | |
|---|---|
| Solyc12g056910.1 | Solyc12g056910.2 |
| Solyc12g087960.1 | Solyc12g087960.2 |
| Solyc12g087970.1 | Solyc12g087970.2 |
| Solyc12g088300.1 | Solyc12g088300.2 |
| Solyc12g094510.1 | Solyc12g094510.2 |
| Solyc12g096170.1 | Solyc12g096170.2 |
| Solyc12g096310.1 | Solyc12g096310.2 |
| Solyc12g098220.1 | Solyc12g098220.2 |
| Solyc12g098450.1 | Solyc12g098450.2 |
| Solyc12g099020.1 | Solyc12g099020.2 |
| Solyc12g099400.1 | Solyc12g099400.2 |
| Solyc12g099500.1 | Solyc12g099500.2 |
| Solyc12g099780.1 | Solyc12g099780.2 |
| Solyc12g099920.1 | Solyc12g099920.2 |
| Solyc12g099940.1 | Solyc12g099940.2 |
| Solyc12g100320.1 | Solyc12g100320.2 |
| Solyc01g005100.2 | Solyc01g005100.4 |
| Solyc01g005570.2 | Solyc01g005570.4 |
| Solyc01g005980.2 | Solyc01g005980.4 |
| Solyc01g006130.2 | Solyc01g006130.4 |
| Solyc01g007790.2 | Solyc01g007790.4 |
| Solyc01g008700.2 | Solyc01g008700.4 |
| Solyc01g009030.2 | Solyc01g009030.4 |
| Solyc01g009740.1 | Solyc01g009740.3 |
| Solyc01g009850.2 | Solyc01g009850.4 |
| Solyc01g015020.2 | Solyc01g015020.4 |
| Solyc01g020240.2 | Solyc01g020240.4 |
| Solyc01g049960.2 | Solyc01g049960.4 |
| Solyc01g058320.2 | Solyc01g058320.4 |
| Solyc01g060260.2 | Solyc01g060260.4 |
| Solyc01g066900.2 | Solyc01g066900.4 |
| Solyc01g067580.2 | Solyc01g067580.4 |
| Solyc01g068070.2 | Solyc01g068070.4 |
| Solyc01g068130.2 | Solyc01g068130.4 |
| Solyc01g068470.2 | Solyc01g068470.4 |
| Solyc01g079640.2 | Solyc01g079640.4 |
| Solyc01g080760.2 | Solyc01g080760.4 |
| Solyc01g086730.2 | Solyc01g086730.4 |
| Solyc01g087080.2 | Solyc01g087080.4 |
| Solyc01g087180.2 | Solyc01g087180.4 |
| Solyc01g090500.2 | Solyc01g090500.4 |
| Solyc01g090770.2 | Solyc01g090770.4 |
| Solyc01g091360.2 | Solyc01g091360.4 |

| | |
|---|---|
| Solyc01g094770.2 | Solyc01g094770.4 |
| Solyc01g094780.2 | Solyc01g094780.4 |
| Solyc01g096240.2 | Solyc01g096240.4 |
| Solyc01g096260.2 | Solyc01g096260.4 |
| Solyc01g097360.2 | Solyc01g097360.4 |
| Solyc01g097860.2 | Solyc01g097860.4 |
| Solyc01g098450.2 | Solyc01g098450.4 |
| Solyc01g098990.2 | Solyc01g098990.4 |
| Solyc01g101010.2 | Solyc01g101010.4 |
| Solyc01g102460.2 | Solyc01g102460.4 |
| Solyc01g103020.2 | Solyc01g103020.4 |
| Solyc01g103200.2 | Solyc01g103200.4 |
| Solyc01g103400.2 | Solyc01g103400.4 |
| Solyc01g103500.2 | Solyc01g103500.4 |
| Solyc01g104160.2 | Solyc01g104160.4 |
| Solyc01g104430.1 | Solyc01g104430.3 |
| Solyc01g104460.2 | Solyc01g104460.4 |
| Solyc01g104480.2 | Solyc01g104480.4 |
| Solyc01g105130.2 | Solyc01g105130.4 |
| Solyc01g105570.2 | Solyc01g105570.4 |
| Solyc01g106190.2 | Solyc01g106190.4 |
| Solyc01g106540.2 | Solyc01g106540.4 |
| Solyc01g106560.2 | Solyc01g106560.4 |
| Solyc01g106680.2 | Solyc01g106680.4 |
| Solyc01g106810.2 | Solyc01g106810.4 |
| Solyc01g106910.2 | Solyc01g106910.4 |
| Solyc01g107210.2 | Solyc01g107210.4 |
| Solyc01g107260.2 | Solyc01g107260.4 |
| Solyc01g107450.2 | Solyc01g107450.4 |
| Solyc01g108910.2 | Solyc01g108910.4 |
| Solyc01g109030.2 | Solyc01g109030.4 |
| Solyc01g110160.2 | Solyc01g110160.4 |
| Solyc01g111190.2 | Solyc01g111190.4 |
| Solyc01g111670.2 | Solyc01g111670.4 |
| Solyc01g112260.2 | Solyc01g112260.4 |
| Solyc02g020890.2 | Solyc02g020890.4 |
| Solyc02g022870.2 | Solyc02g022870.4 |
| Solyc02g031940.1 | Solyc02g031940.3 |
| Solyc02g049070.2 | Solyc02g049070.4 |
| Solyc02g055440.2 | Solyc02g055440.4 |
| Solyc02g055460.1 | Solyc02g055460.3 |
| Solyc02g063280.2 | Solyc02g063280.4 |
| Solyc02g065500.2 | Solyc02g065500.4 |

| | |
|---|---|
| Solyc02g067070.2 | Solyc02g067070.4 |
| Solyc02g067120.2 | Solyc02g067120.4 |
| Solyc02g067150.2 | Solyc02g067150.4 |
| Solyc02g067740.2 | Solyc02g067740.4 |
| Solyc02g068270.2 | Solyc02g068270.4 |
| Solyc02g071600.2 | Solyc02g071600.4 |
| Solyc02g071830.1 | Solyc02g071830.3 |
| Solyc02g076870.2 | Solyc02g076870.4 |
| Solyc02g078010.2 | Solyc02g078010.4 |
| Solyc02g078290.2 | Solyc02g078290.4 |
| Solyc02g078720.2 | Solyc02g078720.4 |
| Solyc02g080400.1 | Solyc02g080400.3 |
| Solyc02g080680.1 | Solyc02g080680.3 |
| Solyc02g081440.2 | Solyc02g081440.4 |
| Solyc02g081840.2 | Solyc02g081840.4 |
| Solyc02g082210.2 | Solyc02g082210.4 |
| Solyc02g082570.1 | Solyc02g082570.3 |
| Solyc02g082680.2 | Solyc02g082680.4 |
| Solyc02g083890.2 | Solyc02g083890.4 |
| Solyc02g084250.2 | Solyc02g084250.4 |
| Solyc02g085080.2 | Solyc02g085080.4 |
| Solyc02g086250.2 | Solyc02g086250.4 |
| Solyc02g087420.1 | Solyc02g087420.3 |
| Solyc02g087470.2 | Solyc02g087470.4 |
| Solyc02g087650.2 | Solyc02g087650.4 |
| Solyc02g087940.2 | Solyc02g087940.4 |
| Solyc02g088200.2 | Solyc02g088200.4 |
| Solyc02g088350.2 | Solyc02g088350.4 |
| Solyc02g088830.2 | Solyc02g088830.4 |
| Solyc02g088930.2 | Solyc02g088930.4 |
| Solyc02g090260.1 | Solyc02g090260.3 |
| Solyc02g090650.2 | Solyc02g090650.4 |
| Solyc02g090760.2 | Solyc02g090760.4 |
| Solyc02g091320.2 | Solyc02g091320.4 |
| Solyc02g091620.1 | Solyc02g091620.3 |
| Solyc02g092830.2 | Solyc02g092830.4 |
| Solyc02g094490.2 | Solyc02g094490.4 |
| Solyc02g094540.2 | Solyc02g094540.4 |
| Solyc03g006280.2 | Solyc03g006280.4 |
| Solyc03g007340.2 | Solyc03g007340.4 |
| Solyc03g007550.2 | Solyc03g007550.4 |
| Solyc03g013240.2 | Solyc03g013240.4 |
| Solyc03g013250.2 | Solyc03g013250.4 |

| | |
|---|---|
| Solyc03g019840.2 | Solyc03g019840.4 |
| Solyc03g019930.2 | Solyc03g019930.4 |
| Solyc03g025690.2 | Solyc03g025690.4 |
| Solyc03g025820.1 | Solyc03g025820.3 |
| Solyc03g026010.1 | Solyc03g026010.3 |
| Solyc03g026330.2 | Solyc03g026330.4 |
| Solyc03g031950.1 | Solyc03g031950.3 |
| Solyc03g032110.1 | Solyc03g032110.3 |
| Solyc03g033720.2 | Solyc03g033720.4 |
| Solyc03g034170.2 | Solyc03g034170.4 |
| Solyc03g034250.2 | Solyc03g034250.4 |
| Solyc03g043970.1 | Solyc03g043970.3 |
| Solyc03g044080.2 | Solyc03g044080.4 |
| Solyc03g053000.2 | Solyc03g053000.4 |
| Solyc03g053010.2 | Solyc03g053010.4 |
| Solyc03g053020.2 | Solyc03g053020.4 |
| Solyc03g062700.2 | Solyc03g062700.4 |
| Solyc03g062930.1 | Solyc03g062930.3 |
| Solyc03g081200.1 | Solyc03g081200.3 |
| Solyc03g082490.2 | Solyc03g082490.4 |
| Solyc03g082540.2 | Solyc03g082540.4 |
| Solyc03g082680.2 | Solyc03g082680.4 |
| Solyc03g083120.1 | Solyc03g083120.3 |
| Solyc03g083190.1 | Solyc03g083190.3 |
| Solyc03g083200.2 | Solyc03g083200.4 |
| Solyc03g083420.2 | Solyc03g083420.4 |
| Solyc03g093340.2 | Solyc03g093340.4 |
| Solyc03g096840.2 | Solyc03g096840.4 |
| Solyc03g097710.2 | Solyc03g097710.4 |
| Solyc03g098110.2 | Solyc03g098110.4 |
| Solyc03g110980.2 | Solyc03g110980.4 |
| Solyc03g113640.2 | Solyc03g113640.4 |
| Solyc03g113660.2 | Solyc03g113660.4 |
| Solyc03g115100.2 | Solyc03g115100.4 |
| Solyc03g115460.1 | Solyc03g115460.3 |
| Solyc03g116800.2 | Solyc03g116800.4 |
| Solyc03g117210.2 | Solyc03g117210.4 |
| Solyc03g117280.2 | Solyc03g117280.4 |
| Solyc03g117660.2 | Solyc03g117660.4 |
| Solyc03g117840.2 | Solyc03g117840.4 |
| Solyc03g118440.1 | Solyc03g118440.3 |
| Solyc03g118520.2 | Solyc03g118520.4 |
| Solyc03g119110.2 | Solyc03g119110.4 |

| | |
|---|---|
| Solyc03g119350.1 | Solyc03g119350.3 |
| Solyc03g119480.2 | Solyc03g119480.4 |
| Solyc03g119750.2 | Solyc03g119750.4 |
| Solyc03g122370.2 | Solyc03g122370.4 |
| Solyc03g123580.2 | Solyc03g123580.4 |
| Solyc03g123700.2 | Solyc03g123700.4 |
| Solyc04g007110.2 | Solyc04g007110.4 |
| Solyc04g007220.2 | Solyc04g007220.4 |
| Solyc04g007340.2 | Solyc04g007340.4 |
| Solyc04g007410.2 | Solyc04g007410.4 |
| Solyc04g009050.2 | Solyc04g009050.4 |
| Solyc04g009370.2 | Solyc04g009370.4 |
| Solyc04g011460.1 | Solyc04g011460.3 |
| Solyc04g015080.2 | Solyc04g015080.4 |
| Solyc04g015430.2 | Solyc04g015430.4 |
| Solyc04g015730.1 | Solyc04g015730.3 |
| Solyc04g045580.2 | Solyc04g045580.4 |
| Solyc04g049660.1 | Solyc04g049660.3 |
| Solyc04g049970.2 | Solyc04g049970.4 |
| Solyc04g050160.2 | Solyc04g050160.4 |
| Solyc04g051120.2 | Solyc04g051120.4 |
| Solyc04g051280.2 | Solyc04g051280.4 |
| Solyc04g051310.2 | Solyc04g051310.4 |
| Solyc04g051570.2 | Solyc04g051570.4 |
| Solyc04g051820.2 | Solyc04g051820.4 |
| Solyc04g054180.2 | Solyc04g054180.4 |
| Solyc04g055250.2 | Solyc04g055250.4 |
| Solyc04g071500.2 | Solyc04g071500.4 |
| Solyc04g072050.2 | Solyc04g072050.4 |
| Solyc04g072210.2 | Solyc04g072210.4 |
| Solyc04g072400.2 | Solyc04g072400.4 |
| Solyc04g072630.2 | Solyc04g072630.4 |
| Solyc04g072940.2 | Solyc04g072940.4 |
| Solyc04g074140.2 | Solyc04g074140.4 |
| Solyc04g074660.1 | Solyc04g074660.3 |
| Solyc04g076100.2 | Solyc04g076100.4 |
| Solyc04g076360.2 | Solyc04g076360.4 |
| Solyc04g077420.2 | Solyc04g077420.4 |
| Solyc04g077500.2 | Solyc04g077500.4 |
| Solyc04g077820.2 | Solyc04g077820.4 |
| Solyc04g077940.2 | Solyc04g077940.4 |
| Solyc04g078010.2 | Solyc04g078010.4 |
| Solyc04g078160.2 | Solyc04g078160.4 |

| | |
|---|---|
| Solyc04g079160.2 | Solyc04g079160.4 |
| Solyc04g079170.2 | Solyc04g079170.4 |
| Solyc04g080140.2 | Solyc04g080140.4 |
| Solyc04g080200.2 | Solyc04g080200.4 |
| Solyc04g080270.2 | Solyc04g080270.4 |
| Solyc04g080390.2 | Solyc04g080390.4 |
| Solyc04g081420.2 | Solyc04g081420.4 |
| Solyc04g081470.2 | Solyc04g081470.4 |
| Solyc04g081850.2 | Solyc04g081850.4 |
| Solyc04g082290.2 | Solyc04g082290.4 |
| Solyc04g082970.2 | Solyc04g082970.4 |
| Solyc05g005030.2 | Solyc05g005030.4 |
| Solyc05g005730.2 | Solyc05g005730.4 |
| Solyc05g006040.2 | Solyc05g006040.4 |
| Solyc05g006440.2 | Solyc05g006440.4 |
| Solyc05g007030.2 | Solyc05g007030.4 |
| Solyc05g007570.2 | Solyc05g007570.4 |
| Solyc05g008380.2 | Solyc05g008380.4 |
| Solyc05g009200.1 | Solyc05g009200.3 |
| Solyc05g009360.2 | Solyc05g009360.4 |
| Solyc05g009760.1 | Solyc05g009760.3 |
| Solyc05g009940.2 | Solyc05g009940.4 |
| Solyc05g010290.1 | Solyc05g010290.3 |
| Solyc05g010460.2 | Solyc05g010460.4 |
| Solyc05g010760.1 | Solyc05g010760.3 |
| Solyc05g010790.1 | Solyc05g010790.3 |
| Solyc05g011830.2 | Solyc05g011830.4 |
| Solyc05g012230.2 | Solyc05g012230.4 |
| Solyc05g012240.1 | Solyc05g012240.3 |
| Solyc05g012620.2 | Solyc05g012620.4 |
| Solyc05g013060.2 | Solyc05g013060.4 |
| Solyc05g013240.2 | Solyc05g013240.4 |
| Solyc05g013480.2 | Solyc05g013480.4 |
| Solyc05g014190.2 | Solyc05g014190.4 |
| Solyc05g015890.2 | Solyc05g015890.4 |
| Solyc05g026060.1 | Solyc05g026060.3 |
| Solyc05g026080.2 | Solyc05g026080.4 |
| Solyc05g050550.2 | Solyc05g050550.4 |
| Solyc05g050720.2 | Solyc05g050720.4 |
| Solyc05g051710.2 | Solyc05g051710.4 |
| Solyc05g052170.2 | Solyc05g052170.4 |
| Solyc05g052630.2 | Solyc05g052630.4 |
| Solyc05g053880.2 | Solyc05g053880.4 |

| | |
|---|---|
| Solyc05g055360.2 | Solyc05g055360.4 |
| Solyc05g055680.2 | Solyc05g055680.4 |
| Solyc05g056080.2 | Solyc05g056080.4 |
| Solyc06g005660.1 | Solyc06g005660.3 |
| Solyc06g007990.2 | Solyc06g007990.4 |
| Solyc06g009690.1 | Solyc06g009690.3 |
| Solyc06g016760.1 | Solyc06g016760.3 |
| Solyc06g036460.2 | Solyc06g036460.4 |
| Solyc06g036640.2 | Solyc06g036640.4 |
| Solyc06g042960.1 | Solyc06g042960.3 |
| Solyc06g043070.2 | Solyc06g043070.4 |
| Solyc06g051140.2 | Solyc06g051140.4 |
| Solyc06g051790.2 | Solyc06g051790.4 |
| Solyc06g053230.2 | Solyc06g053230.4 |
| Solyc06g053920.2 | Solyc06g053920.4 |
| Solyc06g054140.2 | Solyc06g054140.4 |
| Solyc06g054200.1 | Solyc06g054200.3 |
| Solyc06g054450.1 | Solyc06g054450.3 |
| Solyc06g060160.1 | Solyc06g060160.3 |
| Solyc06g060890.2 | Solyc06g060890.4 |
| Solyc06g061130.2 | Solyc06g061130.4 |
| Solyc06g062970.1 | Solyc06g062970.3 |
| Solyc06g063010.2 | Solyc06g063010.4 |
| Solyc06g063050.2 | Solyc06g063050.4 |
| Solyc06g064560.2 | Solyc06g064560.4 |
| Solyc06g068690.2 | Solyc06g068690.4 |
| Solyc06g069170.2 | Solyc06g069170.4 |
| Solyc06g069630.2 | Solyc06g069630.4 |
| Solyc06g071010.2 | Solyc06g071010.4 |
| Solyc06g071040.2 | Solyc06g071040.4 |
| Solyc06g071170.2 | Solyc06g071170.4 |
| Solyc06g072940.1 | Solyc06g072940.3 |
| Solyc06g073160.2 | Solyc06g073160.4 |
| Solyc06g073450.2 | Solyc06g073450.4 |
| Solyc06g074260.2 | Solyc06g074260.4 |
| Solyc06g074540.2 | Solyc06g074540.4 |
| Solyc06g074900.1 | Solyc06g074900.3 |
| Solyc06g075410.2 | Solyc06g075410.4 |
| Solyc06g076370.2 | Solyc06g076370.4 |
| Solyc06g082320.2 | Solyc06g082320.4 |
| Solyc06g082850.2 | Solyc06g082850.4 |
| Solyc06g084470.2 | Solyc06g084470.4 |
| Solyc07g005550.2 | Solyc07g005550.4 |

| | |
|---|---|
| Solyc07g006320.2 | Solyc07g006320.4 |
| Solyc07g008100.2 | Solyc07g008100.4 |
| Solyc07g008280.2 | Solyc07g008280.4 |
| Solyc07g008770.2 | Solyc07g008770.4 |
| Solyc07g017290.2 | Solyc07g017290.4 |
| Solyc07g017670.1 | Solyc07g017670.3 |
| Solyc07g018310.2 | Solyc07g018310.4 |
| Solyc07g021170.1 | Solyc07g021170.3 |
| Solyc07g025390.2 | Solyc07g025390.4 |
| Solyc07g026640.1 | Solyc07g026640.3 |
| Solyc07g032250.2 | Solyc07g032250.4 |
| Solyc07g032290.1 | Solyc07g032290.3 |
| Solyc07g032710.2 | Solyc07g032710.4 |
| Solyc07g039190.2 | Solyc07g039190.4 |
| Solyc07g041330.1 | Solyc07g041330.3 |
| Solyc07g042500.2 | Solyc07g042500.4 |
| Solyc07g042700.2 | Solyc07g042700.4 |
| Solyc07g042910.2 | Solyc07g042910.4 |
| Solyc07g045140.2 | Solyc07g045140.4 |
| Solyc07g045260.1 | Solyc07g045260.3 |
| Solyc07g045300.2 | Solyc07g045300.4 |
| Solyc07g047700.2 | Solyc07g047700.4 |
| Solyc07g047890.2 | Solyc07g047890.4 |
| Solyc07g049200.2 | Solyc07g049200.4 |
| Solyc07g053320.2 | Solyc07g053320.4 |
| Solyc07g053440.2 | Solyc07g053440.4 |
| Solyc07g055910.2 | Solyc07g055910.4 |
| Solyc07g056060.2 | Solyc07g056060.4 |
| Solyc07g062090.1 | Solyc07g062090.3 |
| Solyc07g062410.2 | Solyc07g062410.4 |
| Solyc07g062740.2 | Solyc07g062740.4 |
| Solyc07g062750.2 | Solyc07g062750.4 |
| Solyc07g062920.2 | Solyc07g062920.4 |
| Solyc07g064080.2 | Solyc07g064080.4 |
| Solyc07g064320.1 | Solyc07g064320.3 |
| Solyc07g064350.1 | Solyc07g064350.3 |
| Solyc07g064540.1 | Solyc07g064540.3 |
| Solyc07g065720.2 | Solyc07g065720.4 |
| Solyc07g066200.2 | Solyc07g066200.4 |
| Solyc07g066530.2 | Solyc07g066530.4 |
| Solyc08g006100.2 | Solyc08g006100.4 |
| Solyc08g006800.1 | Solyc08g006800.3 |
| Solyc08g008180.1 | Solyc08g008180.3 |

| | |
|---|---|
| Solyc08g008270.2 | Solyc08g008270.4 |
| Solyc08g061450.1 | Solyc08g061450.3 |
| Solyc08g065260.2 | Solyc08g065260.4 |
| Solyc08g065440.2 | Solyc08g065440.4 |
| Solyc08g067430.2 | Solyc08g067430.4 |
| Solyc08g067880.2 | Solyc08g067880.4 |
| Solyc08g067980.1 | Solyc08g067980.3 |
| Solyc08g075420.2 | Solyc08g075420.4 |
| Solyc08g075520.2 | Solyc08g075520.4 |
| Solyc08g075680.2 | Solyc08g075680.4 |
| Solyc08g076740.2 | Solyc08g076740.4 |
| Solyc08g076770.2 | Solyc08g076770.4 |
| Solyc08g077250.2 | Solyc08g077250.4 |
| Solyc08g077450.2 | Solyc08g077450.4 |
| Solyc08g078520.2 | Solyc08g078520.4 |
| Solyc08g078610.2 | Solyc08g078610.4 |
| Solyc08g079030.2 | Solyc08g079030.4 |
| Solyc08g079240.2 | Solyc08g079240.4 |
| Solyc08g079590.1 | Solyc08g079590.3 |
| Solyc08g079690.2 | Solyc08g079690.4 |
| Solyc08g079810.2 | Solyc08g079810.4 |
| Solyc08g080860.2 | Solyc08g080860.4 |
| Solyc08g081130.2 | Solyc08g081130.4 |
| Solyc08g081760.1 | Solyc08g081760.3 |
| Solyc08g082000.2 | Solyc08g082000.4 |
| Solyc08g082410.2 | Solyc08g082410.4 |
| Solyc08g082570.2 | Solyc08g082570.4 |
| Solyc08g082720.2 | Solyc08g082720.4 |
| Solyc08g083030.2 | Solyc08g083030.4 |
| Solyc09g005890.2 | Solyc09g005890.4 |
| Solyc09g005930.2 | Solyc09g005930.4 |
| Solyc09g007460.2 | Solyc09g007460.4 |
| Solyc09g007950.1 | Solyc09g007950.3 |
| Solyc09g008680.2 | Solyc09g008680.4 |
| Solyc09g009220.2 | Solyc09g009220.4 |
| Solyc09g009710.2 | Solyc09g009710.4 |
| Solyc09g009990.2 | Solyc09g009990.4 |
| Solyc09g010550.2 | Solyc09g010550.4 |
| Solyc09g010710.2 | Solyc09g010710.4 |
| Solyc09g011020.2 | Solyc09g011020.4 |
| Solyc09g013120.2 | Solyc09g013120.4 |
| Solyc09g014490.2 | Solyc09g014490.4 |
| Solyc09g015460.1 | Solyc09g015460.3 |

| | |
|---|---|
| Solyc09g018310.2 | Solyc09g018310.4 |
| Solyc09g018370.2 | Solyc09g018370.4 |
| Solyc09g020150.2 | Solyc09g020150.4 |
| Solyc09g031900.2 | Solyc09g031900.4 |
| Solyc09g056160.2 | Solyc09g056160.4 |
| Solyc09g059450.2 | Solyc09g059450.4 |
| Solyc09g059470.2 | Solyc09g059470.4 |
| Solyc09g059560.1 | Solyc09g059560.3 |
| Solyc09g059910.1 | Solyc09g059910.3 |
| Solyc09g061440.2 | Solyc09g061440.4 |
| Solyc09g063090.2 | Solyc09g063090.4 |
| Solyc09g064420.2 | Solyc09g064420.4 |
| Solyc09g064760.1 | Solyc09g064760.3 |
| Solyc09g065110.2 | Solyc09g065110.4 |
| Solyc09g065160.2 | Solyc09g065160.4 |
| Solyc09g065300.2 | Solyc09g065300.4 |
| Solyc09g065930.2 | Solyc09g065930.4 |
| Solyc09g072840.1 | Solyc09g072840.3 |
| Solyc09g074170.1 | Solyc09g074170.3 |
| Solyc09g074840.2 | Solyc09g074840.4 |
| Solyc09g075600.1 | Solyc09g075600.3 |
| Solyc09g082130.2 | Solyc09g082130.4 |
| Solyc09g082160.2 | Solyc09g082160.4 |
| Solyc09g083390.1 | Solyc09g083390.3 |
| Solyc09g090050.2 | Solyc09g090050.4 |
| Solyc09g090340.1 | Solyc09g090340.3 |
| Solyc09g090690.2 | Solyc09g090690.4 |
| Solyc09g090820.2 | Solyc09g090820.4 |
| Solyc09g091270.1 | Solyc09g091270.3 |
| Solyc09g091370.2 | Solyc09g091370.4 |
| Solyc09g091820.2 | Solyc09g091820.4 |
| Solyc09g091830.2 | Solyc09g091830.4 |
| Solyc09g092550.2 | Solyc09g092550.4 |
| Solyc09g098140.2 | Solyc09g098140.4 |
| Solyc09g098530.2 | Solyc09g098530.4 |
| Solyc10g006430.1 | Solyc10g006430.3 |
| Solyc10g007120.2 | Solyc10g007120.4 |
| Solyc10g007670.2 | Solyc10g007670.4 |
| Solyc10g008350.2 | Solyc10g008350.4 |
| Solyc10g009040.2 | Solyc10g009040.4 |
| Solyc10g012030.2 | Solyc10g012030.4 |
| Solyc10g037890.1 | Solyc10g037890.3 |
| Solyc10g039430.1 | Solyc10g039430.3 |

| | |
|---|---|
| Solyc10g047540.1 | Solyc10g047540.3 |
| Solyc10g047960.1 | Solyc10g047960.3 |
| Solyc10g052650.1 | Solyc10g052650.3 |
| Solyc10g078900.1 | Solyc10g078900.3 |
| Solyc10g079010.1 | Solyc10g079010.3 |
| Solyc10g080450.1 | Solyc10g080450.3 |
| Solyc10g081700.1 | Solyc10g081700.3 |
| Solyc10g083910.1 | Solyc10g083910.3 |
| Solyc10g084620.1 | Solyc10g084620.3 |
| Solyc10g085080.1 | Solyc10g085080.3 |
| Solyc10g085930.1 | Solyc10g085930.3 |
| Solyc11g005390.1 | Solyc11g005390.3 |
| Solyc11g006880.1 | Solyc11g006880.3 |
| Solyc11g007810.1 | Solyc11g007810.3 |
| Solyc11g007840.1 | Solyc11g007840.3 |
| Solyc11g008100.1 | Solyc11g008100.3 |
| Solyc11g008840.1 | Solyc11g008840.3 |
| Solyc11g010490.1 | Solyc11g010490.3 |
| Solyc11g010620.1 | Solyc11g010620.3 |
| Solyc11g011060.1 | Solyc11g011060.3 |
| Solyc11g011760.1 | Solyc11g011760.3 |
| Solyc11g012600.1 | Solyc11g012600.3 |
| Solyc11g012920.1 | Solyc11g012920.3 |
| Solyc11g013330.1 | Solyc11g013330.3 |
| Solyc11g016980.1 | Solyc11g016980.3 |
| Solyc11g017430.1 | Solyc11g017430.3 |
| Solyc11g018470.1 | Solyc11g018470.3 |
| Solyc11g020910.1 | Solyc11g020910.3 |
| Solyc11g030790.1 | Solyc11g030790.3 |
| Solyc11g032090.1 | Solyc11g032090.3 |
| Solyc11g045270.1 | Solyc11g045270.3 |
| Solyc11g045510.1 | Solyc11g045510.3 |
| Solyc11g056620.1 | Solyc11g056620.3 |
| Solyc11g056640.1 | Solyc11g056640.3 |
| Solyc11g064930.1 | Solyc11g064930.3 |
| Solyc11g065630.1 | Solyc11g065630.3 |
| Solyc11g066240.1 | Solyc11g066240.3 |
| Solyc11g066870.1 | Solyc11g066870.3 |
| Solyc11g069080.1 | Solyc11g069080.3 |
| Solyc11g070020.1 | Solyc11g070020.3 |
| Solyc11g071190.1 | Solyc11g071190.3 |
| Solyc11g071950.1 | Solyc11g071950.3 |
| Solyc11g072430.1 | Solyc11g072430.3 |

| | |
|---|---|
| Solyc12g005090.1 | Solyc12g005090.3 |
| Solyc12g005820.1 | Solyc12g005820.3 |
| Solyc12g006230.1 | Solyc12g006230.3 |
| Solyc12g006670.1 | Solyc12g006670.3 |
| Solyc12g007140.1 | Solyc12g007140.3 |
| Solyc12g007240.1 | Solyc12g007240.3 |
| Solyc12g009900.1 | Solyc12g009900.3 |
| Solyc12g010150.1 | Solyc12g010150.3 |
| Solyc12g010640.1 | Solyc12g010640.3 |
| Solyc12g014220.1 | Solyc12g014220.3 |
| Solyc12g015730.1 | Solyc12g015730.3 |
| Solyc12g019040.1 | Solyc12g019040.3 |
| Solyc12g020000.1 | Solyc12g020000.3 |
| Solyc12g027740.1 | Solyc12g027740.3 |
| Solyc12g038240.1 | Solyc12g038240.3 |
| Solyc12g038250.1 | Solyc12g038250.3 |
| Solyc12g042120.1 | Solyc12g042120.3 |
| Solyc12g042700.1 | Solyc12g042700.3 |
| Solyc12g042720.1 | Solyc12g042720.3 |
| Solyc12g042960.1 | Solyc12g042960.3 |
| Solyc12g042990.1 | Solyc12g042990.3 |
| Solyc12g044760.1 | Solyc12g044760.3 |
| Solyc12g049380.1 | Solyc12g049380.3 |
| Solyc12g049510.1 | Solyc12g049510.3 |
| Solyc12g055700.1 | Solyc12g055700.3 |
| Solyc12g055940.1 | Solyc12g055940.3 |
| Solyc12g056060.1 | Solyc12g056060.3 |
| Solyc12g057130.1 | Solyc12g057130.3 |
| Solyc12g087990.1 | Solyc12g087990.3 |
| Solyc12g088410.1 | Solyc12g088410.3 |
| Solyc12g088450.1 | Solyc12g088450.3 |
| Solyc12g094480.1 | Solyc12g094480.3 |
| Solyc12g094580.1 | Solyc12g094580.3 |
| Solyc12g099490.1 | Solyc12g099490.3 |
